# Electrocardiogram feature extraction and interval measurements using optimal representative cycles from persistent homology

**DOI:** 10.1101/2022.02.01.478609

**Authors:** Hunter Dlugas

**Author notes:** 1104 Faculty Administration Building, Wayne State University, Detroit, MI, USA.

## Abstract

Cardiovascular diseases are among the leading causes of death, and their early detection and treatment is important for lowering their prevalence and mortality rate. Electrocardiograms (ECGs) record electrical activity of the heart to provide information used to diagnose and treat various cardiovascular diseases. Many approaches to computer-aided ECG analysis have been performed, including Fourier analysis, principal component analysis, analyzing morphological changes, and machine learning. Due to the high accuracy required of ECG-analysis software, there is no universally-agreed upon algorithm to identify P,Q,R,S, and T-waves and measure intervals of interest. Topological data analysis uses tools from algebraic topology to quantify hole-like shapes within data, and methods using persistence statistics and fractal dimension with machine learning have been applied to ECG signals in the context of detecting arrhythmias within recent years. To our knowledge, there does not exist a method of identifying P,Q,S, and T-waves and measuring intervals of interest which relies on topological features of the data, and we propose a novel topological method for performing these aspects of ECG analysis. Specifically, we establish criteria to identify cardinality-minimal and area-minimal 1-cycles with certain properties as P,Q,S, and T-waves. This yields a procedure for measuring the PR-interval, QT-interval, ST-segment, QRS-duration, P-wave duration, and T-wave duration in Lead II ECG data. We apply our procedure to 400 sets of simulated Lead II ECG signals and compare with the interval values set by the model. Additionally, the algorithm is used to identify cardinality-minimal and area-minimal 1-cycles as P,Q,S, and T-waves in two sets of 200 randomly sampled Lead II ECG signals of real patients with 11 common rhythms. Analysis of optimal 1-cycles identified as P,Q,S, and T-waves and comparison of interval measurements shows that 1-cycle reconstructions can provide useful information about the ECG signal and could hold utility in characterizing arrhythmias.

**Author summary:** Topological data analysis (TDA) has been a rapidly growing field within the past 15 years and has found applications across many fields. In the context of TDA, several algorithms primarily using persistence barcode statistics and machine learning have been applied to electrocardiogram (ECG) signals in recent years. We use a topological data-analytic method to identify subsets of an ECG signal which are representative of certain topological features in the ECG signal, and we propose that those subsets coincide with the P,Q,S, and T-waves in the ECG signal. We then use information about these subsets of the signal identified as P,Q,S, and T-waves to measure the PR-interval, QT-interval, ST-segment, QRS-duration, P-wave duration, and T-wave duration. We demonstrate our method on both simulated and real Lead II ECG data. These results show how identifying subsets of an ECG signal with certain topological properties could be used in analyzing the morphology of the signal over time and in arrhythmia-detection algorithms.

## Introduction

Cardiovascular diseases are among the leading causes of death due to their high prevalence and mortality rate [1] [2] [3]. Electrocardiograms (ECGs) provide a non-invasive measure of the heart’s electrical activity and are used in diagnosing and managing various cardiovascular diseases. Thus the analysis of ECGs is important for accurate diagnosis and proper treatment of cardiovascular diseases. Several approaches to automated ECG analysis have been performed, including machine learning [4] [5] [6] [7] [8] [9] [10] [11], wavelet transforms [12] [13] [14] [15] [16], and persistent homology [17] [18] [19] [20] [21] [22] [23]. Due to the high accuracy required of ECG-analysis software and the fact that the bulk of ECG analysis is carried out by healthcare providers, the development of algorithms that identifies P,Q,R,S, and T-waves, measures intervals of interest, and/or detects arrhythmias is an active area of research.

Topological data analysis (TDA) is concerned with the study of shapes constructed from a dataset which are invariant under continuous deformations such as stretching and twisting. Applications of TDA to ECG signals have used persistence statistics [20] [18], fractal dimension [18], and machine learning [17] [19]. Using cycle reconstructions has shown utility in various applications outside of ECG analysis such as analyzing structures on the atomic scale [24] and in structural engineering [25]. To our knowledge, nobody has performed an “inverse” analysis of using information from cycle reconstructions to analyze ECG signals. We focus our attention on the “rhythm lead”, i.e. Lead II, and for the remainder of the paper, any reference to an ECG signal, whether simulated or real, refers to a Lead II ECG signal.

Suppose we are given an *n*-dimensional dataset, that is, a set of points in ℝ^*n*^. There are *n* distinct types of topological features associated with that dataset: roughly, these”types” correspond to what one would intuitively regard as “holes” or “gaps” of differing dimensions in the dataset. To apply these methods to ECG data, we treat an ECG signal as two dimensional with one temporal dimension and one amplitude dimension rather than one dimensional with a specified sampling frequency. Consequently, from this perspective, there are two distinct types of topological features associated with ECG data: 0-dimensional homology features represent equivalence classes of connected components, while 1-dimensional homology features represent equivalence classes of non-contractible loops. The 0-dimensional homology features are useful in analyzing clustering phenomena and have recently shown utility as a metric of heart rate variability when applied to ECG signals [21] [22].

In this paper, our focus is instead on the 1-dimensional homology features, which we use in a novel way to analyze ECG data. Specifically, we identify subsets of an ECG signal as P,Q,S, and T-waves by considering these subsets to be representative cycles of 1-dimensional homology features of the signal with certain properties and then use these subsets identified as individual waves to measure the PR-interval, QT-interval, ST-segment, QRS-duration, P-wave duration, and T-wave duration. To illustrate the intuition behind 1-dimensional homology features and their representative cycles, we first present an example which demonstrates how the 1-dimensional homology features of a simple 2-dimensional dataset is related to the shape of the data in the “Example: a 1-dimensional topological feature in a simple dataset” section. This section describes some geometric intuition of 1-dimensional homology features without going into linear algebra details. These details are then given and the ideas from this section are formalized in the “Background on topological data analysis” section, where we define key terms in algebraic and computational topology to provide minimal background and establish terms that will be used in identifying features of ECG signals.

### Example: a 1-dimensional topological feature in a simple dataset

Consider the set of points in the Cartesian plane ℝ^2^ shown in Fig 1A. Then consider a circle drawn around each datum, each with the same radius *r*. The geometric Čech complex of radius *r* is defined as the union of the interiors of these circles. It is a subset of ℝ^2^, and as *r* increases, the geometric Čech complex of radius *r* is a larger and larger subset of ℝ^2^. Fig 1B-E depicts the geometric Čech complexes of radius 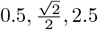, and 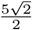 as the region shaded in blue. Notice that for *r <* 0.5, none of the circles overlap, whereas for 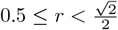, the circles centered at the data comprising the smaller square overlap such that there is a “void” of non-overlapping space enclosed by their region of overlap. Hence for 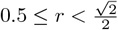, there exists a non-contractible loop within the geometric Čech complex of radius *r*. “Non-contractible” here means that the loop drawn around the void of non-overlapping space cannot be continuously deformed down to a single point without leaving the geometric Čech complex: the loop gets “stuck” on the void encircled by the geometric Čech complex, like trying to pull a rubber band off of a broomstick of infinite length. Notice that for *r <* 0.5, none of the circles of the example dataset in Fig 1A overlap, let alone overlap in such a way that they encircle some non-overlapping space. Consequently, we are unable to construct non-contractible loops within the geometric Čech complex of radius *r* for *r <* 0.5.

**Fig 1.**
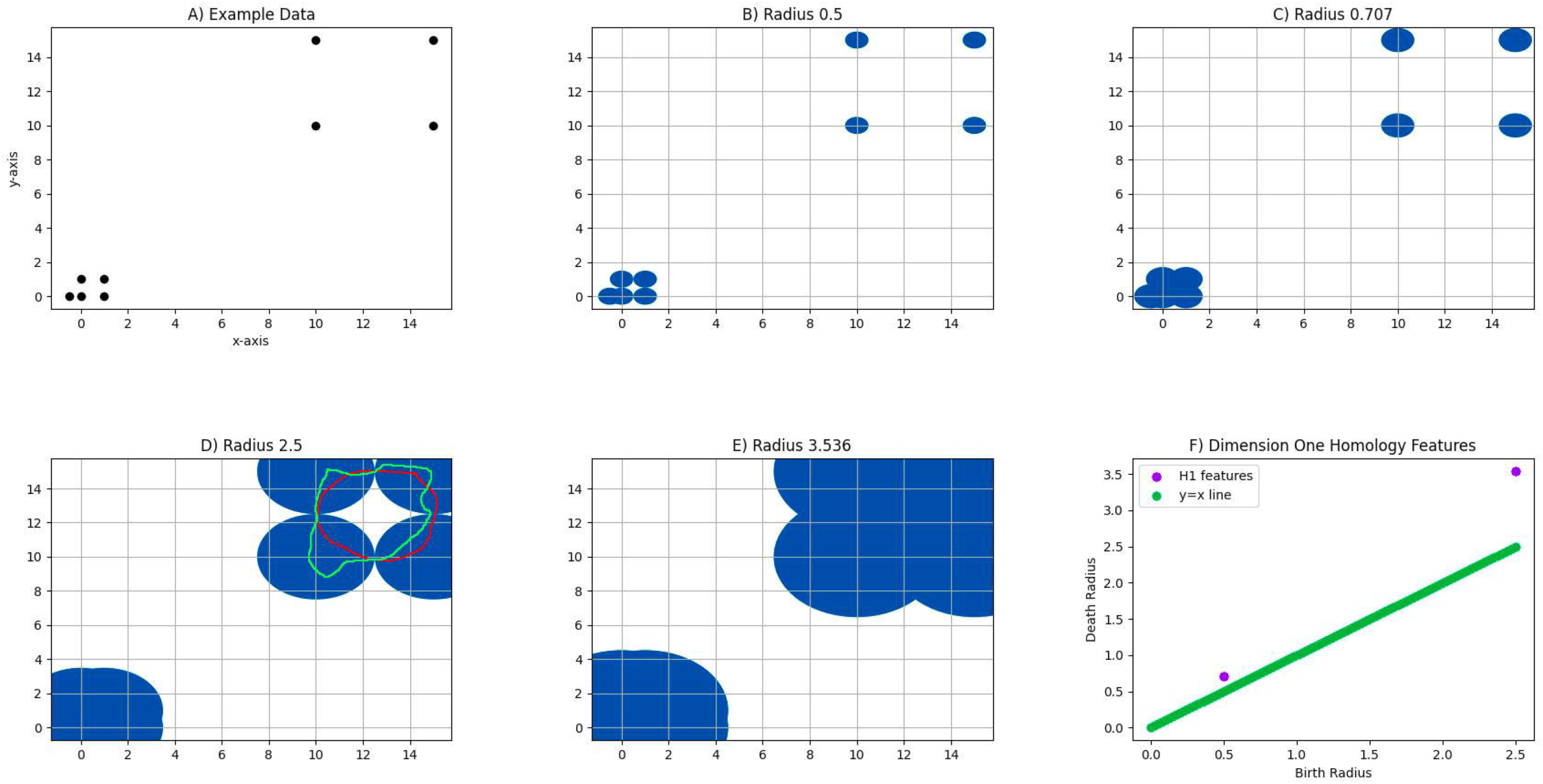
1-dimensional homological features of example dataset in ℝ^2^. A: Plot of example data. B,C,D,E: The blue region depicts the geometric Čech complex of radius 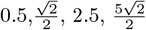, respectively. F: Persistence diagram: each blue triangle represents a 1-dimensional homological feature, i.e. an equivalence class of non-contractible loops, with coordinates (birth radius, death radius); the y=x line is drawn to depict the persistence distribution of all 1-dimensional homological features over a set of *r* values by noting that the more persistent a 1-dimensional homological feature is, i.e. the larger the difference between its death radius and birth radius, the more ‘above’ the y=x line it will be.

Recall that if we can draw a non-contractible loop within the geometric Čech complex of radius *r* for some *r >* 0, then this loop must be “stuck” around some void encircled by the geometric Čech complex. This non-contractible loop can be continuously deformed to construct another non-contractible loop “stuck” around the same void. We say the two loops are homotopic. As an example, the red and green loops shown in Fig 1D are homotopic. The set of all possible non-contractible loops “stuck” around some void encircled by the geometric Čech complex forms an equivalence class of non-contractible loops, i.e. a set of non-contractible loops where any two non-contractible loops in the set are homotopic. In practice, rather than homotopy, we use a weaker but more technically-involved equivalence relation on loops called homology to utilize efficient algorithms such as Ripser [26] and GUDHI [27] in computing topological features. For a rigorous treatment of homotopy and homology, see [28].

Suppose we are given a dataset *X* and a positive real number *r*. We write 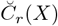 for the geometric Čech complex of *X* of radius *r*. For a given two-dimensional dataset such that there exists a non-contractible loop 𝓁 within its geometric Čech complex of radius *r*, we define the *birth radius* of 𝓁 as the smallest real number *b* such that some loop in 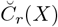 which is equivalent to 𝓁 and which is contained in the subset 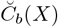of 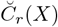 exists. It follows immediately from this definition that *b*≤ *r*. Similarly, we define the *death radius* of 𝓁 as the smallest real number *d* such that *r* ≤ *d* and such that 𝓁 is contractible when regarded as a loop in 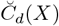. That is, the birth radius of a non-contractible loop is the smallest radius at which the equivalence class of that non-contractible loop forms, and the death radius is the smallest radius at which it vanishes (i.e., becomes contractible). For *r* ∈ [birth radius, death radius], the equivalence class of non-contractible loops ‘persists,’ and this motivates the definition of the *persistence* of an equivalence class of non-contractible loops: the persistence is the difference between the death radius and the birth radius.

The data comprising the larger square in the example dataset were included to compare two 1-dimensional homology features of different persistence within a given 2-dimensional dataset. As previously noted, the birth radius and death radius of the 1-dimensional homology feature constructed from the data comprising the smaller square are 0.5 and 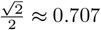, respectively, so the persistence of this feature is approximately 0.707-0.5=0.207. The birth radius of the 1-dimensional homology feature corresponding to the larger square is 2.5, and the death radius is 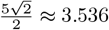. Thus its persistence is approximately 3.536-2.5=1.036. Since 1.036-0.207=0.829, the blue triangle representing the equivalence class of non-contractible loops formed from the subset of the data comprising the larger square lies a distance 0.829 further above the y=x line than the blue triangle representing the equivalence class of non-contractible loops formed from the subset of data comprising the smaller square. This illustrates how a highly persistent 1-dimensional homology feature has a larger “loop-like” structure than a 1-dimensional homology feature of lower persistence.

Also note how there are two choices of subsets of the dataset representative of the equivalence class of non-contractible loops with a smaller persistence, namely the four points that comprise the smaller square and the set of five points where four points comprise the smaller square and the other point is nearby. These choices of data points are referred to as *representative 1-cycles* and can be chosen such that the representative 1-cycle is minimal with respect to either i) the number of data points it consists of or ii) the area it spans. In the first case, we have a *cardinality-minimal* representative 1-cycle for the equivalence class; in the second case, we have a *area-minimal* representative 1-cycle for that equivalence class. In this example, both the cardinality-minimal and area-minimal representative 1-cycle of the equivalence class of non-contractible loops with a lower persistence consists of the four data points comprising the smaller square.

### Background on topological data analysis

We now set out to formalize the notion of “equivalence classes of non-contractible loops that persist for a given range of radius values.” Given a set of data X represented as a finite set of points in ℝ^2^, we construct a simplicial complex as a topological space that approximates the structure of the data.

#### Definition 0.1

*A* simplicial complex *is a collection K of subsets of a finite set V such that:*

- {*v*} ∈ *K for all v* ∈ *V, and*
- *if τ* ⊂ *σ for σ* ∈ *K, then τ* ∈ *K*.

*An element of V is referred to as a* vertex, *and an element of K with cardinality n* + 1 *is referred to as an n*-simplex.

There are several ways to construct a simplicial complex given a finite set of points in ℝ^2^, and to be consistent with the geometry of the simple dataset example, we consider the radius *r* Vietoris-Rips complex, a simplicial complex constructed by considering a circle of radius 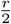 around each point in our dataset and then including *S* ⊂*X* as a simplex if the intersection of the balls of radius 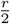 for each point in *S* is nonempty. An example of constructing the Vietoris-Rips complex for several values of *r* is shown in Fig 2.

**Fig 2.**
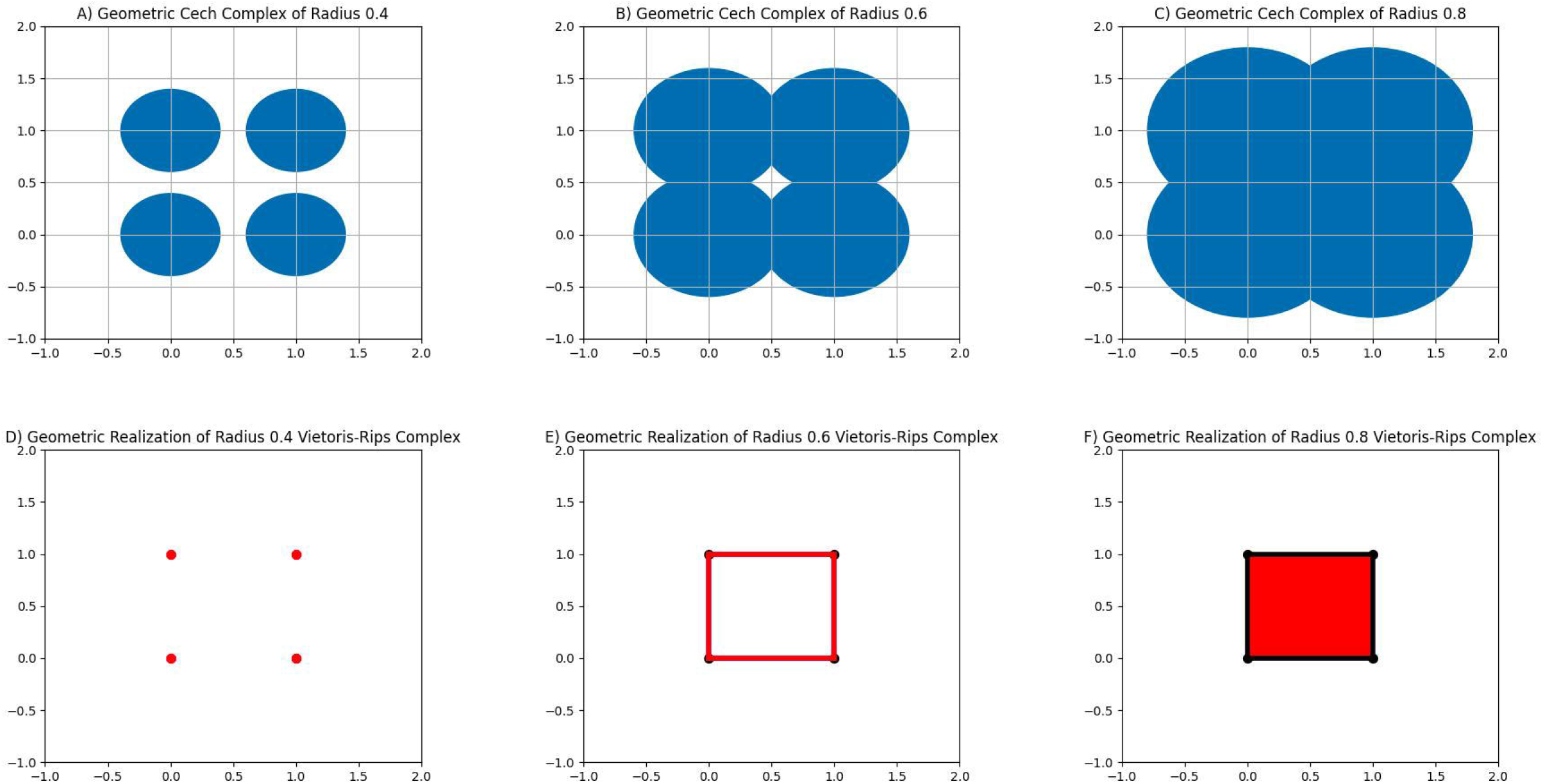
Relation between the geometric Čech complex of radius *r* and the radius *r* Vietoris-Rips Complex. A-C: Geometric Čech complex of radius 0.4, 0.6, 0.8 shown in blue. D-E: Geometric realization of radius 0.4, 0.6, 0.8 Vietoris-Rips complex; a point indicates a 0-simplex, a line indicates a 1-simplex whose members are the endpoints of the line, and a filled-in region indicates a n-simplex whose n+1 members are the data on the boundary of the region; the red regions in each plot indicates the simplicies which are born at r=0, r=0.5, and 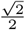, from left to right.

#### Definition 0.2

*Given a dataset X represented as a finite subset of* ℝ^2^, *and given a positive real number r, the* radius *r* Vietoris-Rips complex of *X, denoted V R*_*r*_(*X*), *is the simplicial complex given by the collection of all subsets U of X with the property that if x*_1_, *x*_2_ ∈ *U, then* |*x*_1_ − *x*_2_| *< r*.

Note that if *S*⊂ *U* for *U* ∈ *V R*_*r*_(*X*), then |*x*_1_− *x*_2_| *< r* for all *x*_1_, *x*_2_ ∈*U* implies |*x*_1_ −*x*_2_ |*< r* for all *x*_1_, *x*_2_ ∈ *S*. Thus the radius *r* Vietoris Rips complex of a finite subset of ℝ^2^ defines a simplicial complex.

We are now in a position to be more concrete about the notion of an “equivalence class of non-contractible loops” within the geometric Čech complex, as discussed in the “Example: a 1-dimensional topological feature in a simple dataset” section. By an “equivalence class of non-contractible loops,” we are referring to an element of the 1-dimensional homology group of some radius *r* Vietoris-Rips complex, which we now set out to define.

Let *X* be a finite subset of ℝ^2^, let *r* be a positive real number, and let *C*_*n*_ be the vector space over 𝔽_2_ with basis consisting of the elements of *V R*_*r*_(*X*) of cardinality *n* + 1 for *n* = 0, 1, 2. Furthermore, suppose there is an ordering on *V R*_*r*_(*X*). Consider 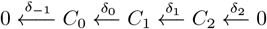 where 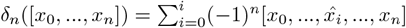 and 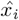 indicates that *x*_*i*_ is omitted from the ordered simplex. The elements of *C*_1_ are referred to as 1-chains, the elements of ker(*δ*_0_) are referred to as 1-cycles, and elements of im(*δ*_1_) are referred to as 1-boundaries. Since *δ*_0_(*δ*_1_(*v*)) = 0 for all *v* ∈ *C*_2_, every 1-boundary is an 1-cycle. However, it is not necessarily true that every 1-cycle is an 1-boundary. Intuitively, if we think of *X* as a cloud of points in the plane, the 1-dimensional homology group of *V R*_*r*_(*X*) is defined such that its dimension over 𝔽_2_ counts the number of “holes” in that cloud.

#### Definition 0.3

*Given r >* 0 *and V R*_*r*_(*X*) *where X is a finite subset of* ℝ^2^, *we follow the construction of* 𝔽_2_*-vector spaces C*_0_, *C*_1_, *C*_2_ *and linear transformations δ*_−1_, *δ*_0_, *δ*_1_, *δ*_2_ *as outlined above and define the first homology group of V R*_*r*_(*X*) *as the quotient vector space H*_1_(*V R*_*r*_(*X*)) = *ker*(*δ*_0_)*/im*(*δ*_1_). *The* 𝔽_2_*-vector space dimension β*_1_ = *dim*(*H*_1_(*V R*_*r*_(*X*))) = *dim*(*ker*(*δ*_0_)) − *dim*(*im*(*δ*_1_)) *of H*_1_(*V R*_*r*_(*X*)) *is called the* first Betti number.

By increasing *r*, we create a sequence of Vietoris-Rips Complexes where *V R*_*r*_(*X*) ⊂ *V R*_*r*′_ (*X*) for *r < r*^′^. We then construct

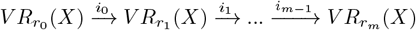

where 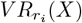 is a proper subset of 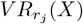 for *i < j* and *i*_0_, *i*_1_,…,*i*_*m*−1_ are inclusion homomorphisms. This induces a sequence of 𝔽_2_-linear functions 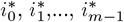 such that

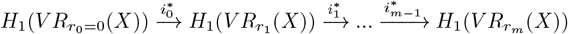

and 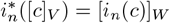 for 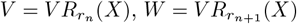, and all *n* = 0, 1, …, *m* −1. We now give a name to the “smallest” and “largest” *r >* 0 such that a given 1-cycle belongs to *H*_1_(*V R*_*r*_(*X*)).

#### Definition 0.4

*Let* [*c*] ∈ *H*_1_(*V R*_*r*_(*X*)) *for some r >* 0. *The birth filtration of* [*c*] *is defined as the greatest lower bound of the set of all ϵ >* 0 *such that* [*c*] *is in the range of the* 𝔽_2_*-linear function H*_1_(*V R*_*ϵ*_(*X*)) → *H*_1_(*V R*_*r*_(*X*)). *Similarly, the death filtration of* [*c*] *is defined as the least upper bound of the set of all ϵ >* 0 *such that* [*c*] *maps to zero under the* 𝔽_2_*-linear function H*_1_(*V R*_*r*_(*X*)) → *H*_1_(*V R*_*ϵ*_(*X*)). *The persistence of* [*c*] *is defined as the difference between the death filtration and the birth filtration*.

Up to a scaling factor in the variable *r*, the geometric Čech complex of radius *r* is homotopy equivalent to the radius *r* Vietoris-Rips complex due to the Nerve Lemma [29]. Consequently, the definitions of the birth and death radii of an equivalence class of non-contractible loops presented in the “Example: a 1-dimensional topological feature in a simple dataset” section are equivalent to the definitions of the birth and death filtration of a class [*c*] ∈ *H*_1_(*V R*_*r*_(*X*)) given in Definition 0.4. For a more thorough treatment of persistent homology, see [30].

## Methods

In this section, we first describe how ECG data are processed to be in a form such that our topological approach can be applied consistently. We then describe our topological method of identifying P,Q,S, and T-waves and measuring the PR-interval, QT-interval, ST-segment, QRS-duration, P-wave duration, and T-wave duration. Lastly, we present the methods used to simulate ECG signals with the PR-interval, QT-interval, ST-segment, QRS-duration, P-wave duration, and T-wave duration determined by parameters of the simulation.

### Data processing

Before an ECG signal can be analyzed using our topological approach, the signal is processed such that the topological approach can be consistently applied. An outline of this processing pipeline is depicted in Fig 3. This processing pipeline is the same regardless of whether the ECG signal is simulated or real.

**Fig 3.**
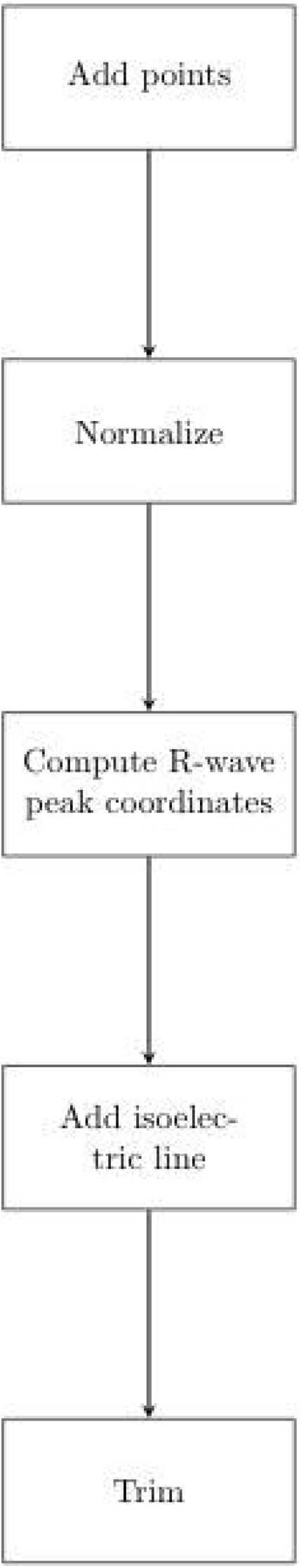
ECG Signal Processing Pipeline. The above flowchart depicts a processing pipeline which is applied to an ECG signal prior to computing topological features of the signal.

Given an ECG signal (*D, f*) where *D* denotes the set of time indices and *f* : *D* →ℝ is such that *f* (*t*) represents the amplitude of the signal at time *t*, we first nearly double the number of points comprising the signal by including the point 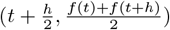 for every pair of adjacent points (*t, f* (*t*)) and (*t* + *h, f* (*t* + *h*)) of the signal. The signal is then normalized by mapping every ordered pair (*t, f* (*t*)) to 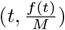 where 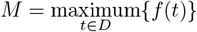. The coordinates of the R-wave peaks are used in identifying P,Q,S, and T-waves, and the coordinate pair of the i-th R-wave peak is computed as the ordered pair (*t*_*R,i*_, *f* (*t*_*R,i*_)) where 0.6 *< f* (*t*_*R,i*_), *f* (*t*_*R,i*_) is a local maximum of the signal, and there are exactly *i* − 1 local maxima *f* (*t*_*R,j*_) such that *t*_*R,j*_ *< t*_*R,i*_ and 0.6 *< f* (*t*_*R,j*_), *j* = 1, 2, …, *i* − 1 [31]. Next, an isoelectric line is incorporated into the signal by mapping 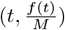 to (*t, baseline*) where *baseline* is the median value of the set {*f* ^*^(*t*) |*t* ∈ *D*} and *f* ^*^(*t*) is *f* (*t*) rounded to the nearest hundredth} for all *t* ∈ *D* such that *t* = *h*(2*k*) for some integer *k*. Lastly, we consider the restriction of the processed ECG signal such that the signal begins and ends with an R-wave peak as the input of our topological method.

### Extracting features of processed ECG signals using TDA

Given a processed ECG signal (*D, f*), we are interested in i) identifying subsets of the signal as P,Q,S, and T-waves using cycle reconstructions of 1-dimensional homology classes and ii) using the identified waves to measure the PR-interval, QT-interval, ST-segment, QRS-duration, P-wave duration, and T-wave duration. To describe how P,Q,S, and T-waves are identified as 1-cycles with certain properties, if 1-cycles with these properties exist, we first give a general description of five properties of ECG signals that are used in this process:

- **Persistence:** Due to the the addition of the isoelectric line, specifically under the P,T-waves and over the Q,S-waves, there is a loop-like structure of data comprising the P,Q,S, and T-waves as shown in Fig 4A. Thus for a 1-cycle to be identified as a P,Q,S, or T-wave, we impose that it must have a persistence within some bounded interval depending on the specific wave.
- **Birth filtration upper bound:** It is a property of ECG signals that adjacent points (*t, f* (*t*)), (*t* + *h, f* (*t* + *h*)) within each P,Q,S, and T-wave are within some distance *δ* from one another where *δ* is small relative to the distance between (*t, f* (*t*)) and a point (*t*^′^, *f* (*t*^′^)) on a different wave. Thus we impose an upper bound on the birth filtration of each potential 1-cycle to exclude 1-cycles constructed from points corresponding to multiple waves, e.g. a 1-cycle constructed from both a P-wave and an R-wave as shown in Fig 4B.
- **Time coordinate of centroid of 1-cycle:** It is a property of ECG signals that a T-wave follows a QRS-complex. We also expect a detected P-wave to necessarily precede a QRS-complex due to our simulations not featuring atrial activity with missed ventricular beats. Thus we expect that the centroids of 1-cycles comprising a T-wave and P-wave lie between a pair of adjacent R-waves such that the T-wave is some distance after the preceding R-wave, the P-wave is some distance prior to the following R-wave, and the P-wave follows the T-wave. Similarly, we impose that a 1-cycle comprising a detected Q-wave and S-wave slightly precedes an R-wave and shortly follows an R-wave, respectively.
- **Amplitude coordinate of centroid of 1-cycle:** It is expected that over a given heartbeat, the average amplitude of the signal in the P,T-wave be larger than the signal amplitude at baseline. Additionally, we expect there to be a positive upper bound on the average amplitude of the signal in the P,T-wave. Conversely, it is expected that the average amplitude of the signal in the Q,S-wave be less than the signal amplitude at baseline.
- **Optimality:** Of the 1-cycles satisfying the persistence, birth filtration, and centroid coordinate conditions above, the 1-cycle representing the given P,Q,S, or T-wave is expected to be optimal in the sense of i) having a minimal number of data points comprising the 1-cycle and/or ii) having a minimal-area convex hull. These optimal cycles were computed using HomCloud [32].

**Fig 4.**
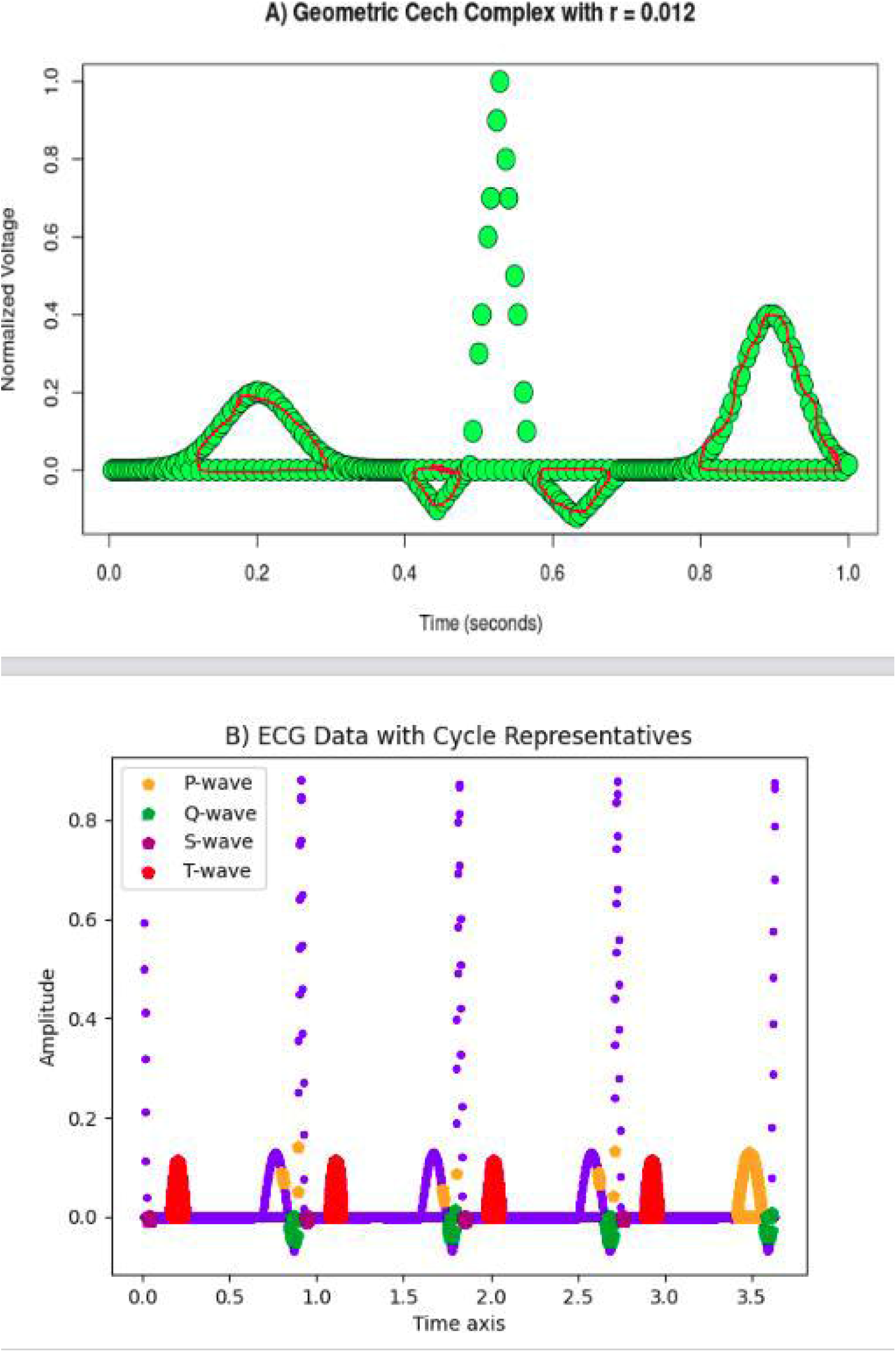
Illustration of properties of an ECG used to detect P,Q,S, and T-waves. A: The red loops illustrate elements from equivalence classes of non-contractible loops corresponding to the P,Q,S, and T-wave. Notice that without the addition of the isoelectric baseline, we would be unable to draw non-contractible loops corresponding to the P,Q,S, and T-waves within the geometric Čech complex. B: Example of output without imposing an upper bound on the birth filtration of a 1-cycle identified as a P-wave.

We now present the specific criteria that identifies 1-cycles with certain properties related to those described above as being a P-wave or T-wave. We note that differences in the classification criteria for P-waves and T-waves involves changes in the bounds of the persistence range and changes to the upper bound of the birth filtration.

Additionally, the centroid of a 1-cycle identified as a P-wave is expected to be closer to the subsequent R-wave than the preceding R-wave, and vice versa for the centroid of a 1-cycle identified as a T-wave.

### P-wave Identification

An optimal 1-cycle satisfying the following conditions is identified as a P-wave (*U*_*P*_, *f*) with collection of ordered pairs { (*t, f* (*t*))| *t* ∈ *U*_*P*_} and time index set *U*_*P*_ :

- the persistence of the 1-cycle is in [0.001,0.2]
- the birth filtration of the 1-cycle is less than 0.03
- 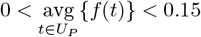
- R_peak_time_coordinate[i] − 0.35 * RR_interval 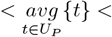 R_peak_time_coordinate[i] − 0.06 * RR_interval for some i.

### T-wave Identification

An optimal 1-cycle satisfying the following conditions is identified as a T-wave (*U*_*T*_, *f*) with collection of ordered pairs{ (*t, f* (*t*)) |*t* ∈ *U*_*T*_} and time index set *U*_*T*_ :

- the persistence of the 1-cycle is in [0.01,0.6]
- the birth filtration of the 1-cycle is less than 0.04
- 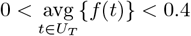
- R_peak_time_coordinate[i] + 0.15 * RR_interval 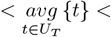 R_peak_time_coordinate[i] + 0.5 * RR_interval for some i.

We continue with describing the criteria for a 1-cycle to be identified as a Q-wave or S-wave. The interval which the persistence of a 1-cycle must belong to for the 1-cycle to be classified as a Q-wave or S-wave includes a smaller lower bound and smaller upper bound compared to the interval of valid persistence values for a 1-cycle to be classified as a P,T-wave. This is due to the data comprising Q-waves and S-waves having less of a loop-like structure than data comprising the P-waves and T-waves. We also impose that the centroid of a 1-cycle identified as a Q-wave or S-wave more closely precede or follow the nearby R-wave peak relative to a P-wave or T-wave, respectively.

### Q-wave Identification

An optimal 1-cycle satisfying the following conditions is identified as a Q-wave (*U*_*Q*_, *f*) with collection of ordered pairs {(*t, f* (*t*))| *t* ∈*U*_*Q*_} and time index set *U*_*Q*_:

- the persistence of the 1-cycle is in [0.007,0.1]
- the birth filtration of the 1-cycle is less than 0.06
- 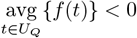
- R_peak_time_coordinate[i] −0.12 * RR_interval 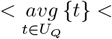 R_peak_time_coordinate[i] for some i.

### S-wave Identification

An optimal 1-cycle satisfying the following conditions is identified as a S-wave (*U*_*S*_, *f*) with collection of ordered pairs {(*t, f* (*t*)) |*t* ∈*U*_*S*_} and time index set *U*_*S*_:

- the persistence of the 1-cycle is in [0.007,0.1]
- the birth filtration of the 1-cycle is less than 0.06
- 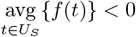
- R_peak_time_coordinate[i] 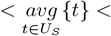 R_peak_time_coordinate[i] + 0.12 * RR_interval for some i.

In the above criteria, “optimal 1-cycle” refers to a 1-cycle computed as being minimal with respect to i) the cardinality of the set of data points comprising the 1-cycle or ii) the area spanned by the convex hull of the 1-cycle. Both types of optimal 1-cycles were computed in our analysis and are compared. An example of area-minimal 1-cycles identified as P,Q,S, and T-waves using the above criteria depicted on an ECG signal is shown in Fig 5. Cardinality-minimal and area-minimal 1-cycles identified as P,Q,S, and T-waves for the 400 simulated ECG signals is provided as Supplementary File 1 and Supplementary File 2. Additionally, cardinality-minimal and area-minimal 1-cycles identified as P,Q,S, and T-waves for 200 Lead II ECG signals randomly sampled from a pool of 10646 denoised signals with 11 common rhythms as described in [33] are shown in Supplementary File 3 and Supplementary File 4, respectively.

**Fig 5.**
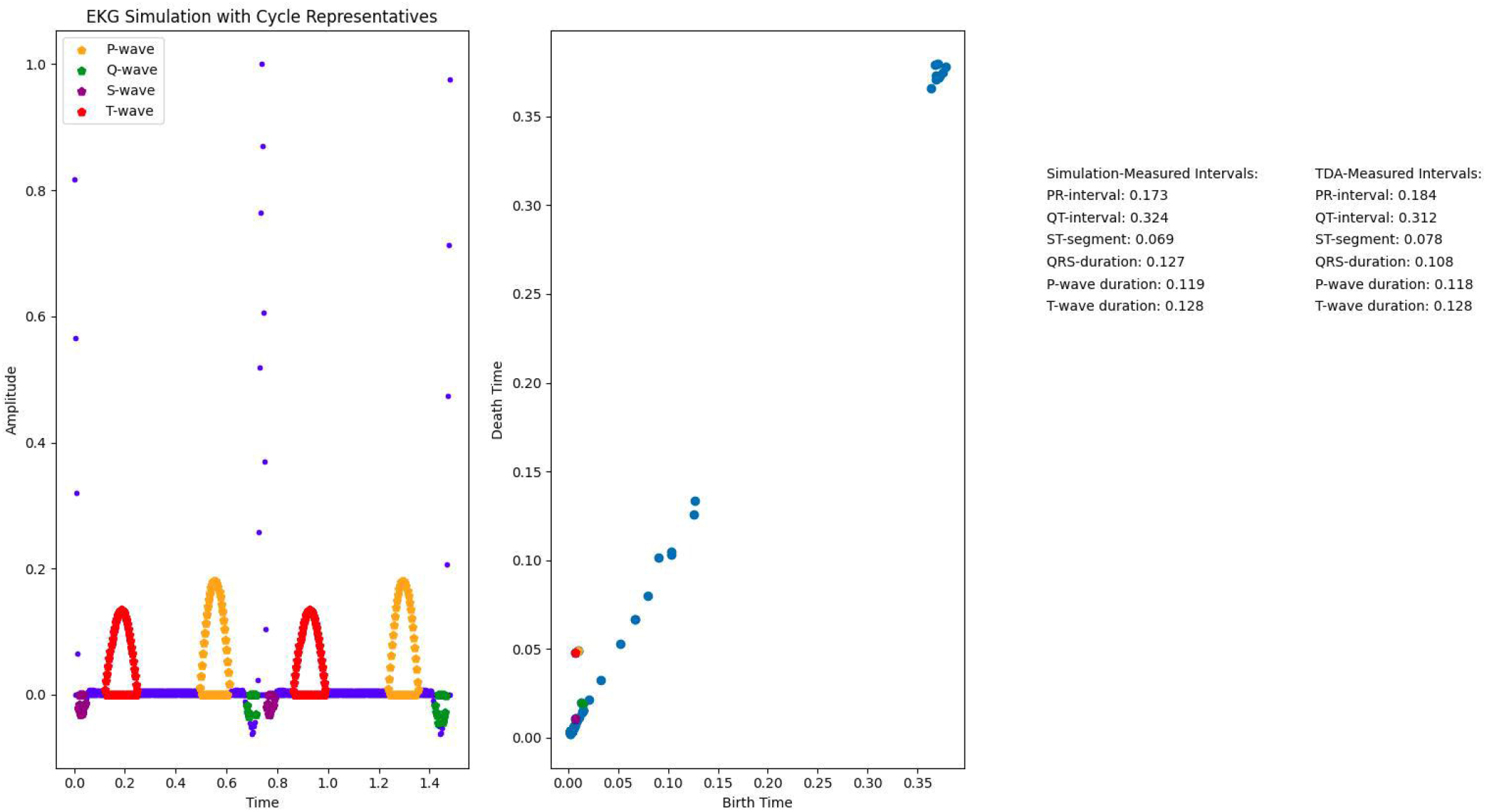
Example of simulated ECG signal and persistence diagram with area-optimal 1-cycles identified as P,Q,S, and T-waves. The points comprising P,Q,S, and T-waves were identified as area-optimal 1-cycles with certain properties depending on their birth filtration, persistence, and centroid. The Q-wave and S-wave 1-cycles drawn have the left-most and right-most centroid of all Q-wave and S-waves detected within a given heartbeat, respectively. This is consistent with the choice of Q-waves and S-waves used to measure intervals of interest. The color of 1-cycles identified as P,Q,S, and T-waves matches the color of the corresponding points in the persistence diagram.

We now describe how the PR-interval, QT-interval, ST-segment, QRS-duration, P-wave duration, and T-wave duration are measured using 1-cycles identified as P,Q,S, and T-waves. To measure these intervals solely using topological features of the data, the PR-interval and QR-interval of a given heartbeat are only measured if a Q-wave is detected. Similarly, the ST-segment of a given heartbeat is only measured if an S-wave is detected. The QRS-duration is only measured if both a Q-wave and an S-wave are detected. The intervals of interest are computed as the difference between their offset and onset. Given a heartbeat of a processed ECG signal with at least one detected Q,S-wave and one detected P,T-wave, we measure the onset and offset of the intervals of interest as follows:

- The onset of the PR-interval and onset of the P-wave duration is minimum{*U*_*P*_}.
- The offset of the PR-interval, onset of the QT-interval, and onset of the QRS-duration is given by minimum 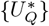 where 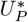 is the set of time indices of the 1-cycle satisfying the Q-wave criteria such that the time coordinate of its centroid is less than the time coordinate of the centroid of every other 1-cycle identified as a Q-wave within the heartbeat.
- The onset of the ST-segment and offset of the QRS-duration is given by minimum 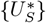 where 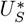 is the set of time indices of the 1-cycle satisfying the S-wave criteria such that the time coordinate of its centroid is greater than the time coordinate of the centroid of every other 1-cycle identified as an S-wave within the heartbeat.
- The offset of the QT-interval and offset of the T-wave duration is minimum{*U*_*T*_ }.
- The offset of the ST-segment and onset of the T-wave duration is minimum{*U*_*T*_ }.
- The offset of the P-wave duration is maximum{*U*_*P*_ }.

The average PR-interval, QT-interval, ST-segment, QRS-duration, P-wave duration, and T-wave duration were calculated using i) the model’s parameters, ii) cardinality-minimal 1-cycle reconstructions, and iii) area-minimal 1-cycle reconstructions for all 1000 simulated ECG signals. The percent difference between the model’s measurement of each interval and the optimal 1-cycle’s measurement of each interval for both cardinality-minimal and area-minimal 1-cycles were calculated for each simulation.

### ECG simulation

ECG signals were generated with a sampling frequency of 500Hz using an ECG Simulator in MATLAB [34]. The set of parameters that completely characterizes what we refer to as a simulated ECG signal includes the heart rate, the amplitude and duration of the R-wave, and the amplitude, duration, and location with respect to i) the subsequent R-wave for the P,Q-wave and ii) the preceding R-wave for the S,T-wave. We now go into detail about how this parameter set is used to construct a simulated ECG signal with a given PR-interval, QT-interval, ST-segment, QRS-duration, P-wave duration, and T-wave duration.

Let 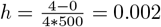 be the time-step and *D* = {*x*∈ [0, 4] |*x* = *hk* for *k* ∈ℤ} be the set of time indices of a simulated ECG signal. A simulated ECG signal (*D, f*) is given by a set of ordered pairs (*t, f* (*t*)) where *t ∈ D* and *f* : [0, 4] → ℝ is such that *f* (*t*) represents the amplitude of the signal at time *t*. Furthermore, the simulated ECG signal is periodic, i.e. *f* (*t*) = *f* (*t* + *rr*_*interval*) where 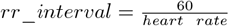. Thus to completely characterize a simulated ECG signal, we need only know how *f* is defined on a contiguous subset *E* of *D* which ranges over one period of *f*. We also impose the property of a simulated ECG signal such that any choice of *E* contains five subsets of the simulated ECG signal which we refer to as the P,Q,R,S, and T-waves. Depending on the choice of *E* and the parameters of the model, the P,Q,R,S, and T-waves are either connected by an isoelectric segment on which *f* is constant or intersect one another.

We now briefly describe the P,Q,R,S, and T-waves. The QRS complex consists of six line segments, and the shape of the P and T-wave is dictated by a Fourier series [34]. The amplitude and duration of the X-wave is given by the parameters amplitude_X_wave and duration_X_wave, respectively, for X=P,Q,R,S,T. The distance along the time-axis from the centroid of the ordered pairs comprising the P,Q-wave to the coordinate pair of the following R-wave peak is given by the parameter location_P,Q_wave. Similarly, the distance along the time-axis from the coordinate pair of an R-wave peak to the centroid of the ordered pairs (*t, f* (*t*)) comprising the subsequent S,T-wave is given by the parameter location_S,T_wave.

We now describe how the intervals of interest, i.e. the PR-interval, QT-interval, ST-segment, QRS-duration, P-wave duration, and T-wave duration, are constructed from the model parameters. We consider the location on the time axis corresponding to the R-wave peak as the origin and measure distance as being positive in both directions along the time axis. Thus, both the onset and offset of the R-wave is given by 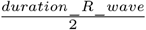. The onset of the P,Q-wave and the offset of the S,T-wave is taken to be 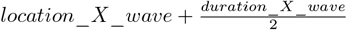 for X=P,Q,S,T. Similarly, the offset of the P,Q-wave and onset of the S,T-wave is given as 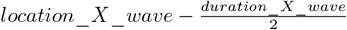 for X=P,Q,S,T. We now define the simulation’s measure of these intervals as

- PR-interval = onset of P-wave - onset of Q-wave
- QT-interval = onset of Q-wave + offset of T-wave
- ST-segment = onset of T-wave - offset of S-wave
- QRS-duration = onset of Q-wave + offset of S-wave
- P-wave duration = onset of P-wave - offset of P-wave
- T-wave duration = offset of T-wave - onset of T-wave

where if the amplitude of the Q-wave or S-wave is below 0.02, then the onset of the Q-wave or offset of the S-wave is given by the onset or offset of the R-wave, respectively.

By uniformly sampling the heart rate from [60,100] and the parameter values of the P,Q,R,S, and T-waves from the ranges in Table 1, 1000 simulated ECGs were generated.

**Table 1.**
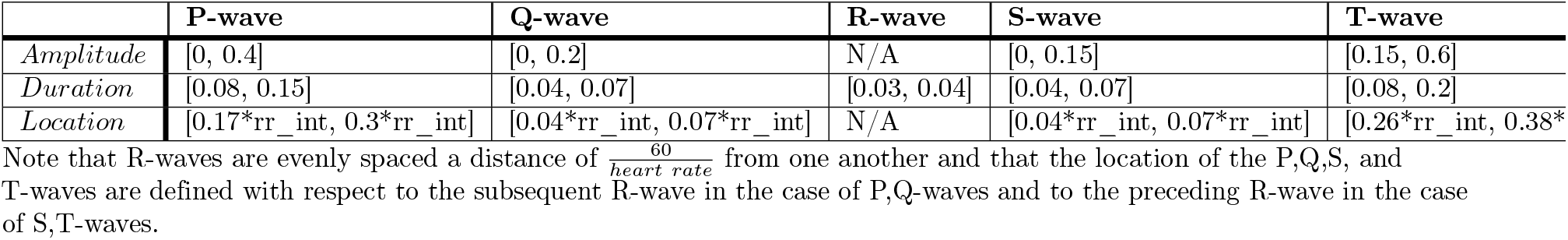
Parameter ranges of simulated ECG signals.

## Results/discussion

The PR-interval, QT-interval, ST-segment, QRS-duration, P-wave duration, and T-wave duration of 1000 simulated ECG signals were measured using i) cardinality-minimal 1-cycle reconstructions and ii) area-minimal 1-cycle reconstructions and compared with the interval measurements made by model parameters. For each simulation, the percent difference between the model’s measurement of each interval and the measurement of each interval performed using optimal 1-cycle reconstructions was calculated. The distribution of these measurements using cardinality-minimal and area-minimal 1-cycles are shown in Fig 6 and Fig 7, respectively. The average percent difference and the standard deviation of the percent difference measurements for each interval of interest across the 1000 simulated ECG signals is shown in Table 2.

**Table 2.**
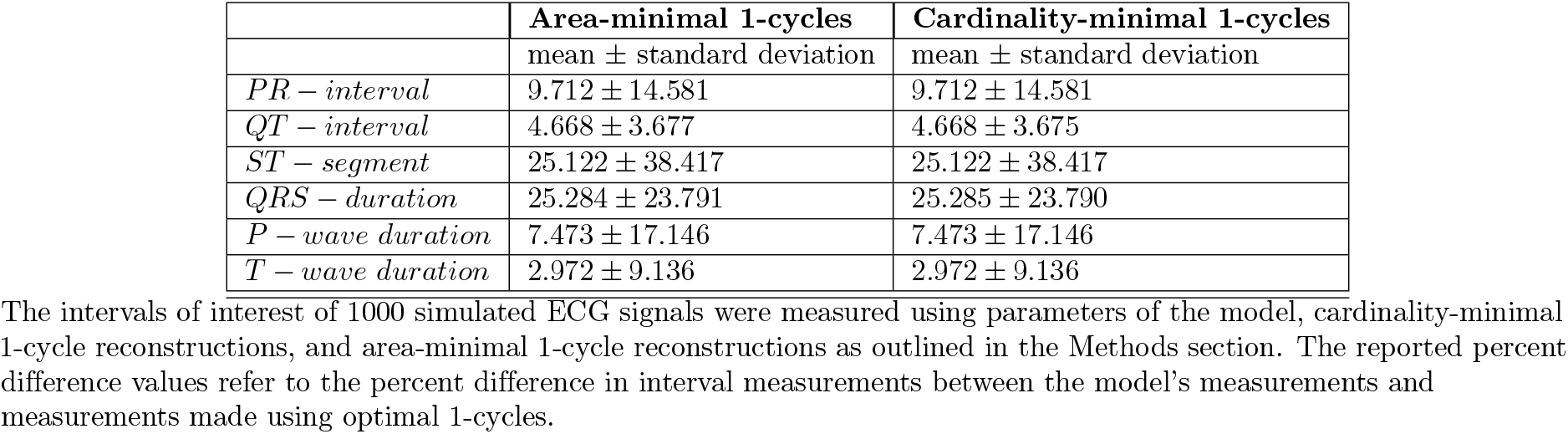
Average percent difference of interval measurements.

**Fig 6.**
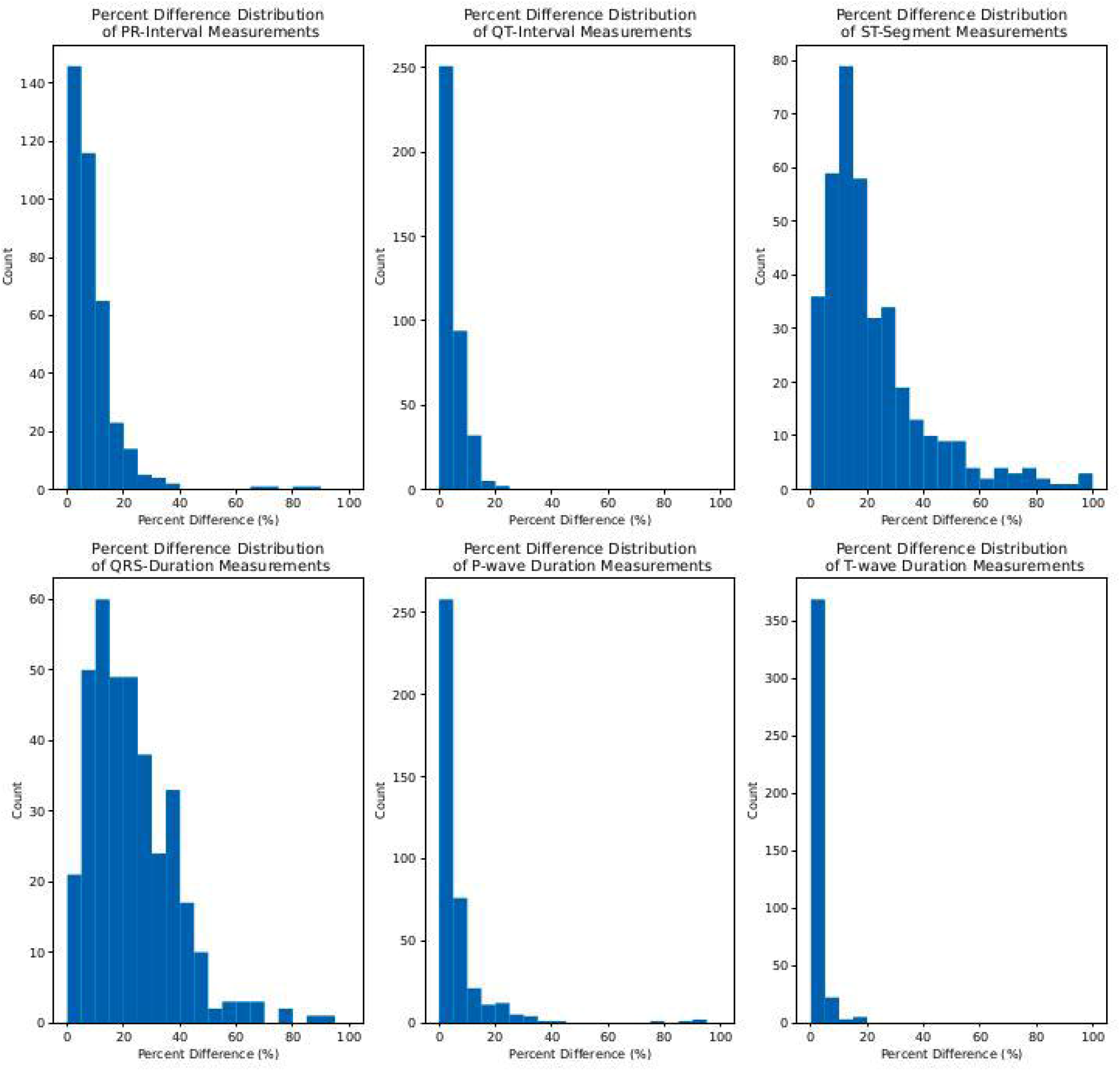
Distribution of percent difference measurements using cardinality-optimal 1-cycles. Each histogram depicts the distribution of percent difference measurements for a single interval of interest. Left to right across the top row: PR-interval, QT-interval, ST-segment. Left to right across the bottom row: QRS-duration, P-wave duration, T-wave duration.

**Fig 7.**
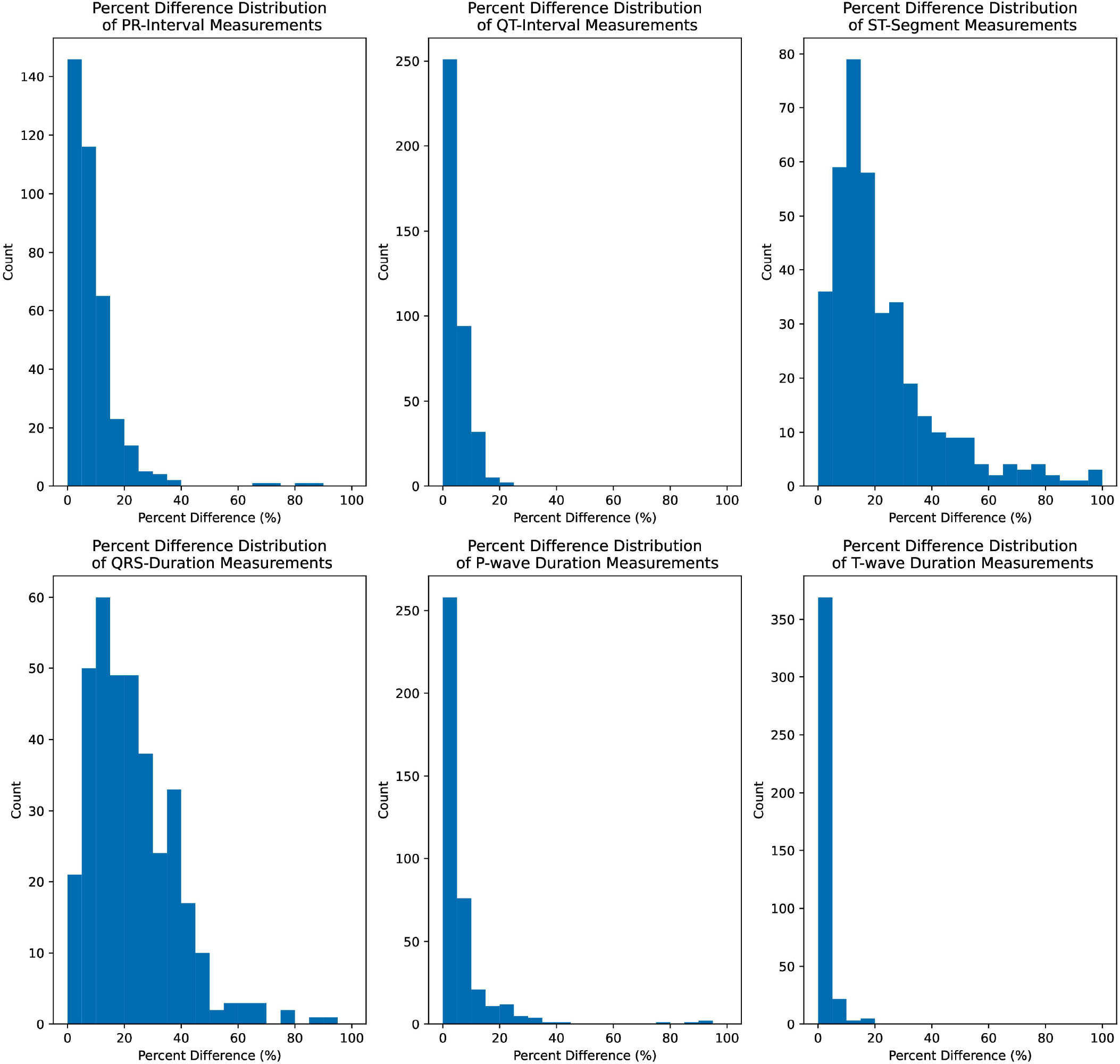
Distribution of percent difference measurements using area-optimal 1-cycles. Each histogram depicts the distribution of percent difference measurements for a single interval of interest. Left to right across the top row: PR-interval, QT-interval, ST-segment. Left to right across the bottom row: QRS-duration, P-wave duration, T-wave duration.

From comparing cardinality-minimal 1-cycles identified as P,Q,S, and T-waves to area-minimal 1-cycles identified as P,Q,S, and T-waves in Supplementary File 1 and Supplementary File 2, we see that these cycles are often “close” to one another, likely resulting in the similar percent error distributions seen in Fig 6, Fig 7, and Table 2. It is also evident from Supplementary File 1 and Supplementary File 2 that the algorithm could be improved by using a more stable method to include an isoelectric baseline in the signal. Additionally, to ensure that P,Q,S, and T-wave identifications and interval measurements are not compromised due to misidentified or undetected R-wave peaks, a more robust R-wave peak detection algorithm could be used such as those in [35], [36], [37].

The parameters in the criteria used to identify 1-cycles as P,Q,S, and T-waves are currently determined from several iterations of identifying 1-cycles as P,Q,S, and T-waves and adjusting the parameter values to obtain more accurate identification based on visual inspection. A more robust method of optimizing this parameter set could result in more accurate identification of 1-cycles which better represent the shape and/or endpoints of the P,Q,S, and T-waves. A difficulty in this is that it may be difficult to know what a “correct” subset of a signal is in order for it to be considered a P,Q,S, or T-wave. Additionally, information about optimal 1-cycles identified as P,Q,S, and T-waves could be combined with other existing approaches such as analyzing other persistent homology statistics, wavelet decompositions, and machine learning for automated arrhythmia detection in future work [12] [13] [17] [19] [21].

## Supporting information

The code used to produce simulated ECG signals is available at https://www.mathworks.com/matlabcentral/fileexchange/10858-ecg-simulation-using-matlab/ [34]. The real ECG data used in this study is publicly available and can be freely downloaded at https://www.nature.com/articles/s41597-020-0386-x/ [33]. The code used to identify P,Q,S, and T-waves and measure intervals of interest based off of topological features of the ECG signal can be found in the GitHub repository at https://github.com/hdlugas/ekg_tda.

**Supplementary File 1 Cardinality-minimal 1-cycle reconstructions identified as P**,**Q**,**S**,**T-waves for 400 simulated ECG signals**

**Supplementary File 2 Area-minimal 1-cycle reconstructions identified as P**,**Q**,**S**,**T-waves for 400 simulated ECG signals**

**Supplementary File 3 Cardinality-minimal 1-cycle reconstructions identified as P**,**Q**,**S**,**T-waves for 200 ECG signals randomly sampled from the pooled described in ‘Methods-Extracting Features of Processed ECG Signals using TDA’**

**Supplementary File 4 Area-minimal 1-cycle reconstructions identified as P**,**Q**,**S**,**T-waves for 200 ECG signals randomly sampled from the pooled described in ‘Methods-Extracting Features of Processed ECG Signals using TDA’**

**Supplementary File 5 File names of the randomly sampled ECG signals analyzed using cardinality-minimal 1-cycles**.

**Supplementary File 6 File names of the randomly sampled ECG signals analyzed using cardinality-minimal 1-cycles**.

## Acknowledgments

The author would like to thank Andrew Salch for discussions on interpreting topological features of data and for guidance towards submitting this paper. The author would also like to thank Hassan Abdallah for discussions on i) persistent homology software and ii) a statistical analysis comparing interval measurements performed using cardinality-minimal and area-minimal 1-cycle reconstructions.

