## Supplementary File 2 for "Electrocardiogram feature extraction and interval measurements using optimal representative cycles from persistent homology"

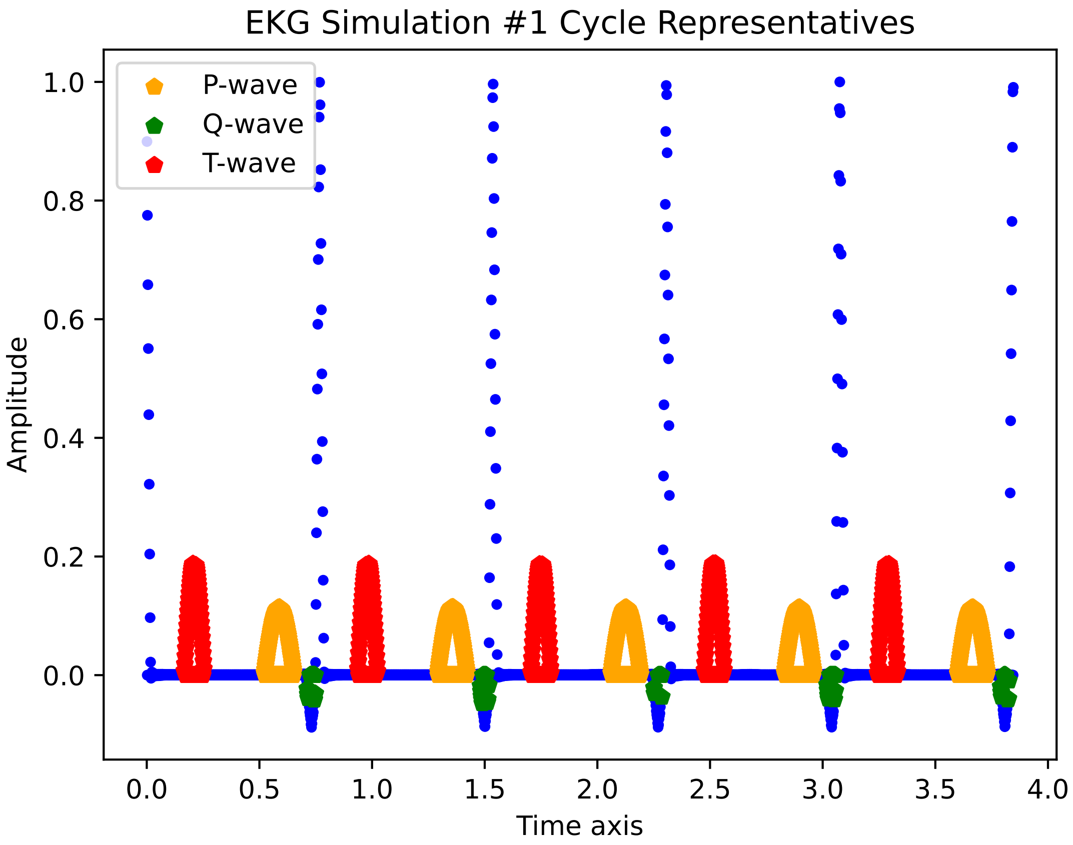

EKG Simulation #1 Cycle Representatives
P-wave
Q-wave
T-wave
Amplitude
1.0
0.8
0.6
0.4
0.2
0.0
0.0
0.5
1.0
1.5
2.0
2.5
3.0
3.5
4.0
Time axis

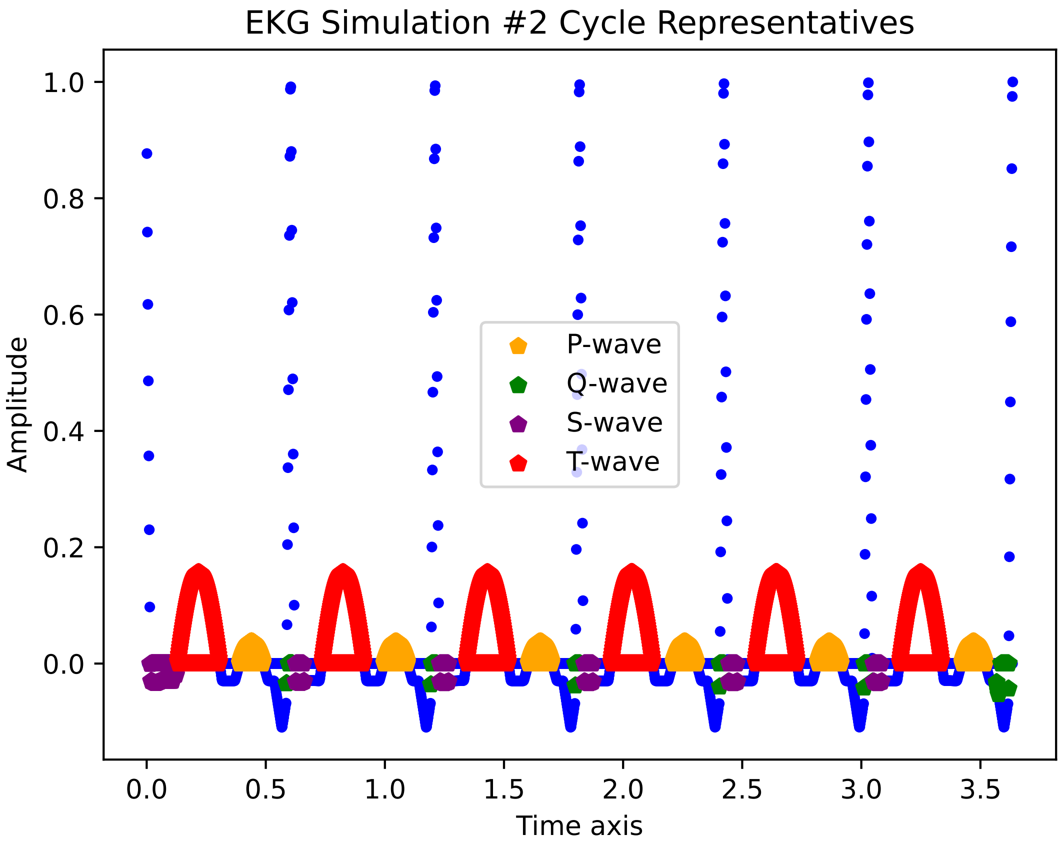
EKG Simulation #2 Cycle Representatives
P-wave
Q-wave
S-wave
T-wave
Amplitude
Time axis
1.0
0.8
0.6
0.4
0.2
0.0
0.0
0.5
1.0
1.5
2.0
2.5
3.0
3.5

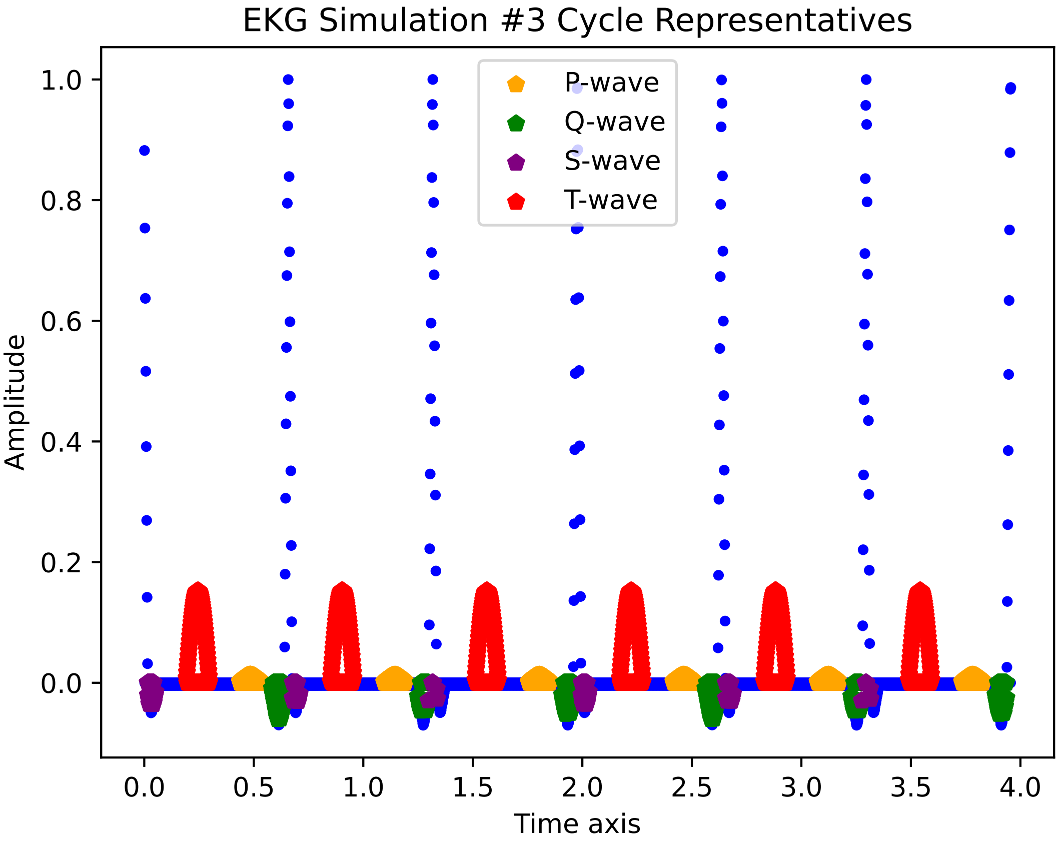
EKG Simulation #3 Cycle Representatives
P-wave
Q-wave
S-wave
T-wave
Amplitude
Time axis
1.0
0.8
0.6
0.4
0.2
0.0
0.0
0.5
1.0
1.5
2.0
2.5
3.0
3.5
4.0

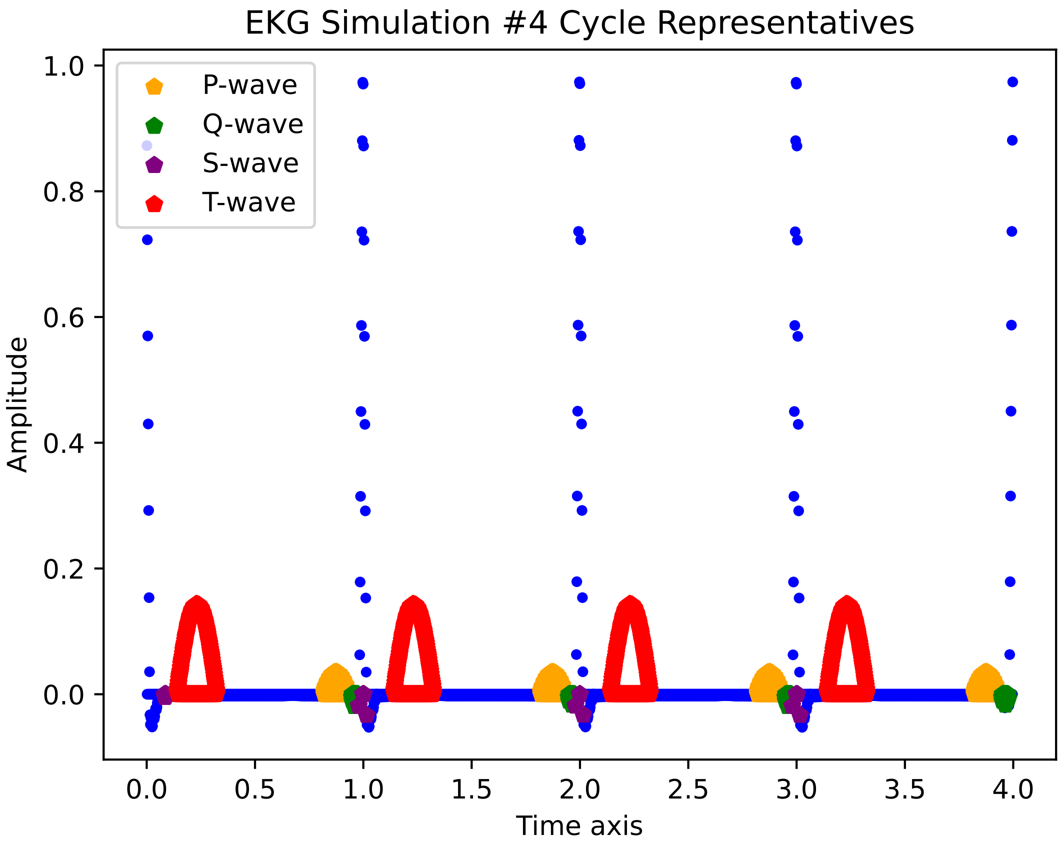

EKG Simulation #4 Cycle Representatives
P-wave
Q-wave
S-wave
T-wave
Amplitude
Time axis
1.0
0.8
0.6
0.4
0.2
0.0
0.0
0.5
1.0
1.5
2.0
2.5
3.0
3.5
4.0

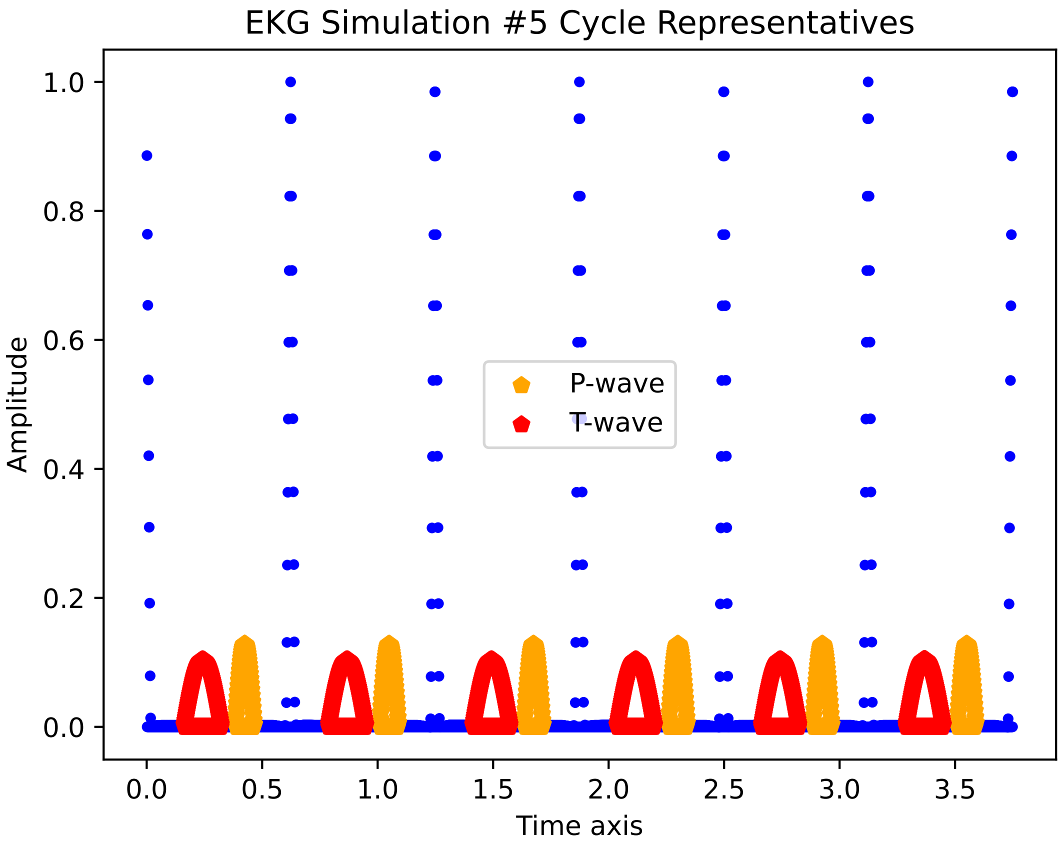

EKG Simulation #5 Cycle Representatives
Amplitude
Time axis
P-wave
T-wave
1.0
0.8
0.6
0.4
0.2
0.0
0.0
0.5
1.0
1.5
2.0
2.5
3.0
3.5

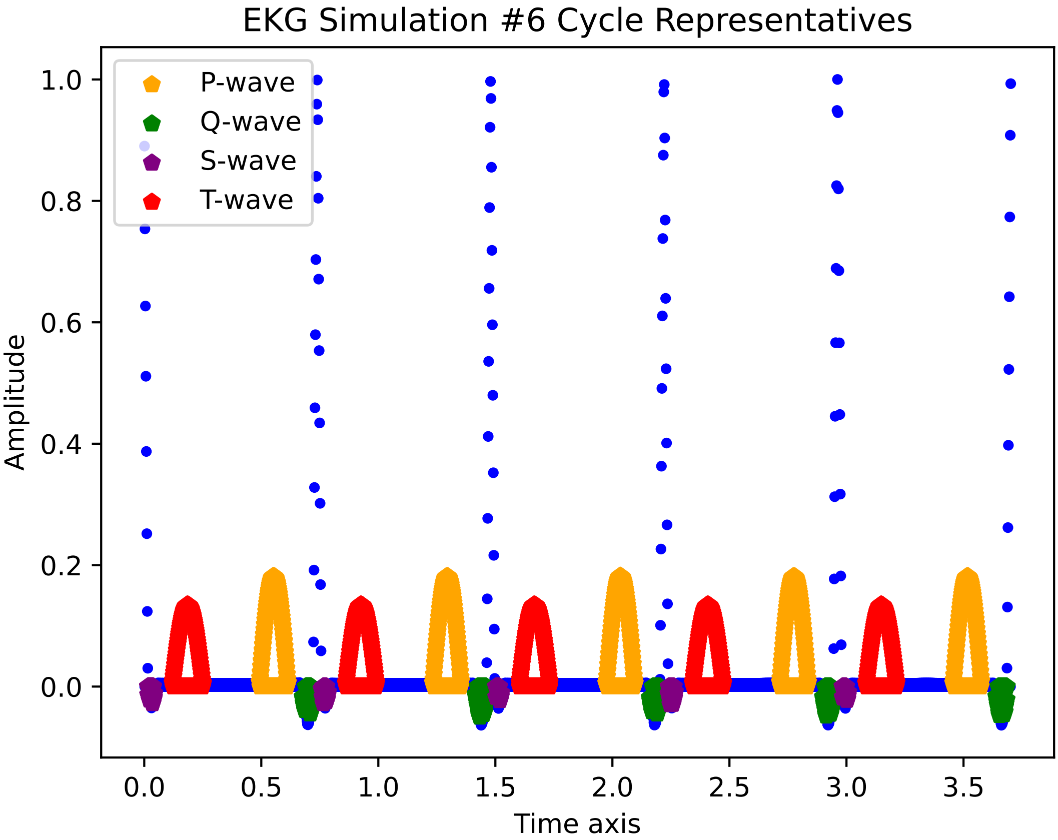

EKG Simulation #6 Cycle Representatives
P-wave
Q-wave
S-wave
T-wave
Amplitude
1.0
0.8
0.6
0.4
0.2
0.0
0.0
0.5
1.0
1.5
2.0
2.5
3.0
3.5
Time axis

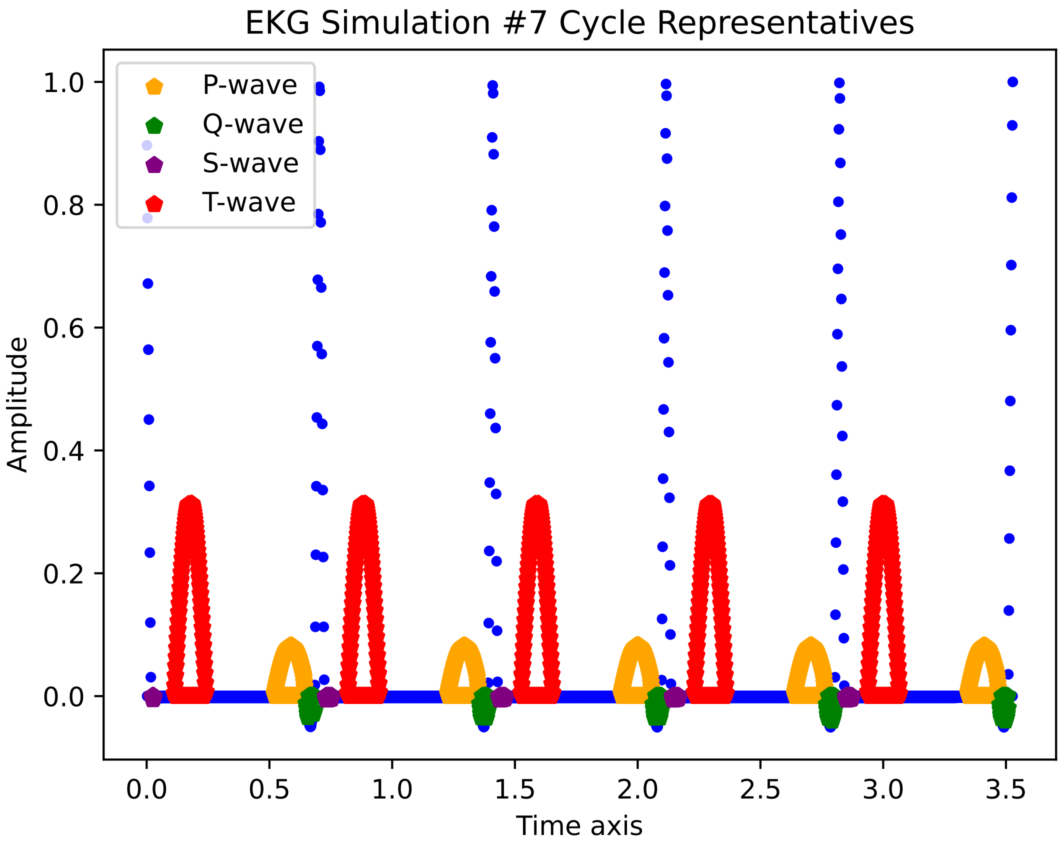

EKG Simulation #7 Cycle Representatives
P-wave
Q-wave
S-wave
T-wave
Amplitude
Time axis
1.0
0.8
0.6
0.4
0.2
0.0
0.0
0.5
1.0
1.5
2.0
2.5
3.0
3.5

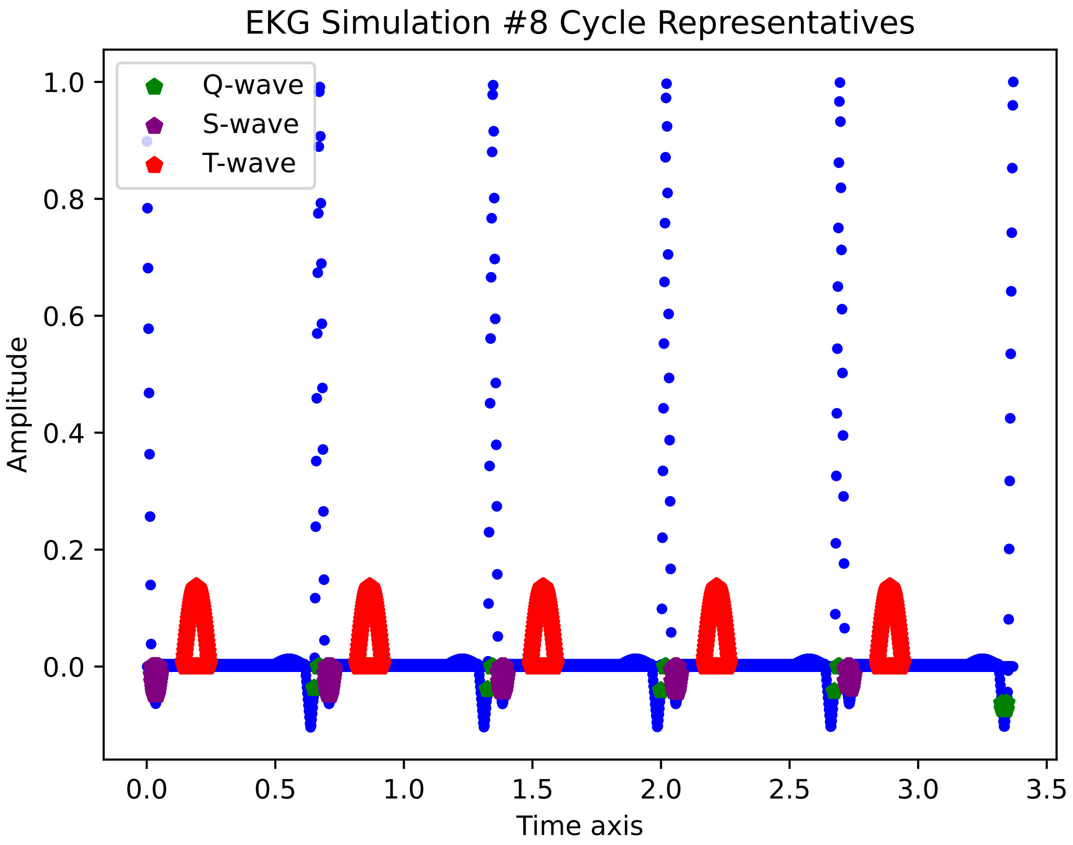
EKG Simulation #8 Cycle Representatives
Q-wave
S-wave
T-wave
Amplitude
1.0
0.8
0.6
0.4
0.2
0.0
0.0
0.5
1.0
1.5
2.0
2.5
3.0
3.5
Time axis

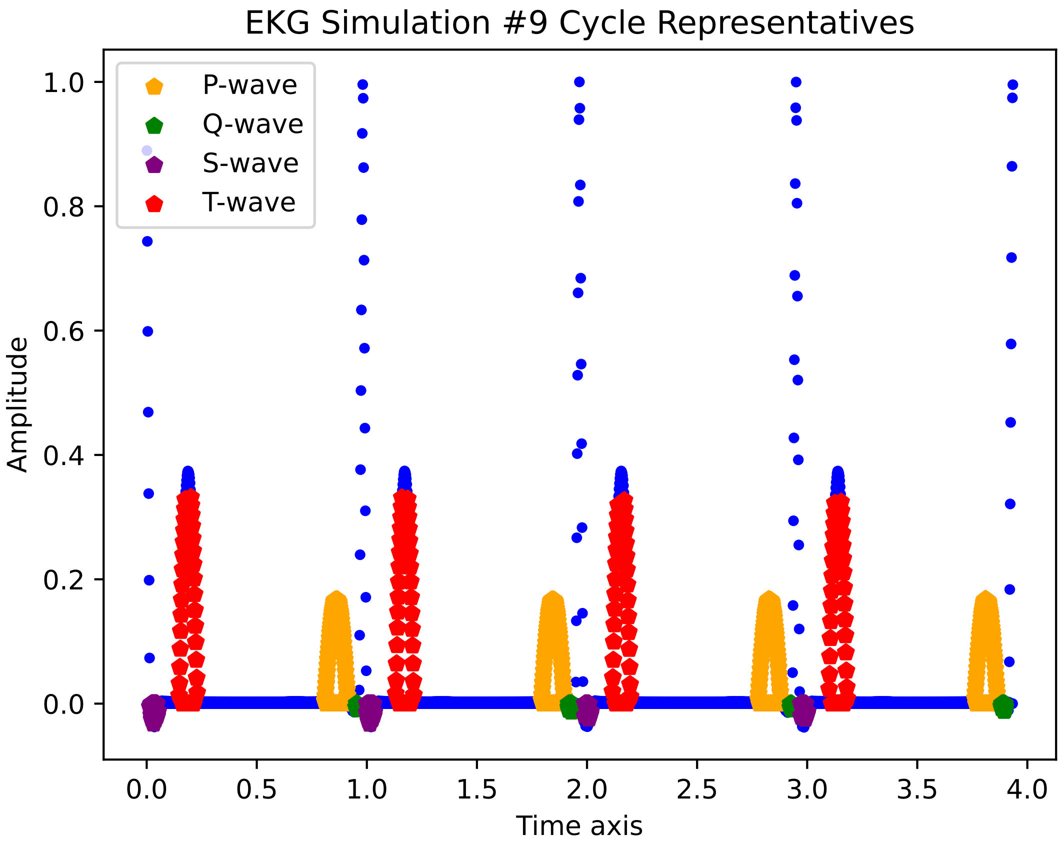
EKG Simulation #9 Cycle Representatives
P-wave
Q-wave
S-wave
T-wave
Amplitude
Time axis
1.0
0.8
0.6
0.4
0.2
0.0
0.0
0.5
1.0
1.5
2.0
2.5
3.0
3.5
4.0

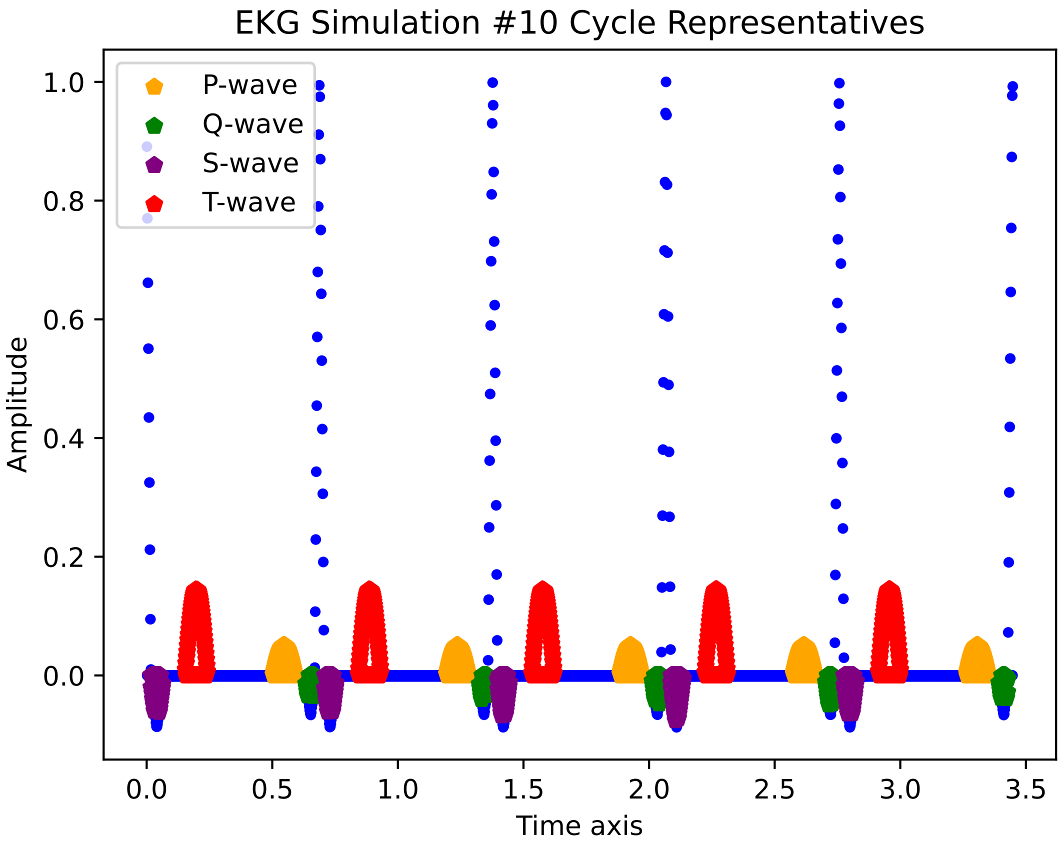

EKG Simulation #10 Cycle Representatives
P-wave
Q-wave
S-wave
T-wave
Amplitude
Time axis
1.0
0.8
0.6
0.4
0.2
0.0
0.0
0.5
1.0
1.5
2.0
2.5
3.0
3.5

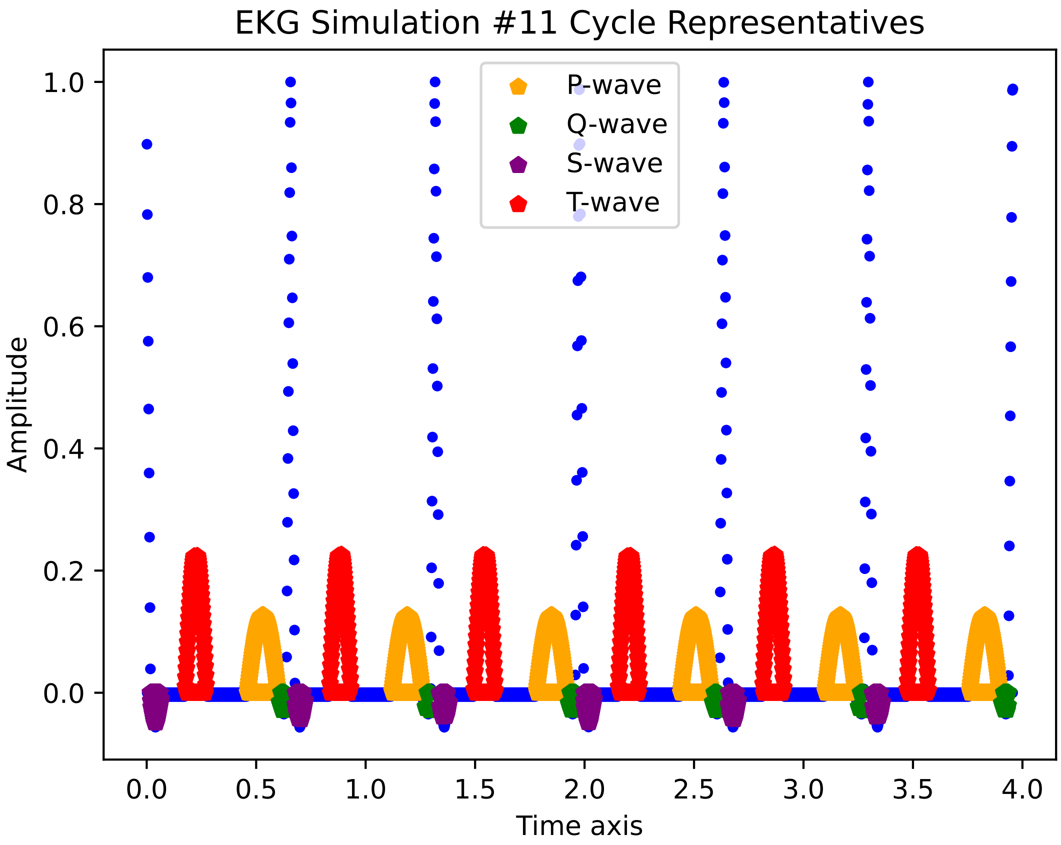
EKG Simulation #11 Cycle Representatives
P-wave
Q-wave
S-wave
T-wave
Amplitude
Time axis
1.0
0.8
0.6
0.4
0.2
0.0
0.0
0.5
1.0
1.5
2.0
2.5
3.0
3.5
4.0

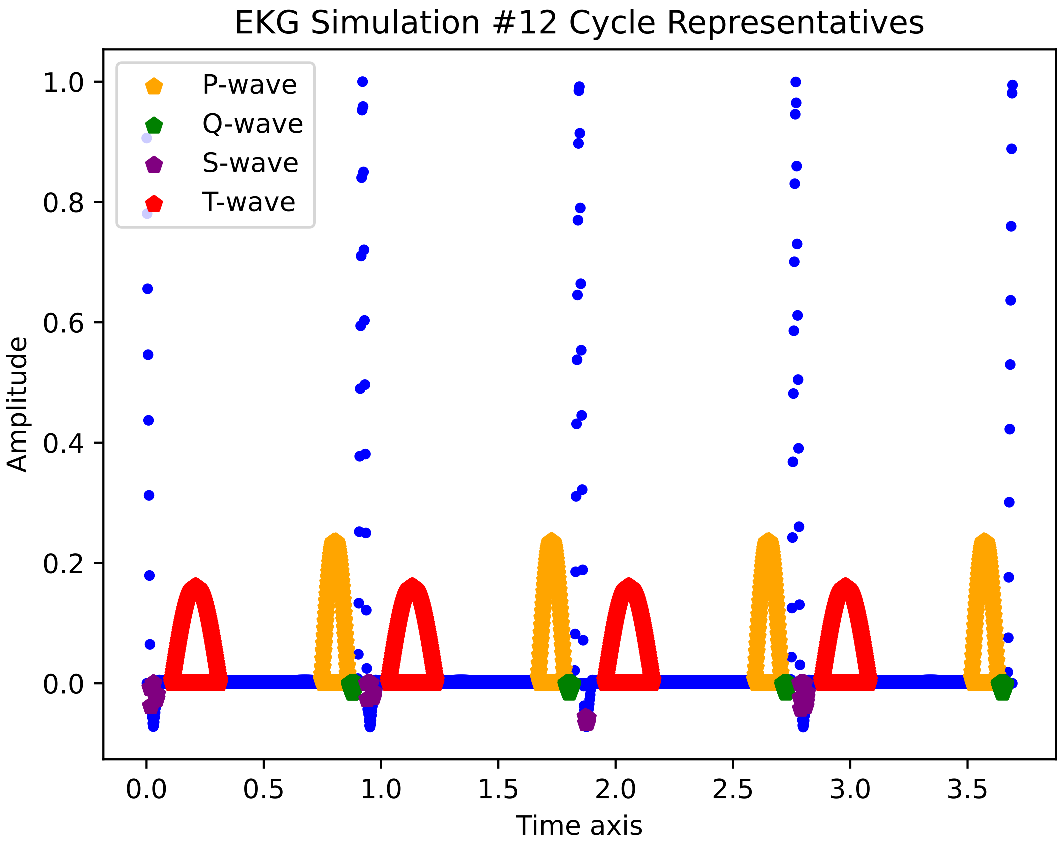

EKG Simulation #12 Cycle Representatives
P-wave
Q-wave
S-wave
T-wave
Amplitude
Time axis
1.0
0.8
0.6
0.4
0.2
0.0
0.0
0.5
1.0
1.5
2.0
2.5
3.0
3.5

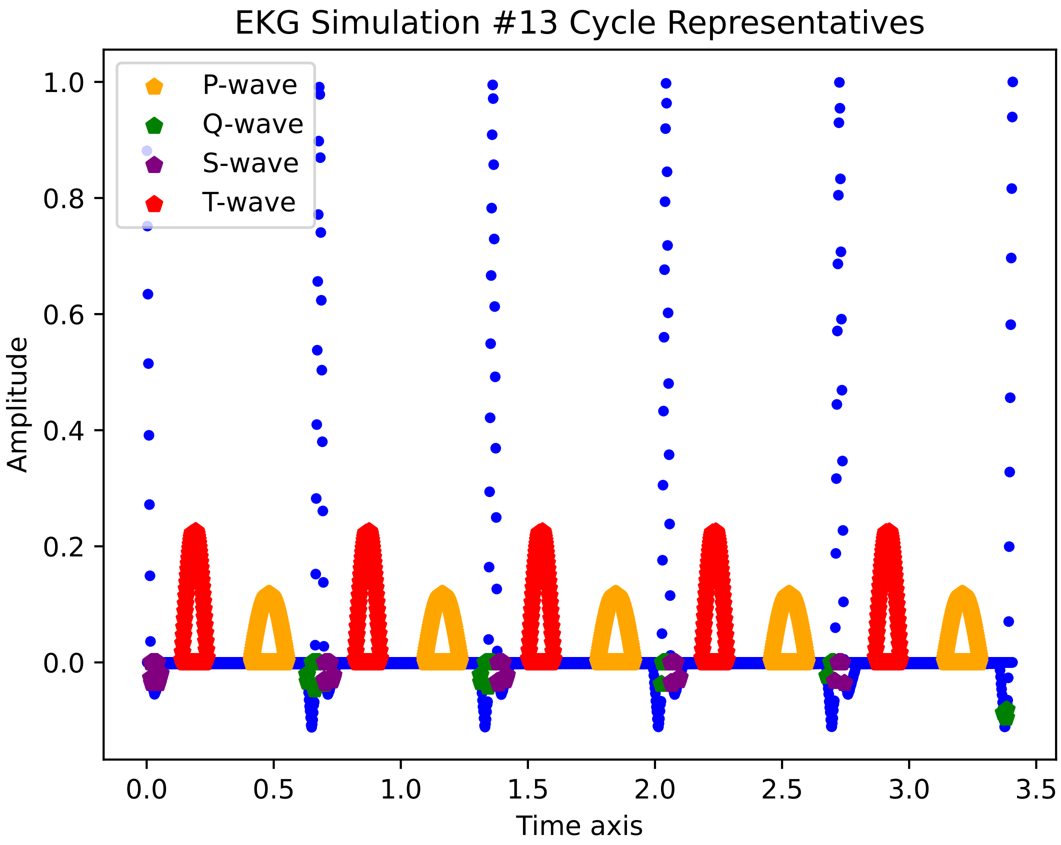

EKG Simulation #13 Cycle Representatives
P-wave
Q-wave
S-wave
T-wave
Amplitude
1.0
0.8
0.6
0.4
0.2
0.0
0.0
0.5
1.0
1.5
2.0
2.5
3.0
3.5
Time axis

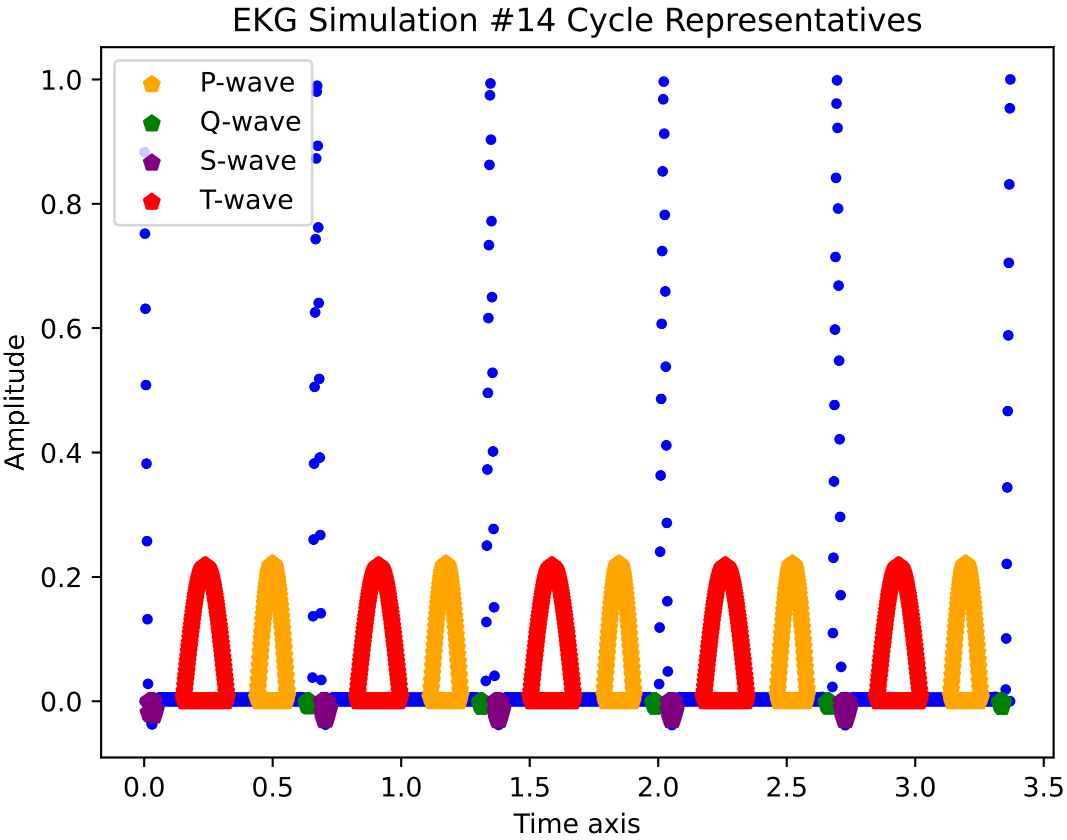
EKG Simulation #14 Cycle Representatives
P-wave
Q-wave
S-wave
T-wave
1.0
0.8
0.6
0.4
0.2
0.0
Amplitude
0.0
0.5
1.0
1.5
2.0
2.5
3.0
3.5
Time axis

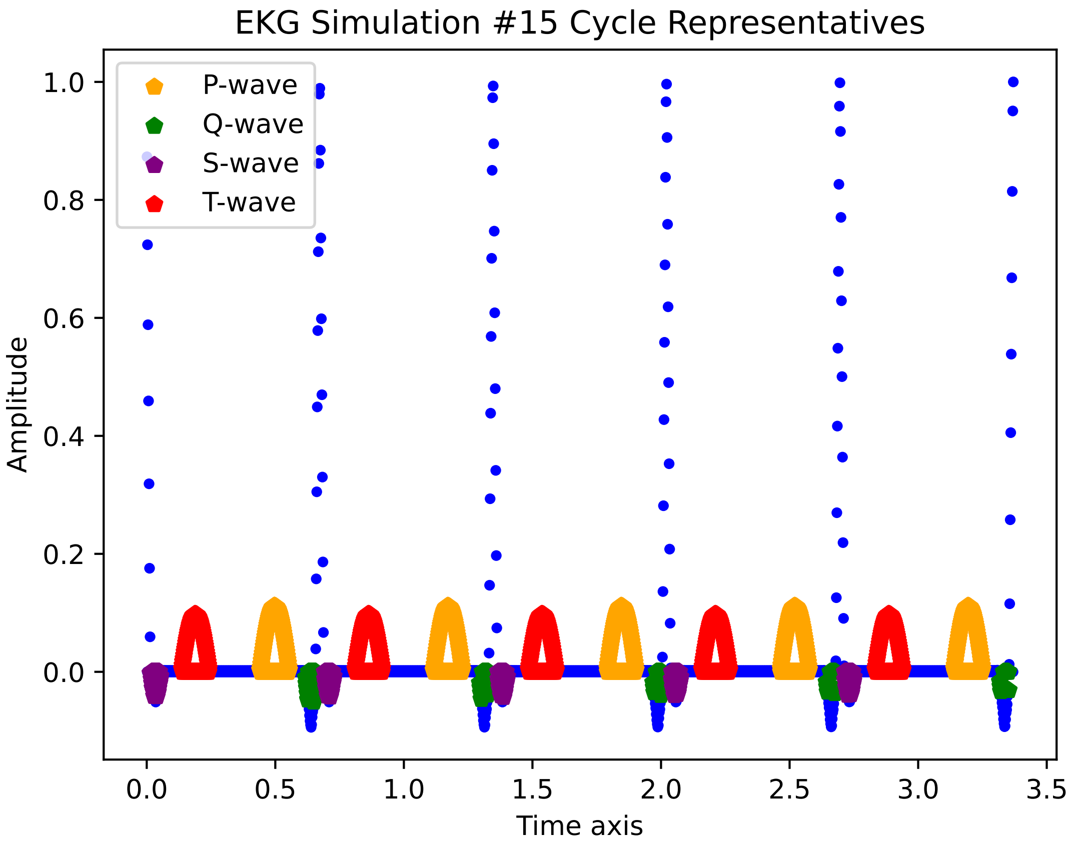

EKG Simulation #15 Cycle Representatives
P-wave
Q-wave
S-wave
T-wave
Amplitude
Time axis
1.0
0.8
0.6
0.4
0.2
0.0
0.0
0.5
1.0
1.5
2.0
2.5
3.0
3.5

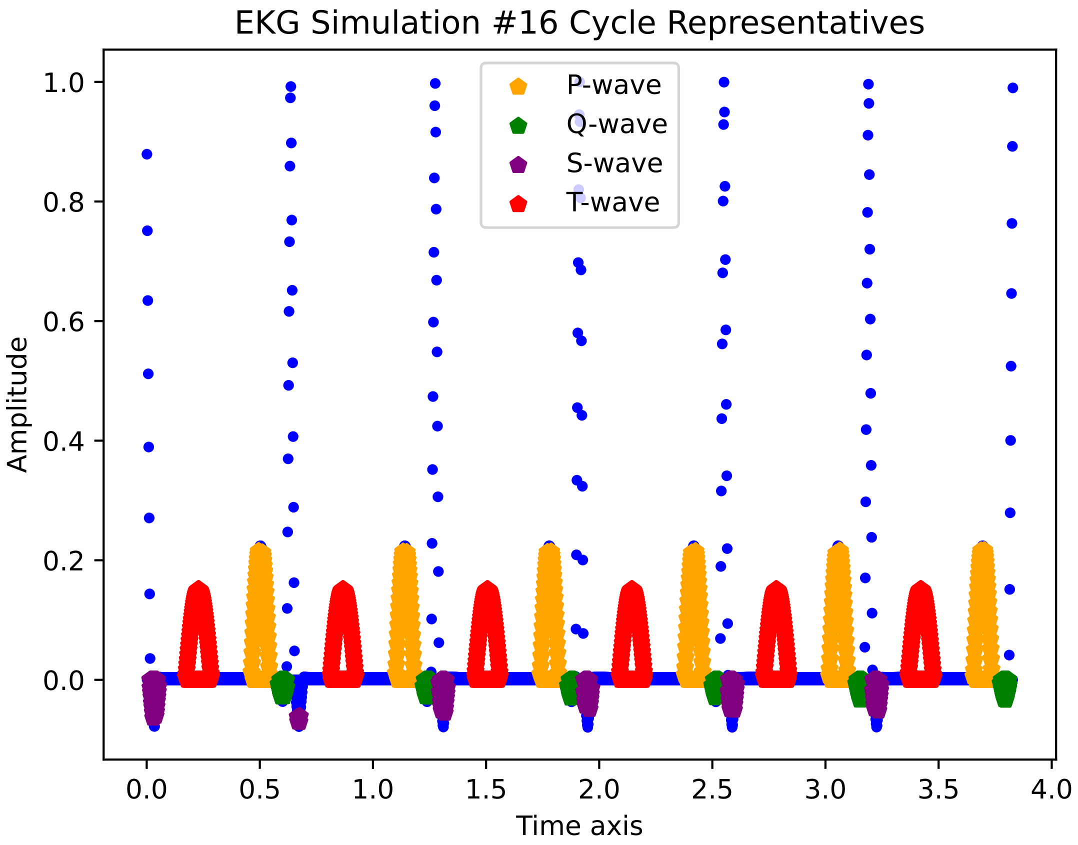
EKG Simulation #16 Cycle Representatives
P-wave
Q-wave
S-wave
T-wave
Amplitude
1.0
0.8
0.6
0.4
0.2
0.0
0.0
0.5
1.0
1.5
2.0
2.5
3.0
3.5
4.0
Time axis

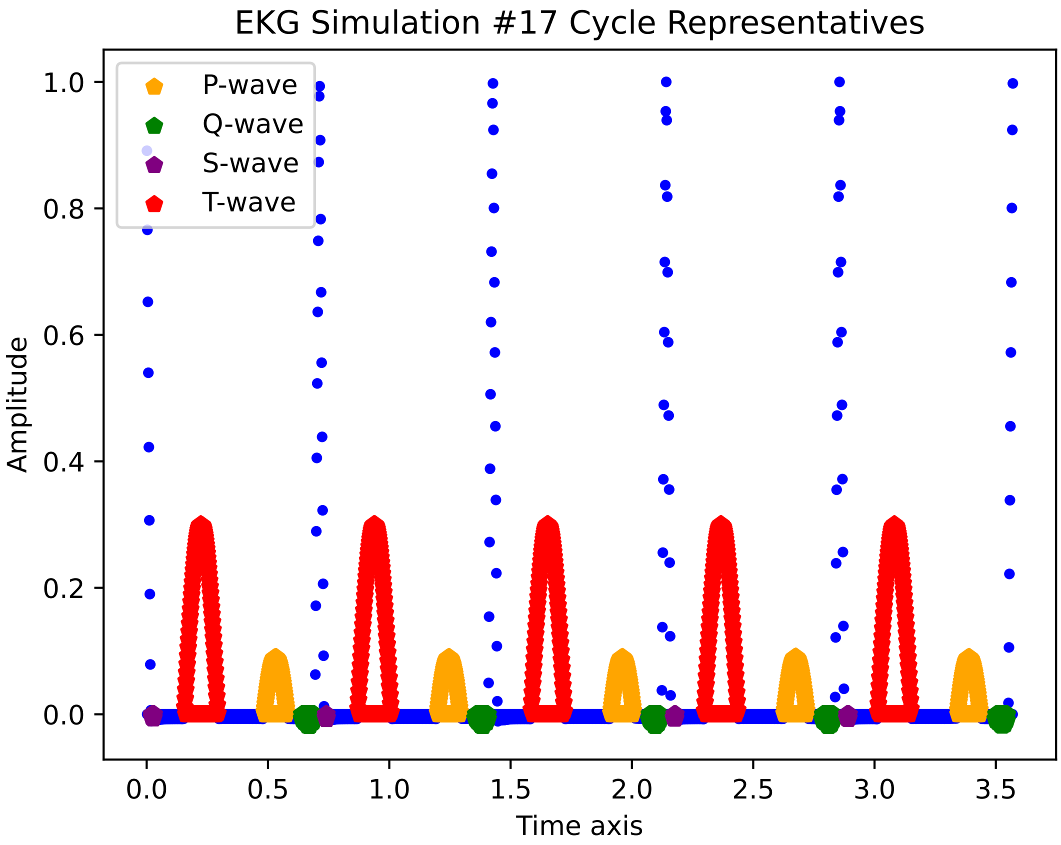
EKG Simulation #17 Cycle Representatives
P-wave
Q-wave
S-wave
T-wave
Amplitude
Time axis
0.0
0.2
0.4
0.6
0.8
1.0
0.0
0.5
1.0
1.5
2.0
2.5
3.0
3.5

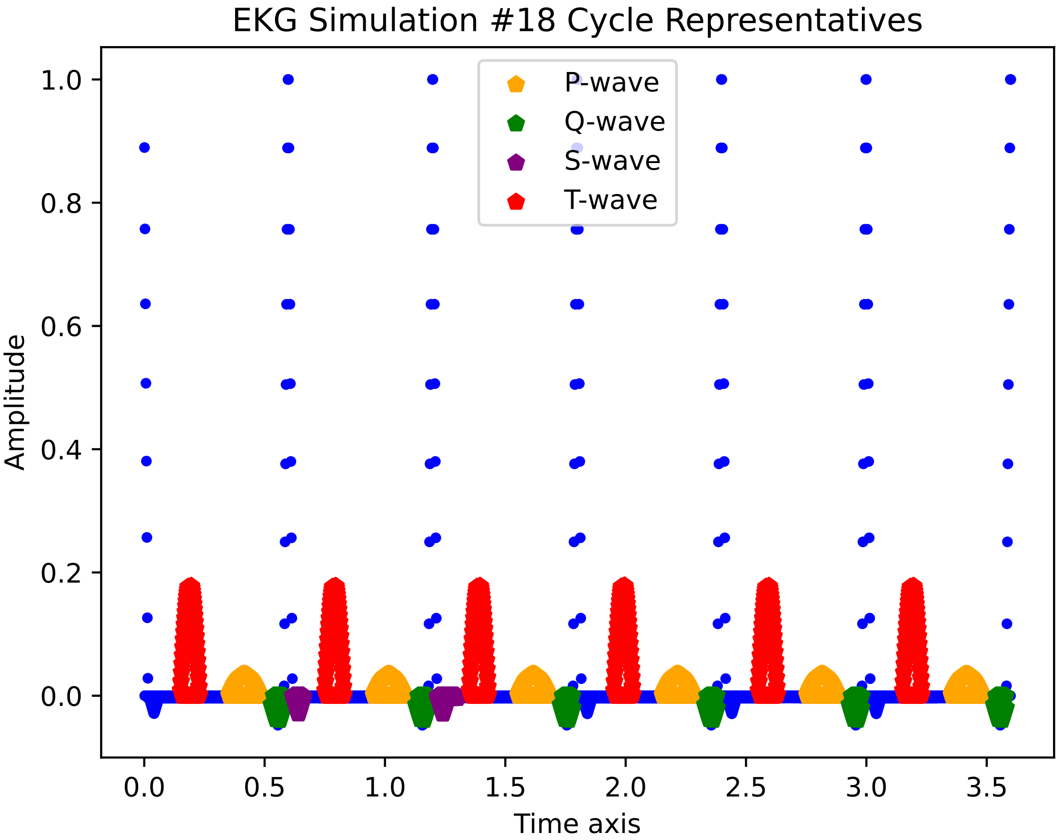

EKG Simulation #18 Cycle Representatives
P-wave
Q-wave
S-wave
T-wave
1.0
0.8
0.6
0.4
0.2
0.0
Amplitude
0.0
0.5
1.0
1.5
2.0
2.5
3.0
3.5
Time axis

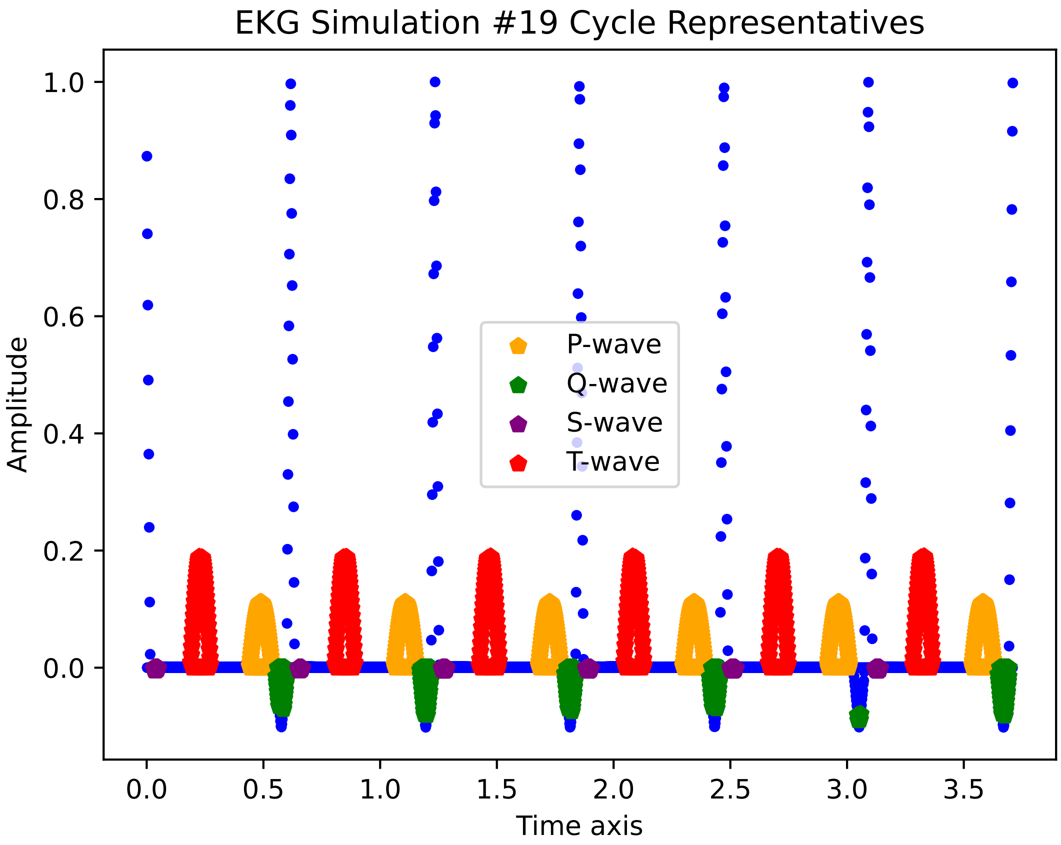
EKG Simulation #19 Cycle Representatives
Amplitude
Time axis
1.0
0.8
0.6
0.4
0.2
0.0
0.0
0.5
1.0
1.5
2.0
2.5
3.0
3.5
P-wave
Q-wave
S-wave
T-wave

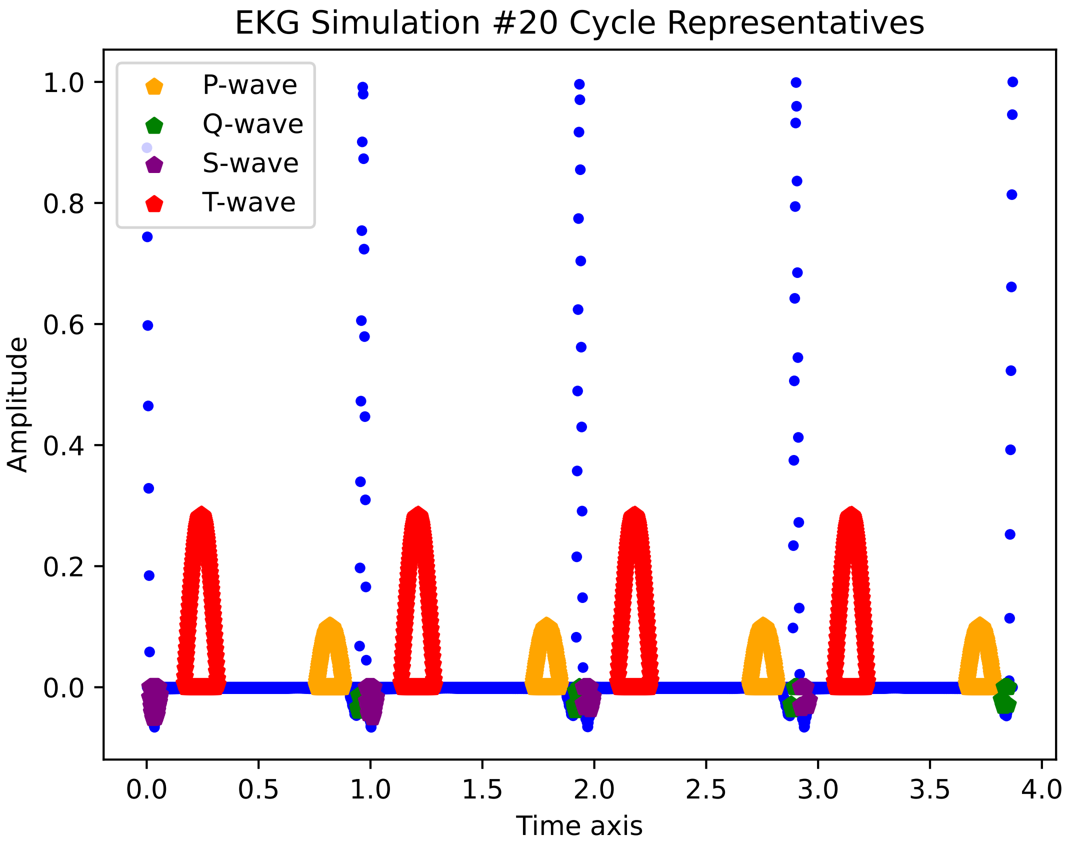
EKG Simulation #20 Cycle Representatives
P-wave
Q-wave
S-wave
T-wave
Amplitude
1.0
0.8
0.6
0.4
0.2
0.0
0.0
0.5
1.0
1.5
2.0
2.5
3.0
3.5
4.0
Time axis

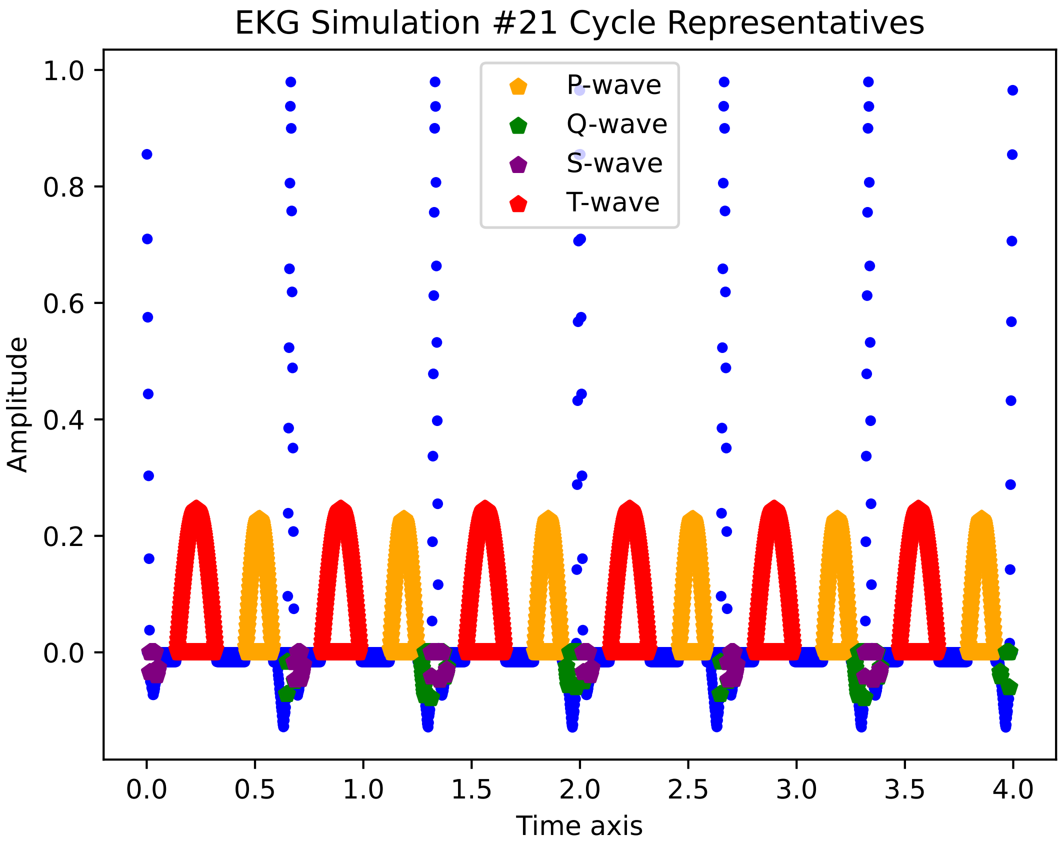

EKG Simulation #21 Cycle Representatives
P-wave
Q-wave
S-wave
T-wave
Amplitude
Time axis
1.0
0.8
0.6
0.4
0.2
0.0
0.0
0.5
1.0
1.5
2.0
2.5
3.0
3.5
4.0

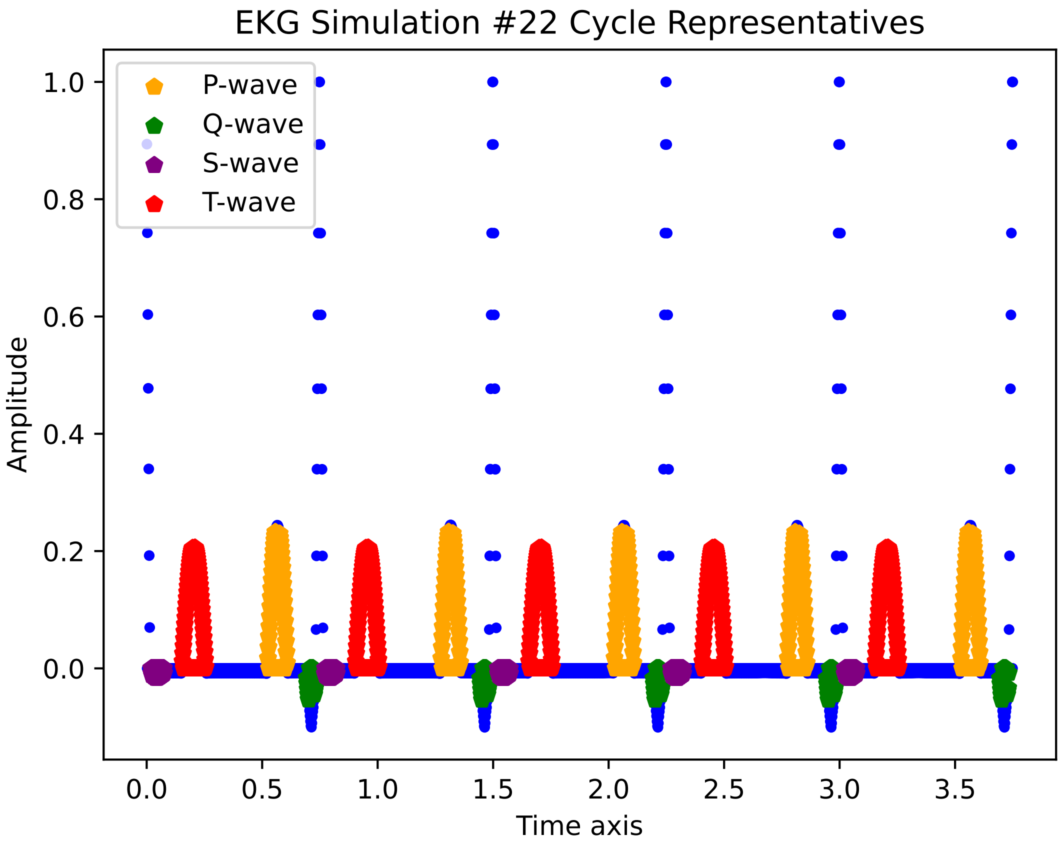

EKG Simulation #22 Cycle Representatives
P-wave
Q-wave
S-wave
T-wave
Amplitude
1.0
0.8
0.6
0.4
0.2
0.0
0.0
0.5
1.0
1.5
2.0
2.5
3.0
3.5
Time axis

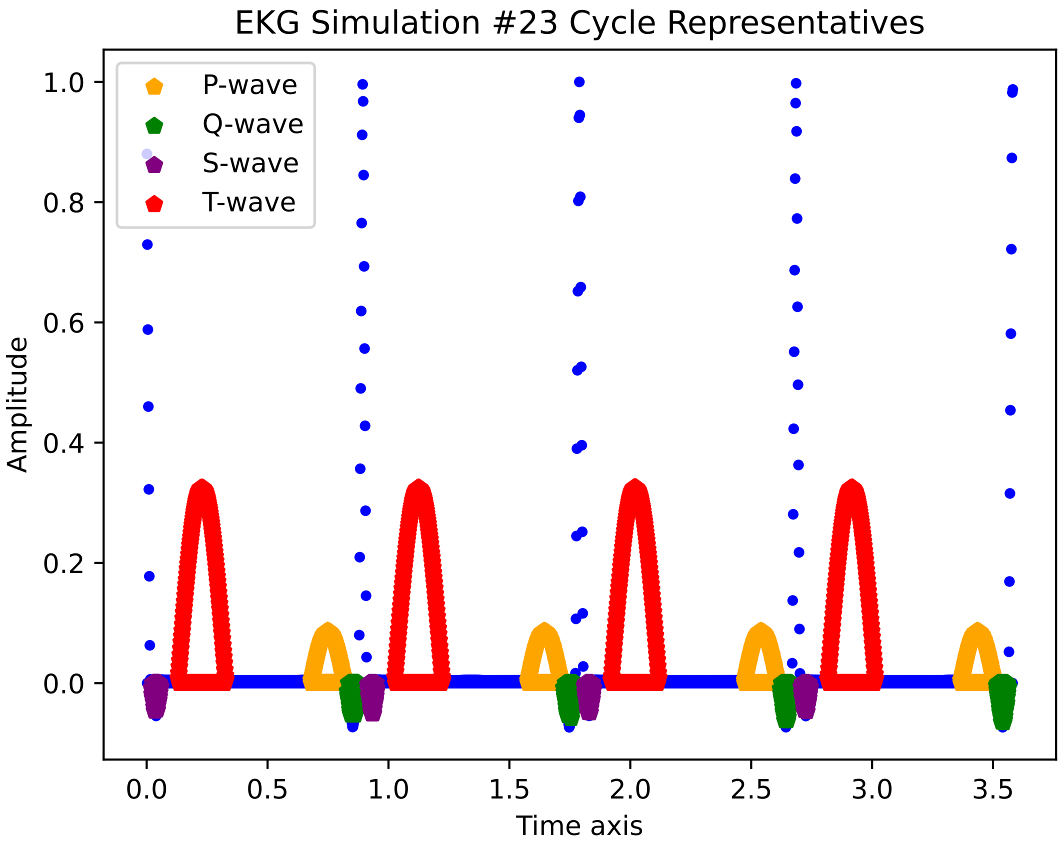
EKG Simulation #23 Cycle Representatives
P-wave
Q-wave
S-wave
T-wave
1.0
0.8
0.6
0.4
0.2
0.0
Amplitude
0.0
0.5
1.0
1.5
2.0
2.5
3.0
3.5
Time axis

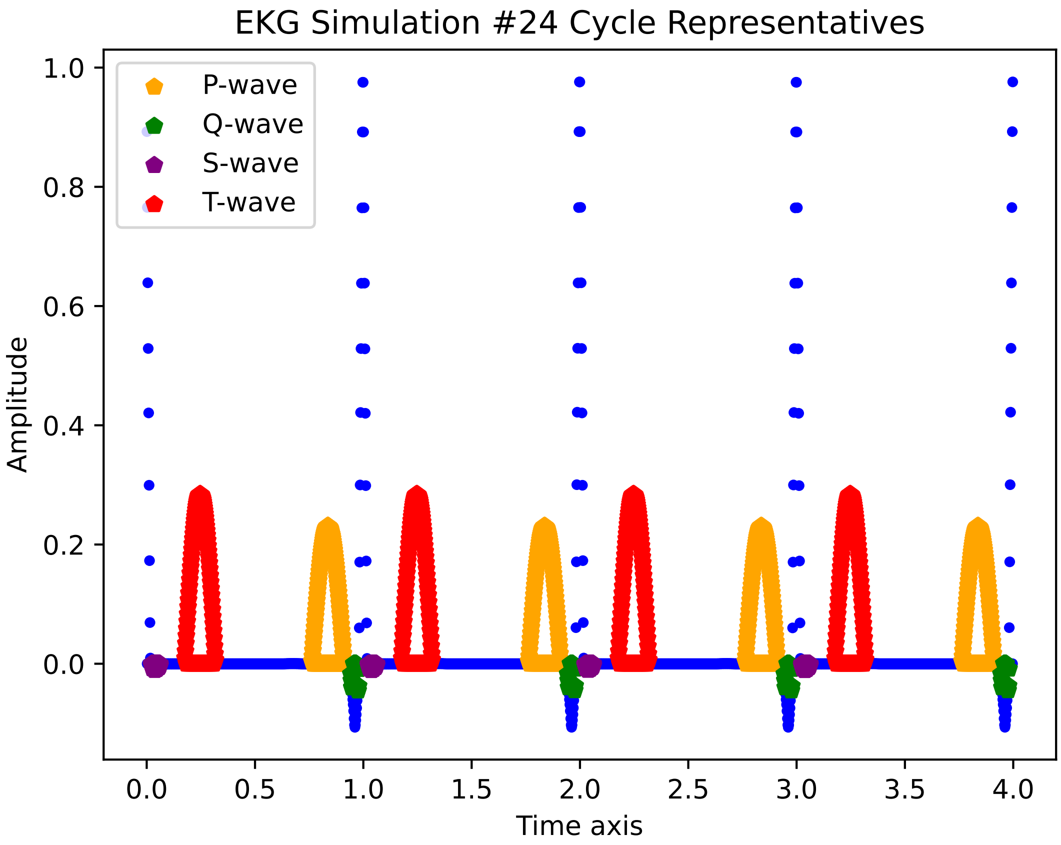

EKG Simulation #24 Cycle Representatives
P-wave
Q-wave
S-wave
T-wave
Amplitude
1.0
0.8
0.6
0.4
0.2
0.0
0.0
0.5
1.0
1.5
2.0
2.5
3.0
3.5
4.0
Time axis

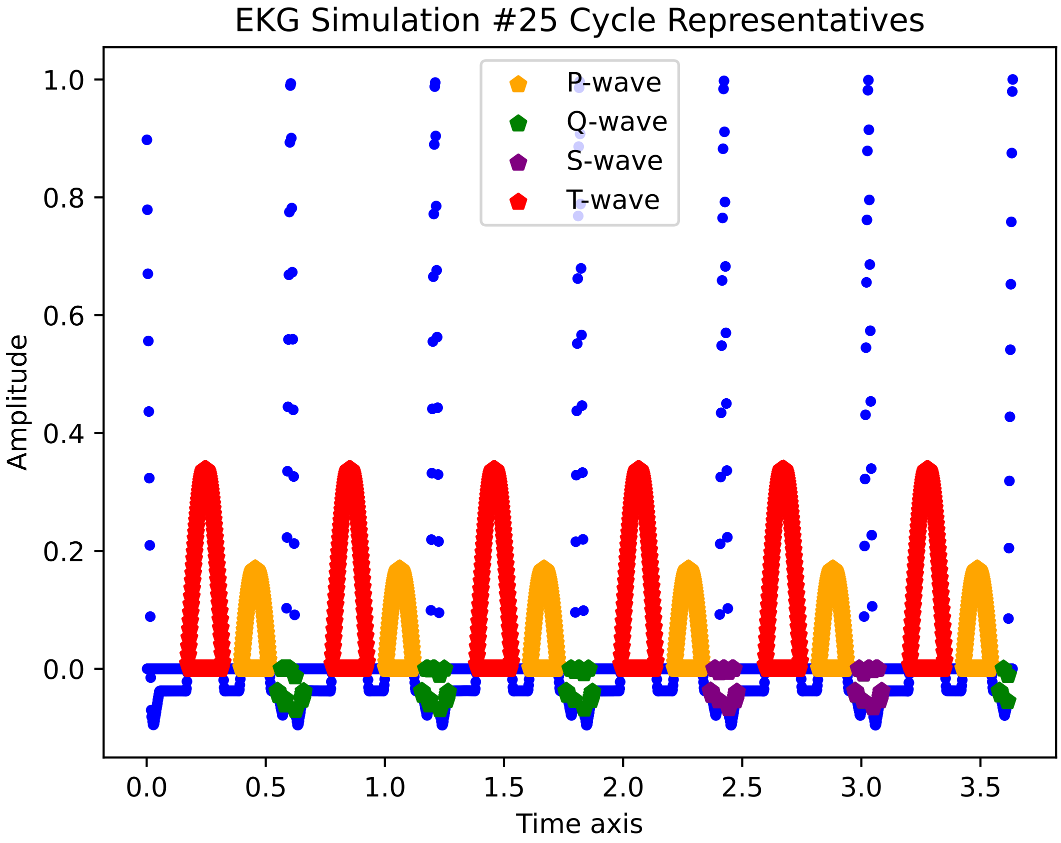
EKG Simulation #25 Cycle Representatives
P-wave
Q-wave
S-wave
T-wave
Amplitude
1.0
0.8
0.6
0.4
0.2
0.0
0.0
0.5
1.0
1.5
2.0
2.5
3.0
3.5
Time axis

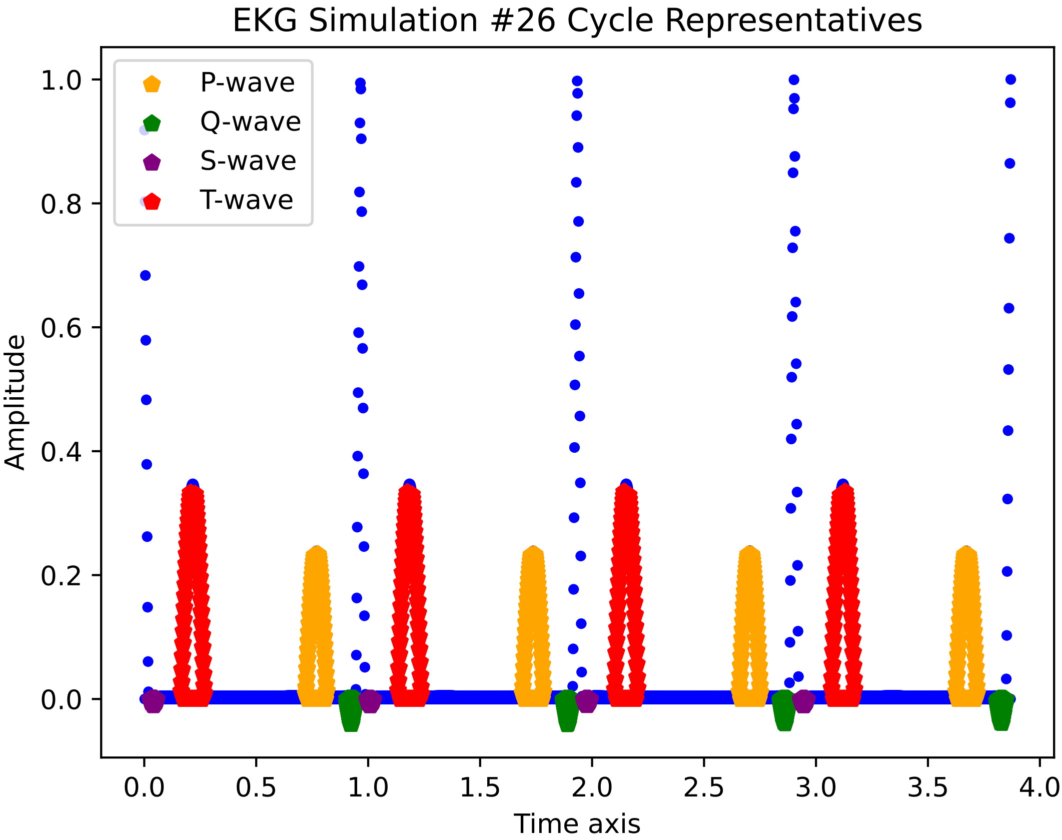
EKG Simulation #26 Cycle Representatives
P-wave
Q-wave
S-wave
T-wave
Amplitude
Time axis
0.0
0.2
0.4
0.6
0.8
1.0
0.0
0.5
1.0
1.5
2.0
2.5
3.0
3.5
4.0

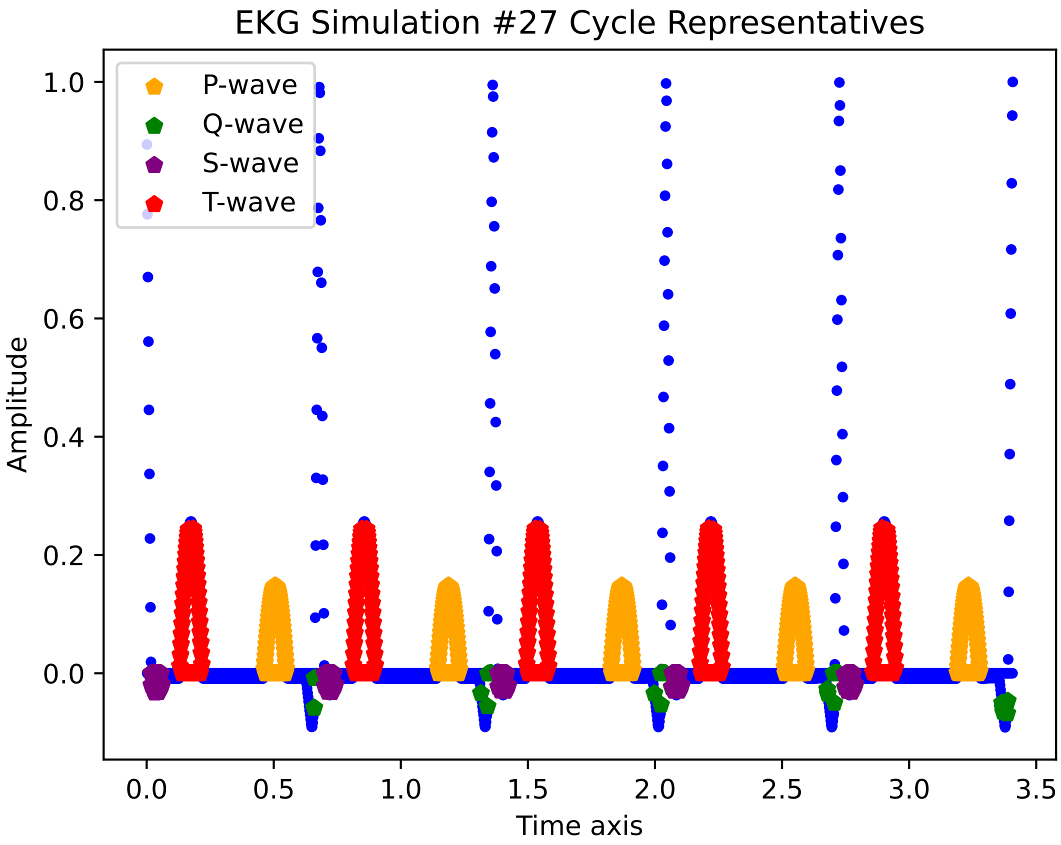
EKG Simulation #27 Cycle Representatives
P-wave
Q-wave
S-wave
T-wave
1.0
0.8
0.6
0.4
0.2
0.0
Amplitude
0.0
0.5
1.0
1.5
2.0
2.5
3.0
3.5
Time axis

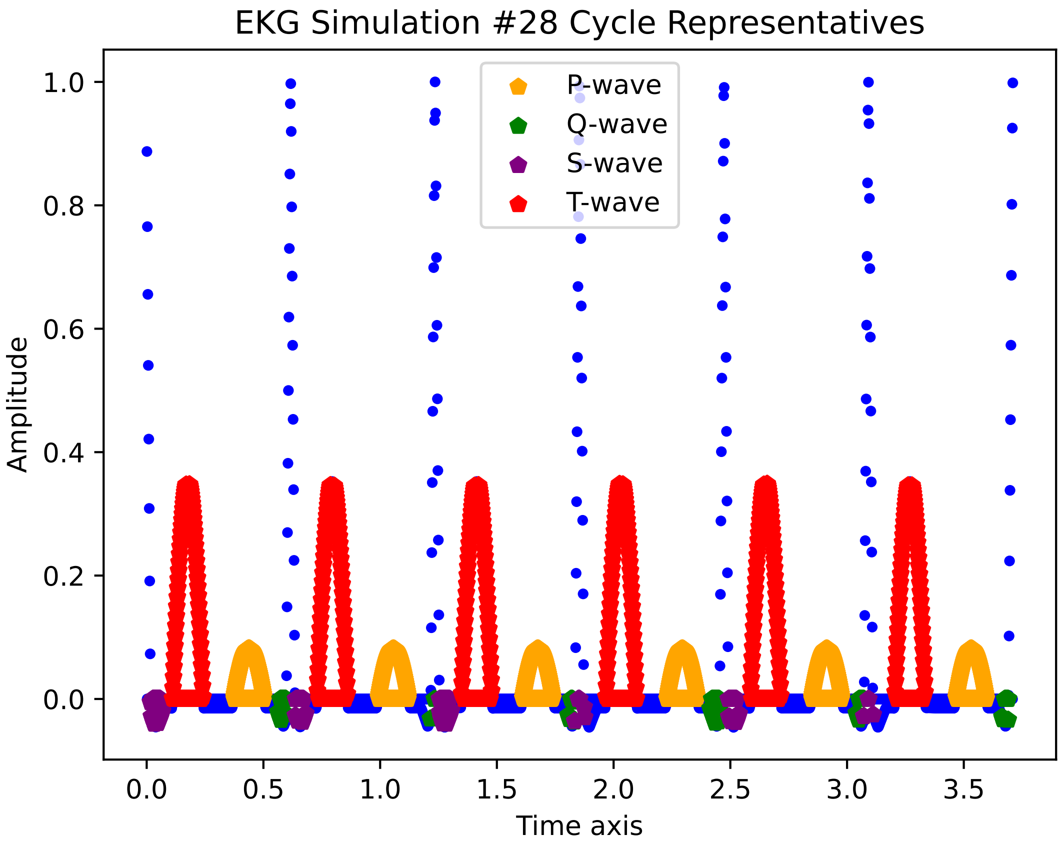

EKG Simulation #28 Cycle Representatives
P-wave
Q-wave
S-wave
T-wave
Amplitude
Time axis
0.0
0.2
0.4
0.6
0.8
1.0
0.0
0.5
1.0
1.5
2.0
2.5
3.0
3.5

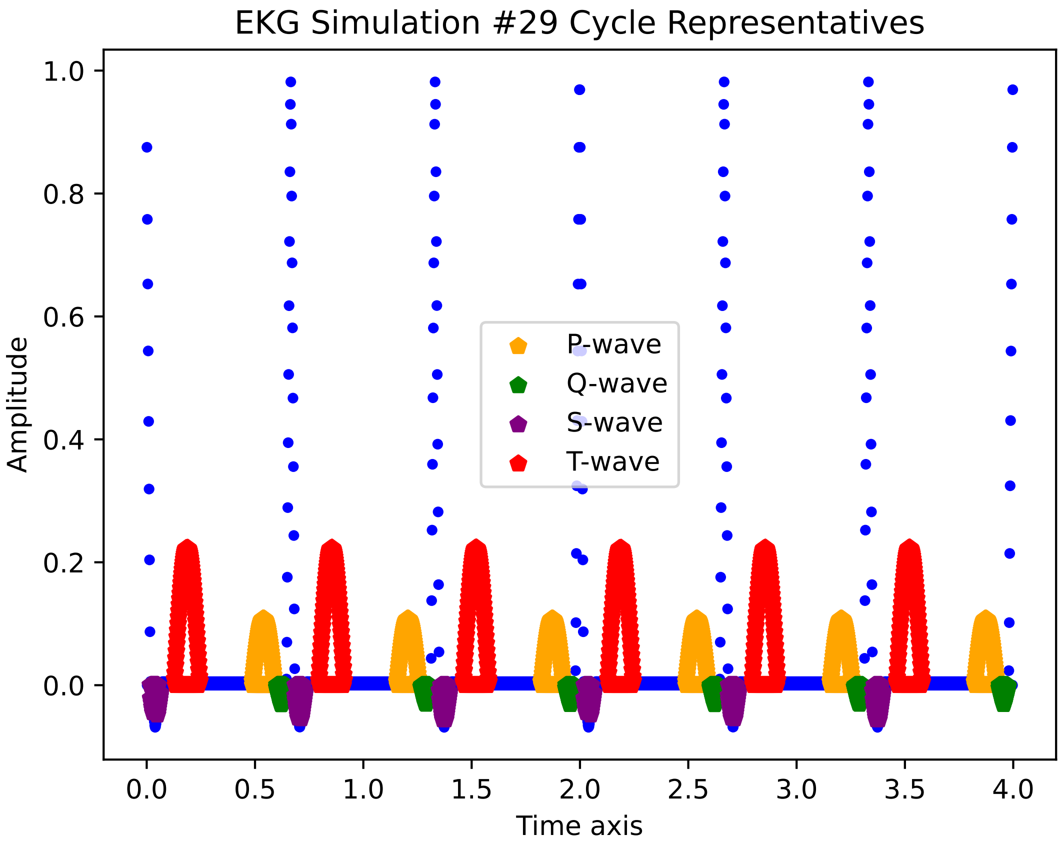
EKG Simulation #29 Cycle Representatives
Amplitude
Time axis
P-wave
Q-wave
S-wave
T-wave
1.0
0.8
0.6
0.4
0.2
0.0
0.0
0.5
1.0
1.5
2.0
2.5
3.0
3.5
4.0

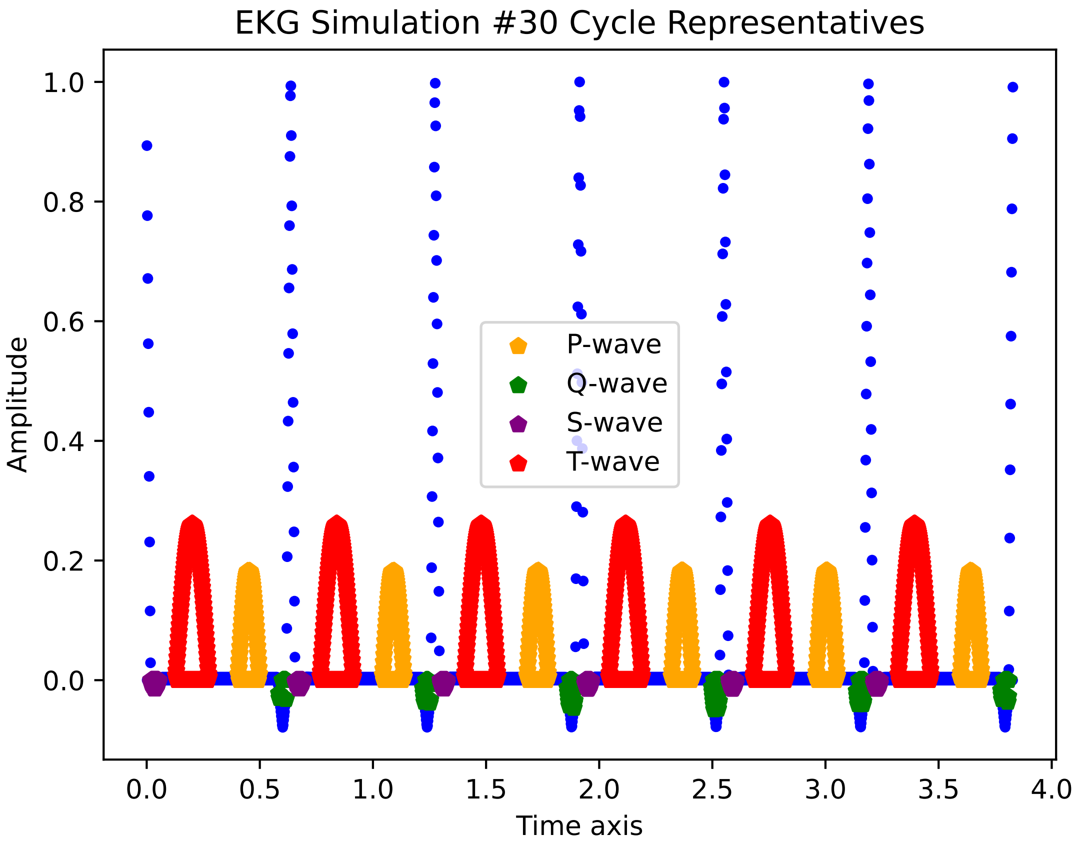

EKG Simulation #30 Cycle Representatives
Amplitude
Time axis
1.0
0.8
0.6
0.4
0.2
0.0
0.0
0.5
1.0
1.5
2.0
2.5
3.0
3.5
4.0
P-wave
Q-wave
S-wave
T-wave

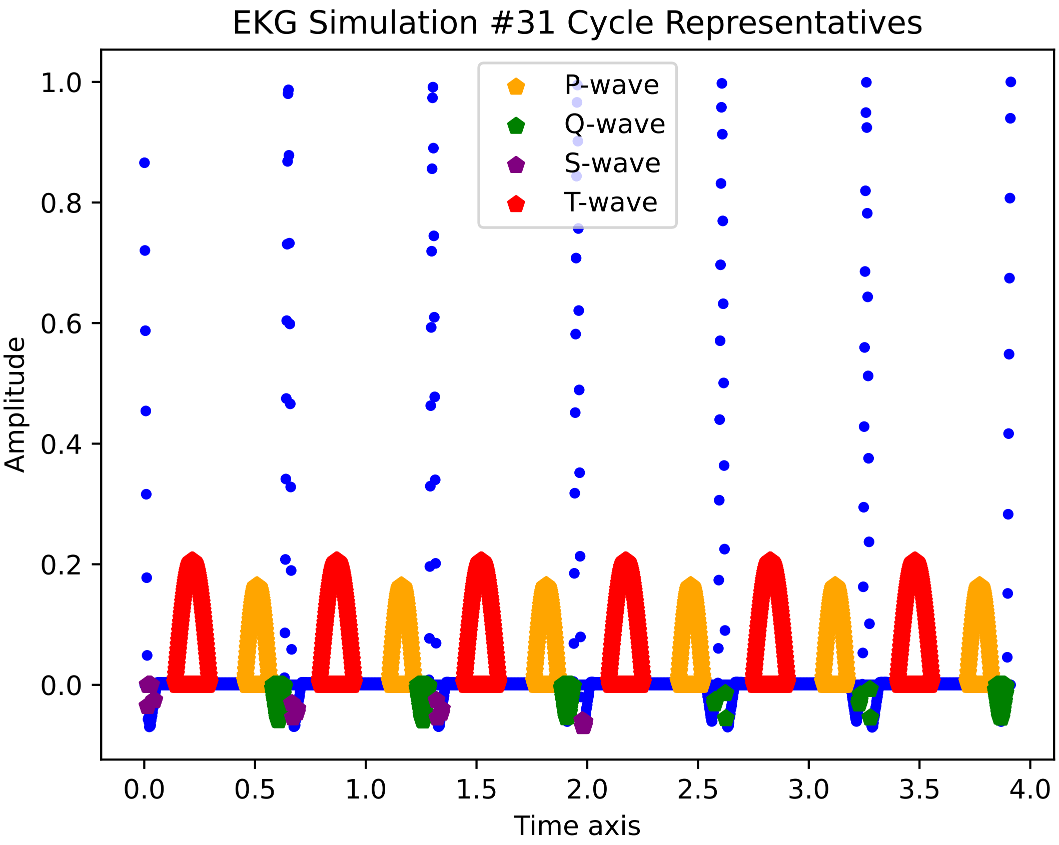

EKG Simulation #31 Cycle Representatives
P-wave
Q-wave
S-wave
T-wave
Amplitude
Time axis
1.0
0.8
0.6
0.4
0.2
0.0
0.0
0.5
1.0
1.5
2.0
2.5
3.0
3.5
4.0

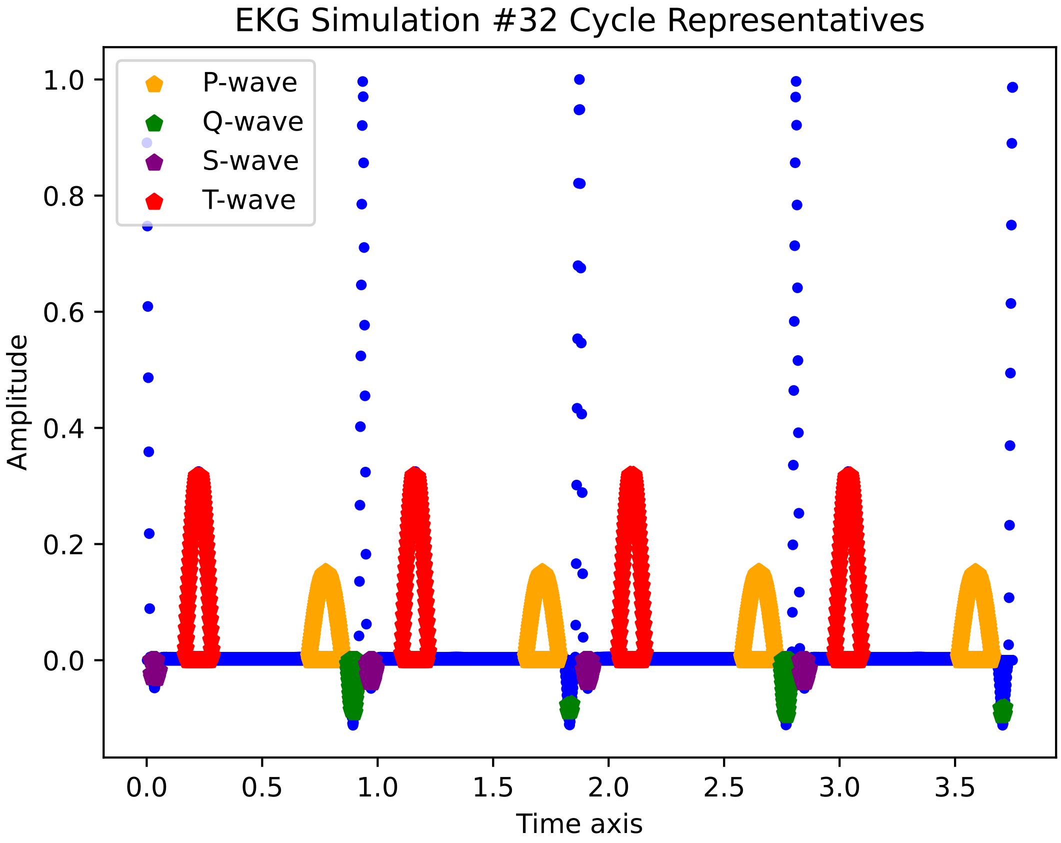

EKG Simulation #32 Cycle Representatives
P-wave
Q-wave
S-wave
T-wave
1.0
0.8
0.6
0.4
0.2
0.0
Amplitude
0.0
0.5
1.0
1.5
2.0
2.5
3.0
3.5
Time axis

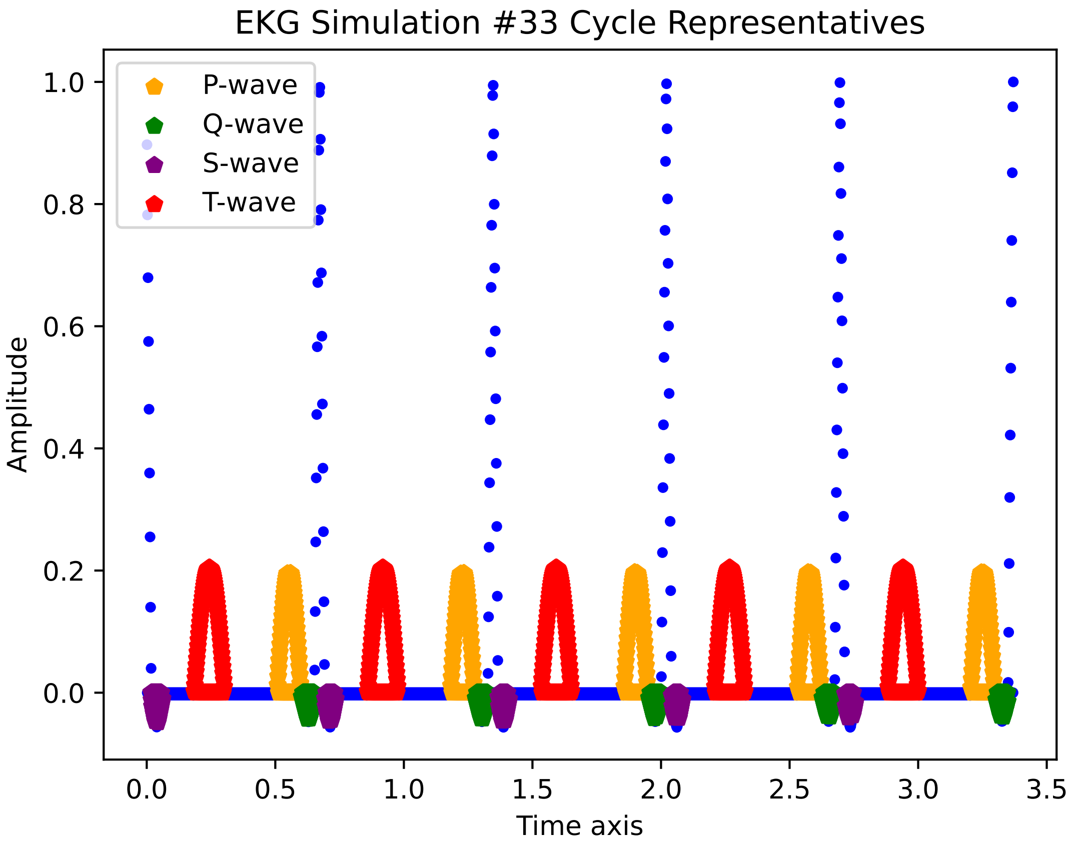
EKG Simulation #33 Cycle Representatives
P-wave
Q-wave
S-wave
T-wave
1.0
0.8
0.6
0.4
0.2
0.0
Amplitude
0.0
0.5
1.0
1.5
2.0
2.5
3.0
3.5
Time axis

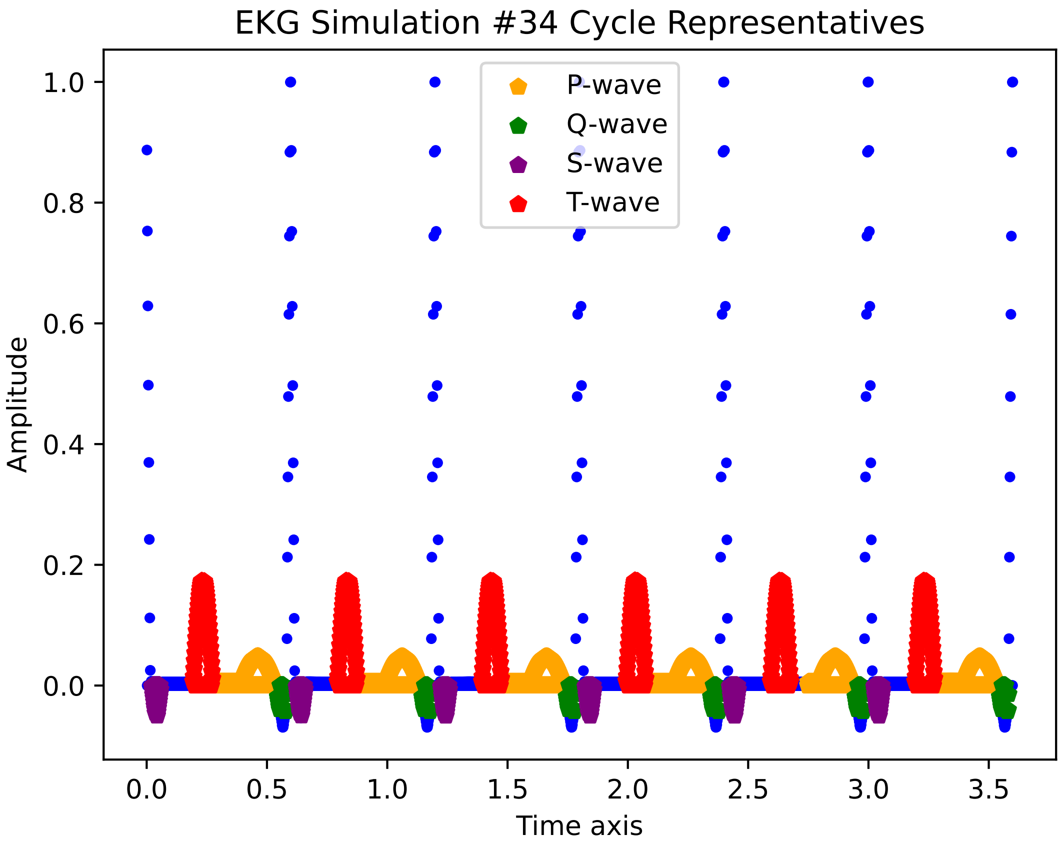
EKG Simulation #34 Cycle Representatives
P-wave
Q-wave
S-wave
T-wave
Amplitude
Time axis
1.0
0.8
0.6
0.4
0.2
0.0
0.0
0.5
1.0
1.5
2.0
2.5
3.0
3.5

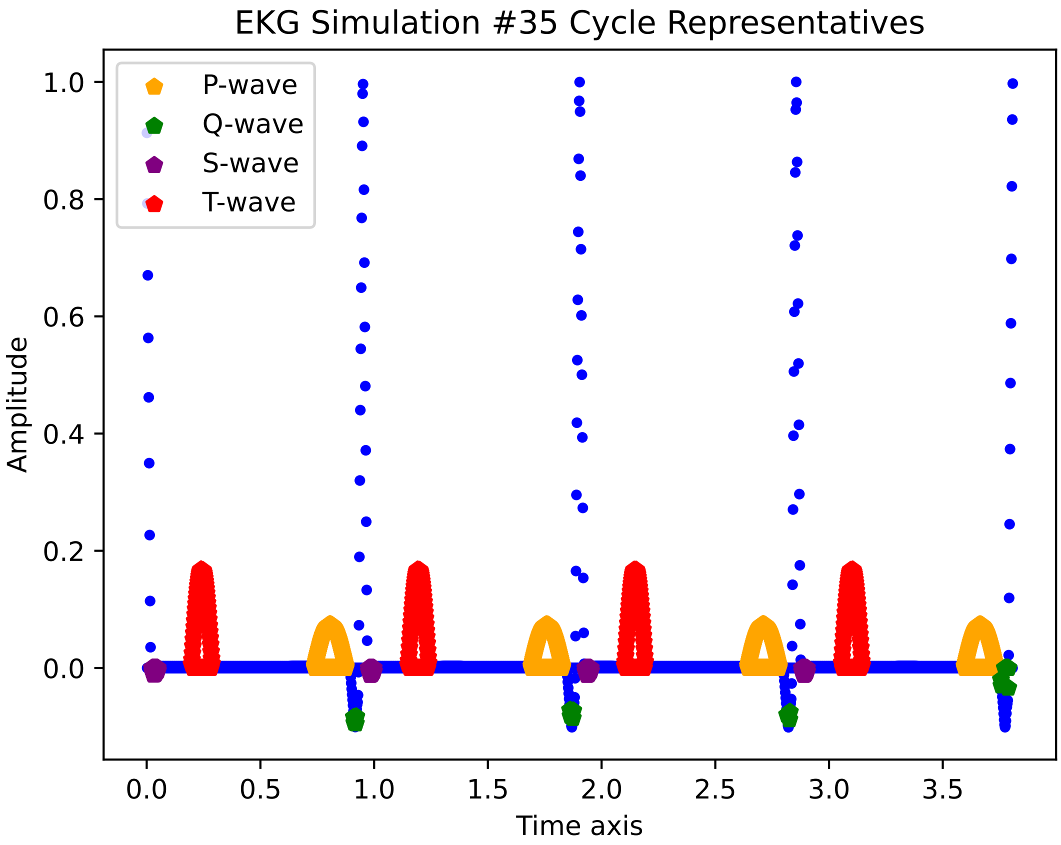

EKG Simulation #35 Cycle Representatives
P-wave
Q-wave
S-wave
T-wave
Amplitude
Time axis
1.0
0.8
0.6
0.4
0.2
0.0
0.0
0.5
1.0
1.5
2.0
2.5
3.0
3.5

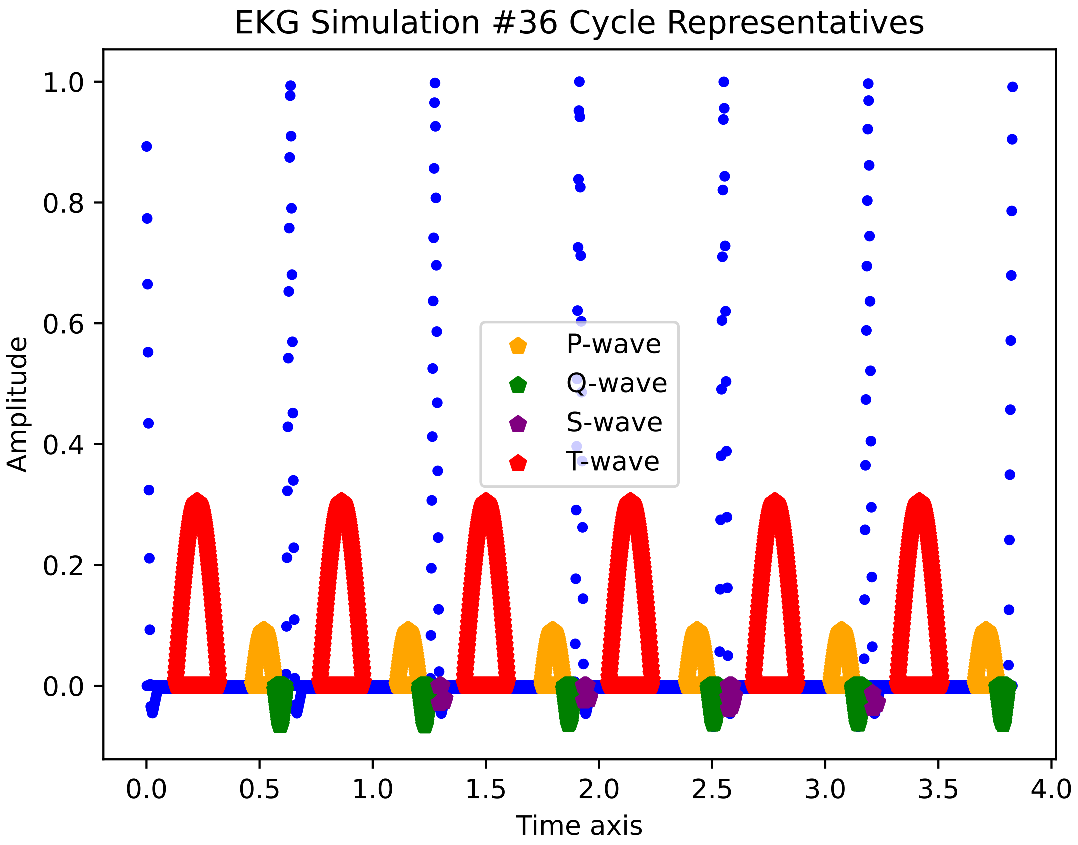
EKG Simulation #36 Cycle Representatives
Amplitude
Time axis
P-wave
Q-wave
S-wave
T-wave
0.0
0.2
0.4
0.6
0.8
1.0
0.0
0.5
1.0
1.5
2.0
2.5
3.0
3.5
4.0

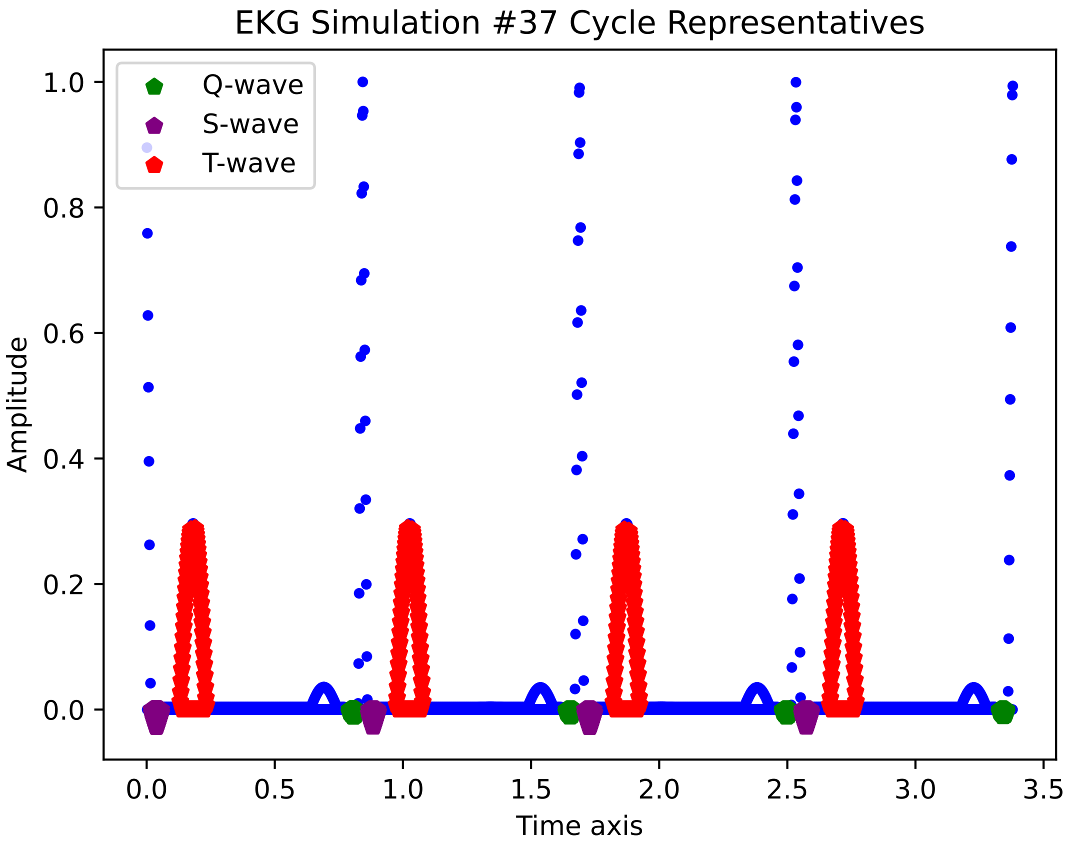
EKG Simulation #37 Cycle Representatives
Q-wave
S-wave
T-wave
1.0
0.8
0.6
0.4
0.2
0.0
Amplitude
0.0
0.5
1.0
1.5
2.0
2.5
3.0
3.5
Time axis

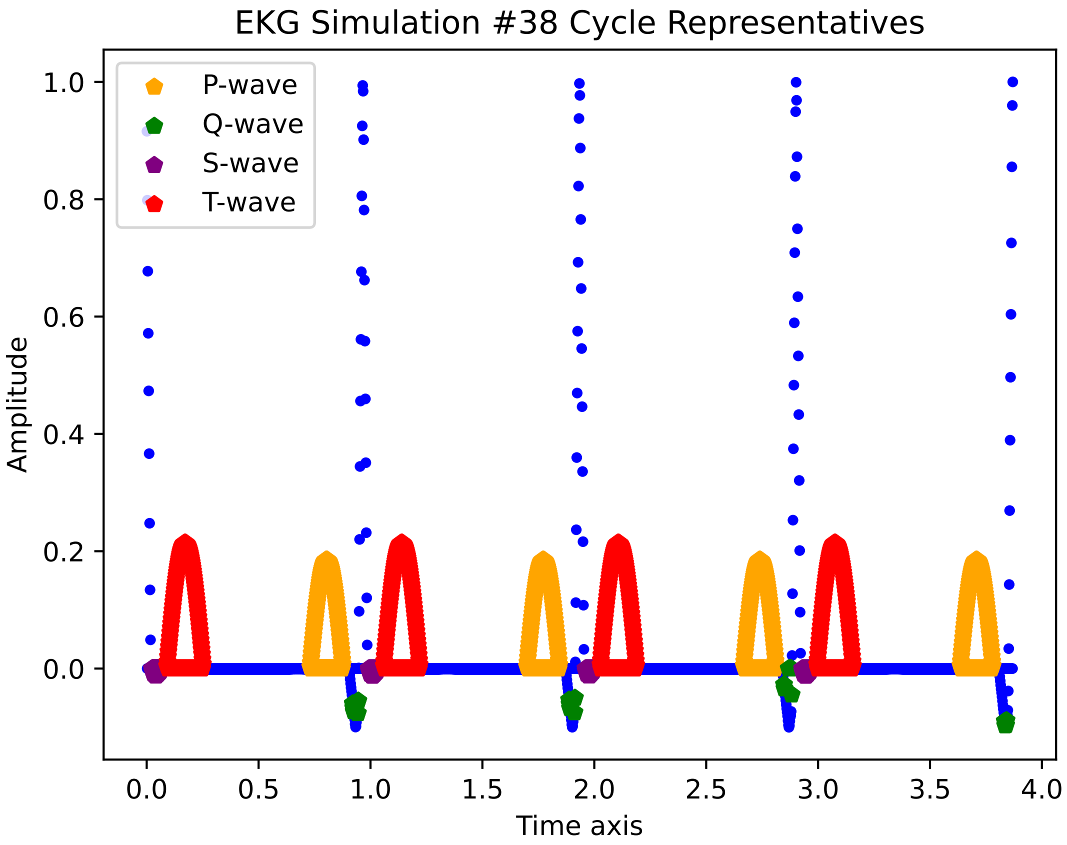
EKG Simulation #38 Cycle Representatives
P-wave
Q-wave
S-wave
T-wave
Amplitude
Time axis
1.0
0.8
0.6
0.4
0.2
0.0
0.0
0.5
1.0
1.5
2.0
2.5
3.0
3.5
4.0

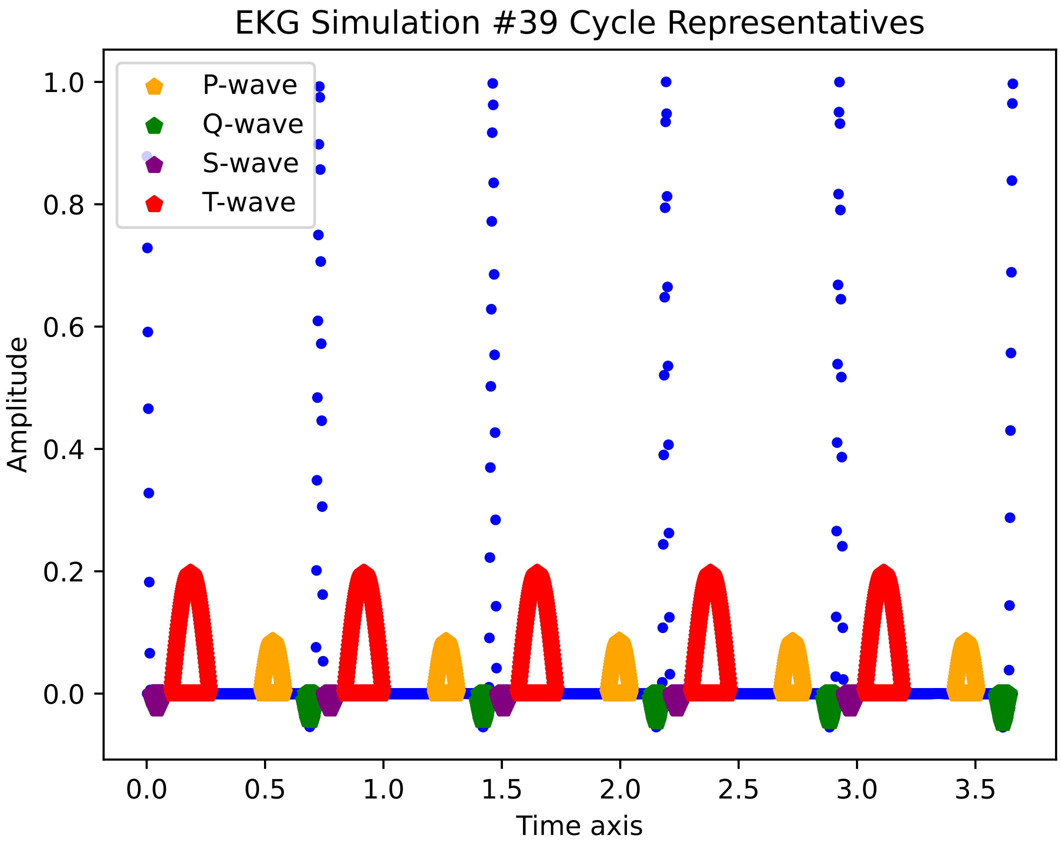

EKG Simulation #39 Cycle Representatives
P-wave
Q-wave
S-wave
T-wave
Amplitude
1.0
0.8
0.6
0.4
0.2
0.0
0.0
0.5
1.0
1.5
2.0
2.5
3.0
3.5
Time axis

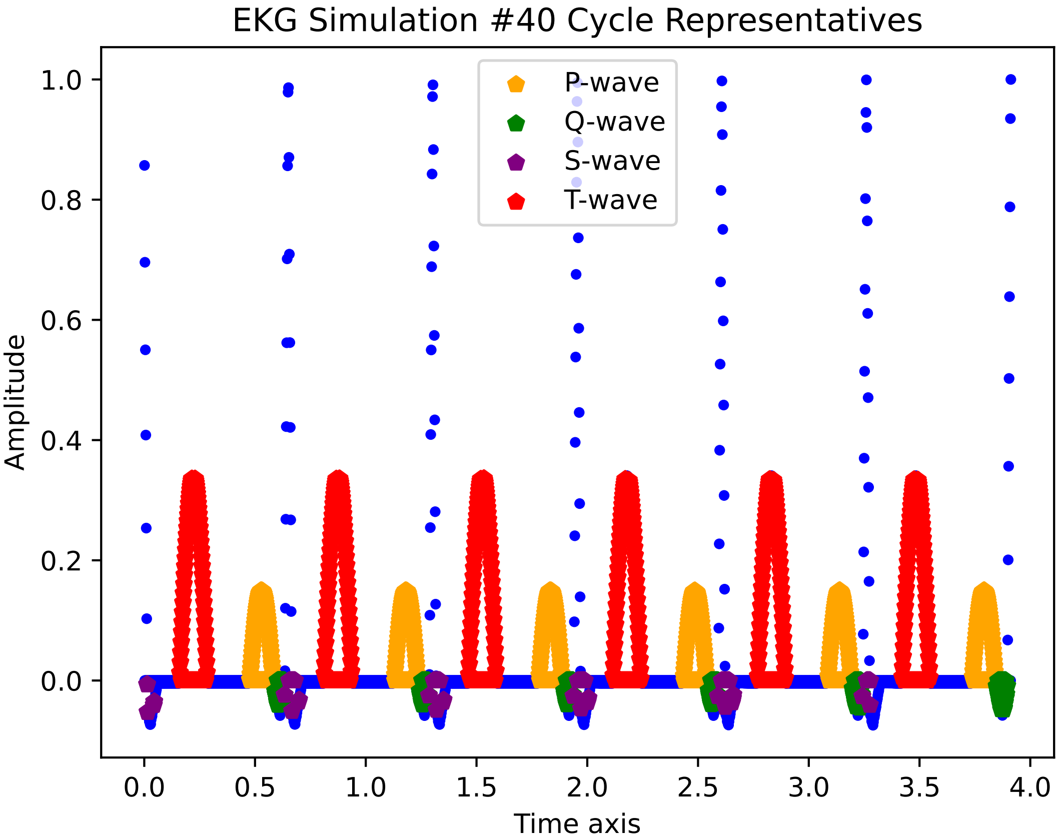

EKG Simulation #40 Cycle Representatives
P-wave
Q-wave
S-wave
T-wave
Amplitude
Time axis
1.0
0.8
0.6
0.4
0.2
0.0
0.0
0.5
1.0
1.5
2.0
2.5
3.0
3.5
4.0

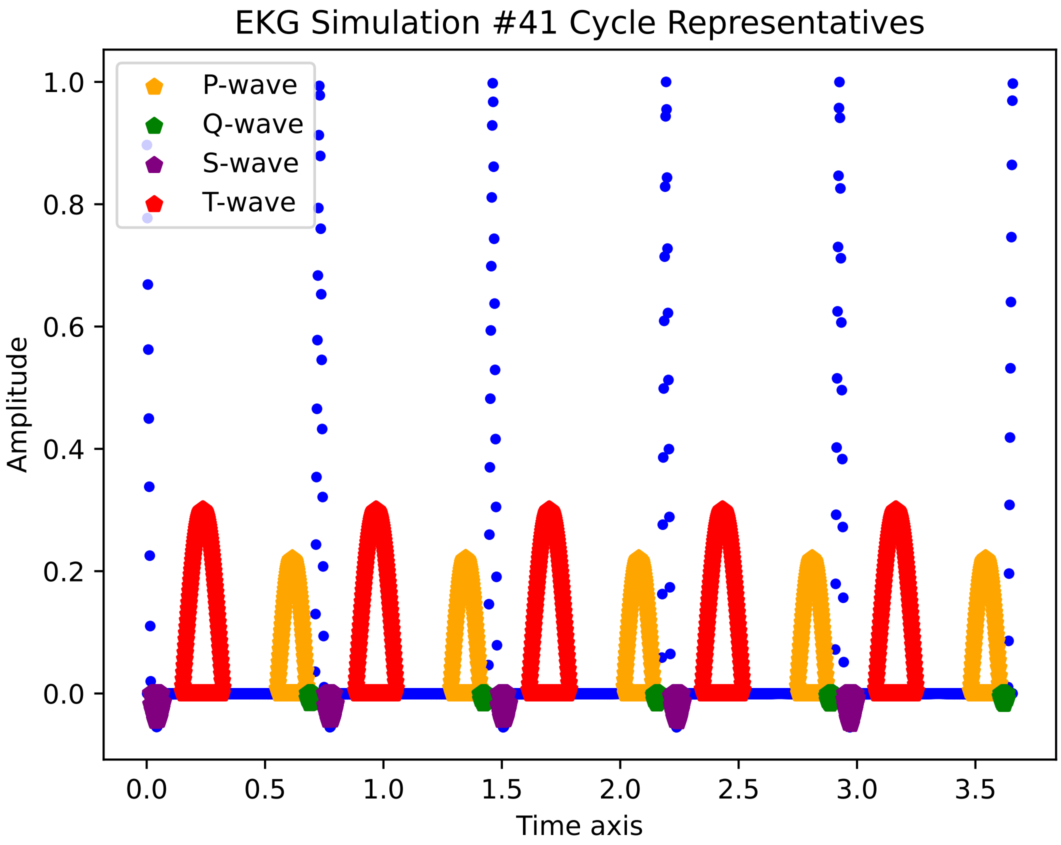
EKG Simulation #41 Cycle Representatives
P-wave
Q-wave
S-wave
T-wave
1.0
0.8
0.6
0.4
0.2
0.0
Amplitude
0.0
0.5
1.0
1.5
2.0
2.5
3.0
3.5
Time axis

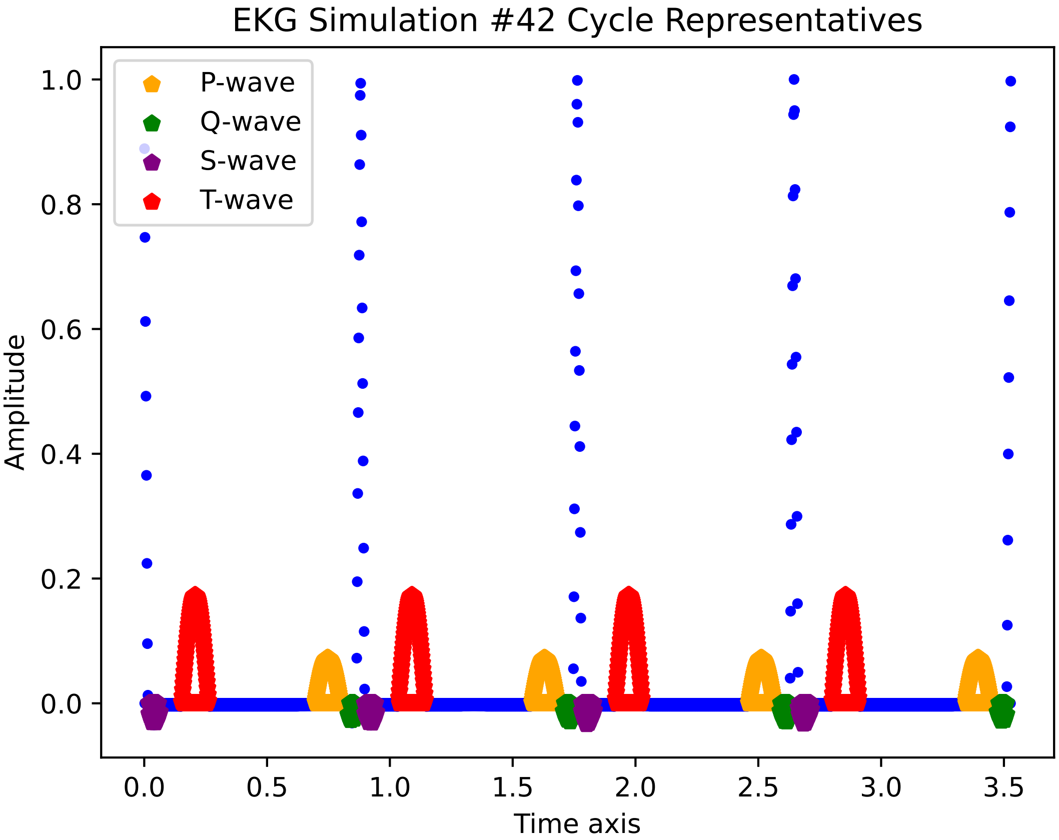
EKG Simulation #42 Cycle Representatives
P-wave
Q-wave
S-wave
T-wave
Amplitude
1.0
0.8
0.6
0.4
0.2
0.0
0.0
0.5
1.0
1.5
2.0
2.5
3.0
3.5
Time axis

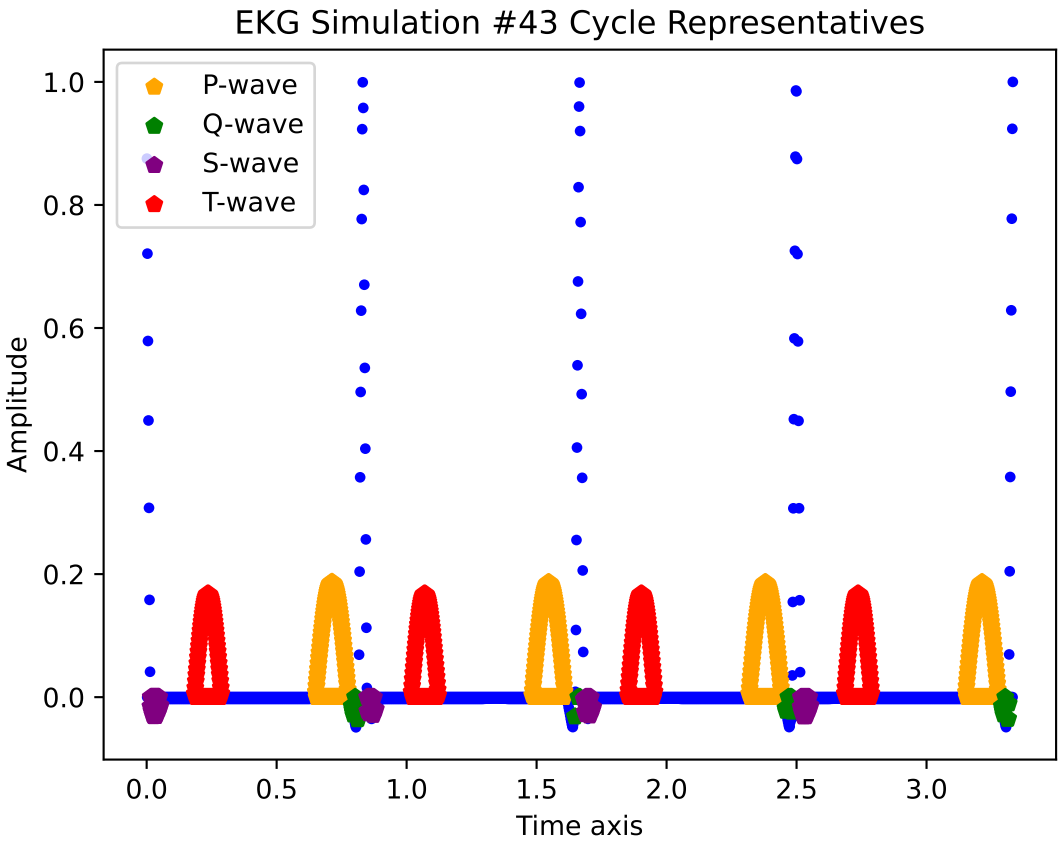
EKG Simulation #43 Cycle Representatives
P-wave
Q-wave
S-wave
T-wave
Amplitude
1.0
0.8
0.6
0.4
0.2
0.0
0.0
0.5
1.0
1.5
2.0
2.5
3.0
Time axis

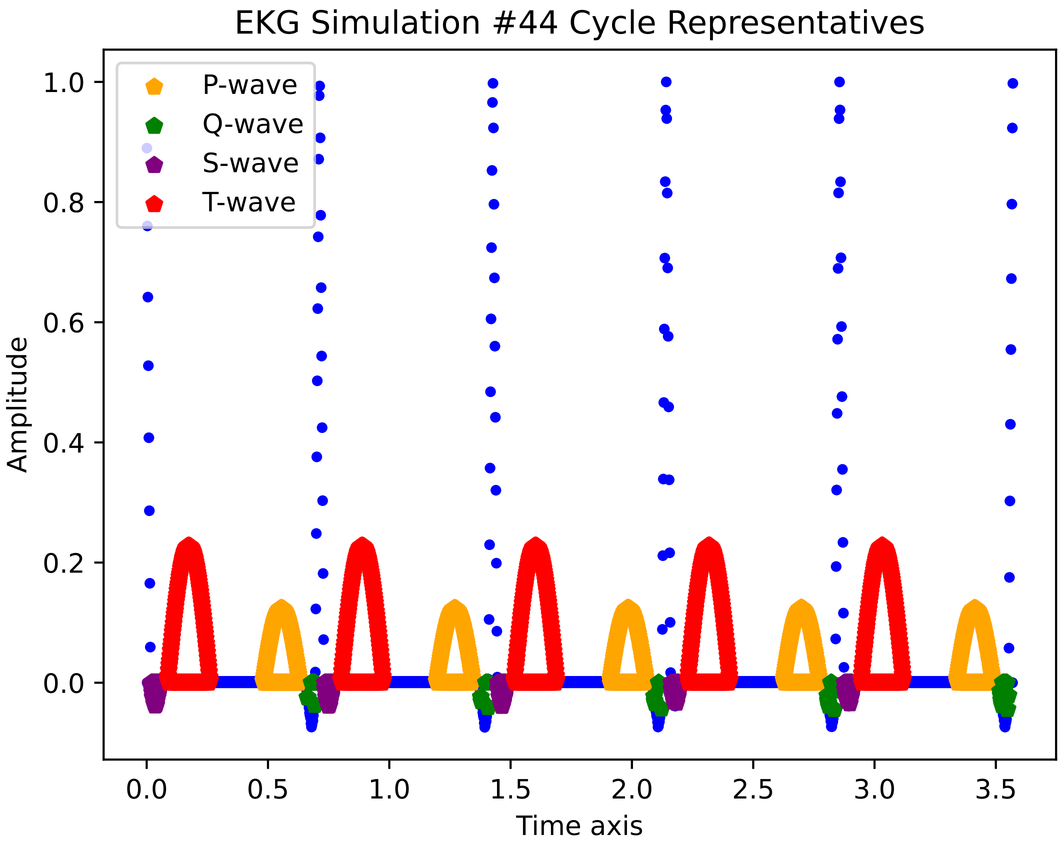
EKG Simulation #44 Cycle Representatives
P-wave
Q-wave
S-wave
T-wave
Amplitude
Time axis
1.0
0.8
0.6
0.4
0.2
0.0
0.0
0.5
1.0
1.5
2.0
2.5
3.0
3.5

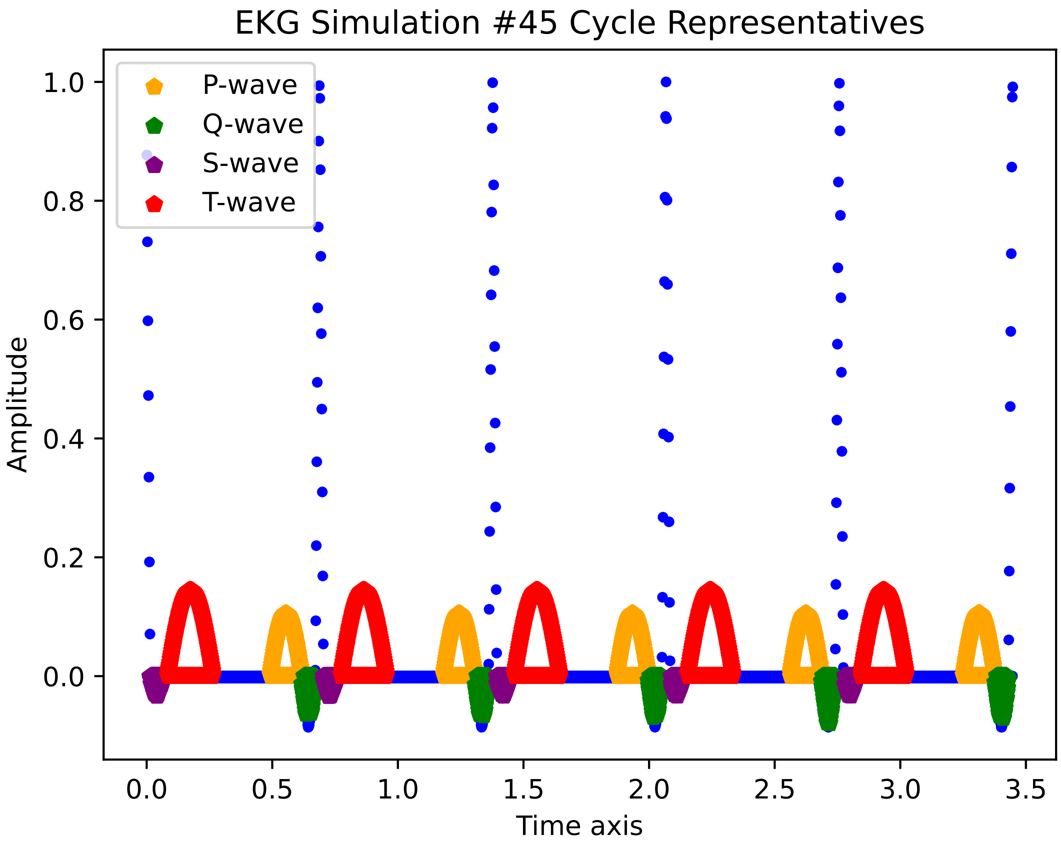

EKG Simulation #45 Cycle Representatives
P-wave
Q-wave
S-wave
T-wave
1.0
0.8
0.6
0.4
0.2
0.0
Amplitude
0.0
0.5
1.0
1.5
2.0
2.5
3.0
3.5
Time axis

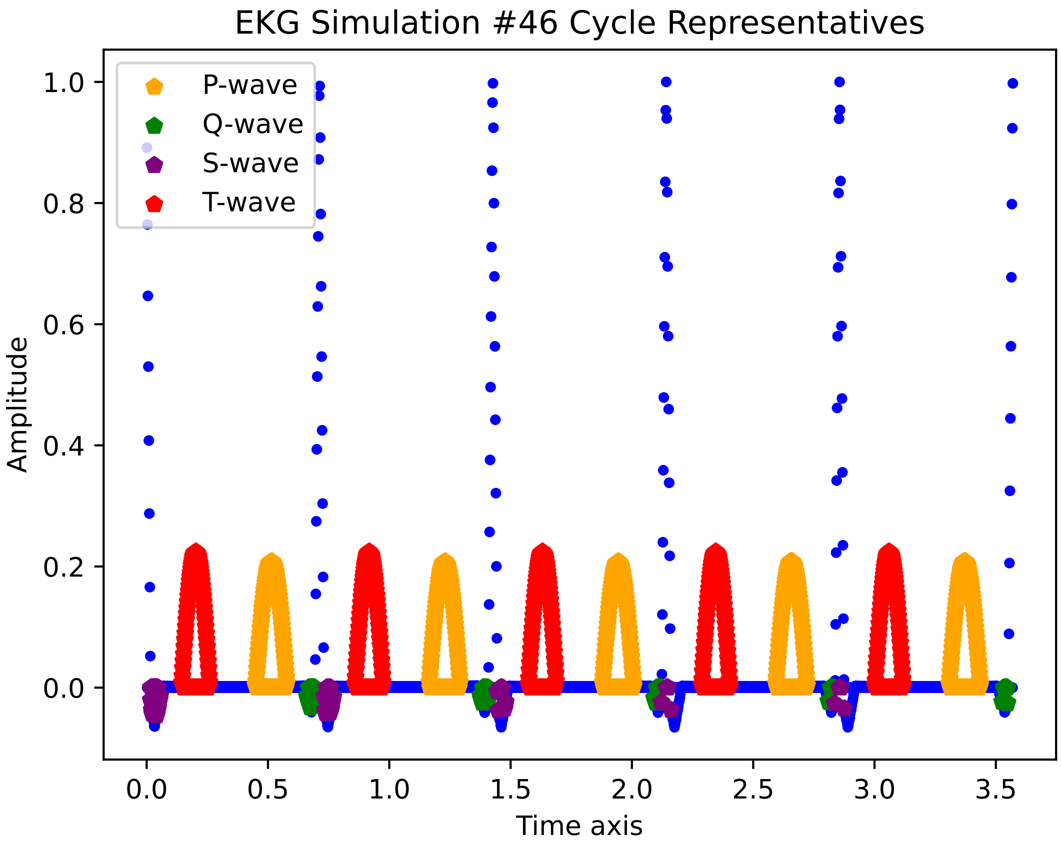
EKG Simulation #46 Cycle Representatives
P-wave
Q-wave
S-wave
T-wave
Amplitude
1.0
0.8
0.6
0.4
0.2
0.0
0.0
0.5
1.0
1.5
2.0
2.5
3.0
3.5
Time axis

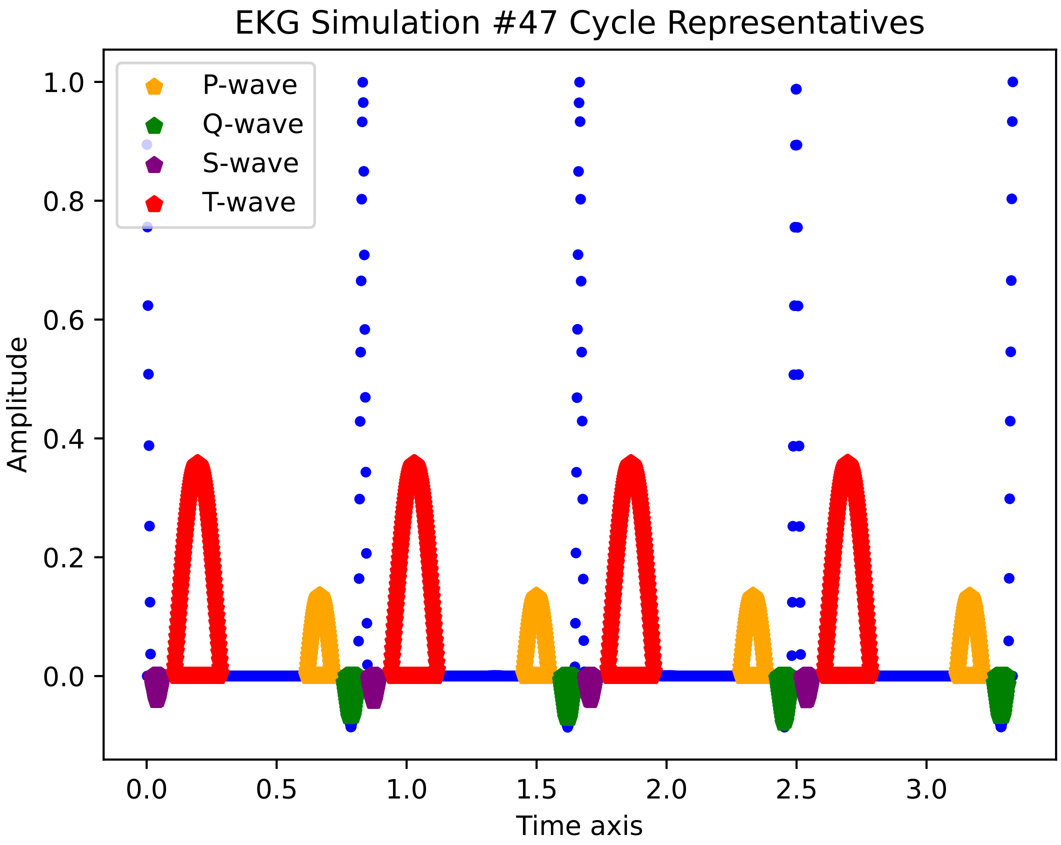
EKG Simulation #47 Cycle Representatives
P-wave
Q-wave
S-wave
T-wave
Amplitude
1.0
0.8
0.6
0.4
0.2
0.0
0.0
0.5
1.0
1.5
2.0
2.5
3.0
Time axis

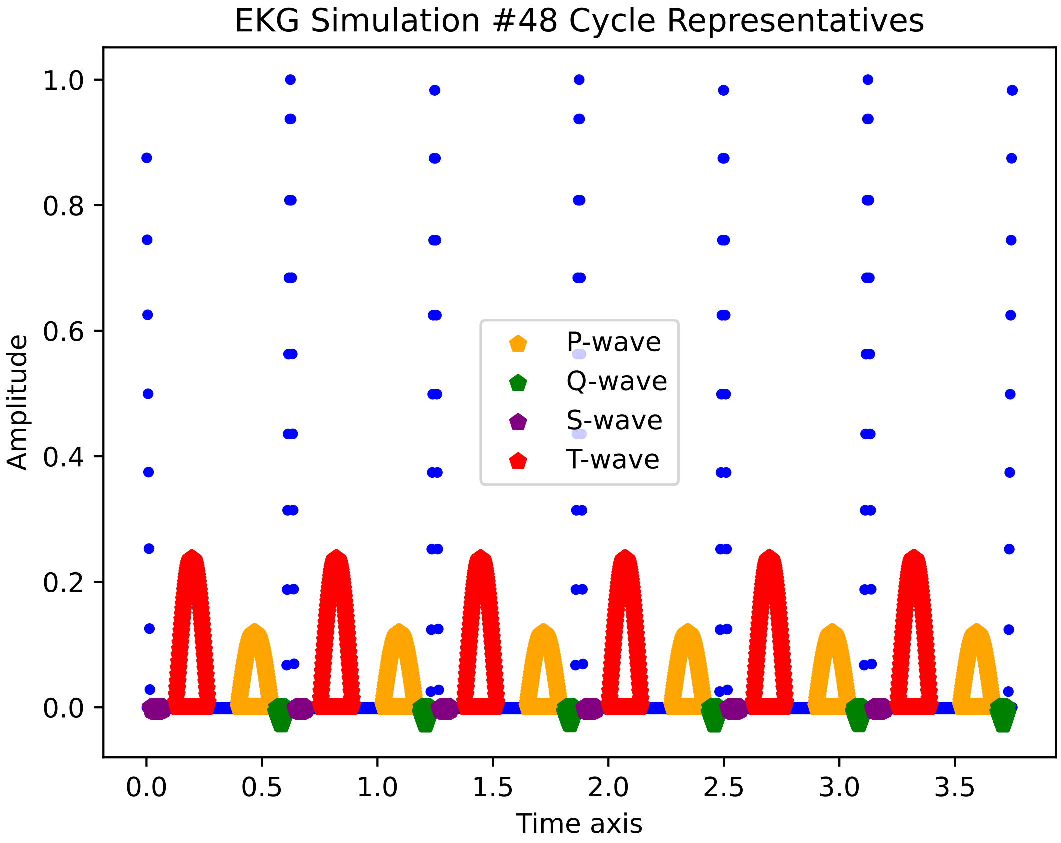

EKG Simulation #48 Cycle Representatives
Amplitude
Time axis
P-wave
Q-wave
S-wave
T-wave
1.0
0.8
0.6
0.4
0.2
0.0
0.0
0.5
1.0
1.5
2.0
2.5
3.0
3.5

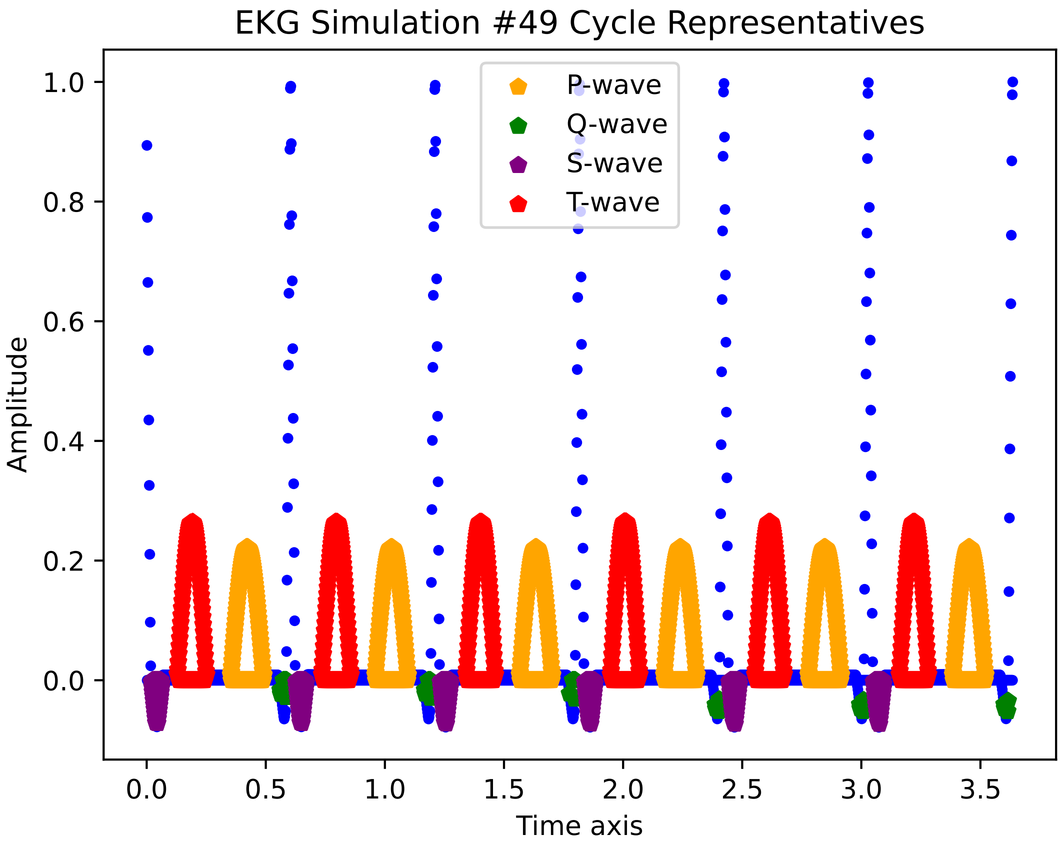
EKG Simulation #49 Cycle Representatives
P-wave
Q-wave
S-wave
T-wave
Amplitude
Time axis
1.0
0.8
0.6
0.4
0.2
0.0
0.0
0.5
1.0
1.5
2.0
2.5
3.0
3.5

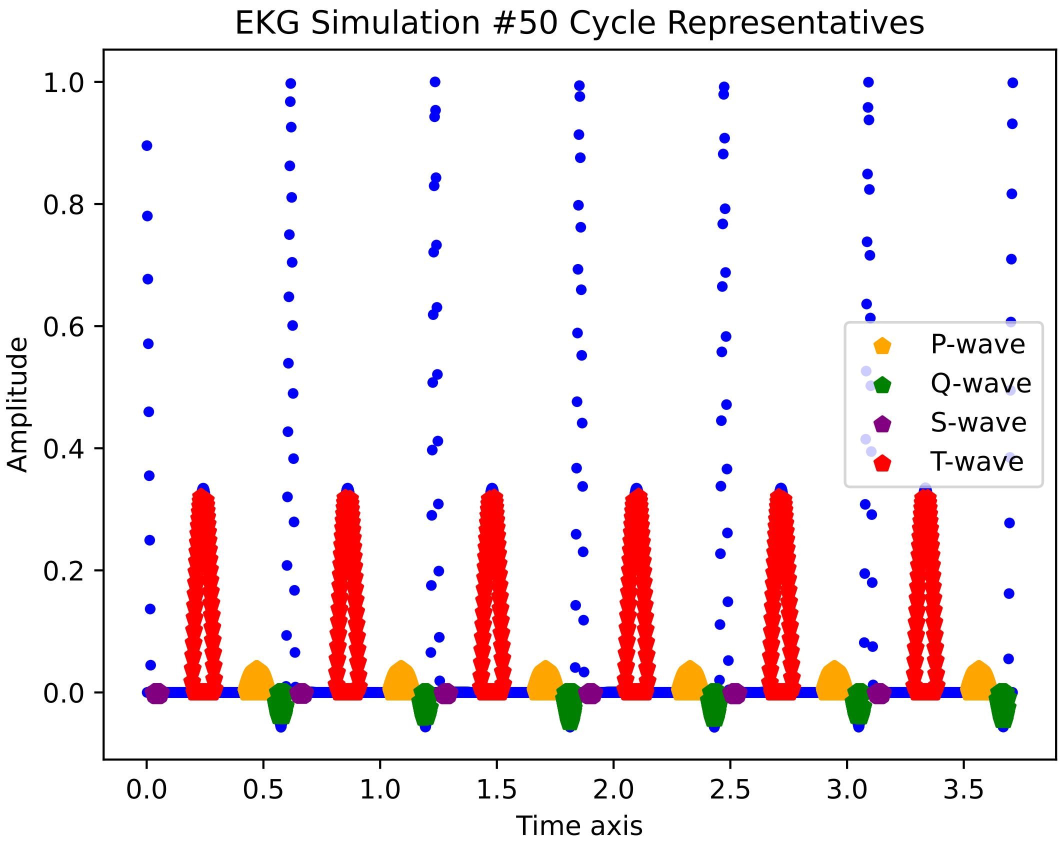

EKG Simulation #50 Cycle Representatives
Amplitude
Time axis
1.0
0.8
0.6
0.4
0.2
0.0
0.0
0.5
1.0
1.5
2.0
2.5
3.0
3.5
P-wave
Q-wave
S-wave
T-wave

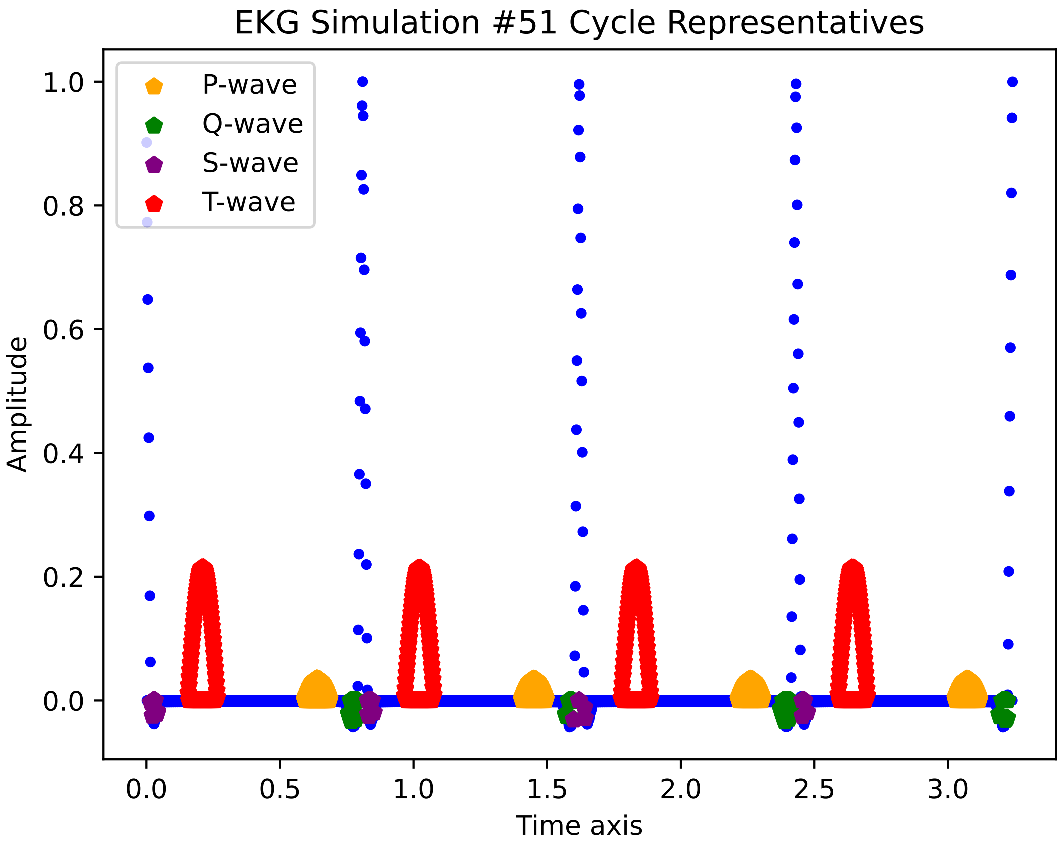
EKG Simulation #51 Cycle Representatives
P-wave
Q-wave
S-wave
T-wave
Amplitude
Time axis
0.0
0.2
0.4
0.6
0.8
1.0
0.0
0.5
1.0
1.5
2.0
2.5
3.0

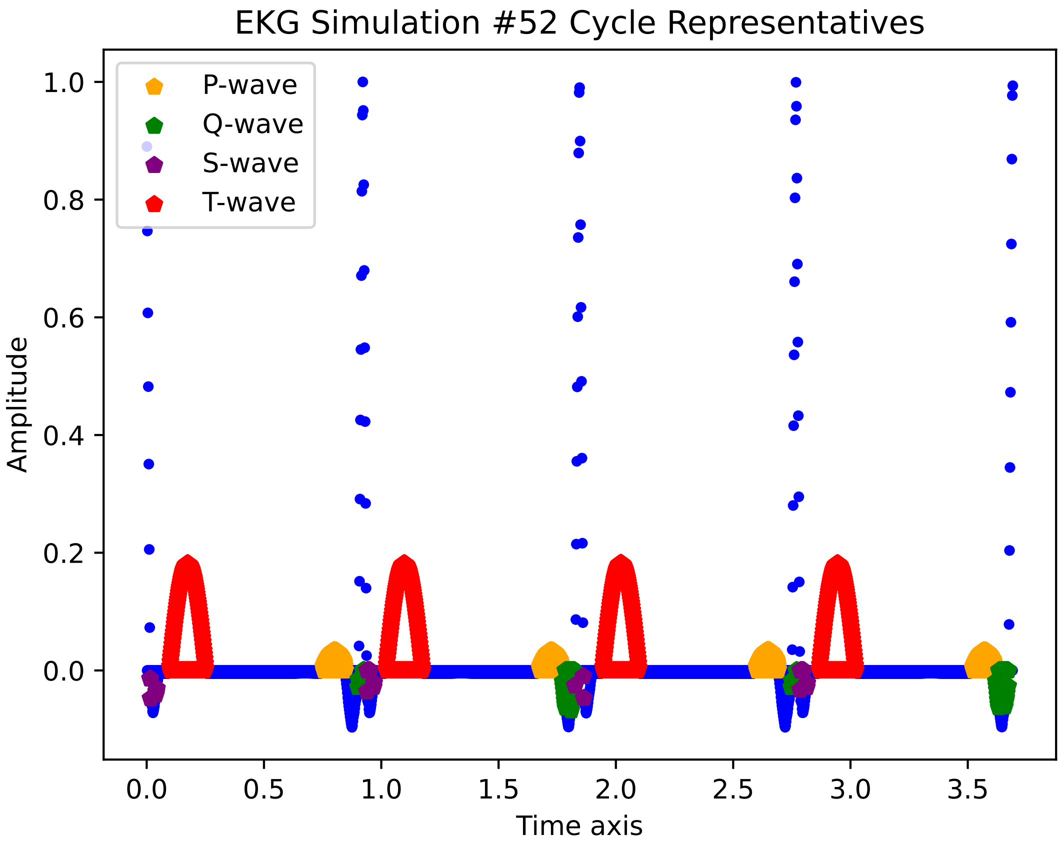
EKG Simulation #52 Cycle Representatives
P-wave
Q-wave
S-wave
T-wave
Amplitude
1.0
0.8
0.6
0.4
0.2
0.0
0.0
0.5
1.0
1.5
2.0
2.5
3.0
3.5
Time axis

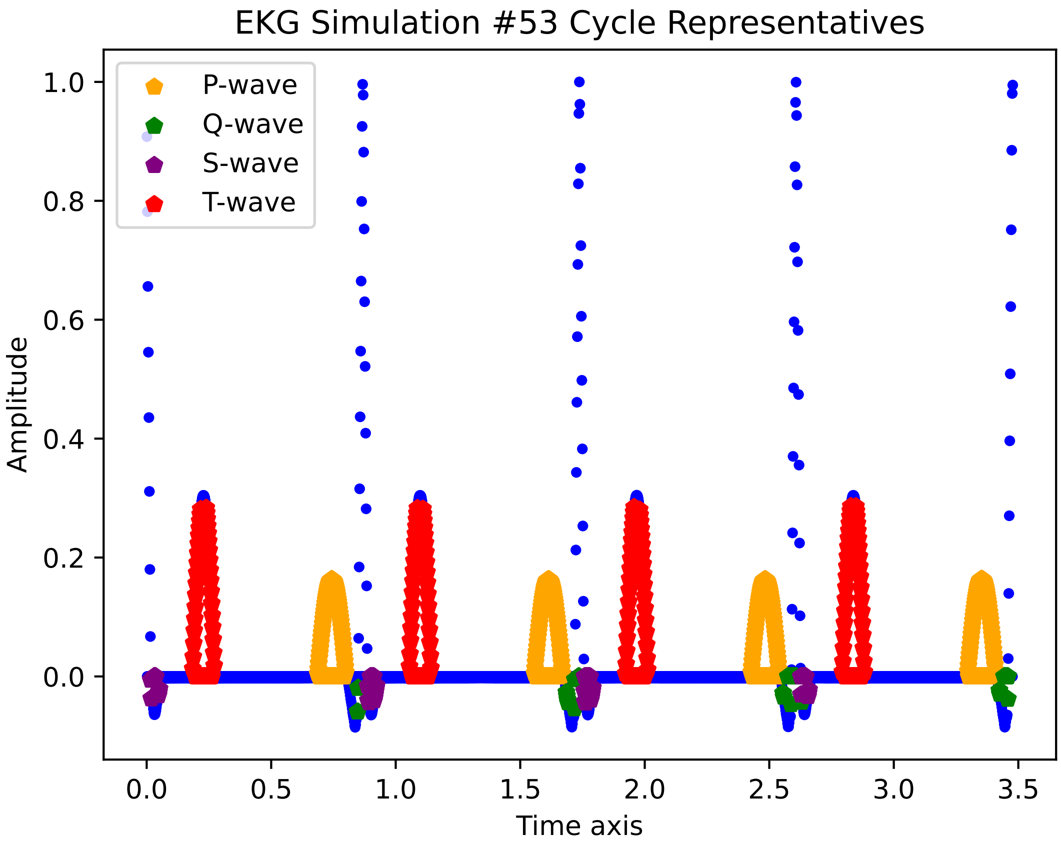
EKG Simulation #53 Cycle Representatives
P-wave
Q-wave
S-wave
T-wave
Amplitude
1.0
0.8
0.6
0.4
0.2
0.0
0.0
0.5
1.0
1.5
2.0
2.5
3.0
3.5
Time axis

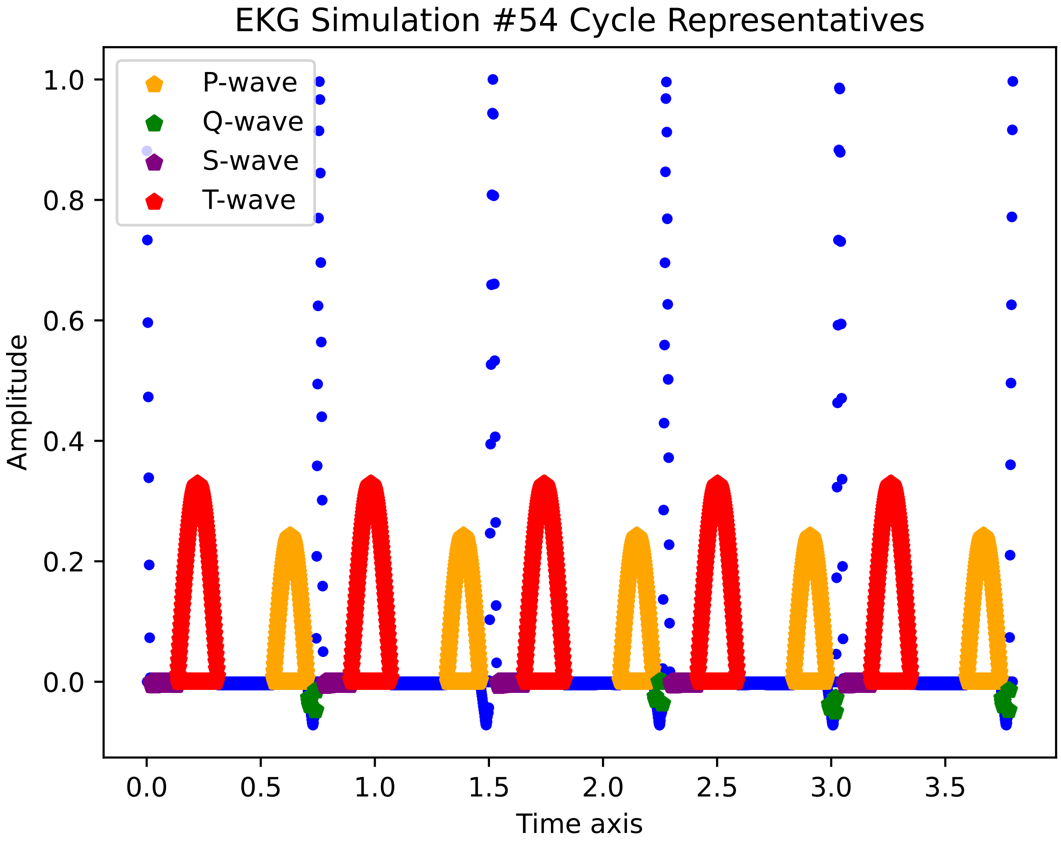

EKG Simulation #54 Cycle Representatives
P-wave
Q-wave
S-wave
T-wave
Amplitude
Time axis
1.0
0.8
0.6
0.4
0.2
0.0
0.0
0.5
1.0
1.5
2.0
2.5
3.0
3.5

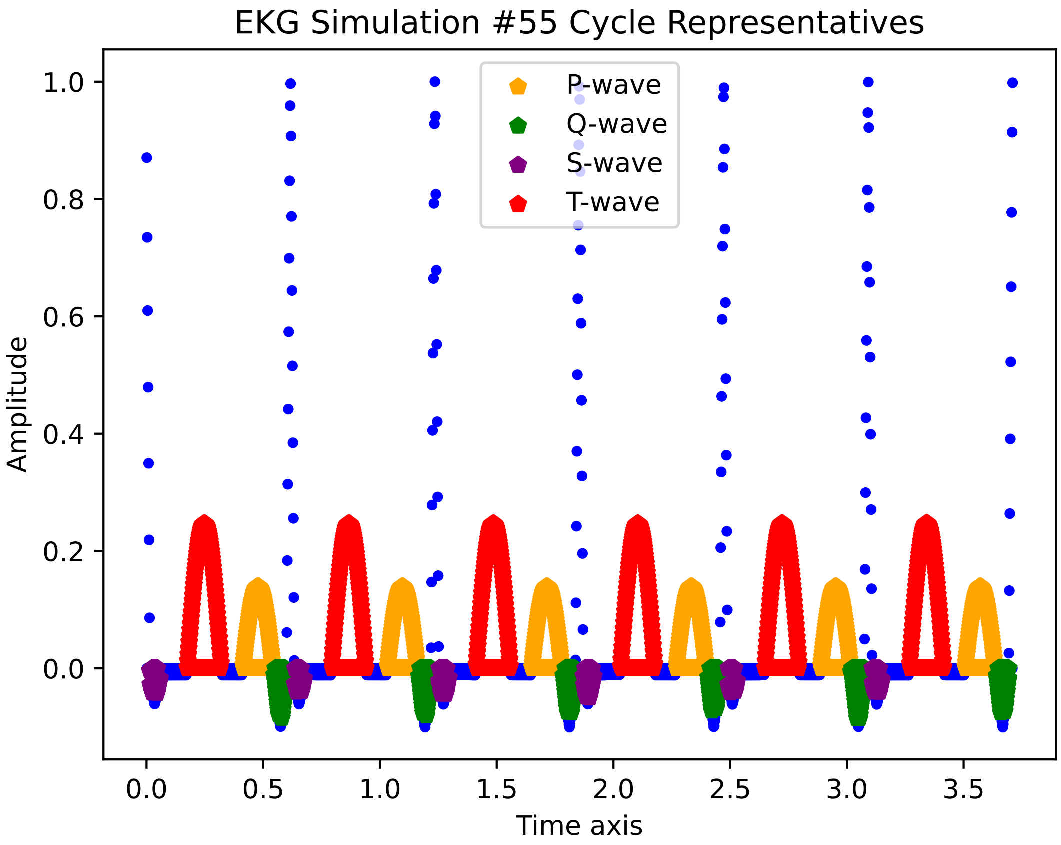

EKG Simulation #55 Cycle Representatives
P-wave
Q-wave
S-wave
T-wave
Amplitude
Time axis
1.0
0.8
0.6
0.4
0.2
0.0
0.0
0.5
1.0
1.5
2.0
2.5
3.0
3.5

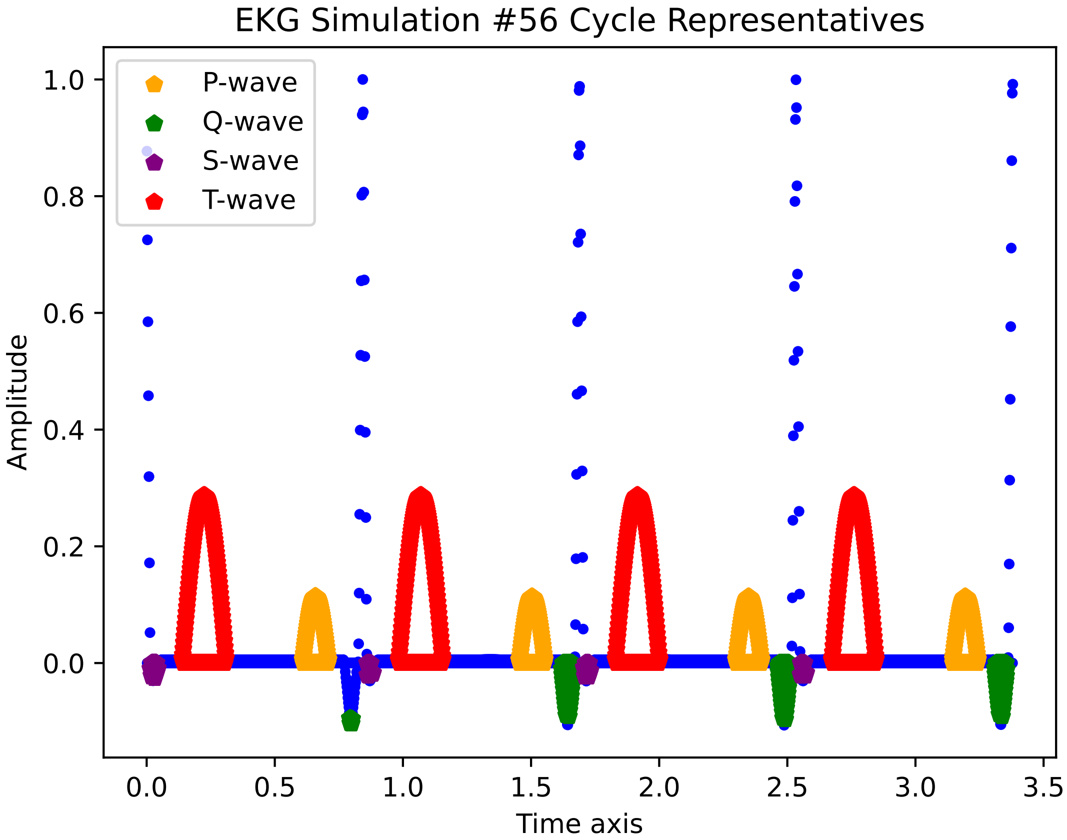
EKG Simulation #56 Cycle Representatives
P-wave
Q-wave
S-wave
T-wave
Amplitude
Time axis
1.0
0.8
0.6
0.4
0.2
0.0
0.0
0.5
1.0
1.5
2.0
2.5
3.0
3.5

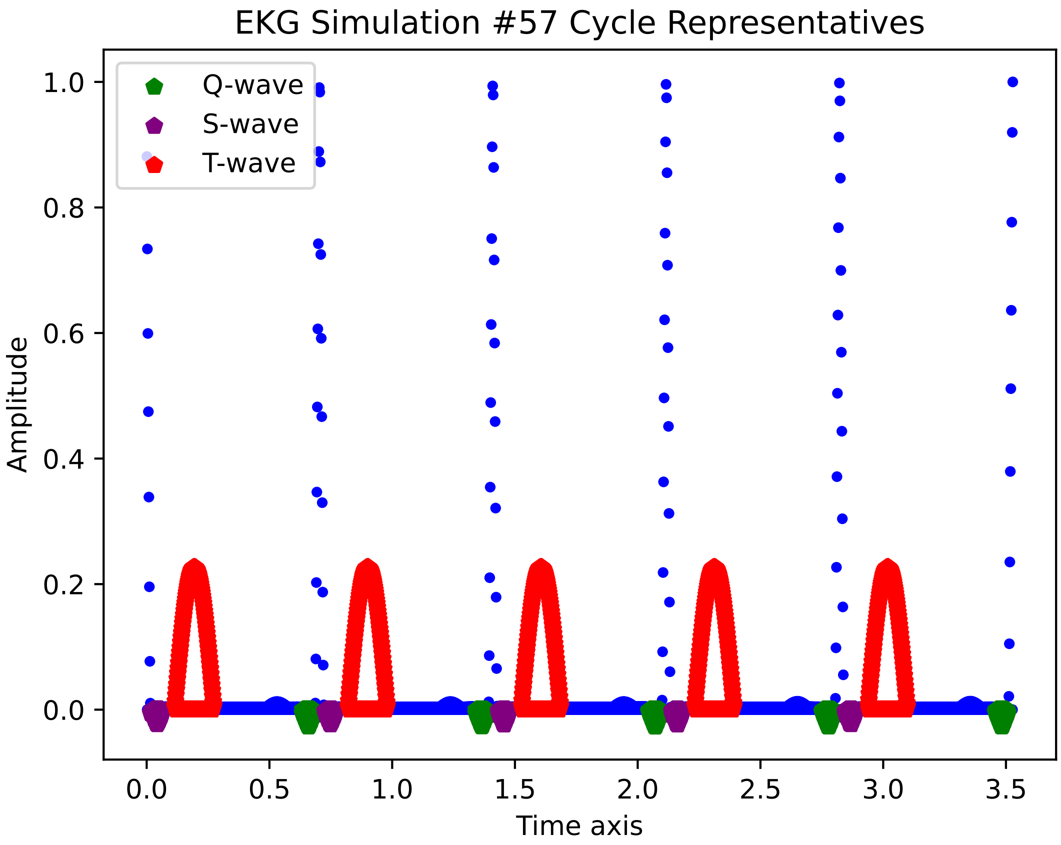

EKG Simulation #57 Cycle Representatives
Q-wave
S-wave
T-wave
Amplitude
Time axis
1.0
0.8
0.6
0.4
0.2
0.0
0.0
0.5
1.0
1.5
2.0
2.5
3.0
3.5

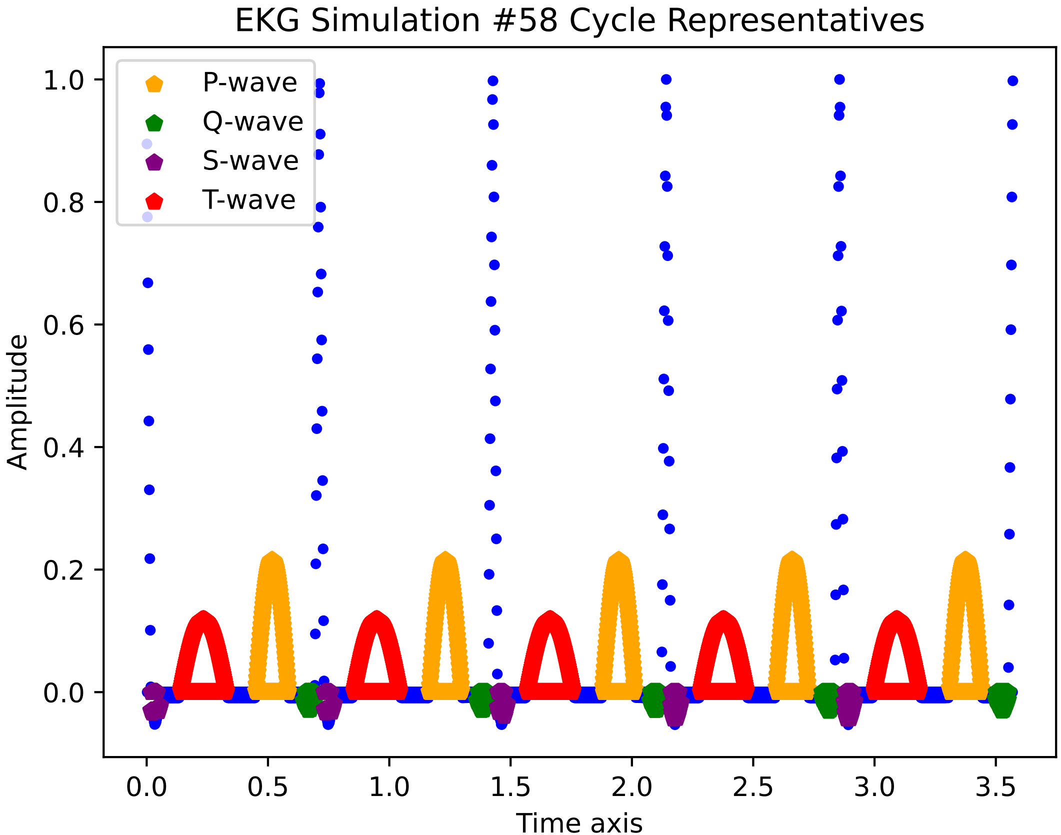

EKG Simulation #58 Cycle Representatives
P-wave
Q-wave
S-wave
T-wave
Amplitude
Time axis
0.0
0.2
0.4
0.6
0.8
1.0
0.0
0.5
1.0
1.5
2.0
2.5
3.0
3.5

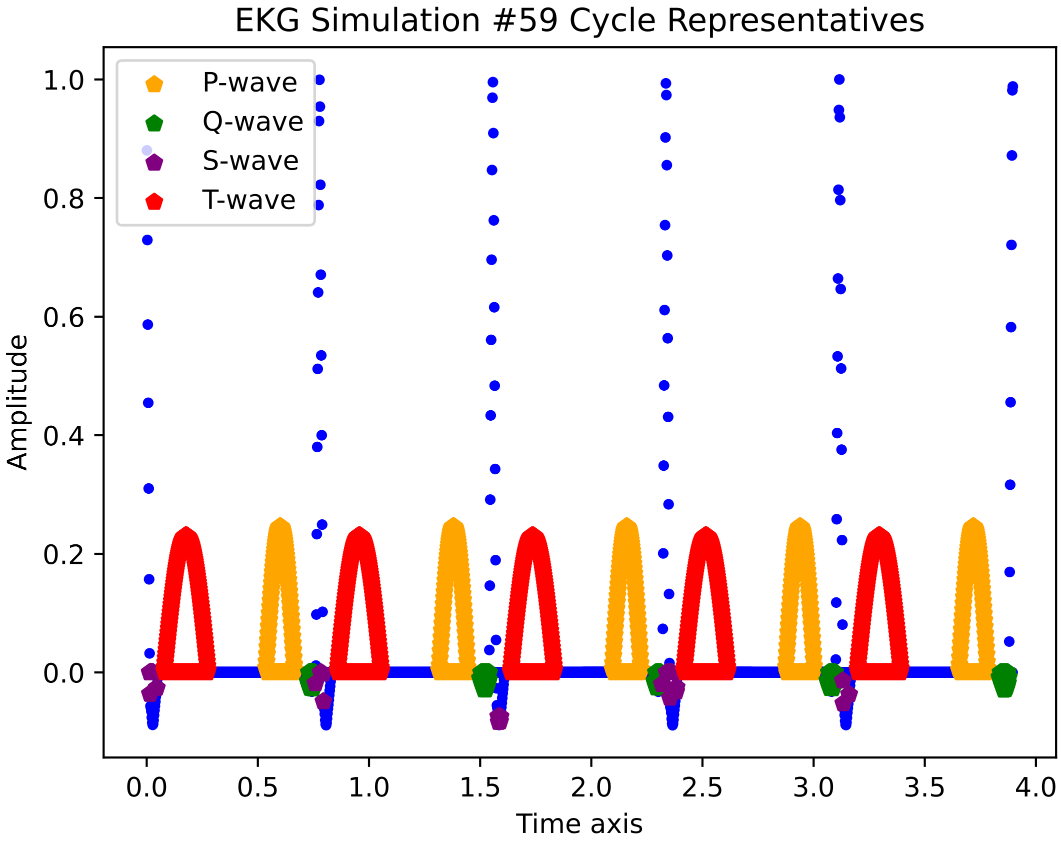

EKG Simulation #59 Cycle Representatives
P-wave
Q-wave
S-wave
T-wave
Amplitude
Time axis
1.0
0.8
0.6
0.4
0.2
0.0
0.0
0.5
1.0
1.5
2.0
2.5
3.0
3.5
4.0

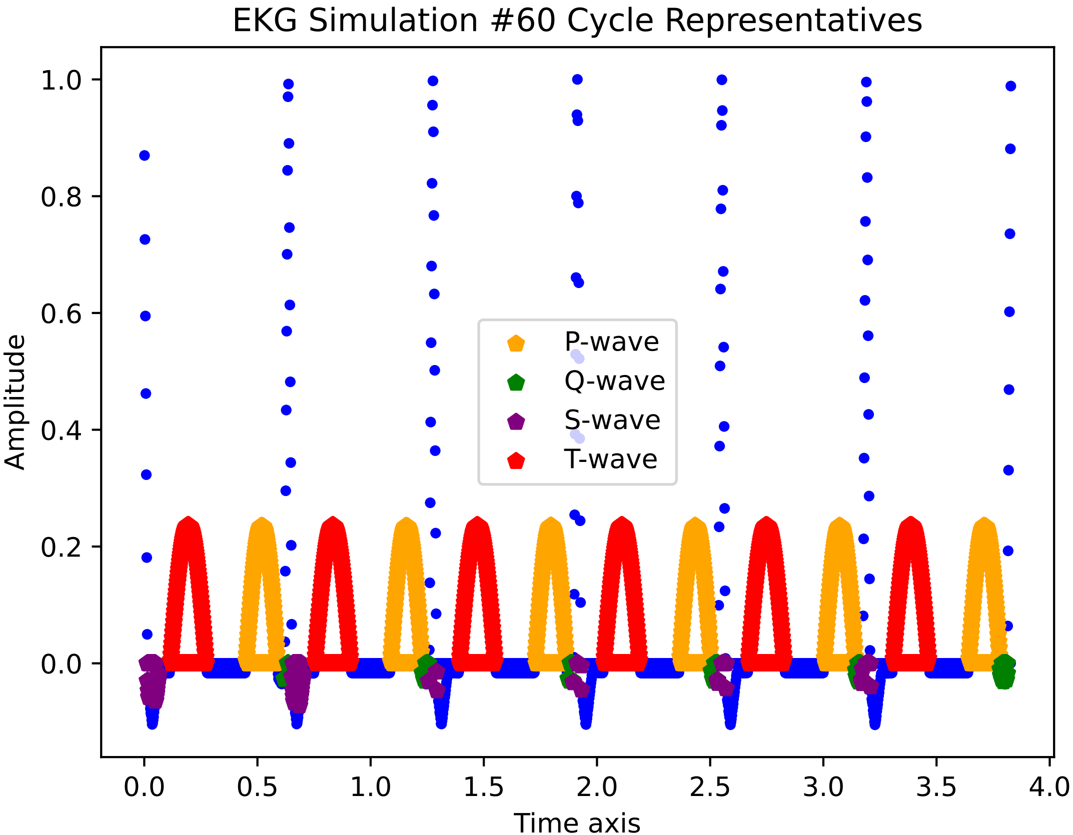

EKG Simulation #60 Cycle Representatives
Amplitude
Time axis
P-wave
Q-wave
S-wave
T-wave
1.0
0.8
0.6
0.4
0.2
0.0
0.0
0.5
1.0
1.5
2.0
2.5
3.0
3.5
4.0

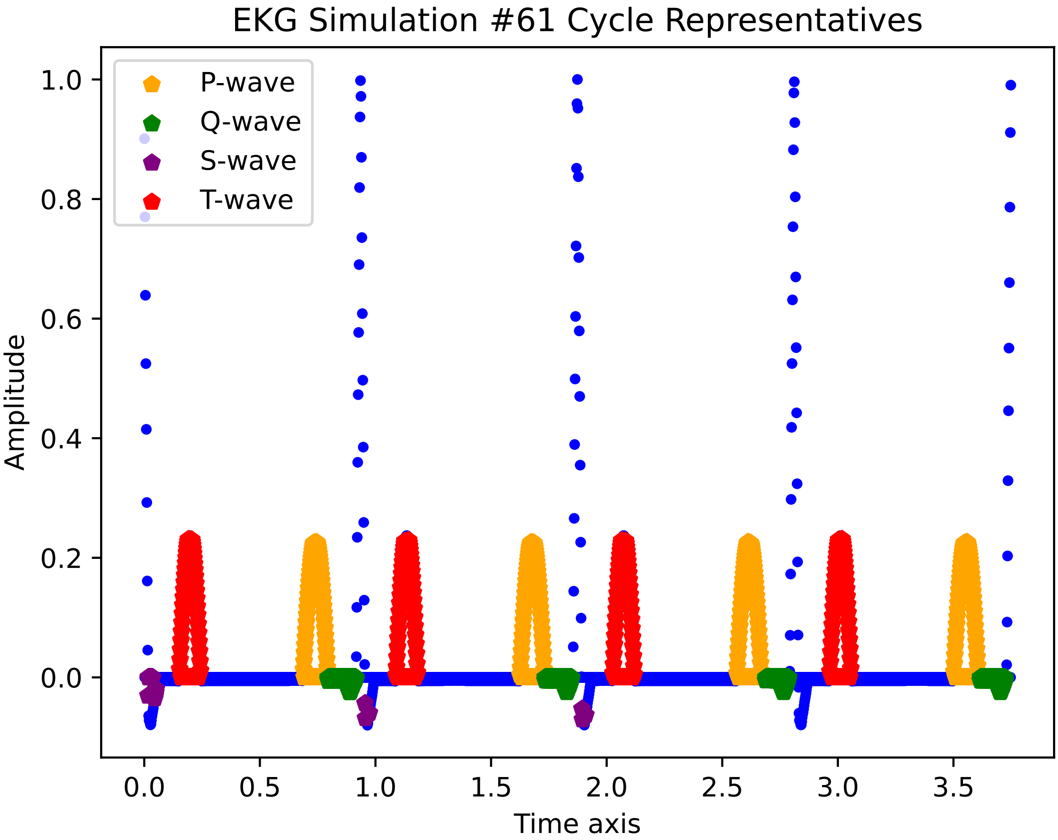

EKG Simulation #61 Cycle Representatives
P-wave
Q-wave
S-wave
T-wave
1.0
0.8
0.6
0.4
0.2
0.0
Amplitude
0.0
0.5
1.0
1.5
2.0
2.5
3.0
3.5
Time axis

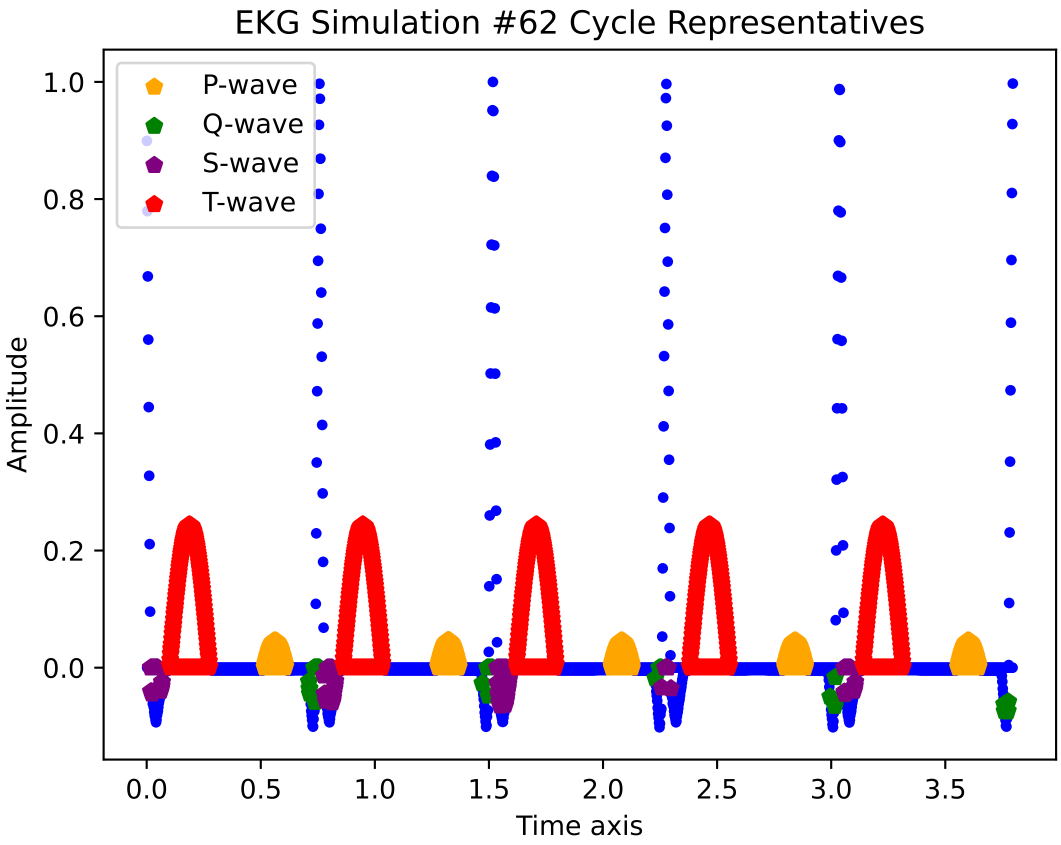
EKG Simulation #62 Cycle Representatives
P-wave
Q-wave
S-wave
T-wave
1.0
0.8
0.6
0.4
0.2
0.0
Amplitude
0.0
0.5
1.0
1.5
2.0
2.5
3.0
3.5
Time axis

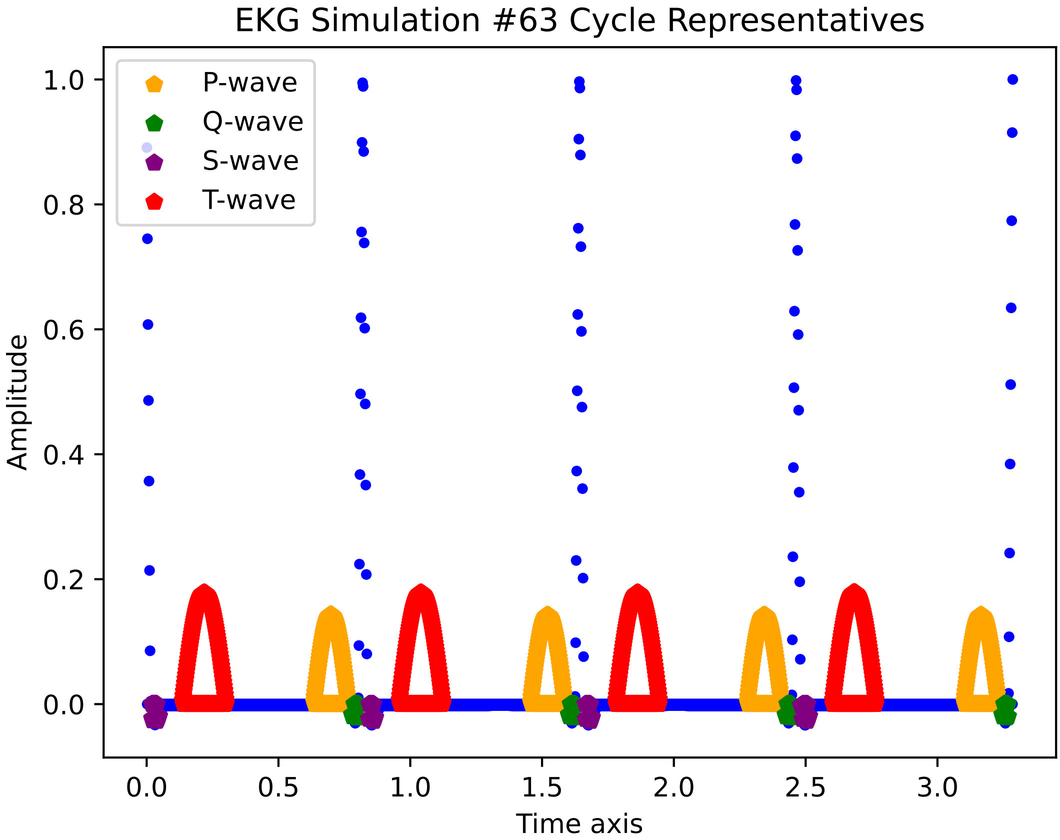

EKG Simulation #63 Cycle Representatives
P-wave
Q-wave
S-wave
T-wave
Amplitude
Time axis
1.0
0.8
0.6
0.4
0.2
0.0
0.0
0.5
1.0
1.5
2.0
2.5
3.0

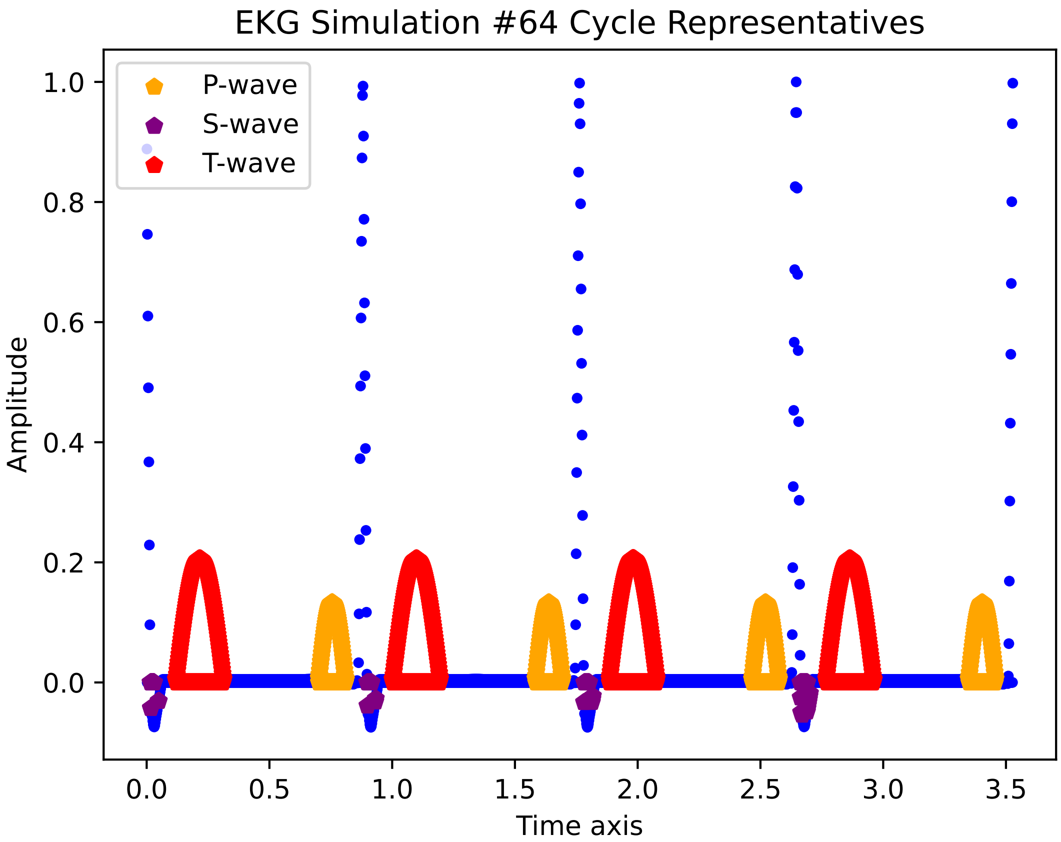

EKG Simulation #64 Cycle Representatives
P-wave
S-wave
T-wave
Amplitude
1.0
0.8
0.6
0.4
0.2
0.0
0.0
0.5
1.0
1.5
2.0
2.5
3.0
3.5
Time axis

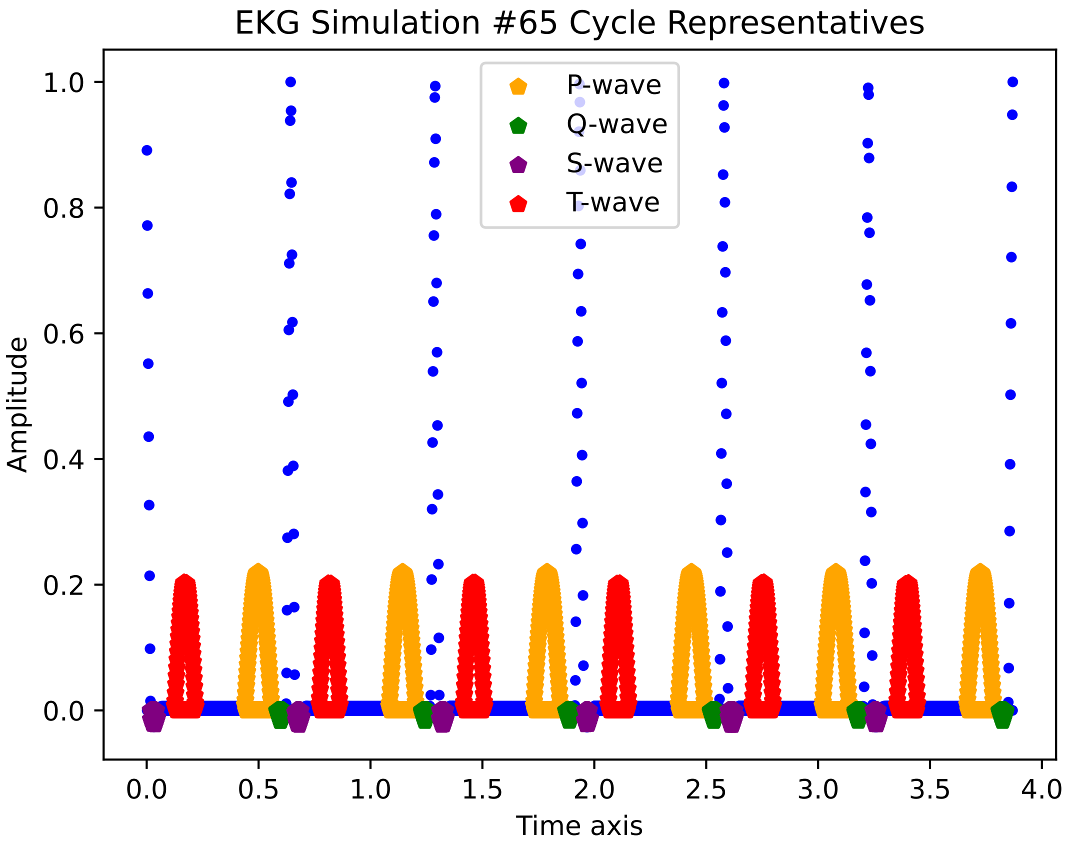

EKG Simulation #65 Cycle Representatives
P-wave
Q-wave
S-wave
T-wave
Amplitude
Time axis
0.0
0.2
0.4
0.6
0.8
1.0
0.0
0.5
1.0
1.5
2.0
2.5
3.0
3.5
4.0

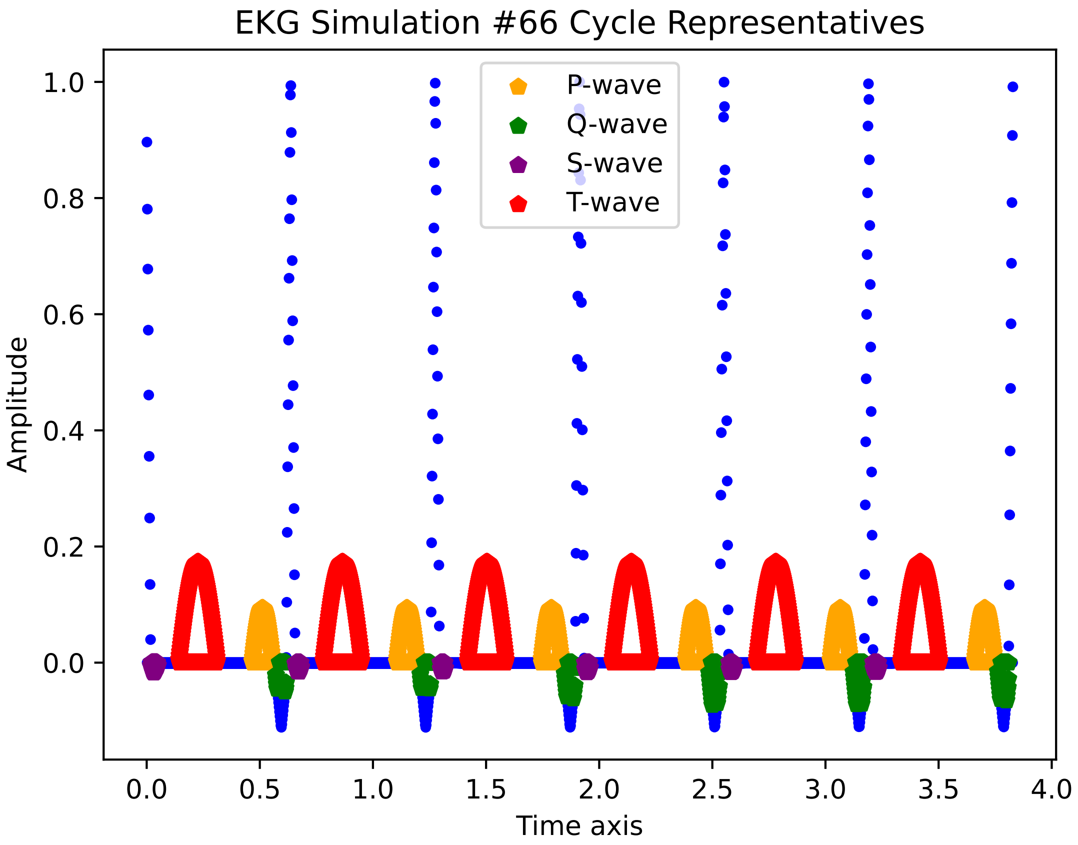
EKG Simulation #66 Cycle Representatives
P-wave
Q-wave
S-wave
T-wave
Amplitude
1.0
0.8
0.6
0.4
0.2
0.0
0.0
0.5
1.0
1.5
2.0
2.5
3.0
3.5
4.0
Time axis

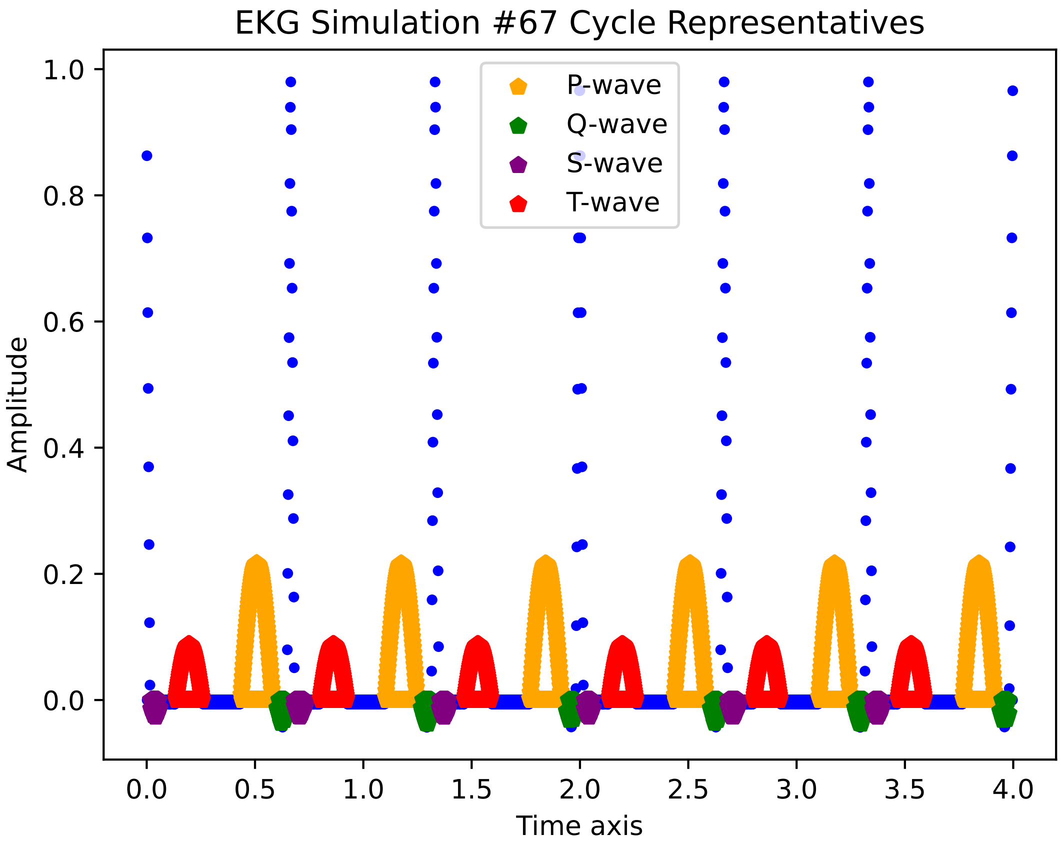
EKG Simulation #67 Cycle Representatives
P-wave
Q-wave
S-wave
T-wave
Amplitude
Time axis
1.0
0.8
0.6
0.4
0.2
0.0
0.0
0.5
1.0
1.5
2.0
2.5
3.0
3.5
4.0

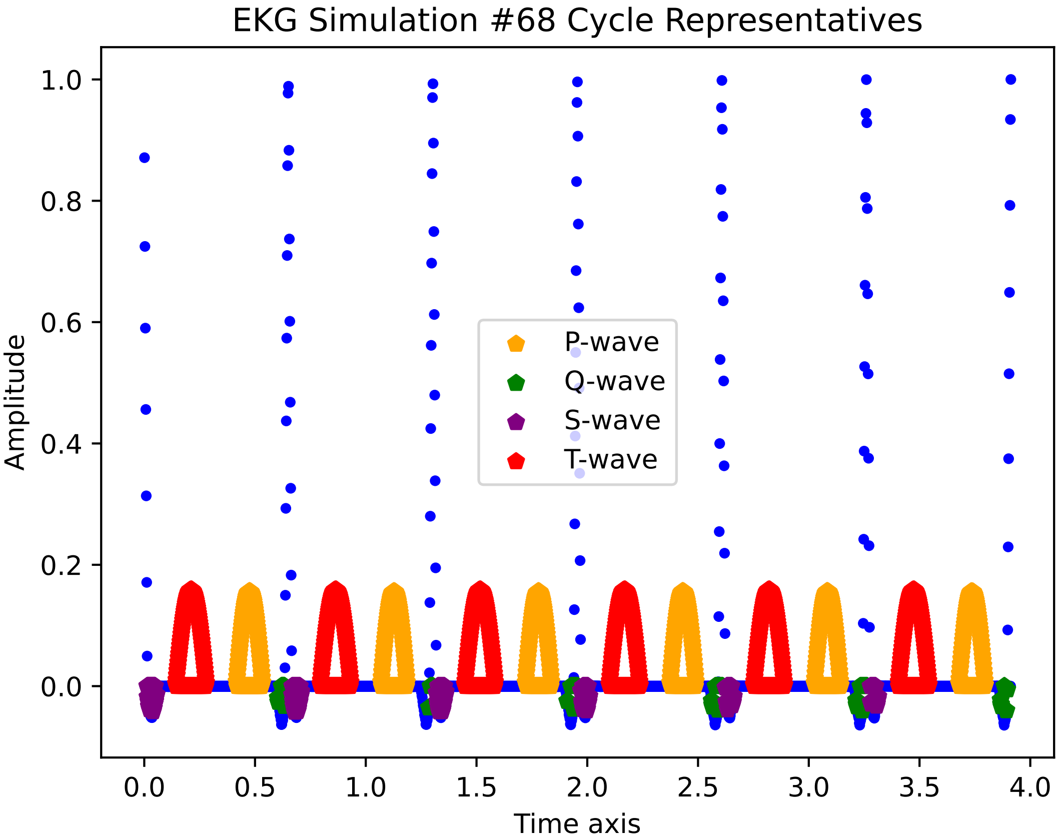

EKG Simulation #68 Cycle Representatives
Amplitude
1.0
0.8
0.6
0.4
0.2
0.0
0.0
0.5
1.0
1.5
2.0
2.5
3.0
3.5
4.0
Time axis
P-wave
Q-wave
S-wave
T-wave

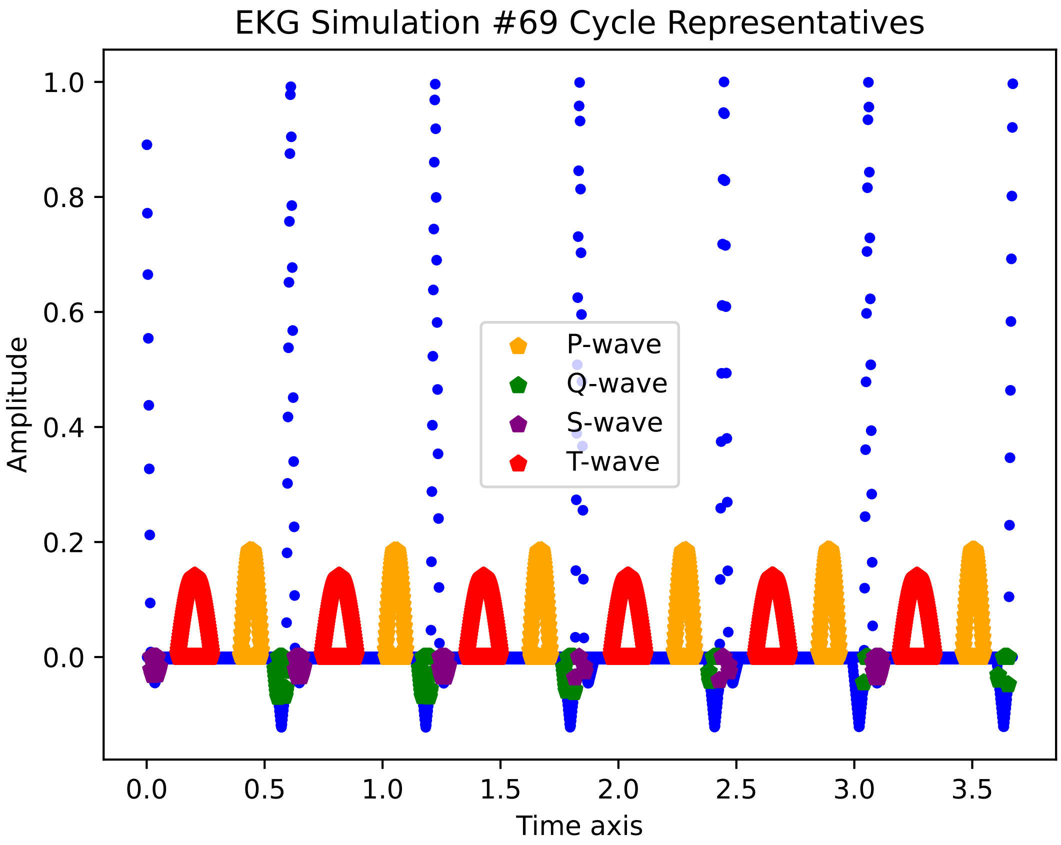

EKG Simulation #69 Cycle Representatives
Amplitude
Time axis
1.0
0.8
0.6
0.4
0.2
0.0
0.0
0.5
1.0
1.5
2.0
2.5
3.0
3.5
P-wave
Q-wave
S-wave
T-wave

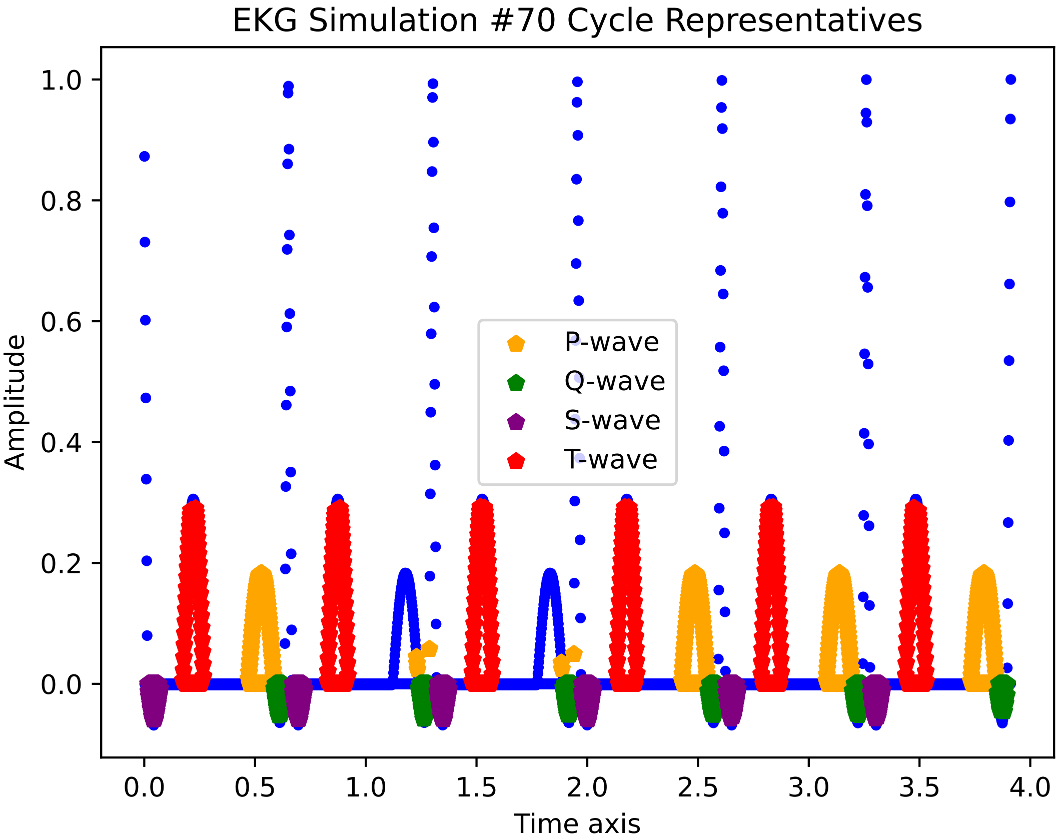
EKG Simulation #70 Cycle Representatives
P-wave
Q-wave
S-wave
T-wave
Amplitude
Time axis
1.0
0.8
0.6
0.4
0.2
0.0
0.0
0.5
1.0
1.5
2.0
2.5
3.0
3.5
4.0

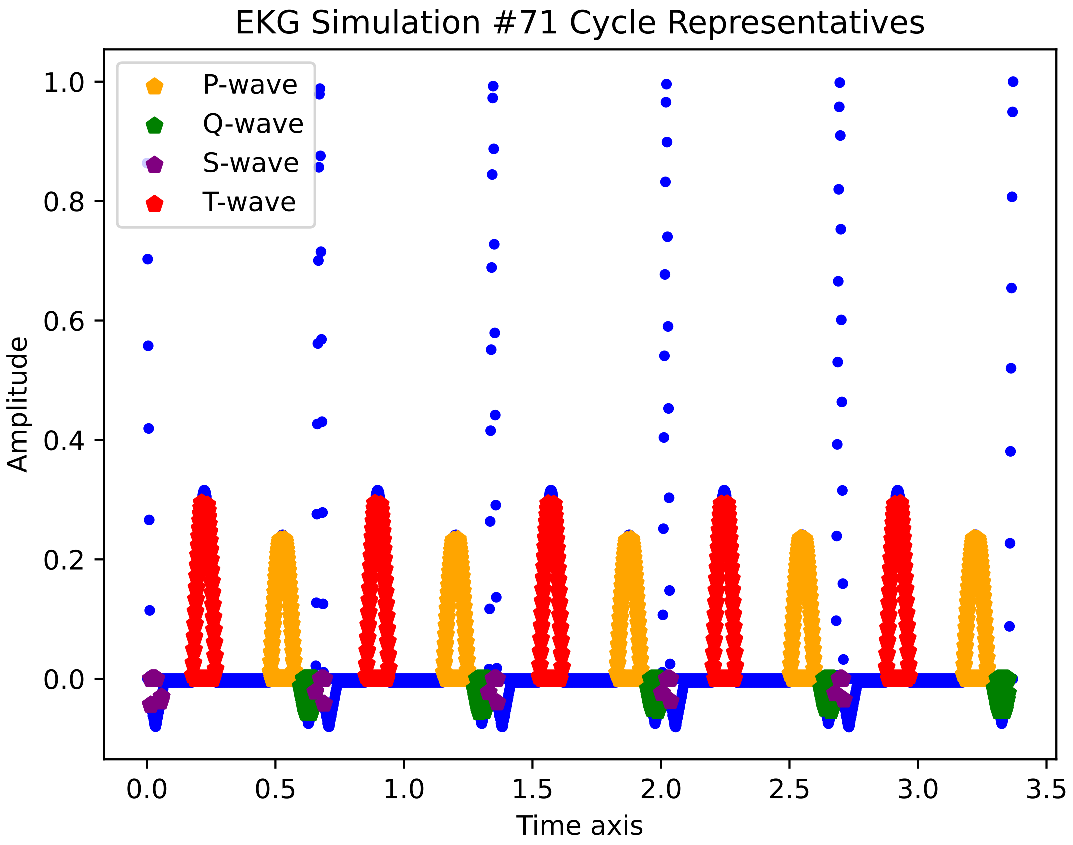

EKG Simulation #71 Cycle Representatives
P-wave
Q-wave
S-wave
T-wave
1.0
0.8
0.6
0.4
0.2
0.0
Amplitude
0.0
0.5
1.0
1.5
2.0
2.5
3.0
3.5
Time axis

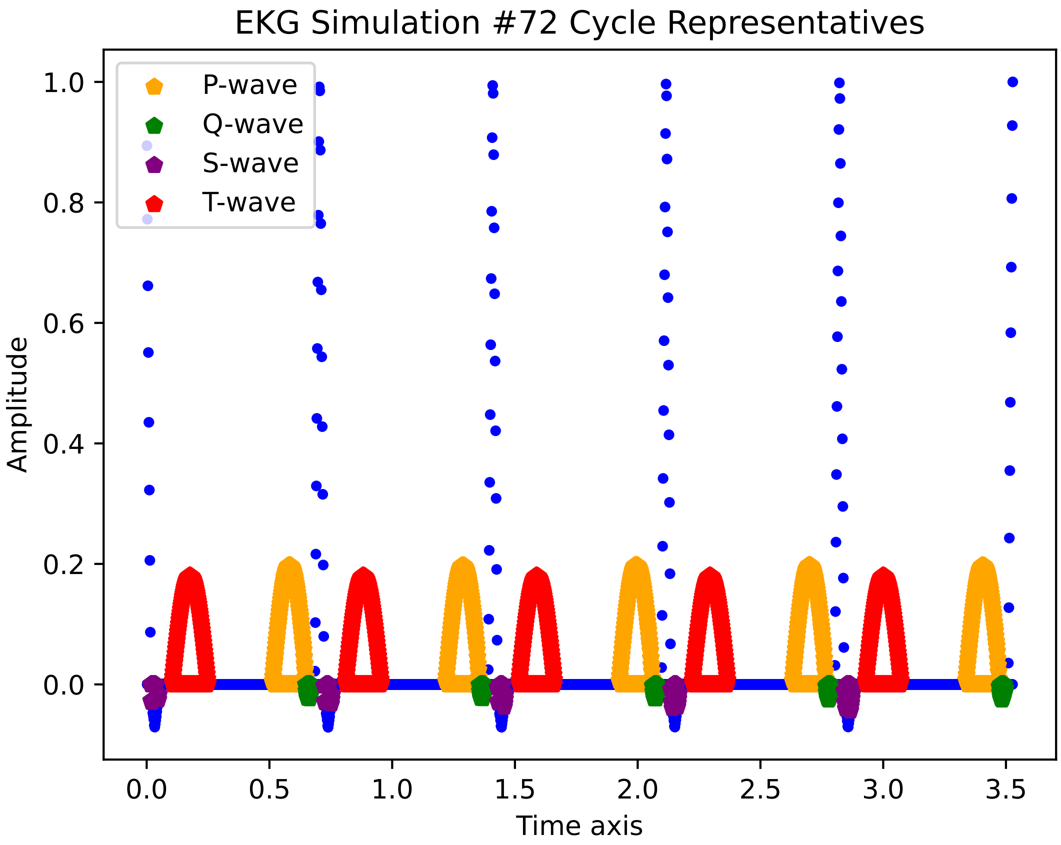

EKG Simulation #72 Cycle Representatives
P-wave
Q-wave
S-wave
T-wave
1.0
0.8
0.6
0.4
0.2
0.0
Amplitude
0.0
0.5
1.0
1.5
2.0
2.5
3.0
3.5
Time axis

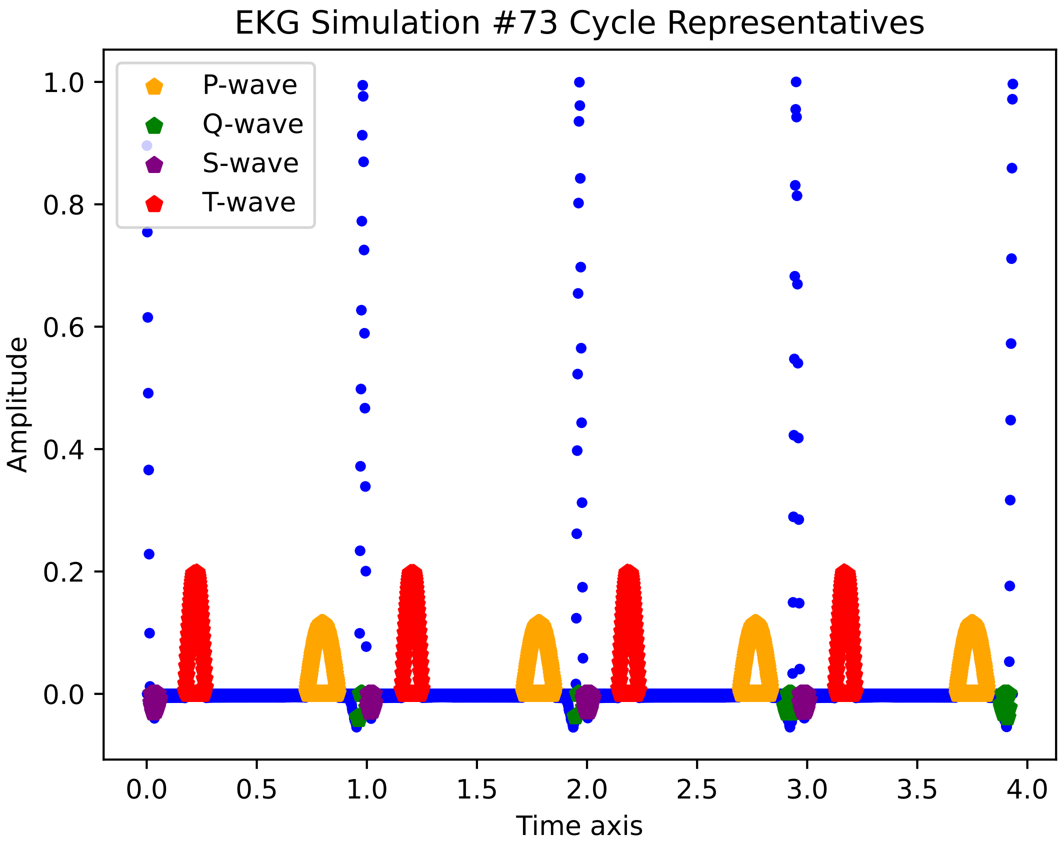

EKG Simulation #73 Cycle Representatives
P-wave
Q-wave
S-wave
T-wave
Amplitude
Time axis
1.0
0.8
0.6
0.4
0.2
0.0
0.0
0.5
1.0
1.5
2.0
2.5
3.0
3.5
4.0

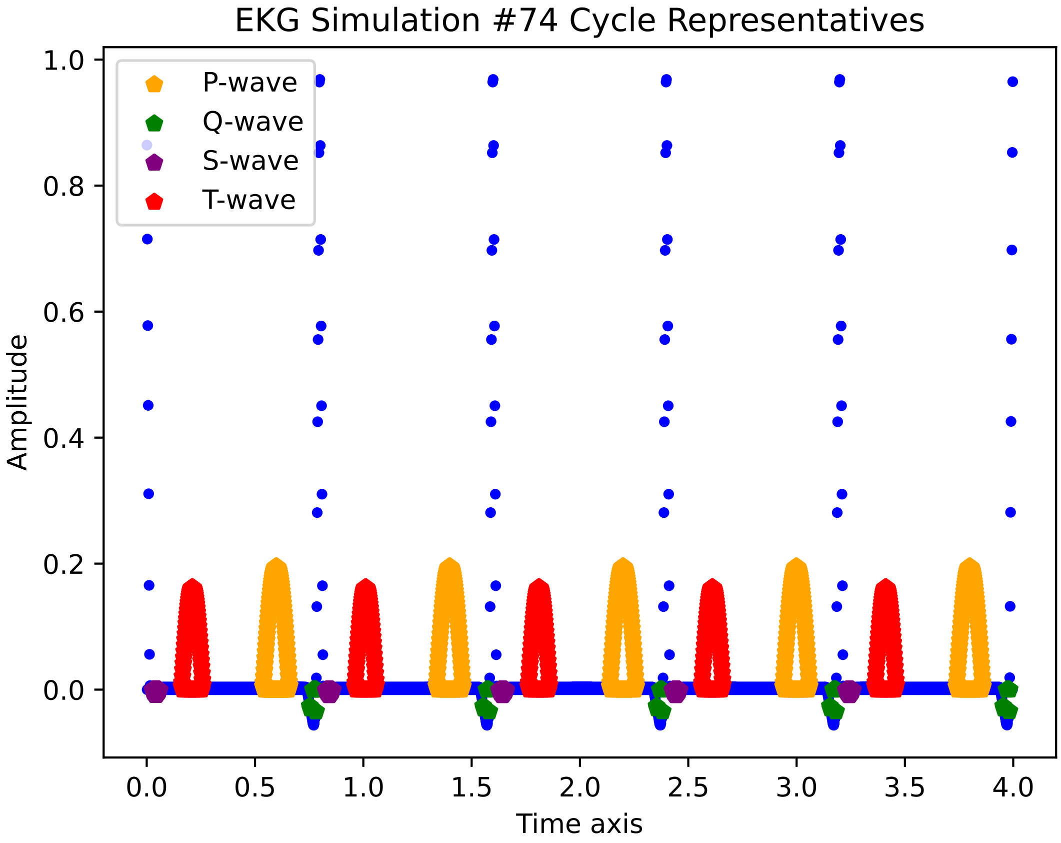

EKG Simulation #74 Cycle Representatives
P-wave
Q-wave
S-wave
T-wave
Amplitude
Time axis
1.0
0.8
0.6
0.4
0.2
0.0
0.0
0.5
1.0
1.5
2.0
2.5
3.0
3.5
4.0

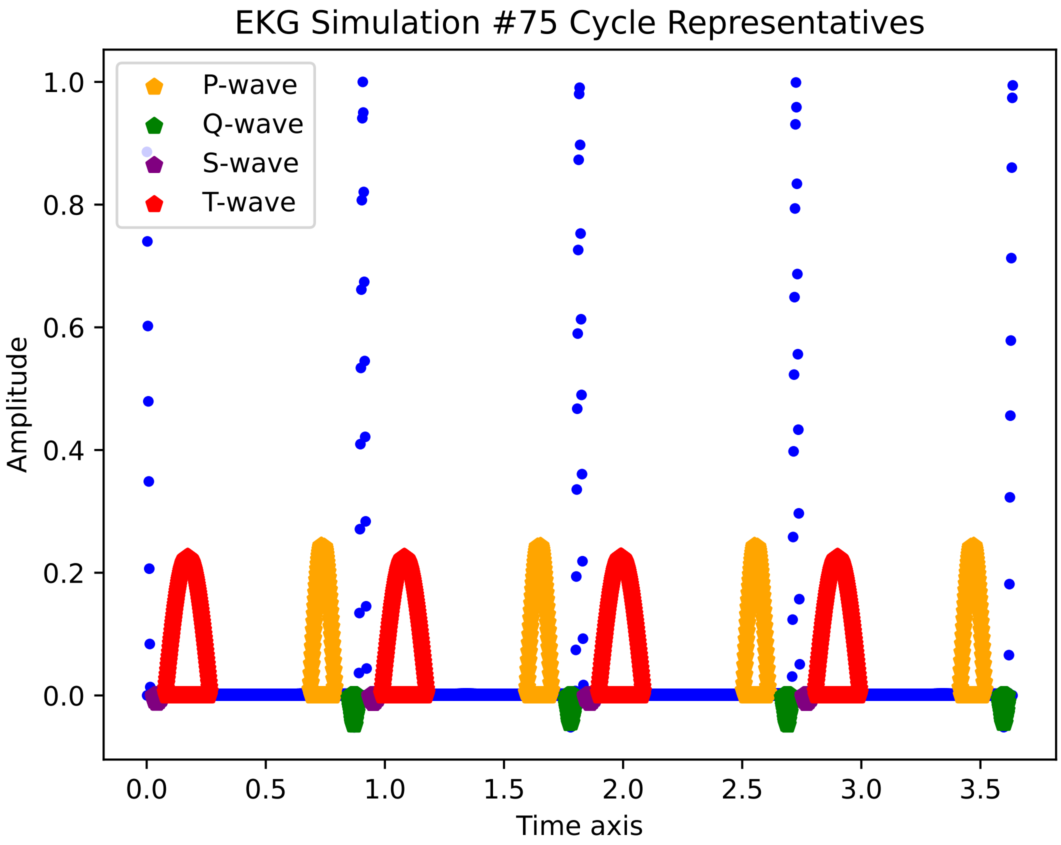
EKG Simulation #75 Cycle Representatives
P-wave
Q-wave
S-wave
T-wave
Amplitude
Time axis
1.0
0.8
0.6
0.4
0.2
0.0
0.0
0.5
1.0
1.5
2.0
2.5
3.0
3.5

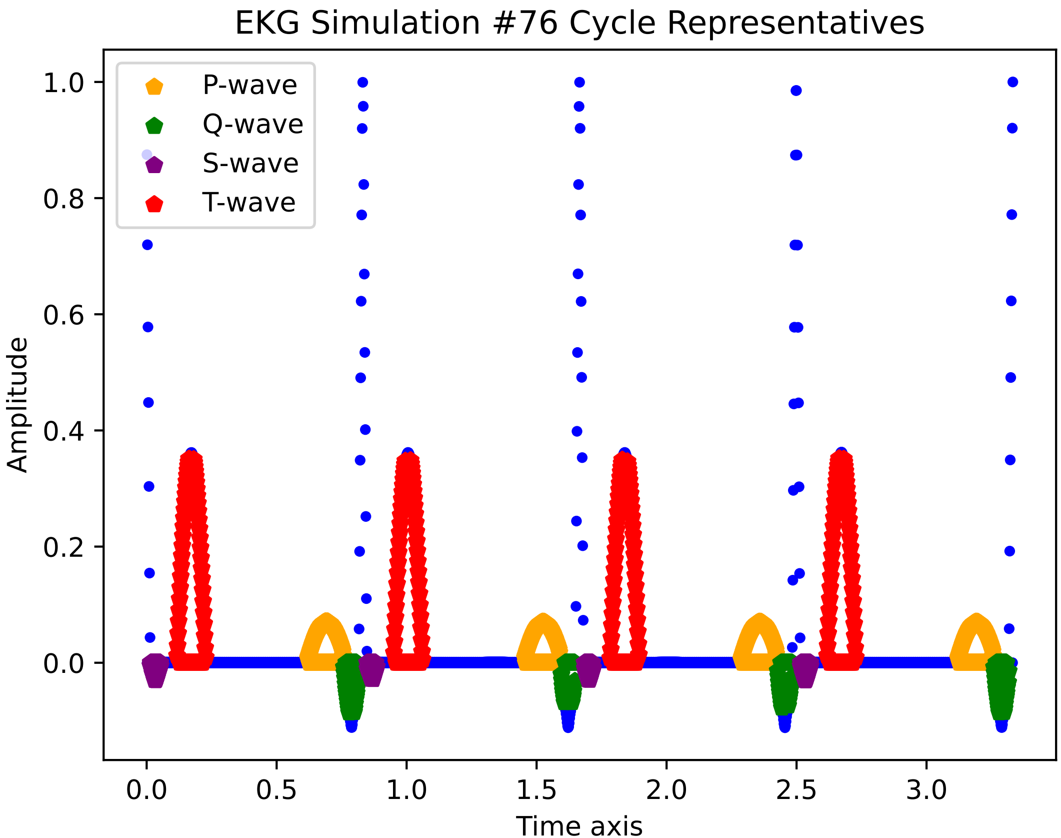

EKG Simulation #76 Cycle Representatives
P-wave
Q-wave
S-wave
T-wave
1.0
0.8
0.6
0.4
0.2
0.0
Amplitude
0.0
0.5
1.0
1.5
2.0
2.5
3.0
Time axis

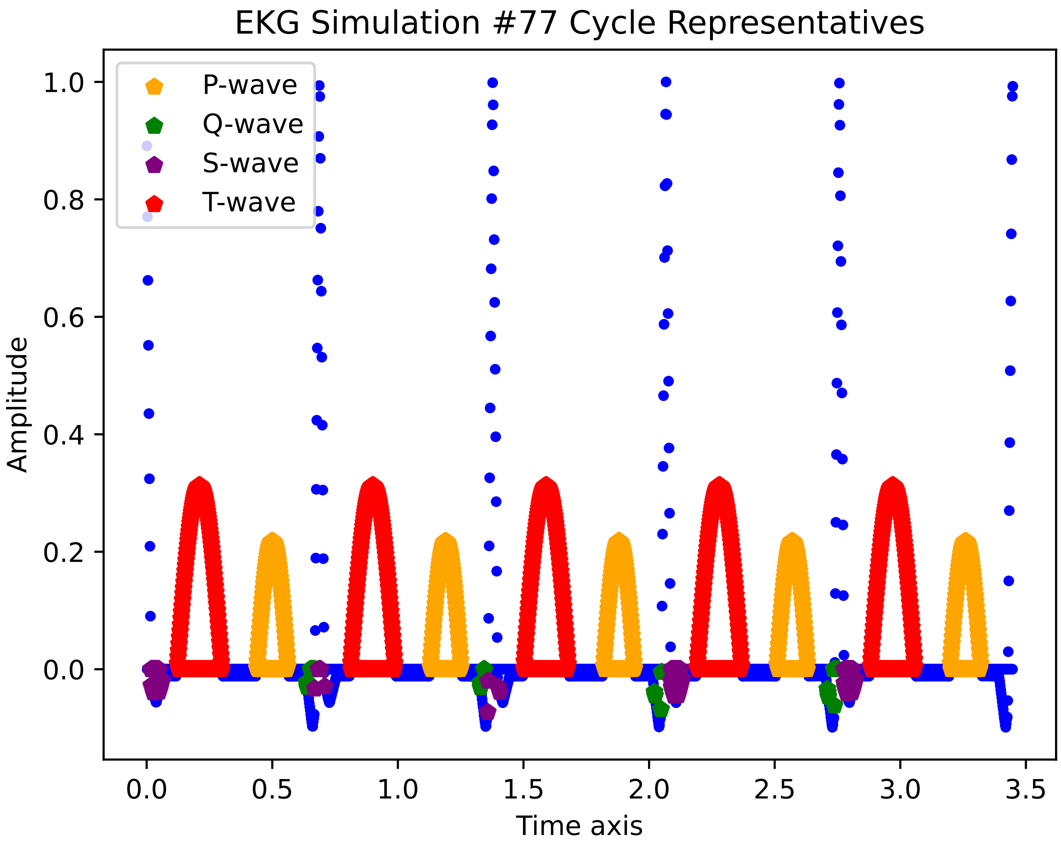

EKG Simulation #77 Cycle Representatives
P-wave
Q-wave
S-wave
T-wave
Amplitude
Time axis
1.0
0.8
0.6
0.4
0.2
0.0
0.0
0.5
1.0
1.5
2.0
2.5
3.0
3.5

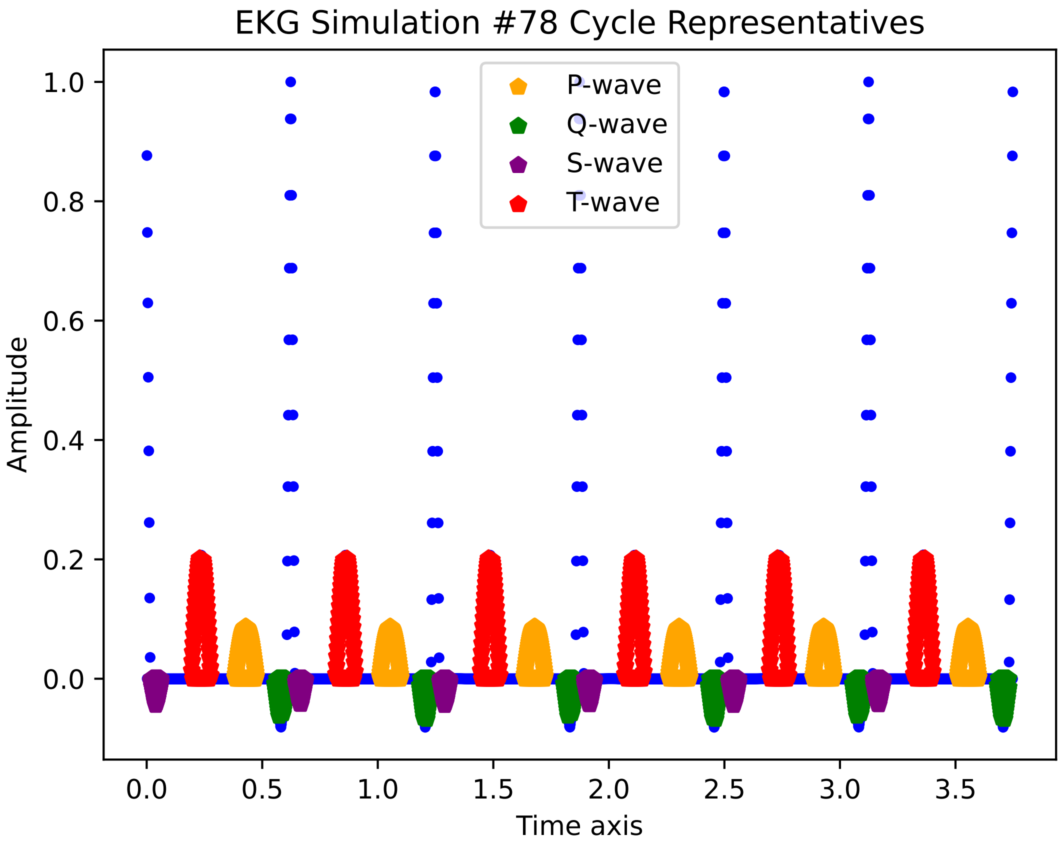
EKG Simulation #78 Cycle Representatives
P-wave
Q-wave
S-wave
T-wave
Amplitude
1.0
0.8
0.6
0.4
0.2
0.0
0.0
0.5
1.0
1.5
2.0
2.5
3.0
3.5
Time axis

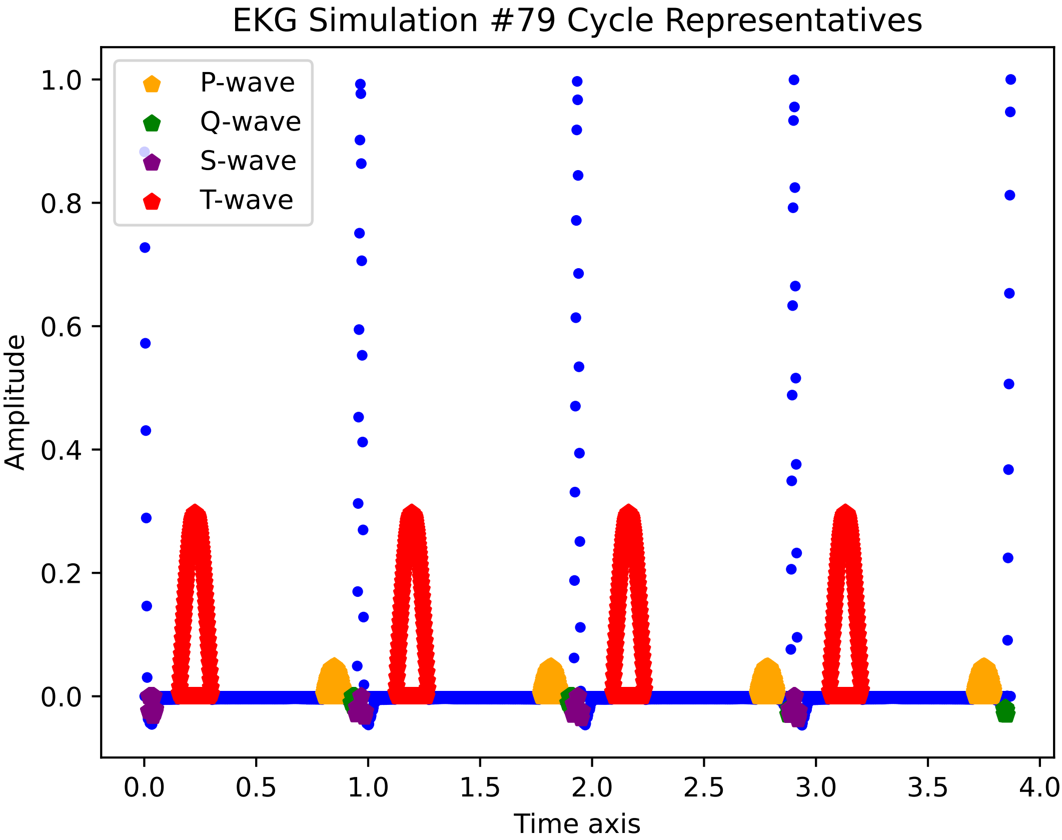

EKG Simulation #79 Cycle Representatives
P-wave
Q-wave
S-wave
T-wave
Amplitude
Time axis
1.0
0.8
0.6
0.4
0.2
0.0
0.0
0.5
1.0
1.5
2.0
2.5
3.0
3.5
4.0

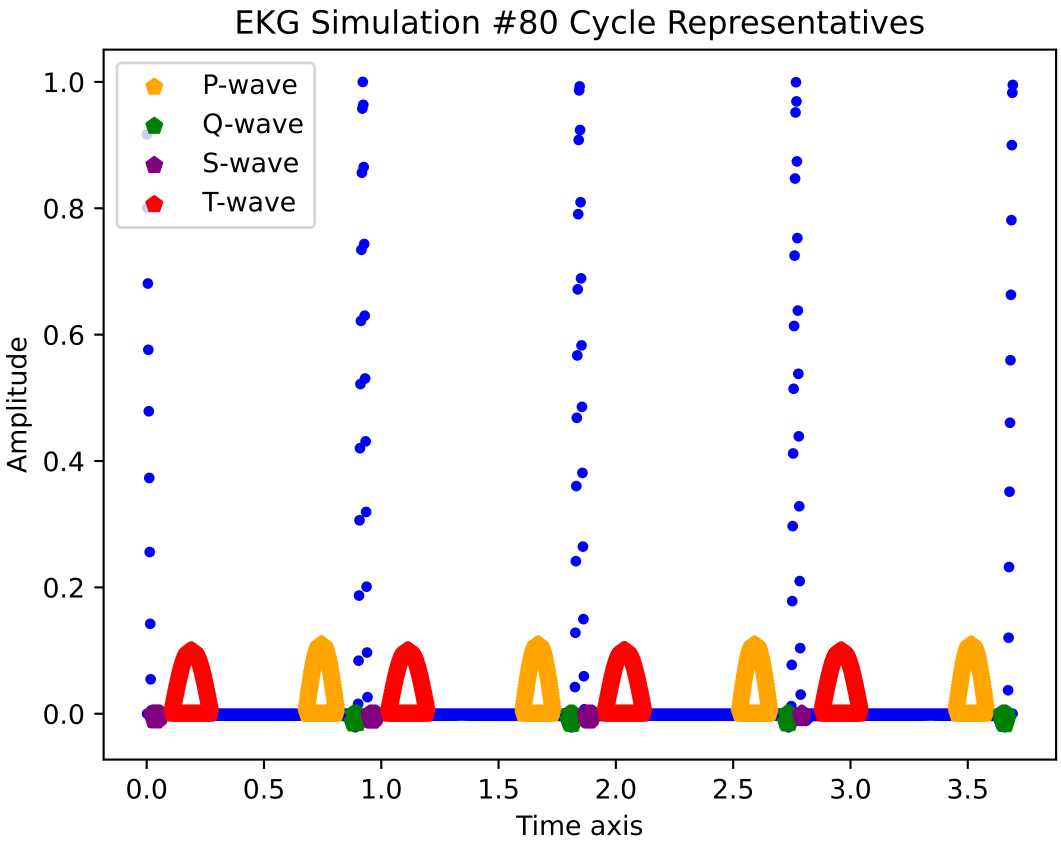
EKG Simulation #80 Cycle Representatives
P-wave
Q-wave
S-wave
T-wave
Amplitude
Time axis
1.0
0.8
0.6
0.4
0.2
0.0
0.0
0.5
1.0
1.5
2.0
2.5
3.0
3.5

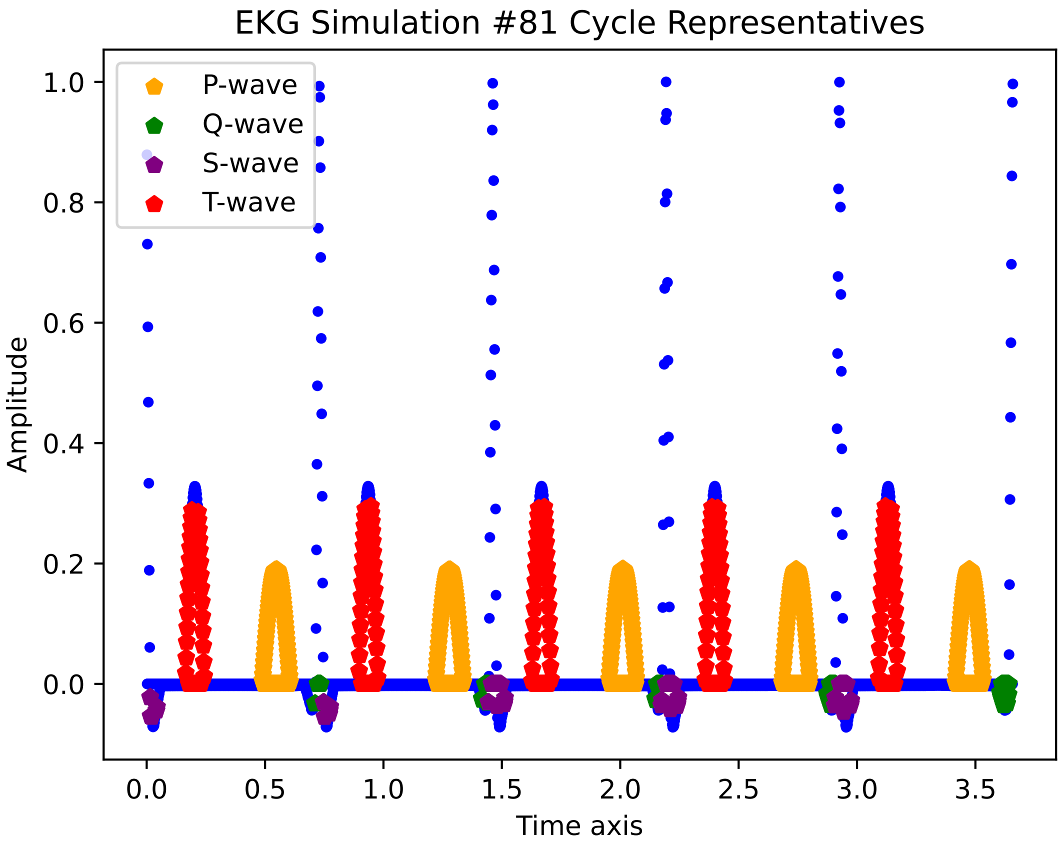
EKG Simulation #81 Cycle Representatives
P-wave
Q-wave
S-wave
T-wave
Amplitude
1.0
0.8
0.6
0.4
0.2
0.0
0.0
0.5
1.0
1.5
2.0
2.5
3.0
3.5
Time axis

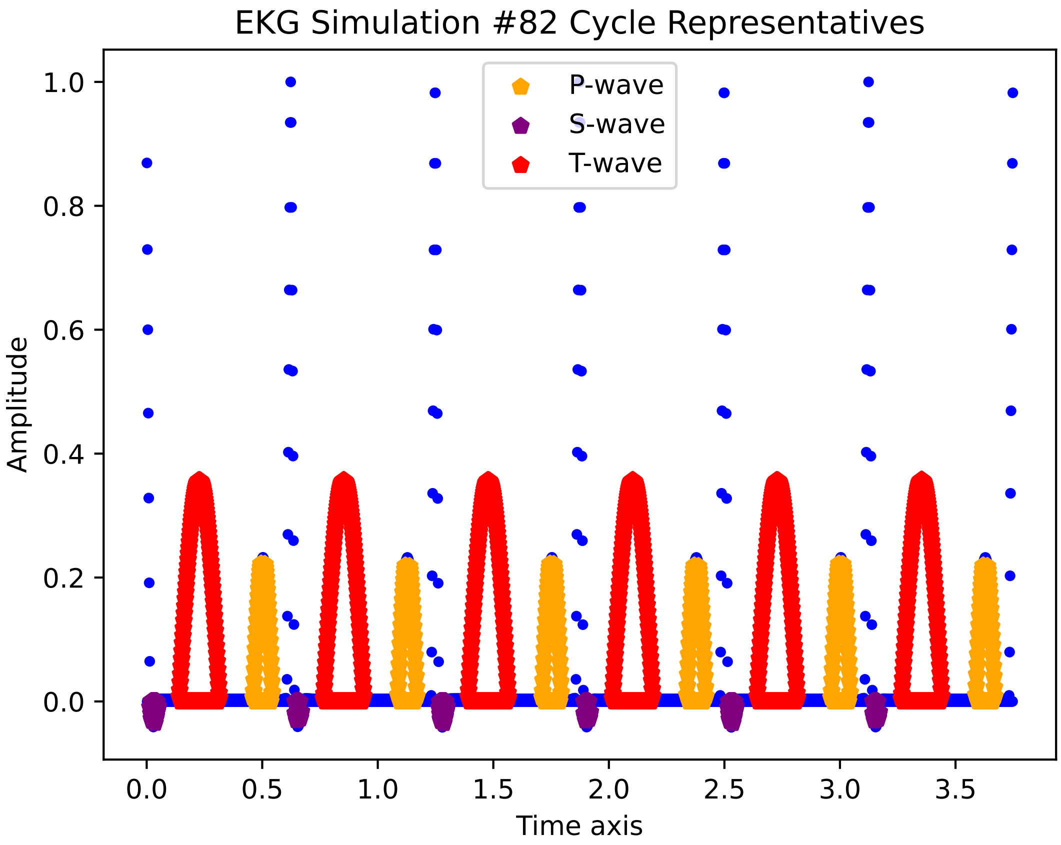

EKG Simulation #82 Cycle Representatives
P-wave
S-wave
T-wave
Amplitude
Time axis
0.0
0.2
0.4
0.6
0.8
1.0
0.0
0.5
1.0
1.5
2.0
2.5
3.0
3.5

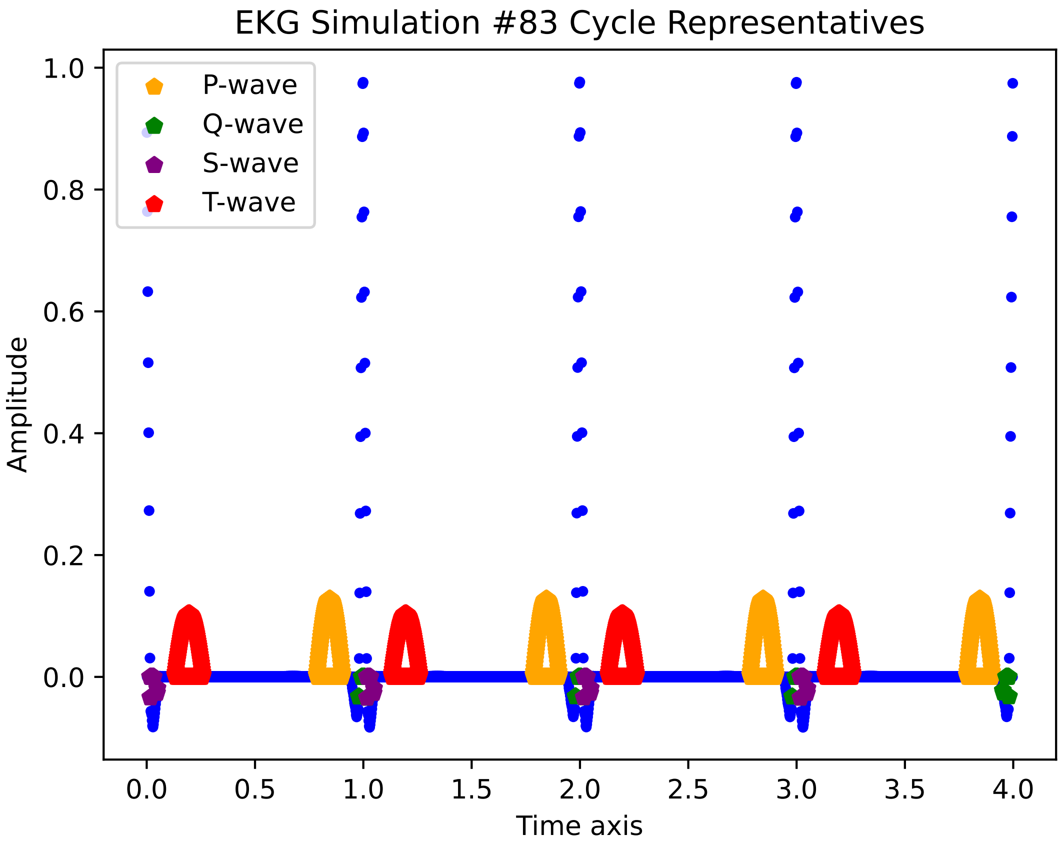

EKG Simulation #83 Cycle Representatives
P-wave
Q-wave
S-wave
T-wave
Amplitude
Time axis
1.0
0.8
0.6
0.4
0.2
0.0
0.0
0.5
1.0
1.5
2.0
2.5
3.0
3.5
4.0

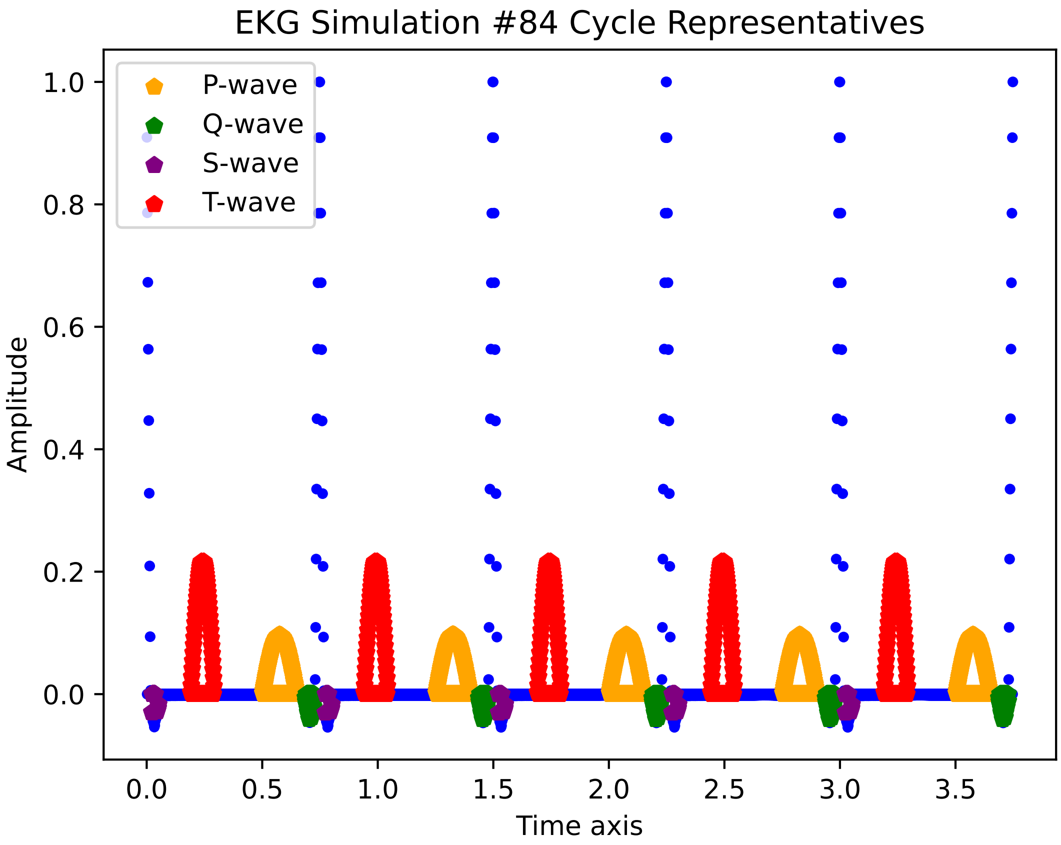
EKG Simulation #84 Cycle Representatives
P-wave
Q-wave
S-wave
T-wave
Amplitude
Time axis
1.0
0.8
0.6
0.4
0.2
0.0
0.0
0.5
1.0
1.5
2.0
2.5
3.0
3.5

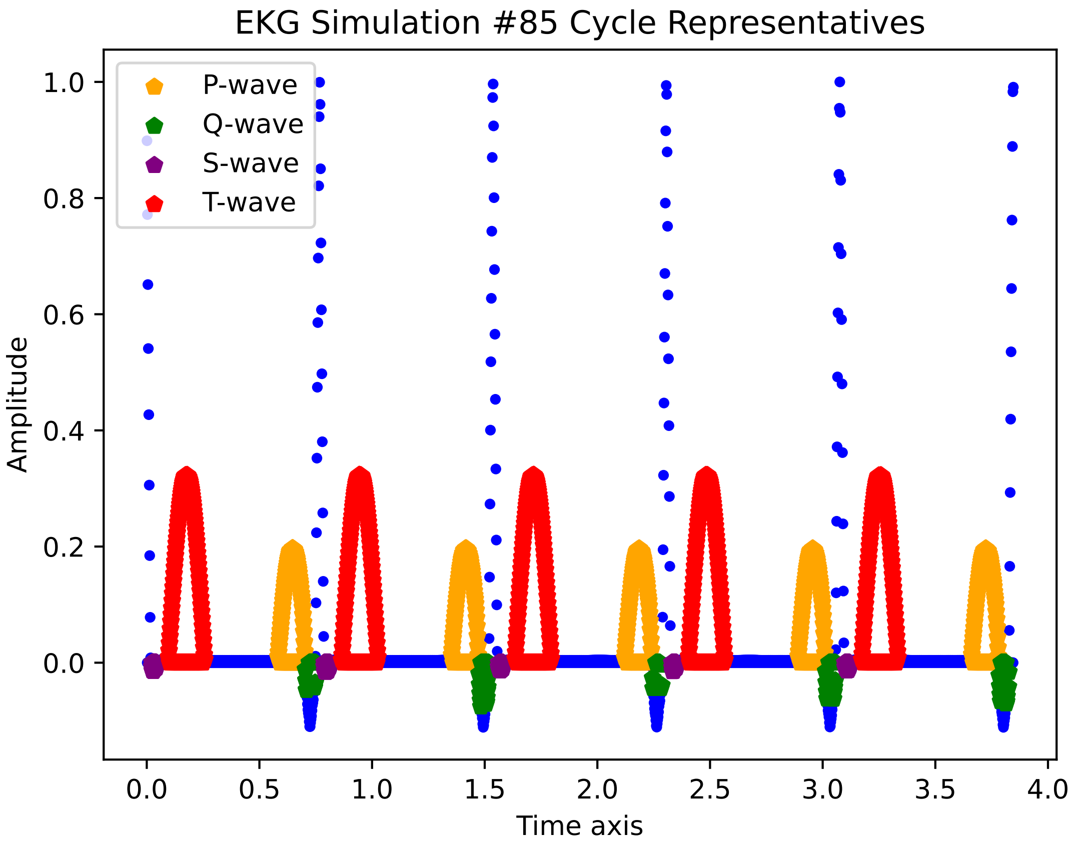
EKG Simulation #85 Cycle Representatives
P-wave
Q-wave
S-wave
T-wave
Amplitude
Time axis
1.0
0.8
0.6
0.4
0.2
0.0
0.0
0.5
1.0
1.5
2.0
2.5
3.0
3.5
4.0

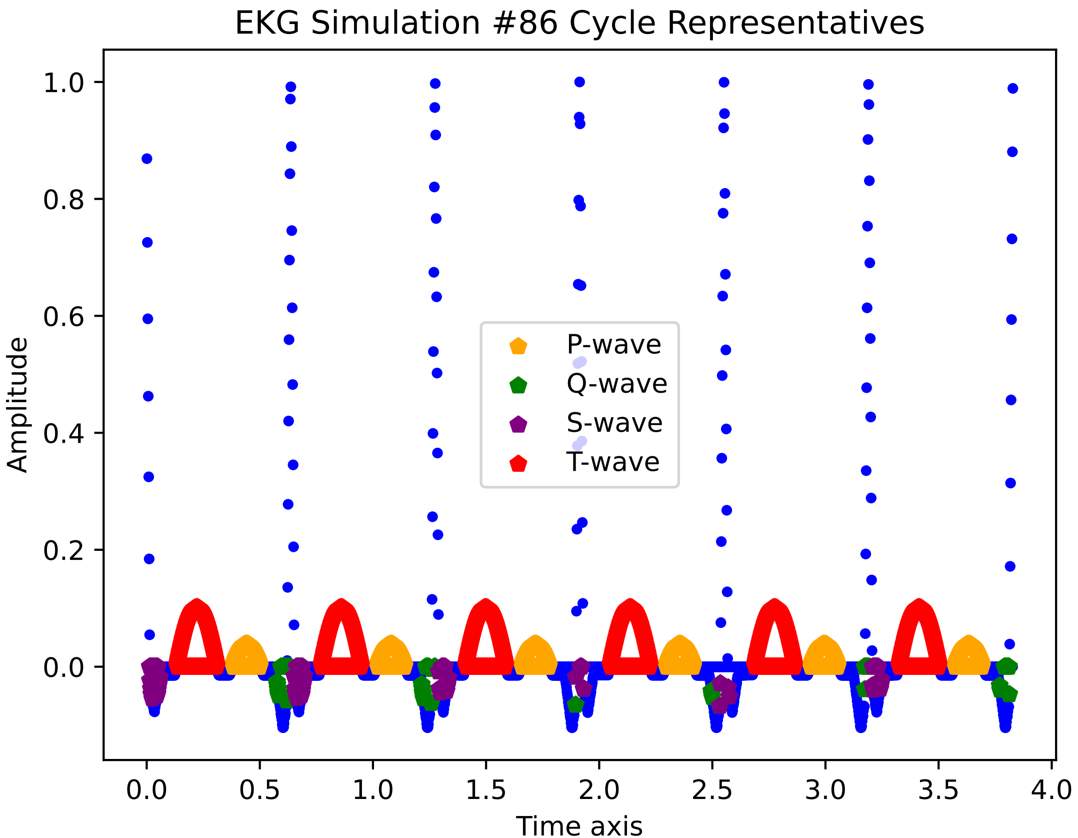
EKG Simulation #86 Cycle Representatives
Amplitude
Time axis
P-wave
Q-wave
S-wave
T-wave
1.0
0.8
0.6
0.4
0.2
0.0
0.0
0.5
1.0
1.5
2.0
2.5
3.0
3.5
4.0

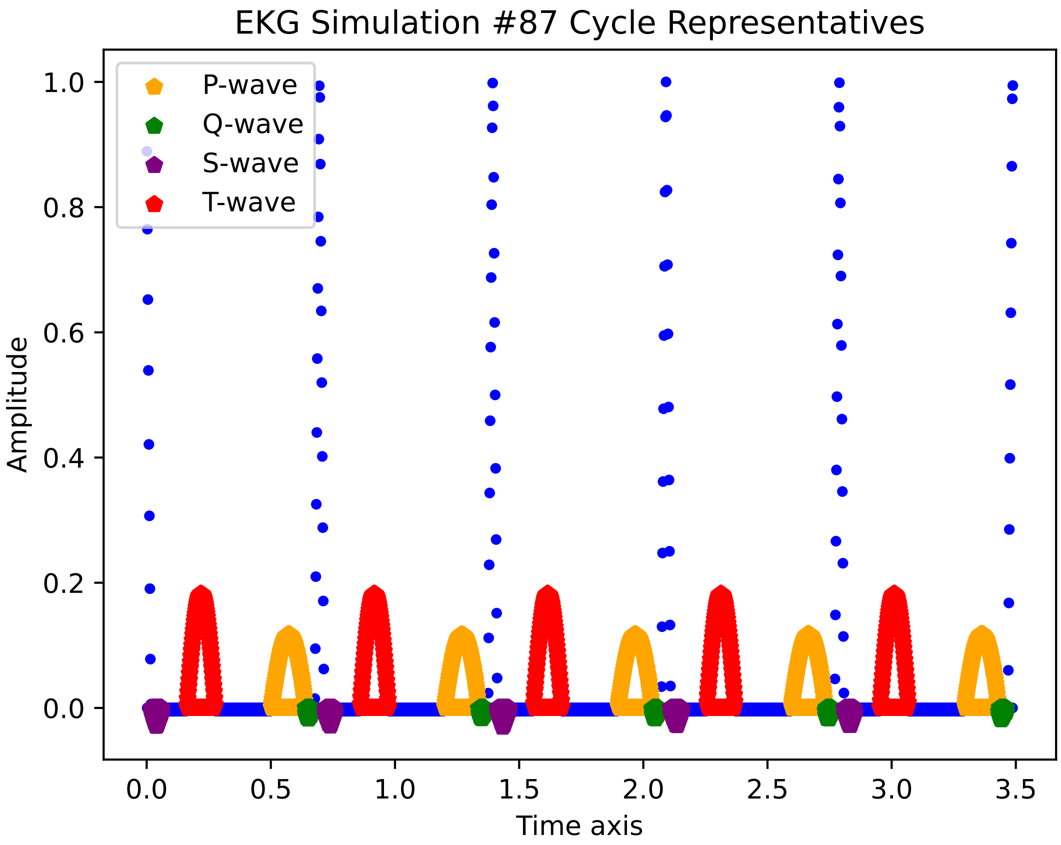
EKG Simulation #87 Cycle Representatives
P-wave
Q-wave
S-wave
T-wave
Amplitude
Time axis
0.0
0.2
0.4
0.6
0.8
1.0
0.0
0.5
1.0
1.5
2.0
2.5
3.0
3.5

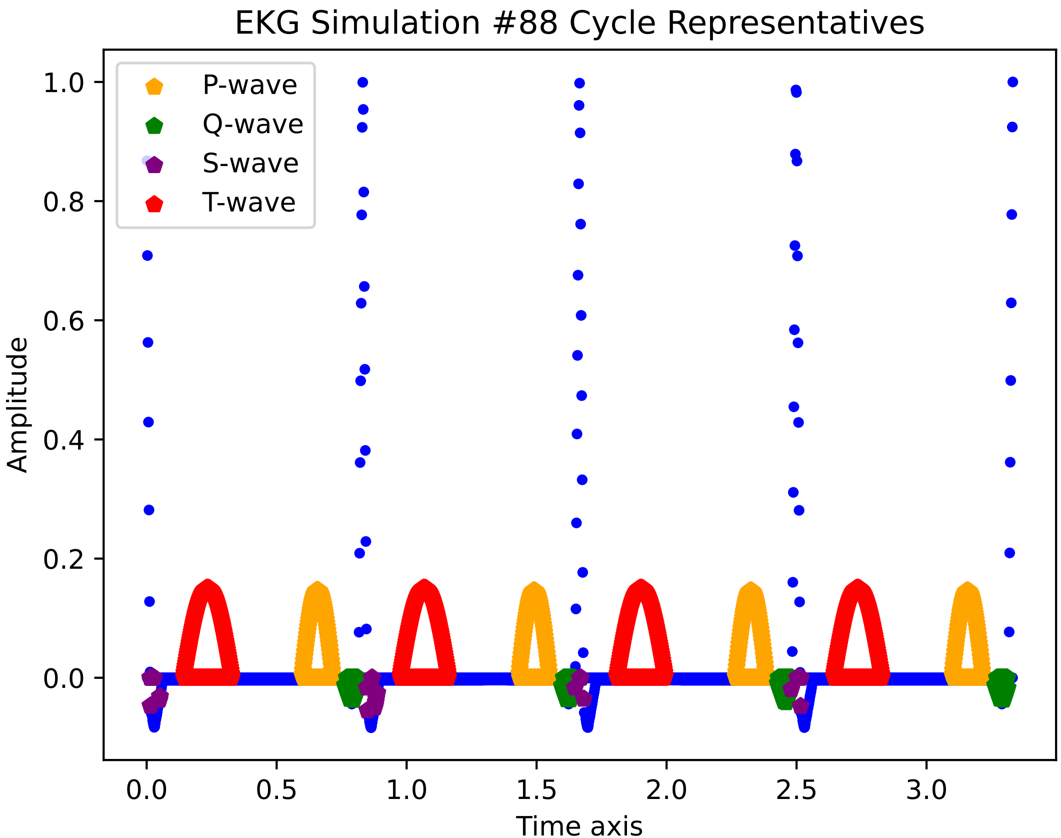
EKG Simulation #88 Cycle Representatives
P-wave
Q-wave
S-wave
T-wave
Amplitude
1.0
0.8
0.6
0.4
0.2
0.0
Time axis
0.0
0.5
1.0
1.5
2.0
2.5
3.0

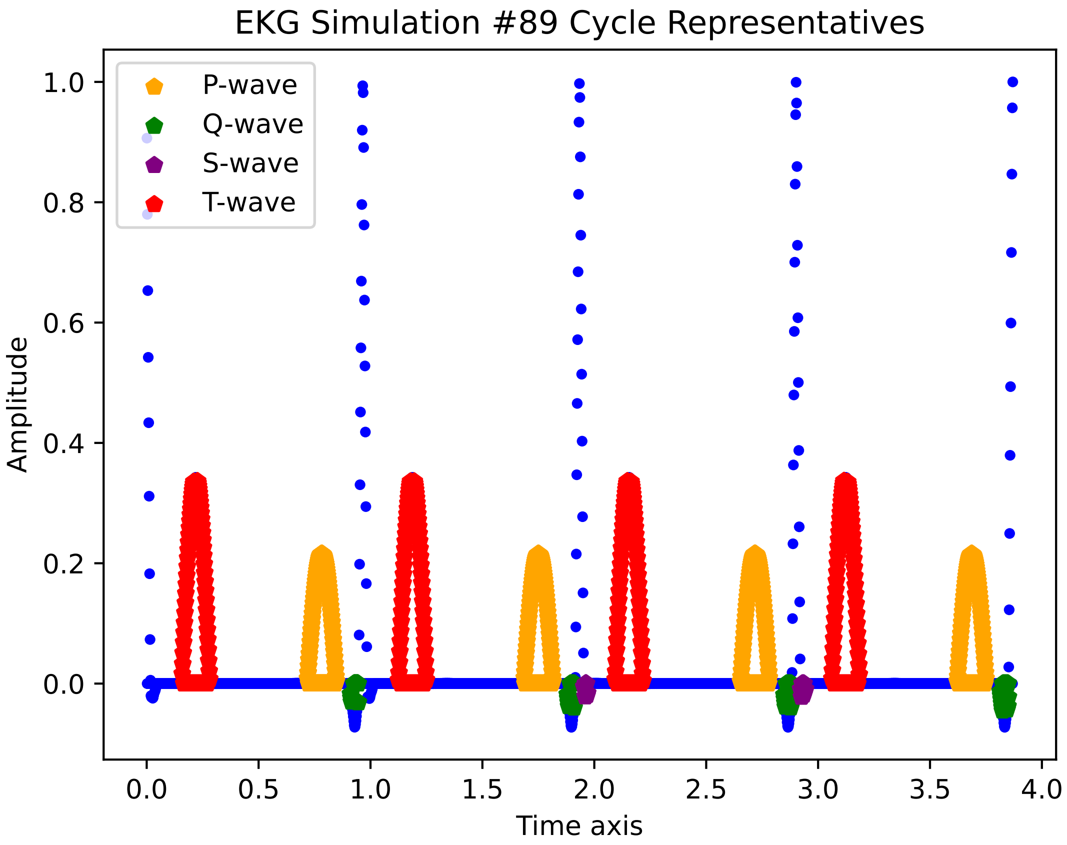
EKG Simulation #89 Cycle Representatives
P-wave
Q-wave
S-wave
T-wave
Amplitude
Time axis
1.0
0.8
0.6
0.4
0.2
0.0
0.0
0.5
1.0
1.5
2.0
2.5
3.0
3.5
4.0

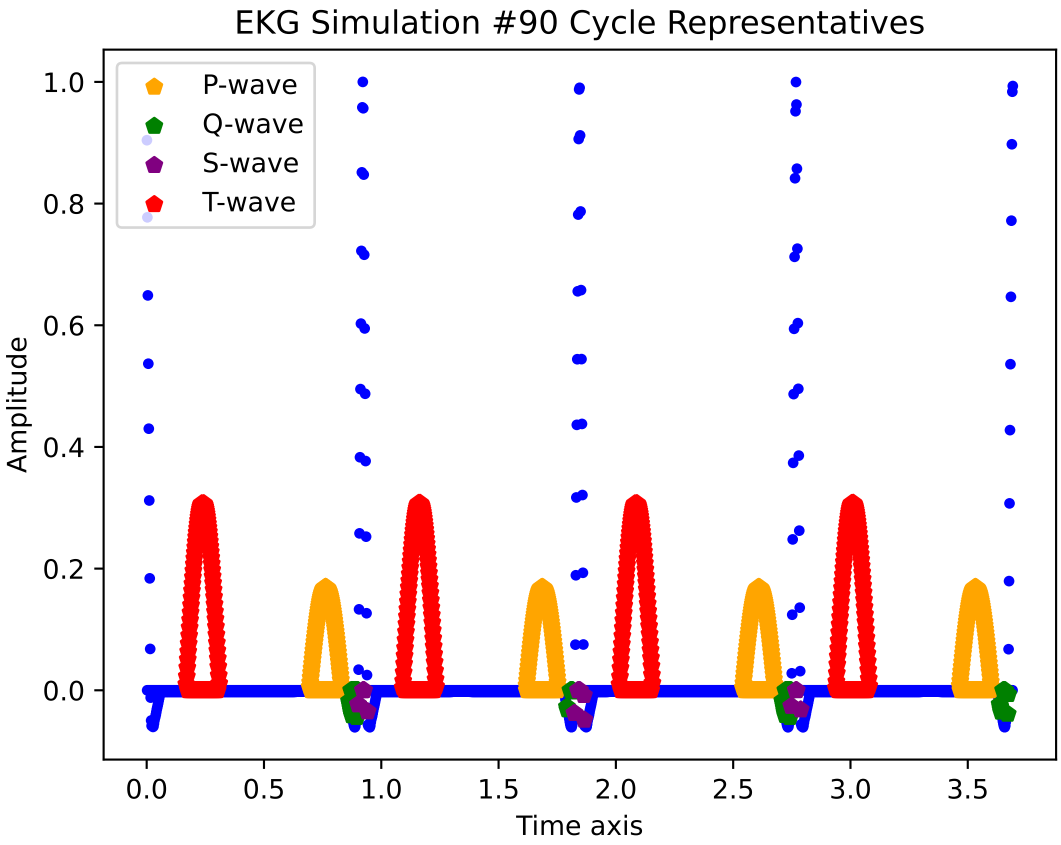

EKG Simulation #90 Cycle Representatives
P-wave
Q-wave
S-wave
T-wave
Amplitude
Time axis
1.0
0.8
0.6
0.4
0.2
0.0
0.0
0.5
1.0
1.5
2.0
2.5
3.0
3.5

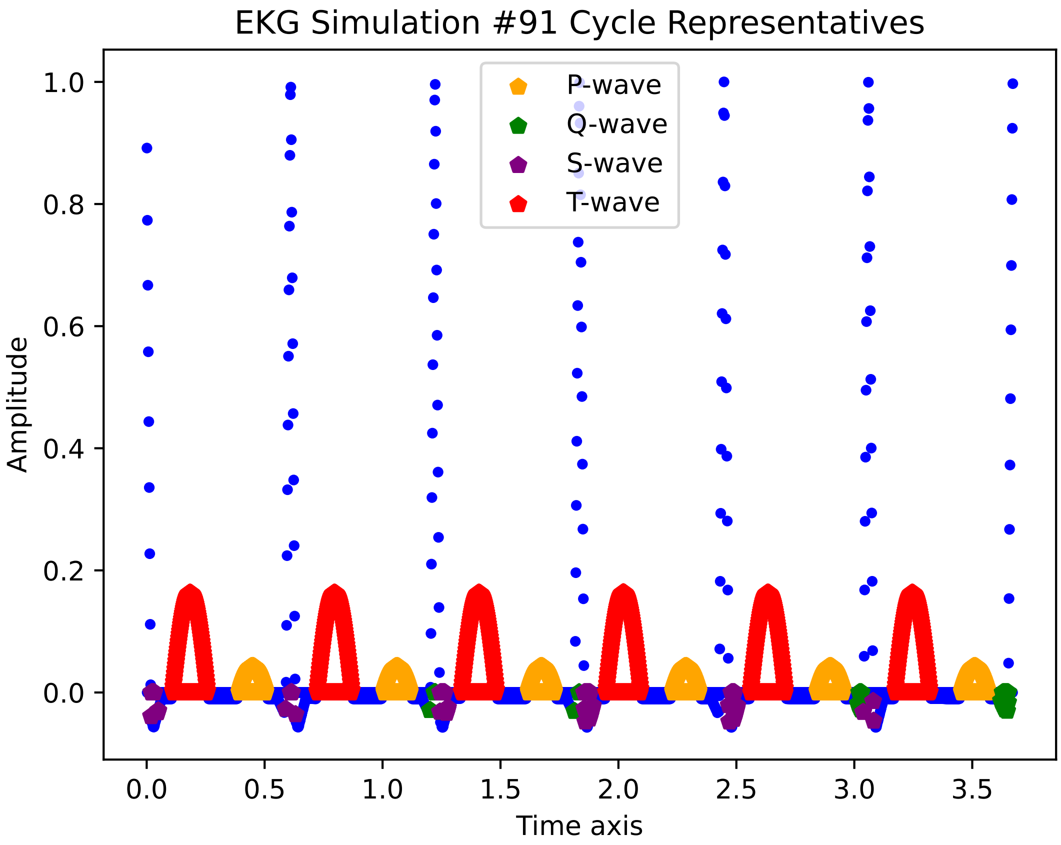

EKG Simulation #91 Cycle Representatives
P-wave
Q-wave
S-wave
T-wave
Amplitude
Time axis
1.0
0.8
0.6
0.4
0.2
0.0
0.0
0.5
1.0
1.5
2.0
2.5
3.0
3.5

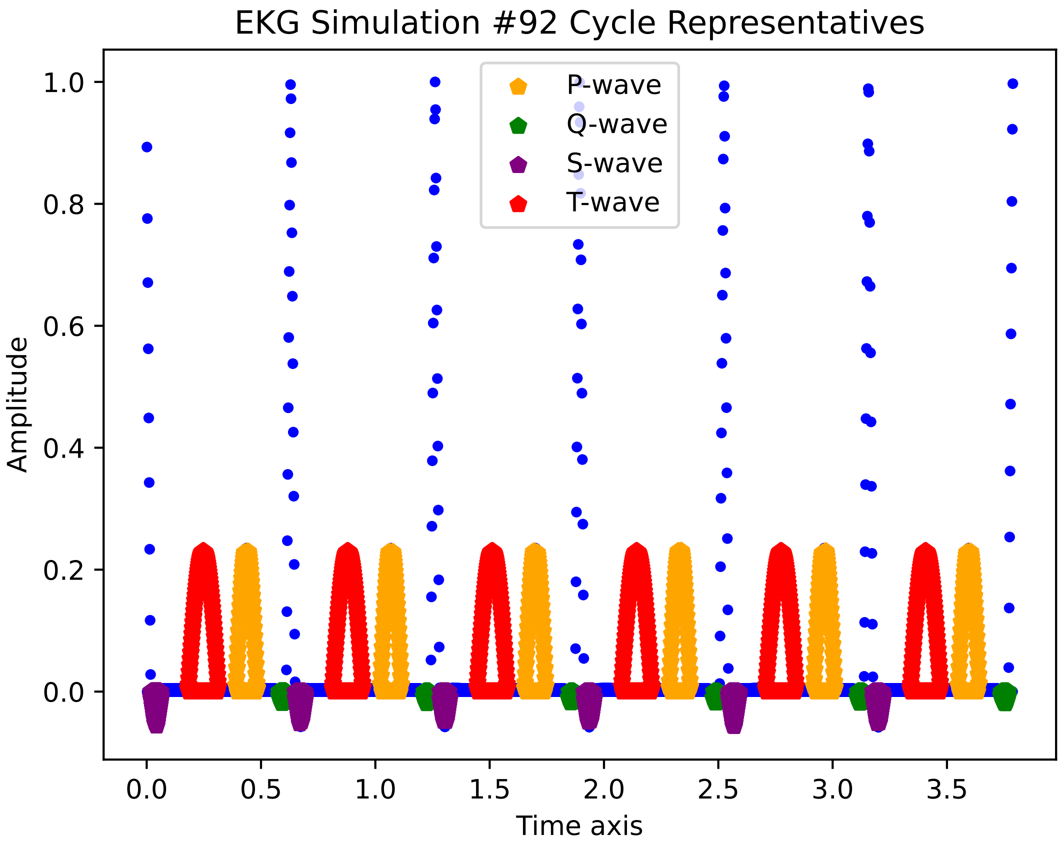

EKG Simulation #92 Cycle Representatives
P-wave
Q-wave
S-wave
T-wave
Amplitude
Time axis
1.0
0.8
0.6
0.4
0.2
0.0
0.0
0.5
1.0
1.5
2.0
2.5
3.0
3.5

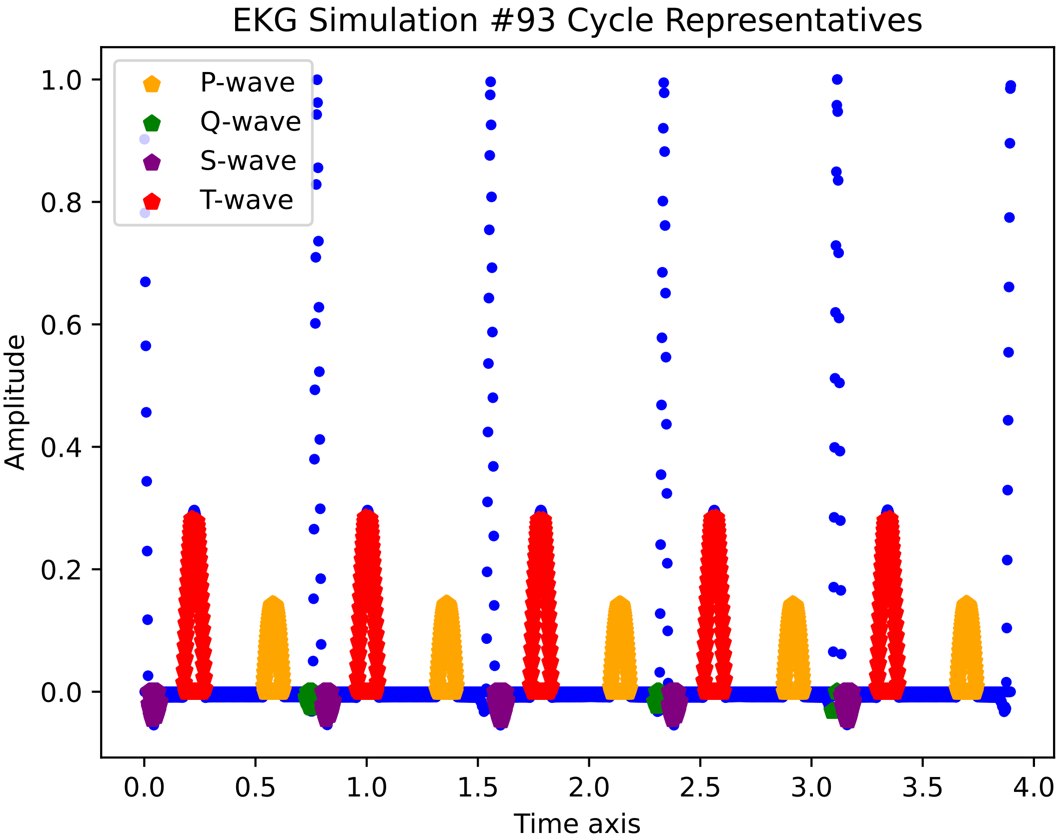
EKG Simulation #93 Cycle Representatives
P-wave
Q-wave
S-wave
T-wave
Amplitude
Time axis
0.0
0.2
0.4
0.6
0.8
1.0
0.0
0.5
1.0
1.5
2.0
2.5
3.0
3.5
4.0

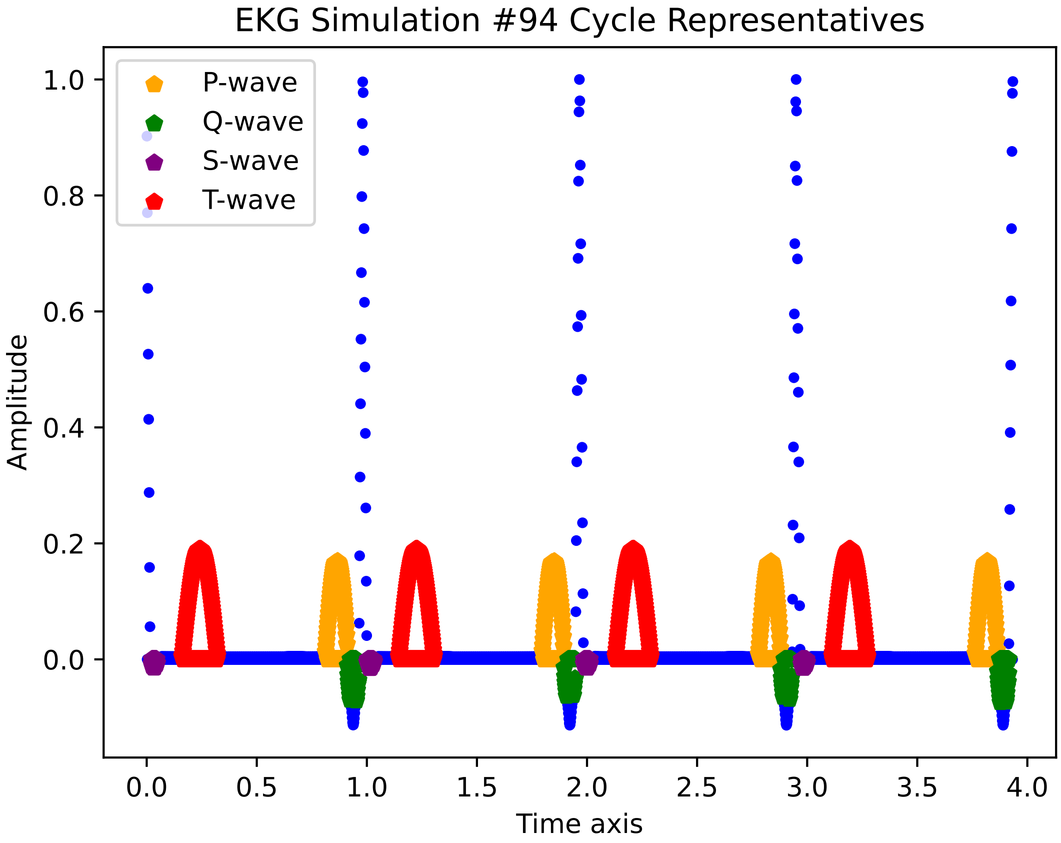
EKG Simulation #94 Cycle Representatives
P-wave
Q-wave
S-wave
T-wave
Amplitude
1.0
0.8
0.6
0.4
0.2
0.0
0.0
0.5
1.0
1.5
2.0
2.5
3.0
3.5
4.0
Time axis

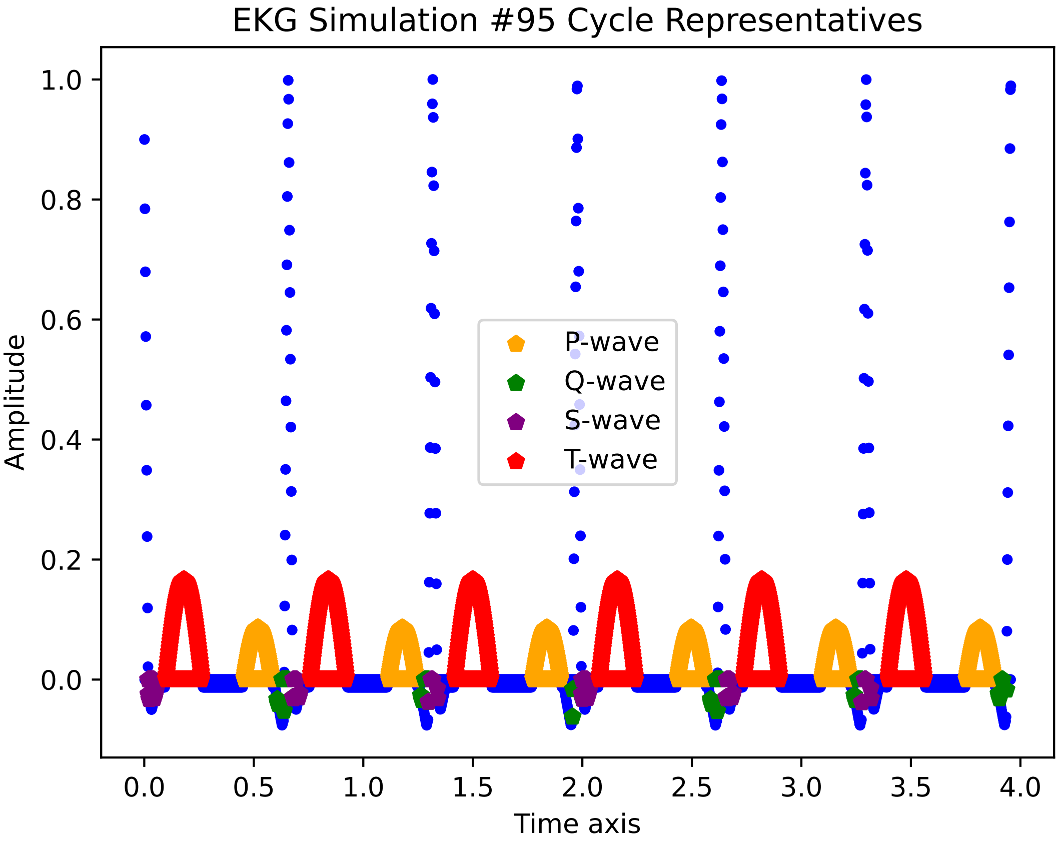
EKG Simulation #95 Cycle Representatives
P-wave
Q-wave
S-wave
T-wave
Amplitude
Time axis
0.0
0.2
0.4
0.6
0.8
1.0
0.0
0.5
1.0
1.5
2.0
2.5
3.0
3.5
4.0

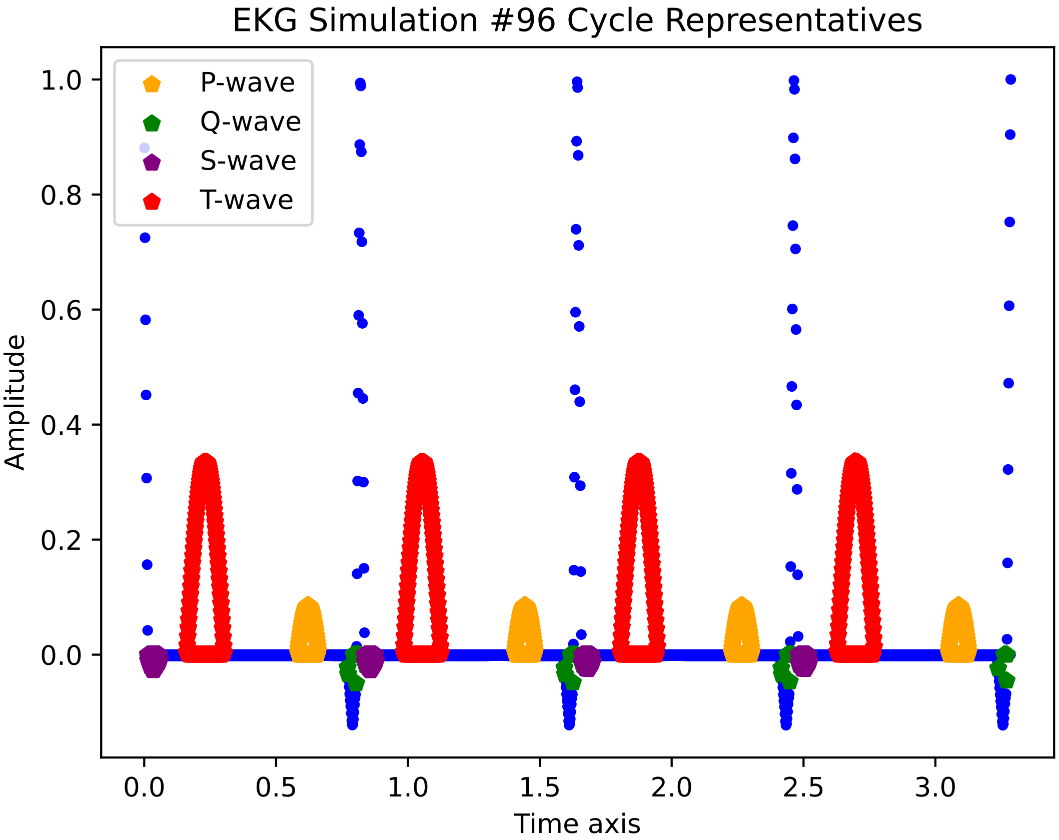

EKG Simulation #96 Cycle Representatives
P-wave
Q-wave
S-wave
T-wave
Amplitude
Time axis
1.0
0.8
0.6
0.4
0.2
0.0
0.0
0.5
1.0
1.5
2.0
2.5
3.0

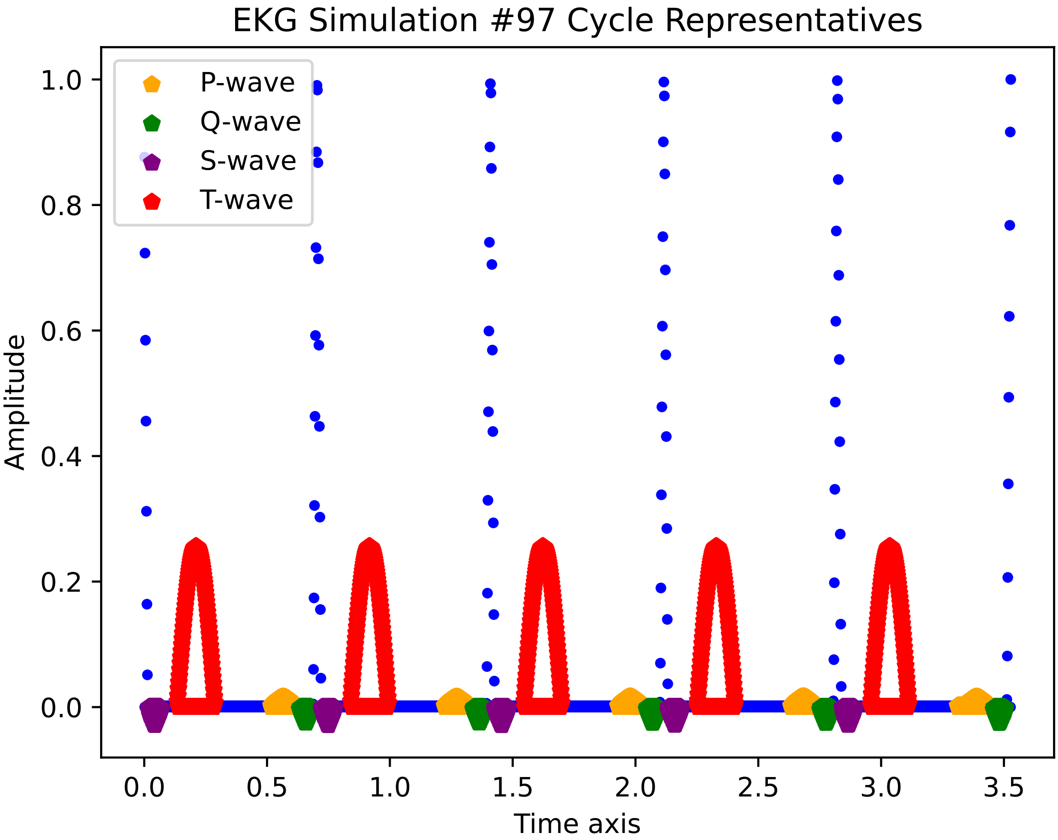

EKG Simulation #97 Cycle Representatives
P-wave
Q-wave
S-wave
T-wave
Amplitude
Time axis
1.0
0.8
0.6
0.4
0.2
0.0
0.0
0.5
1.0
1.5
2.0
2.5
3.0
3.5

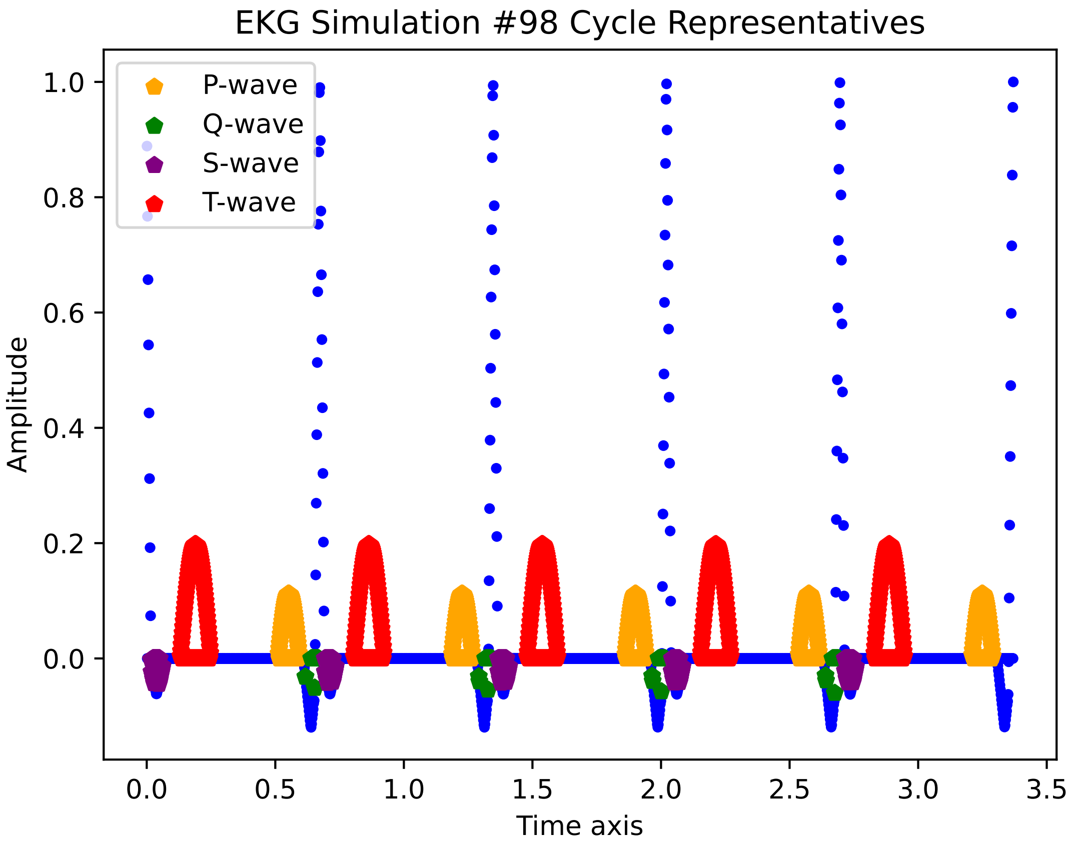

EKG Simulation #98 Cycle Representatives
P-wave
Q-wave
S-wave
T-wave
Amplitude
Time axis
1.0
0.8
0.6
0.4
0.2
0.0
0.0
0.5
1.0
1.5
2.0
2.5
3.0
3.5

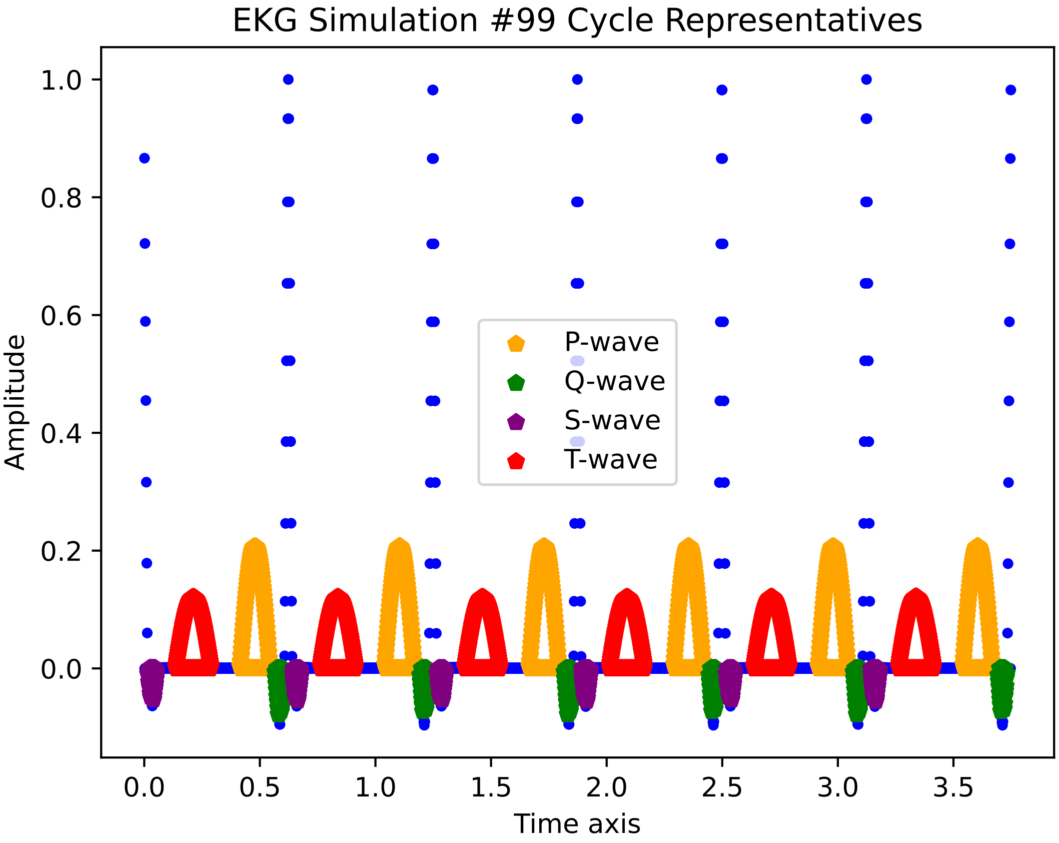
EKG Simulation #99 Cycle Representatives
Amplitude
Time axis
P-wave
Q-wave
S-wave
T-wave
1.0
0.8
0.6
0.4
0.2
0.0
0.0
0.5
1.0
1.5
2.0
2.5
3.0
3.5

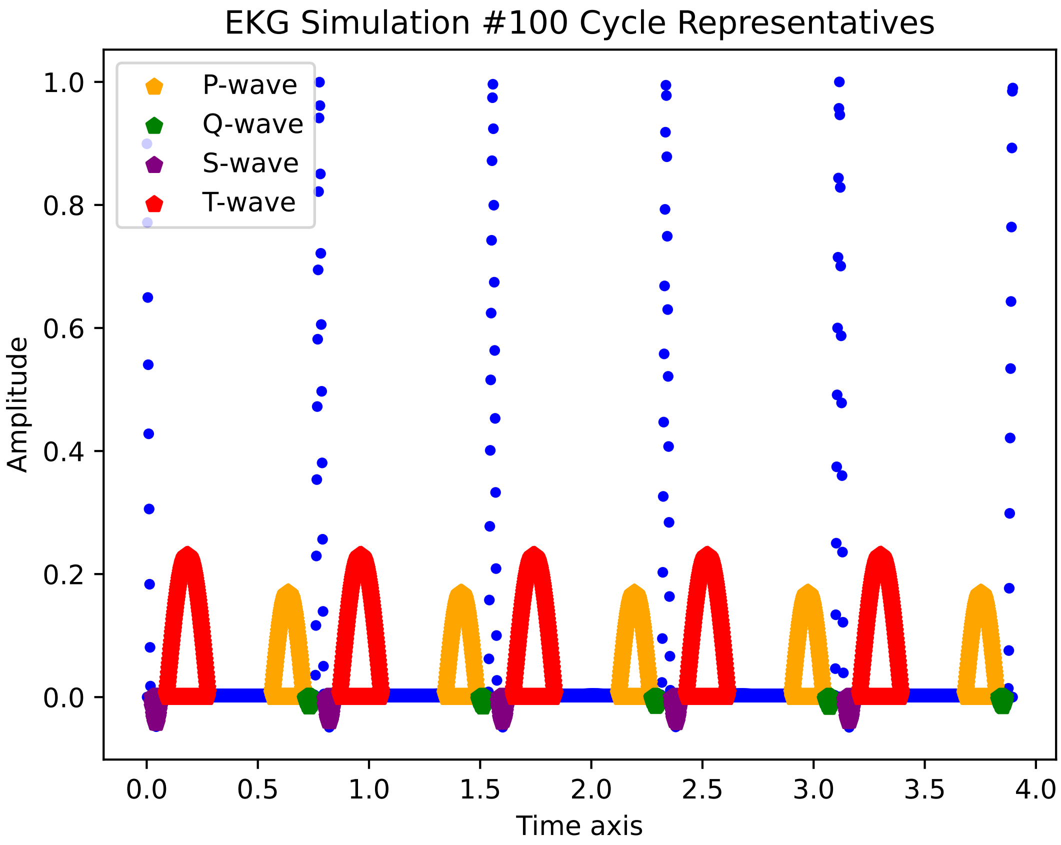
EKG Simulation #100 Cycle Representatives
P-wave
Q-wave
S-wave
T-wave
Amplitude
Time axis
0.0
0.2
0.4
0.6
0.8
1.0
0.0
0.5
1.0
1.5
2.0
2.5
3.0
3.5
4.0

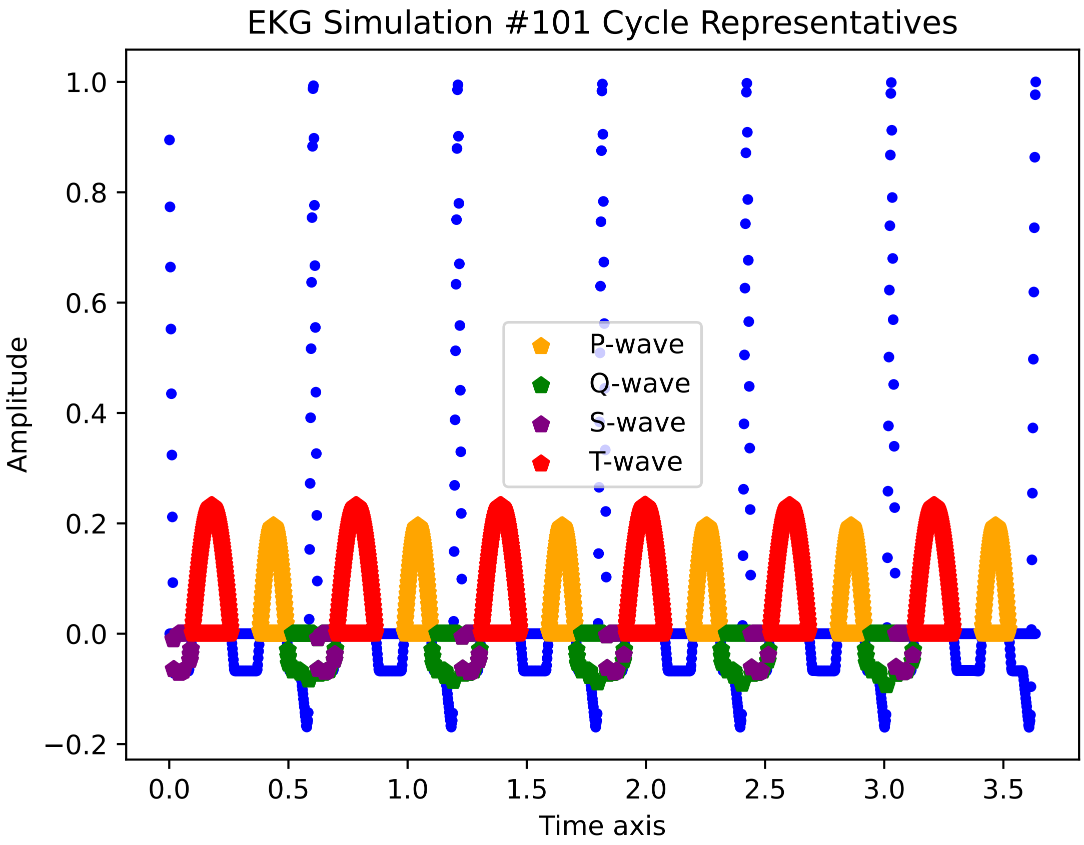

EKG Simulation #101 Cycle Representatives
Amplitude
Time axis
1.0
0.8
0.6
0.4
0.2
0.0
−0.2
0.0
0.5
1.0
1.5
2.0
2.5
3.0
3.5
P-wave
Q-wave
S-wave
T-wave

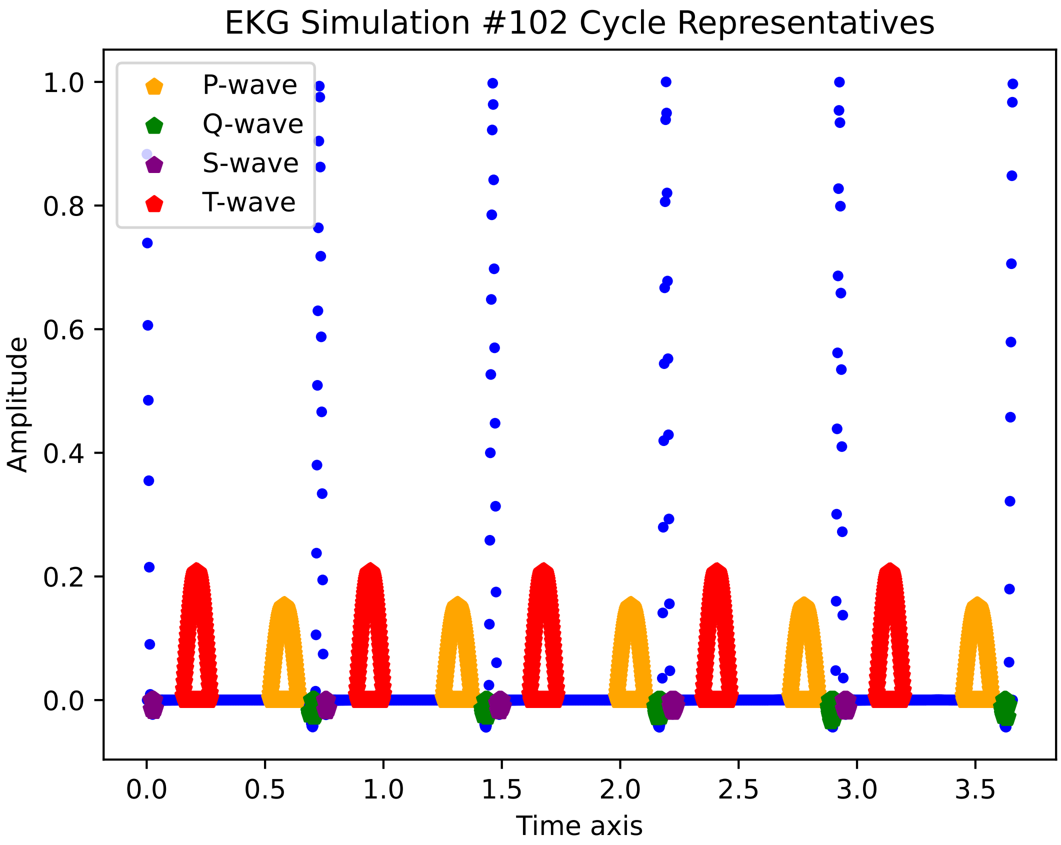
EKG Simulation #102 Cycle Representatives
P-wave
Q-wave
S-wave
T-wave
Amplitude
Time axis
1.0
0.8
0.6
0.4
0.2
0.0
0.0
0.5
1.0
1.5
2.0
2.5
3.0
3.5

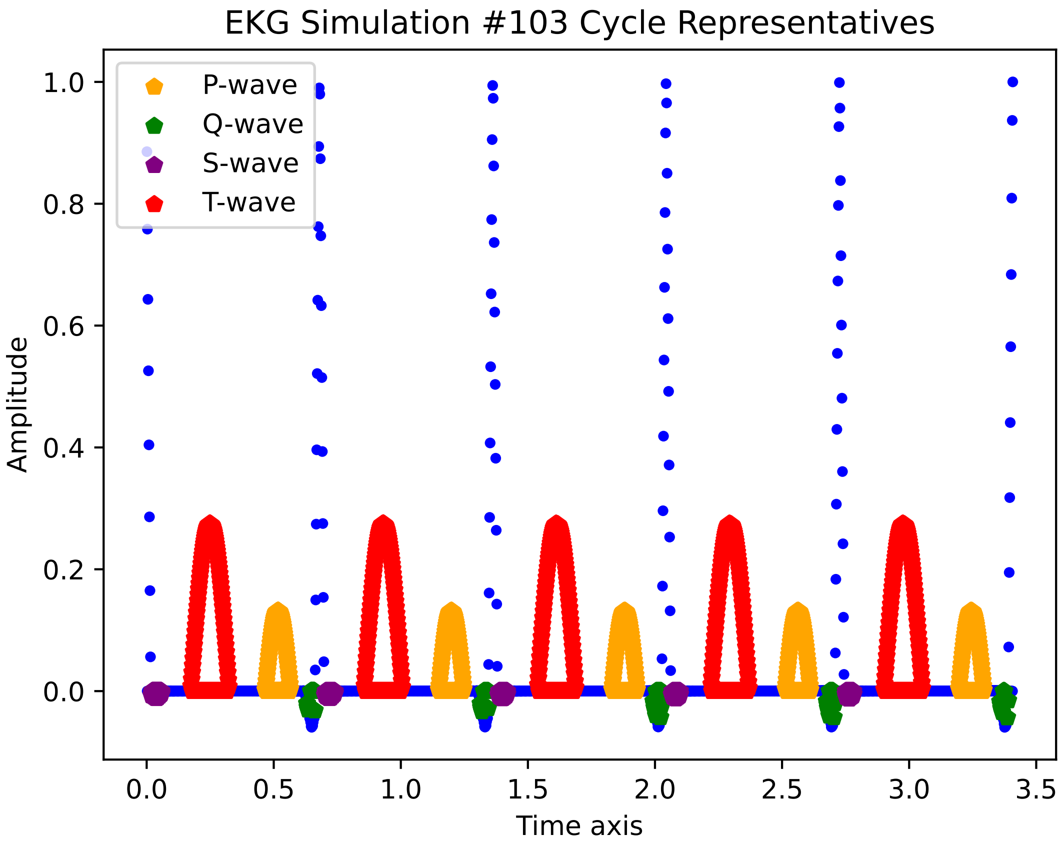
EKG Simulation #103 Cycle Representatives
P-wave
Q-wave
S-wave
T-wave
1.0
0.8
0.6
0.4
0.2
0.0
Amplitude
0.0
0.5
1.0
1.5
2.0
2.5
3.0
3.5
Time axis

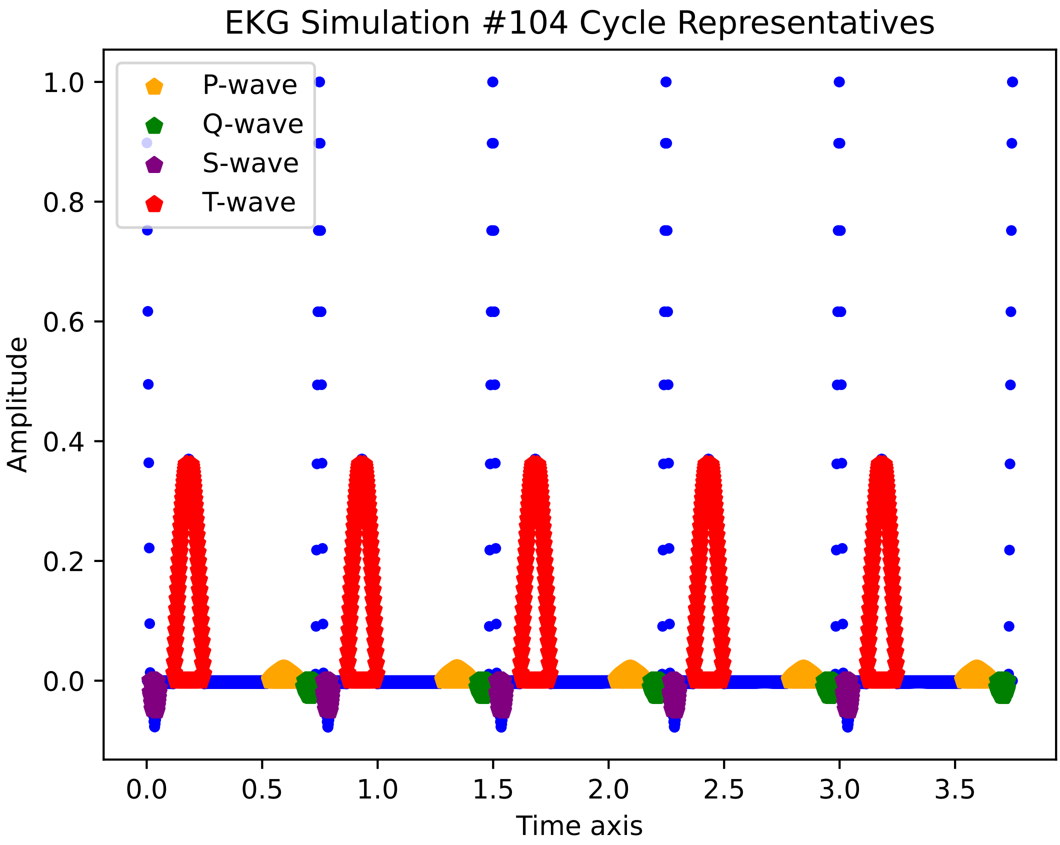
EKG Simulation #104 Cycle Representatives
P-wave
Q-wave
S-wave
T-wave
Amplitude
Time axis
1.0
0.8
0.6
0.4
0.2
0.0
0.0
0.5
1.0
1.5
2.0
2.5
3.0
3.5

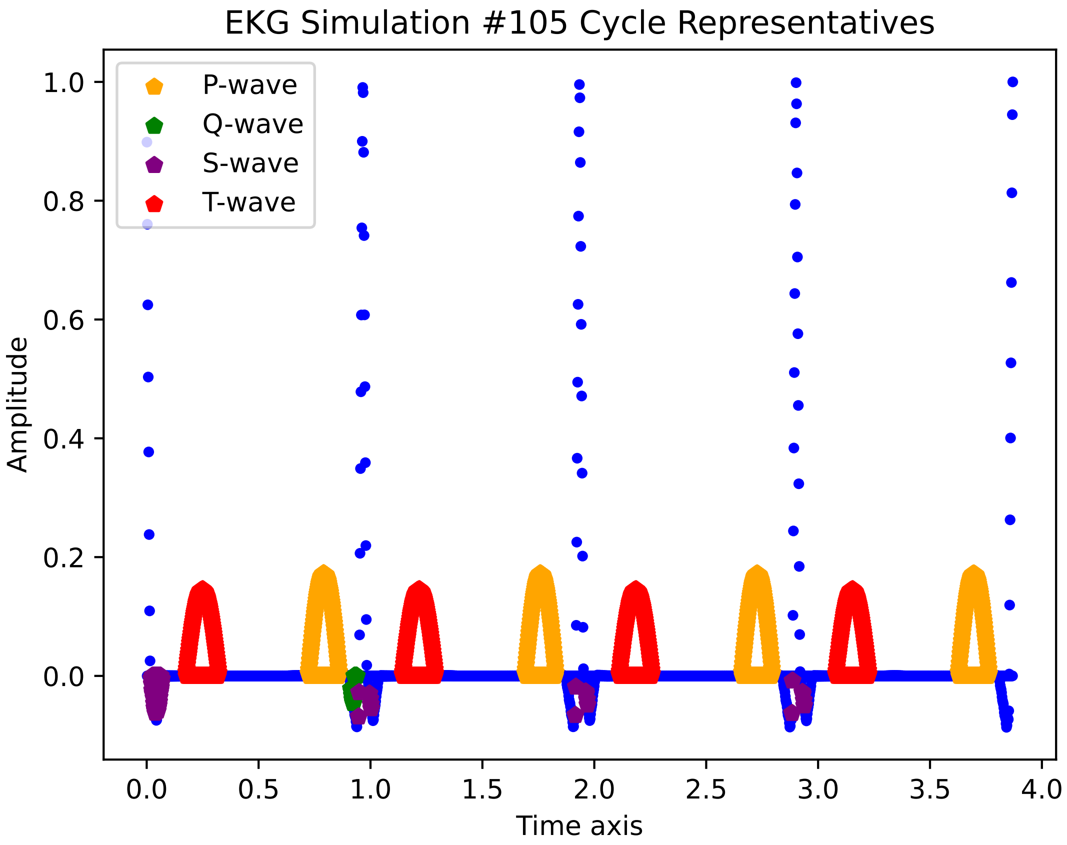
EKG Simulation #105 Cycle Representatives
P-wave
Q-wave
S-wave
T-wave
1.0
0.8
0.6
0.4
0.2
0.0
Amplitude
0.0
0.5
1.0
1.5
2.0
2.5
3.0
3.5
4.0
Time axis

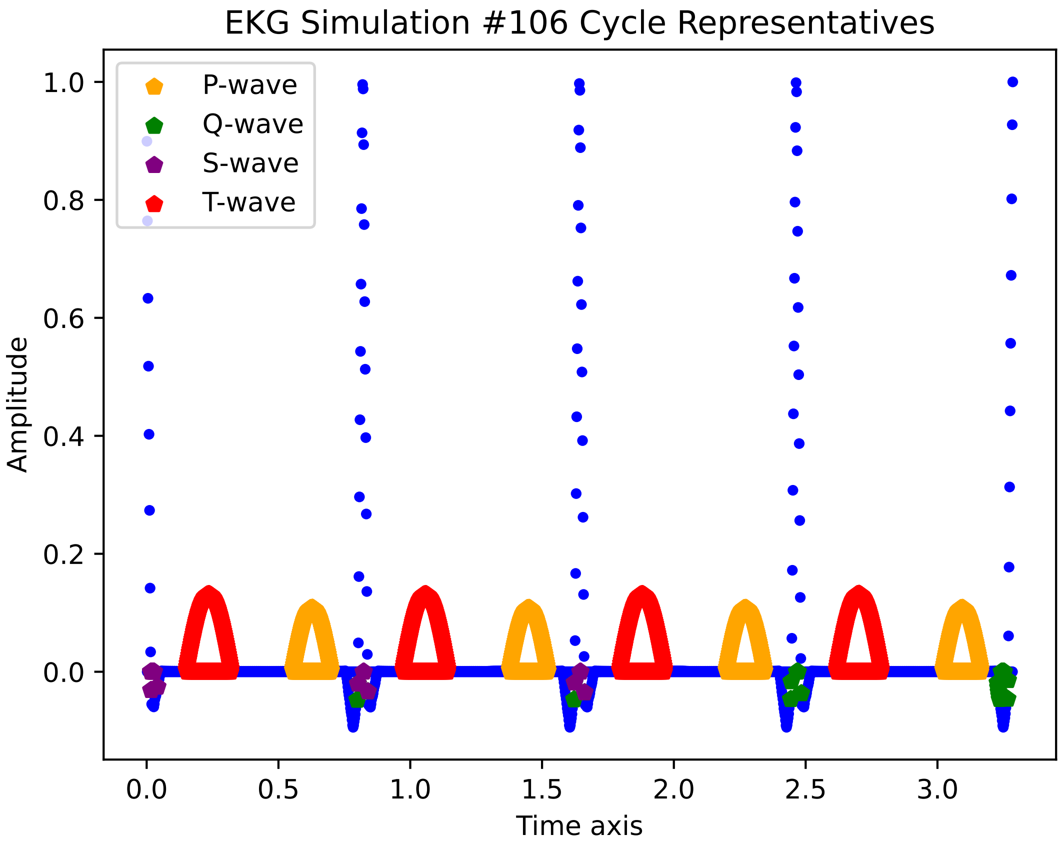
EKG Simulation #106 Cycle Representatives
P-wave
Q-wave
S-wave
T-wave
1.0
0.8
0.6
0.4
0.2
0.0
Amplitude
0.0
0.5
1.0
1.5
2.0
2.5
3.0
Time axis

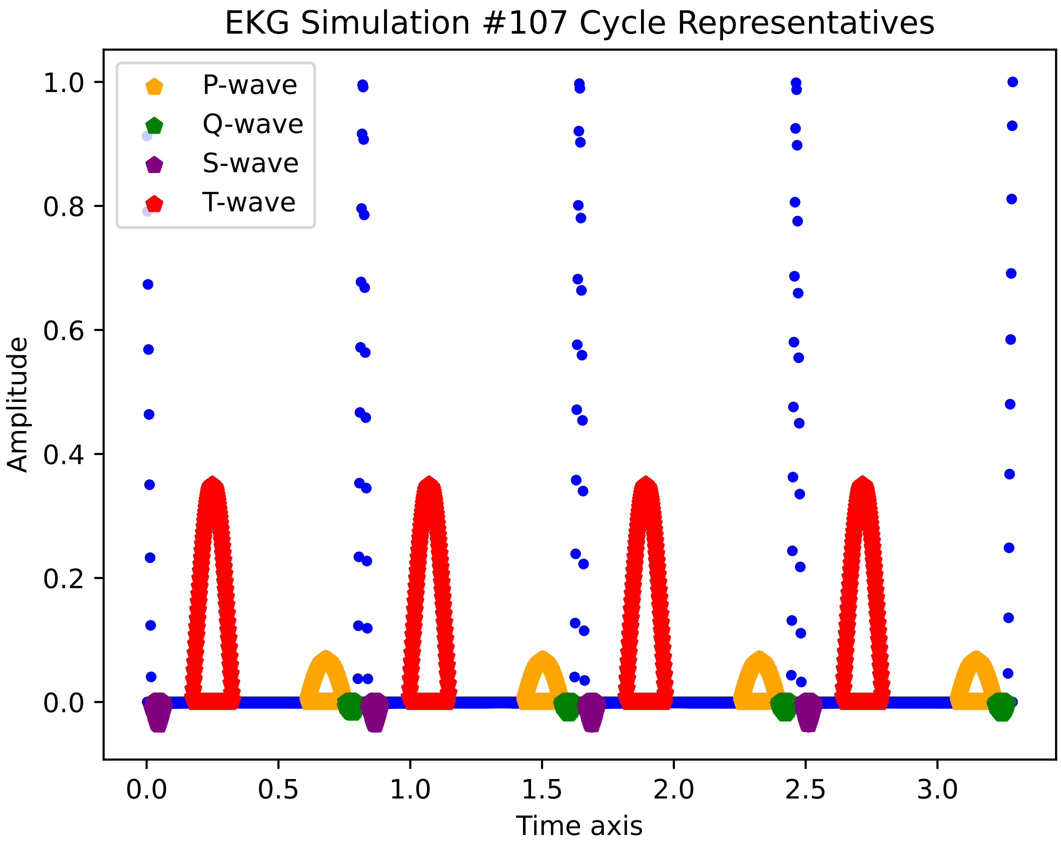
EKG Simulation #107 Cycle Representatives
P-wave
Q-wave
S-wave
T-wave
1.0
0.8
0.6
0.4
0.2
0.0
Amplitude
0.0
0.5
1.0
1.5
2.0
2.5
3.0
Time axis

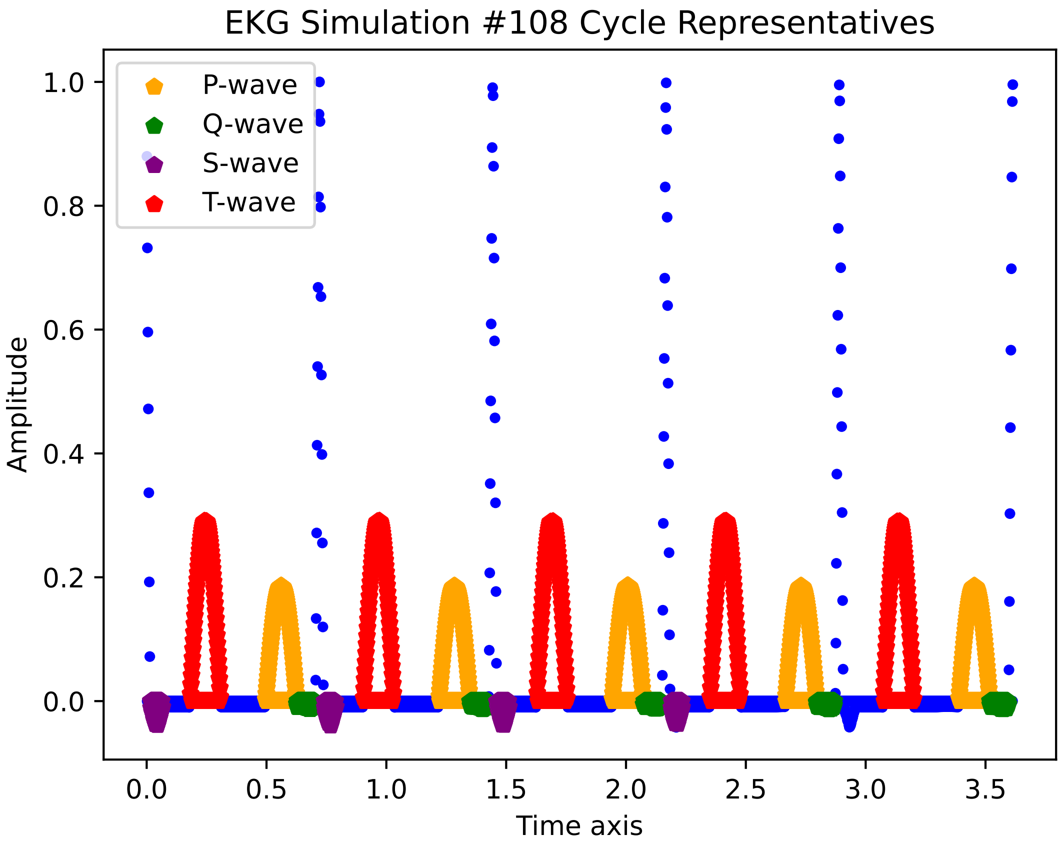

EKG Simulation #108 Cycle Representatives
P-wave
Q-wave
S-wave
T-wave
Amplitude
Time axis
1.0
0.8
0.6
0.4
0.2
0.0
0.0
0.5
1.0
1.5
2.0
2.5
3.0
3.5

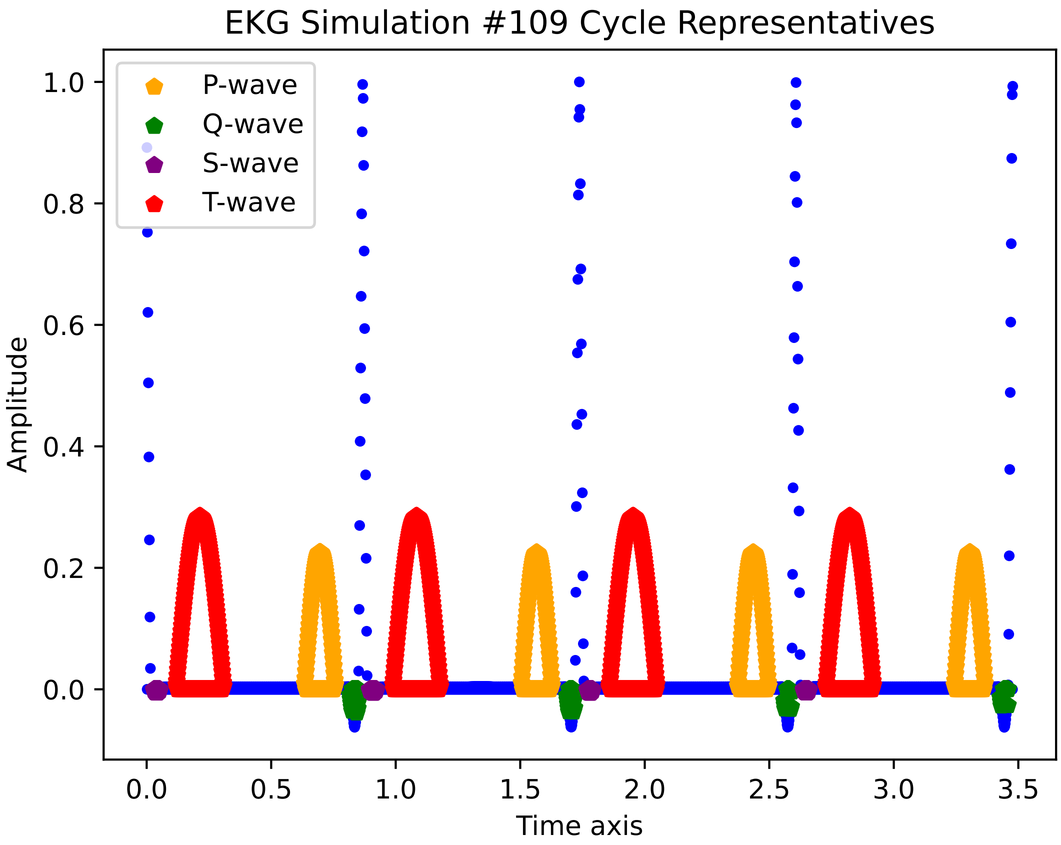
EKG Simulation #109 Cycle Representatives
P-wave
Q-wave
S-wave
T-wave
Amplitude
Time axis
1.0
0.8
0.6
0.4
0.2
0.0
0.0
0.5
1.0
1.5
2.0
2.5
3.0
3.5

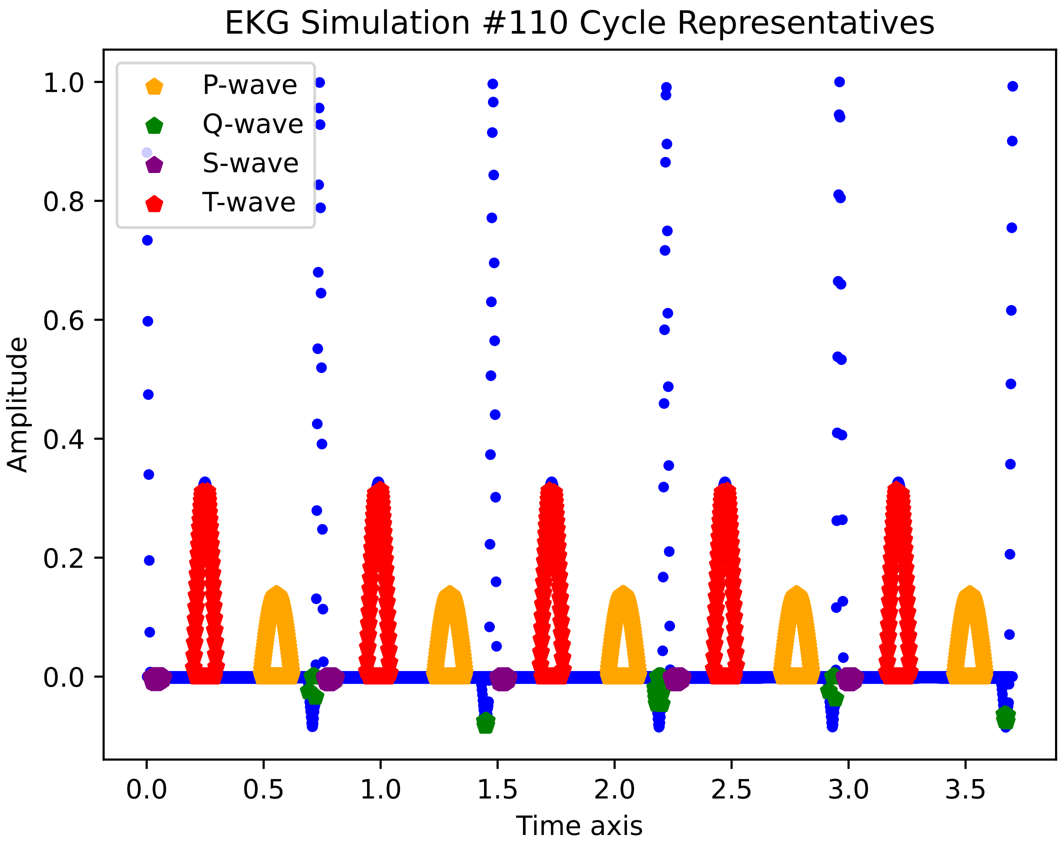

EKG Simulation #110 Cycle Representatives
P-wave
Q-wave
S-wave
T-wave
Amplitude
Time axis
1.0
0.8
0.6
0.4
0.2
0.0
0.0
0.5
1.0
1.5
2.0
2.5
3.0
3.5

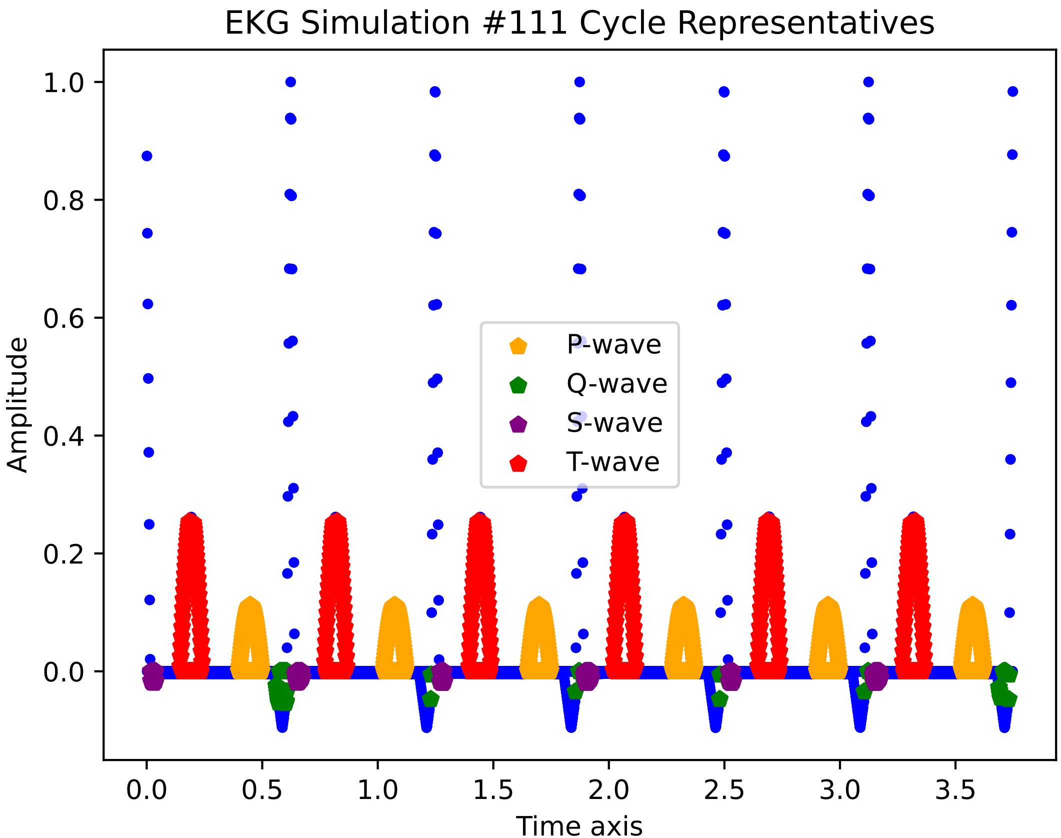

EKG Simulation #111 Cycle Representatives
Amplitude
1.0
0.8
0.6
0.4
0.2
0.0
0.0
0.5
1.0
1.5
2.0
2.5
3.0
3.5
Time axis
P-wave
Q-wave
S-wave
T-wave

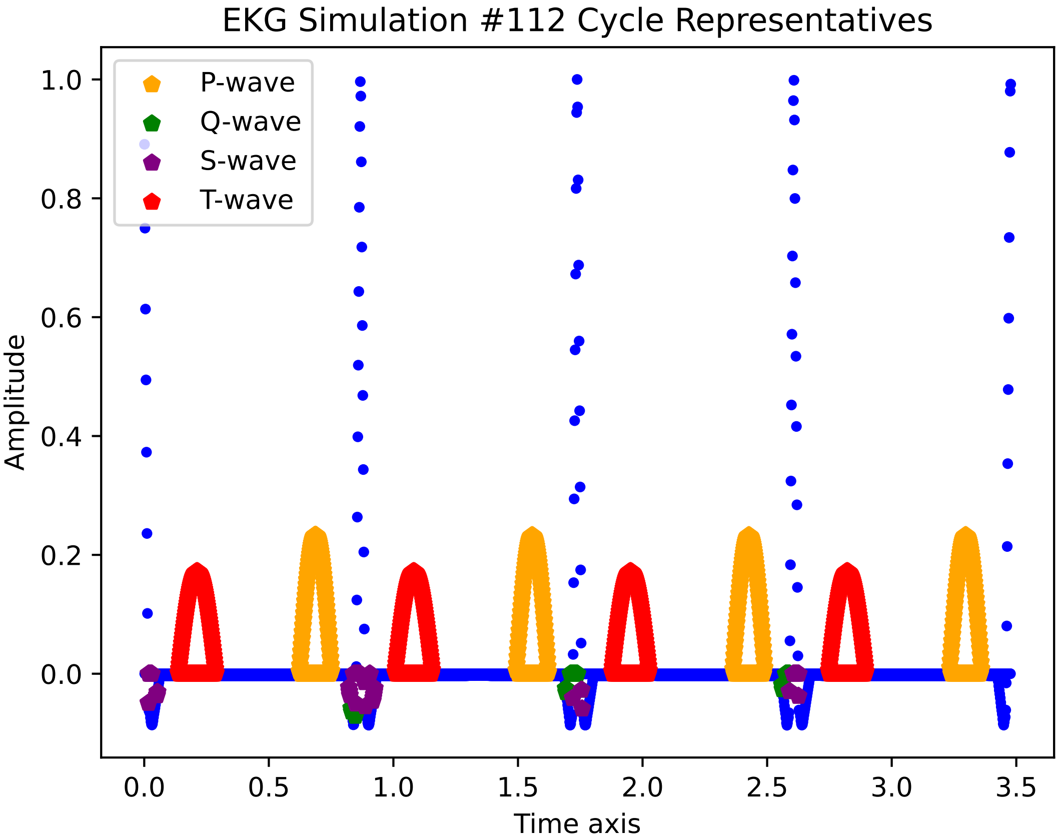
EKG Simulation #112 Cycle Representatives
P-wave
Q-wave
S-wave
T-wave
Amplitude
1.0
0.8
0.6
0.4
0.2
0.0
0.0
0.5
1.0
1.5
2.0
2.5
3.0
3.5
Time axis

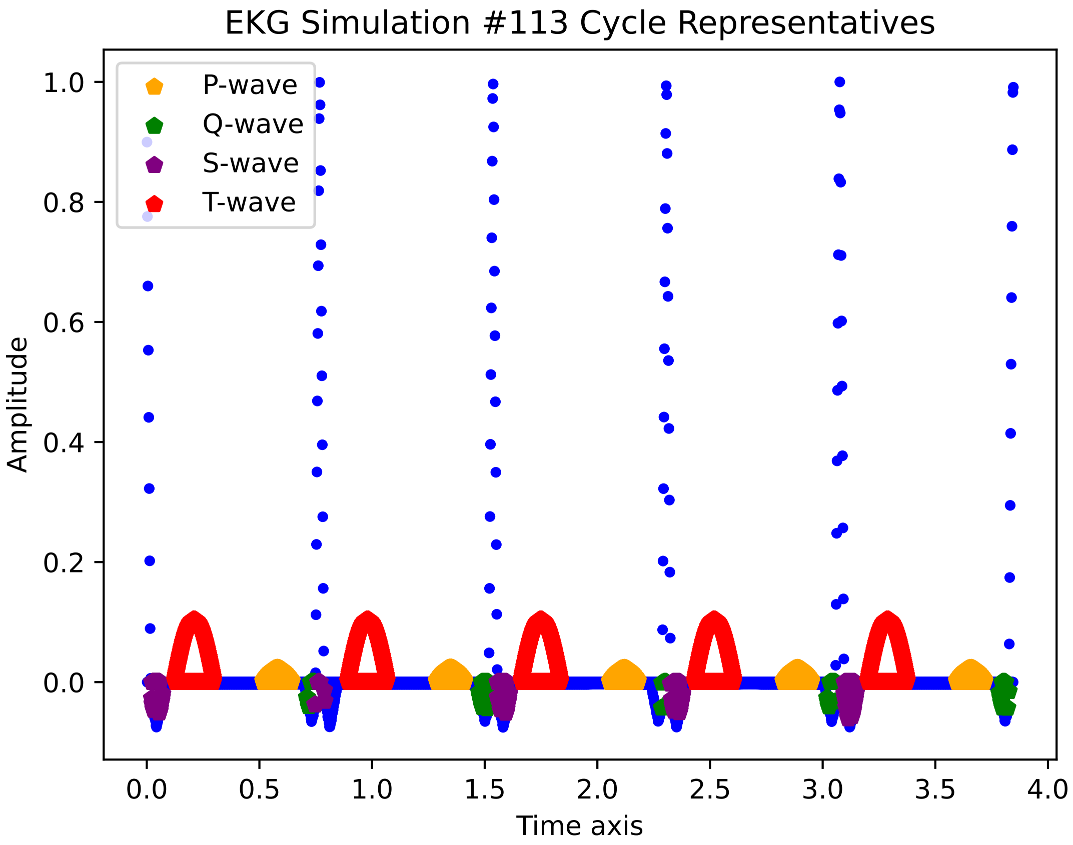

EKG Simulation #113 Cycle Representatives
P-wave
Q-wave
S-wave
T-wave
Amplitude
Time axis
1.0
0.8
0.6
0.4
0.2
0.0
0.0
0.5
1.0
1.5
2.0
2.5
3.0
3.5
4.0

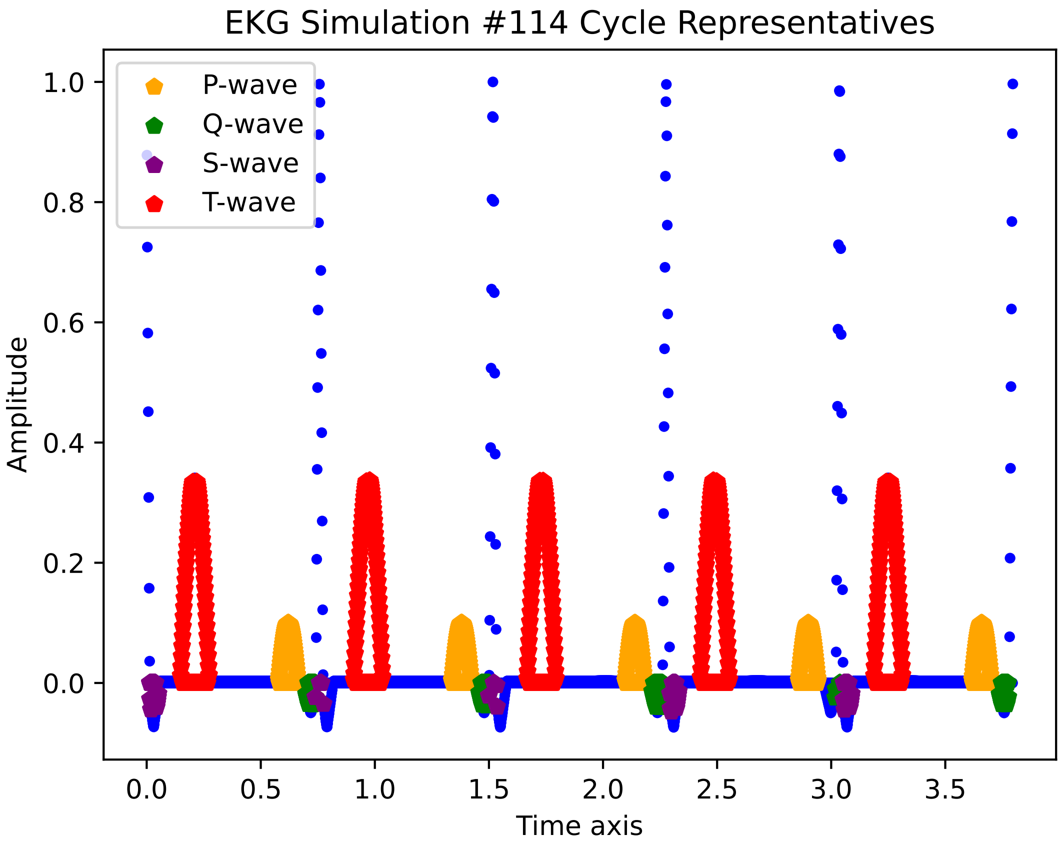
EKG Simulation #114 Cycle Representatives
P-wave
Q-wave
S-wave
T-wave
Amplitude
Time axis
1.0
0.8
0.6
0.4
0.2
0.0
0.0
0.5
1.0
1.5
2.0
2.5
3.0
3.5

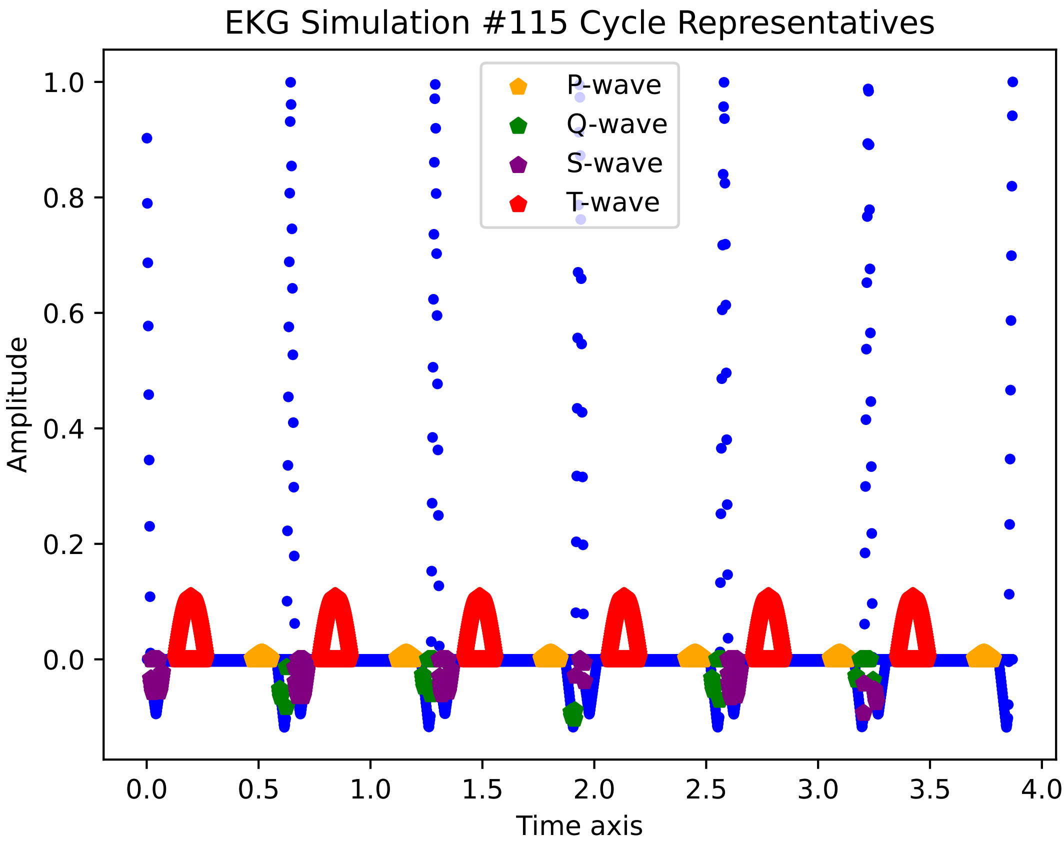
EKG Simulation #115 Cycle Representatives
P-wave
Q-wave
S-wave
T-wave
Amplitude
Time axis
1.0
0.8
0.6
0.4
0.2
0.0
0.0
0.5
1.0
1.5
2.0
2.5
3.0
3.5
4.0

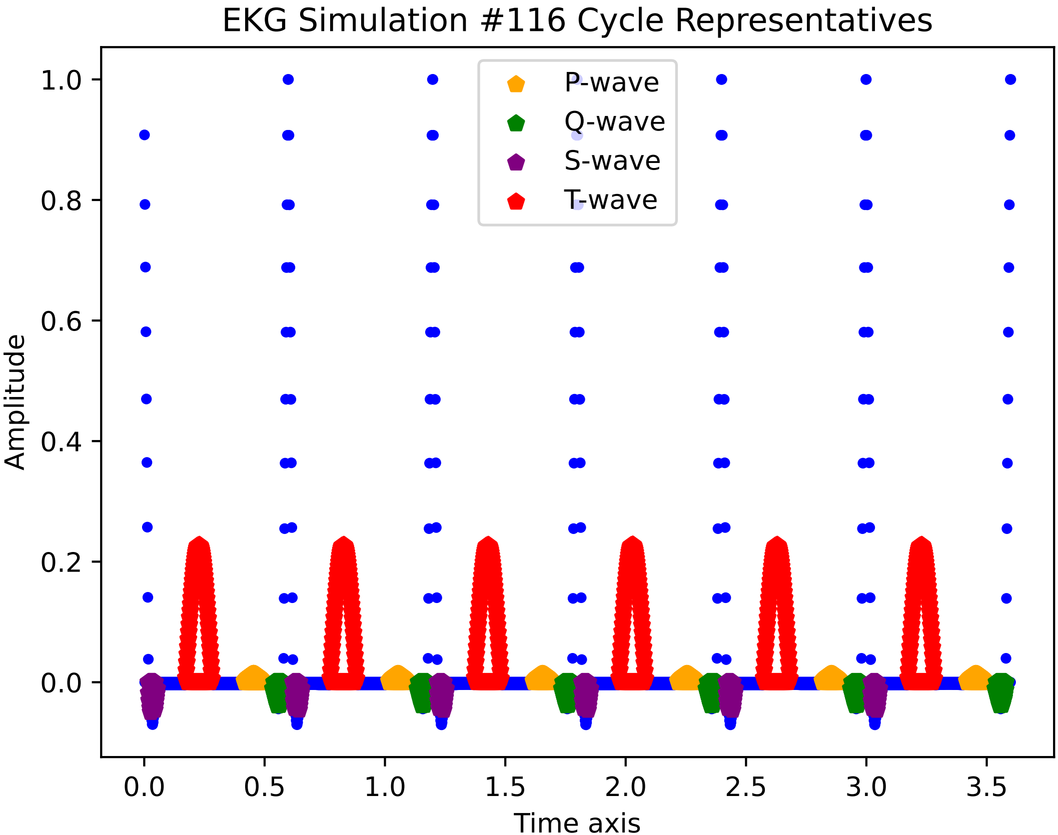
EKG Simulation #116 Cycle Representatives
P-wave
Q-wave
S-wave
T-wave
Amplitude
Time axis
1.0
0.8
0.6
0.4
0.2
0.0
0.0
0.5
1.0
1.5
2.0
2.5
3.0
3.5

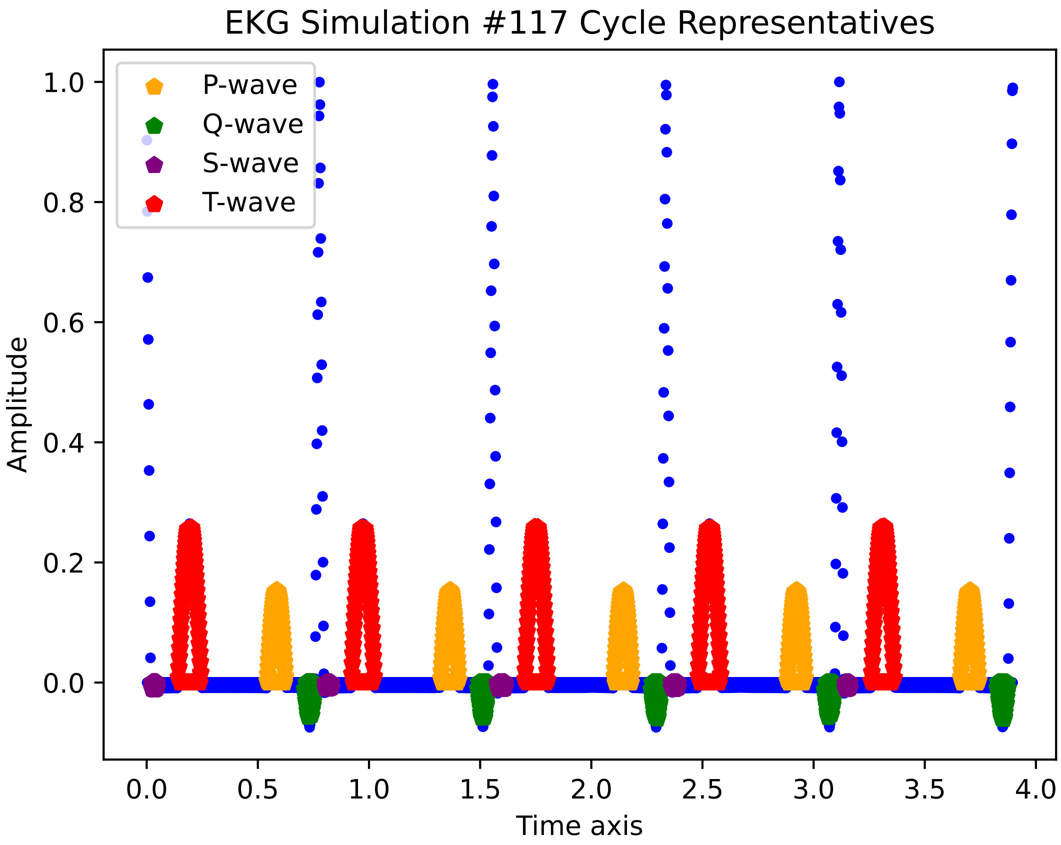
EKG Simulation #117 Cycle Representatives
P-wave
Q-wave
S-wave
T-wave
Amplitude
1.0
0.8
0.6
0.4
0.2
0.0
0.0
0.5
1.0
1.5
2.0
2.5
3.0
3.5
4.0
Time axis

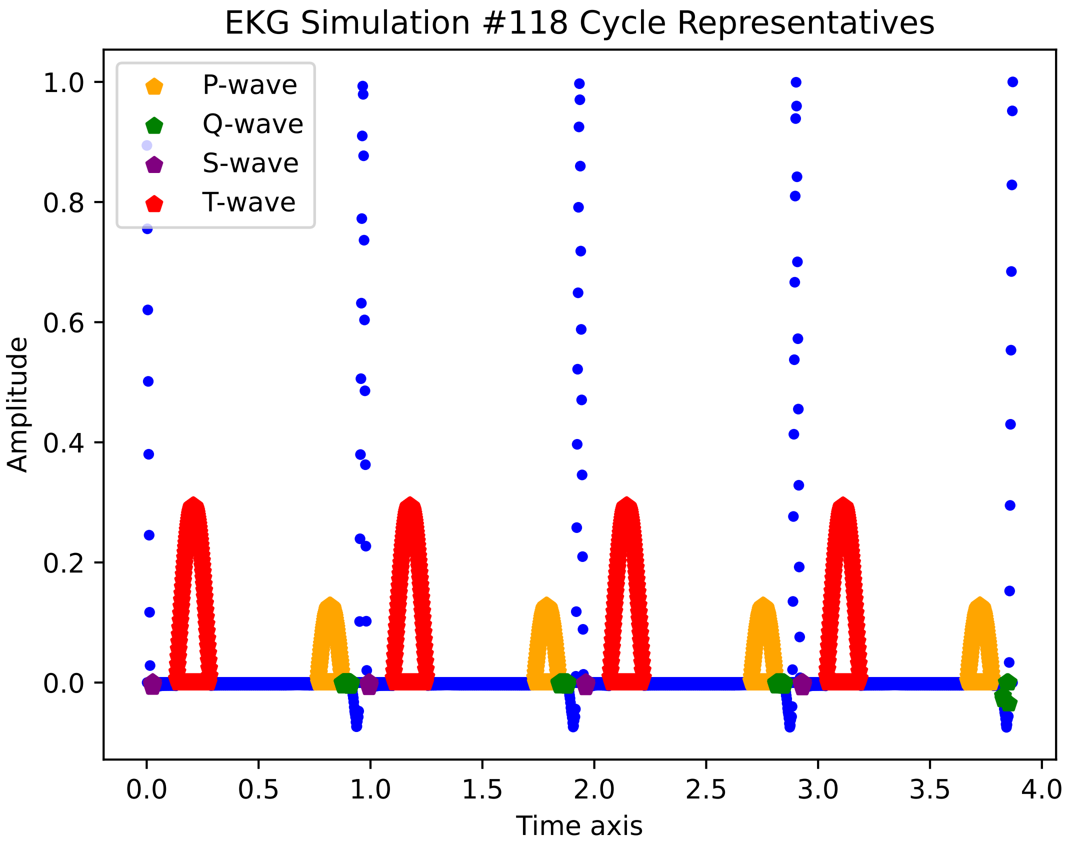
EKG Simulation #118 Cycle Representatives
P-wave
Q-wave
S-wave
T-wave
Amplitude
Time axis
1.0
0.8
0.6
0.4
0.2
0.0
0.0
0.5
1.0
1.5
2.0
2.5
3.0
3.5
4.0

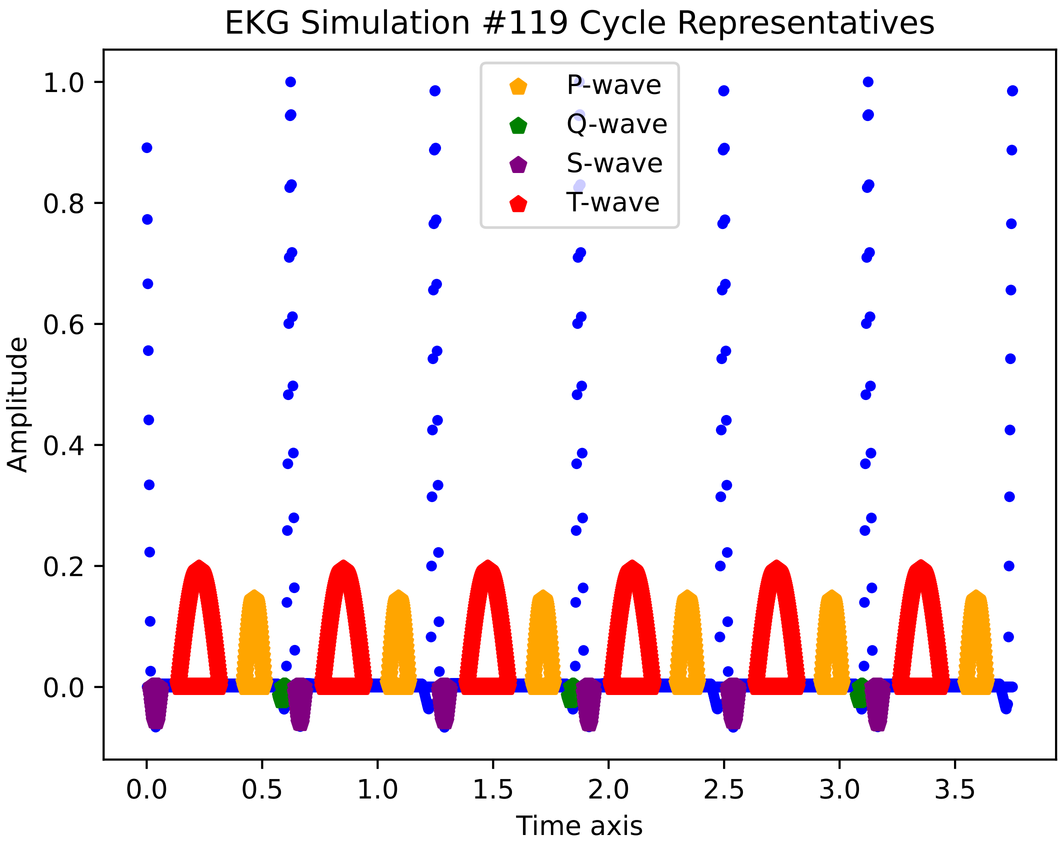
EKG Simulation #119 Cycle Representatives
P-wave
Q-wave
S-wave
T-wave
Amplitude
Time axis
1.0
0.8
0.6
0.4
0.2
0.0
0.0
0.5
1.0
1.5
2.0
2.5
3.0
3.5

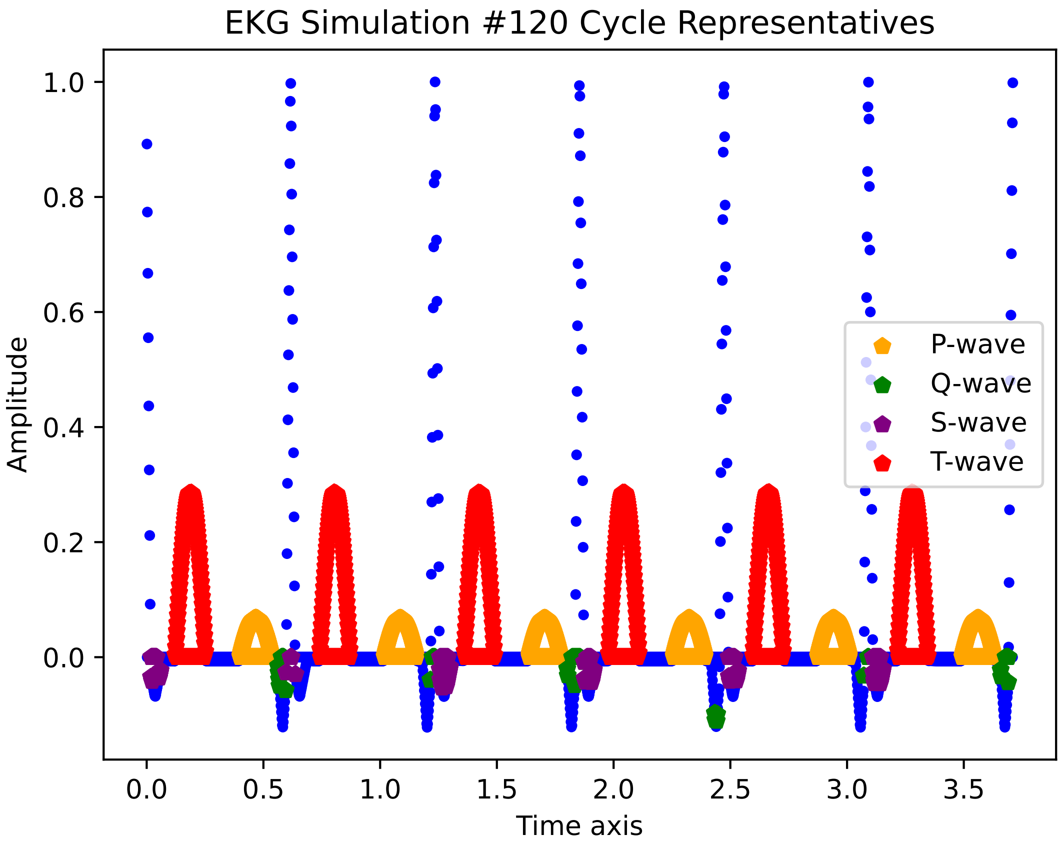

EKG Simulation #120 Cycle Representatives
Amplitude
Time axis
1.0
0.8
0.6
0.4
0.2
0.0
0.0
0.5
1.0
1.5
2.0
2.5
3.0
3.5
P-wave
Q-wave
S-wave
T-wave

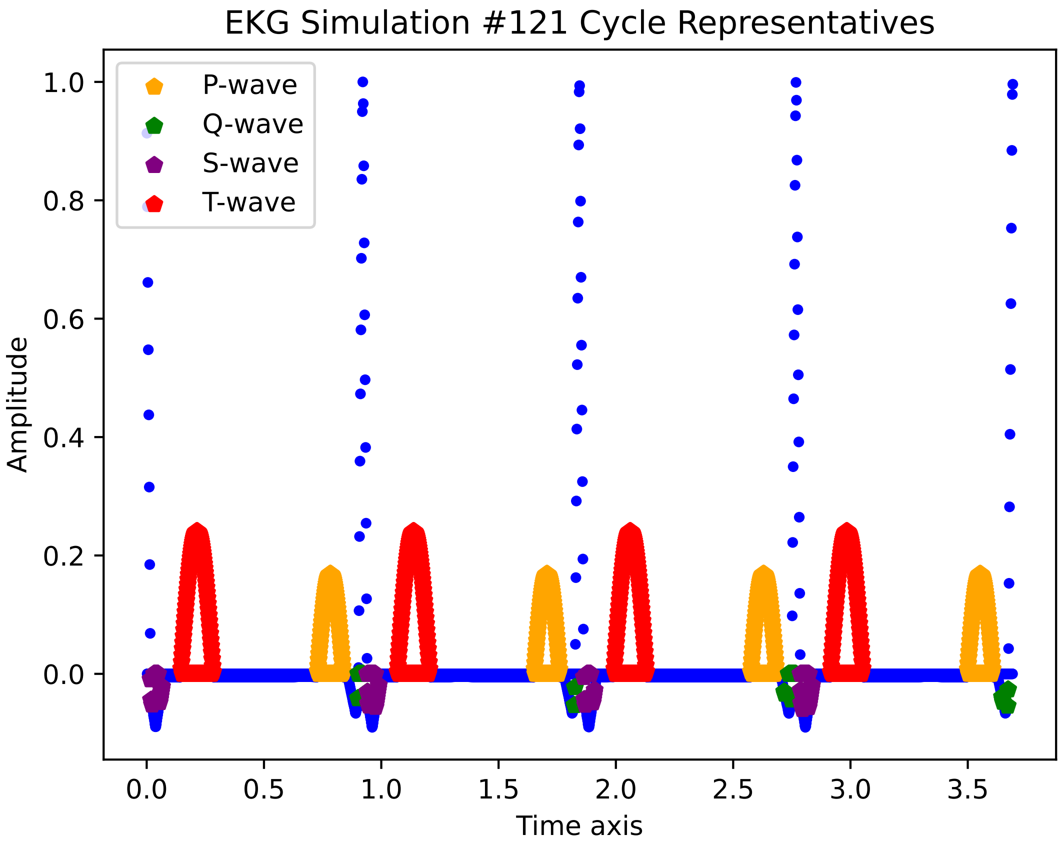

EKG Simulation #121 Cycle Representatives
P-wave
Q-wave
S-wave
T-wave
Amplitude
Time axis
1.0
0.8
0.6
0.4
0.2
0.0
0.0
0.5
1.0
1.5
2.0
2.5
3.0
3.5

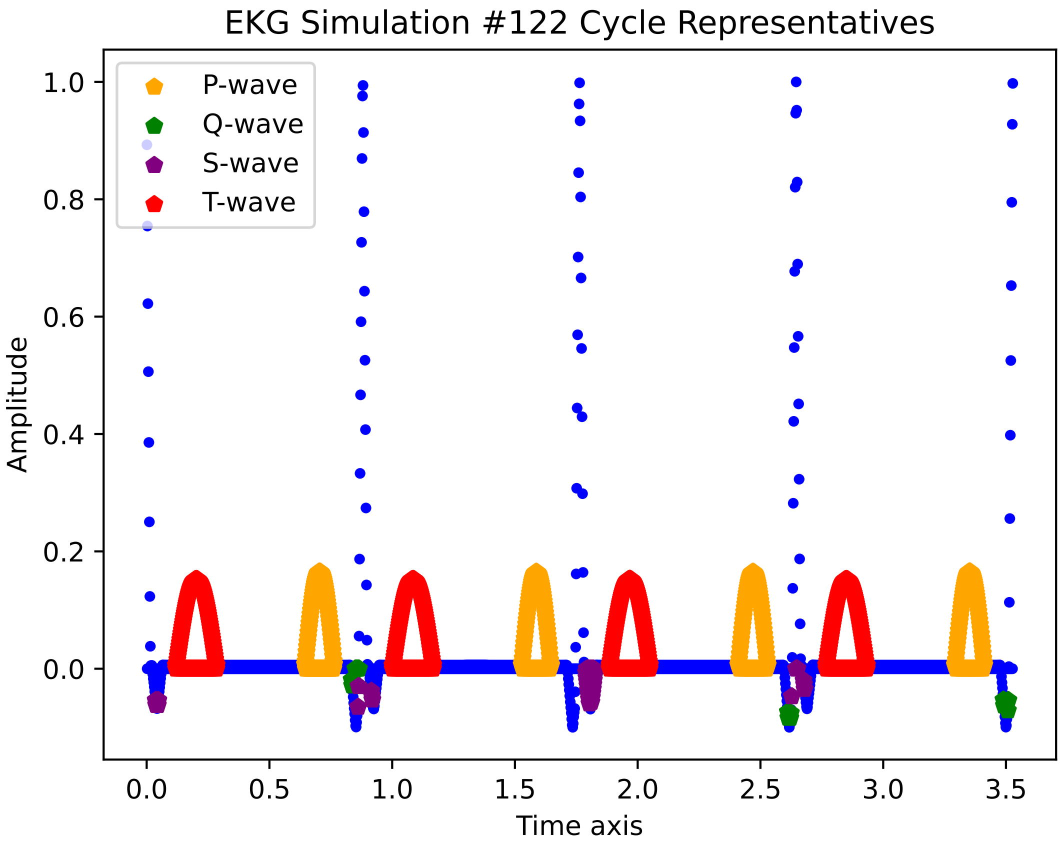

EKG Simulation #122 Cycle Representatives
P-wave
Q-wave
S-wave
T-wave
1.0
0.8
0.6
0.4
0.2
0.0
Amplitude
0.0
0.5
1.0
1.5
2.0
2.5
3.0
3.5
Time axis

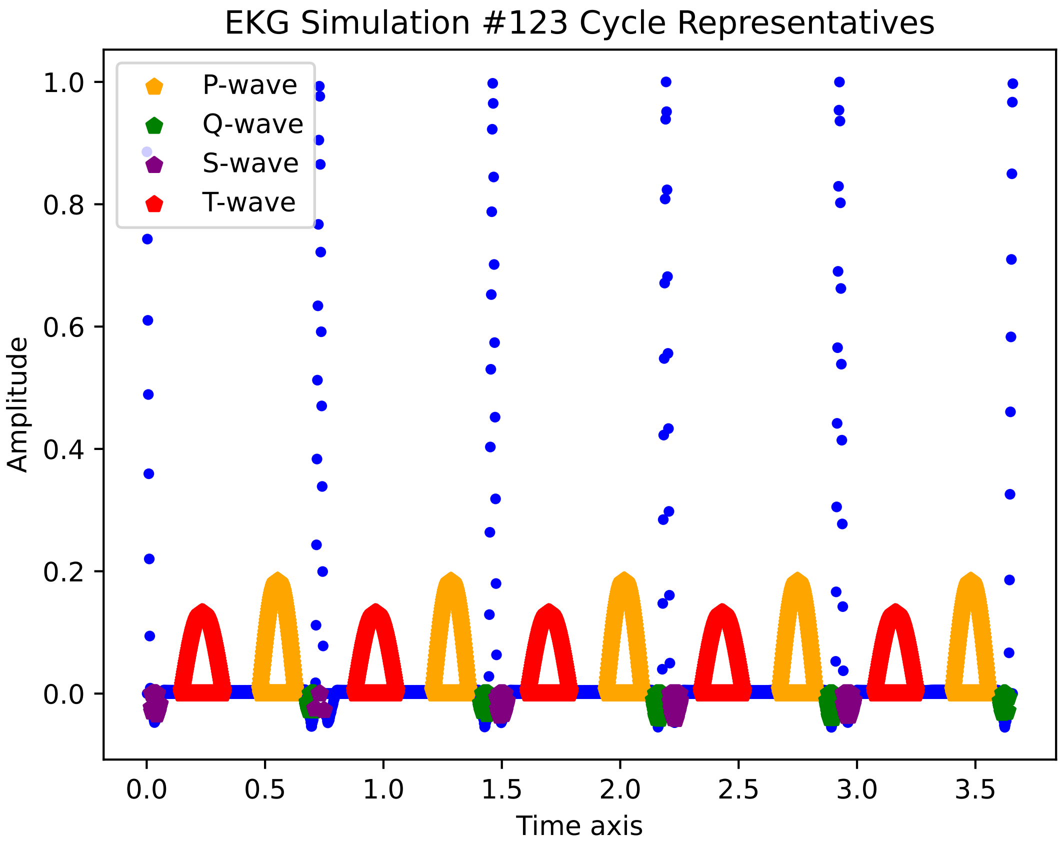

EKG Simulation #123 Cycle Representatives
P-wave
Q-wave
S-wave
T-wave
Amplitude
1.0
0.8
0.6
0.4
0.2
0.0
0.0
0.5
1.0
1.5
2.0
2.5
3.0
3.5
Time axis

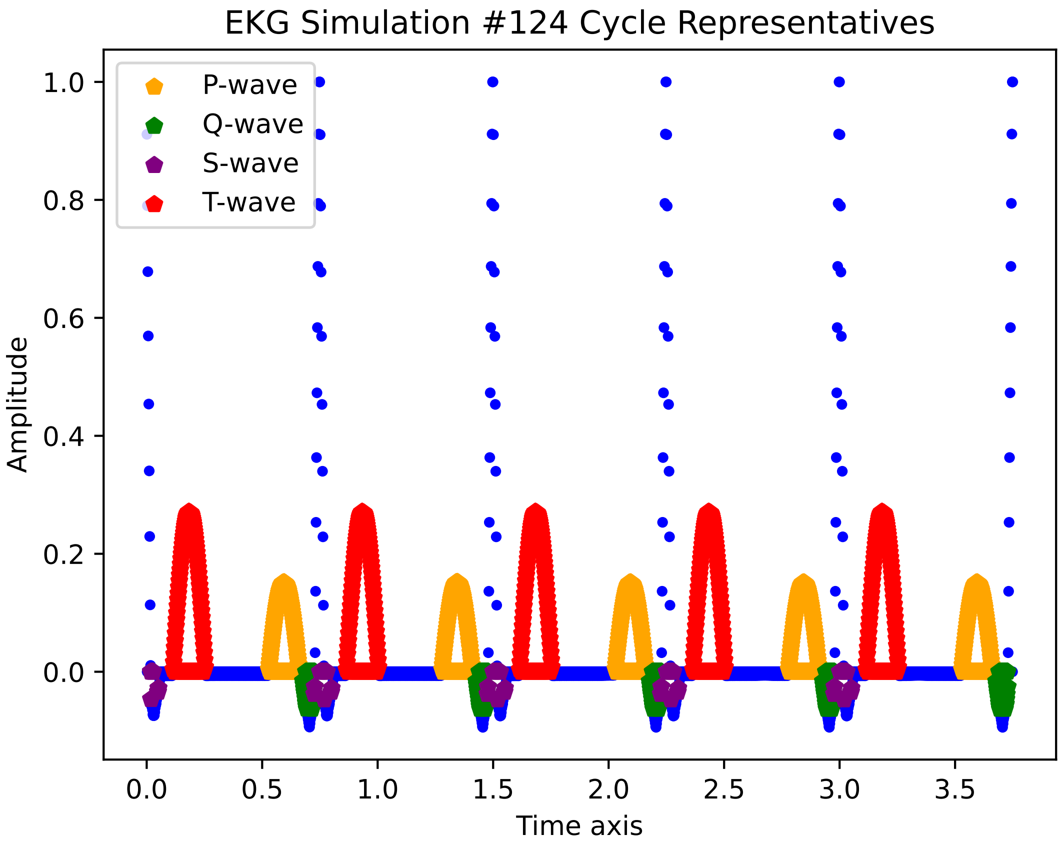

EKG Simulation #124 Cycle Representatives
P-wave
Q-wave
S-wave
T-wave
1.0
0.8
0.6
0.4
0.2
0.0
Amplitude
0.0
0.5
1.0
1.5
2.0
2.5
3.0
3.5
Time axis

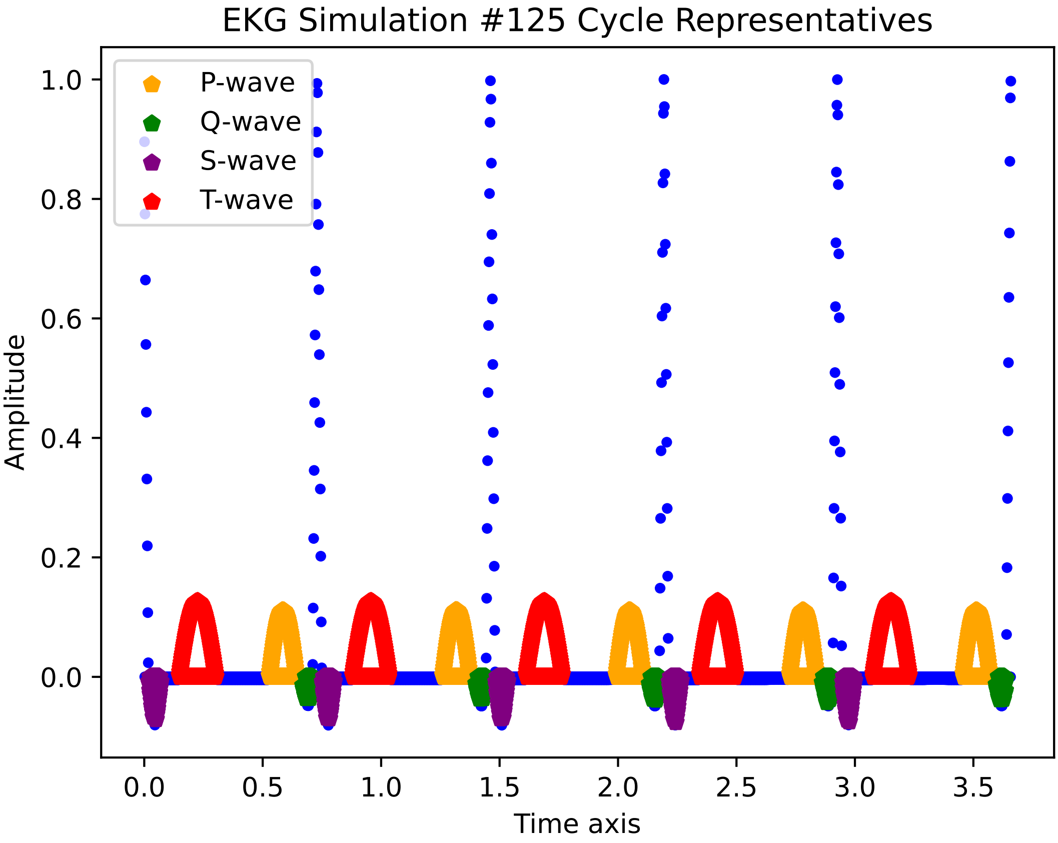

EKG Simulation #125 Cycle Representatives
P-wave
Q-wave
S-wave
T-wave
Amplitude
Time axis
1.0
0.8
0.6
0.4
0.2
0.0
0.0
0.5
1.0
1.5
2.0
2.5
3.0
3.5

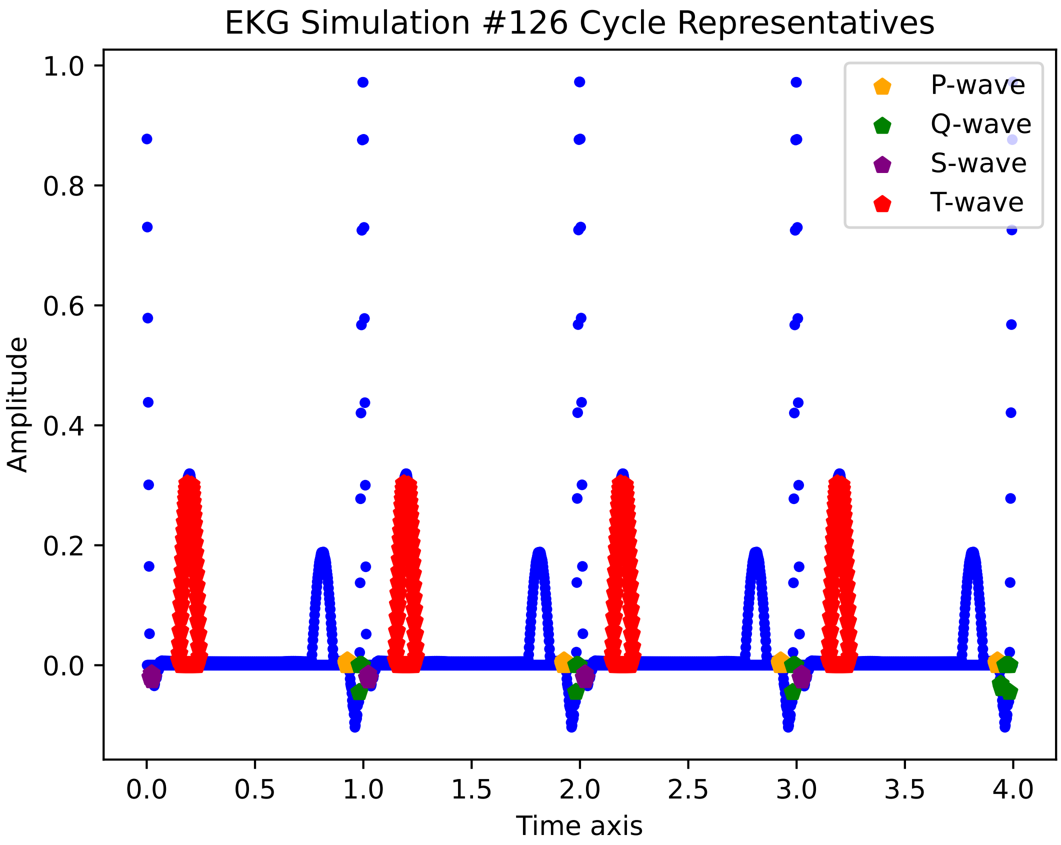
EKG Simulation #126 Cycle Representatives
Amplitude
Time axis
1.0
0.8
0.6
0.4
0.2
0.0
0.0
0.5
1.0
1.5
2.0
2.5
3.0
3.5
4.0
P-wave
Q-wave
S-wave
T-wave

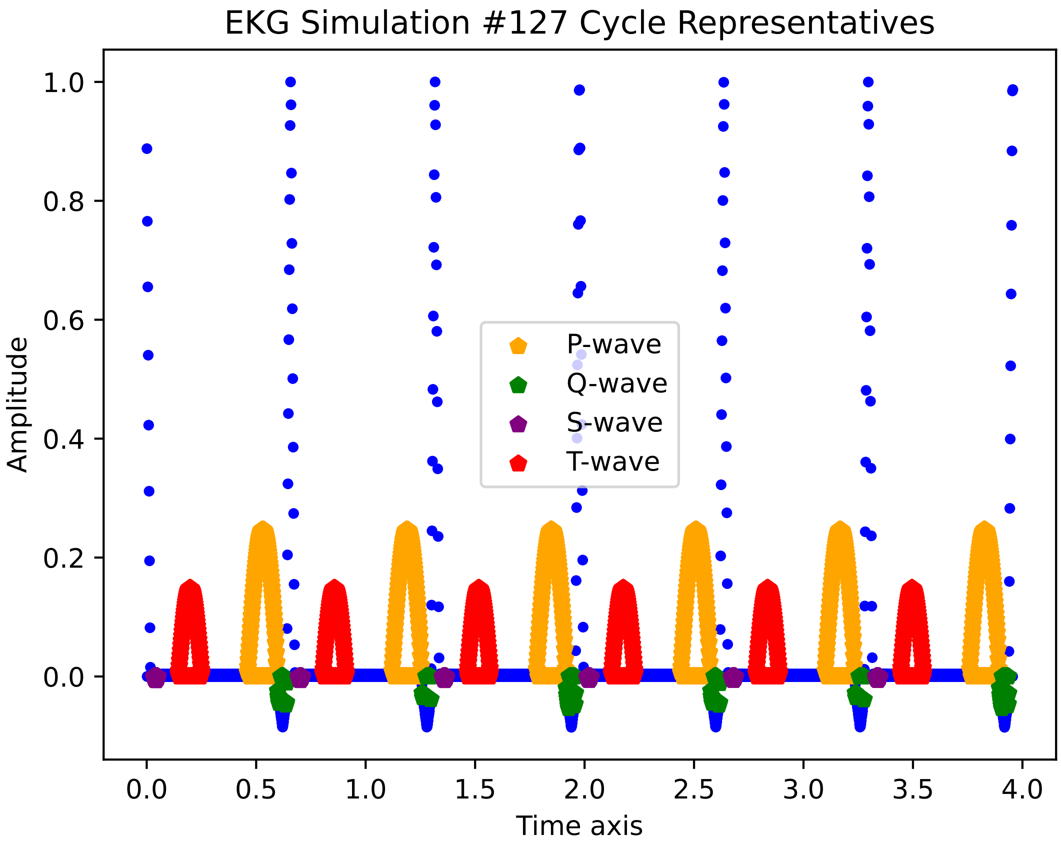

EKG Simulation #127 Cycle Representatives
Amplitude
Time axis
1.0
0.8
0.6
0.4
0.2
0.0
0.0
0.5
1.0
1.5
2.0
2.5
3.0
3.5
4.0
P-wave
Q-wave
S-wave
T-wave

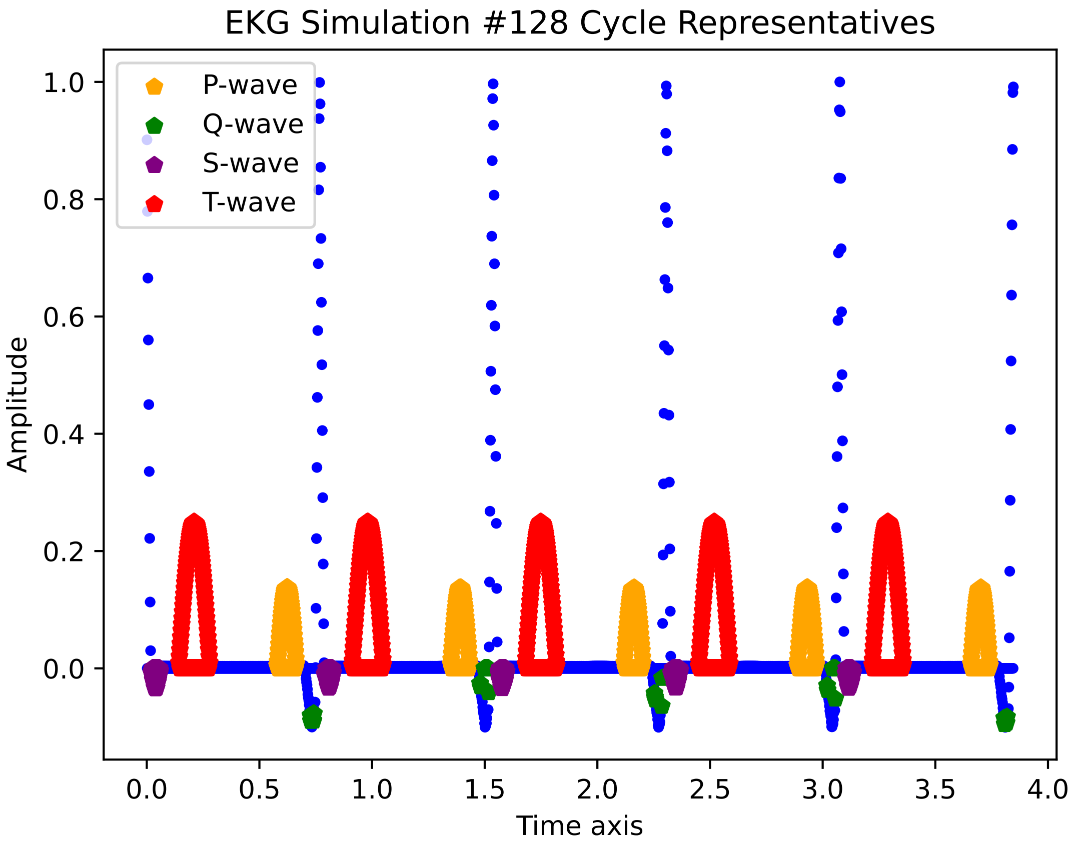
EKG Simulation #128 Cycle Representatives
P-wave
Q-wave
S-wave
T-wave
Amplitude
1.0
0.8
0.6
0.4
0.2
0.0
0.0
0.5
1.0
1.5
2.0
2.5
3.0
3.5
4.0
Time axis

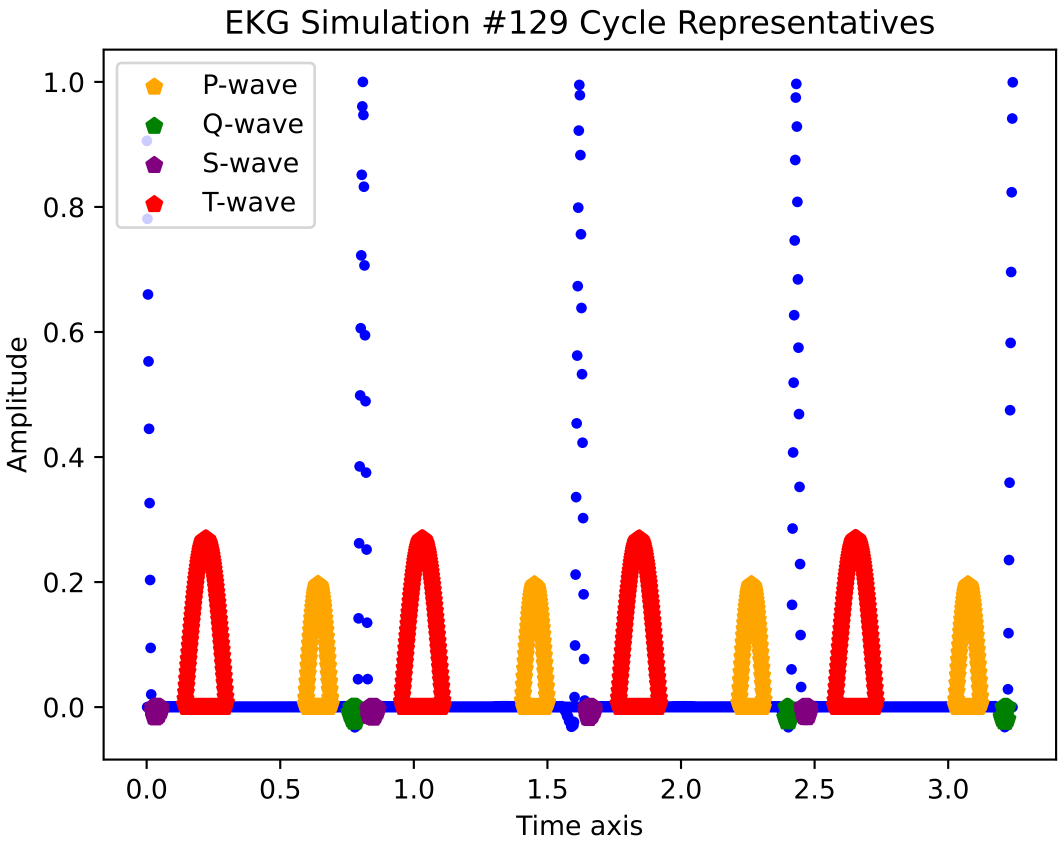
EKG Simulation #129 Cycle Representatives
P-wave
Q-wave
S-wave
T-wave
Amplitude
Time axis
1.0
0.8
0.6
0.4
0.2
0.0
0.0
0.5
1.0
1.5
2.0
2.5
3.0

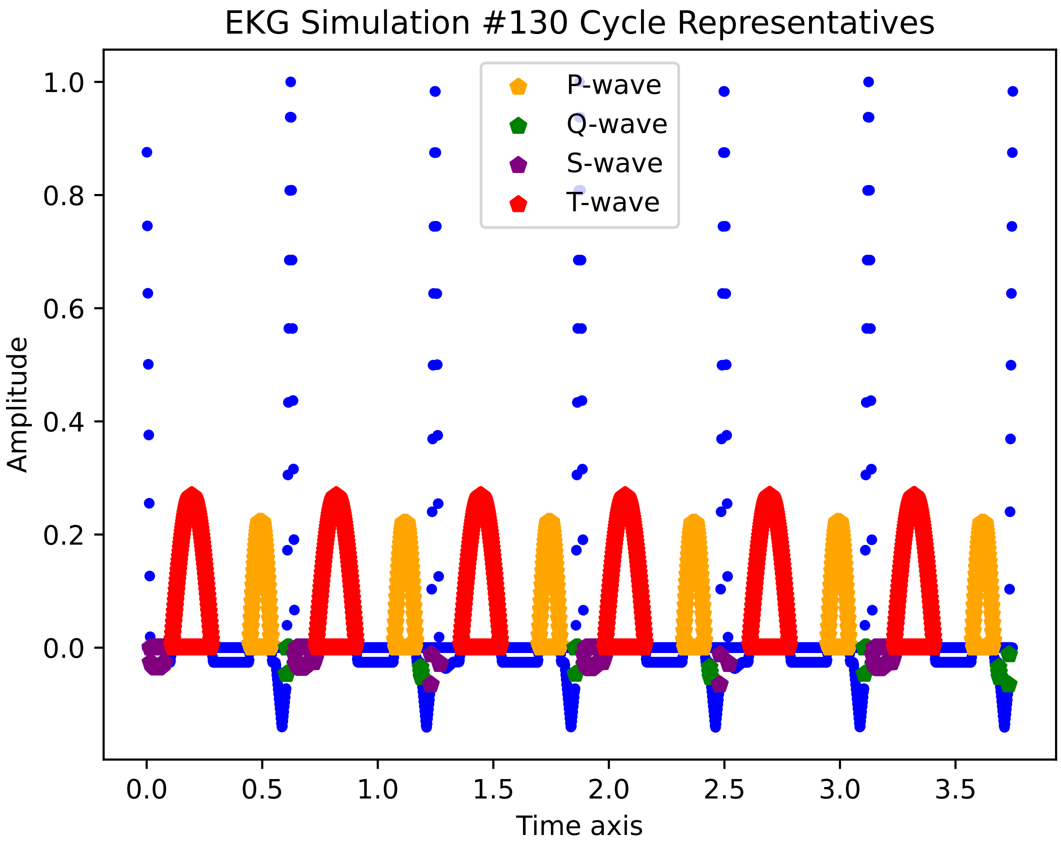
EKG Simulation #130 Cycle Representatives
P-wave
Q-wave
S-wave
T-wave
Amplitude
Time axis
1.0
0.8
0.6
0.4
0.2
0.0
0.0
0.5
1.0
1.5
2.0
2.5
3.0
3.5

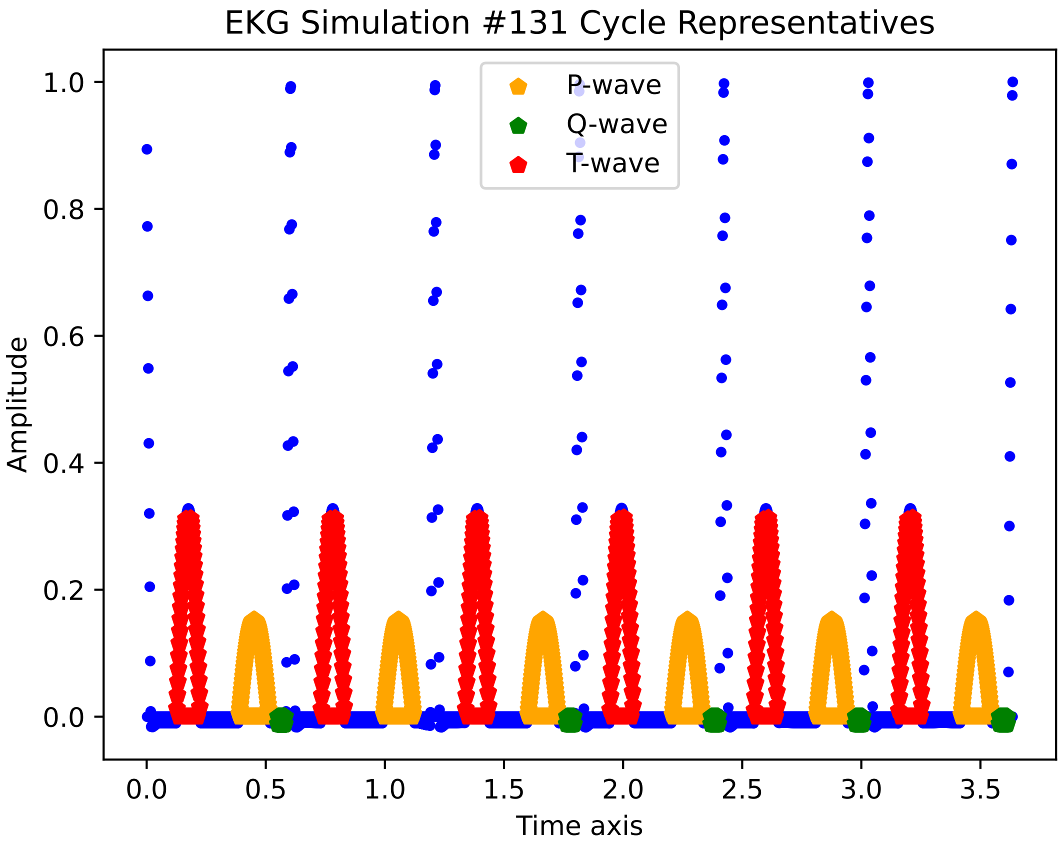
EKG Simulation #131 Cycle Representatives
P-wave
Q-wave
T-wave
Amplitude
Time axis
0.0
0.2
0.4
0.6
0.8
1.0
0.0
0.5
1.0
1.5
2.0
2.5
3.0
3.5

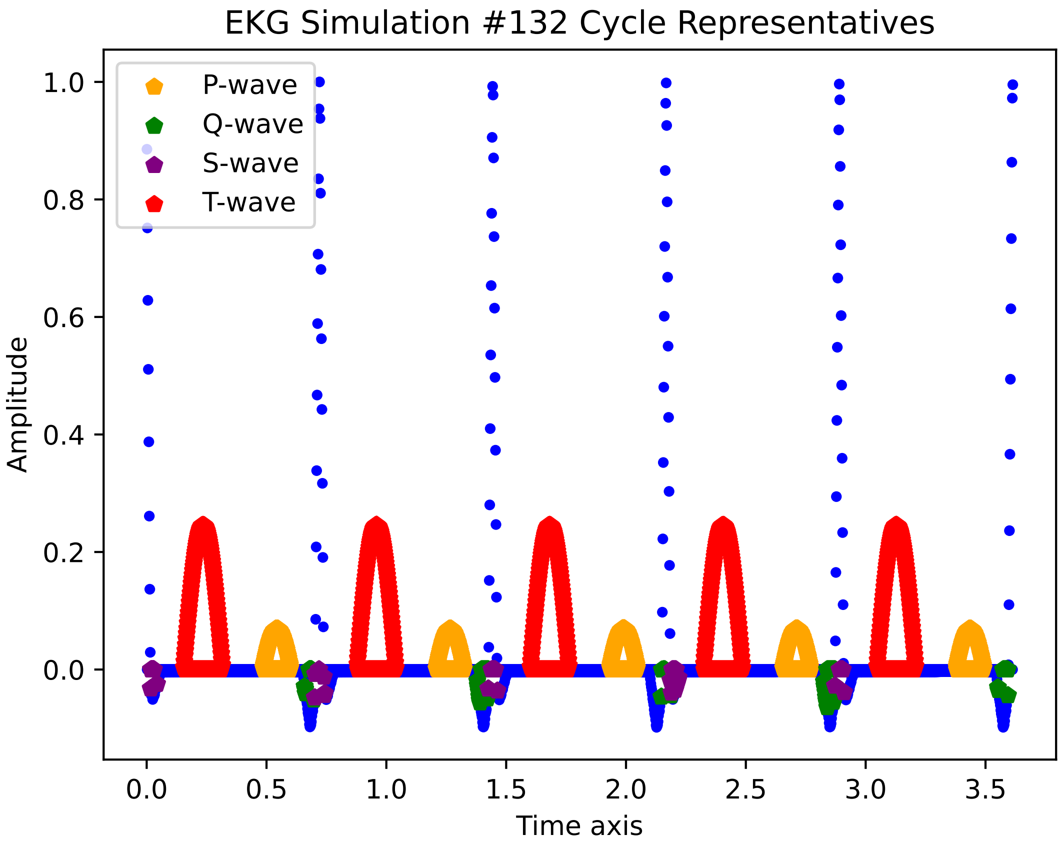

EKG Simulation #132 Cycle Representatives
P-wave
Q-wave
S-wave
T-wave
Amplitude
Time axis
0.0
0.2
0.4
0.6
0.8
1.0
0.0
0.5
1.0
1.5
2.0
2.5
3.0
3.5

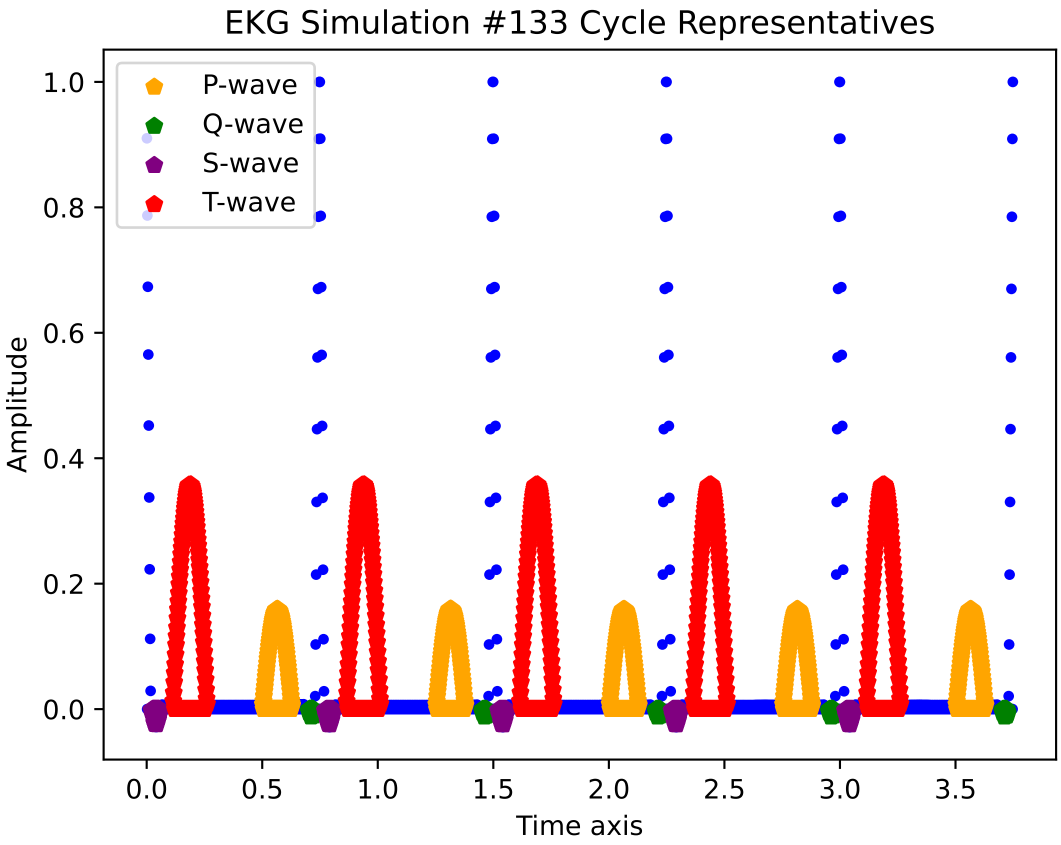

EKG Simulation #133 Cycle Representatives
P-wave
Q-wave
S-wave
T-wave
Amplitude
1.0
0.8
0.6
0.4
0.2
0.0
0.0
0.5
1.0
1.5
2.0
2.5
3.0
3.5
Time axis

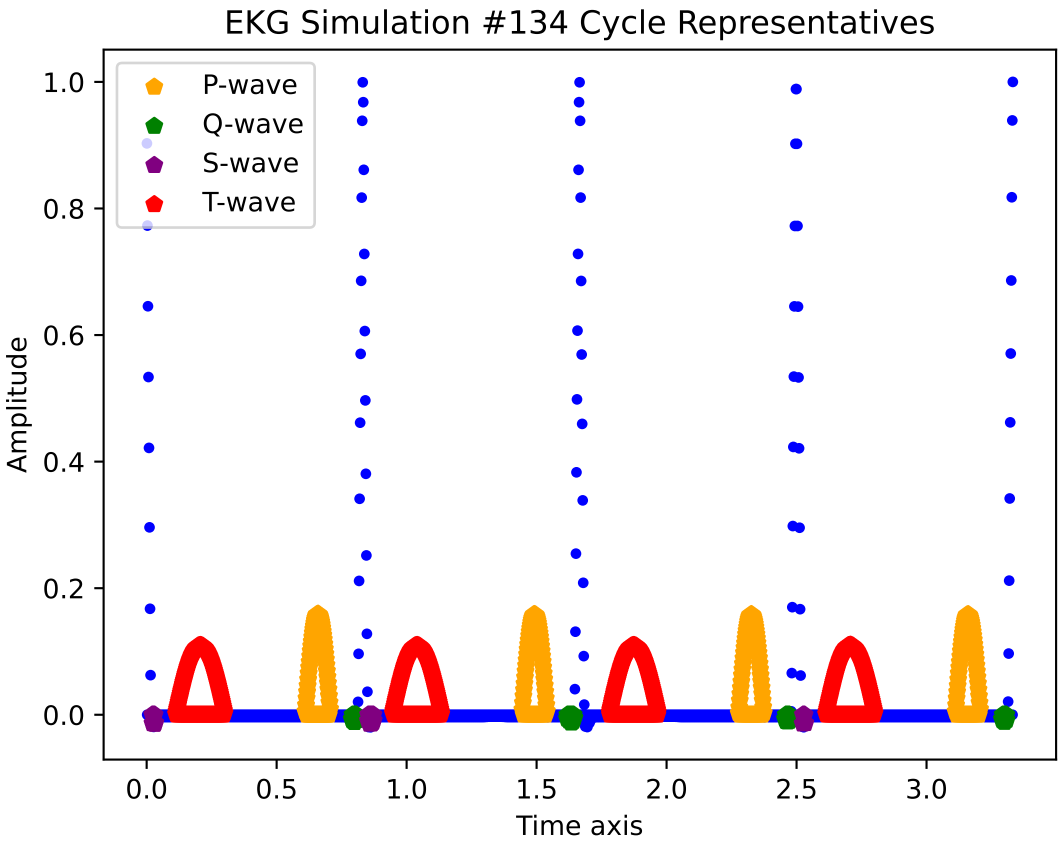
EKG Simulation #134 Cycle Representatives
P-wave
Q-wave
S-wave
T-wave
Amplitude
0.0
0.2
0.4
0.6
0.8
1.0
0.0
0.5
1.0
1.5
2.0
2.5
3.0
Time axis

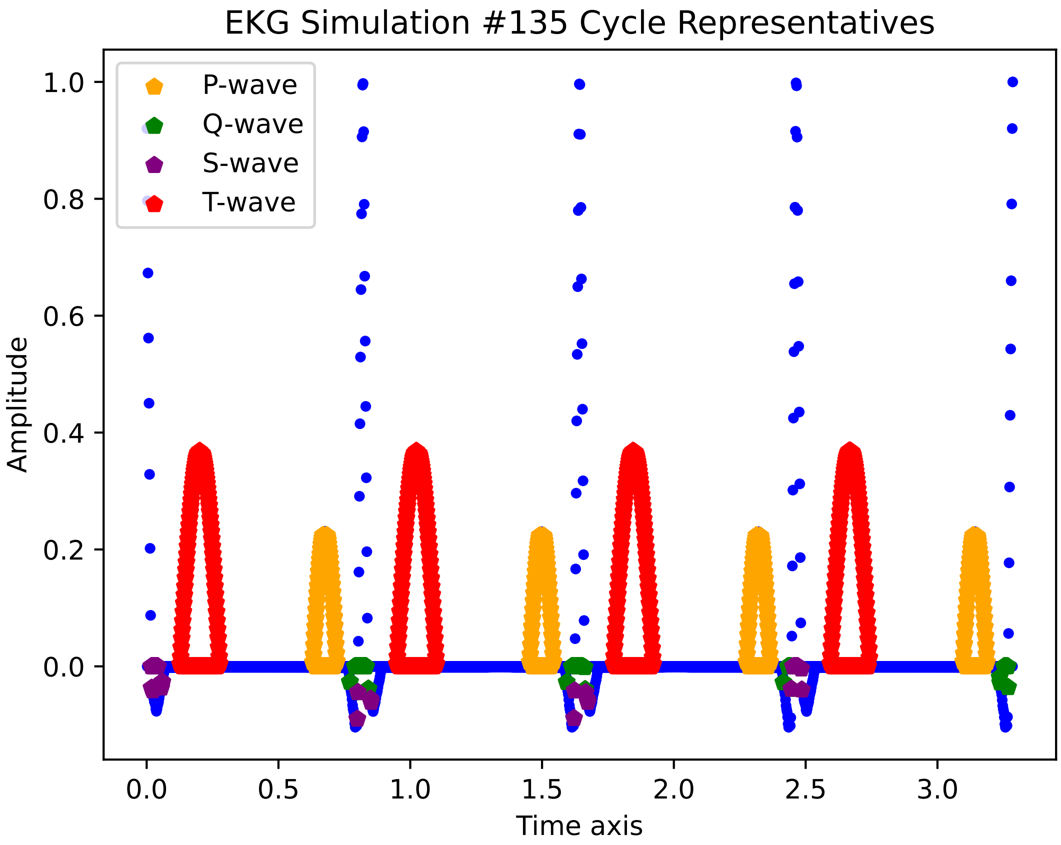

EKG Simulation #135 Cycle Representatives
P-wave
Q-wave
S-wave
T-wave
1.0
0.8
0.6
0.4
0.2
0.0
Amplitude
0.0
0.5
1.0
1.5
2.0
2.5
3.0
Time axis

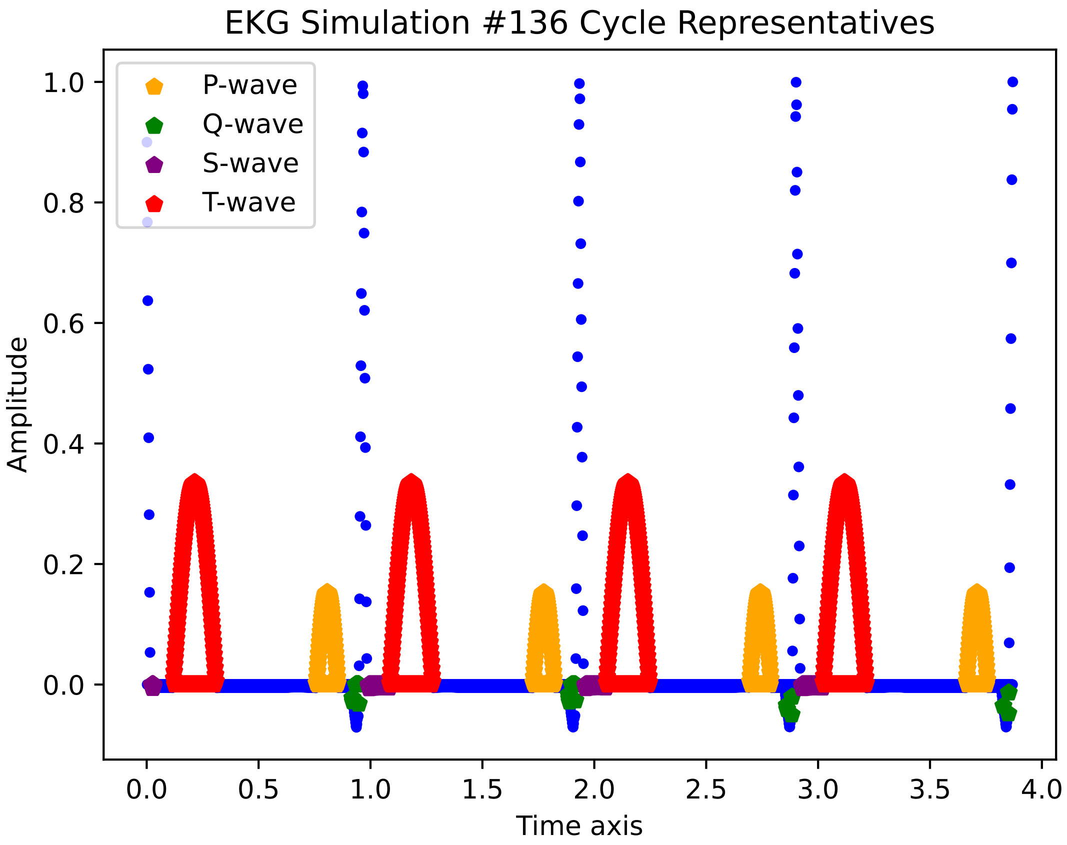
EKG Simulation #136 Cycle Representatives
P-wave
Q-wave
S-wave
T-wave
1.0
0.8
0.6
0.4
0.2
0.0
Amplitude
0.0
0.5
1.0
1.5
2.0
2.5
3.0
3.5
4.0
Time axis

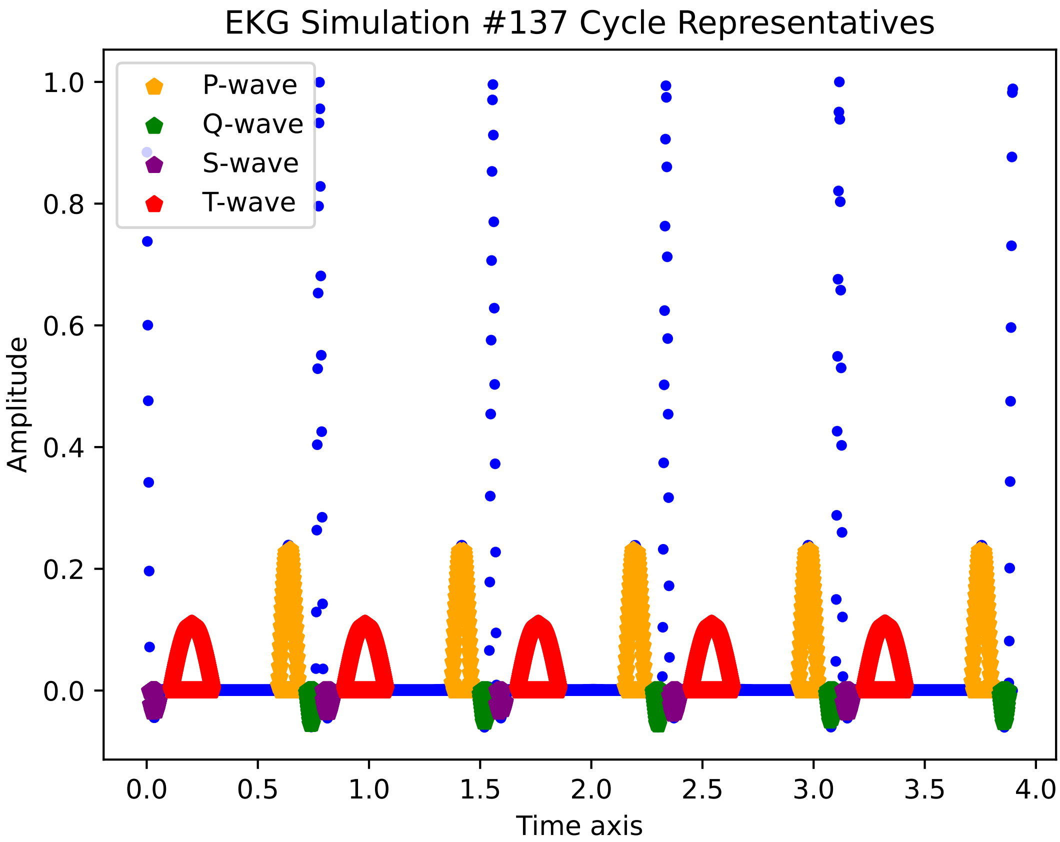
EKG Simulation #137 Cycle Representatives
P-wave
Q-wave
S-wave
T-wave
Amplitude
Time axis
0.0
0.2
0.4
0.6
0.8
1.0
0.0
0.5
1.0
1.5
2.0
2.5
3.0
3.5
4.0

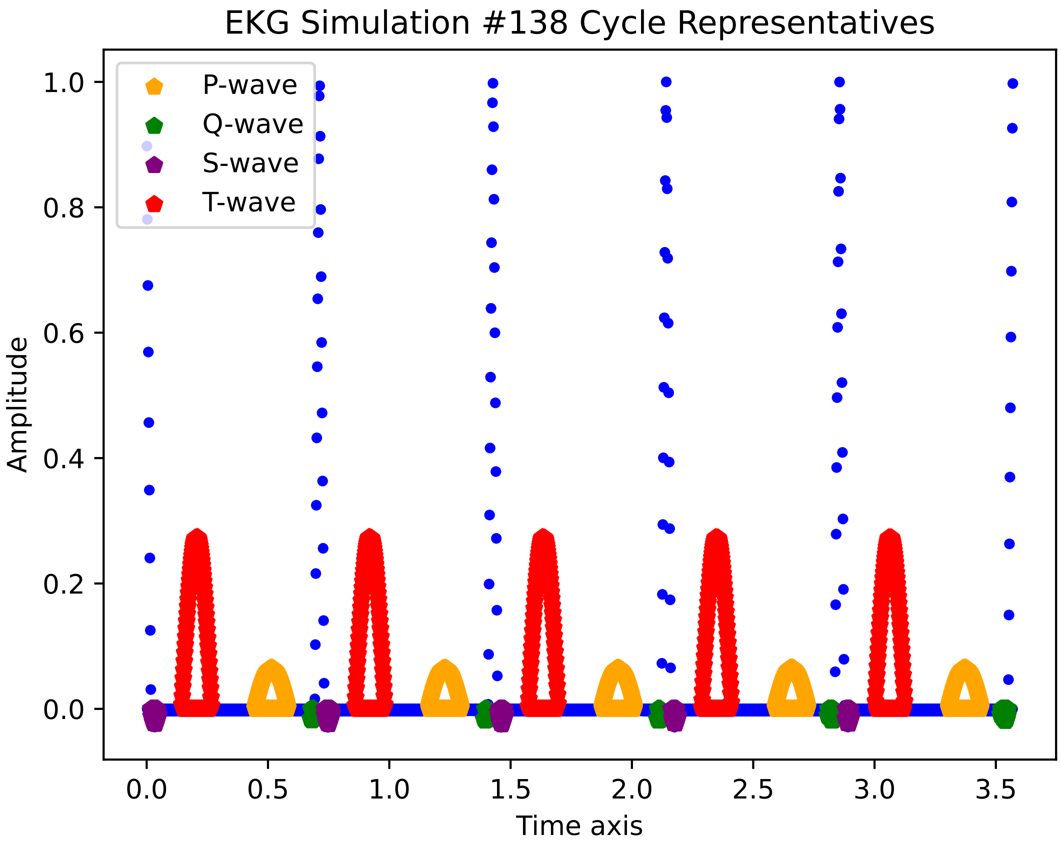
EKG Simulation #138 Cycle Representatives
P-wave
Q-wave
S-wave
T-wave
Amplitude
Time axis
0.0
0.2
0.4
0.6
0.8
1.0
0.0
0.5
1.0
1.5
2.0
2.5
3.0
3.5

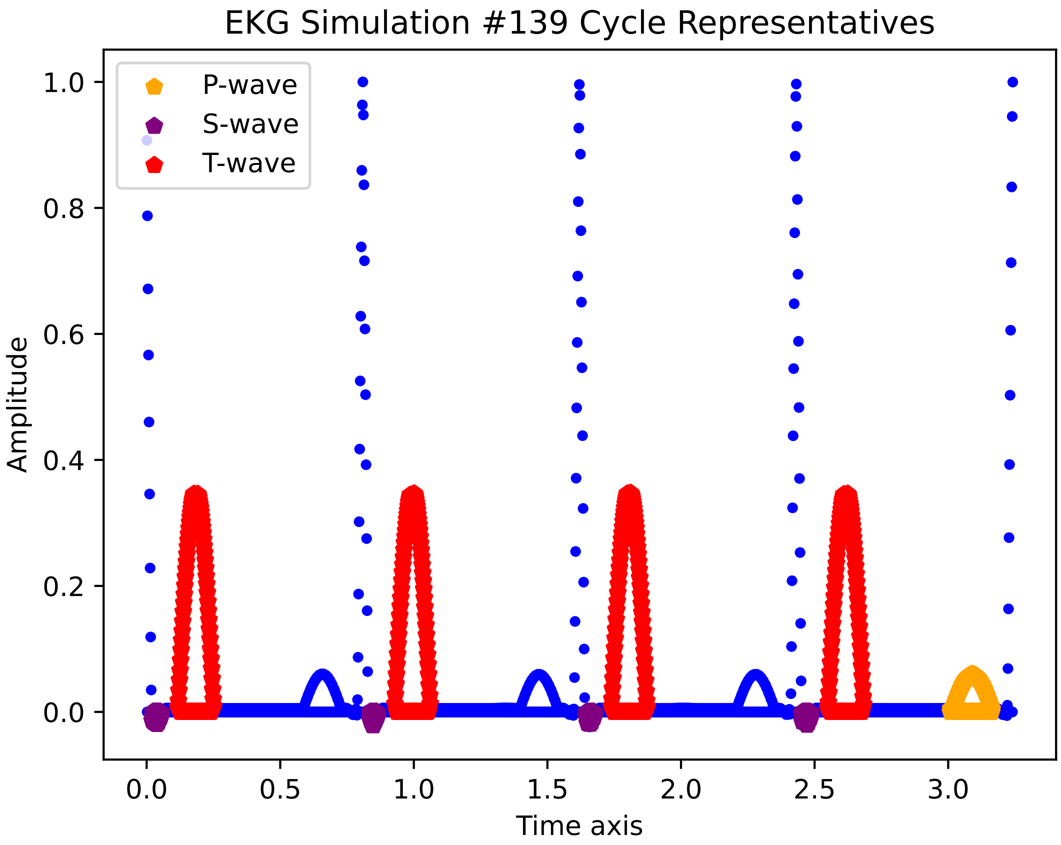
EKG Simulation #139 Cycle Representatives
P-wave
S-wave
T-wave
1.0
0.8
0.6
0.4
0.2
0.0
Amplitude
0.0
0.5
1.0
1.5
2.0
2.5
3.0
Time axis

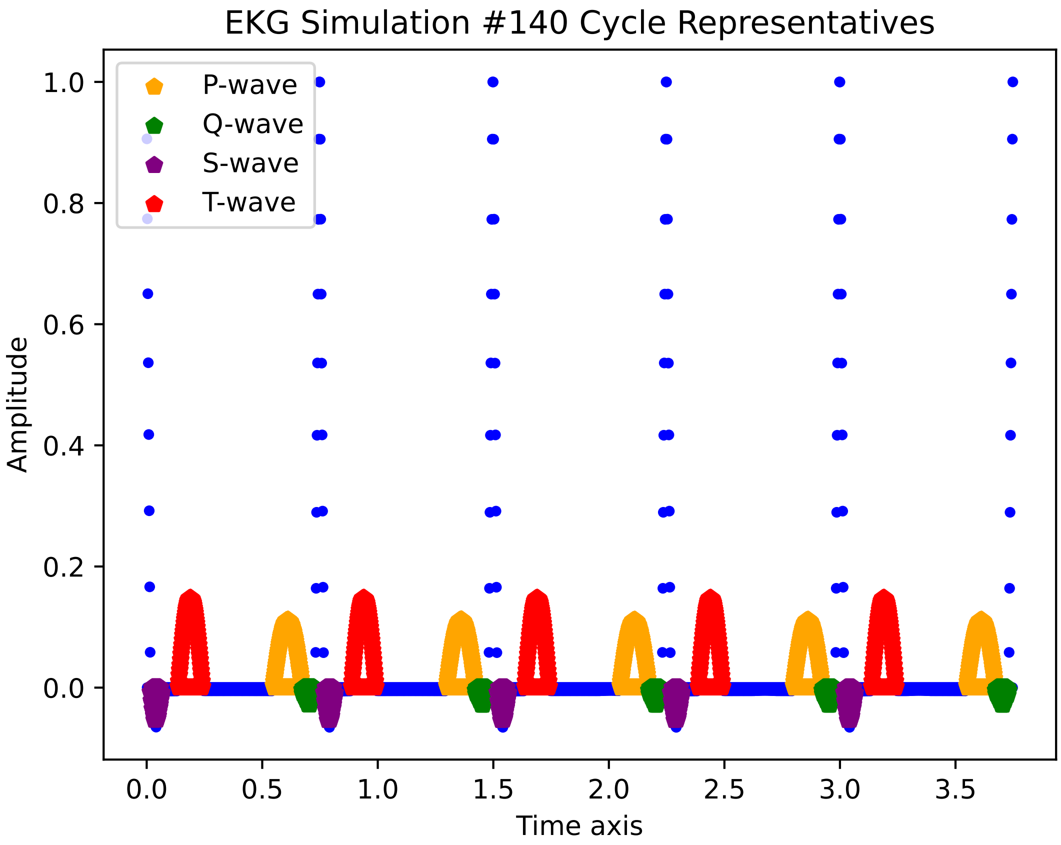
EKG Simulation #140 Cycle Representatives
P-wave
Q-wave
S-wave
T-wave
1.0
0.8
0.6
0.4
0.2
0.0
Amplitude
0.0
0.5
1.0
1.5
2.0
2.5
3.0
3.5
Time axis

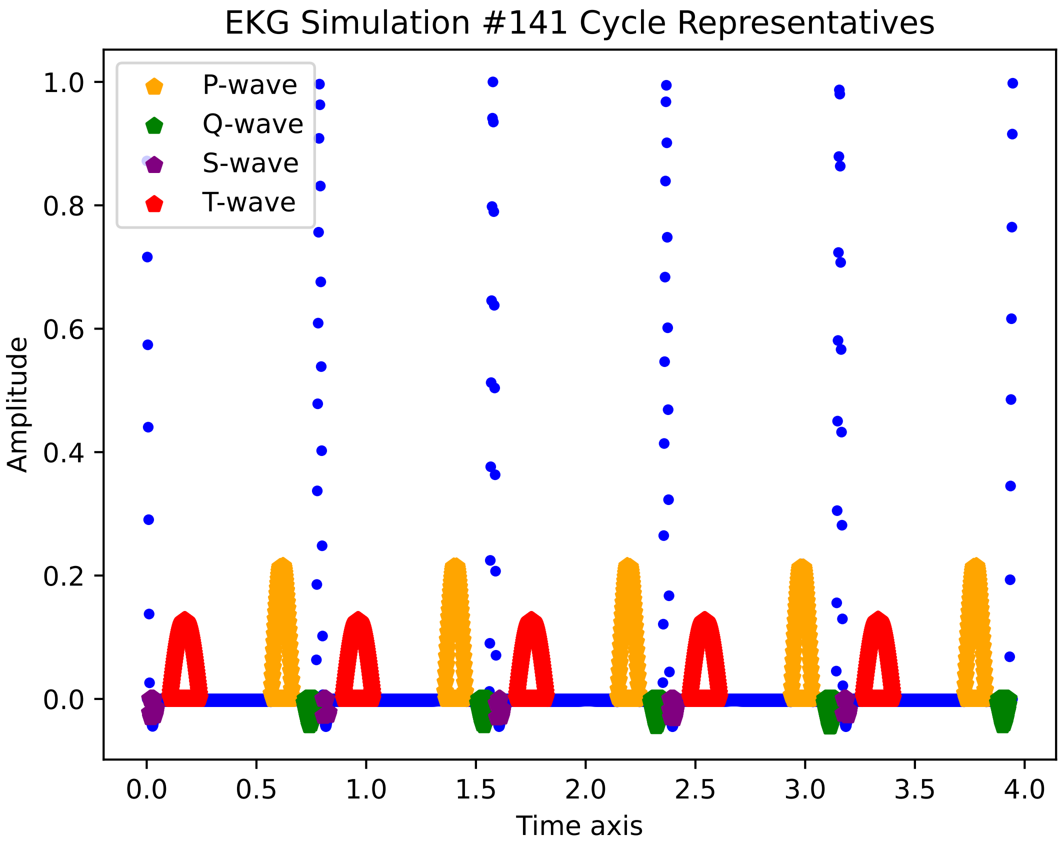

EKG Simulation #141 Cycle Representatives
P-wave
Q-wave
S-wave
T-wave
Amplitude
Time axis
0.0
0.2
0.4
0.6
0.8
1.0
0.0
0.5
1.0
1.5
2.0
2.5
3.0
3.5
4.0

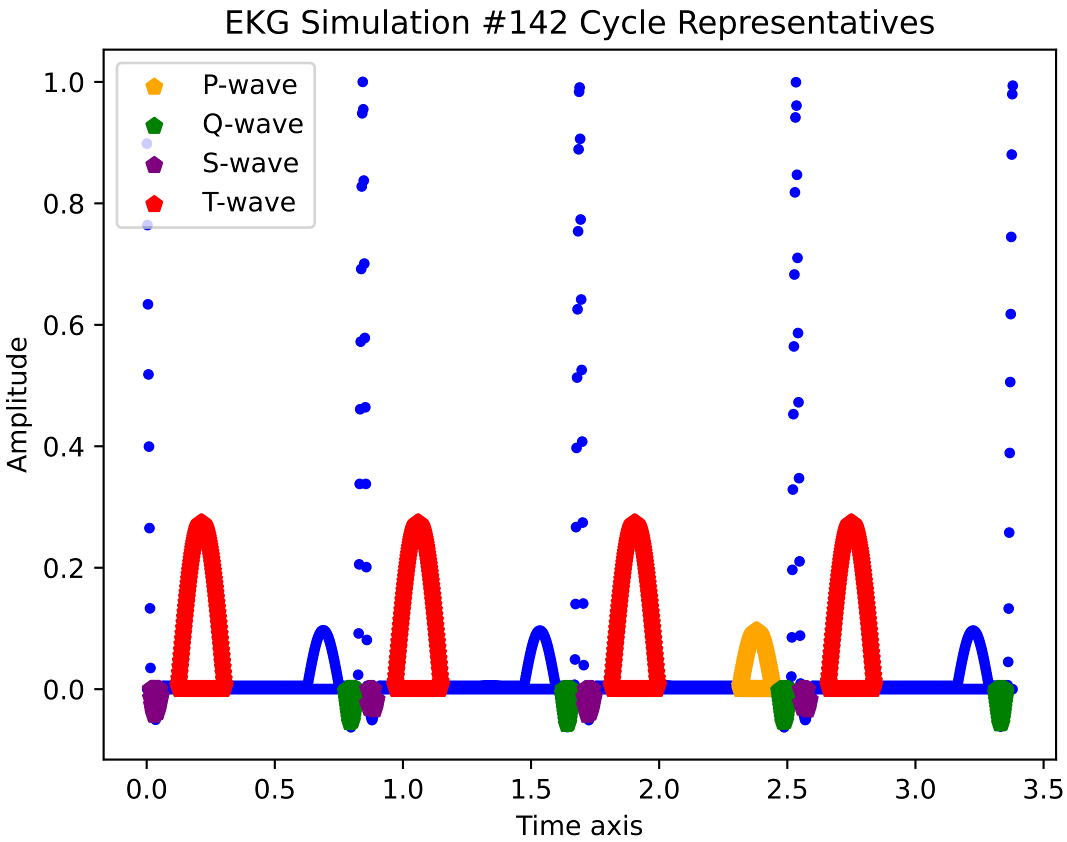

EKG Simulation #142 Cycle Representatives
P-wave
Q-wave
S-wave
T-wave
1.0
0.8
0.6
0.4
0.2
0.0
Amplitude
0.0
0.5
1.0
1.5
2.0
2.5
3.0
3.5
Time axis

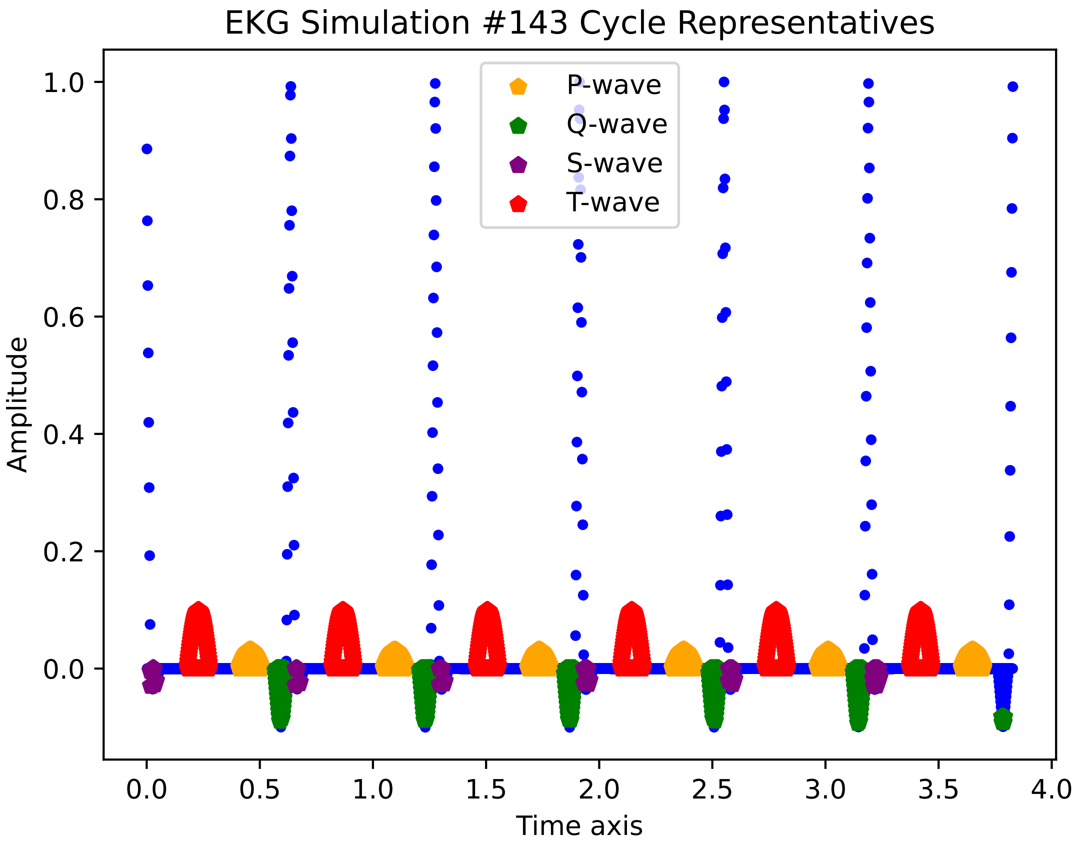
EKG Simulation #143 Cycle Representatives
P-wave
Q-wave
S-wave
T-wave
Amplitude
Time axis
1.0
0.8
0.6
0.4
0.2
0.0
0.0
0.5
1.0
1.5
2.0
2.5
3.0
3.5
4.0

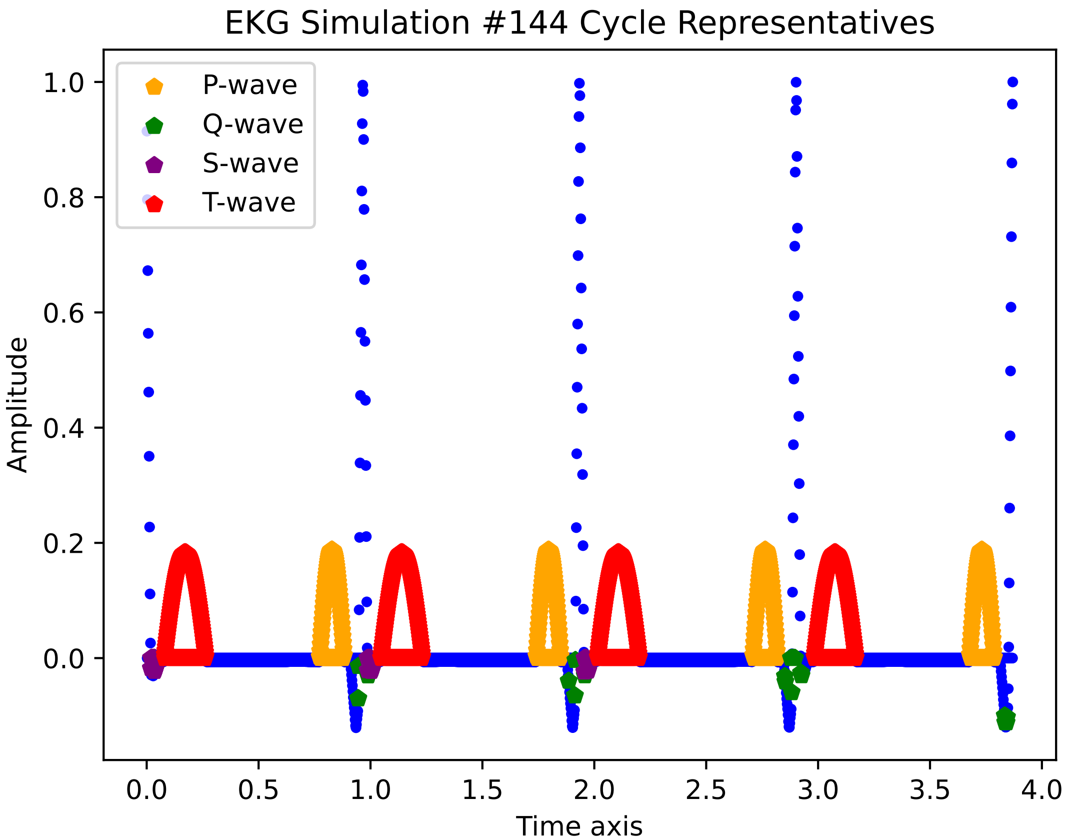
EKG Simulation #144 Cycle Representatives
P-wave
Q-wave
S-wave
T-wave
Amplitude
Time axis
1.0
0.8
0.6
0.4
0.2
0.0
0.0
0.5
1.0
1.5
2.0
2.5
3.0
3.5
4.0

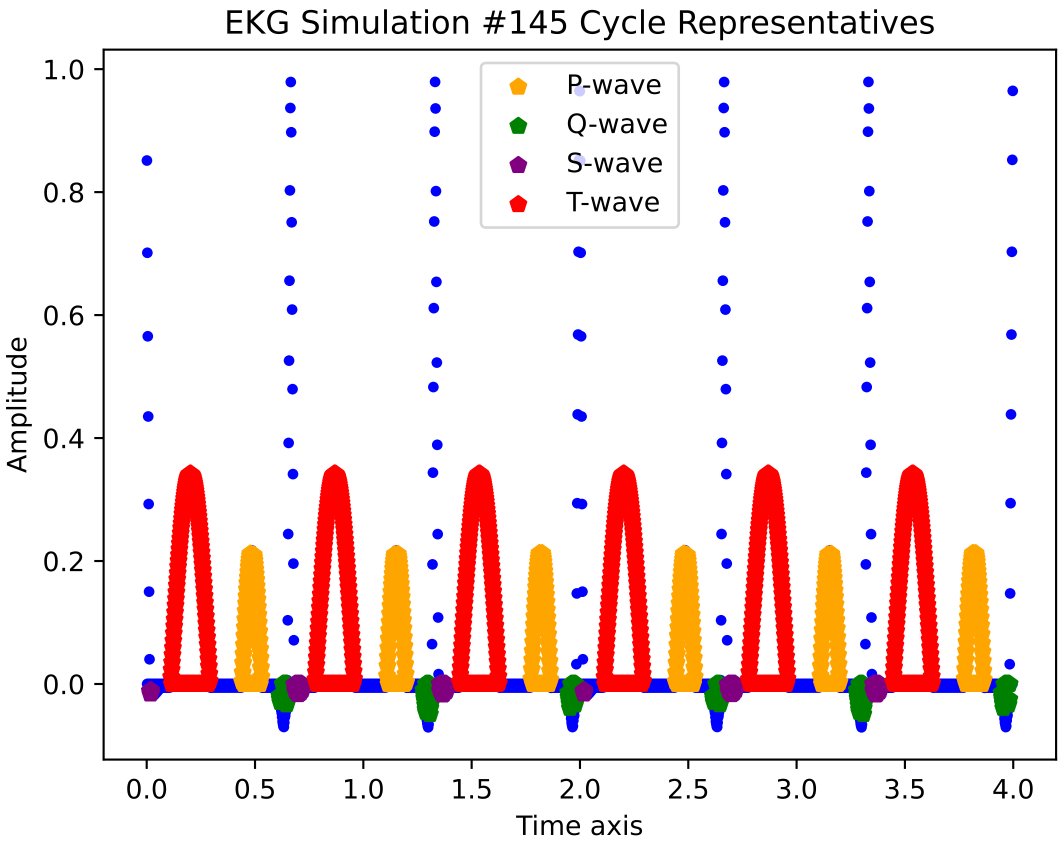
EKG Simulation #145 Cycle Representatives
P-wave
Q-wave
S-wave
T-wave
Amplitude
Time axis
1.0
0.8
0.6
0.4
0.2
0.0
0.0
0.5
1.0
1.5
2.0
2.5
3.0
3.5
4.0

EKG Simulation #146 Cycle Representatives

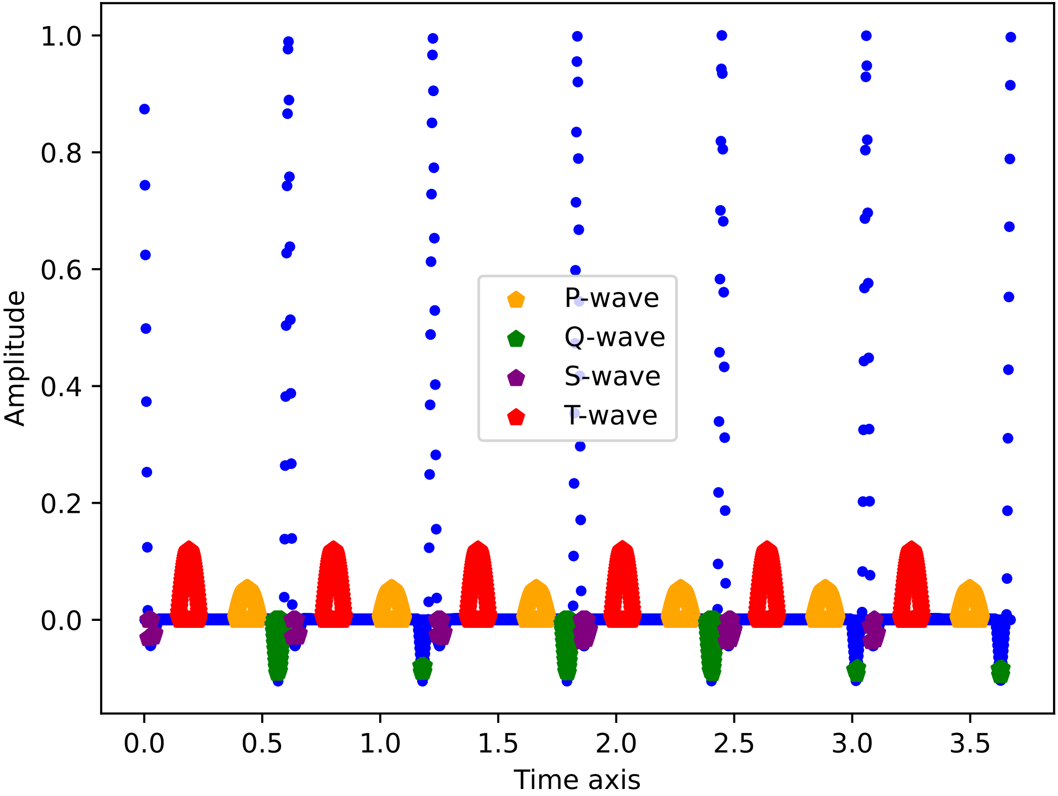

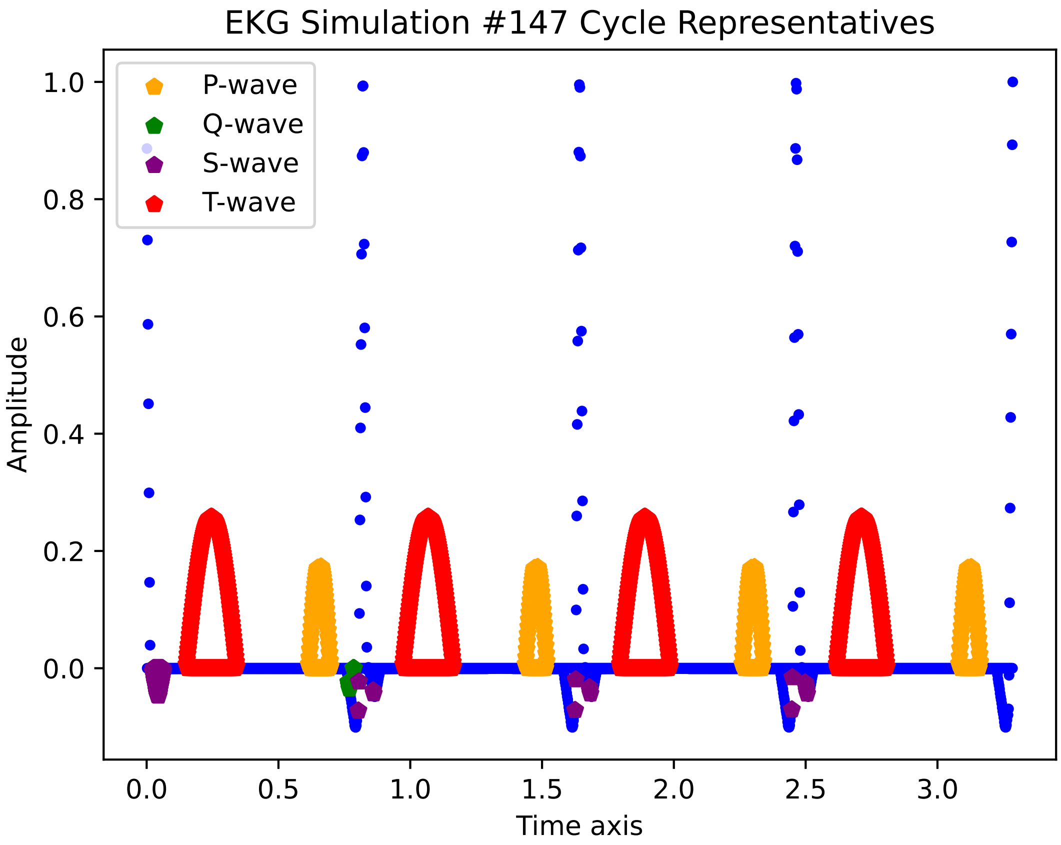
EKG Simulation #147 Cycle Representatives
P-wave
Q-wave
S-wave
T-wave
Amplitude
Time axis
1.0
0.8
0.6
0.4
0.2
0.0
0.0
0.5
1.0
1.5
2.0
2.5
3.0

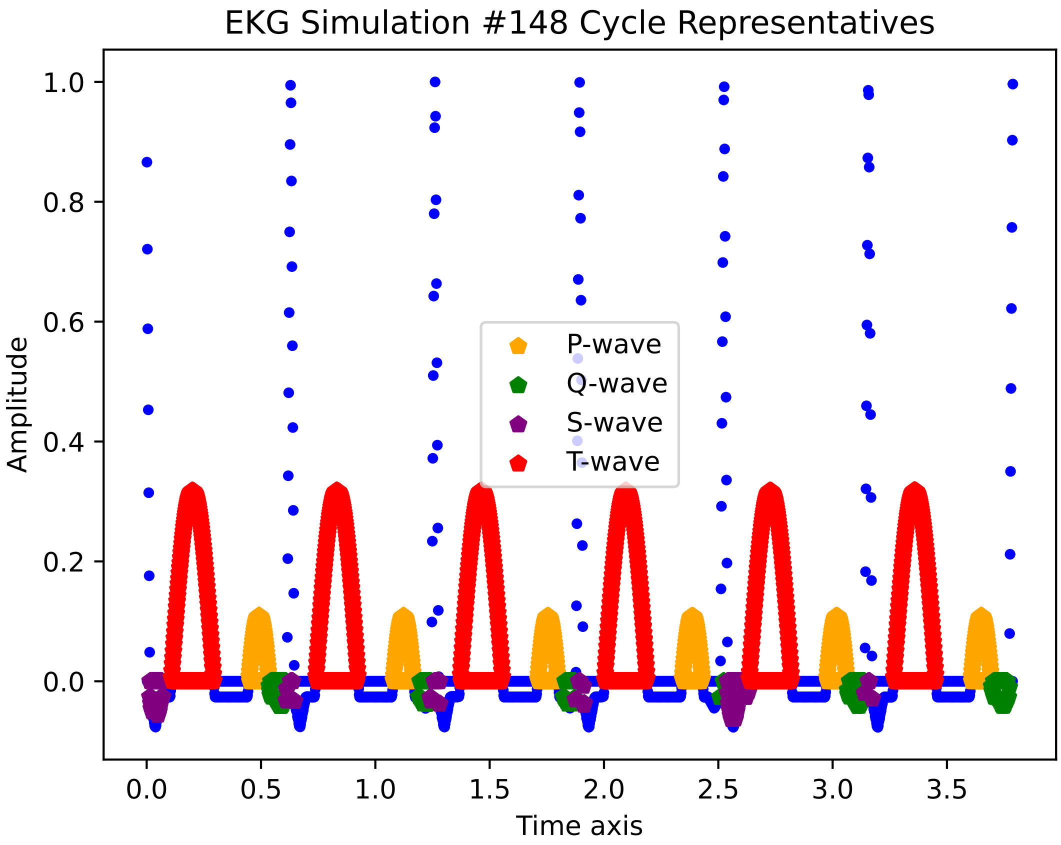
EKG Simulation #148 Cycle Representatives
Amplitude
Time axis
P-wave
Q-wave
S-wave
T-wave
1.0
0.8
0.6
0.4
0.2
0.0
0.0
0.5
1.0
1.5
2.0
2.5
3.0
3.5

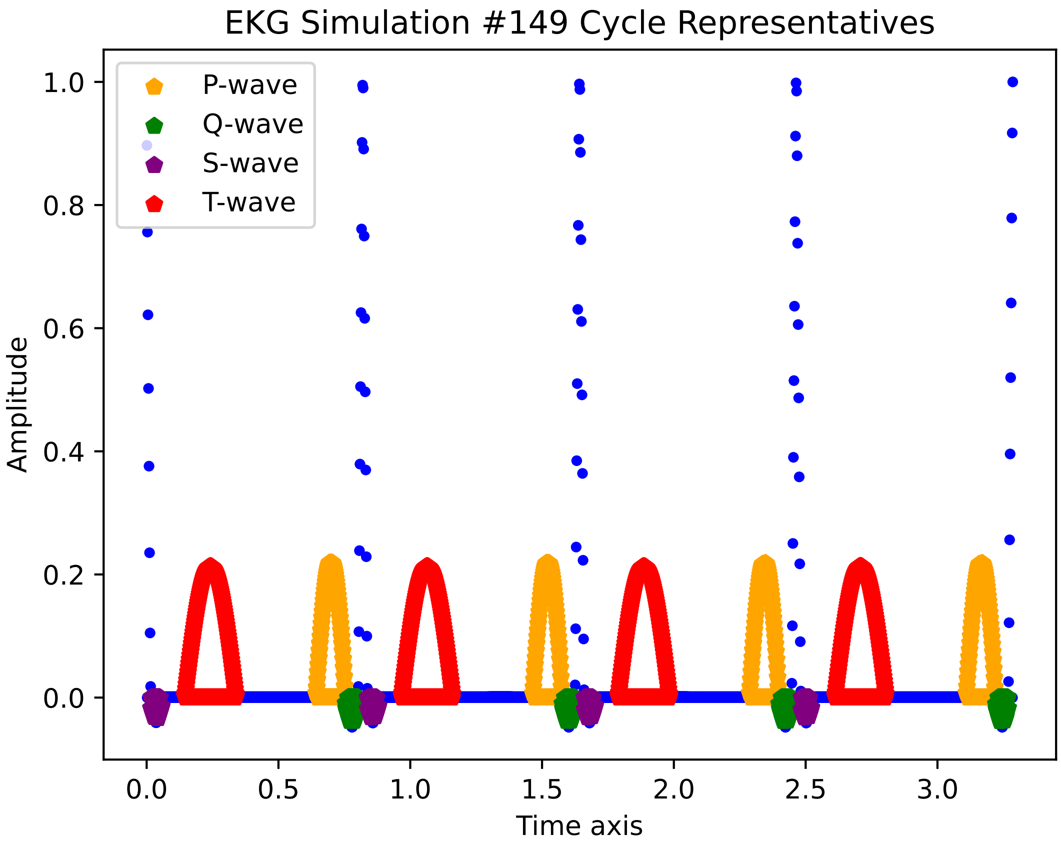

EKG Simulation #149 Cycle Representatives
P-wave
Q-wave
S-wave
T-wave
Amplitude
Time axis
1.0
0.8
0.6
0.4
0.2
0.0
0.0
0.5
1.0
1.5
2.0
2.5
3.0

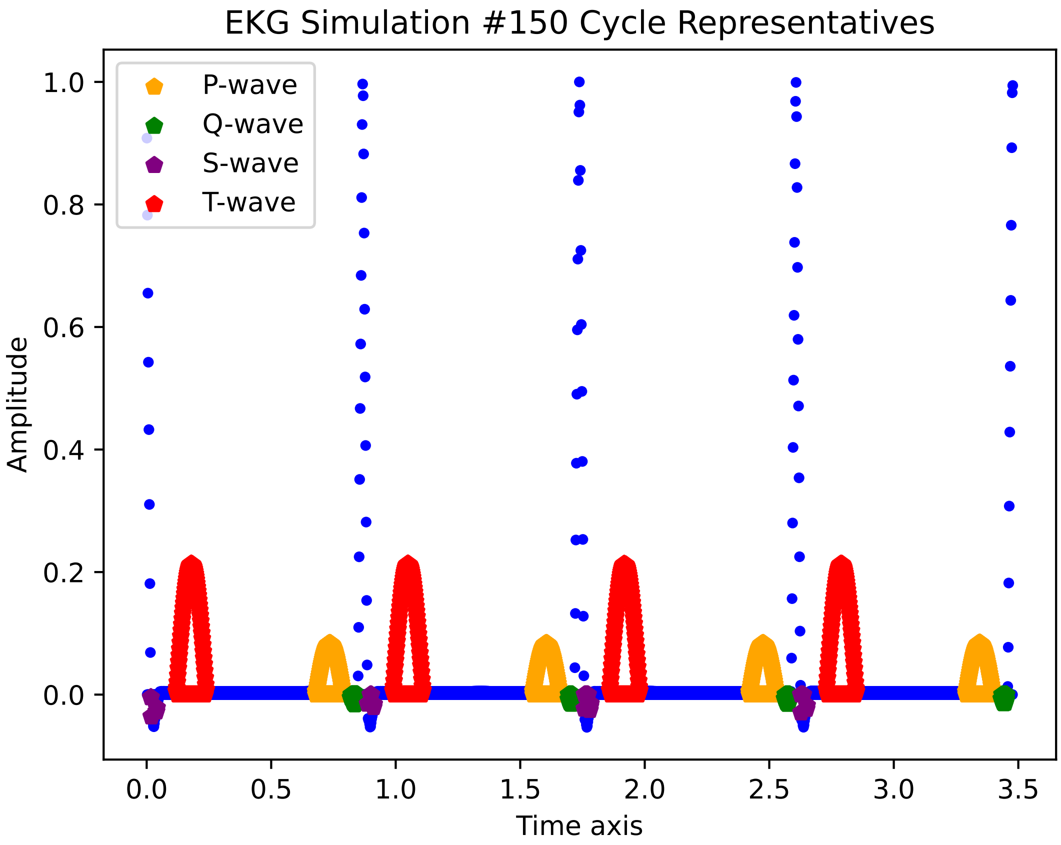

EKG Simulation #150 Cycle Representatives
P-wave
Q-wave
S-wave
T-wave
Amplitude
1.0
0.8
0.6
0.4
0.2
0.0
0.0
0.5
1.0
1.5
2.0
2.5
3.0
3.5
Time axis

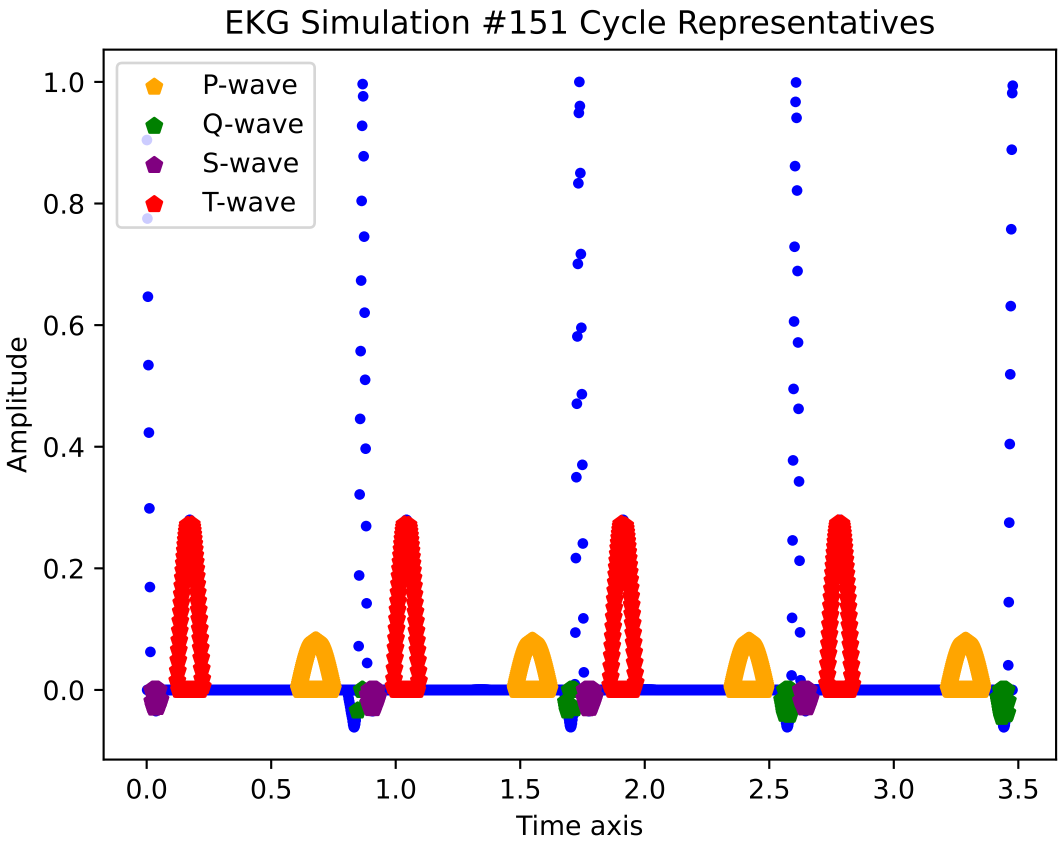

EKG Simulation #151 Cycle Representatives
P-wave
Q-wave
S-wave
T-wave
1.0
0.8
0.6
0.4
0.2
0.0
Amplitude
0.0
0.5
1.0
1.5
2.0
2.5
3.0
3.5
Time axis

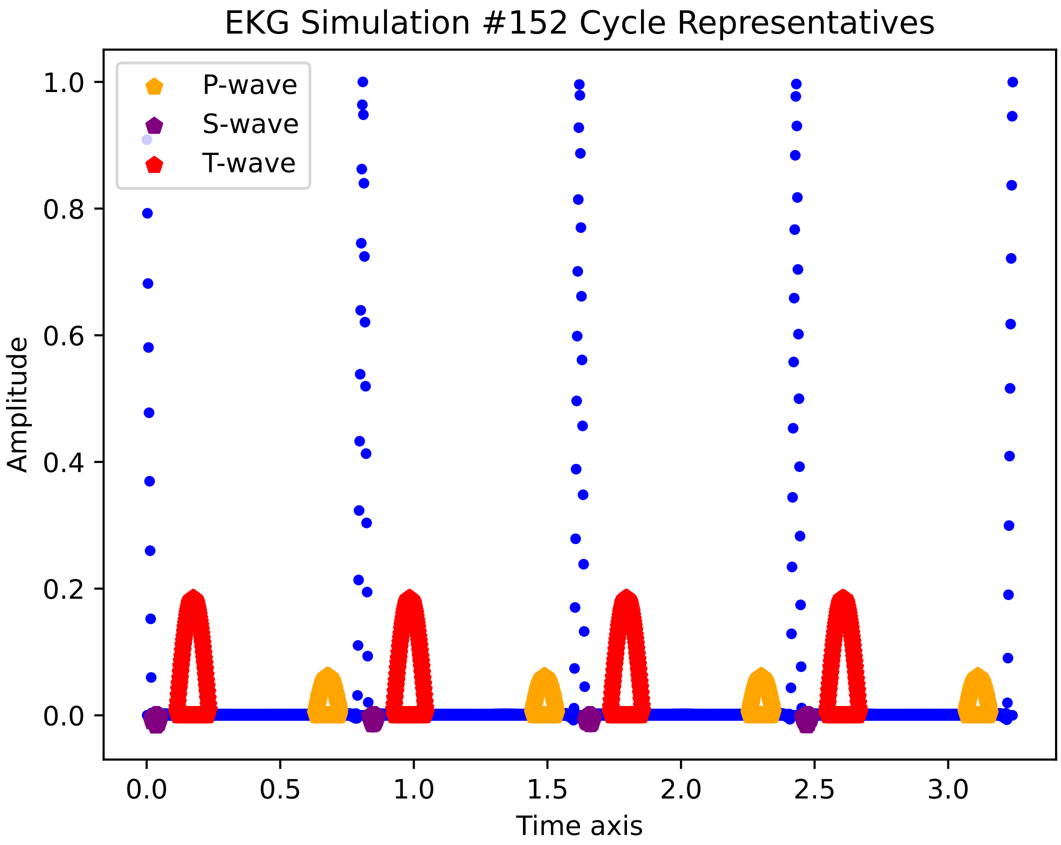
EKG Simulation #152 Cycle Representatives
P-wave
S-wave
T-wave
Amplitude
Time axis
0.0
0.2
0.4
0.6
0.8
1.0
0.0
0.5
1.0
1.5
2.0
2.5
3.0

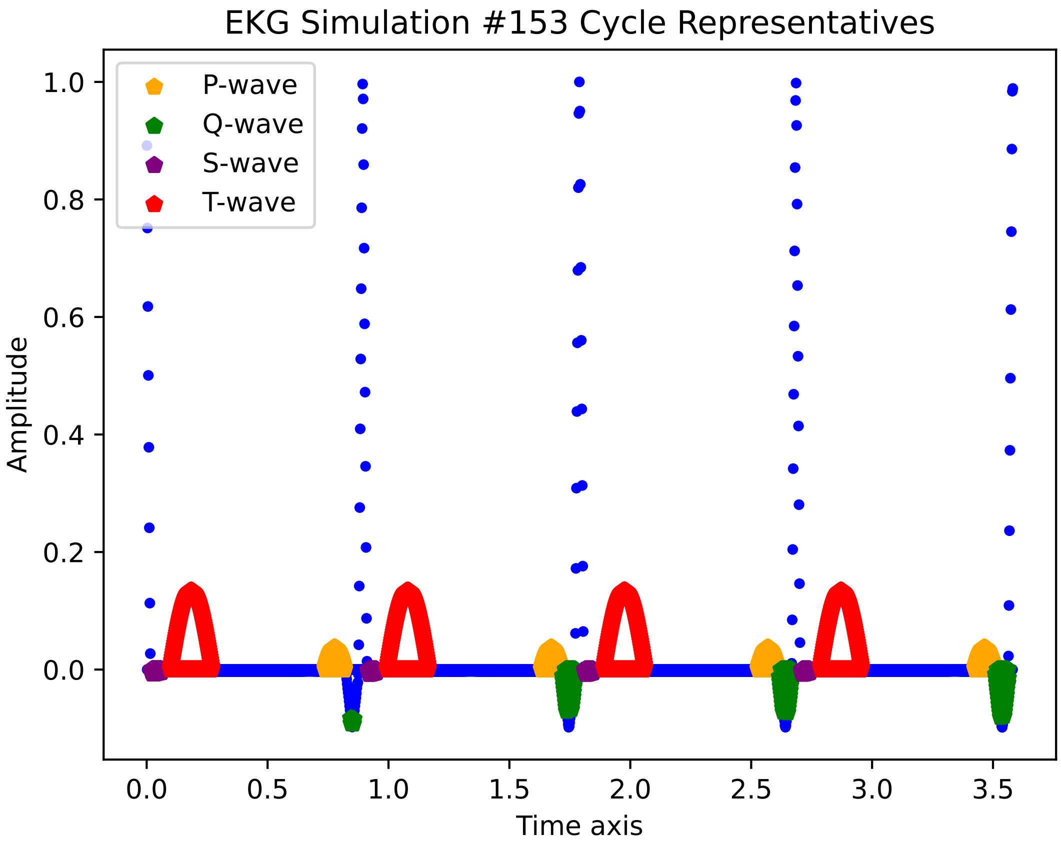

EKG Simulation #153 Cycle Representatives
P-wave
Q-wave
S-wave
T-wave
Amplitude
Time axis
1.0
0.8
0.6
0.4
0.2
0.0
0.0
0.5
1.0
1.5
2.0
2.5
3.0
3.5

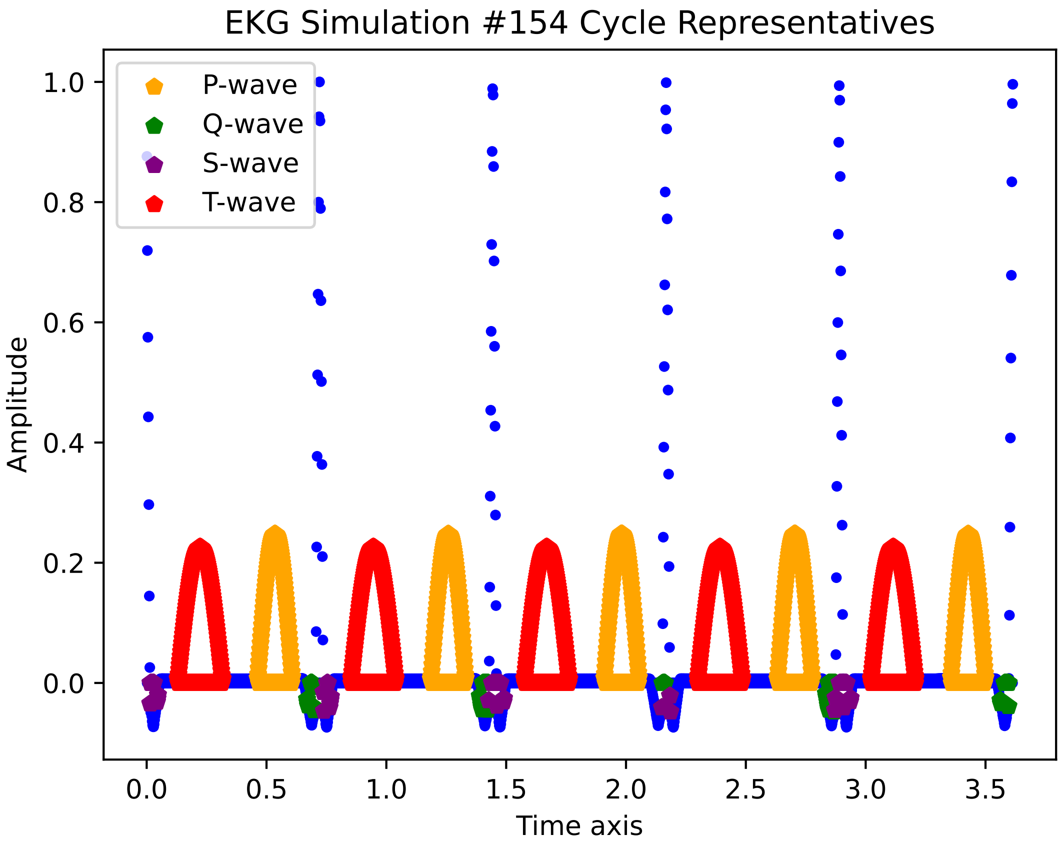

EKG Simulation #154 Cycle Representatives
P-wave
Q-wave
S-wave
T-wave
Amplitude
Time axis
1.0
0.8
0.6
0.4
0.2
0.0
0.0
0.5
1.0
1.5
2.0
2.5
3.0
3.5

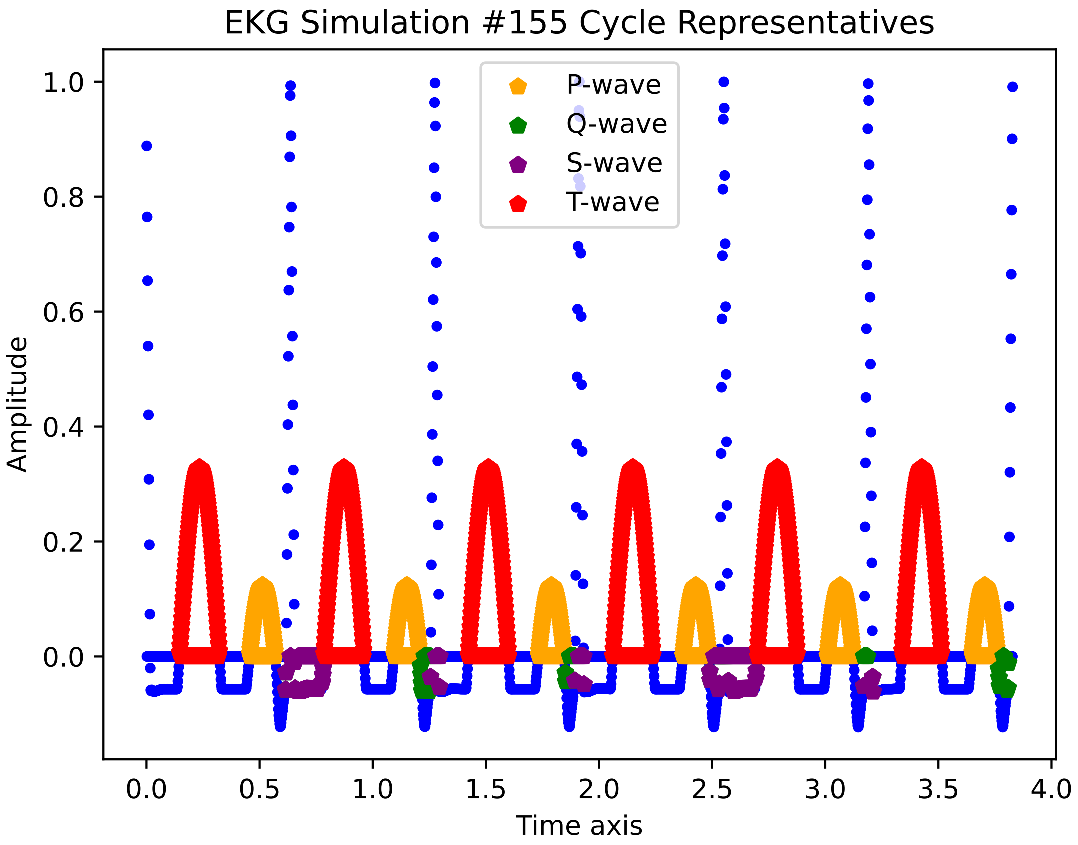

EKG Simulation #155 Cycle Representatives
P-wave
Q-wave
S-wave
T-wave
Amplitude
Time axis
1.0
0.8
0.6
0.4
0.2
0.0
0.0
0.5
1.0
1.5
2.0
2.5
3.0
3.5
4.0

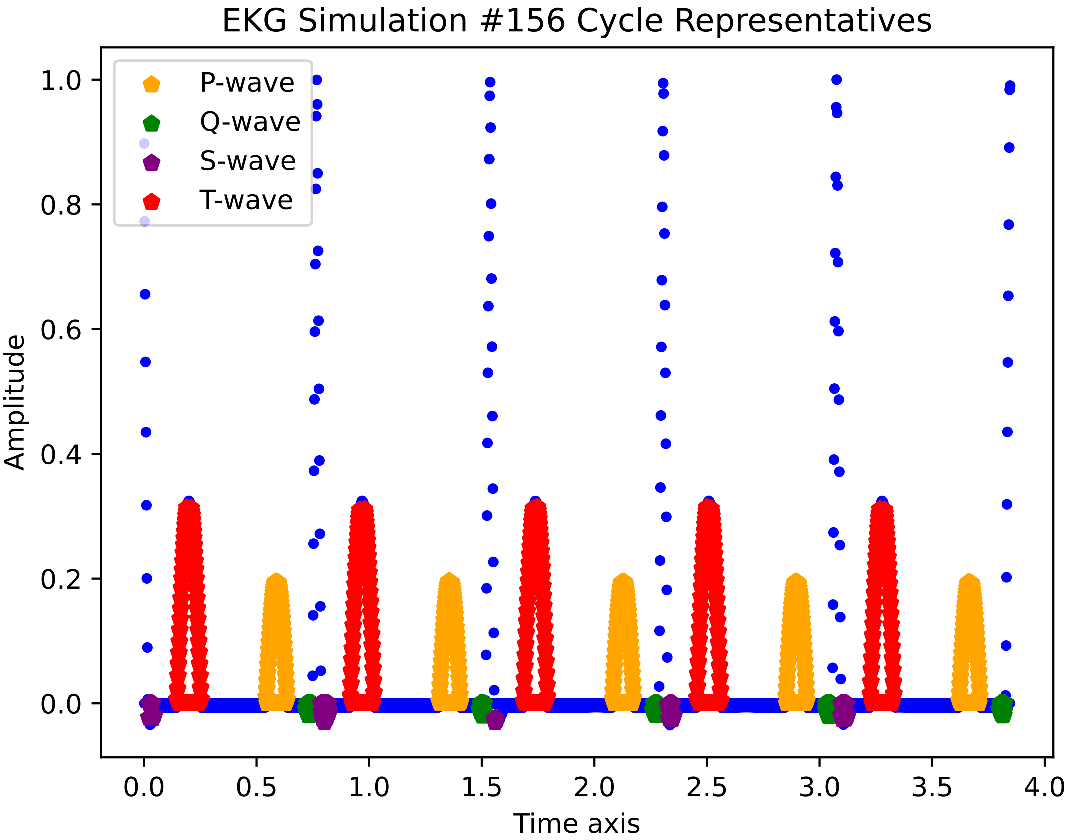
EKG Simulation #156 Cycle Representatives
P-wave
Q-wave
S-wave
T-wave
Amplitude
Time axis
1.0
0.8
0.6
0.4
0.2
0.0
0.0
0.5
1.0
1.5
2.0
2.5
3.0
3.5
4.0

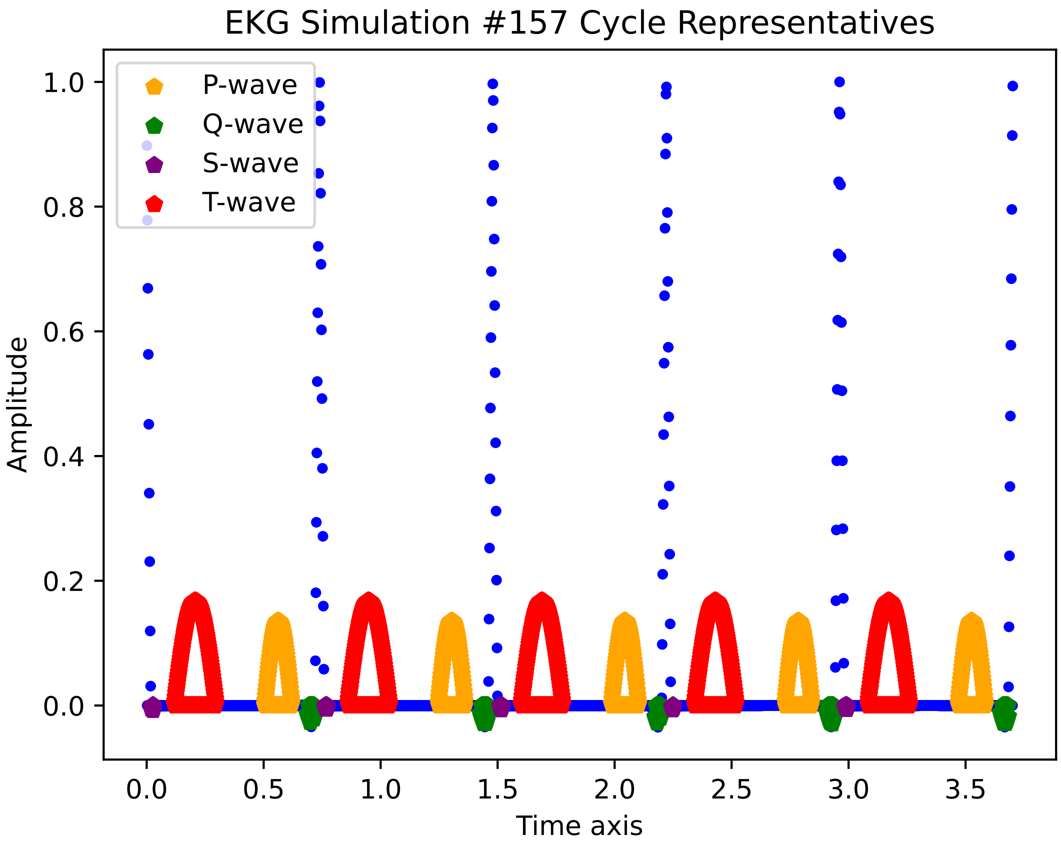
EKG Simulation #157 Cycle Representatives
P-wave
Q-wave
S-wave
T-wave
Amplitude
Time axis
0.0
0.2
0.4
0.6
0.8
1.0
0.0
0.5
1.0
1.5
2.0
2.5
3.0
3.5

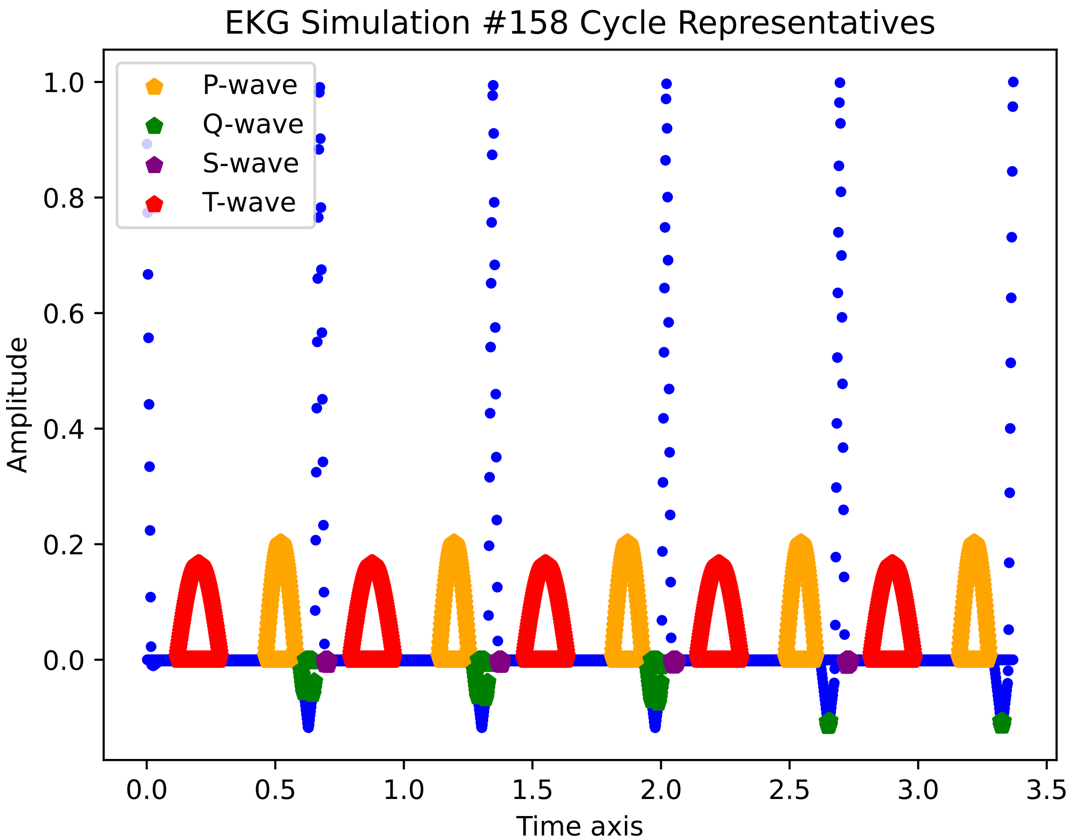

EKG Simulation #158 Cycle Representatives
P-wave
Q-wave
S-wave
T-wave
Amplitude
Time axis
1.0
0.8
0.6
0.4
0.2
0.0
0.0
0.5
1.0
1.5
2.0
2.5
3.0
3.5

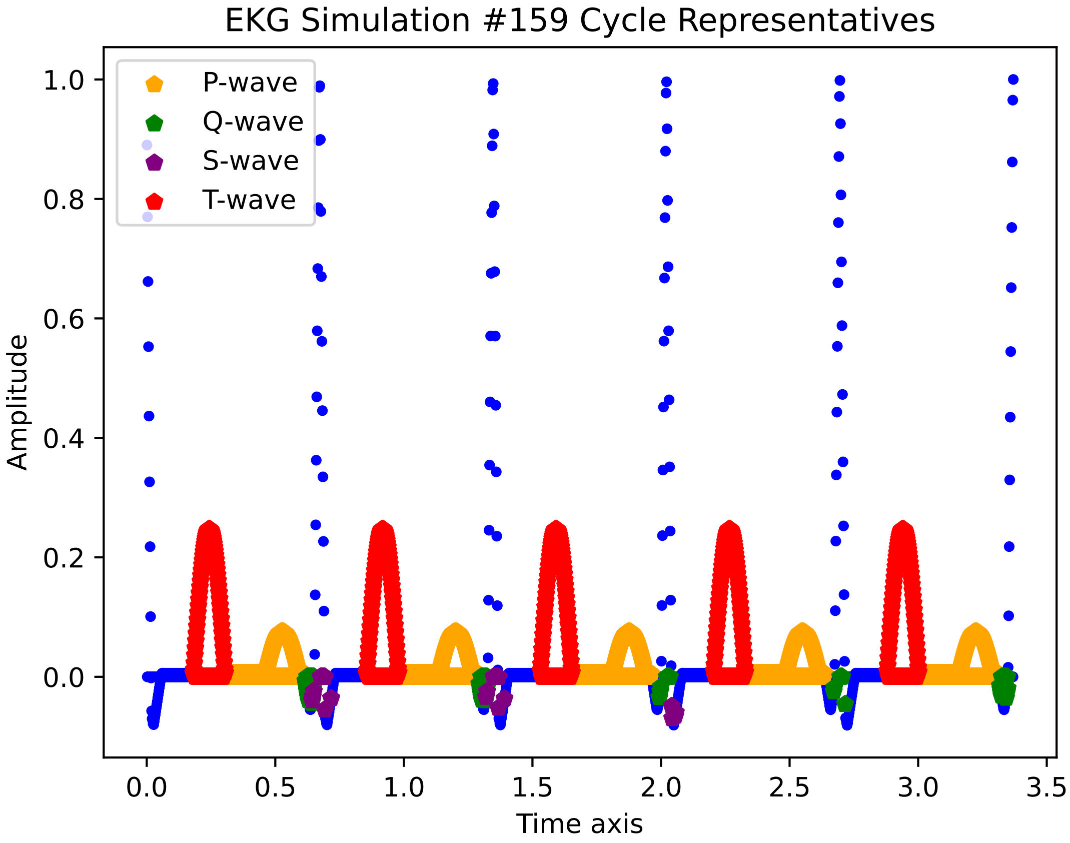

EKG Simulation #159 Cycle Representatives
P-wave
Q-wave
S-wave
T-wave
Amplitude
Time axis
1.0
0.8
0.6
0.4
0.2
0.0
0.0
0.5
1.0
1.5
2.0
2.5
3.0
3.5

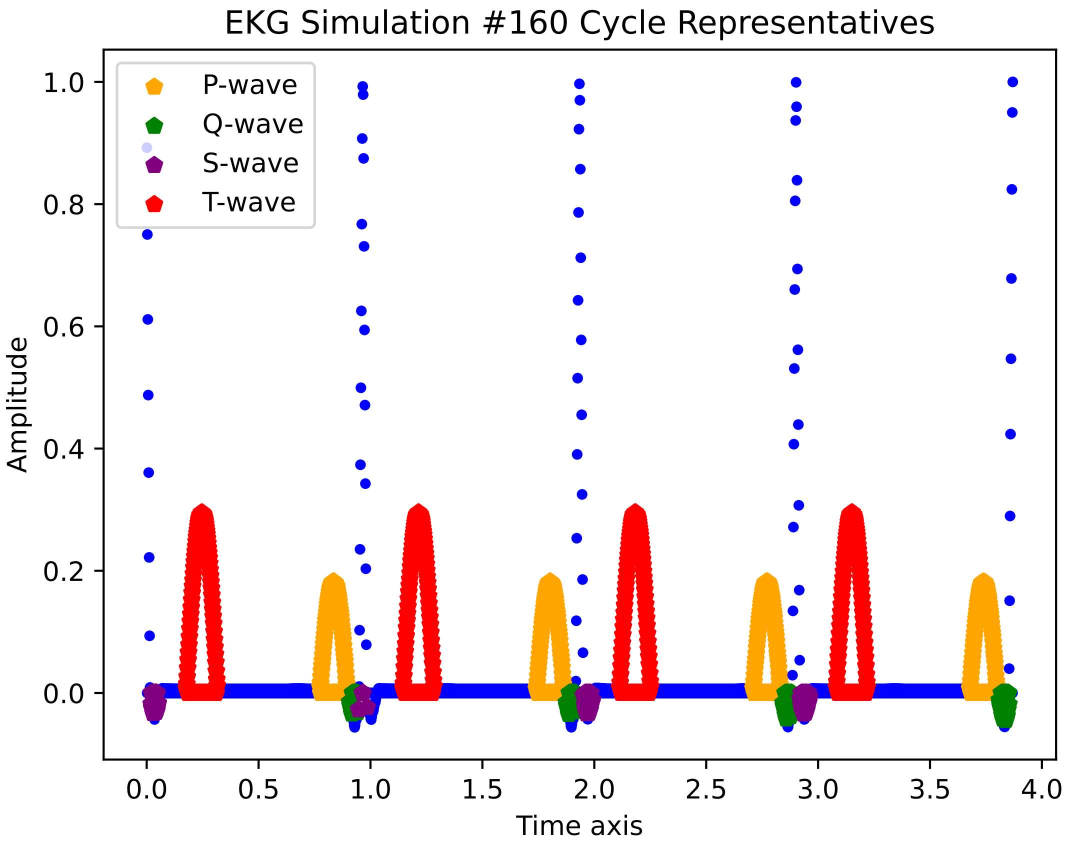
EKG Simulation #160 Cycle Representatives
P-wave
Q-wave
S-wave
T-wave
1.0
0.8
0.6
0.4
0.2
0.0
Amplitude
0.0
0.5
1.0
1.5
2.0
2.5
3.0
3.5
4.0
Time axis

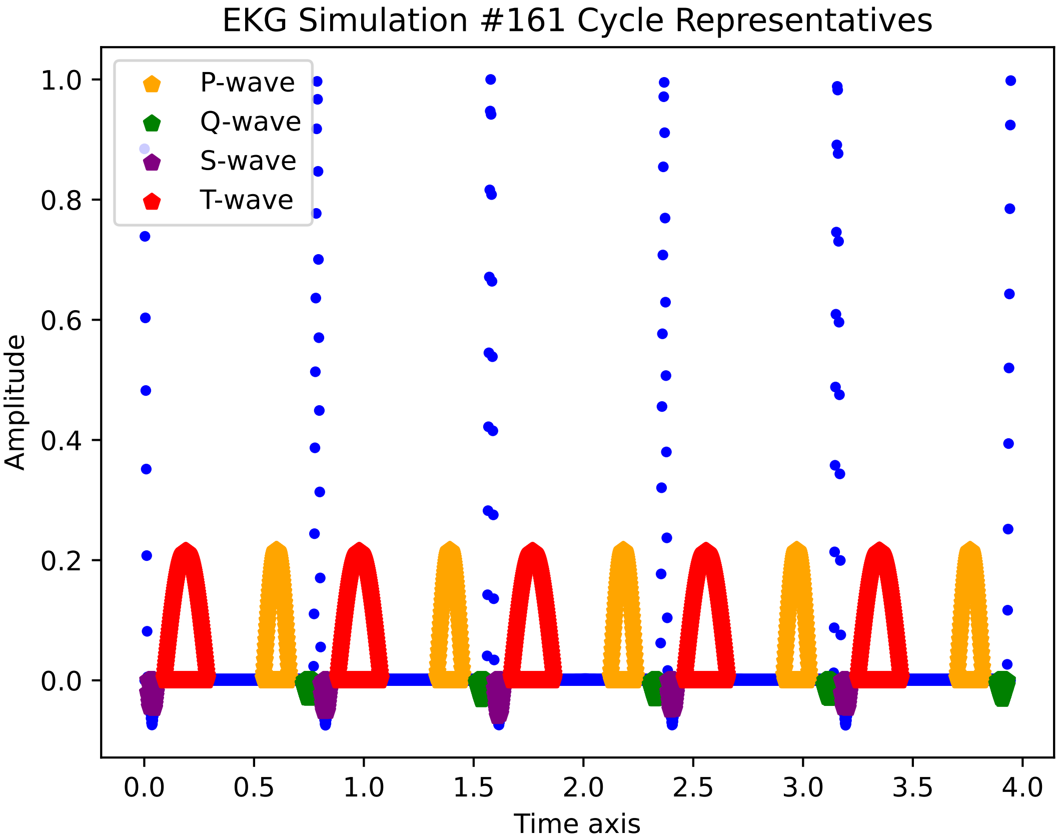
EKG Simulation #161 Cycle Representatives
P-wave
Q-wave
S-wave
T-wave
1.0
0.8
0.6
0.4
0.2
0.0
Amplitude
0.0
0.5
1.0
1.5
2.0
2.5
3.0
3.5
4.0
Time axis

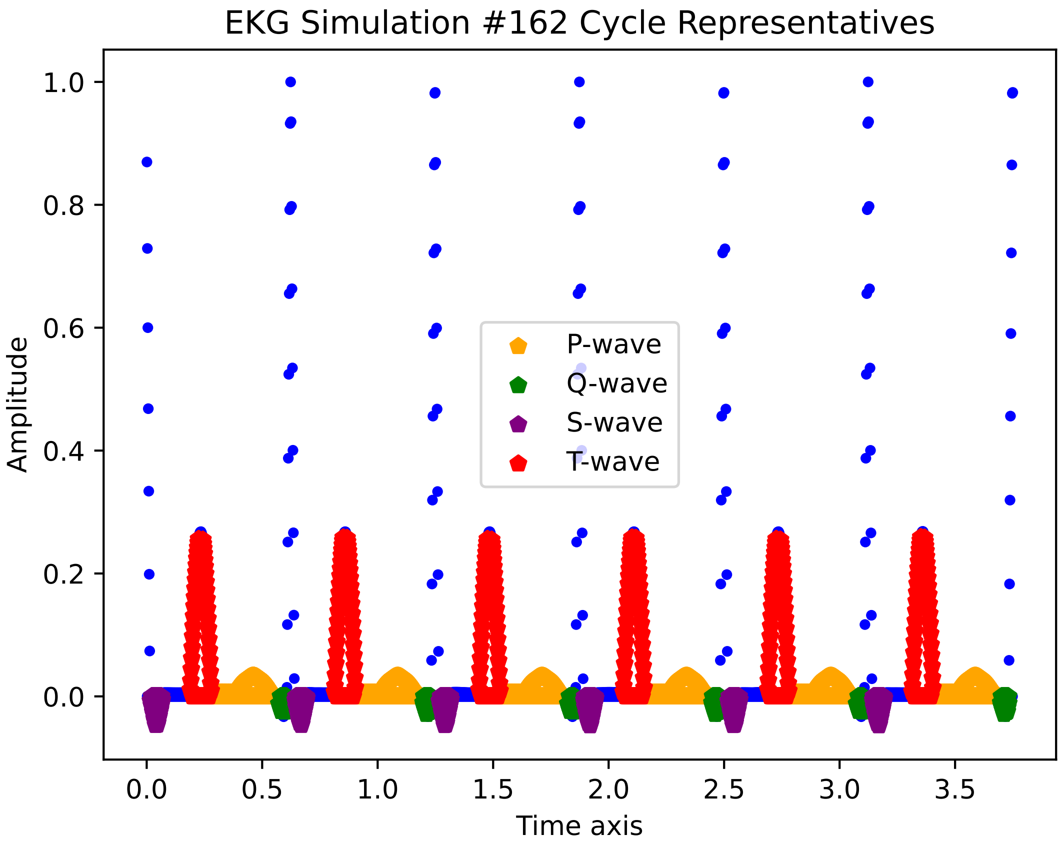

EKG Simulation #162 Cycle Representatives
Amplitude
Time axis
P-wave
Q-wave
S-wave
T-wave
1.0
0.8
0.6
0.4
0.2
0.0
0.0
0.5
1.0
1.5
2.0
2.5
3.0
3.5

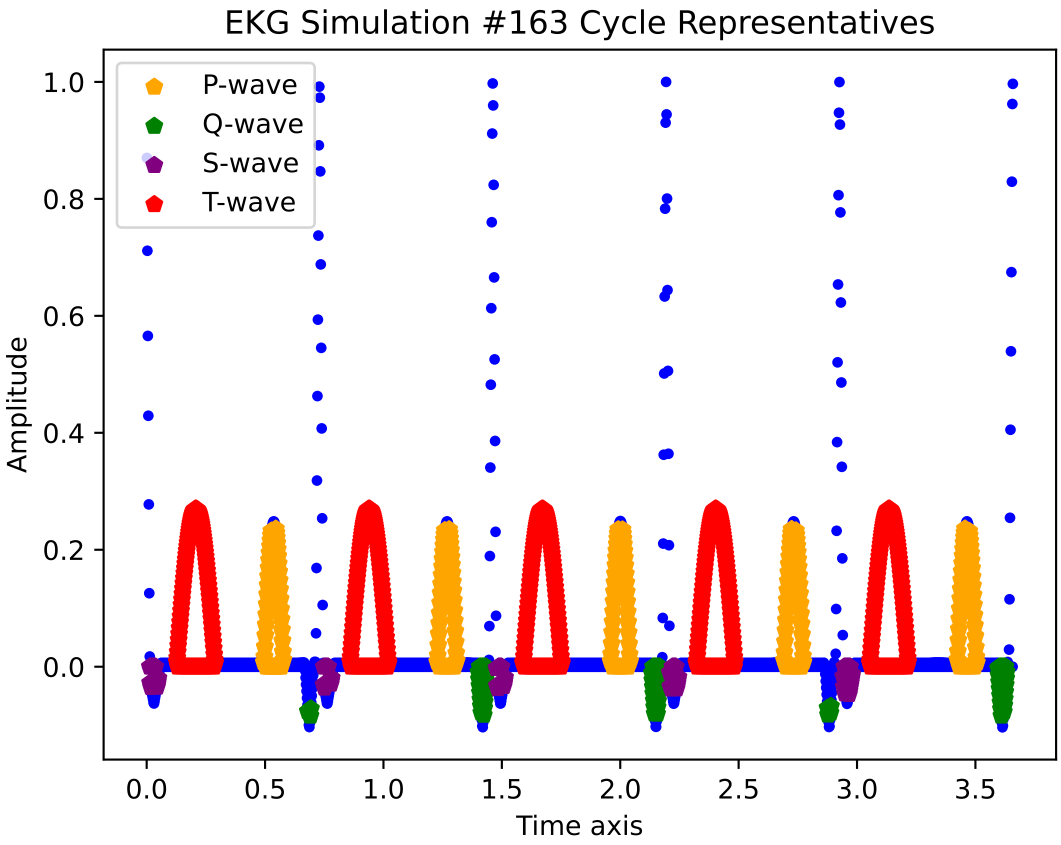

EKG Simulation #163 Cycle Representatives
P-wave
Q-wave
S-wave
T-wave
Amplitude
1.0
0.8
0.6
0.4
0.2
0.0
0.0
0.5
1.0
1.5
2.0
2.5
3.0
3.5
Time axis

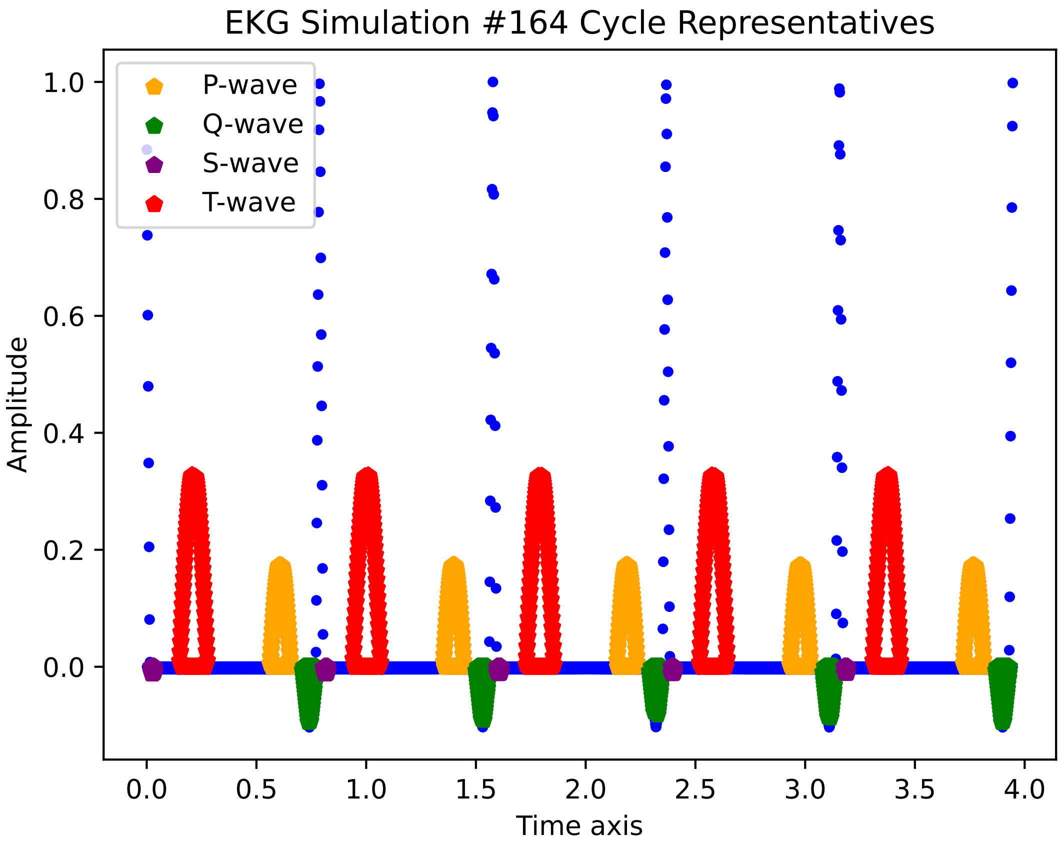
EKG Simulation #164 Cycle Representatives
P-wave
Q-wave
S-wave
T-wave
Amplitude
Time axis
1.0
0.8
0.6
0.4
0.2
0.0
0.0
0.5
1.0
1.5
2.0
2.5
3.0
3.5
4.0

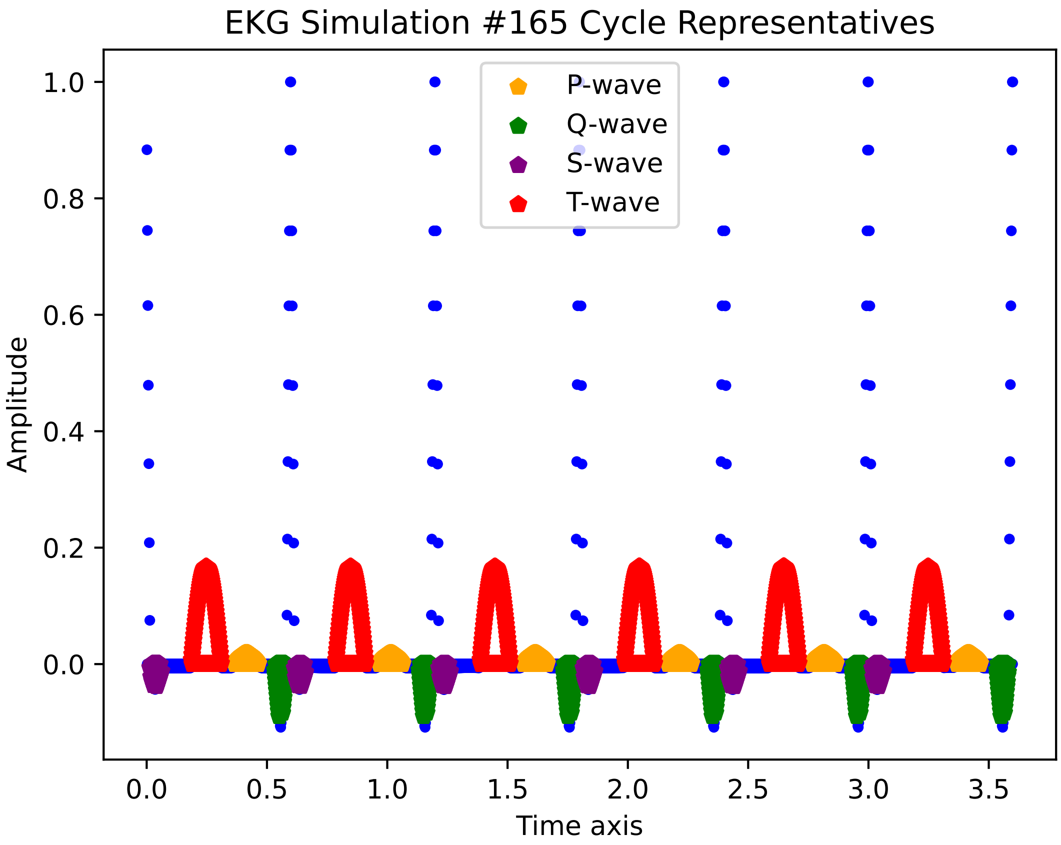
EKG Simulation #165 Cycle Representatives
P-wave
Q-wave
S-wave
T-wave
Amplitude
Time axis
1.0
0.8
0.6
0.4
0.2
0.0
0.0
0.5
1.0
1.5
2.0
2.5
3.0
3.5

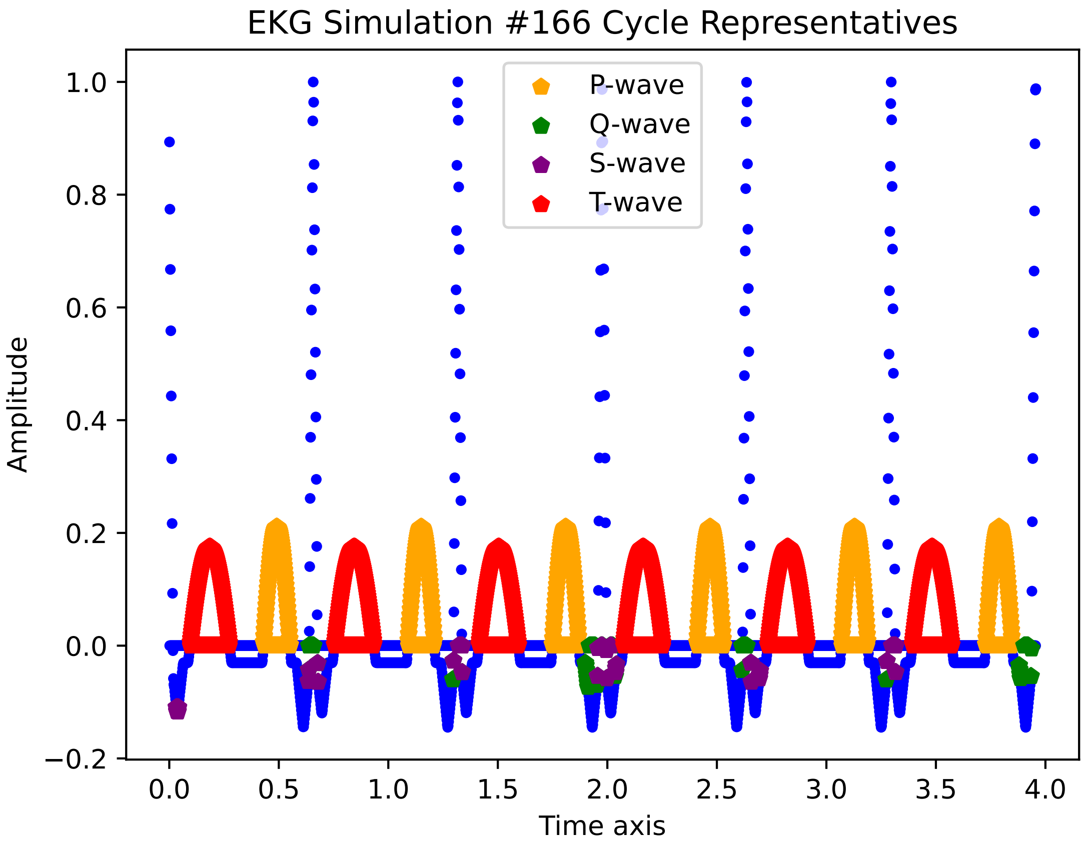

EKG Simulation #166 Cycle Representatives
P-wave
Q-wave
S-wave
T-wave
Amplitude
Time axis
1.0
0.8
0.6
0.4
0.2
0.0
−0.2
0.0
0.5
1.0
1.5
2.0
2.5
3.0
3.5
4.0

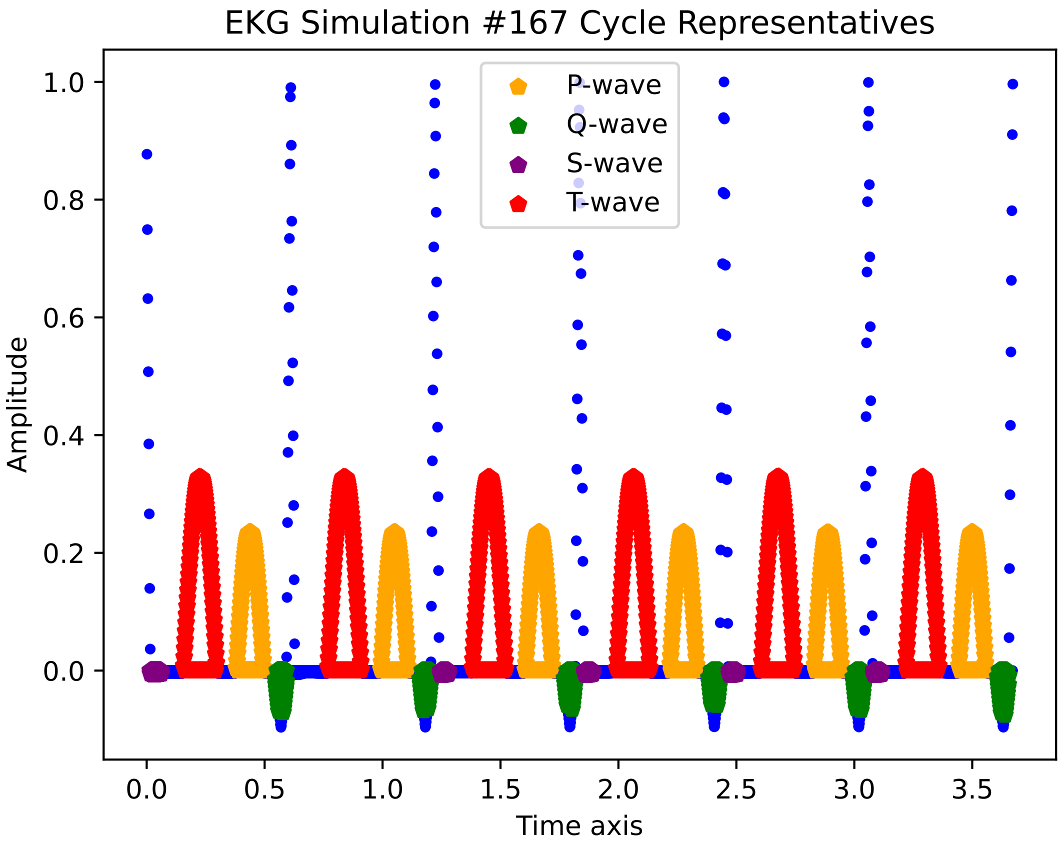
EKG Simulation #167 Cycle Representatives
P-wave
Q-wave
S-wave
T-wave
Amplitude
Time axis
1.0
0.8
0.6
0.4
0.2
0.0
0.0
0.5
1.0
1.5
2.0
2.5
3.0
3.5

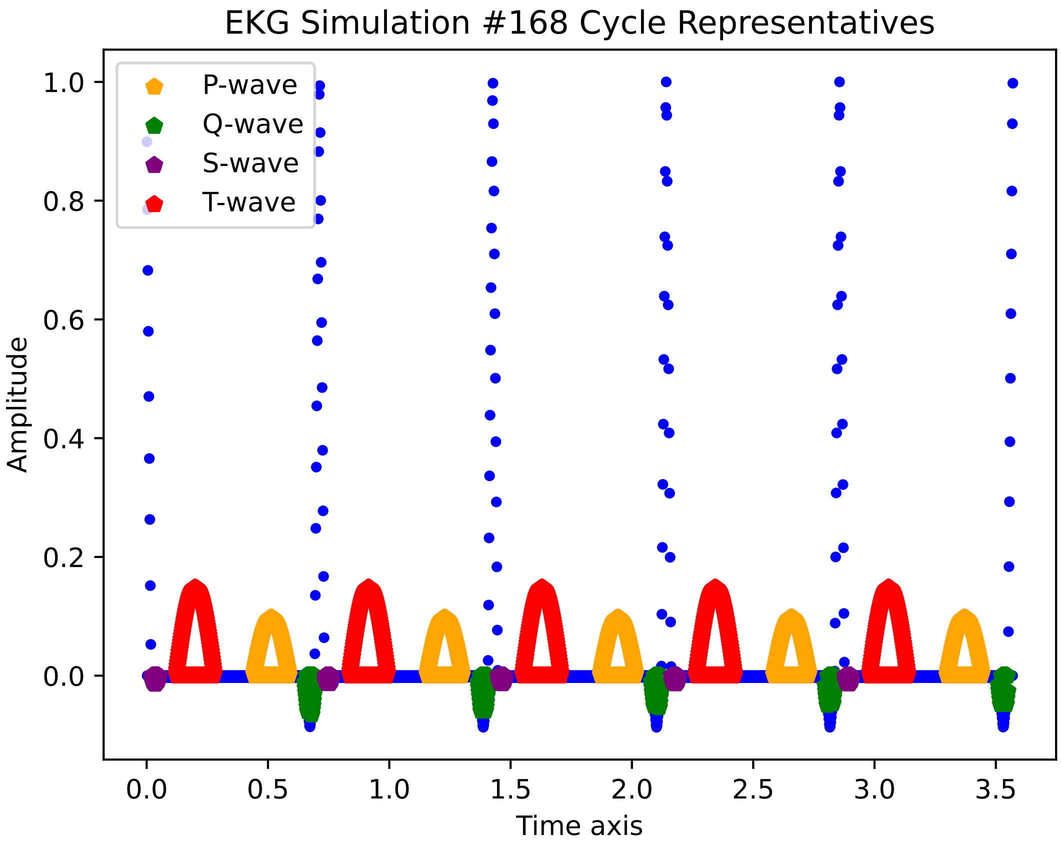
EKG Simulation #168 Cycle Representatives
P-wave
Q-wave
S-wave
T-wave
Amplitude
1.0
0.8
0.6
0.4
0.2
0.0
0.0
0.5
1.0
1.5
2.0
2.5
3.0
3.5
Time axis

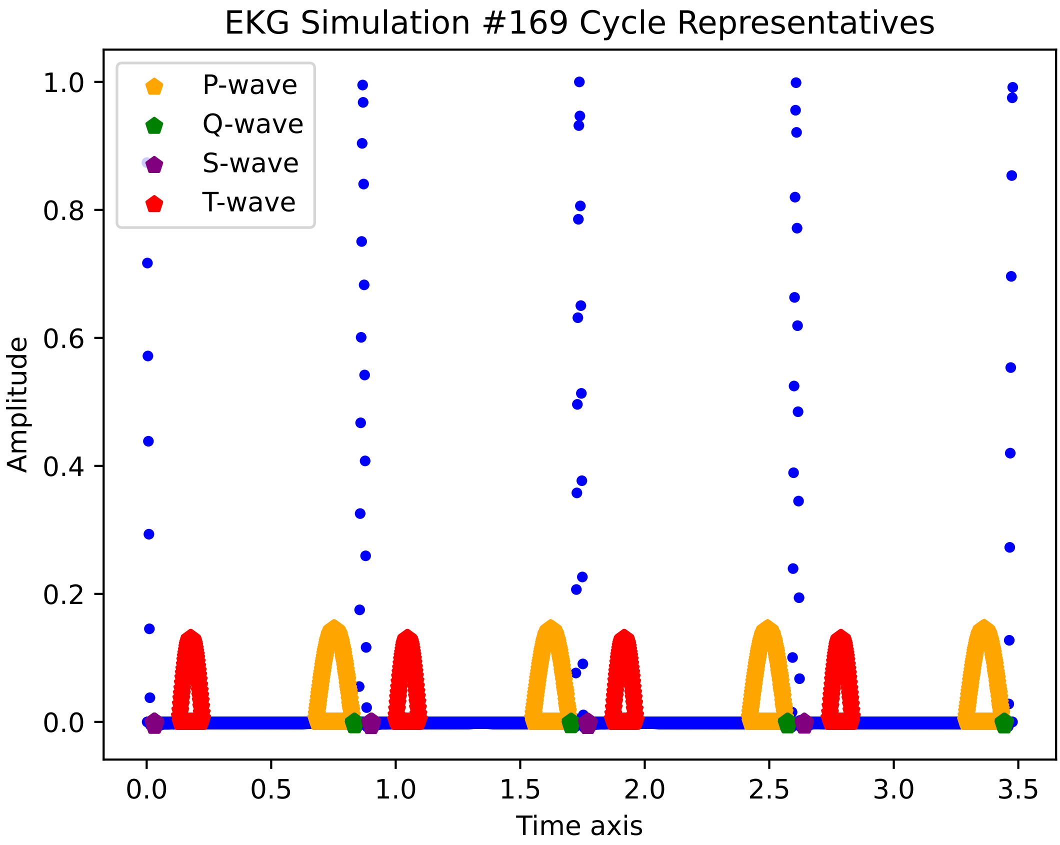

EKG Simulation #169 Cycle Representatives
P-wave
Q-wave
S-wave
T-wave
Amplitude
Time axis
1.0
0.8
0.6
0.4
0.2
0.0
0.0
0.5
1.0
1.5
2.0
2.5
3.0
3.5

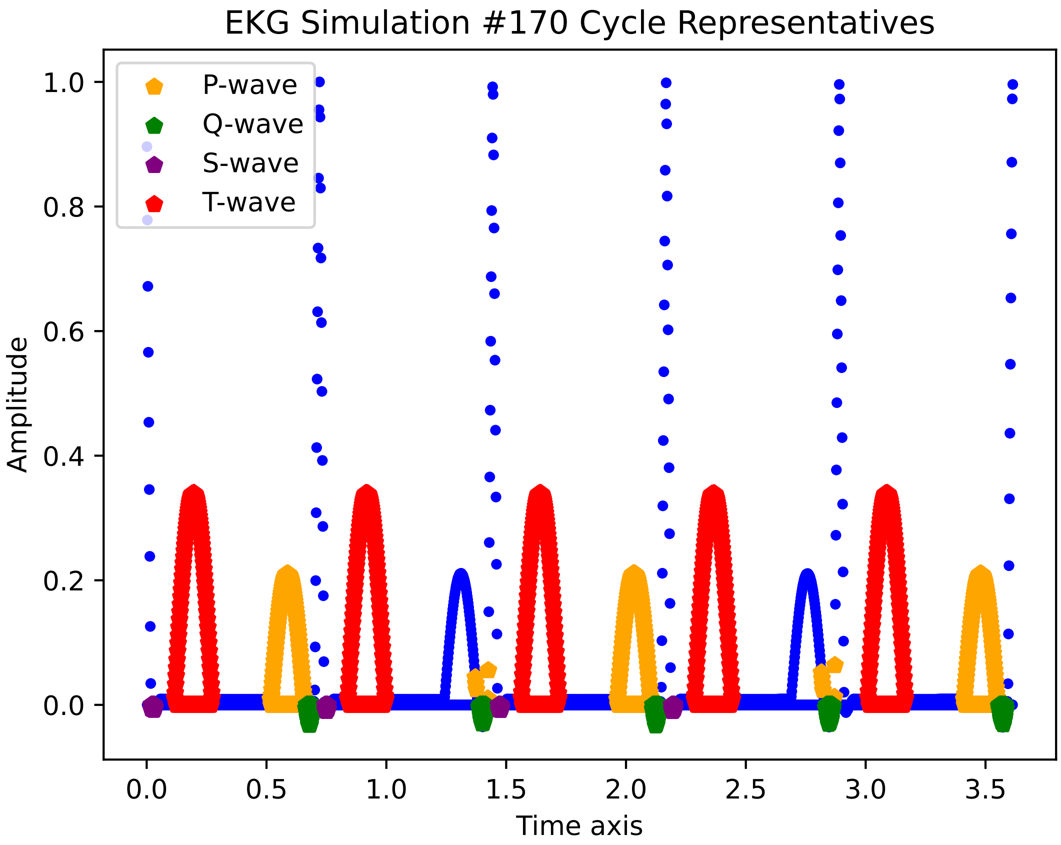

EKG Simulation #170 Cycle Representatives
P-wave
Q-wave
S-wave
T-wave
1.0
0.8
0.6
0.4
0.2
0.0
Amplitude
0.0
0.5
1.0
1.5
2.0
2.5
3.0
3.5
Time axis

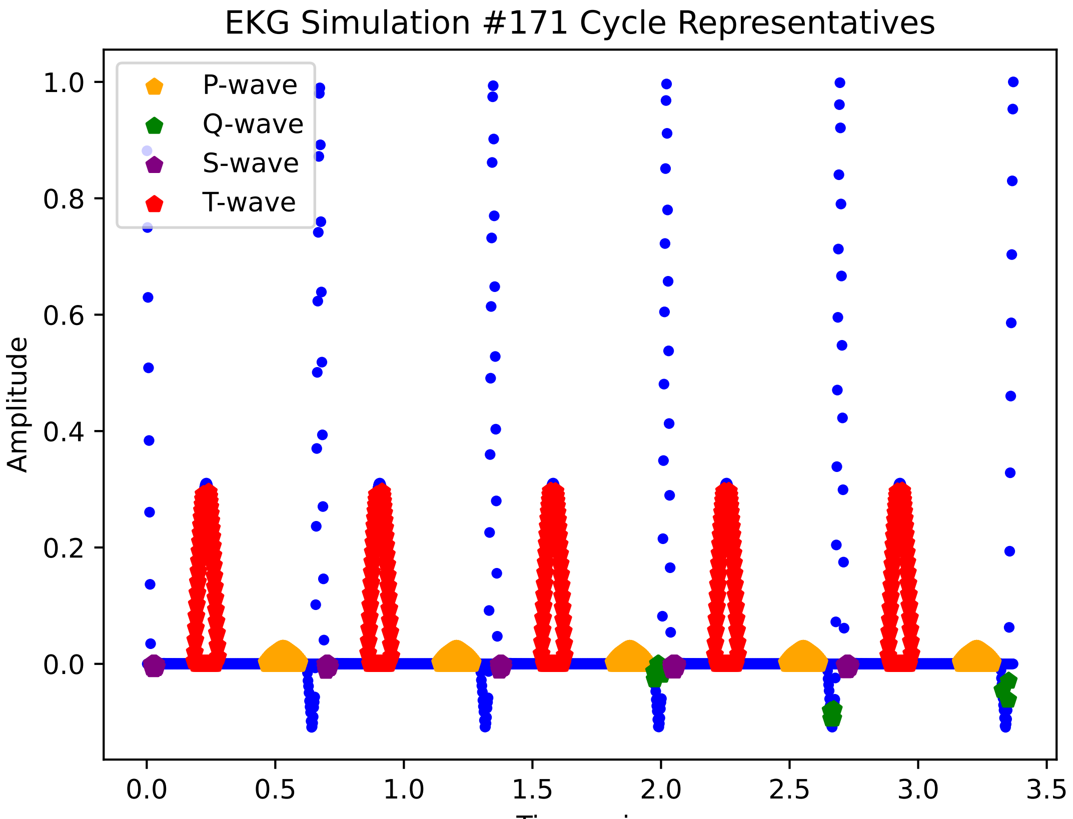
EKG Simulation #171 Cycle Representatives
P-wave
Q-wave
S-wave
T-wave
Amplitude
Time axis
1.0
0.8
0.6
0.4
0.2
0.0
0.0
0.5
1.0
1.5
2.0
2.5
3.0
3.5

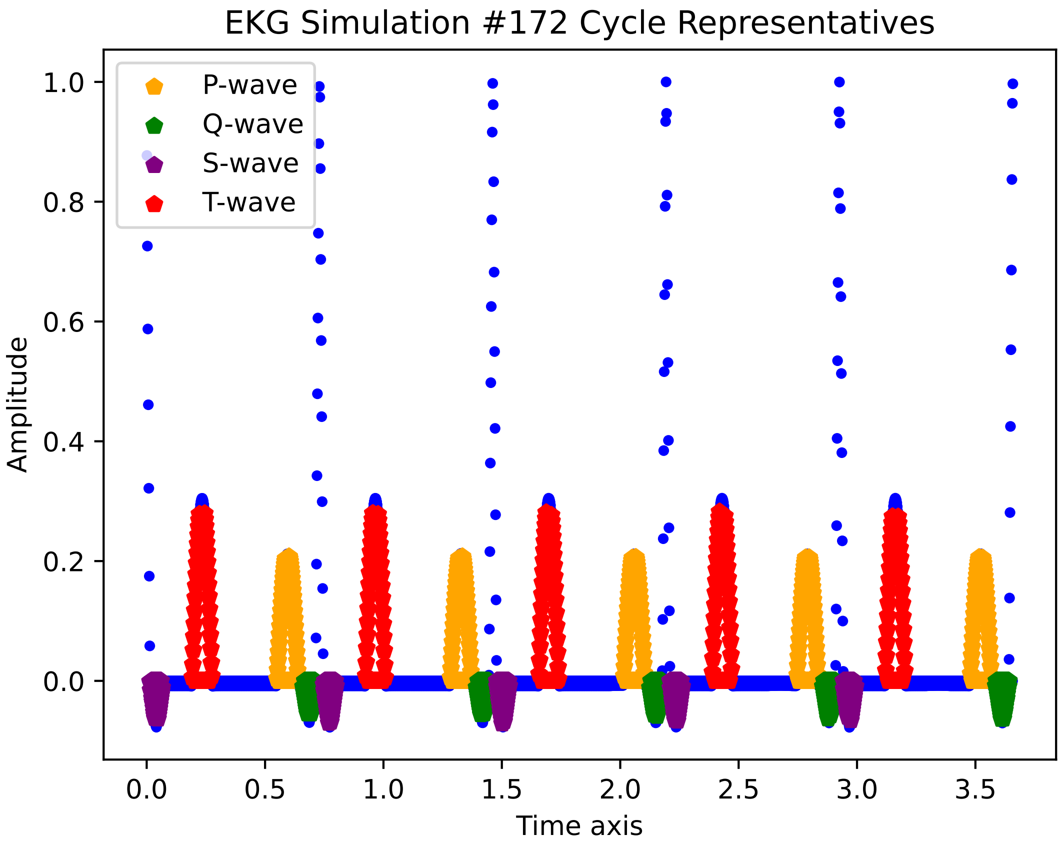
EKG Simulation #172 Cycle Representatives
P-wave
Q-wave
S-wave
T-wave
Amplitude
Time axis
1.0
0.8
0.6
0.4
0.2
0.0
0.0
0.5
1.0
1.5
2.0
2.5
3.0
3.5

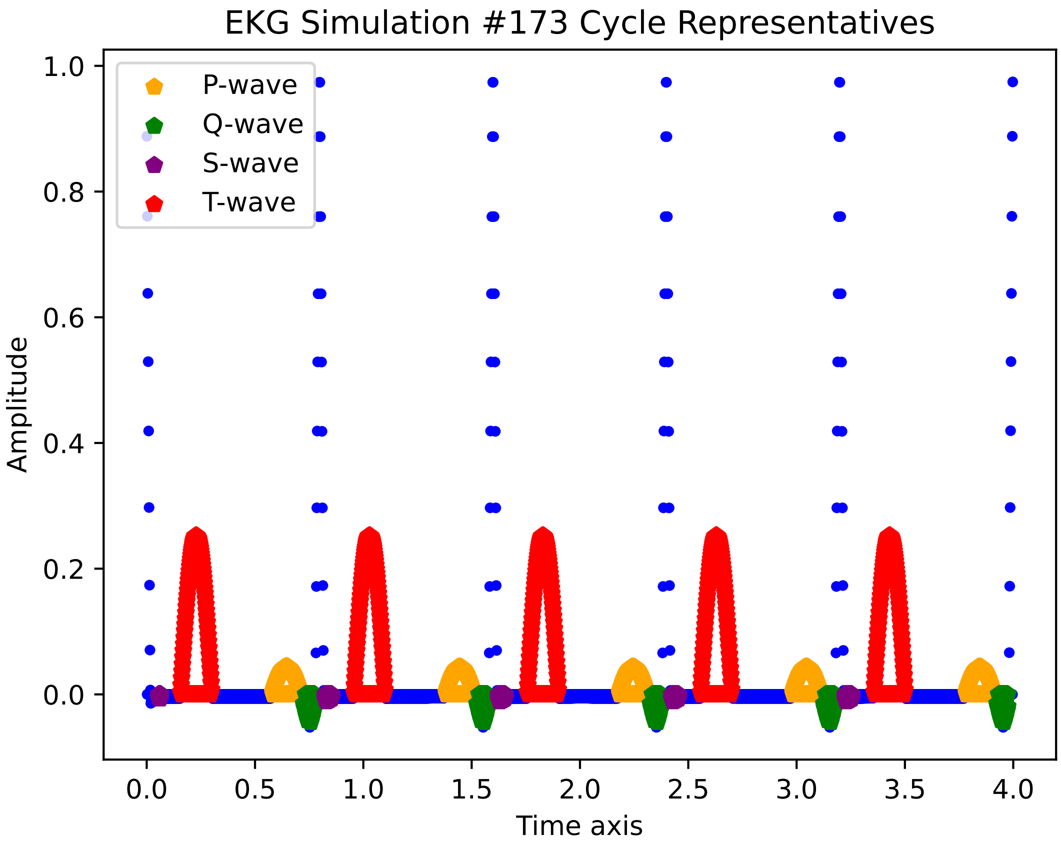

EKG Simulation #173 Cycle Representatives
P-wave
Q-wave
S-wave
T-wave
1.0
0.8
0.6
0.4
0.2
0.0
Amplitude
0.0
0.5
1.0
1.5
2.0
2.5
3.0
3.5
4.0
Time axis

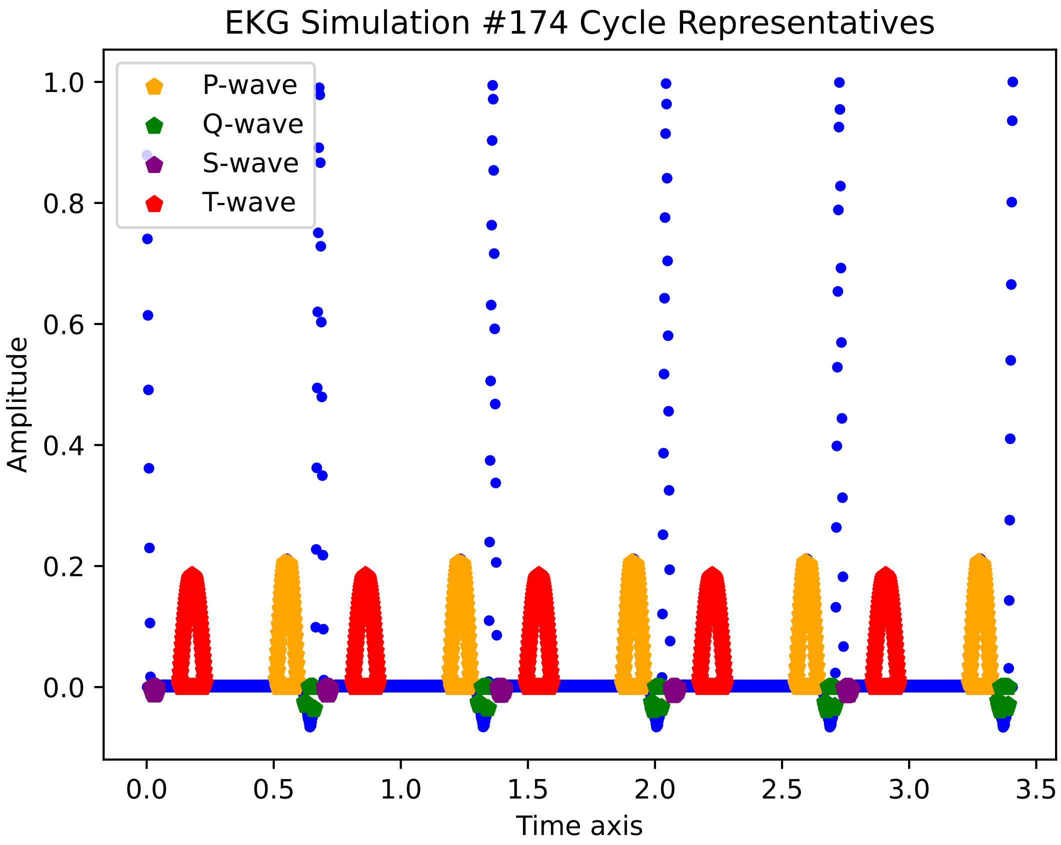

EKG Simulation #174 Cycle Representatives
P-wave
Q-wave
S-wave
T-wave
Amplitude
Time axis
1.0
0.8
0.6
0.4
0.2
0.0
0.0
0.5
1.0
1.5
2.0
2.5
3.0
3.5

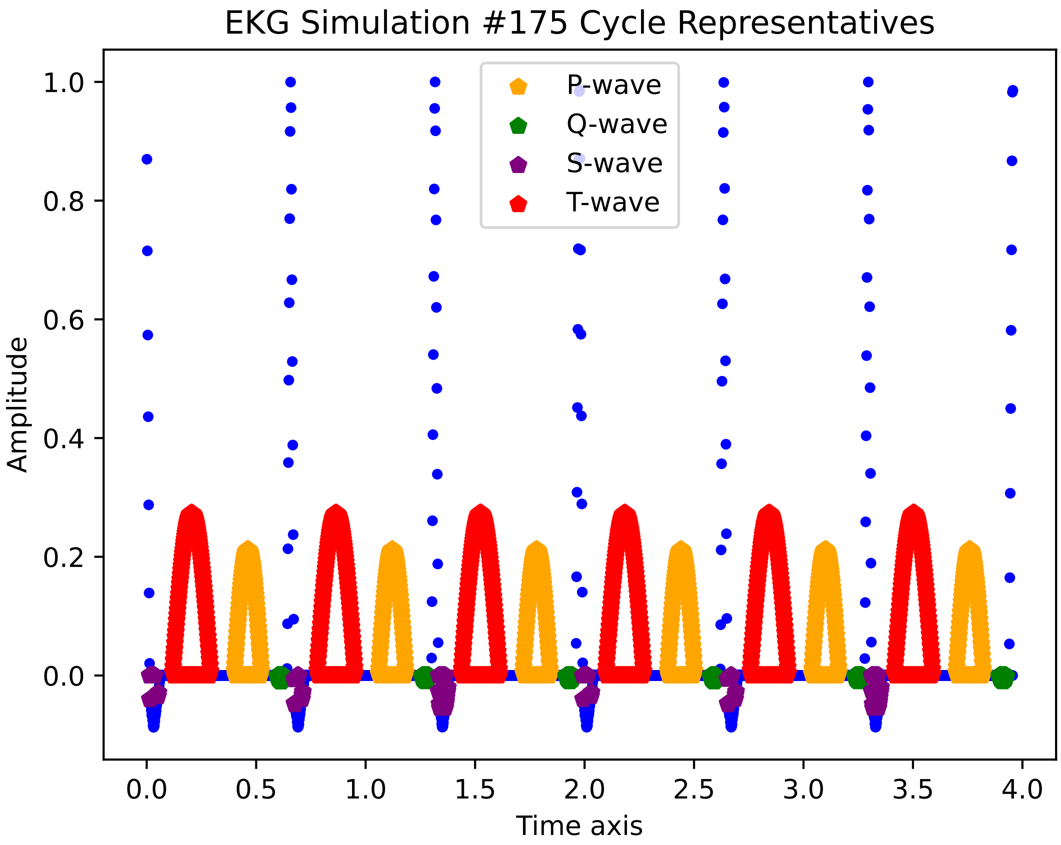
EKG Simulation #175 Cycle Representatives
P-wave
Q-wave
S-wave
T-wave
Amplitude
Time axis
1.0
0.8
0.6
0.4
0.2
0.0
0.0
0.5
1.0
1.5
2.0
2.5
3.0
3.5
4.0

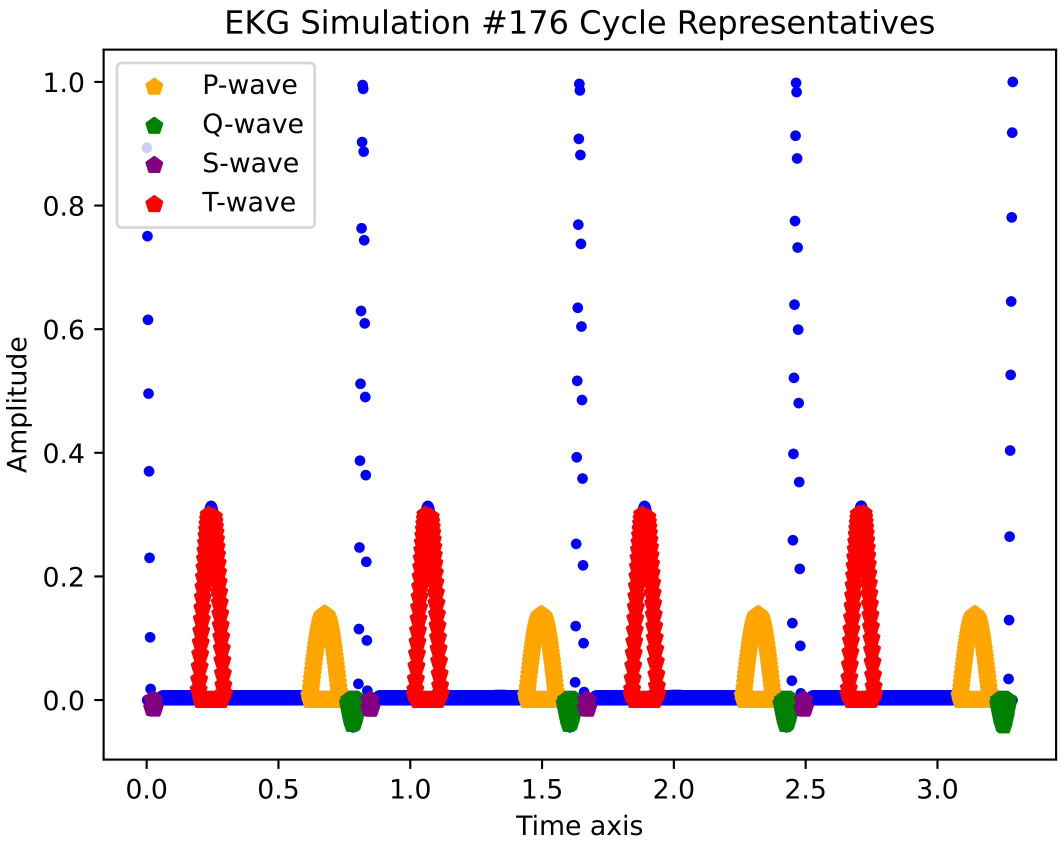
EKG Simulation #176 Cycle Representatives
P-wave
Q-wave
S-wave
T-wave
Amplitude
1.0
0.8
0.6
0.4
0.2
0.0
0.0
0.5
1.0
1.5
2.0
2.5
3.0
Time axis

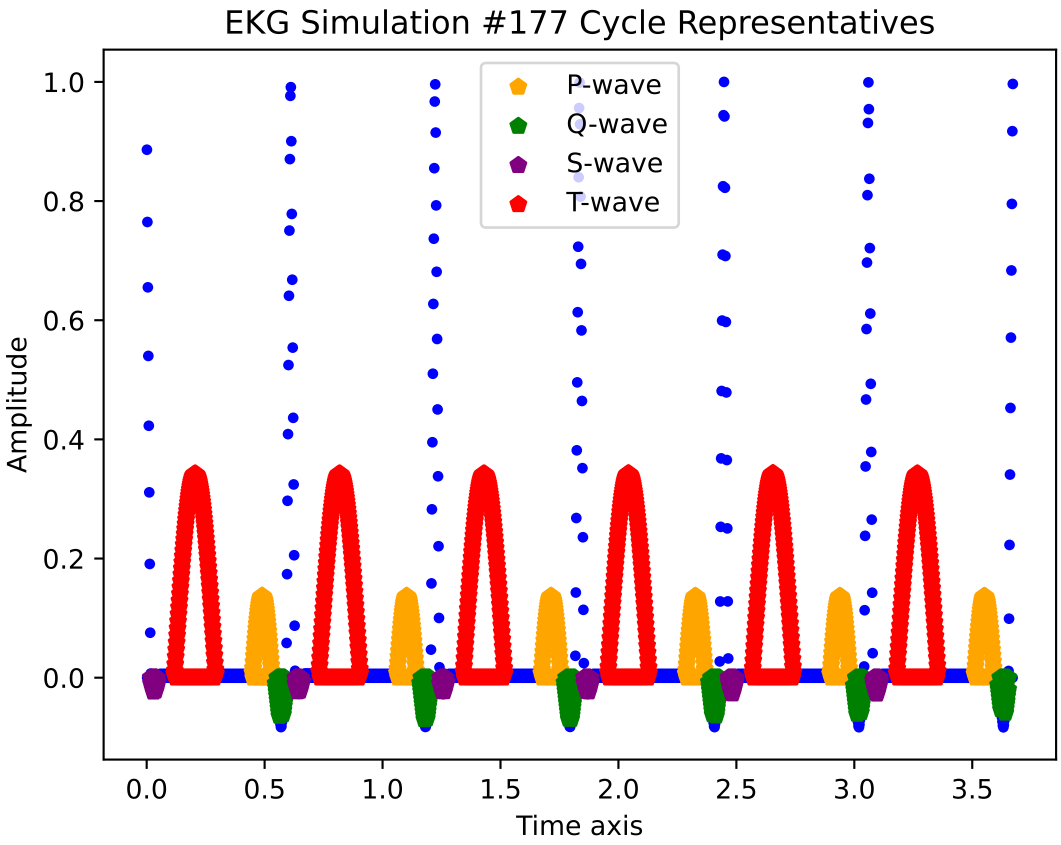
EKG Simulation #177 Cycle Representatives
P-wave
Q-wave
S-wave
T-wave
Amplitude
1.0
0.8
0.6
0.4
0.2
0.0
0.0
0.5
1.0
1.5
2.0
2.5
3.0
3.5
Time axis

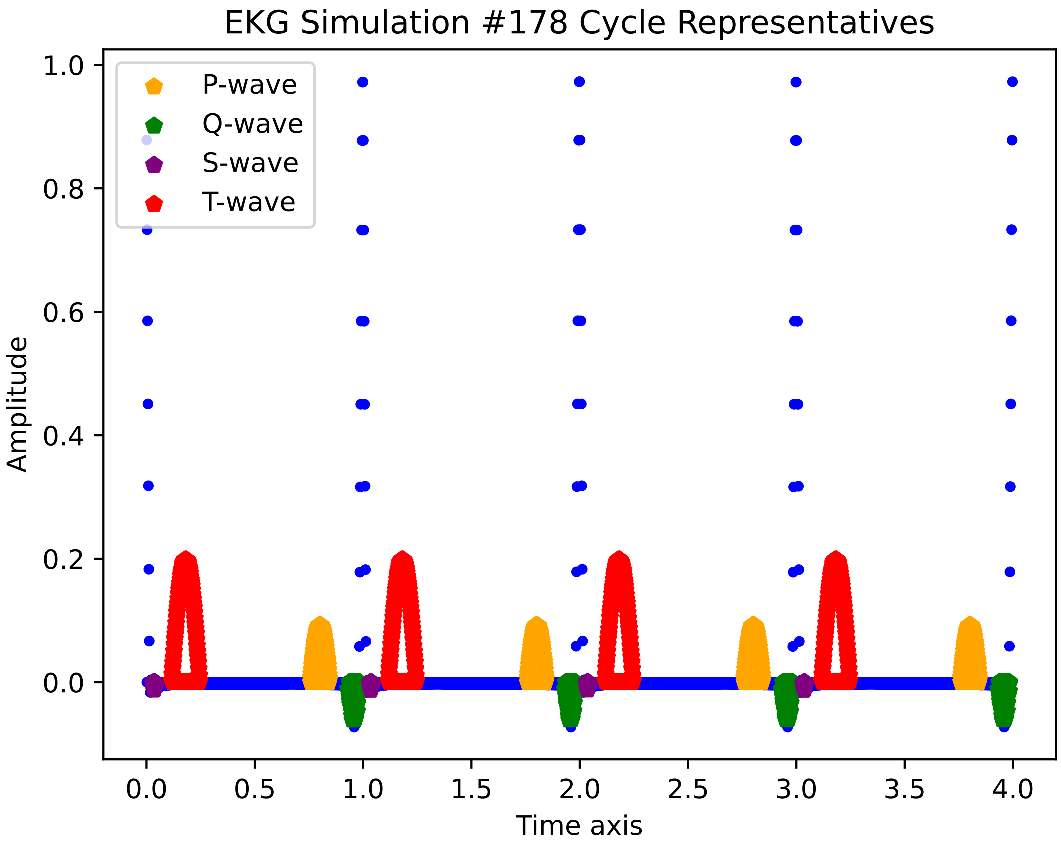

EKG Simulation #178 Cycle Representatives
P-wave
Q-wave
S-wave
T-wave
Amplitude
Time axis
1.0
0.8
0.6
0.4
0.2
0.0
0.0
0.5
1.0
1.5
2.0
2.5
3.0
3.5
4.0

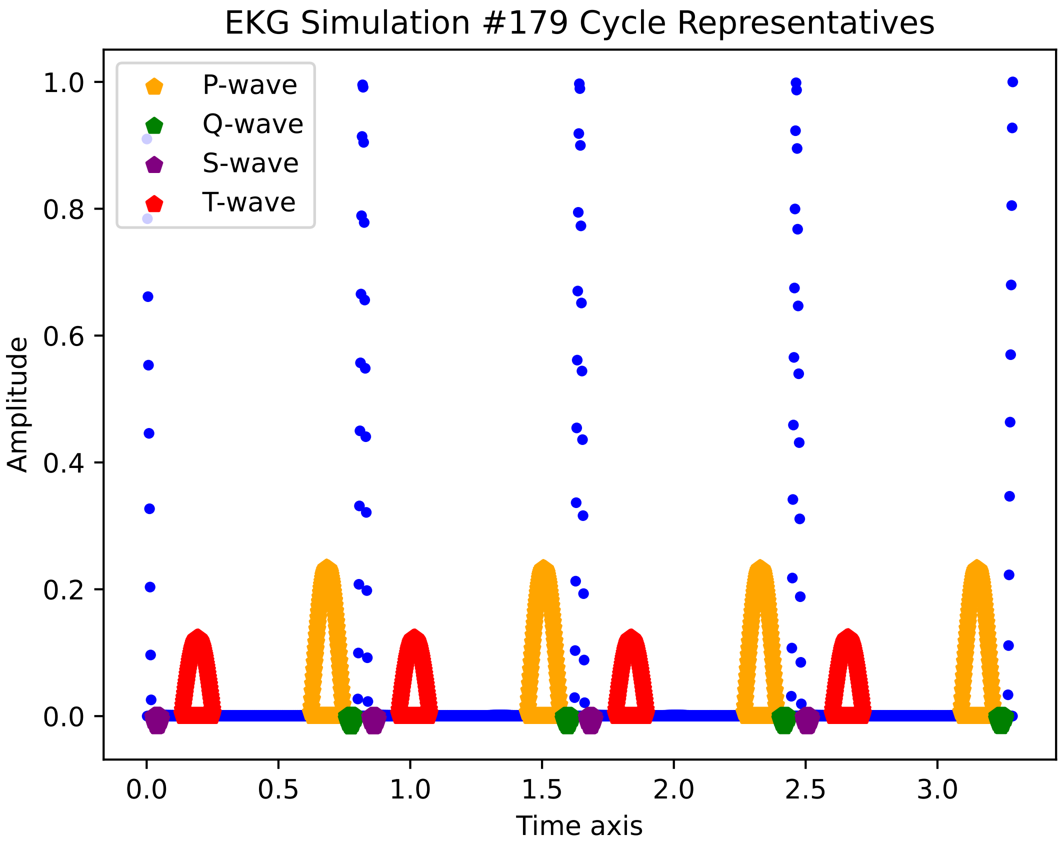
EKG Simulation #179 Cycle Representatives
P-wave
Q-wave
S-wave
T-wave
Amplitude
Time axis
1.0
0.8
0.6
0.4
0.2
0.0
0.0
0.5
1.0
1.5
2.0
2.5
3.0

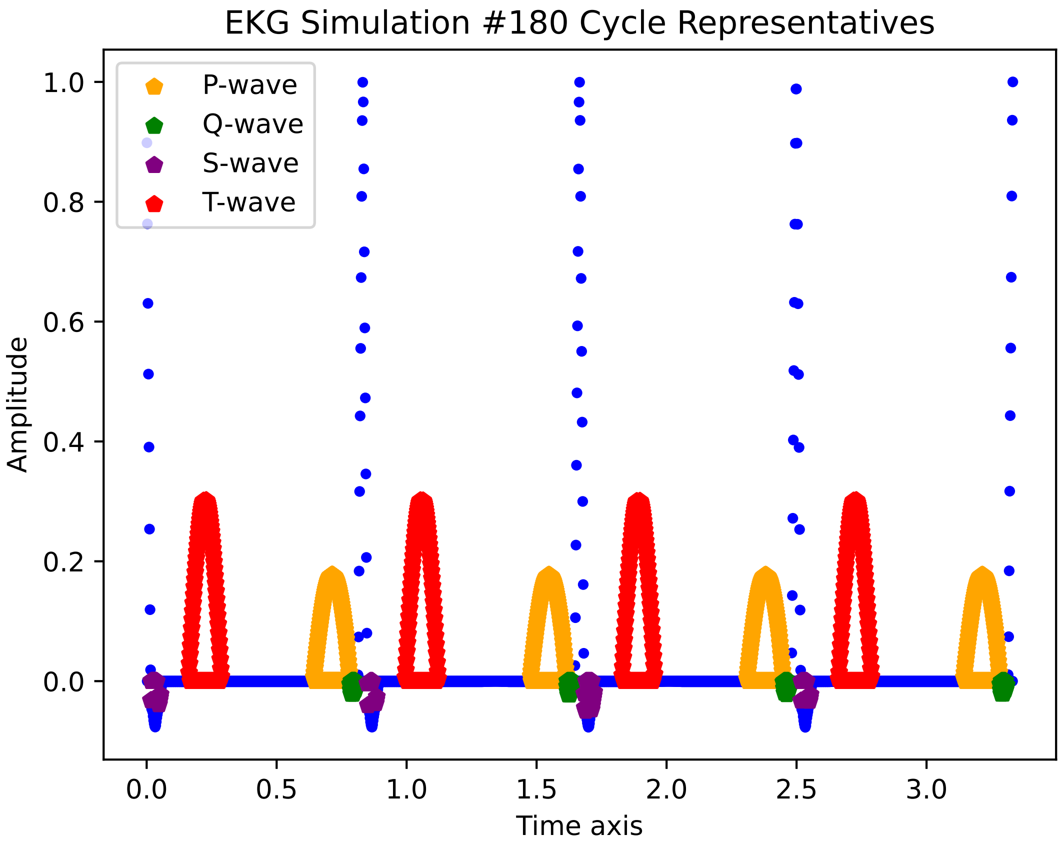
EKG Simulation #180 Cycle Representatives
P-wave
Q-wave
S-wave
T-wave
Amplitude
Time axis
1.0
0.8
0.6
0.4
0.2
0.0
0.0
0.5
1.0
1.5
2.0
2.5
3.0

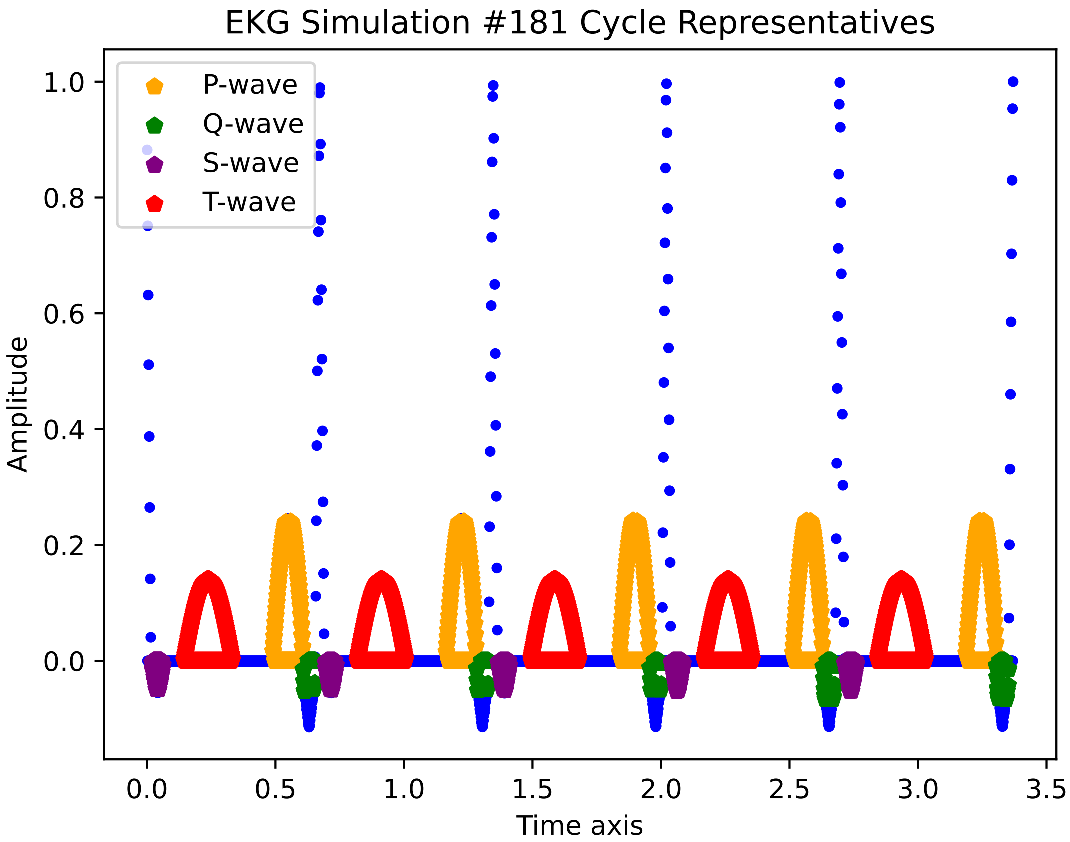
EKG Simulation #181 Cycle Representatives
P-wave
Q-wave
S-wave
T-wave
1.0
0.8
0.6
0.4
0.2
0.0
Amplitude
0.0
0.5
1.0
1.5
2.0
2.5
3.0
3.5
Time axis

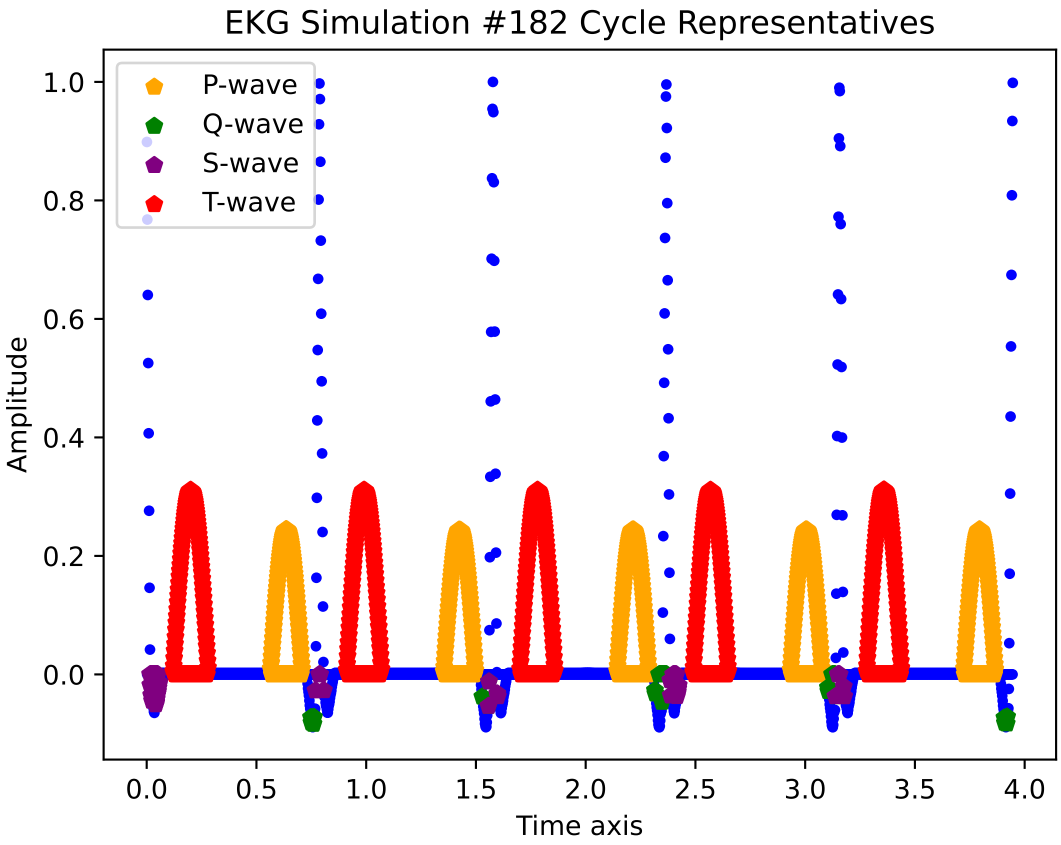
EKG Simulation #182 Cycle Representatives
P-wave
Q-wave
S-wave
T-wave
Amplitude
1.0
0.8
0.6
0.4
0.2
0.0
0.0
0.5
1.0
1.5
2.0
2.5
3.0
3.5
4.0
Time axis

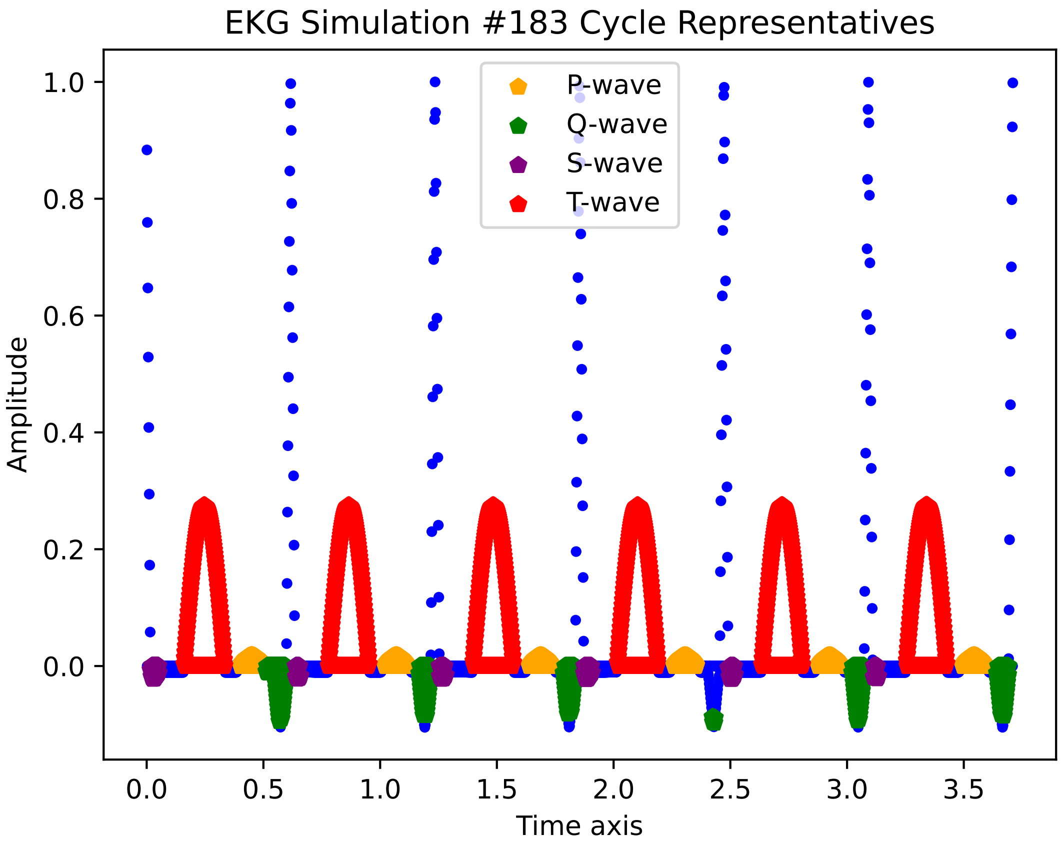
EKG Simulation #183 Cycle Representatives
P-wave
Q-wave
S-wave
T-wave
Amplitude
Time axis
1.0
0.8
0.6
0.4
0.2
0.0
0.0
0.5
1.0
1.5
2.0
2.5
3.0
3.5

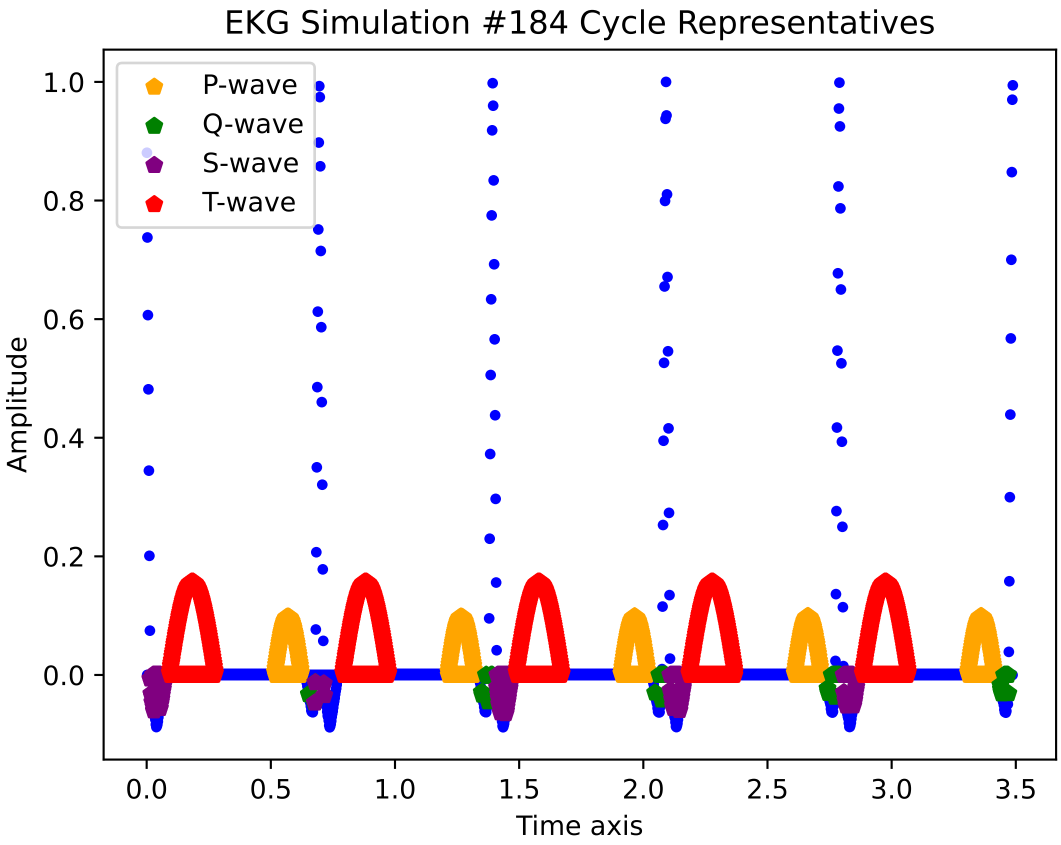
EKG Simulation #184 Cycle Representatives
P-wave
Q-wave
S-wave
T-wave
Amplitude
1.0
0.8
0.6
0.4
0.2
0.0
0.0
0.5
1.0
1.5
2.0
2.5
3.0
3.5
Time axis

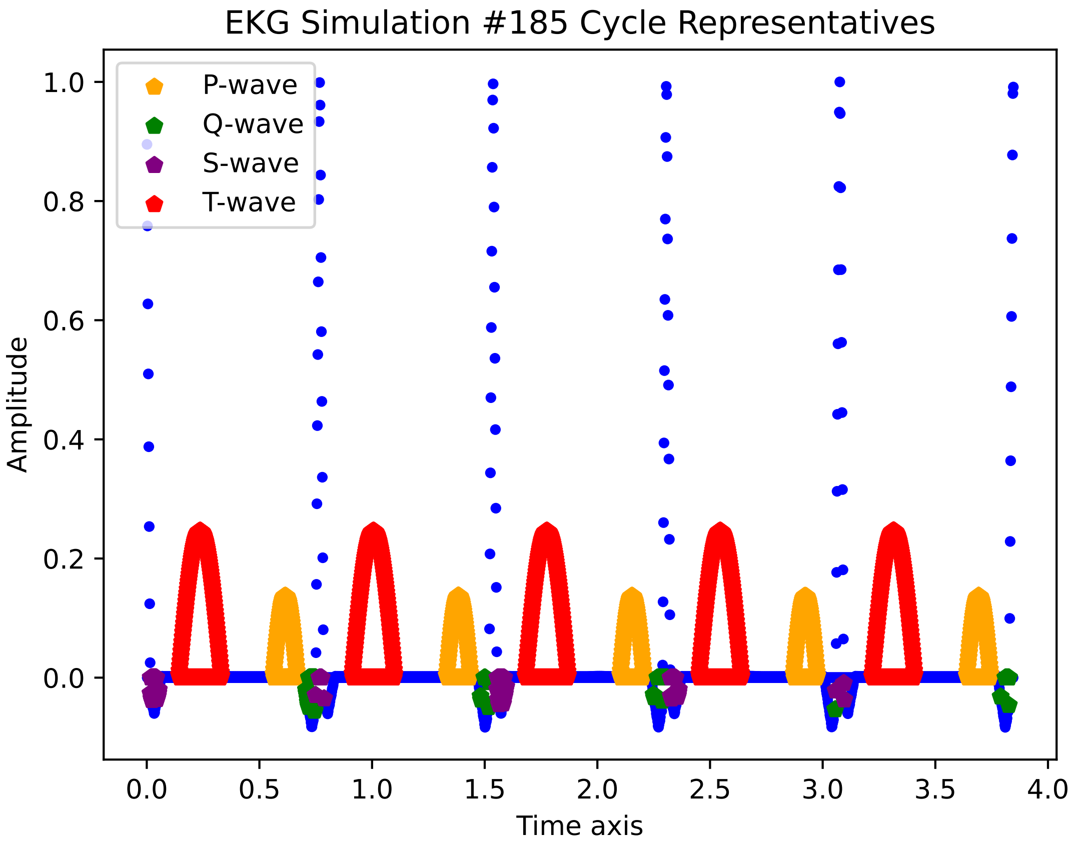
EKG Simulation #185 Cycle Representatives
P-wave
Q-wave
S-wave
T-wave
Amplitude
Time axis
1.0
0.8
0.6
0.4
0.2
0.0
0.0
0.5
1.0
1.5
2.0
2.5
3.0
3.5
4.0

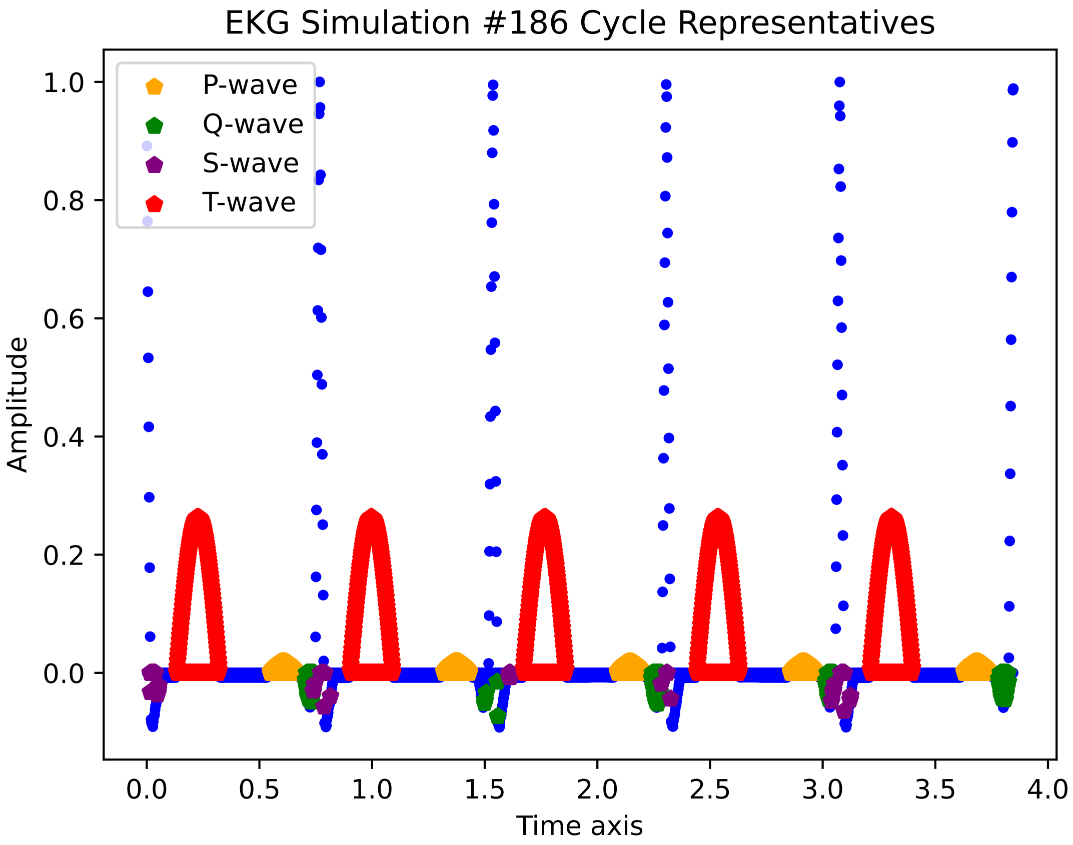
EKG Simulation #186 Cycle Representatives
P-wave
Q-wave
S-wave
T-wave
Amplitude
Time axis
1.0
0.8
0.6
0.4
0.2
0.0
0.0
0.5
1.0
1.5
2.0
2.5
3.0
3.5
4.0

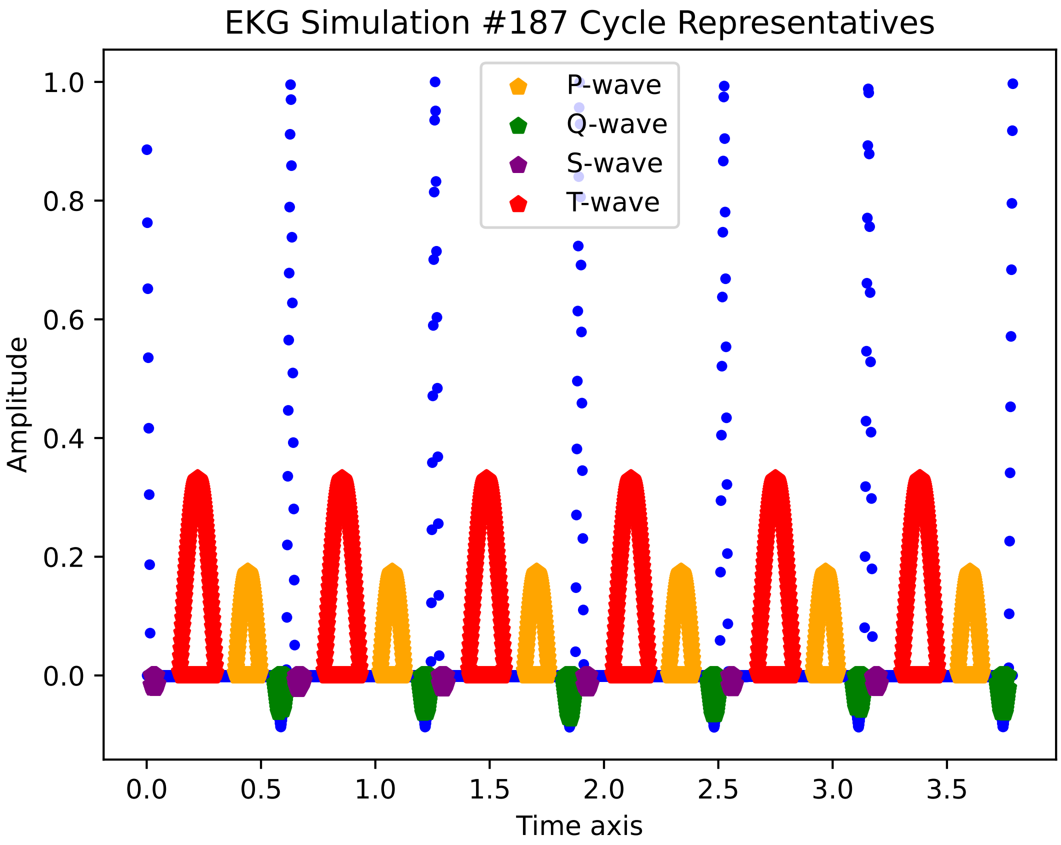

EKG Simulation #187 Cycle Representatives
P-wave
Q-wave
S-wave
T-wave
Amplitude
Time axis
1.0
0.8
0.6
0.4
0.2
0.0
0.0
0.5
1.0
1.5
2.0
2.5
3.0
3.5

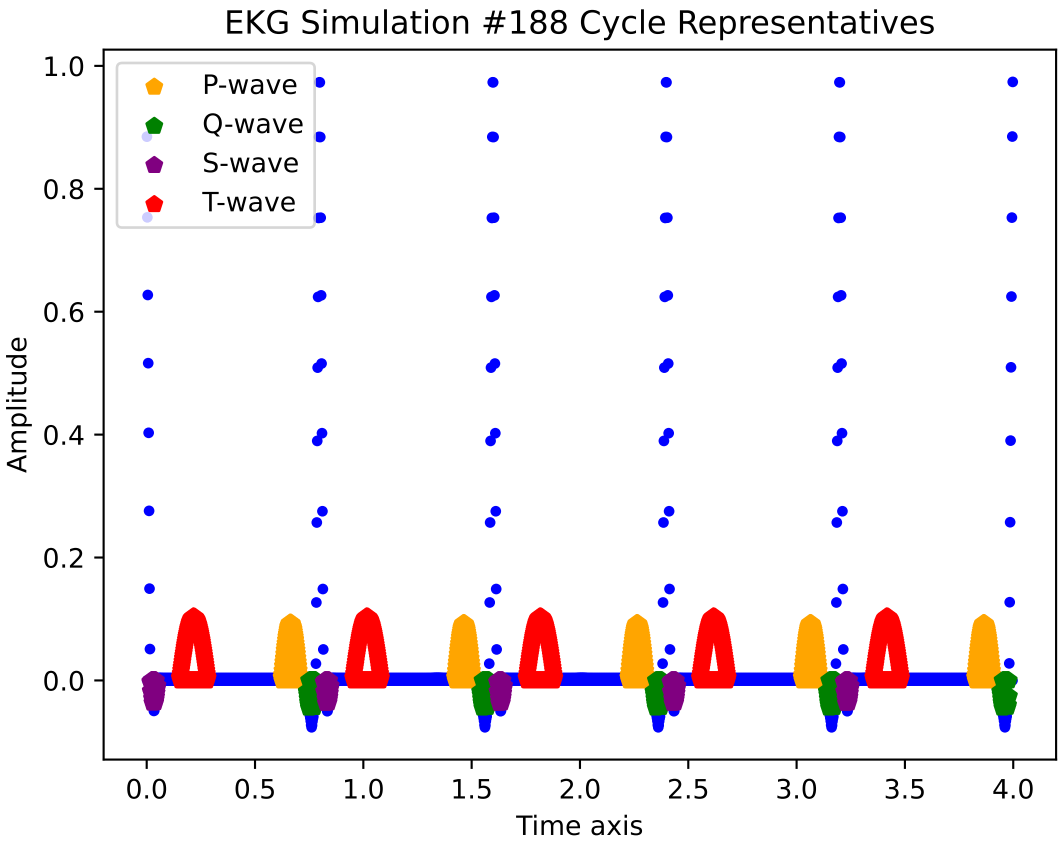
EKG Simulation #188 Cycle Representatives
P-wave
Q-wave
S-wave
T-wave
Amplitude
Time axis
1.0
0.8
0.6
0.4
0.2
0.0
0.0
0.5
1.0
1.5
2.0
2.5
3.0
3.5
4.0

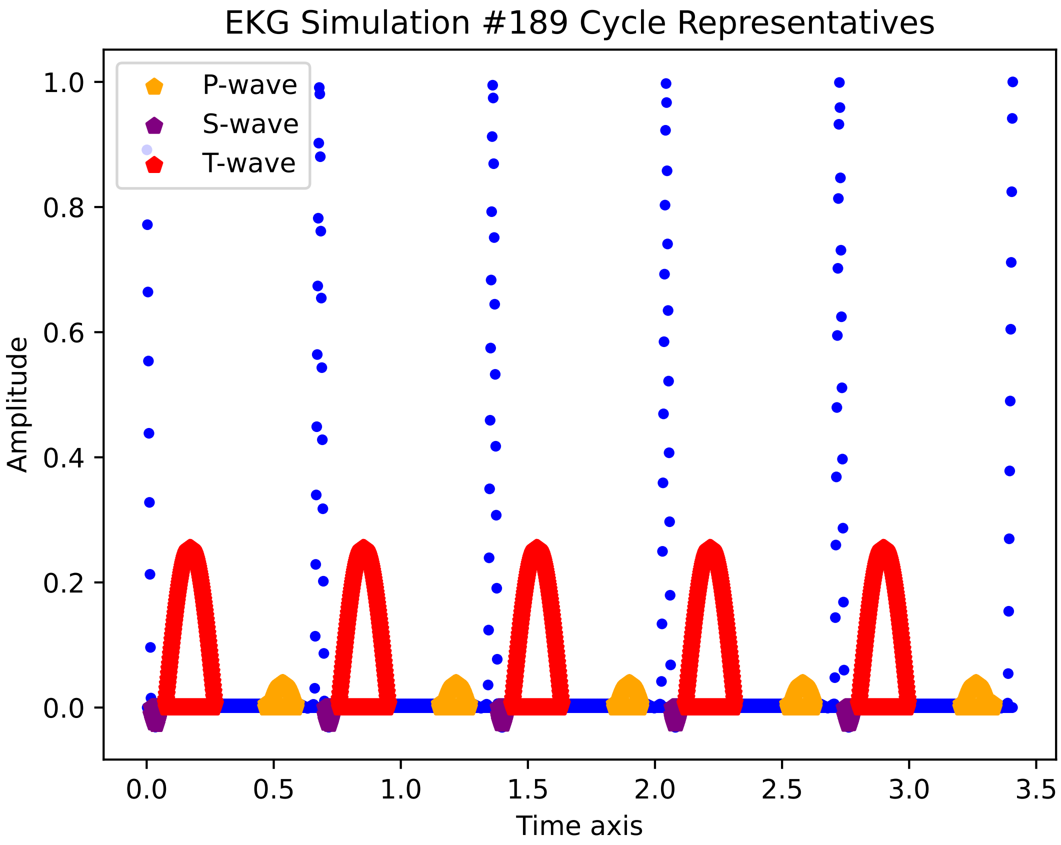

EKG Simulation #189 Cycle Representatives
P-wave
S-wave
T-wave
Amplitude
Time axis
1.0
0.8
0.6
0.4
0.2
0.0
0.0
0.5
1.0
1.5
2.0
2.5
3.0
3.5

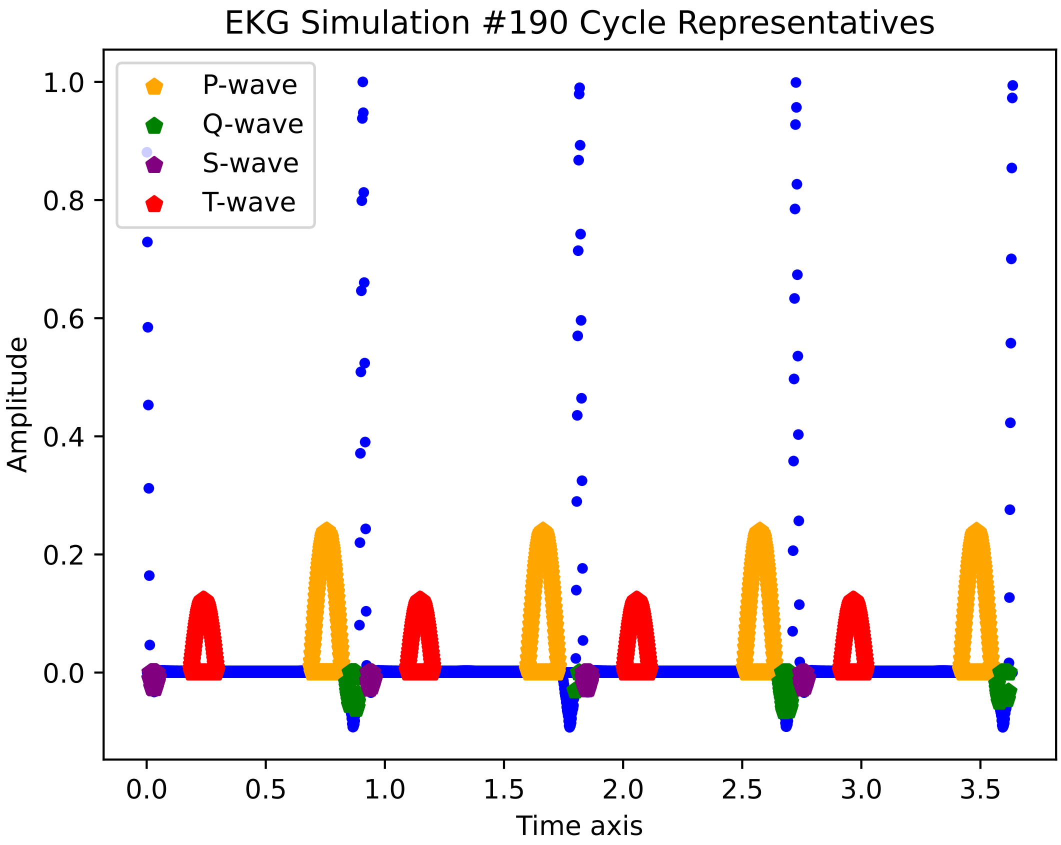
EKG Simulation #190 Cycle Representatives
P-wave
Q-wave
S-wave
T-wave
Amplitude
1.0
0.8
0.6
0.4
0.2
0.0
0.0
0.5
1.0
1.5
2.0
2.5
3.0
3.5
Time axis

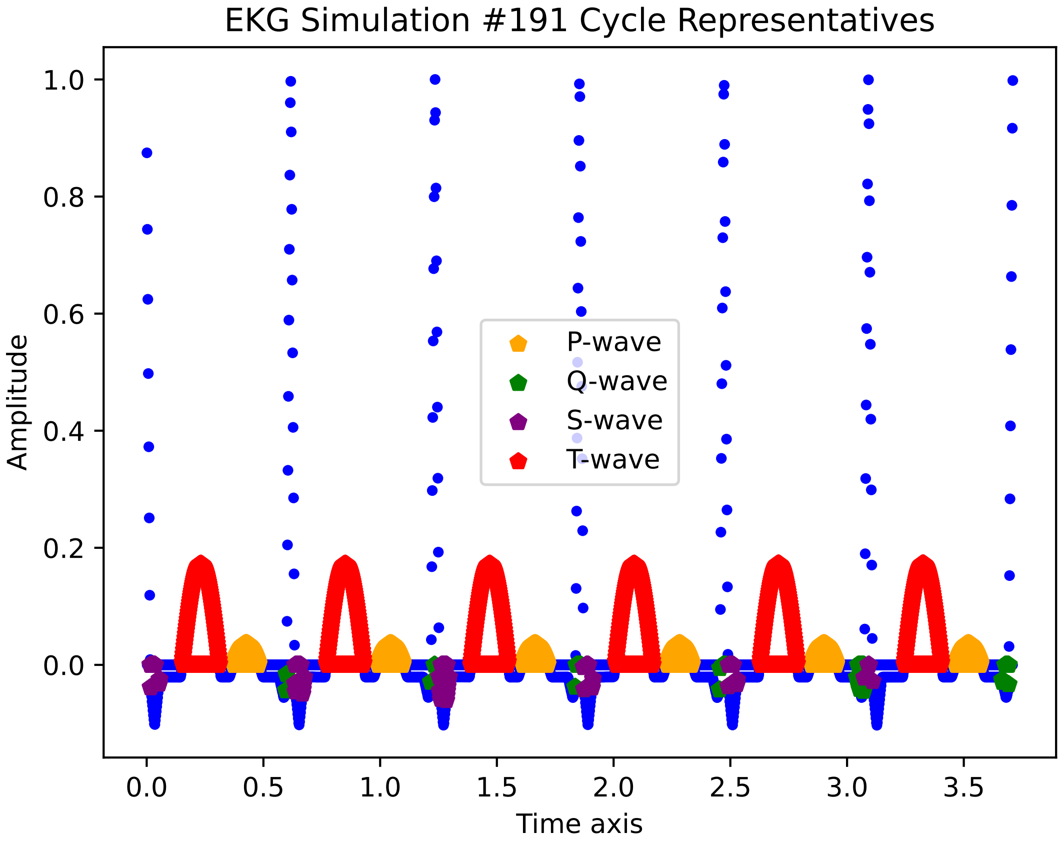
EKG Simulation #191 Cycle Representatives
Amplitude
1.0
0.8
0.6
0.4
0.2
0.0
0.0
0.5
1.0
1.5
2.0
2.5
3.0
3.5
Time axis
P-wave
Q-wave
S-wave
T-wave

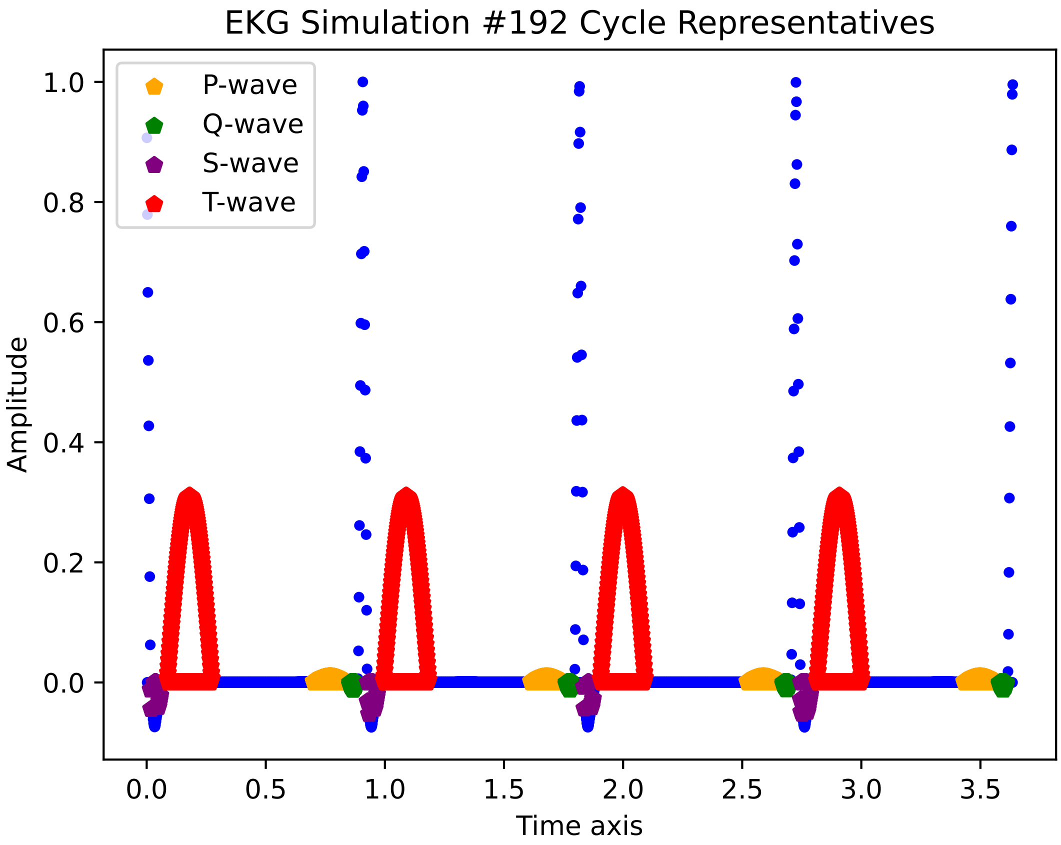
EKG Simulation #192 Cycle Representatives
P-wave
Q-wave
S-wave
T-wave
Amplitude
Time axis
1.0
0.8
0.6
0.4
0.2
0.0
0.0
0.5
1.0
1.5
2.0
2.5
3.0
3.5

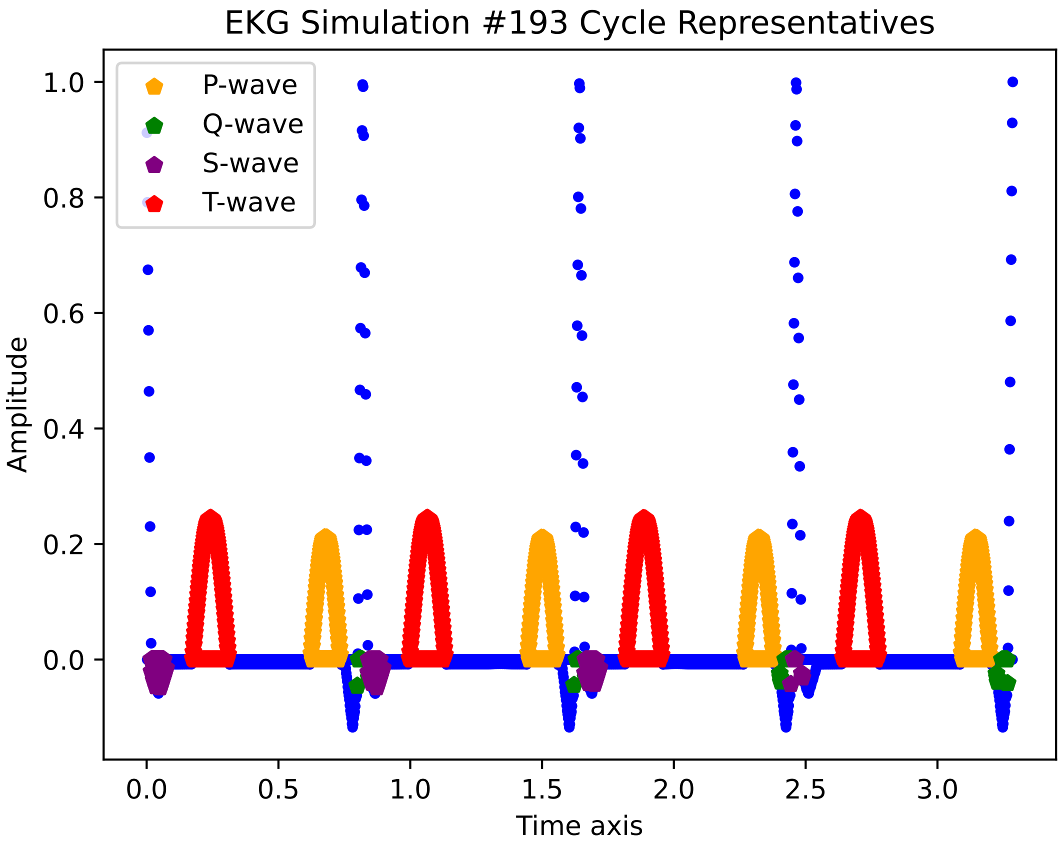

EKG Simulation #193 Cycle Representatives
P-wave
Q-wave
S-wave
T-wave
1.0
0.8
0.6
0.4
0.2
0.0
Amplitude
0.0
0.5
1.0
1.5
2.0
2.5
3.0
Time axis

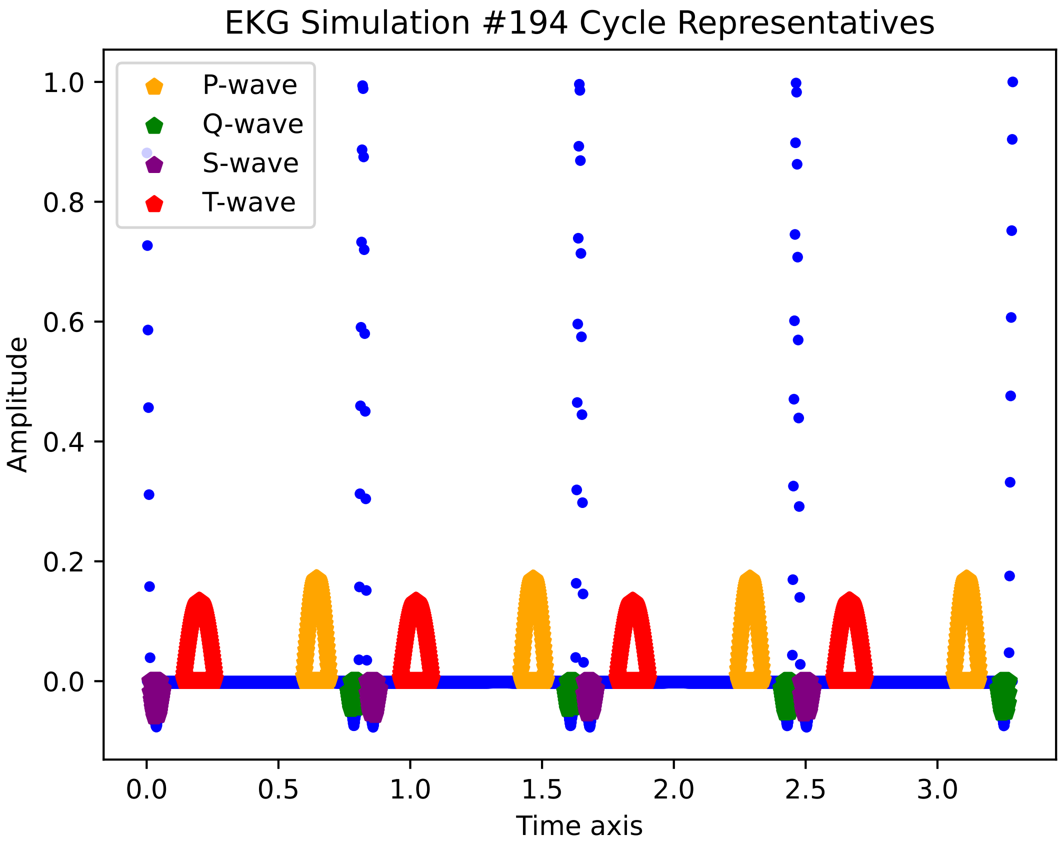

EKG Simulation #194 Cycle Representatives
P-wave
Q-wave
S-wave
T-wave
Amplitude
1.0
0.8
0.6
0.4
0.2
0.0
0.0
0.5
1.0
1.5
2.0
2.5
3.0
Time axis

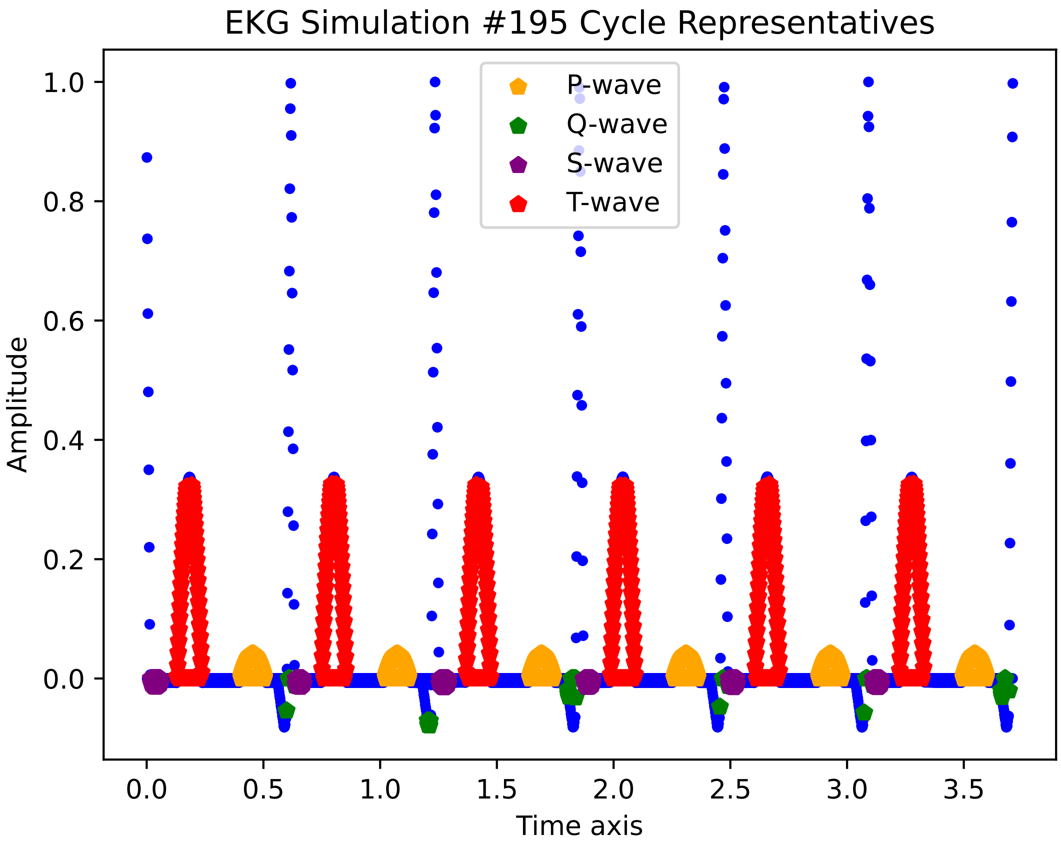

EKG Simulation #195 Cycle Representatives
P-wave
Q-wave
S-wave
T-wave
Amplitude
Time axis
1.0
0.8
0.6
0.4
0.2
0.0
0.0
0.5
1.0
1.5
2.0
2.5
3.0
3.5

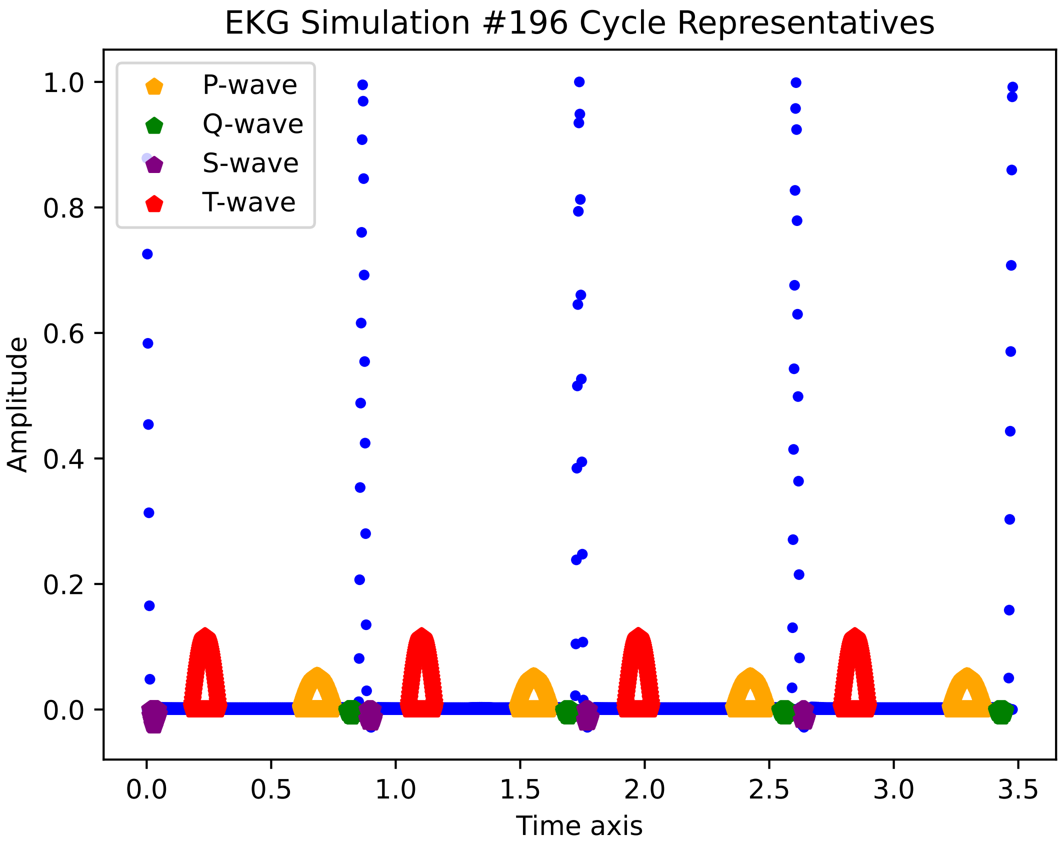
EKG Simulation #196 Cycle Representatives
P-wave
Q-wave
S-wave
T-wave
Amplitude
Time axis
1.0
0.8
0.6
0.4
0.2
0.0
0.0
0.5
1.0
1.5
2.0
2.5
3.0
3.5

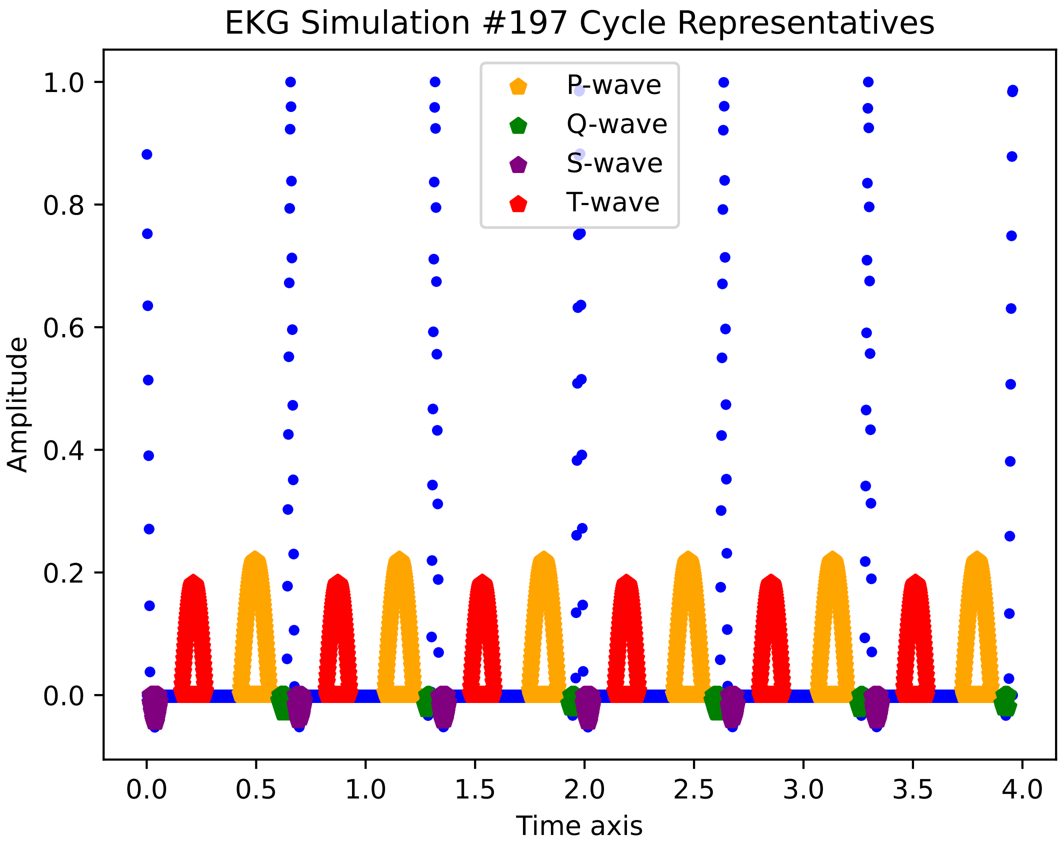
EKG Simulation #197 Cycle Representatives
P-wave
Q-wave
S-wave
T-wave
Amplitude
1.0
0.8
0.6
0.4
0.2
0.0
0.0
0.5
1.0
1.5
2.0
2.5
3.0
3.5
4.0
Time axis

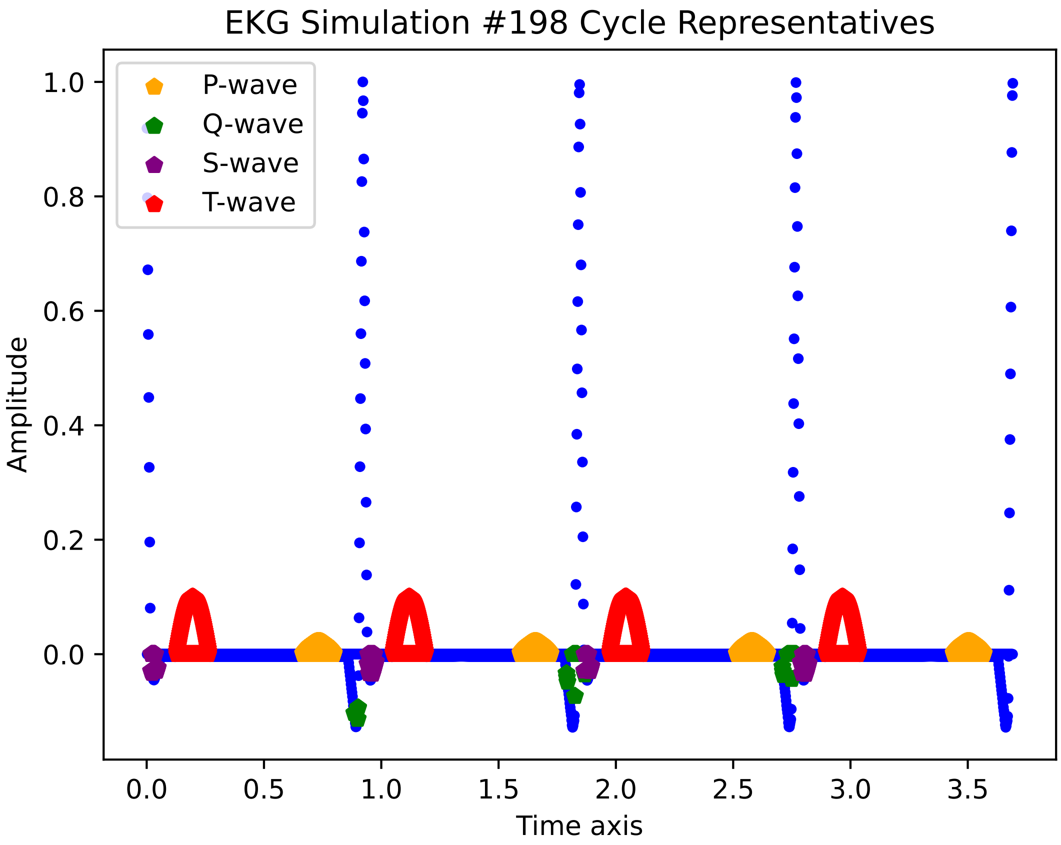

EKG Simulation #198 Cycle Representatives
P-wave
Q-wave
S-wave
T-wave
1.0
0.8
0.6
0.4
0.2
0.0
Amplitude
0.0
0.5
1.0
1.5
2.0
2.5
3.0
3.5
Time axis

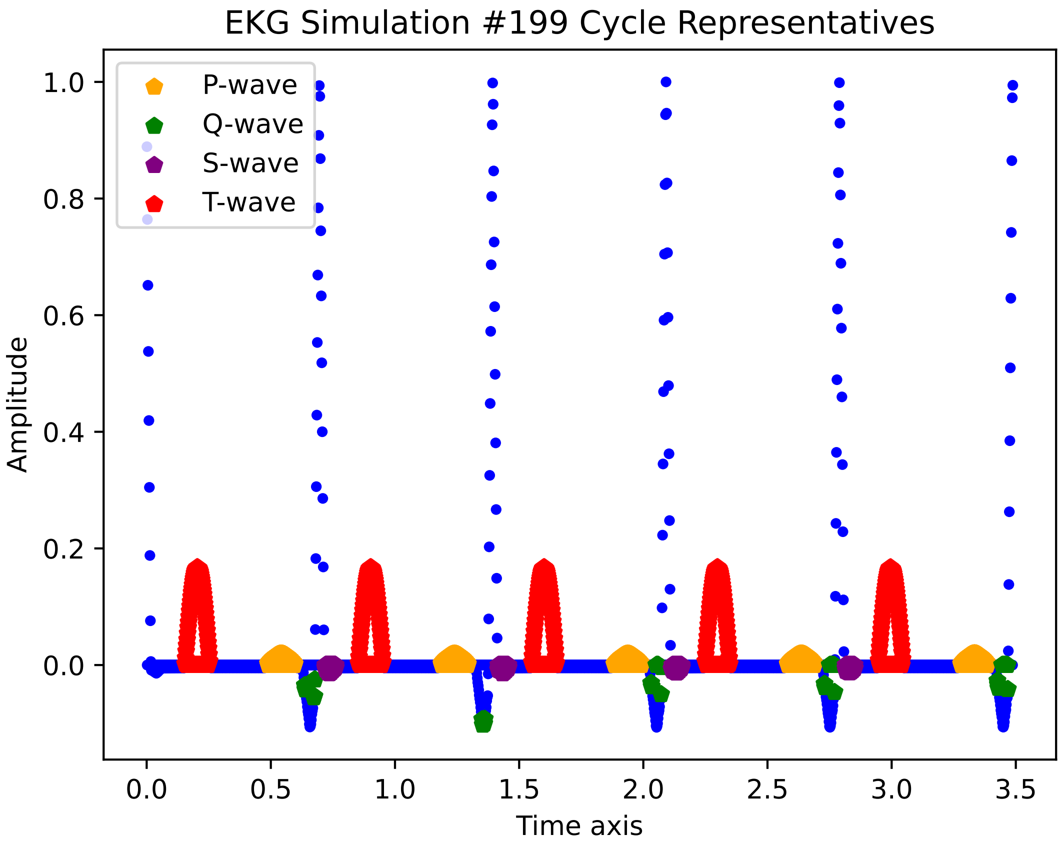
EKG Simulation #199 Cycle Representatives
P-wave
Q-wave
S-wave
T-wave
Amplitude
Time axis
1.0
0.8
0.6
0.4
0.2
0.0
0.0
0.5
1.0
1.5
2.0
2.5
3.0
3.5

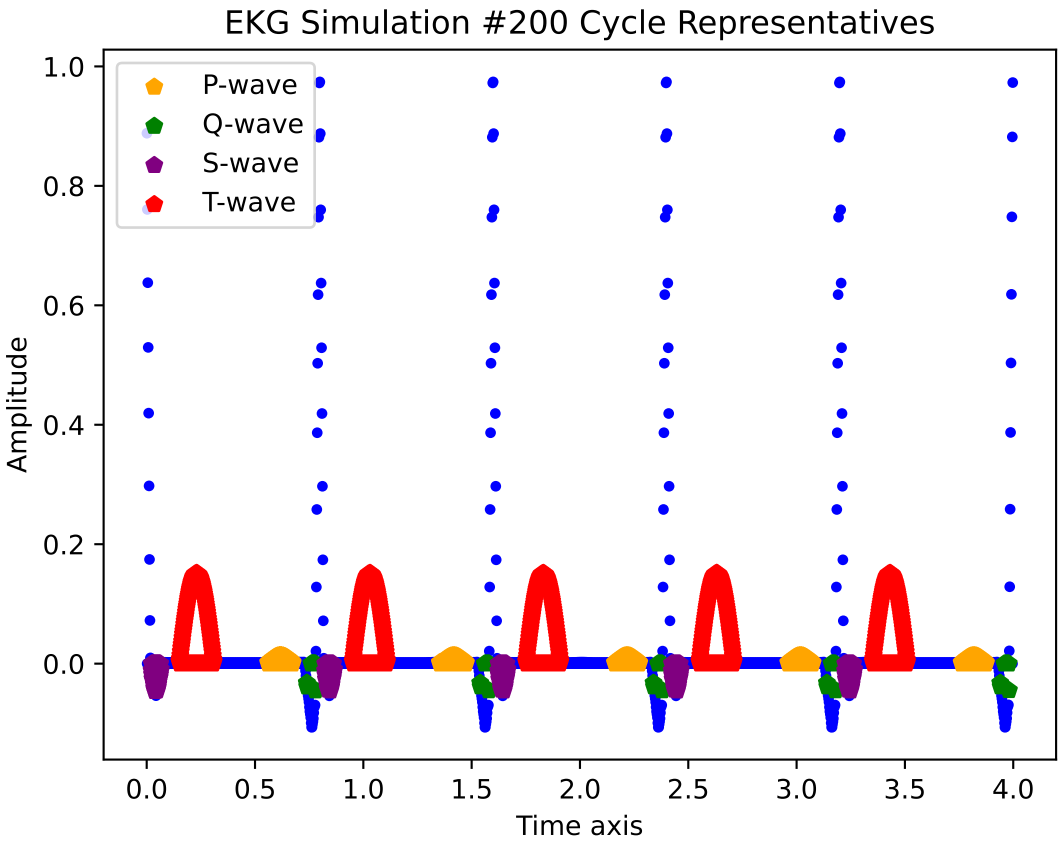

EKG Simulation #200 Cycle Representatives
P-wave
Q-wave
S-wave
T-wave
Amplitude
Time axis
1.0
0.8
0.6
0.4
0.2
0.0
0.0
0.5
1.0
1.5
2.0
2.5
3.0
3.5
4.0

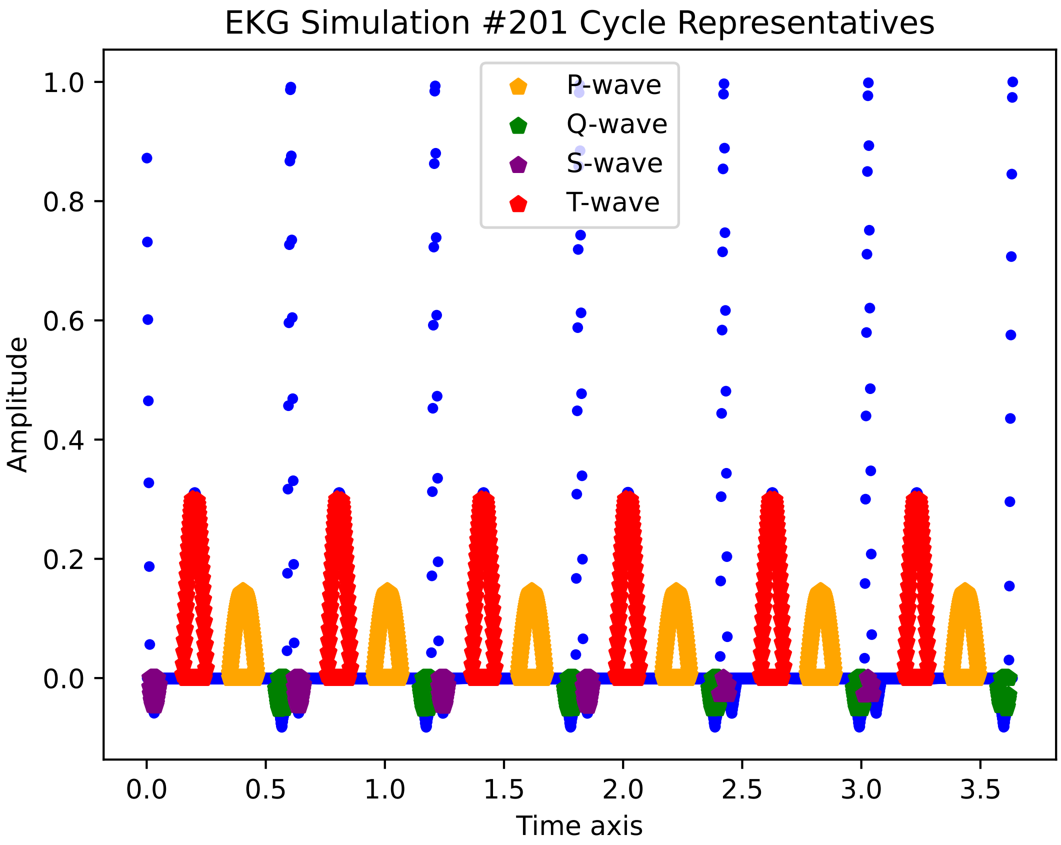

EKG Simulation #201 Cycle Representatives
P-wave
Q-wave
S-wave
T-wave
Amplitude
Time axis
1.0
0.8
0.6
0.4
0.2
0.0
0.0
0.5
1.0
1.5
2.0
2.5
3.0
3.5

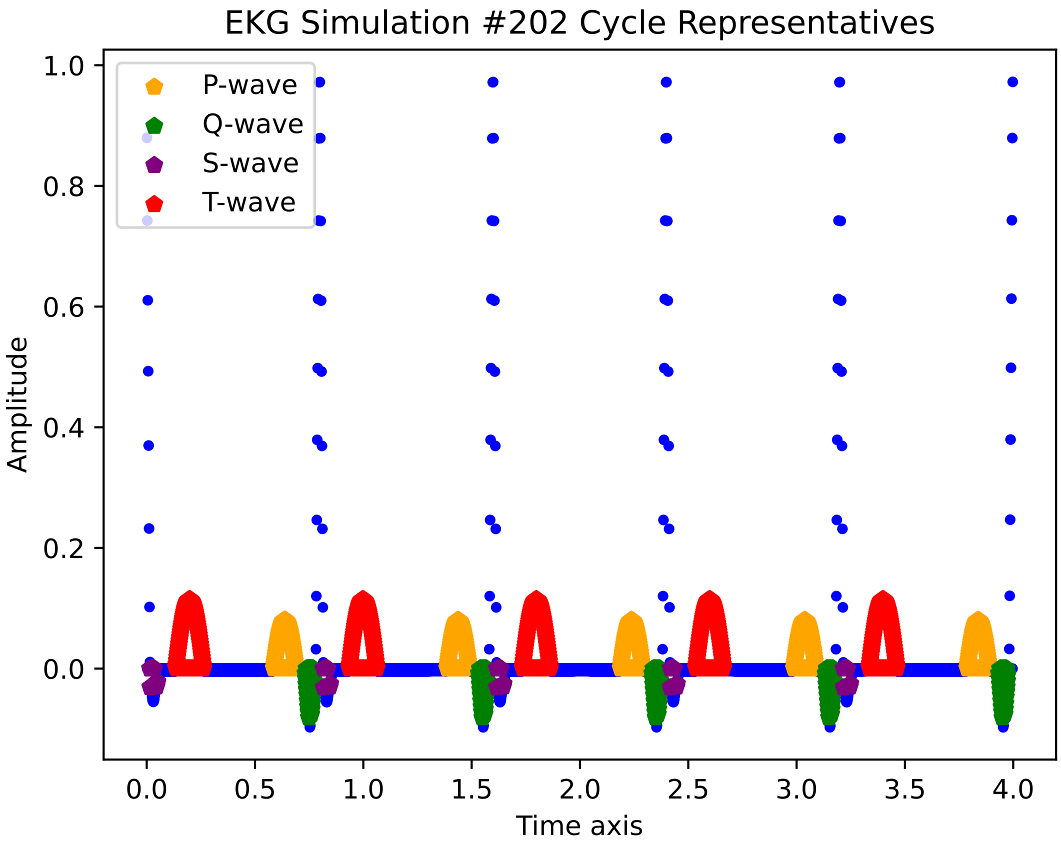

EKG Simulation #202 Cycle Representatives
P-wave
Q-wave
S-wave
T-wave
Amplitude
Time axis
1.0
0.8
0.6
0.4
0.2
0.0
0.0
0.5
1.0
1.5
2.0
2.5
3.0
3.5
4.0

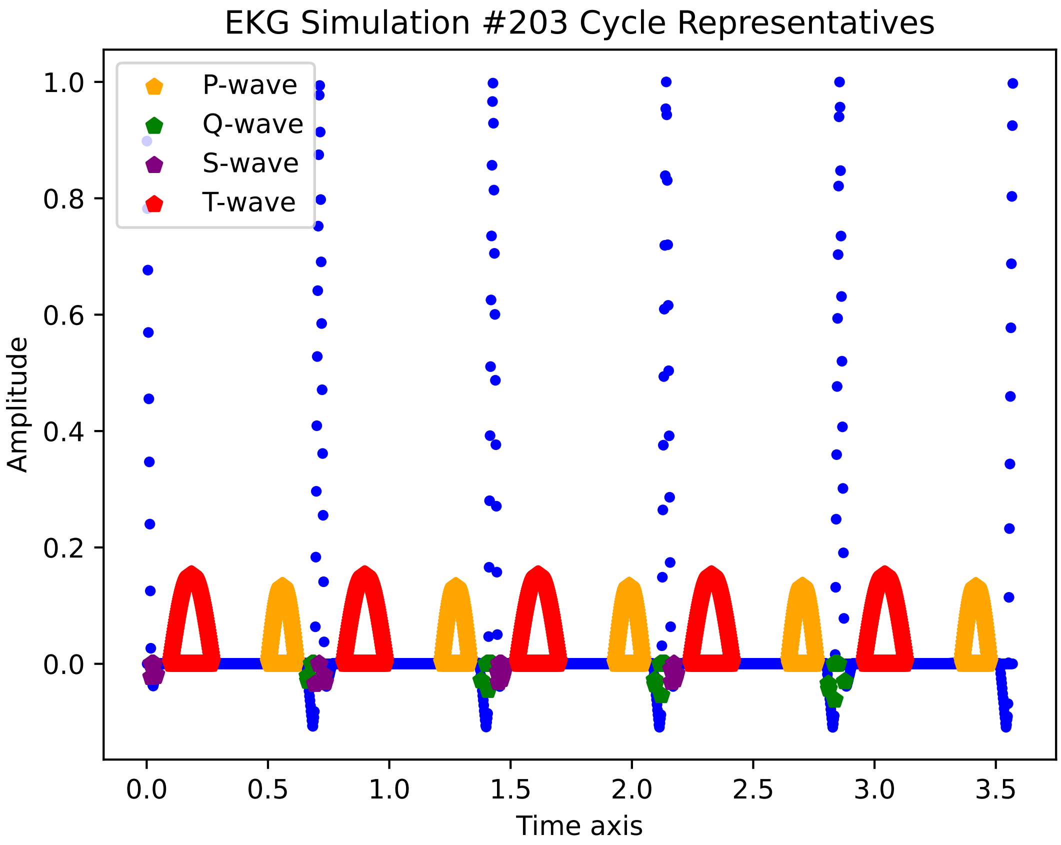

EKG Simulation #203 Cycle Representatives
P-wave
Q-wave
S-wave
T-wave
1.0
0.8
0.6
0.4
0.2
0.0
Amplitude
0.0
0.5
1.0
1.5
2.0
2.5
3.0
3.5
Time axis

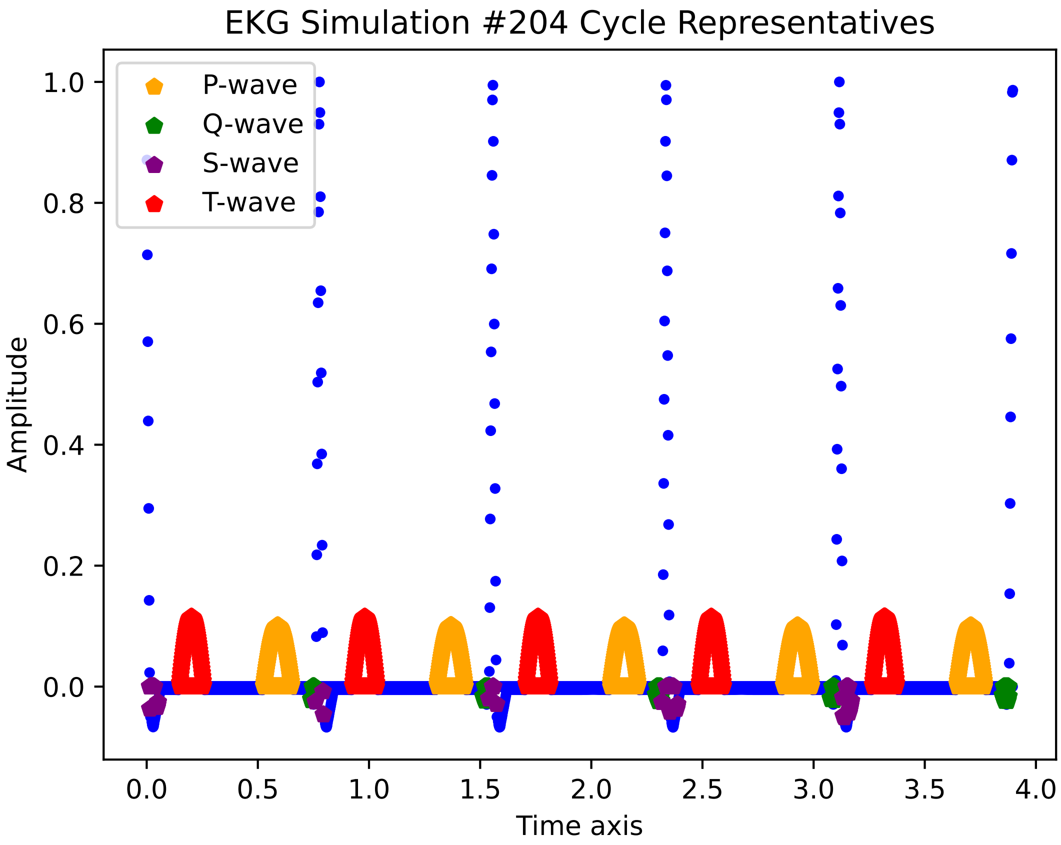
EKG Simulation #204 Cycle Representatives
P-wave
Q-wave
S-wave
T-wave
Amplitude
Time axis
0.0
0.2
0.4
0.6
0.8
1.0
0.0
0.5
1.0
1.5
2.0
2.5
3.0
3.5
4.0

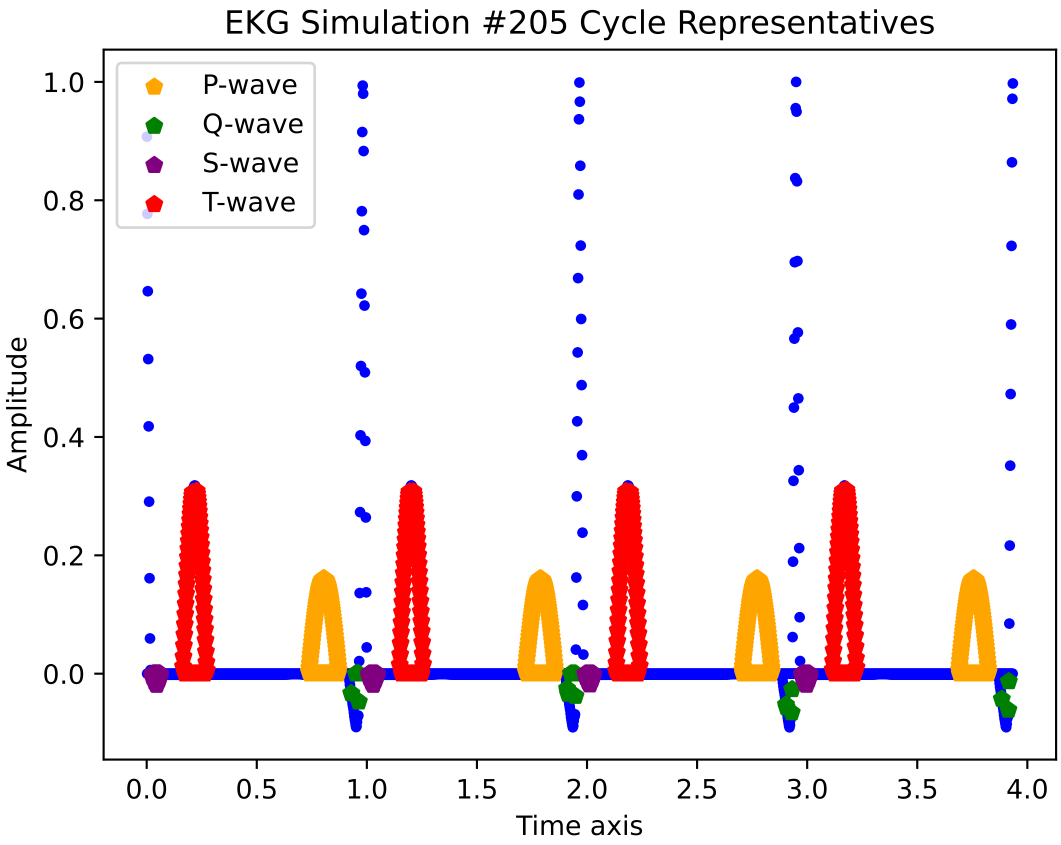
EKG Simulation #205 Cycle Representatives
P-wave
Q-wave
S-wave
T-wave
1.0
0.8
0.6
0.4
0.2
0.0
Amplitude
0.0
0.5
1.0
1.5
2.0
2.5
3.0
3.5
4.0
Time axis

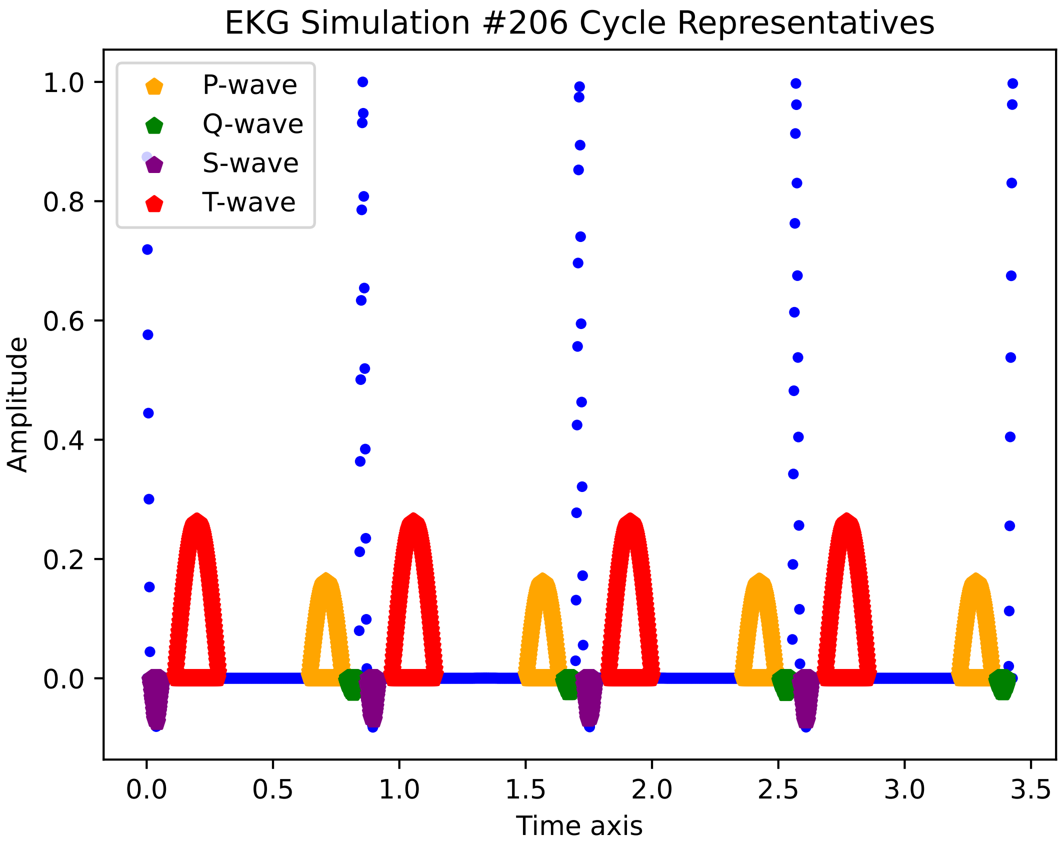
EKG Simulation #206 Cycle Representatives
P-wave
Q-wave
S-wave
T-wave
Amplitude
1.0
0.8
0.6
0.4
0.2
0.0
0.0
0.5
1.0
1.5
2.0
2.5
3.0
3.5
Time axis

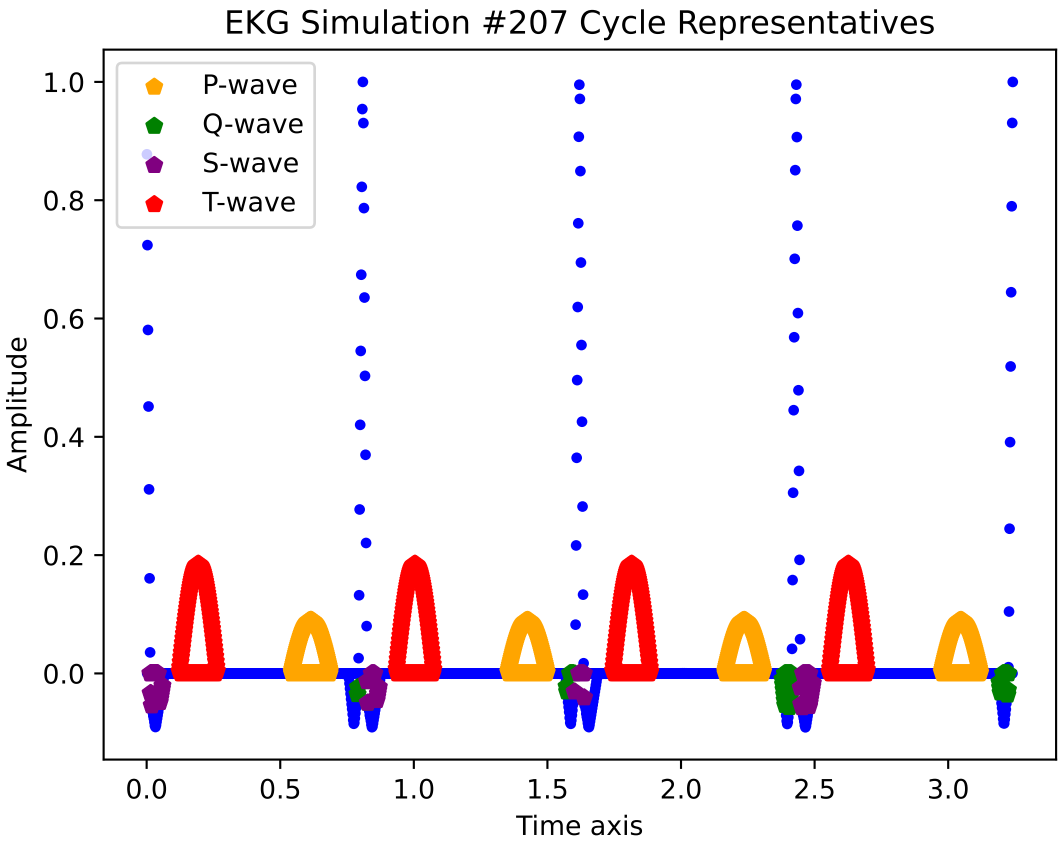
EKG Simulation #207 Cycle Representatives
P-wave
Q-wave
S-wave
T-wave
Amplitude
1.0
0.8
0.6
0.4
0.2
0.0
0.0
0.5
1.0
1.5
2.0
2.5
3.0
Time axis

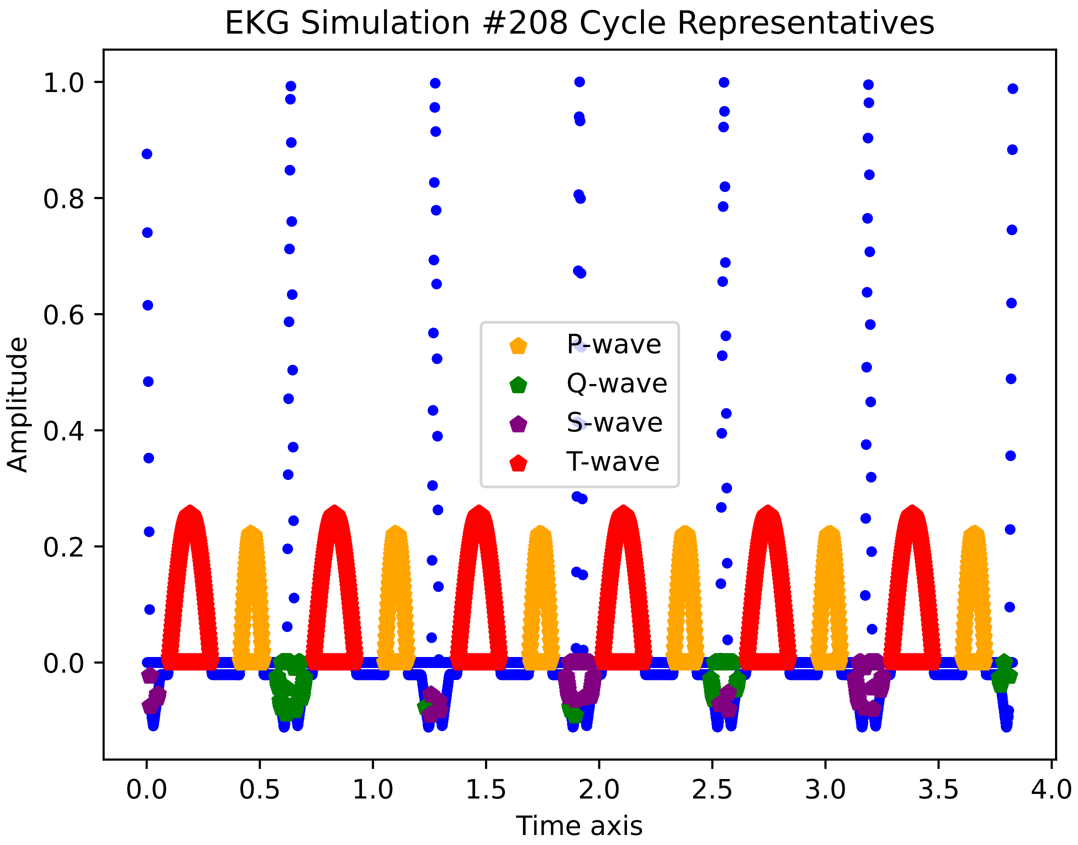

EKG Simulation #208 Cycle Representatives
Amplitude
Time axis
1.0
0.8
0.6
0.4
0.2
0.0
0.0
0.5
1.0
1.5
2.0
2.5
3.0
3.5
4.0
P-wave
Q-wave
S-wave
T-wave

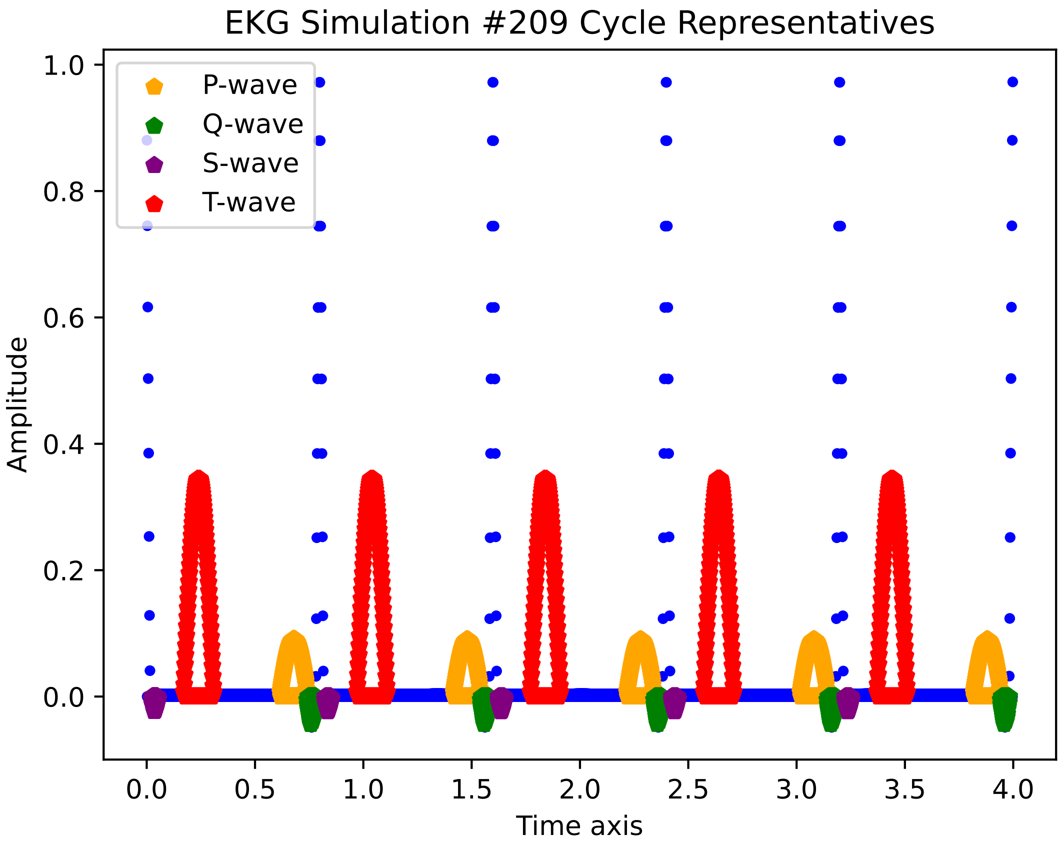
EKG Simulation #209 Cycle Representatives
P-wave
Q-wave
S-wave
T-wave
Amplitude
Time axis
1.0
0.8
0.6
0.4
0.2
0.0
0.0
0.5
1.0
1.5
2.0
2.5
3.0
3.5
4.0

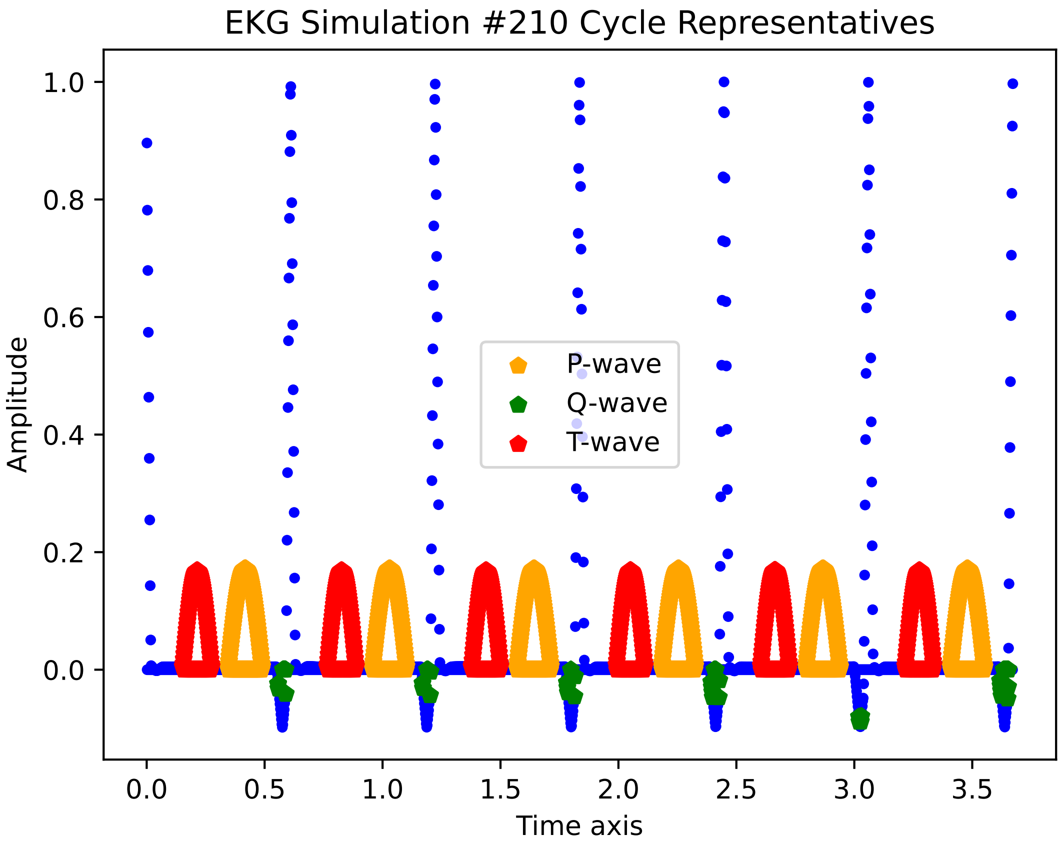

EKG Simulation #210 Cycle Representatives
Amplitude
Time axis
P-wave
Q-wave
T-wave
1.0
0.8
0.6
0.4
0.2
0.0
0.0
0.5
1.0
1.5
2.0
2.5
3.0
3.5

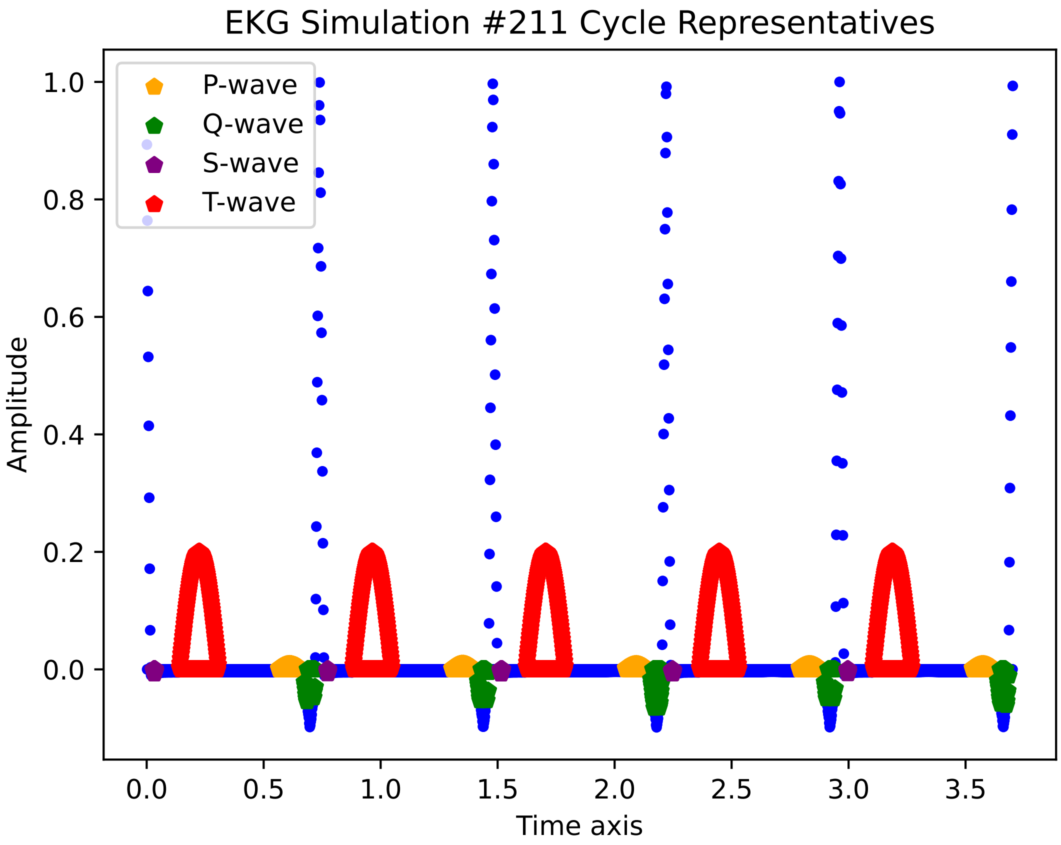
EKG Simulation #211 Cycle Representatives
P-wave
Q-wave
S-wave
T-wave
1.0
0.8
0.6
0.4
0.2
0.0
Amplitude
0.0
0.5
1.0
1.5
2.0
2.5
3.0
3.5
Time axis

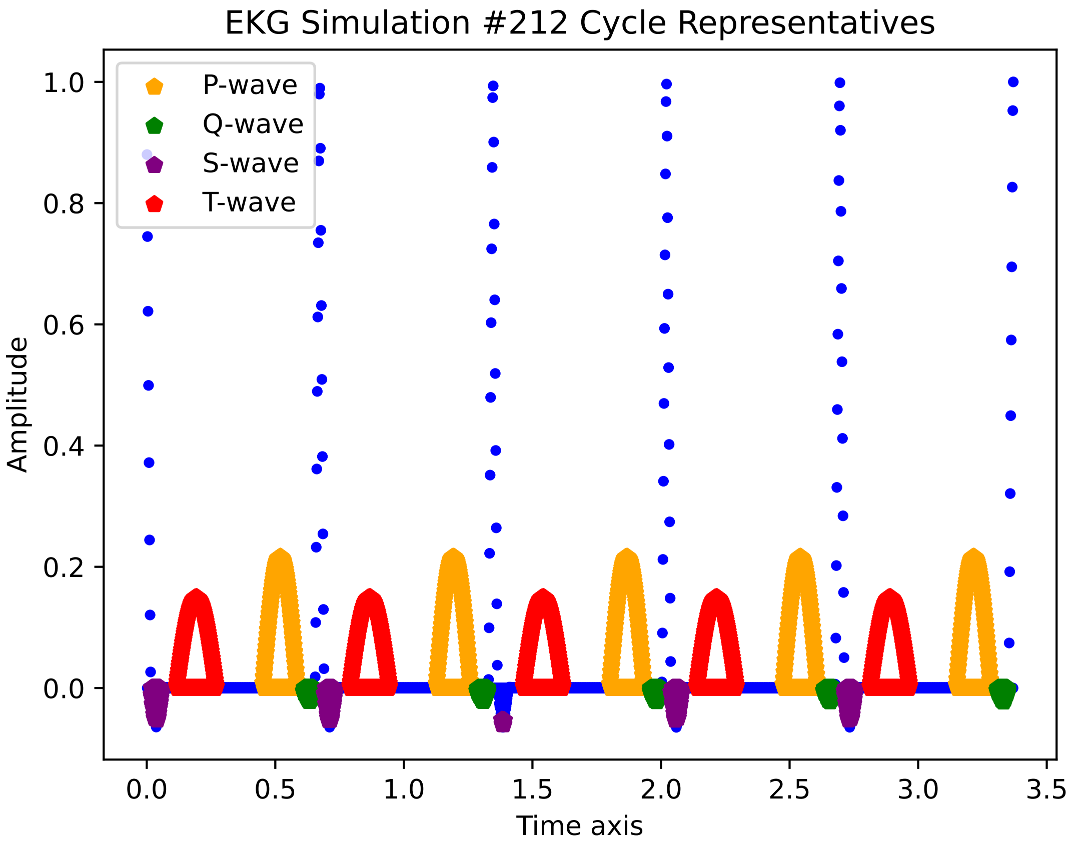

EKG Simulation #212 Cycle Representatives
P-wave
Q-wave
S-wave
T-wave
1.0
0.8
0.6
0.4
0.2
0.0
Amplitude
0.0
0.5
1.0
1.5
2.0
2.5
3.0
3.5
Time axis

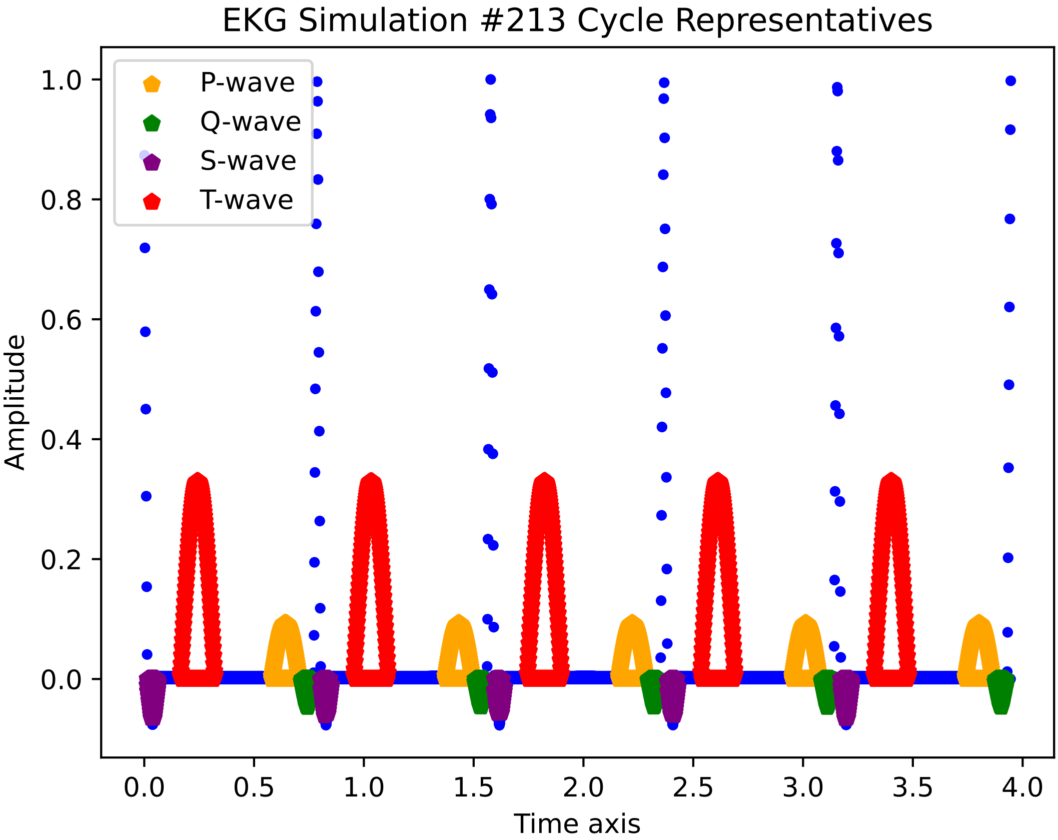

EKG Simulation #213 Cycle Representatives
P-wave
Q-wave
S-wave
T-wave
Amplitude
Time axis
1.0
0.8
0.6
0.4
0.2
0.0
0.0
0.5
1.0
1.5
2.0
2.5
3.0
3.5
4.0

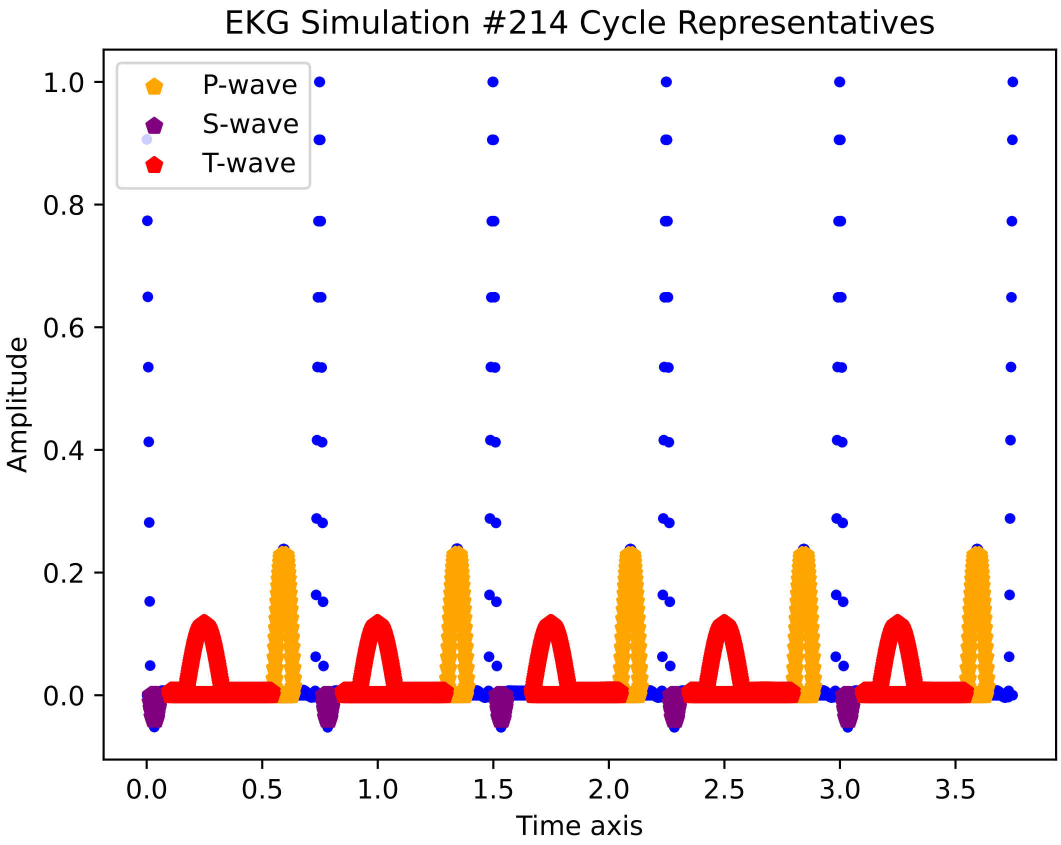
EKG Simulation #214 Cycle Representatives
P-wave
S-wave
T-wave
Amplitude
1.0
0.8
0.6
0.4
0.2
0.0
0.0
0.5
1.0
1.5
2.0
2.5
3.0
3.5
Time axis

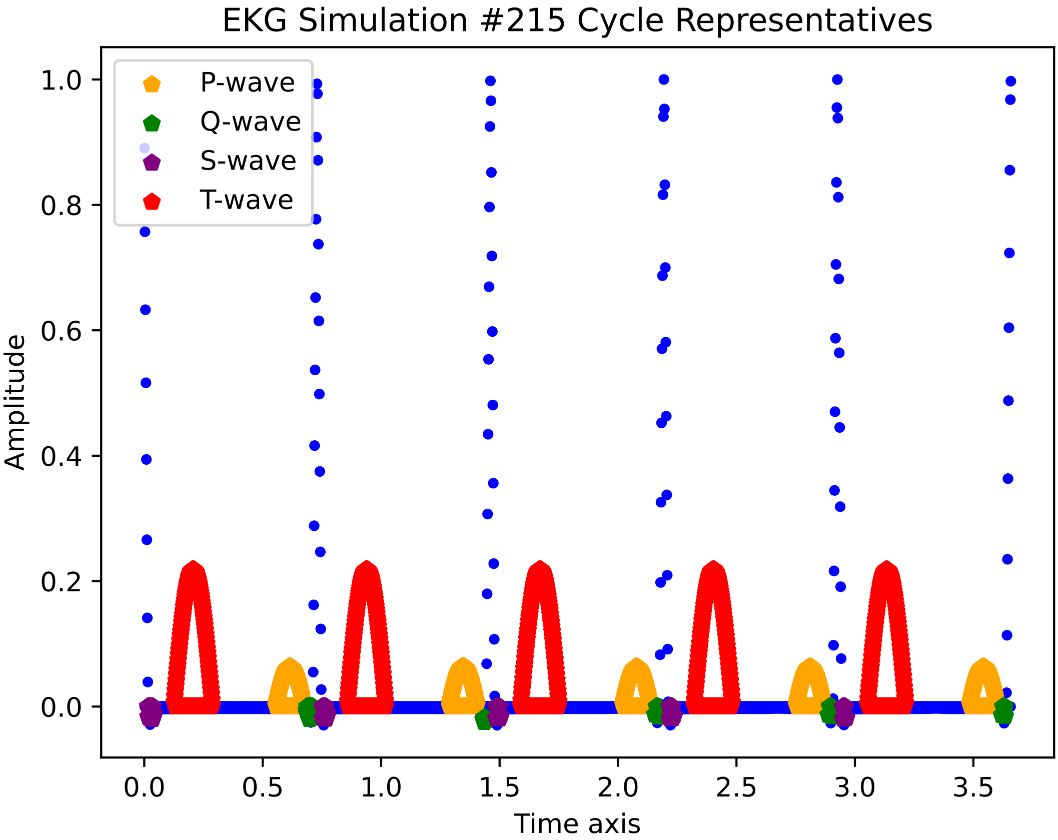

EKG Simulation #215 Cycle Representatives
P-wave
Q-wave
S-wave
T-wave
Amplitude
Time axis
1.0
0.8
0.6
0.4
0.2
0.0
0.0
0.5
1.0
1.5
2.0
2.5
3.0
3.5

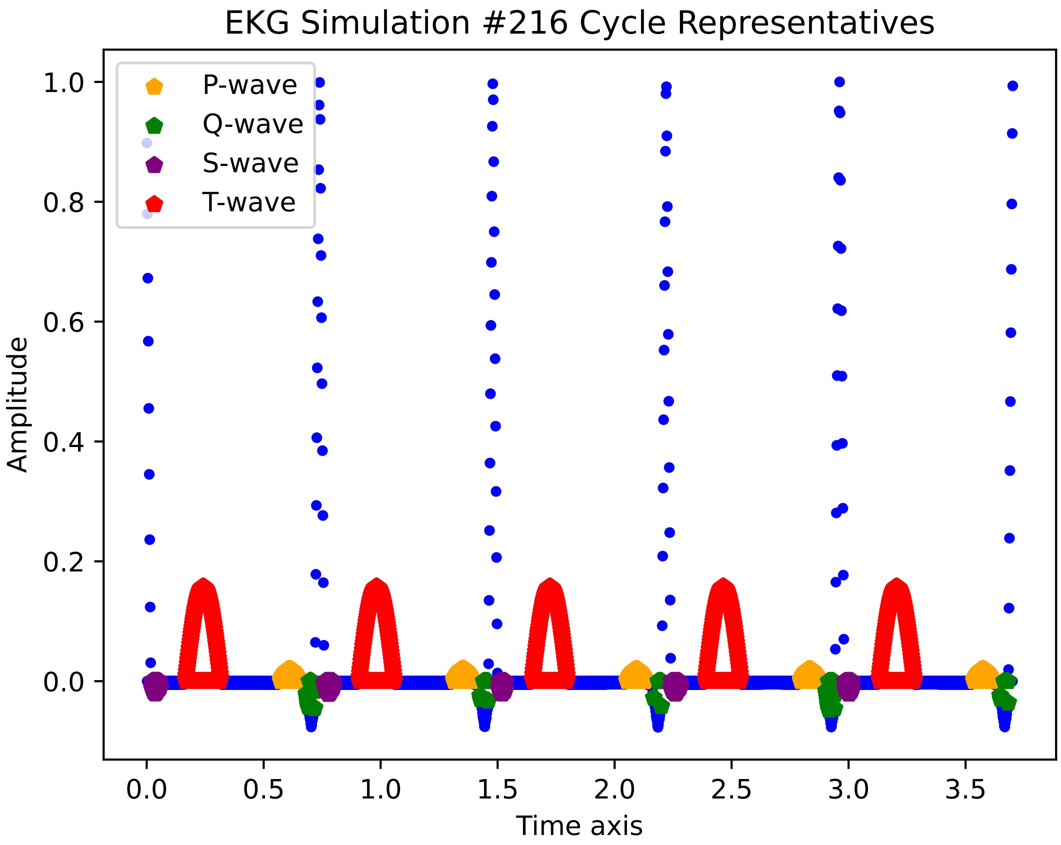

EKG Simulation #216 Cycle Representatives
P-wave
Q-wave
S-wave
T-wave
Amplitude
Time axis
1.0
0.8
0.6
0.4
0.2
0.0
0.0
0.5
1.0
1.5
2.0
2.5
3.0
3.5

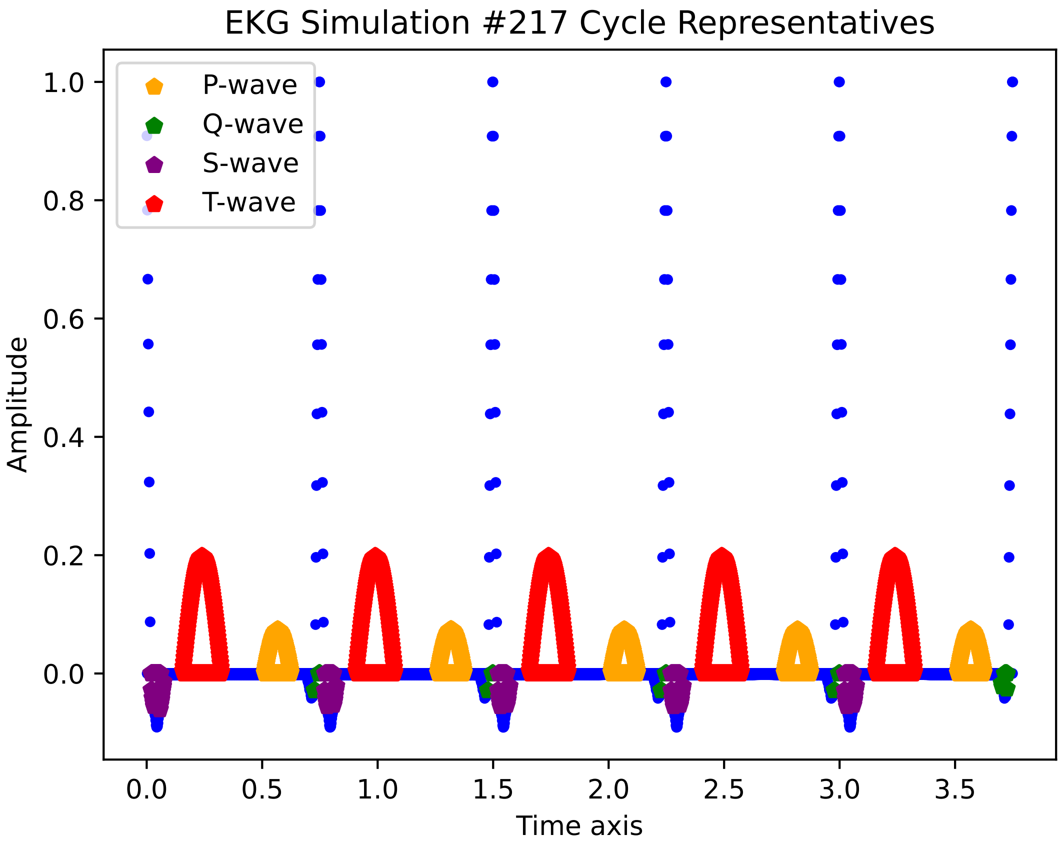
EKG Simulation #217 Cycle Representatives
P-wave
Q-wave
S-wave
T-wave
1.0
0.8
0.6
0.4
0.2
0.0
Amplitude
0.0
0.5
1.0
1.5
2.0
2.5
3.0
3.5
Time axis

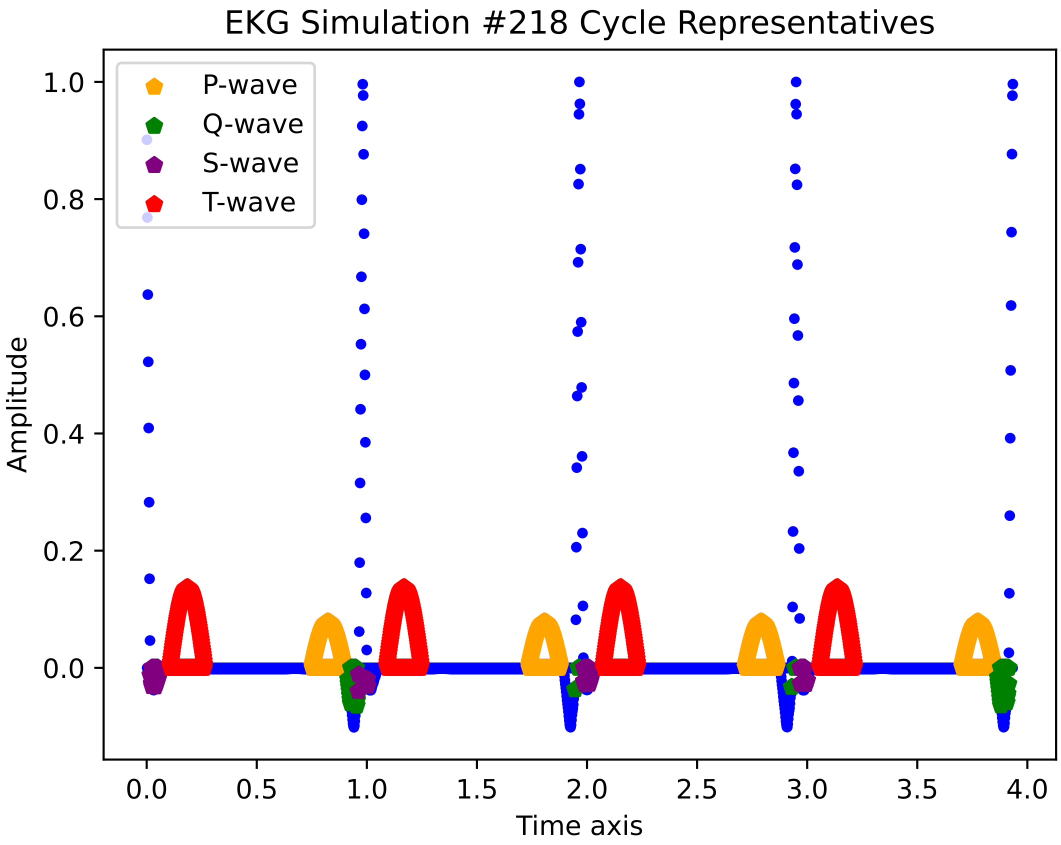

EKG Simulation #218 Cycle Representatives
P-wave
Q-wave
S-wave
T-wave
Amplitude
Time axis
1.0
0.8
0.6
0.4
0.2
0.0
0.0
0.5
1.0
1.5
2.0
2.5
3.0
3.5
4.0

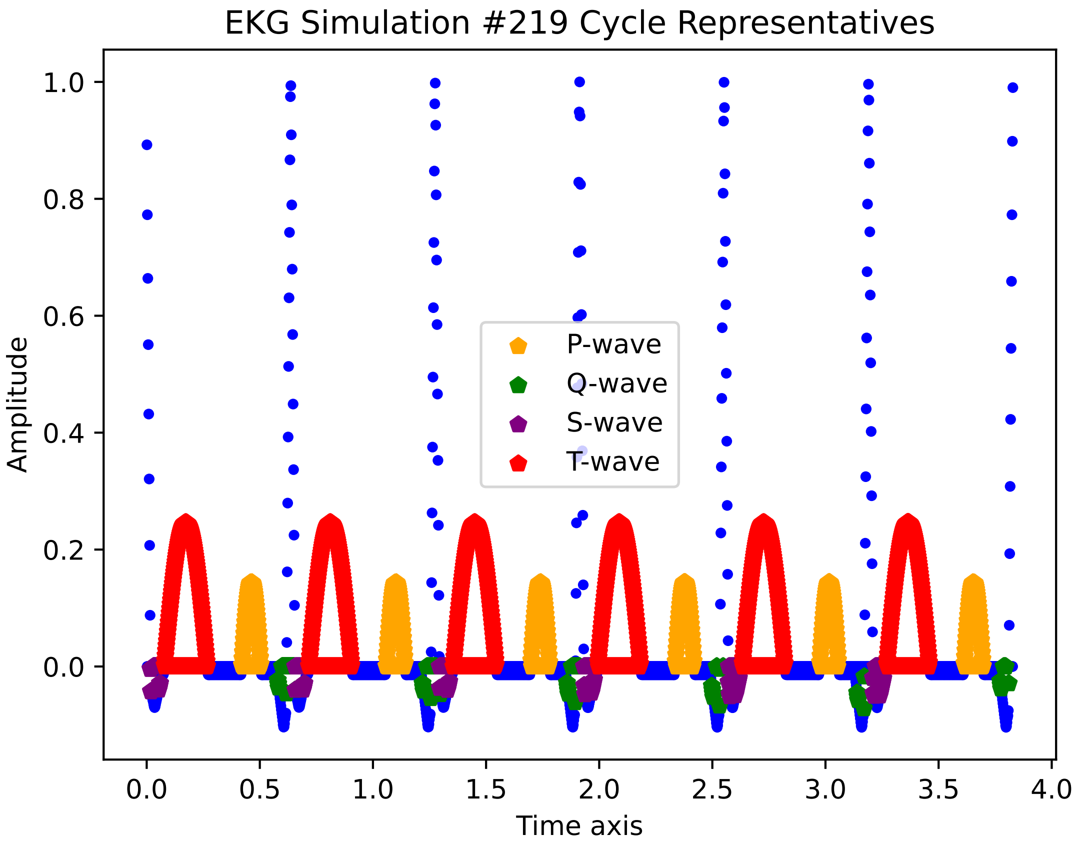

EKG Simulation #219 Cycle Representatives
Amplitude
Time axis
P-wave
Q-wave
S-wave
T-wave
1.0
0.8
0.6
0.4
0.2
0.0
0.0
0.5
1.0
1.5
2.0
2.5
3.0
3.5
4.0

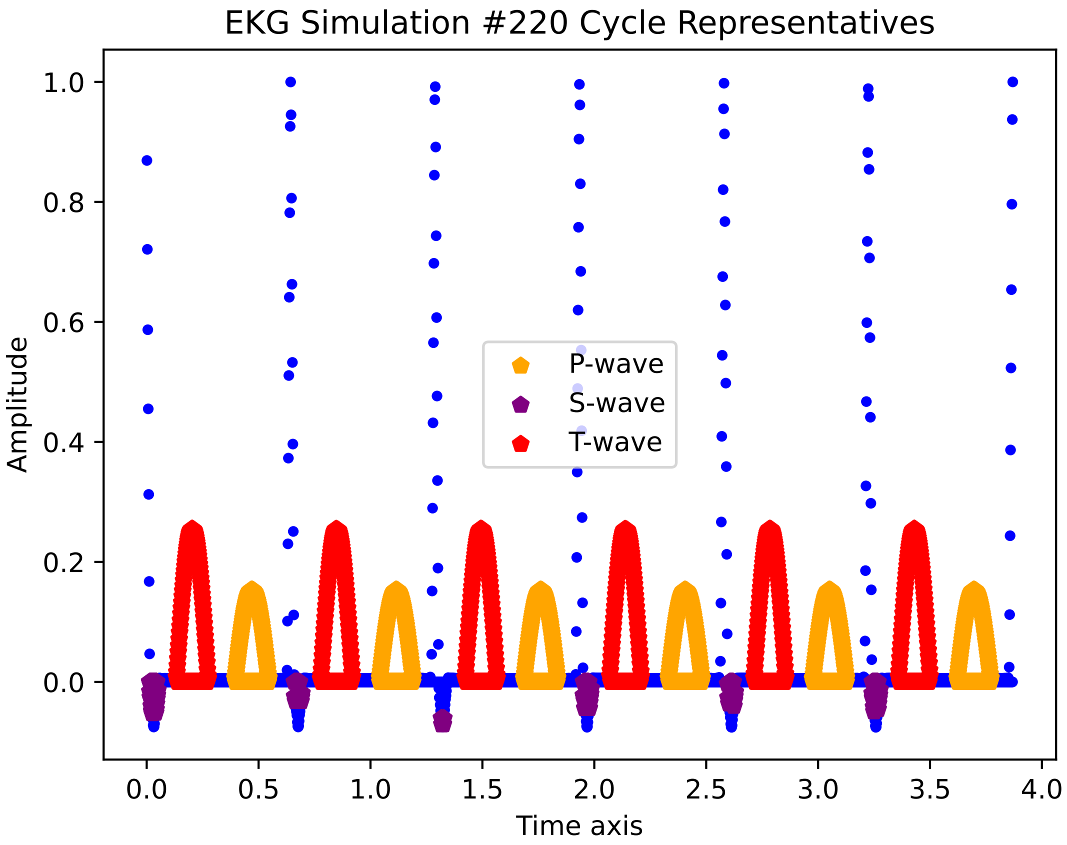
EKG Simulation #220 Cycle Representatives
P-wave
S-wave
T-wave
Amplitude
Time axis
1.0
0.8
0.6
0.4
0.2
0.0
0.0
0.5
1.0
1.5
2.0
2.5
3.0
3.5
4.0

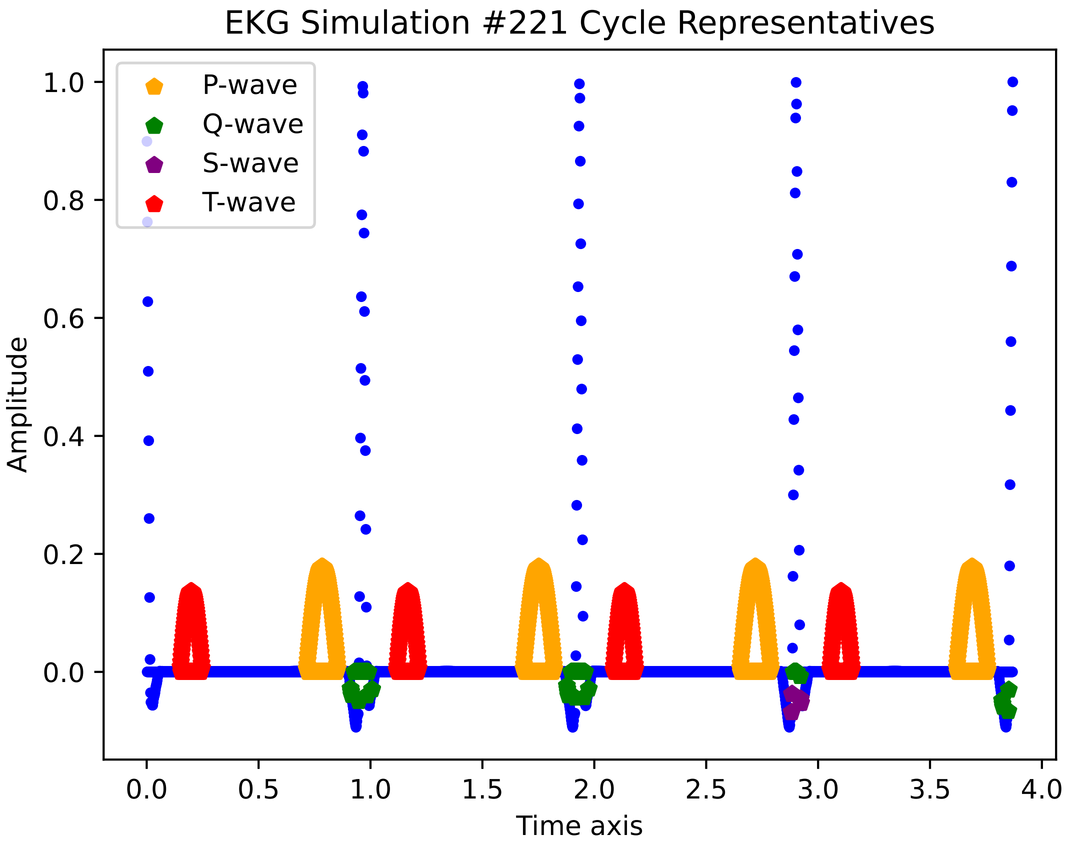

EKG Simulation #221 Cycle Representatives
P-wave
Q-wave
S-wave
T-wave
Amplitude
Time axis
1.0
0.8
0.6
0.4
0.2
0.0
0.0
0.5
1.0
1.5
2.0
2.5
3.0
3.5
4.0

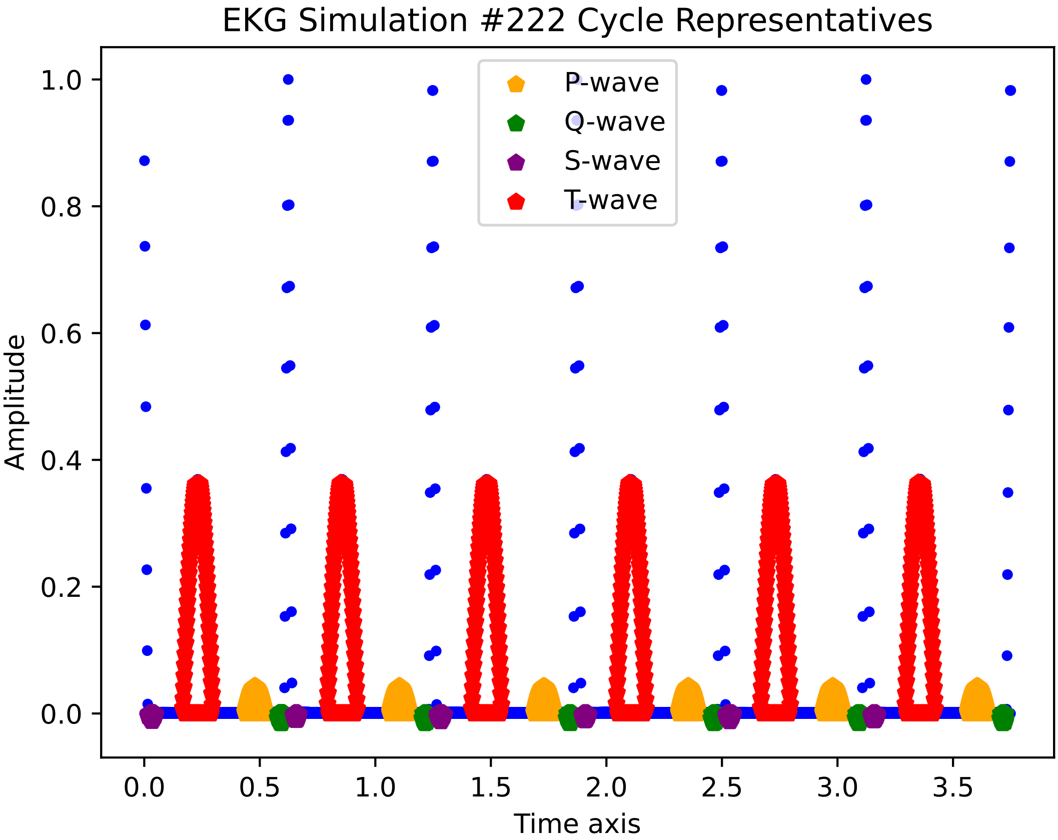
EKG Simulation #222 Cycle Representatives
P-wave
Q-wave
S-wave
T-wave
Amplitude
Time axis
0.0
0.2
0.4
0.6
0.8
1.0
0.0
0.5
1.0
1.5
2.0
2.5
3.0
3.5

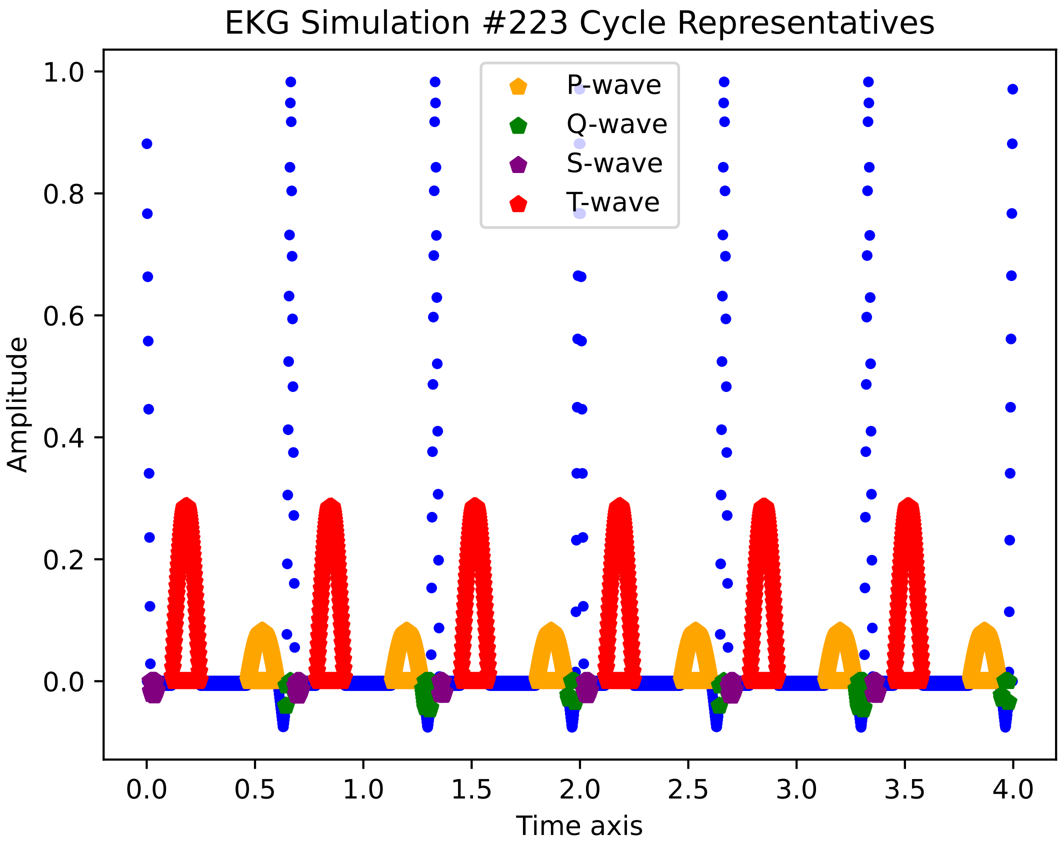
EKG Simulation #223 Cycle Representatives
P-wave
Q-wave
S-wave
T-wave
Amplitude
Time axis
1.0
0.8
0.6
0.4
0.2
0.0
0.0
0.5
1.0
1.5
2.0
2.5
3.0
3.5
4.0

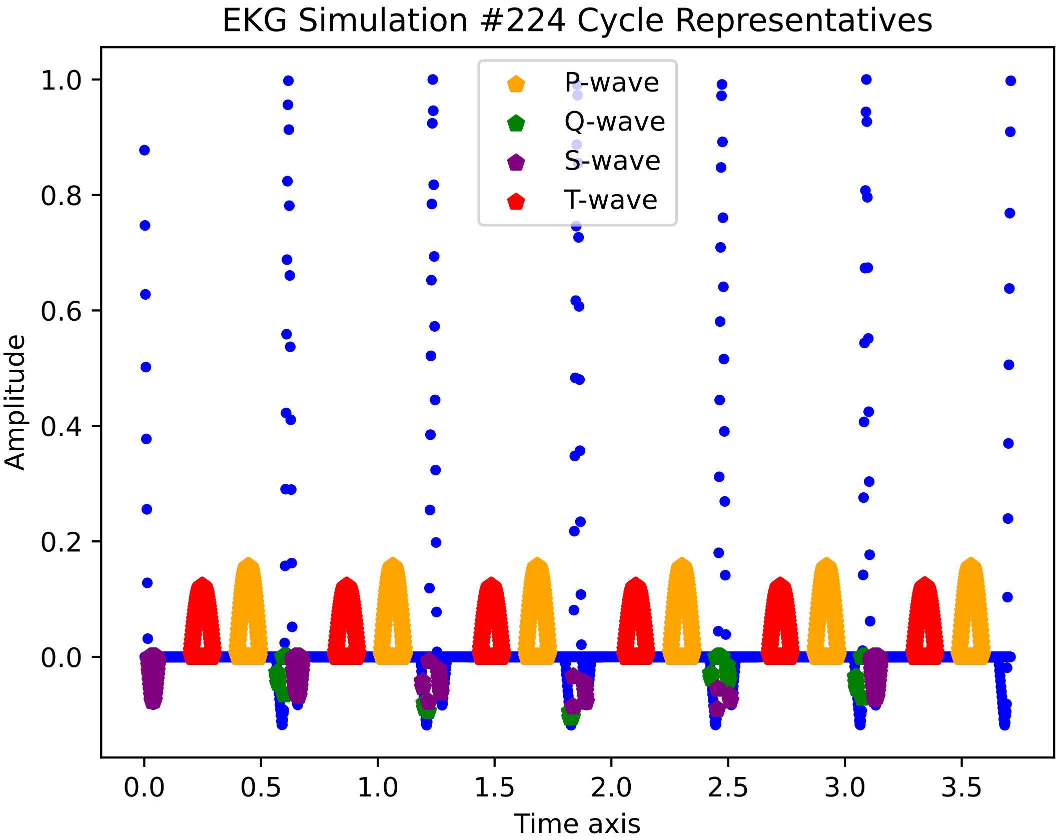

EKG Simulation #224 Cycle Representatives
P-wave
Q-wave
S-wave
T-wave
Amplitude
Time axis
1.0
0.8
0.6
0.4
0.2
0.0
0.0
0.5
1.0
1.5
2.0
2.5
3.0
3.5

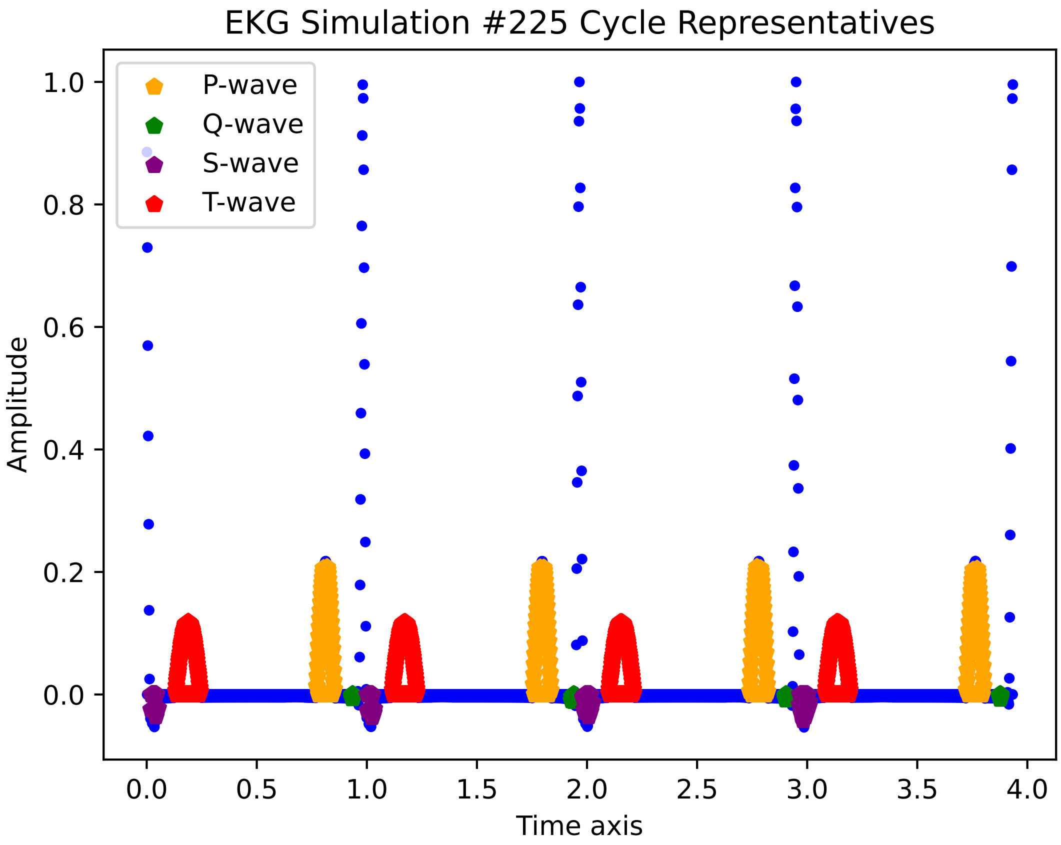

EKG Simulation #225 Cycle Representatives
P-wave
Q-wave
S-wave
T-wave
1.0
0.8
0.6
0.4
0.2
0.0
Amplitude
0.0
0.5
1.0
1.5
2.0
2.5
3.0
3.5
4.0
Time axis

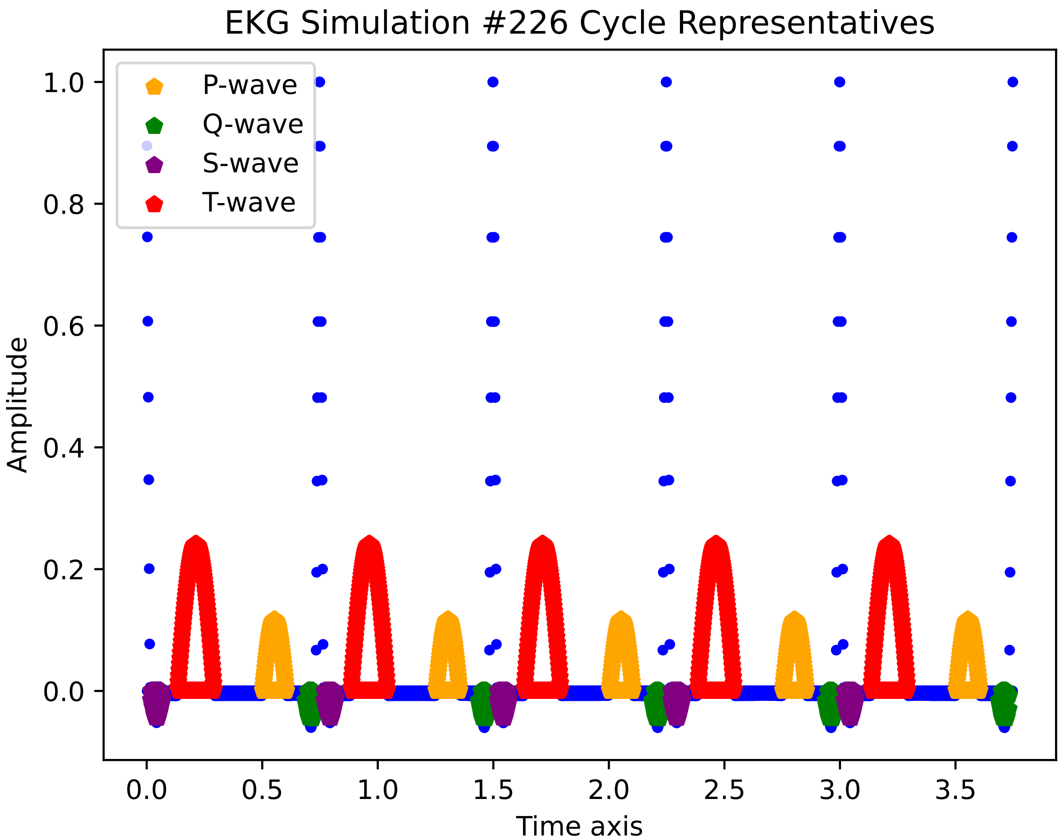
EKG Simulation #226 Cycle Representatives
P-wave
Q-wave
S-wave
T-wave
Amplitude
Time axis
1.0
0.8
0.6
0.4
0.2
0.0
0.0
0.5
1.0
1.5
2.0
2.5
3.0
3.5

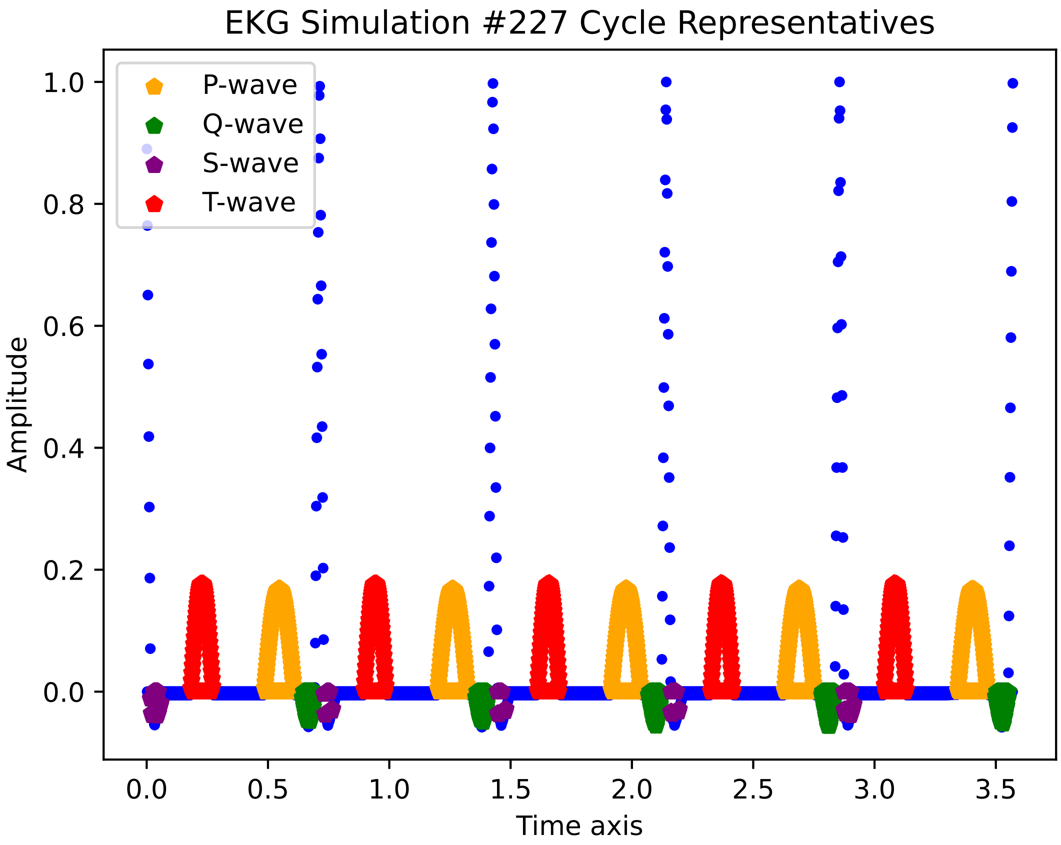
EKG Simulation #227 Cycle Representatives
P-wave
Q-wave
S-wave
T-wave
Amplitude
Time axis
1.0
0.8
0.6
0.4
0.2
0.0
0.0
0.5
1.0
1.5
2.0
2.5
3.0
3.5

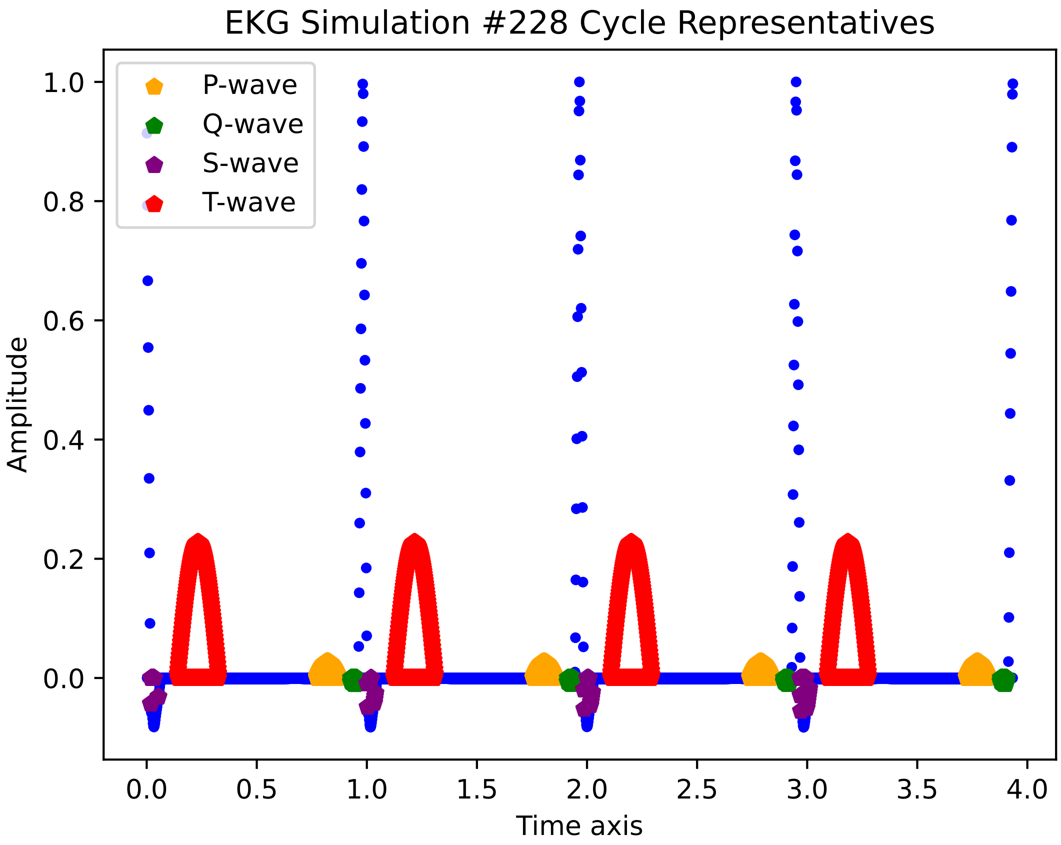

EKG Simulation #228 Cycle Representatives
P-wave
Q-wave
S-wave
T-wave
Amplitude
1.0
0.8
0.6
0.4
0.2
0.0
0.0
0.5
1.0
1.5
2.0
2.5
3.0
3.5
4.0
Time axis

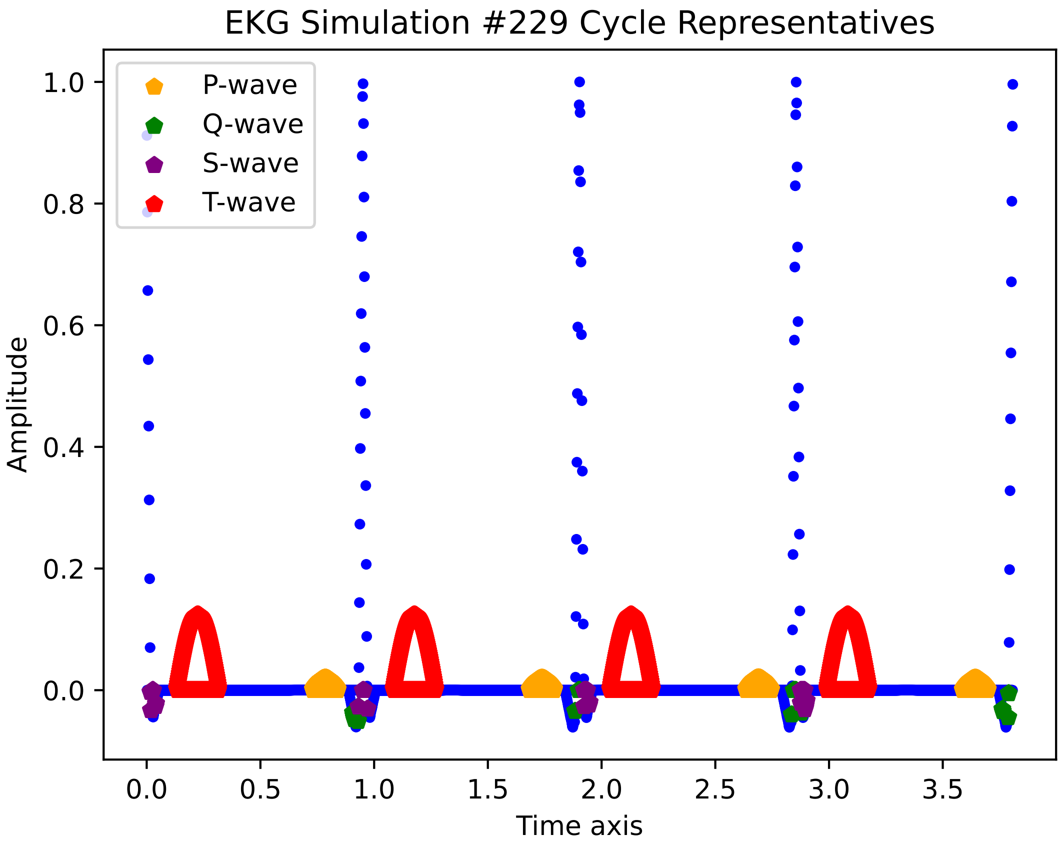
EKG Simulation #229 Cycle Representatives
P-wave
Q-wave
S-wave
T-wave
1.0
0.8
0.6
0.4
0.2
0.0
Amplitude
0.0
0.5
1.0
1.5
2.0
2.5
3.0
3.5
Time axis

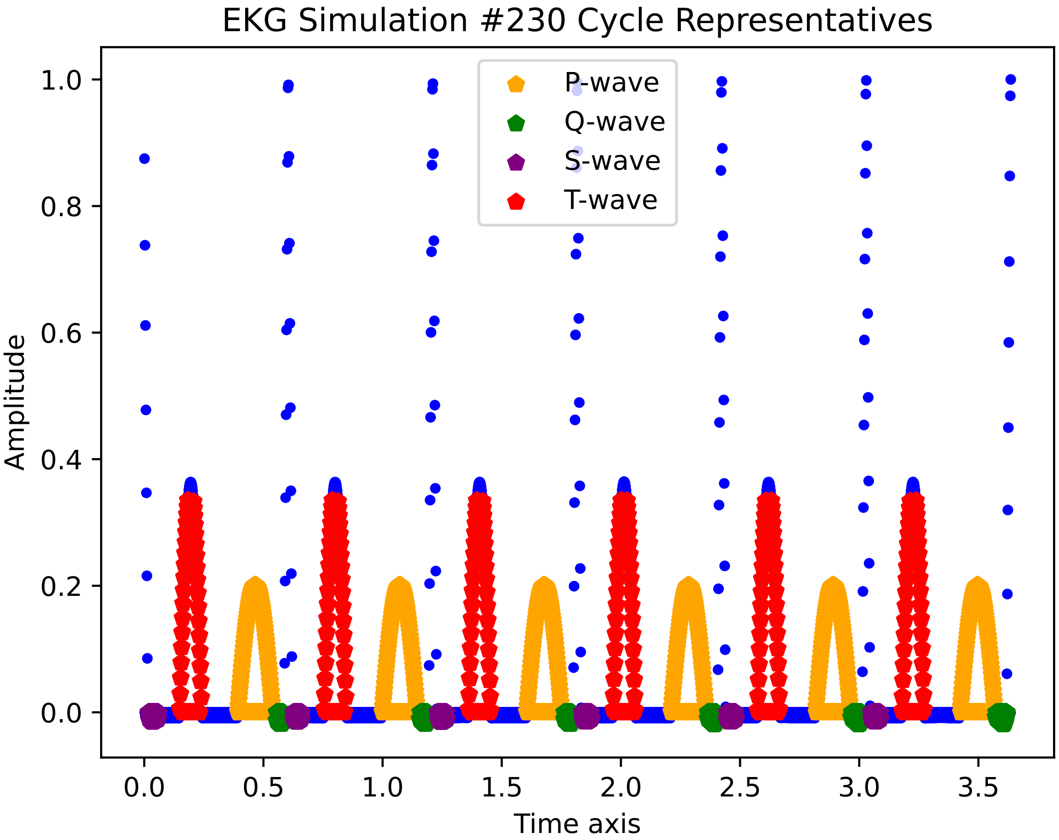

EKG Simulation #230 Cycle Representatives
P-wave
Q-wave
S-wave
T-wave
Amplitude
Time axis
1.0
0.8
0.6
0.4
0.2
0.0
0.0
0.5
1.0
1.5
2.0
2.5
3.0
3.5

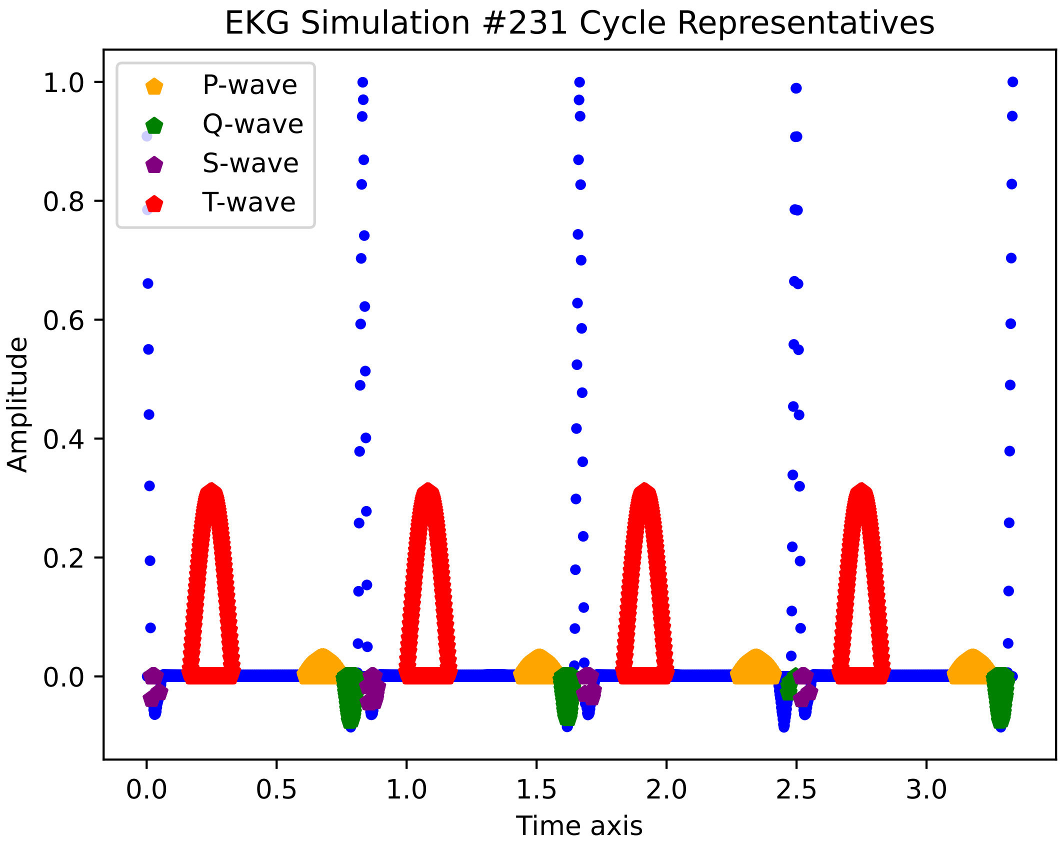

EKG Simulation #231 Cycle Representatives
P-wave
Q-wave
S-wave
T-wave
Amplitude
Time axis
1.0
0.8
0.6
0.4
0.2
0.0
0.0
0.5
1.0
1.5
2.0
2.5
3.0

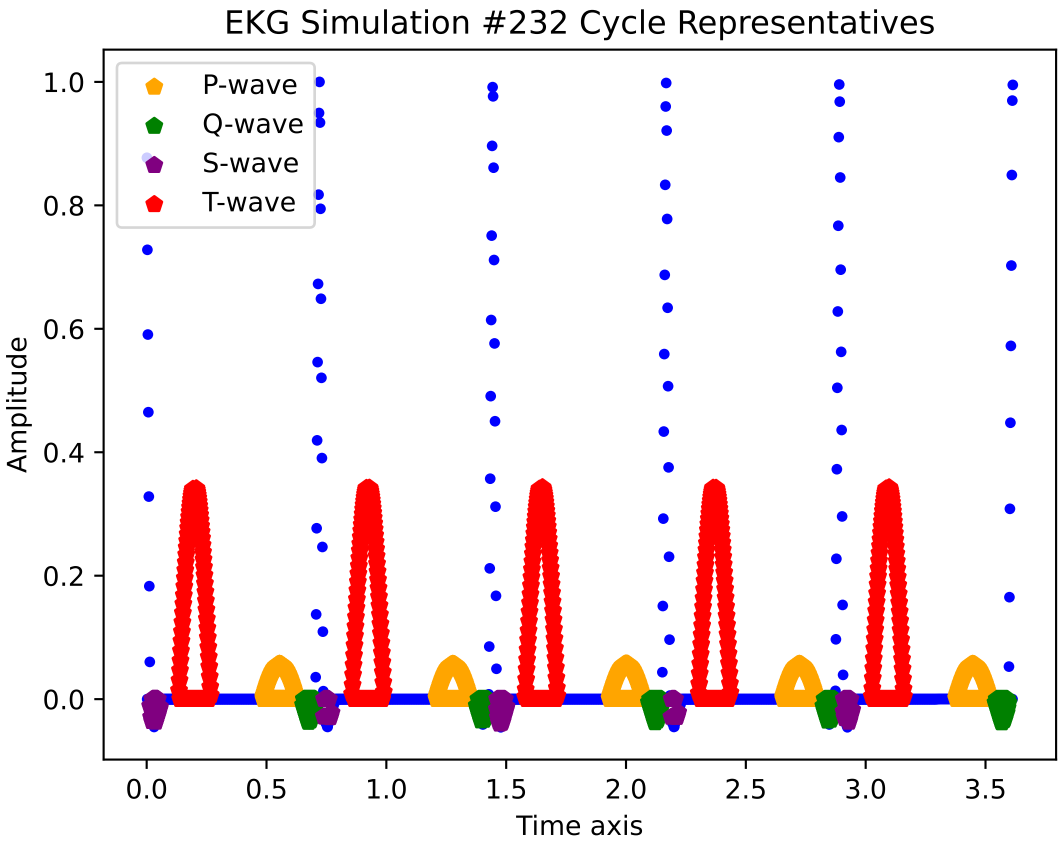
EKG Simulation #232 Cycle Representatives
P-wave
Q-wave
S-wave
T-wave
Amplitude
Time axis
1.0
0.8
0.6
0.4
0.2
0.0
0.0
0.5
1.0
1.5
2.0
2.5
3.0
3.5

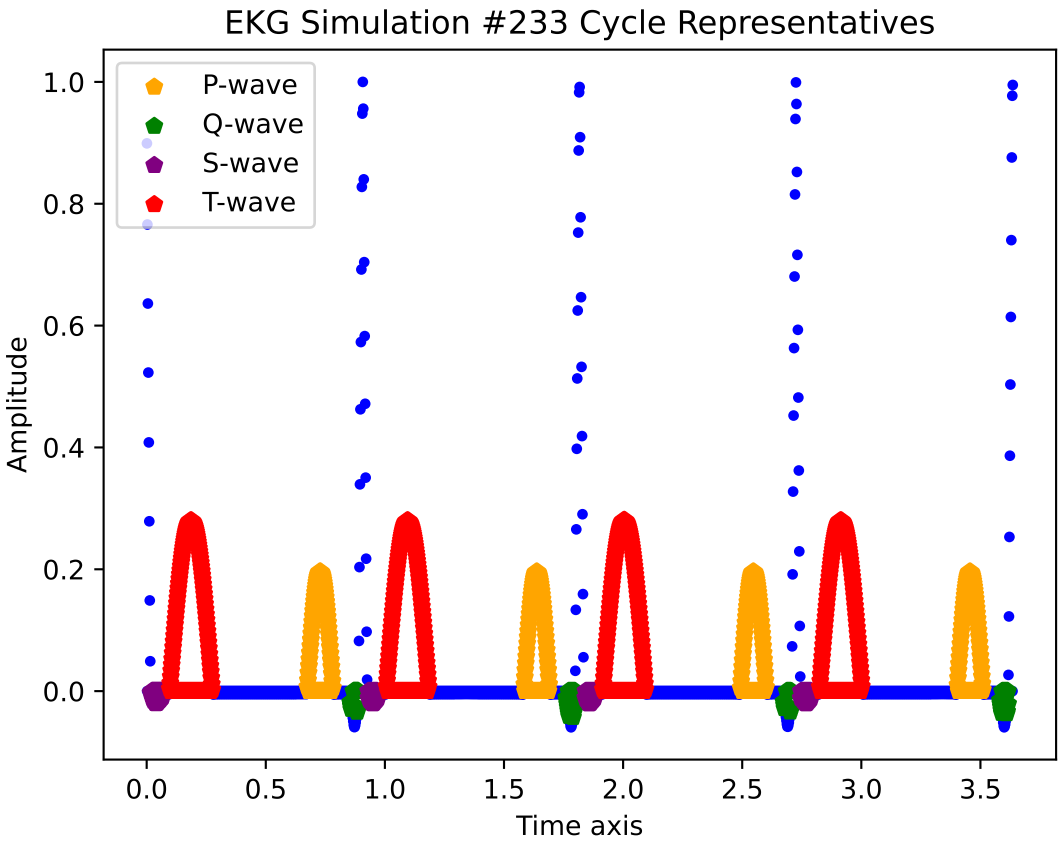
EKG Simulation #233 Cycle Representatives
P-wave
Q-wave
S-wave
T-wave
1.0
0.8
0.6
0.4
0.2
0.0
Amplitude
0.0
0.5
1.0
1.5
2.0
2.5
3.0
3.5
Time axis

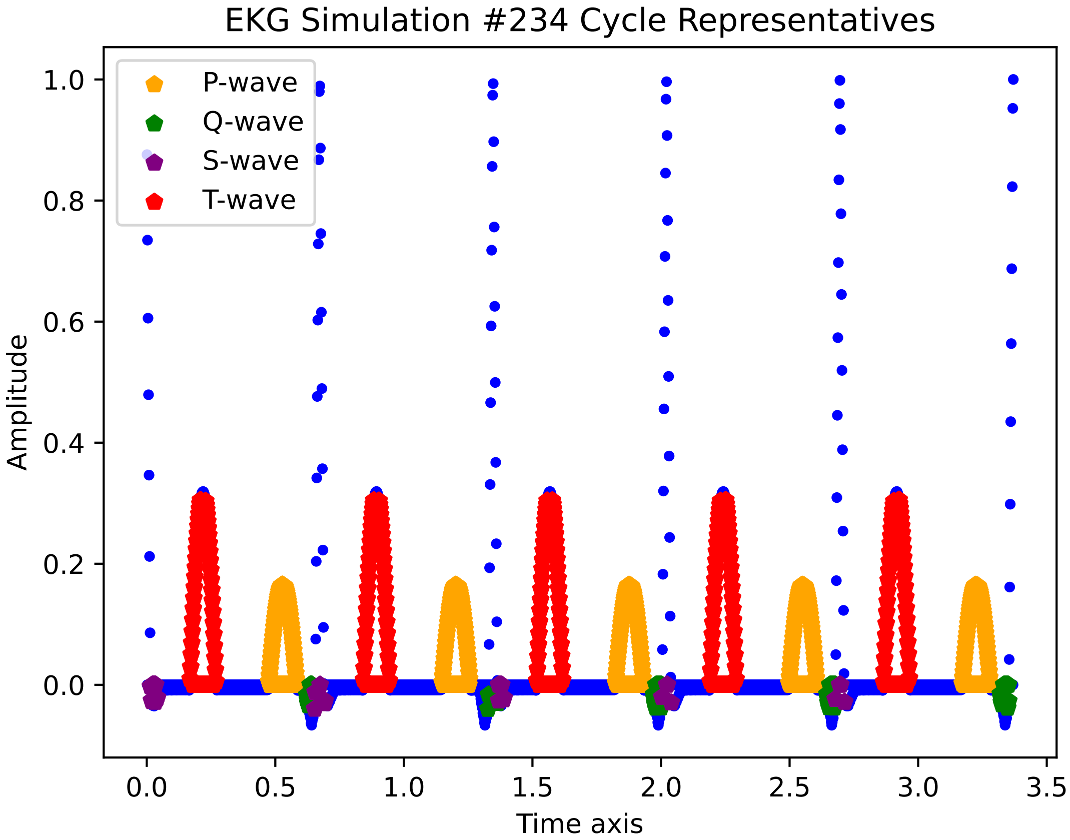
EKG Simulation #234 Cycle Representatives
P-wave
Q-wave
S-wave
T-wave
1.0
0.8
0.6
0.4
0.2
0.0
Amplitude
0.0
0.5
1.0
1.5
2.0
2.5
3.0
3.5
Time axis

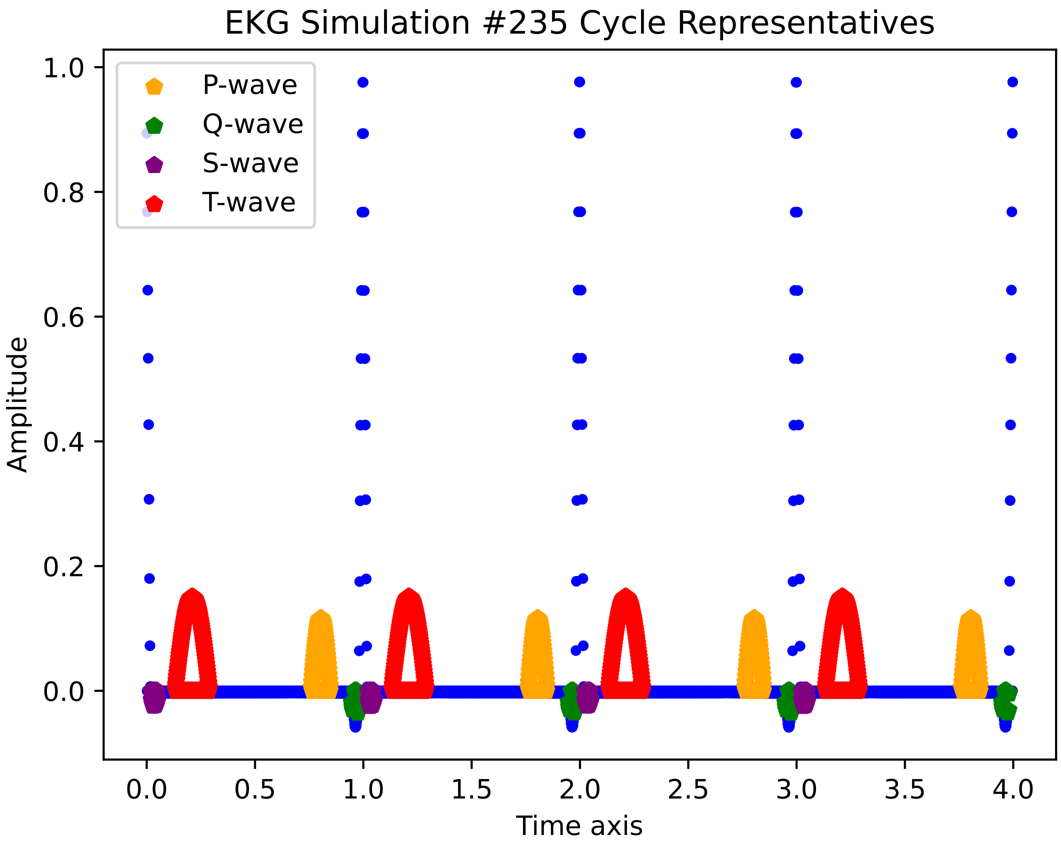
EKG Simulation #235 Cycle Representatives
P-wave
Q-wave
S-wave
T-wave
Amplitude
Time axis
1.0
0.8
0.6
0.4
0.2
0.0
0.0
0.5
1.0
1.5
2.0
2.5
3.0
3.5
4.0

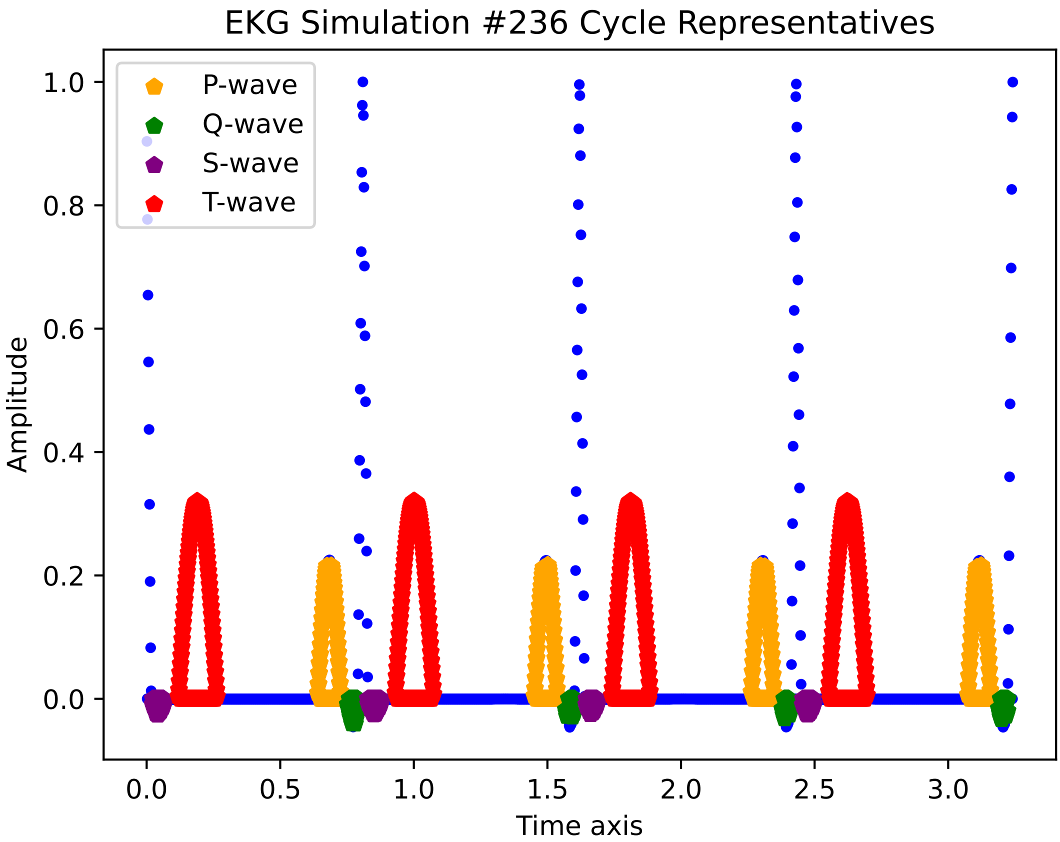
EKG Simulation #236 Cycle Representatives
P-wave
Q-wave
S-wave
T-wave
Amplitude
Time axis
1.0
0.8
0.6
0.4
0.2
0.0
0.0
0.5
1.0
1.5
2.0
2.5
3.0

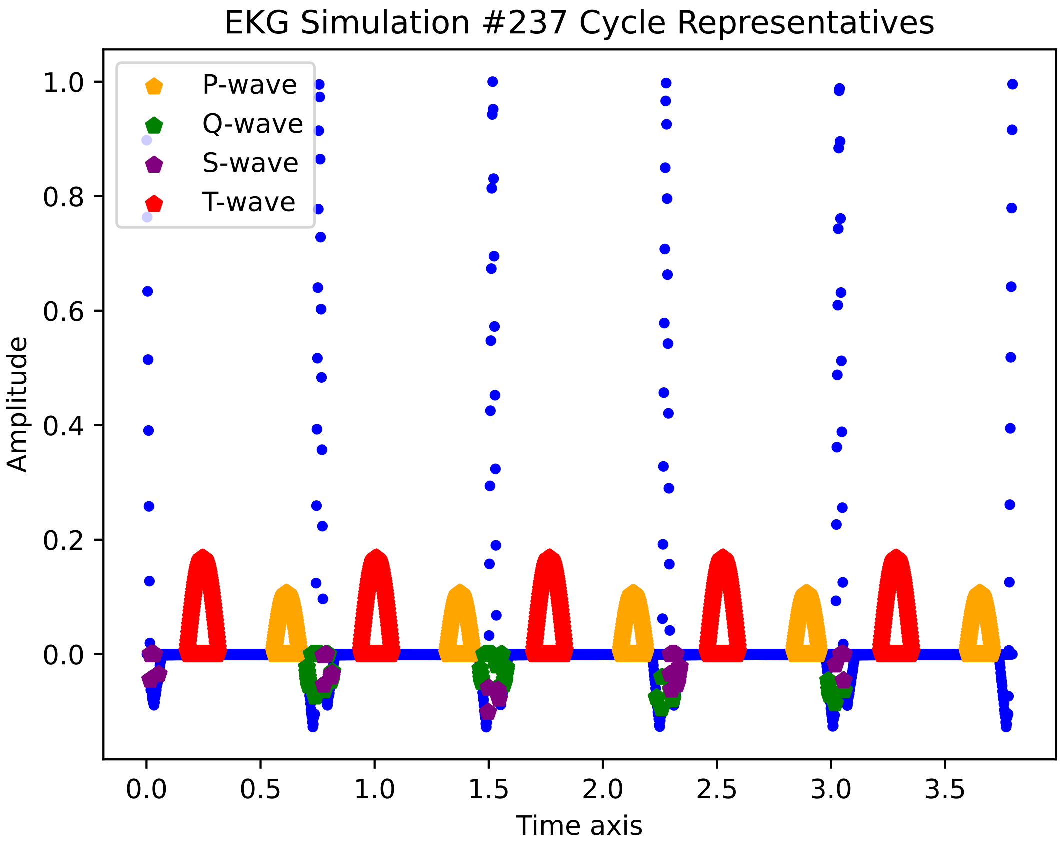

EKG Simulation #237 Cycle Representatives
P-wave
Q-wave
S-wave
T-wave
1.0
0.8
0.6
0.4
0.2
0.0
Amplitude
0.0
0.5
1.0
1.5
2.0
2.5
3.0
3.5
Time axis

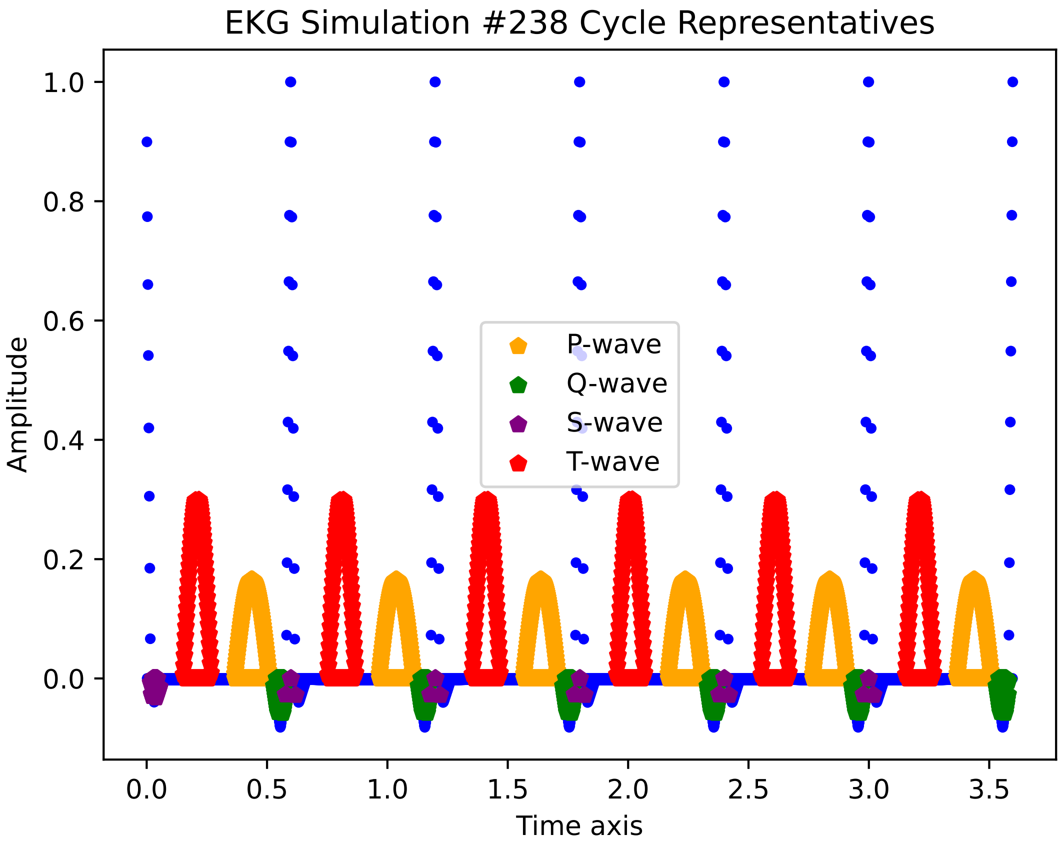

EKG Simulation #238 Cycle Representatives
Amplitude
1.0
0.8
0.6
0.4
0.2
0.0
0.0
0.5
1.0
1.5
2.0
2.5
3.0
3.5
Time axis
P-wave
Q-wave
S-wave
T-wave

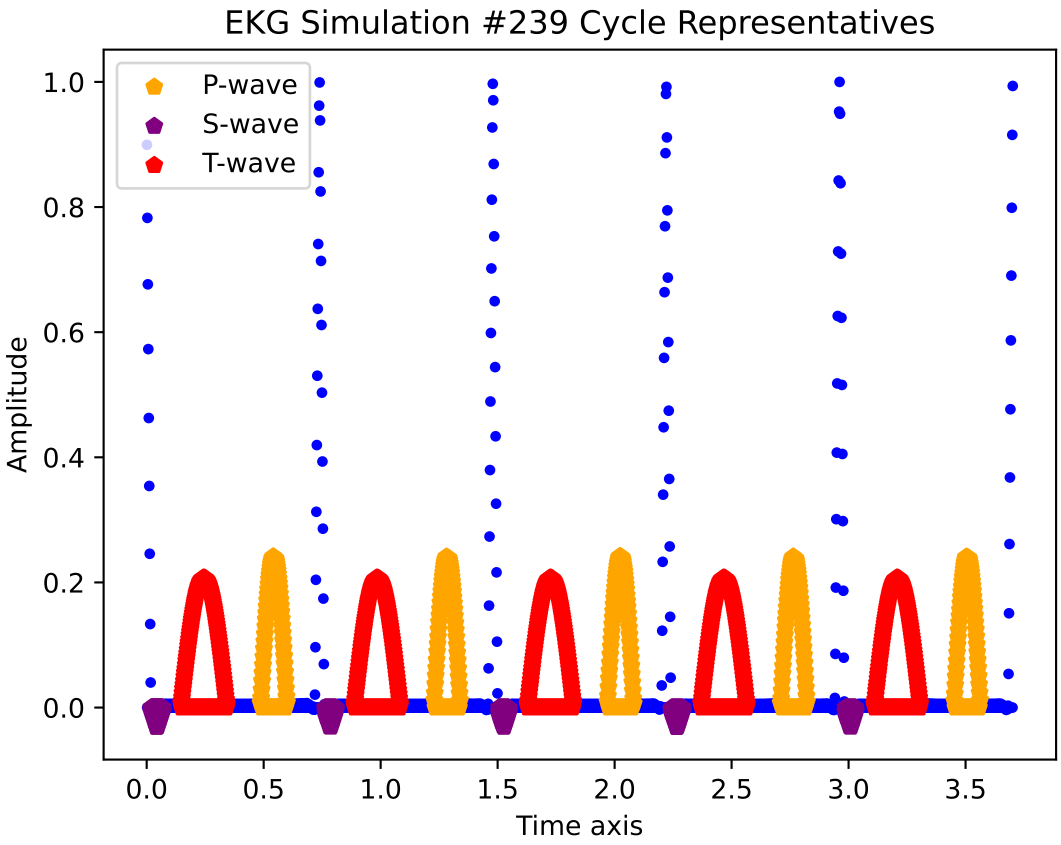
EKG Simulation #239 Cycle Representatives
P-wave
S-wave
T-wave
Amplitude
Time axis
1.0
0.8
0.6
0.4
0.2
0.0
0.0
0.5
1.0
1.5
2.0
2.5
3.0
3.5

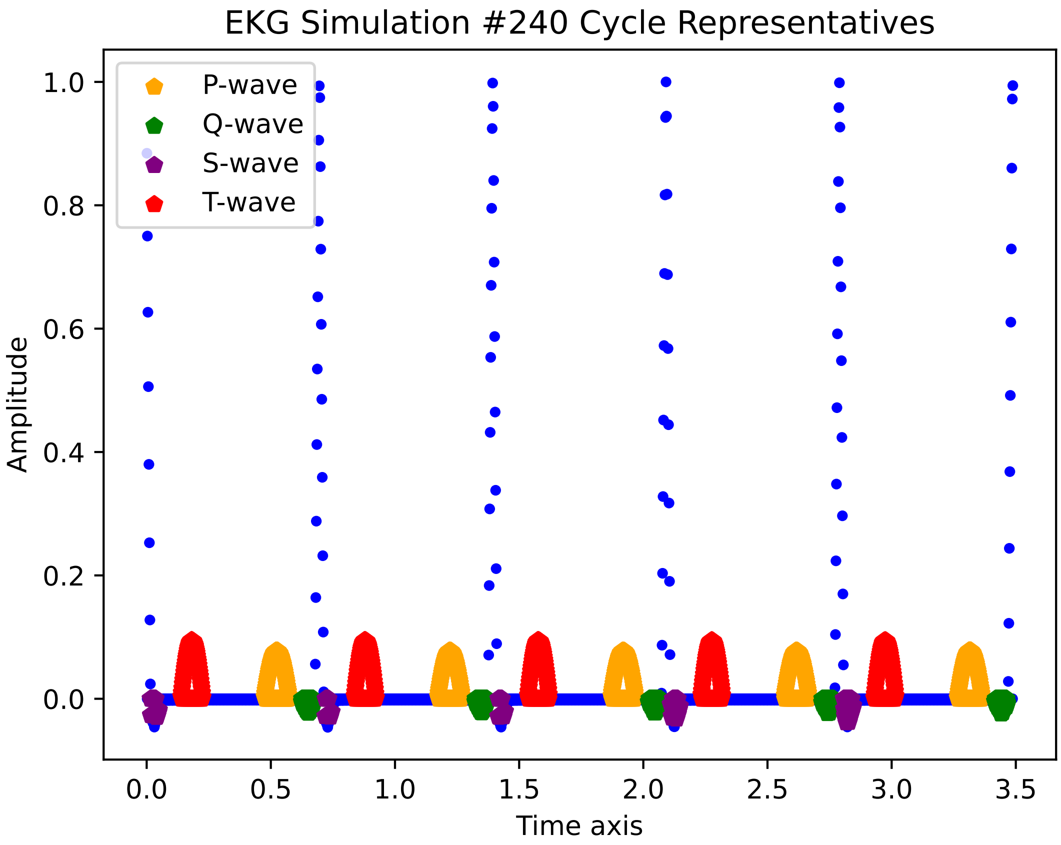

EKG Simulation #240 Cycle Representatives
P-wave
Q-wave
S-wave
T-wave
Amplitude
1.0
0.8
0.6
0.4
0.2
0.0
0.0
0.5
1.0
1.5
2.0
2.5
3.0
3.5
Time axis

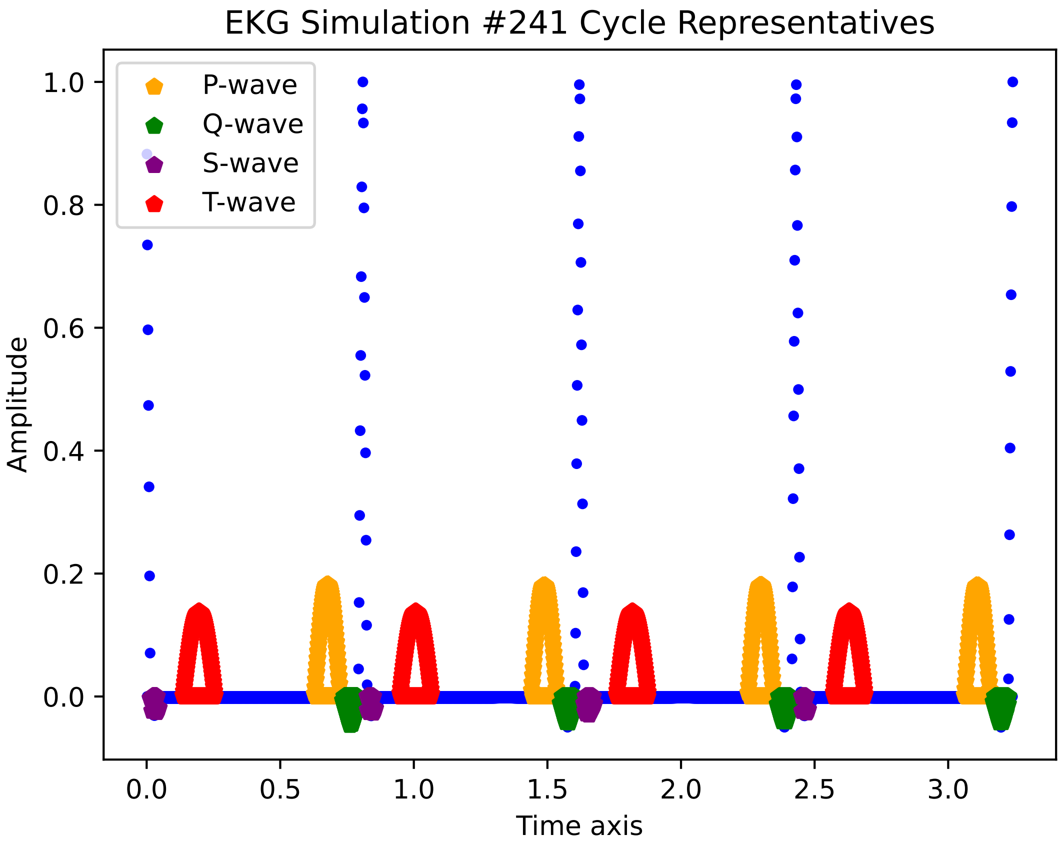
EKG Simulation #241 Cycle Representatives
P-wave
Q-wave
S-wave
T-wave
Amplitude
1.0
0.8
0.6
0.4
0.2
0.0
0.0
0.5
1.0
1.5
2.0
2.5
3.0
Time axis

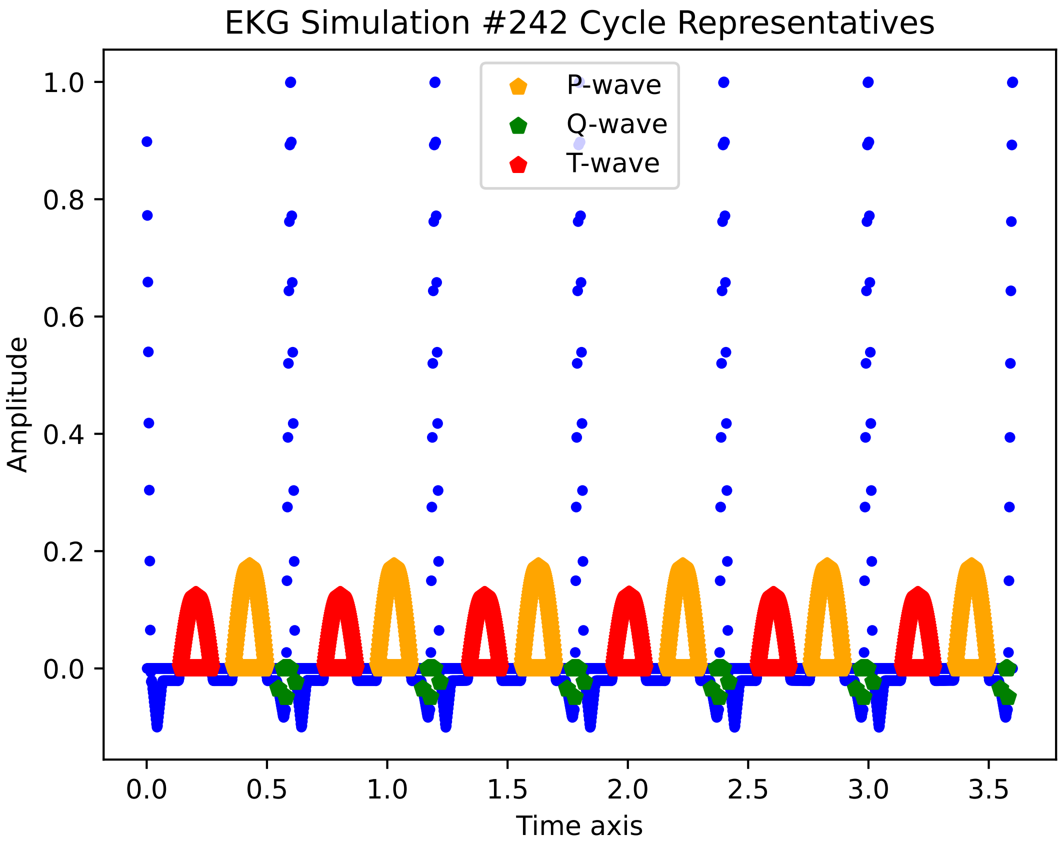

EKG Simulation #242 Cycle Representatives
P-wave
Q-wave
T-wave
Amplitude
Time axis
1.0
0.8
0.6
0.4
0.2
0.0
0.0
0.5
1.0
1.5
2.0
2.5
3.0
3.5

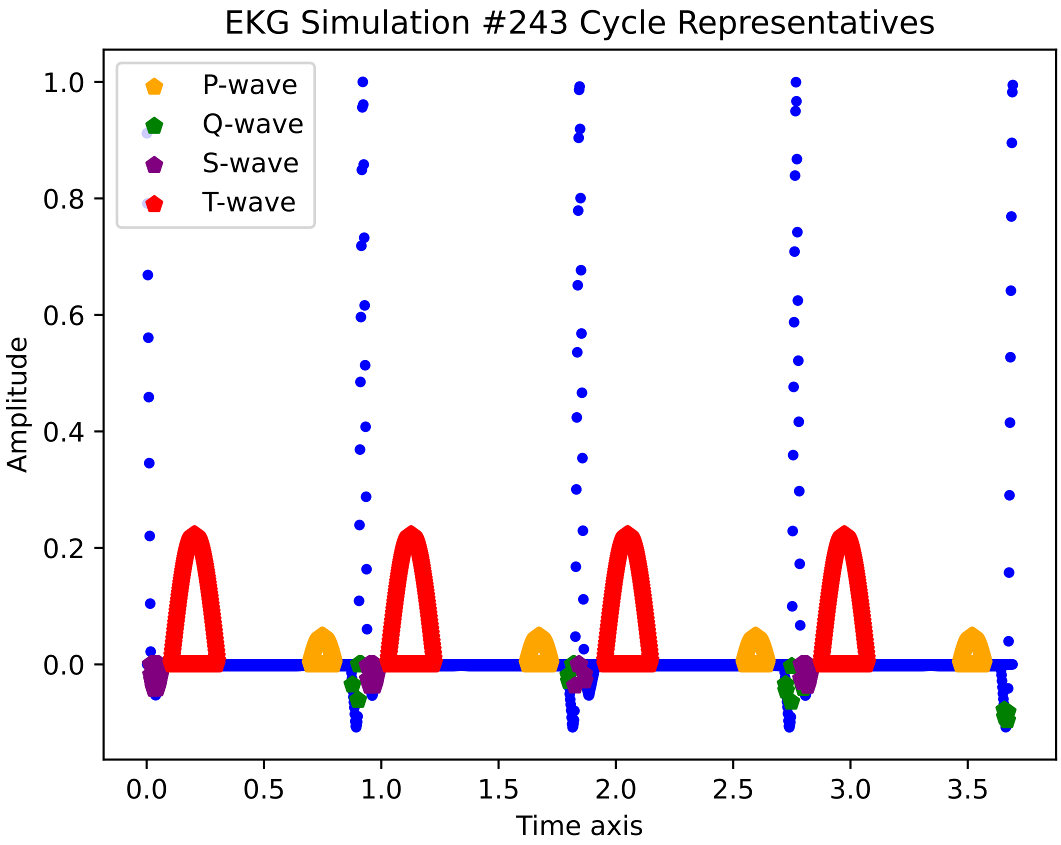

EKG Simulation #243 Cycle Representatives
P-wave
Q-wave
S-wave
T-wave
Amplitude
Time axis
1.0
0.8
0.6
0.4
0.2
0.0
0.0
0.5
1.0
1.5
2.0
2.5
3.0
3.5

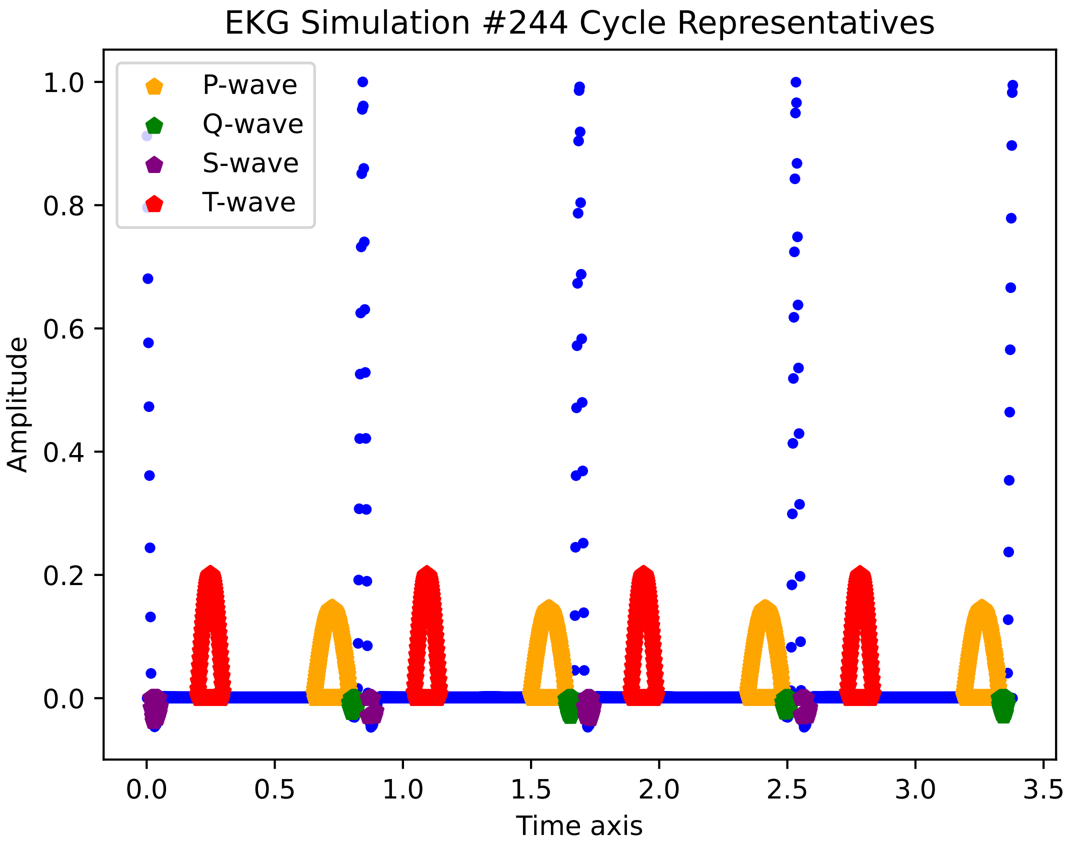

EKG Simulation #244 Cycle Representatives
P-wave
Q-wave
S-wave
T-wave
1.0
0.8
0.6
0.4
0.2
0.0
Amplitude
0.0
0.5
1.0
1.5
2.0
2.5
3.0
3.5
Time axis

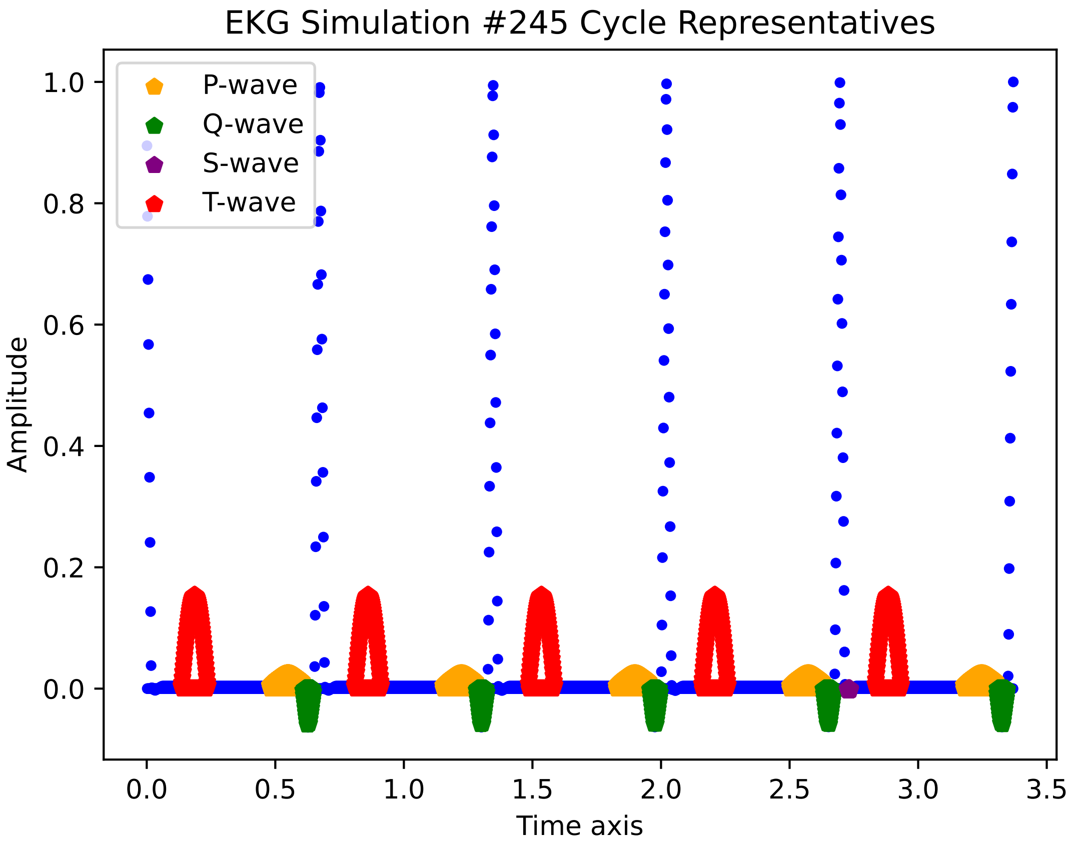

EKG Simulation #245 Cycle Representatives
P-wave
Q-wave
S-wave
T-wave
1.0
0.8
0.6
0.4
0.2
0.0
Amplitude
0.0
0.5
1.0
1.5
2.0
2.5
3.0
3.5
Time axis

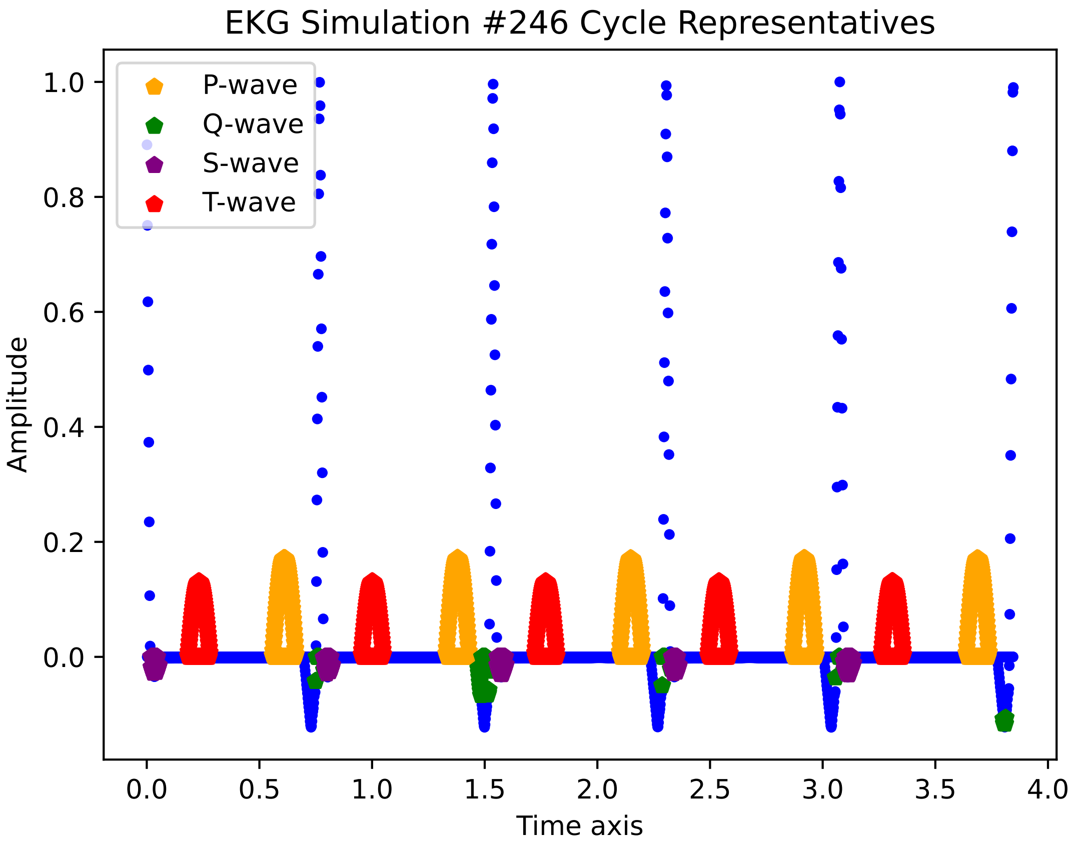

EKG Simulation #246 Cycle Representatives
P-wave
Q-wave
S-wave
T-wave
1.0
0.8
0.6
0.4
0.2
0.0
Amplitude
0.0
0.5
1.0
1.5
2.0
2.5
3.0
3.5
4.0
Time axis

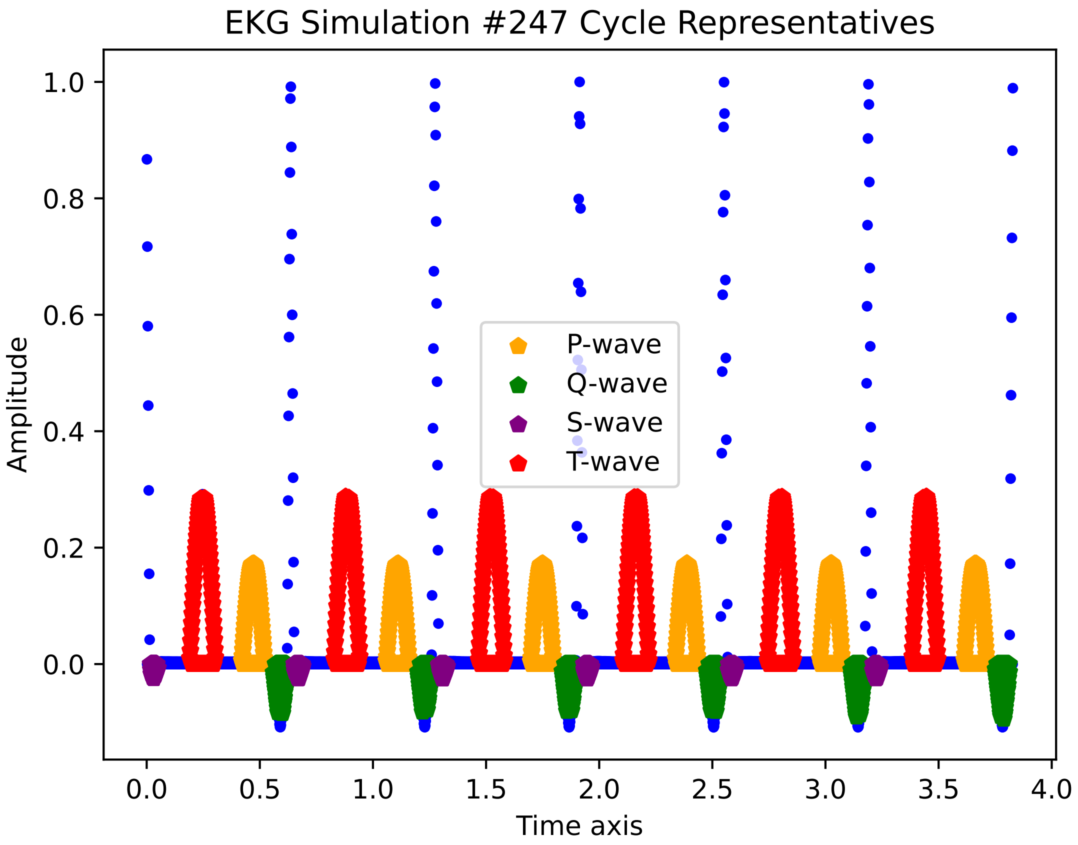

EKG Simulation #247 Cycle Representatives
Amplitude
1.0
0.8
0.6
0.4
0.2
0.0
0.0
0.5
1.0
1.5
2.0
2.5
3.0
3.5
4.0
Time axis
P-wave
Q-wave
S-wave
T-wave

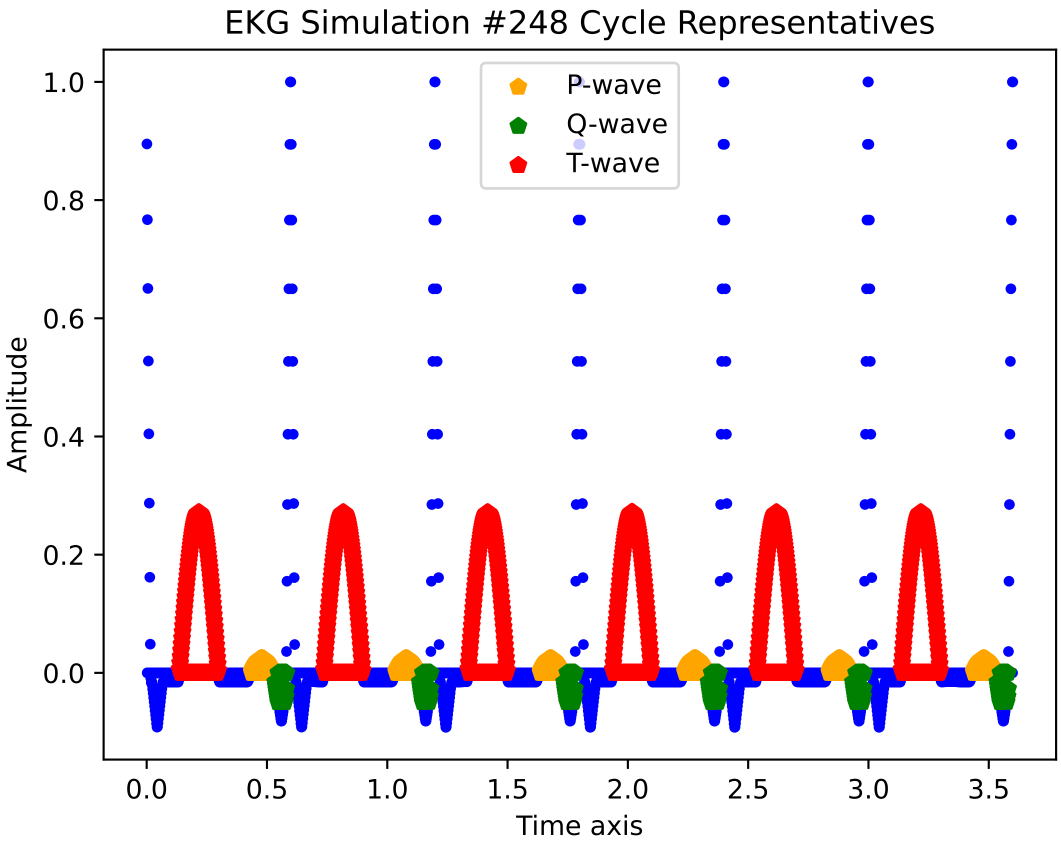
EKG Simulation #248 Cycle Representatives
P-wave
Q-wave
T-wave
Amplitude
Time axis
1.0
0.8
0.6
0.4
0.2
0.0
0.0
0.5
1.0
1.5
2.0
2.5
3.0
3.5

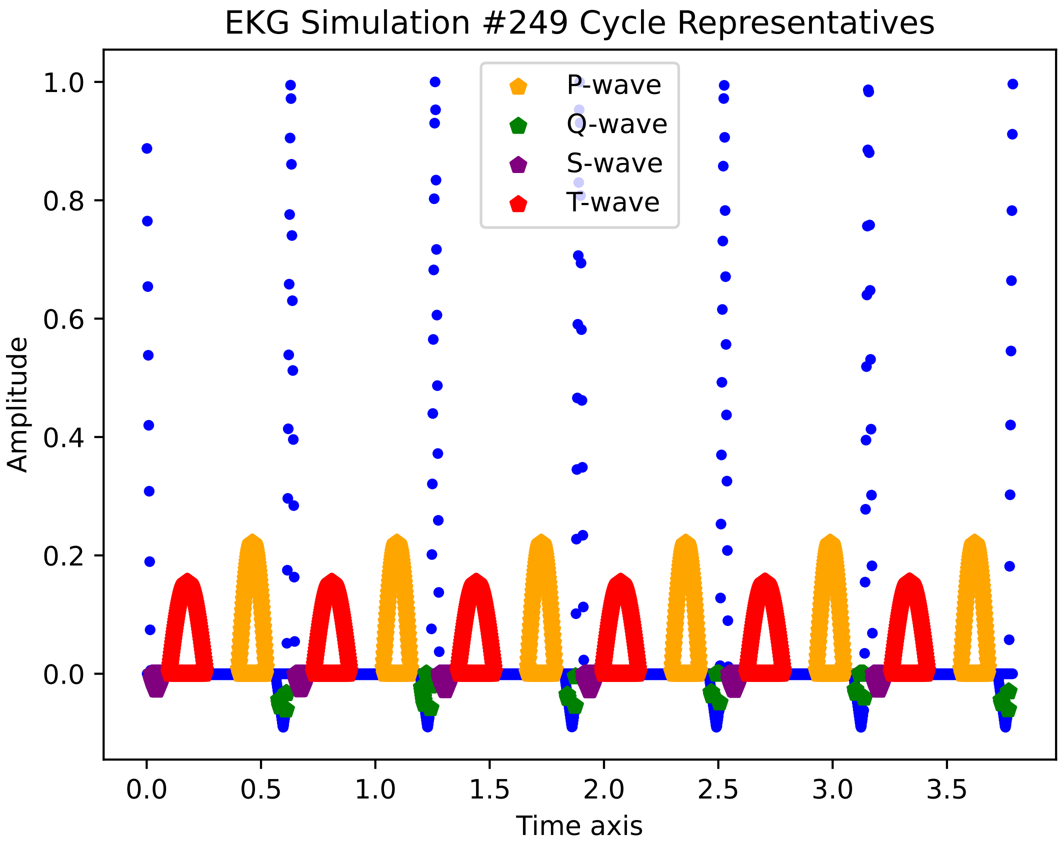
EKG Simulation #249 Cycle Representatives
P-wave
Q-wave
S-wave
T-wave
1.0
0.8
0.6
0.4
0.2
0.0
Amplitude
0.0
0.5
1.0
1.5
2.0
2.5
3.0
3.5
Time axis

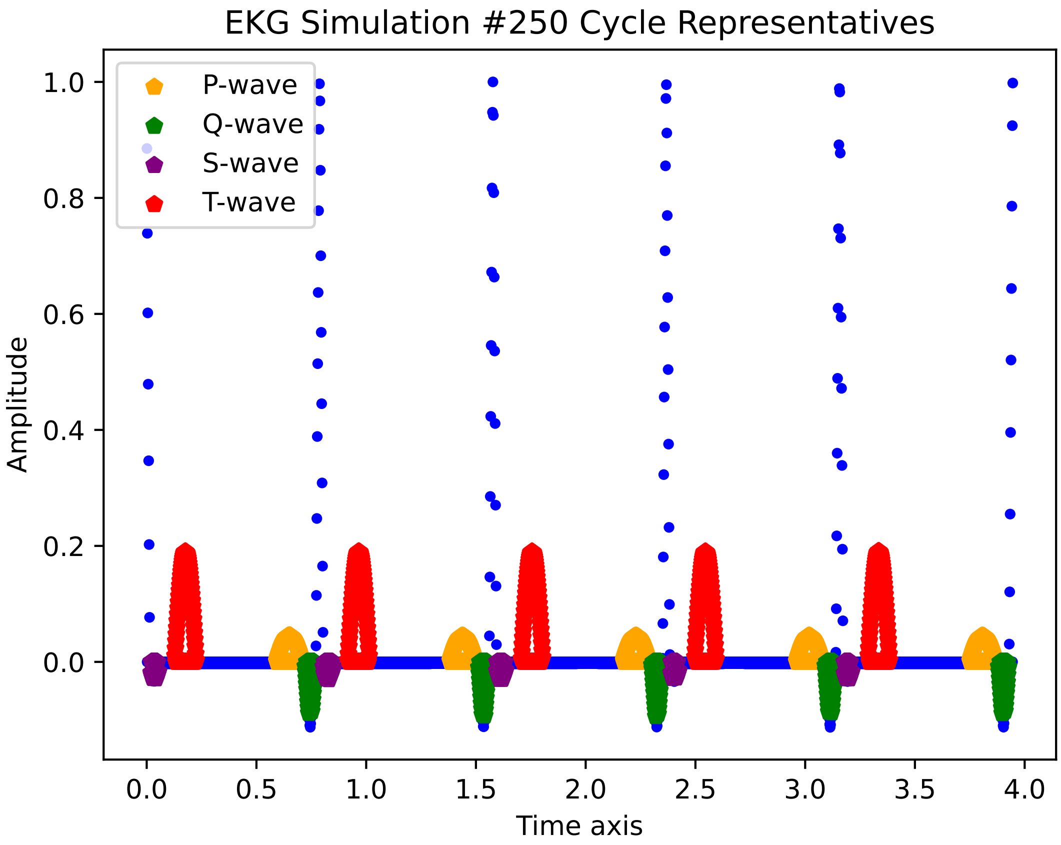
EKG Simulation #250 Cycle Representatives
P-wave
Q-wave
S-wave
T-wave
Amplitude
Time axis
1.0
0.8
0.6
0.4
0.2
0.0
0.0
0.5
1.0
1.5
2.0
2.5
3.0
3.5
4.0

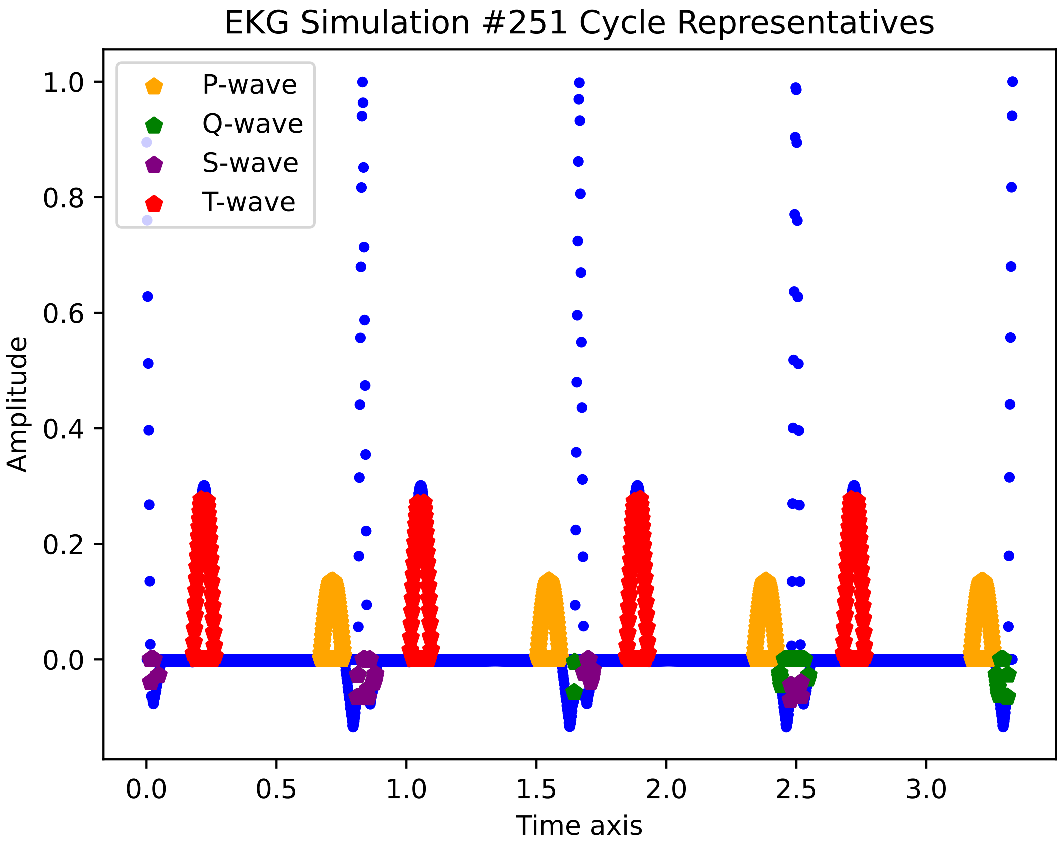

EKG Simulation #251 Cycle Representatives
P-wave
Q-wave
S-wave
T-wave
Amplitude
Time axis
1.0
0.8
0.6
0.4
0.2
0.0
0.0
0.5
1.0
1.5
2.0
2.5
3.0

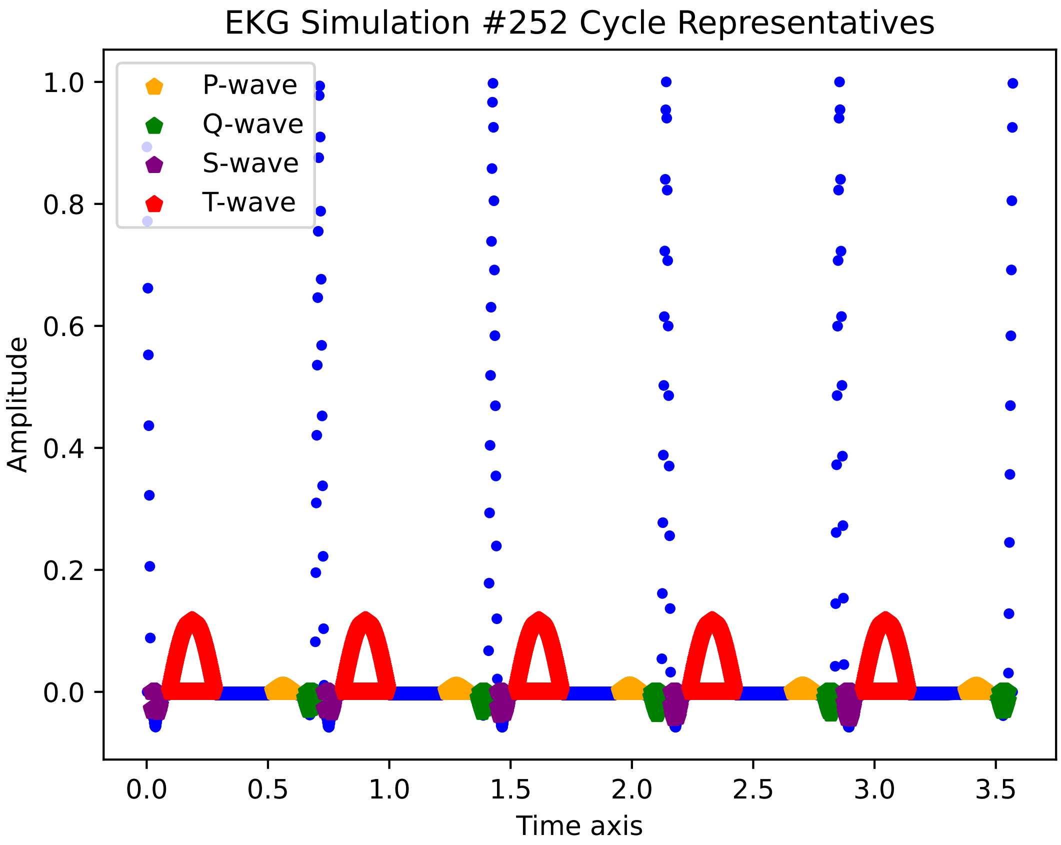
EKG Simulation #252 Cycle Representatives
P-wave
Q-wave
S-wave
T-wave
Amplitude
Time axis
1.0
0.8
0.6
0.4
0.2
0.0
0.0
0.5
1.0
1.5
2.0
2.5
3.0
3.5

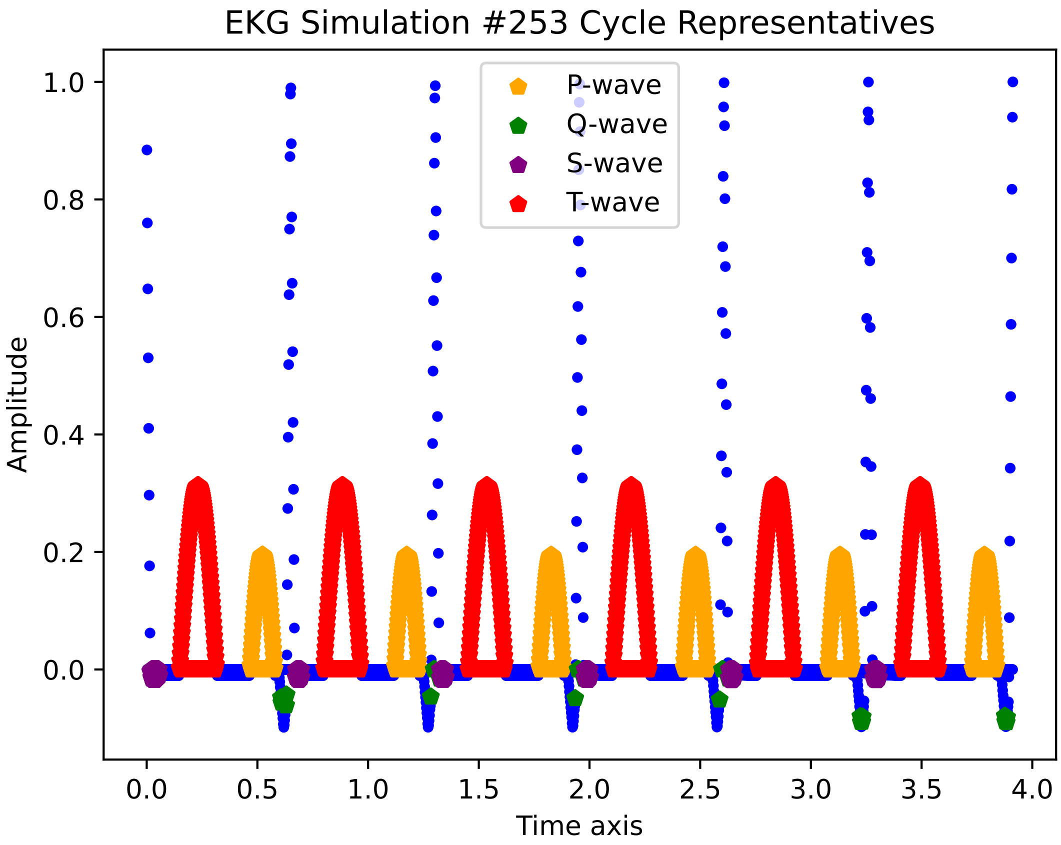

EKG Simulation #253 Cycle Representatives
P-wave
Q-wave
S-wave
T-wave
Amplitude
Time axis
1.0
0.8
0.6
0.4
0.2
0.0
0.0
0.5
1.0
1.5
2.0
2.5
3.0
3.5
4.0

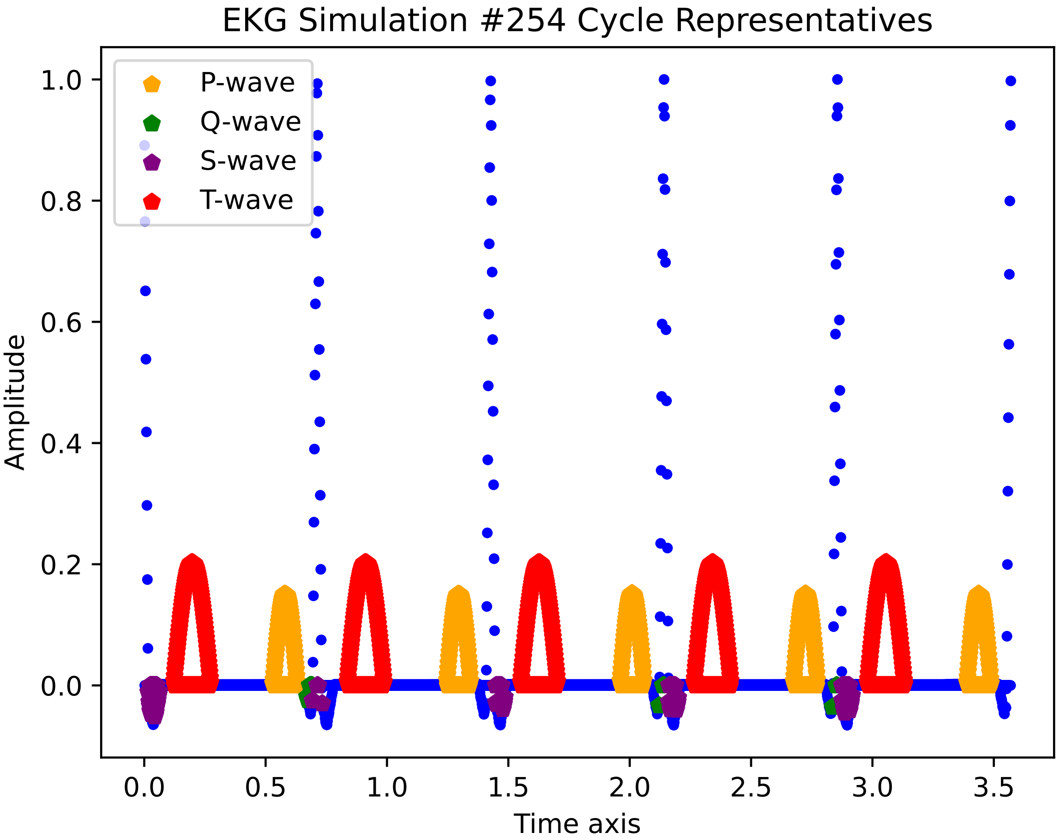

EKG Simulation #254 Cycle Representatives
P-wave
Q-wave
S-wave
T-wave
Amplitude
Time axis
1.0
0.8
0.6
0.4
0.2
0.0
0.0
0.5
1.0
1.5
2.0
2.5
3.0
3.5

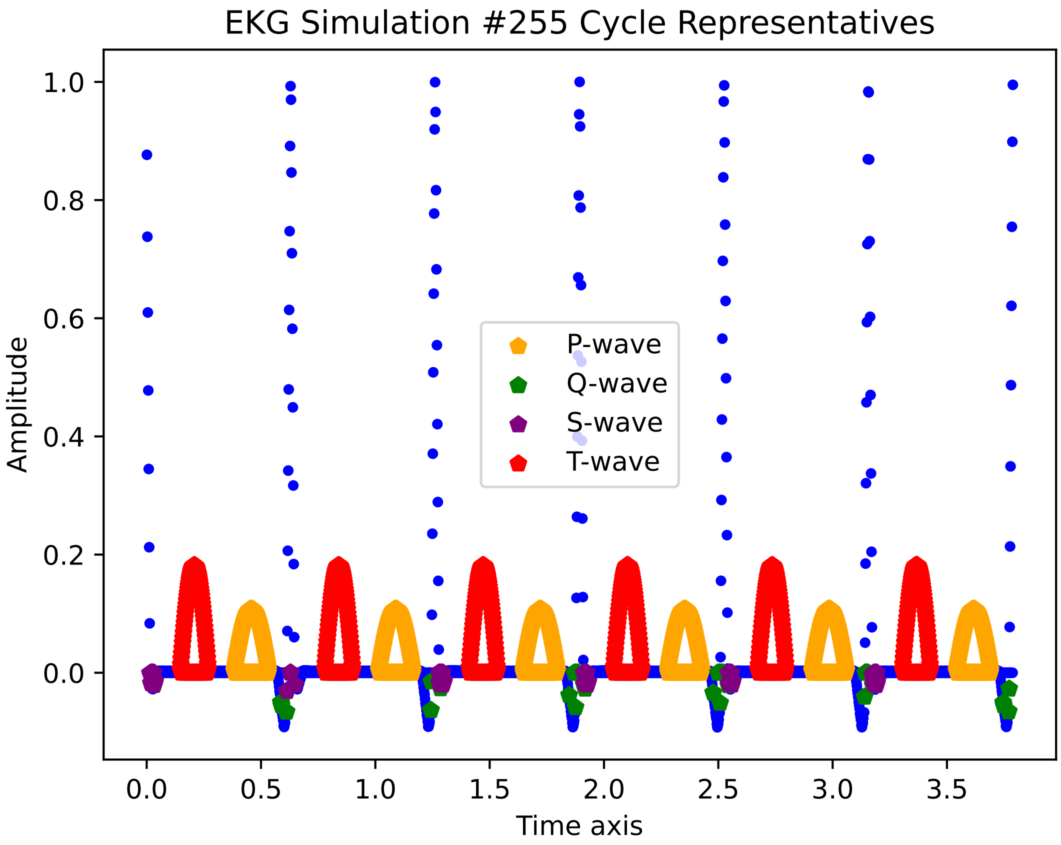

EKG Simulation #255 Cycle Representatives
Amplitude
Time axis
P-wave
Q-wave
S-wave
T-wave
1.0
0.8
0.6
0.4
0.2
0.0
0.0
0.5
1.0
1.5
2.0
2.5
3.0
3.5

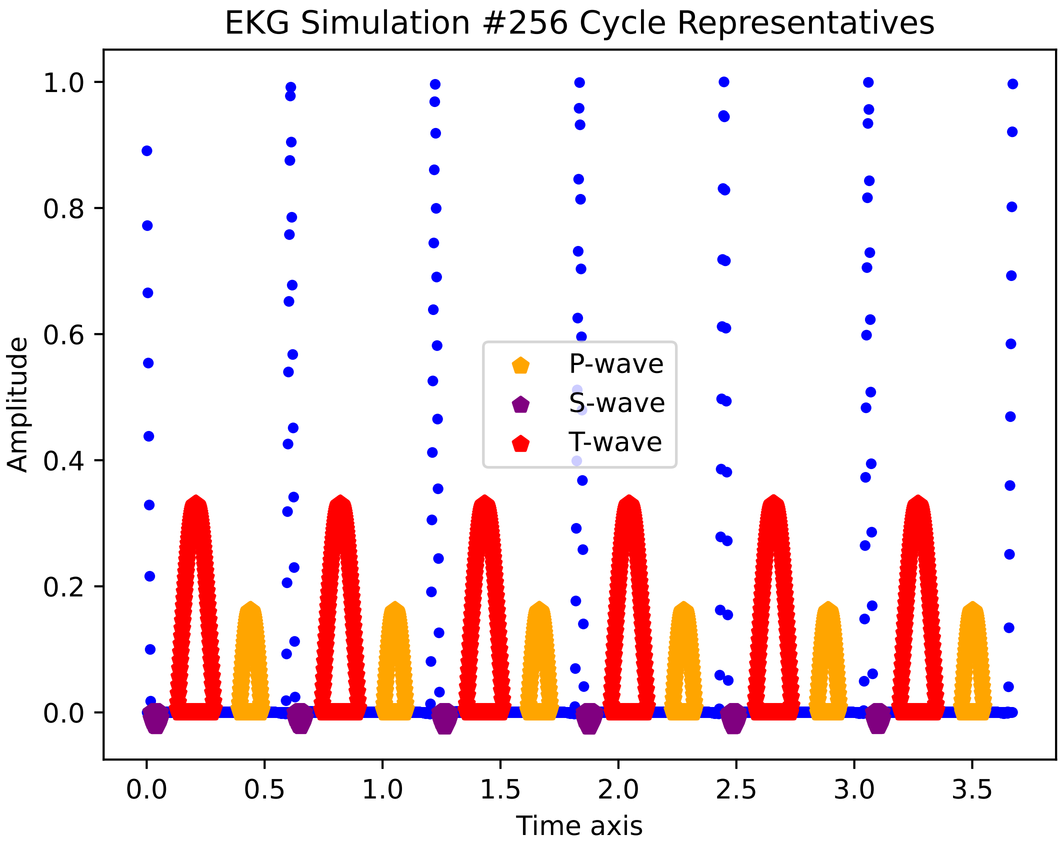
EKG Simulation #256 Cycle Representatives
Amplitude
Time axis
P-wave
S-wave
T-wave
1.0
0.8
0.6
0.4
0.2
0.0
0.0
0.5
1.0
1.5
2.0
2.5
3.0
3.5

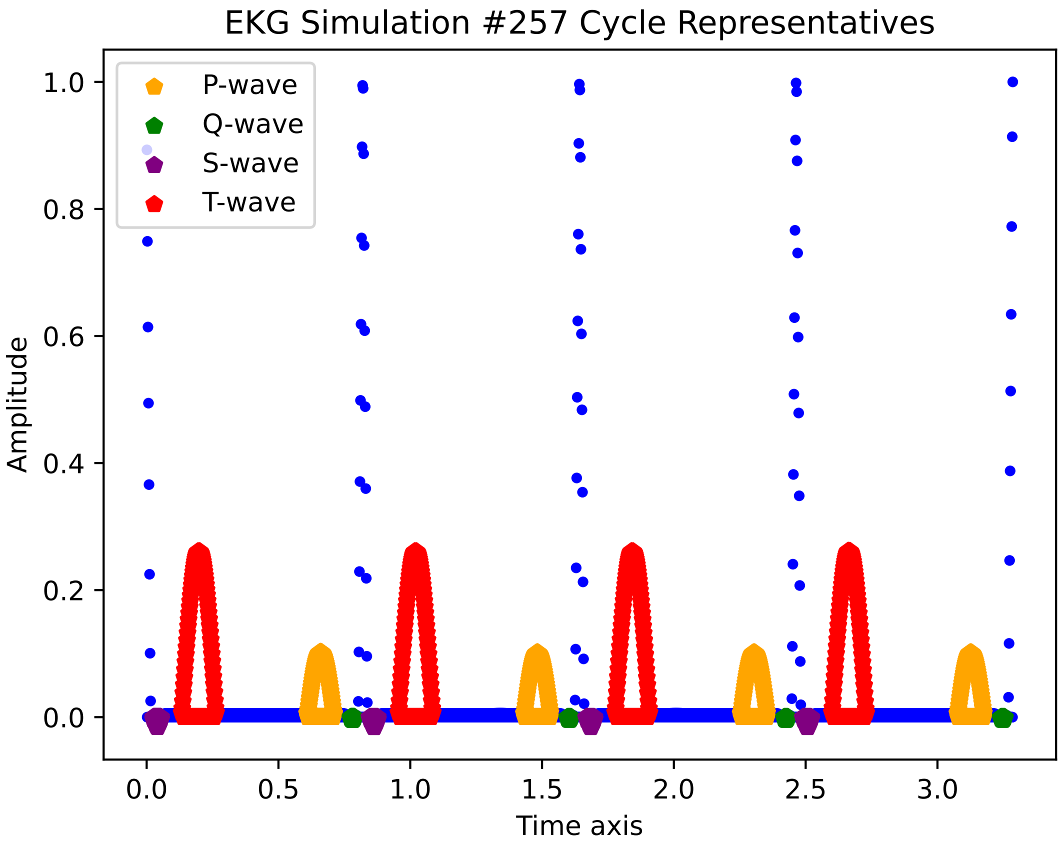
EKG Simulation #257 Cycle Representatives
P-wave
Q-wave
S-wave
T-wave
Amplitude
Time axis
1.0
0.8
0.6
0.4
0.2
0.0
0.0
0.5
1.0
1.5
2.0
2.5
3.0

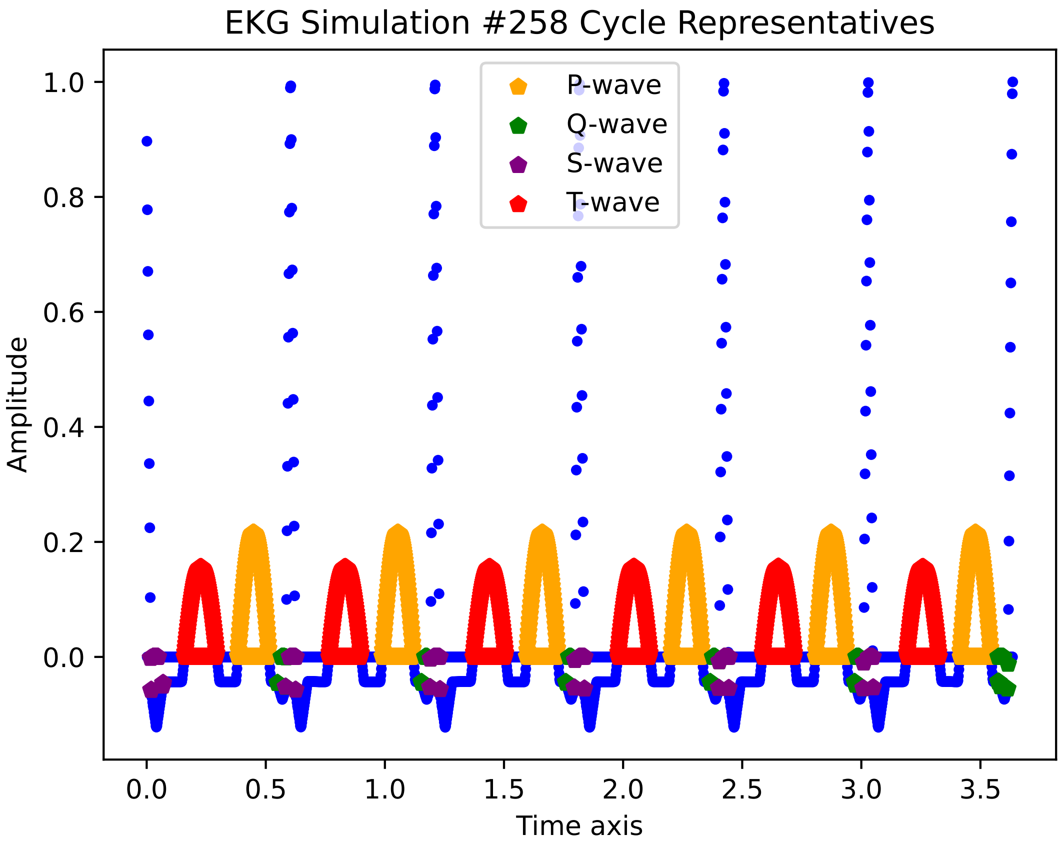
EKG Simulation #258 Cycle Representatives
P-wave
Q-wave
S-wave
T-wave
Amplitude
1.0
0.8
0.6
0.4
0.2
0.0
0.0
0.5
1.0
1.5
2.0
2.5
3.0
3.5
Time axis

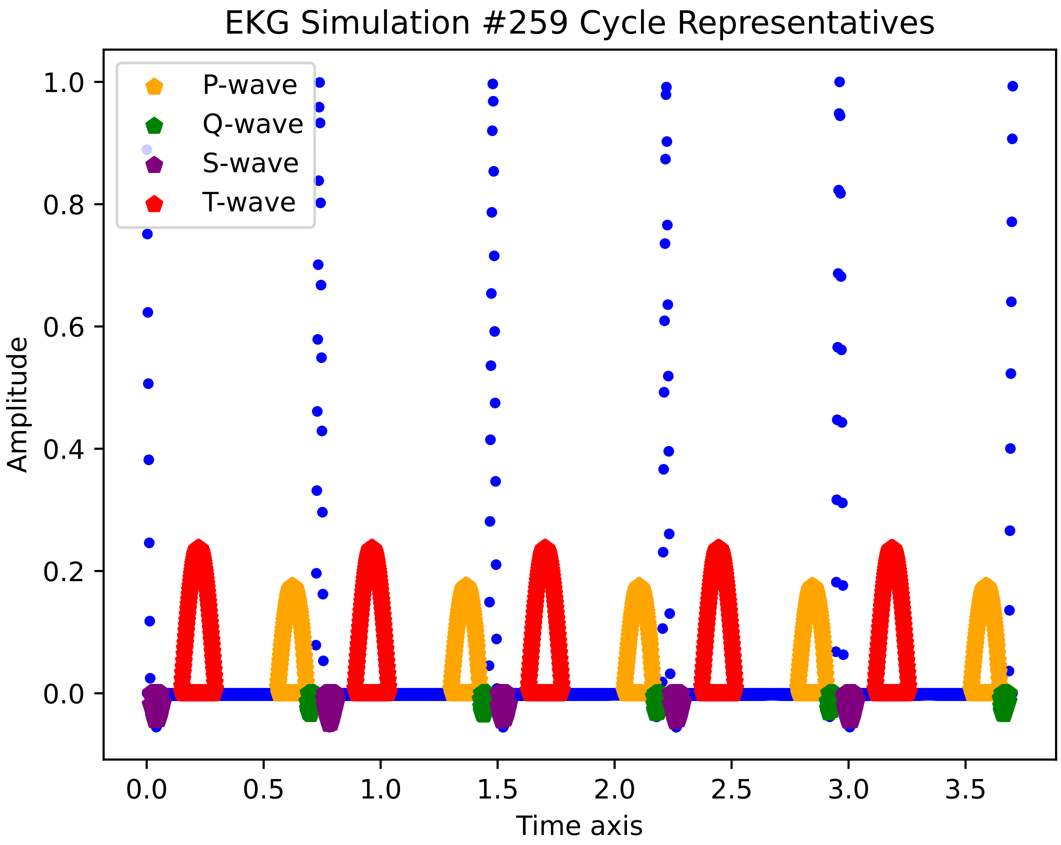
EKG Simulation #259 Cycle Representatives
P-wave
Q-wave
S-wave
T-wave
Amplitude
Time axis
1.0
0.8
0.6
0.4
0.2
0.0
0.0
0.5
1.0
1.5
2.0
2.5
3.0
3.5

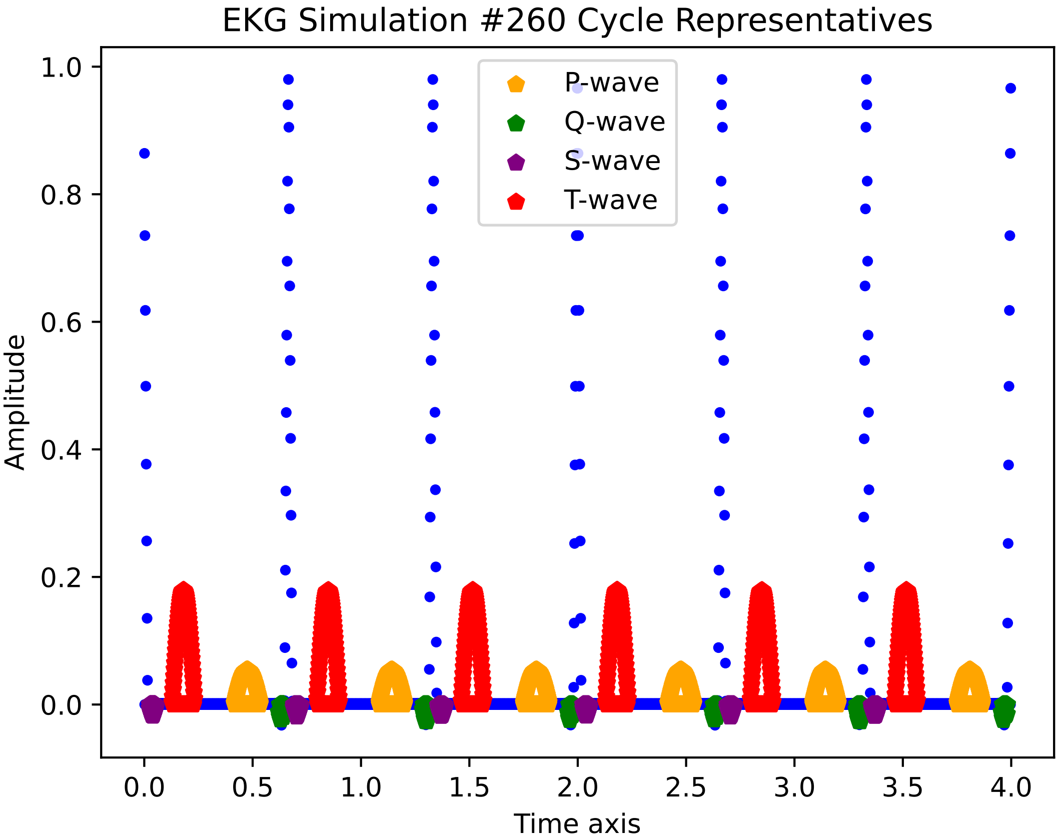
EKG Simulation #260 Cycle Representatives
P-wave
Q-wave
S-wave
T-wave
Amplitude
Time axis
1.0
0.8
0.6
0.4
0.2
0.0
0.0
0.5
1.0
1.5
2.0
2.5
3.0
3.5
4.0

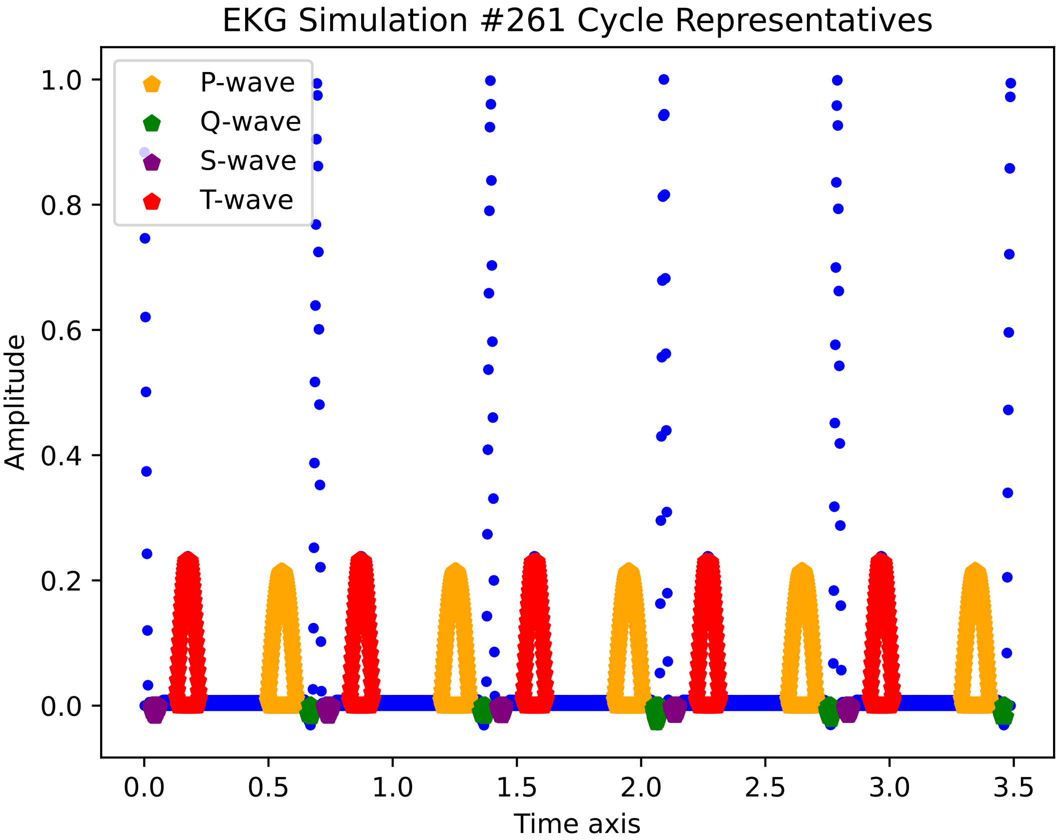
EKG Simulation #261 Cycle Representatives
P-wave
Q-wave
S-wave
T-wave
Amplitude
Time axis
0.0
0.2
0.4
0.6
0.8
1.0
0.5
1.5
2.0
2.5
3.0
3.5

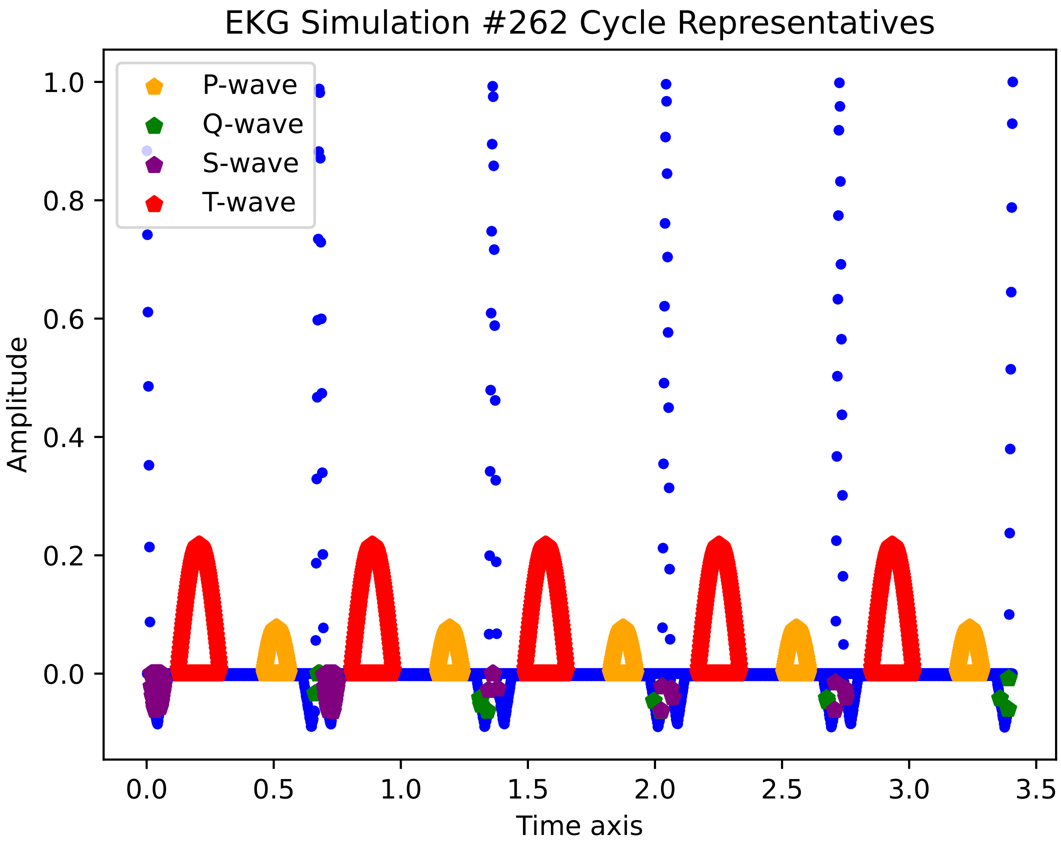

EKG Simulation #262 Cycle Representatives
P-wave
Q-wave
S-wave
T-wave
1.0
0.8
0.6
0.4
0.2
0.0
Amplitude
0.0
0.5
1.0
1.5
2.0
2.5
3.0
3.5
Time axis

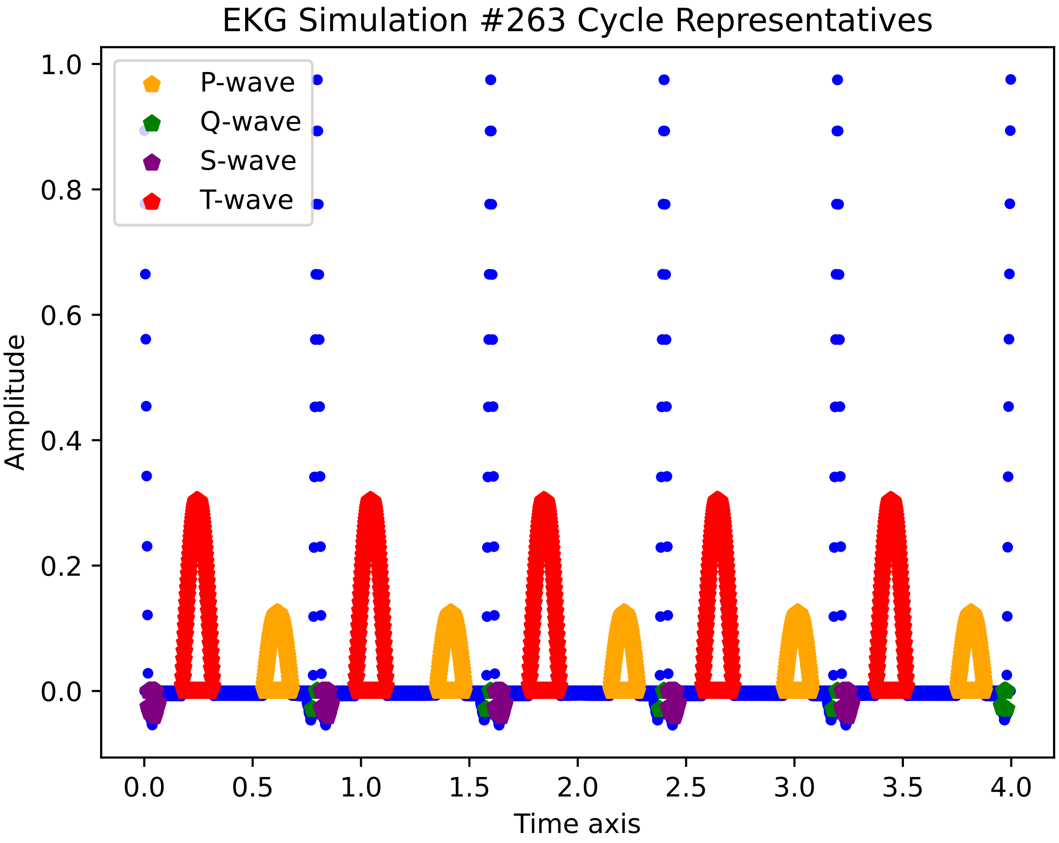

EKG Simulation #263 Cycle Representatives
P-wave
Q-wave
S-wave
T-wave
Amplitude
1.0
0.8
0.6
0.4
0.2
0.0
0.0
0.5
1.0
1.5
2.0
2.5
3.0
3.5
4.0
Time axis

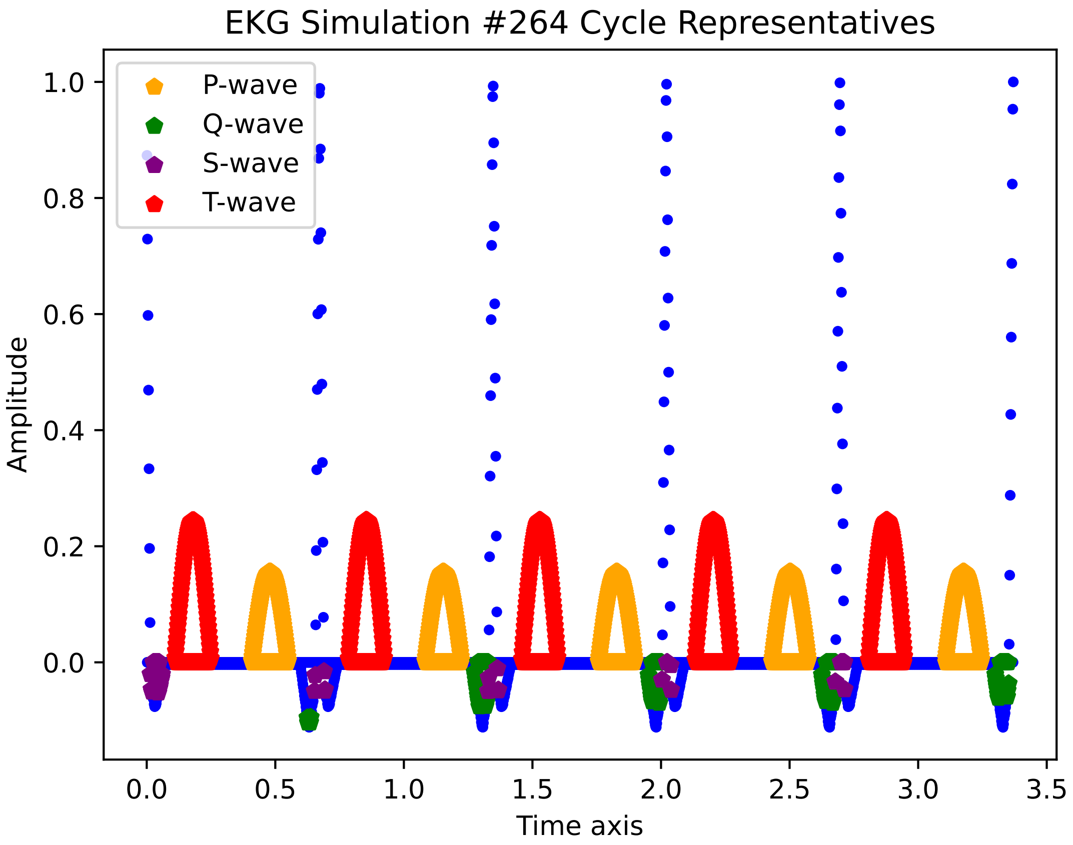
EKG Simulation #264 Cycle Representatives
P-wave
Q-wave
S-wave
T-wave
Amplitude
Time axis
1.0
0.8
0.6
0.4
0.2
0.0
0.0
0.5
1.0
1.5
2.0
2.5
3.0
3.5

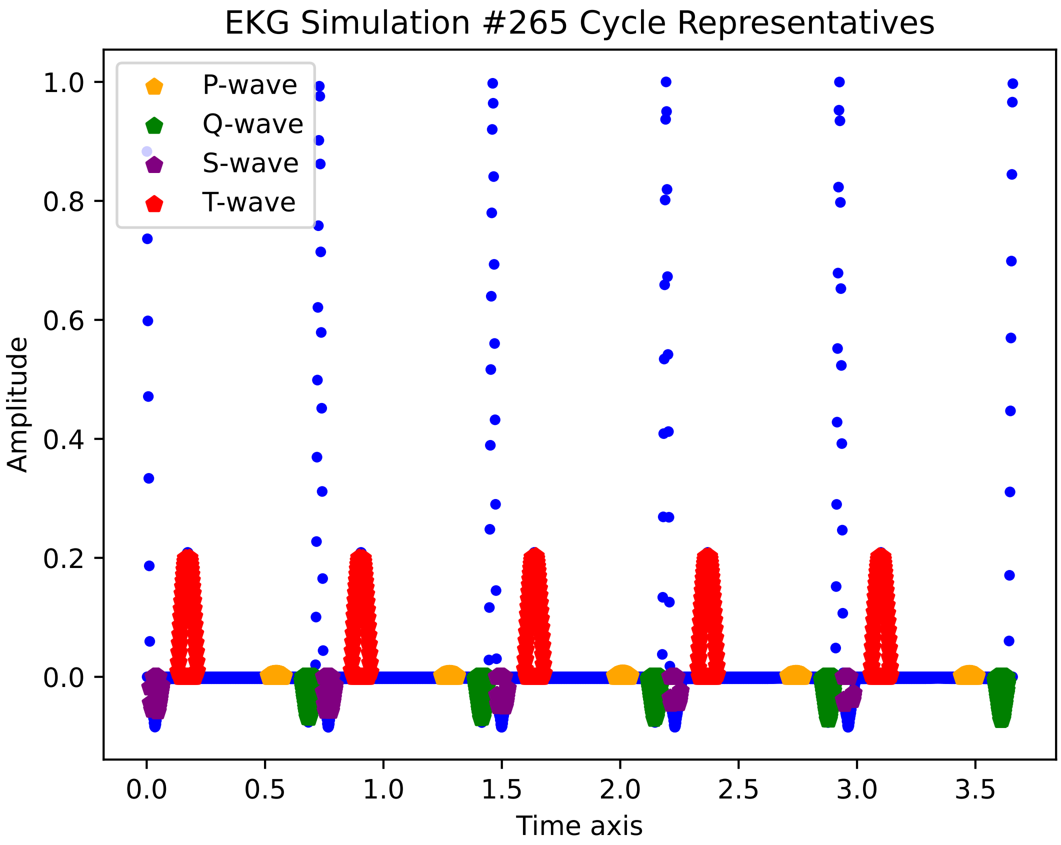

EKG Simulation #265 Cycle Representatives
P-wave
Q-wave
S-wave
T-wave
Amplitude
Time axis
1.0
0.8
0.6
0.4
0.2
0.0
0.0
0.5
1.0
1.5
2.0
2.5
3.0
3.5

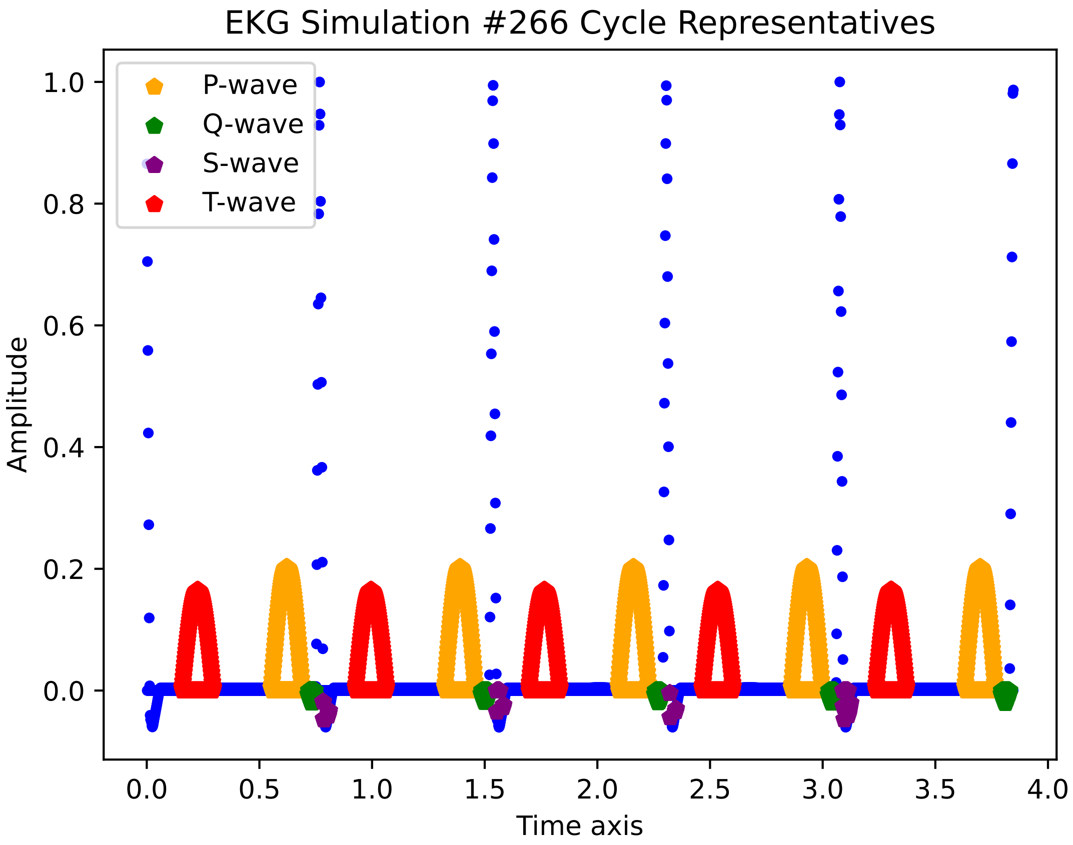
EKG Simulation #266 Cycle Representatives
P-wave
Q-wave
S-wave
T-wave
Amplitude
1.0
0.8
0.6
0.4
0.2
0.0
0.0
0.5
1.0
1.5
2.0
2.5
3.0
3.5
4.0
Time axis

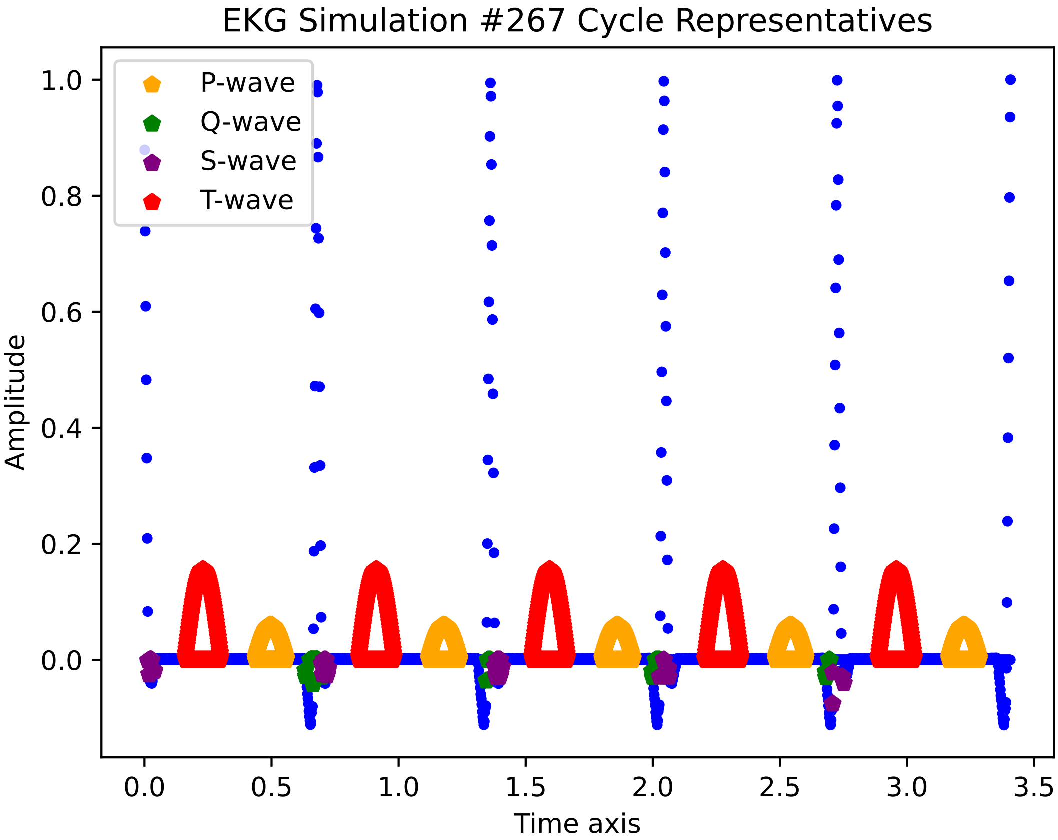

EKG Simulation #267 Cycle Representatives
P-wave
Q-wave
S-wave
T-wave
Amplitude
Time axis
1.0
0.8
0.6
0.4
0.2
0.0
0.0
0.5
1.0
1.5
2.0
2.5
3.0
3.5

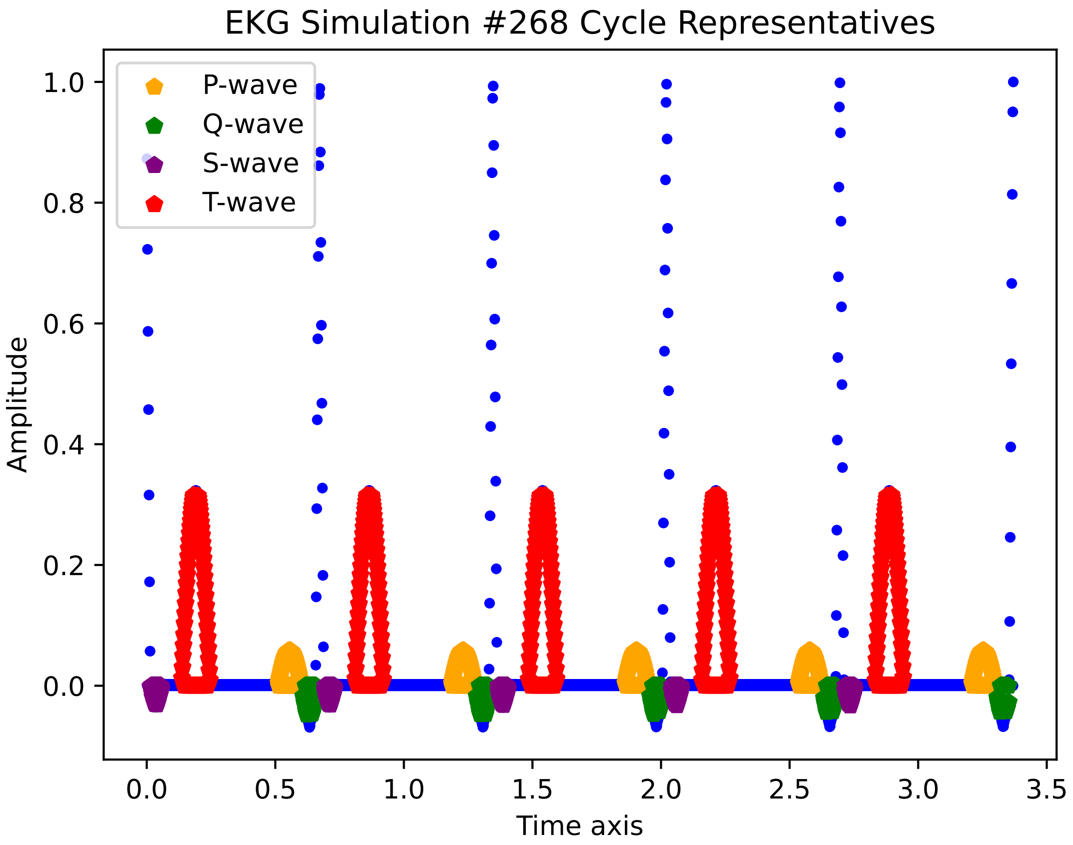
EKG Simulation #268 Cycle Representatives
P-wave
Q-wave
S-wave
T-wave
Amplitude
1.0
0.8
0.6
0.4
0.2
0.0
0.0
0.5
1.0
1.5
2.0
2.5
3.0
3.5
Time axis

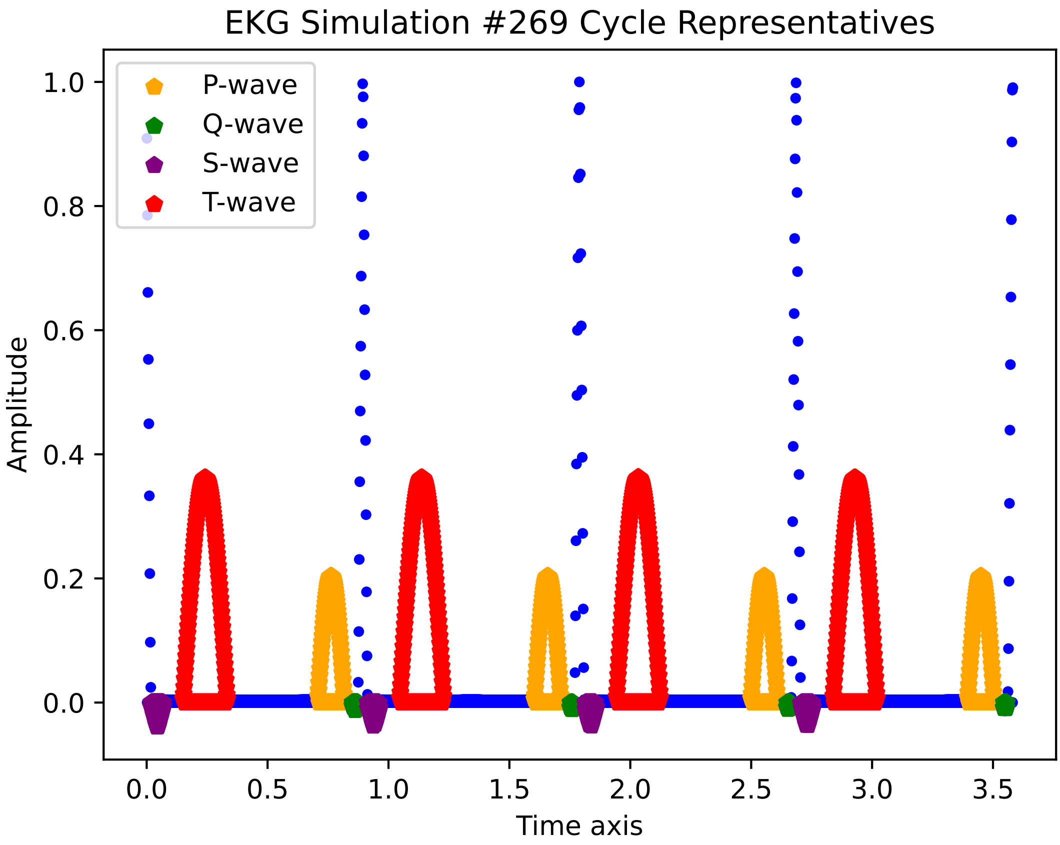
EKG Simulation #269 Cycle Representatives
P-wave
Q-wave
S-wave
T-wave
1.0
0.8
0.6
0.4
0.2
0.0
Amplitude
0.0
0.5
1.0
1.5
2.0
2.5
3.0
3.5
Time axis

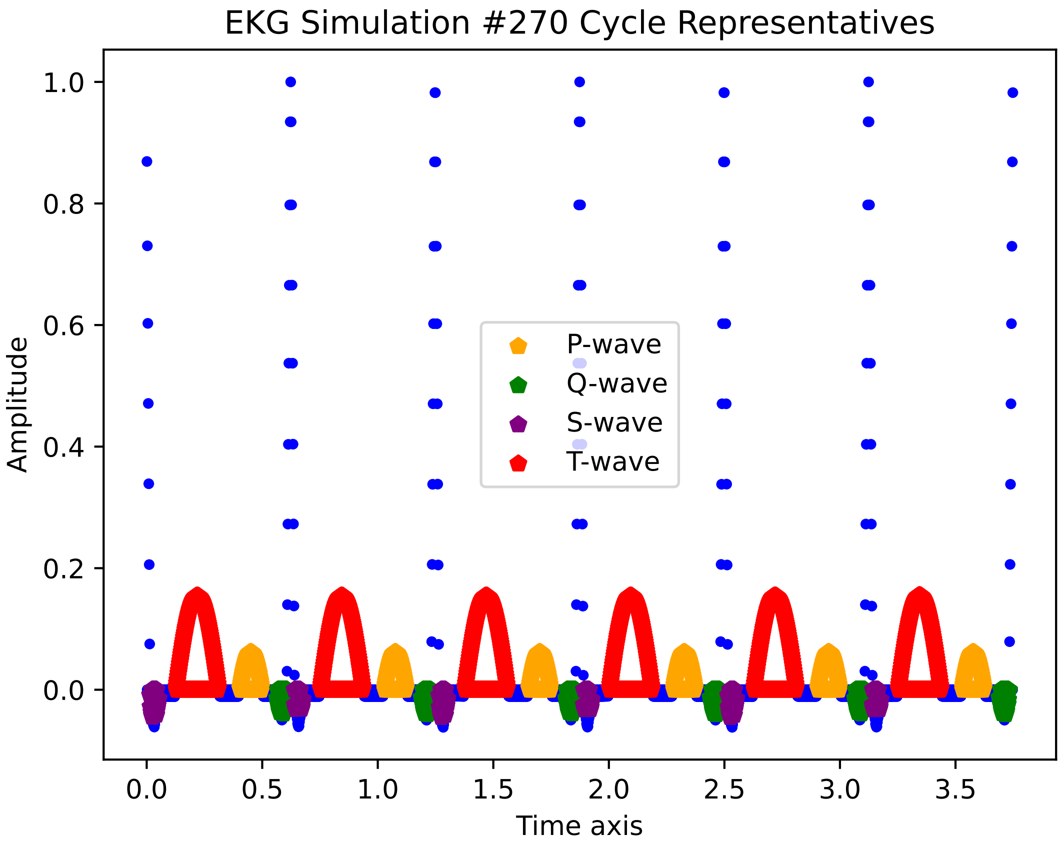
EKG Simulation #270 Cycle Representatives
Amplitude
1.0
0.8
0.6
0.4
0.2
0.0
0.0
0.5
1.0
1.5
2.0
2.5
3.0
3.5
Time axis
P-wave
Q-wave
S-wave
T-wave

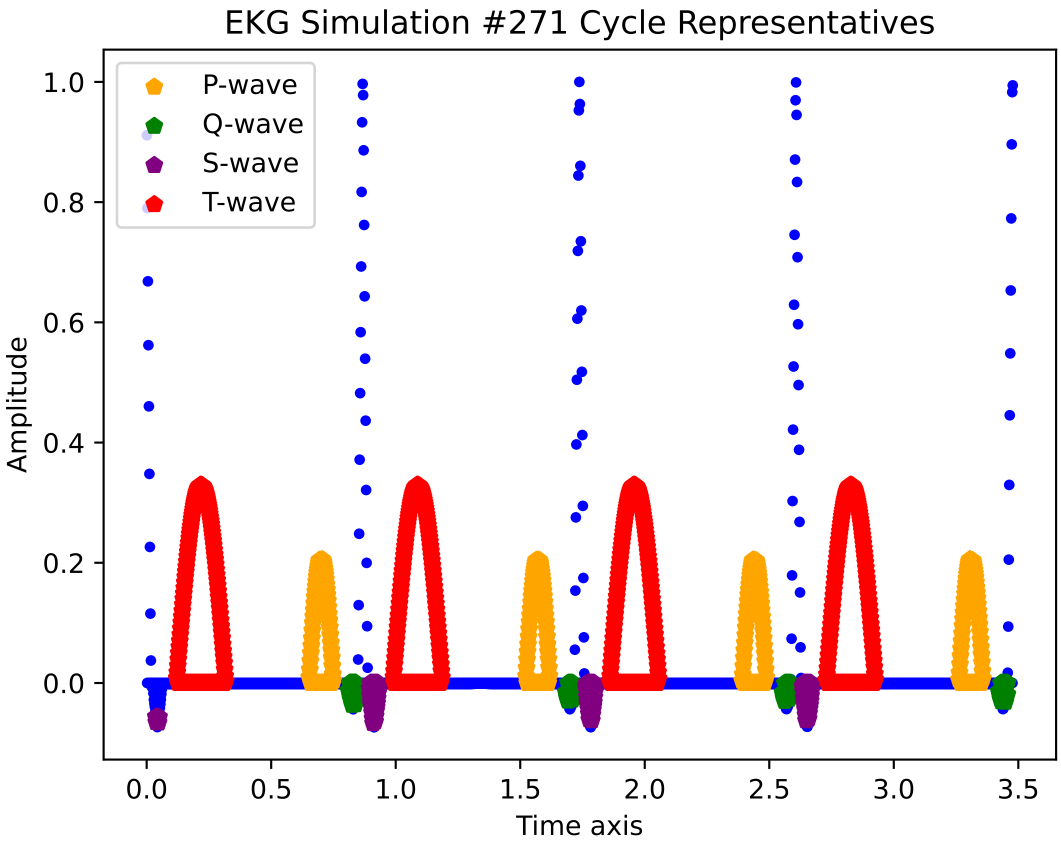

EKG Simulation #271 Cycle Representatives
P-wave
Q-wave
S-wave
T-wave
1.0
0.8
0.6
0.4
0.2
0.0
Amplitude
0.0
0.5
1.0
1.5
2.0
2.5
3.0
3.5
Time axis

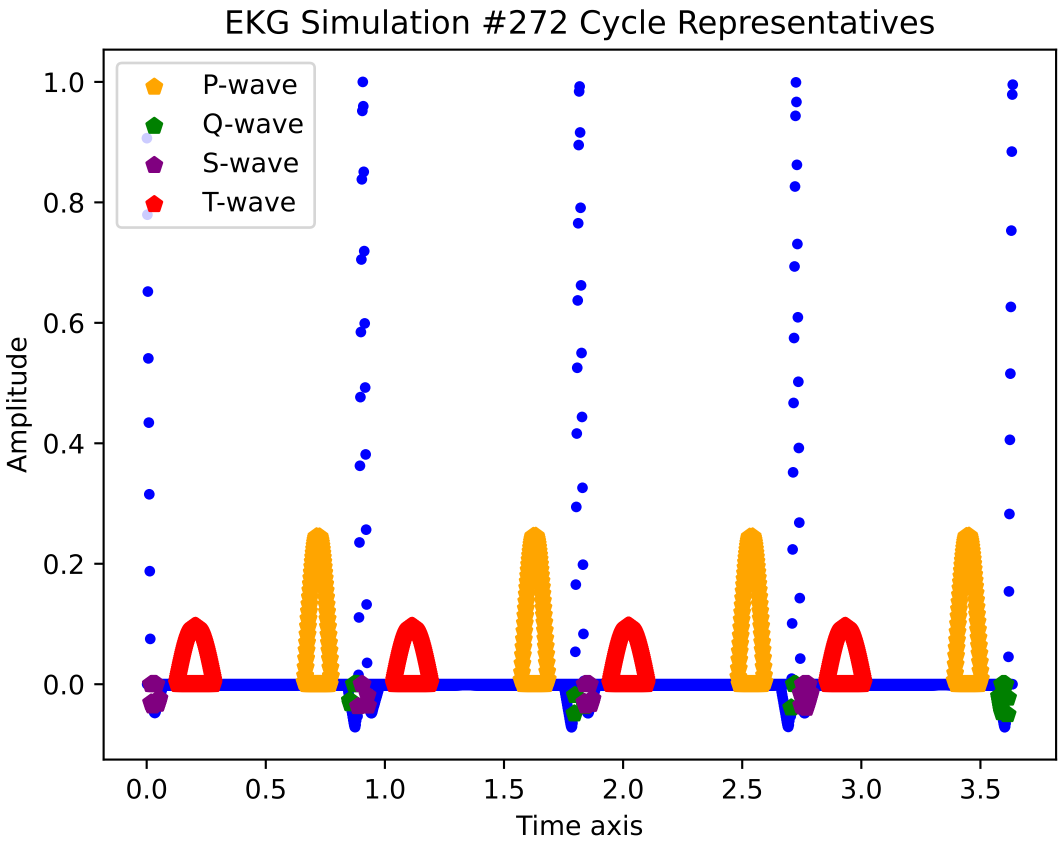
EKG Simulation #272 Cycle Representatives
P-wave
Q-wave
S-wave
T-wave
Amplitude
Time axis
1.0
0.8
0.6
0.4
0.2
0.0
0.0
0.5
1.0
1.5
2.0
2.5
3.0
3.5

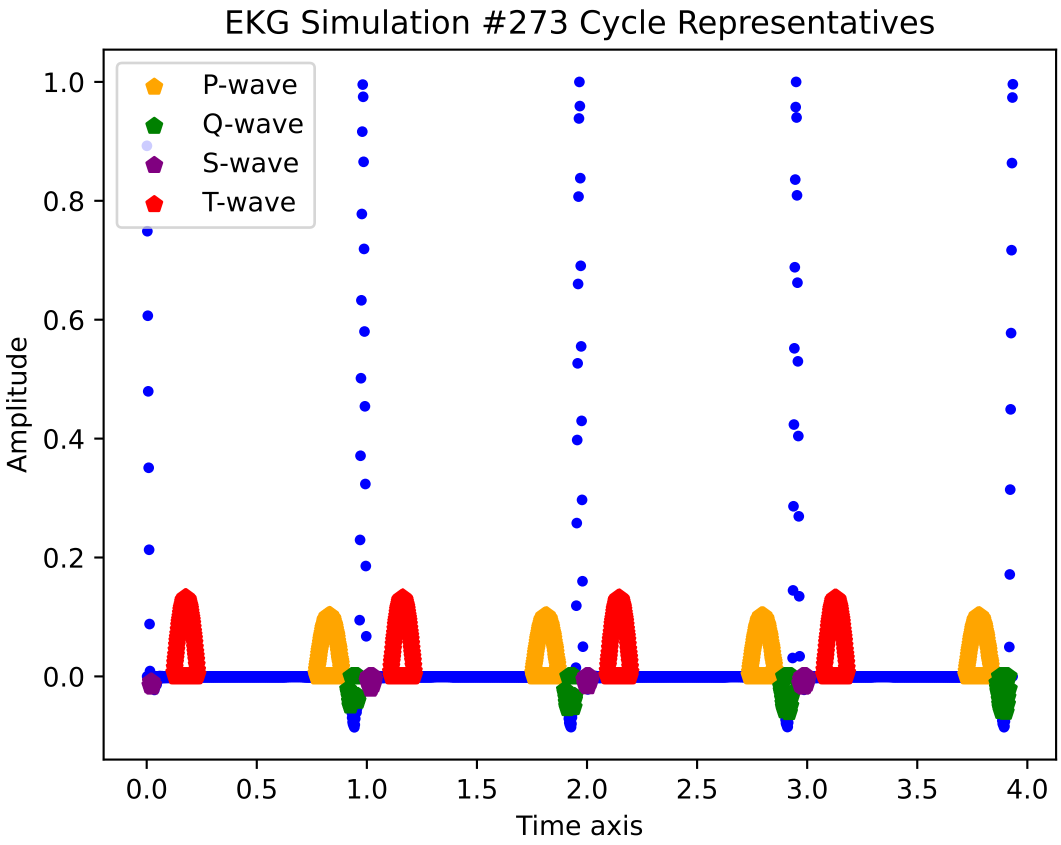

EKG Simulation #273 Cycle Representatives
P-wave
Q-wave
S-wave
T-wave
Amplitude
Time axis
1.0
0.8
0.6
0.4
0.2
0.0
0.0
0.5
1.0
1.5
2.0
2.5
3.0
3.5
4.0

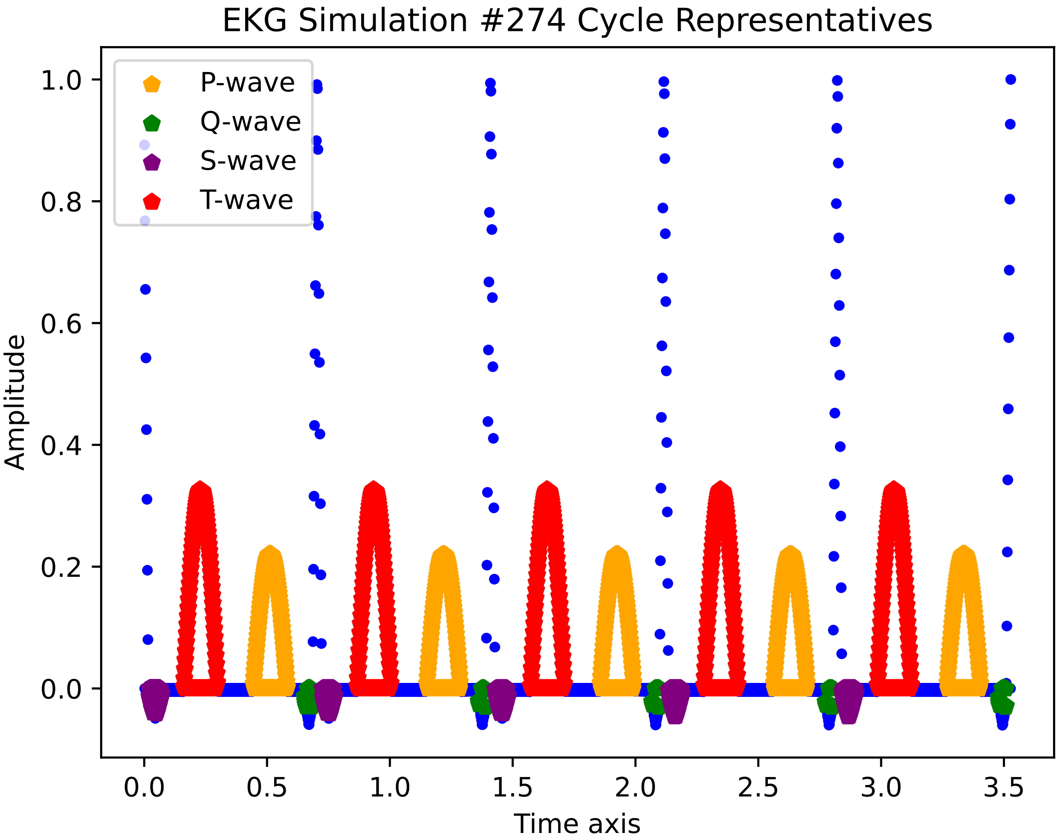

EKG Simulation #274 Cycle Representatives
P-wave
Q-wave
S-wave
T-wave
1.0
0.8
0.6
0.4
0.2
0.0
Amplitude
0.0
0.5
1.0
1.5
2.0
2.5
3.0
3.5
Time axis

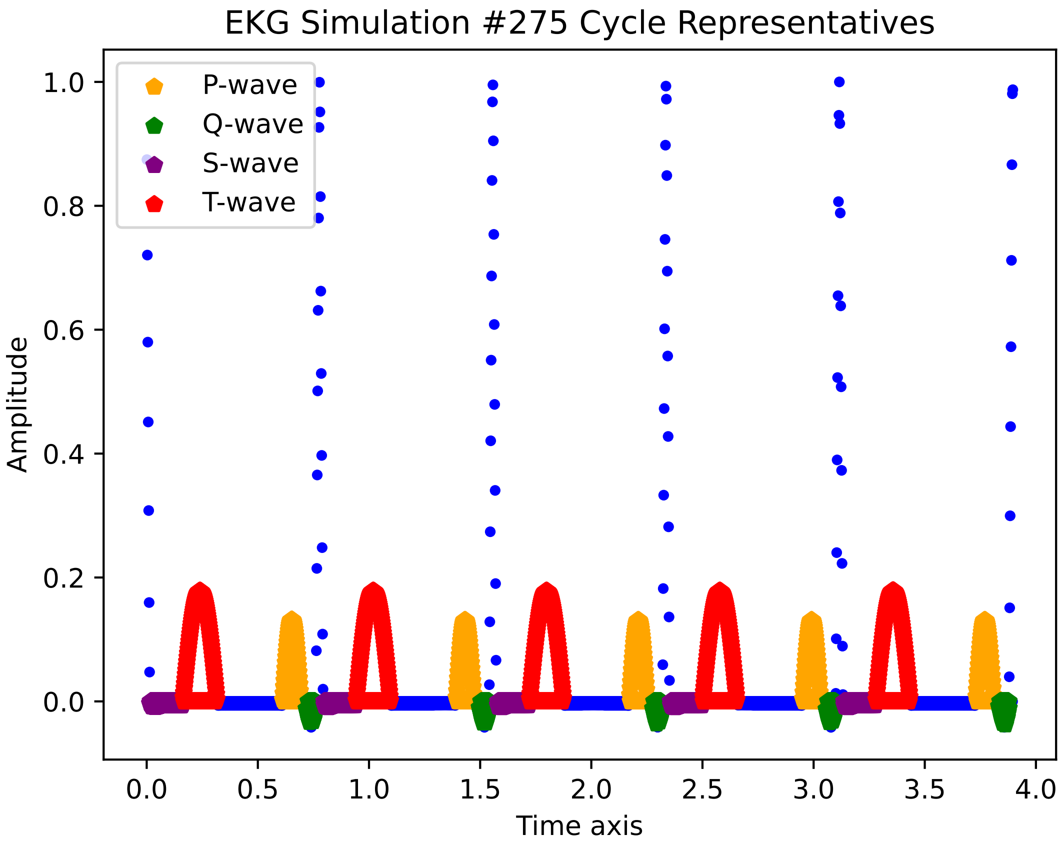
EKG Simulation #275 Cycle Representatives
P-wave
Q-wave
S-wave
T-wave
Amplitude
Time axis
1.0
0.8
0.6
0.4
0.2
0.0
0.0
0.5
1.0
1.5
2.0
2.5
3.0
3.5
4.0

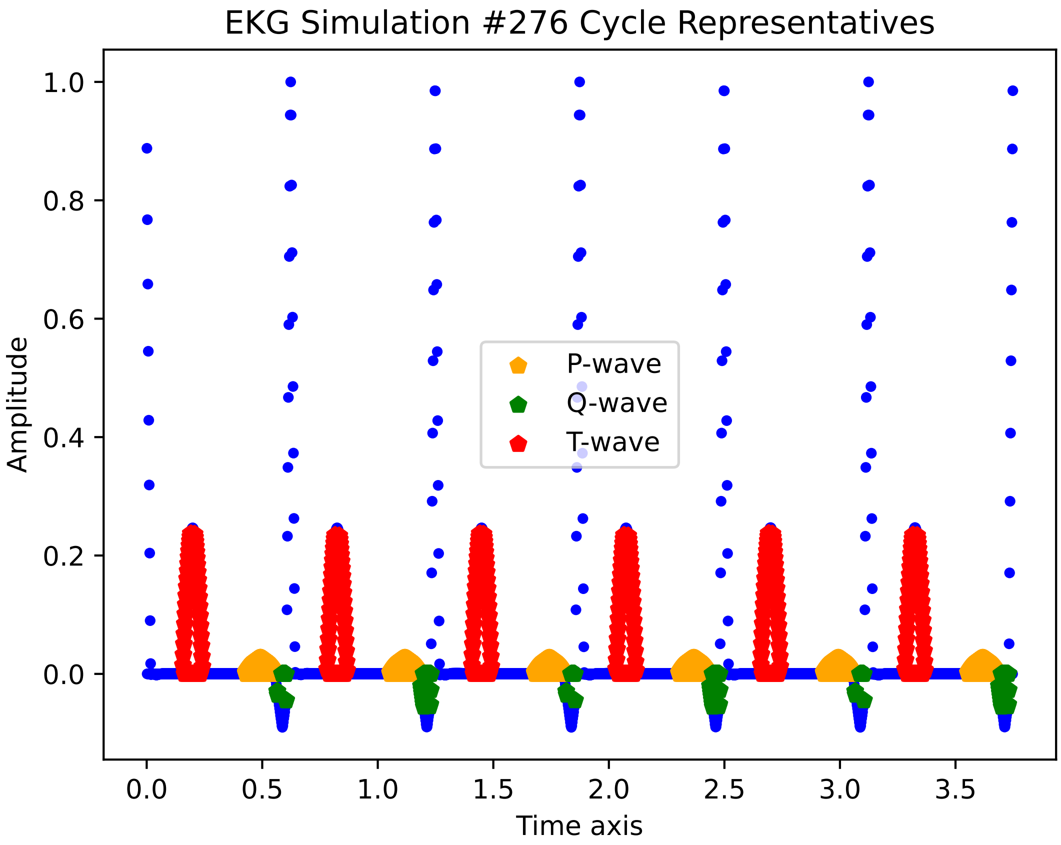
EKG Simulation #276 Cycle Representatives
Amplitude
1.0
0.8
0.6
0.4
0.2
0.0
0.0
0.5
1.0
1.5
2.0
2.5
3.0
3.5
Time axis
P-wave
Q-wave
T-wave

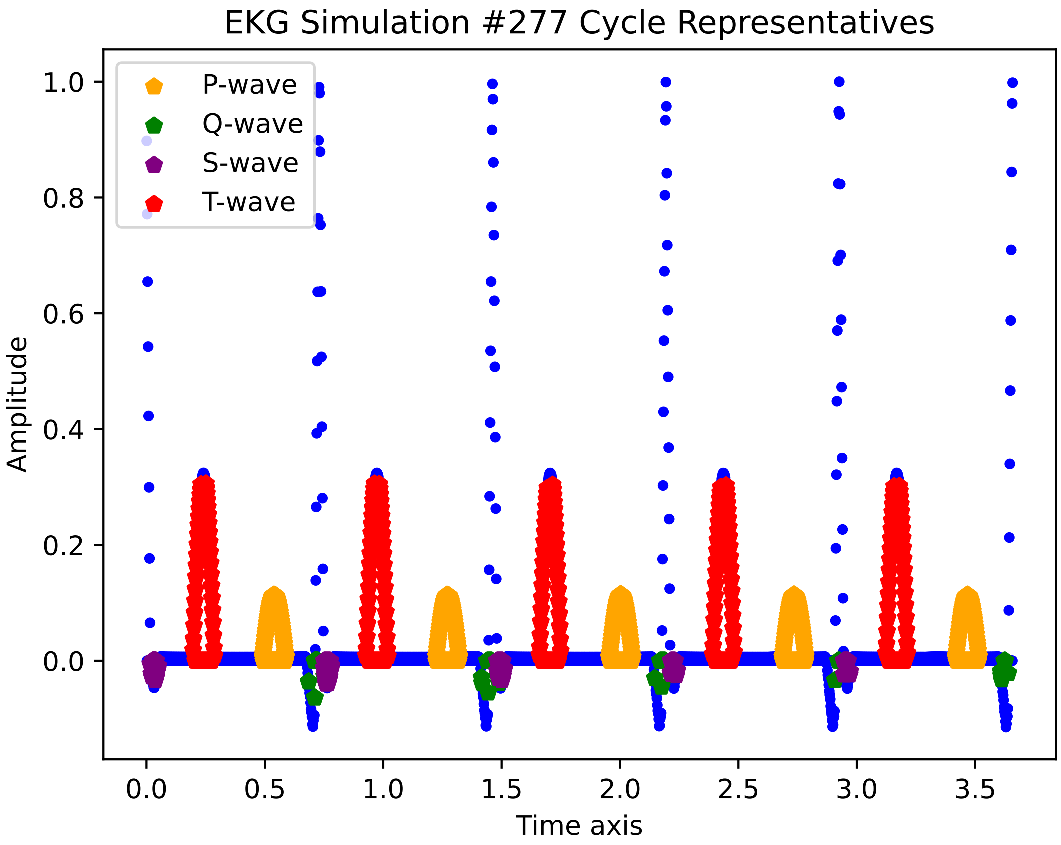

EKG Simulation #277 Cycle Representatives
P-wave
Q-wave
S-wave
T-wave
Amplitude
1.0
0.8
0.6
0.4
0.2
0.0
0.0
0.5
1.0
1.5
2.0
2.5
3.0
3.5
Time axis

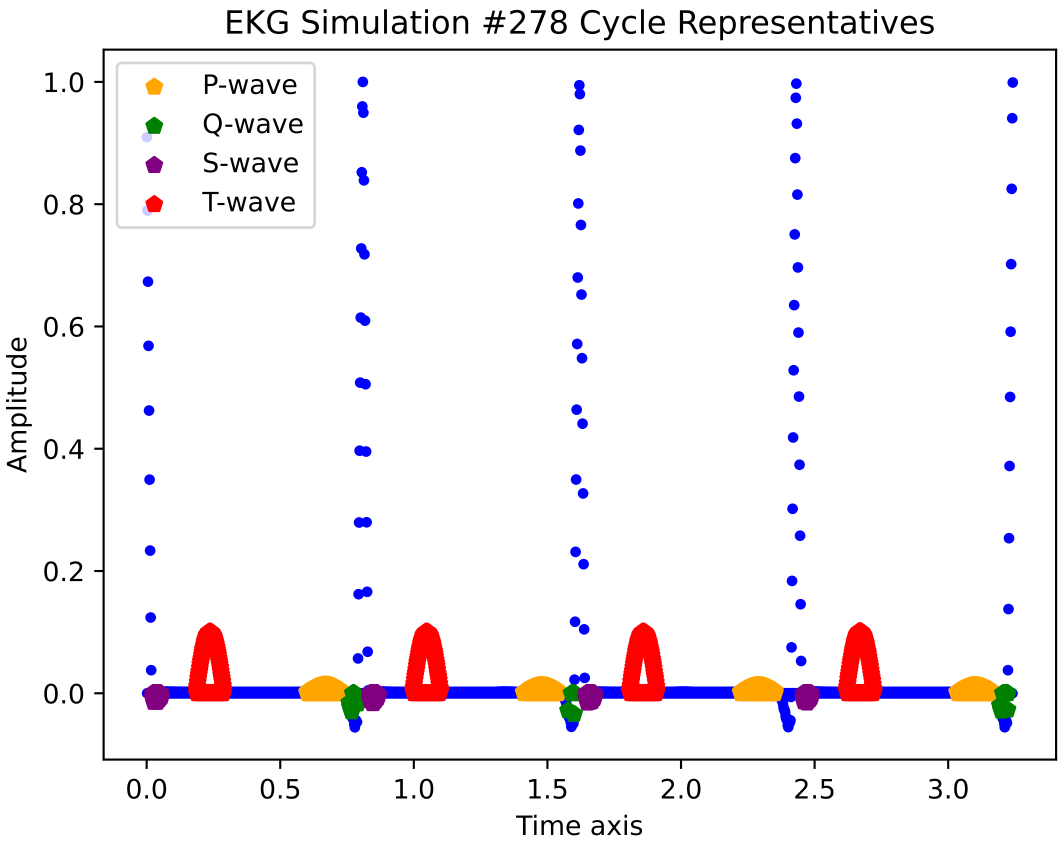

EKG Simulation #278 Cycle Representatives
P-wave
Q-wave
S-wave
T-wave
Amplitude
Time axis
1.0
0.8
0.6
0.4
0.2
0.0
0.0
0.5
1.0
1.5
2.0
2.5
3.0

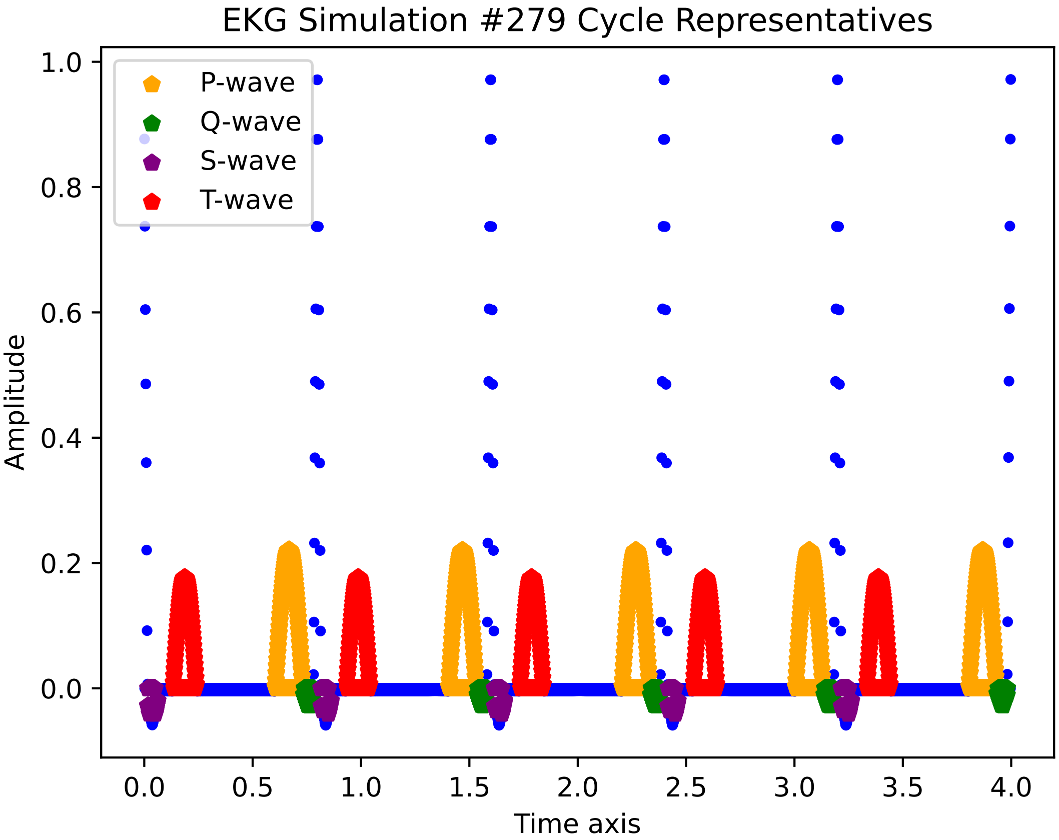
EKG Simulation #279 Cycle Representatives
P-wave
Q-wave
S-wave
T-wave
Amplitude
Time axis
1.0
0.8
0.6
0.4
0.2
0.0
0.0
0.5
1.0
1.5
2.0
2.5
3.0
3.5
4.0

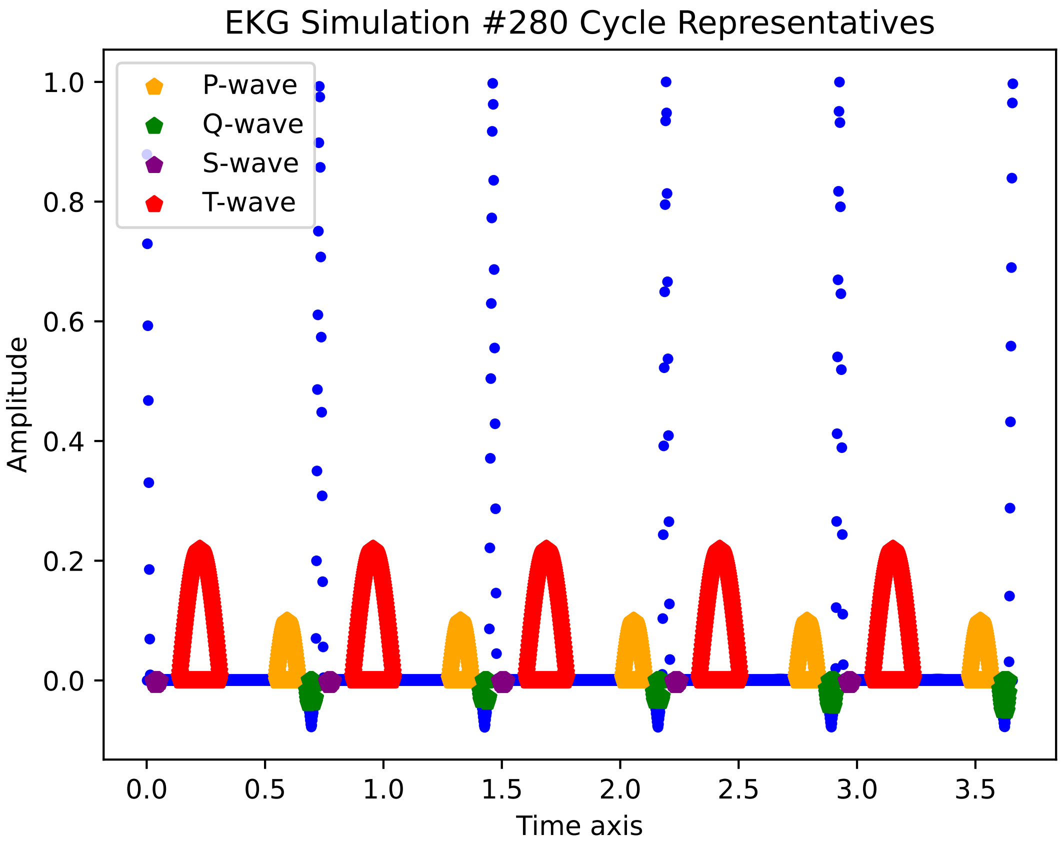
EKG Simulation #280 Cycle Representatives
P-wave
Q-wave
S-wave
T-wave
Amplitude
1.0
0.8
0.6
0.4
0.2
0.0
0.0
0.5
1.0
1.5
2.0
2.5
3.0
3.5
Time axis

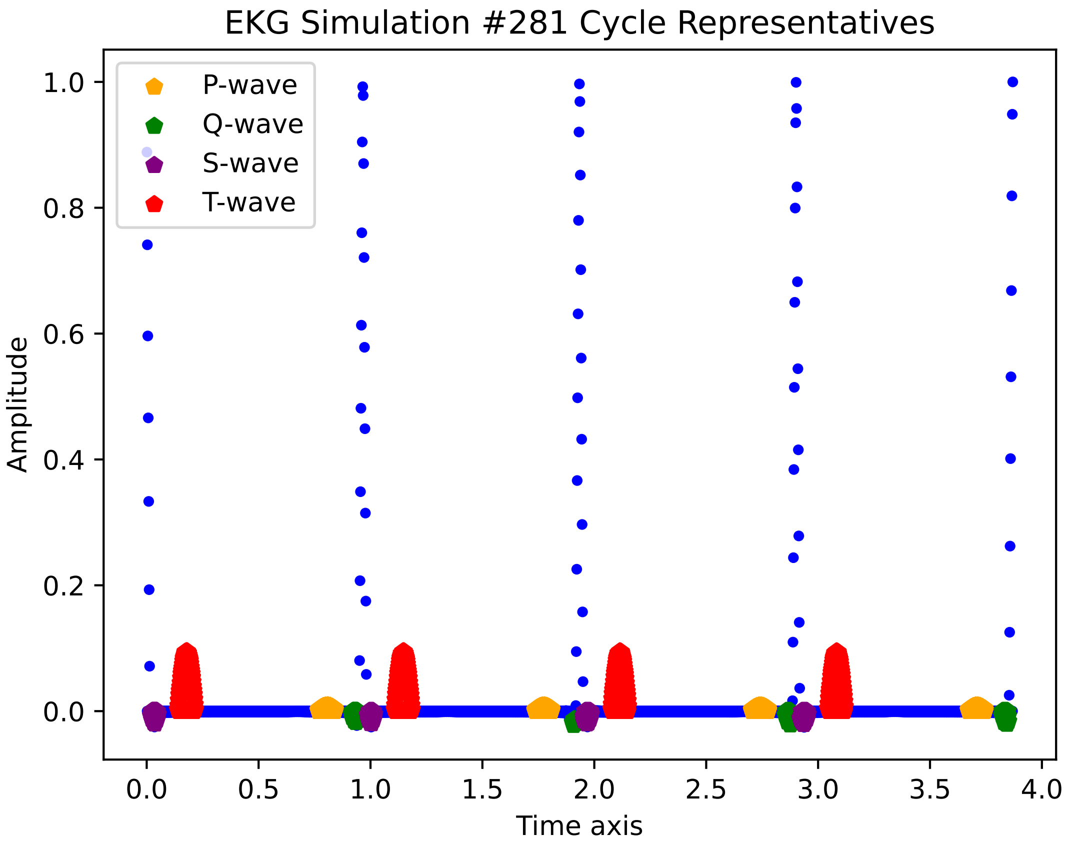

EKG Simulation #281 Cycle Representatives
P-wave
Q-wave
S-wave
T-wave
Amplitude
1.0
0.8
0.6
0.4
0.2
0.0
0.0
0.5
1.0
1.5
2.0
2.5
3.0
3.5
4.0
Time axis

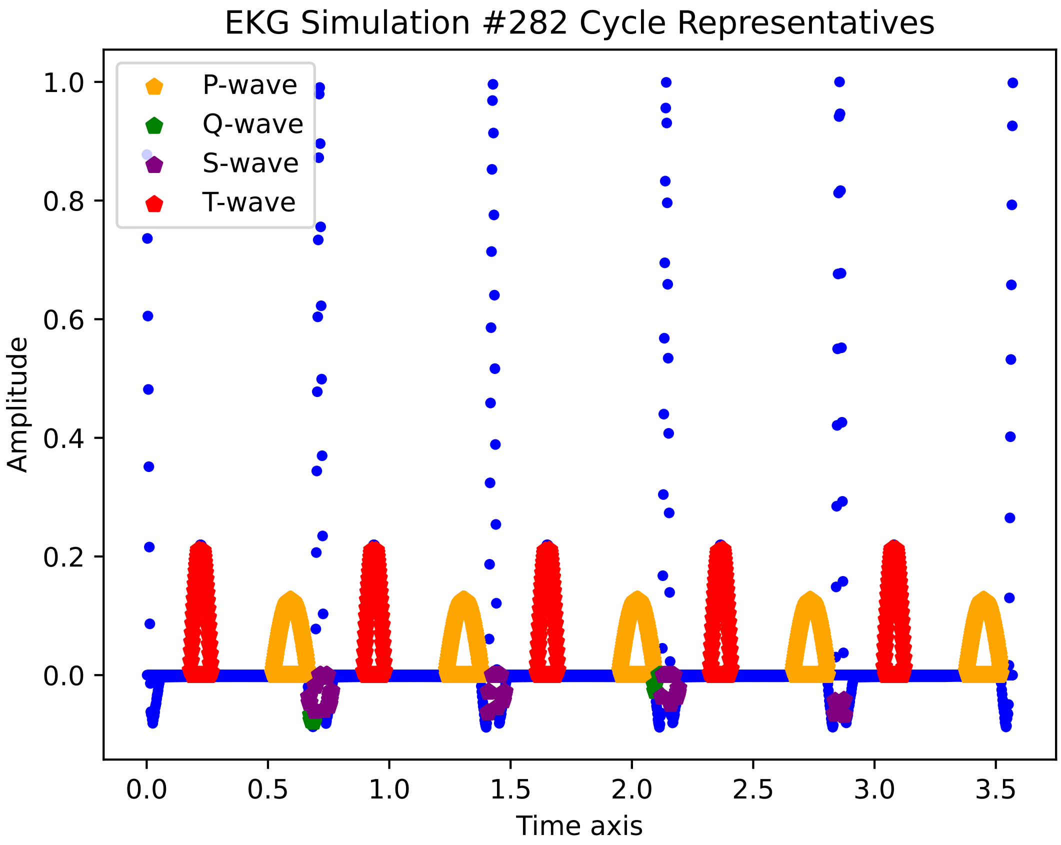

EKG Simulation #282 Cycle Representatives
P-wave
Q-wave
S-wave
T-wave
Amplitude
Time axis
1.0
0.8
0.6
0.4
0.2
0.0
0.0
0.5
1.0
1.5
2.0
2.5
3.0
3.5

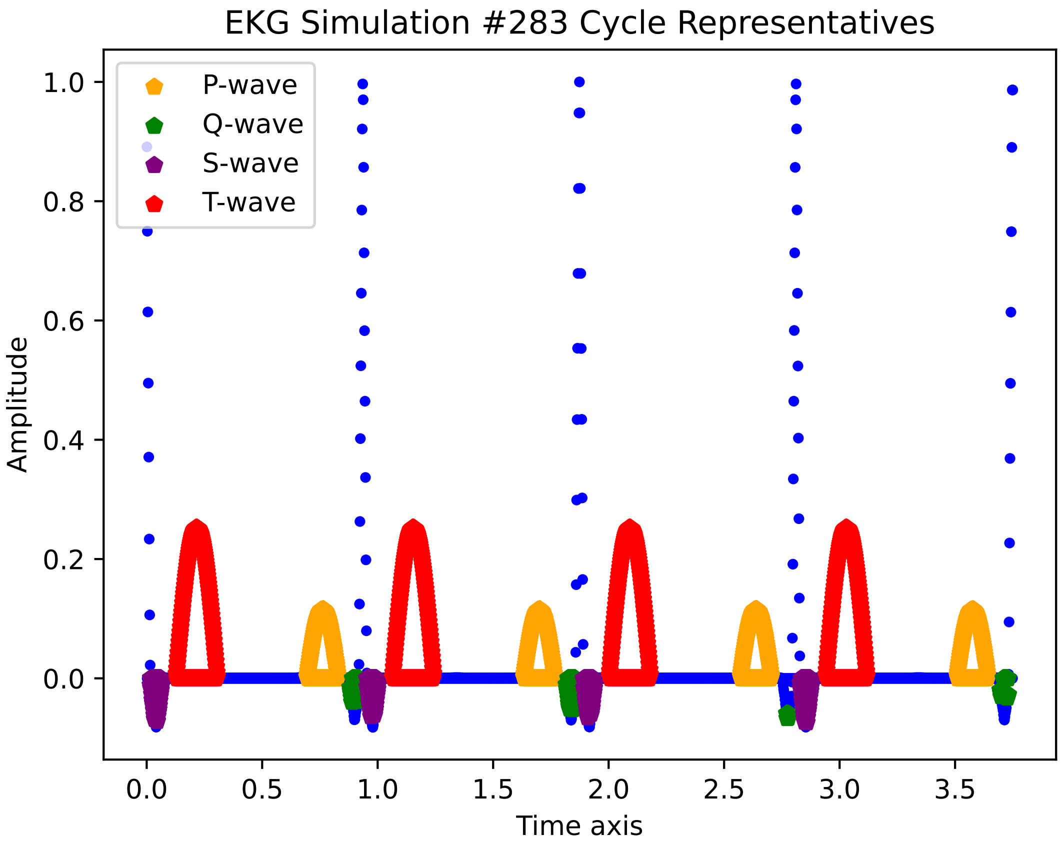
EKG Simulation #283 Cycle Representatives
P-wave
Q-wave
S-wave
T-wave
1.0
0.8
0.6
0.4
0.2
0.0
Amplitude
0.0
0.5
1.0
1.5
2.0
2.5
3.0
3.5
Time axis

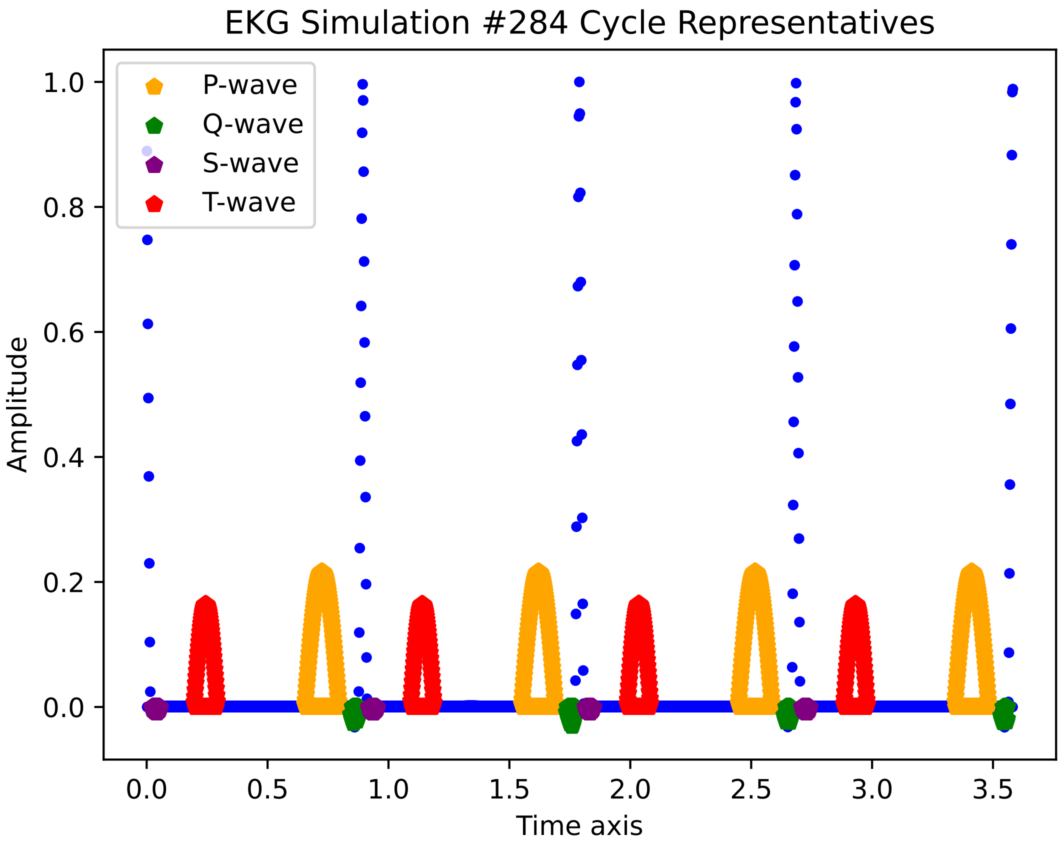
EKG Simulation #284 Cycle Representatives
P-wave
Q-wave
S-wave
T-wave
Amplitude
Time axis
0.0
0.2
0.4
0.6
0.8
1.0
0.0
0.5
1.0
1.5
2.0
2.5
3.0
3.5

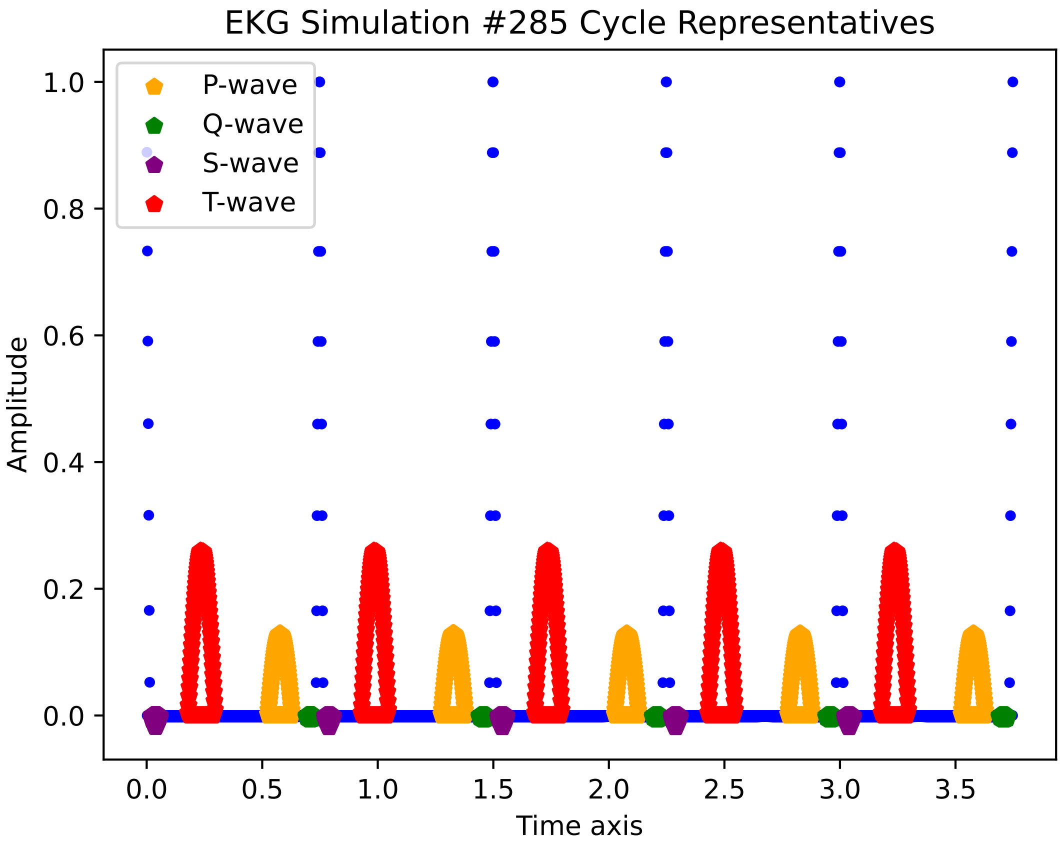

EKG Simulation #285 Cycle Representatives
P-wave
Q-wave
S-wave
T-wave
Amplitude
Time axis
0.0
0.2
0.4
0.6
0.8
1.0
0.0
0.5
1.0
1.5
2.0
2.5
3.0
3.5

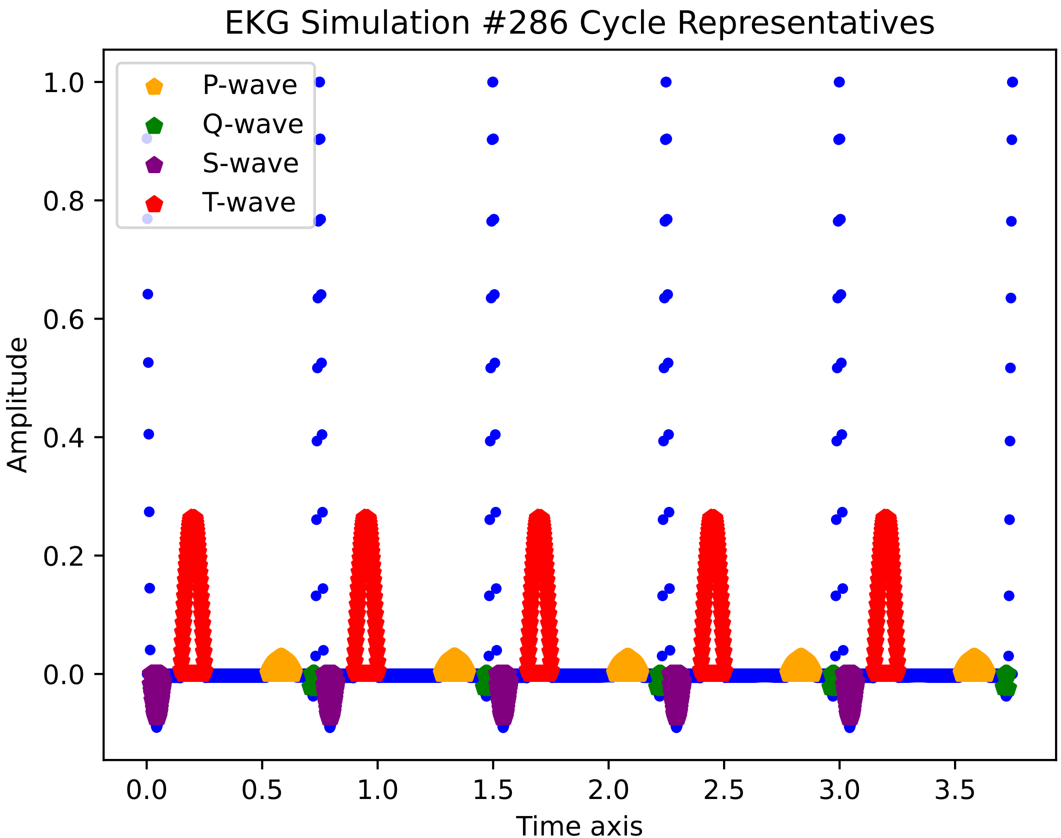
EKG Simulation #286 Cycle Representatives
P-wave
Q-wave
S-wave
T-wave
Amplitude
Time axis
1.0
0.8
0.6
0.4
0.2
0.0
0.0
0.5
1.0
1.5
2.0
2.5
3.0
3.5

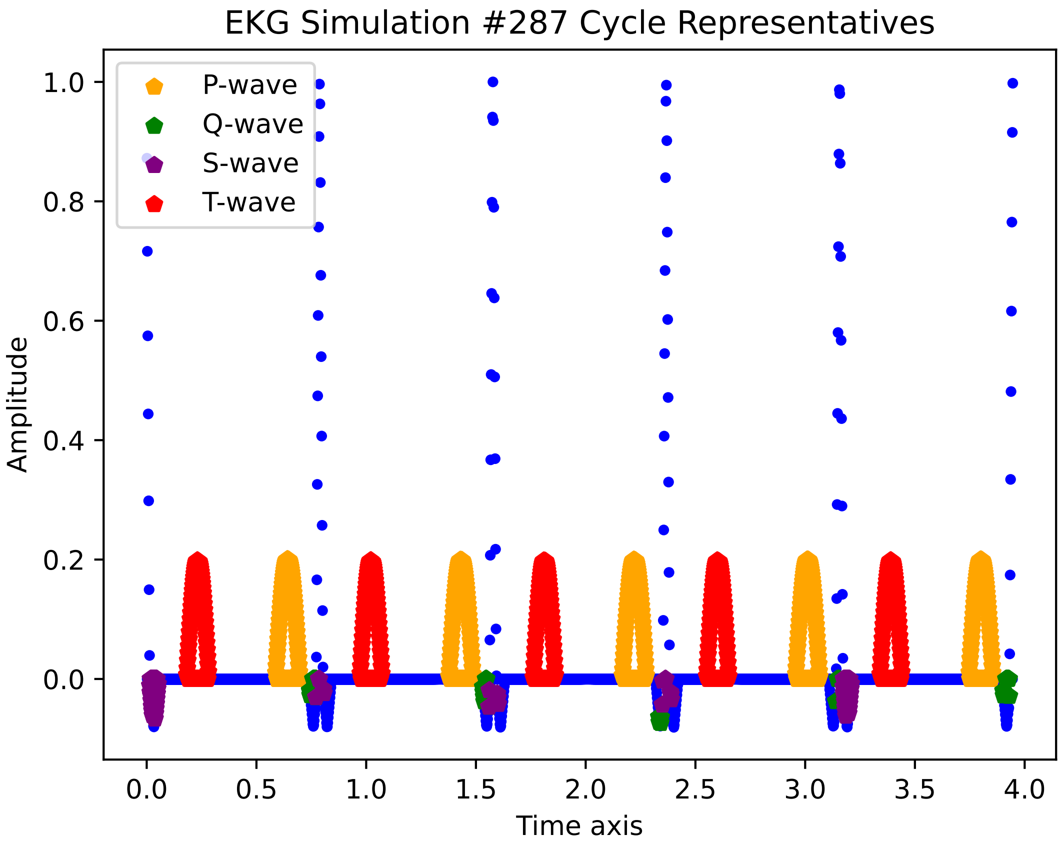

EKG Simulation #287 Cycle Representatives
P-wave
Q-wave
S-wave
T-wave
Amplitude
1.0
0.8
0.6
0.4
0.2
0.0
0.0
0.5
1.0
1.5
2.0
2.5
3.0
3.5
4.0
Time axis

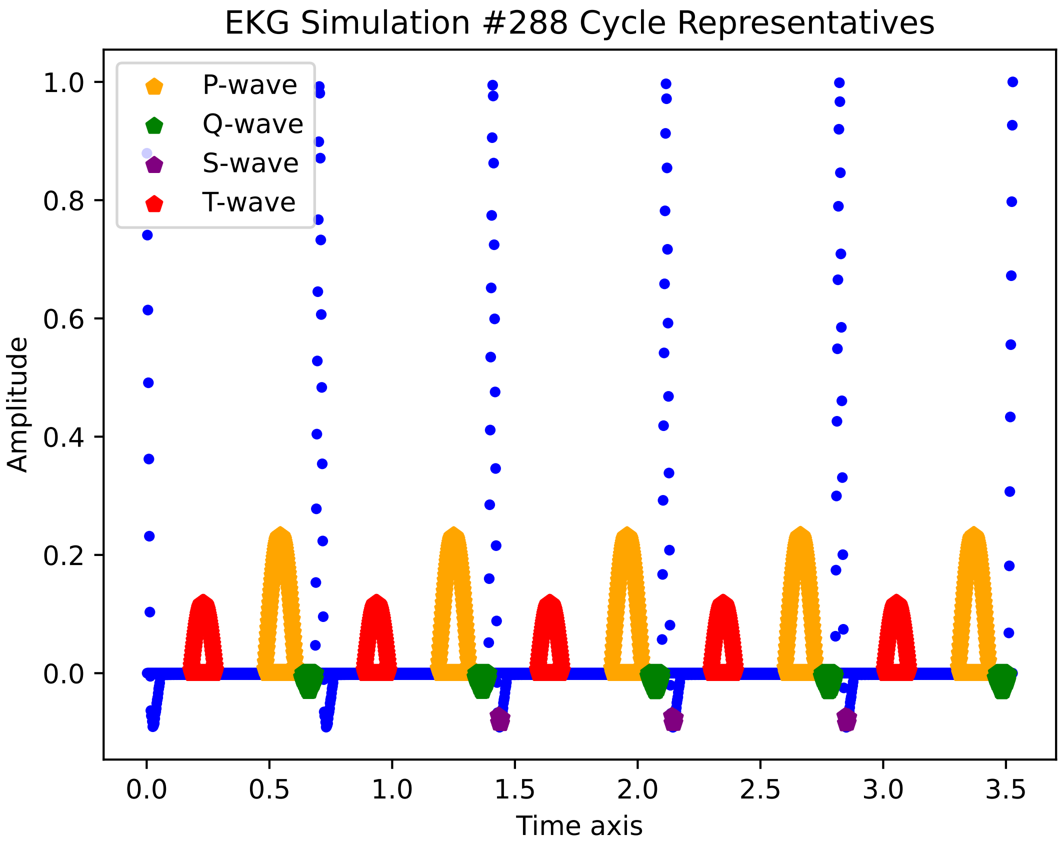

EKG Simulation #288 Cycle Representatives
P-wave
Q-wave
S-wave
T-wave
Amplitude
Time axis
1.0
0.8
0.6
0.4
0.2
0.0
0.0
0.5
1.0
1.5
2.0
2.5
3.0
3.5

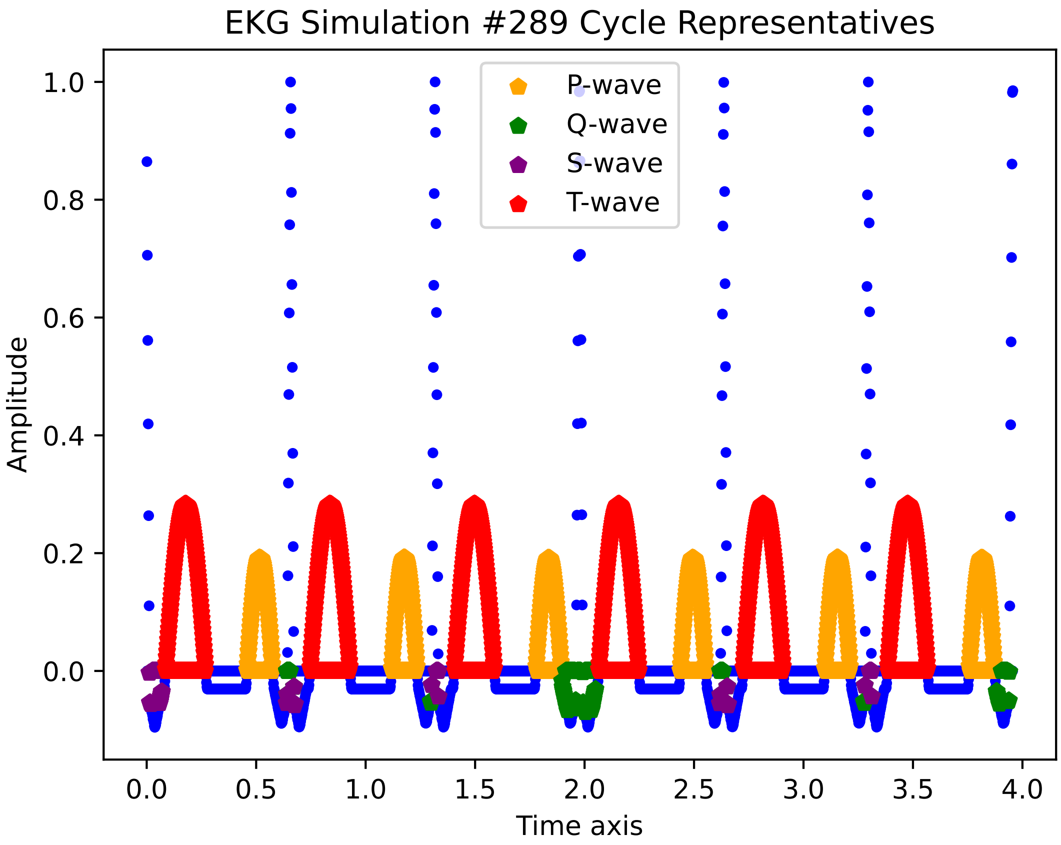

EKG Simulation #289 Cycle Representatives
P-wave
Q-wave
S-wave
T-wave
Amplitude
Time axis
1.0
0.8
0.6
0.4
0.2
0.0
0.0
0.5
1.0
1.5
2.0
2.5
3.0
3.5
4.0

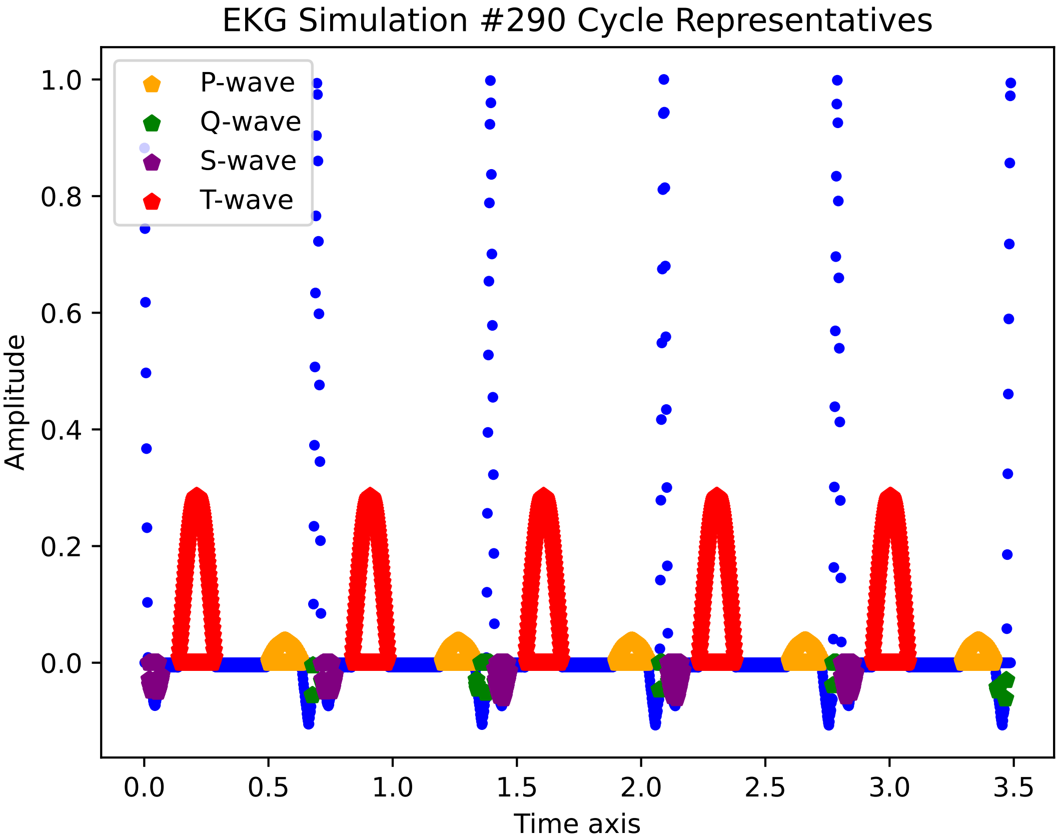

EKG Simulation #290 Cycle Representatives
P-wave
Q-wave
S-wave
T-wave
Amplitude
1.0
0.8
0.6
0.4
0.2
0.0
0.0
0.5
1.0
1.5
2.0
2.5
3.0
3.5
Time axis

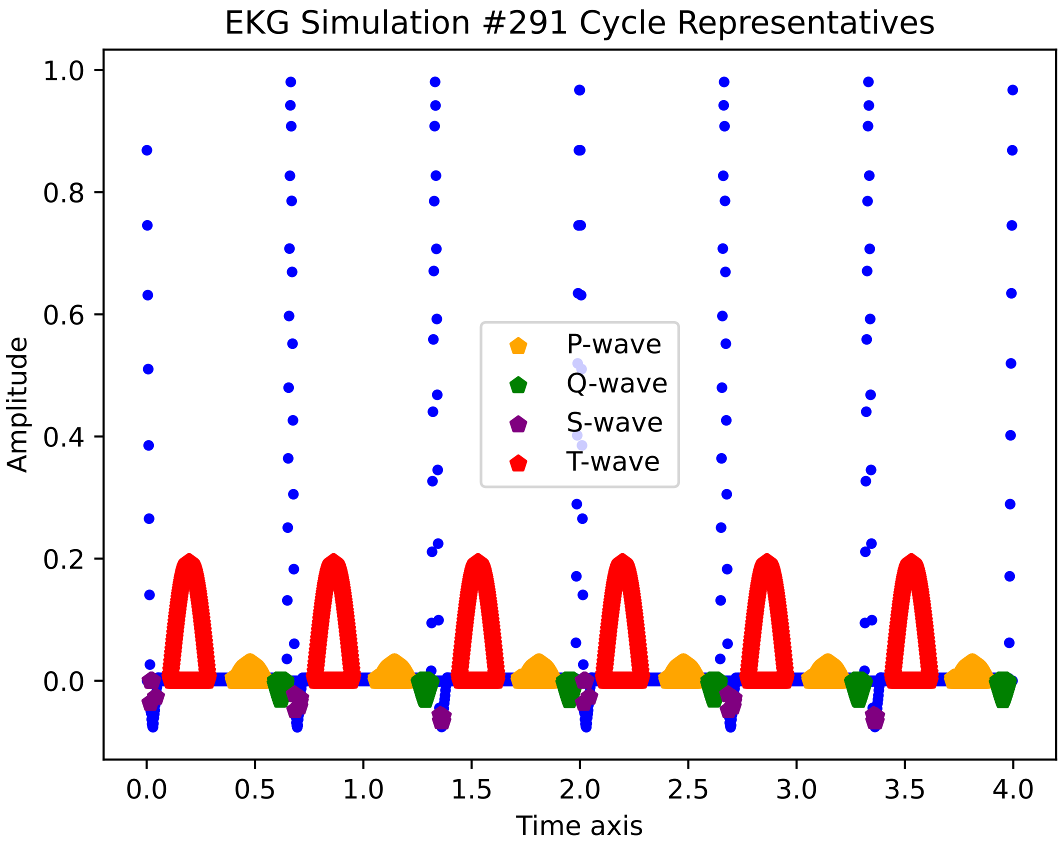

EKG Simulation #291 Cycle Representatives
Amplitude
Time axis
P-wave
Q-wave
S-wave
T-wave
1.0
0.8
0.6
0.4
0.2
0.0
0.0
0.5
1.0
1.5
2.0
2.5
3.0
3.5
4.0

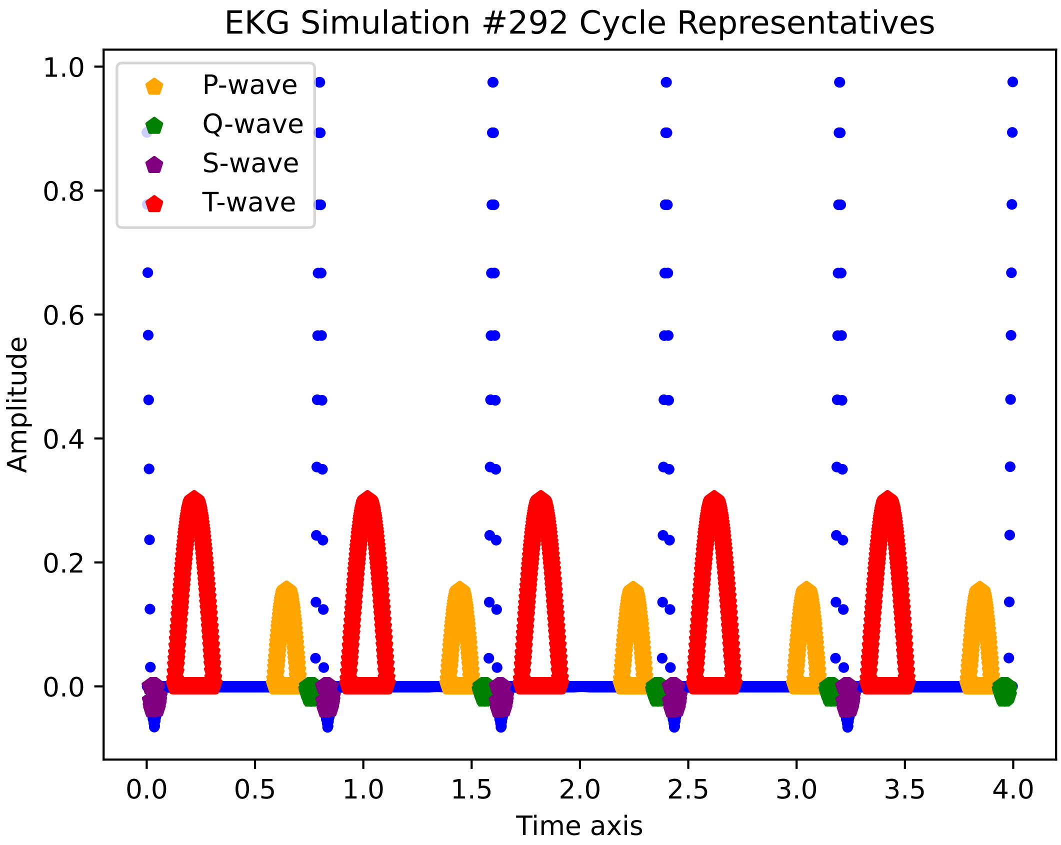
EKG Simulation #292 Cycle Representatives
P-wave
Q-wave
S-wave
T-wave
Amplitude
Time axis
1.0
0.8
0.6
0.4
0.2
0.0
0.0
0.5
1.0
1.5
2.0
2.5
3.0
3.5
4.0

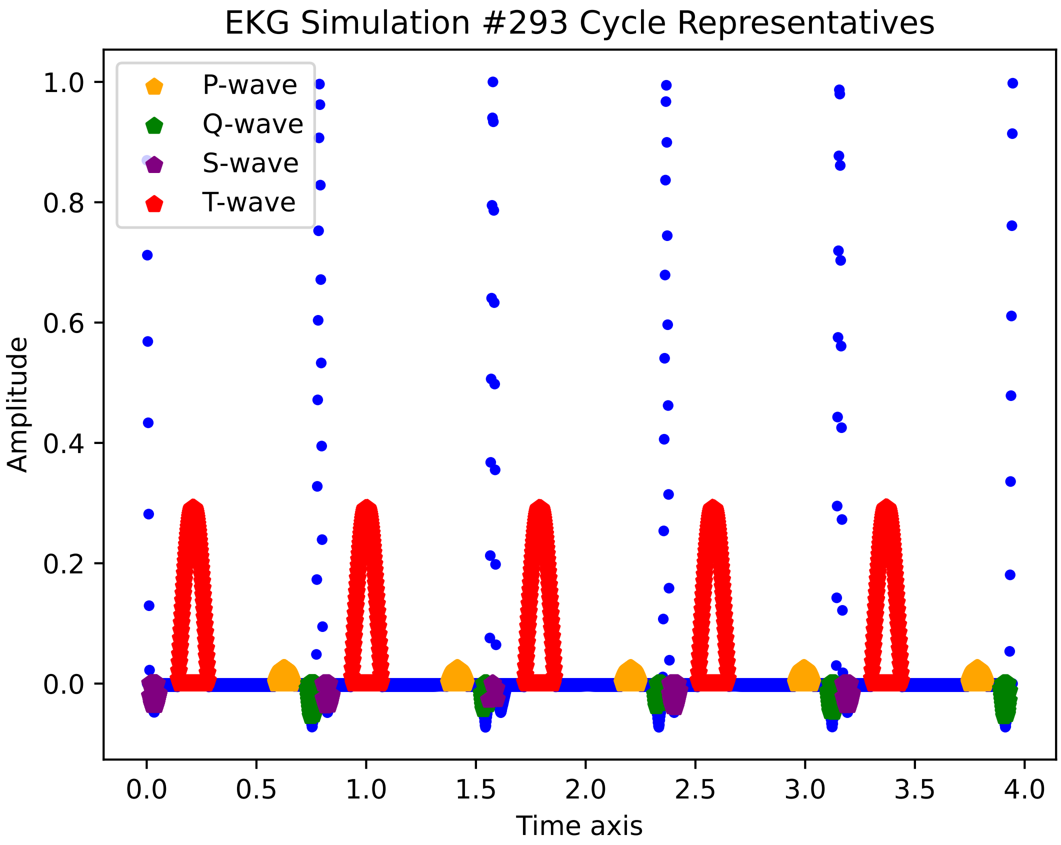
EKG Simulation #293 Cycle Representatives
P-wave
Q-wave
S-wave
T-wave
1.0
0.8
0.6
0.4
0.2
0.0
Amplitude
0.0
0.5
1.0
1.5
2.0
2.5
3.0
3.5
4.0
Time axis

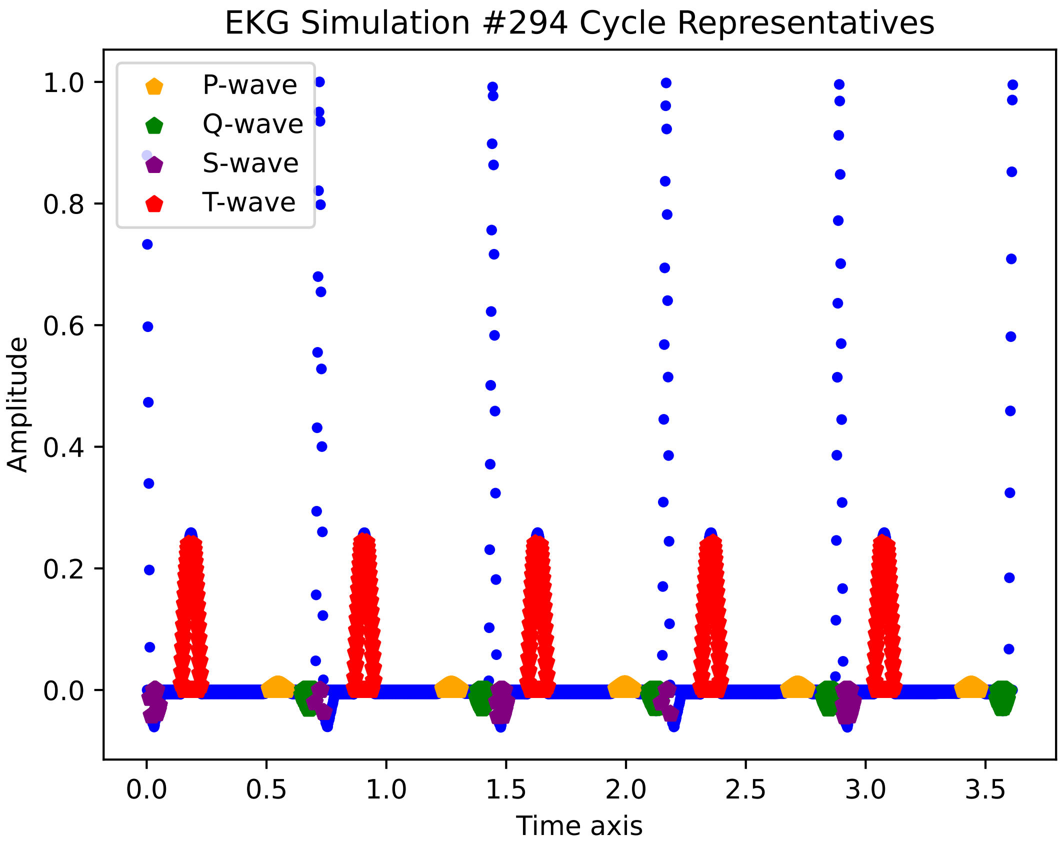
EKG Simulation #294 Cycle Representatives
P-wave
Q-wave
S-wave
T-wave
Amplitude
Time axis
1.0
0.8
0.6
0.4
0.2
0.0
0.0
0.5
1.0
1.5
2.0
2.5
3.0
3.5

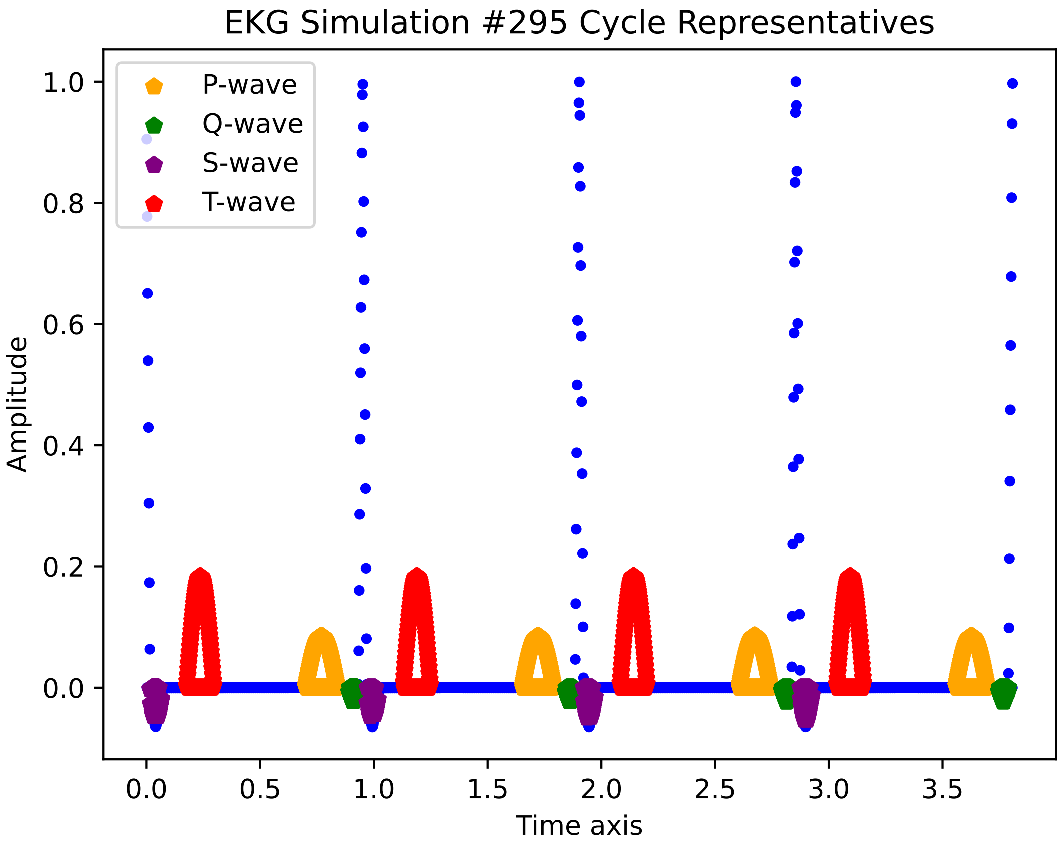

EKG Simulation #295 Cycle Representatives
P-wave
Q-wave
S-wave
T-wave
Amplitude
Time axis
1.0
0.8
0.6
0.4
0.2
0.0
0.0
0.5
1.0
1.5
2.0
2.5
3.0
3.5

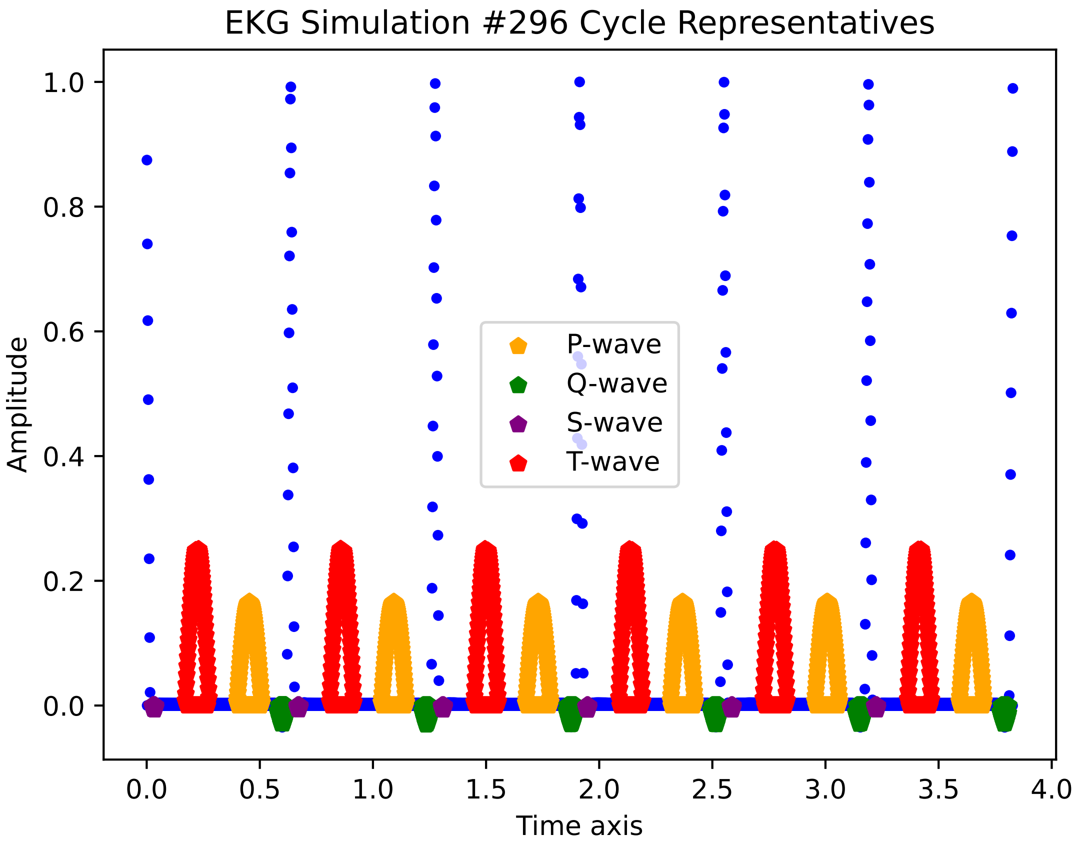

EKG Simulation #296 Cycle Representatives
Amplitude
Time axis
1.0
0.8
0.6
0.4
0.2
0.0
0.0
0.5
1.0
1.5
2.0
2.5
3.0
3.5
4.0
P-wave
Q-wave
S-wave
T-wave

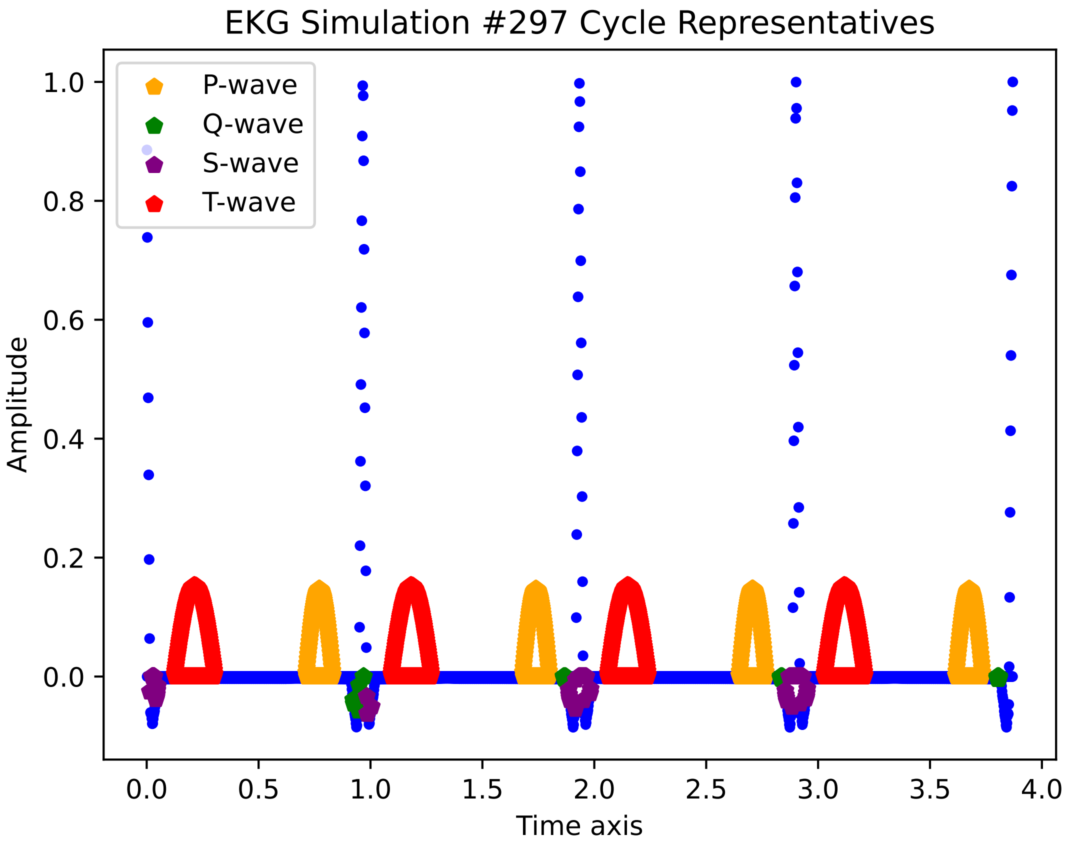
EKG Simulation #297 Cycle Representatives
P-wave
Q-wave
S-wave
T-wave
1.0
0.8
0.6
0.4
0.2
0.0
Amplitude
0.0
0.5
1.0
1.5
2.0
2.5
3.0
3.5
4.0
Time axis

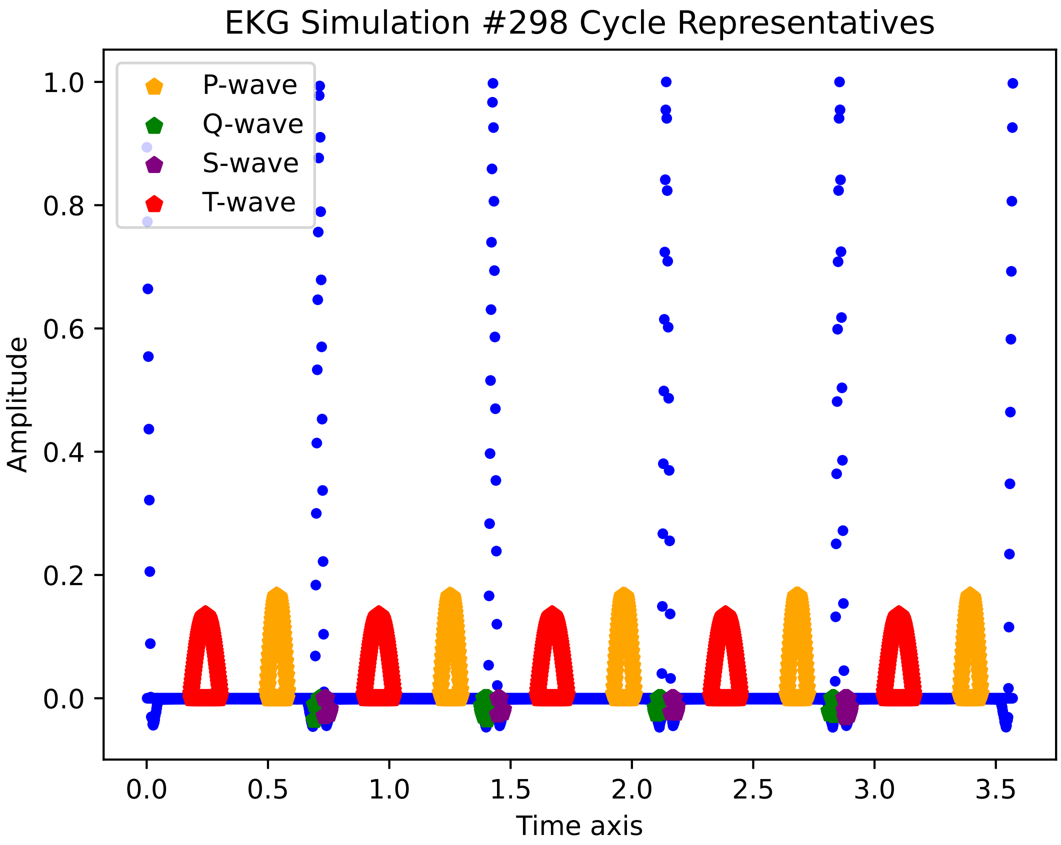

EKG Simulation #298 Cycle Representatives
P-wave
Q-wave
S-wave
T-wave
Amplitude
Time axis
1.0
0.8
0.6
0.4
0.2
0.0
0.0
0.5
1.0
1.5
2.0
2.5
3.0
3.5

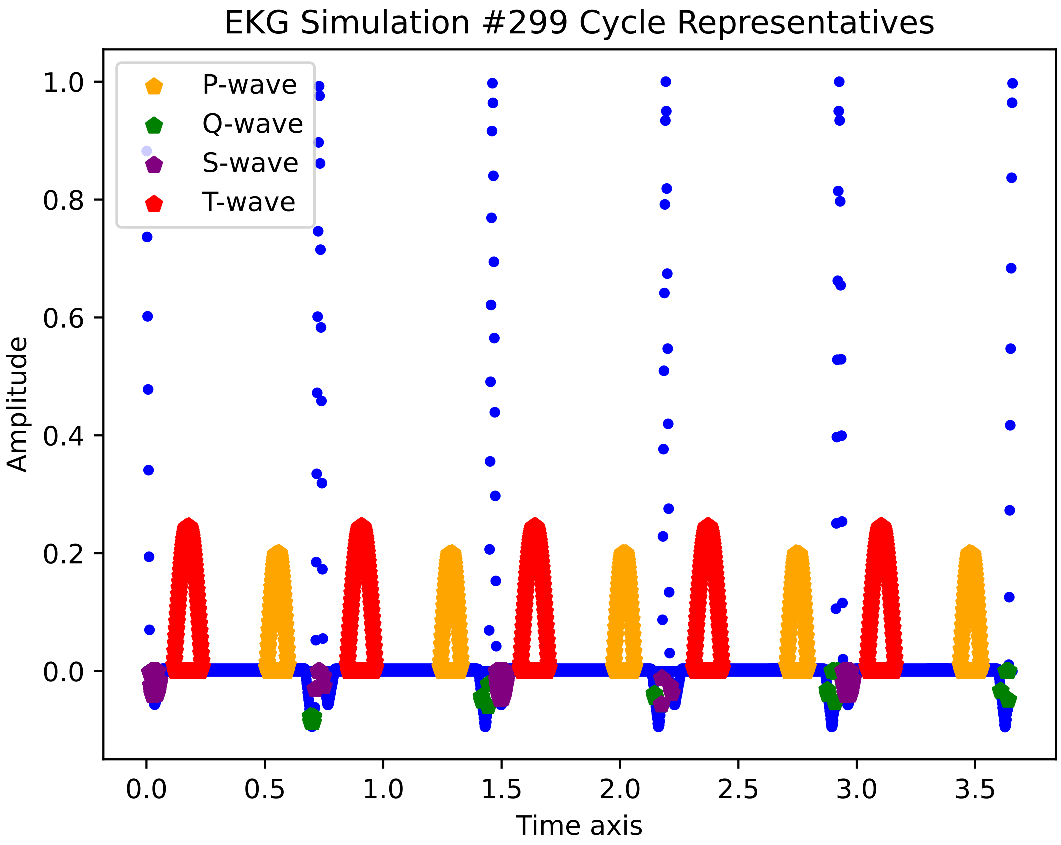
EKG Simulation #299 Cycle Representatives
P-wave
Q-wave
S-wave
T-wave
Amplitude
Time axis
1.0
0.8
0.6
0.4
0.2
0.0
0.0
0.5
1.0
1.5
2.0
2.5
3.0
3.5

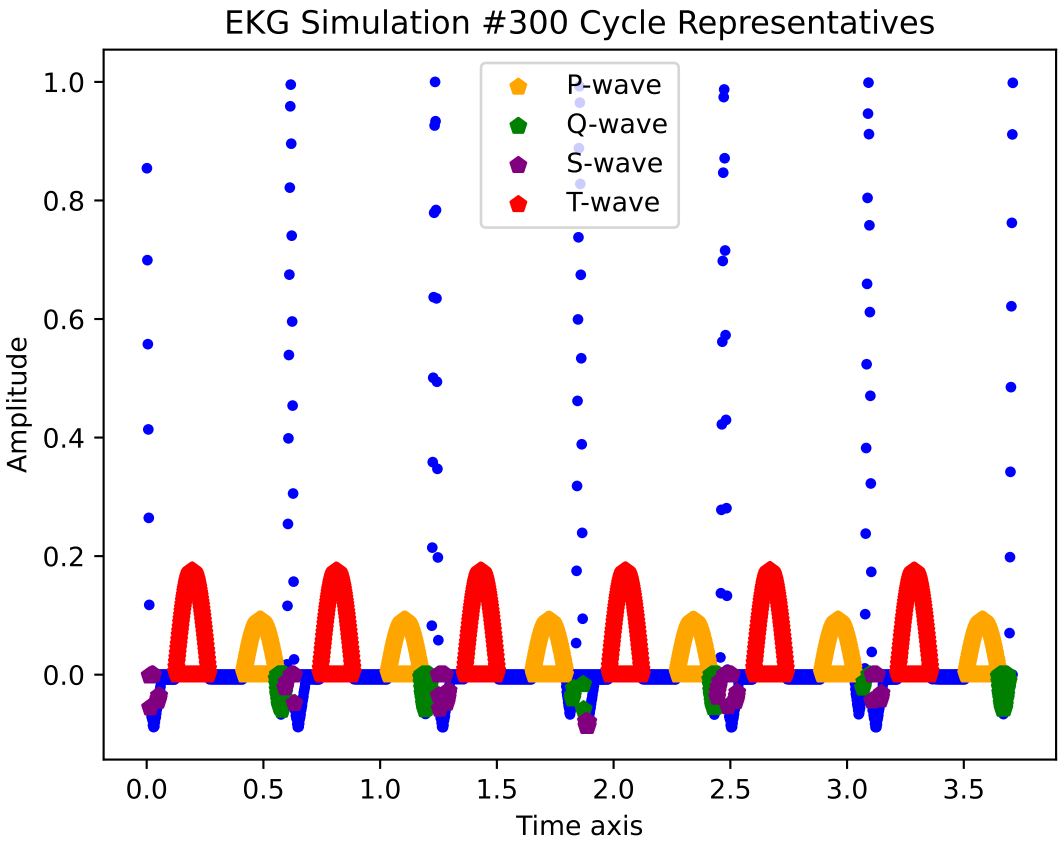

EKG Simulation #300 Cycle Representatives
P-wave
Q-wave
S-wave
T-wave
Amplitude
Time axis
1.0
0.8
0.6
0.4
0.2
0.0
0.0
0.5
1.0
1.5
2.0
2.5
3.0
3.5

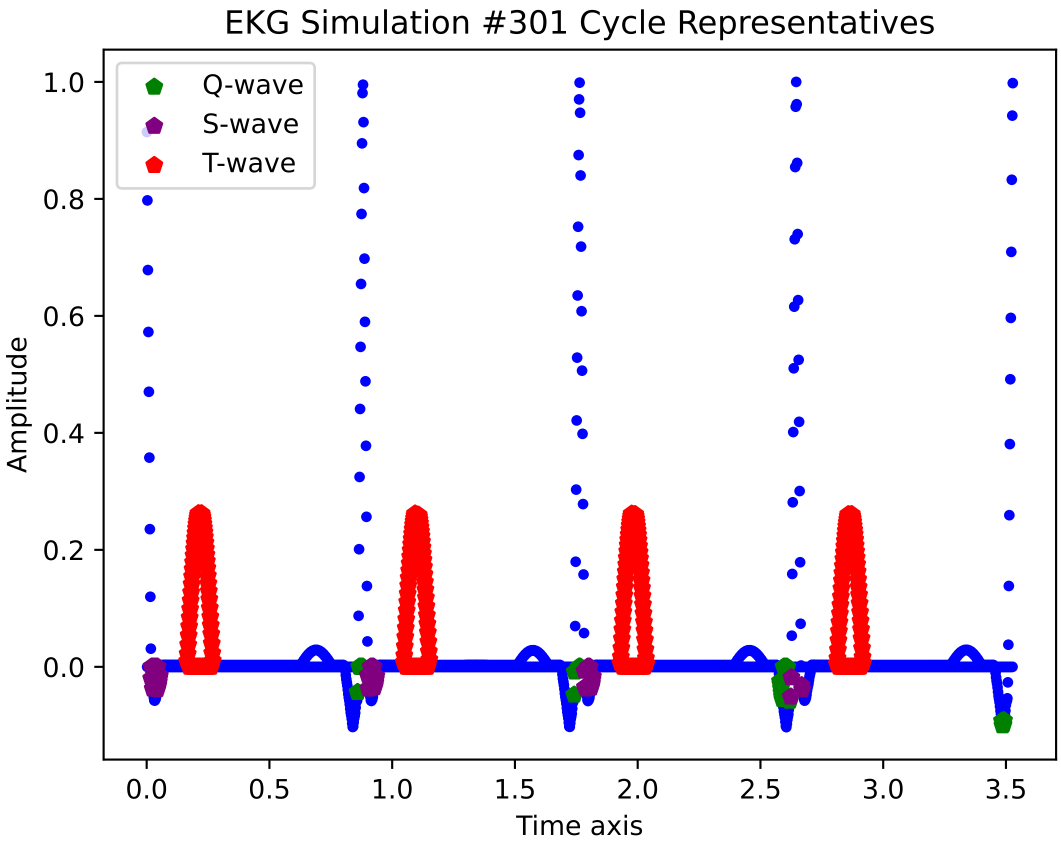

EKG Simulation #301 Cycle Representatives
Q-wave
S-wave
T-wave
1.0
0.8
0.6
0.4
0.2
0.0
Amplitude
0.0
0.5
1.0
1.5
2.0
2.5
3.0
3.5
Time axis

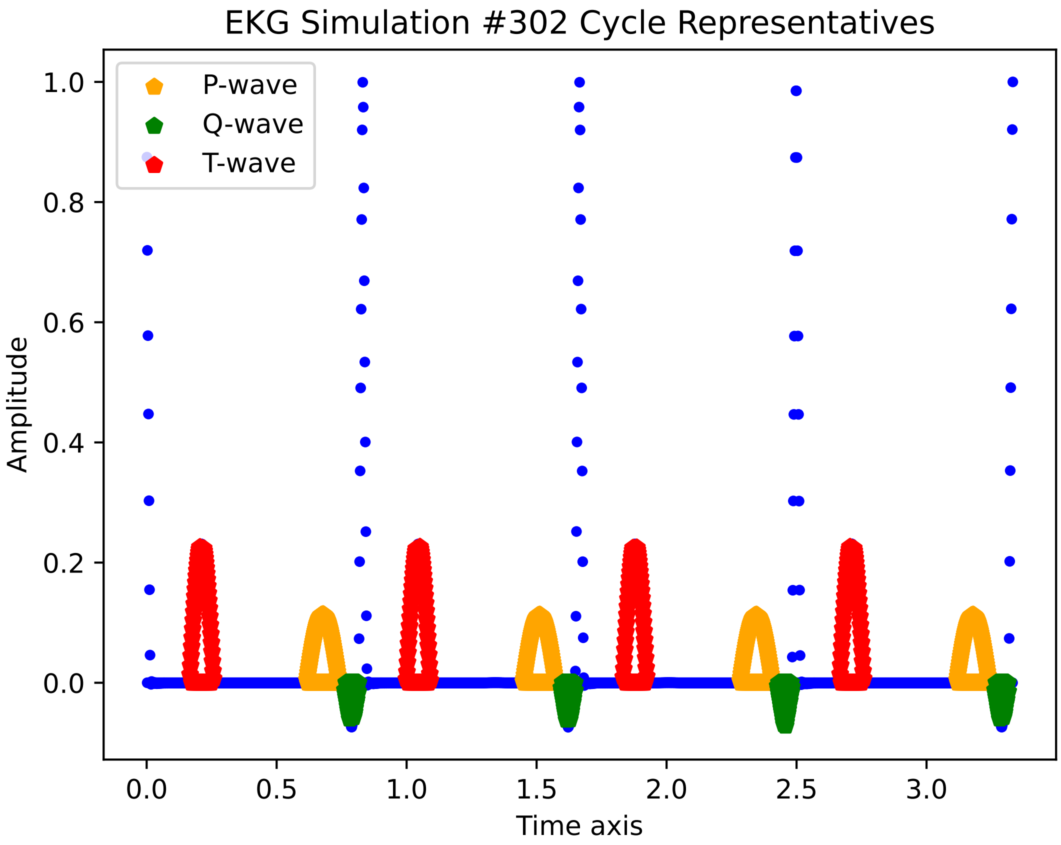

EKG Simulation #302 Cycle Representatives
P-wave
Q-wave
T-wave
Amplitude
Time axis
1.0
0.8
0.6
0.4
0.2
0.0
0.0
0.5
1.0
1.5
2.0
2.5
3.0

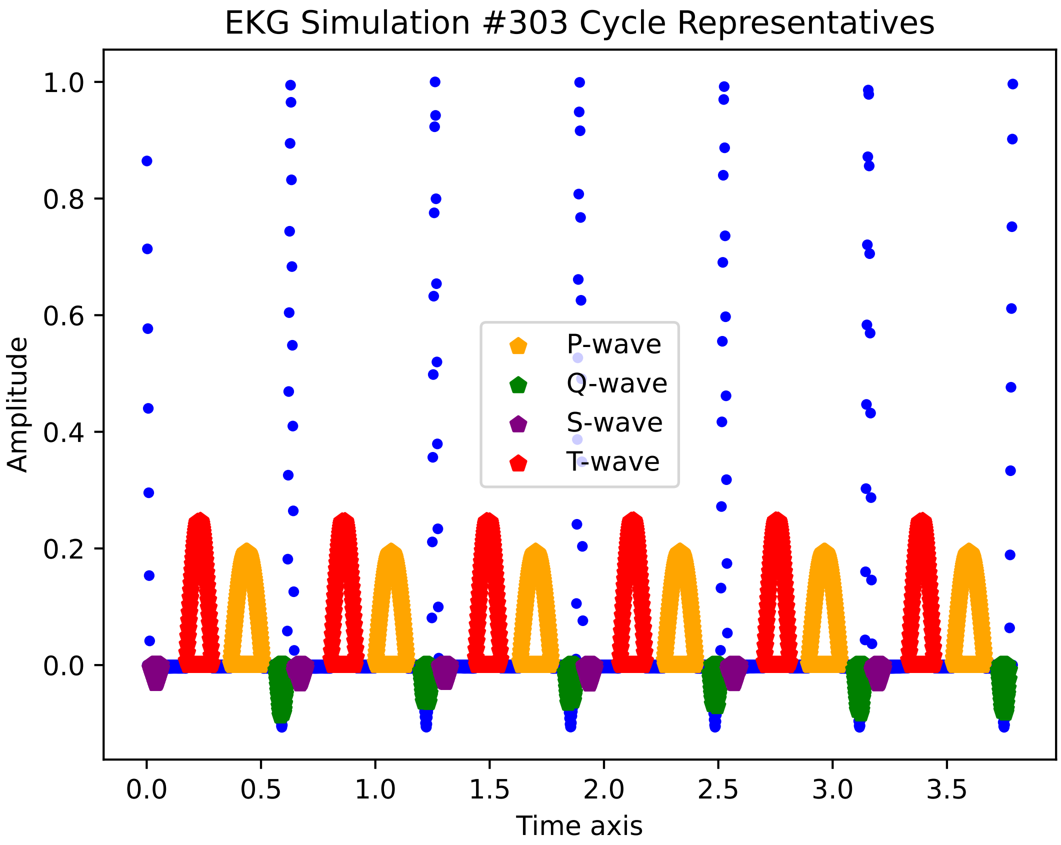

EKG Simulation #303 Cycle Representatives
Amplitude
Time axis
1.0
0.8
0.6
0.4
0.2
0.0
0.0
0.5
1.0
1.5
2.0
2.5
3.0
3.5
P-wave
Q-wave
S-wave
T-wave

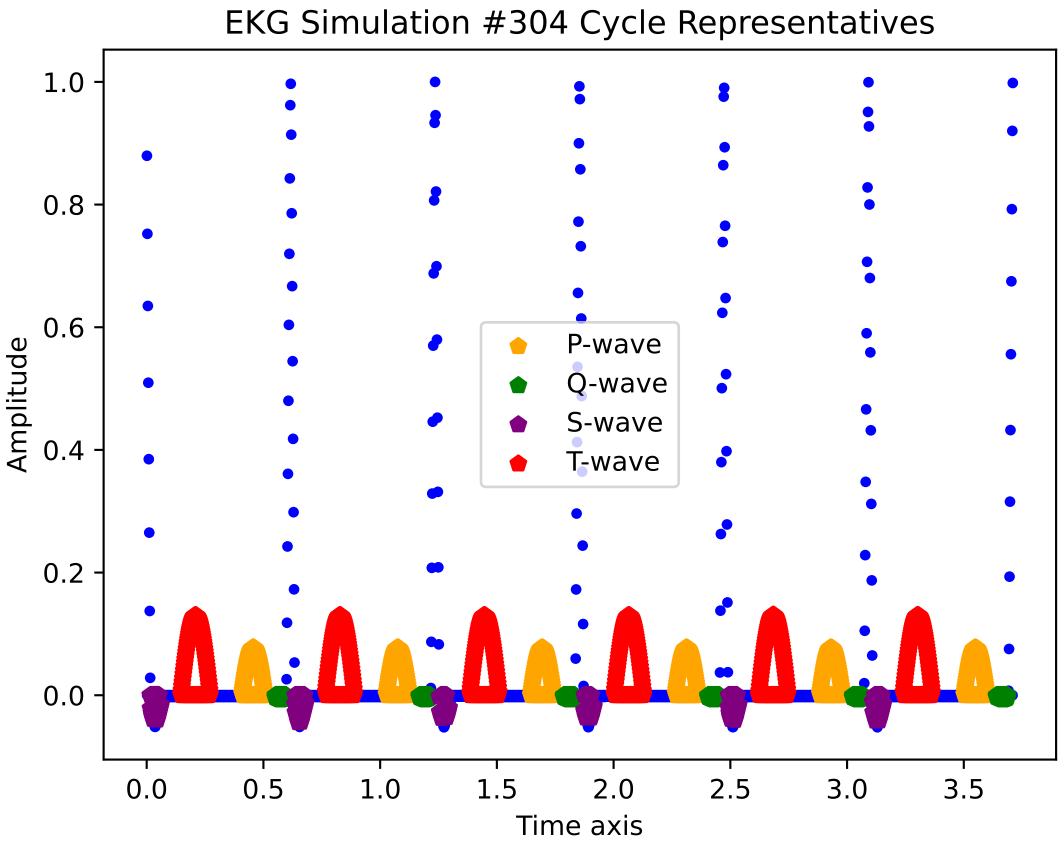
EKG Simulation #304 Cycle Representatives
P-wave
Q-wave
S-wave
T-wave
Amplitude
Time axis
0.0
0.2
0.4
0.6
0.8
1.0
0.0
0.5
1.0
1.5
2.0
2.5
3.0
3.5

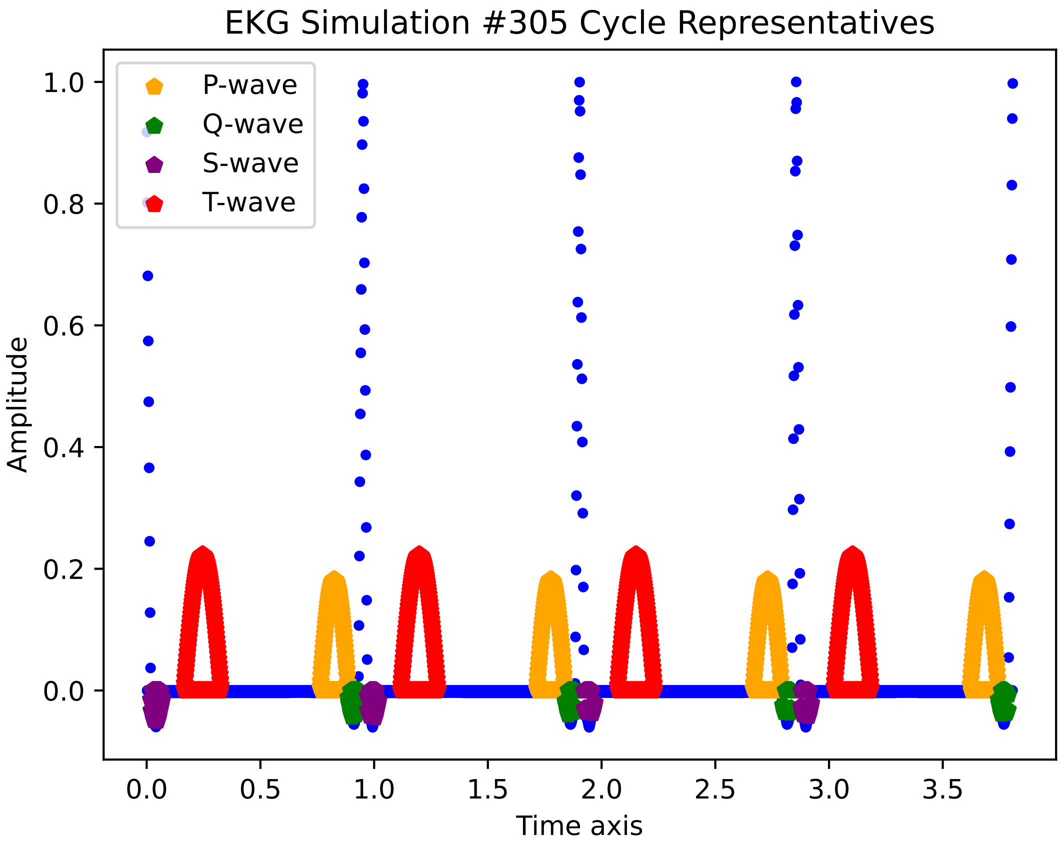

EKG Simulation #305 Cycle Representatives
P-wave
Q-wave
S-wave
T-wave
Amplitude
Time axis
1.0
0.8
0.6
0.4
0.2
0.0
0.0
0.5
1.0
1.5
2.0
2.5
3.0
3.5

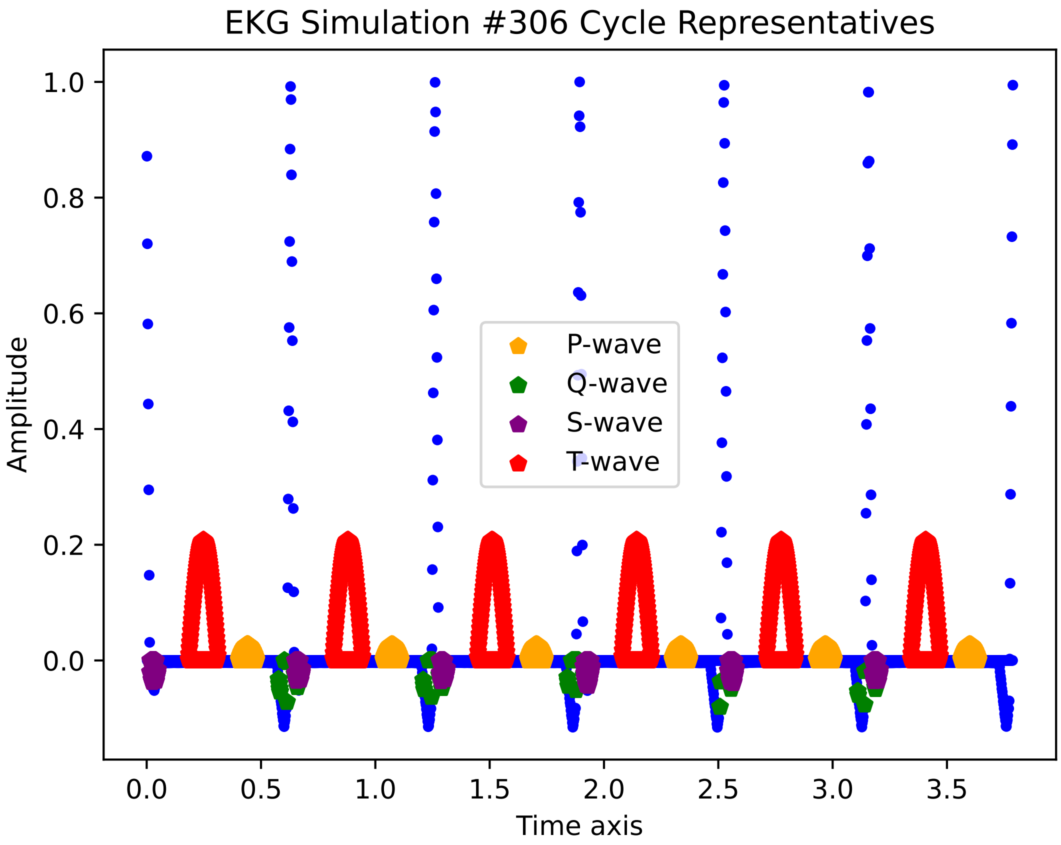

EKG Simulation #306 Cycle Representatives
P-wave
Q-wave
S-wave
T-wave
Amplitude
Time axis
1.0
0.8
0.6
0.4
0.2
0.0
0.0
0.5
1.0
1.5
2.0
2.5
3.0
3.5

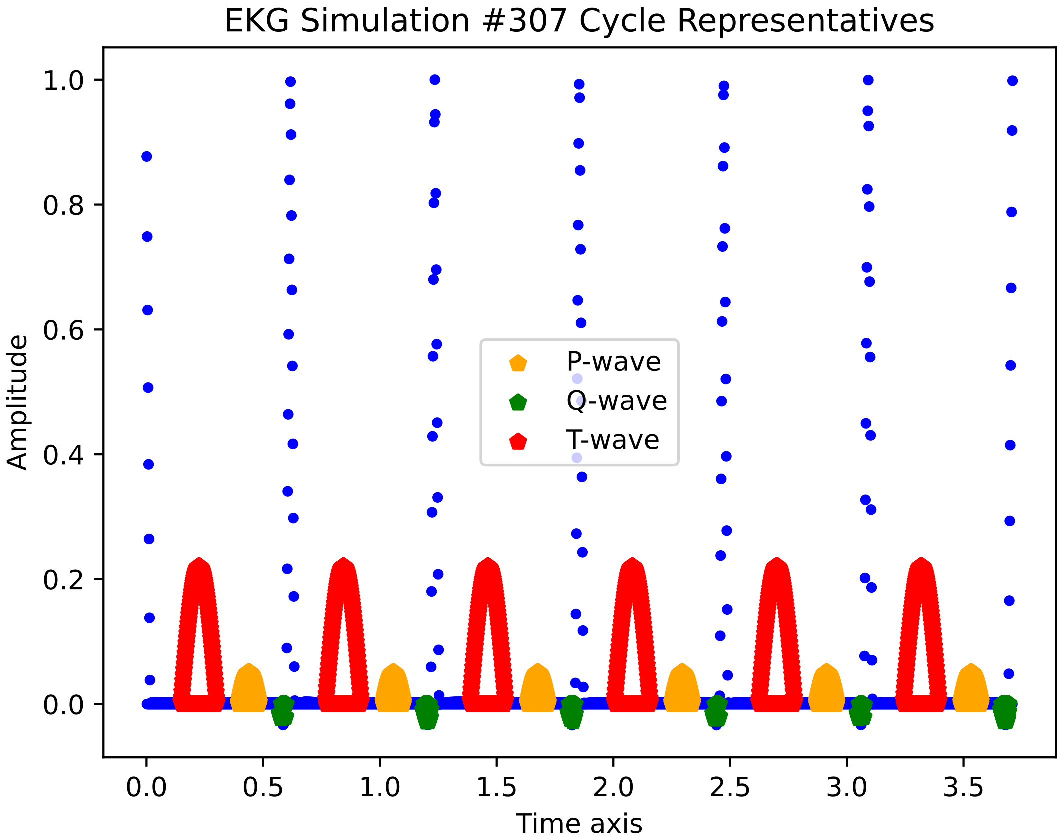

EKG Simulation #307 Cycle Representatives
Amplitude
Time axis
P-wave
Q-wave
T-wave
1.0
0.8
0.6
0.4
0.2
0.0
0.0
0.5
1.0
1.5
2.0
2.5
3.0
3.5

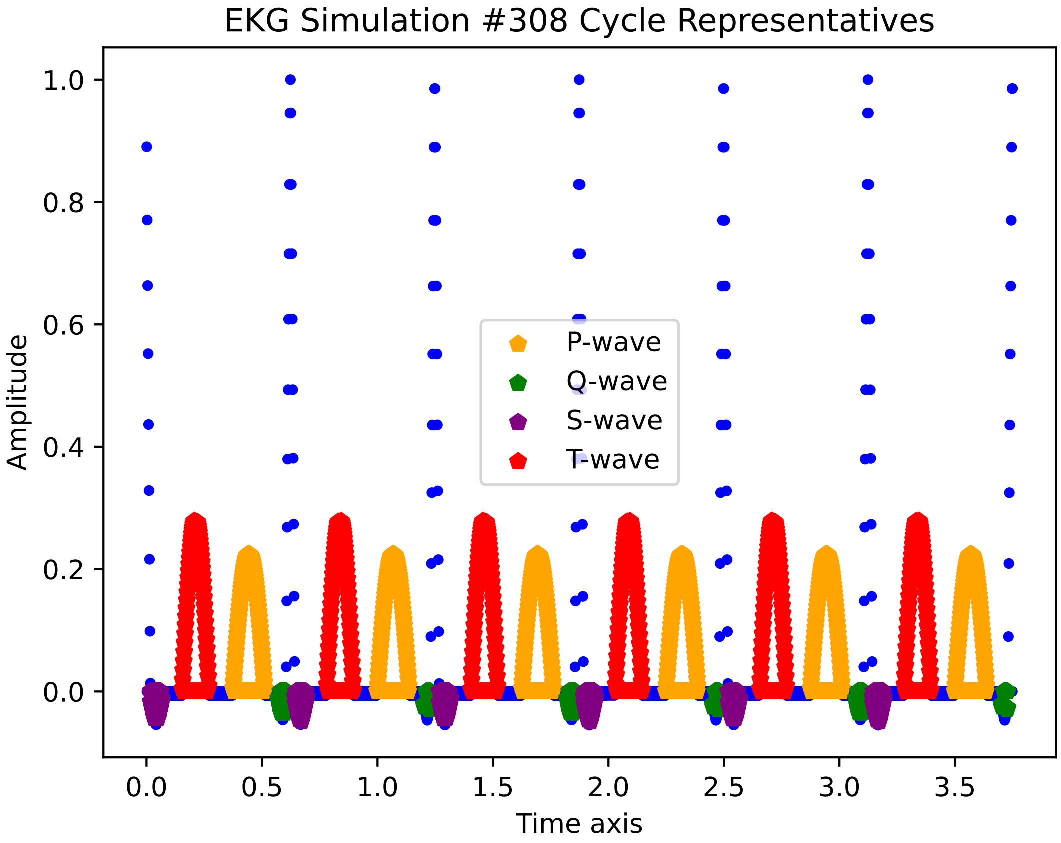
EKG Simulation #308 Cycle Representatives
Amplitude
Time axis
P-wave
Q-wave
S-wave
T-wave
1.0
0.8
0.6
0.4
0.2
0.0
0.0
0.5
1.0
1.5
2.0
2.5
3.0
3.5

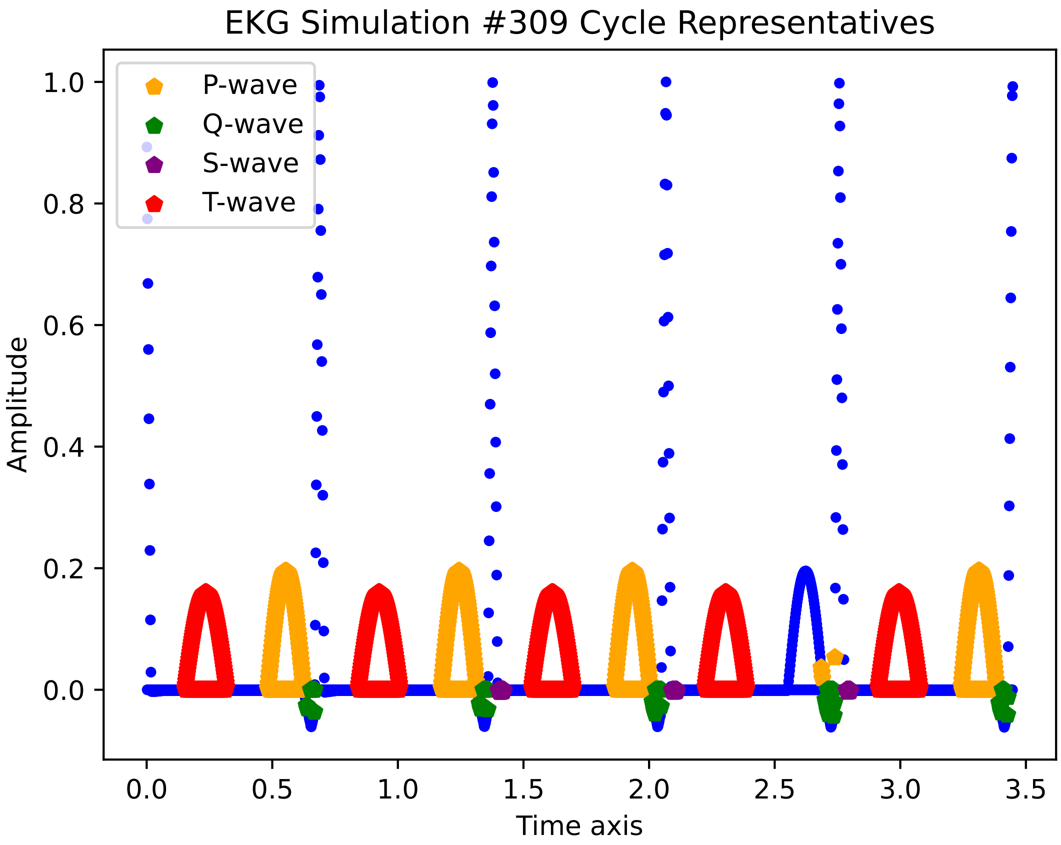
EKG Simulation #309 Cycle Representatives
P-wave
Q-wave
S-wave
T-wave
Amplitude
Time axis
1.0
0.8
0.6
0.4
0.2
0.0
0.0
0.5
1.0
1.5
2.0
2.5
3.0
3.5

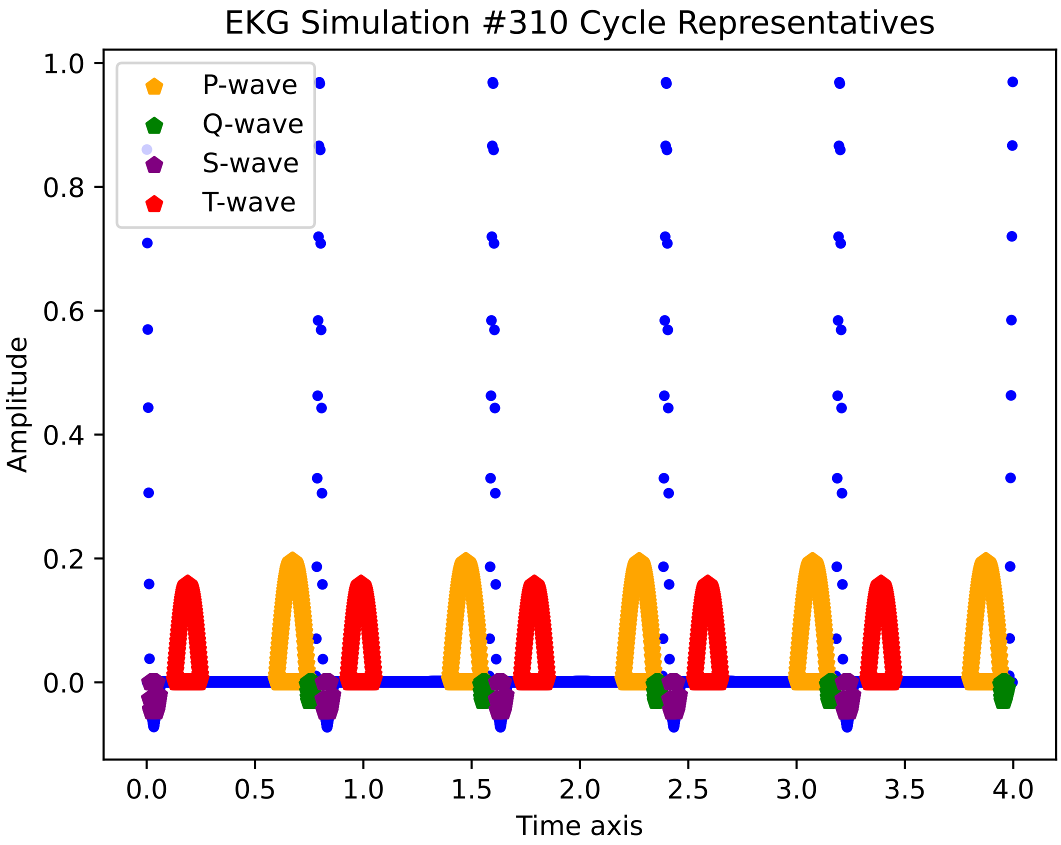

EKG Simulation #310 Cycle Representatives
P-wave
Q-wave
S-wave
T-wave
Amplitude
Time axis
1.0
0.8
0.6
0.4
0.2
0.0
0.0
0.5
1.0
1.5
2.0
2.5
3.0
3.5
4.0

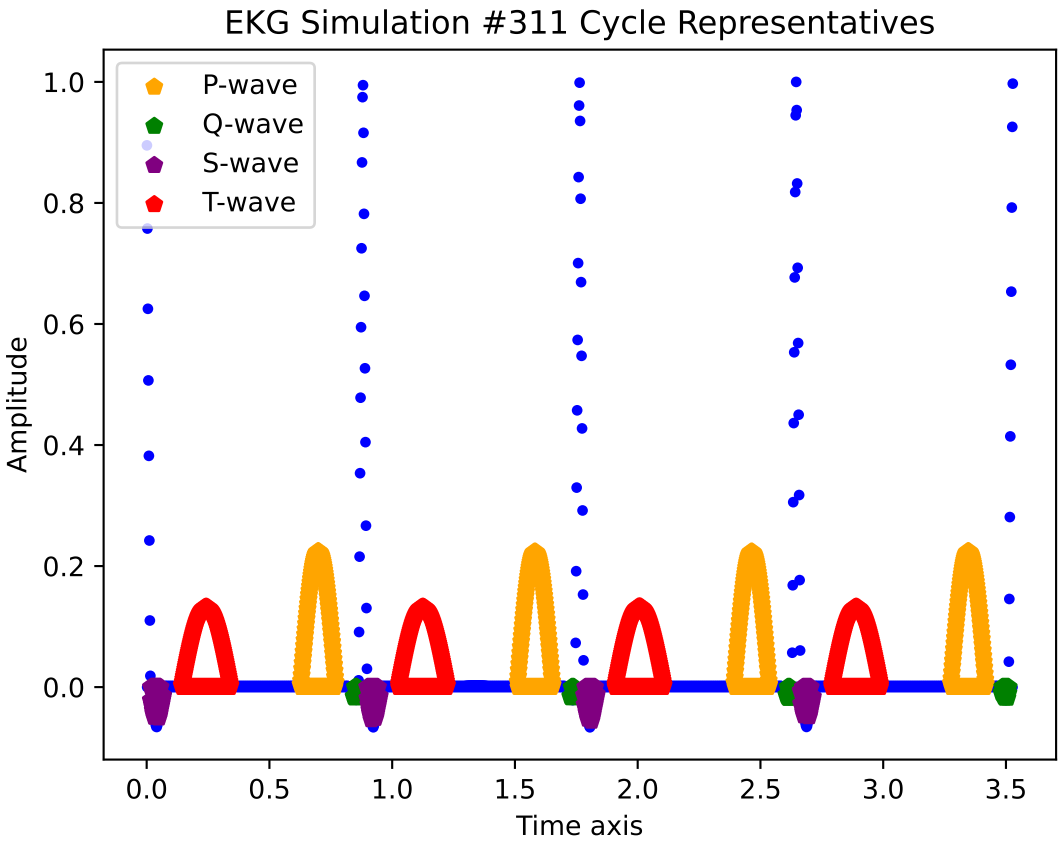
EKG Simulation #311 Cycle Representatives
P-wave
Q-wave
S-wave
T-wave
1.0
0.8
0.6
0.4
0.2
0.0
Amplitude
0.0
0.5
1.0
1.5
2.0
2.5
3.0
3.5
Time axis

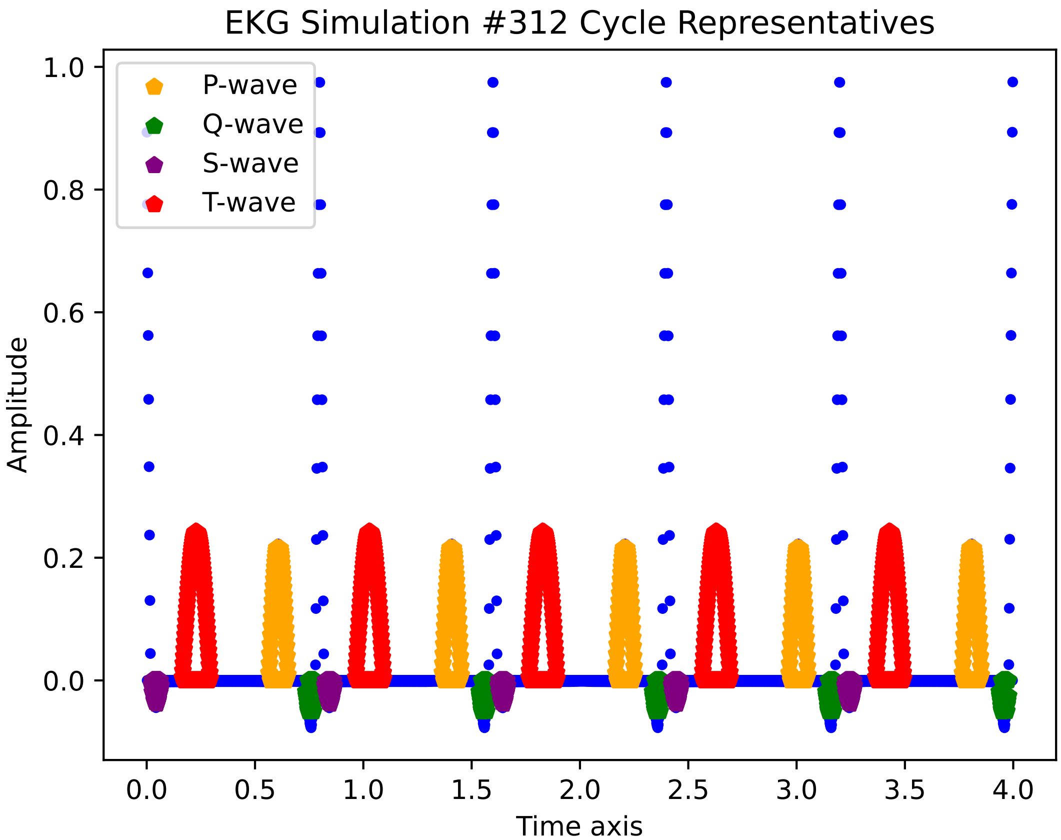
EKG Simulation #312 Cycle Representatives
P-wave
Q-wave
S-wave
T-wave
Amplitude
Time axis
1.0
0.8
0.6
0.4
0.2
0.0
0.0
0.5
1.0
1.5
2.0
2.5
3.0
3.5
4.0

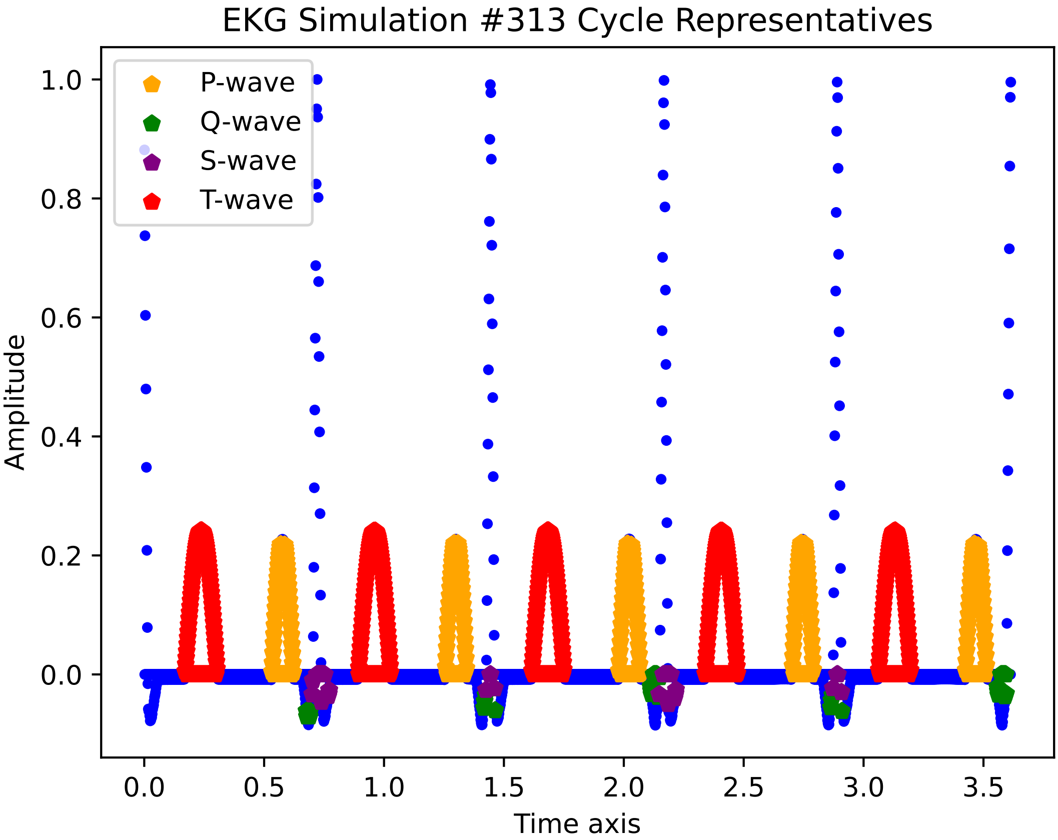

EKG Simulation #313 Cycle Representatives
P-wave
Q-wave
S-wave
T-wave
Amplitude
Time axis
1.0
0.8
0.6
0.4
0.2
0.0
0.0
0.5
1.0
1.5
2.0
2.5
3.0
3.5

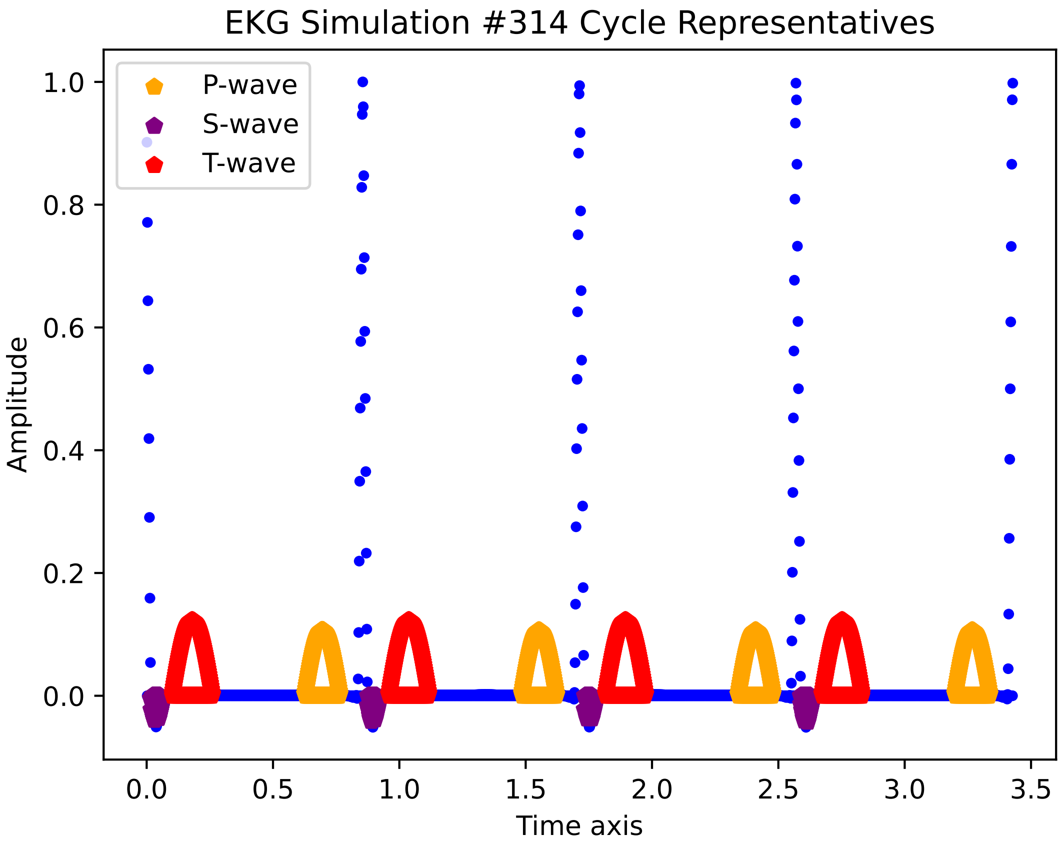
EKG Simulation #314 Cycle Representatives
P-wave
S-wave
T-wave
Amplitude
Time axis
1.0
0.8
0.6
0.4
0.2
0.0
0.0
0.5
1.0
1.5
2.0
2.5
3.0
3.5

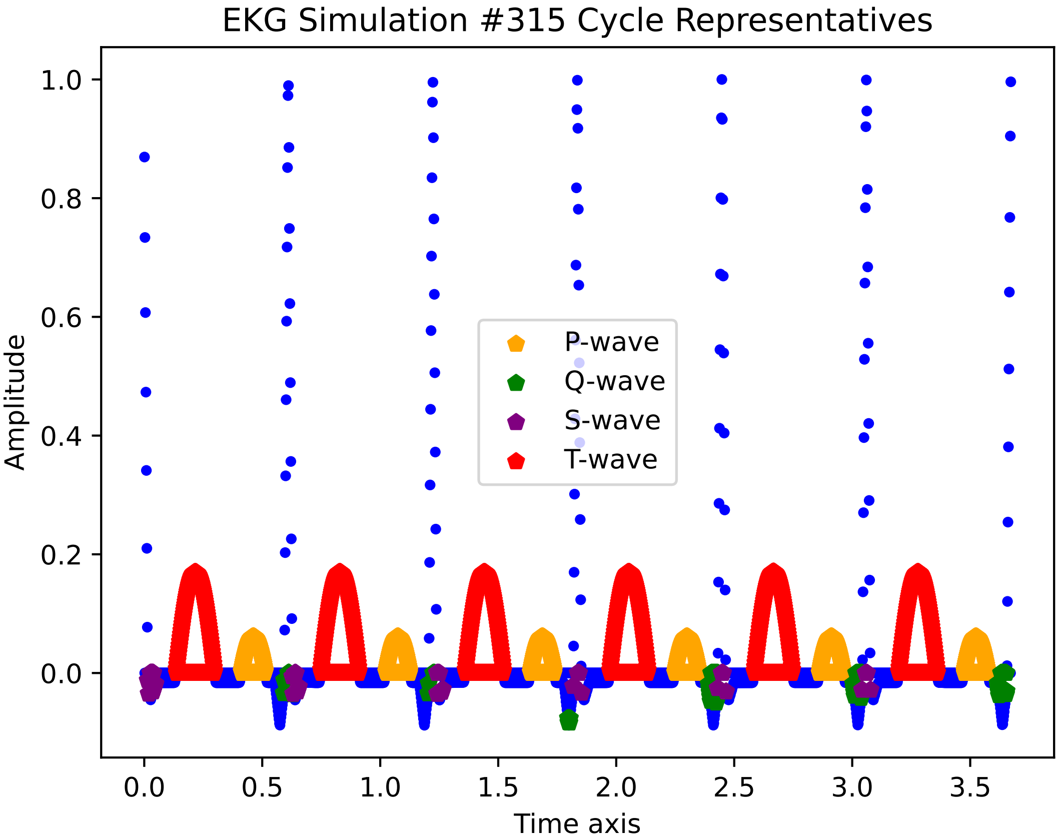

EKG Simulation #315 Cycle Representatives
Amplitude
Time axis
P-wave
Q-wave
S-wave
T-wave
1.0
0.8
0.6
0.4
0.2
0.0
0.0
0.5
1.0
1.5
2.0
2.5
3.0
3.5

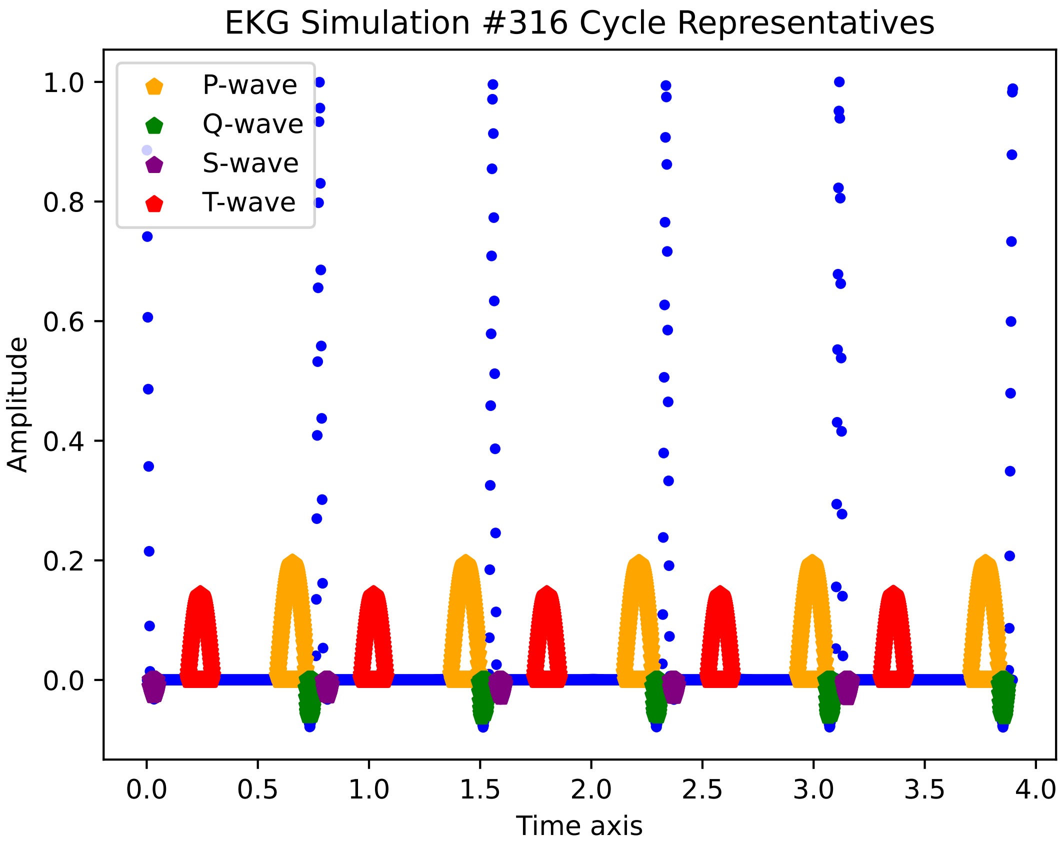
EKG Simulation #316 Cycle Representatives
P-wave
Q-wave
S-wave
T-wave
Amplitude
Time axis
1.0
0.8
0.6
0.4
0.2
0.0
0.0
0.5
1.0
1.5
2.0
2.5
3.0
3.5
4.0

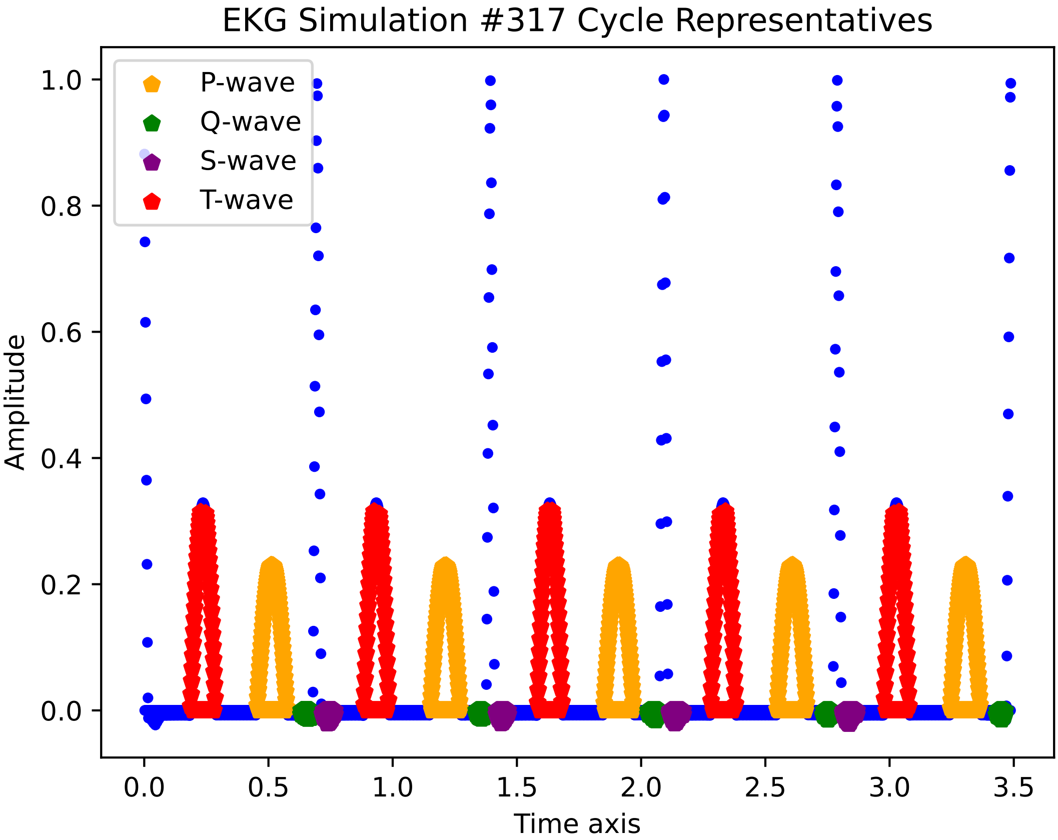
EKG Simulation #317 Cycle Representatives
P-wave
Q-wave
S-wave
T-wave
Amplitude
Time axis
1.0
0.8
0.6
0.4
0.2
0.0
0.0
0.5
1.0
1.5
2.0
2.5
3.0
3.5

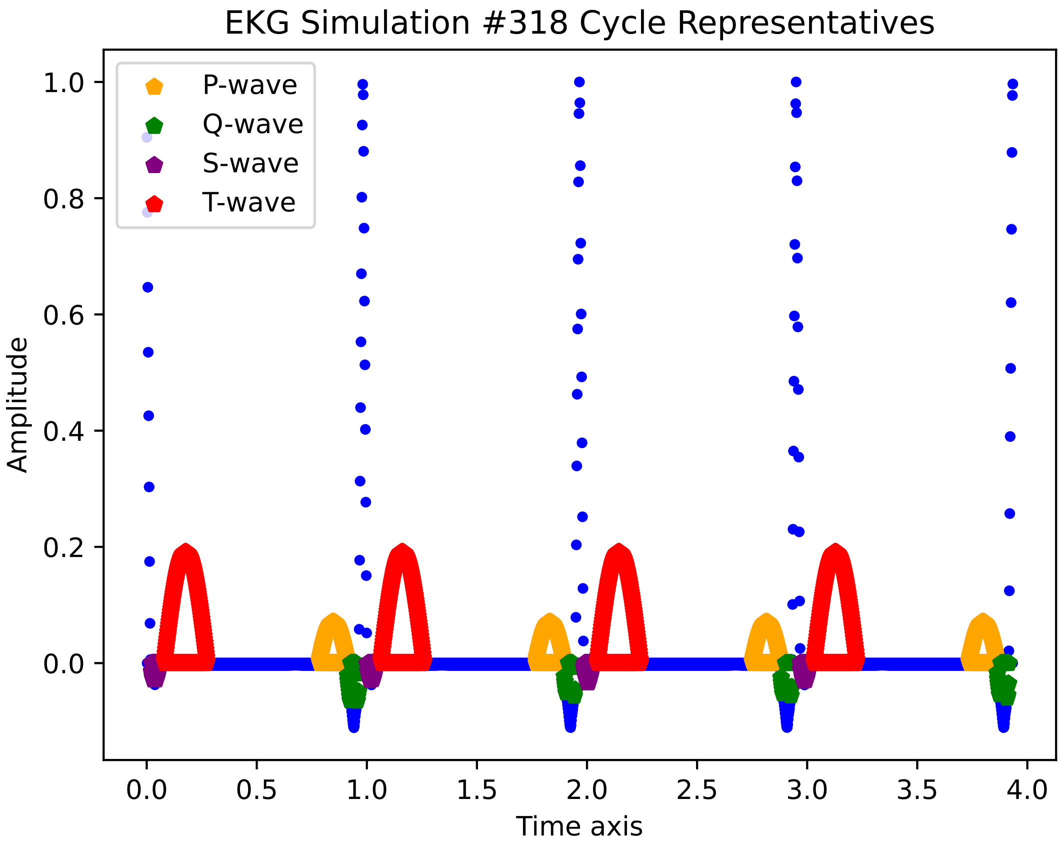

EKG Simulation #318 Cycle Representatives
P-wave
Q-wave
S-wave
T-wave
Amplitude
1.0
0.8
0.6
0.4
0.2
0.0
0.0
0.5
1.0
1.5
2.0
2.5
3.0
3.5
4.0
Time axis

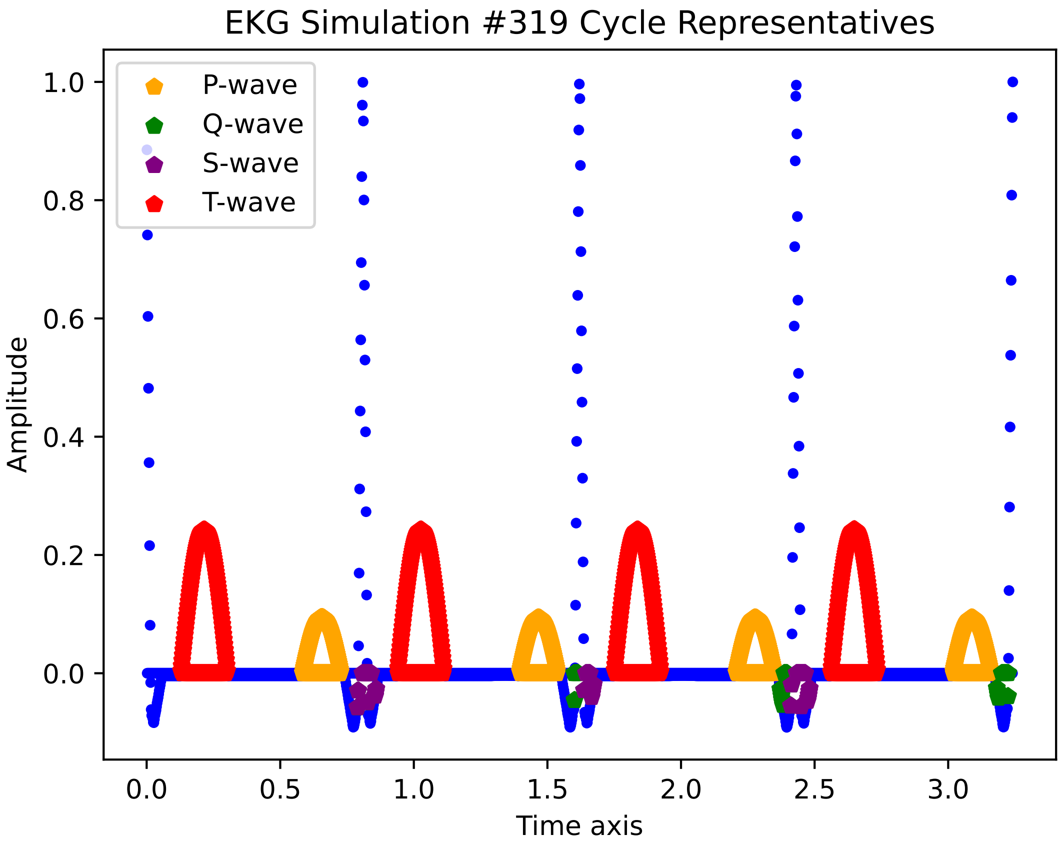
EKG Simulation #319 Cycle Representatives
P-wave
Q-wave
S-wave
T-wave
Amplitude
1.0
0.8
0.6
0.4
0.2
0.0
0.0
0.5
1.0
1.5
2.0
2.5
3.0
Time axis

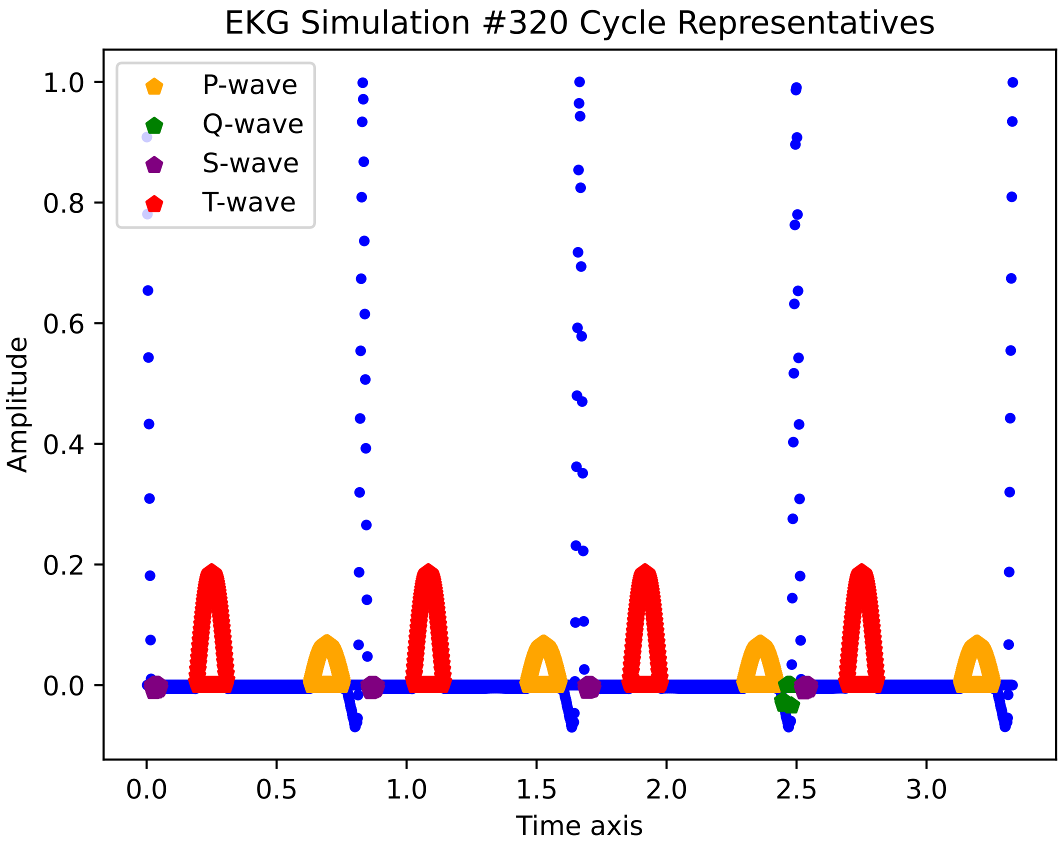
EKG Simulation #320 Cycle Representatives
P-wave
Q-wave
S-wave
T-wave
1.0
0.8
0.6
0.4
0.2
0.0
Amplitude
0.0
0.5
1.0
1.5
2.0
2.5
3.0
Time axis

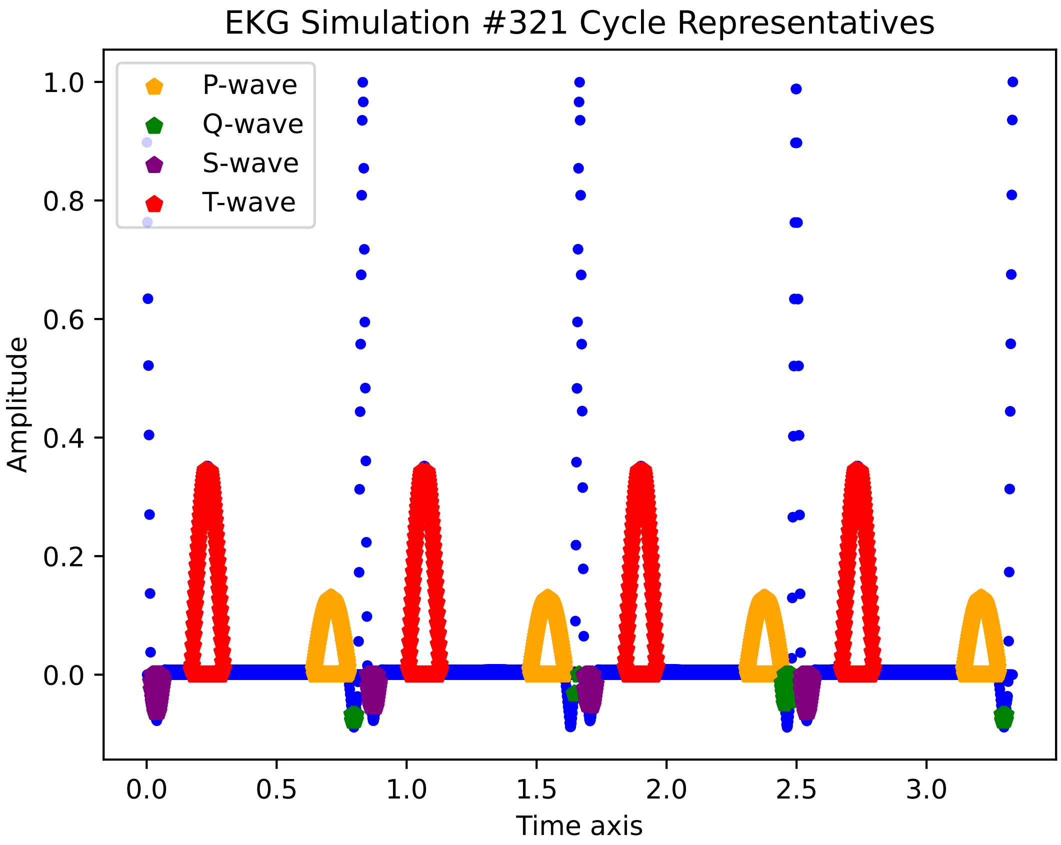

EKG Simulation #321 Cycle Representatives
P-wave
Q-wave
S-wave
T-wave
Amplitude
Time axis
1.0
0.8
0.6
0.4
0.2
0.0
0.0
0.5
1.0
1.5
2.0
2.5
3.0

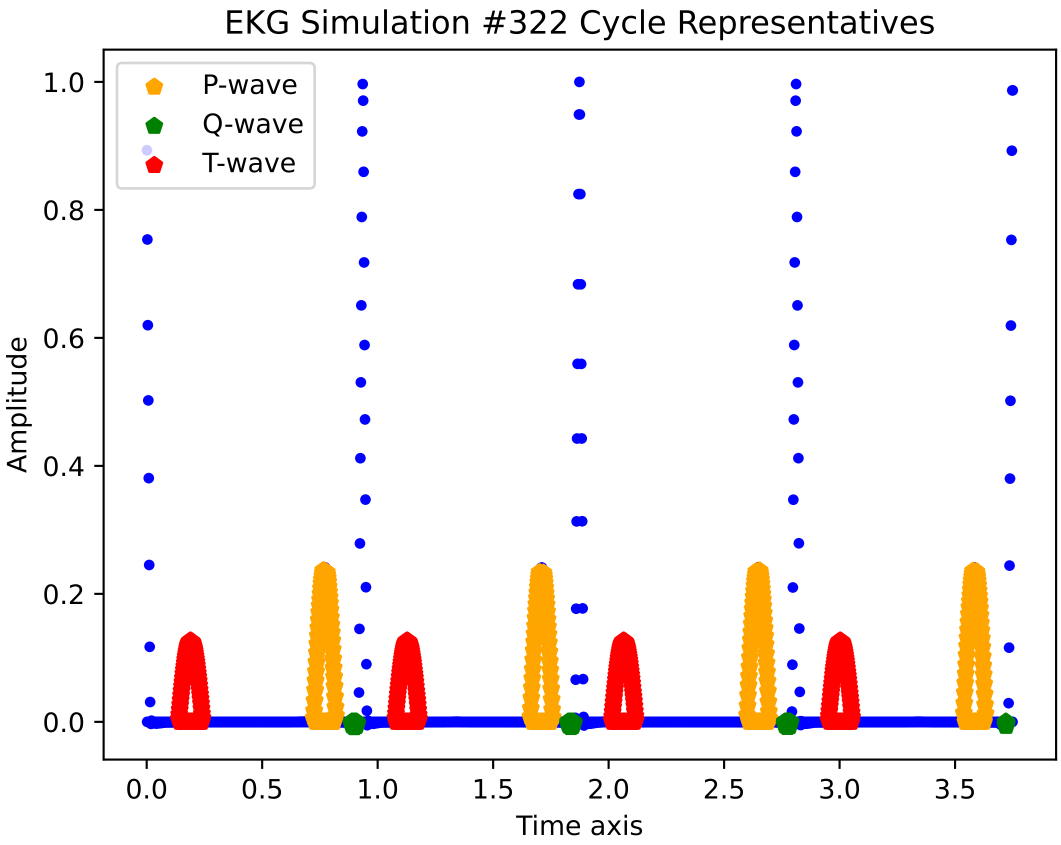
EKG Simulation #322 Cycle Representatives
P-wave
Q-wave
T-wave
Amplitude
1.0
0.8
0.6
0.4
0.2
0.0
0.0
0.5
1.0
1.5
2.0
2.5
3.0
3.5
Time axis

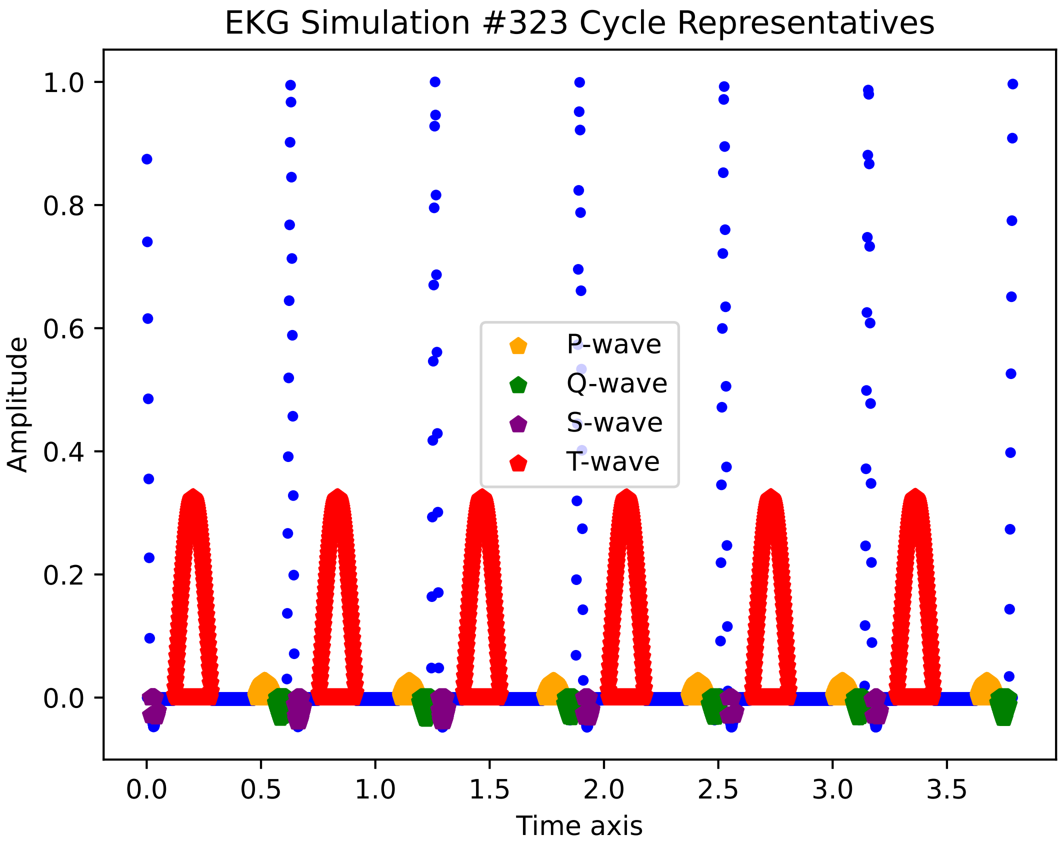

EKG Simulation #323 Cycle Representatives
Amplitude
Time axis
1.0
0.8
0.6
0.4
0.2
0.0
0.0
0.5
1.0
1.5
2.0
2.5
3.0
3.5
P-wave
Q-wave
S-wave
T-wave

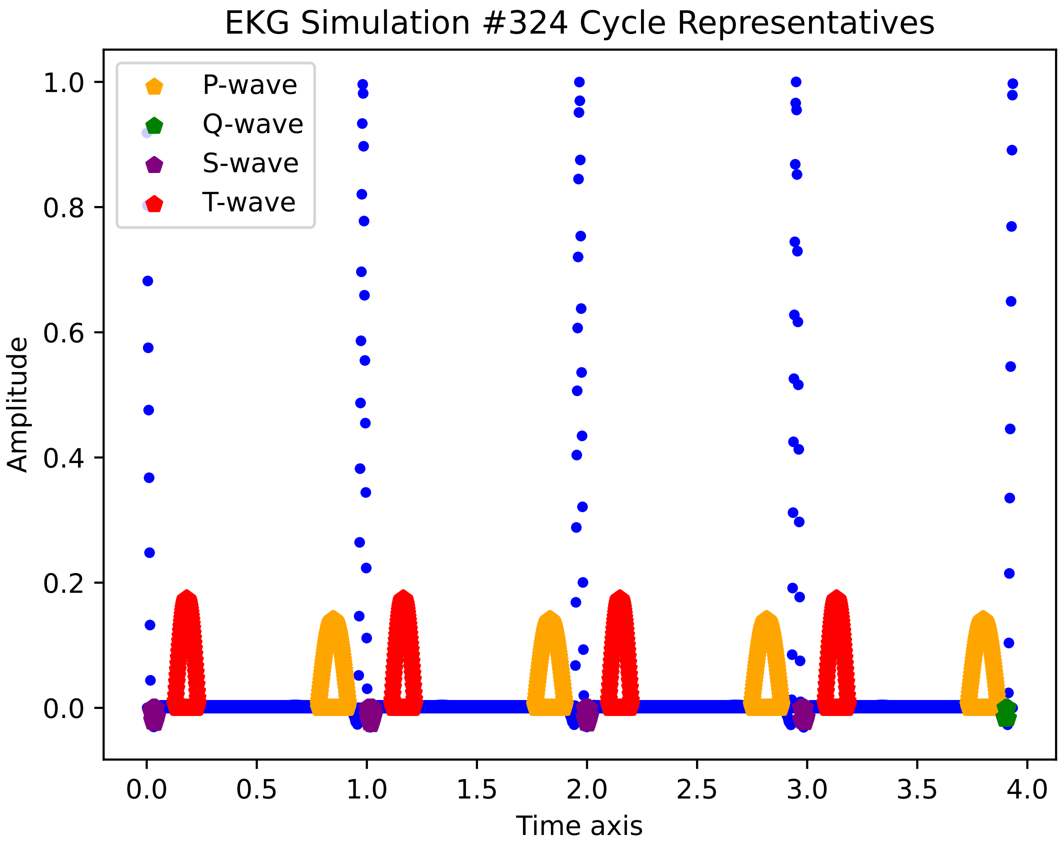
EKG Simulation #324 Cycle Representatives
P-wave
Q-wave
S-wave
T-wave
Amplitude
Time axis
1.0
0.8
0.6
0.4
0.2
0.0
0.0
0.5
1.0
1.5
2.0
2.5
3.0
3.5
4.0

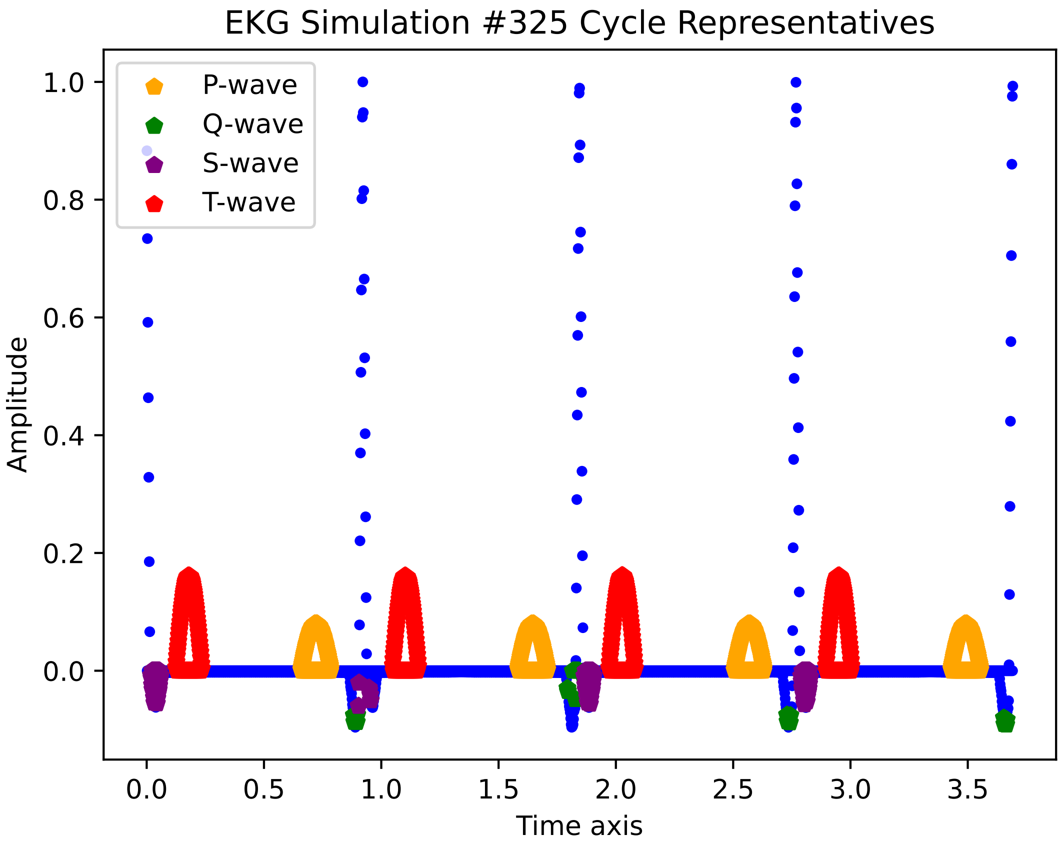

EKG Simulation #325 Cycle Representatives
P-wave
Q-wave
S-wave
T-wave
1.0
0.8
0.6
0.4
0.2
0.0
Amplitude
0.0
0.5
1.0
1.5
2.0
2.5
3.0
3.5
Time axis

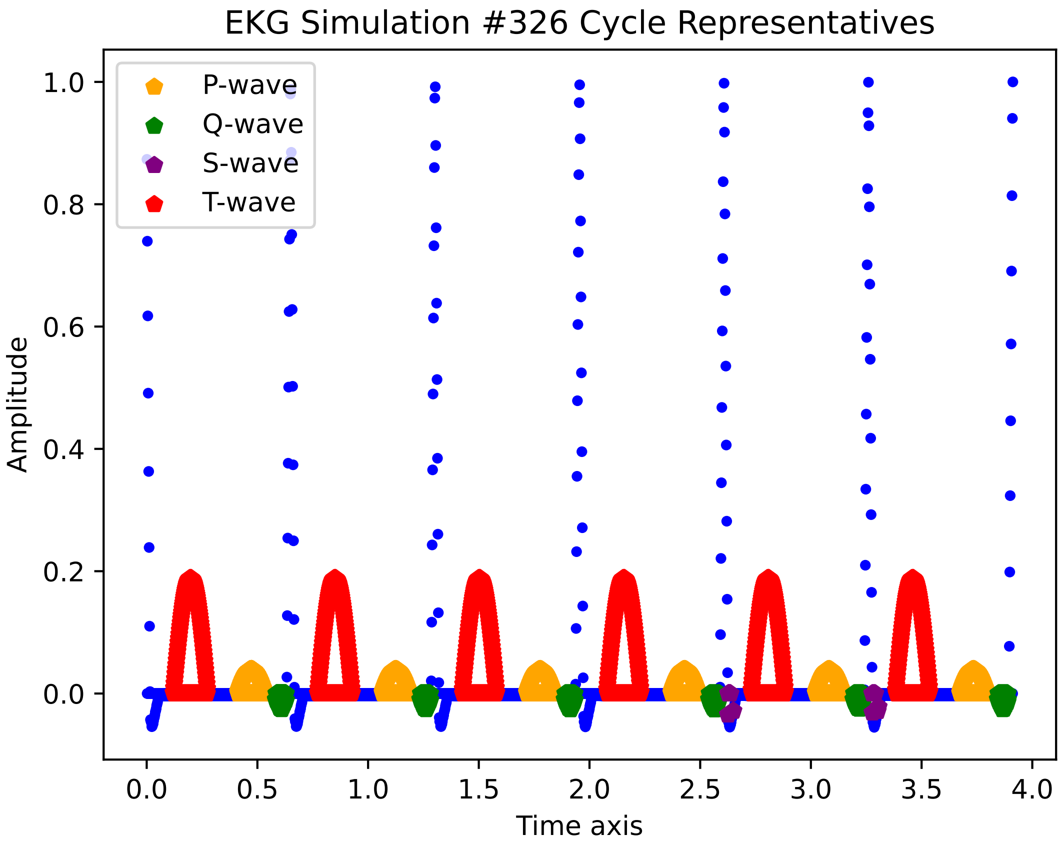

EKG Simulation #326 Cycle Representatives
P-wave
Q-wave
S-wave
T-wave
Amplitude
Time axis
1.0
0.8
0.6
0.4
0.2
0.0
0.0
0.5
1.0
1.5
2.0
2.5
3.0
3.5
4.0

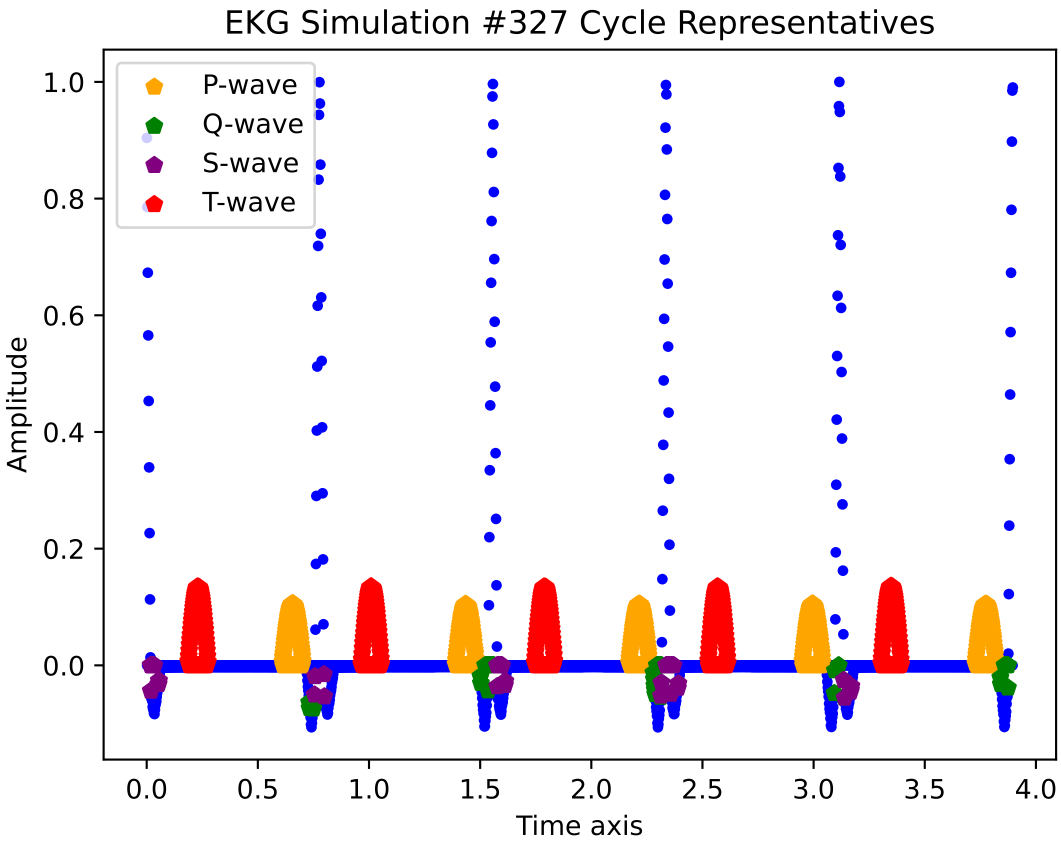
EKG Simulation #327 Cycle Representatives
P-wave
Q-wave
S-wave
T-wave
Amplitude
Time axis
1.0
0.8
0.6
0.4
0.2
0.0
0.0
0.5
1.0
1.5
2.0
2.5
3.0
3.5
4.0

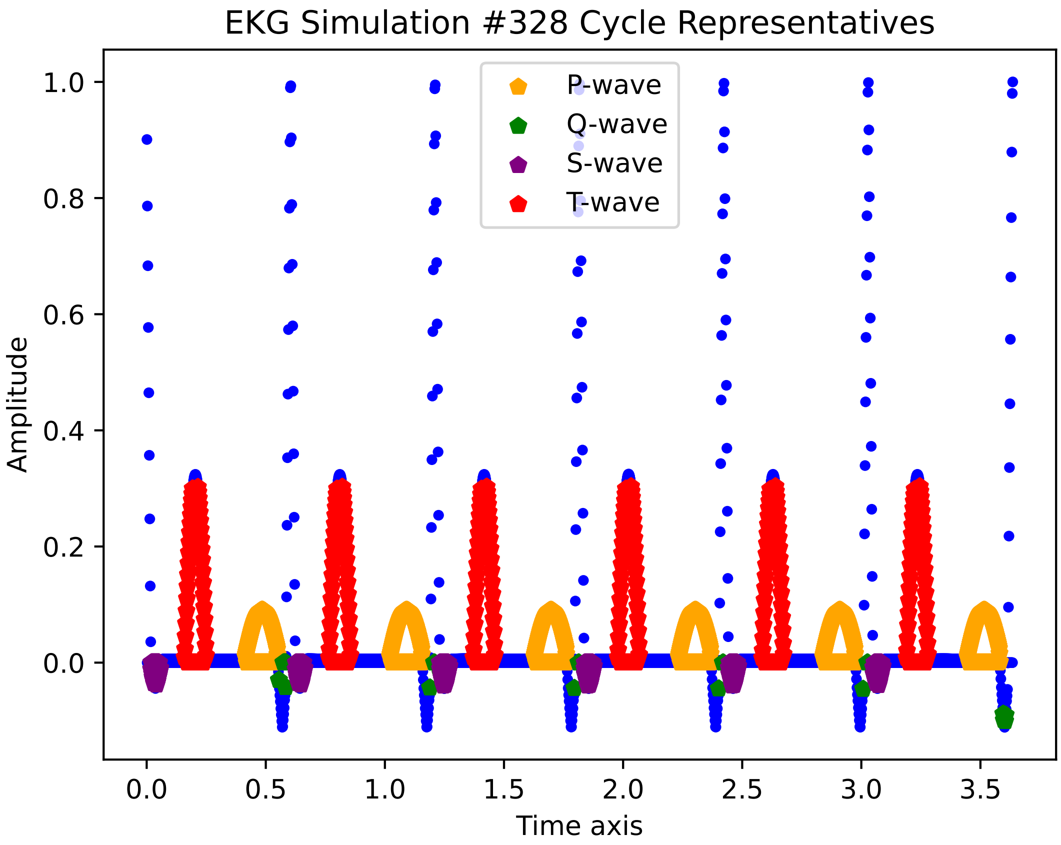
EKG Simulation #328 Cycle Representatives
P-wave
Q-wave
S-wave
T-wave
Amplitude
Time axis
1.0
0.8
0.6
0.4
0.2
0.0
0.0
0.5
1.0
1.5
2.0
2.5
3.0
3.5

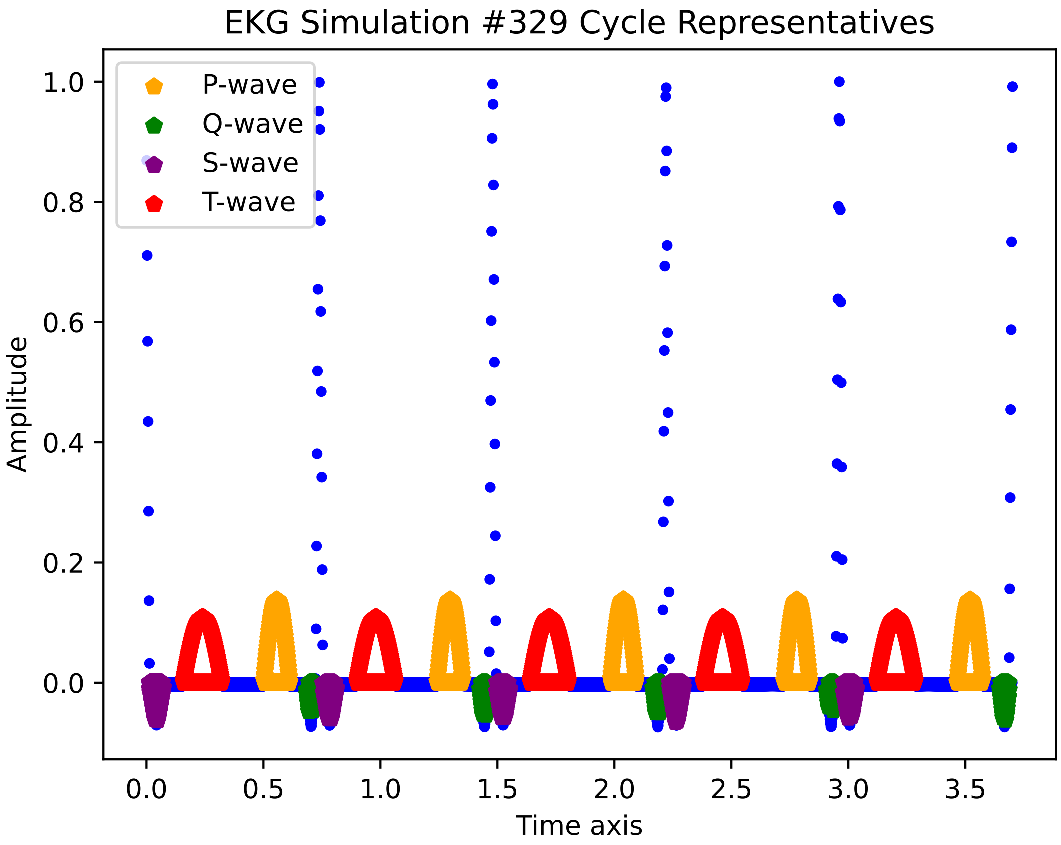
EKG Simulation #329 Cycle Representatives
P-wave
Q-wave
S-wave
T-wave
Amplitude
1.0
0.8
0.6
0.4
0.2
0.0
0.0
0.5
1.0
1.5
2.0
2.5
3.0
3.5
Time axis

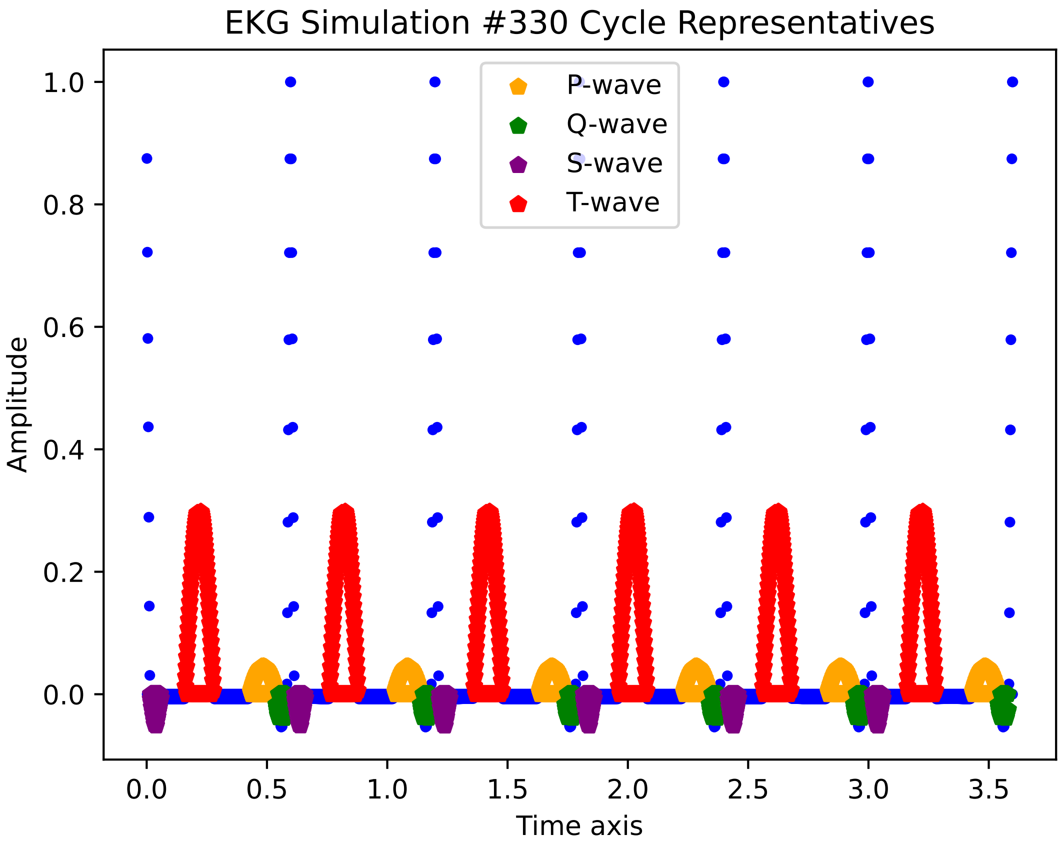
EKG Simulation #330 Cycle Representatives
P-wave
Q-wave
S-wave
T-wave
Amplitude
Time axis
1.0
0.8
0.6
0.4
0.2
0.0
0.0
0.5
1.0
1.5
2.0
2.5
3.0
3.5

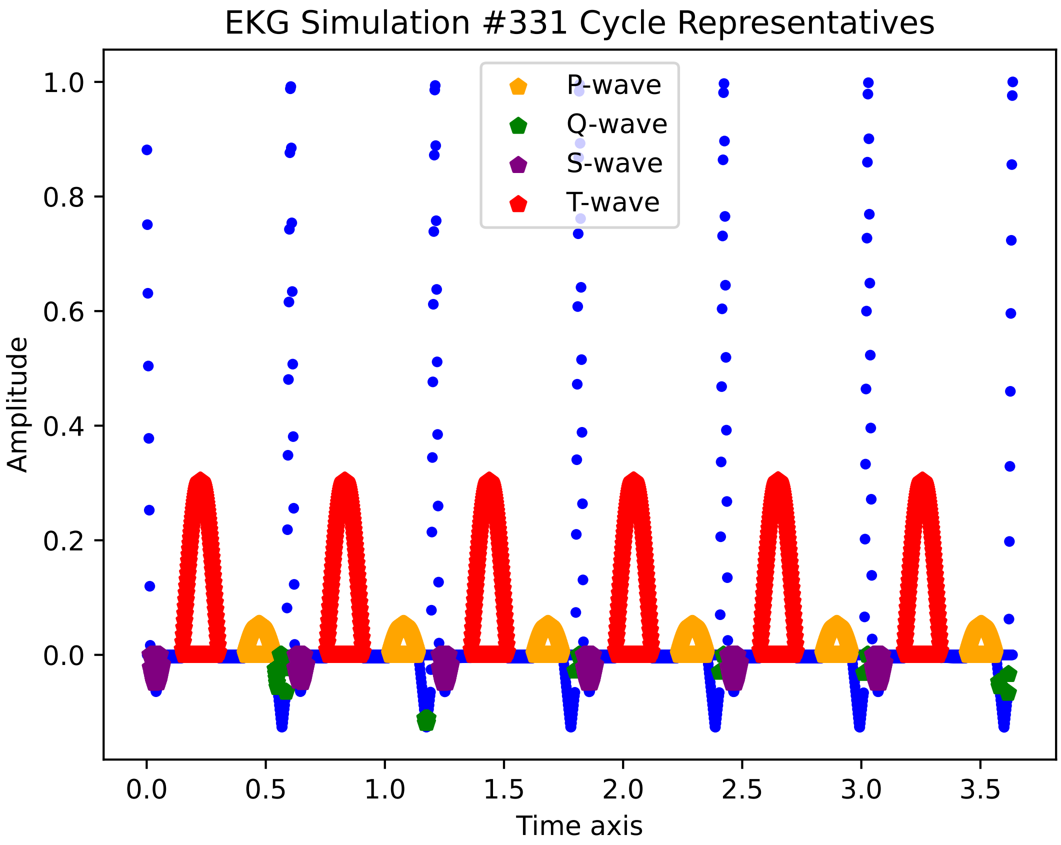

EKG Simulation #331 Cycle Representatives
P-wave
Q-wave
S-wave
T-wave
Amplitude
Time axis
1.0
0.8
0.6
0.4
0.2
0.0
0.0
0.5
1.0
1.5
2.0
2.5
3.0
3.5

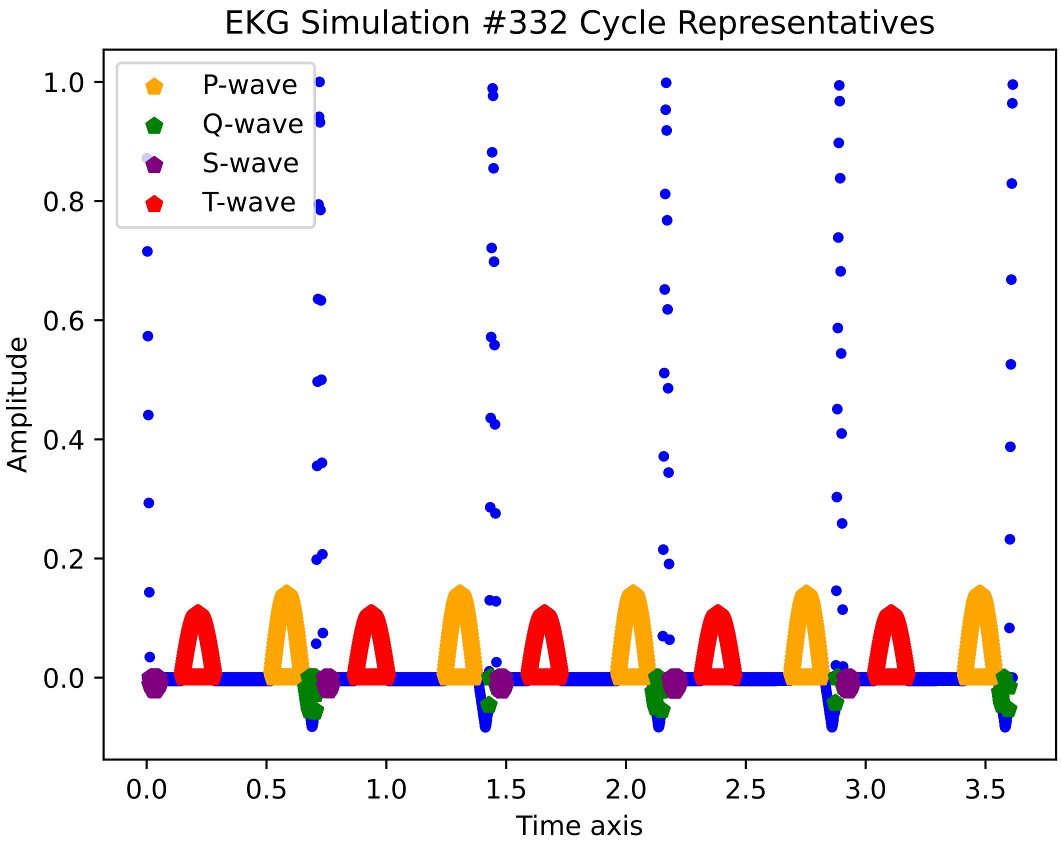

EKG Simulation #332 Cycle Representatives
P-wave
Q-wave
S-wave
T-wave
Amplitude
Time axis
1.0
0.8
0.6
0.4
0.2
0.0
0.0
0.5
1.0
1.5
2.0
2.5
3.0
3.5

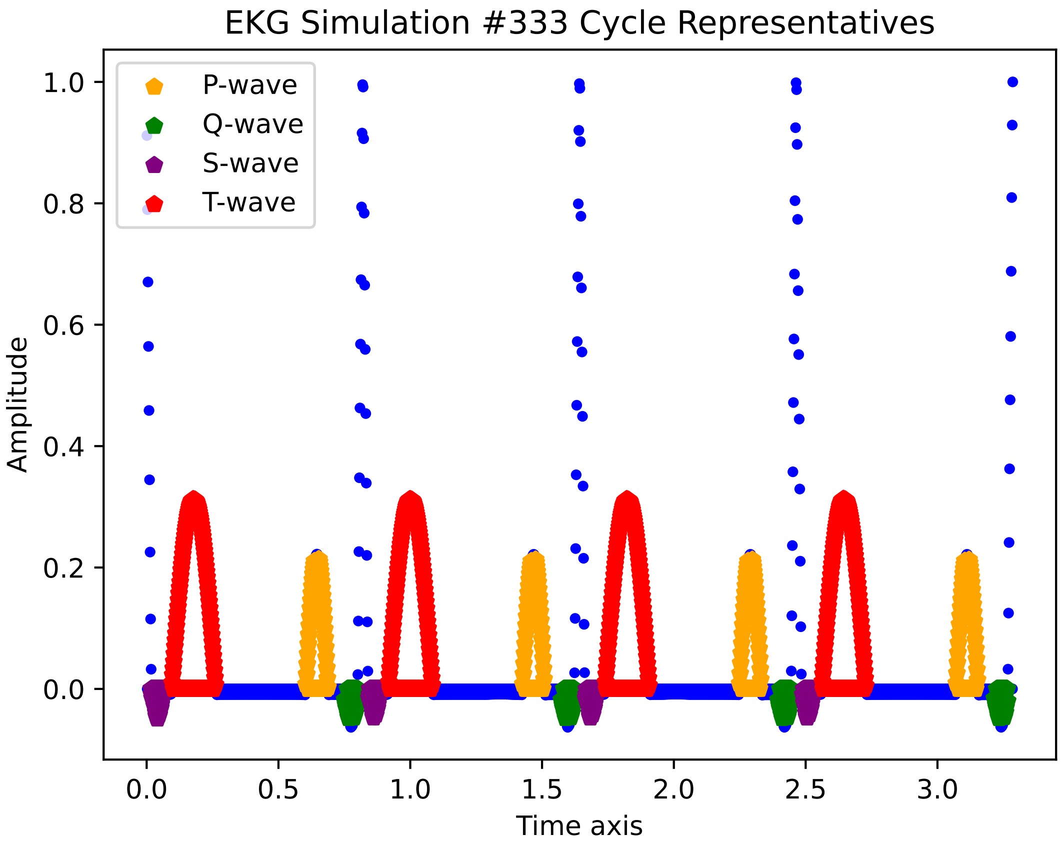
EKG Simulation #333 Cycle Representatives
P-wave
Q-wave
S-wave
T-wave
Amplitude
Time axis
1.0
0.8
0.6
0.4
0.2
0.0
0.0
0.5
1.0
1.5
2.0
2.5
3.0

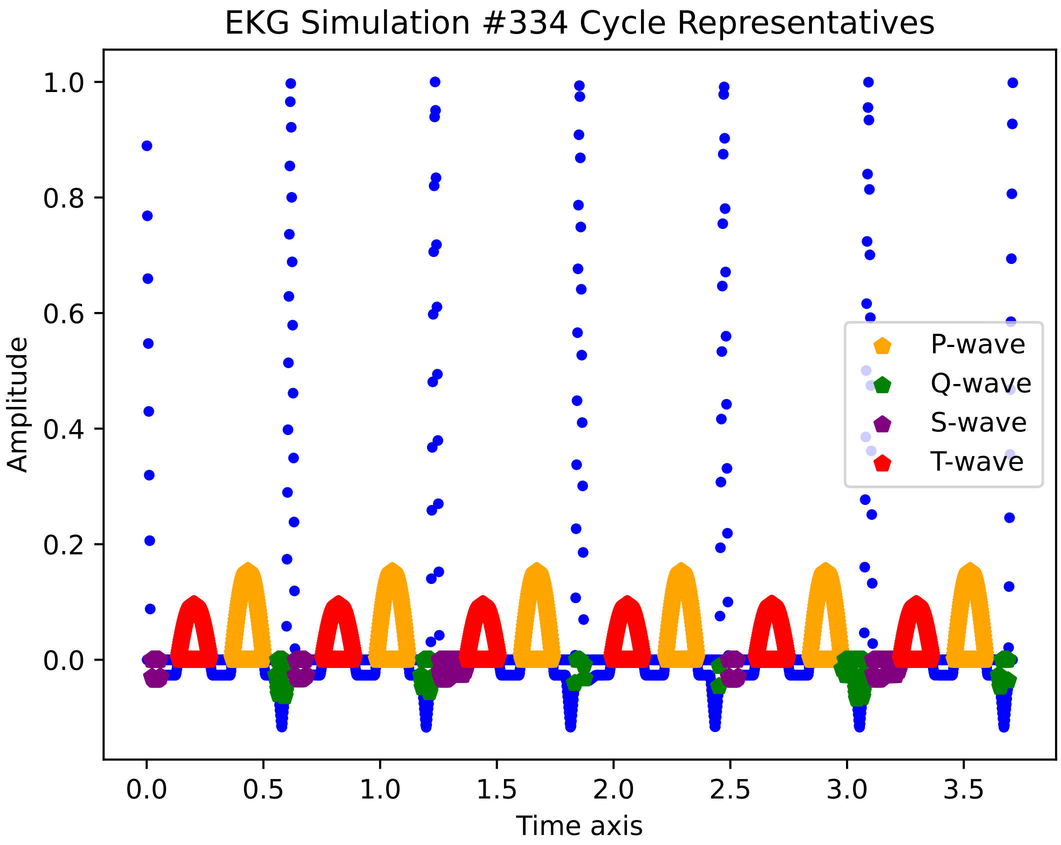

EKG Simulation #334 Cycle Representatives
Amplitude
Time axis
1.0
0.8
0.6
0.4
0.2
0.0
0.0
0.5
1.0
1.5
2.0
2.5
3.0
3.5
P-wave
Q-wave
S-wave
T-wave

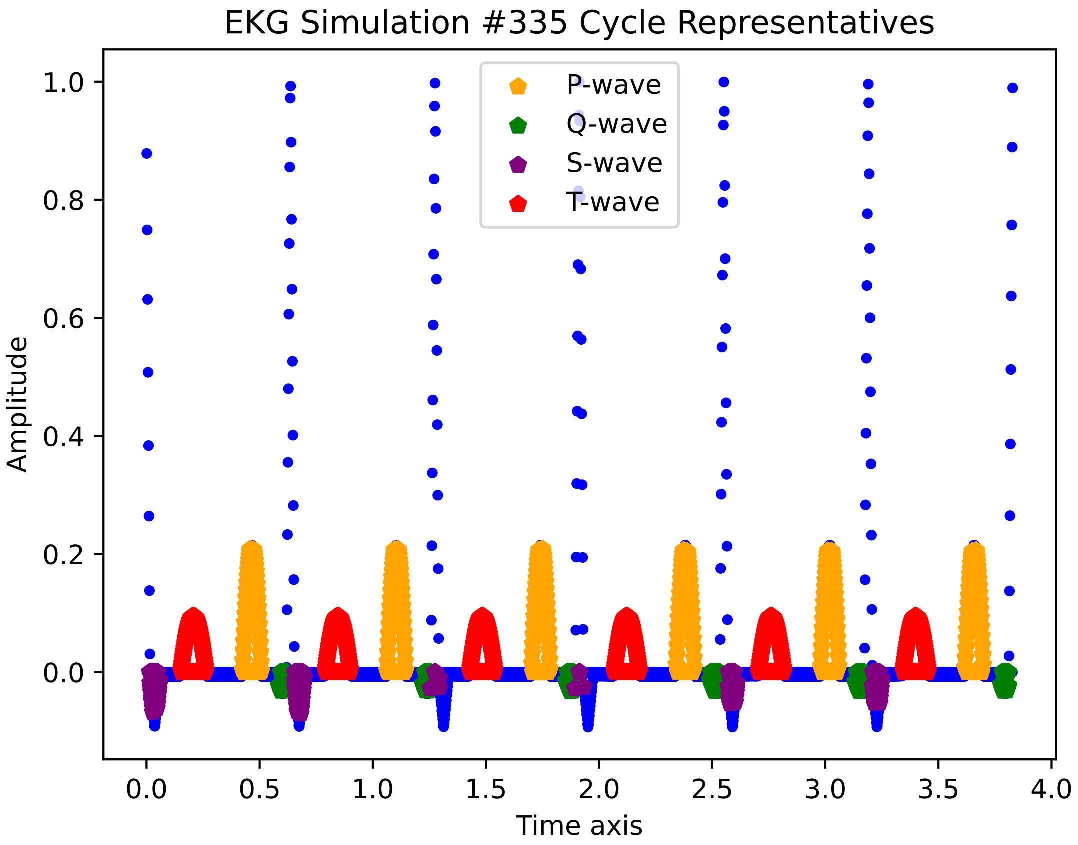

EKG Simulation #335 Cycle Representatives
P-wave
Q-wave
S-wave
T-wave
Amplitude
Time axis
1.0
0.8
0.6
0.4
0.2
0.0
0.0
0.5
1.0
1.5
2.0
2.5
3.0
3.5
4.0

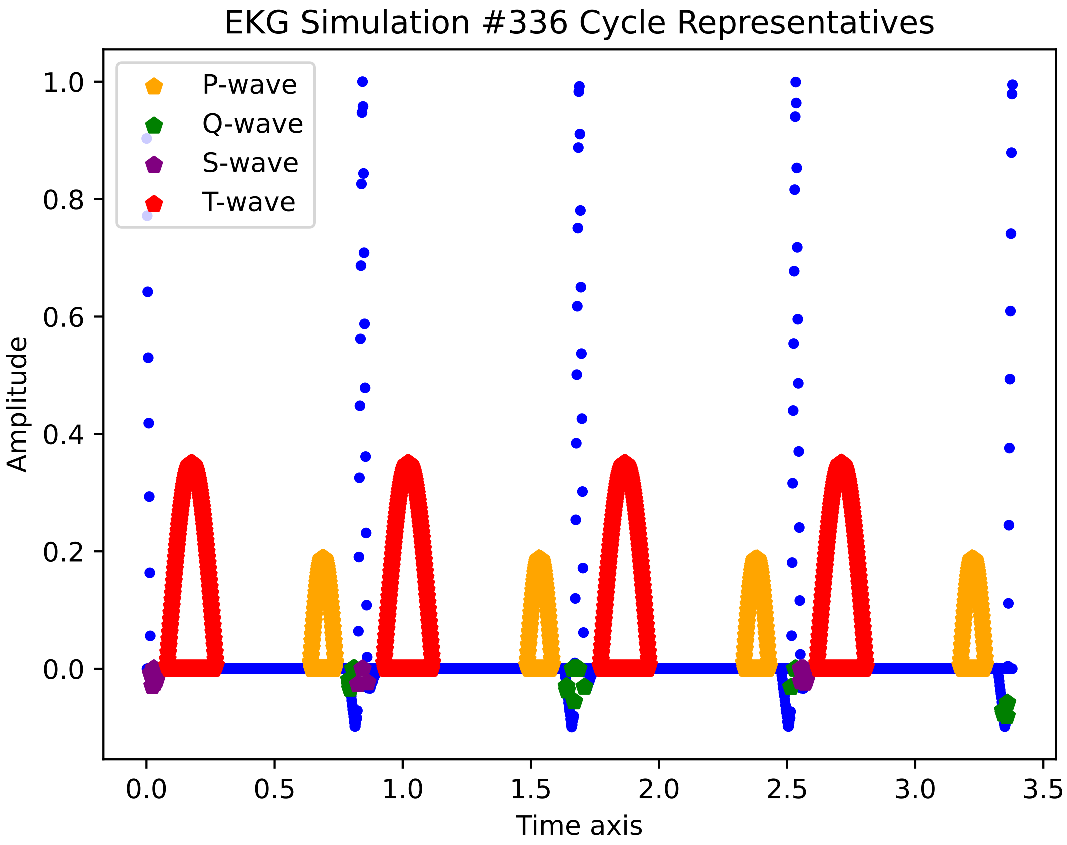

EKG Simulation #336 Cycle Representatives
P-wave
Q-wave
S-wave
T-wave
Amplitude
Time axis
1.0
0.8
0.6
0.4
0.2
0.0
0.0
0.5
1.0
1.5
2.0
2.5
3.0
3.5

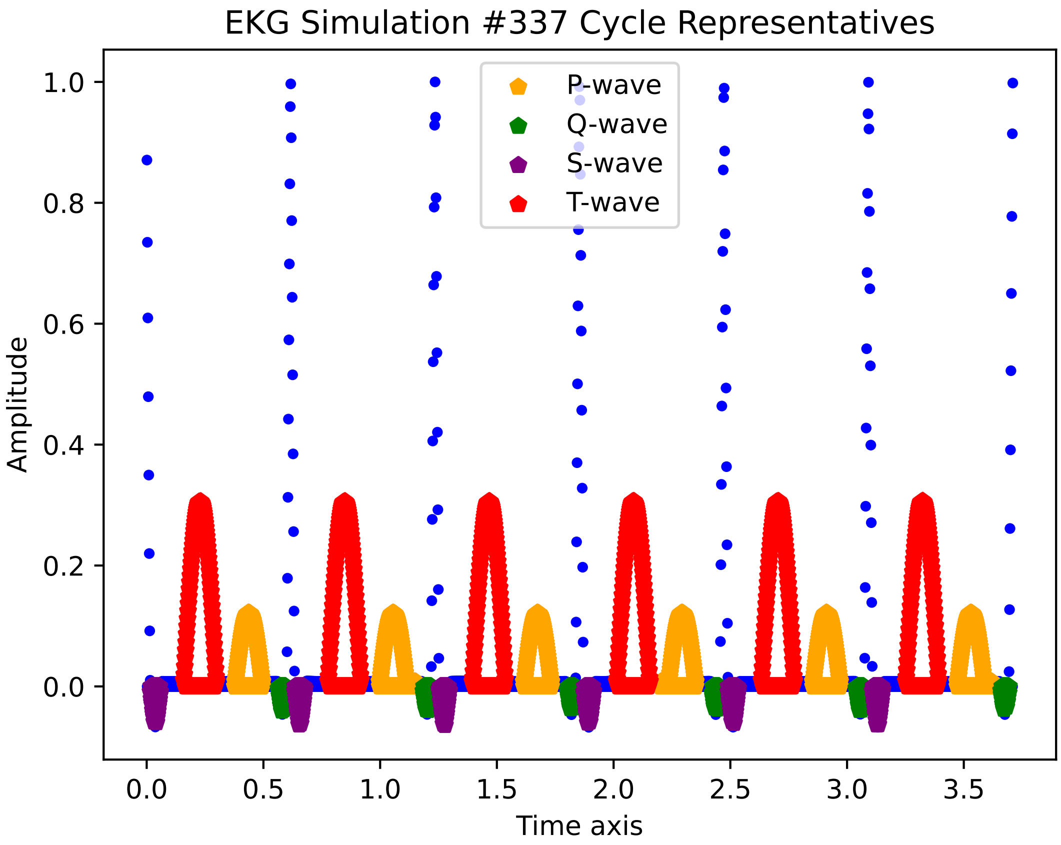

EKG Simulation #337 Cycle Representatives
P-wave
Q-wave
S-wave
T-wave
Amplitude
Time axis
1.0
0.8
0.6
0.4
0.2
0.0
0.0
0.5
1.0
1.5
2.0
2.5
3.0
3.5

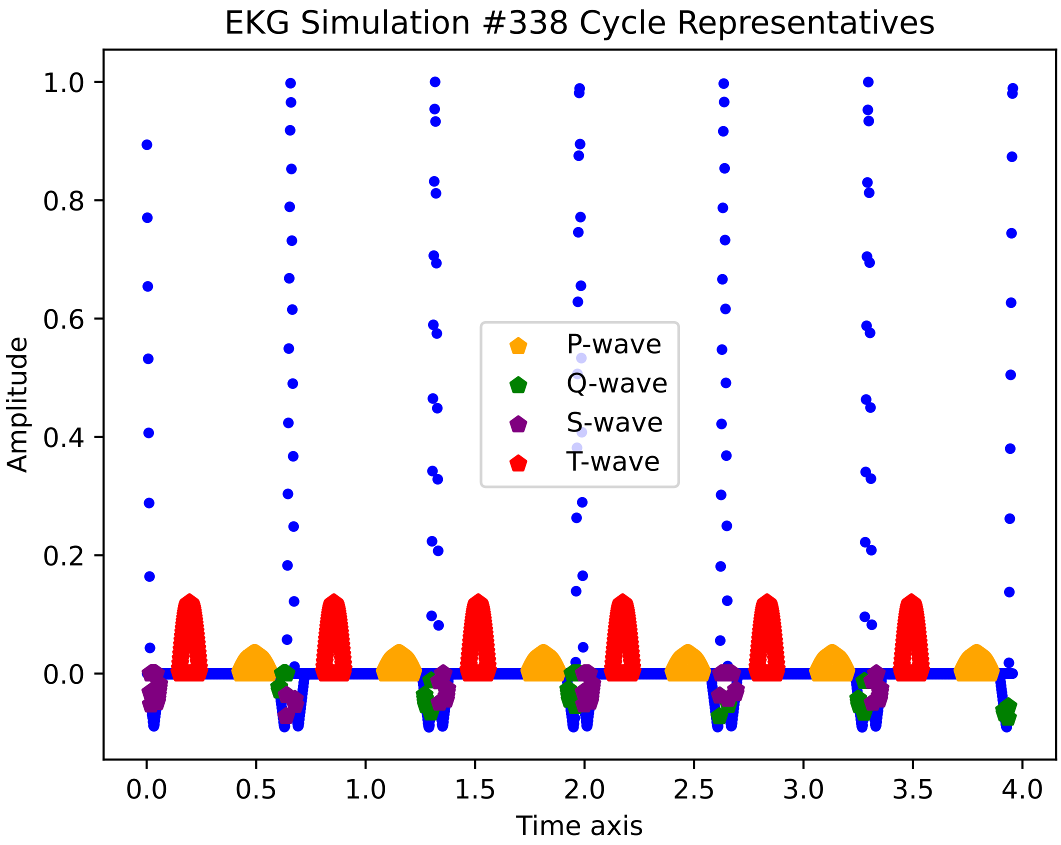

EKG Simulation #338 Cycle Representatives
Amplitude
Time axis
1.0
0.8
0.6
0.4
0.2
0.0
0.0
0.5
1.0
1.5
2.0
2.5
3.0
3.5
4.0
P-wave
Q-wave
S-wave
T-wave

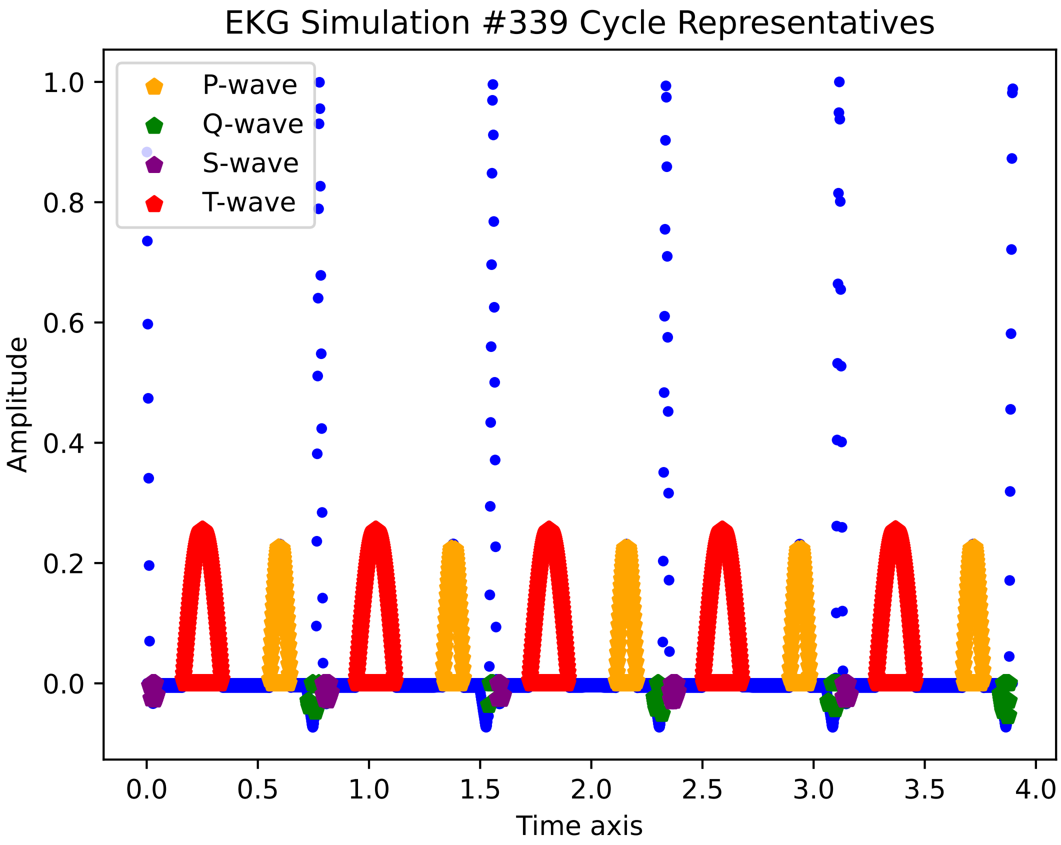
EKG Simulation #339 Cycle Representatives
P-wave
Q-wave
S-wave
T-wave
Amplitude
1.0
0.8
0.6
0.4
0.2
0.0
0.0
0.5
1.0
1.5
2.0
2.5
3.0
3.5
4.0
Time axis

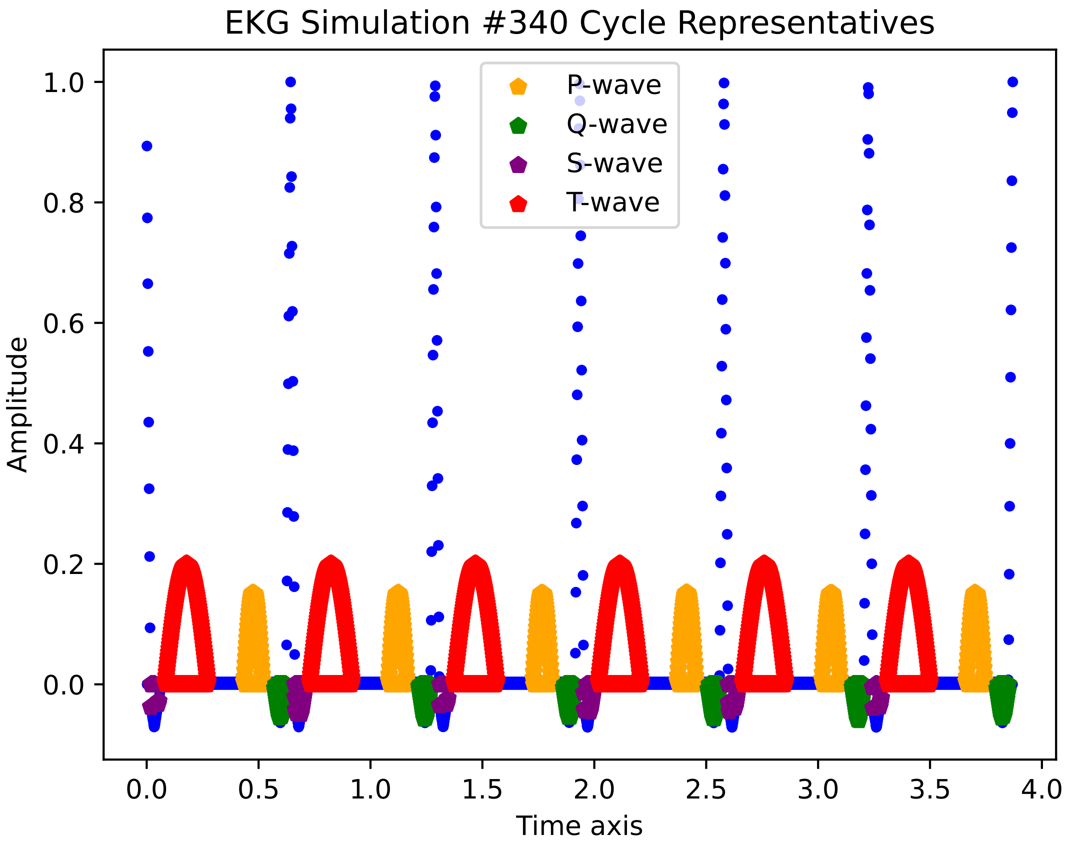
EKG Simulation #340 Cycle Representatives
P-wave
Q-wave
S-wave
T-wave
Amplitude
1.0
0.8
0.6
0.4
0.2
0.0
0.0
0.5
1.0
1.5
2.0
2.5
3.0
3.5
4.0
Time axis

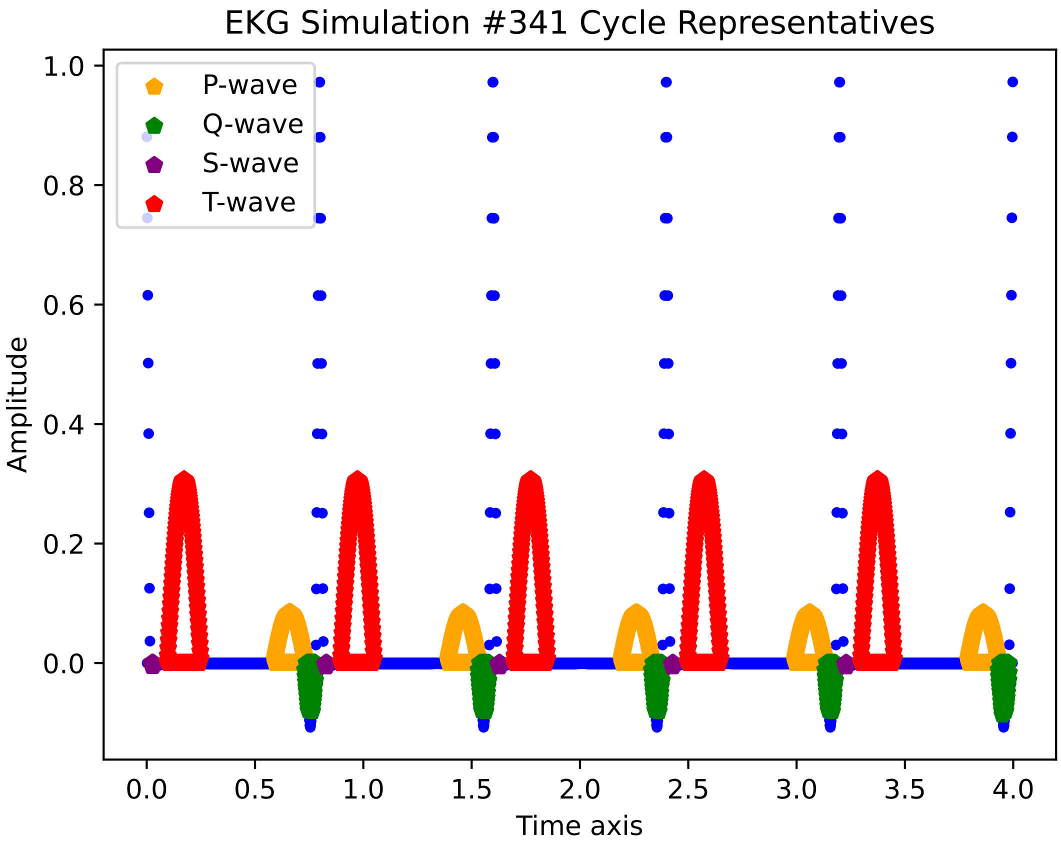
EKG Simulation #341 Cycle Representatives
P-wave
Q-wave
S-wave
T-wave
Amplitude
Time axis
1.0
0.8
0.6
0.4
0.2
0.0
0.0
0.5
1.0
1.5
2.0
2.5
3.0
3.5
4.0

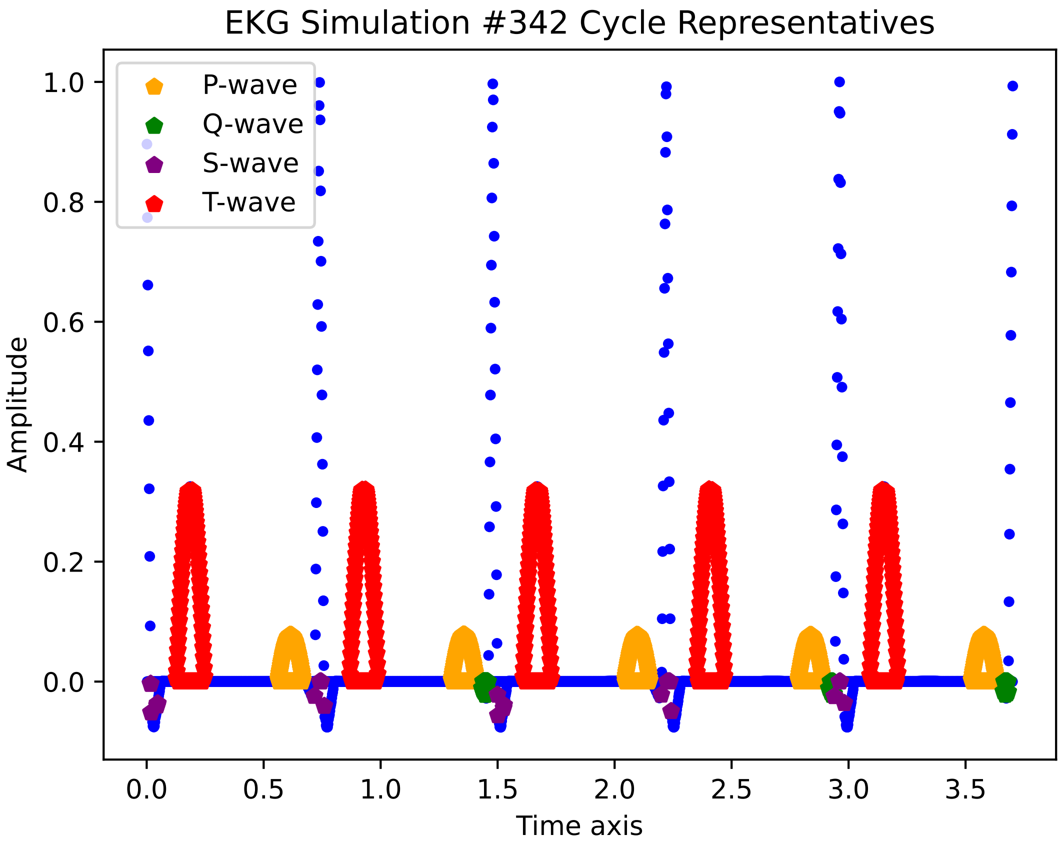
EKG Simulation #342 Cycle Representatives
P-wave
Q-wave
S-wave
T-wave
Amplitude
Time axis
1.0
0.8
0.6
0.4
0.2
0.0
0.0
0.5
1.0
1.5
2.0
2.5
3.0
3.5

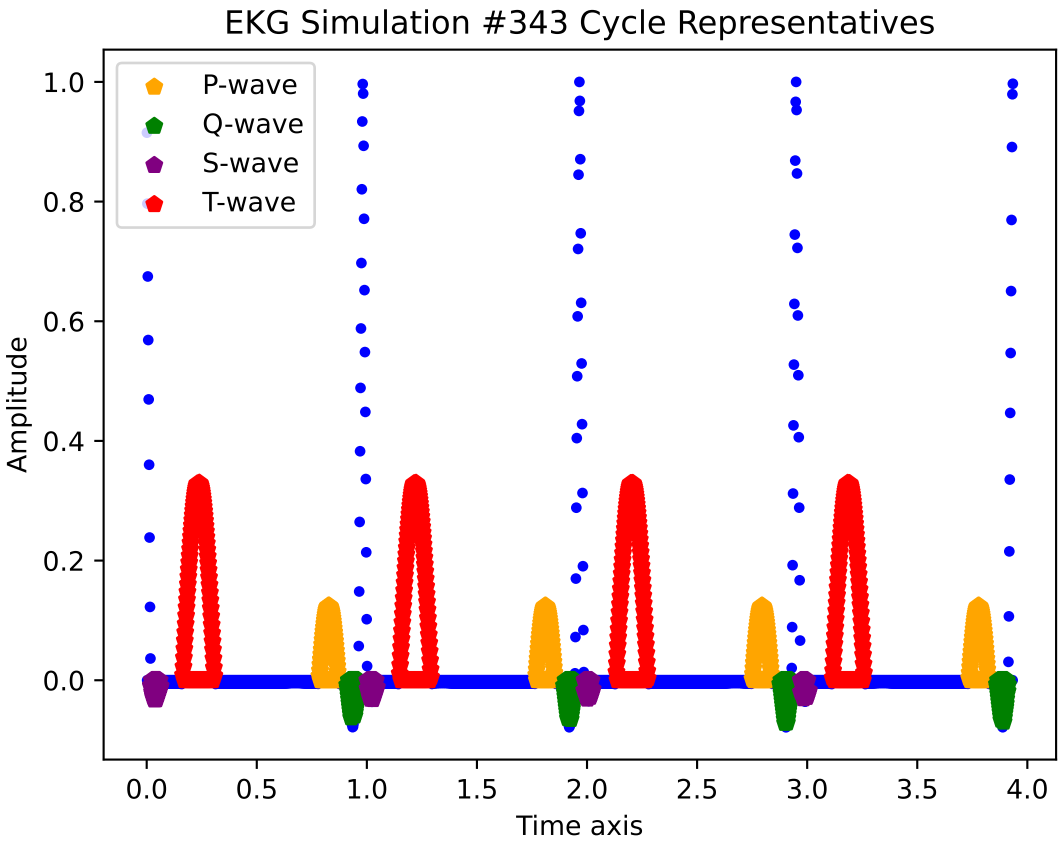
EKG Simulation #343 Cycle Representatives
P-wave
Q-wave
S-wave
T-wave
Amplitude
Time axis
1.0
0.8
0.6
0.4
0.2
0.0
0.0
0.5
1.0
1.5
2.0
2.5
3.0
3.5
4.0

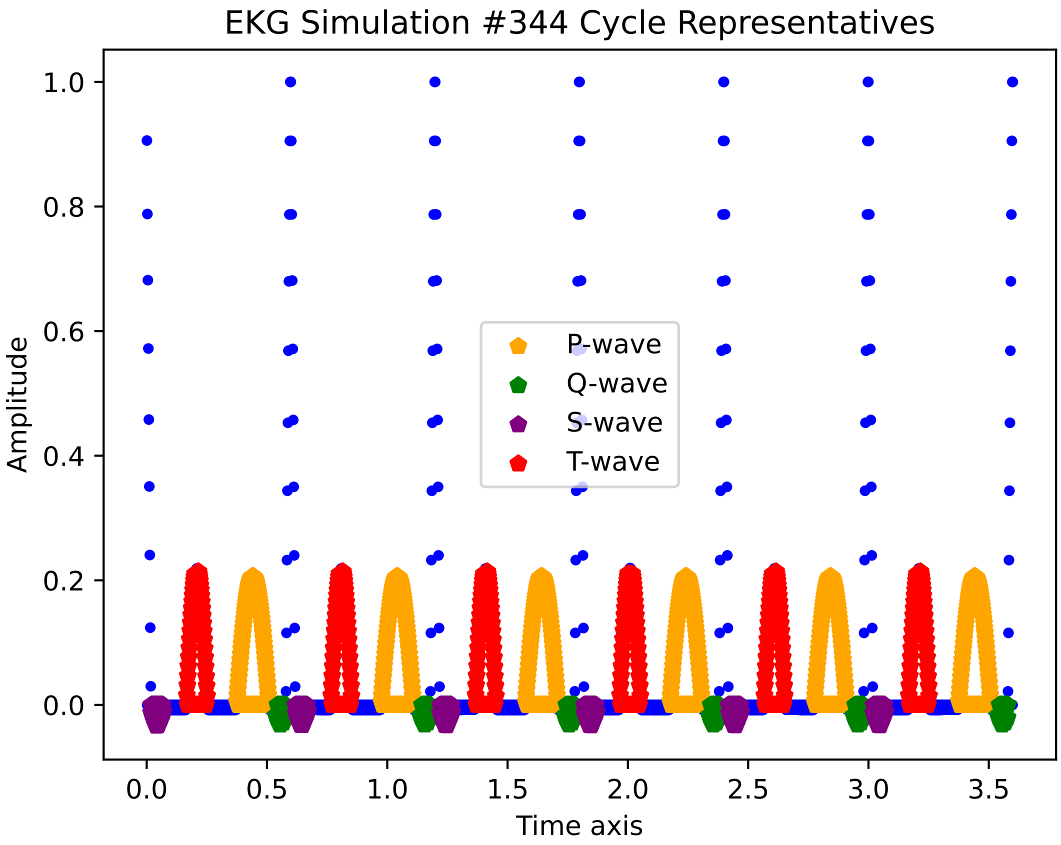

EKG Simulation #344 Cycle Representatives
Amplitude
Time axis
P-wave
Q-wave
S-wave
T-wave
1.0
0.8
0.6
0.4
0.2
0.0
0.0
0.5
1.0
1.5
2.0
2.5
3.0
3.5

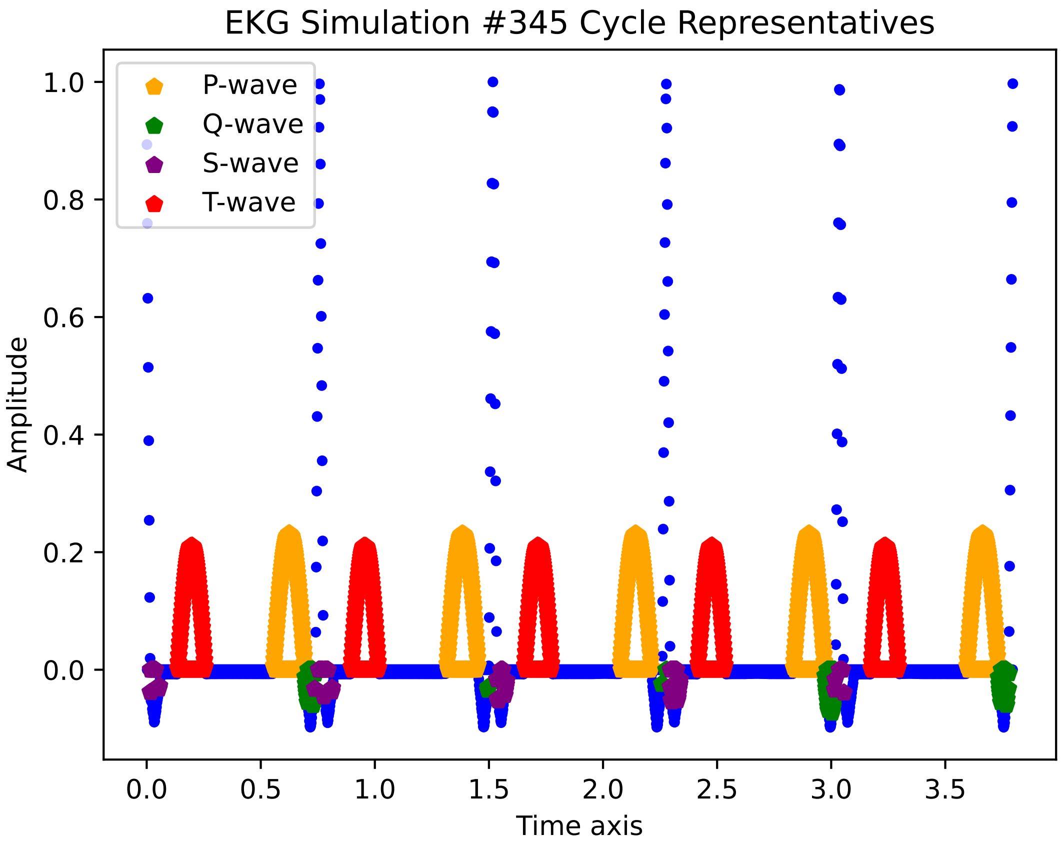
EKG Simulation #345 Cycle Representatives
P-wave
Q-wave
S-wave
T-wave
Amplitude
Time axis
1.0
0.8
0.6
0.4
0.2
0.0
0.0
0.5
1.0
1.5
2.0
2.5
3.0
3.5

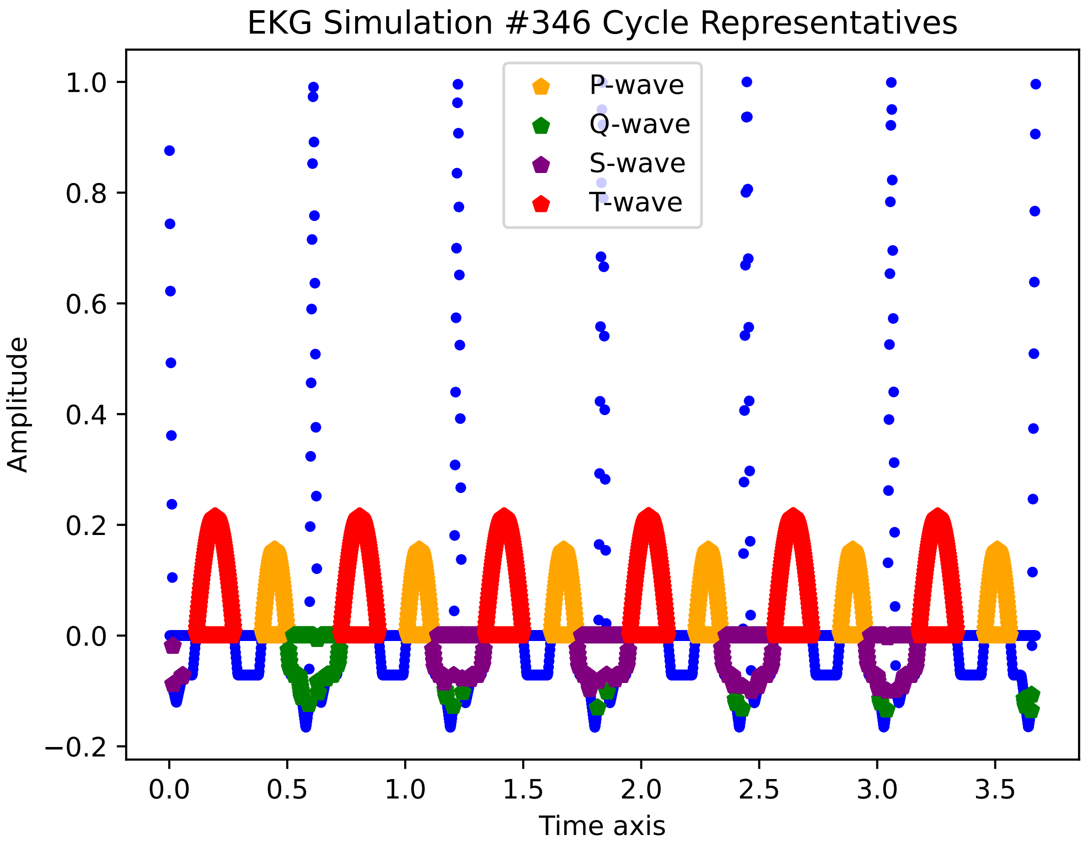

EKG Simulation #346 Cycle Representatives
P-wave
Q-wave
S-wave
T-wave
Amplitude
Time axis
1.0
0.8
0.6
0.4
0.2
0.0
−0.2
0.0
0.5
1.0
1.5
2.0
2.5
3.0
3.5

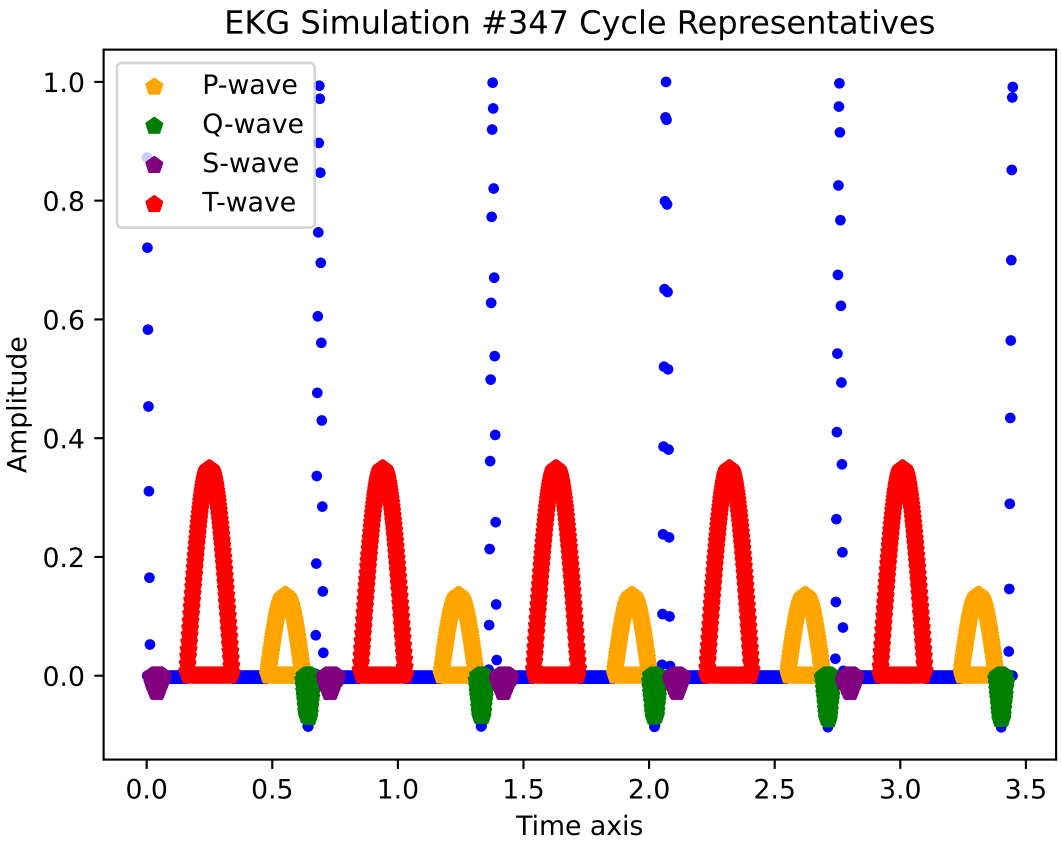
EKG Simulation #347 Cycle Representatives
P-wave
Q-wave
S-wave
T-wave
1.0
0.8
0.6
0.4
0.2
0.0
Amplitude
0.0
0.5
1.0
1.5
2.0
2.5
3.0
3.5
Time axis

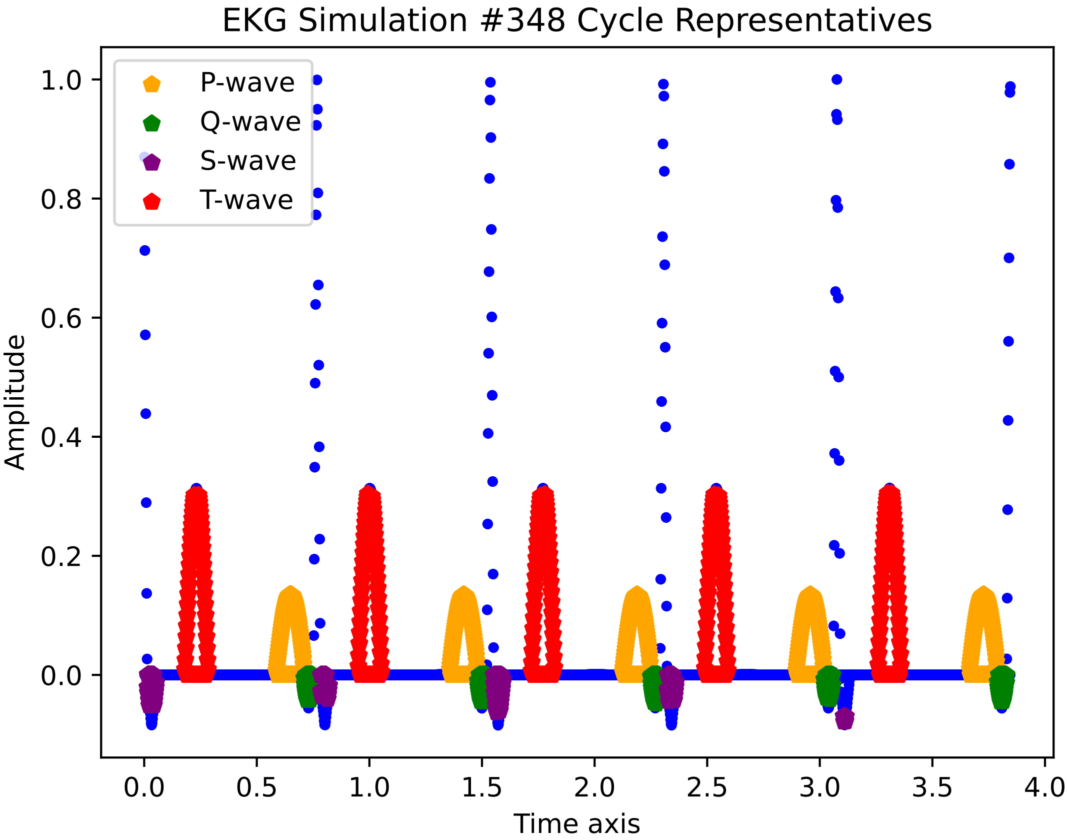

EKG Simulation #348 Cycle Representatives
P-wave
Q-wave
S-wave
T-wave
1.0
0.8
0.6
0.4
0.2
0.0
Amplitude
0.0
0.5
1.0
1.5
2.0
2.5
3.0
3.5
4.0
Time axis

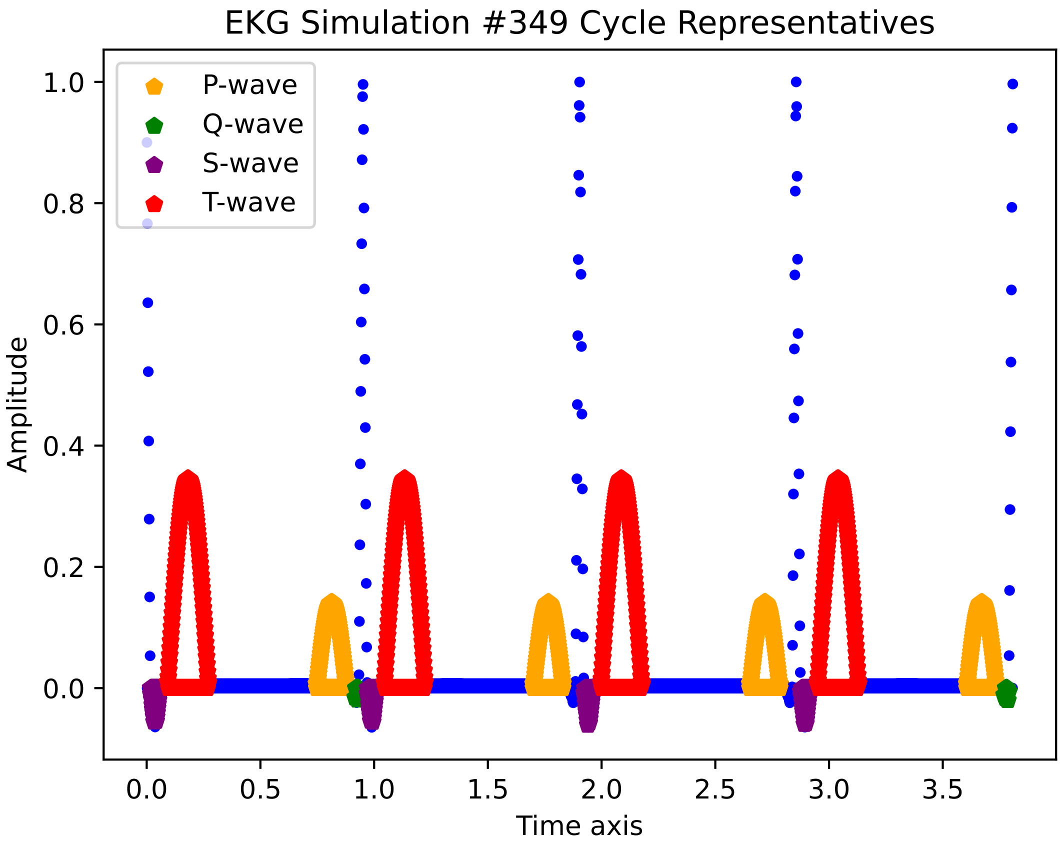
EKG Simulation #349 Cycle Representatives
P-wave
Q-wave
S-wave
T-wave
1.0
0.8
0.6
0.4
0.2
0.0
Amplitude
0.0
0.5
1.0
1.5
2.0
2.5
3.0
3.5
Time axis

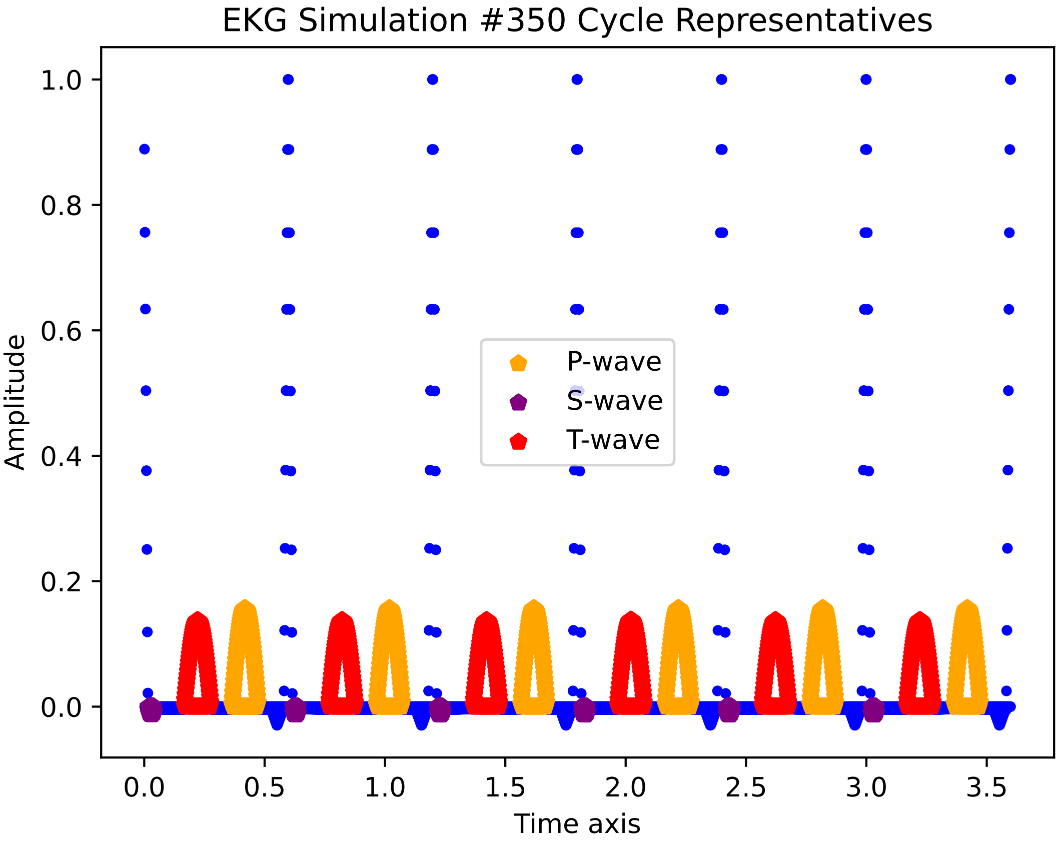

EKG Simulation #350 Cycle Representatives
Amplitude
Time axis
P-wave
S-wave
T-wave
1.0
0.8
0.6
0.4
0.2
0.0
0.0
0.5
1.0
1.5
2.0
2.5
3.0
3.5

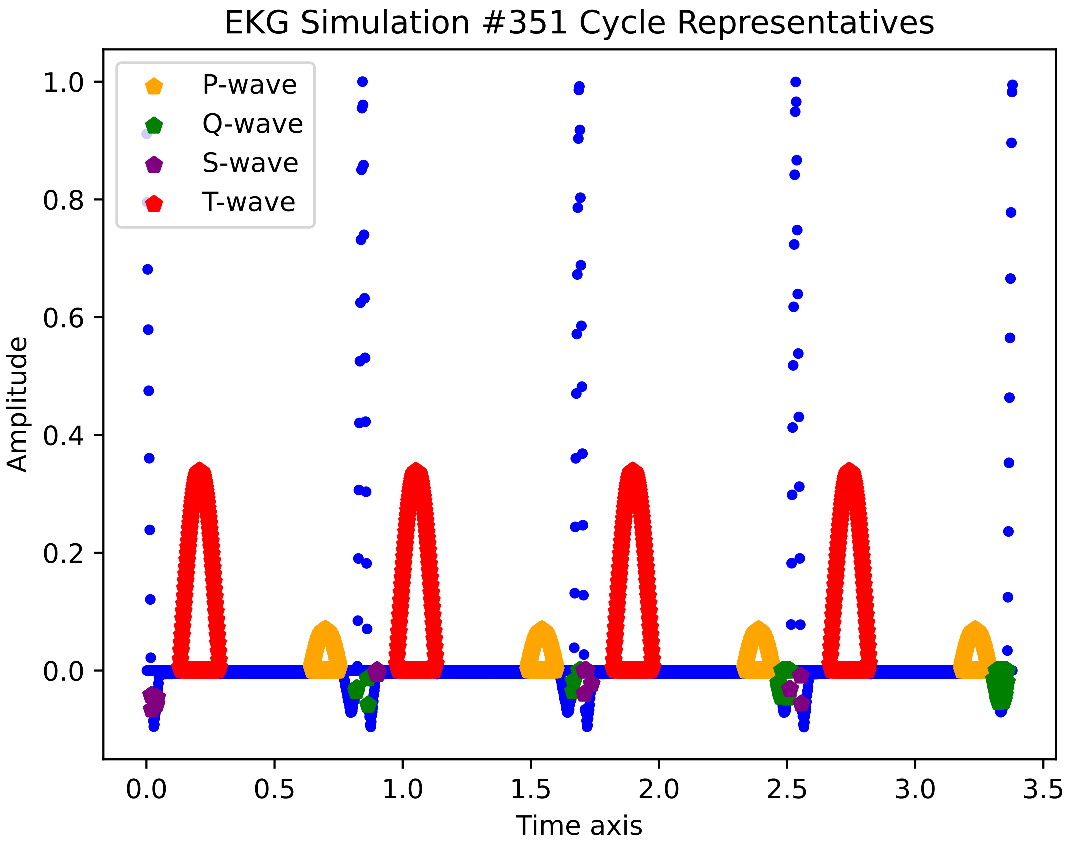
EKG Simulation #351 Cycle Representatives
P-wave
Q-wave
S-wave
T-wave
Amplitude
Time axis
1.0
0.8
0.6
0.4
0.2
0.0
0.0
0.5
1.0
1.5
2.0
2.5
3.0
3.5

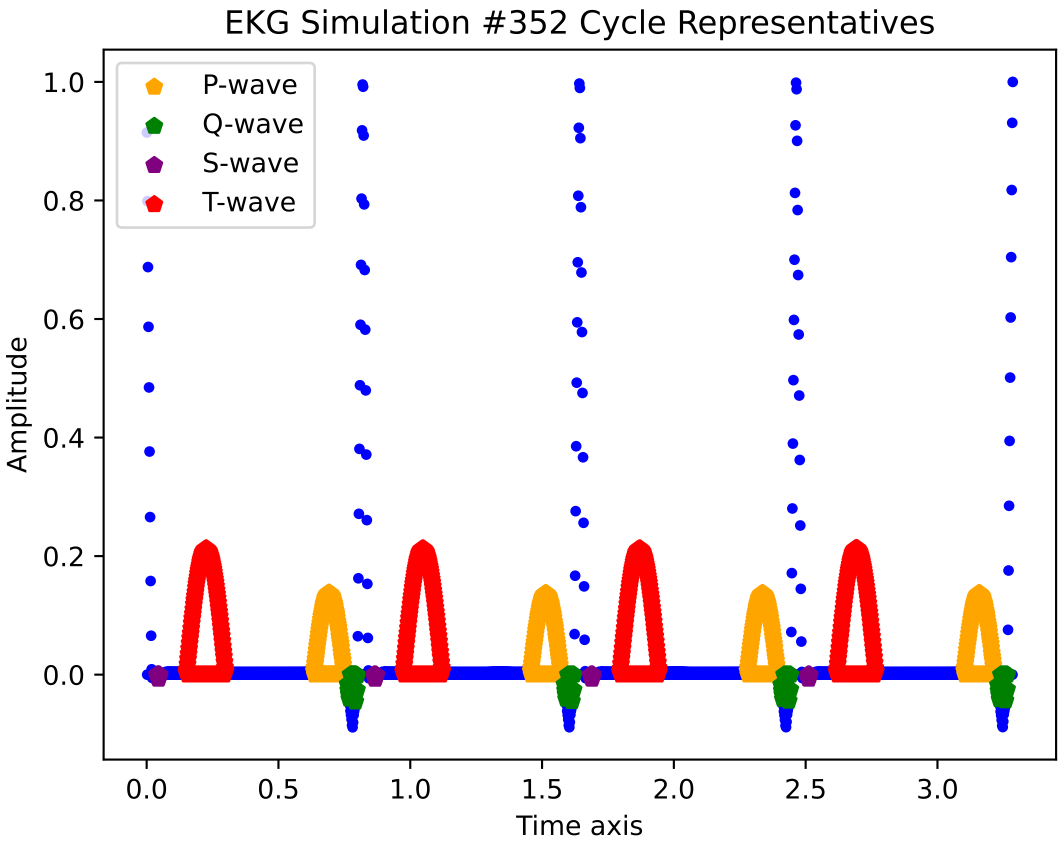

EKG Simulation #352 Cycle Representatives
P-wave
Q-wave
S-wave
T-wave
1.0
0.8
0.6
0.4
0.2
0.0
Amplitude
0.0
0.5
1.0
1.5
2.0
2.5
3.0
Time axis

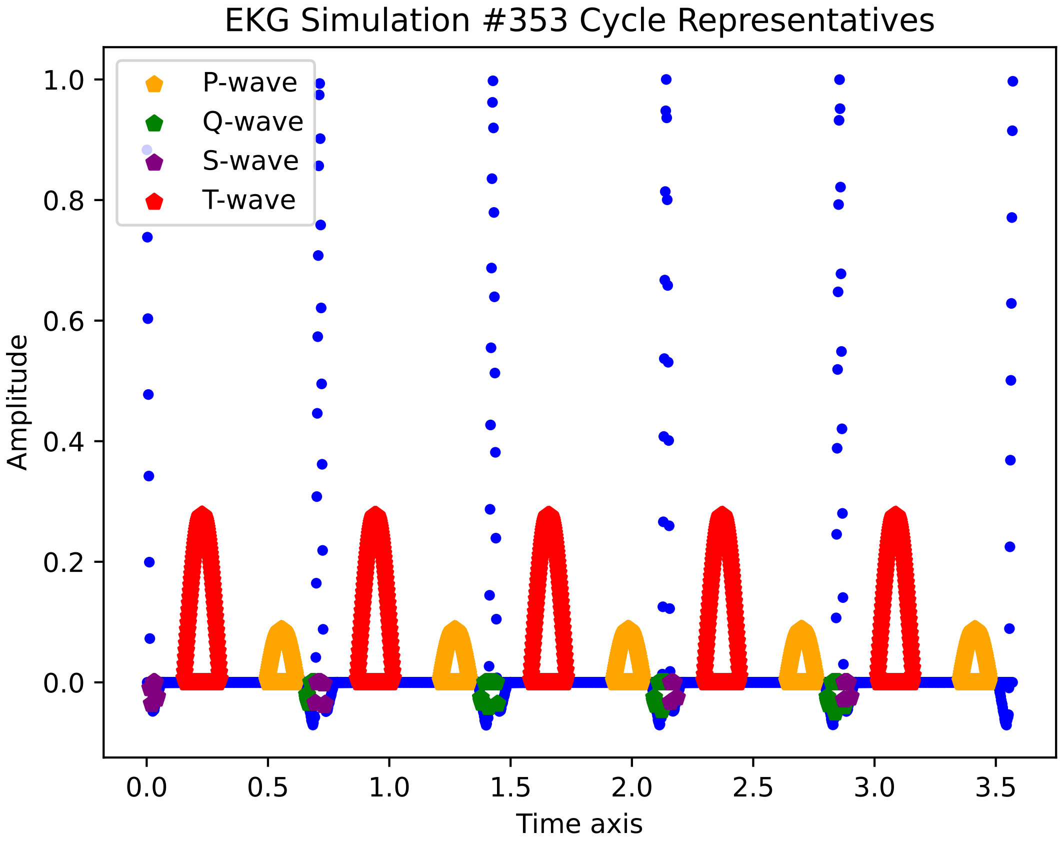
EKG Simulation #353 Cycle Representatives
P-wave
Q-wave
S-wave
T-wave
Amplitude
1.0
0.8
0.6
0.4
0.2
0.0
0.0
0.5
1.0
1.5
2.0
2.5
3.0
3.5
Time axis

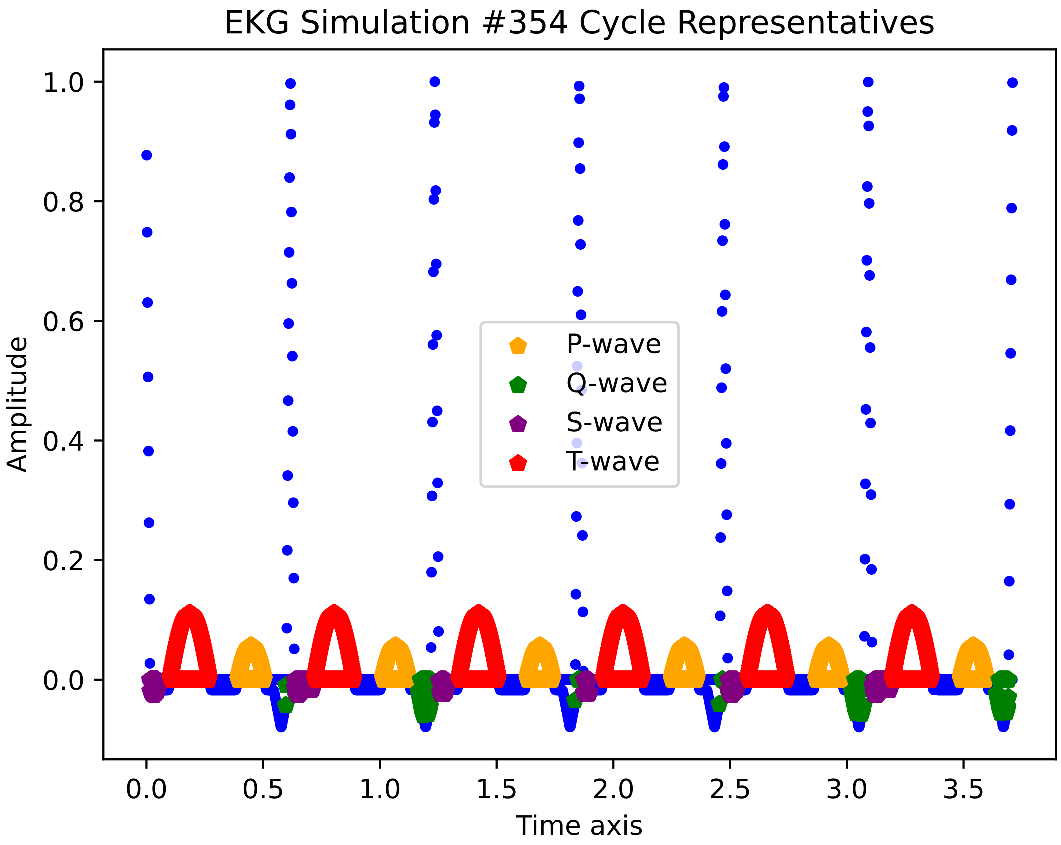
EKG Simulation #354 Cycle Representatives
Amplitude
Time axis
P-wave
Q-wave
S-wave
T-wave
1.0
0.8
0.6
0.4
0.2
0.0
0.0
0.5
1.0
1.5
2.0
2.5
3.0
3.5

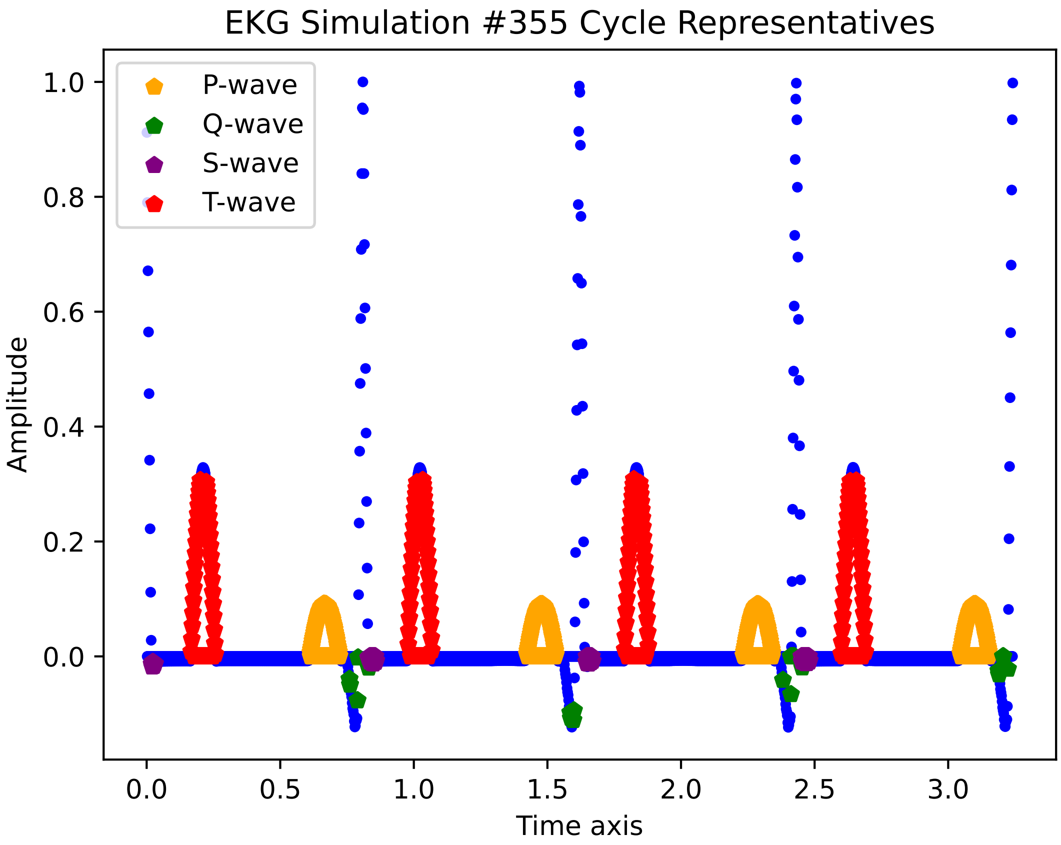
EKG Simulation #355 Cycle Representatives
P-wave
Q-wave
S-wave
T-wave
Amplitude
Time axis
1.0
0.8
0.6
0.4
0.2
0.0
0.0
0.5
1.0
1.5
2.0
2.5
3.0

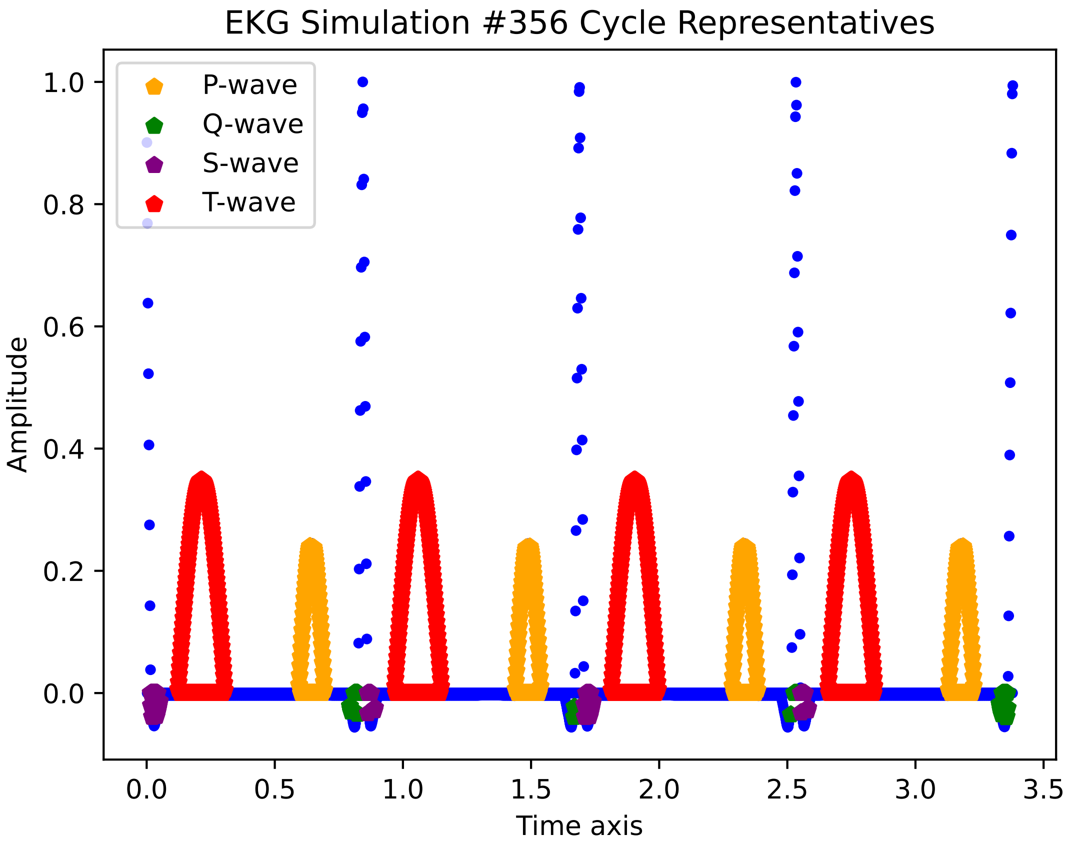

EKG Simulation #356 Cycle Representatives
P-wave
Q-wave
S-wave
T-wave
Amplitude
Time axis
0.0
0.2
0.4
0.6
0.8
1.0
0.0
0.5
1.0
1.5
2.0
2.5
3.0
3.5

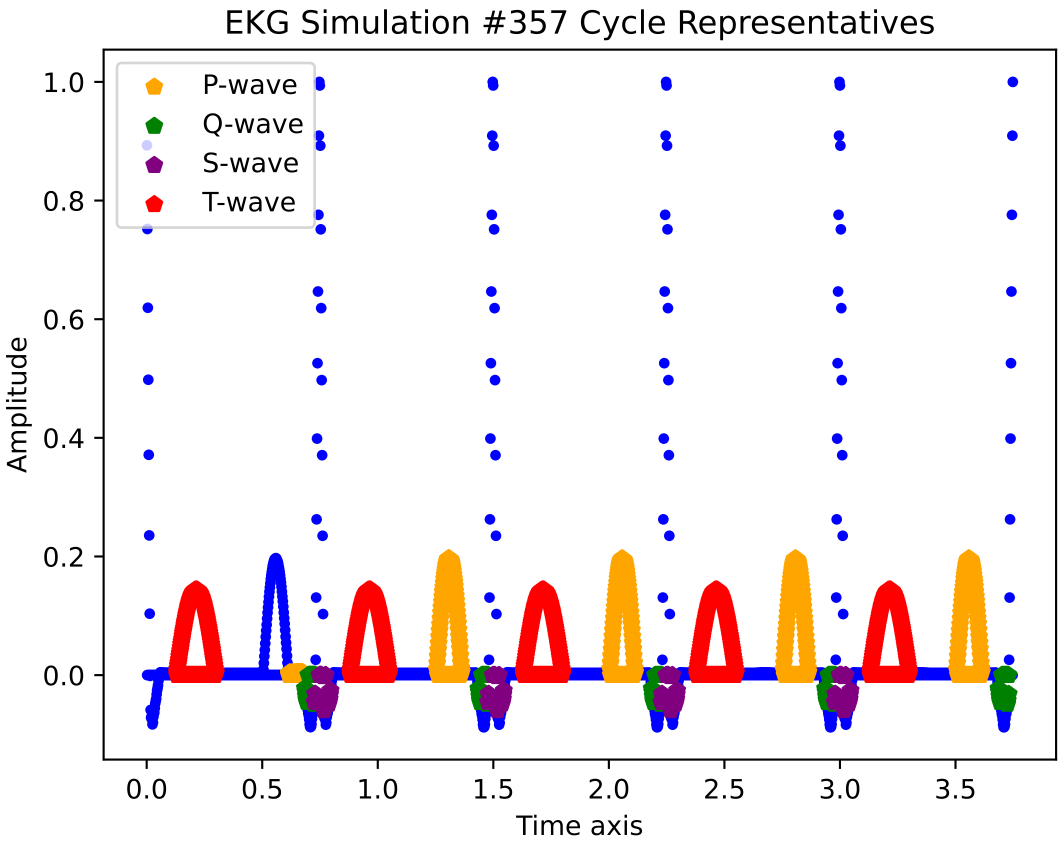

EKG Simulation #357 Cycle Representatives
P-wave
Q-wave
S-wave
T-wave
Amplitude
Time axis
1.0
0.8
0.6
0.4
0.2
0.0
0.0
0.5
1.0
1.5
2.0
2.5
3.0
3.5

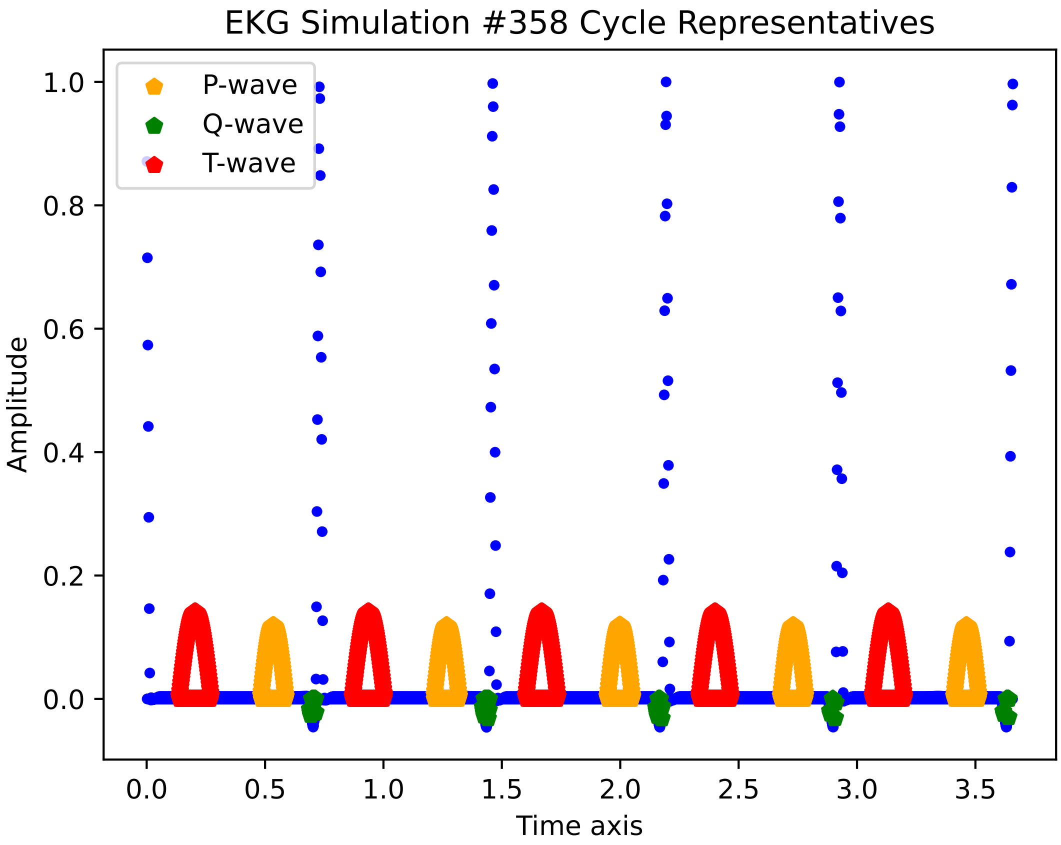

EKG Simulation #358 Cycle Representatives
P-wave
Q-wave
T-wave
Amplitude
Time axis
1.0
0.8
0.6
0.4
0.2
0.0
0.0
0.5
1.0
1.5
2.0
2.5
3.0
3.5

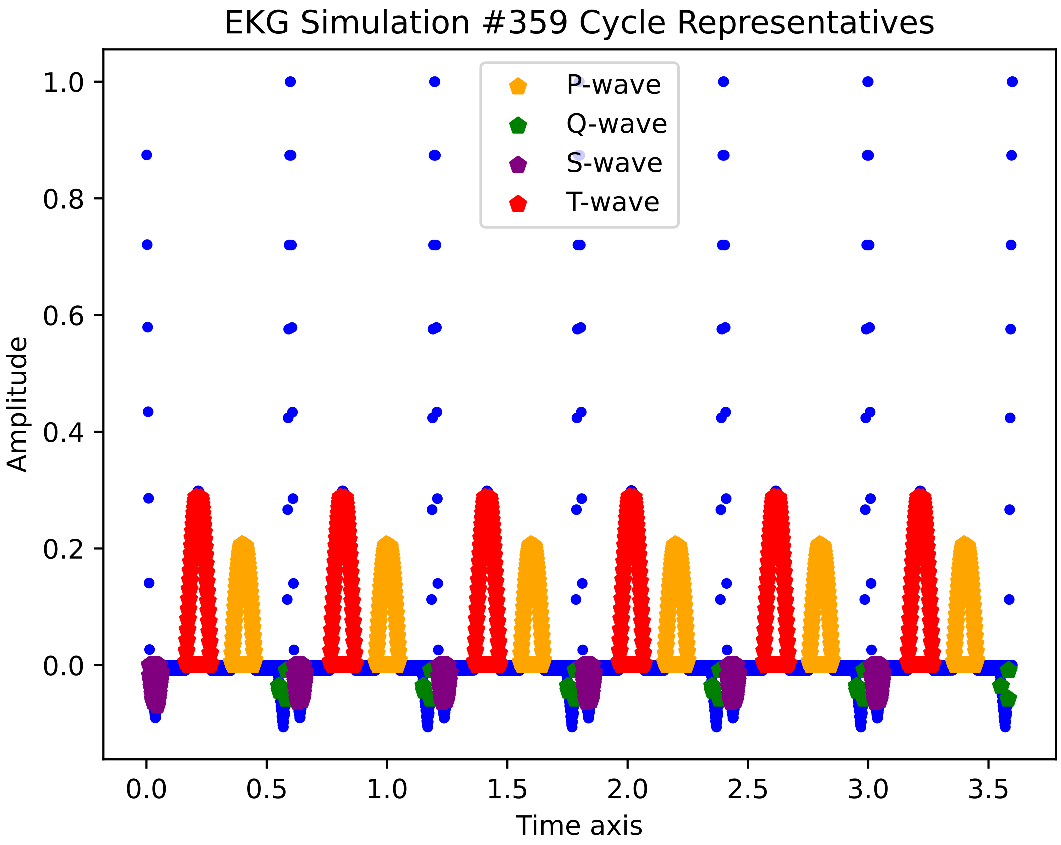

EKG Simulation #359 Cycle Representatives
P-wave
Q-wave
S-wave
T-wave
Amplitude
Time axis
1.0
0.8
0.6
0.4
0.2
0.0
0.0
0.5
1.0
1.5
2.0
2.5
3.0
3.5

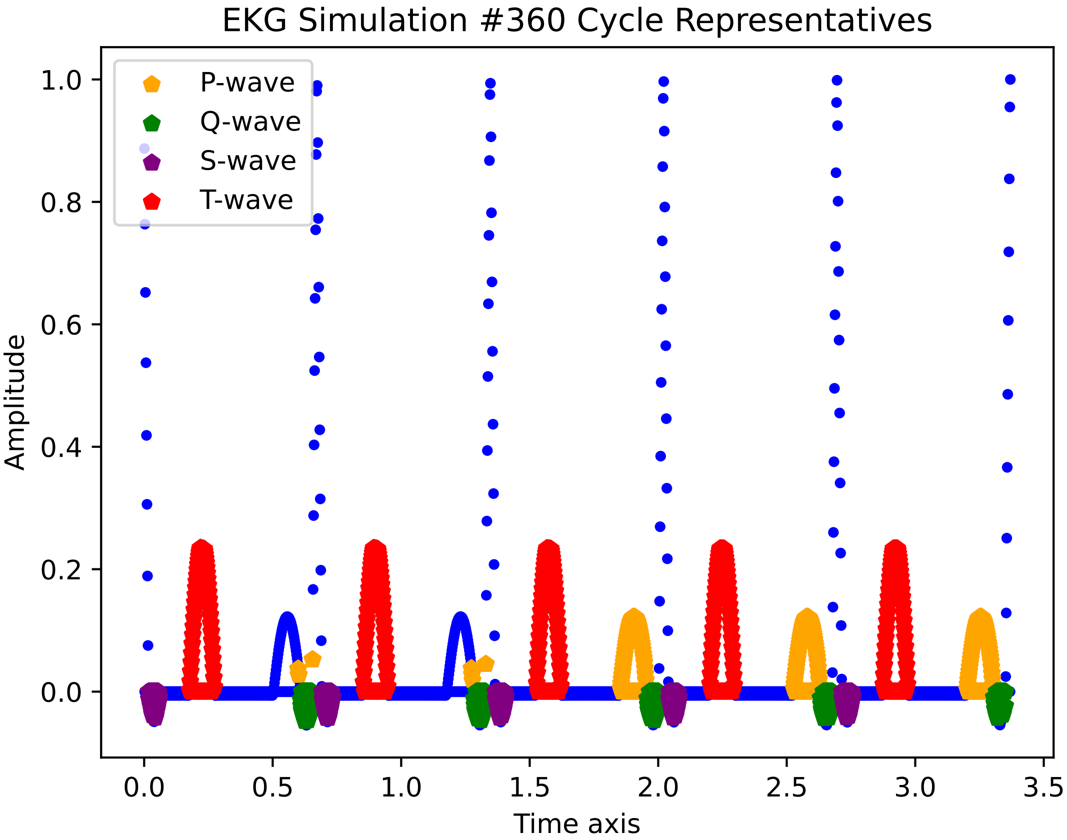

EKG Simulation #360 Cycle Representatives
P-wave
Q-wave
S-wave
T-wave
Amplitude
Time axis
1.0
0.8
0.6
0.4
0.2
0.0
0.0
0.5
1.0
1.5
2.0
2.5
3.0
3.5

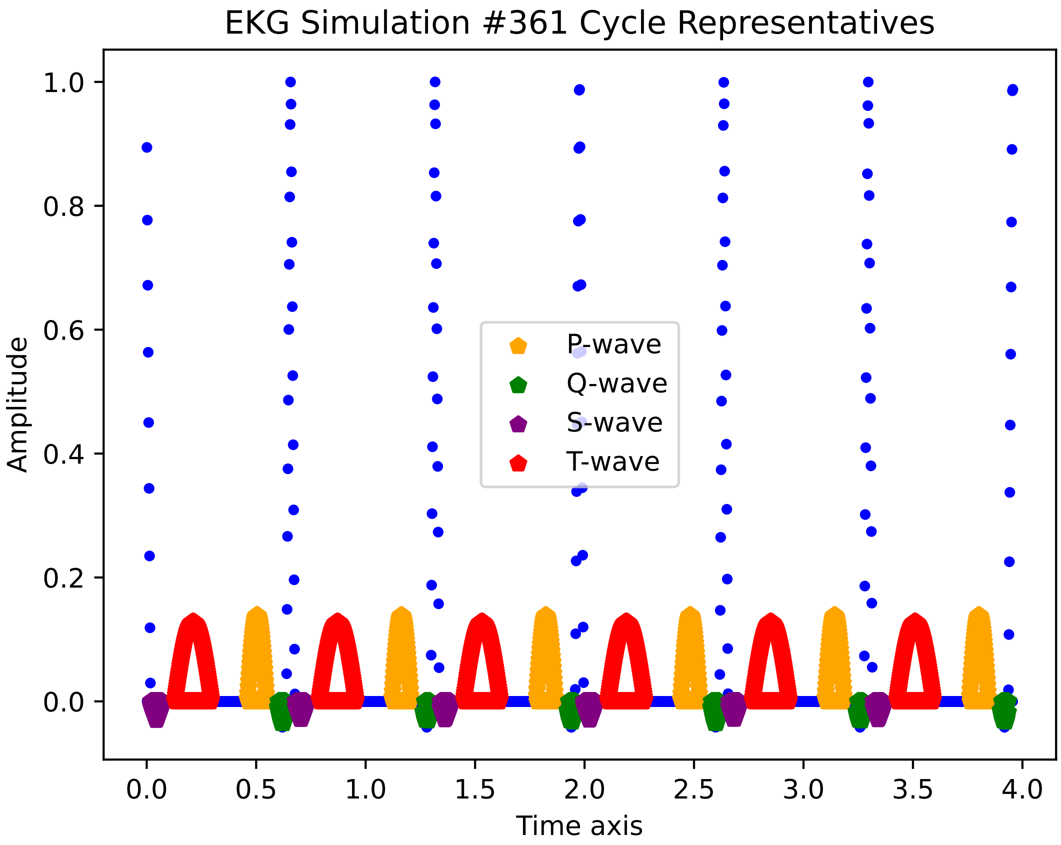
EKG Simulation #361 Cycle Representatives
Amplitude
Time axis
P-wave
Q-wave
S-wave
T-wave
1.0
0.8
0.6
0.4
0.2
0.0
0.0
0.5
1.0
1.5
2.0
2.5
3.0
3.5
4.0

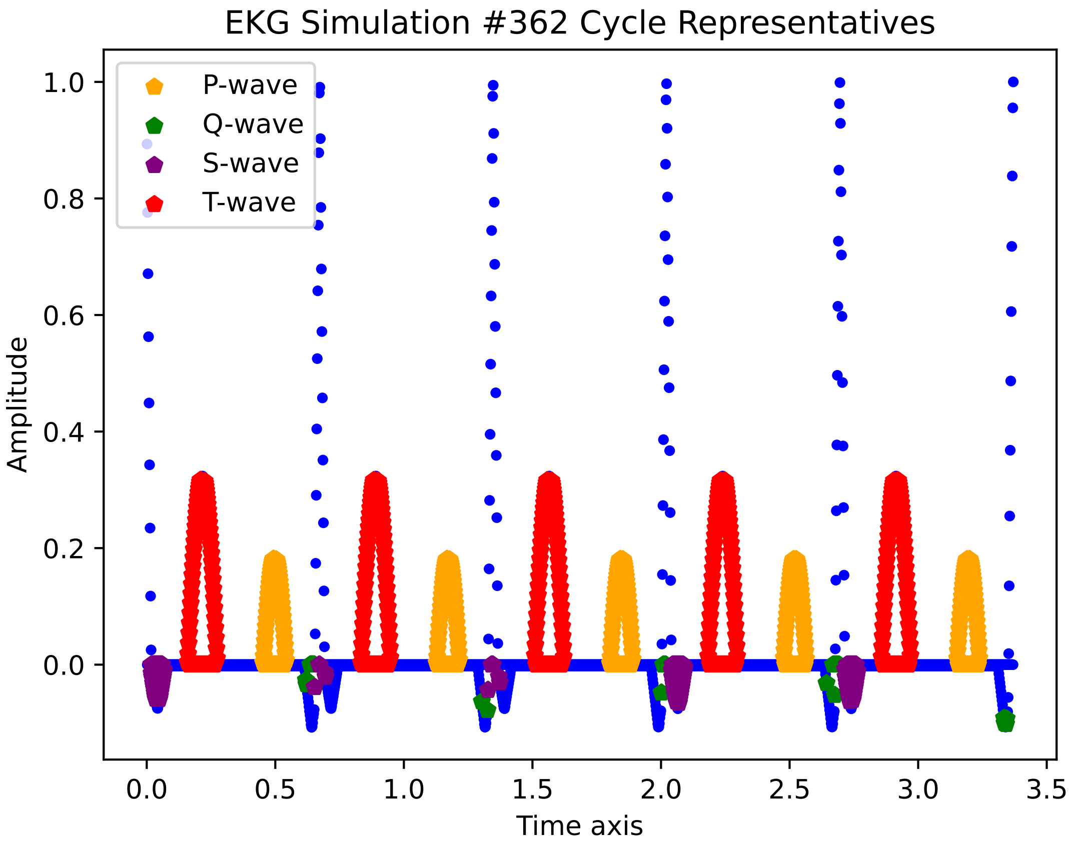

EKG Simulation #362 Cycle Representatives
P-wave
Q-wave
S-wave
T-wave
Amplitude
Time axis
1.0
0.8
0.6
0.4
0.2
0.0
0.0
0.5
1.0
1.5
2.0
2.5
3.0
3.5

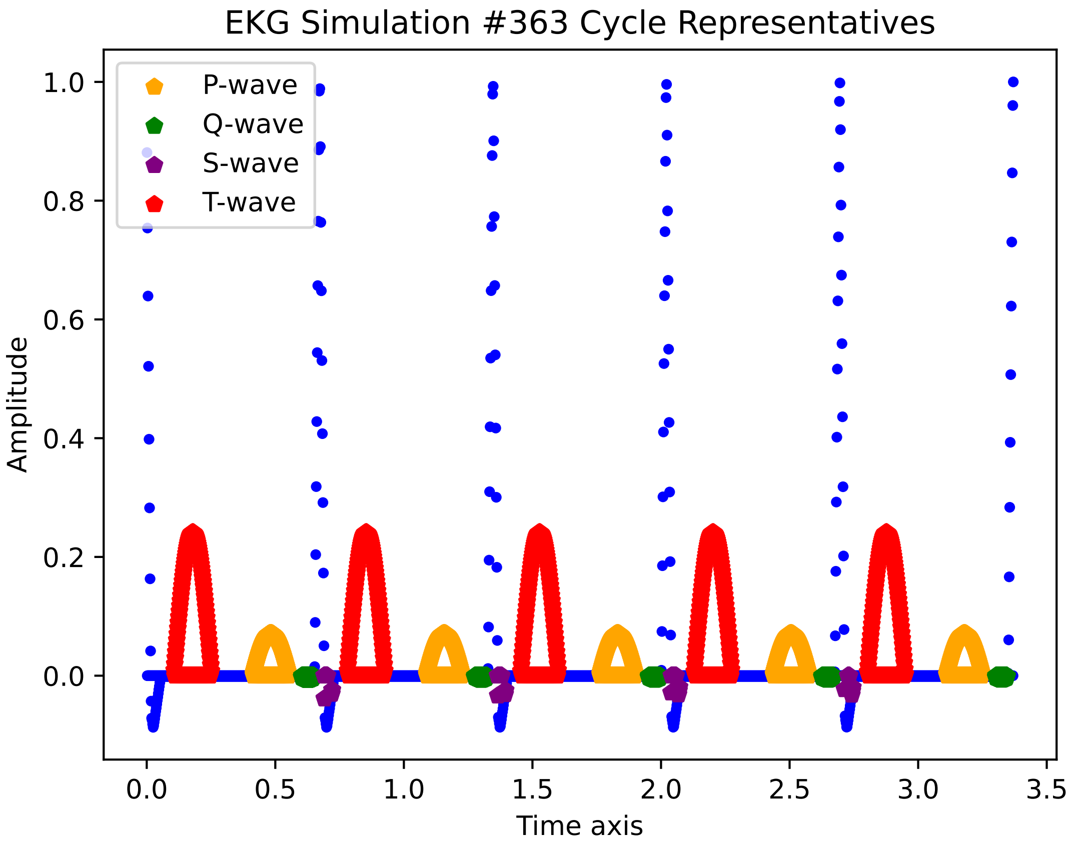
EKG Simulation #363 Cycle Representatives
P-wave
Q-wave
S-wave
T-wave
1.0
0.8
0.6
0.4
0.2
0.0
Amplitude
0.0
0.5
1.0
1.5
2.0
2.5
3.0
3.5
Time axis

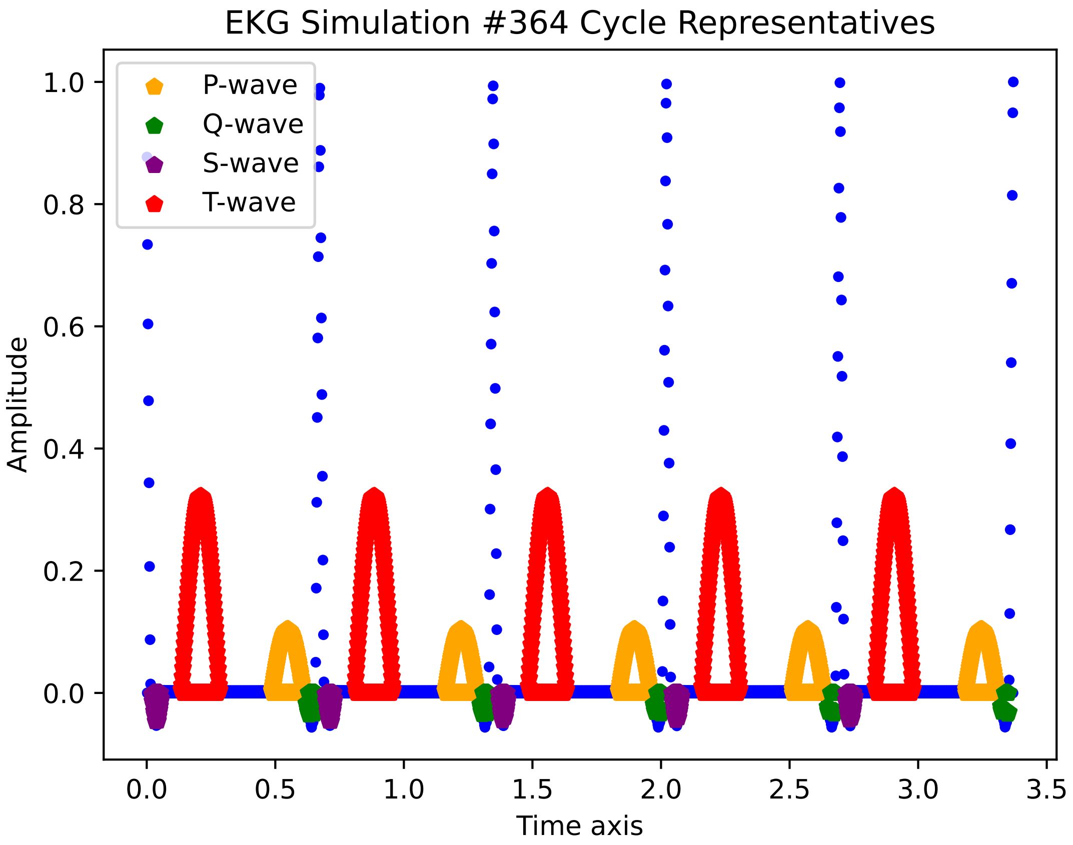
EKG Simulation #364 Cycle Representatives
P-wave
Q-wave
S-wave
T-wave
Amplitude
Time axis
0.0
0.2
0.4
0.6
0.8
1.0
0.0
0.5
1.0
1.5
2.0
2.5
3.0
3.5

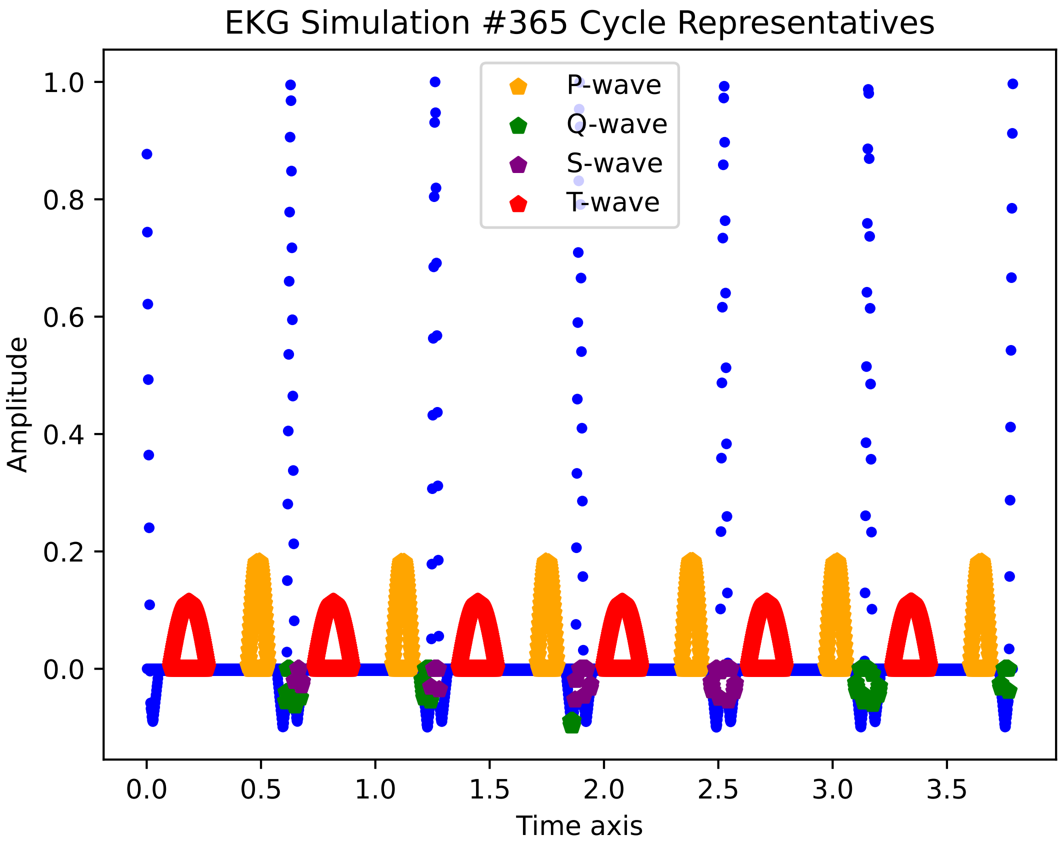
EKG Simulation #365 Cycle Representatives
P-wave
Q-wave
S-wave
T-wave
Amplitude
1.0
0.8
0.6
0.4
0.2
0.0
0.0
0.5
1.0
1.5
2.0
2.5
3.0
3.5
Time axis

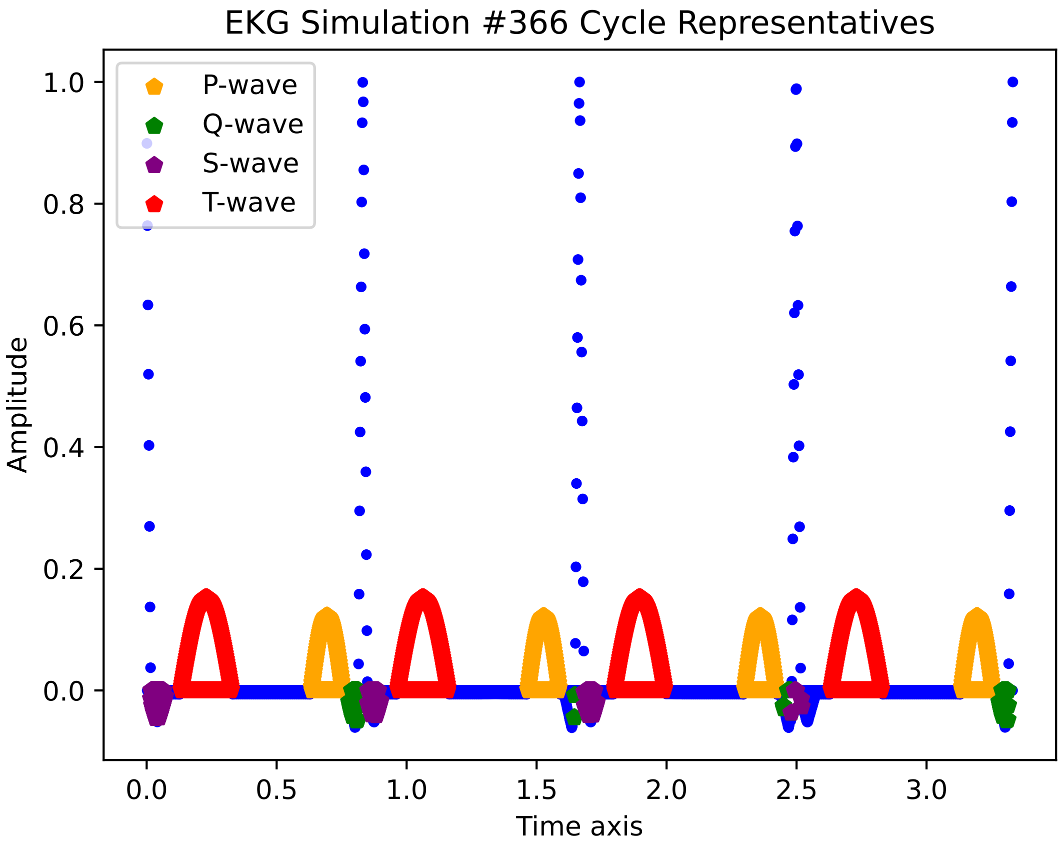

EKG Simulation #366 Cycle Representatives
P-wave
Q-wave
S-wave
T-wave
1.0
0.8
0.6
0.4
0.2
0.0
Amplitude
0.0
0.5
1.0
1.5
2.0
2.5
3.0
Time axis

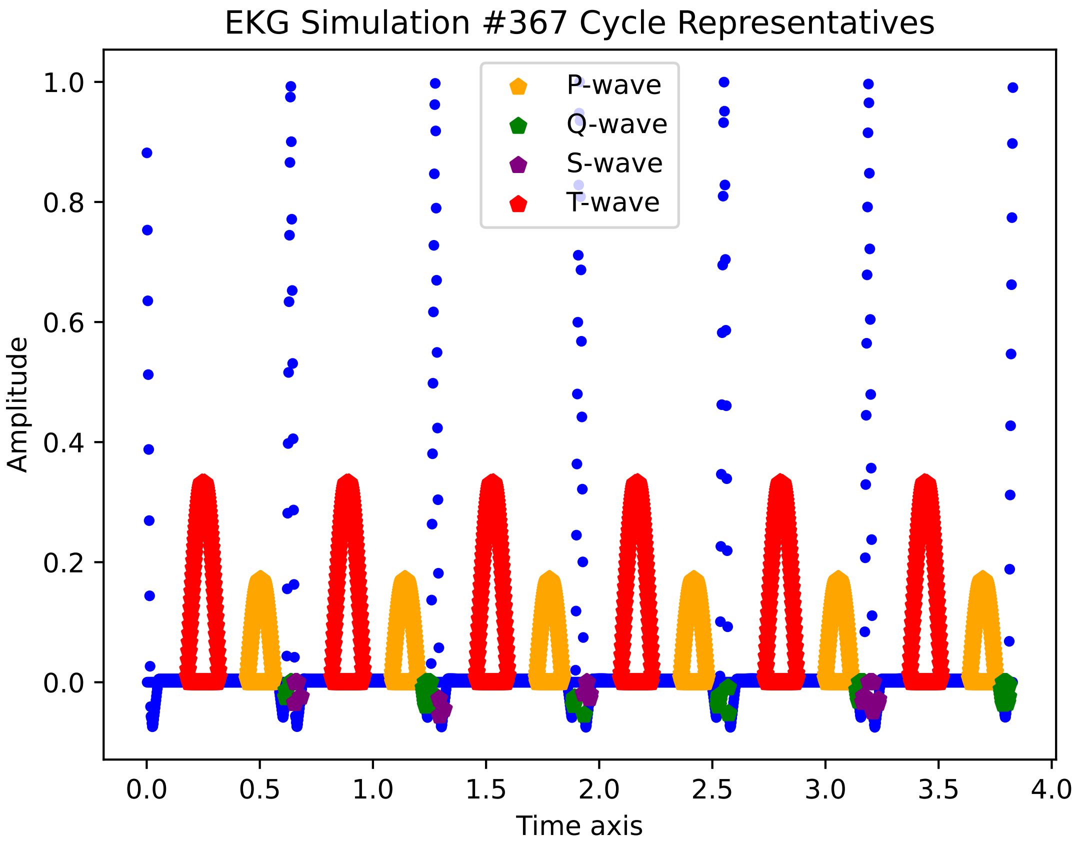

EKG Simulation #367 Cycle Representatives
P-wave
Q-wave
S-wave
T-wave
Amplitude
Time axis
1.0
0.8
0.6
0.4
0.2
0.0
0.0
0.5
1.0
1.5
2.0
2.5
3.0
3.5
4.0

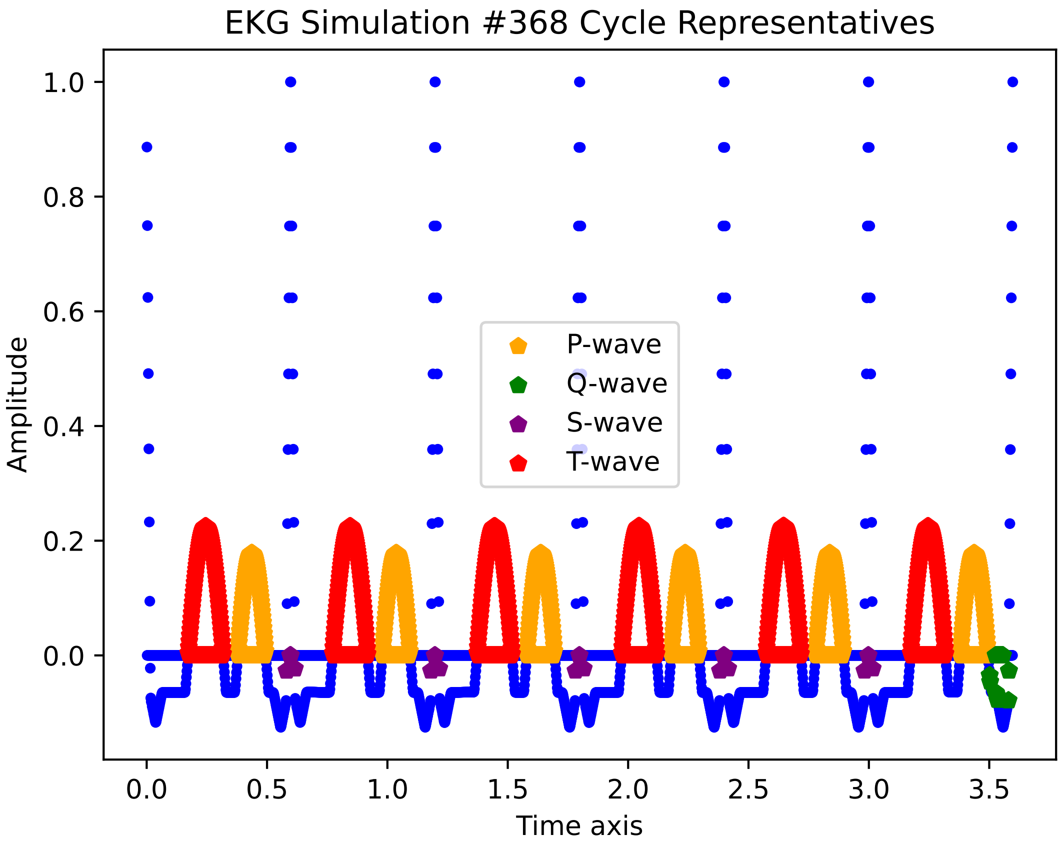
EKG Simulation #368 Cycle Representatives
Amplitude
Time axis
1.0
0.8
0.6
0.4
0.2
0.0
0.0
0.5
1.0
1.5
2.0
2.5
3.0
3.5
P-wave
Q-wave
S-wave
T-wave

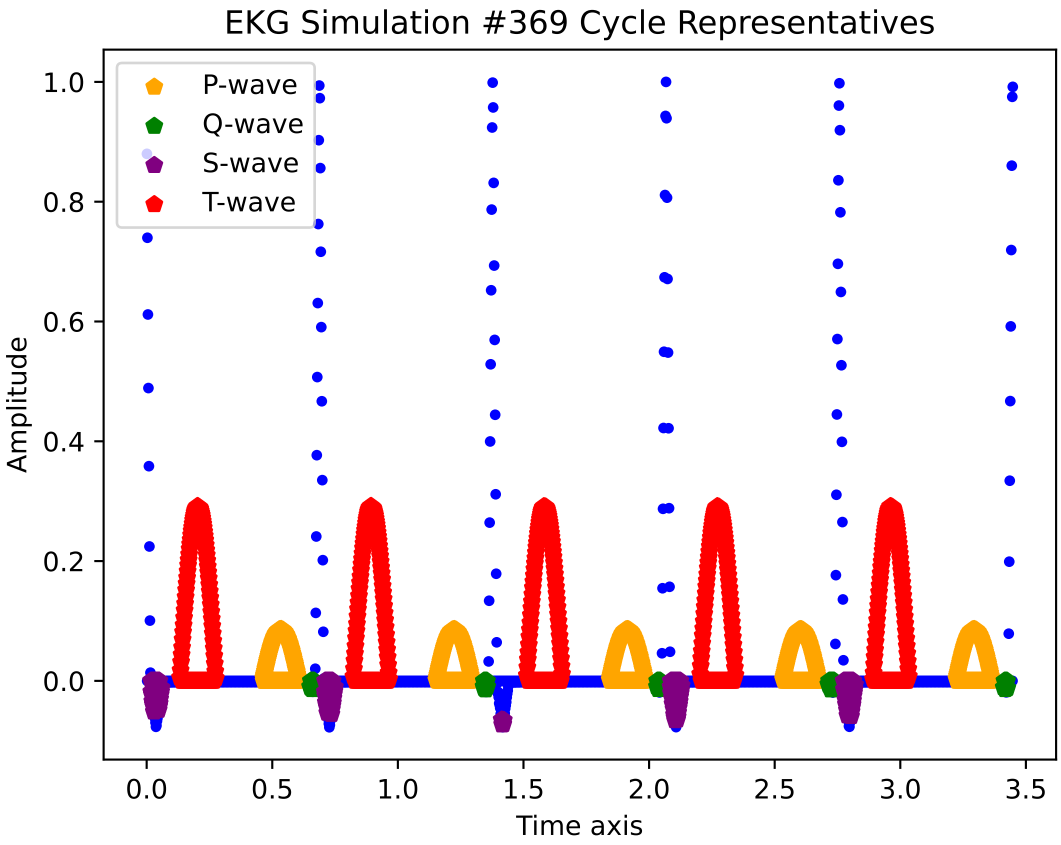

EKG Simulation #369 Cycle Representatives
P-wave
Q-wave
S-wave
T-wave
Amplitude
Time axis
1.0
0.8
0.6
0.4
0.2
0.0
0.0
0.5
1.0
1.5
2.0
2.5
3.0
3.5

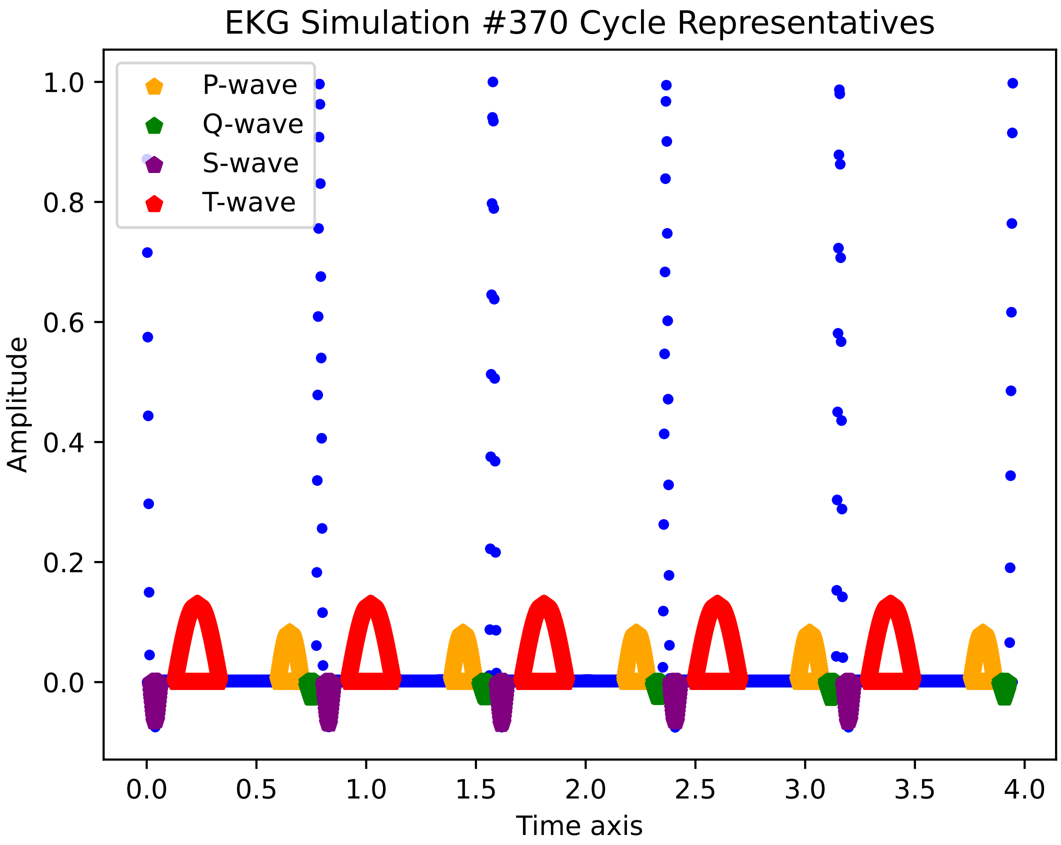
EKG Simulation #370 Cycle Representatives
P-wave
Q-wave
S-wave
T-wave
Amplitude
Time axis
1.0
0.8
0.6
0.4
0.2
0.0
0.0
0.5
1.0
1.5
2.0
2.5
3.0
3.5
4.0

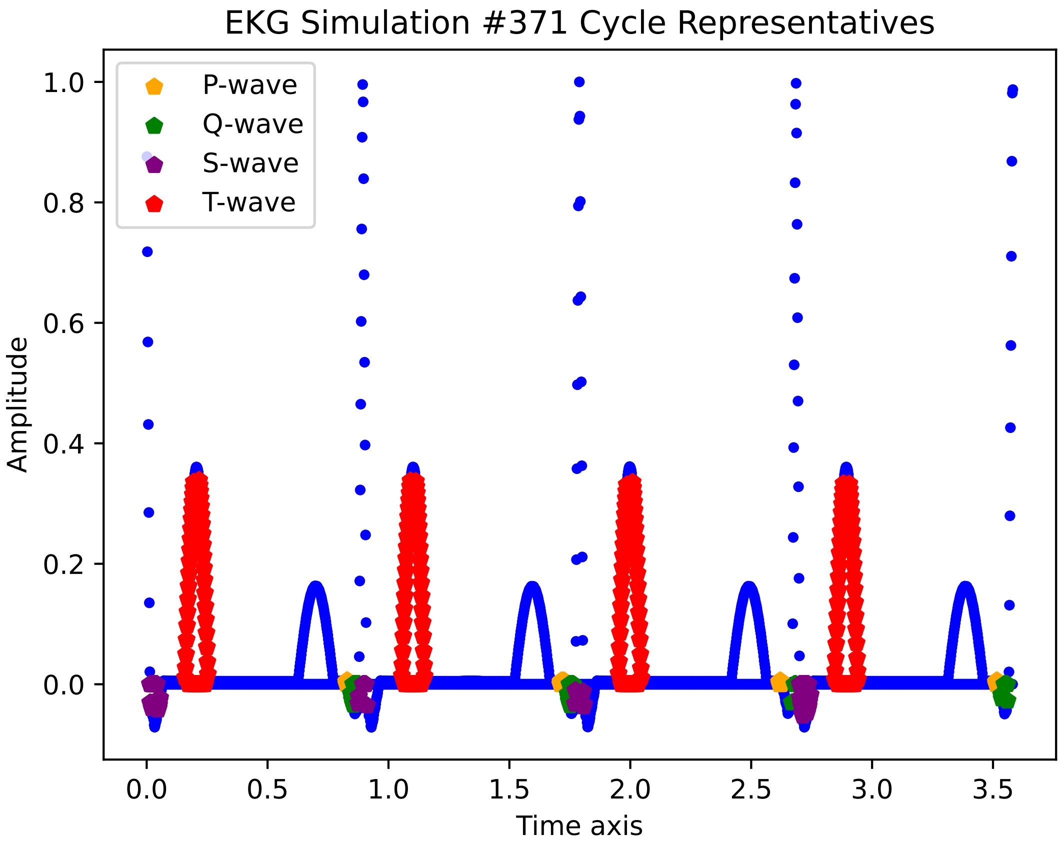

EKG Simulation #371 Cycle Representatives
P-wave
Q-wave
S-wave
T-wave
Amplitude
Time axis
1.0
0.8
0.6
0.4
0.2
0.0
0.0
0.5
1.0
1.5
2.0
2.5
3.0
3.5

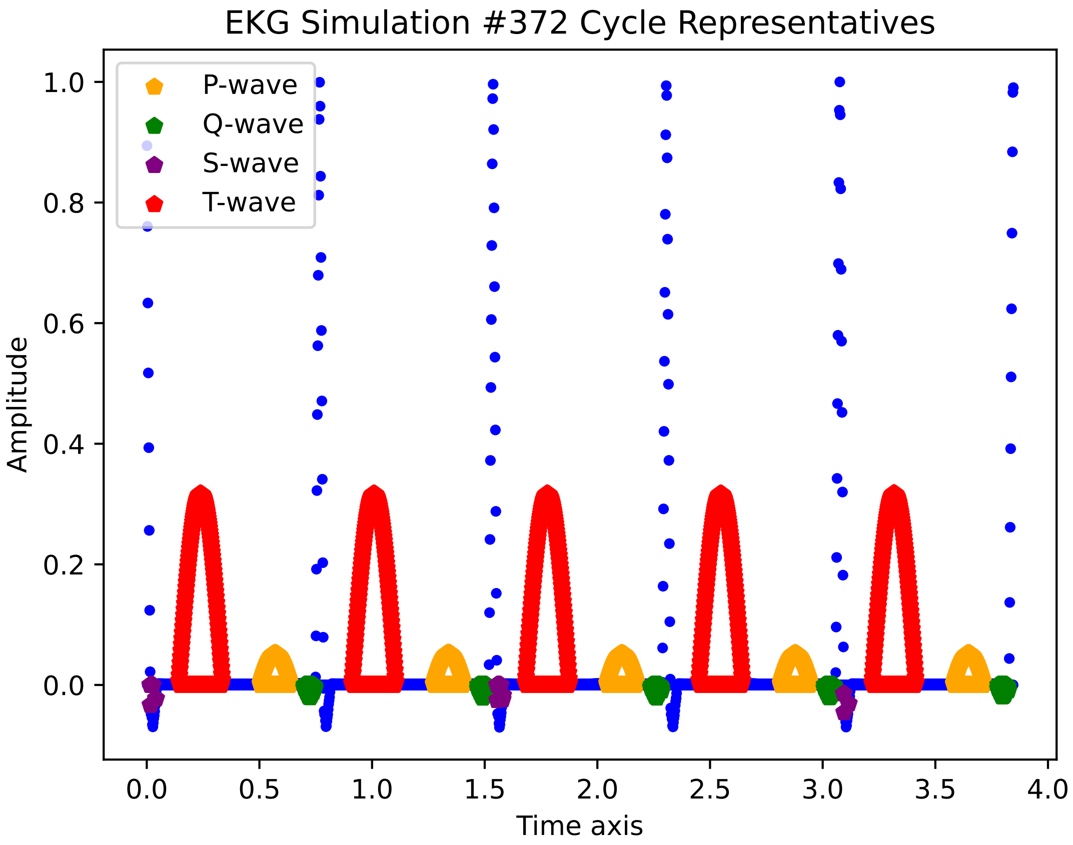
EKG Simulation #372 Cycle Representatives
P-wave
Q-wave
S-wave
T-wave
Amplitude
Time axis
1.0
0.8
0.6
0.4
0.2
0.0
0.0
0.5
1.0
1.5
2.0
2.5
3.0
3.5
4.0

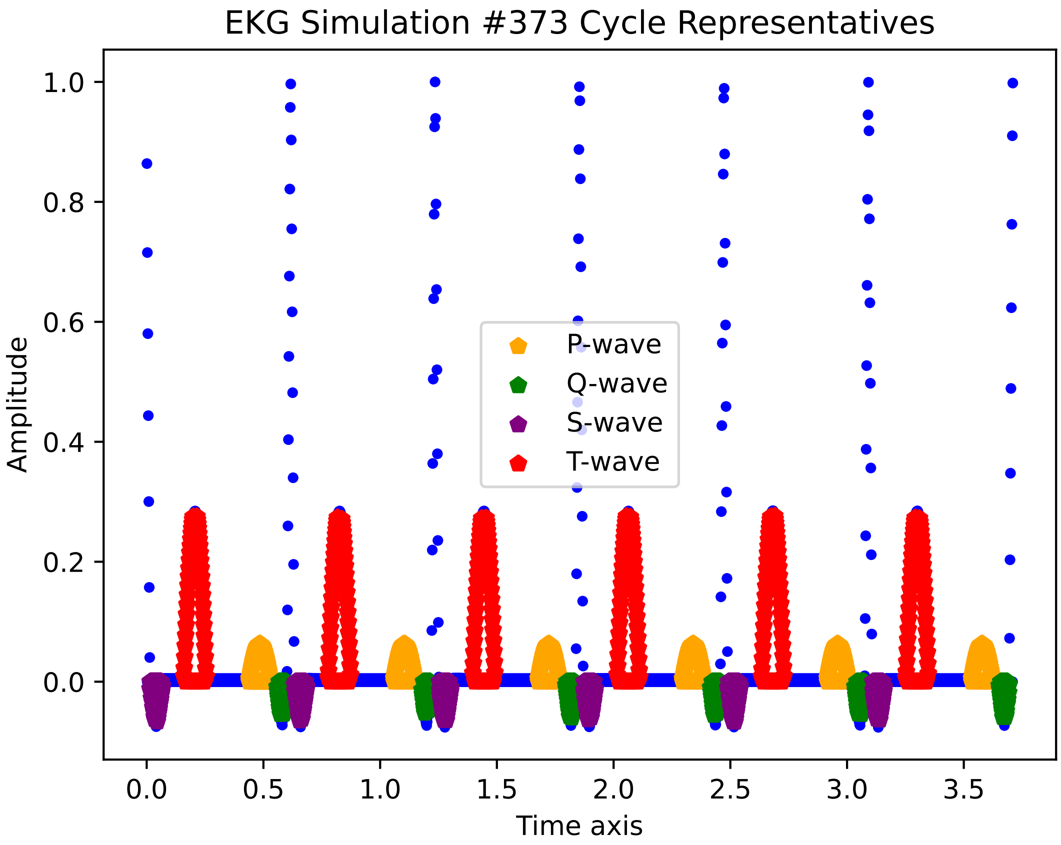
EKG Simulation #373 Cycle Representatives
1.0
0.8
0.6
0.4
0.2
0.0
Amplitude
0.0
0.5
1.0
1.5
2.0
2.5
3.0
3.5
Time axis
P-wave
Q-wave
S-wave
T-wave

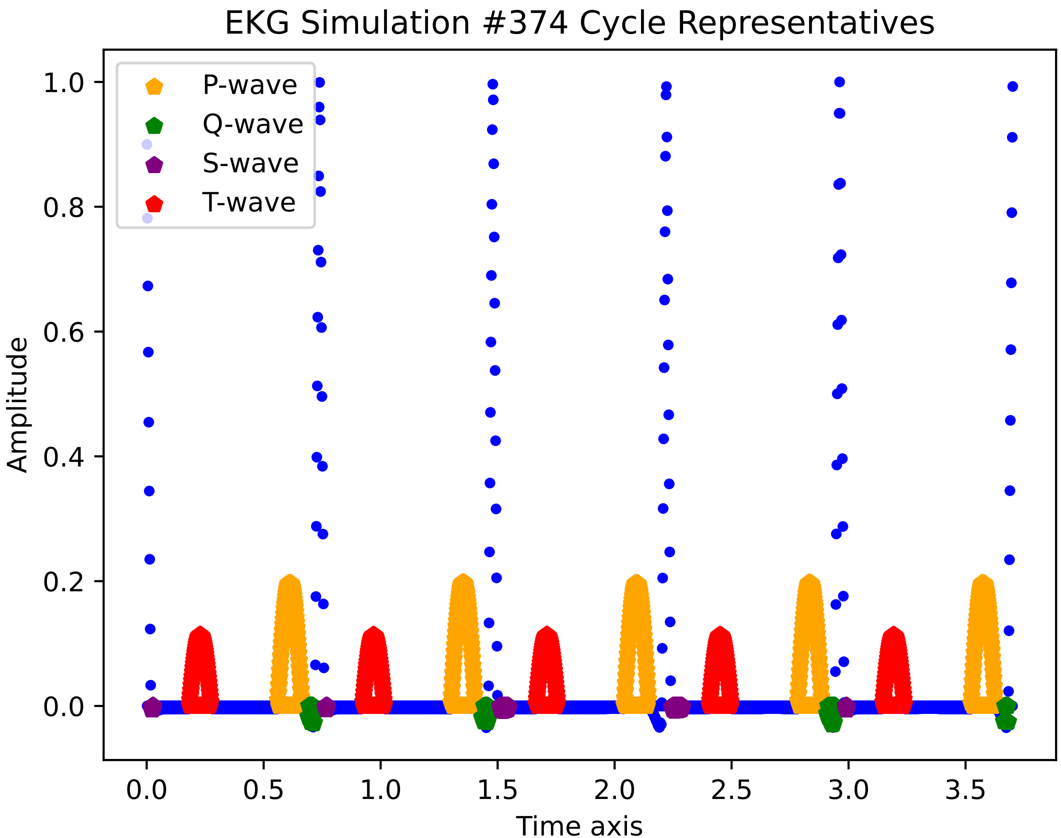

EKG Simulation #374 Cycle Representatives
P-wave
Q-wave
S-wave
T-wave
Amplitude
Time axis
1.0
0.8
0.6
0.4
0.2
0.0
0.0
0.5
1.0
1.5
2.0
2.5
3.0
3.5

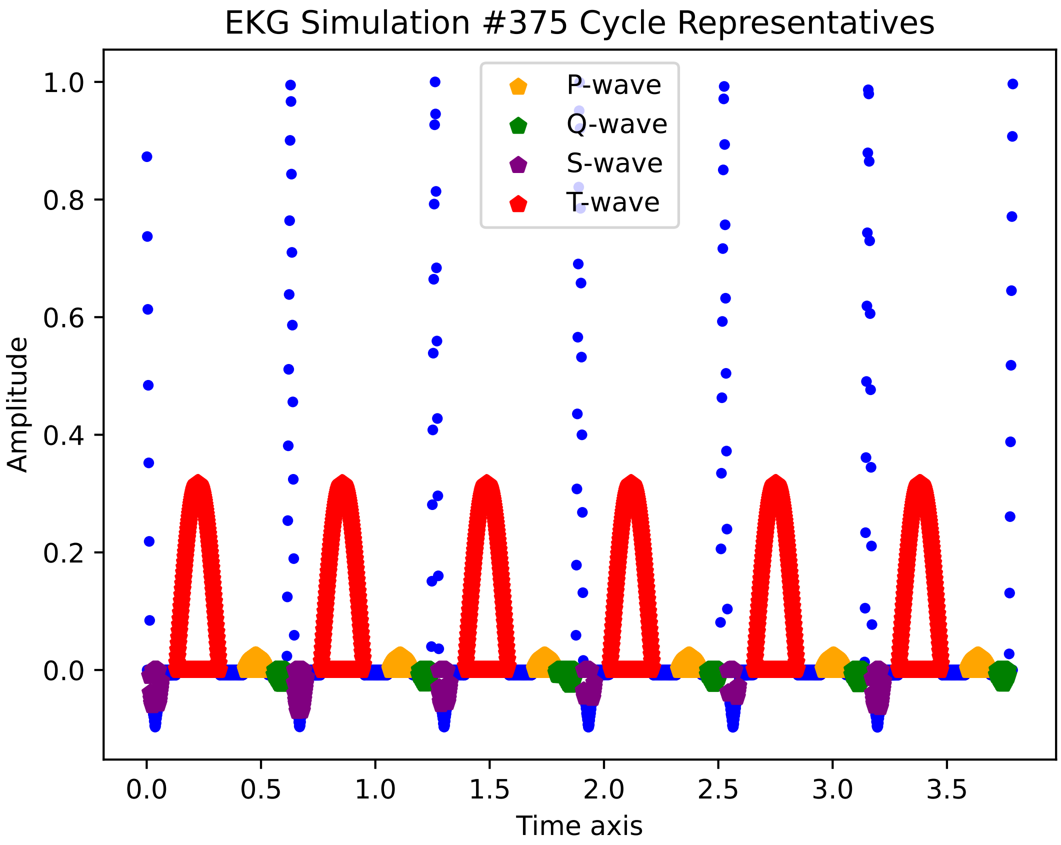

EKG Simulation #375 Cycle Representatives
P-wave
Q-wave
S-wave
T-wave
Amplitude
Time axis
1.0
0.8
0.6
0.4
0.2
0.0
0.0
0.5
1.0
1.5
2.0
2.5
3.0
3.5

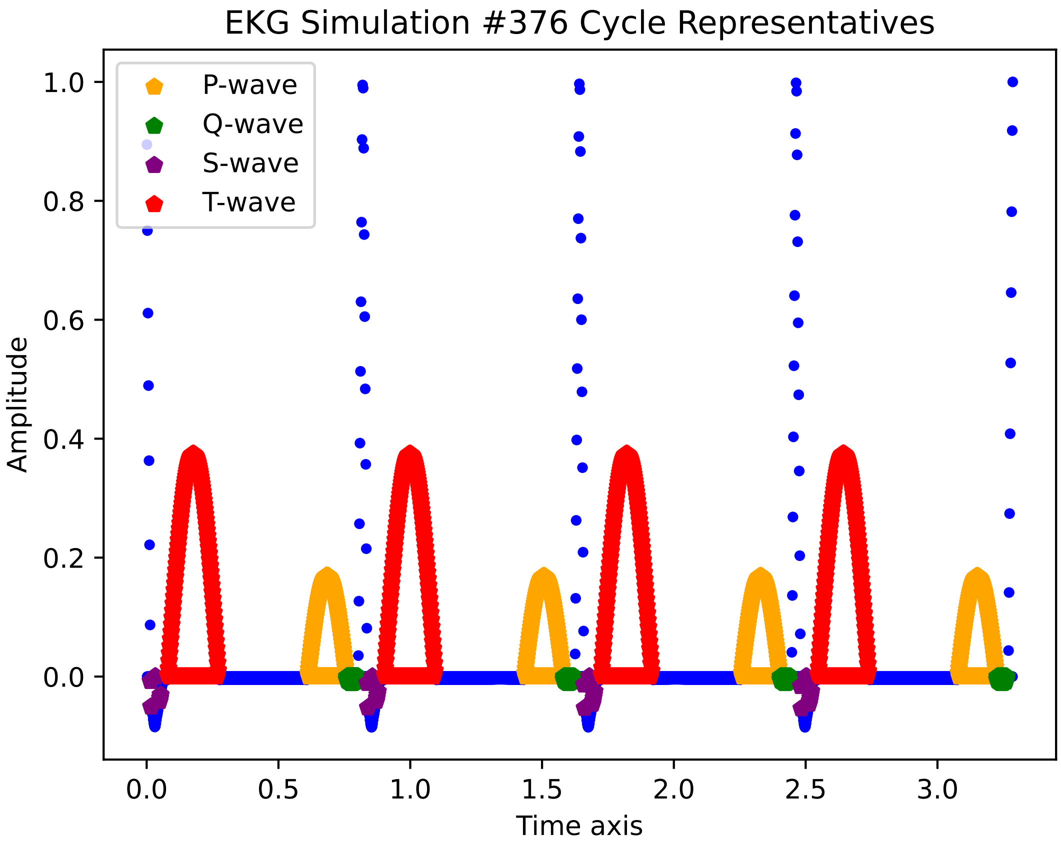
EKG Simulation #376 Cycle Representatives
P-wave
Q-wave
S-wave
T-wave
Amplitude
Time axis
1.0
0.8
0.6
0.4
0.2
0.0
0.0
0.5
1.0
1.5
2.0
2.5
3.0

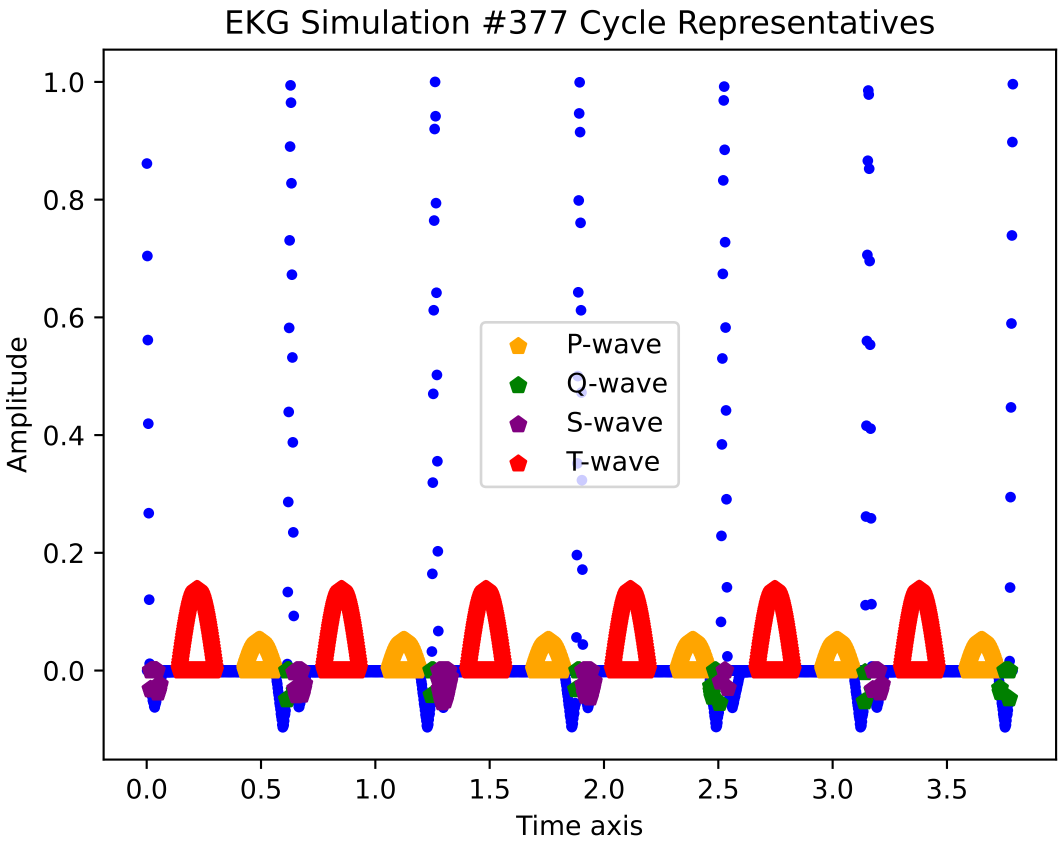

EKG Simulation #377 Cycle Representatives
Amplitude
Time axis
1.0
0.8
0.6
0.4
0.2
0.0
0.0
0.5
1.0
1.5
2.0
2.5
3.0
3.5
P-wave
Q-wave
S-wave
T-wave

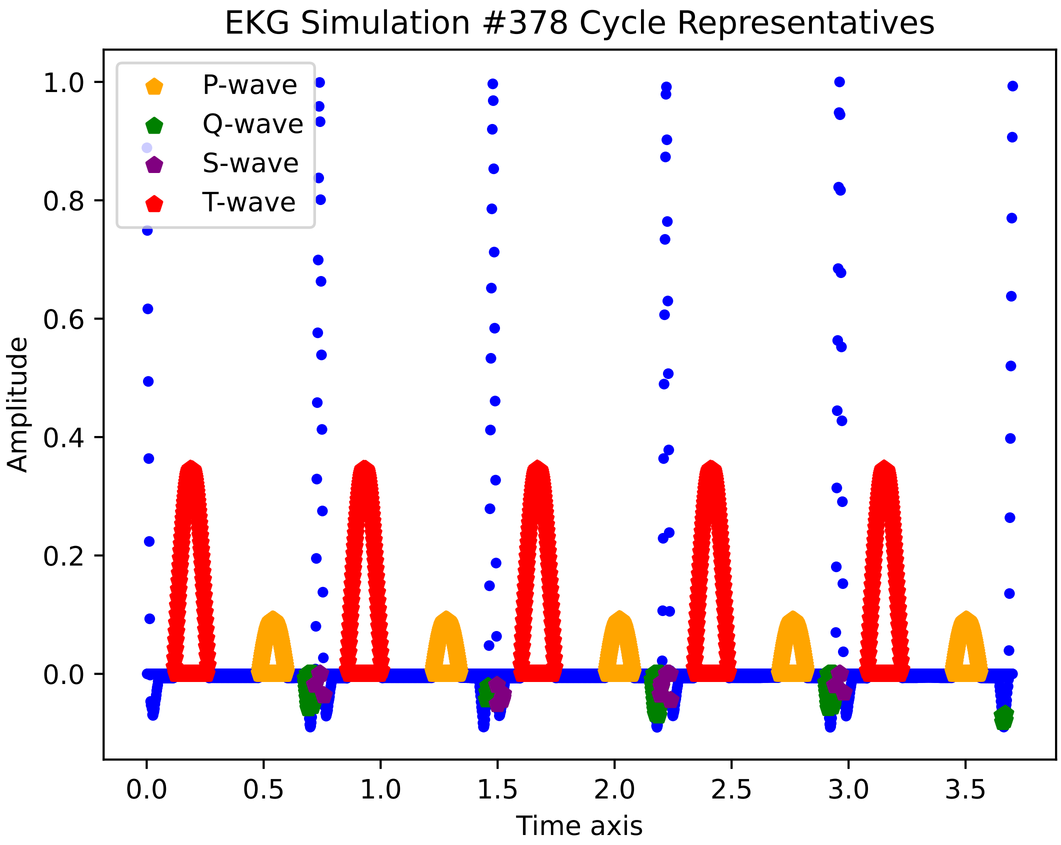
EKG Simulation #378 Cycle Representatives
P-wave
Q-wave
S-wave
T-wave
Amplitude
Time axis
1.0
0.8
0.6
0.4
0.2
0.0
0.0
0.5
1.0
1.5
2.0
2.5
3.0
3.5

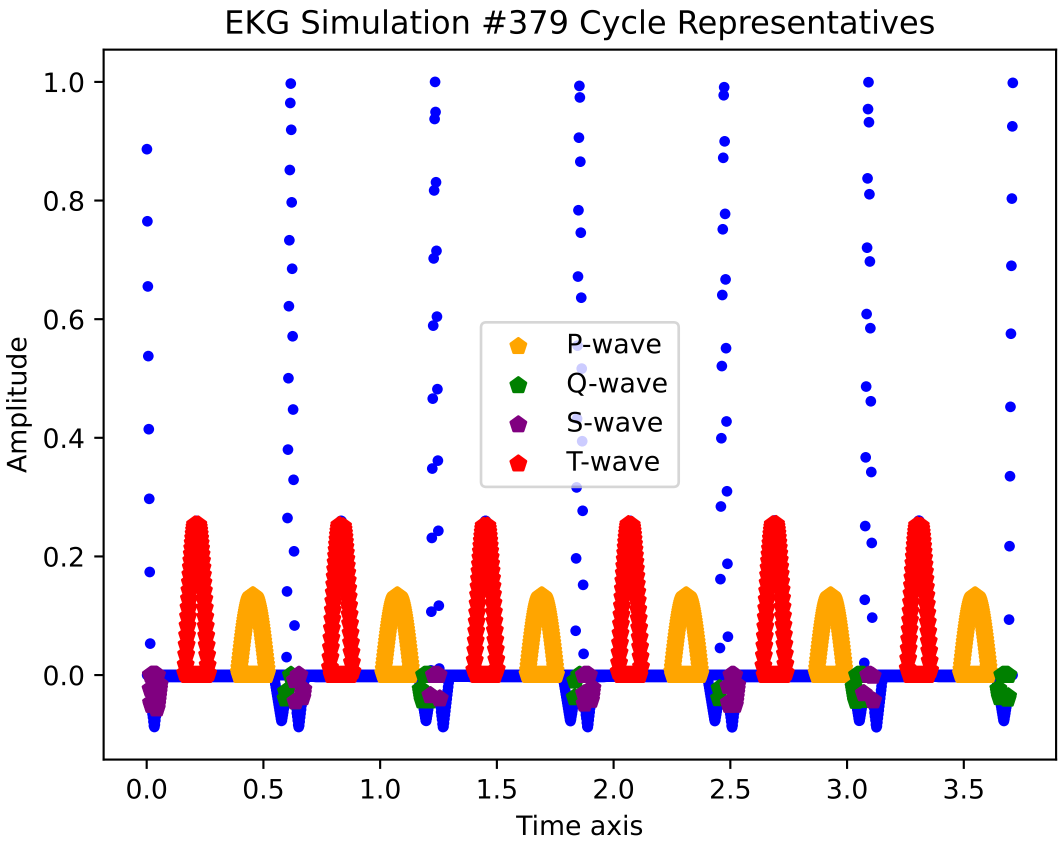

EKG Simulation #379 Cycle Representatives
Amplitude
Time axis
P-wave
Q-wave
S-wave
T-wave
1.0
0.8
0.6
0.4
0.2
0.0
0.0
0.5
1.0
1.5
2.0
2.5
3.0
3.5

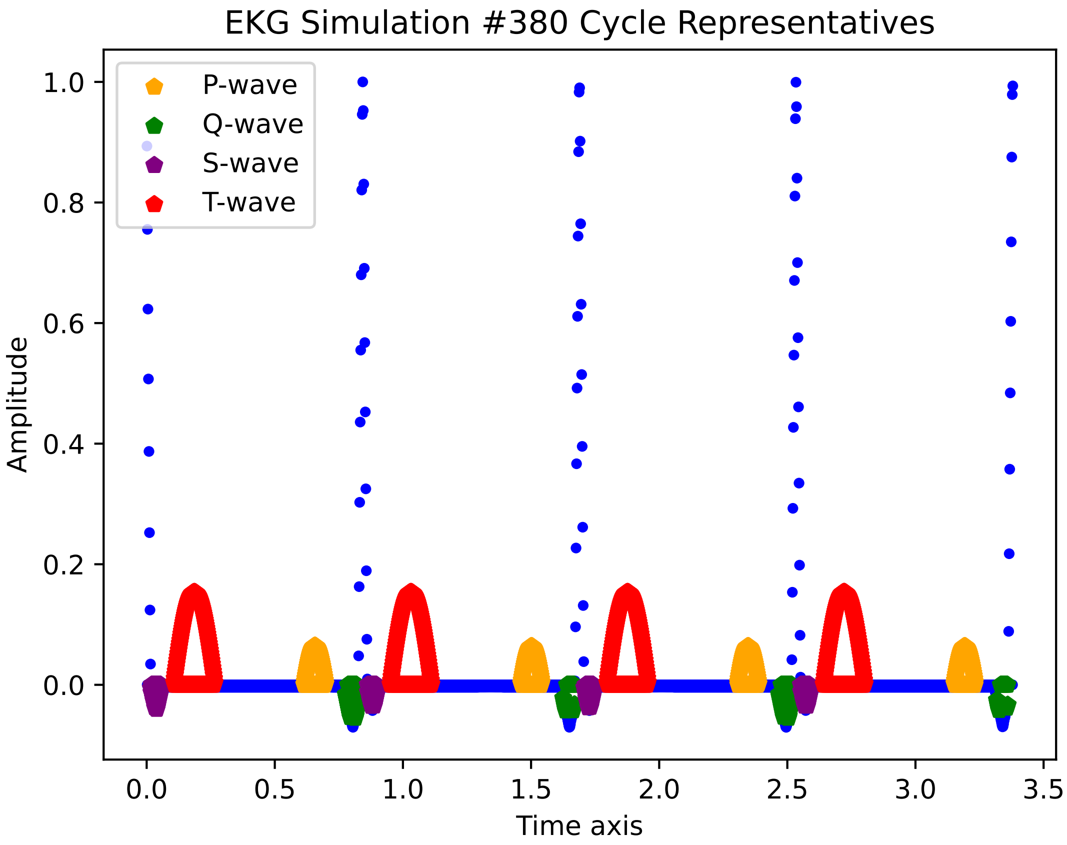

EKG Simulation #380 Cycle Representatives
P-wave
Q-wave
S-wave
T-wave
Amplitude
Time axis
1.0
0.8
0.6
0.4
0.2
0.0
0.0
0.5
1.0
1.5
2.0
2.5
3.0
3.5

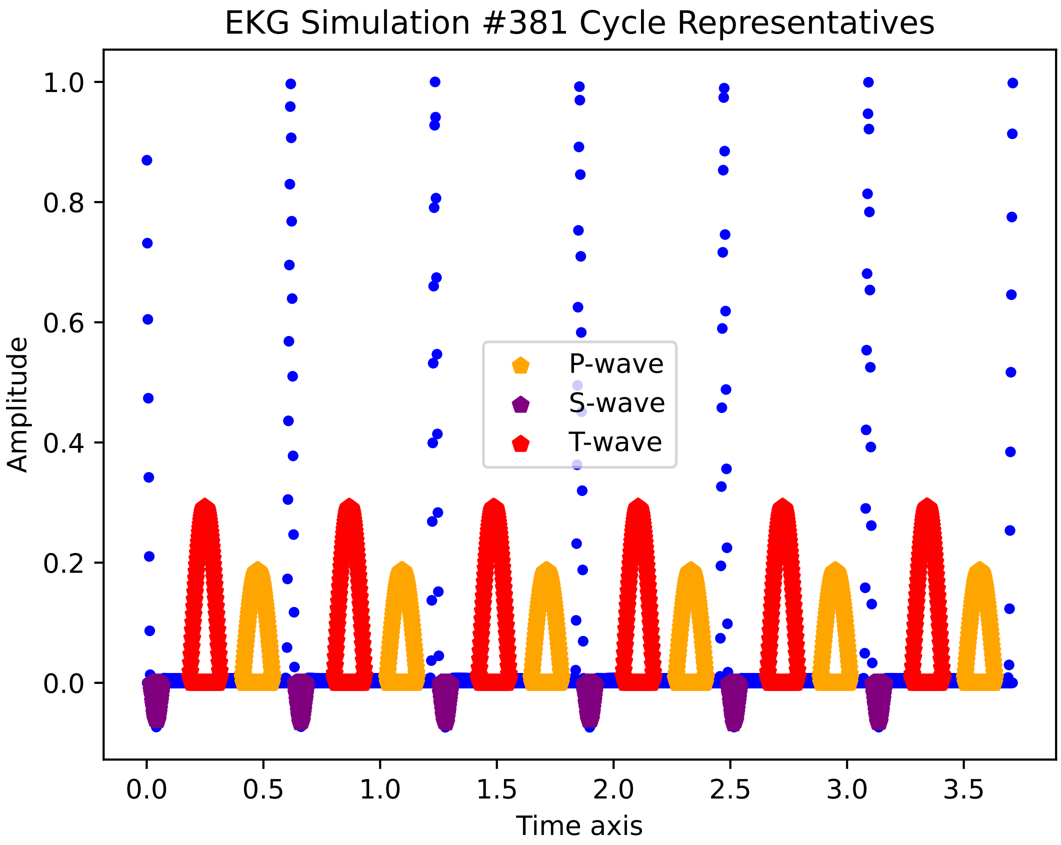

EKG Simulation #381 Cycle Representatives
Amplitude
Time axis
P-wave
S-wave
T-wave
1.0
0.8
0.6
0.4
0.2
0.0
0.0
0.5
1.0
1.5
2.0
2.5
3.0
3.5

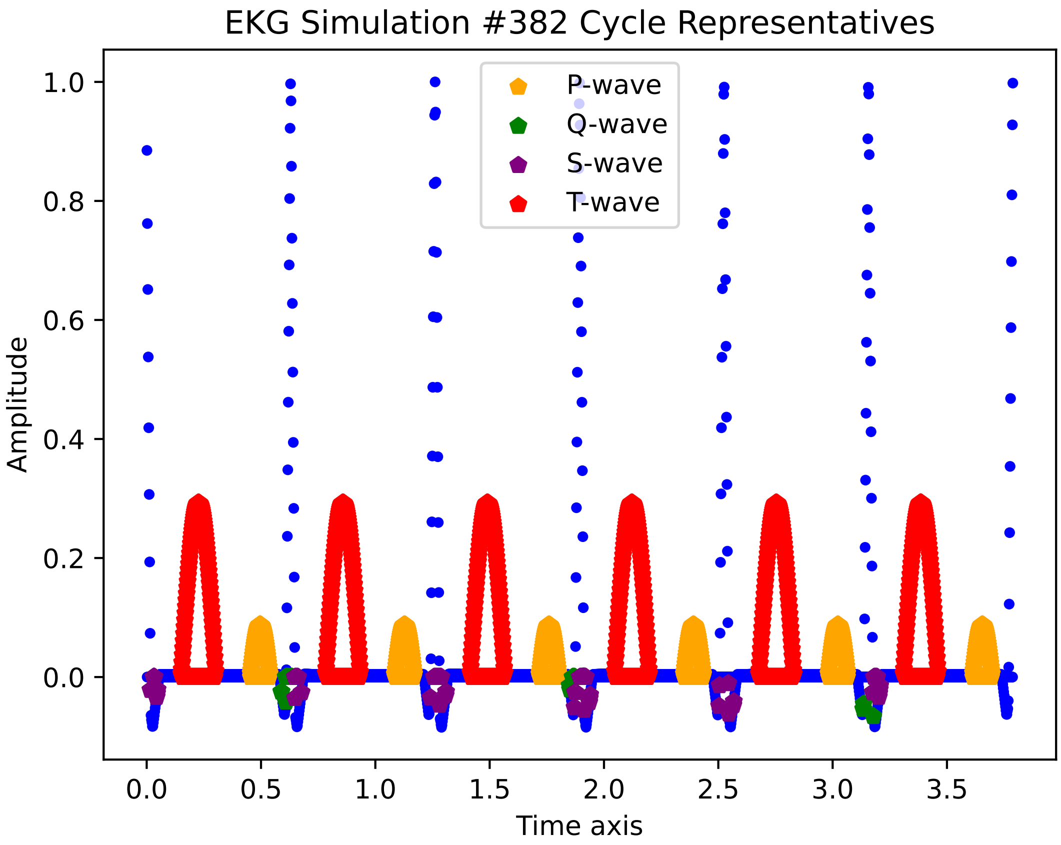

EKG Simulation #382 Cycle Representatives
P-wave
Q-wave
S-wave
T-wave
Amplitude
Time axis
1.0
0.8
0.6
0.4
0.2
0.0
0.0
0.5
1.0
1.5
2.0
2.5
3.0
3.5

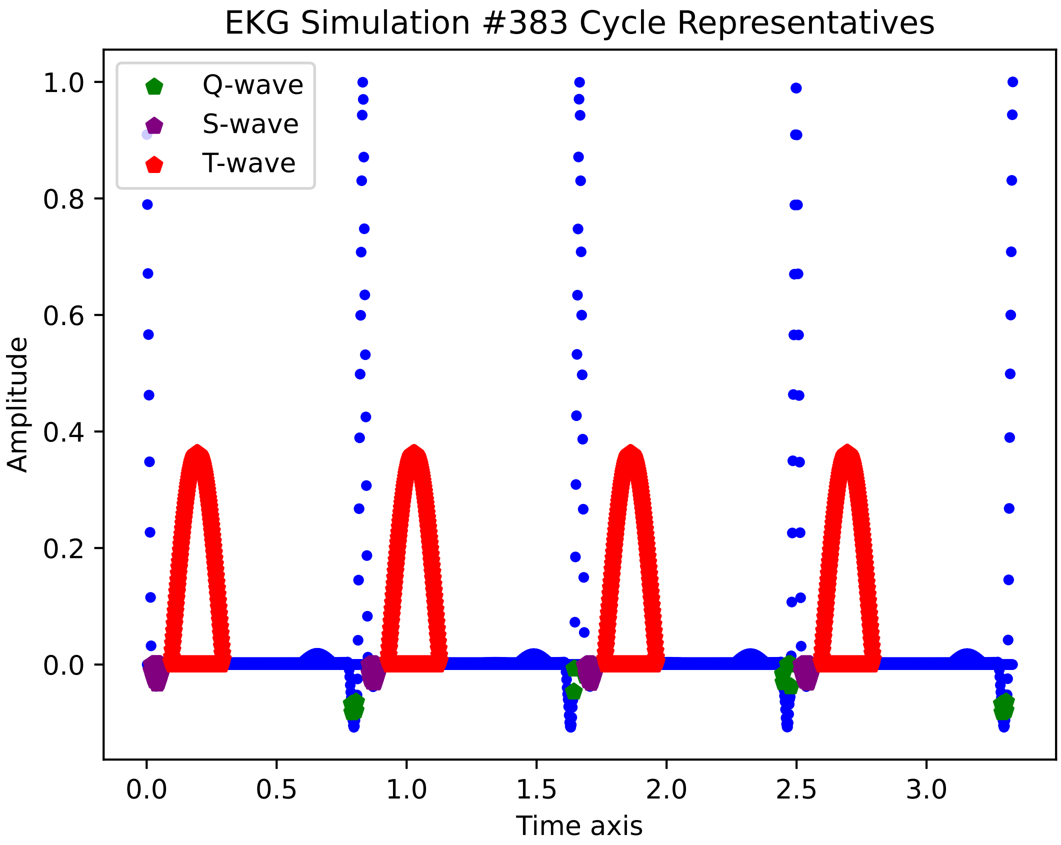

EKG Simulation #383 Cycle Representatives
Q-wave
S-wave
T-wave
Amplitude
Time axis
1.0
0.8
0.6
0.4
0.2
0.0
0.0
0.5
1.0
1.5
2.0
2.5
3.0

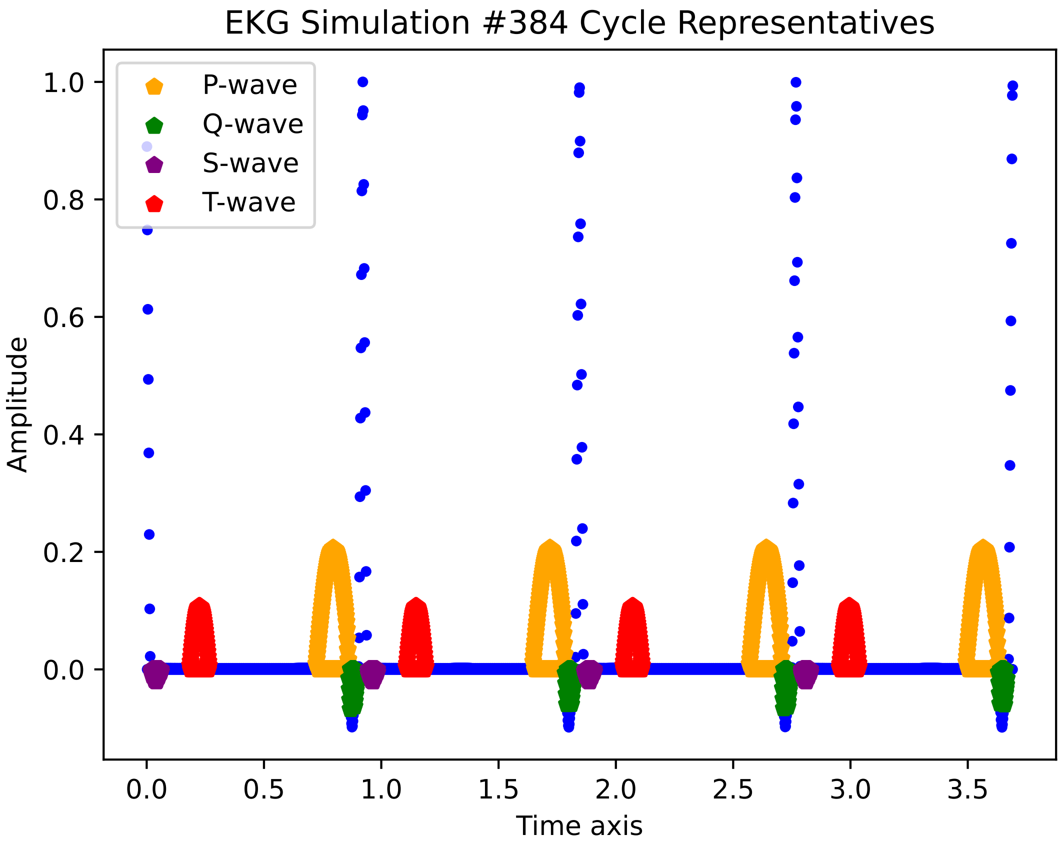
EKG Simulation #384 Cycle Representatives
P-wave
Q-wave
S-wave
T-wave
1.0
0.8
0.6
0.4
0.2
0.0
Amplitude
0.0
0.5
1.0
1.5
2.0
2.5
3.0
3.5
Time axis

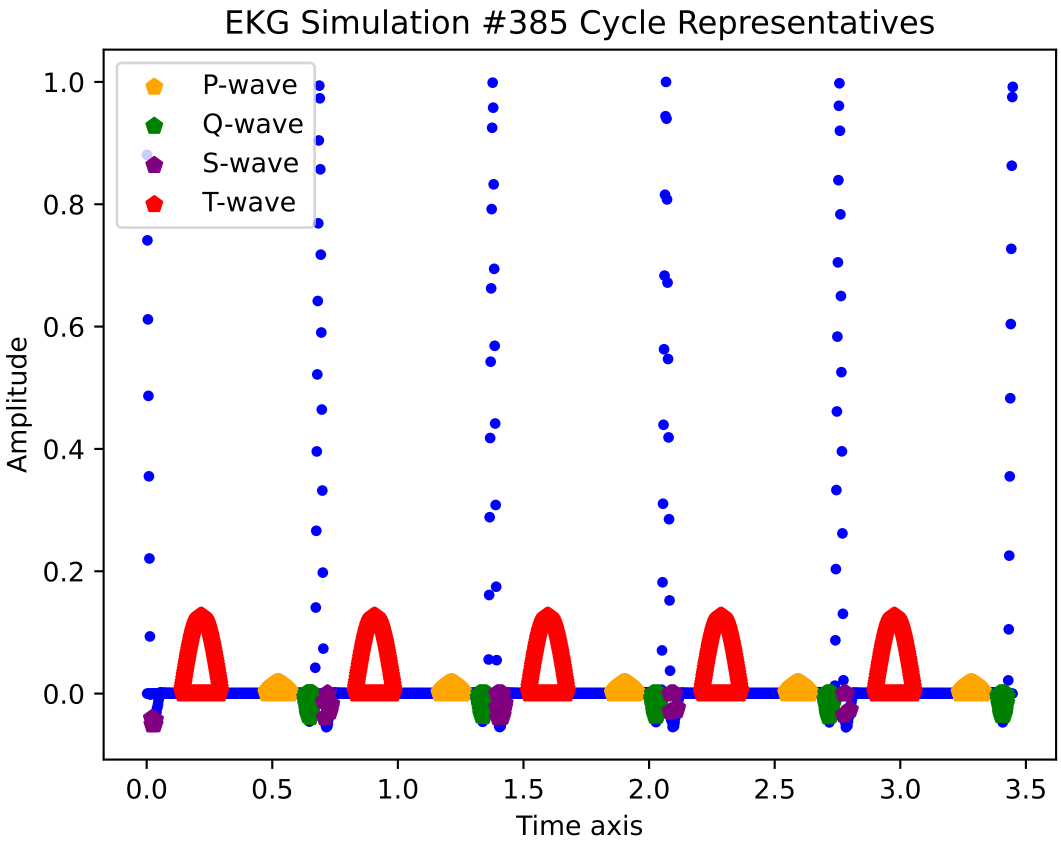
EKG Simulation #385 Cycle Representatives
P-wave
Q-wave
S-wave
T-wave
Amplitude
Time axis
1.0
0.8
0.6
0.4
0.2
0.0
0.0
0.5
1.0
1.5
2.0
2.5
3.0
3.5

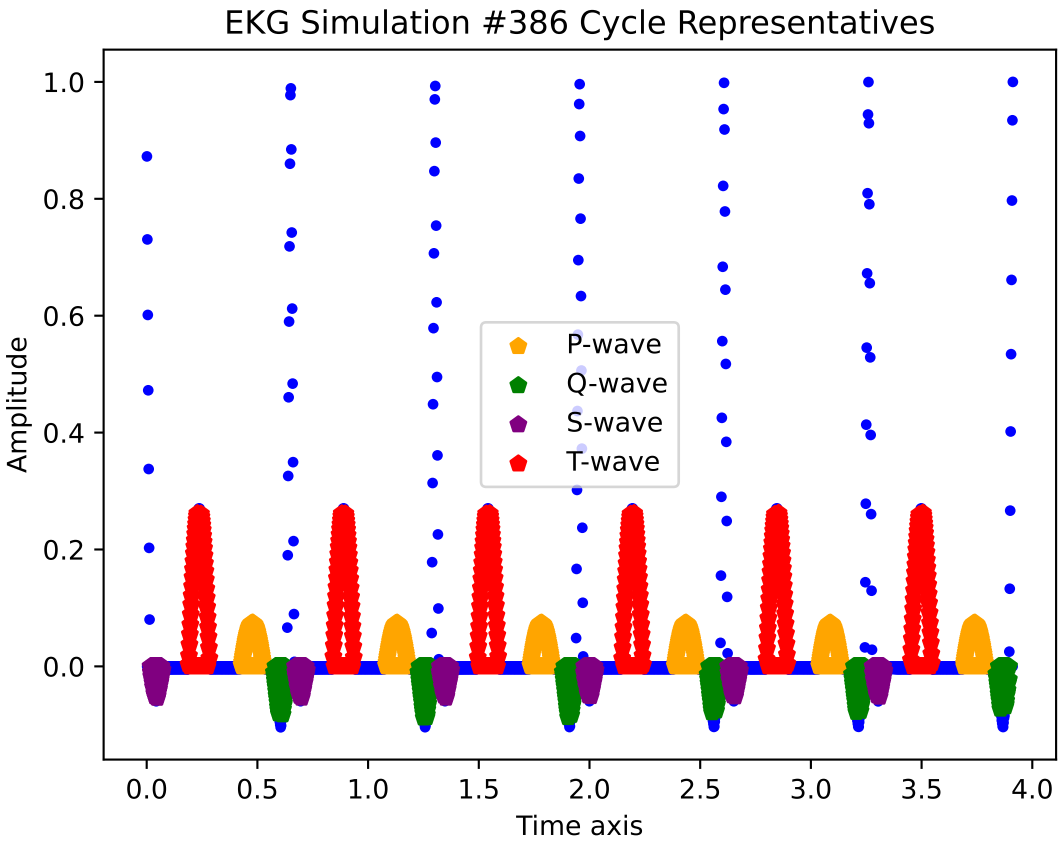

EKG Simulation #386 Cycle Representatives
Amplitude
Time axis
P-wave
Q-wave
S-wave
T-wave
0.0
0.2
0.4
0.6
0.8
1.0
0.0
0.5
1.0
1.5
2.0
2.5
3.0
3.5
4.0

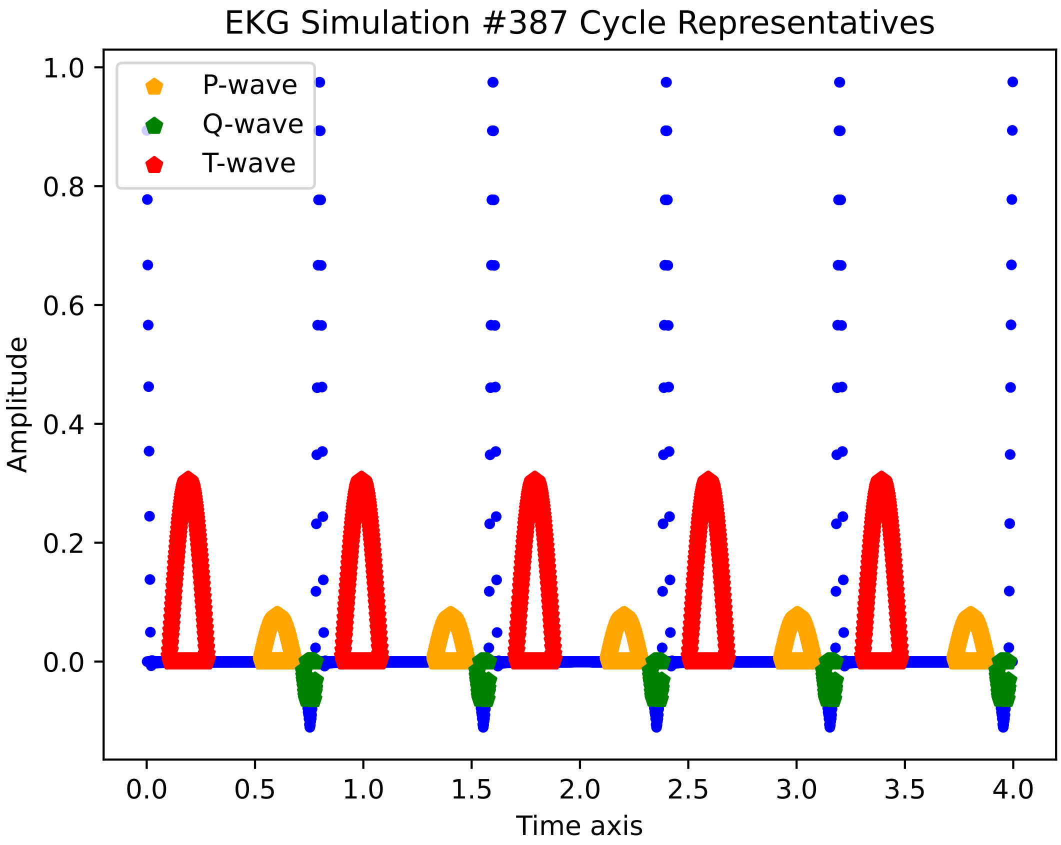
EKG Simulation #387 Cycle Representatives
P-wave
Q-wave
T-wave
Amplitude
Time axis
1.0
0.8
0.6
0.4
0.2
0.0
0.0
0.5
1.0
1.5
2.0
2.5
3.0
3.5
4.0

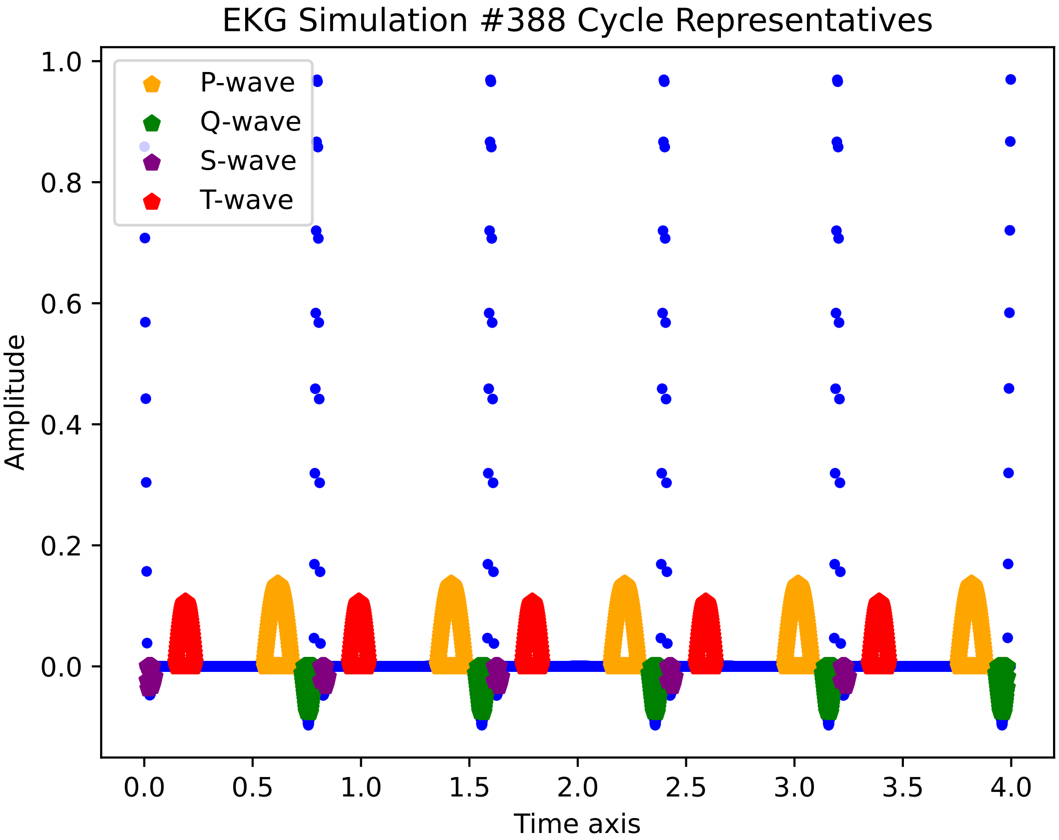
EKG Simulation #388 Cycle Representatives
P-wave
Q-wave
S-wave
T-wave
Amplitude
Time axis
1.0
0.8
0.6
0.4
0.2
0.0
0.0
0.5
1.0
1.5
2.0
2.5
3.0
3.5
4.0

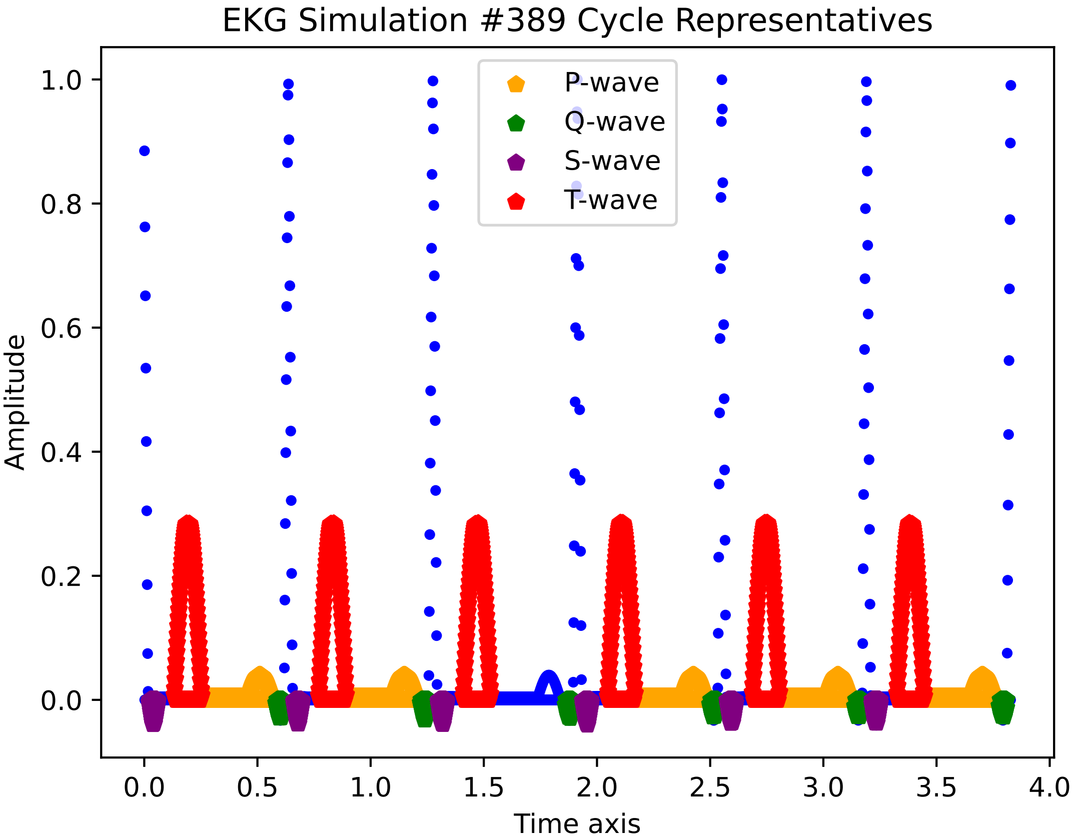
EKG Simulation #389 Cycle Representatives
P-wave
Q-wave
S-wave
T-wave
Amplitude
0.0
0.2
0.4
0.6
0.8
1.0
0.5
1.5
2.0
2.5
3.0
3.5
4.0
Time axis

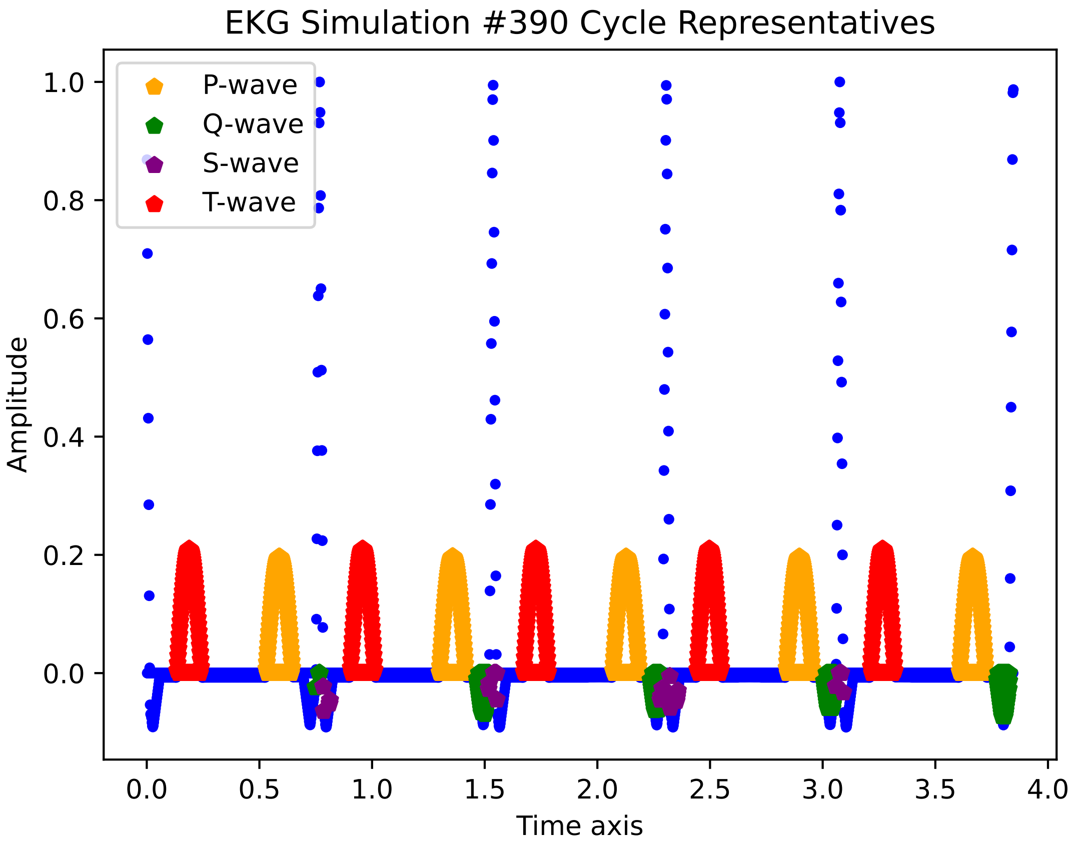

EKG Simulation #390 Cycle Representatives
P-wave
Q-wave
S-wave
T-wave
Amplitude
Time axis
1.0
0.8
0.6
0.4
0.2
0.0
0.0
0.5
1.0
1.5
2.0
2.5
3.0
3.5
4.0

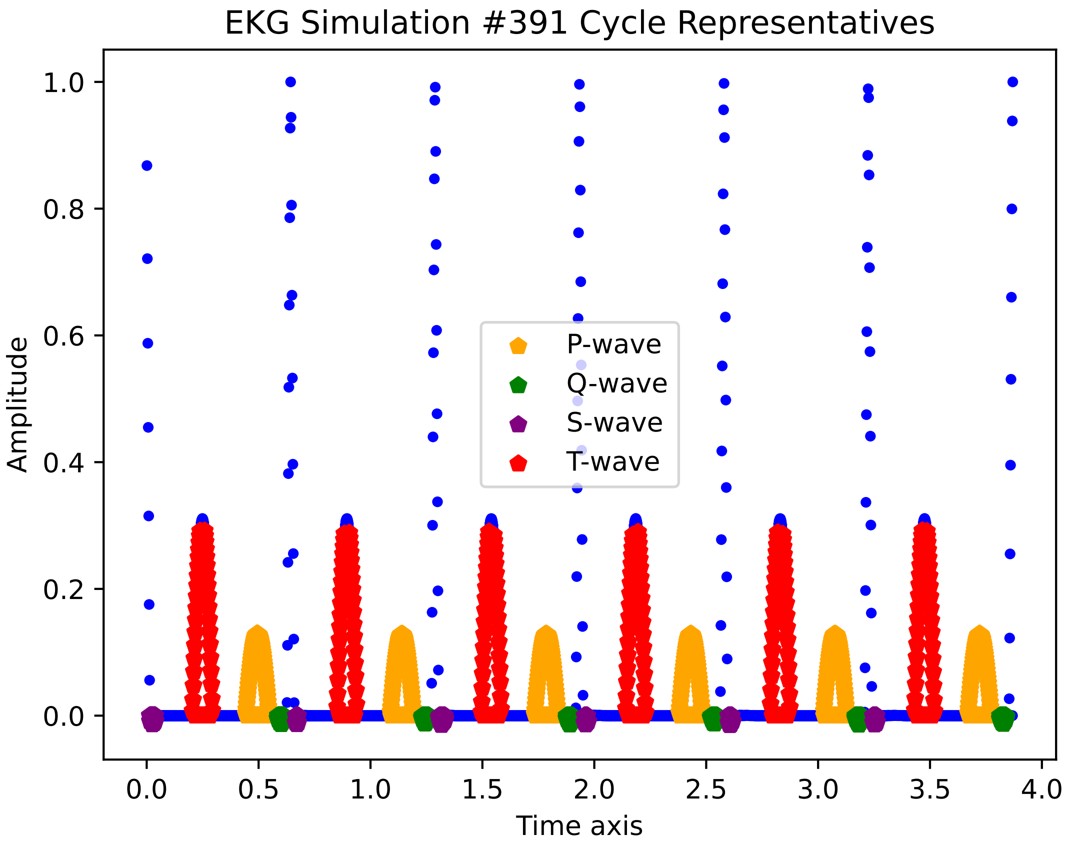
EKG Simulation #391 Cycle Representatives
Amplitude
1.0
0.8
0.6
0.4
0.2
0.0
0.0
0.5
1.0
1.5
2.0
2.5
3.0
3.5
4.0
Time axis
P-wave
Q-wave
S-wave
T-wave

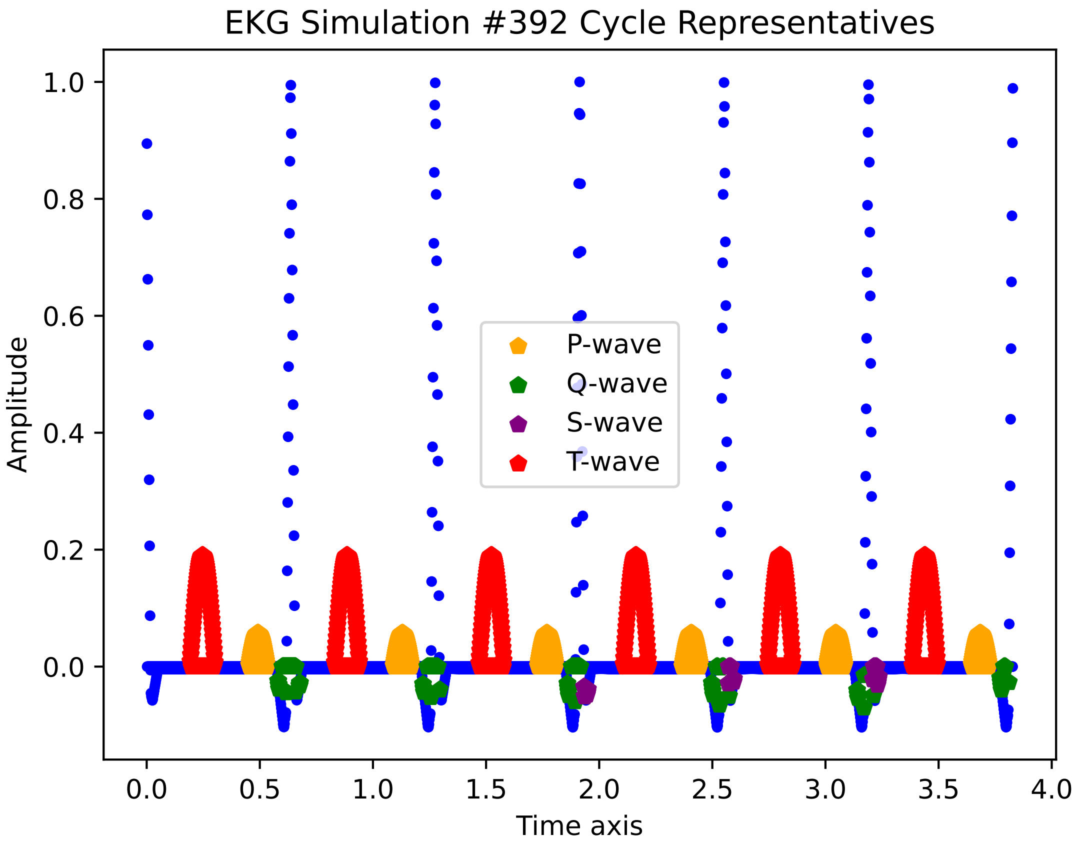

EKG Simulation #392 Cycle Representatives
Amplitude
Time axis
P-wave
Q-wave
S-wave
T-wave
1.0
0.8
0.6
0.4
0.2
0.0
0.0
0.5
1.0
1.5
2.0
2.5
3.0
3.5
4.0

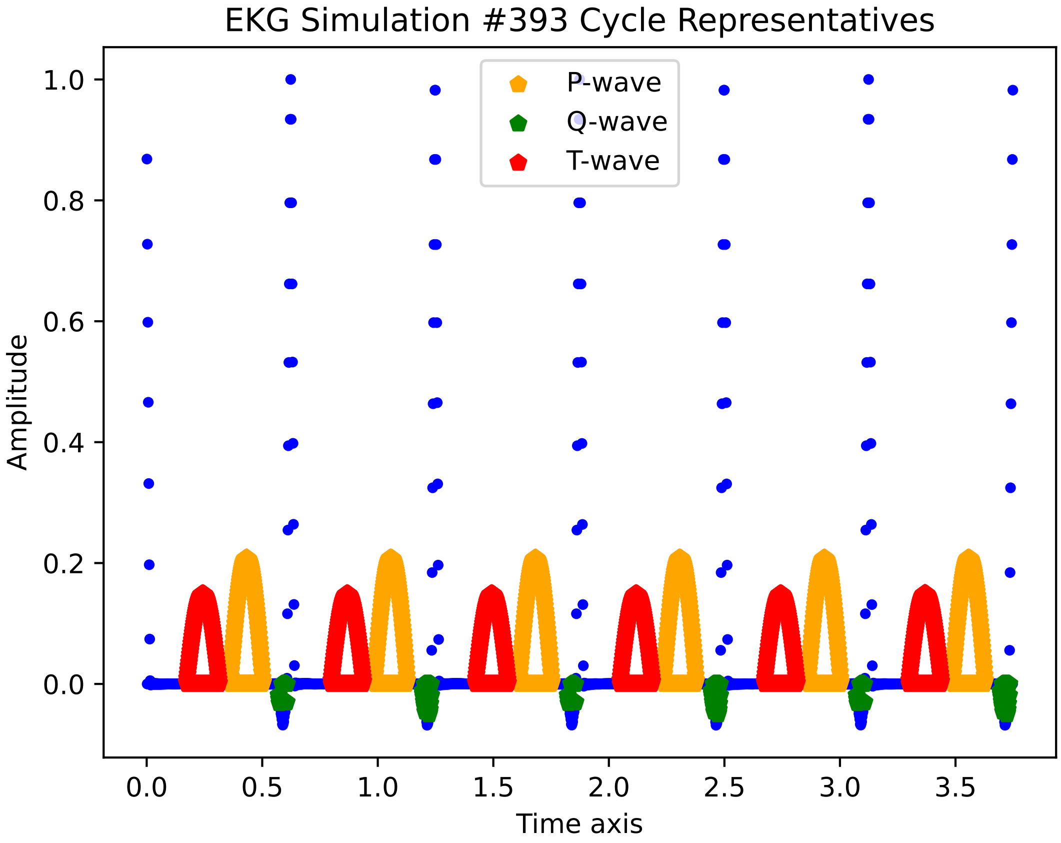
EKG Simulation #393 Cycle Representatives
P-wave
Q-wave
T-wave
Amplitude
Time axis
1.0
0.8
0.6
0.4
0.2
0.0
0.0
0.5
1.0
1.5
2.0
2.5
3.0
3.5

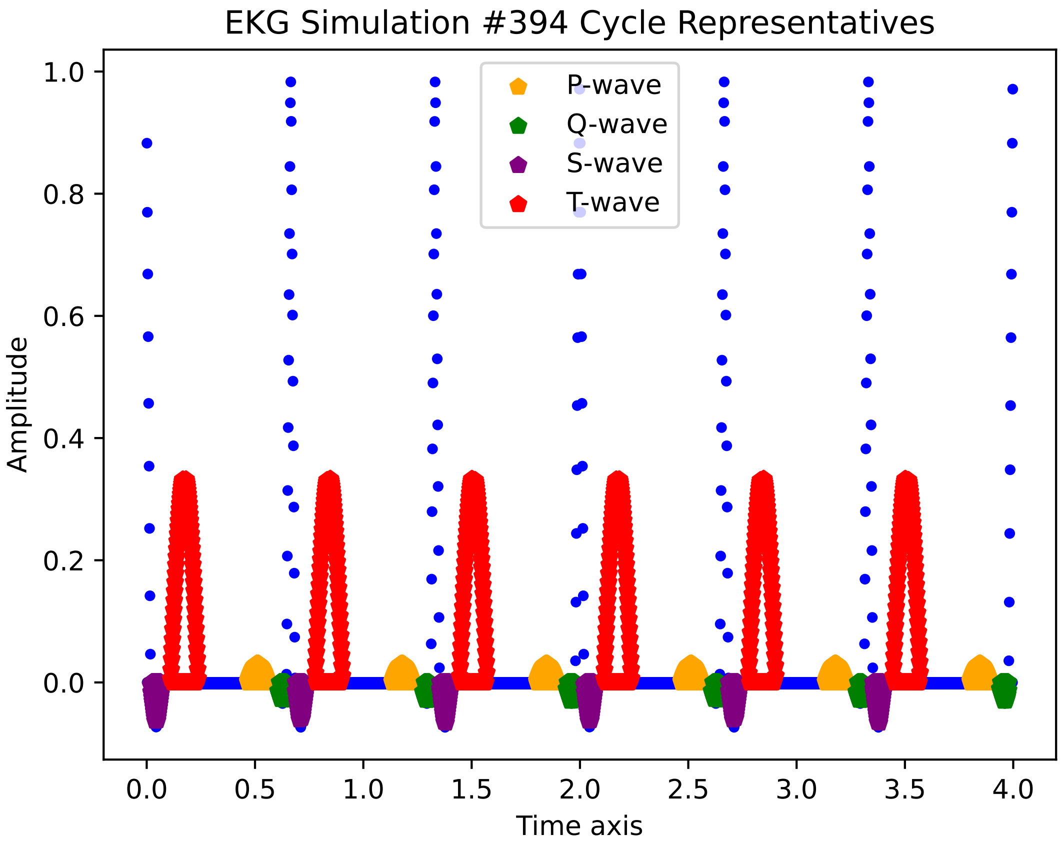
EKG Simulation #394 Cycle Representatives
P-wave
Q-wave
S-wave
T-wave
Amplitude
Time axis
0.0
0.2
0.4
0.6
0.8
1.0
0.0
0.5
1.0
1.5
2.0
2.5
3.0
3.5
4.0

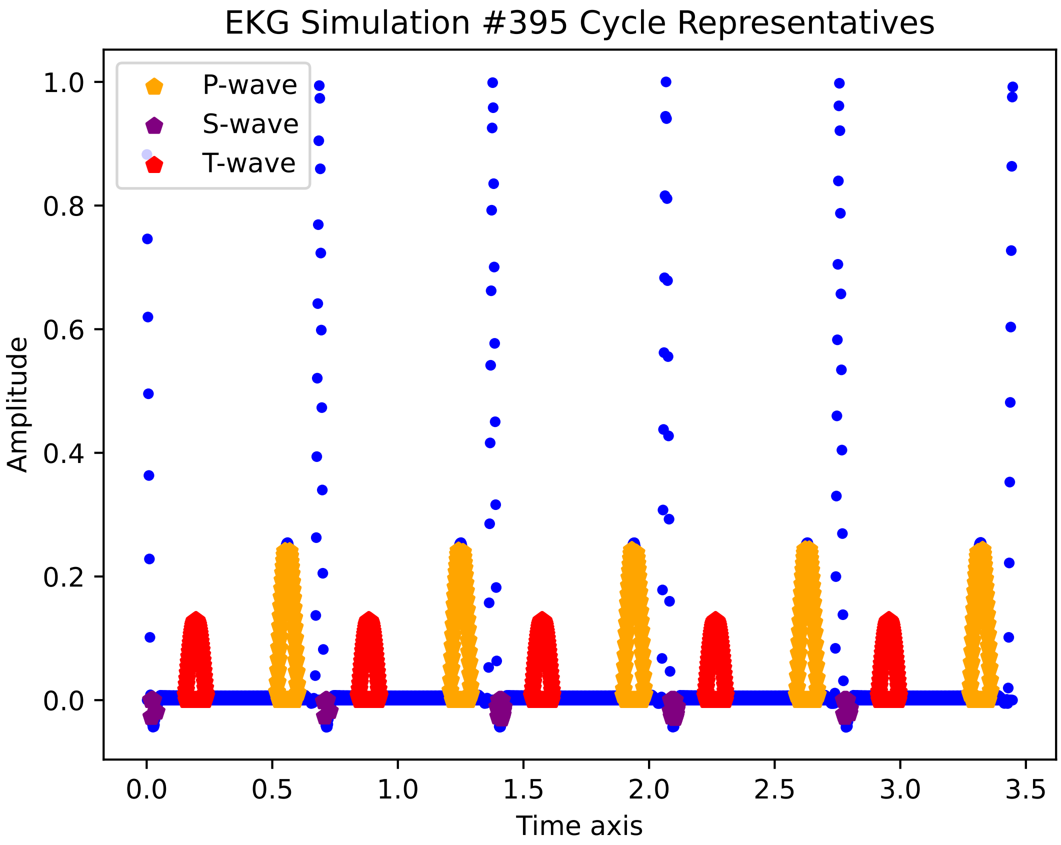
EKG Simulation #395 Cycle Representatives
P-wave
S-wave
T-wave
Amplitude
Time axis
1.0
0.8
0.6
0.4
0.2
0.0
0.0
0.5
1.0
1.5
2.0
2.5
3.0
3.5

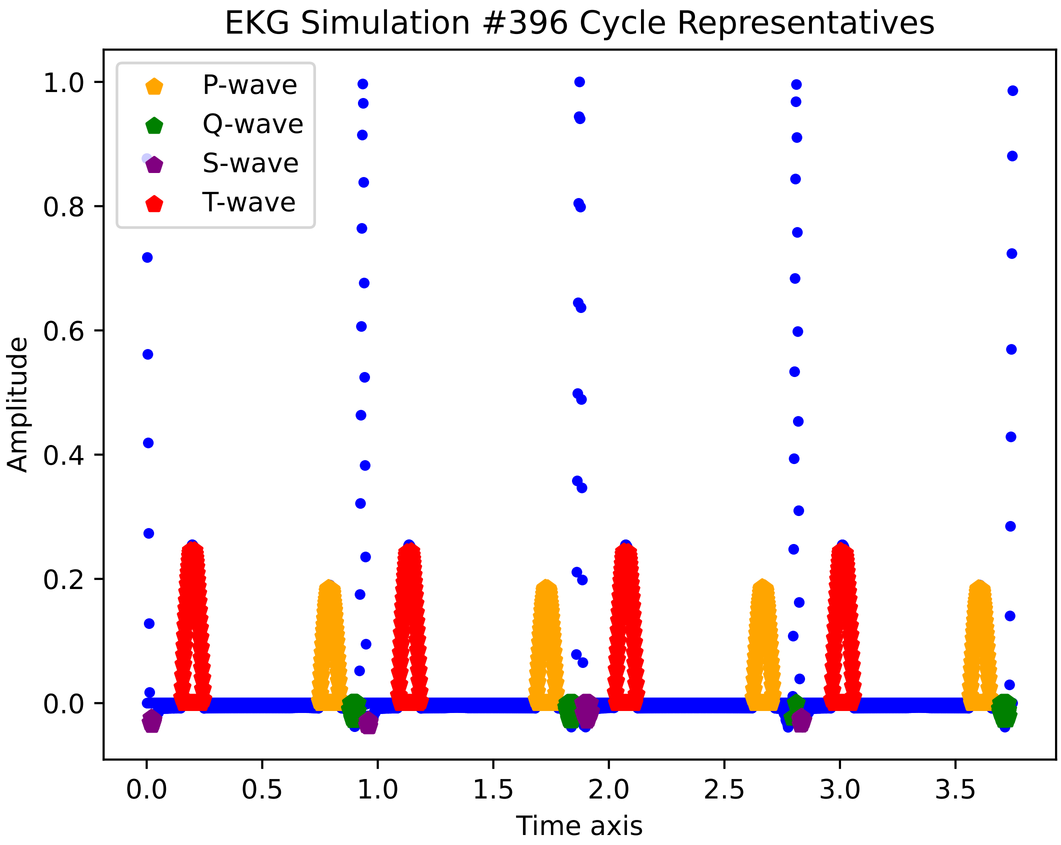

EKG Simulation #396 Cycle Representatives
P-wave
Q-wave
S-wave
T-wave
Amplitude
Time axis
1.0
0.8
0.6
0.4
0.2
0.0
0.0
0.5
1.0
1.5
2.0
2.5
3.0
3.5

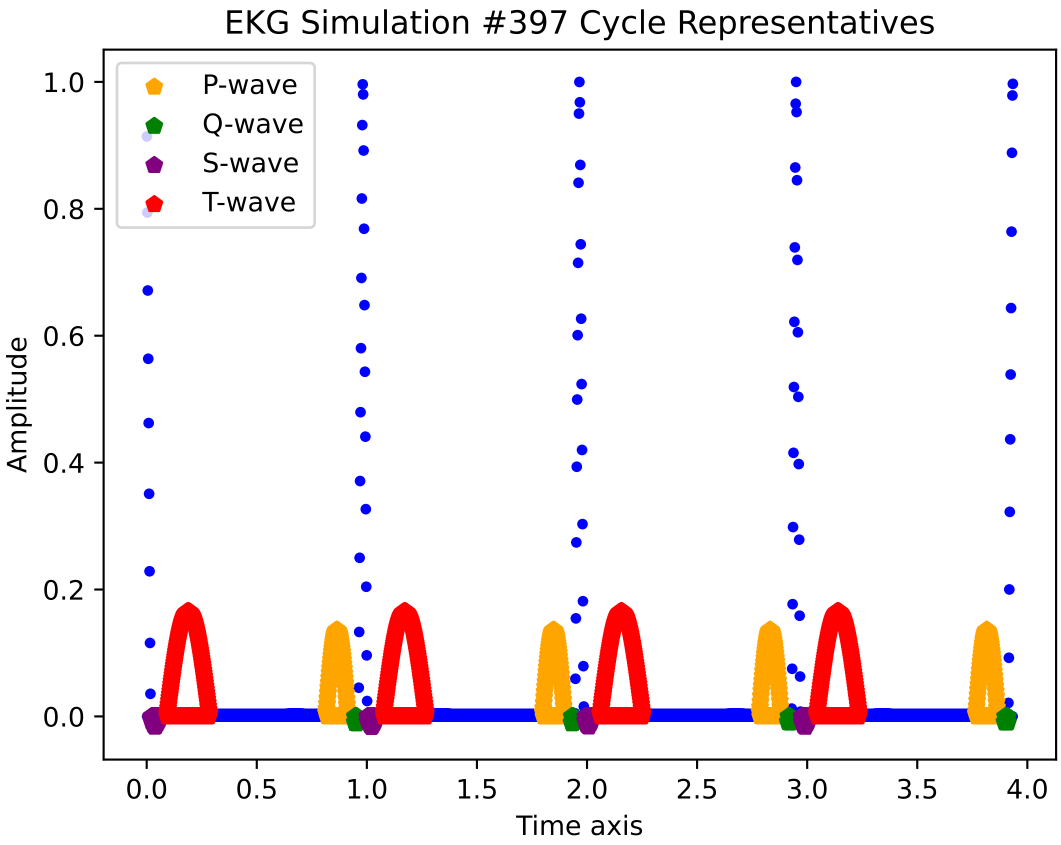

EKG Simulation #397 Cycle Representatives
P-wave
Q-wave
S-wave
T-wave
Amplitude
1.0
0.8
0.6
0.4
0.2
0.0
0.0
0.5
1.0
1.5
2.0
2.5
3.0
3.5
4.0
Time axis

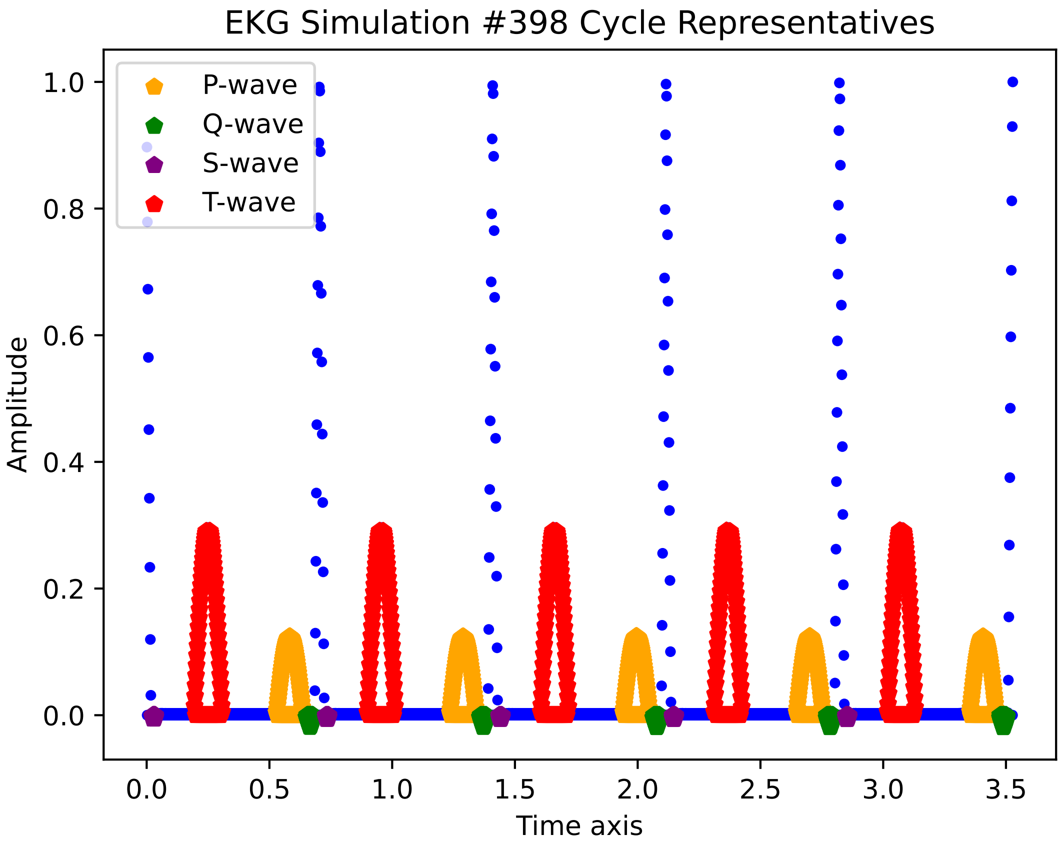
EKG Simulation #398 Cycle Representatives
P-wave
Q-wave
S-wave
T-wave
Amplitude
1.0
0.8
0.6
0.4
0.2
0.0
0.0
0.5
1.0
1.5
2.0
2.5
3.0
3.5
Time axis

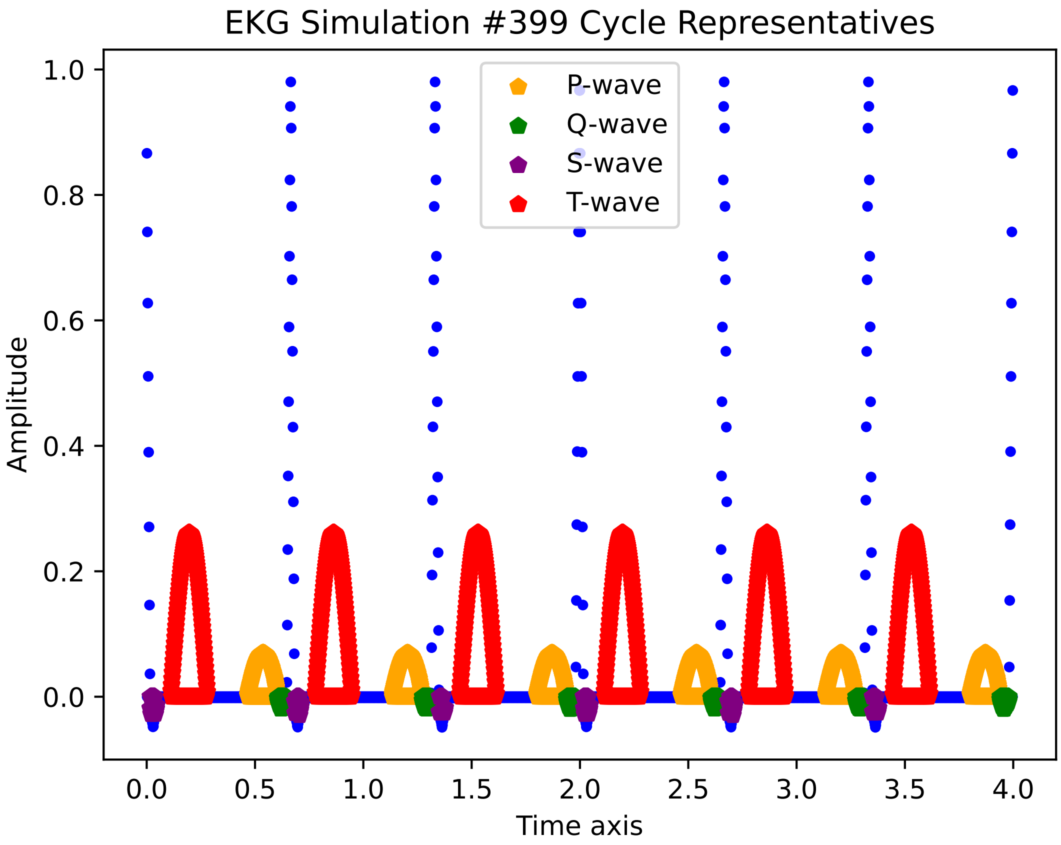
EKG Simulation #399 Cycle Representatives
P-wave
Q-wave
S-wave
T-wave
Amplitude
Time axis
0.0
0.2
0.4
0.6
0.8
1.0
0.0
0.5
1.0
1.5
2.0
2.5
3.0
3.5
4.0

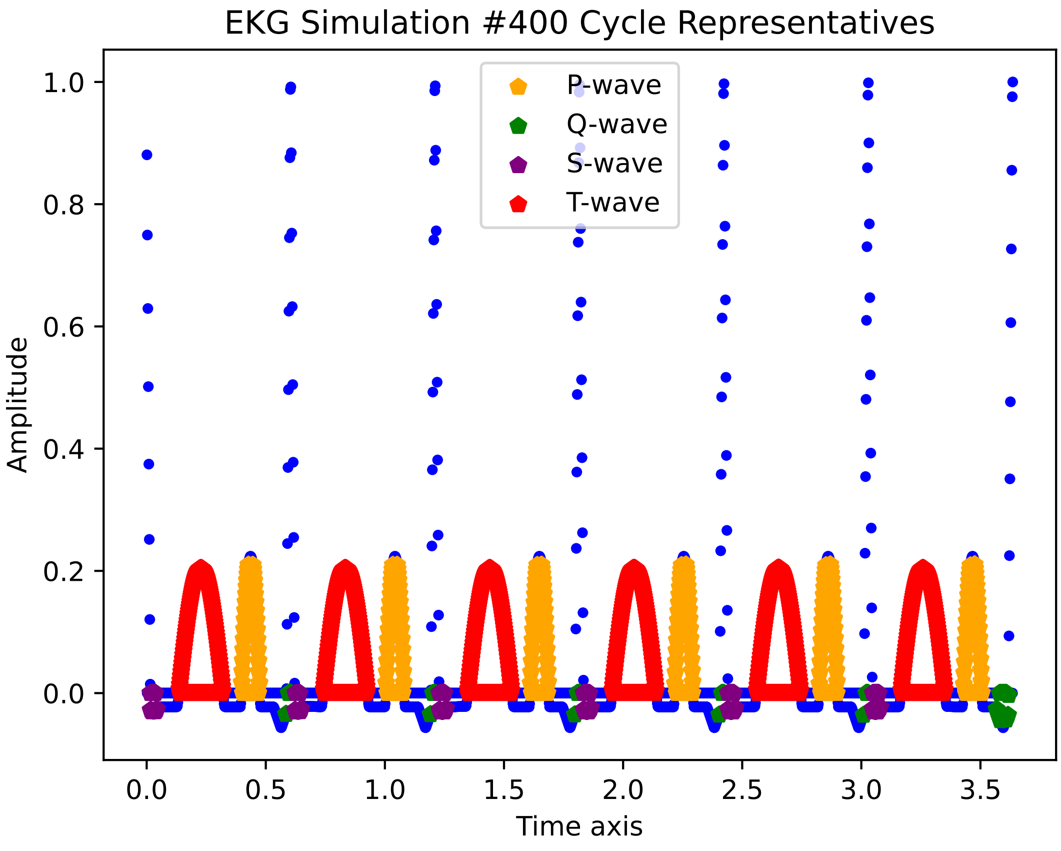

EKG Simulation #400 Cycle Representatives
P-wave
Q-wave
S-wave
T-wave
Amplitude
Time axis
1.0
0.8
0.6
0.4
0.2
0.0
0.0
0.5
1.0
1.5
2.0
2.5
3.0
3.5
