## Supplementary File 4 for "Electrocardiogram feature extraction and interval measurements using optimal representative cycles from persistent homology"

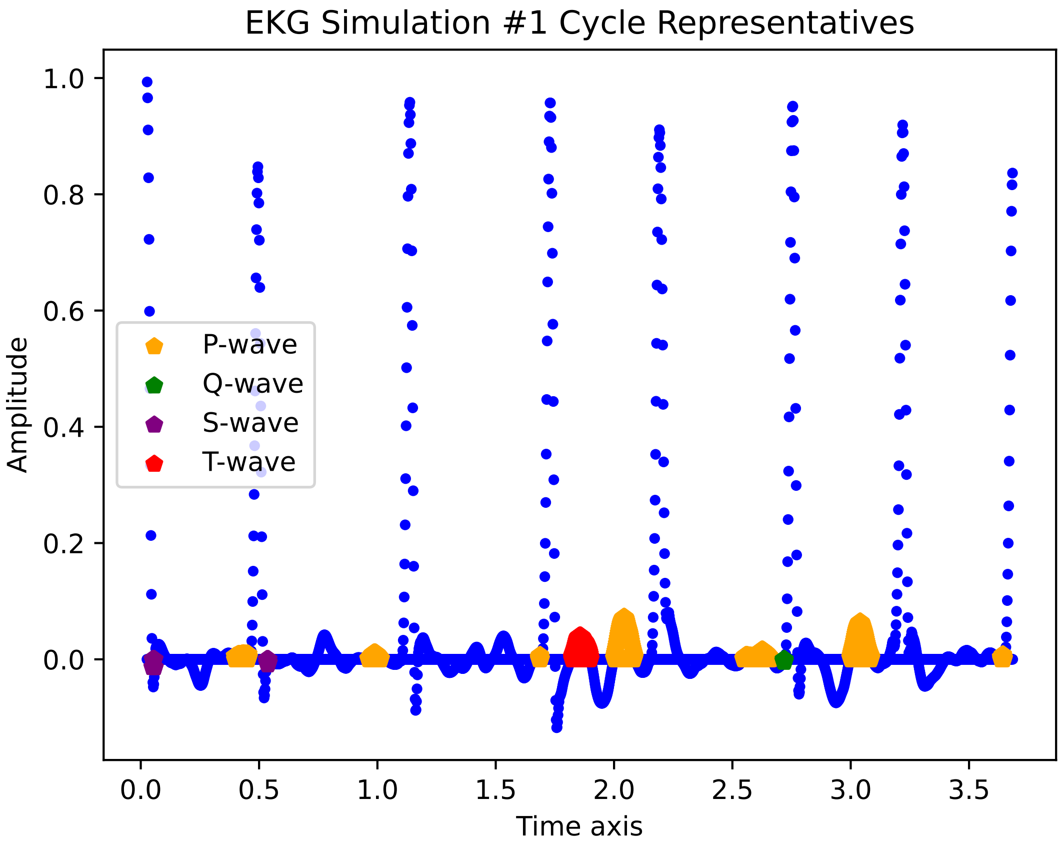

EKG Simulation #1 Cycle Representatives
Amplitude
1.0
0.8
0.6
0.4
0.2
0.0
0.0
0.5
1.0
1.5
2.0
2.5
3.0
3.5
Time axis
P-wave
Q-wave
S-wave
T-wave

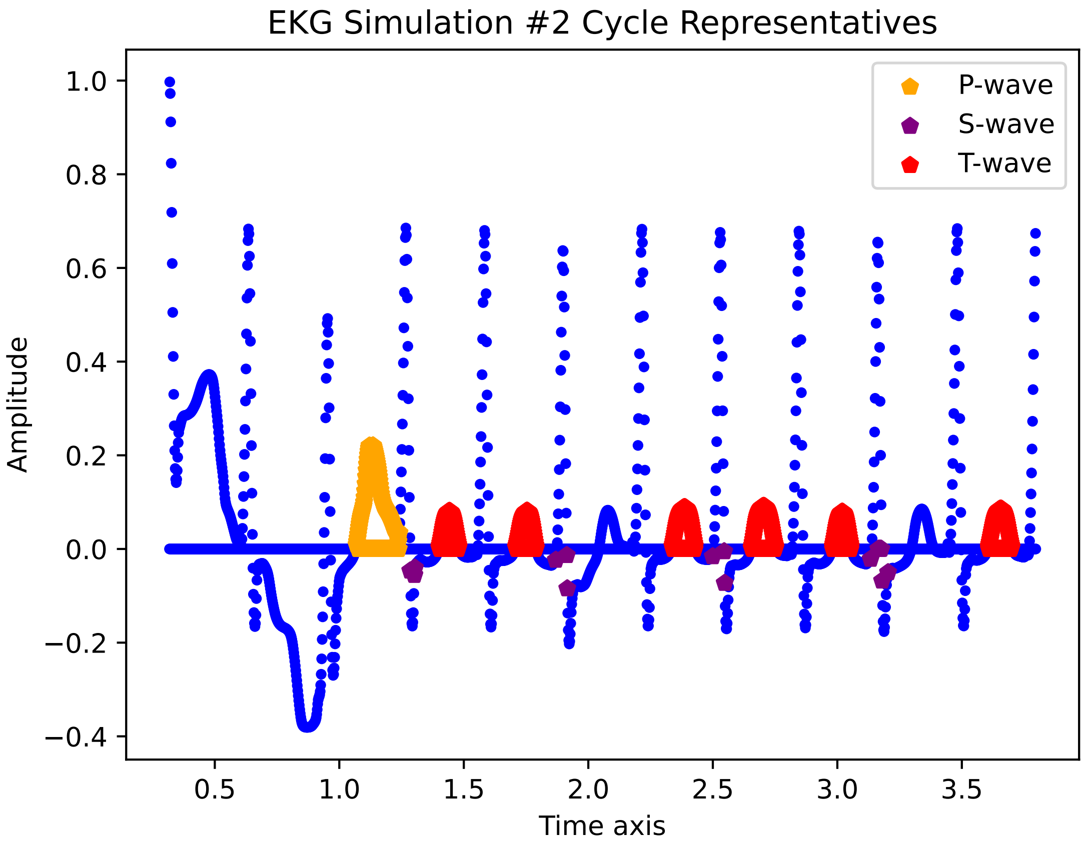
EKG Simulation #2 Cycle Representatives
P-wave
S-wave
T-wave
Amplitude
Time axis
1.0
0.8
0.6
0.4
0.2
0.0
−0.2
−0.4
0.5
1.0
1.5
2.0
2.5
3.0
3.5

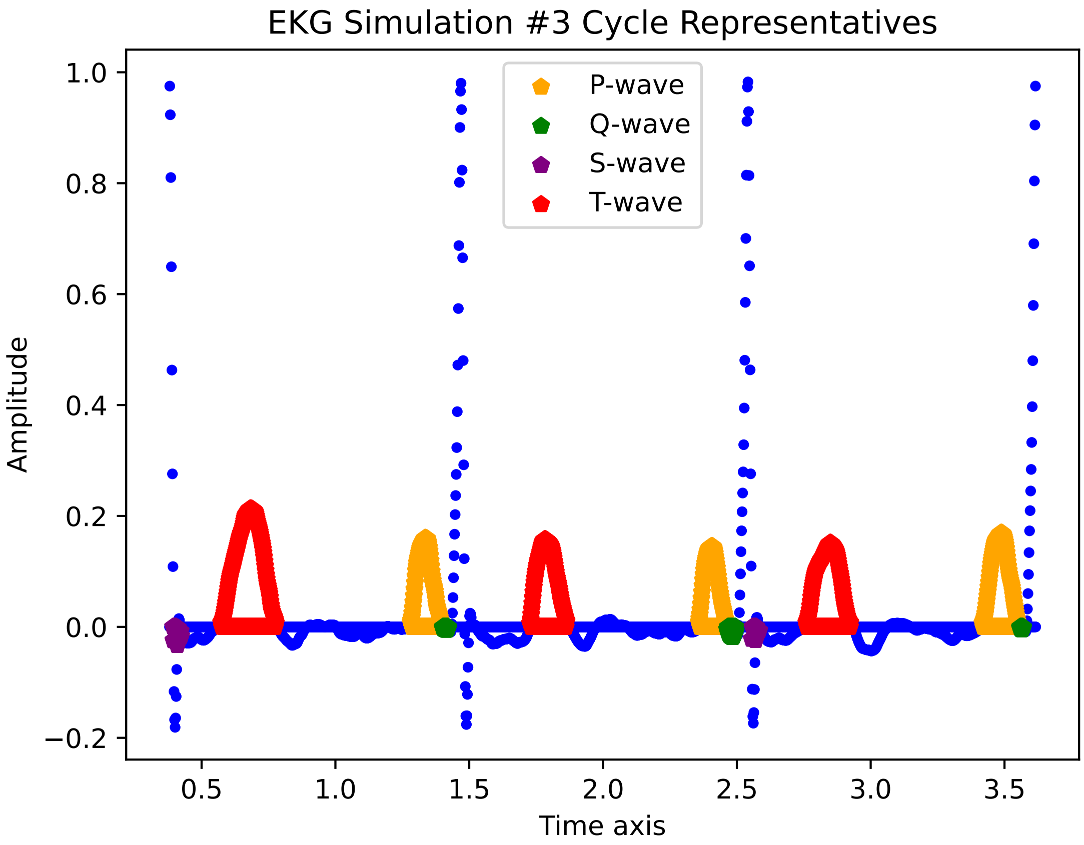
EKG Simulation #3 Cycle Representatives
P-wave
Q-wave
S-wave
T-wave
Amplitude
Time axis
1.0
0.8
0.6
0.4
0.2
0.0
−0.2
0.5
1.0
1.5
2.0
2.5
3.0
3.5

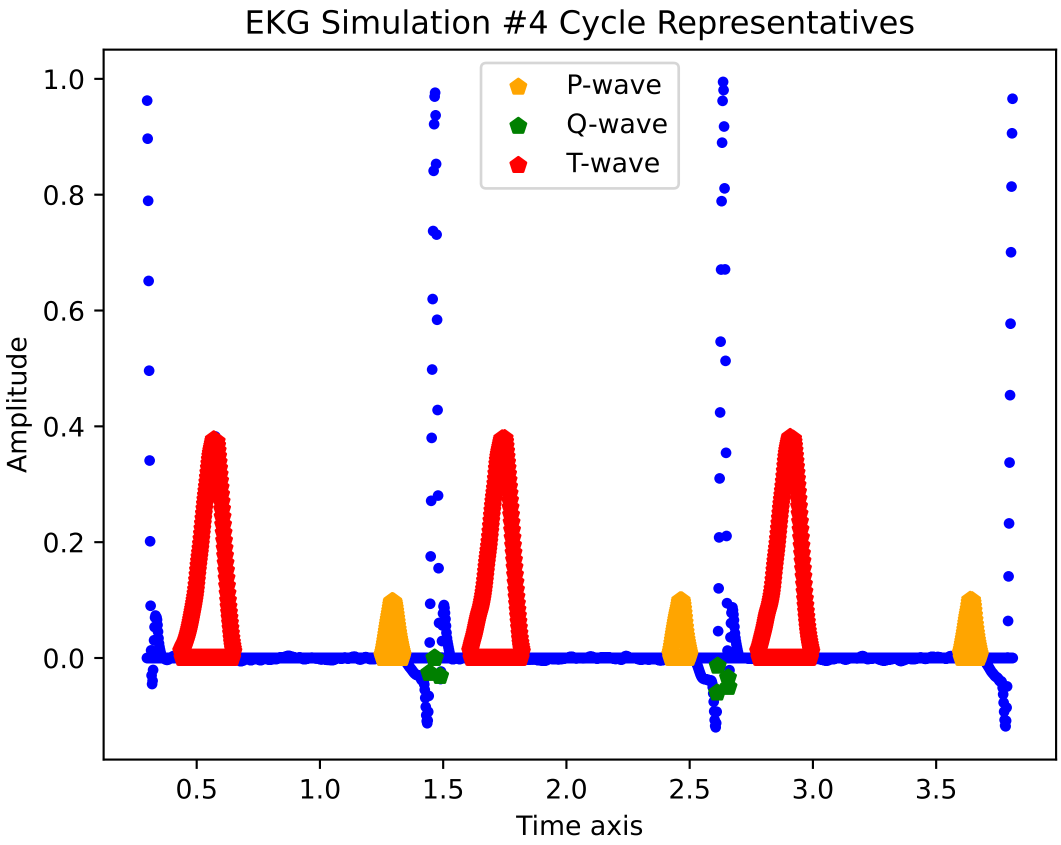
EKG Simulation #4 Cycle Representatives
P-wave
Q-wave
T-wave
Amplitude
Time axis
1.0
0.8
0.6
0.4
0.2
0.0
0.5
1.0
1.5
2.0
2.5
3.0
3.5

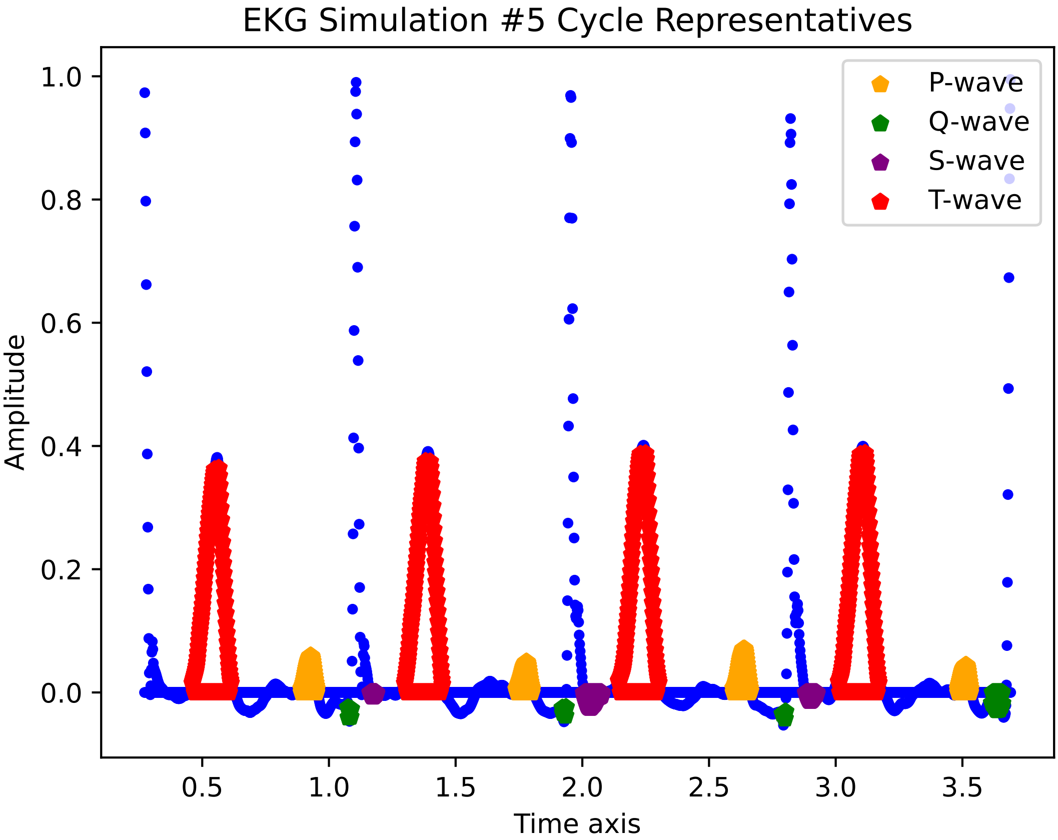
EKG Simulation #5 Cycle Representatives
P-wave
Q-wave
S-wave
T-wave
Amplitude
1.0
0.8
0.6
0.4
0.2
0.0
0.5
1.0
1.5
2.0
2.5
3.0
3.5
Time axis

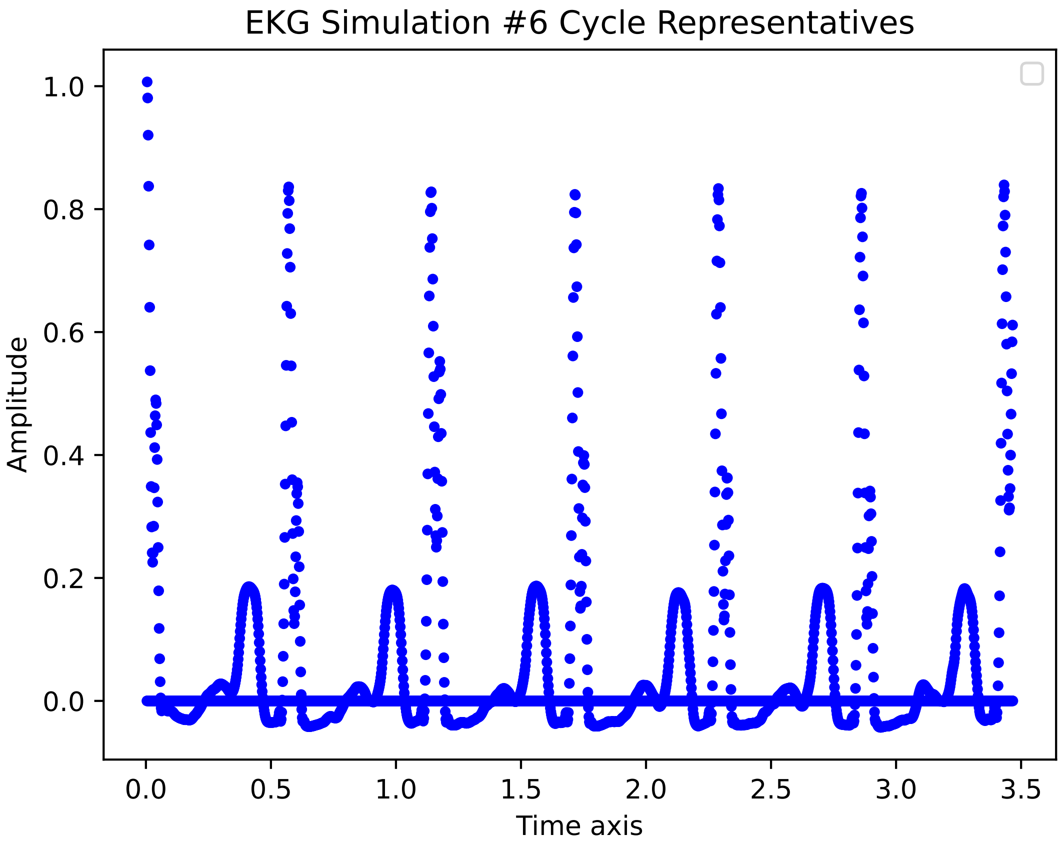

EKG Simulation #6 Cycle Representatives
Amplitude
Time axis
1.0
0.8
0.6
0.4
0.2
0.0
0.0
0.5
1.0
1.5
2.0
2.5
3.0
3.5

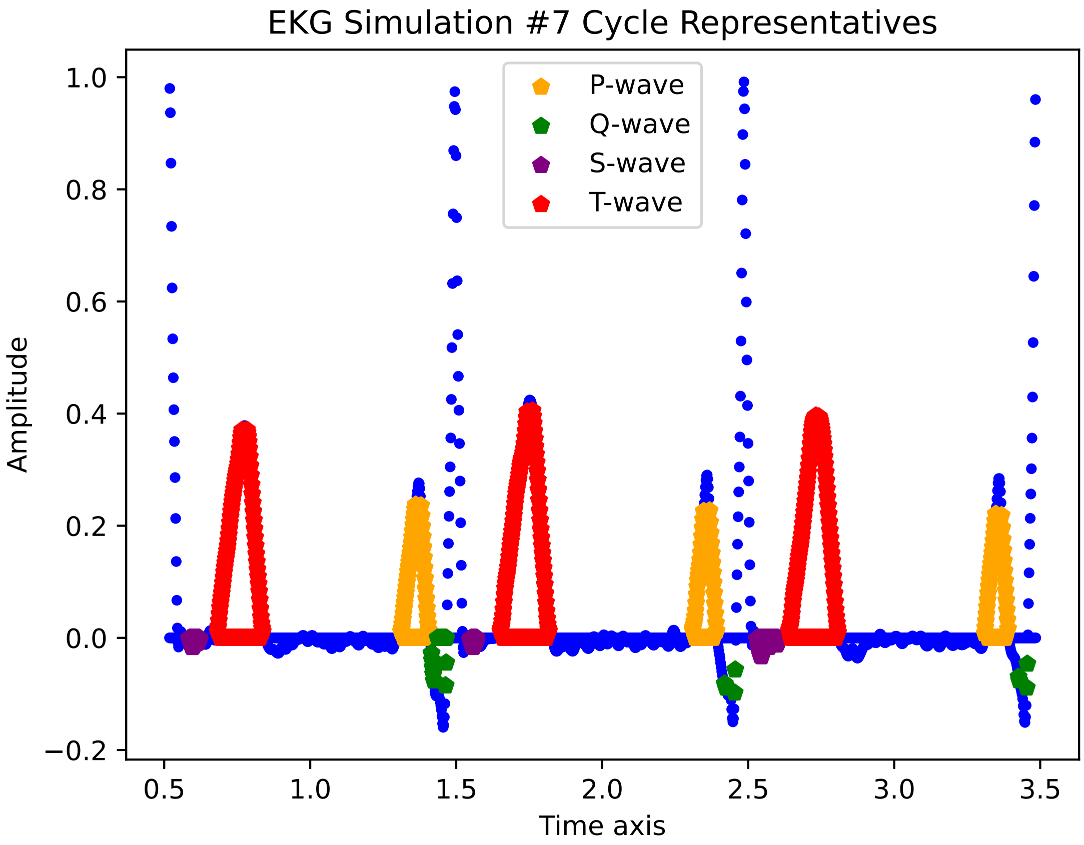
EKG Simulation #7 Cycle Representatives
P-wave
Q-wave
S-wave
T-wave
1.0
0.8
0.6
0.4
0.2
0.0
−0.2
Amplitude
0.5
1.0
1.5
2.0
2.5
3.0
3.5
Time axis

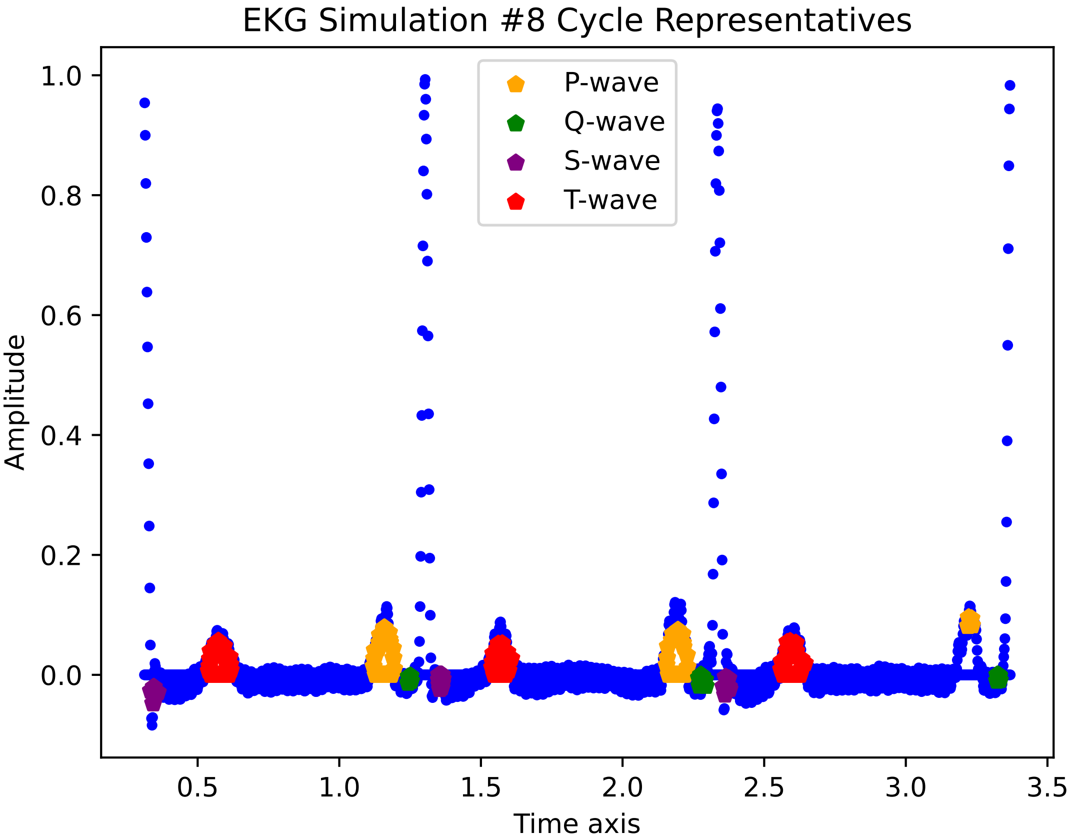

EKG Simulation #8 Cycle Representatives
P-wave
Q-wave
S-wave
T-wave
Amplitude
Time axis
1.0
0.8
0.6
0.4
0.2
0.0
0.5
1.0
1.5
2.0
2.5
3.0
3.5

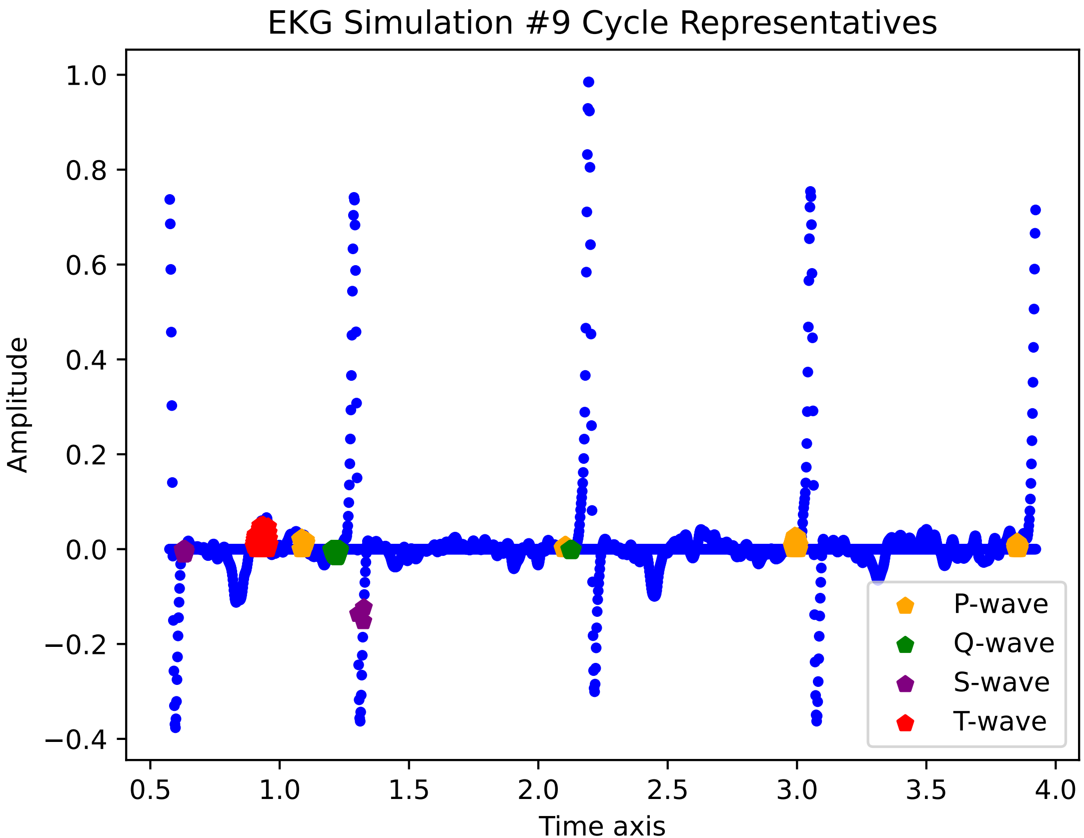
EKG Simulation #9 Cycle Representatives
Amplitude
Time axis
P-wave
Q-wave
S-wave
T-wave
1.0
0.8
0.6
0.4
0.2
0.0
−0.2
−0.4
0.5
1.0
1.5
2.0
2.5
3.0
3.5
4.0

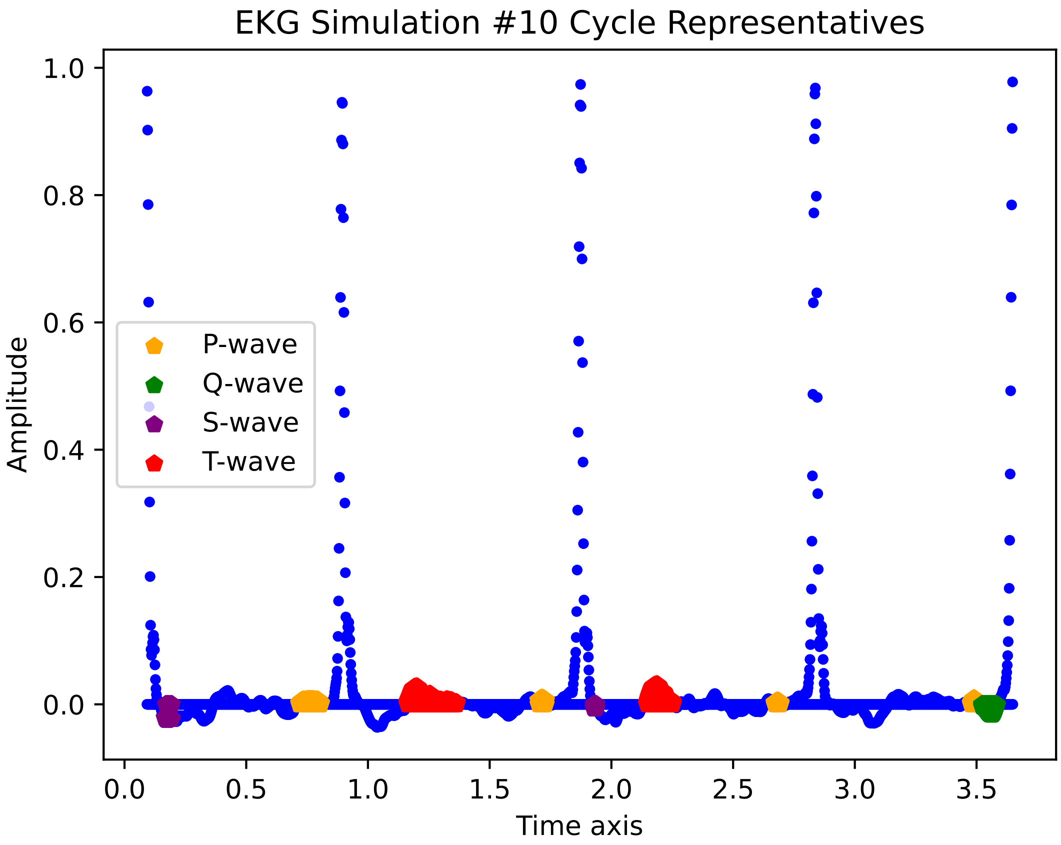

EKG Simulation #10 Cycle Representatives
Amplitude
Time axis
P-wave
Q-wave
S-wave
T-wave
0.0
0.2
0.4
0.6
0.8
1.0
0.0
0.5
1.0
1.5
2.0
2.5
3.0
3.5

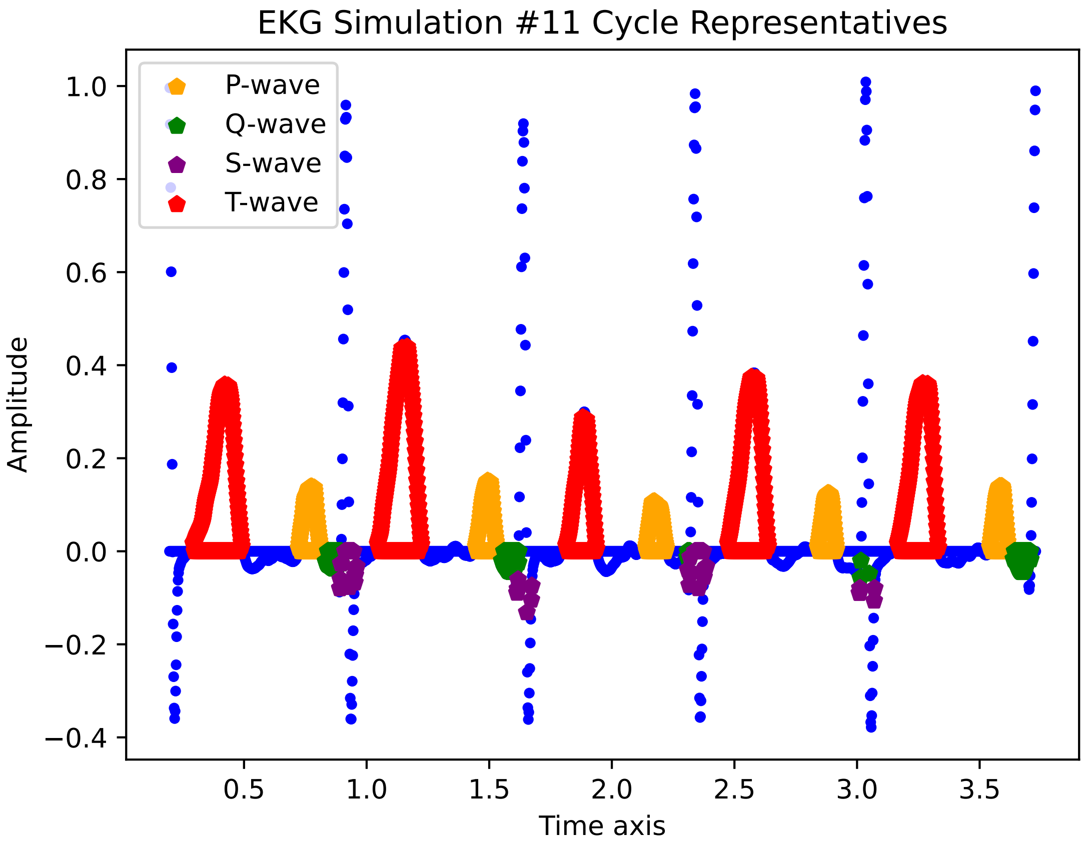

EKG Simulation #11 Cycle Representatives
P-wave
Q-wave
S-wave
T-wave
Amplitude
Time axis
1.0
0.8
0.6
0.4
0.2
0.0
−0.2
−0.4
0.5
1.0
1.5
2.0
2.5
3.0
3.5

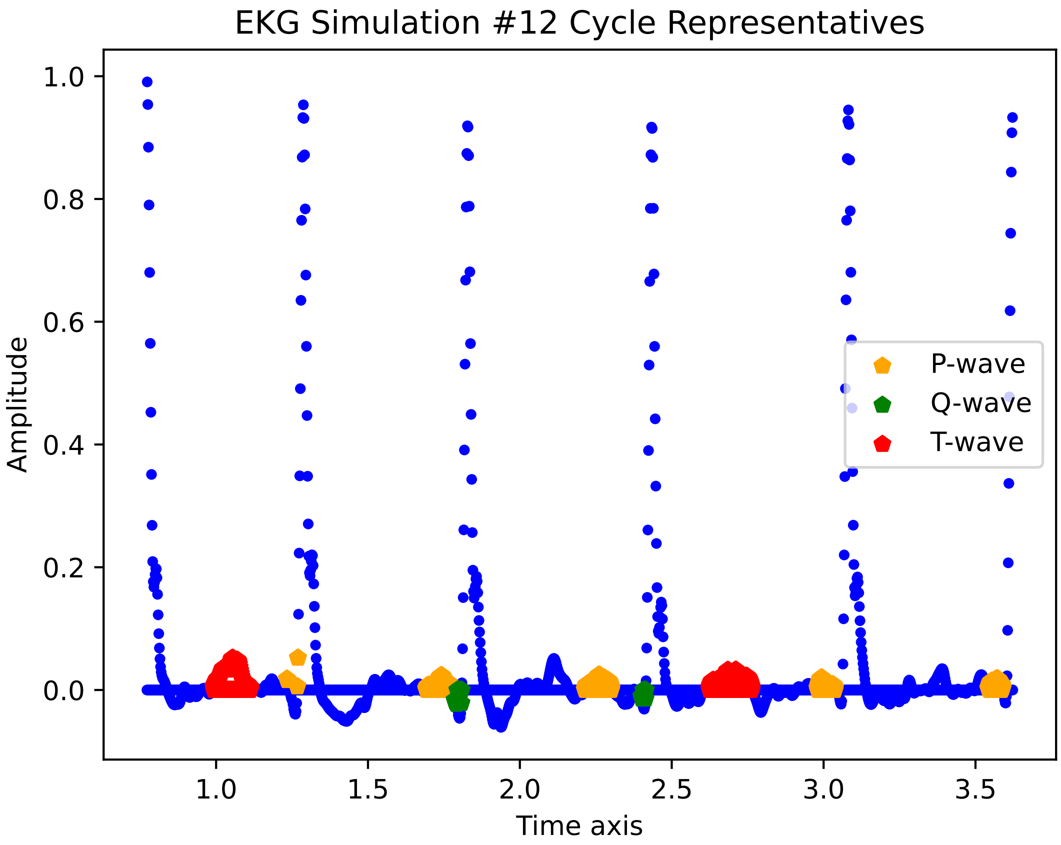

EKG Simulation #12 Cycle Representatives
Amplitude
Time axis
1.0
0.8
0.6
0.4
0.2
0.0
1.0
1.5
2.0
2.5
3.0
3.5
P-wave
Q-wave
T-wave

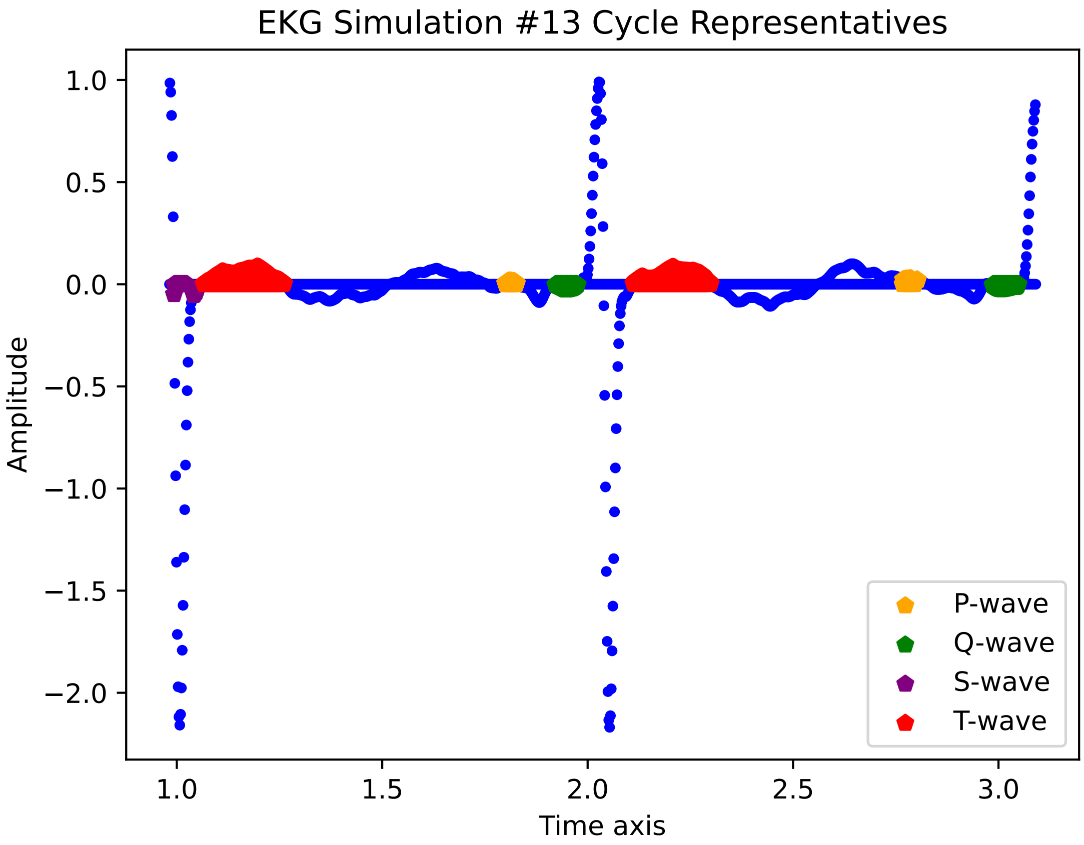

EKG Simulation #13 Cycle Representatives
Amplitude
Time axis
1.0
0.5
0.0
−0.5
−1.0
−1.5
−2.0
1.0
1.5
2.0
2.5
3.0
P-wave
Q-wave
S-wave
T-wave

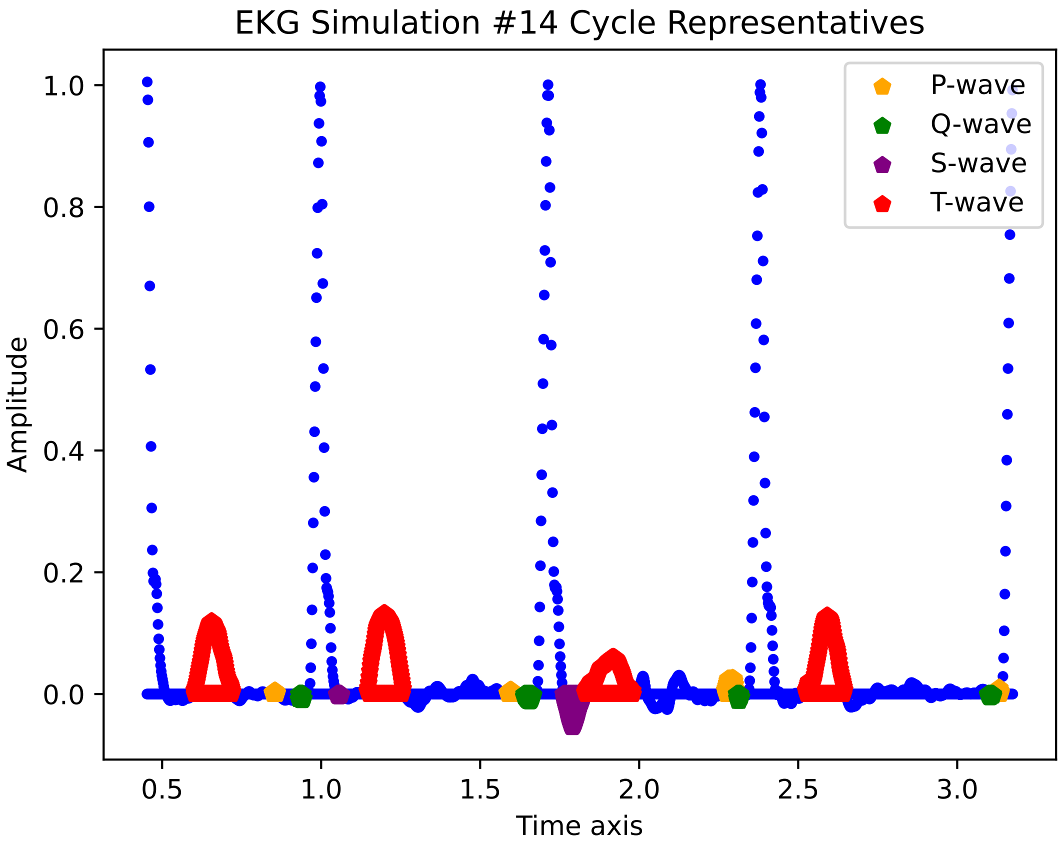
EKG Simulation #14 Cycle Representatives
P-wave
Q-wave
S-wave
T-wave
Amplitude
1.0
0.8
0.6
0.4
0.2
0.0
0.5
1.0
1.5
2.0
2.5
3.0
Time axis

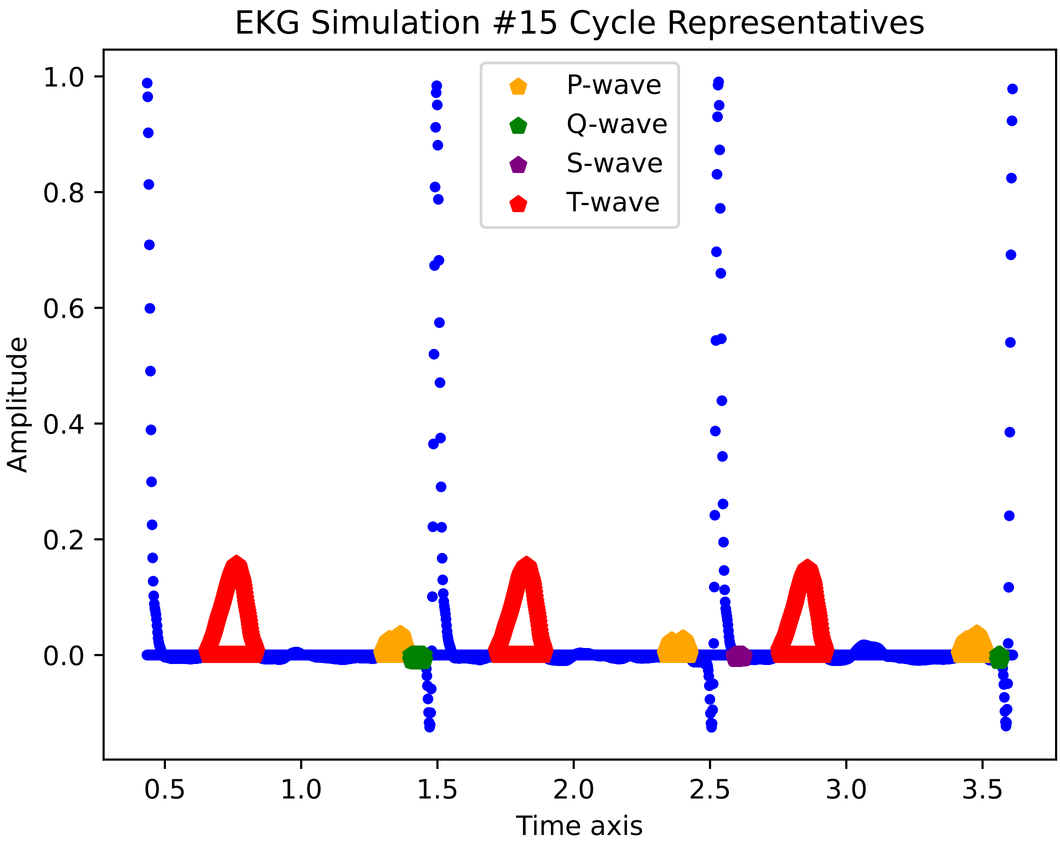

EKG Simulation #15 Cycle Representatives
P-wave
Q-wave
S-wave
T-wave
Amplitude
Time axis
1.0
0.8
0.6
0.4
0.2
0.0
0.5
1.0
1.5
2.0
2.5
3.0
3.5

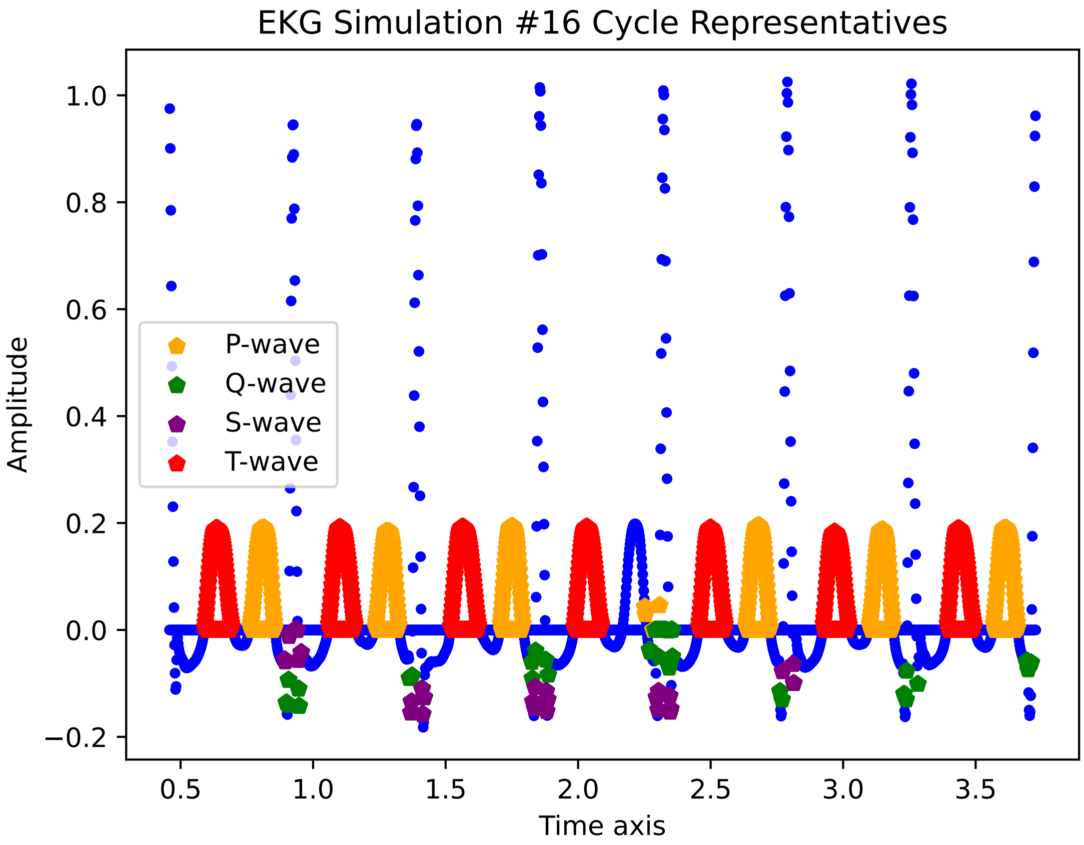

EKG Simulation #16 Cycle Representatives
Amplitude
Time axis
P-wave
Q-wave
S-wave
T-wave
1.0
0.8
0.6
0.4
0.2
0.0
−0.2
0.5
1.0
1.5
2.0
2.5
3.0
3.5

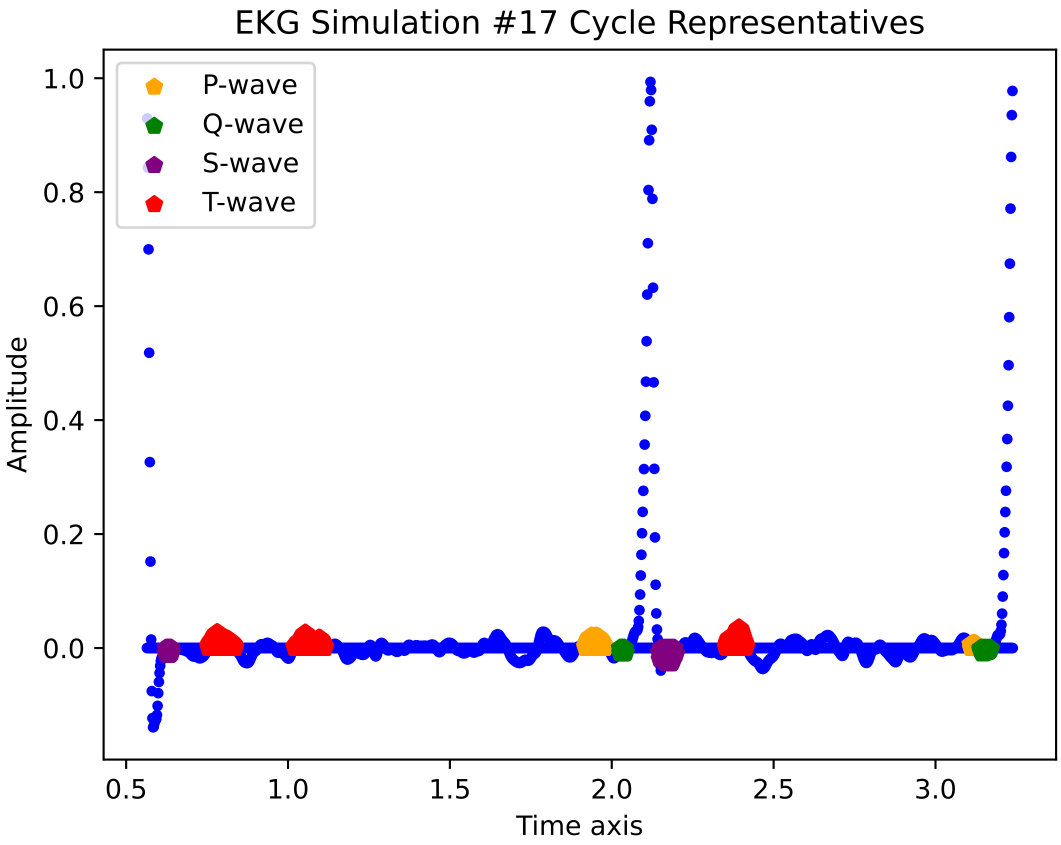
EKG Simulation #17 Cycle Representatives
P-wave
Q-wave
S-wave
T-wave
1.0
0.8
0.6
0.4
0.2
0.0
Amplitude
0.5
1.0
1.5
2.0
2.5
3.0
Time axis

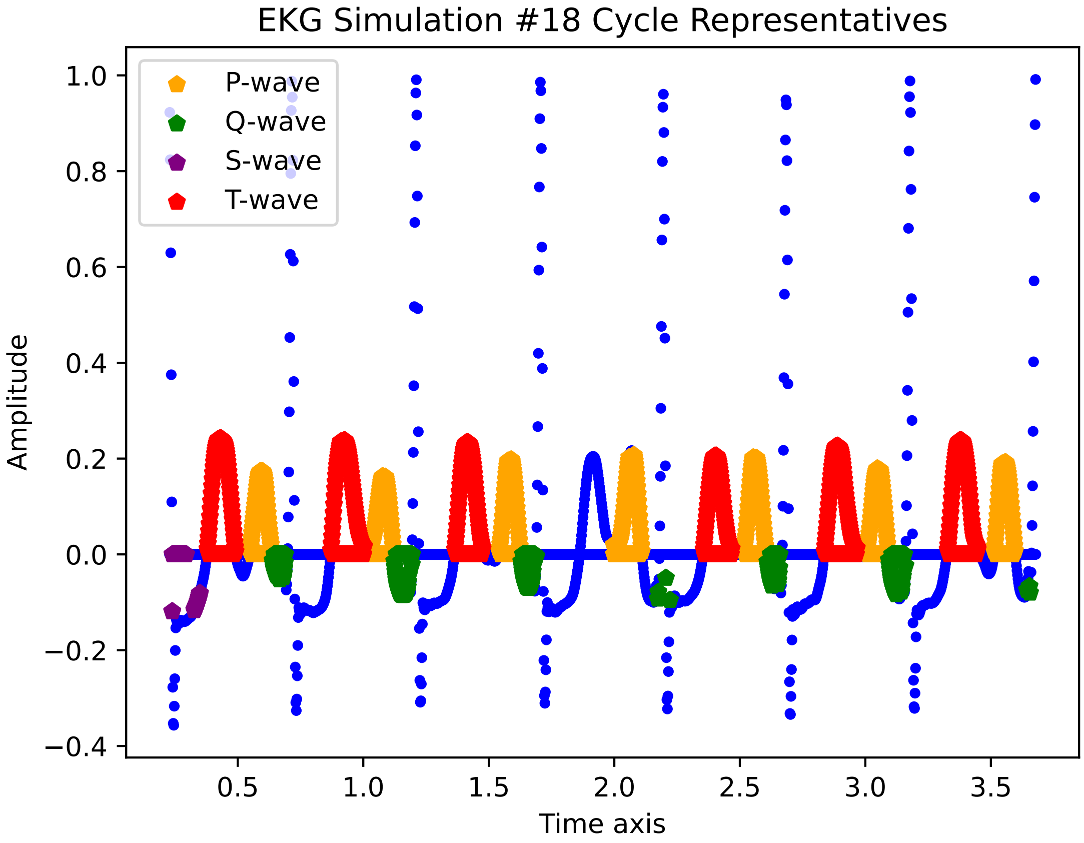
EKG Simulation #18 Cycle Representatives
P-wave
Q-wave
S-wave
T-wave
Amplitude
Time axis
1.0
0.8
0.6
0.4
0.2
0.0
−0.2
−0.4
0.5
1.0
1.5
2.0
2.5
3.0
3.5

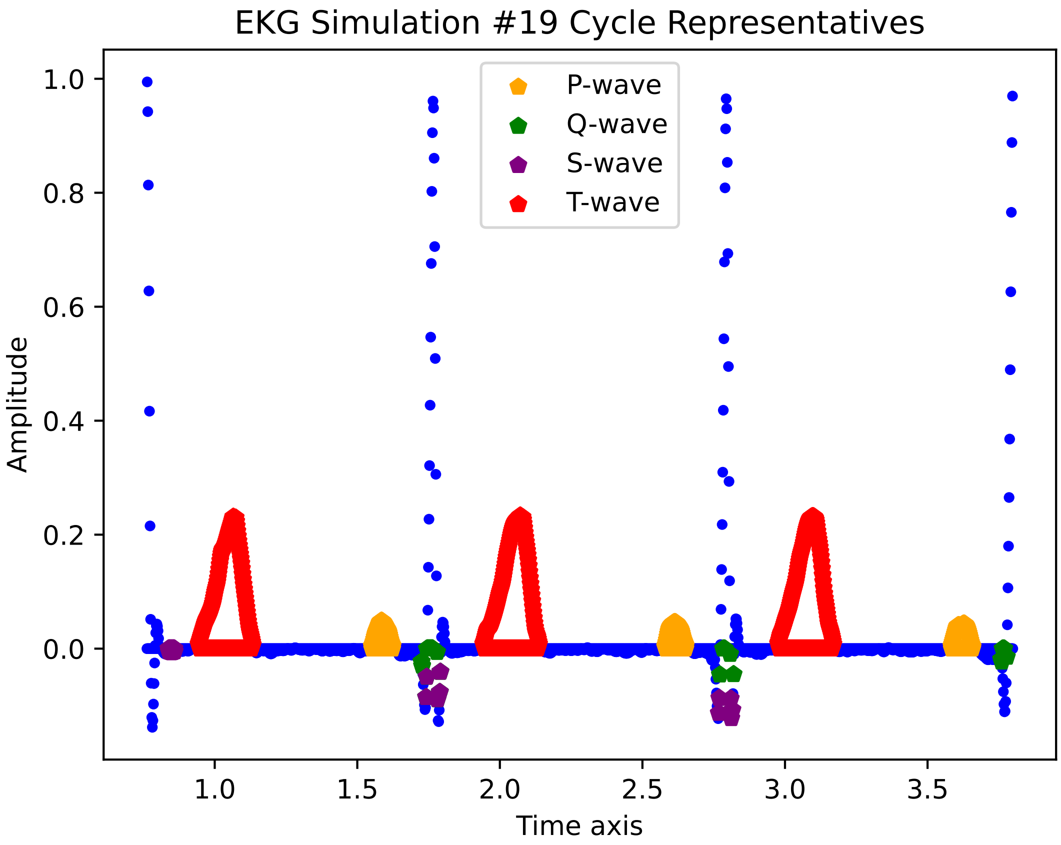
EKG Simulation #19 Cycle Representatives
P-wave
Q-wave
S-wave
T-wave
Amplitude
1.0
0.8
0.6
0.4
0.2
0.0
1.0
1.5
2.0
2.5
3.0
3.5
Time axis

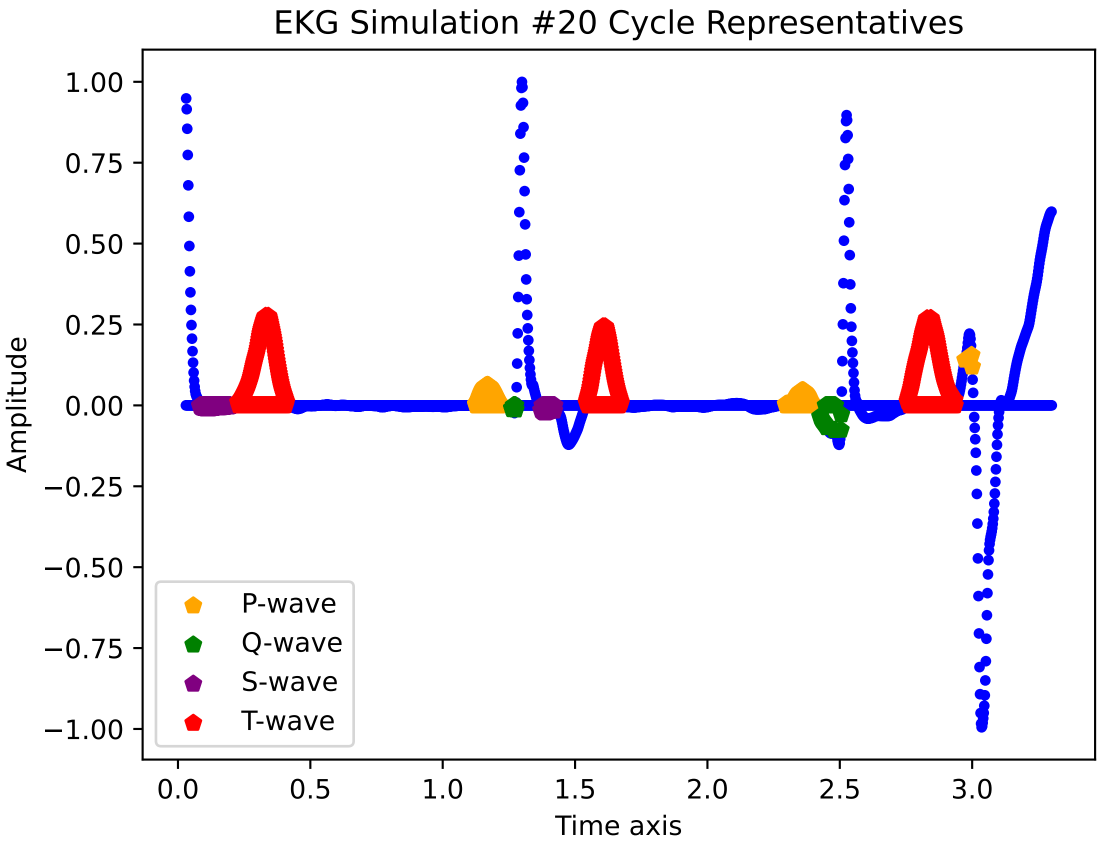

EKG Simulation #20 Cycle Representatives
Amplitude
Time axis
1.00
0.75
0.50
0.25
0.00
−0.25
−0.50
−0.75
−1.00
0.0
0.5
1.0
1.5
2.0
2.5
3.0
P-wave
Q-wave
S-wave
T-wave

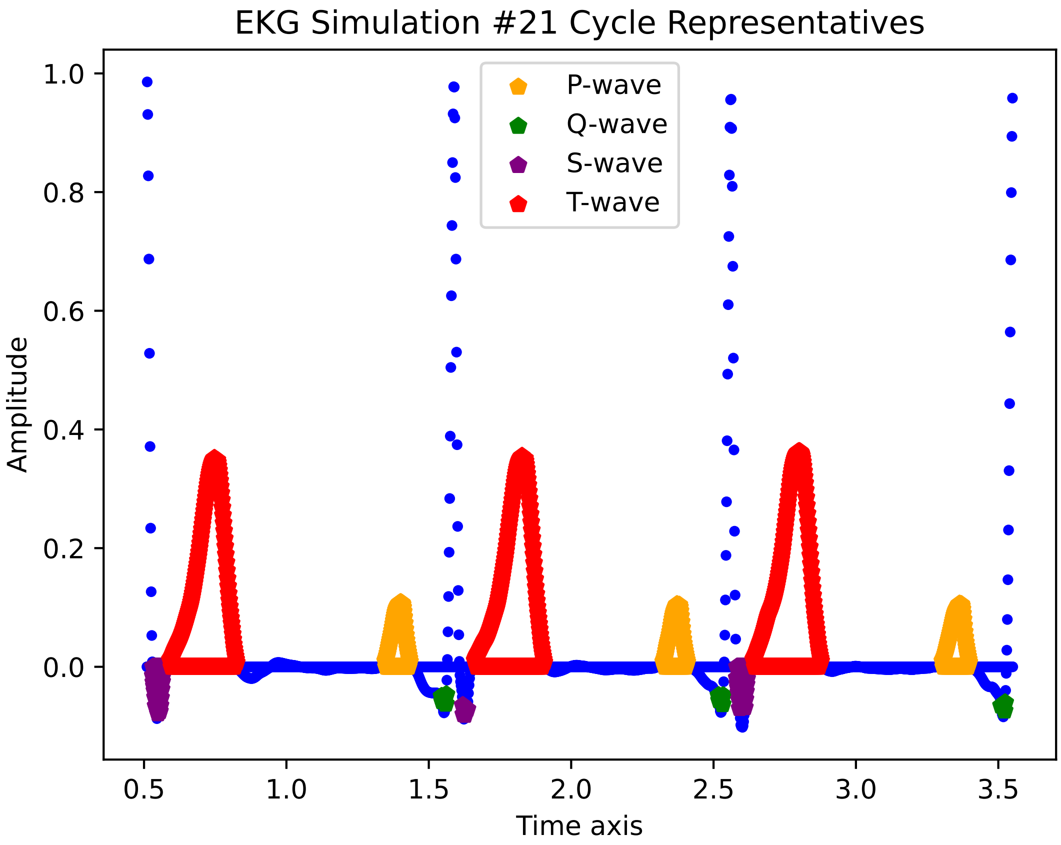
EKG Simulation #21 Cycle Representatives
P-wave
Q-wave
S-wave
T-wave
Amplitude
Time axis
1.0
0.8
0.6
0.4
0.2
0.0
0.5
1.0
1.5
2.0
2.5
3.0
3.5

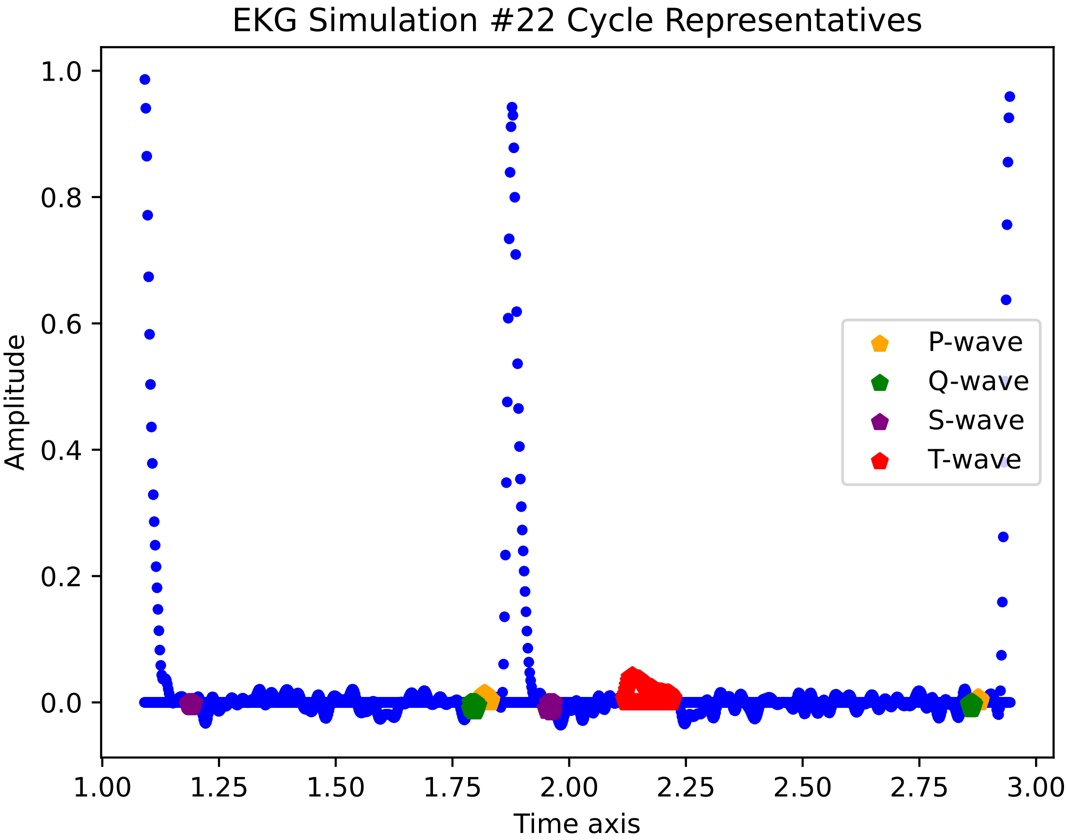
EKG Simulation #22 Cycle Representatives
Amplitude
Time axis
1.0
0.8
0.6
0.4
0.2
0.0
1.00
1.25
1.50
1.75
2.00
2.25
2.50
2.75
3.00
P-wave
Q-wave
S-wave
T-wave

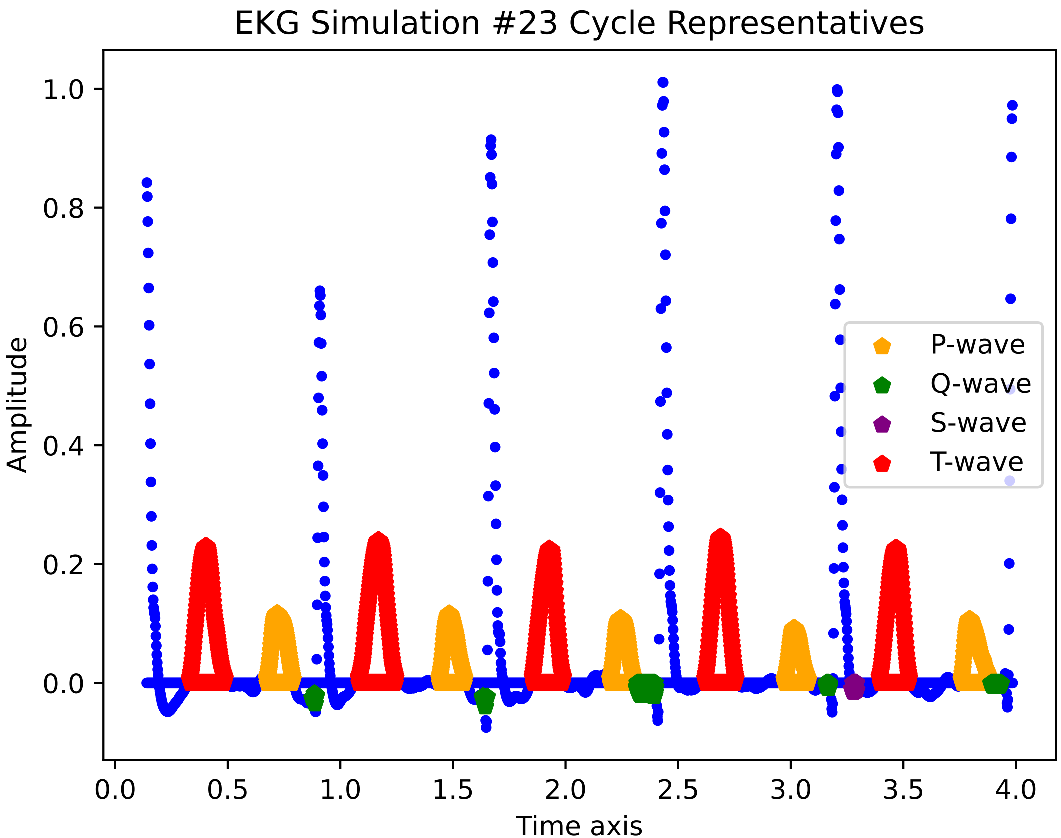
EKG Simulation #23 Cycle Representatives
Amplitude
Time axis
1.0
0.8
0.6
0.4
0.2
0.0
0.0
0.5
1.0
1.5
2.0
2.5
3.0
3.5
4.0
P-wave
Q-wave
S-wave
T-wave

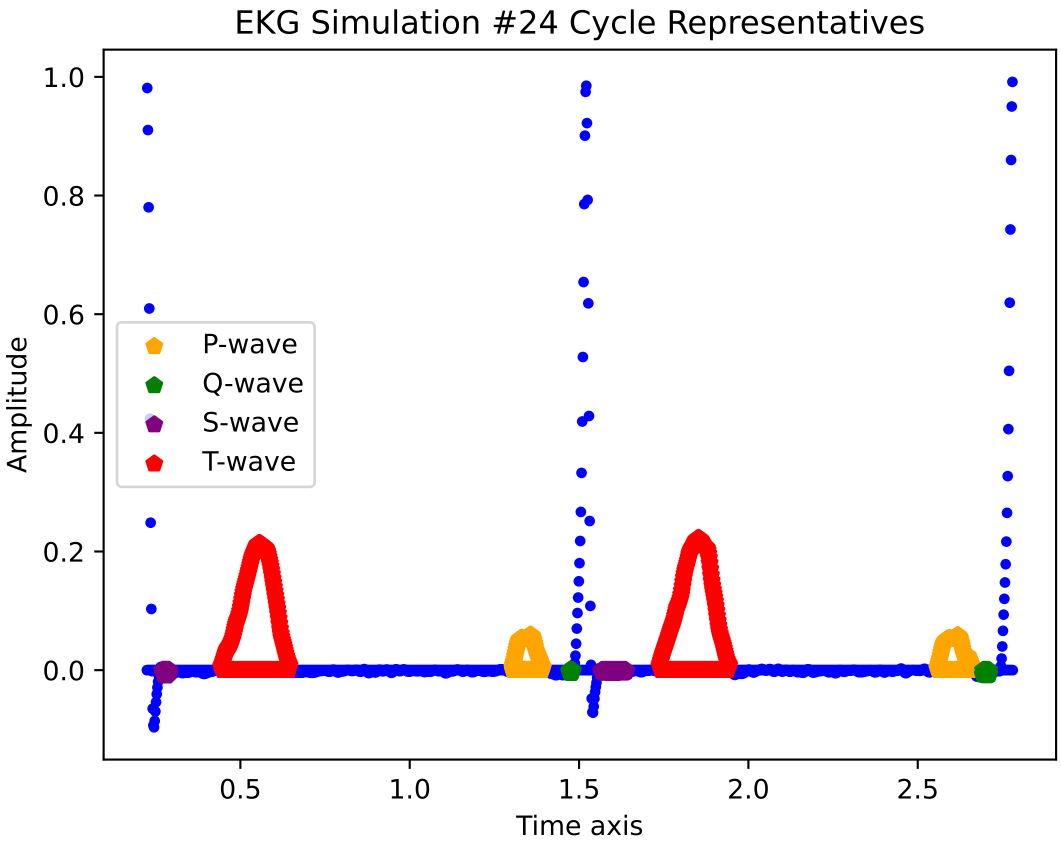
EKG Simulation #24 Cycle Representatives
Amplitude
Time axis
1.0
0.8
0.6
0.4
0.2
0.0
0.5
1.0
1.5
2.0
2.5
P-wave
Q-wave
S-wave
T-wave

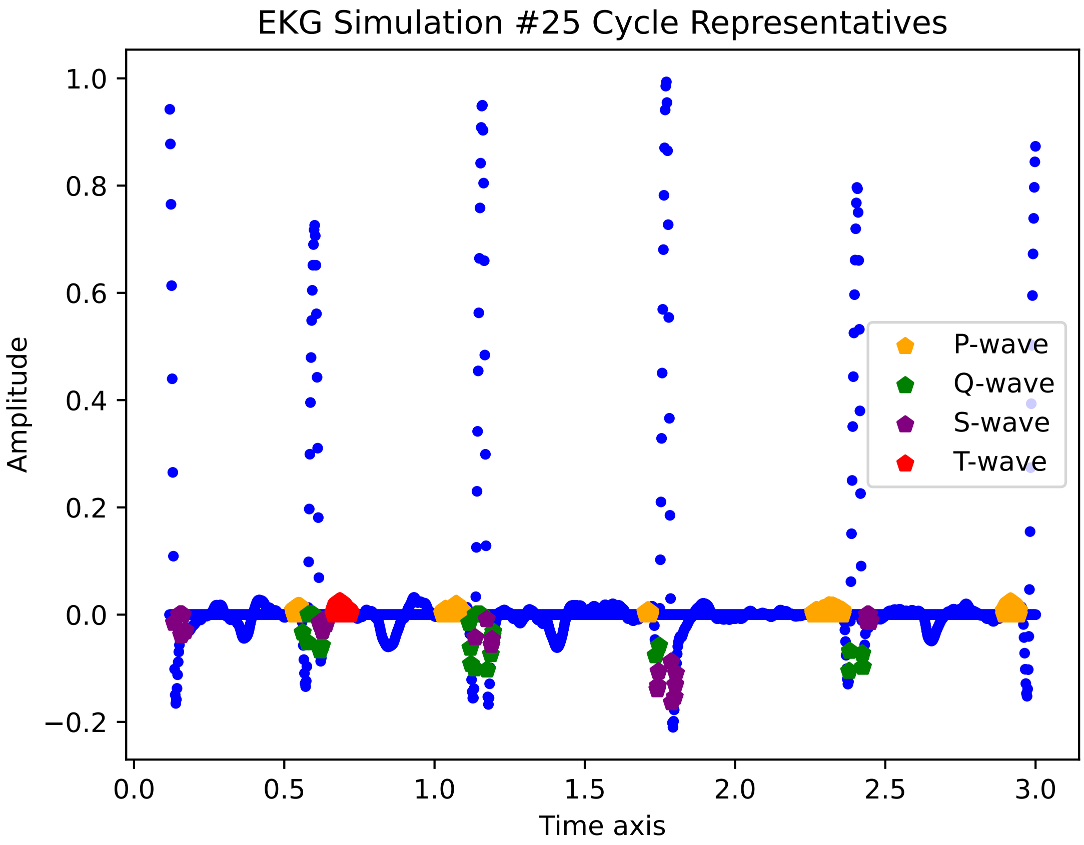
EKG Simulation #25 Cycle Representatives
Amplitude
Time axis
1.0
0.8
0.6
0.4
0.2
0.0
−0.2
0.0
0.5
1.0
1.5
2.0
2.5
3.0
P-wave
Q-wave
S-wave
T-wave

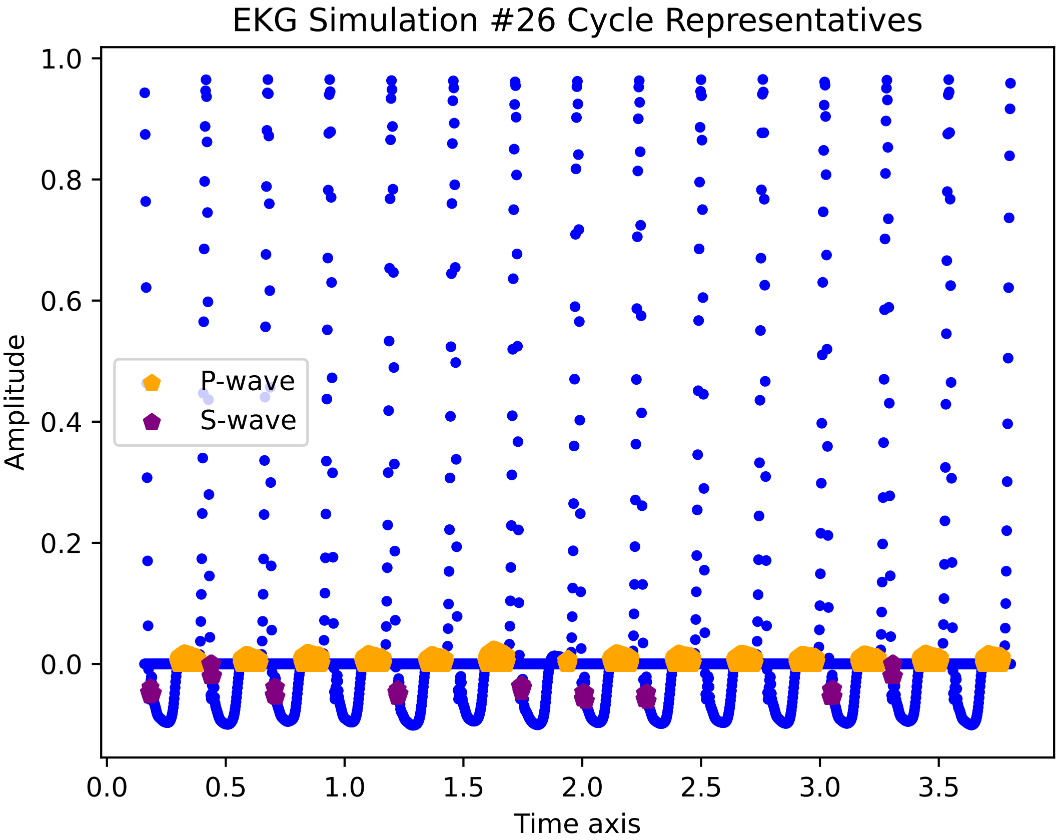

EKG Simulation #26 Cycle Representatives
1.0
0.8
0.6
0.4
0.2
0.0
Amplitude
P-wave
S-wave
0.0
0.5
1.0
1.5
2.0
2.5
3.0
3.5
Time axis

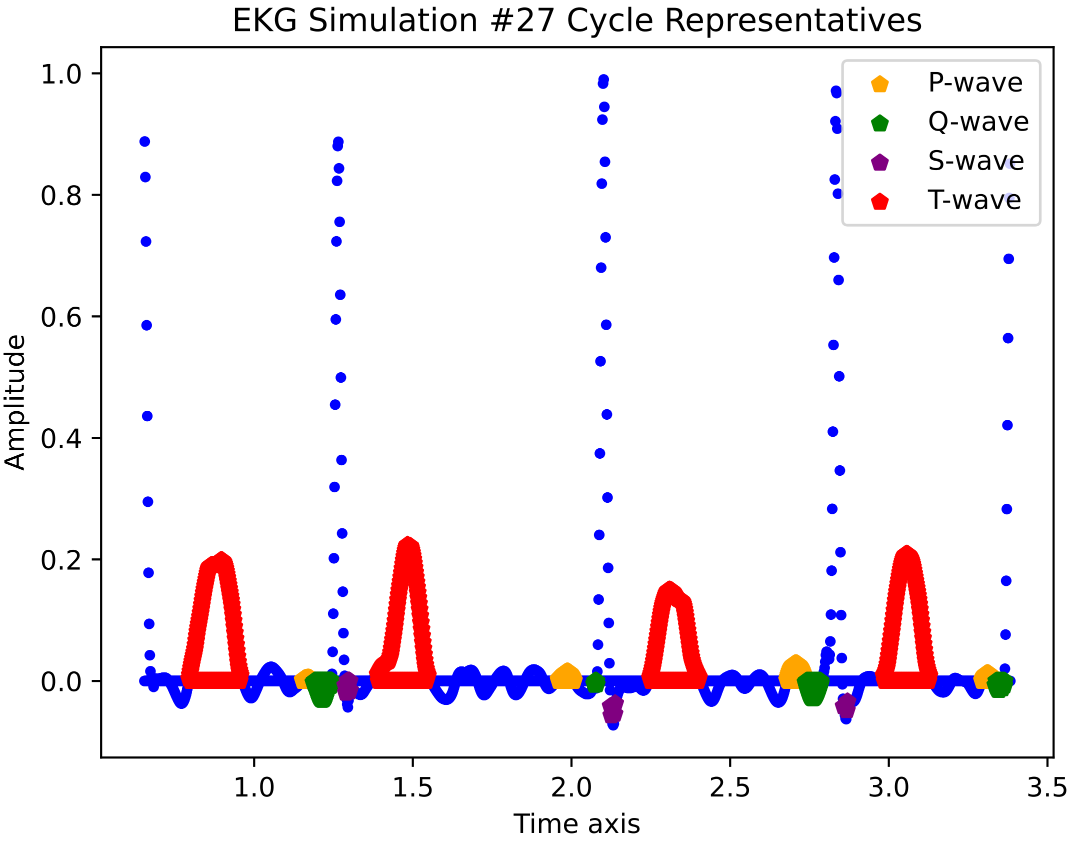
EKG Simulation #27 Cycle Representatives
P-wave
Q-wave
S-wave
T-wave
Amplitude
Time axis
1.0
0.8
0.6
0.4
0.2
0.0
1.0
1.5
2.0
2.5
3.0
3.5

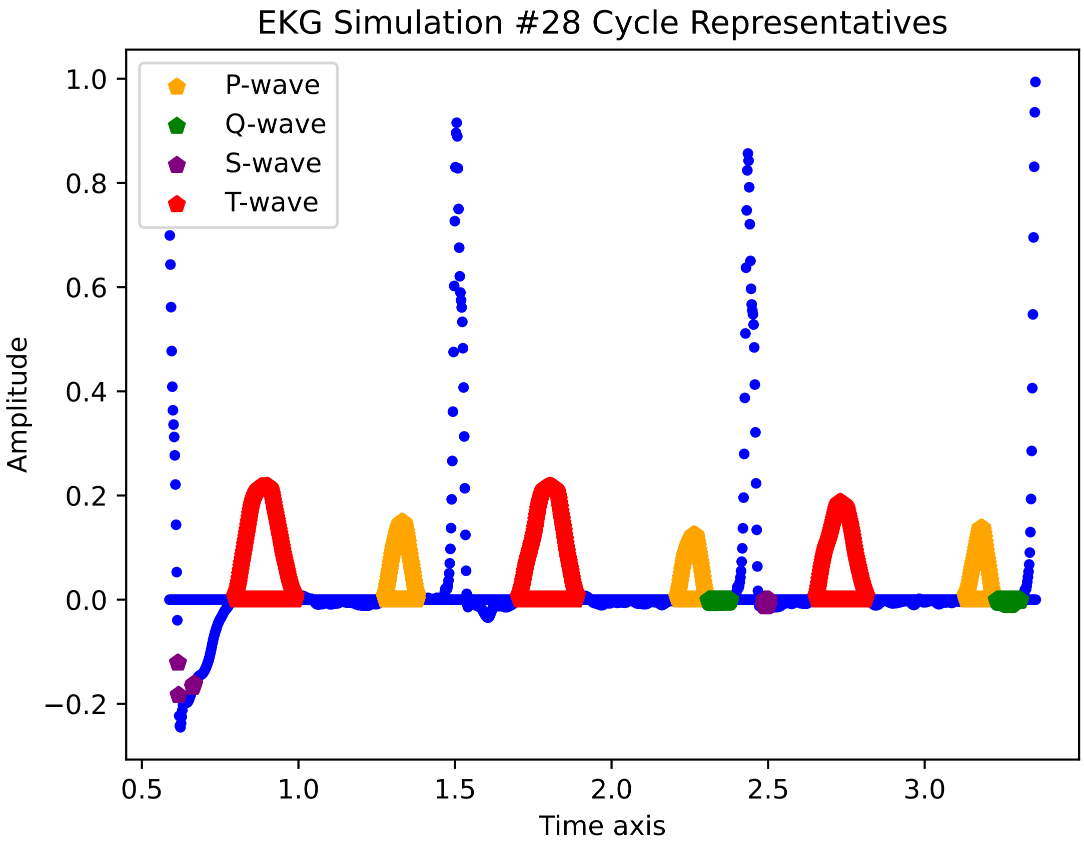
EKG Simulation #28 Cycle Representatives
P-wave
Q-wave
S-wave
T-wave
Amplitude
Time axis
1.0
0.8
0.6
0.4
0.2
0.0
−0.2
0.5
1.0
1.5
2.0
2.5
3.0

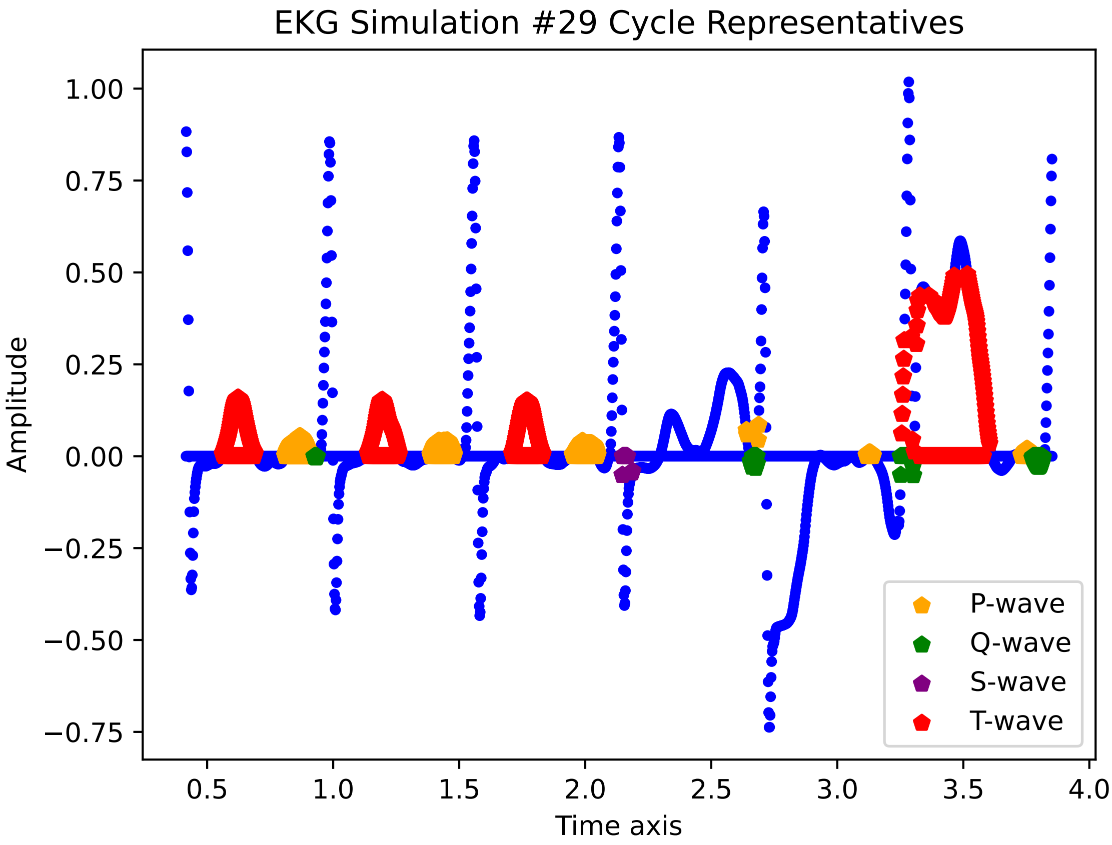
EKG Simulation #29 Cycle Representatives
Amplitude
1.00
0.75
0.50
0.25
0.00
−0.25
−0.50
−0.75
0.5
1.0
1.5
2.0
2.5
3.0
3.5
4.0
Time axis
P-wave
Q-wave
S-wave
T-wave

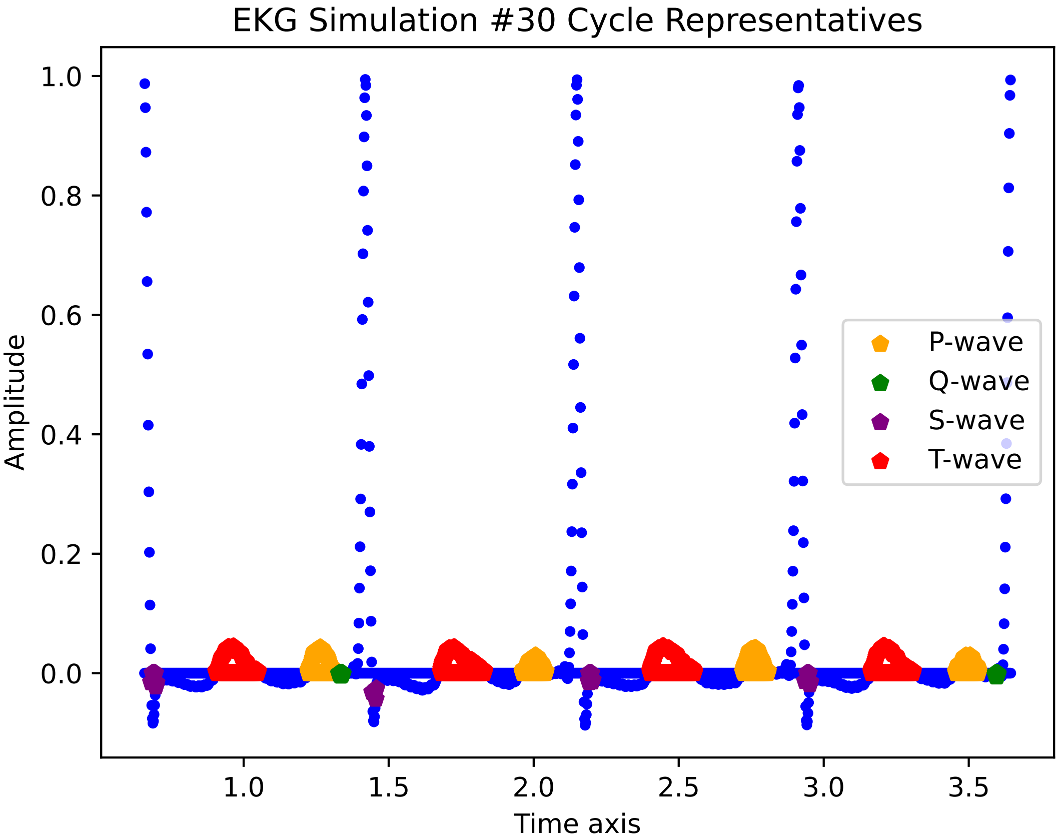
EKG Simulation #30 Cycle Representatives
Amplitude
Time axis
1.0
0.8
0.6
0.4
0.2
0.0
1.0
1.5
2.0
2.5
3.0
3.5
P-wave
Q-wave
S-wave
T-wave

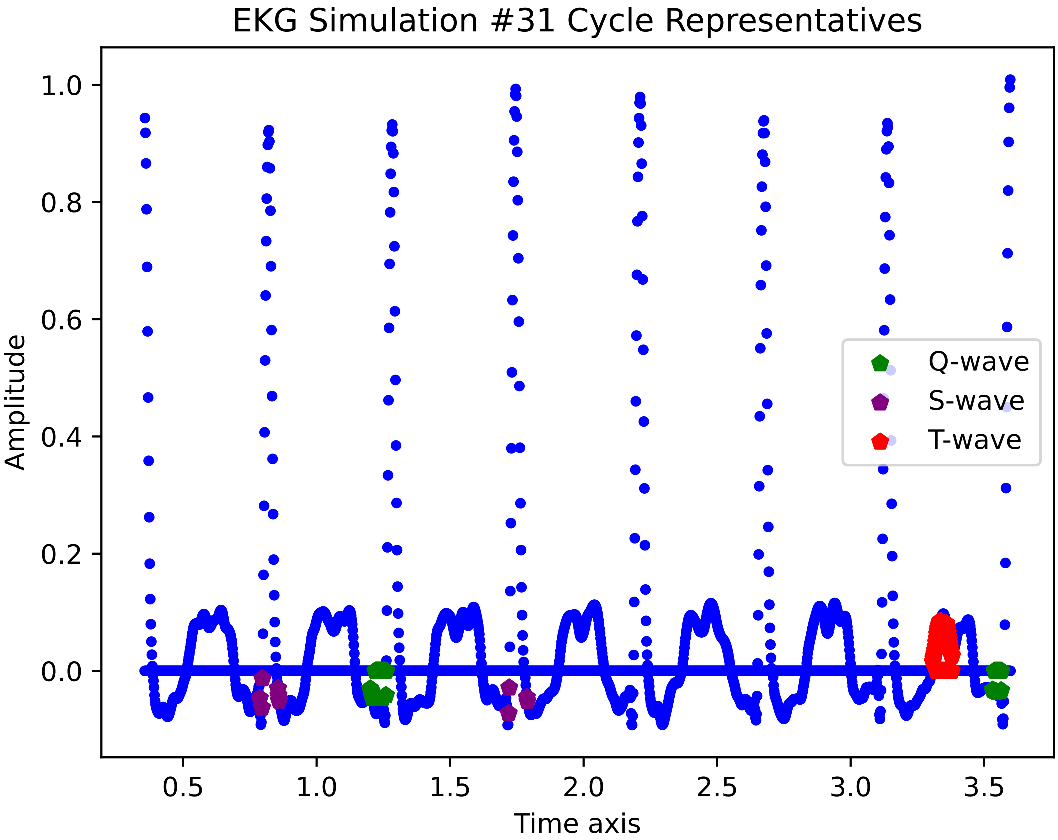
EKG Simulation #31 Cycle Representatives
Amplitude
Time axis
1.0
0.8
0.6
0.4
0.2
0.0
0.5
1.0
1.5
2.0
2.5
3.0
3.5
Q-wave
S-wave
T-wave

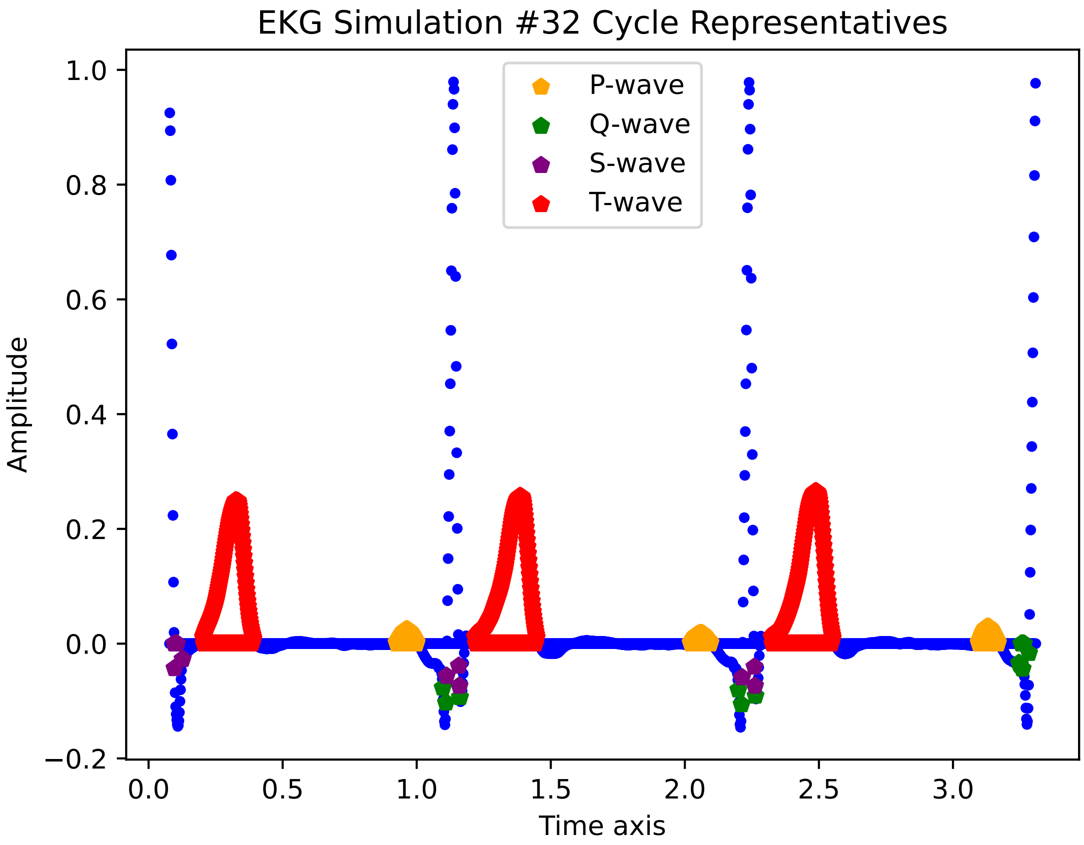
EKG Simulation #32 Cycle Representatives
P-wave
Q-wave
S-wave
T-wave
Amplitude
Time axis
1.0
0.8
0.6
0.4
0.2
0.0
−0.2
0.0
0.5
1.0
1.5
2.0
2.5
3.0

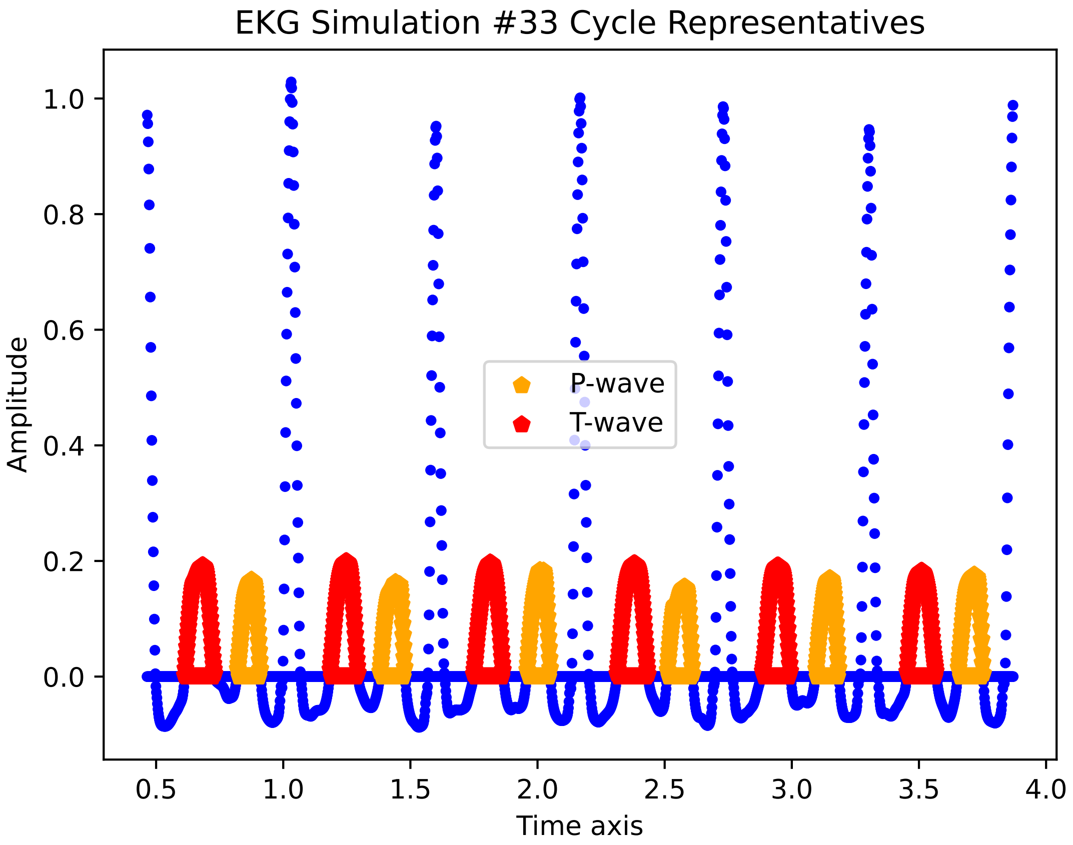
EKG Simulation #33 Cycle Representatives
Amplitude
Time axis
P-wave
T-wave
1.0
0.8
0.6
0.4
0.2
0.0
0.5
1.0
1.5
2.0
2.5
3.0
3.5
4.0

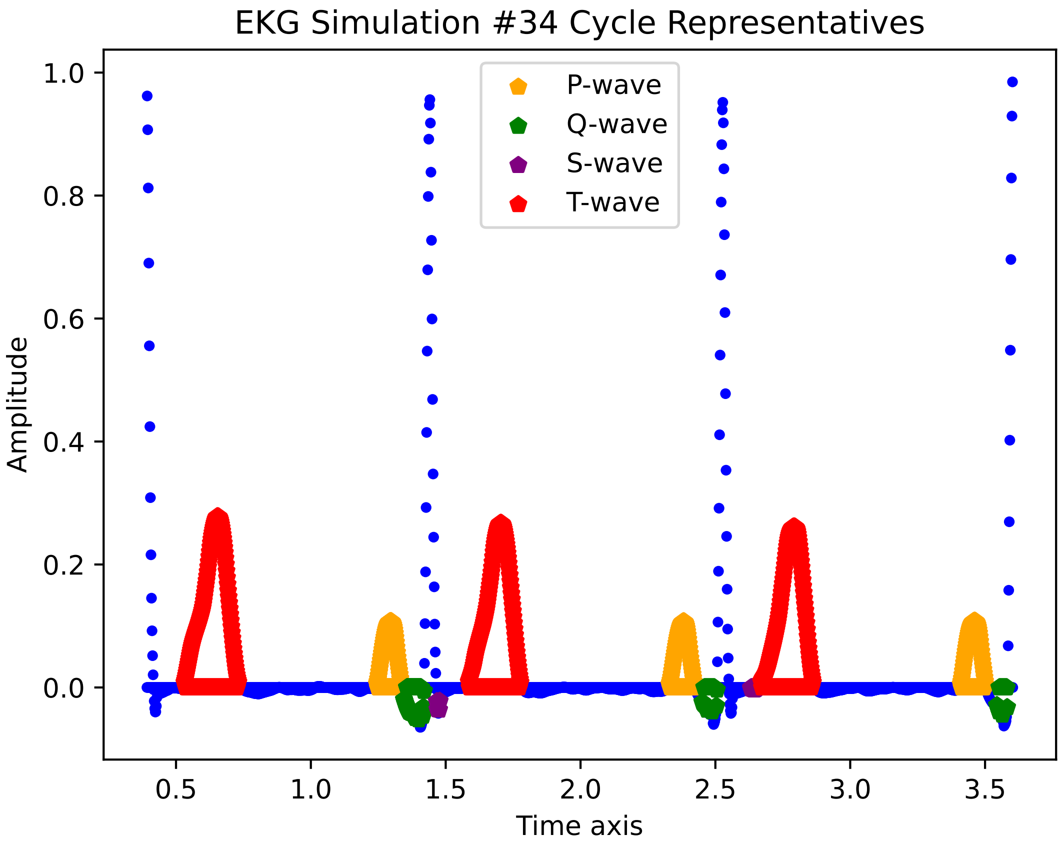

EKG Simulation #34 Cycle Representatives
P-wave
Q-wave
S-wave
T-wave
Amplitude
1.0
0.8
0.6
0.4
0.2
0.0
0.5
1.0
1.5
2.0
2.5
3.0
3.5
Time axis

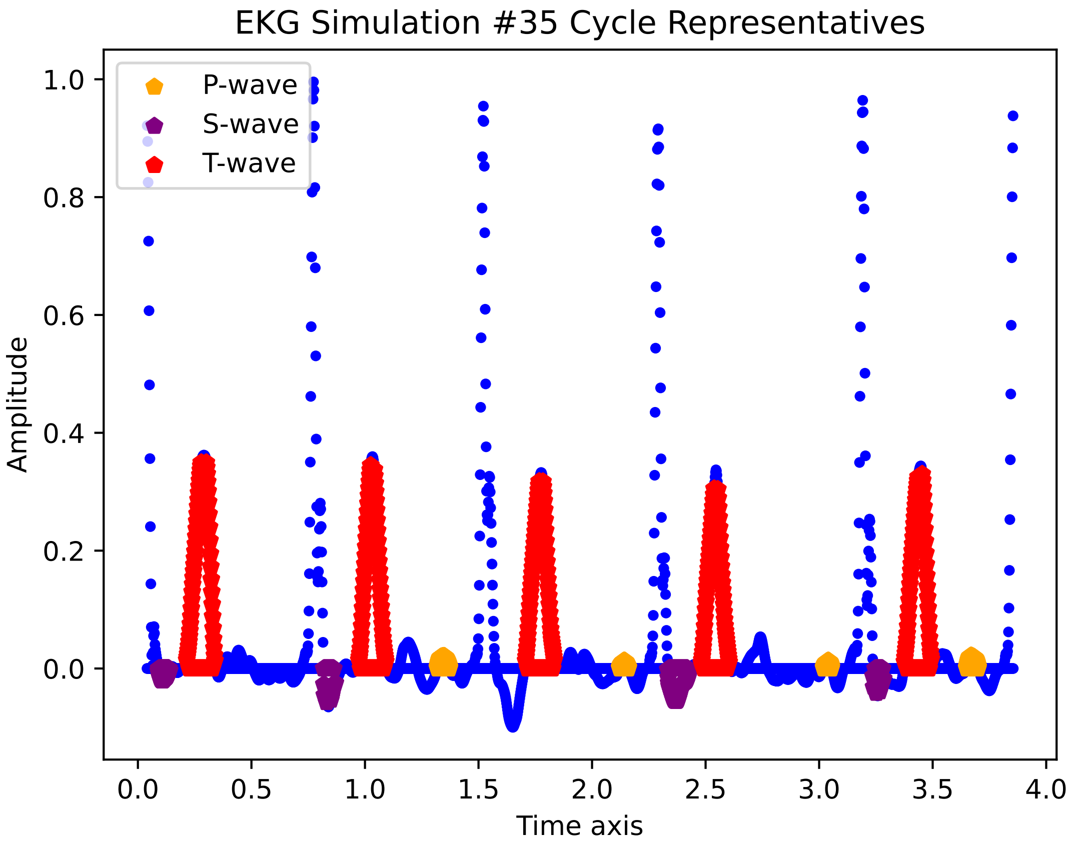
EKG Simulation #35 Cycle Representatives
P-wave
S-wave
T-wave
Amplitude
Time axis
1.0
0.8
0.6
0.4
0.2
0.0
0.0
0.5
1.0
1.5
2.0
2.5
3.0
3.5
4.0

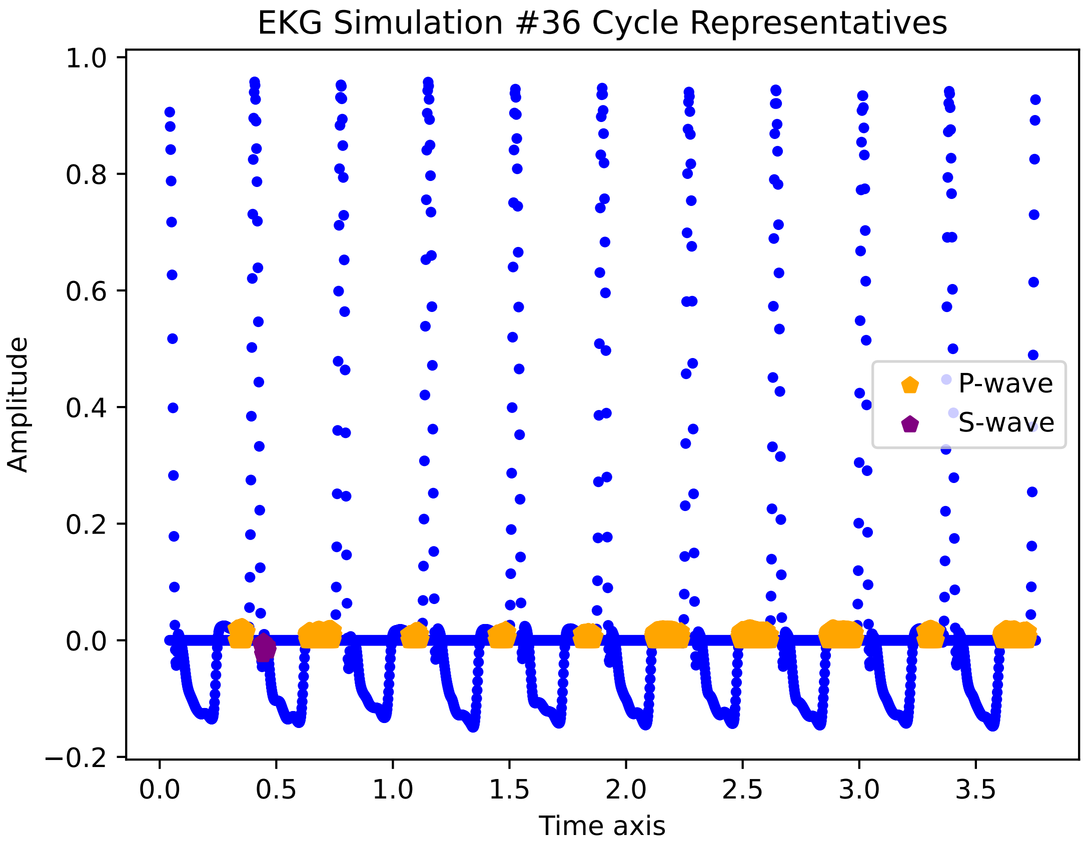

EKG Simulation #36 Cycle Representatives
Amplitude
Time axis
P-wave
S-wave
1.0
0.8
0.6
0.4
0.2
0.0
−0.2
0.0
0.5
1.0
1.5
2.0
2.5
3.0
3.5

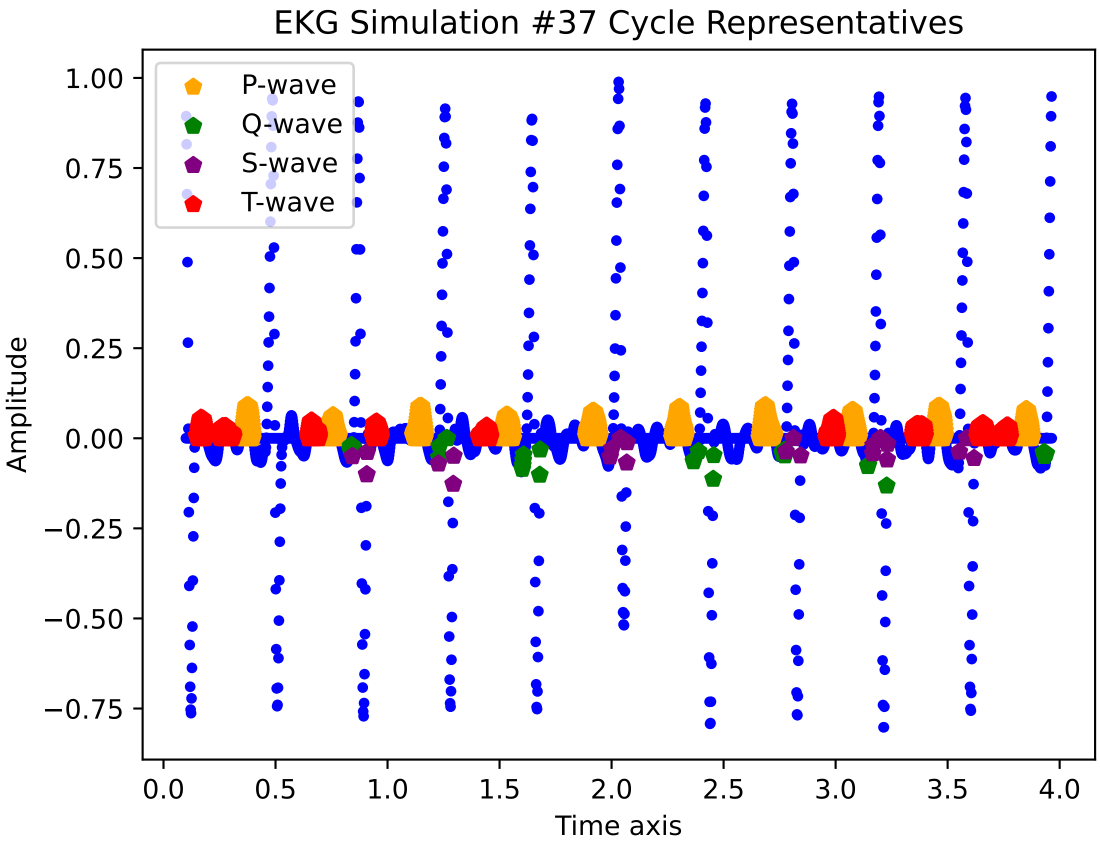
EKG Simulation #37 Cycle Representatives
P-wave
Q-wave
S-wave
T-wave
Amplitude
Time axis
1.00
0.75
0.50
0.25
0.00
−0.25
−0.50
−0.75
0.0
0.5
1.0
1.5
2.0
2.5
3.0
3.5
4.0

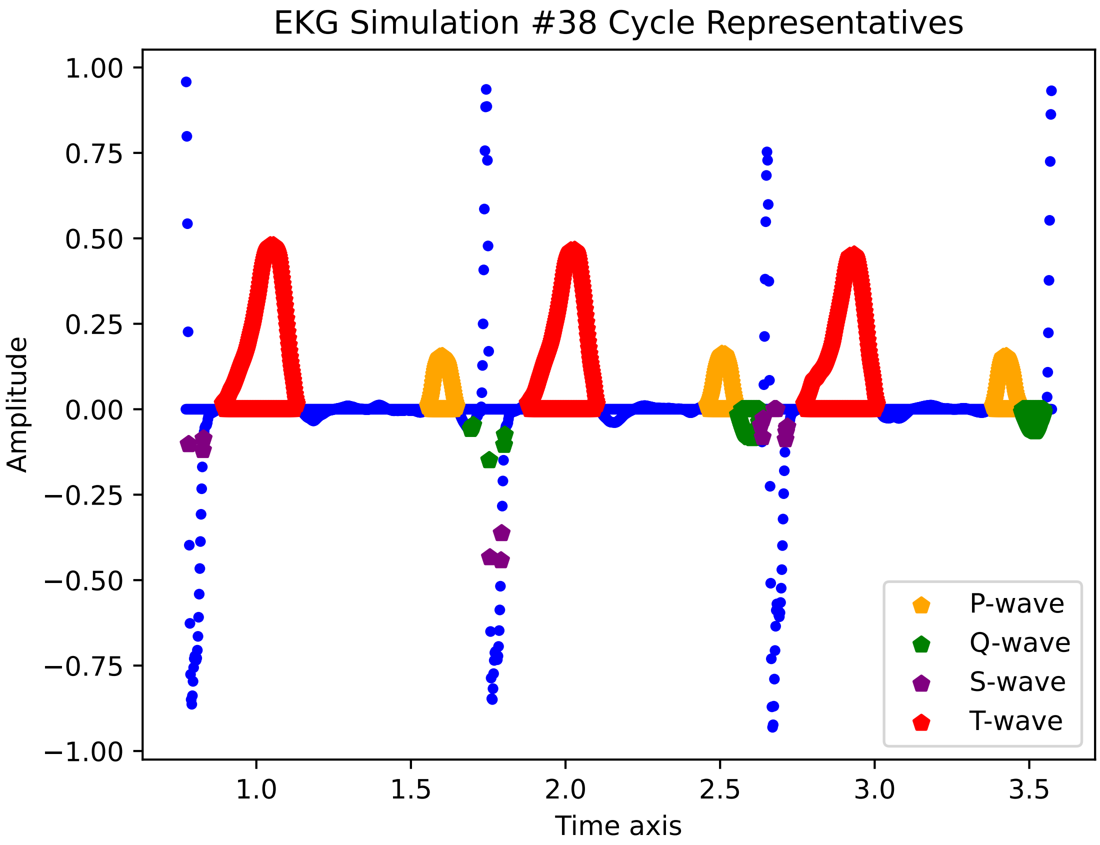
EKG Simulation #38 Cycle Representatives
Amplitude
Time axis
1.00
0.75
0.50
0.25
0.00
−0.25
−0.50
−0.75
−1.00
1.0
1.5
2.0
2.5
3.0
3.5
P-wave
Q-wave
S-wave
T-wave

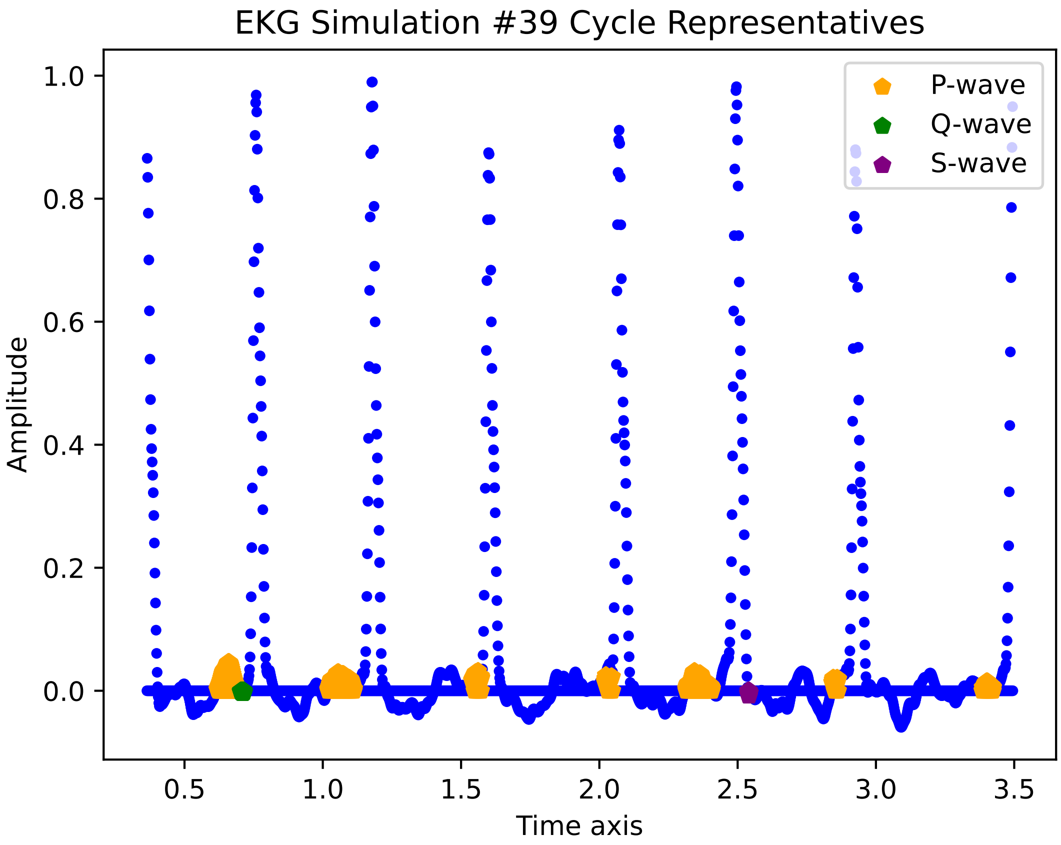
EKG Simulation #39 Cycle Representatives
P-wave
Q-wave
S-wave
Amplitude
1.0
0.8
0.6
0.4
0.2
0.0
0.5
1.0
1.5
2.0
2.5
3.0
3.5
Time axis

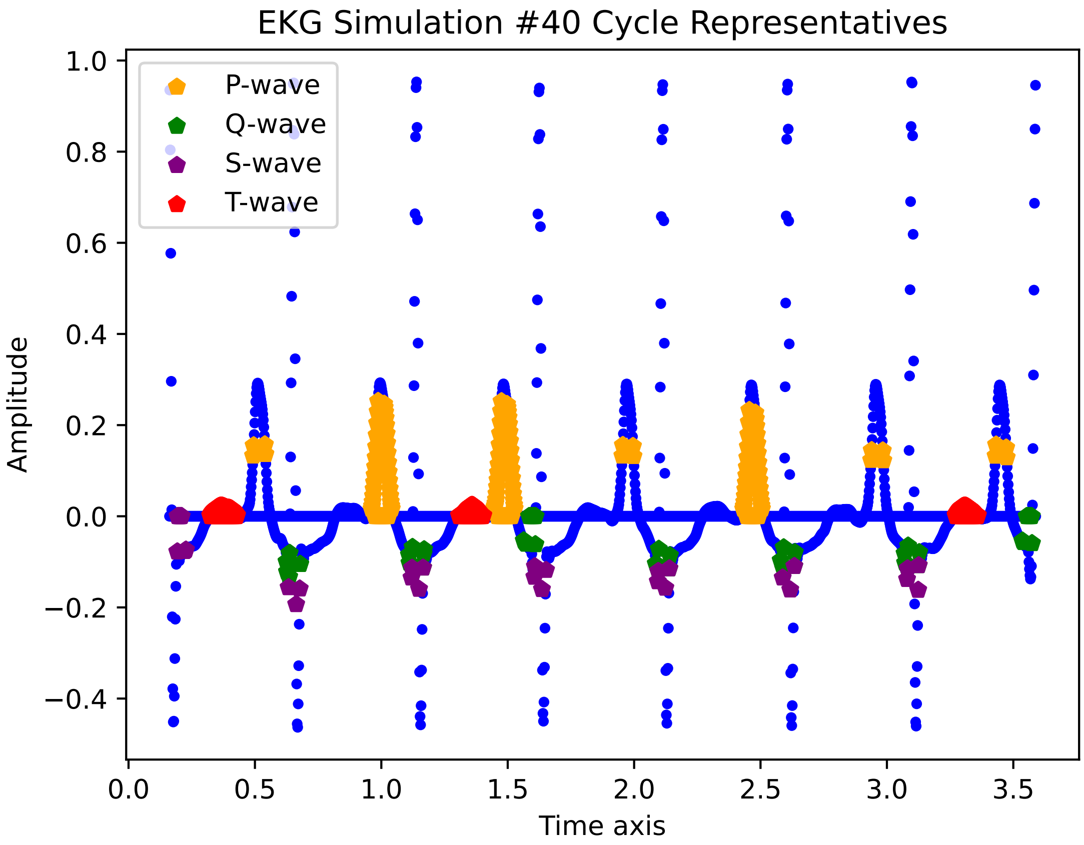
EKG Simulation #40 Cycle Representatives
P-wave
Q-wave
S-wave
T-wave
Amplitude
Time axis
1.0
0.8
0.6
0.4
0.2
0.0
−0.2
−0.4
0.0
0.5
1.0
1.5
2.0
2.5
3.0
3.5

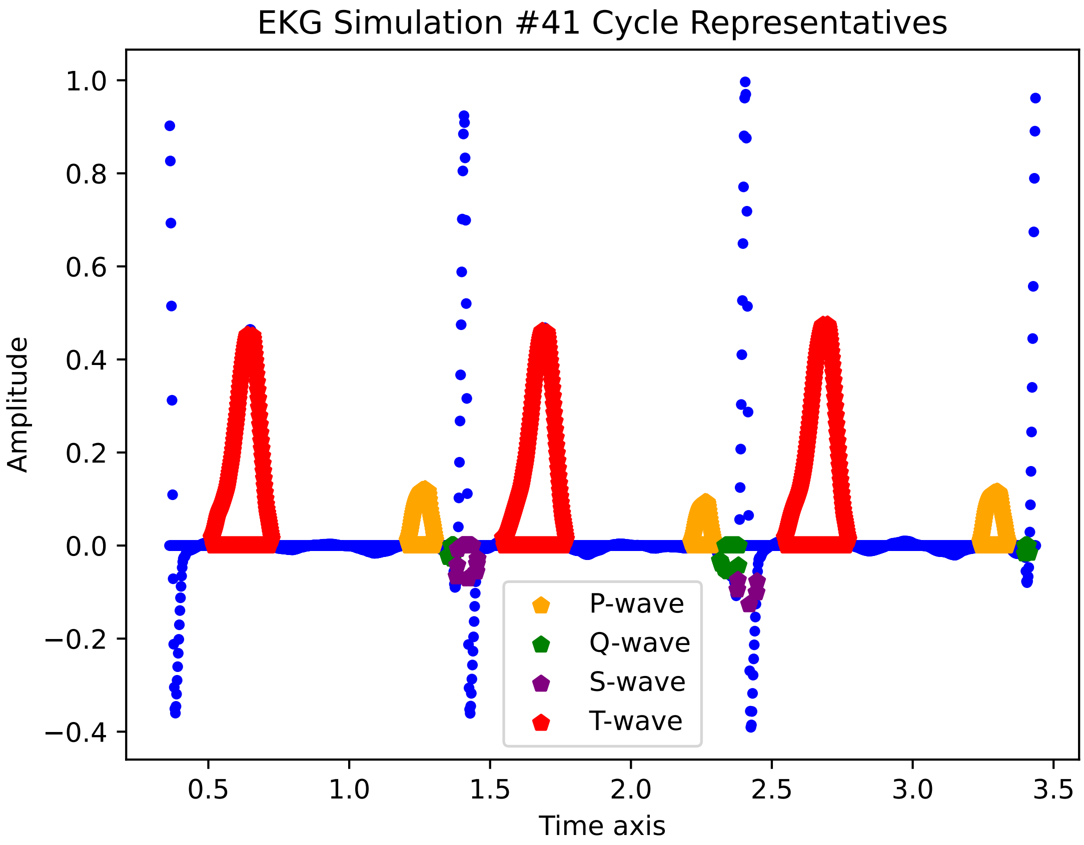
EKG Simulation #41 Cycle Representatives
1.0
0.8
0.6
0.4
0.2
0.0
−0.2
−0.4
Amplitude
0.5
1.0
1.5
2.0
2.5
3.0
3.5
Time axis
P-wave
Q-wave
S-wave
T-wave

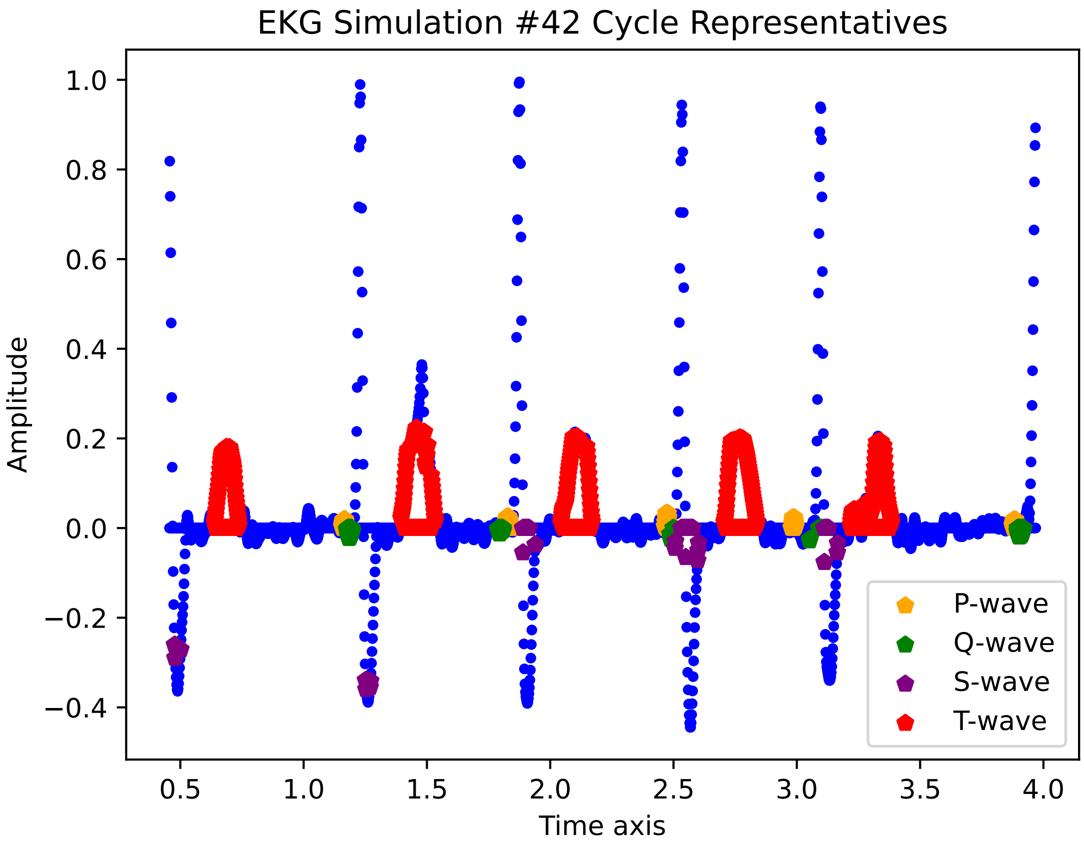
EKG Simulation #42 Cycle Representatives
Amplitude
Time axis
1.0
0.8
0.6
0.4
0.2
0.0
−0.2
−0.4
0.5
1.0
1.5
2.0
2.5
3.0
3.5
4.0
P-wave
Q-wave
S-wave
T-wave

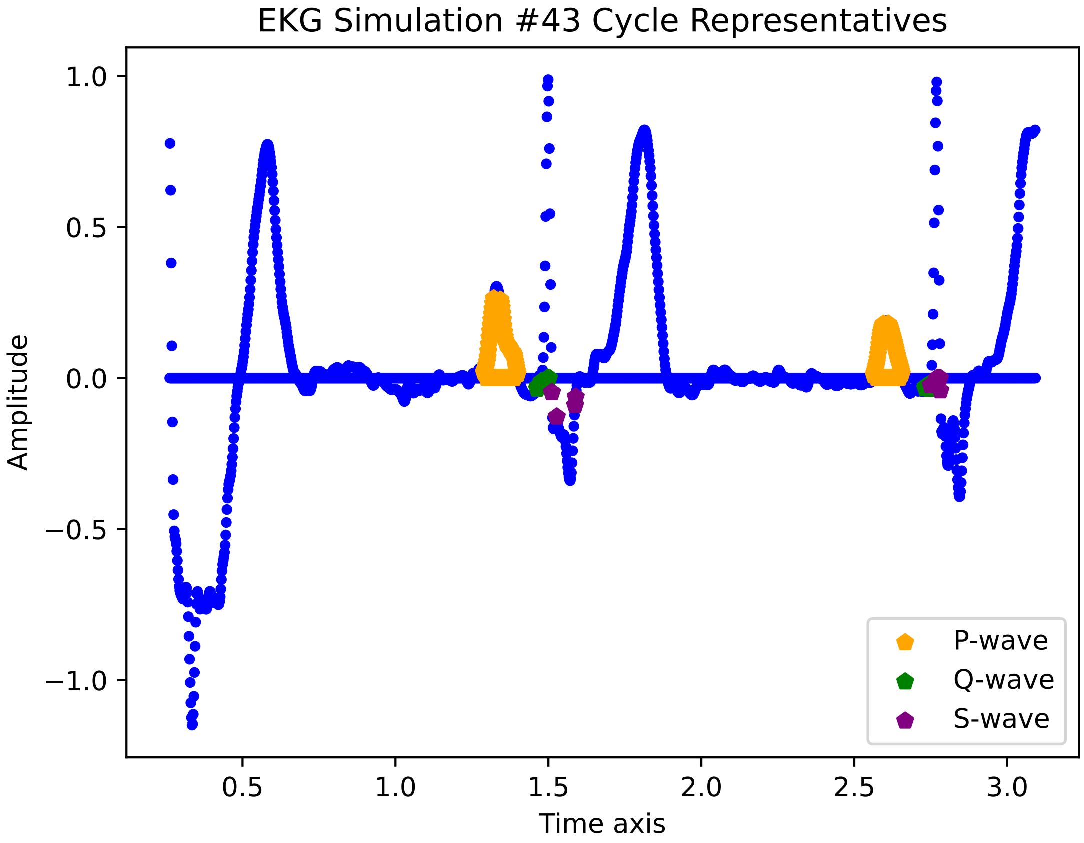
EKG Simulation #43 Cycle Representatives
1.0
0.5
0.0
−0.5
−1.0
Amplitude
0.5
1.0
1.5
2.0
2.5
3.0
Time axis
P-wave
Q-wave
S-wave

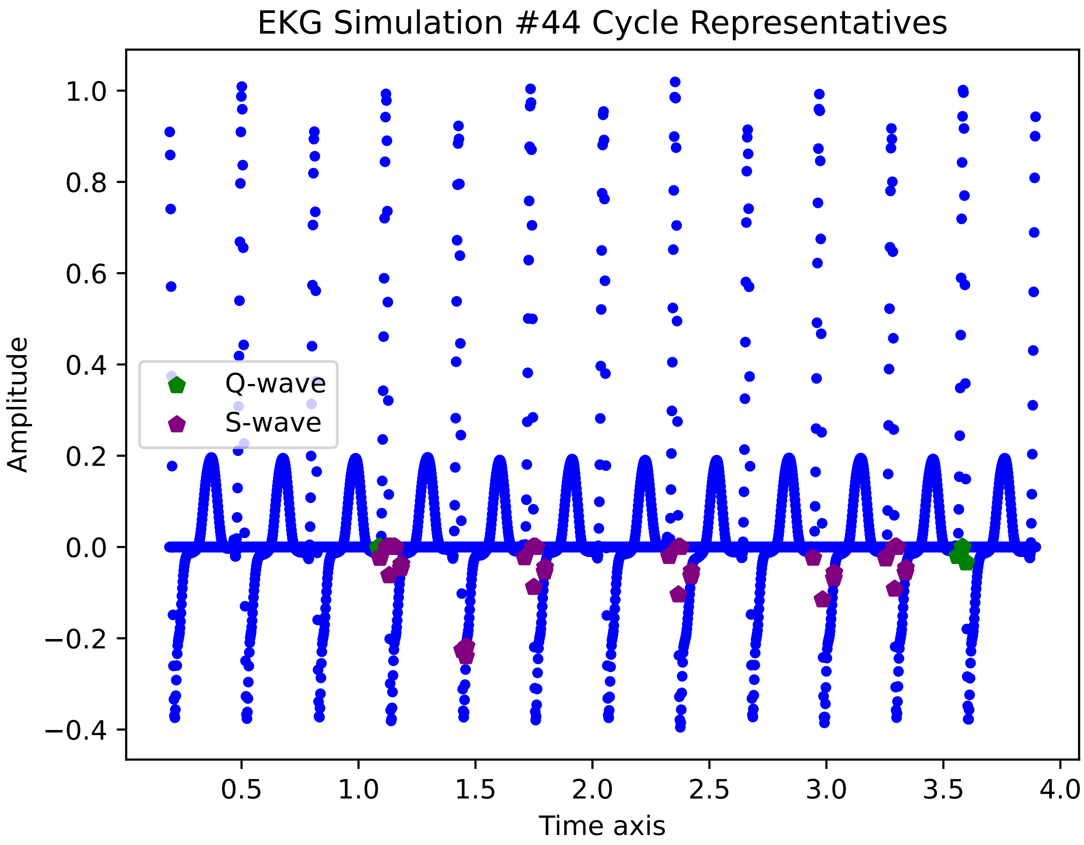
EKG Simulation #44 Cycle Representatives
Amplitude
Time axis
Q-wave
S-wave
1.0
0.8
0.6
0.4
0.2
0.0
−0.2
−0.4
0.5
1.0
1.5
2.0
2.5
3.0
3.5
4.0

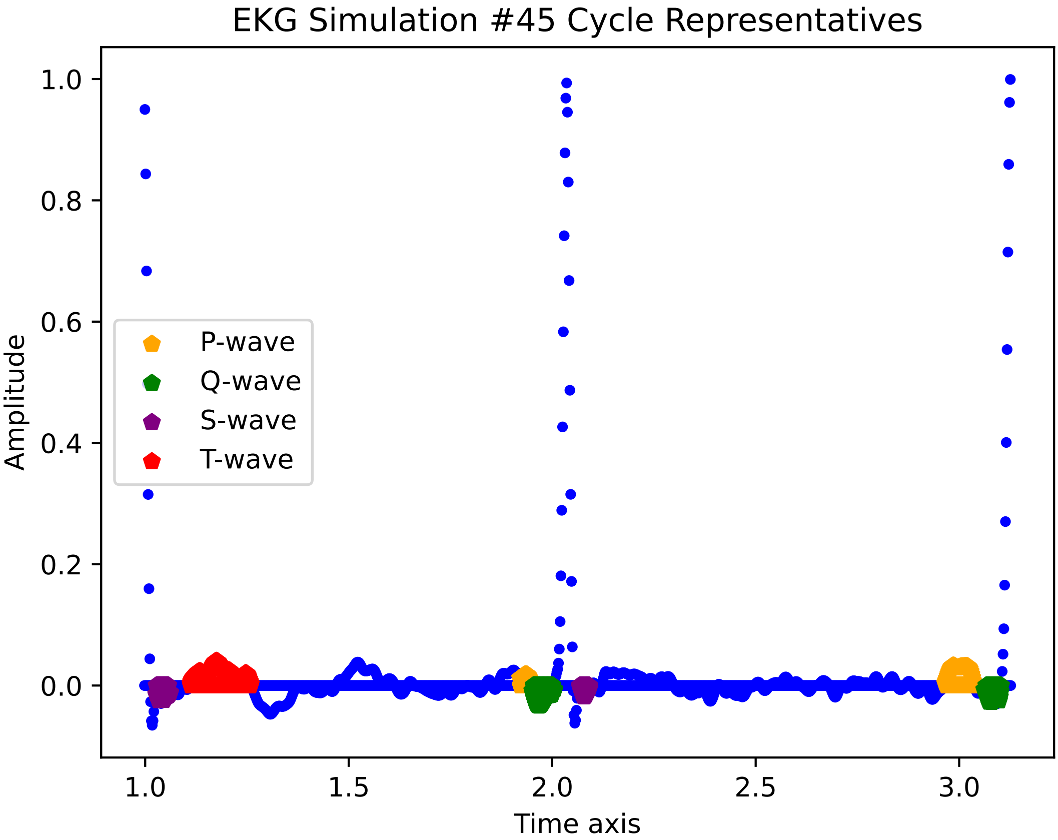
EKG Simulation #45 Cycle Representatives
Amplitude
Time axis
P-wave
Q-wave
S-wave
T-wave
1.0
0.8
0.6
0.4
0.2
0.0
1.0
1.5
2.0
2.5
3.0

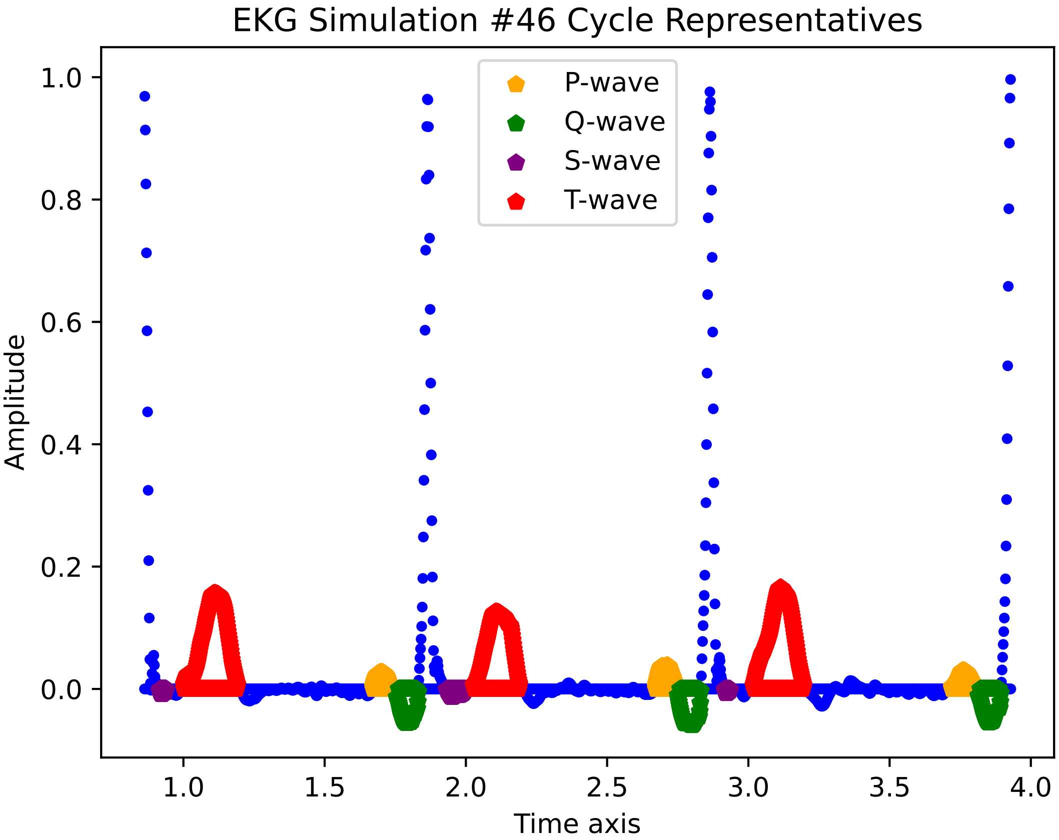

EKG Simulation #46 Cycle Representatives
P-wave
Q-wave
S-wave
T-wave
Amplitude
Time axis
1.0
0.8
0.6
0.4
0.2
0.0
1.0
1.5
2.0
2.5
3.0
3.5
4.0

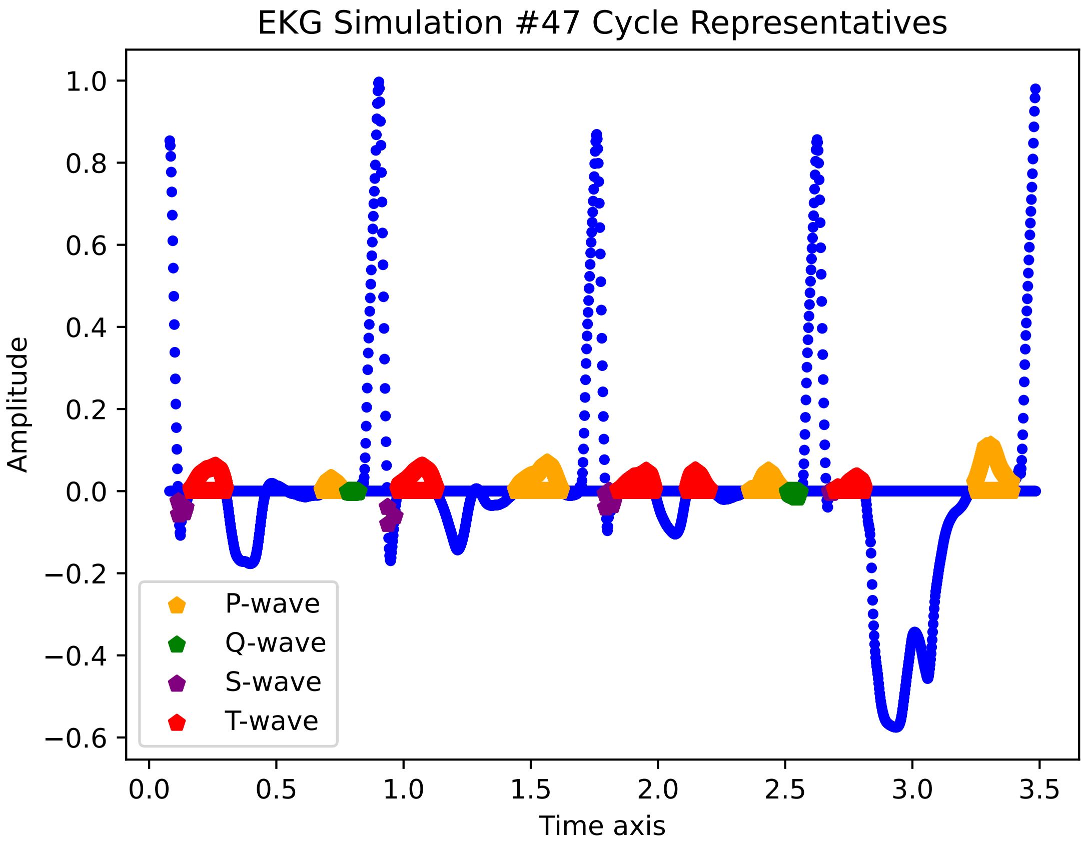
EKG Simulation #47 Cycle Representatives
Amplitude
Time axis
1.0
0.8
0.6
0.4
0.2
0.0
−0.2
−0.4
−0.6
0.0
0.5
1.0
1.5
2.0
2.5
3.0
3.5
P-wave
Q-wave
S-wave
T-wave

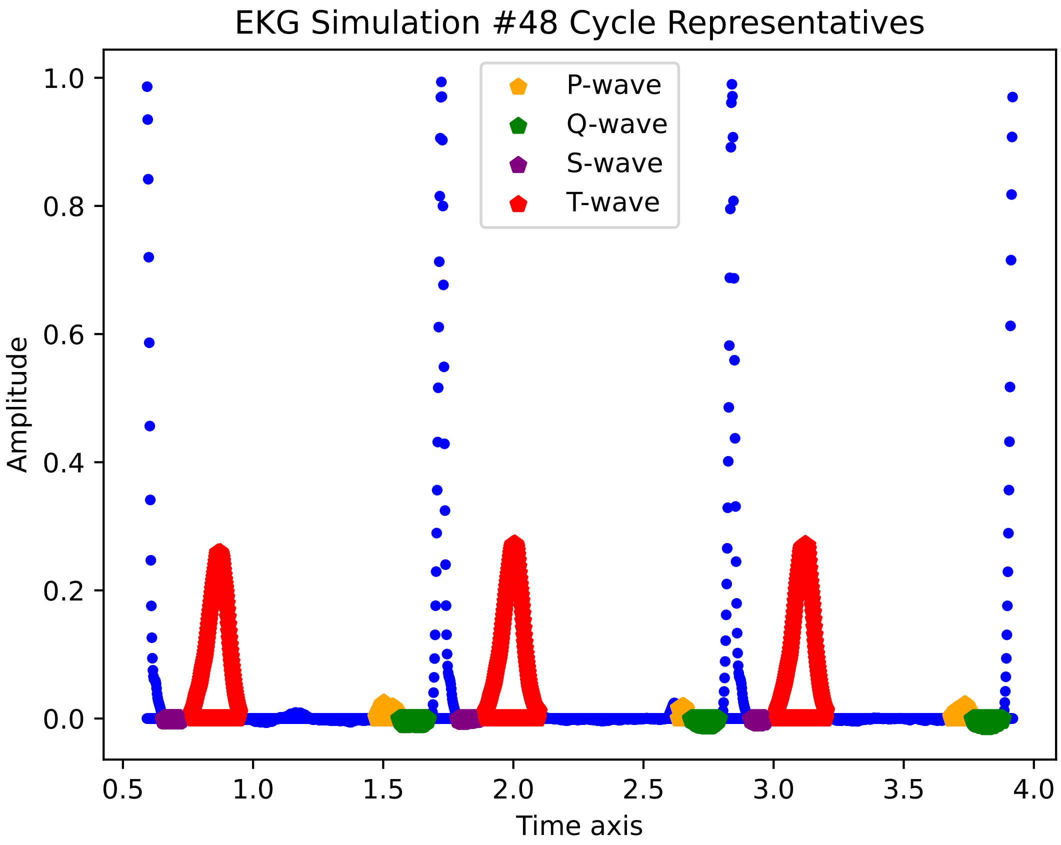
EKG Simulation #48 Cycle Representatives
P-wave
Q-wave
S-wave
T-wave
Amplitude
Time axis
0.0
0.2
0.4
0.6
0.8
1.0
0.5
1.0
1.5
2.0
2.5
3.0
3.5
4.0

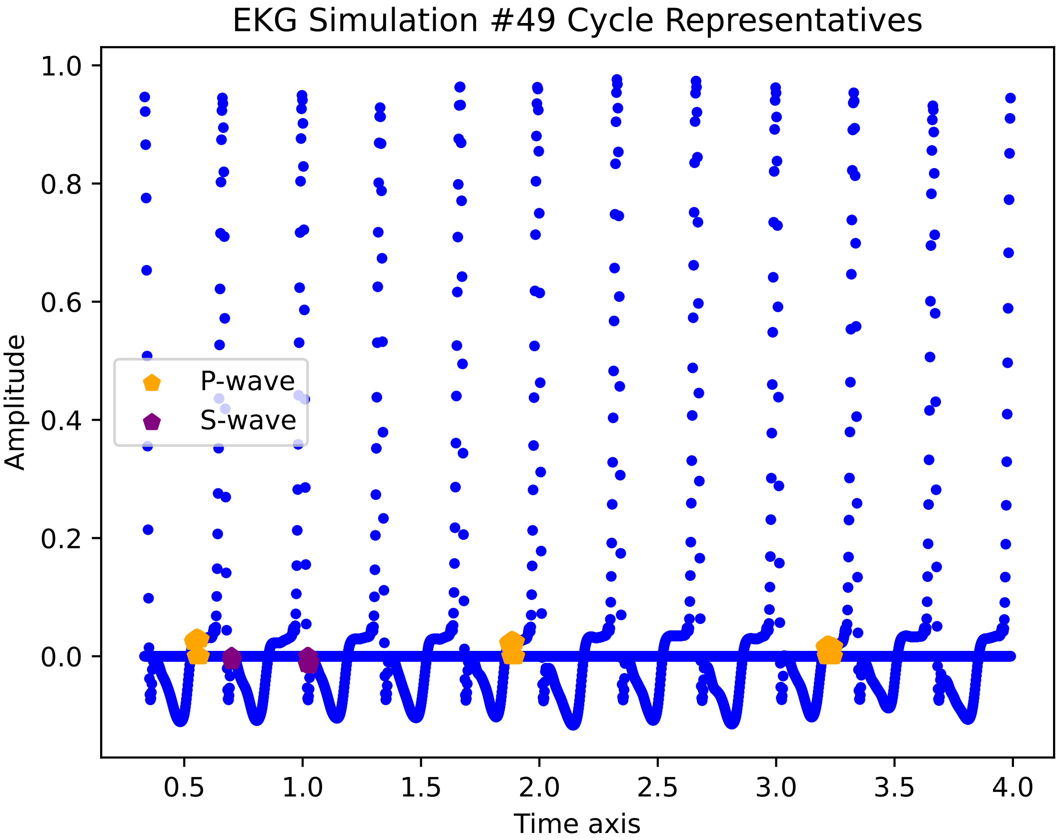

EKG Simulation #49 Cycle Representatives
Amplitude
Time axis
P-wave
S-wave
1.0
0.8
0.6
0.4
0.2
0.0
0.5
1.0
1.5
2.0
2.5
3.0
3.5
4.0

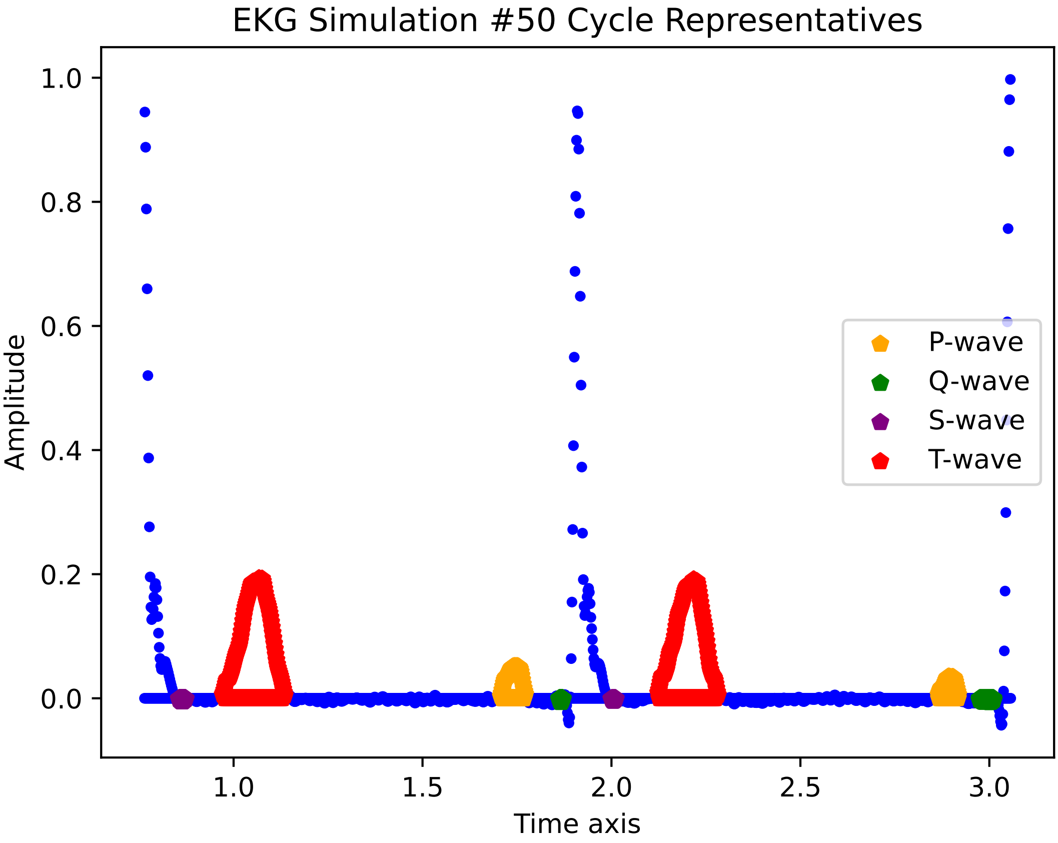

EKG Simulation #50 Cycle Representatives
Amplitude
Time axis
1.0
0.8
0.6
0.4
0.2
0.0
1.0
1.5
2.0
2.5
3.0
P-wave
Q-wave
S-wave
T-wave

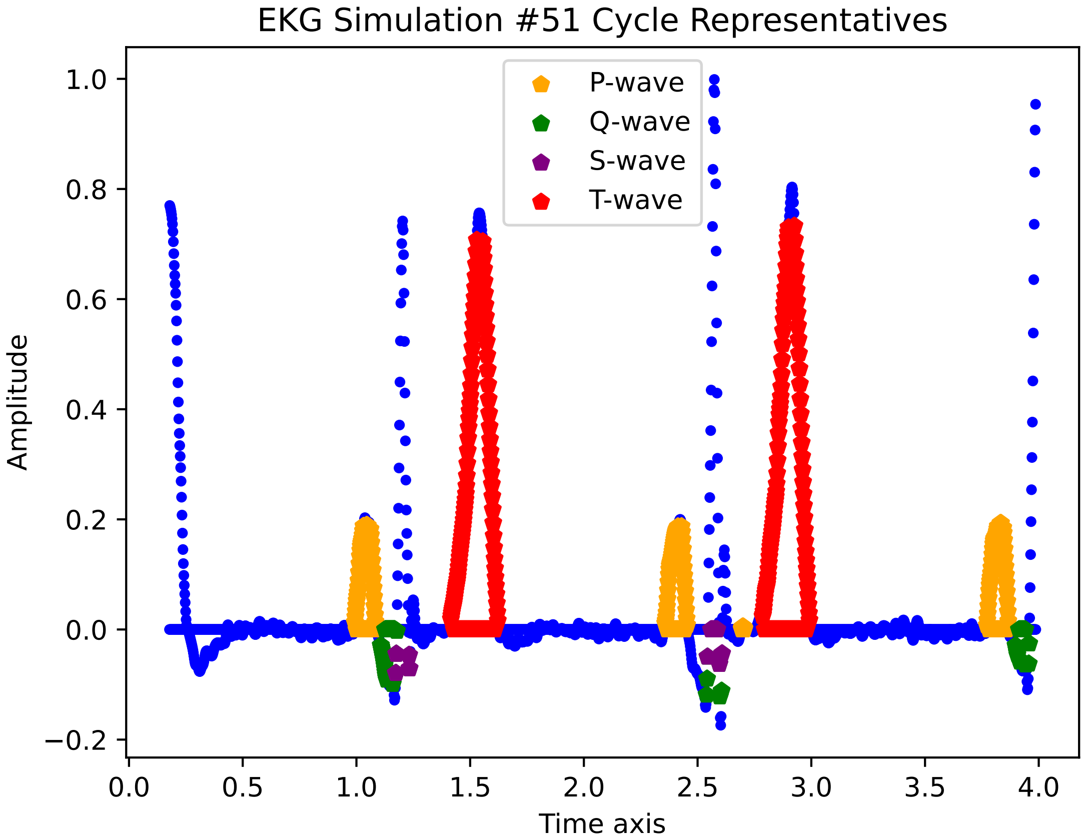
EKG Simulation #51 Cycle Representatives
P-wave
Q-wave
S-wave
T-wave
1.0
0.8
0.6
0.4
0.2
0.0
−0.2
Amplitude
0.0
0.5
1.0
1.5
2.0
2.5
3.0
3.5
4.0
Time axis

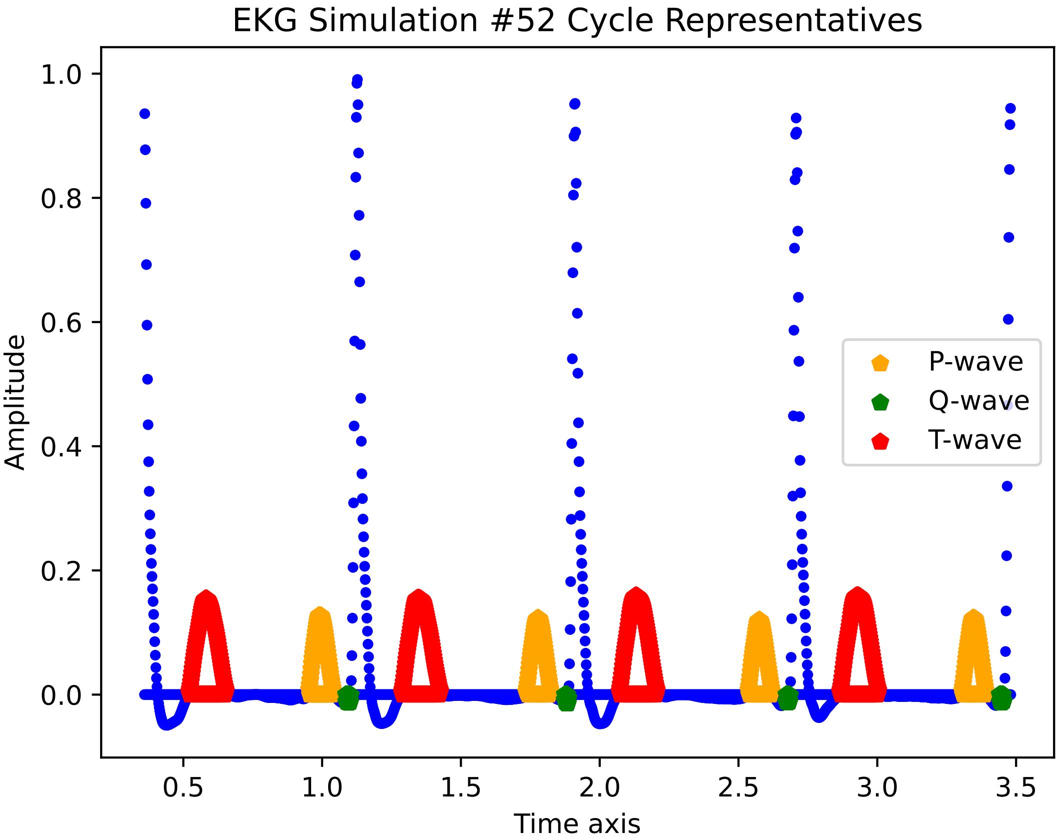

EKG Simulation #52 Cycle Representatives
Amplitude
1.0
0.8
0.6
0.4
0.2
0.0
0.5
1.0
1.5
2.0
2.5
3.0
3.5
Time axis
P-wave
Q-wave
T-wave

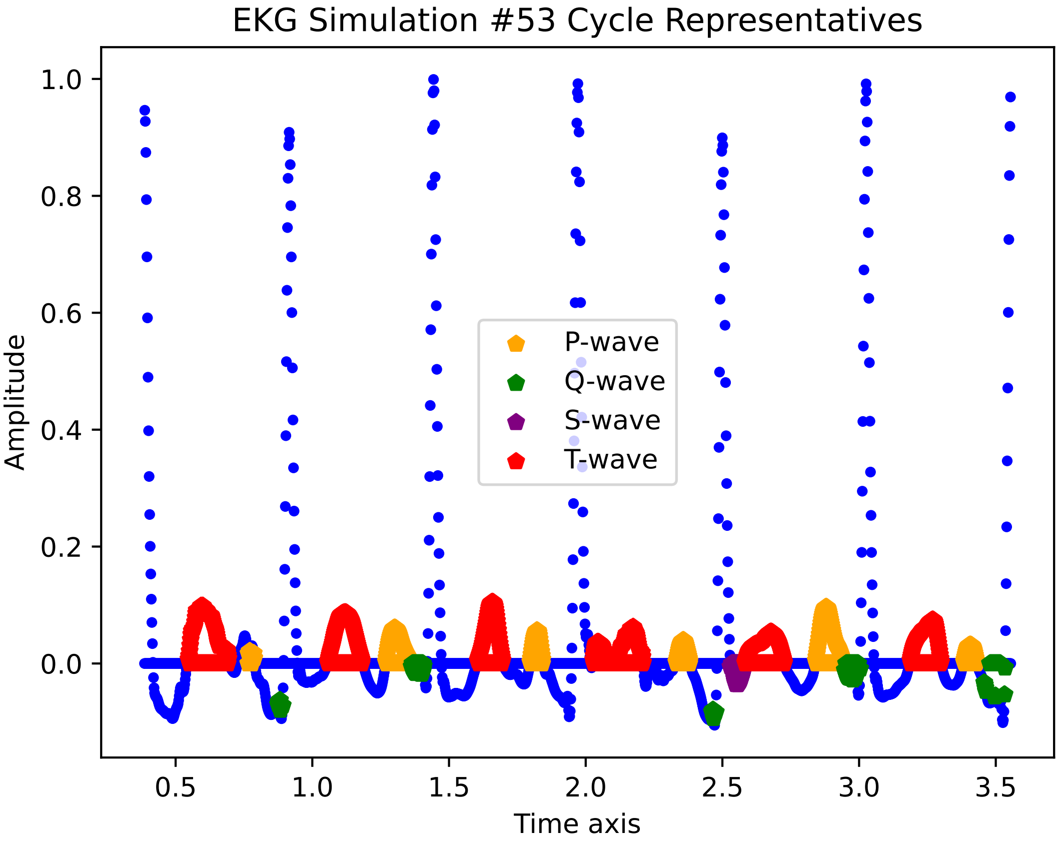
EKG Simulation #53 Cycle Representatives
Amplitude
Time axis
P-wave
Q-wave
S-wave
T-wave
1.0
0.8
0.6
0.4
0.2
0.0
0.5
1.0
1.5
2.0
2.5
3.0
3.5

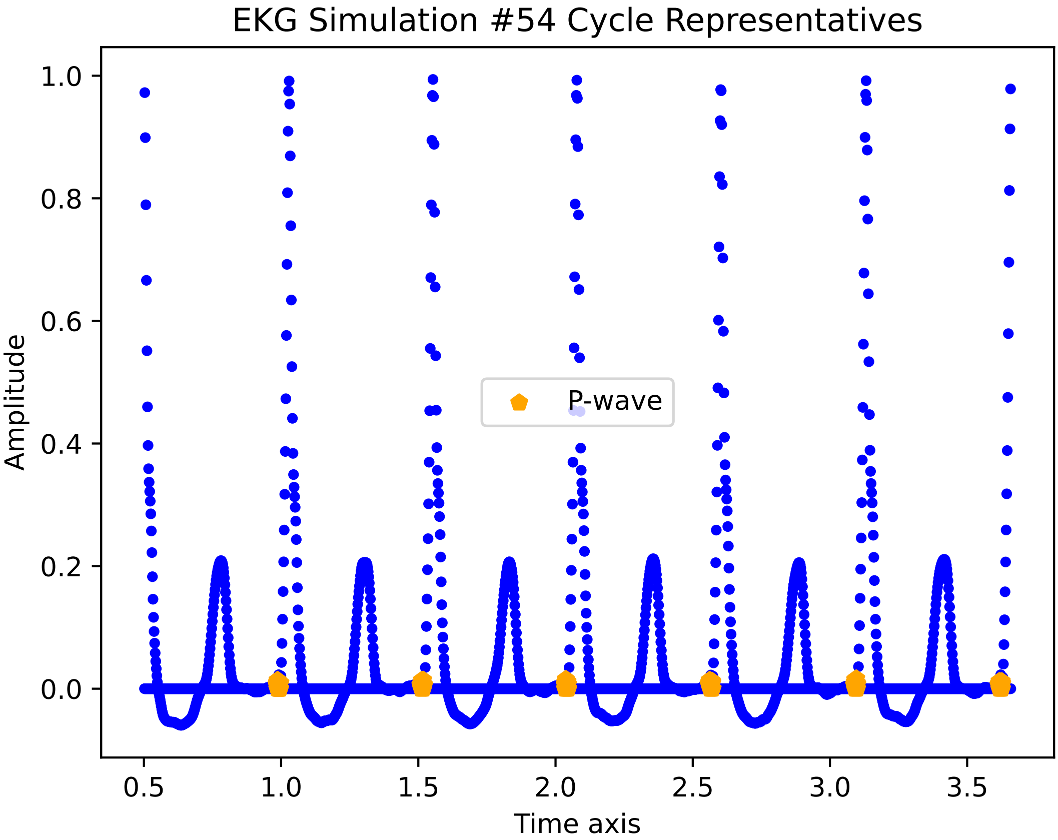
EKG Simulation #54 Cycle Representatives
P-wave
Amplitude
Time axis
1.0
0.8
0.6
0.4
0.2
0.0
0.5
1.0
1.5
2.0
2.5
3.0
3.5

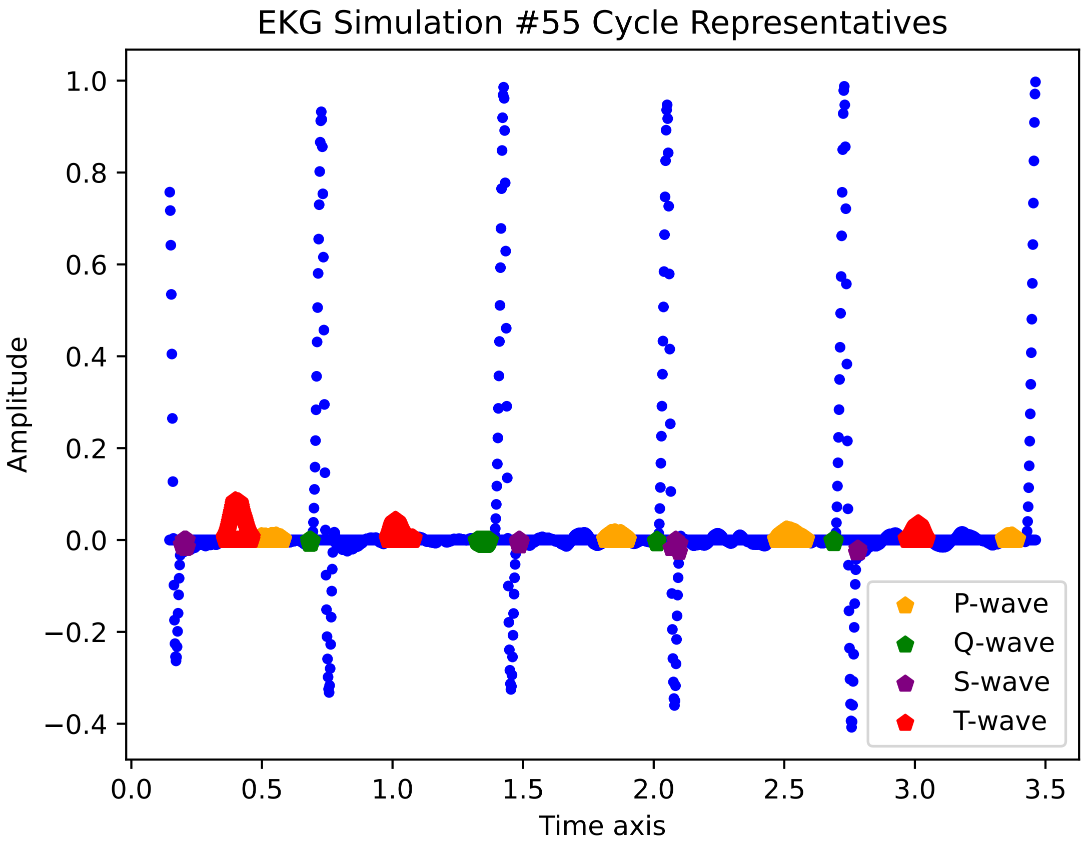
EKG Simulation #55 Cycle Representatives
Amplitude
Time axis
P-wave
Q-wave
S-wave
T-wave
1.0
0.8
0.6
0.4
0.2
0.0
−0.2
−0.4
0.0
0.5
1.0
1.5
2.0
2.5
3.0
3.5

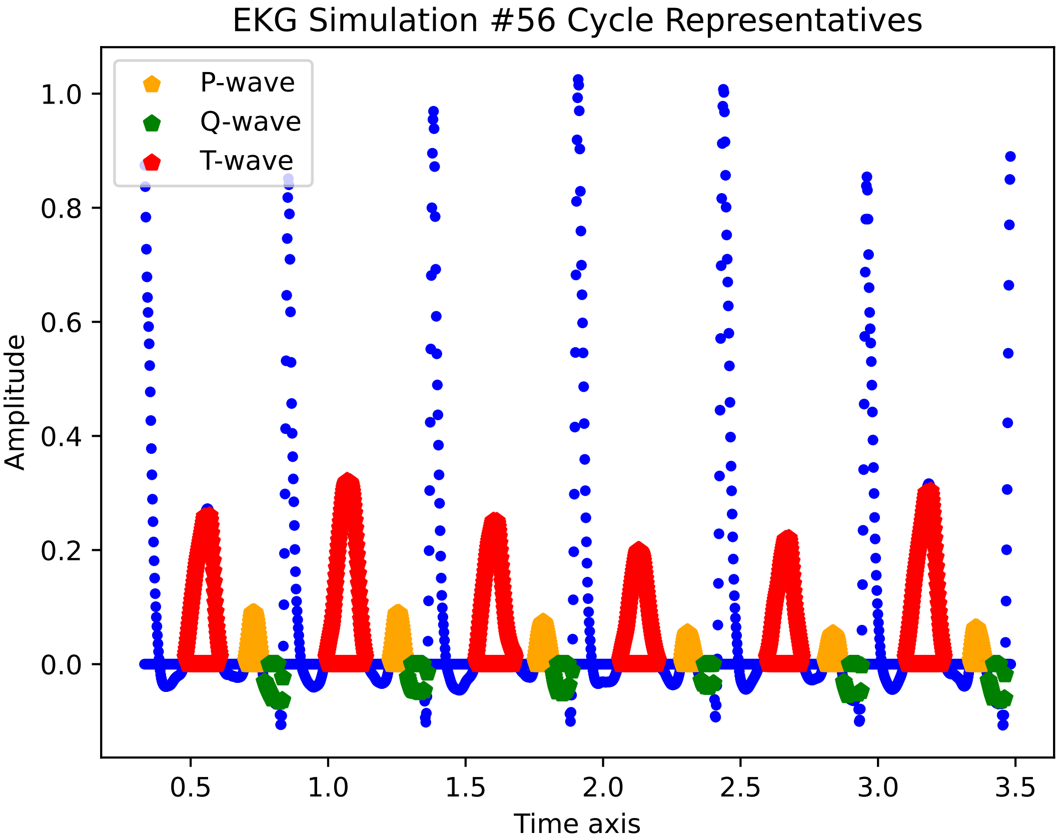

EKG Simulation #56 Cycle Representatives
P-wave
Q-wave
T-wave
Amplitude
Time axis
1.0
0.8
0.6
0.4
0.2
0.0
0.5
1.0
1.5
2.0
2.5
3.0
3.5

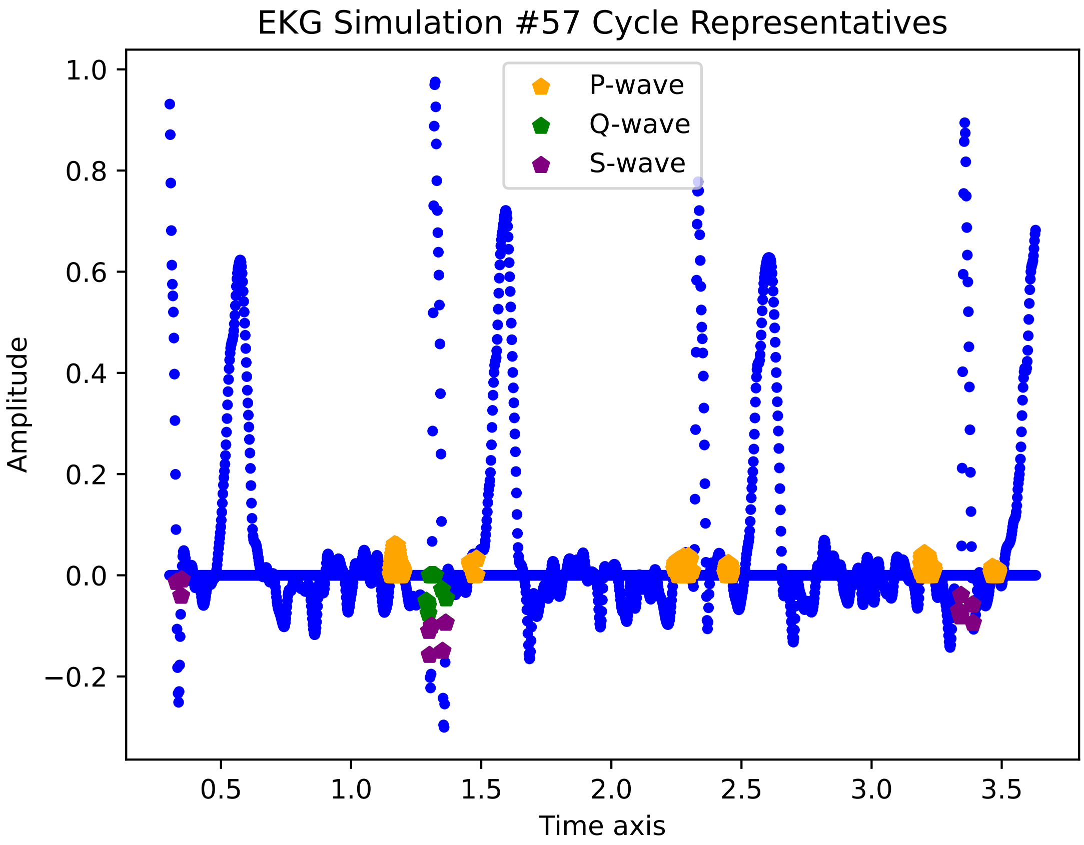
EKG Simulation #57 Cycle Representatives
P-wave
Q-wave
S-wave
Amplitude
Time axis
1.0
0.8
0.6
0.4
0.2
0.0
−0.2
0.5
1.0
1.5
2.0
2.5
3.0
3.5

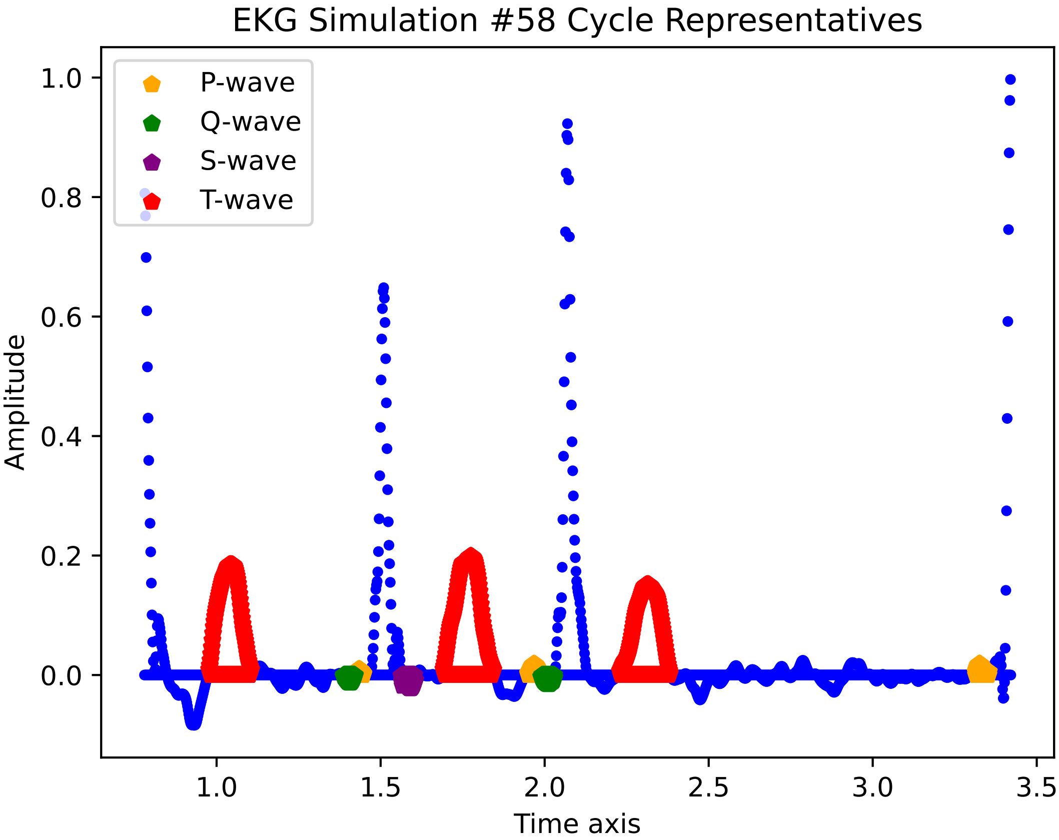

EKG Simulation #58 Cycle Representatives
P-wave
Q-wave
S-wave
T-wave
Amplitude
Time axis
1.0
0.8
0.6
0.4
0.2
0.0
1.0
1.5
2.0
2.5
3.0
3.5

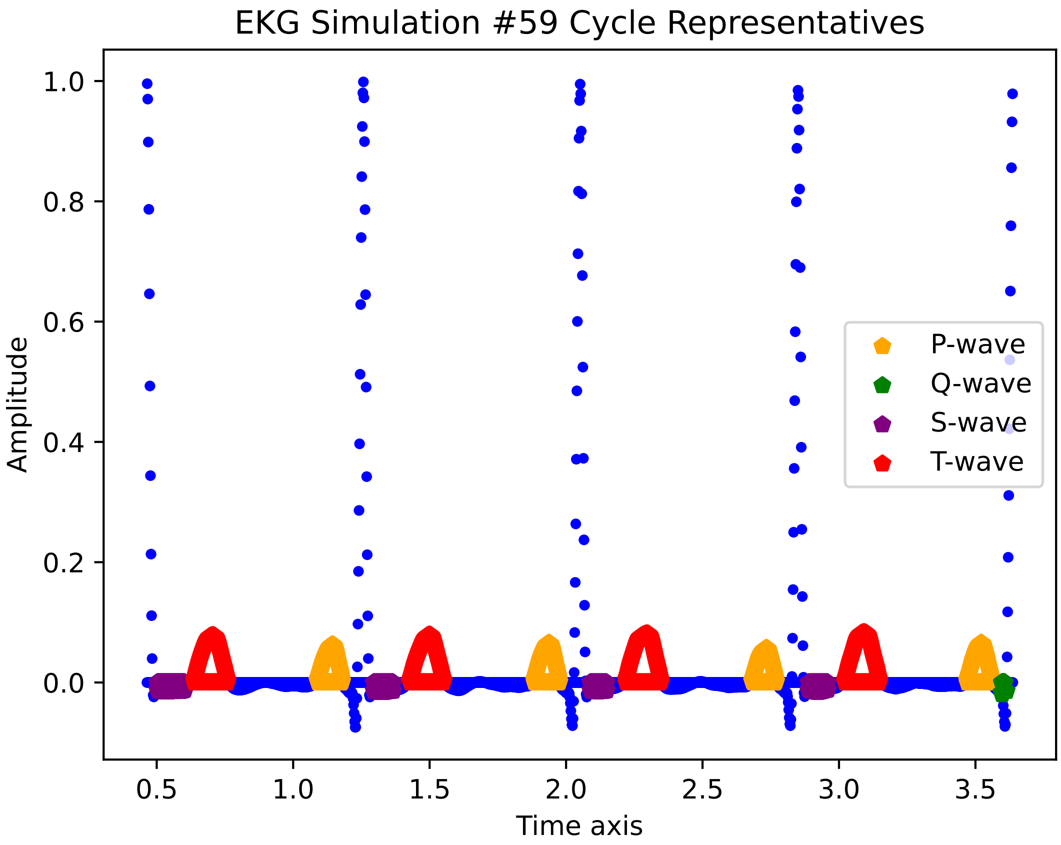
EKG Simulation #59 Cycle Representatives
Amplitude
Time axis
1.0
0.8
0.6
0.4
0.2
0.0
0.5
1.0
1.5
2.0
2.5
3.0
3.5
P-wave
Q-wave
S-wave
T-wave

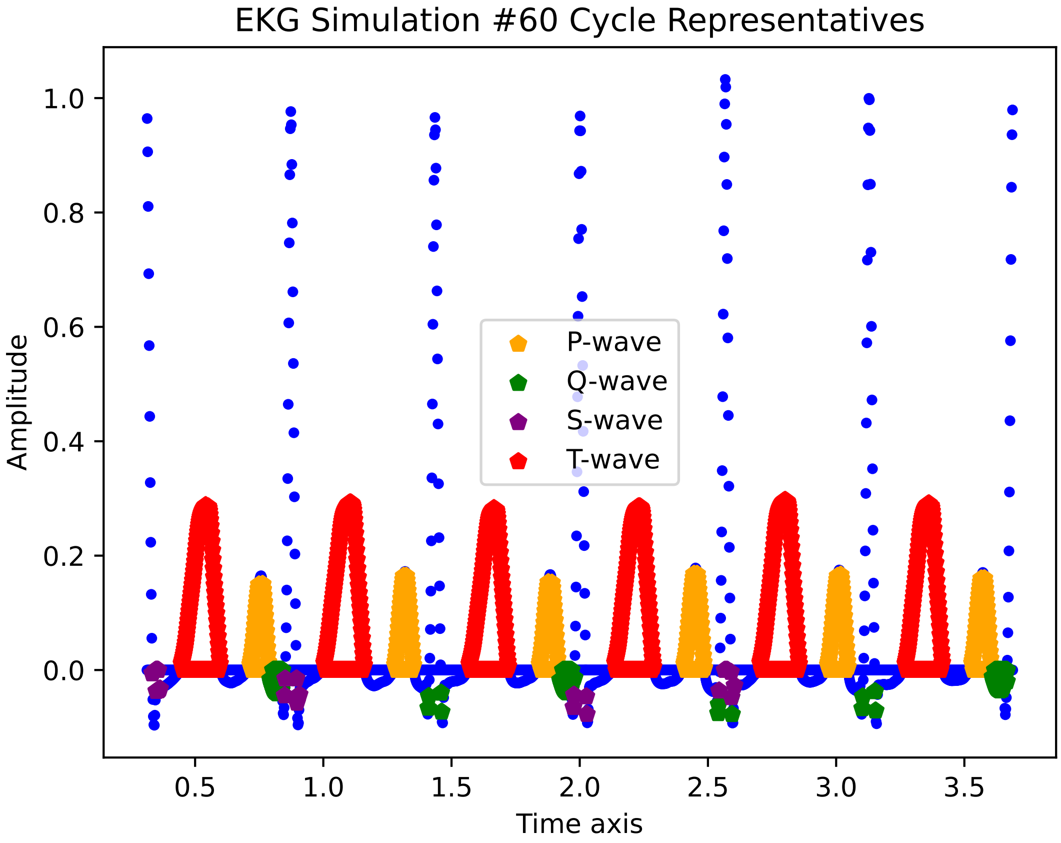
EKG Simulation #60 Cycle Representatives
Amplitude
Time axis
P-wave
Q-wave
S-wave
T-wave
1.0
0.8
0.6
0.4
0.2
0.0
0.5
1.0
1.5
2.0
2.5
3.0
3.5

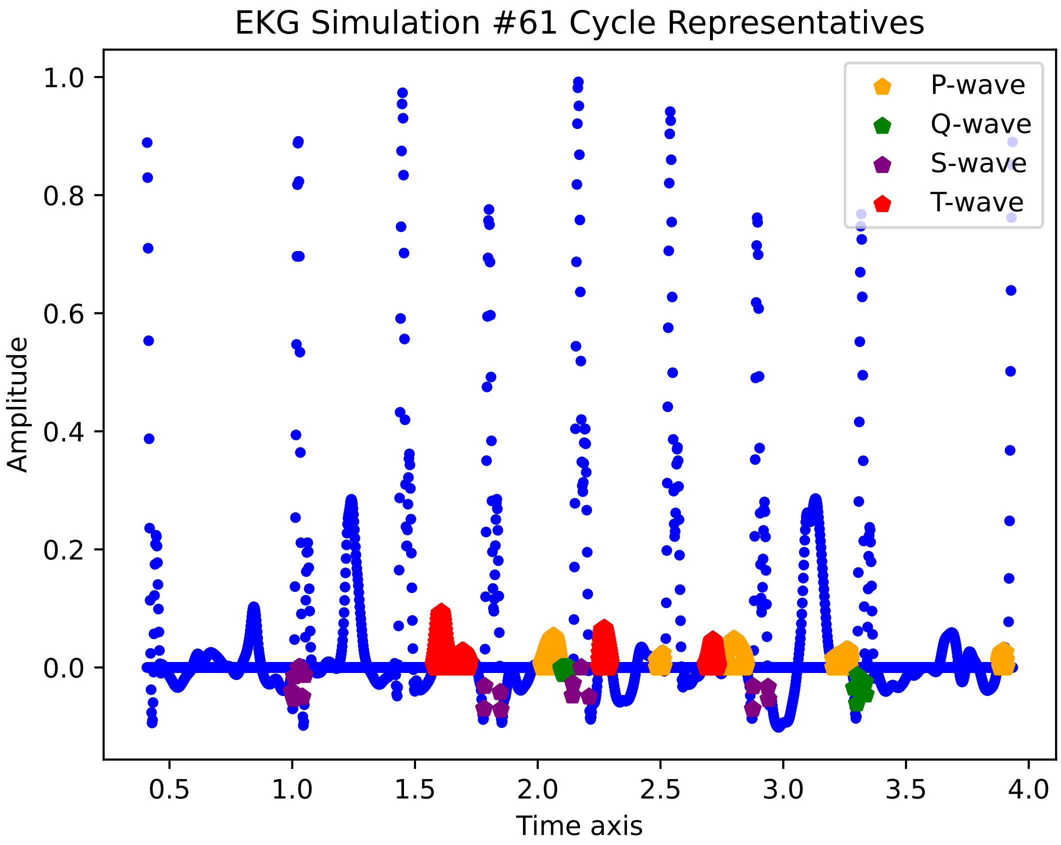

EKG Simulation #61 Cycle Representatives
Amplitude
Time axis
P-wave
Q-wave
S-wave
T-wave
1.0
0.8
0.6
0.4
0.2
0.0
0.5
1.0
1.5
2.0
2.5
3.0
3.5
4.0

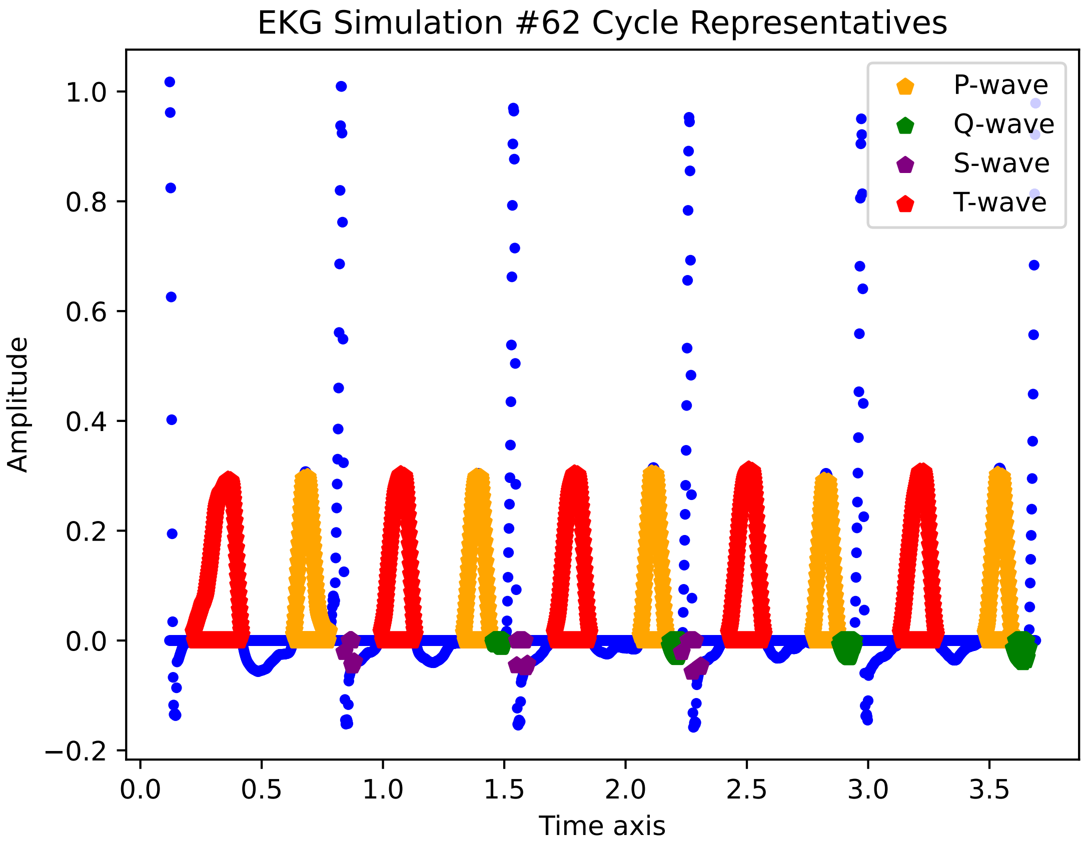
EKG Simulation #62 Cycle Representatives
P-wave
Q-wave
S-wave
T-wave
Amplitude
Time axis
1.0
0.8
0.6
0.4
0.2
0.0
−0.2
0.0
0.5
1.0
1.5
2.0
2.5
3.0
3.5

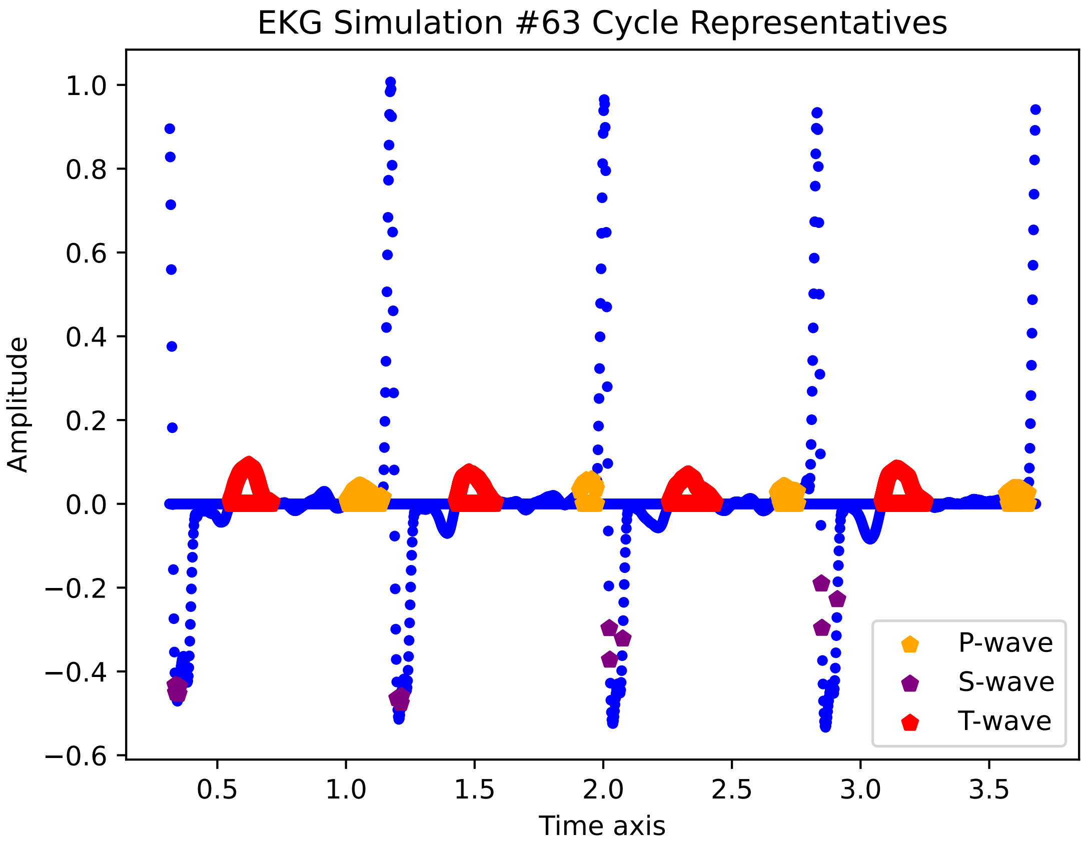
EKG Simulation #63 Cycle Representatives
Amplitude
Time axis
1.0
0.8
0.6
0.4
0.2
0.0
−0.2
−0.4
−0.6
0.5
1.0
1.5
2.0
2.5
3.0
3.5
P-wave
S-wave
T-wave

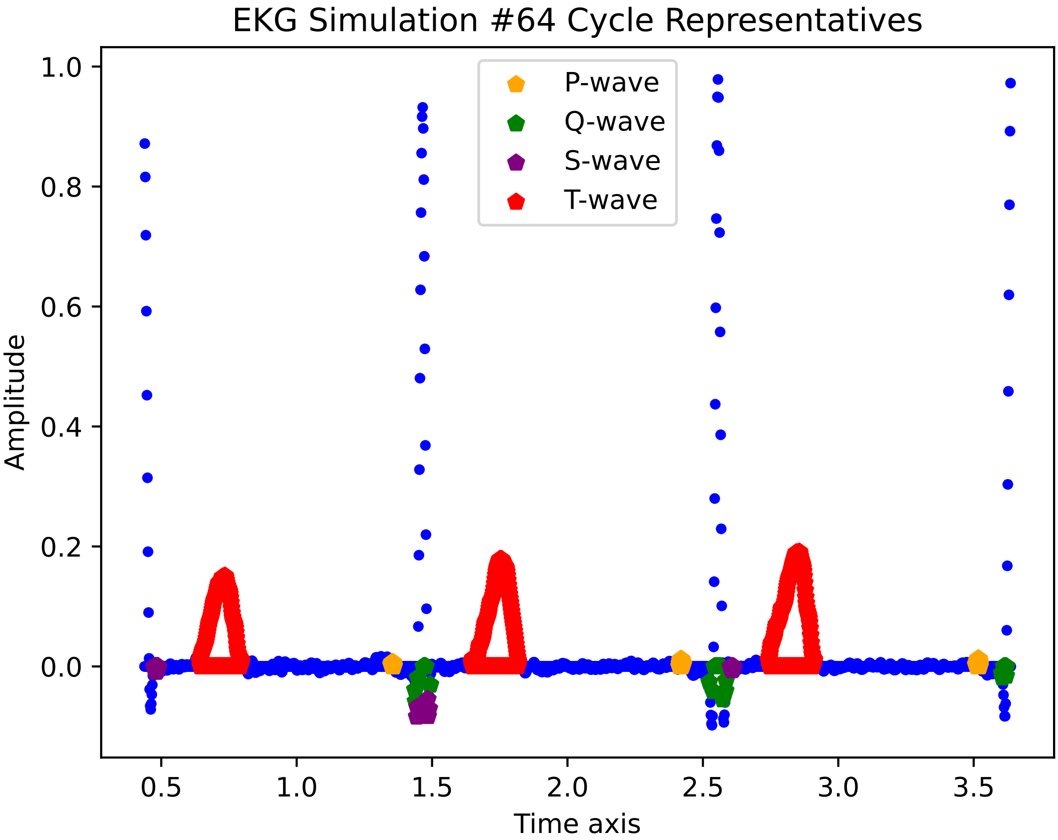

EKG Simulation #64 Cycle Representatives
P-wave
Q-wave
S-wave
T-wave
Amplitude
Time axis
1.0
0.8
0.6
0.4
0.2
0.0
0.5
1.0
1.5
2.0
2.5
3.0
3.5

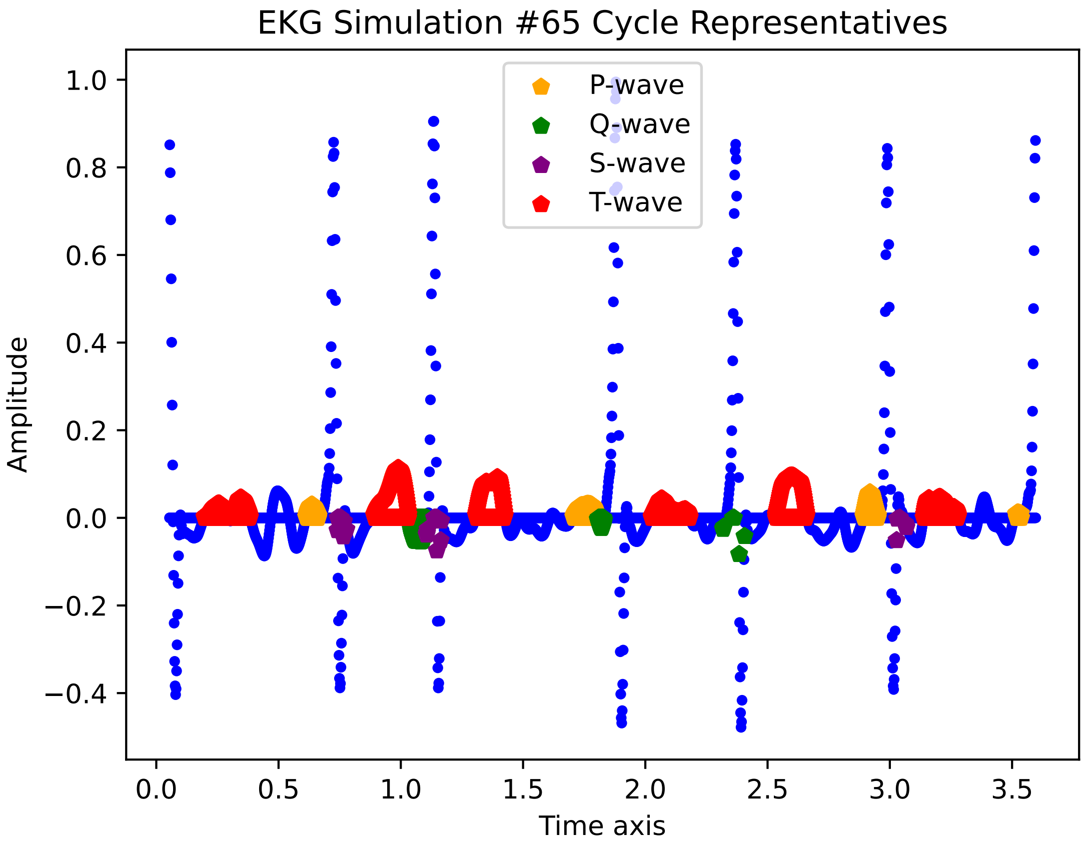

EKG Simulation #65 Cycle Representatives
P-wave
Q-wave
S-wave
T-wave
Amplitude
Time axis
1.0
0.8
0.6
0.4
0.2
0.0
−0.2
−0.4
0.0
0.5
1.0
1.5
2.0
2.5
3.0
3.5

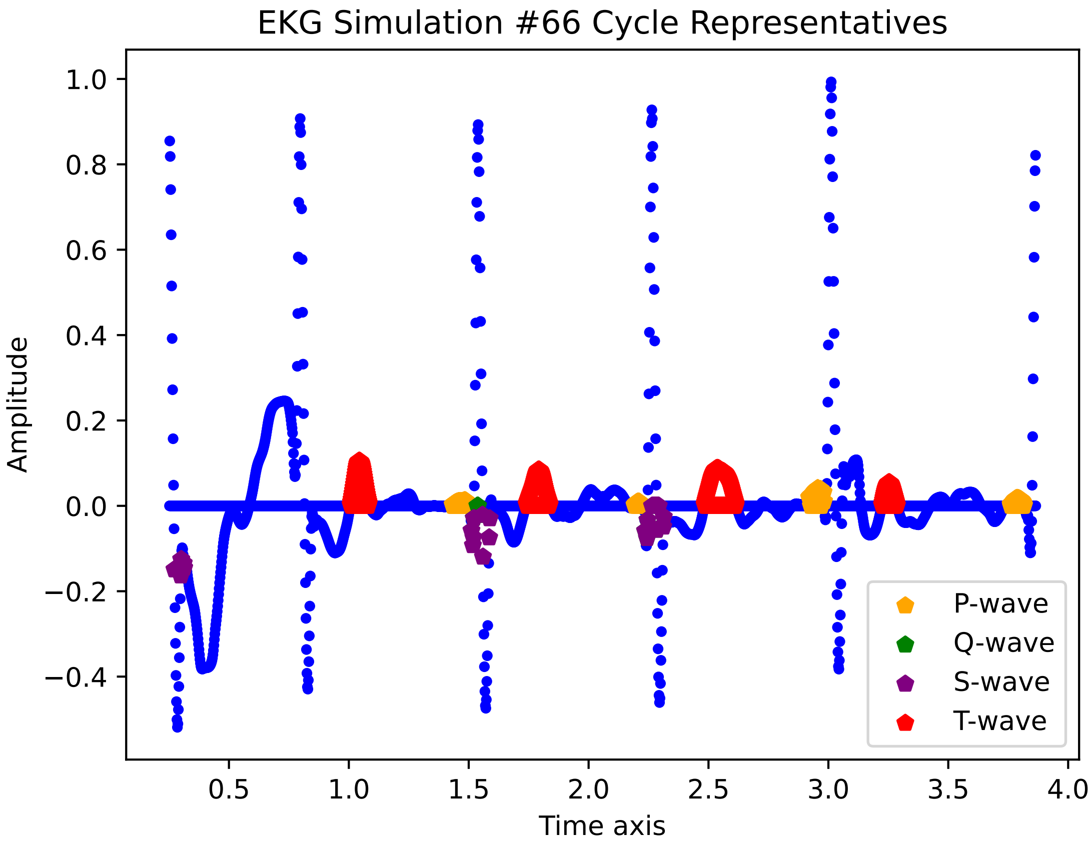

EKG Simulation #66 Cycle Representatives
Amplitude
Time axis
1.0
0.8
0.6
0.4
0.2
0.0
−0.2
−0.4
0.5
1.0
1.5
2.0
2.5
3.0
3.5
4.0
P-wave
Q-wave
S-wave
T-wave

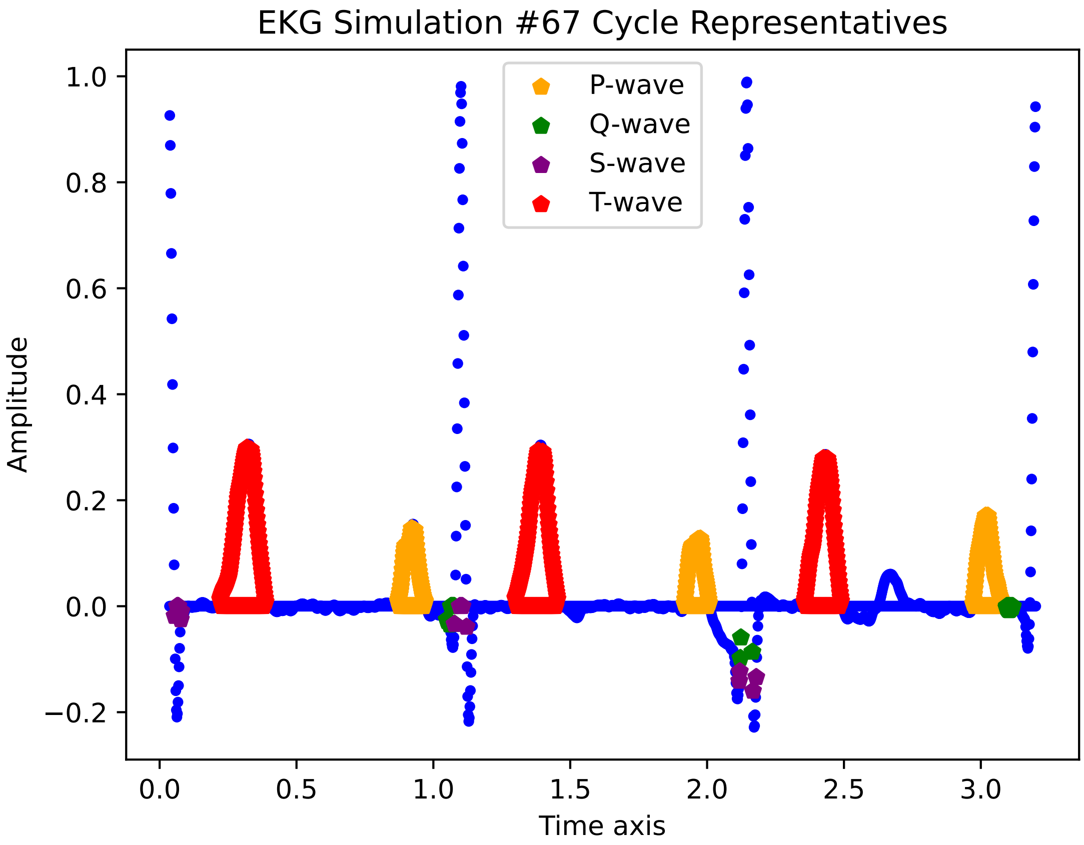

EKG Simulation #67 Cycle Representatives
P-wave
Q-wave
S-wave
T-wave
Amplitude
Time axis
1.0
0.8
0.6
0.4
0.2
0.0
−0.2
0.0
0.5
1.0
1.5
2.0
2.5
3.0

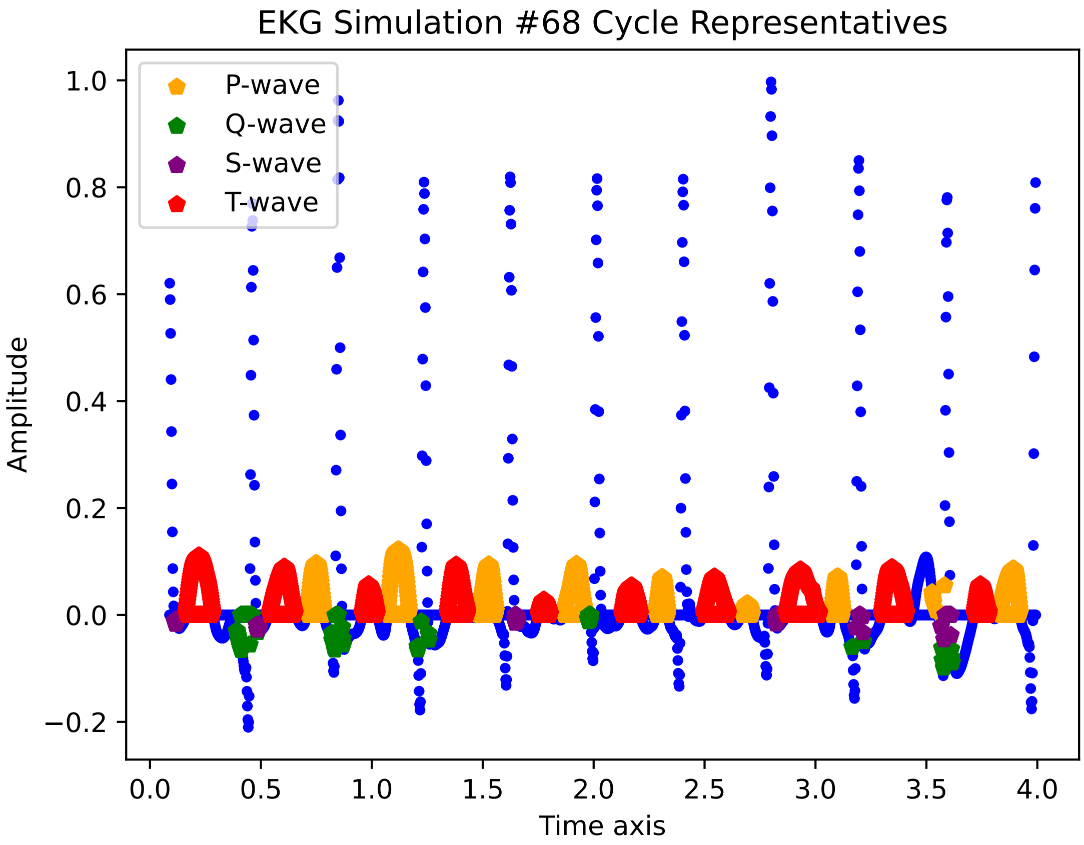

EKG Simulation #68 Cycle Representatives
P-wave
Q-wave
S-wave
T-wave
Amplitude
Time axis
1.0
0.8
0.6
0.4
0.2
0.0
−0.2
0.0
0.5
1.0
1.5
2.0
2.5
3.0
3.5
4.0

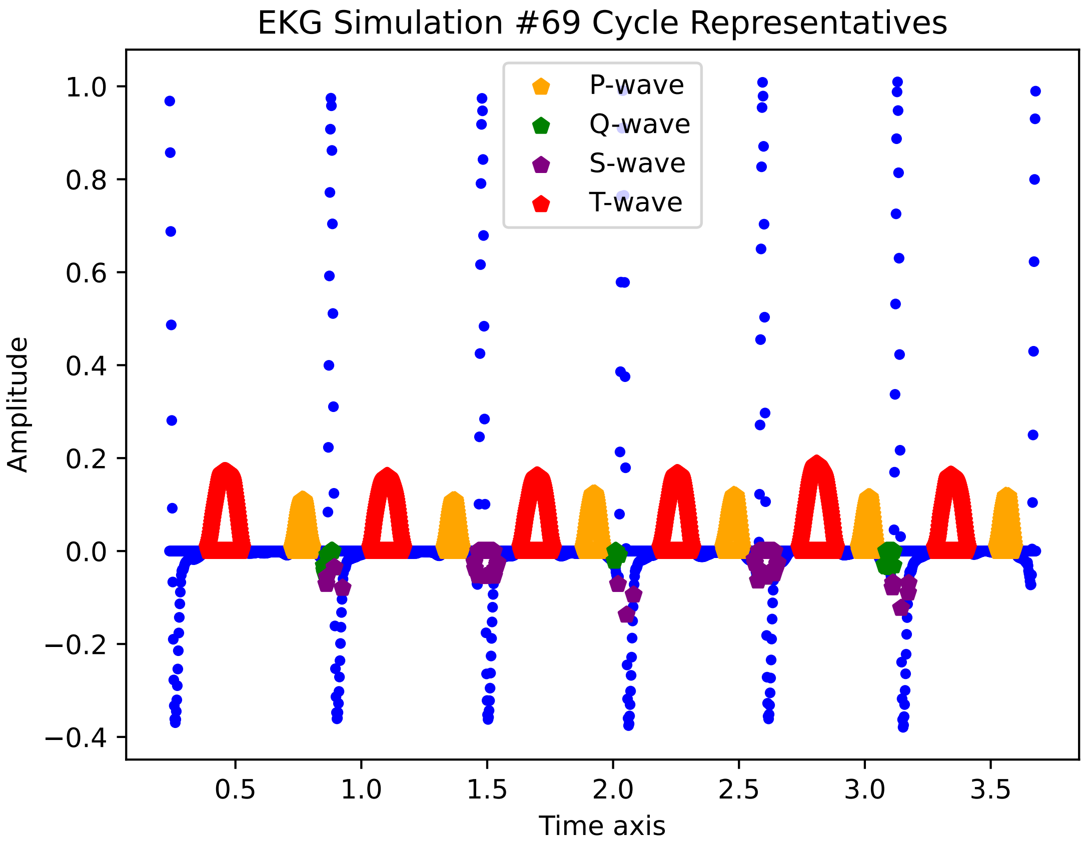

EKG Simulation #69 Cycle Representatives
P-wave
Q-wave
S-wave
T-wave
Amplitude
Time axis
1.0
0.8
0.6
0.4
0.2
0.0
−0.2
−0.4
0.5
1.0
1.5
2.0
2.5
3.0
3.5

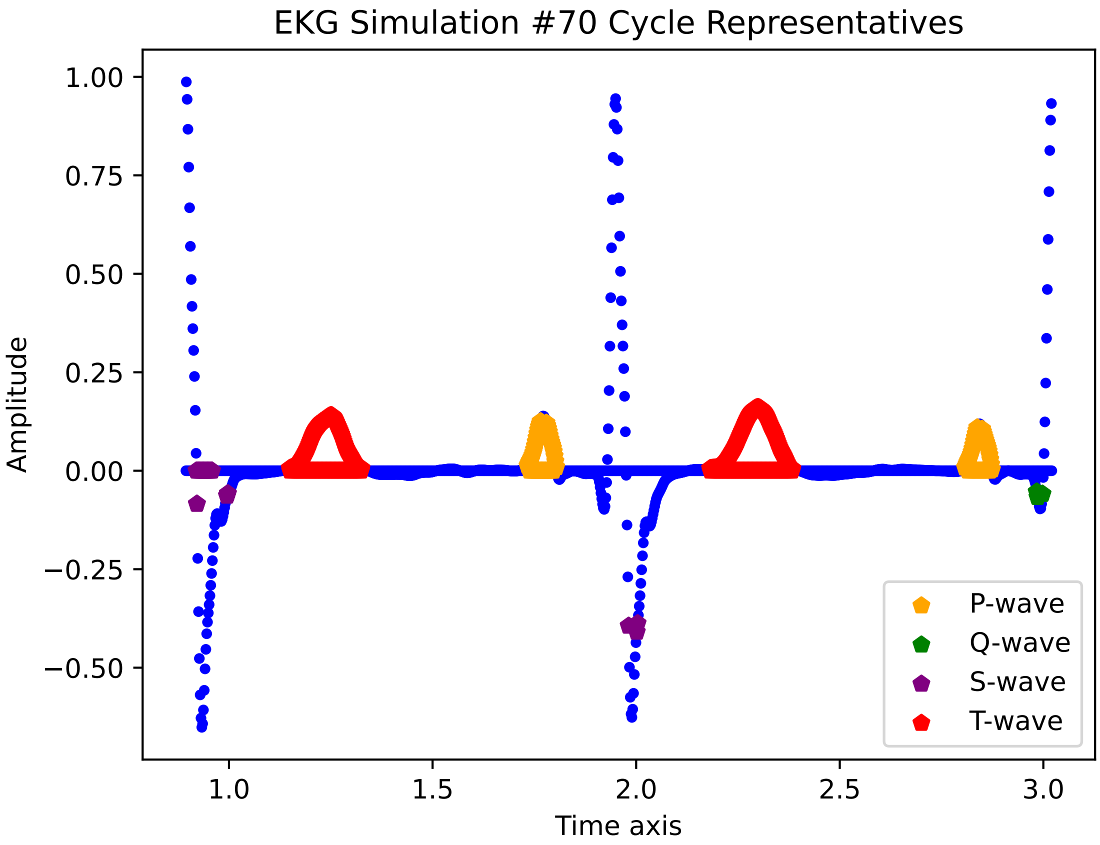
EKG Simulation #70 Cycle Representatives
Amplitude
1.00
0.75
0.50
0.25
0.00
−0.25
−0.50
1.0
1.5
2.0
2.5
3.0
Time axis
P-wave
Q-wave
S-wave
T-wave

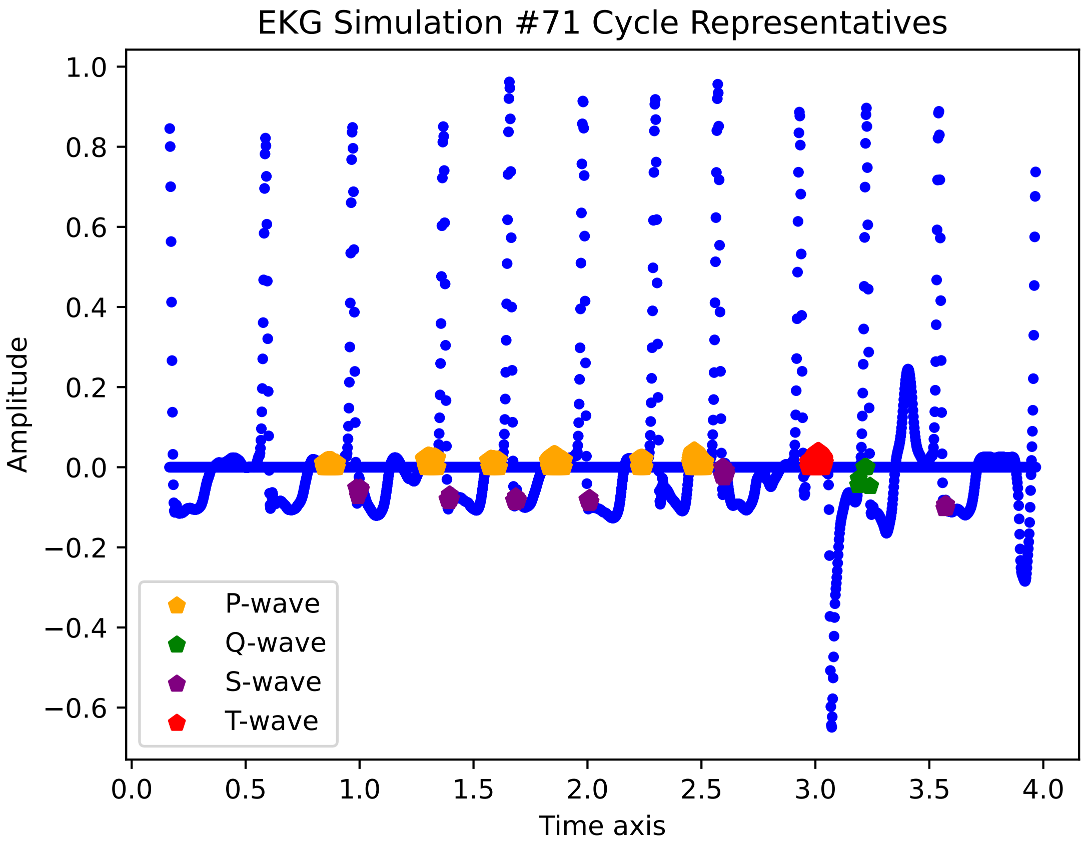

EKG Simulation #71 Cycle Representatives
Amplitude
Time axis
P-wave
Q-wave
S-wave
T-wave
1.0
0.8
0.6
0.4
0.2
0.0
−0.2
−0.4
−0.6
0.0
0.5
1.0
1.5
2.0
2.5
3.0
3.5
4.0

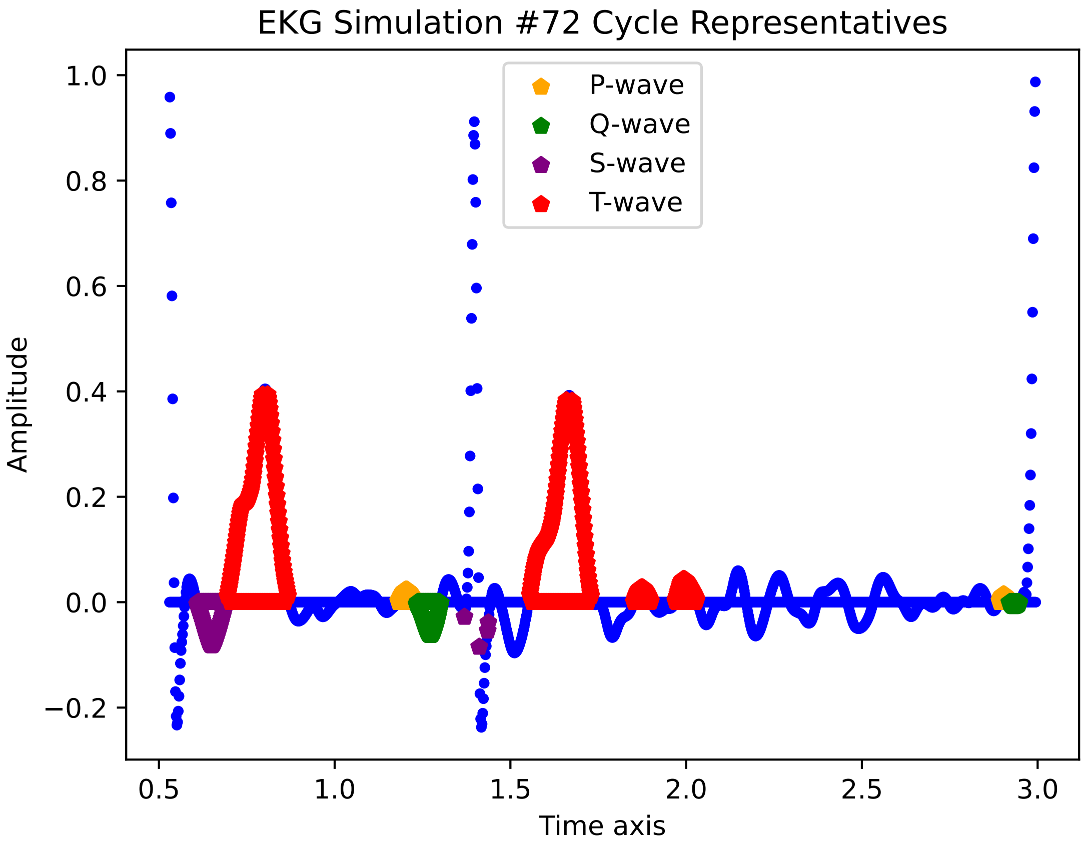

EKG Simulation #72 Cycle Representatives
P-wave
Q-wave
S-wave
T-wave
Amplitude
Time axis
1.0
0.8
0.6
0.4
0.2
0.0
−0.2
0.5
1.0
1.5
2.0
2.5
3.0

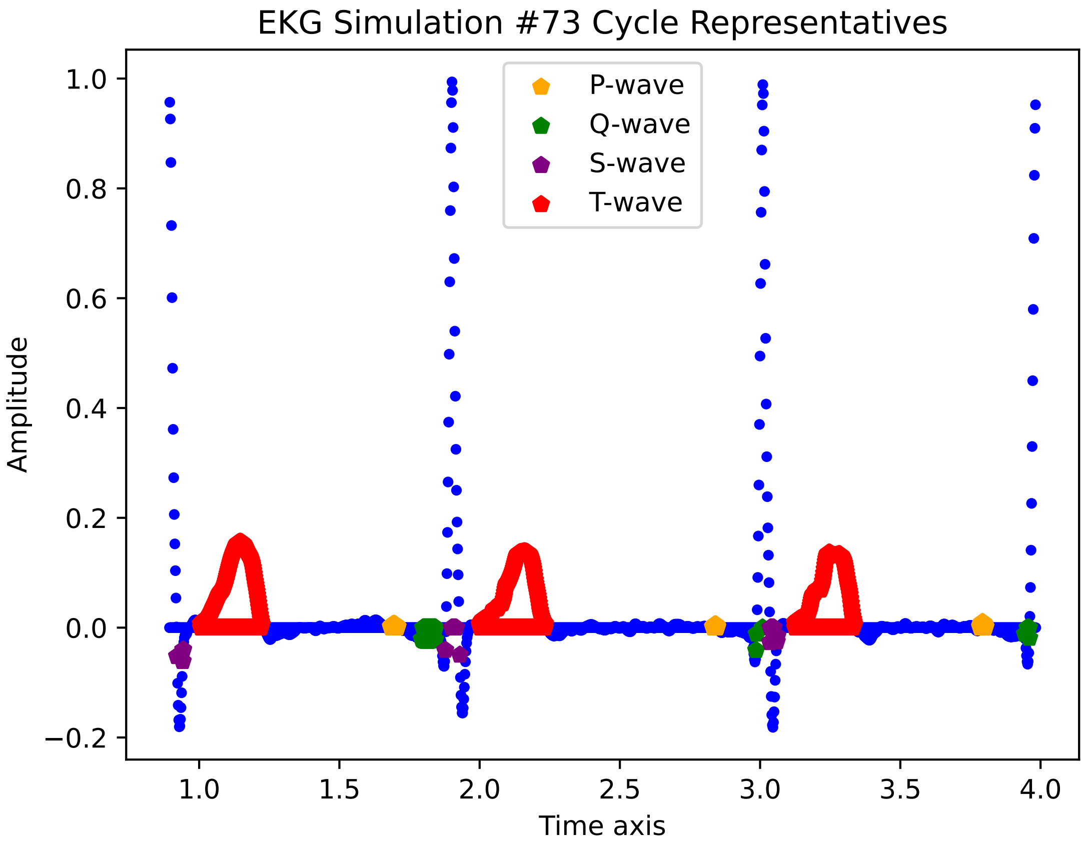

EKG Simulation #73 Cycle Representatives
P-wave
Q-wave
S-wave
T-wave
Amplitude
1.0
0.8
0.6
0.4
0.2
0.0
−0.2
1.0
1.5
2.0
2.5
3.0
3.5
4.0
Time axis

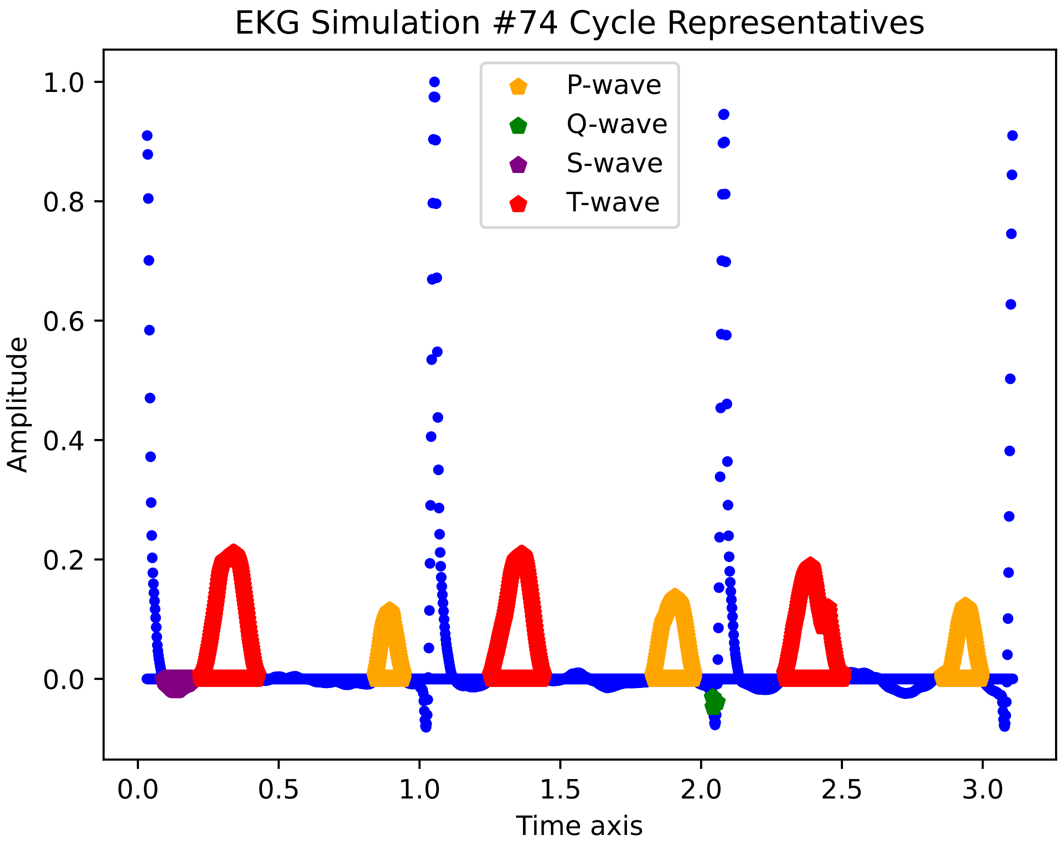

EKG Simulation #74 Cycle Representatives
P-wave
Q-wave
S-wave
T-wave
Amplitude
Time axis
1.0
0.8
0.6
0.4
0.2
0.0
0.0
0.5
1.0
1.5
2.0
2.5
3.0

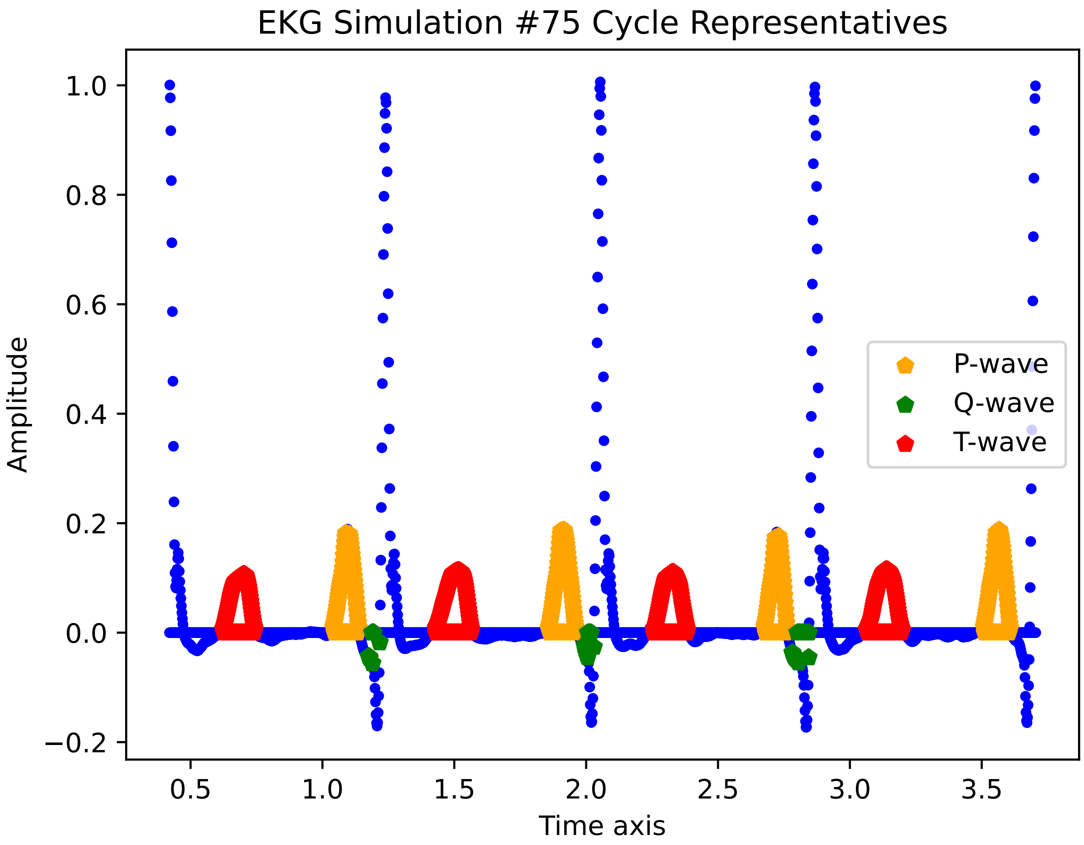

EKG Simulation #75 Cycle Representatives
Amplitude
Time axis
1.0
0.8
0.6
0.4
0.2
0.0
−0.2
0.5
1.0
1.5
2.0
2.5
3.0
3.5
P-wave
Q-wave
T-wave

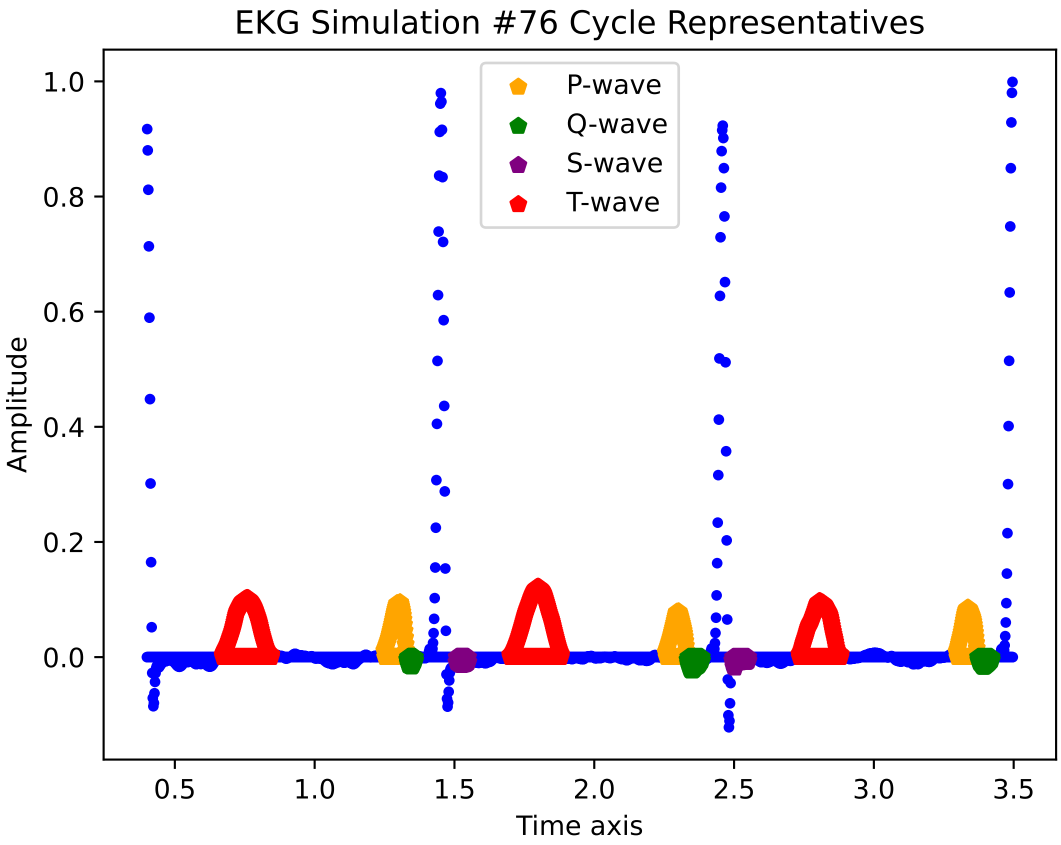

EKG Simulation #76 Cycle Representatives
P-wave
Q-wave
S-wave
T-wave
Amplitude
Time axis
1.0
0.8
0.6
0.4
0.2
0.0
0.5
1.0
1.5
2.0
2.5
3.0
3.5

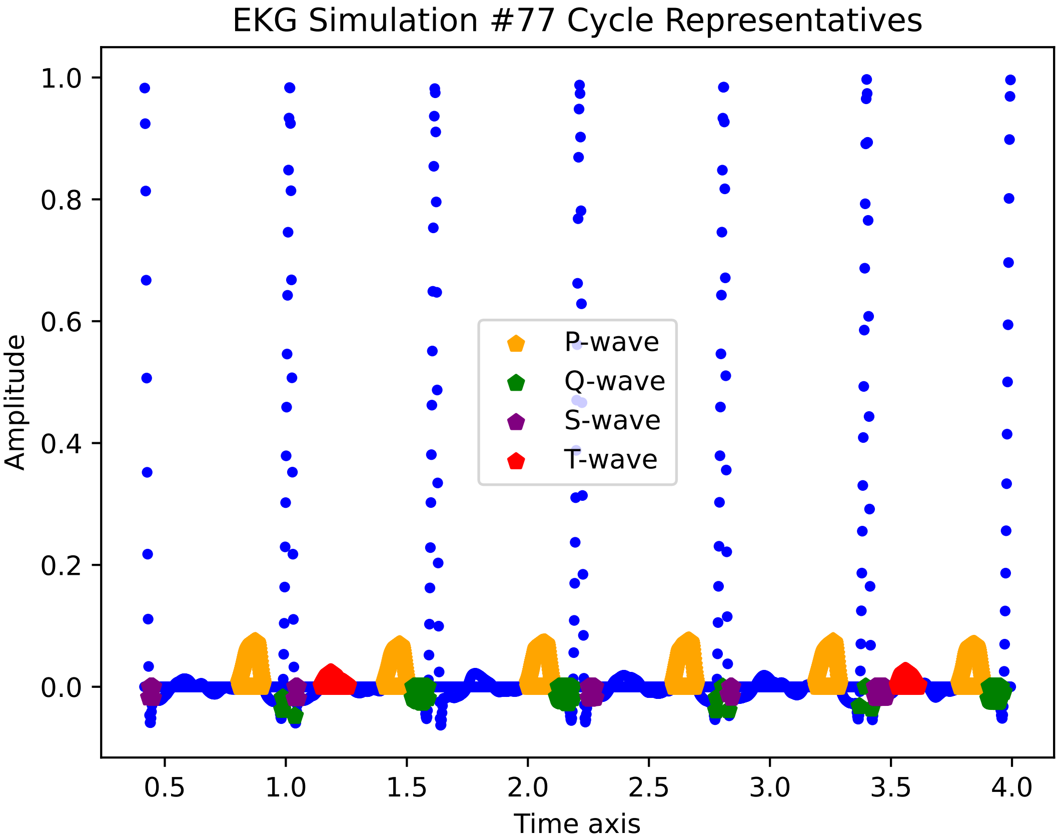

EKG Simulation #77 Cycle Representatives
Amplitude
Time axis
P-wave
Q-wave
S-wave
T-wave
1.0
0.8
0.6
0.4
0.2
0.0
0.5
1.0
1.5
2.0
2.5
3.0
3.5
4.0

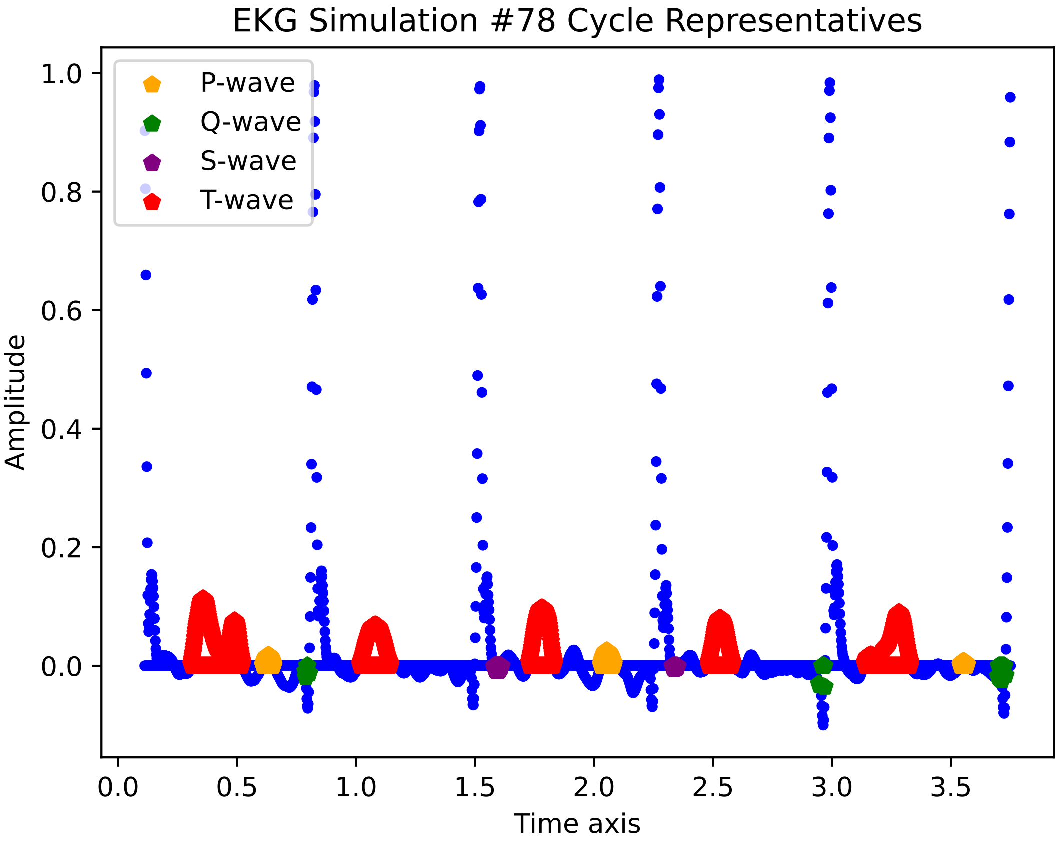

EKG Simulation #78 Cycle Representatives
P-wave
Q-wave
S-wave
T-wave
Amplitude
Time axis
1.0
0.8
0.6
0.4
0.2
0.0
0.0
0.5
1.0
1.5
2.0
2.5
3.0
3.5

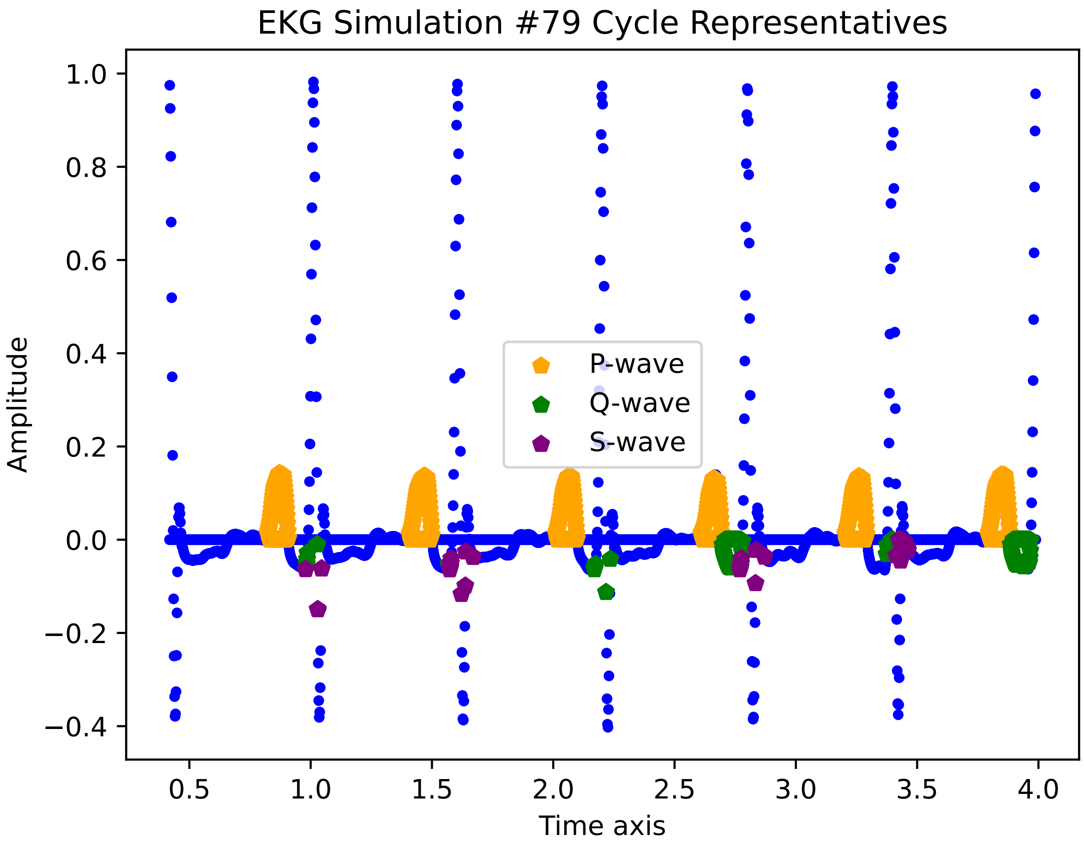
EKG Simulation #79 Cycle Representatives
Amplitude
Time axis
P-wave
Q-wave
S-wave
1.0
0.8
0.6
0.4
0.2
0.0
−0.2
−0.4
0.5
1.0
1.5
2.0
2.5
3.0
3.5
4.0

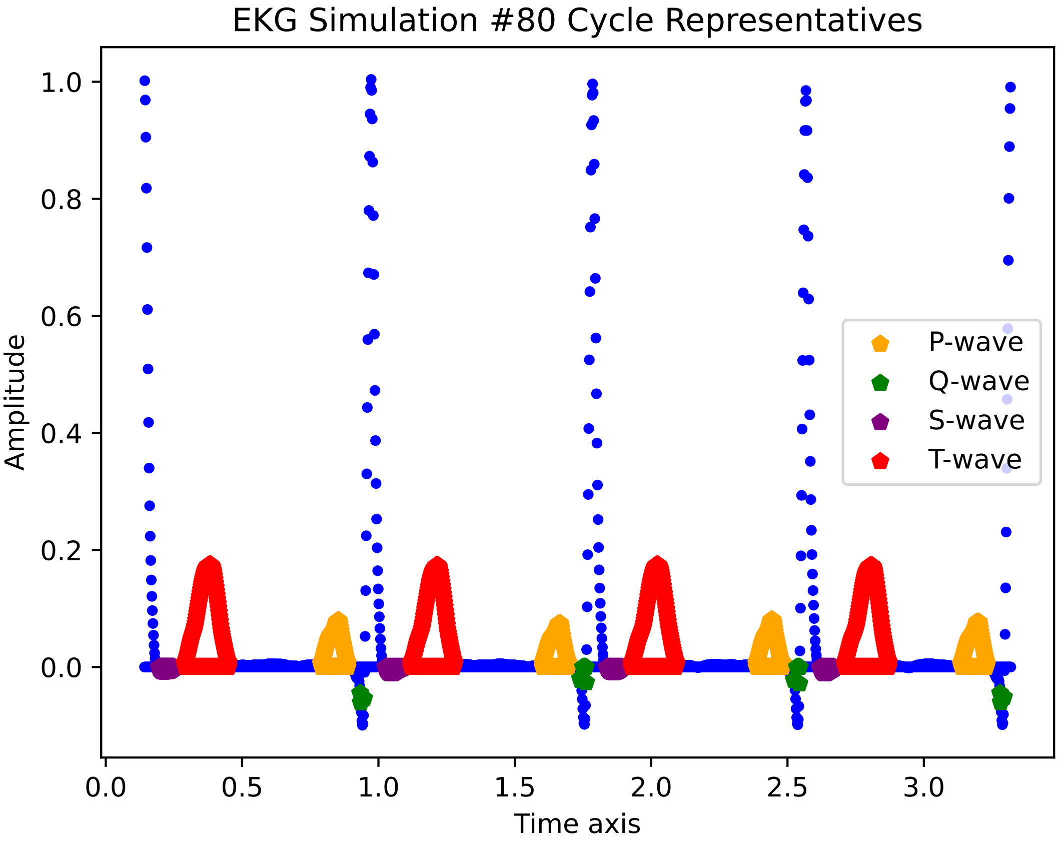

EKG Simulation #80 Cycle Representatives
Amplitude
Time axis
P-wave
Q-wave
S-wave
T-wave
1.0
0.8
0.6
0.4
0.2
0.0
0.0
0.5
1.0
1.5
2.0
2.5
3.0

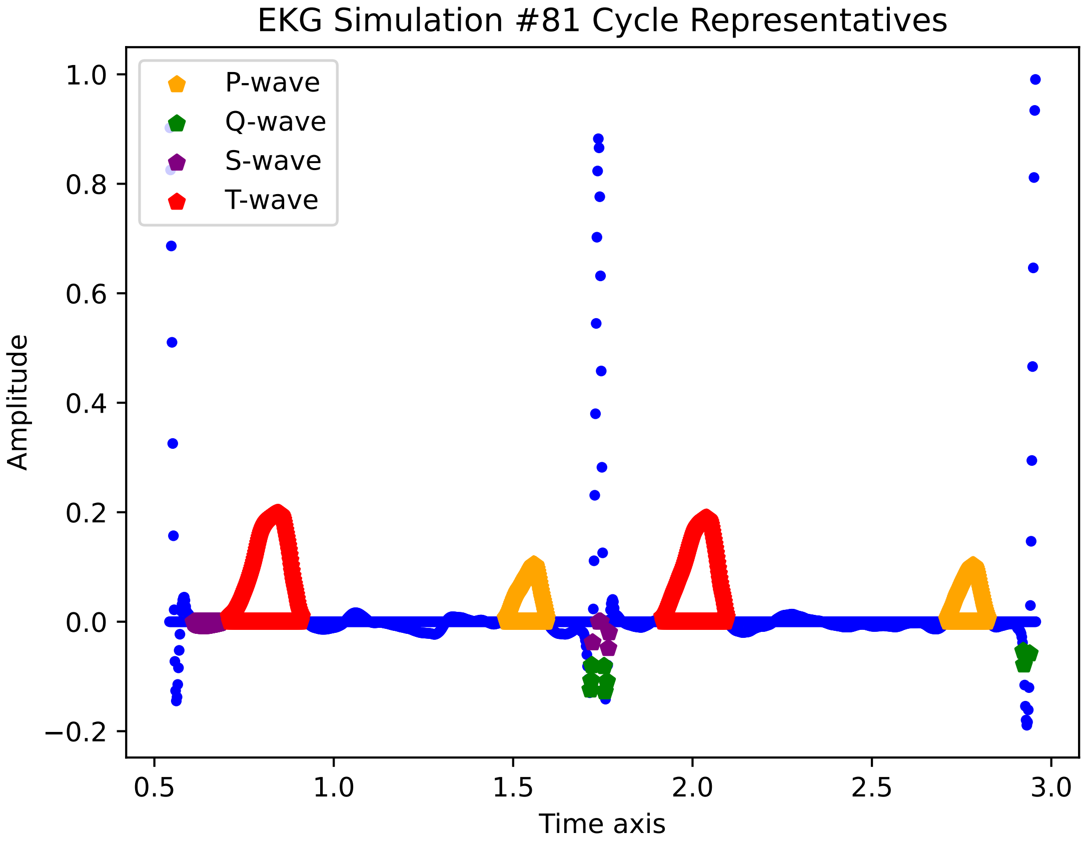

EKG Simulation #81 Cycle Representatives
P-wave
Q-wave
S-wave
T-wave
Amplitude
Time axis
1.0
0.8
0.6
0.4
0.2
0.0
−0.2
0.5
1.0
1.5
2.0
2.5
3.0

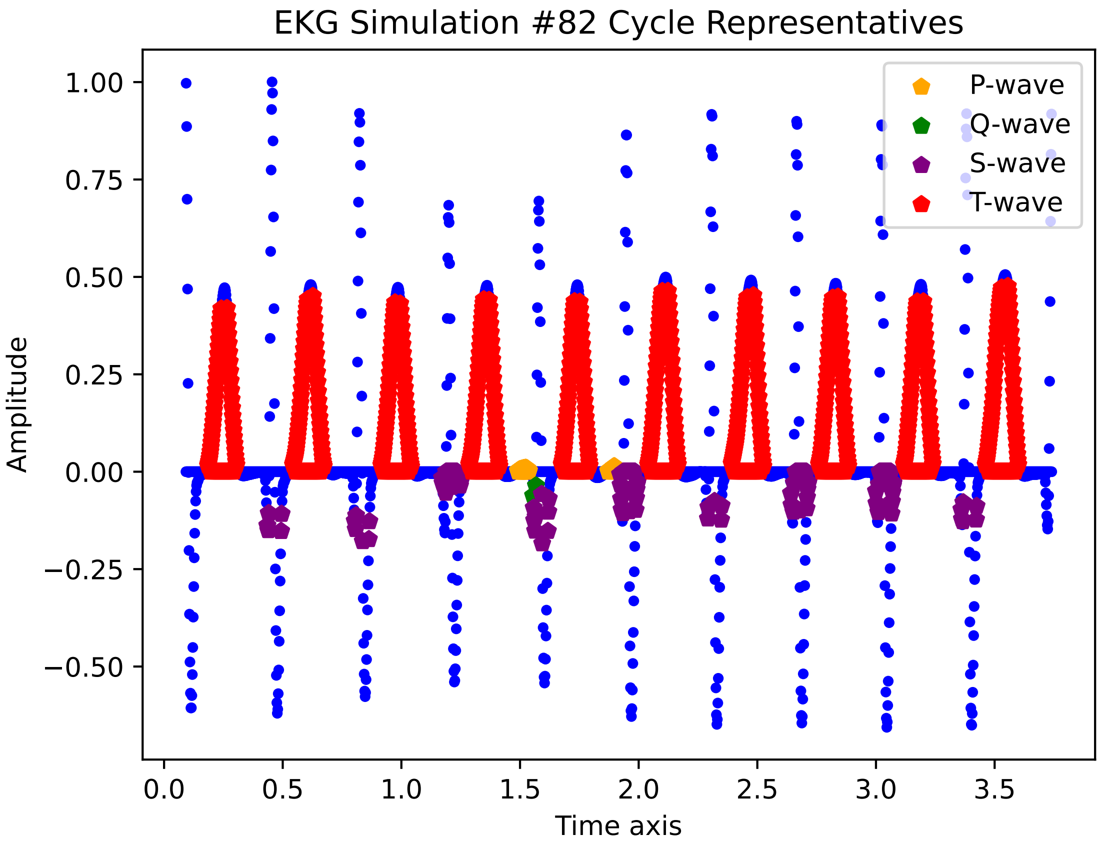

EKG Simulation #82 Cycle Representatives
1.00
0.75
0.50
0.25
0.00
−0.25
−0.50
Amplitude
0.0
0.5
1.0
1.5
2.0
2.5
3.0
3.5
Time axis
P-wave
Q-wave
S-wave
T-wave

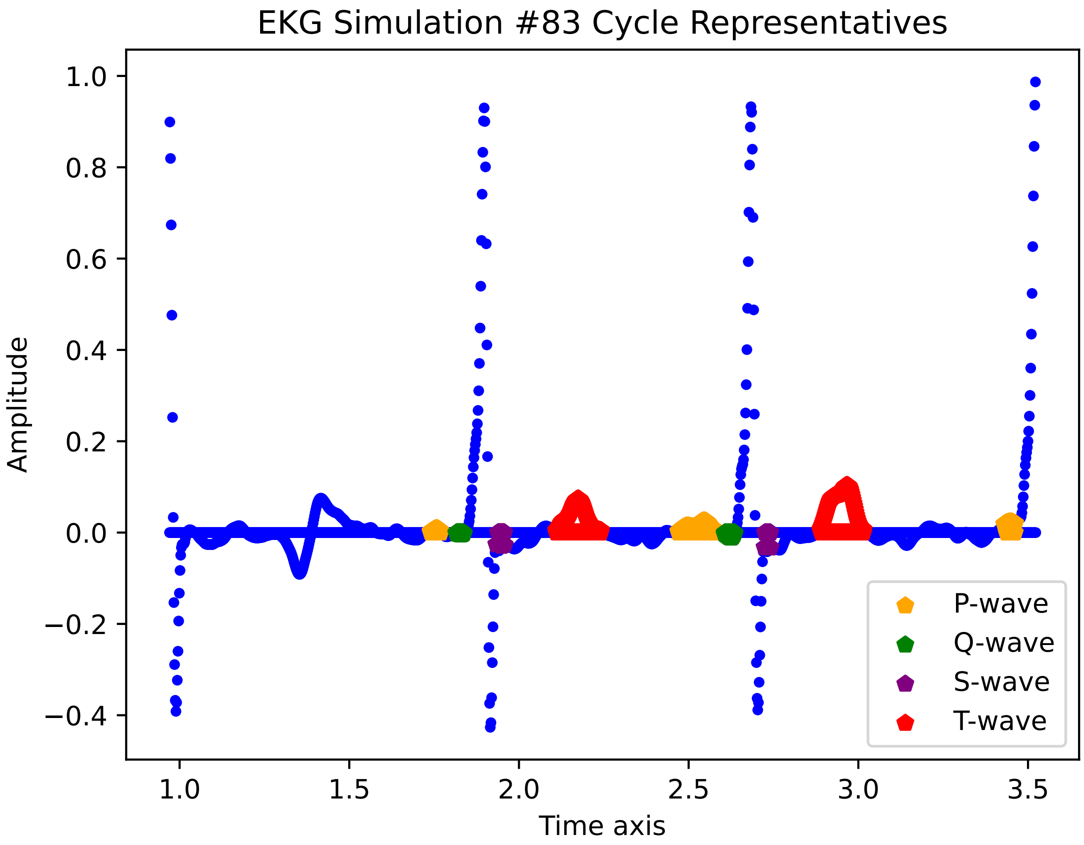

EKG Simulation #83 Cycle Representatives
Amplitude
Time axis
1.0
0.8
0.6
0.4
0.2
0.0
−0.2
−0.4
1.0
1.5
2.0
2.5
3.0
3.5
P-wave
Q-wave
S-wave
T-wave

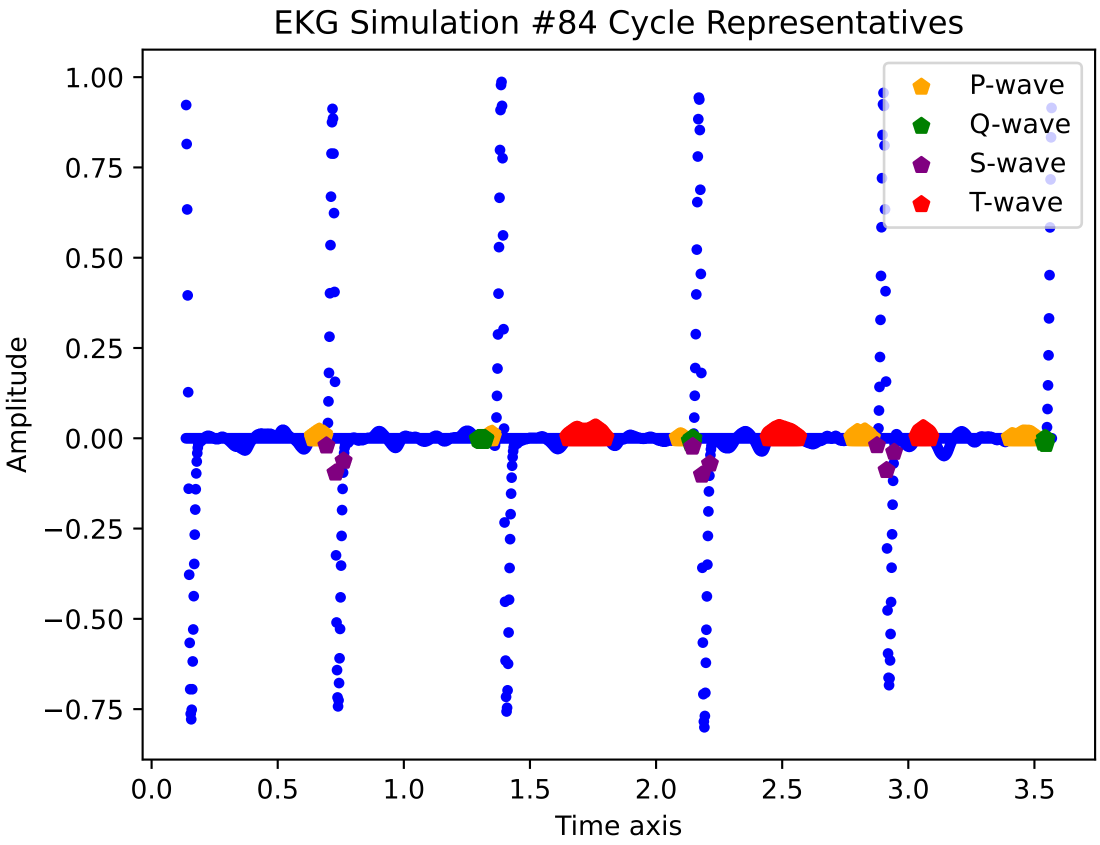
EKG Simulation #84 Cycle Representatives
P-wave
Q-wave
S-wave
T-wave
Amplitude
Time axis
1.00
0.75
0.50
0.25
0.00
−0.25
−0.50
−0.75
0.0
0.5
1.0
1.5
2.0
2.5
3.0
3.5

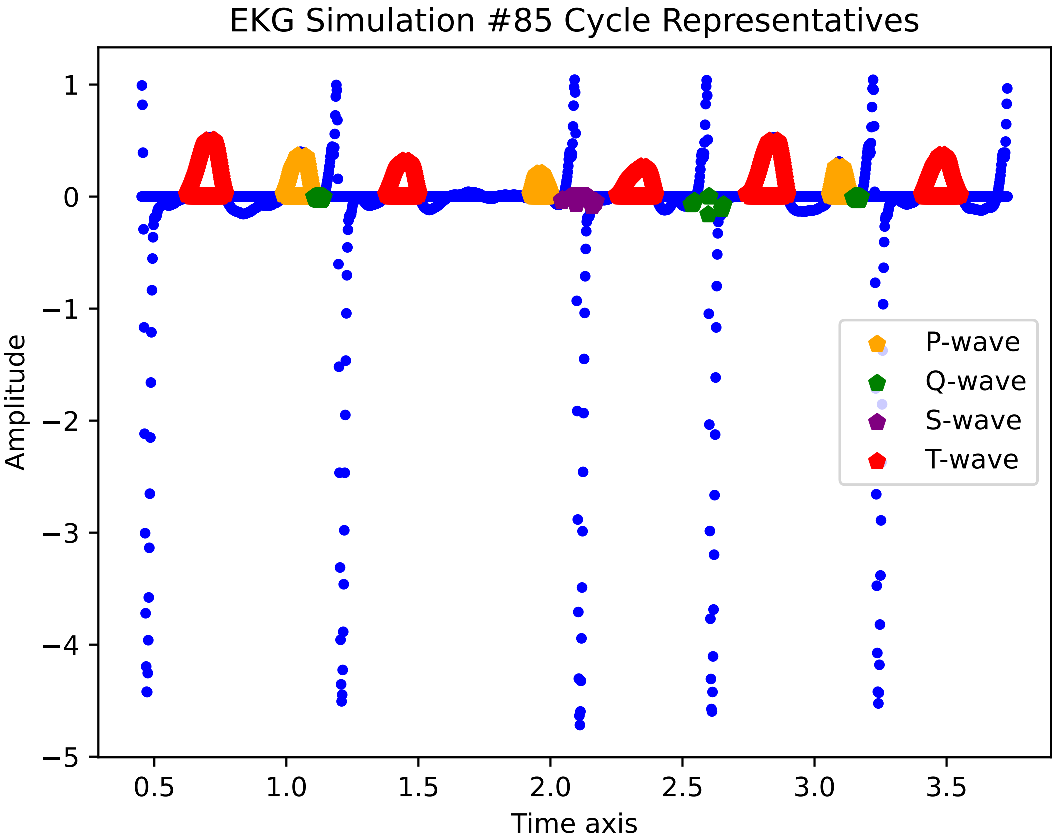
EKG Simulation #85 Cycle Representatives
Amplitude
Time axis
1
0
−1
−2
−3
−4
−5
0.5
1.0
1.5
2.0
2.5
3.0
3.5
P-wave
Q-wave
S-wave
T-wave

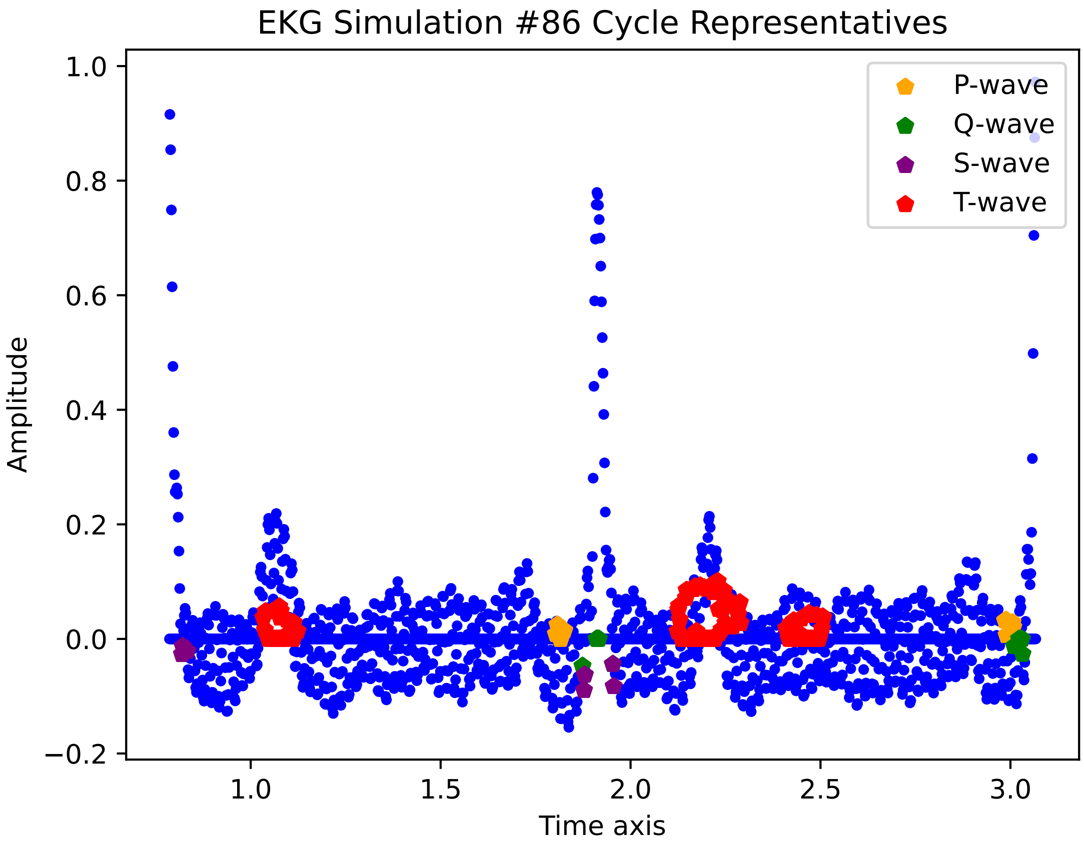

EKG Simulation #86 Cycle Representatives
Amplitude
Time axis
P-wave
Q-wave
S-wave
T-wave
1.0
0.8
0.6
0.4
0.2
0.0
−0.2
1.0
1.5
2.0
2.5
3.0

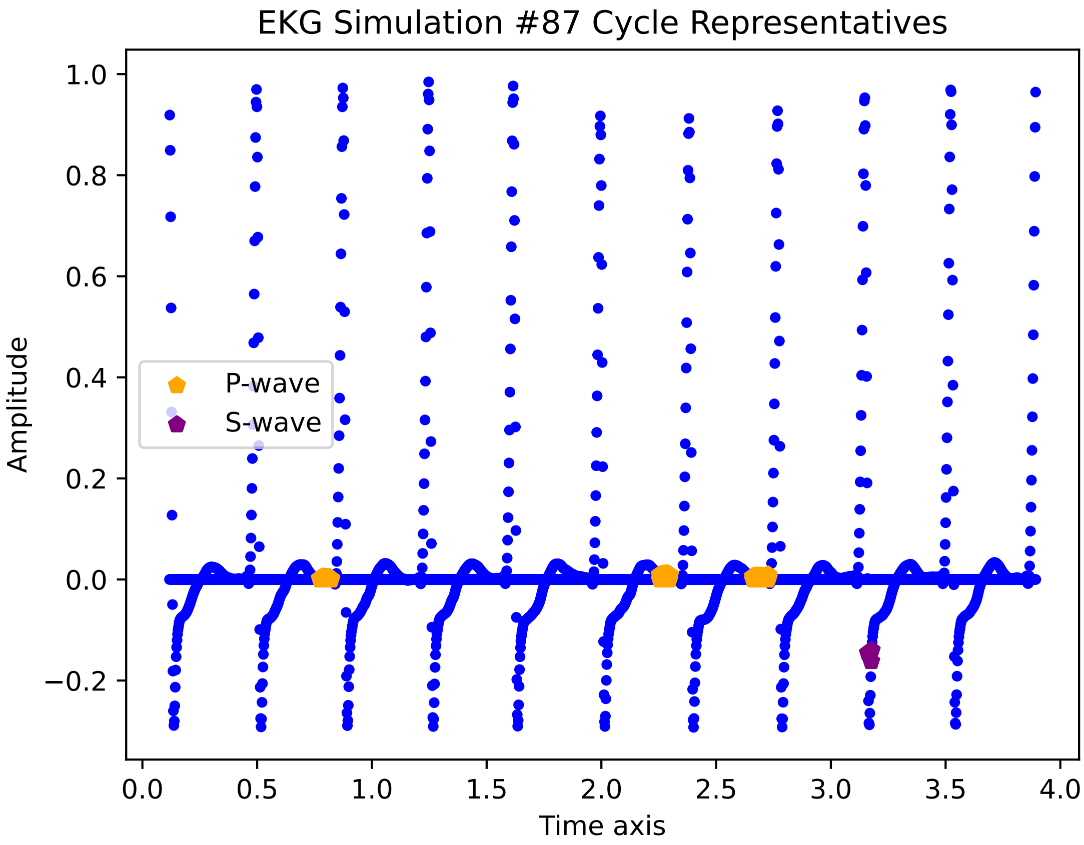

EKG Simulation #87 Cycle Representatives
1.0
0.8
0.6
0.4
0.2
0.0
−0.2
Amplitude
0.0
0.5
1.0
1.5
2.0
2.5
3.0
3.5
4.0
Time axis
P-wave
S-wave

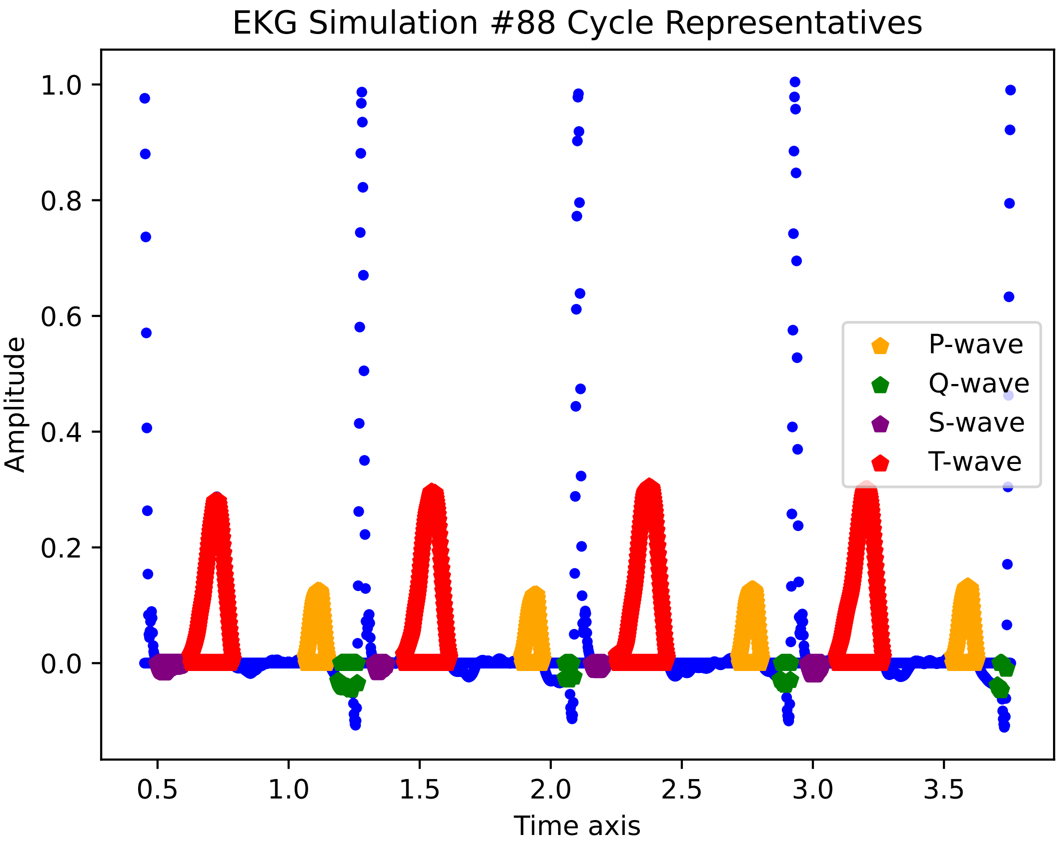
EKG Simulation #88 Cycle Representatives
Amplitude
Time axis
1.0
0.8
0.6
0.4
0.2
0.0
0.5
1.0
1.5
2.0
2.5
3.0
3.5
P-wave
Q-wave
S-wave
T-wave

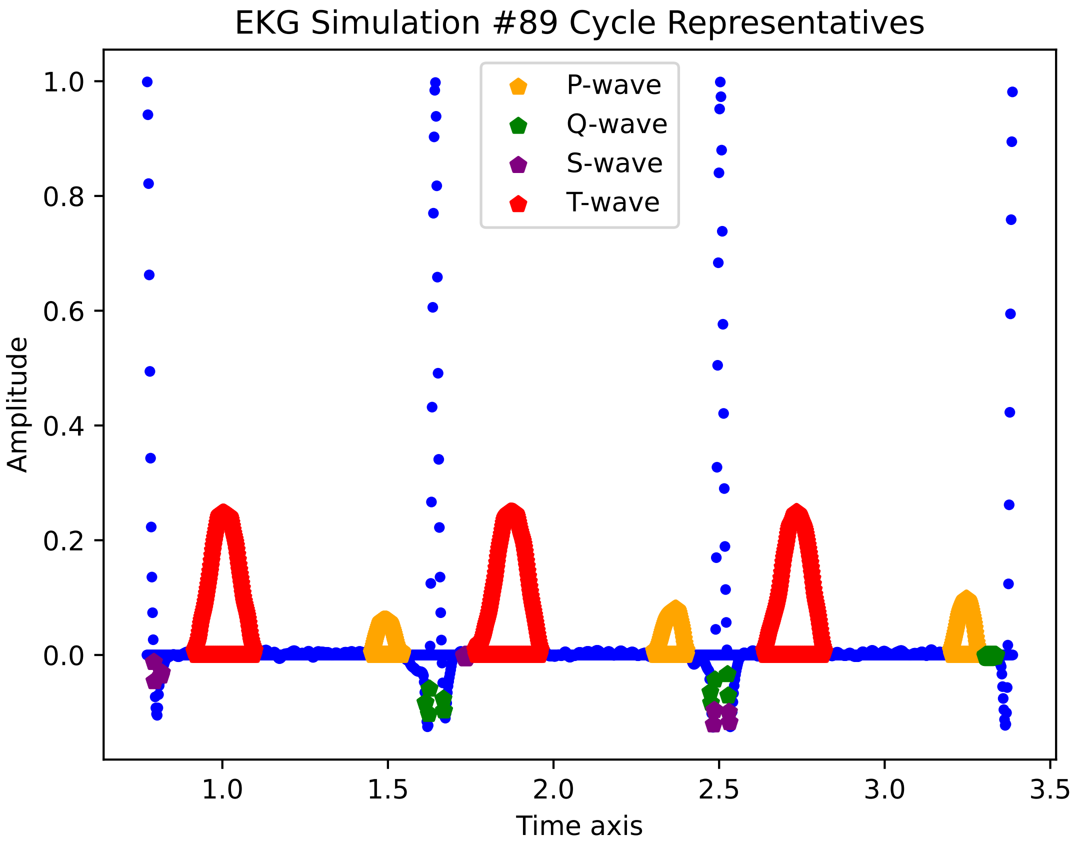
EKG Simulation #89 Cycle Representatives
P-wave
Q-wave
S-wave
T-wave
Amplitude
Time axis
1.0
0.8
0.6
0.4
0.2
0.0
1.0
1.5
2.0
2.5
3.0
3.5

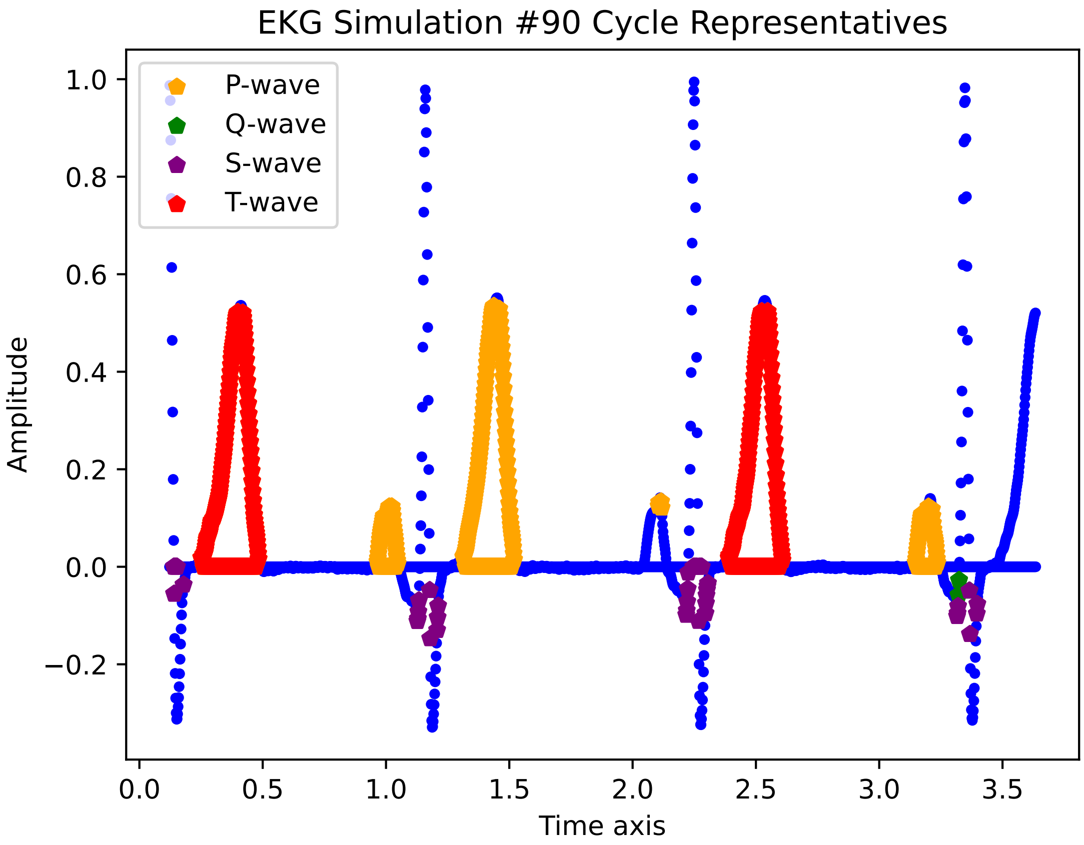

EKG Simulation #90 Cycle Representatives
P-wave
Q-wave
S-wave
T-wave
Amplitude
Time axis
1.0
0.8
0.6
0.4
0.2
0.0
−0.2
0.0
0.5
1.0
1.5
2.0
2.5
3.0
3.5

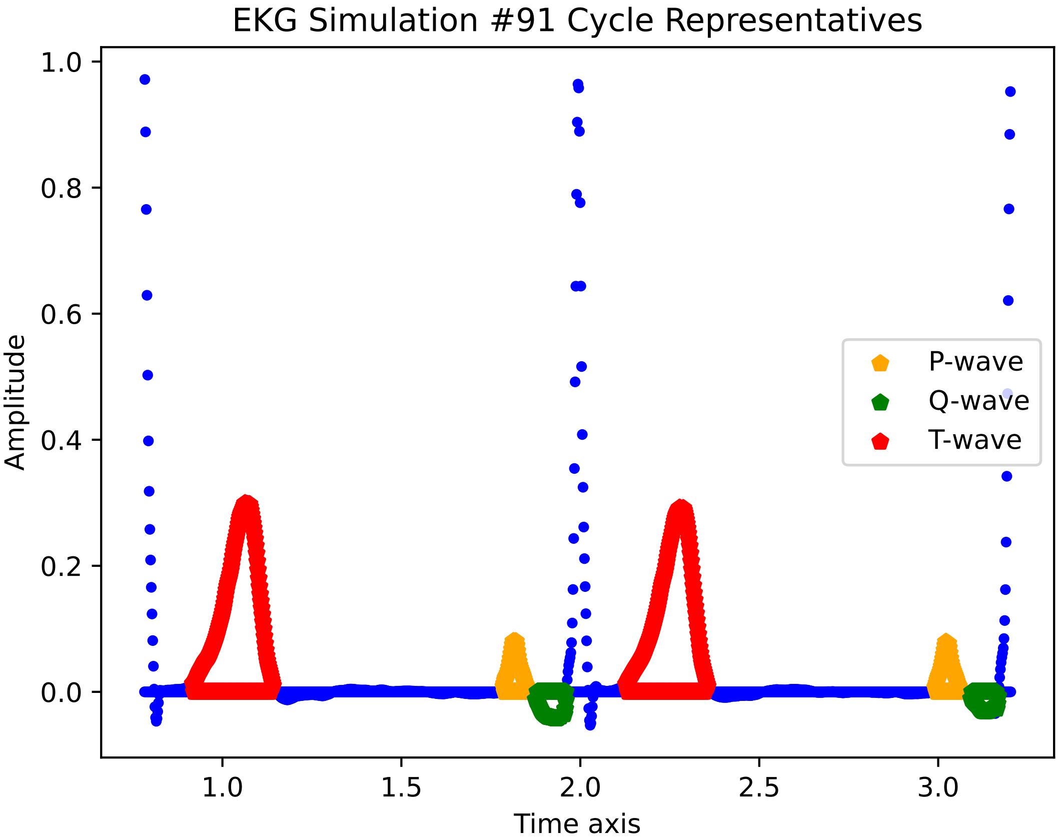

EKG Simulation #91 Cycle Representatives
Amplitude
Time axis
1.0
0.8
0.6
0.4
0.2
0.0
1.0
1.5
2.0
2.5
3.0
P-wave
Q-wave
T-wave

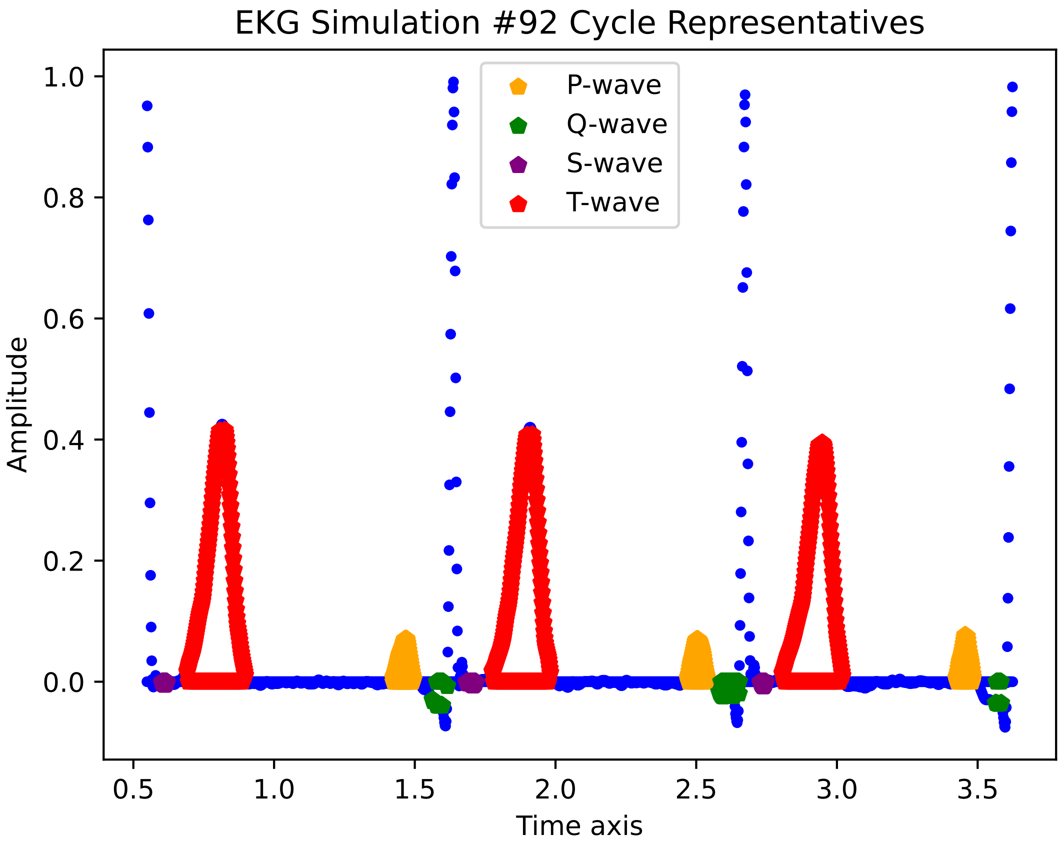
EKG Simulation #92 Cycle Representatives
P-wave
Q-wave
S-wave
T-wave
Amplitude
1.0
0.8
0.6
0.4
0.2
0.0
0.5
1.0
1.5
2.0
2.5
3.0
3.5
Time axis

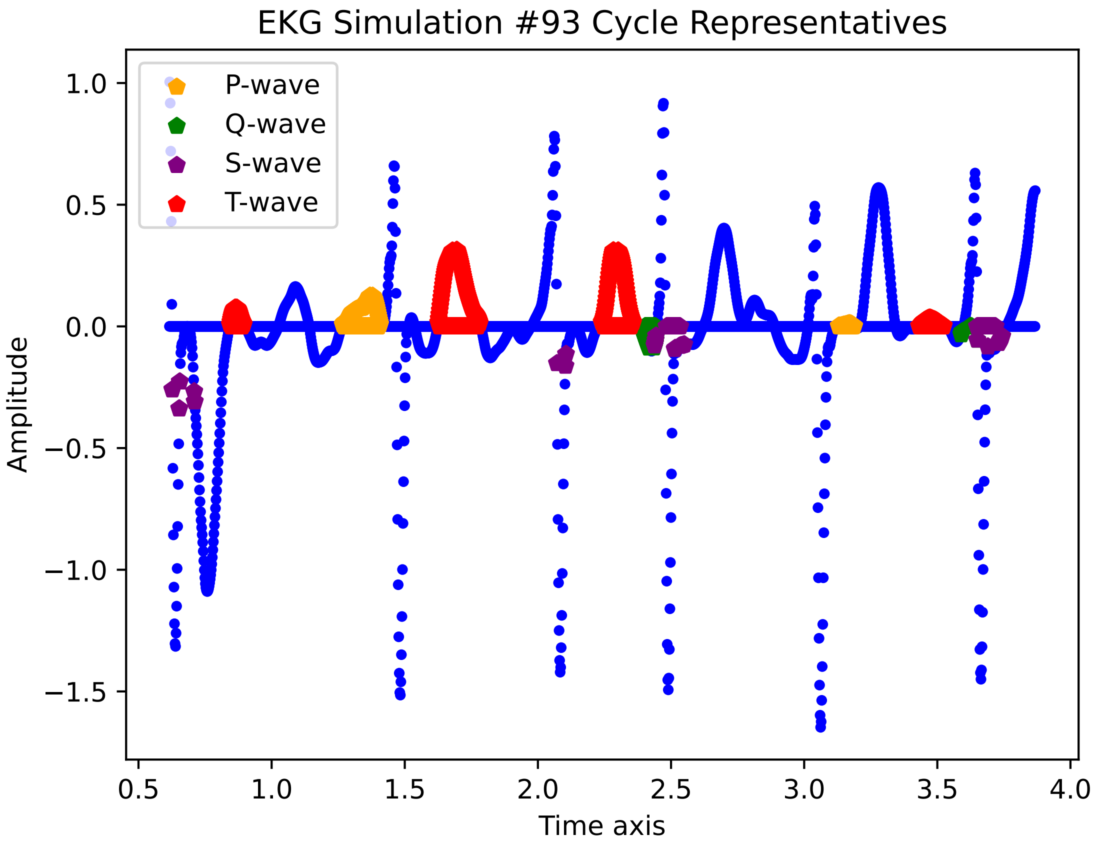

EKG Simulation #93 Cycle Representatives
P-wave
Q-wave
S-wave
T-wave
1.0
0.5
0.0
−0.5
−1.0
−1.5
Amplitude
0.5
1.0
1.5
2.0
2.5
3.0
3.5
4.0
Time axis

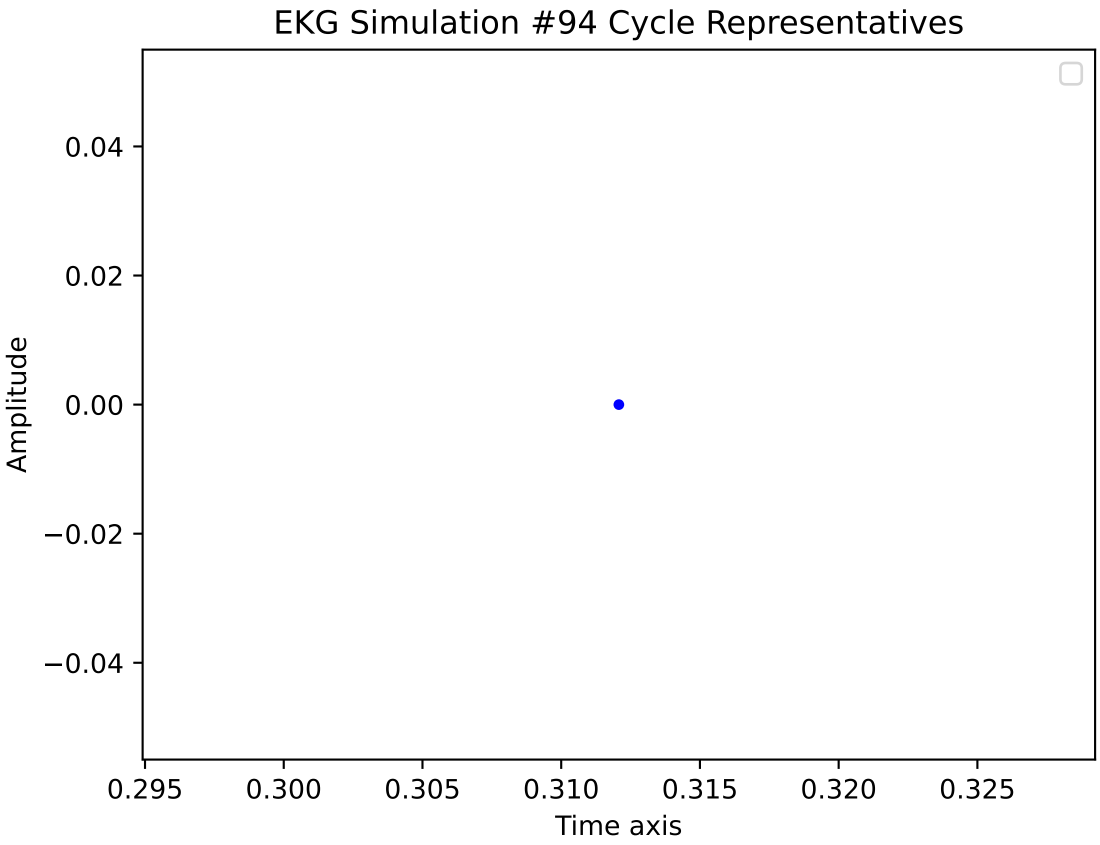
EKG Simulation #94 Cycle Representatives
Amplitude
0.04
0.02
0.00
−0.02
−0.04
Time axis
0.295
0.300
0.305
0.310
0.315
0.320
0.325

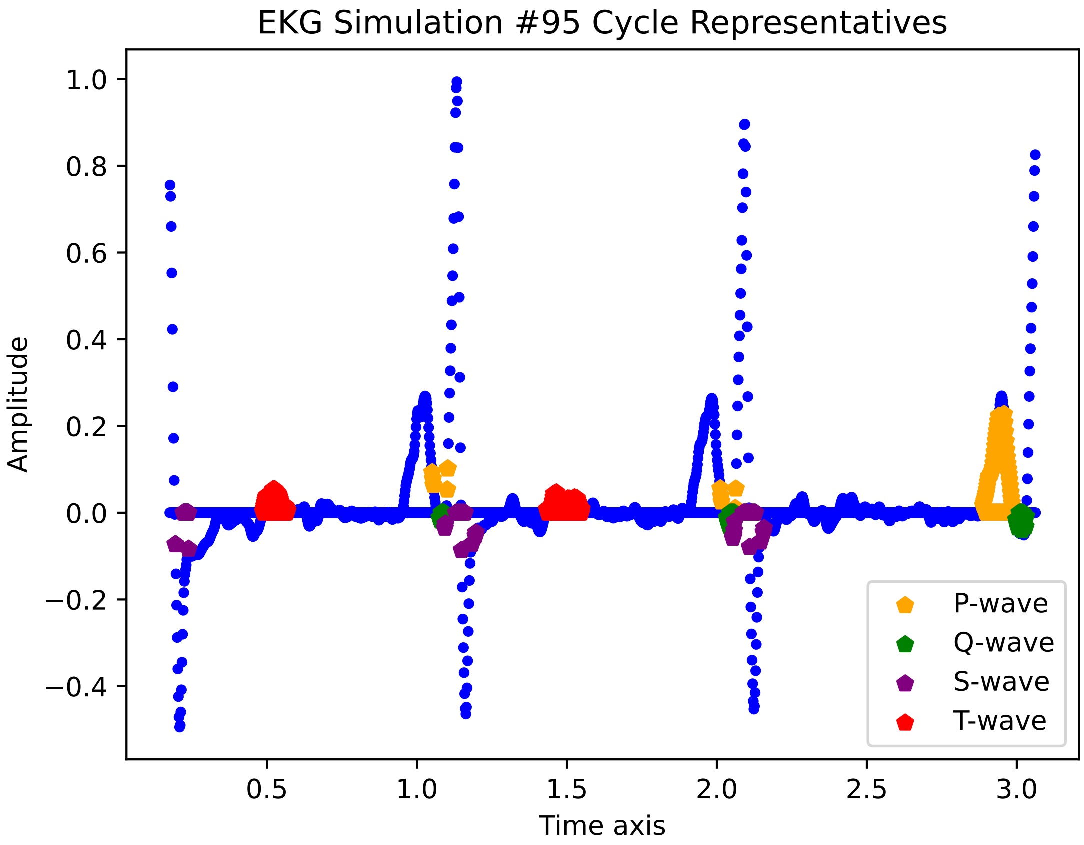
EKG Simulation #95 Cycle Representatives
Amplitude
Time axis
1.0
0.8
0.6
0.4
0.2
0.0
−0.2
−0.4
0.5
1.0
1.5
2.0
2.5
3.0
P-wave
Q-wave
S-wave
T-wave

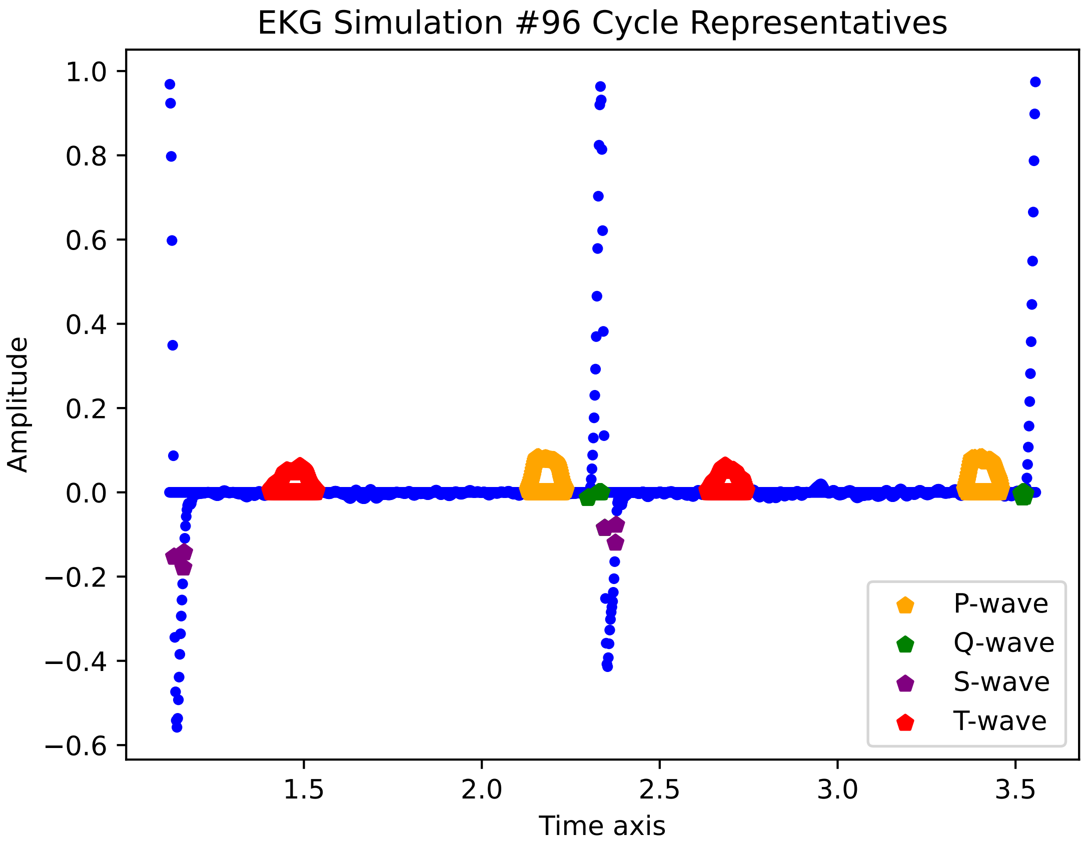
EKG Simulation #96 Cycle Representatives
Amplitude
Time axis
1.0
0.8
0.6
0.4
0.2
0.0
−0.2
−0.4
−0.6
1.5
2.0
2.5
3.0
3.5
P-wave
Q-wave
S-wave
T-wave

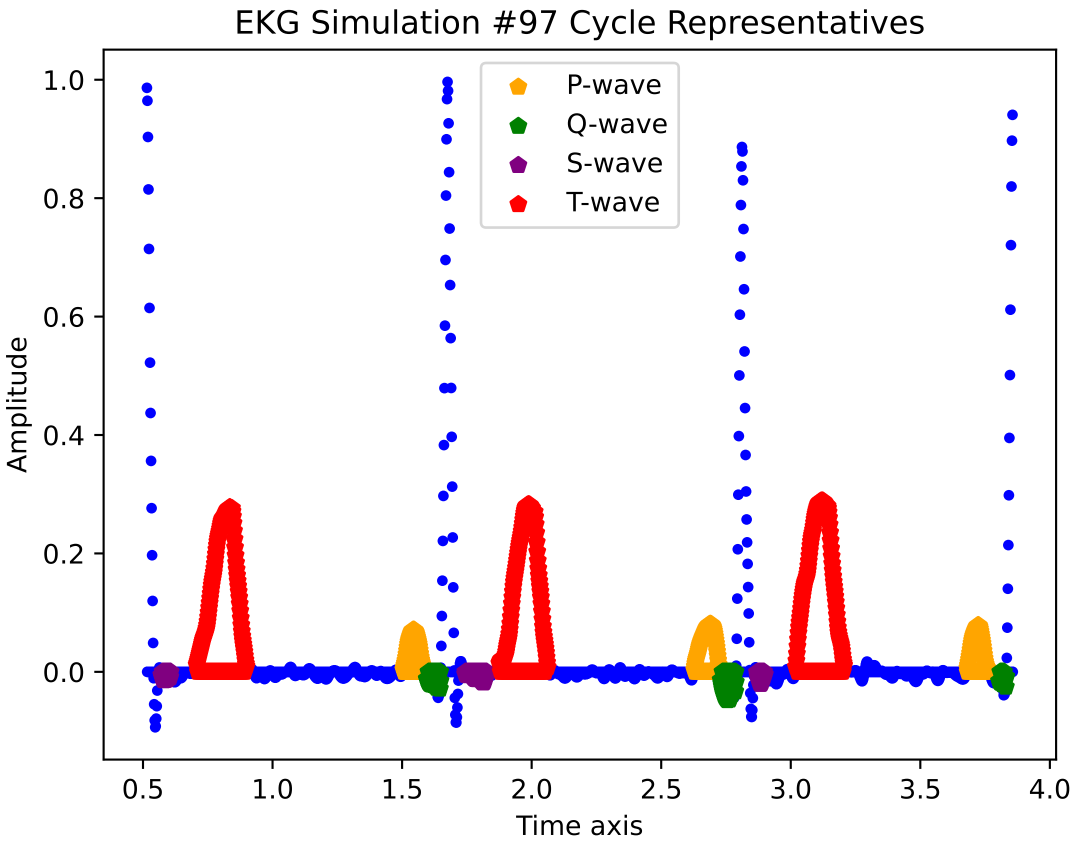
EKG Simulation #97 Cycle Representatives
P-wave
Q-wave
S-wave
T-wave
Amplitude
Time axis
1.0
0.8
0.6
0.4
0.2
0.0
0.5
1.0
1.5
2.0
2.5
3.0
3.5
4.0

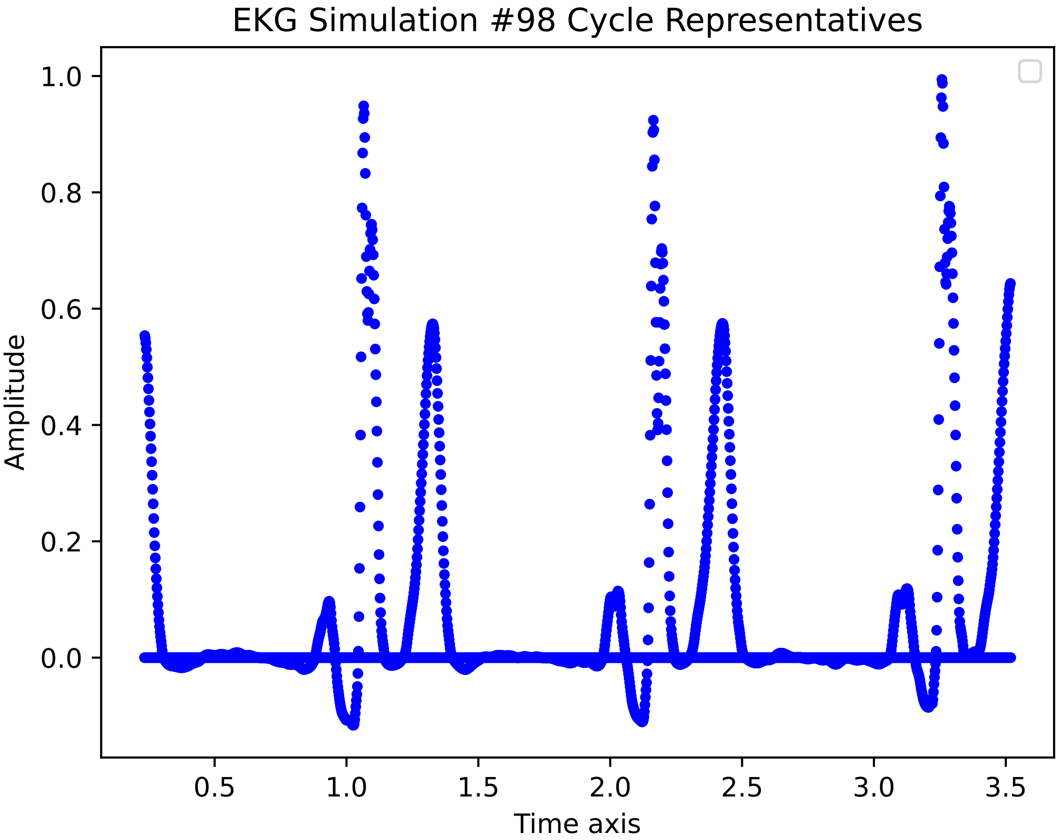
EKG Simulation #98 Cycle Representatives
Amplitude
Time axis
1.0
0.8
0.6
0.4
0.2
0.0
0.5
1.0
1.5
2.0
2.5
3.0
3.5

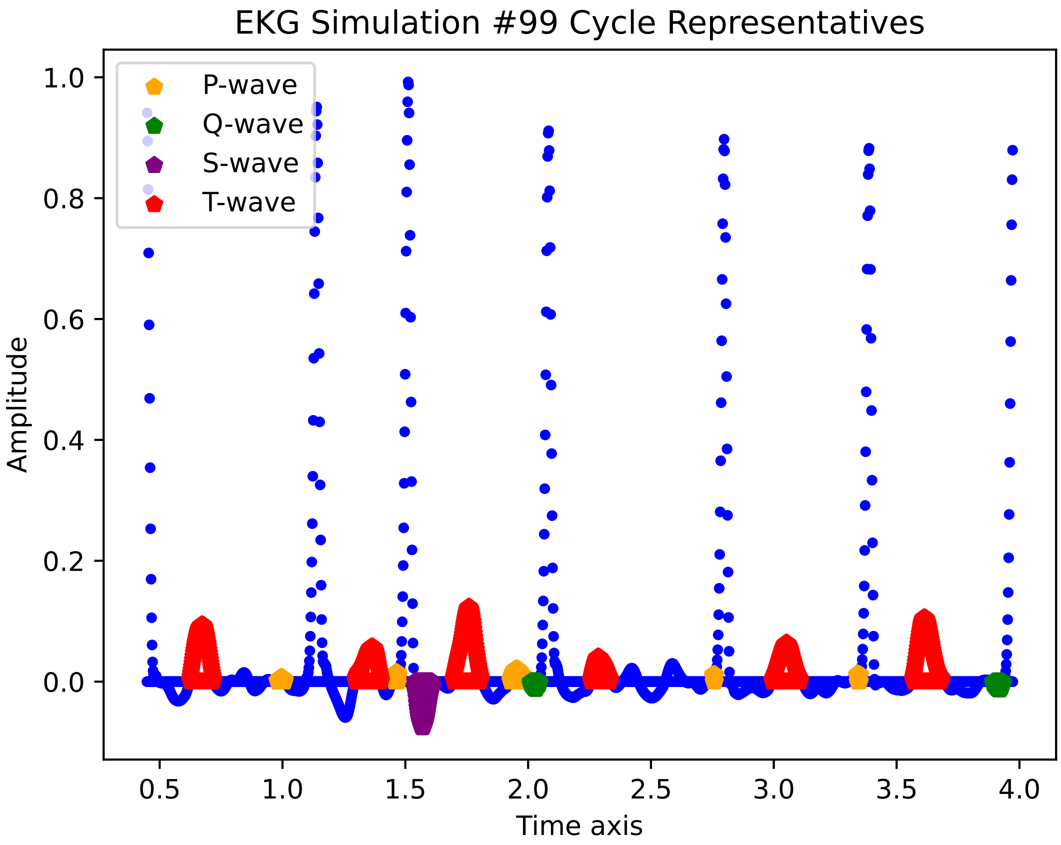

EKG Simulation #99 Cycle Representatives
P-wave
Q-wave
S-wave
T-wave
1.0
0.8
0.6
0.4
0.2
0.0
Amplitude
0.5
1.0
1.5
2.0
2.5
3.0
3.5
4.0
Time axis

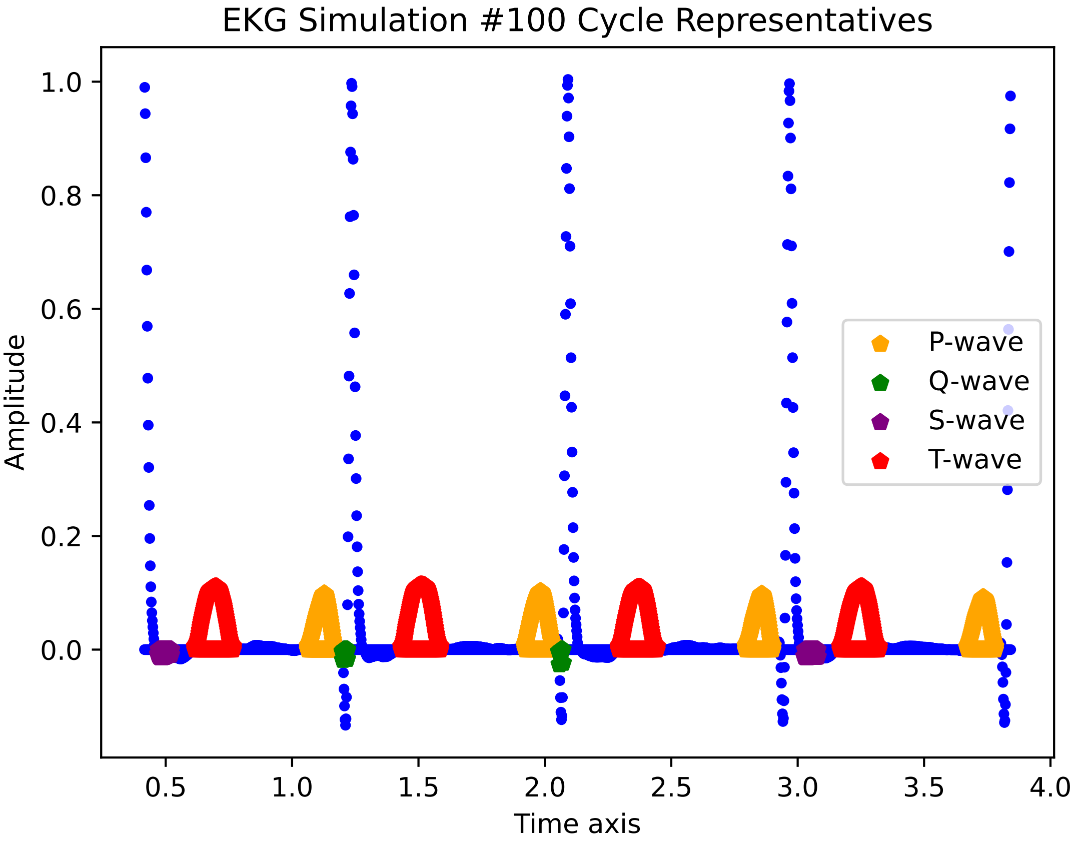

EKG Simulation #100 Cycle Representatives
1.0
0.8
0.6
0.4
0.2
0.0
Amplitude
0.5
1.0
1.5
2.0
2.5
3.0
3.5
4.0
Time axis
P-wave
Q-wave
S-wave
T-wave

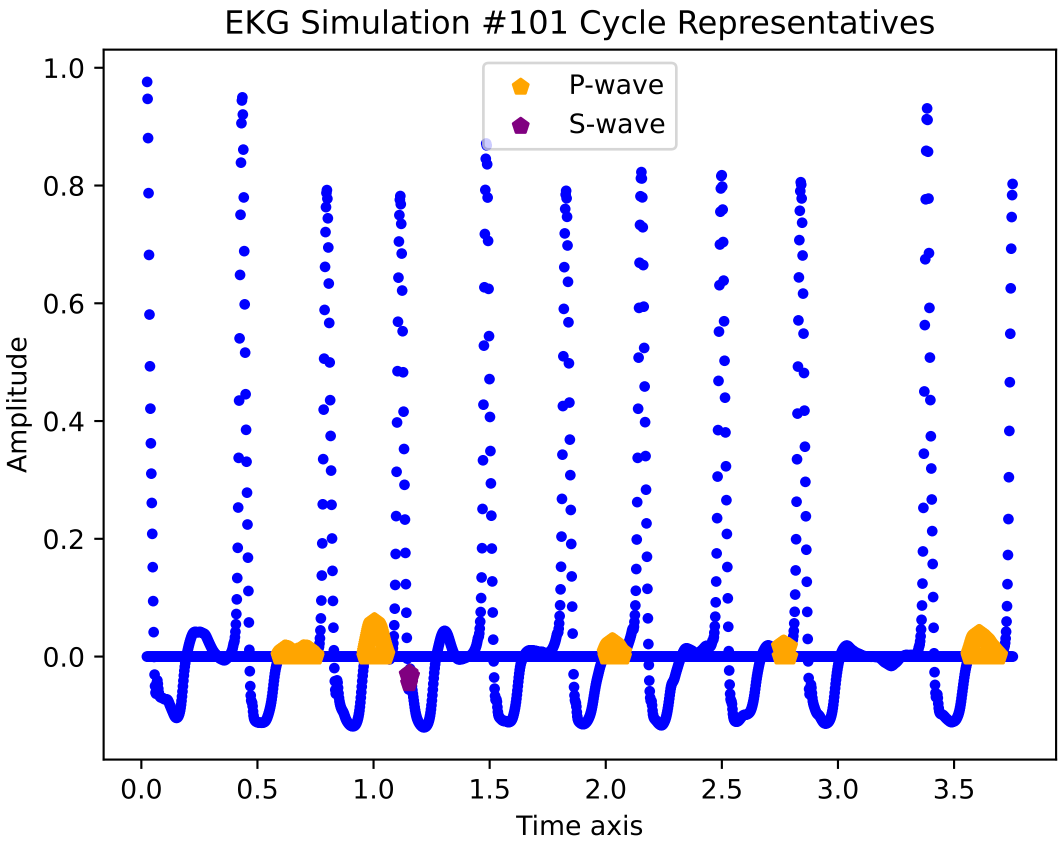
EKG Simulation #101 Cycle Representatives
P-wave
S-wave
Amplitude
1.0
0.8
0.6
0.4
0.2
0.0
0.0
0.5
1.0
1.5
2.0
2.5
3.0
3.5
Time axis

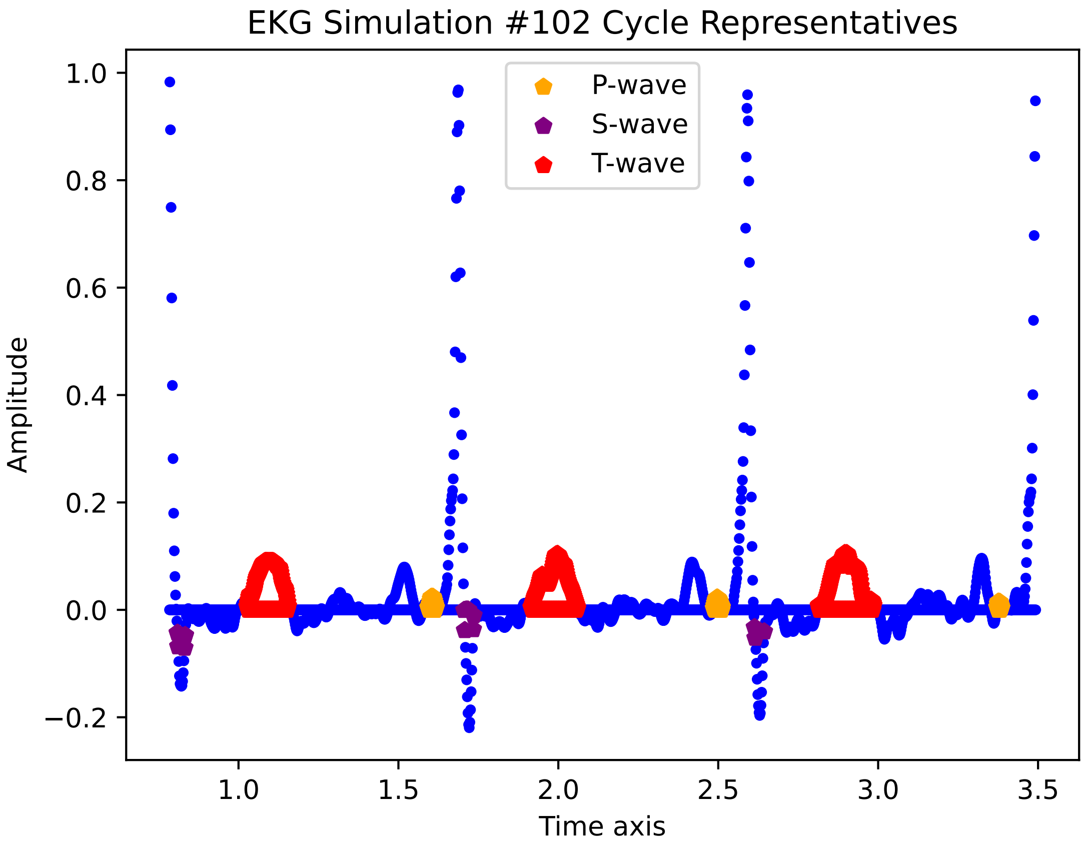

EKG Simulation #102 Cycle Representatives
P-wave
S-wave
T-wave
Amplitude
Time axis
1.0
0.8
0.6
0.4
0.2
0.0
−0.2
1.0
1.5
2.0
2.5
3.0
3.5

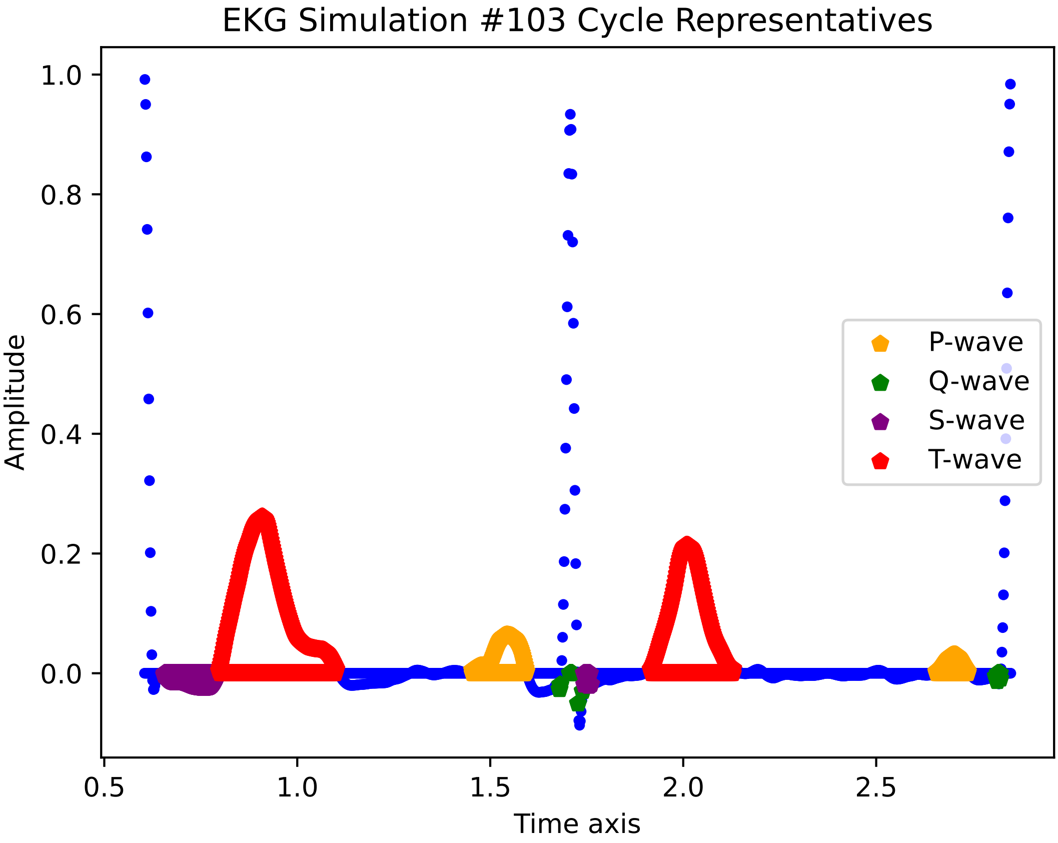

EKG Simulation #103 Cycle Representatives
Amplitude
Time axis
1.0
0.8
0.6
0.4
0.2
0.0
0.5
1.0
1.5
2.0
2.5
P-wave
Q-wave
S-wave
T-wave

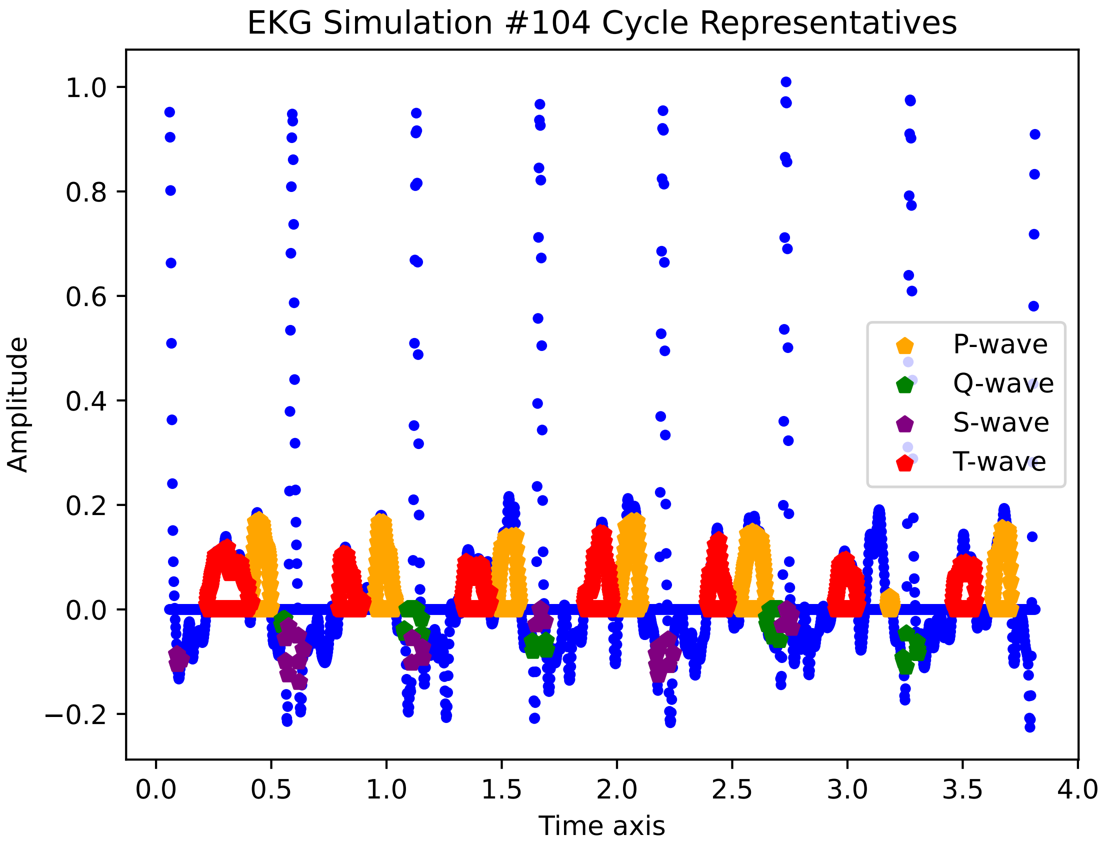

EKG Simulation #104 Cycle Representatives
Amplitude
Time axis
1.0
0.8
0.6
0.4
0.2
0.0
−0.2
0.0
0.5
1.0
1.5
2.0
2.5
3.0
3.5
4.0
P-wave
Q-wave
S-wave
T-wave

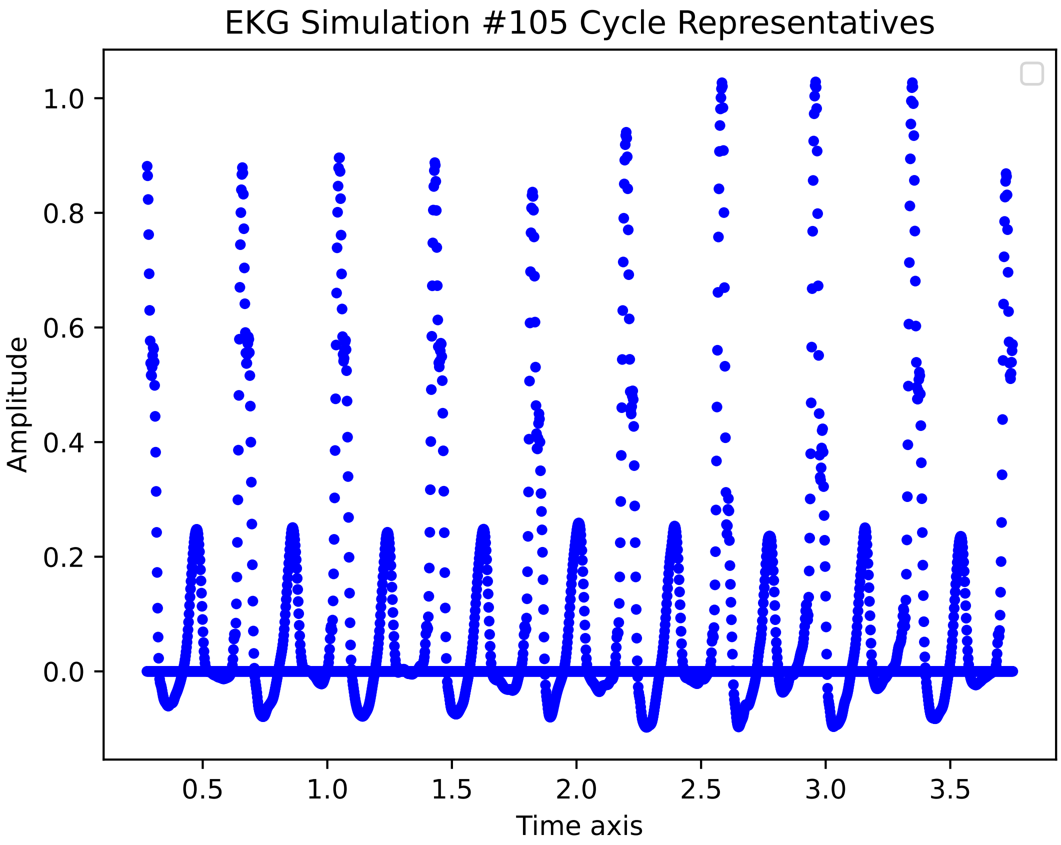
EKG Simulation #105 Cycle Representatives
Amplitude
Time axis
1.0
0.8
0.6
0.4
0.2
0.0
0.5
1.0
1.5
2.0
2.5
3.0
3.5

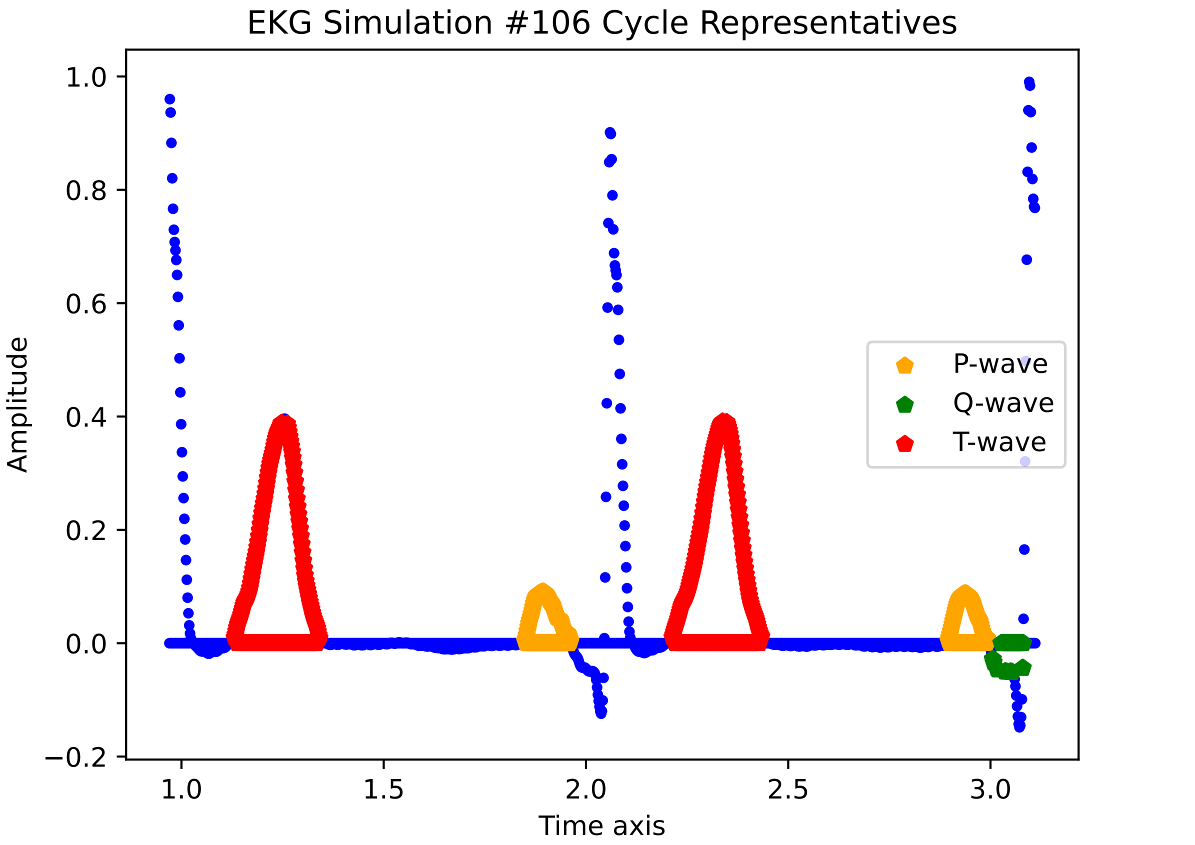

EKG Simulation #106 Cycle Representatives
Amplitude
Time axis
P-wave
Q-wave
T-wave
1.0
0.8
0.6
0.4
0.2
0.0
−0.2
1.0
1.5
2.0
2.5
3.0

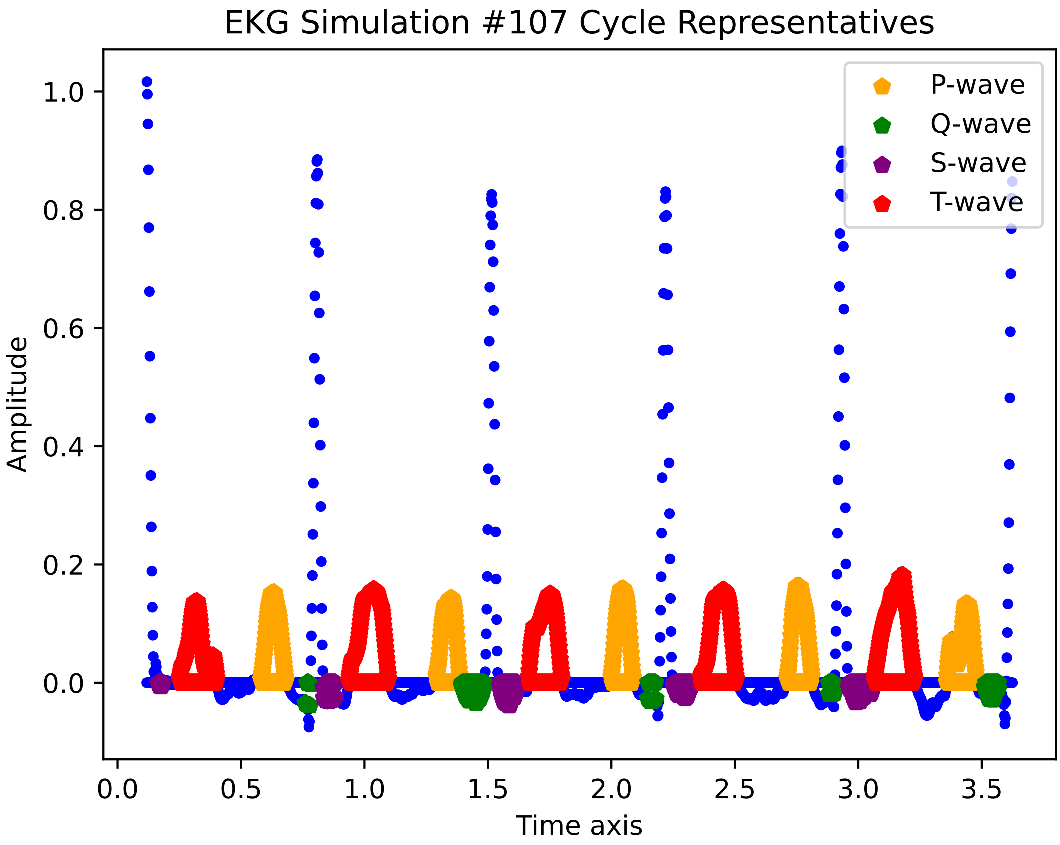
EKG Simulation #107 Cycle Representatives
Amplitude
Time axis
1.0
0.8
0.6
0.4
0.2
0.0
0.0
0.5
1.0
1.5
2.0
2.5
3.0
3.5
P-wave
Q-wave
S-wave
T-wave

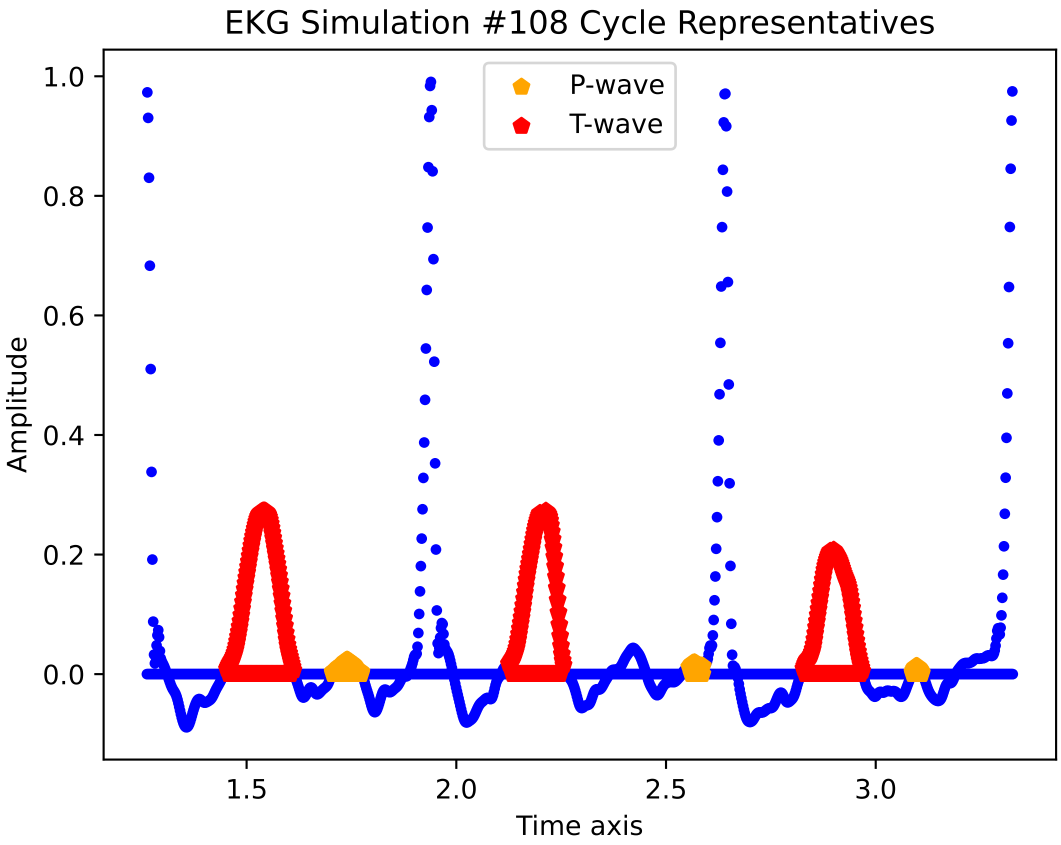
EKG Simulation #108 Cycle Representatives
P-wave
T-wave
Amplitude
Time axis
1.0
0.8
0.6
0.4
0.2
0.0
1.5
2.0
2.5
3.0

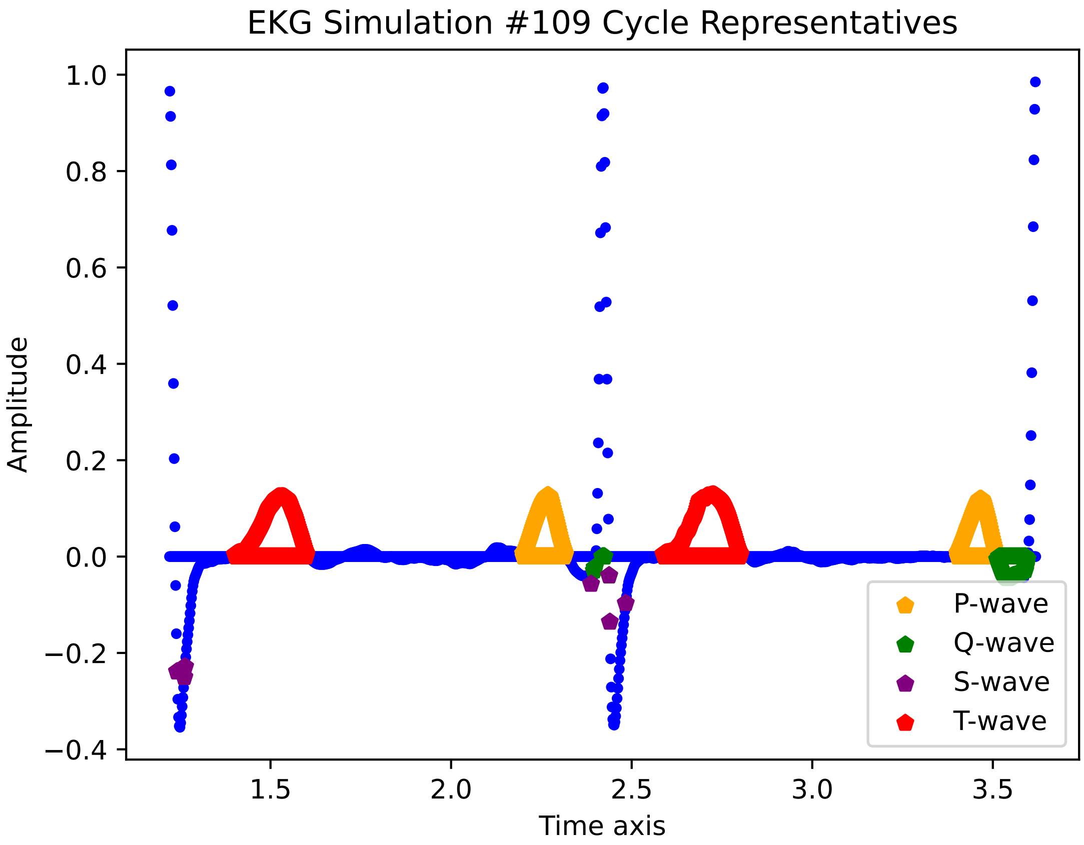

EKG Simulation #109 Cycle Representatives
Amplitude
Time axis
1.0
0.8
0.6
0.4
0.2
0.0
−0.2
−0.4
1.5
2.0
2.5
3.0
3.5
P-wave
Q-wave
S-wave
T-wave

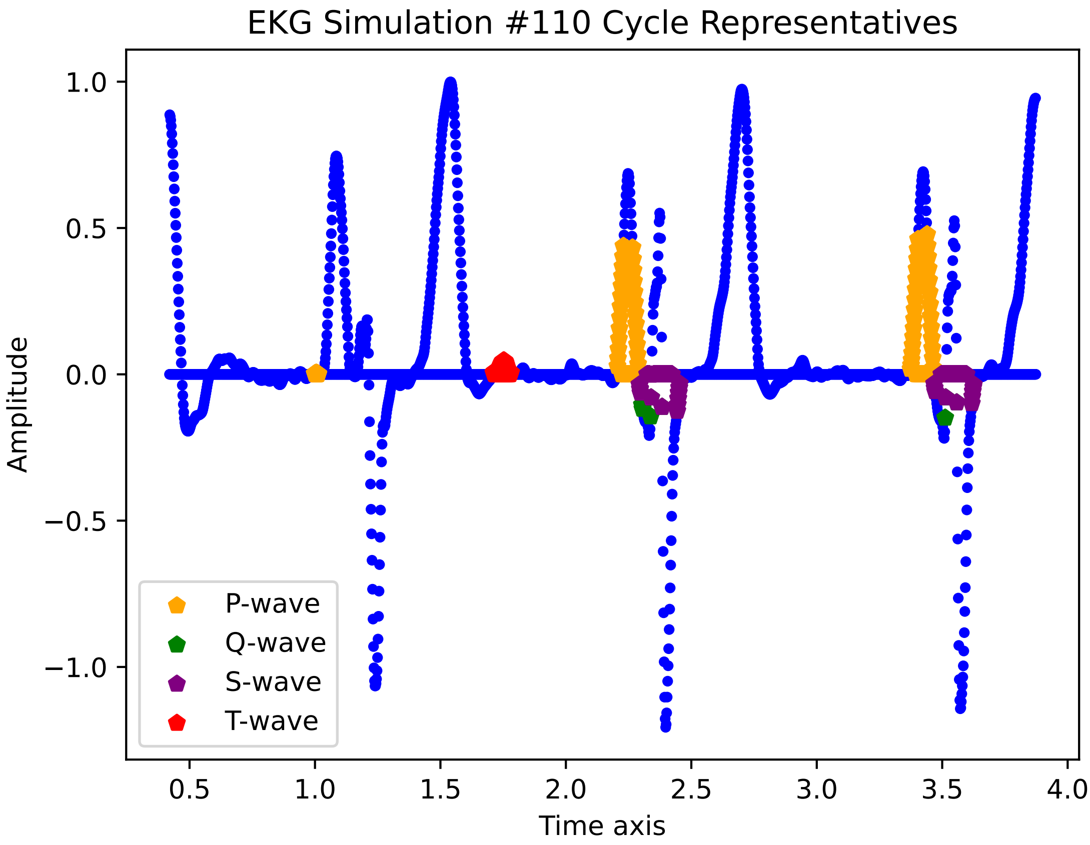
EKG Simulation #110 Cycle Representatives
1.0
0.5
0.0
−0.5
−1.0
Amplitude
0.5
1.0
1.5
2.0
2.5
3.0
3.5
4.0
Time axis
P-wave
Q-wave
S-wave
T-wave

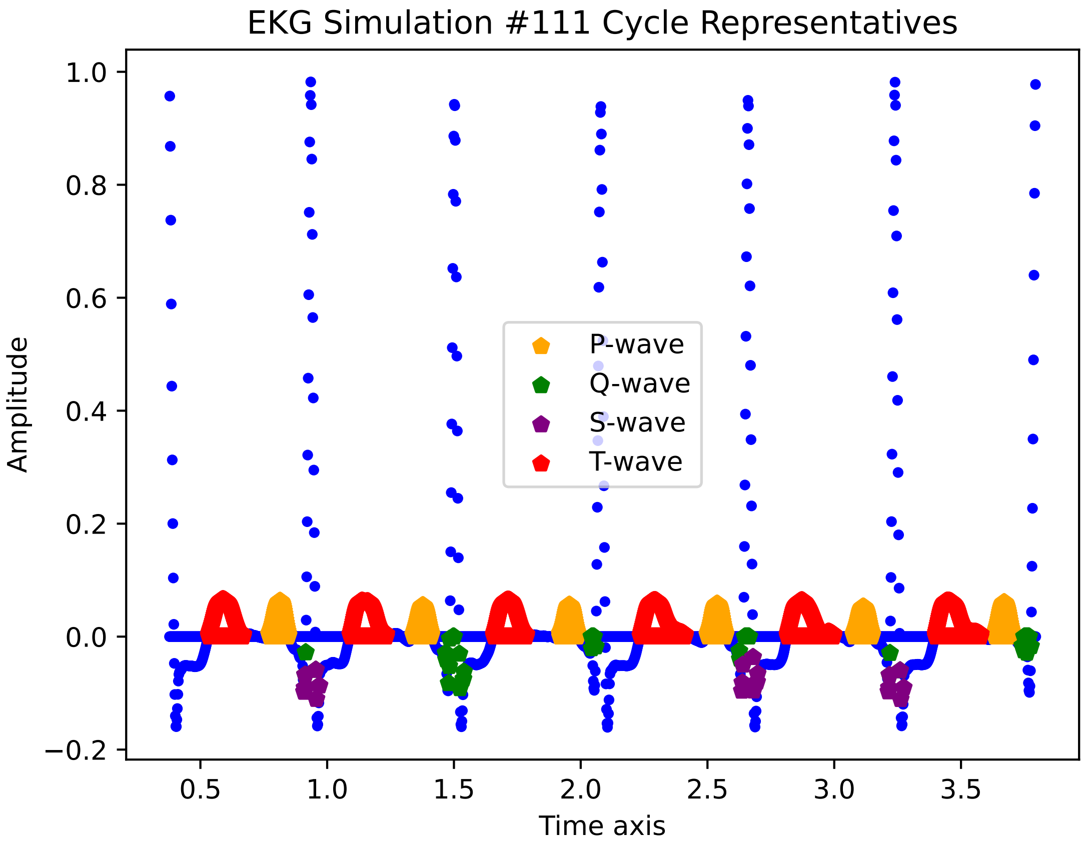

EKG Simulation #111 Cycle Representatives
P-wave
Q-wave
S-wave
T-wave
Amplitude
Time axis
1.0
0.8
0.6
0.4
0.2
0.0
−0.2
0.5
1.0
1.5
2.0
2.5
3.0
3.5

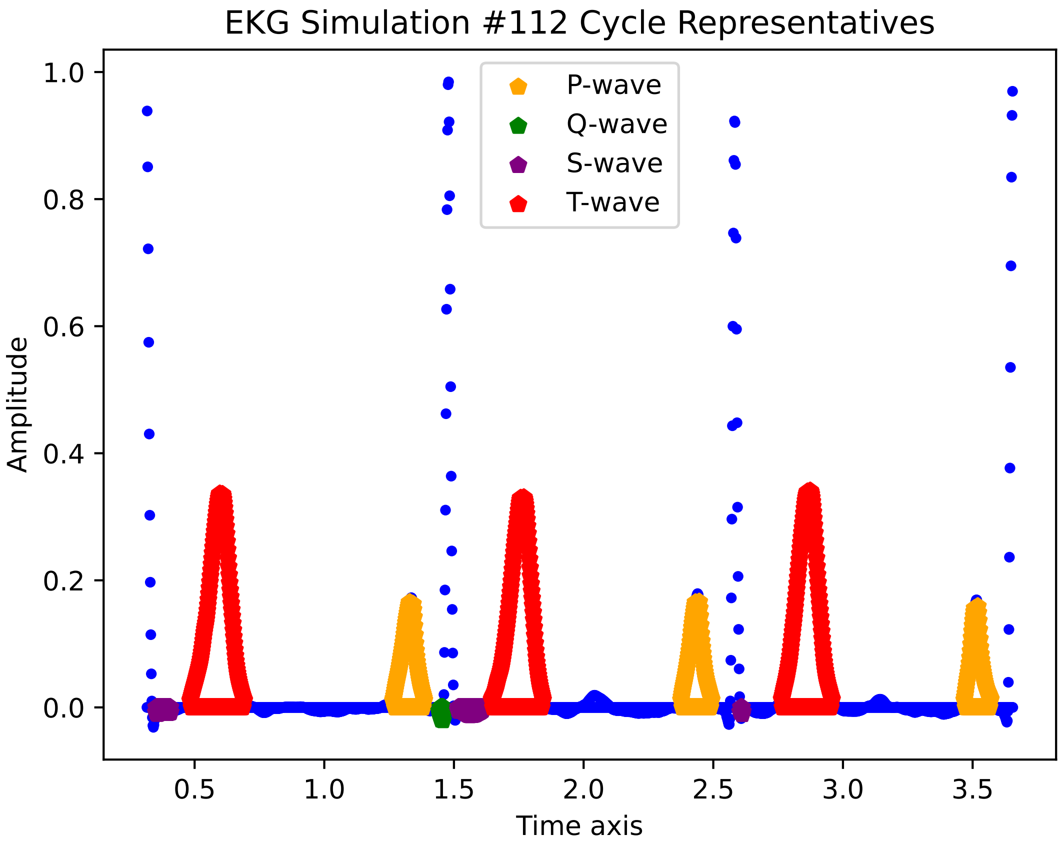
EKG Simulation #112 Cycle Representatives
P-wave
Q-wave
S-wave
T-wave
Amplitude
1.0
0.8
0.6
0.4
0.2
0.0
0.5
1.0
1.5
2.0
2.5
3.0
3.5
Time axis

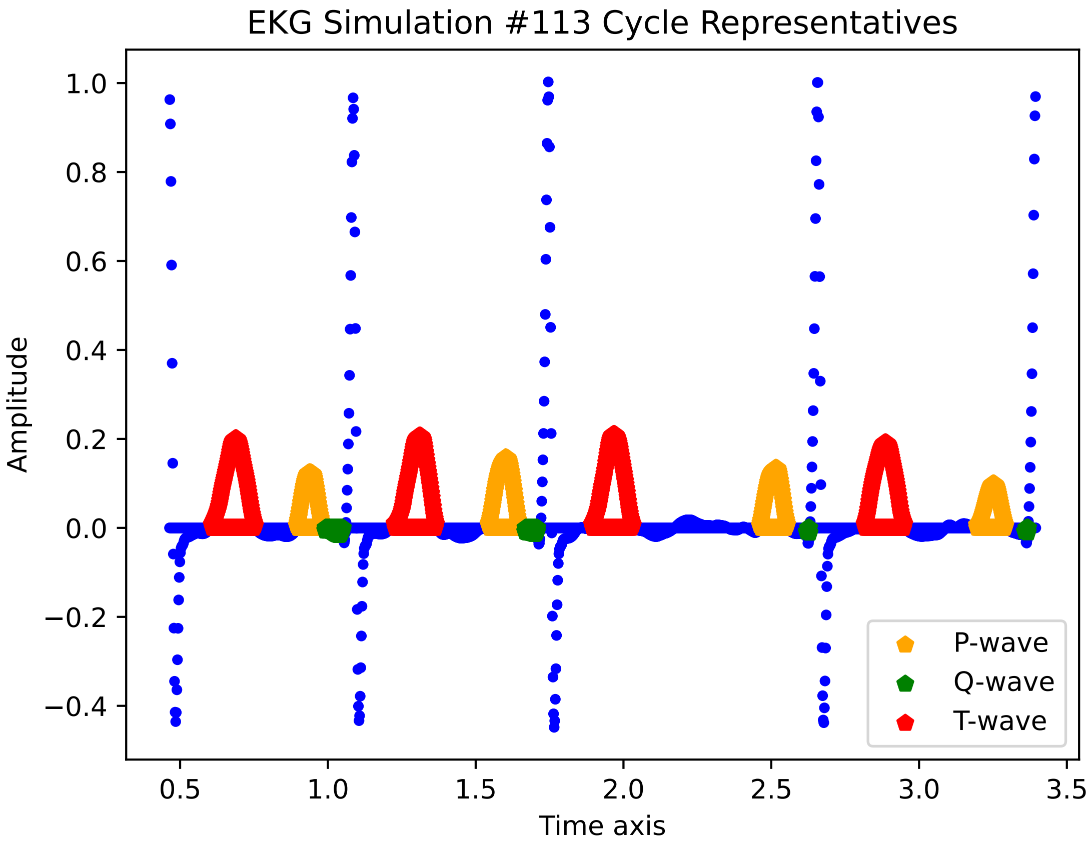
EKG Simulation #113 Cycle Representatives
Amplitude
Time axis
1.0
0.8
0.6
0.4
0.2
0.0
−0.2
−0.4
0.5
1.0
1.5
2.0
2.5
3.0
3.5
P-wave
Q-wave
T-wave

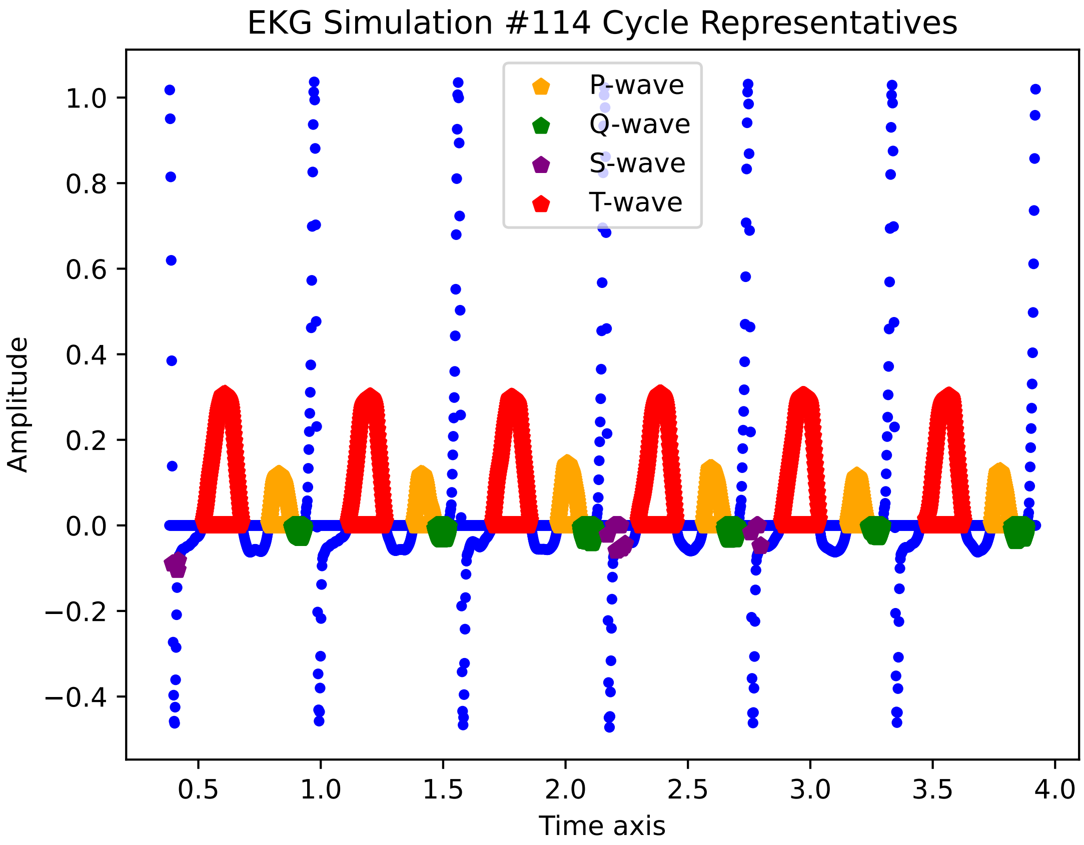

EKG Simulation #114 Cycle Representatives
P-wave
Q-wave
S-wave
T-wave
Amplitude
Time axis
1.0
0.8
0.6
0.4
0.2
0.0
−0.2
−0.4
0.5
1.0
1.5
2.0
2.5
3.0
3.5
4.0

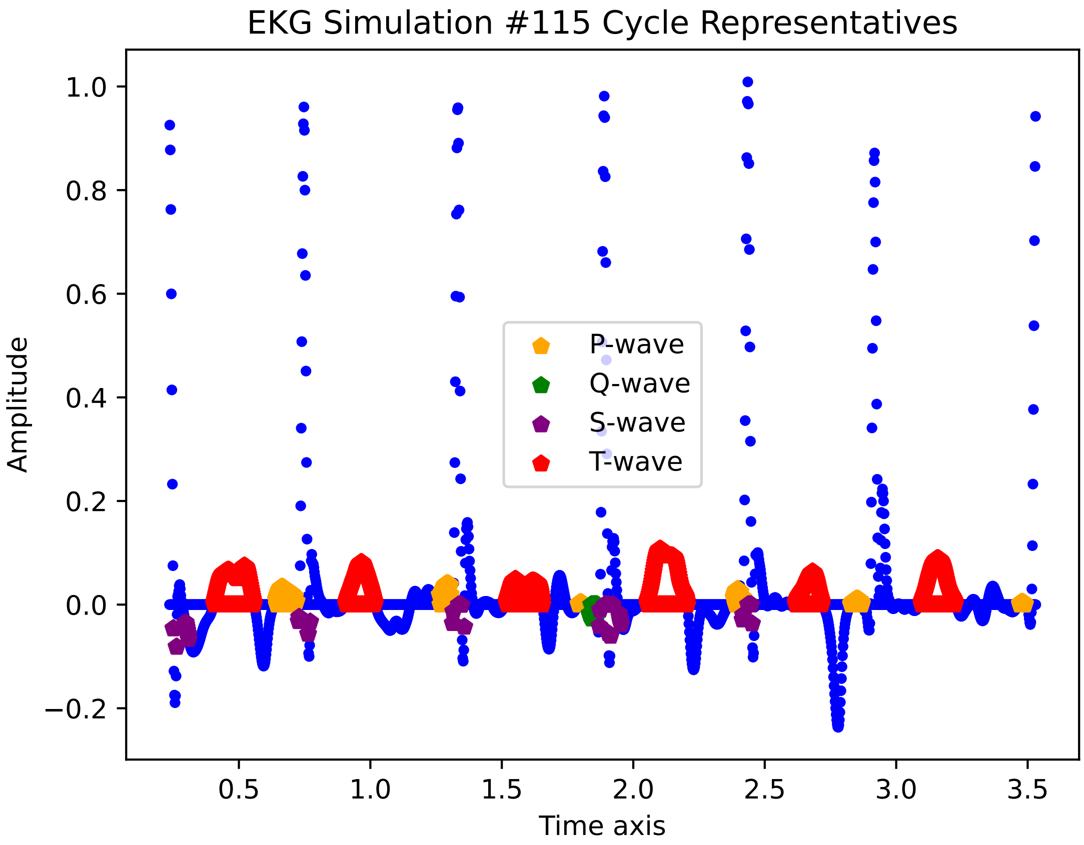

EKG Simulation #115 Cycle Representatives
Amplitude
Time axis
1.0
0.8
0.6
0.4
0.2
0.0
−0.2
0.5
1.0
1.5
2.0
2.5
3.0
3.5
P-wave
Q-wave
S-wave
T-wave

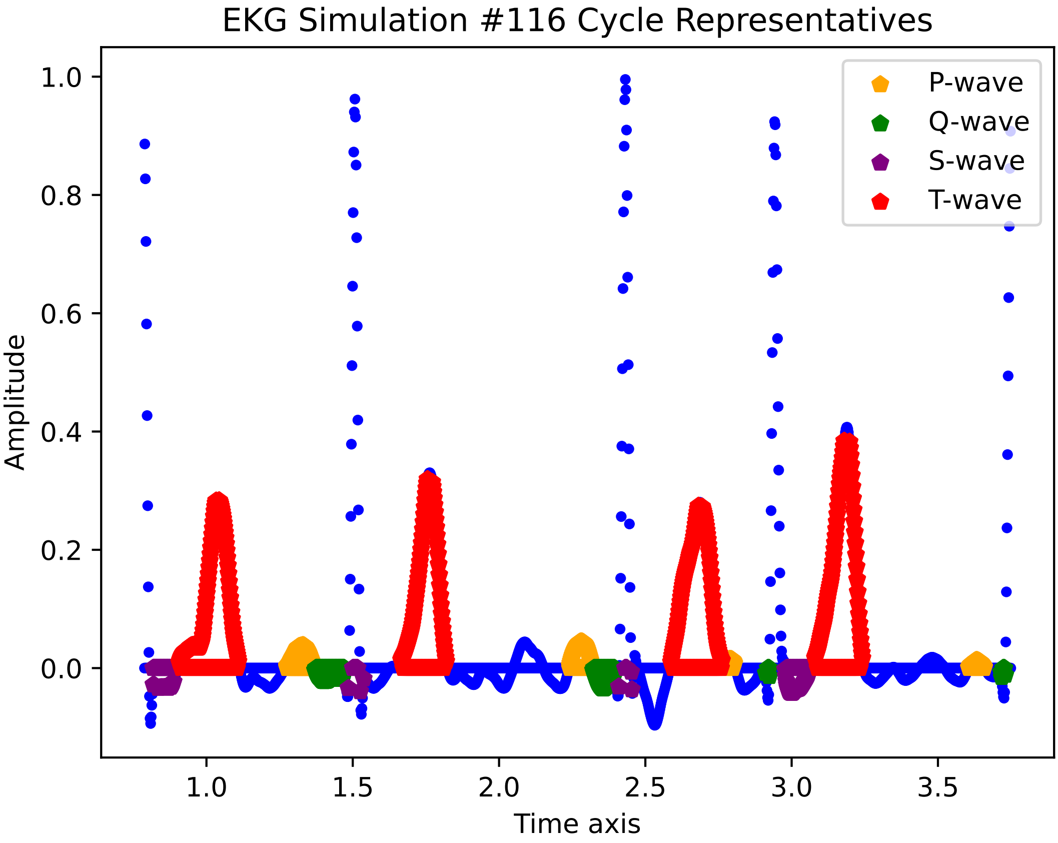

EKG Simulation #116 Cycle Representatives
1.0
0.8
0.6
0.4
0.2
0.0
Amplitude
1.0
1.5
2.0
2.5
3.0
3.5
Time axis
P-wave
Q-wave
S-wave
T-wave

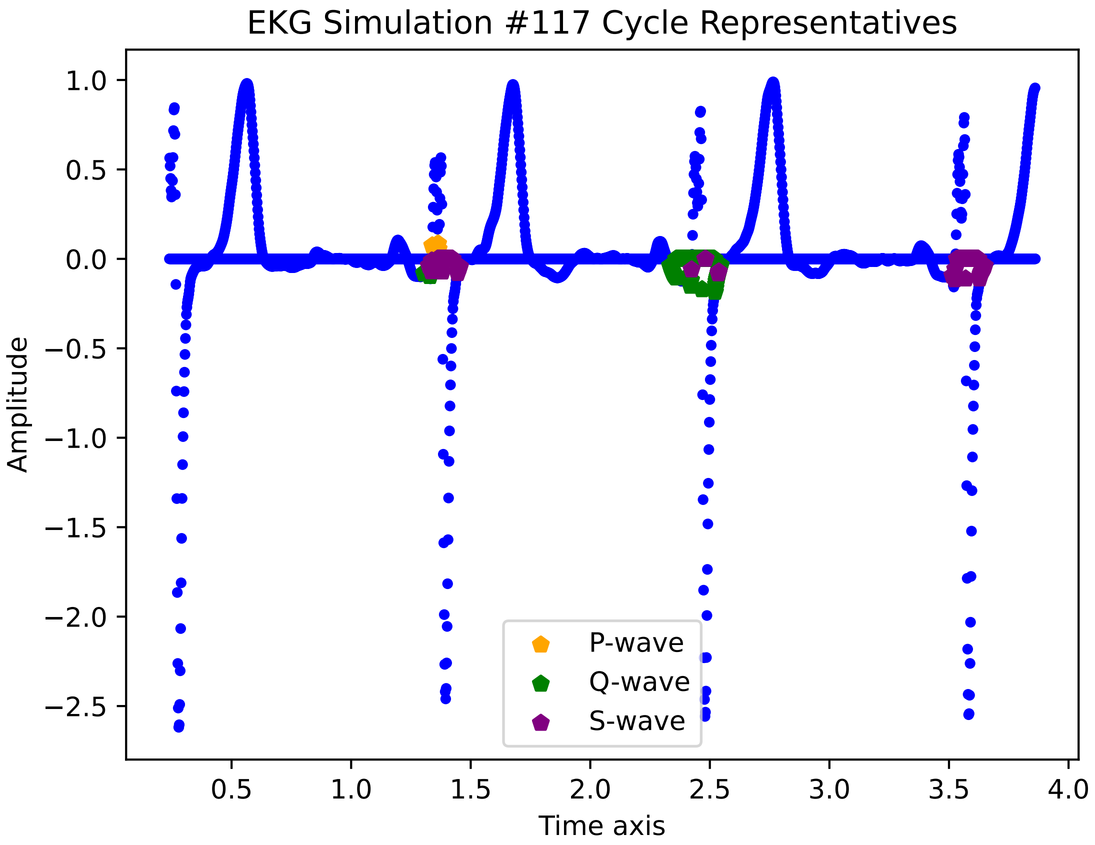
EKG Simulation #117 Cycle Representatives
1.0
0.5
0.0
−0.5
−1.0
−1.5
−2.0
−2.5
Amplitude
0.5
1.0
1.5
2.0
2.5
3.0
3.5
4.0
Time axis
P-wave
Q-wave
S-wave

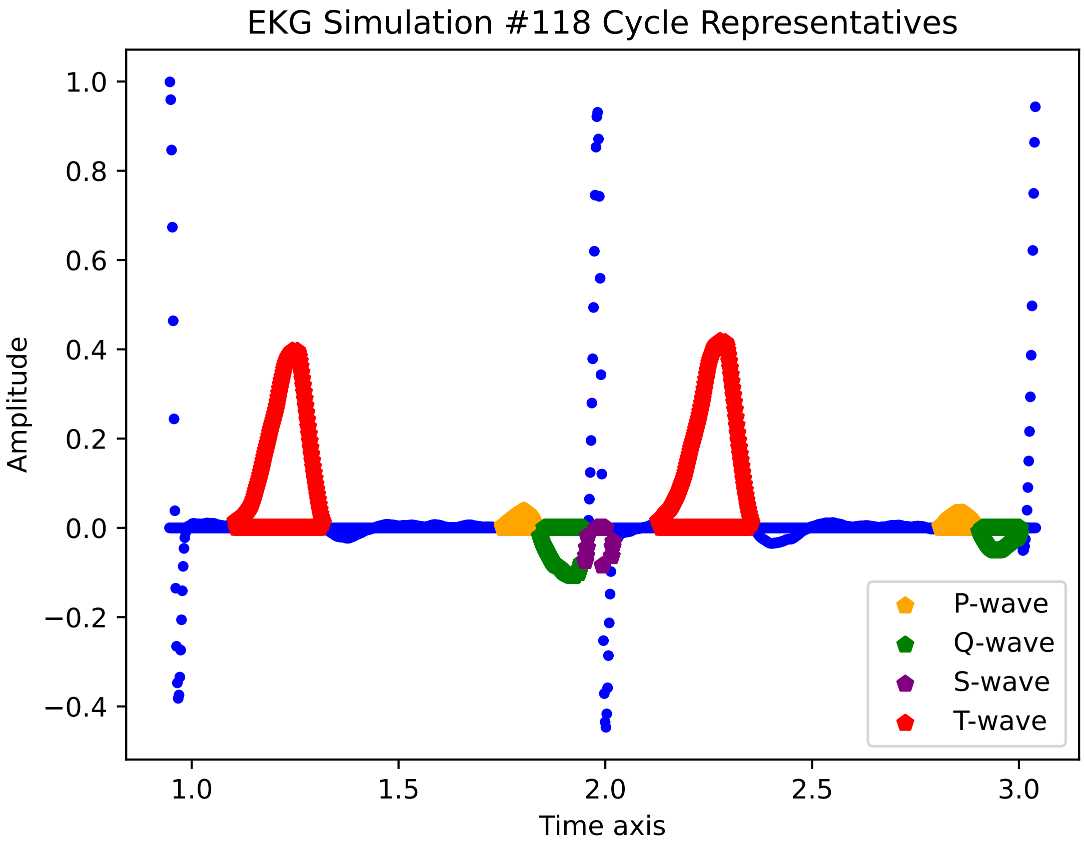

EKG Simulation #118 Cycle Representatives
Amplitude
Time axis
1.0
0.8
0.6
0.4
0.2
0.0
−0.2
−0.4
1.0
1.5
2.0
2.5
3.0
P-wave
Q-wave
S-wave
T-wave

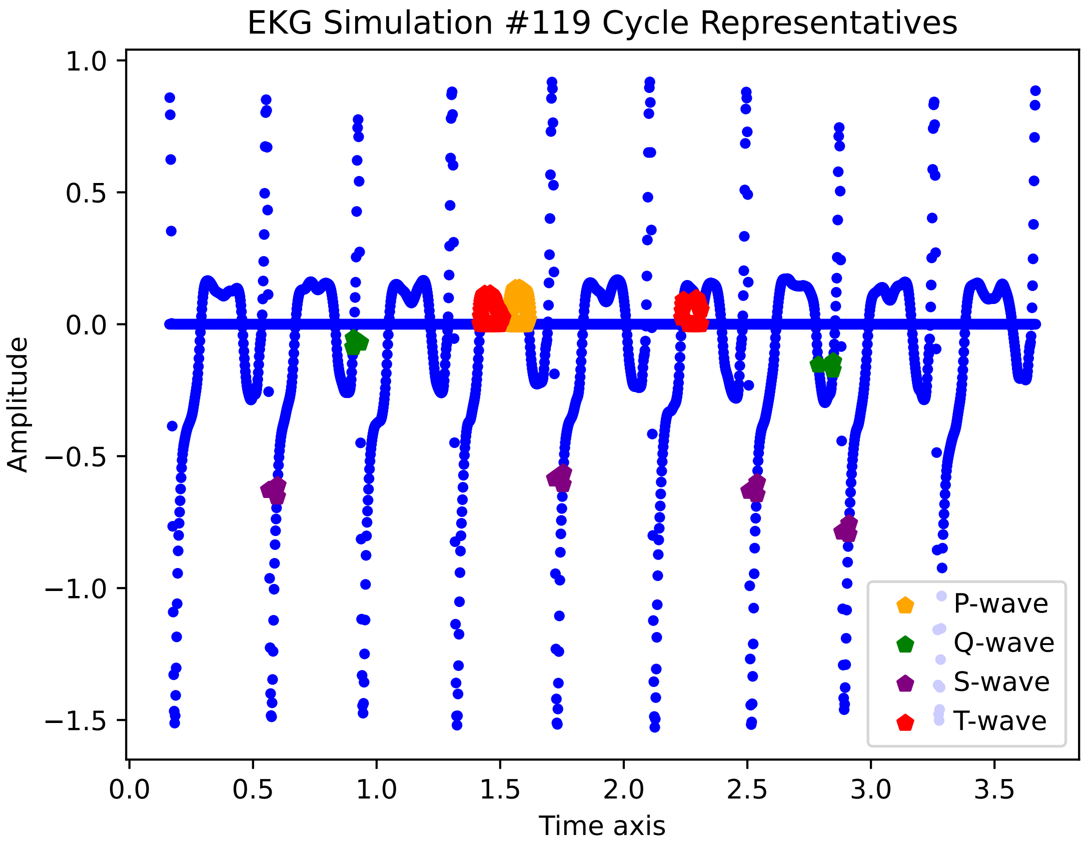
EKG Simulation #119 Cycle Representatives
Amplitude
Time axis
1.0
0.5
0.0
−0.5
−1.0
−1.5
0.0
0.5
1.0
1.5
2.0
2.5
3.0
3.5
P-wave
Q-wave
S-wave
T-wave

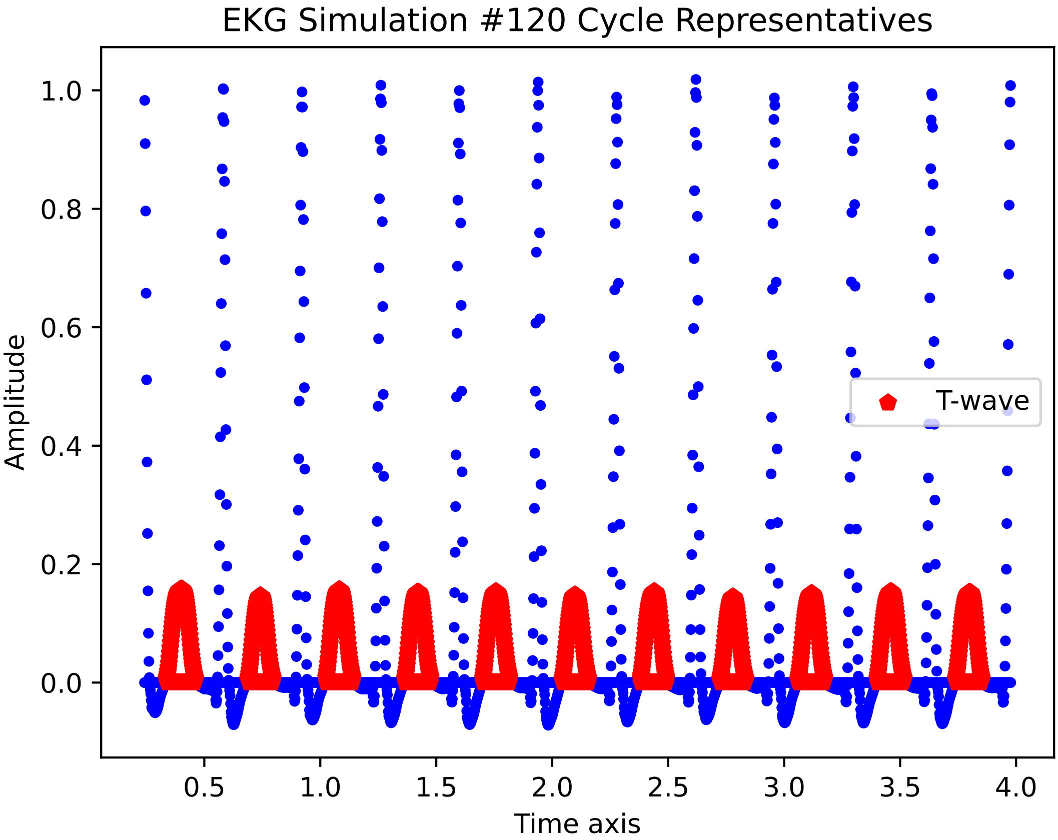

EKG Simulation #120 Cycle Representatives
Amplitude
Time axis
T-wave
1.0
0.8
0.6
0.4
0.2
0.0
0.5
1.0
1.5
2.0
2.5
3.0
3.5
4.0

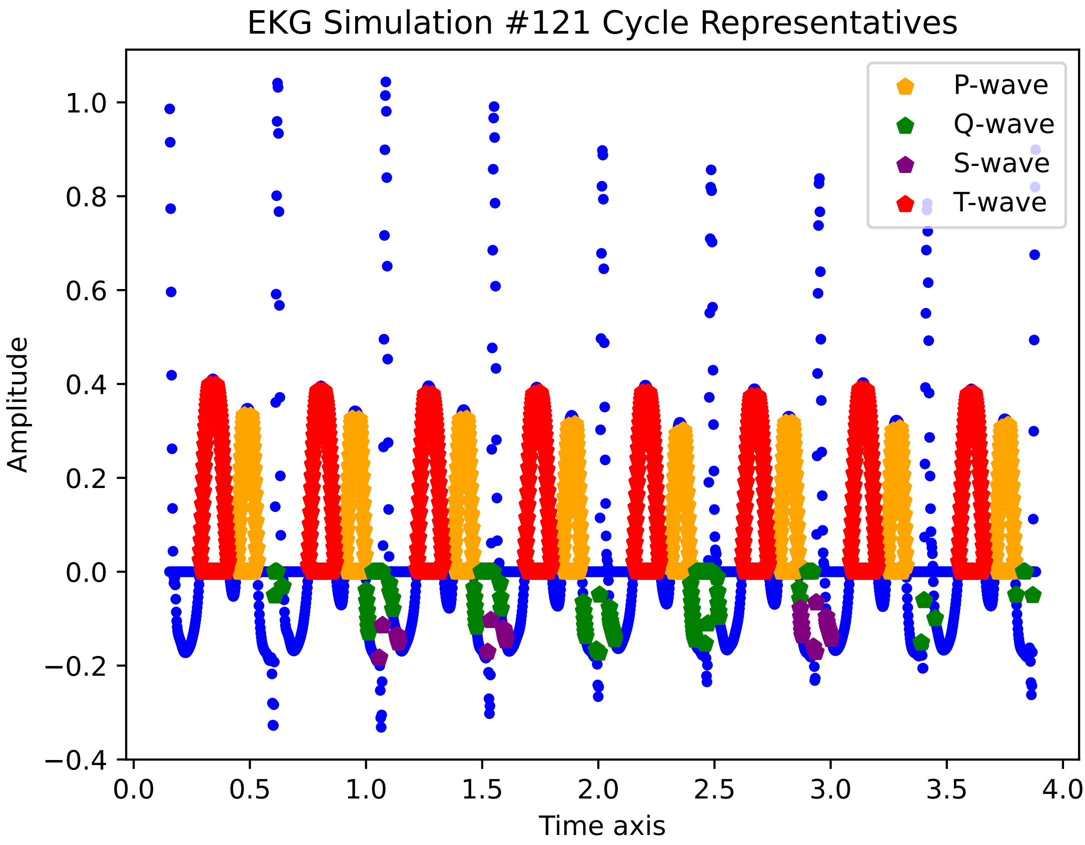
EKG Simulation #121 Cycle Representatives
P-wave
Q-wave
S-wave
T-wave
Amplitude
Time axis
1.0
0.8
0.6
0.4
0.2
0.0
−0.2
−0.4
0.0
0.5
1.0
1.5
2.0
2.5
3.0
3.5
4.0

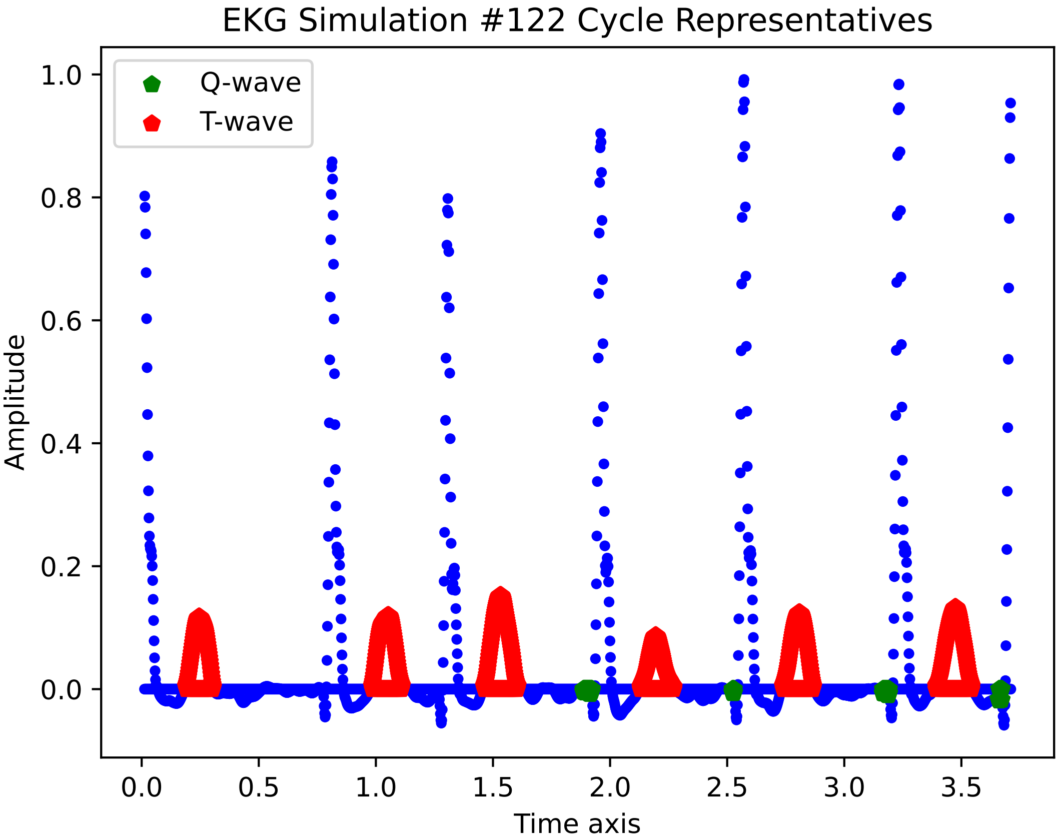

EKG Simulation #122 Cycle Representatives
Q-wave
T-wave
Amplitude
Time axis
1.0
0.8
0.6
0.4
0.2
0.0
0.0
0.5
1.0
1.5
2.0
2.5
3.0
3.5

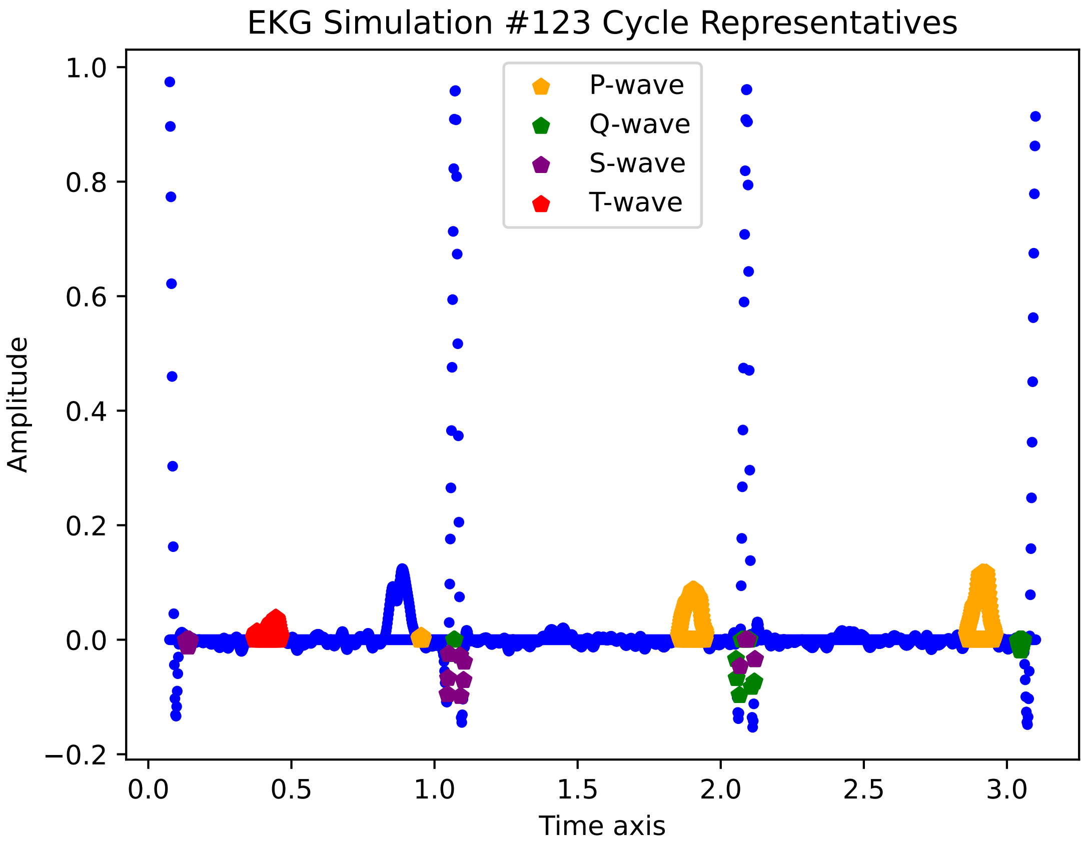

EKG Simulation #123 Cycle Representatives
P-wave
Q-wave
S-wave
T-wave
Amplitude
Time axis
1.0
0.8
0.6
0.4
0.2
0.0
−0.2
0.0
0.5
1.0
1.5
2.0
2.5
3.0

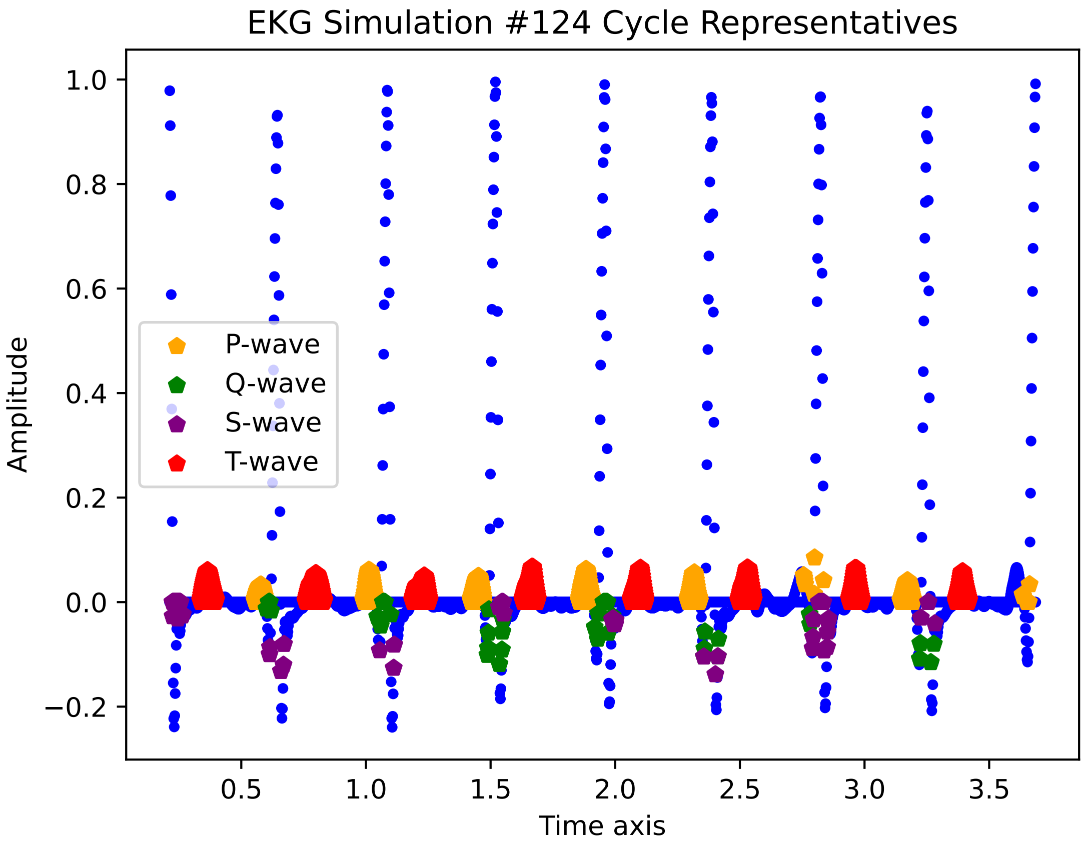
EKG Simulation #124 Cycle Representatives
Amplitude
Time axis
P-wave
Q-wave
S-wave
T-wave
1.0
0.8
0.6
0.4
0.2
0.0
−0.2
0.5
1.0
1.5
2.0
2.5
3.0
3.5

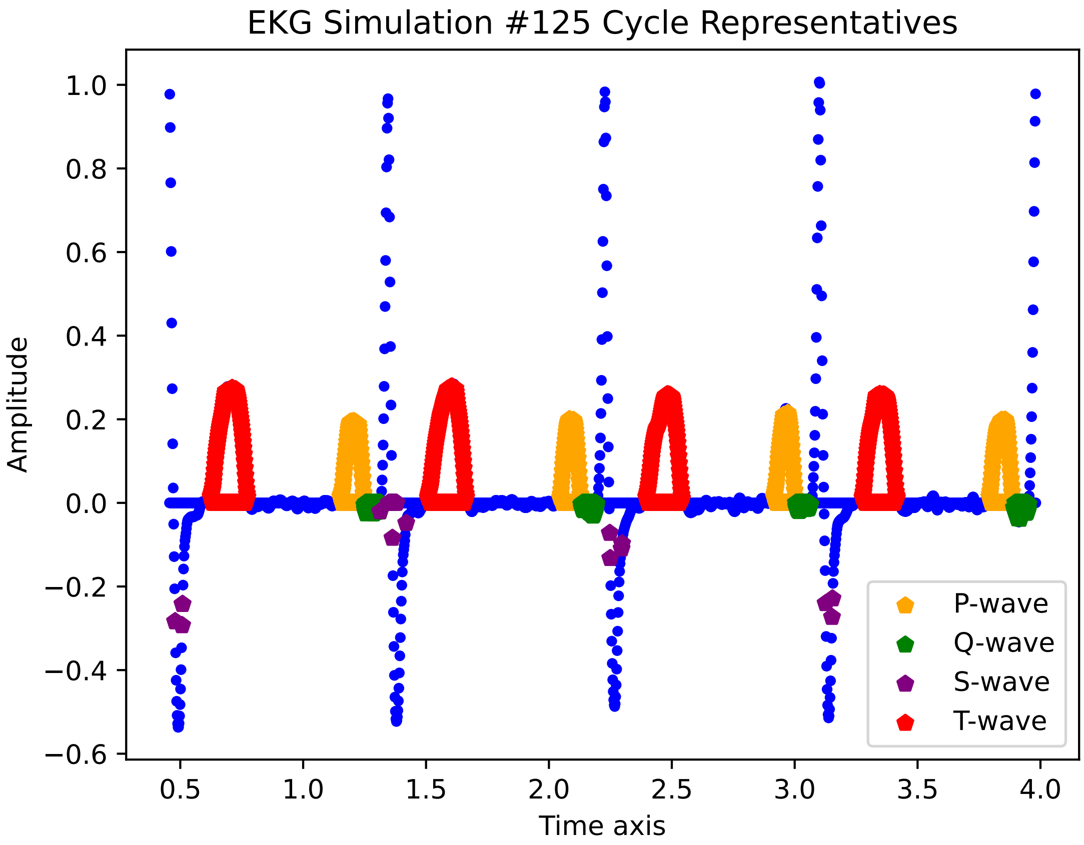
EKG Simulation #125 Cycle Representatives
Amplitude
Time axis
1.0
0.8
0.6
0.4
0.2
0.0
−0.2
−0.4
−0.6
0.5
1.0
1.5
2.0
2.5
3.0
3.5
4.0
P-wave
Q-wave
S-wave
T-wave

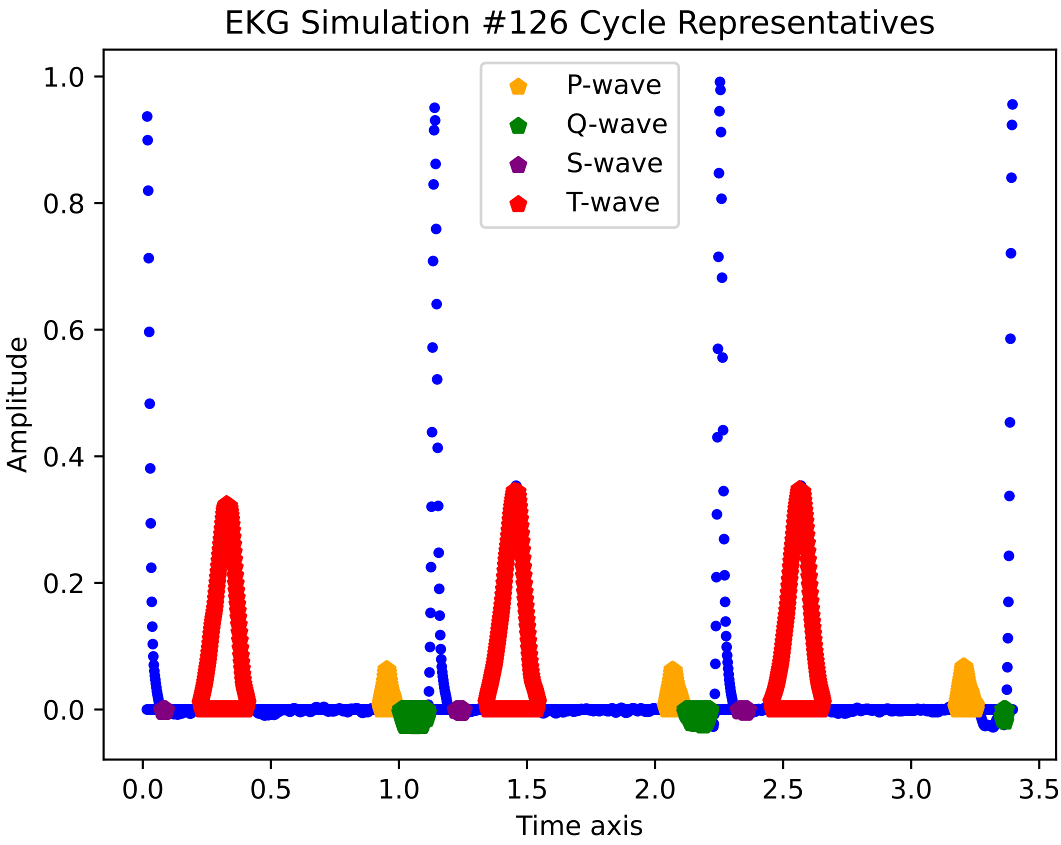

EKG Simulation #126 Cycle Representatives
P-wave
Q-wave
S-wave
T-wave
Amplitude
Time axis
1.0
0.8
0.6
0.4
0.2
0.0
0.0
0.5
1.0
1.5
2.0
2.5
3.0
3.5

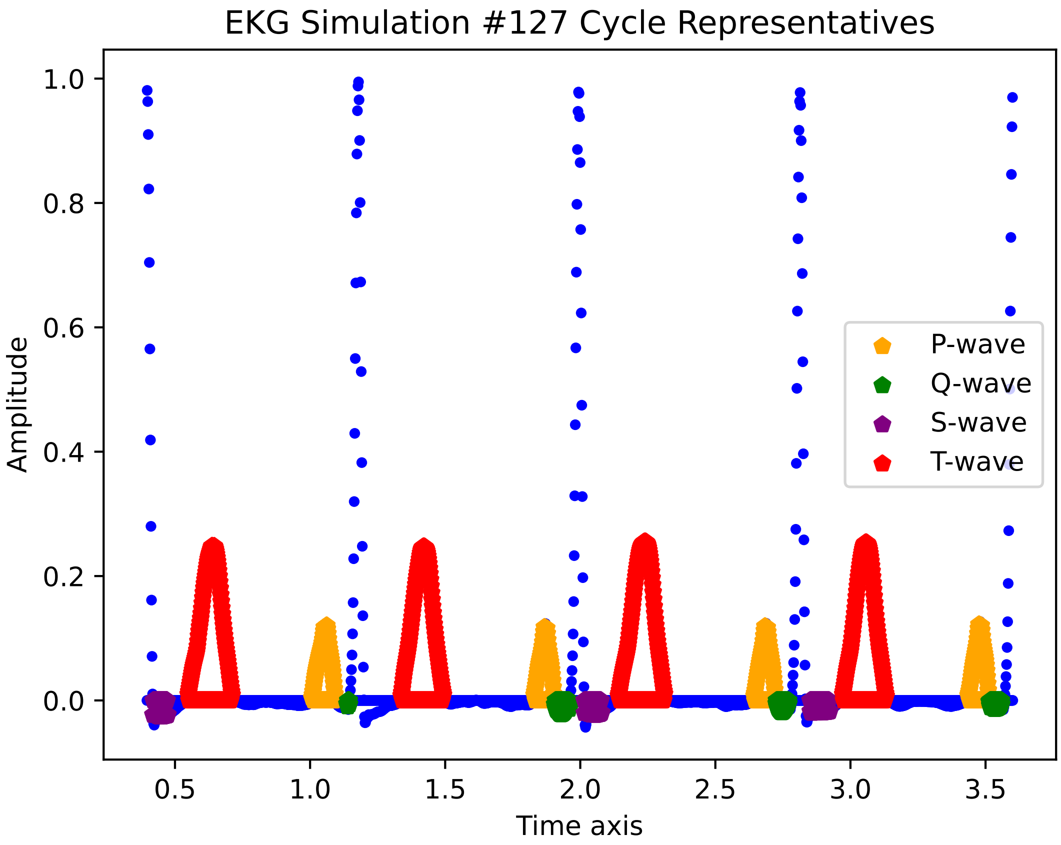
EKG Simulation #127 Cycle Representatives
Amplitude
Time axis
1.0
0.8
0.6
0.4
0.2
0.0
0.5
1.0
1.5
2.0
2.5
3.0
3.5
P-wave
Q-wave
S-wave
T-wave

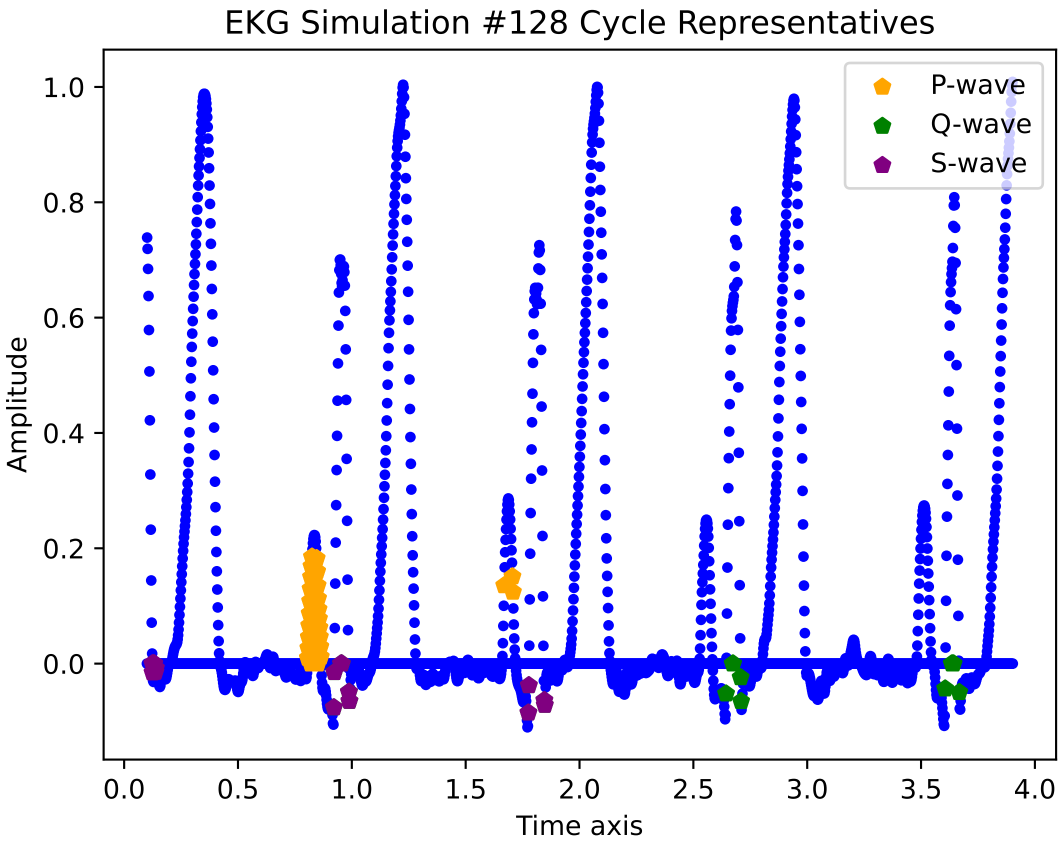
EKG Simulation #128 Cycle Representatives
P-wave
Q-wave
S-wave
Amplitude
Time axis
1.0
0.8
0.6
0.4
0.2
0.0
0.0
0.5
1.0
1.5
2.0
2.5
3.0
3.5
4.0

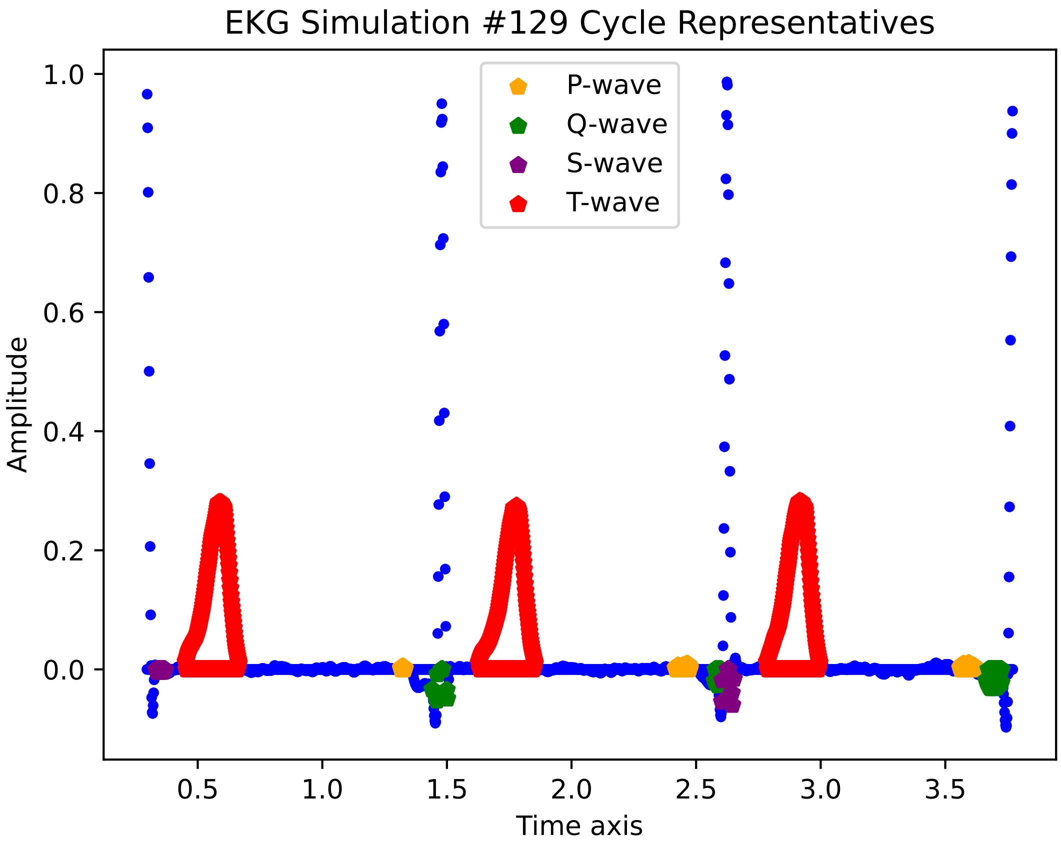
EKG Simulation #129 Cycle Representatives
P-wave
Q-wave
S-wave
T-wave
Amplitude
Time axis
1.0
0.8
0.6
0.4
0.2
0.0
0.5
1.0
1.5
2.0
2.5
3.0
3.5

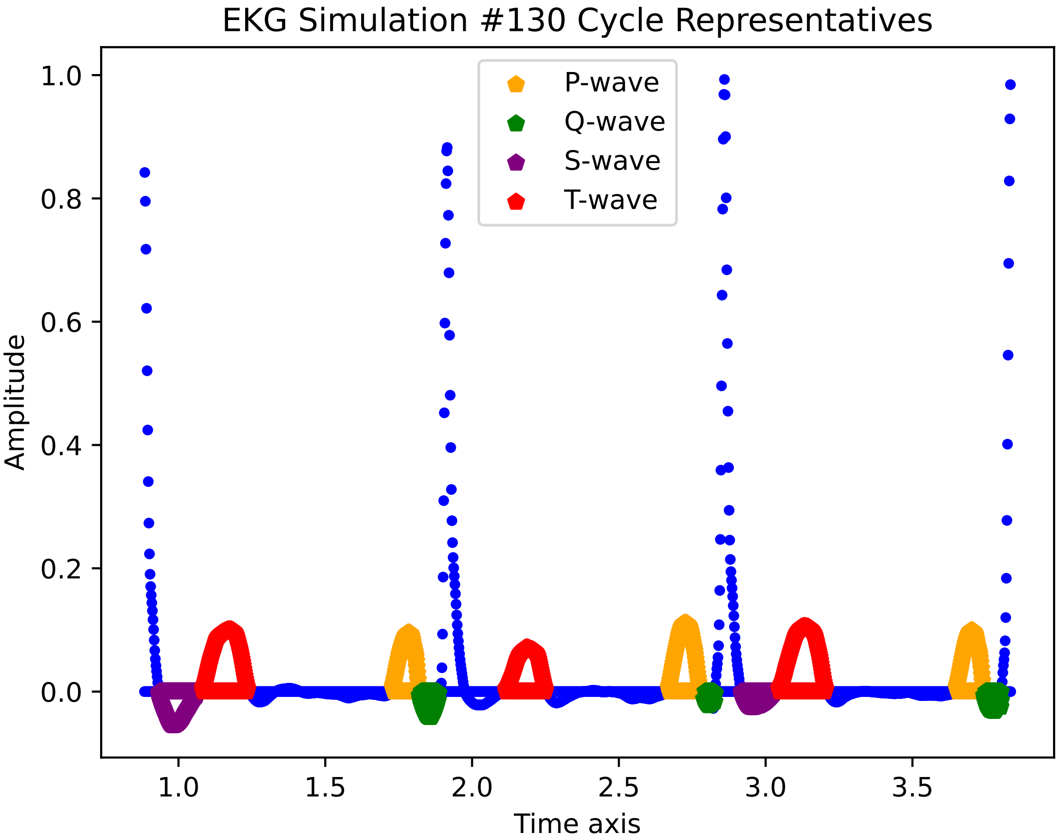
EKG Simulation #130 Cycle Representatives
P-wave
Q-wave
S-wave
T-wave
Amplitude
Time axis
1.0
0.8
0.6
0.4
0.2
0.0
1.0
1.5
2.0
2.5
3.0
3.5

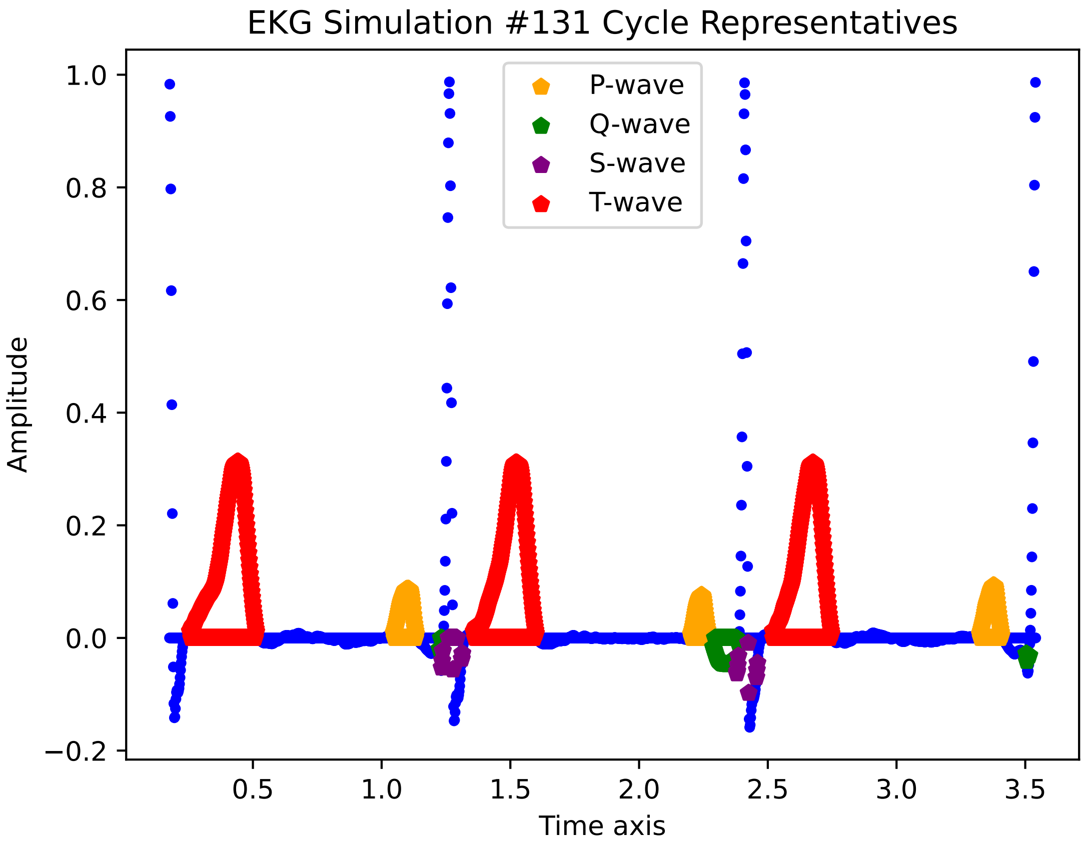

EKG Simulation #131 Cycle Representatives
P-wave
Q-wave
S-wave
T-wave
Amplitude
1.0
0.8
0.6
0.4
0.2
0.0
−0.2
0.5
1.0
1.5
2.0
2.5
3.0
3.5
Time axis

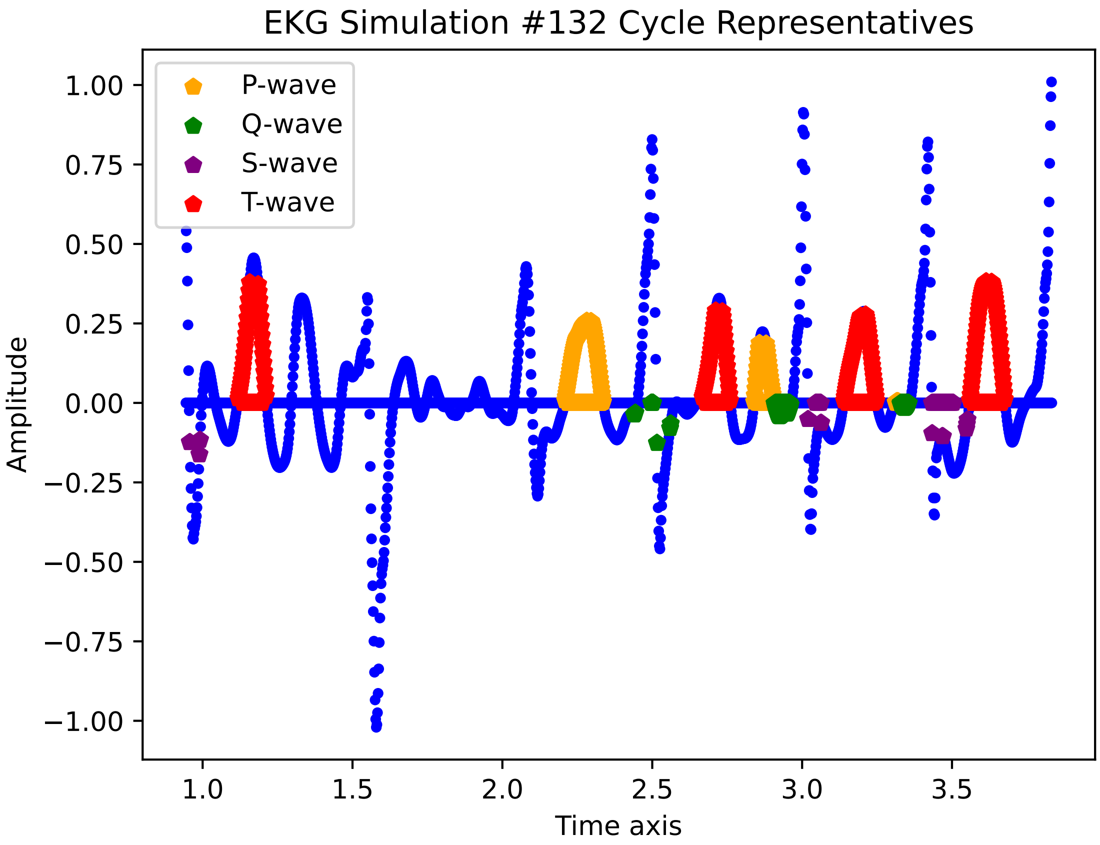

EKG Simulation #132 Cycle Representatives
P-wave
Q-wave
S-wave
T-wave
1.00
0.75
0.50
0.25
0.00
−0.25
−0.50
−0.75
−1.00
Amplitude
1.0
1.5
2.0
2.5
3.0
3.5
Time axis

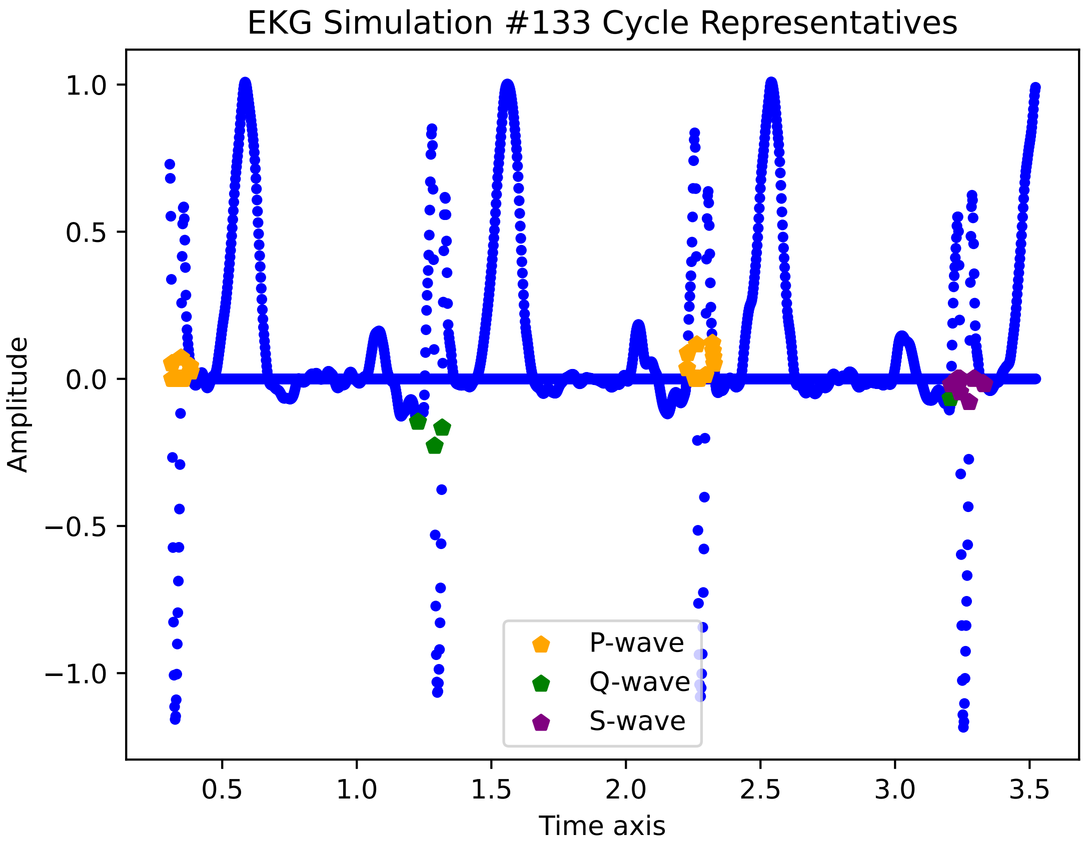
EKG Simulation #133 Cycle Representatives
Amplitude
Time axis
1.0
0.5
0.0
−0.5
−1.0
0.5
1.0
1.5
2.0
2.5
3.0
3.5
P-wave
Q-wave
S-wave

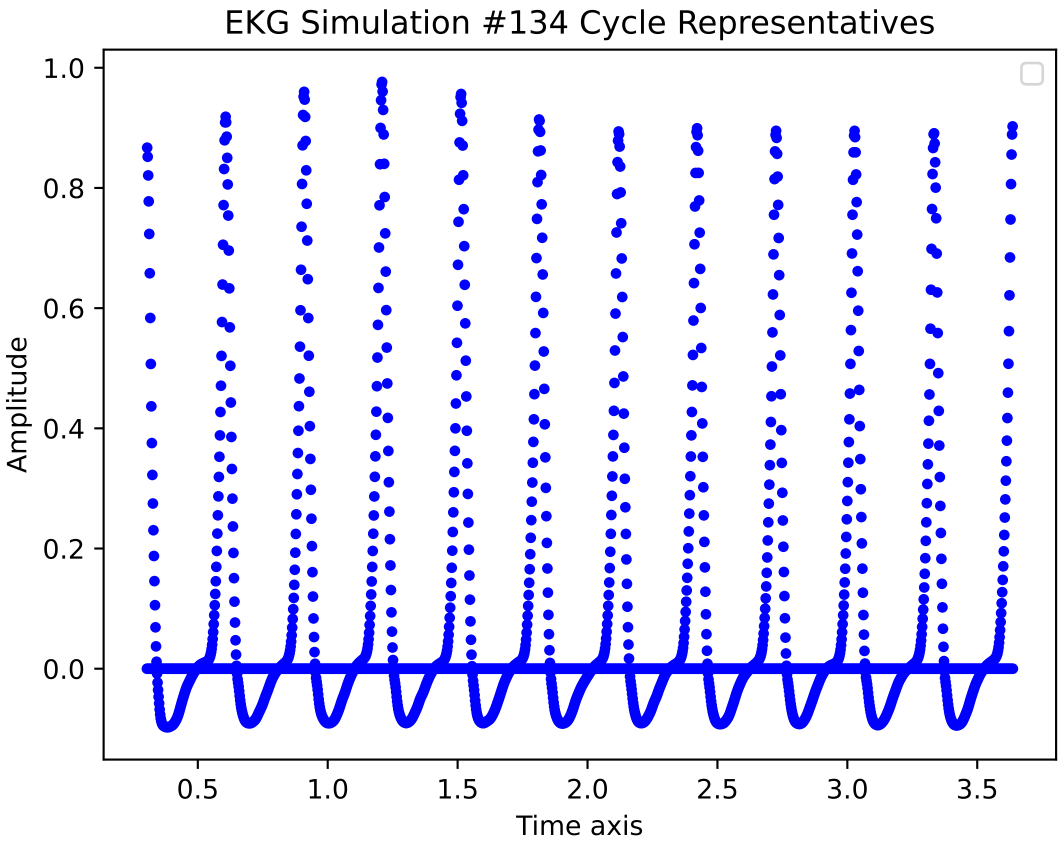
EKG Simulation #134 Cycle Representatives
Amplitude
Time axis
1.0
0.8
0.6
0.4
0.2
0.0
0.5
1.0
1.5
2.0
2.5
3.0
3.5

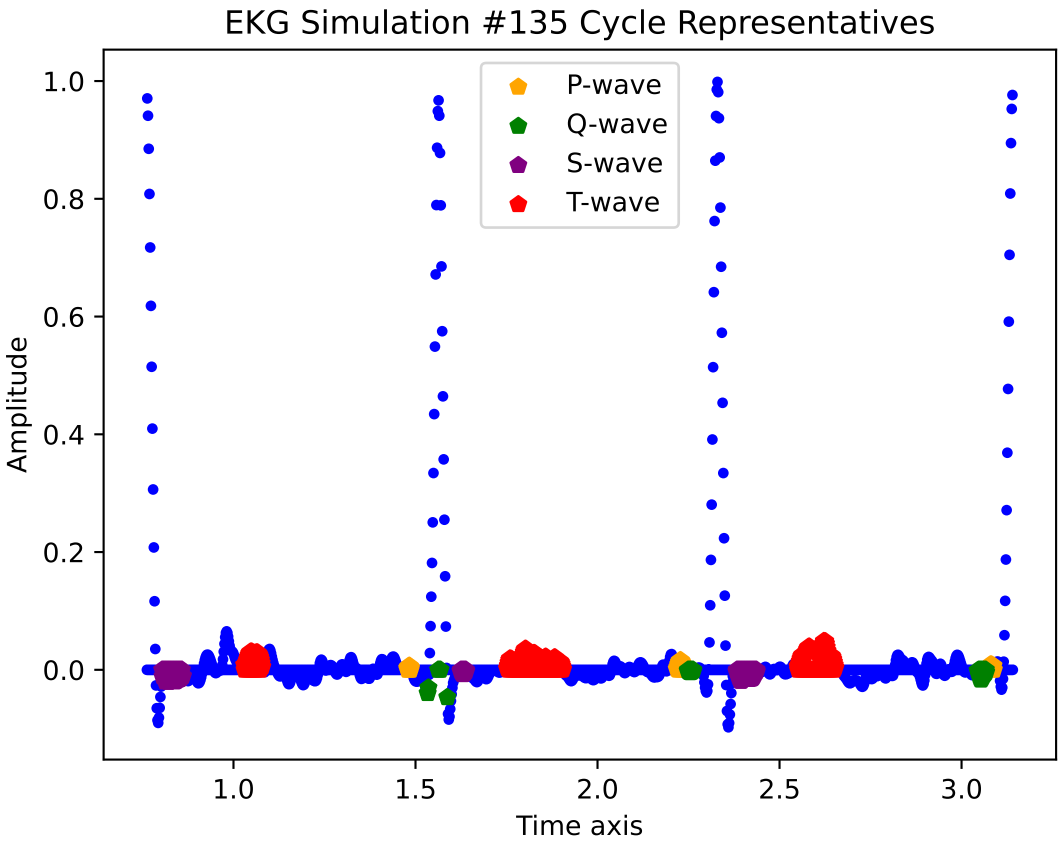
EKG Simulation #135 Cycle Representatives
P-wave
Q-wave
S-wave
T-wave
Amplitude
Time axis
1.0
0.8
0.6
0.4
0.2
0.0
1.0
1.5
2.0
2.5
3.0

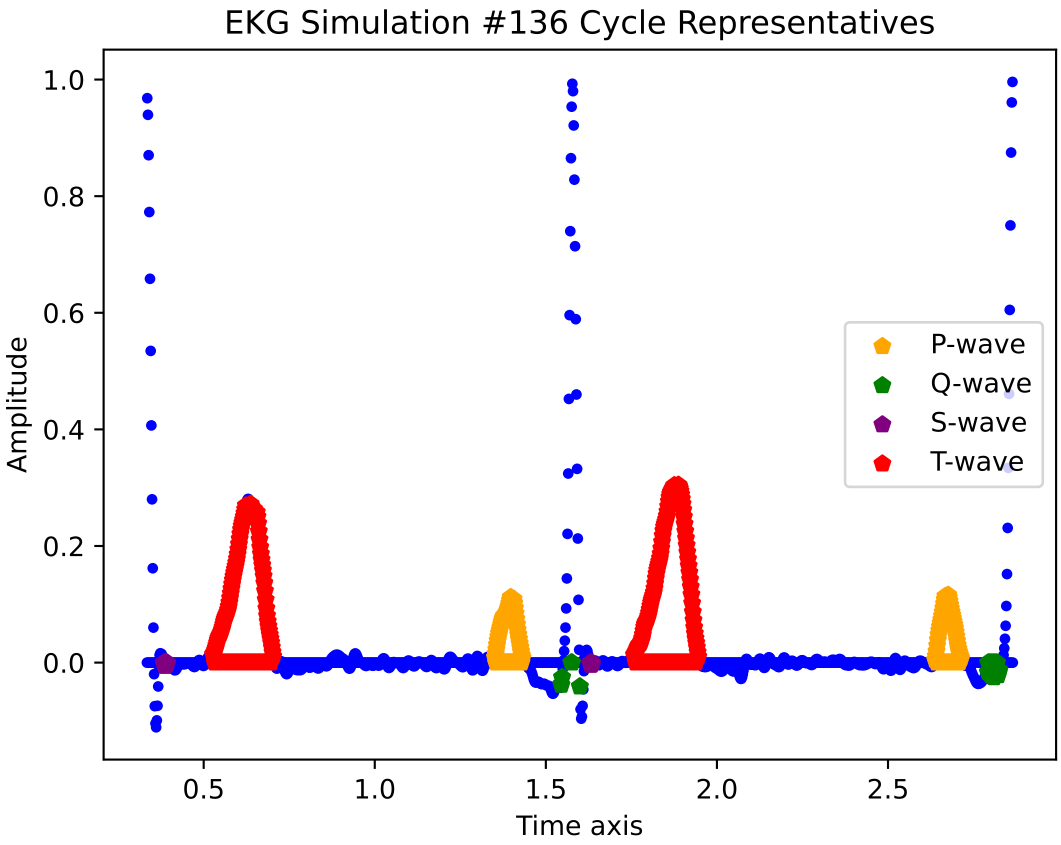

EKG Simulation #136 Cycle Representatives
Amplitude
Time axis
P-wave
Q-wave
S-wave
T-wave
1.0
0.8
0.6
0.4
0.2
0.0
0.5
1.0
1.5
2.0
2.5

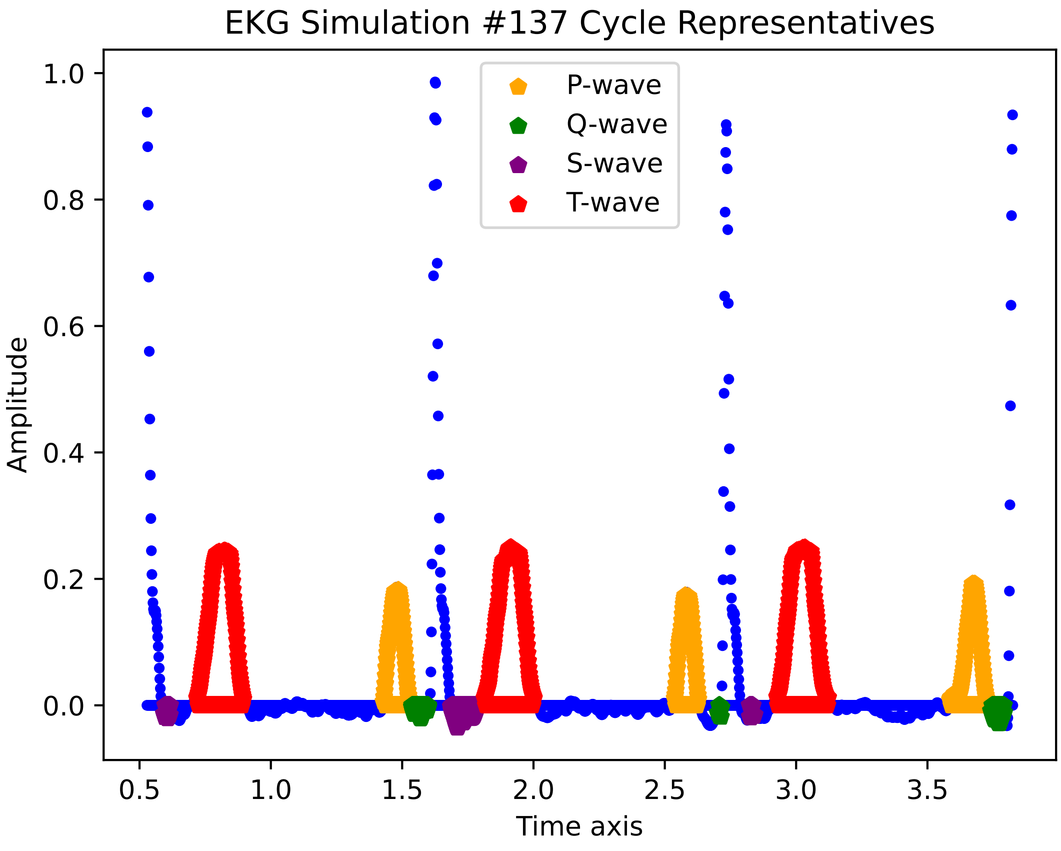

EKG Simulation #137 Cycle Representatives
P-wave
Q-wave
S-wave
T-wave
Amplitude
1.0
0.8
0.6
0.4
0.2
0.0
0.5
1.0
1.5
2.0
2.5
3.0
3.5
Time axis

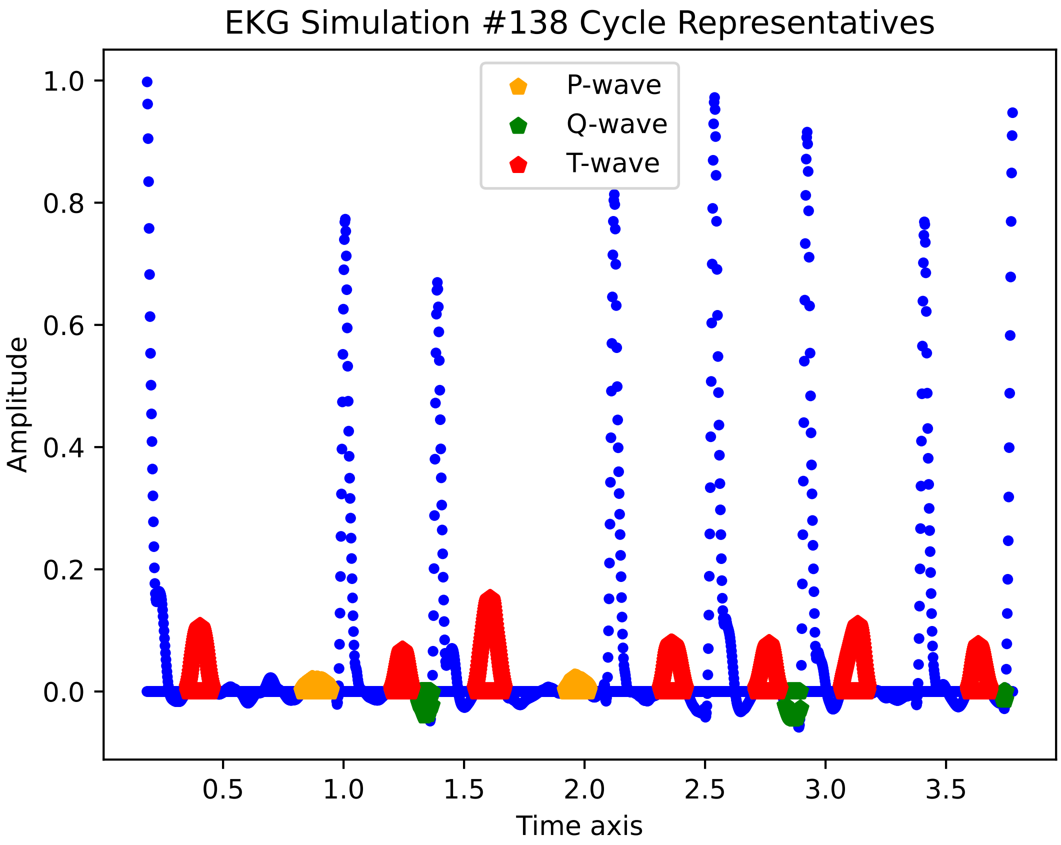

EKG Simulation #138 Cycle Representatives
P-wave
Q-wave
T-wave
Amplitude
Time axis
1.0
0.8
0.6
0.4
0.2
0.0
0.5
1.0
1.5
2.0
2.5
3.0
3.5

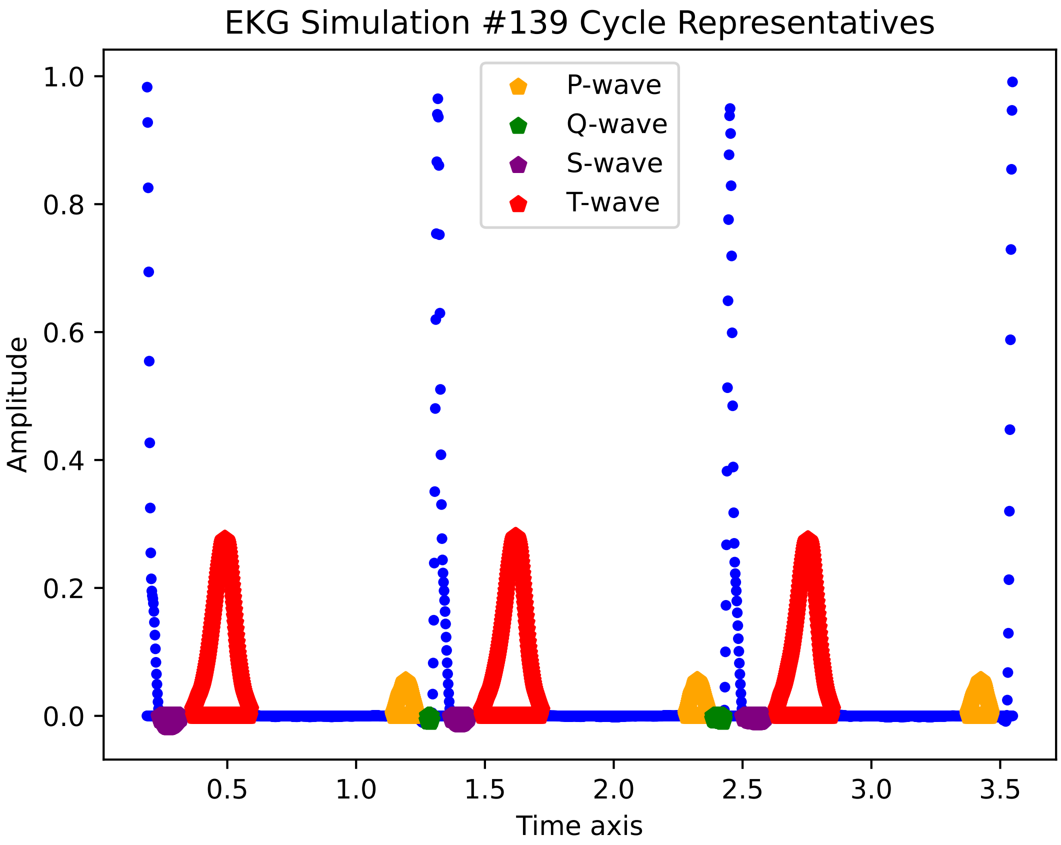

EKG Simulation #139 Cycle Representatives
P-wave
Q-wave
S-wave
T-wave
Amplitude
Time axis
1.0
0.8
0.6
0.4
0.2
0.0
0.5
1.0
1.5
2.0
2.5
3.0
3.5

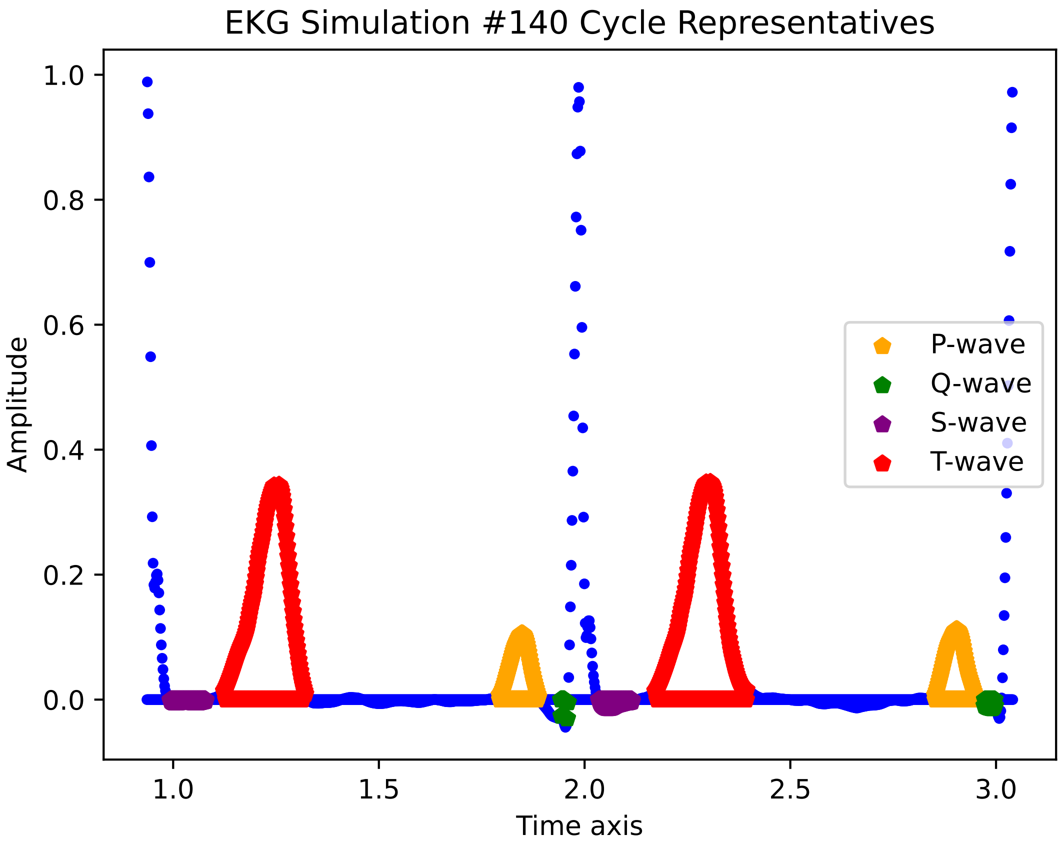
EKG Simulation #140 Cycle Representatives
Amplitude
Time axis
1.0
0.8
0.6
0.4
0.2
0.0
1.0
1.5
2.0
2.5
3.0
P-wave
Q-wave
S-wave
T-wave

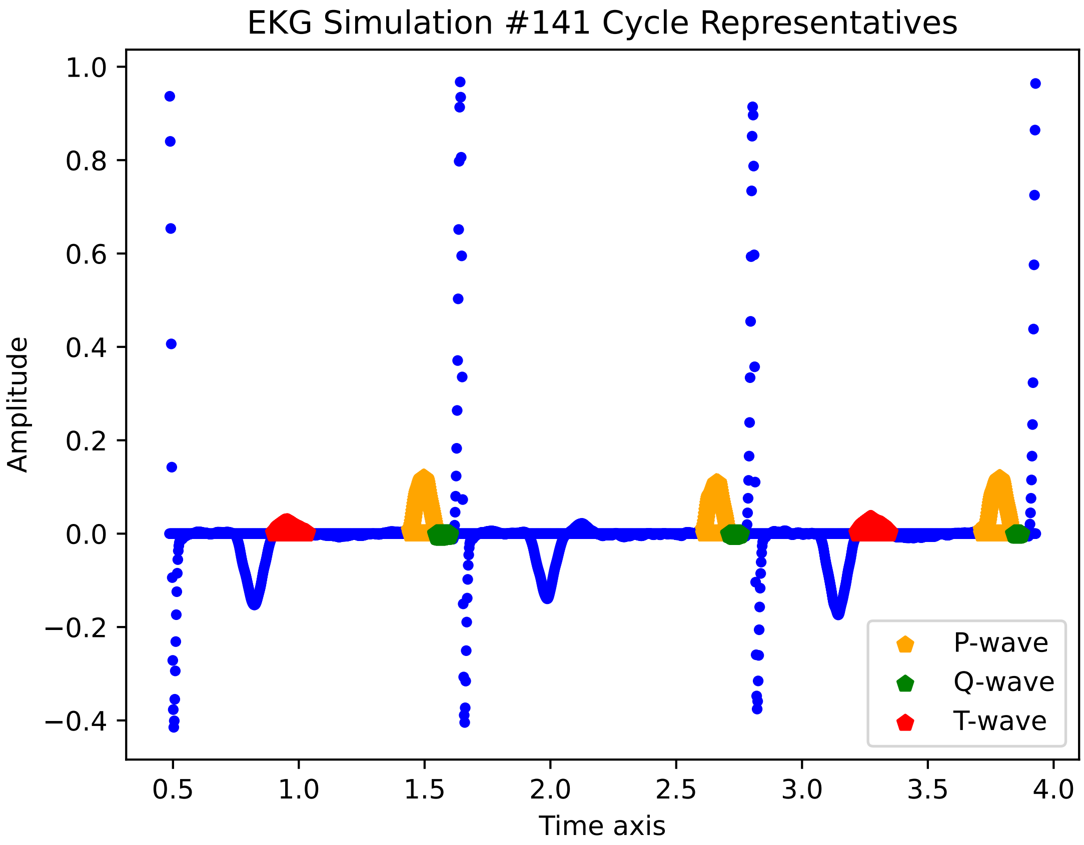
EKG Simulation #141 Cycle Representatives
Amplitude
Time axis
1.0
0.8
0.6
0.4
0.2
0.0
−0.2
−0.4
0.5
1.0
1.5
2.0
2.5
3.0
3.5
4.0
P-wave
Q-wave
T-wave

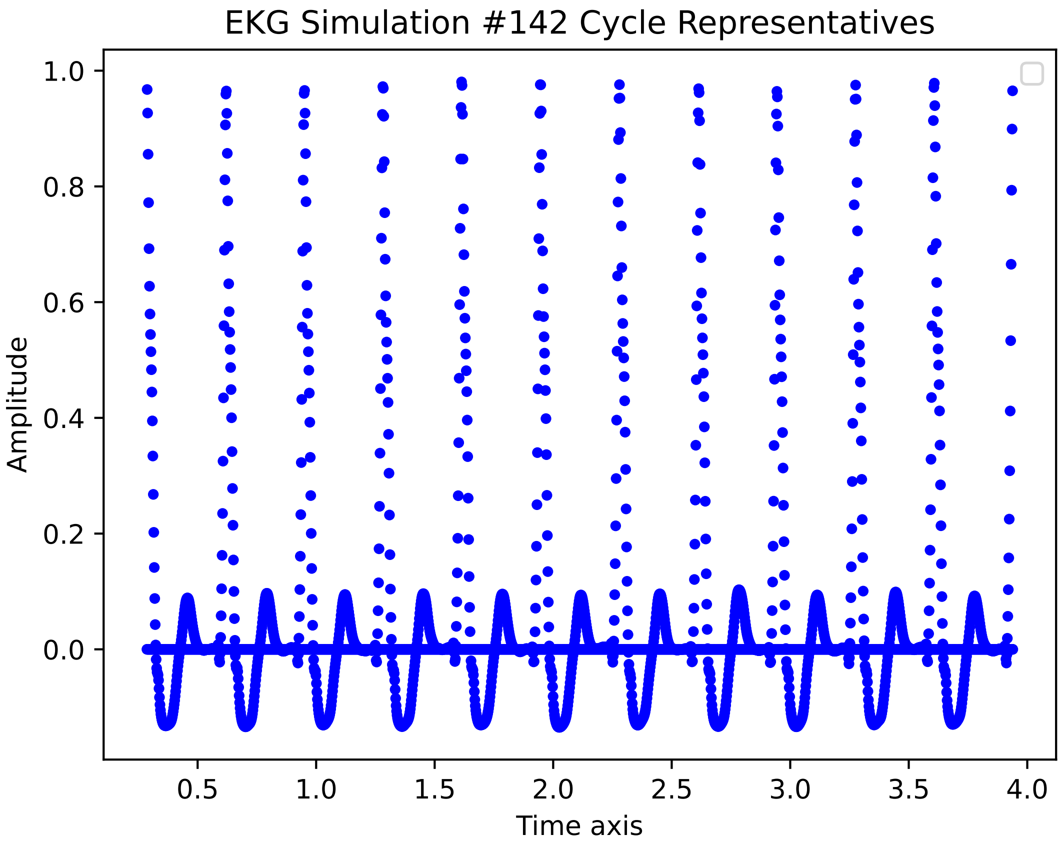
EKG Simulation #142 Cycle Representatives
Amplitude
Time axis
1.0
0.8
0.6
0.4
0.2
0.0
0.5
1.0
1.5
2.0
2.5
3.0
3.5
4.0

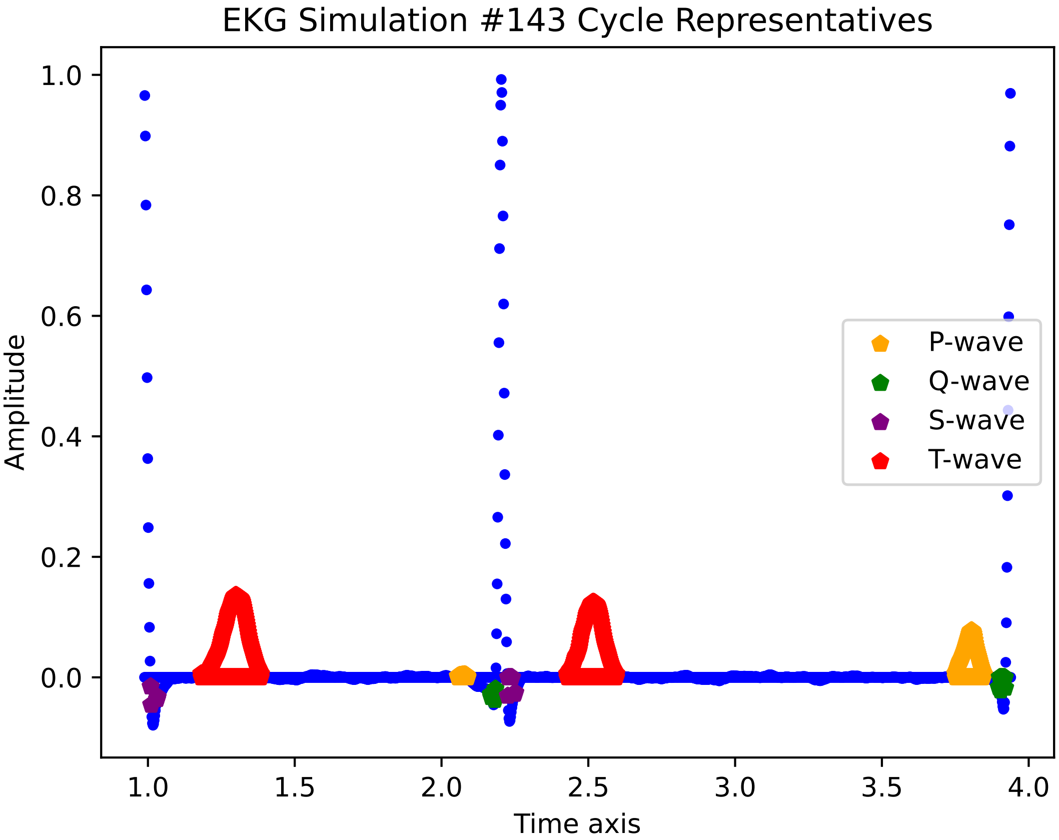

EKG Simulation #143 Cycle Representatives
Amplitude
Time axis
1.0
0.8
0.6
0.4
0.2
0.0
1.0
1.5
2.0
2.5
3.0
3.5
4.0
P-wave
Q-wave
S-wave
T-wave

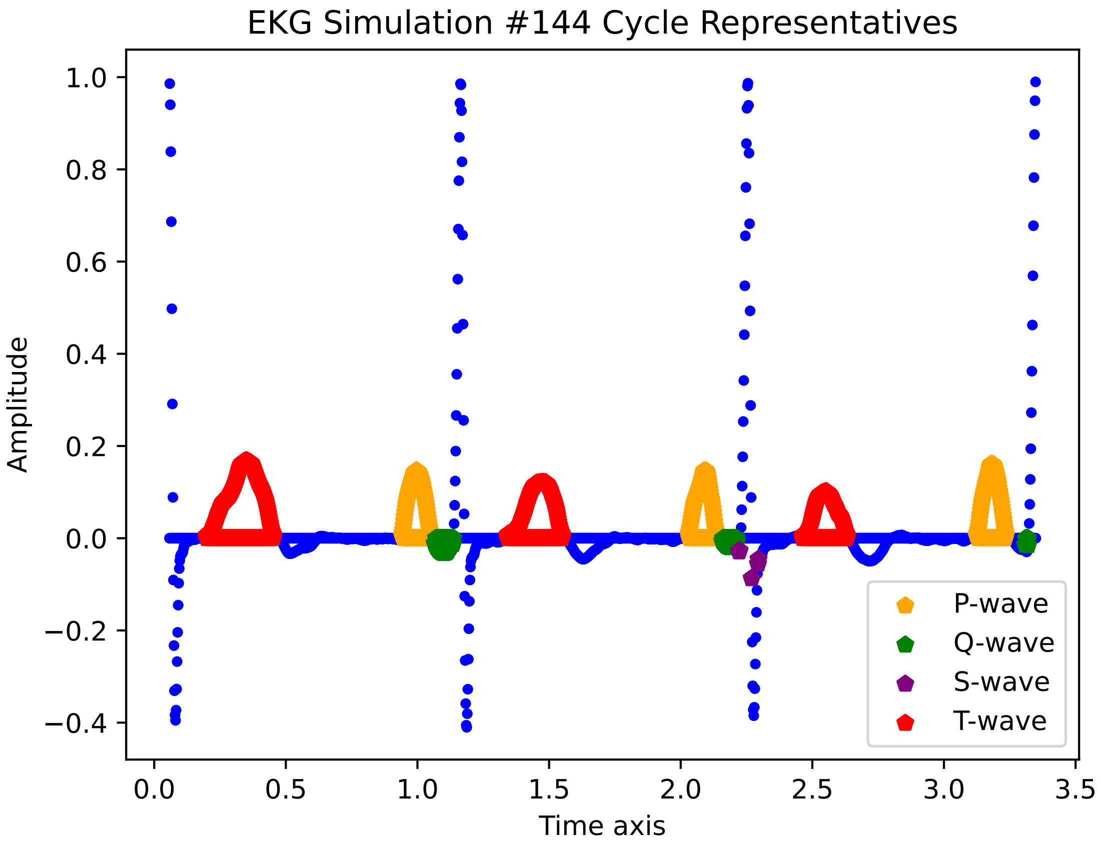
EKG Simulation #144 Cycle Representatives
Amplitude
Time axis
1.0
0.8
0.6
0.4
0.2
0.0
−0.2
−0.4
0.0
0.5
1.0
1.5
2.0
2.5
3.0
3.5
P-wave
Q-wave
S-wave
T-wave

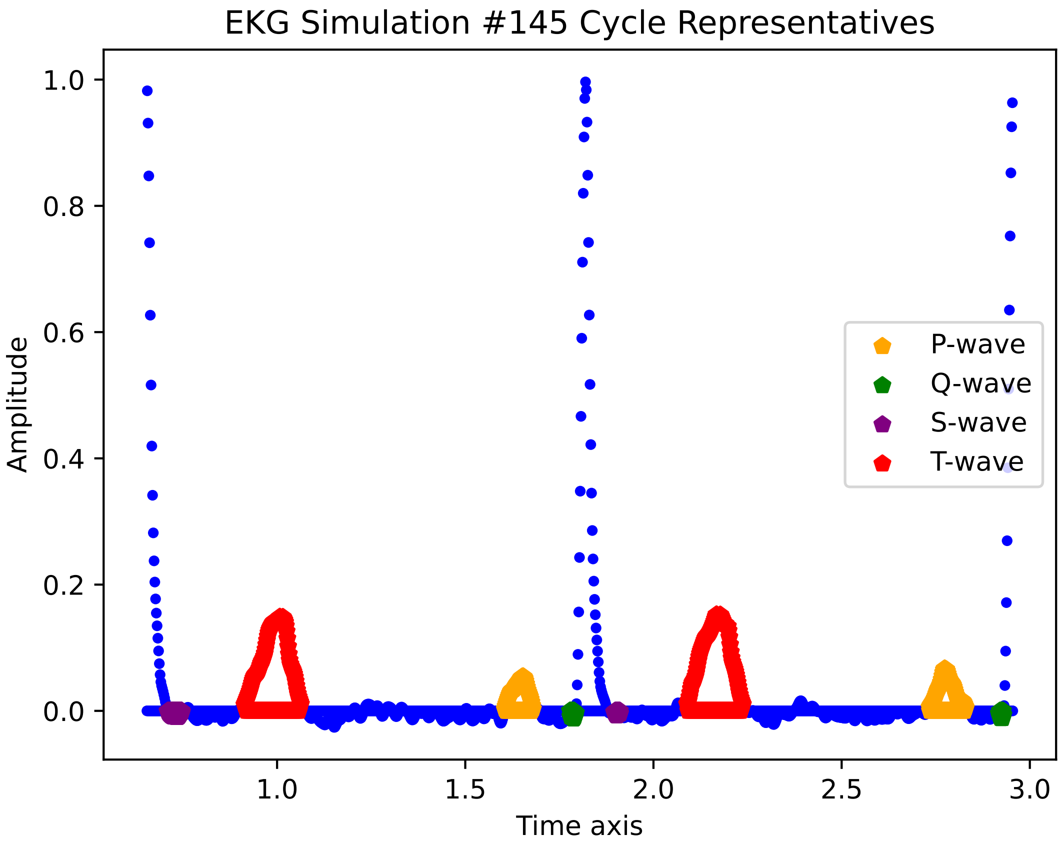
EKG Simulation #145 Cycle Representatives
Amplitude
Time axis
1.0
0.8
0.6
0.4
0.2
0.0
1.0
1.5
2.0
2.5
3.0
P-wave
Q-wave
S-wave
T-wave

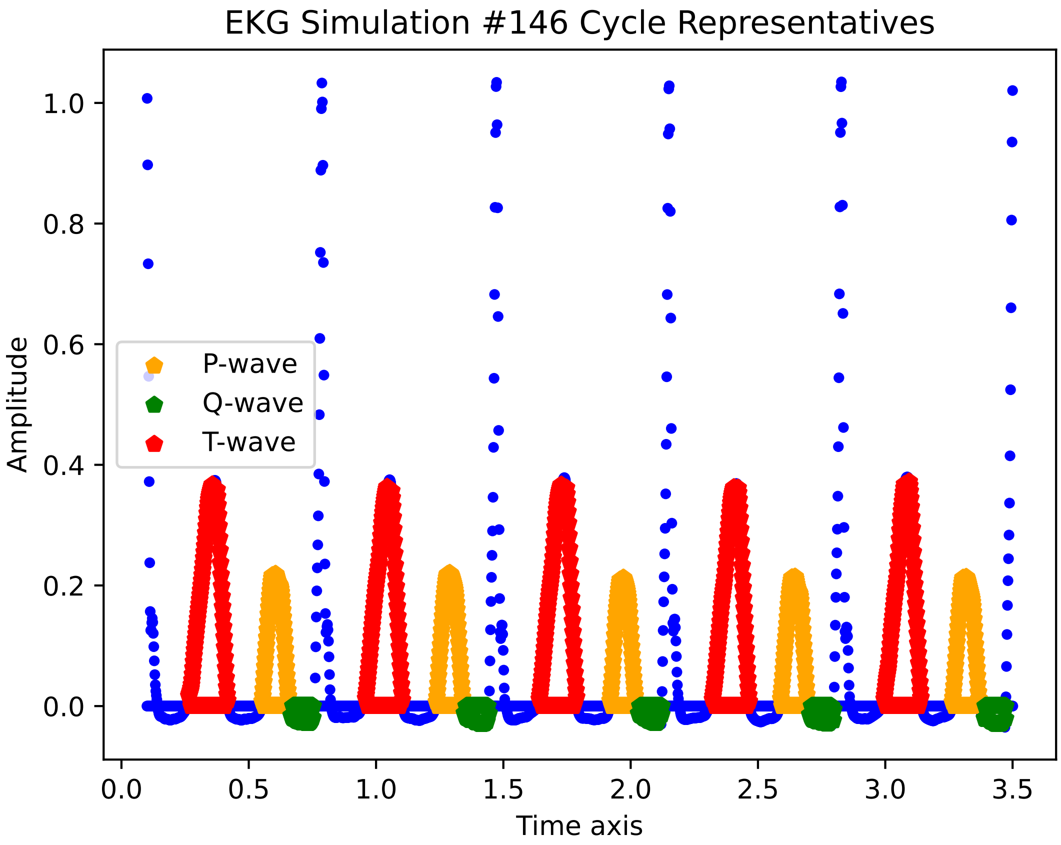

EKG Simulation #146 Cycle Representatives
Amplitude
Time axis
P-wave
Q-wave
T-wave
1.0
0.8
0.6
0.4
0.2
0.0
0.0
0.5
1.0
1.5
2.0
2.5
3.0
3.5

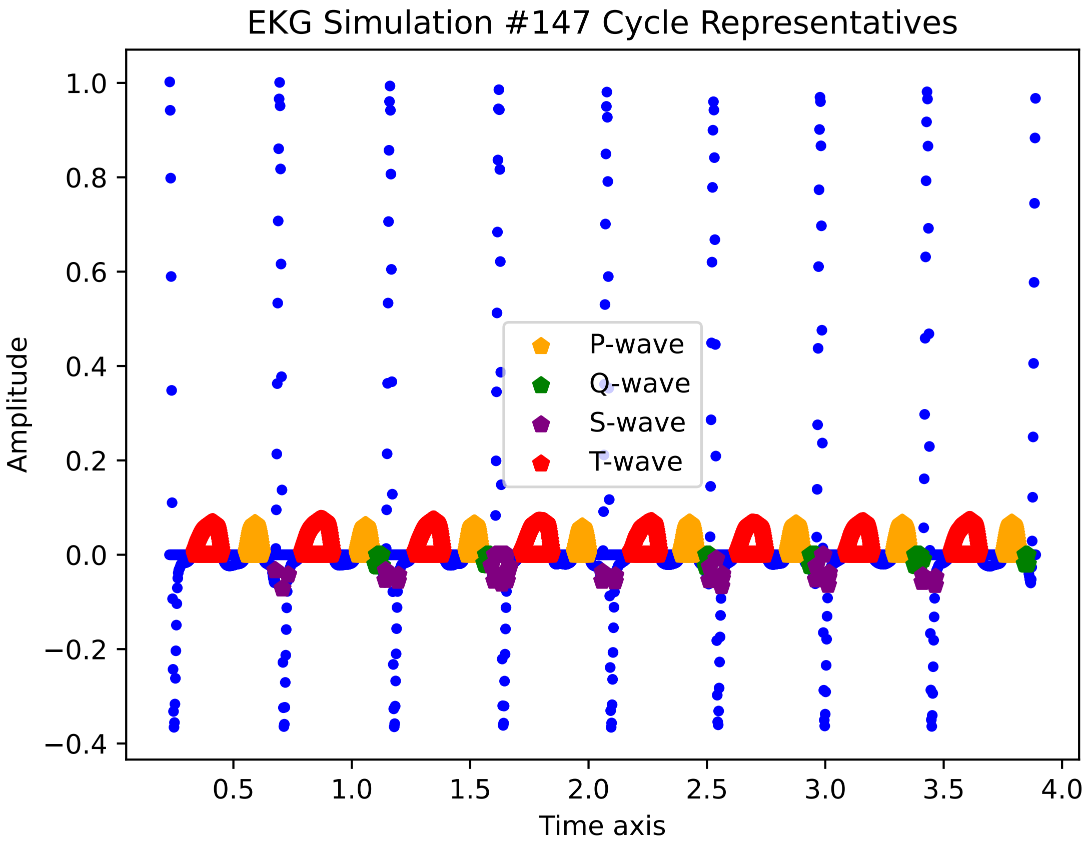

EKG Simulation #147 Cycle Representatives
Amplitude
Time axis
P-wave
Q-wave
S-wave
T-wave
1.0
0.8
0.6
0.4
0.2
0.0
−0.2
−0.4
0.5
1.0
1.5
2.0
2.5
3.0
3.5
4.0

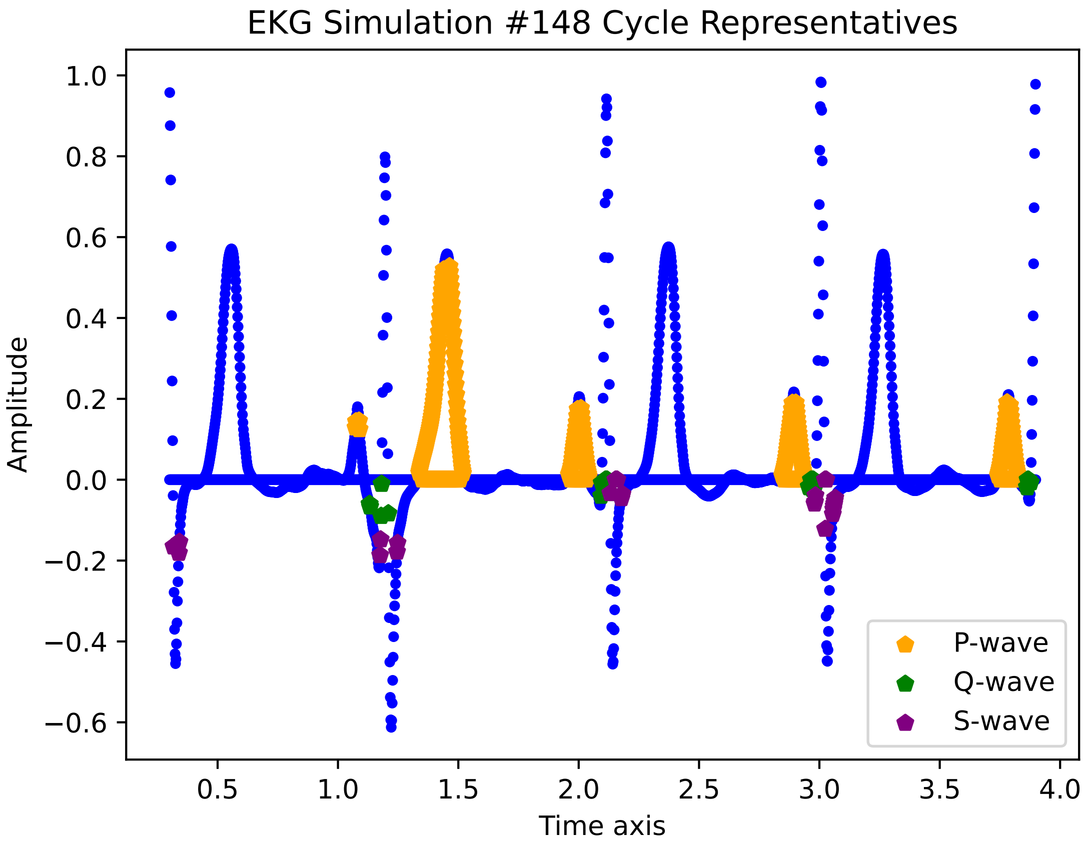

EKG Simulation #148 Cycle Representatives
Amplitude
Time axis
1.0
0.8
0.6
0.4
0.2
0.0
−0.2
−0.4
−0.6
0.5
1.0
1.5
2.0
2.5
3.0
3.5
4.0
P-wave
Q-wave
S-wave

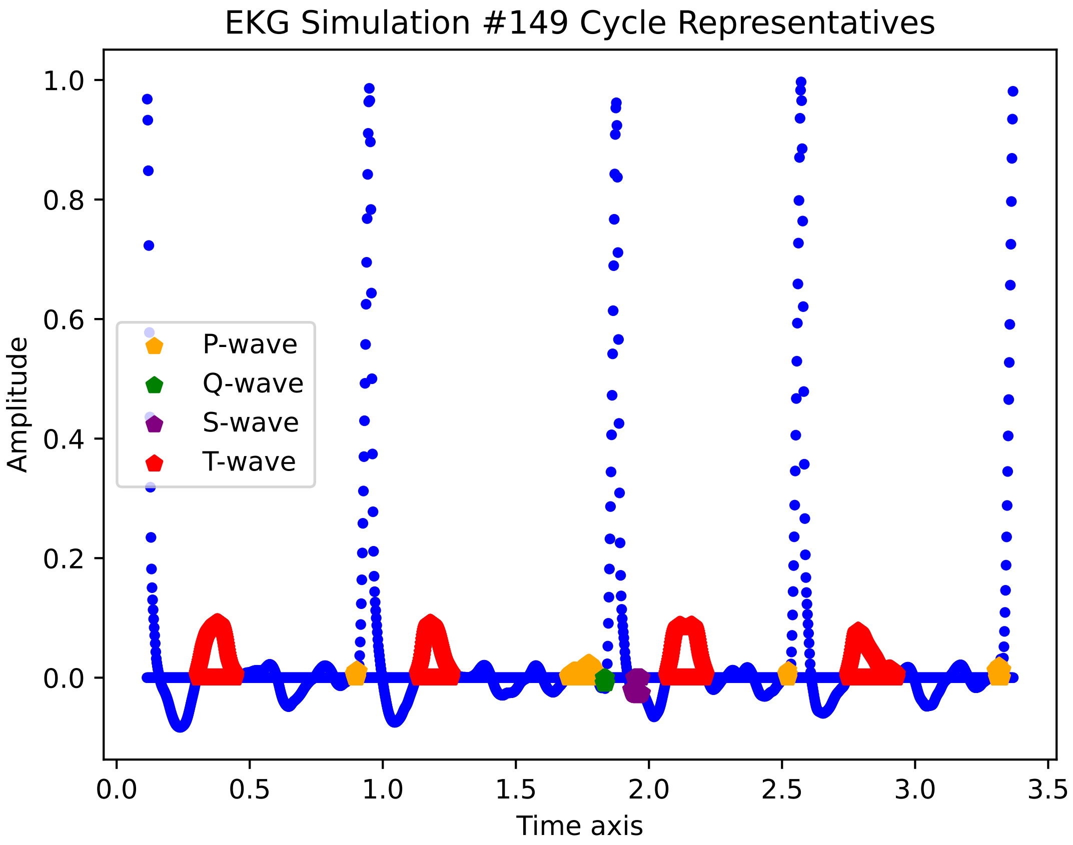

EKG Simulation #149 Cycle Representatives
Amplitude
Time axis
P-wave
Q-wave
S-wave
T-wave
1.0
0.8
0.6
0.4
0.2
0.0
0.0
0.5
1.0
1.5
2.0
2.5
3.0
3.5

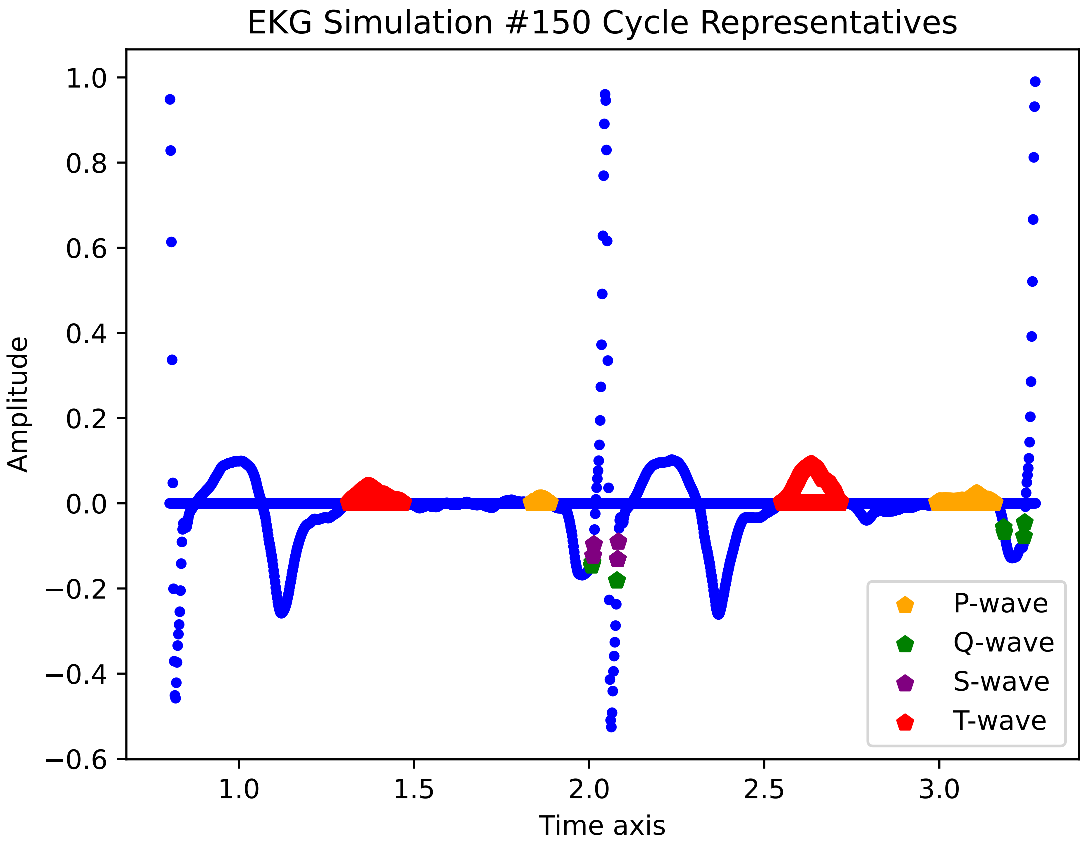

EKG Simulation #150 Cycle Representatives
Amplitude
Time axis
1.0
0.8
0.6
0.4
0.2
0.0
−0.2
−0.4
−0.6
1.0
1.5
2.0
2.5
3.0
P-wave
Q-wave
S-wave
T-wave

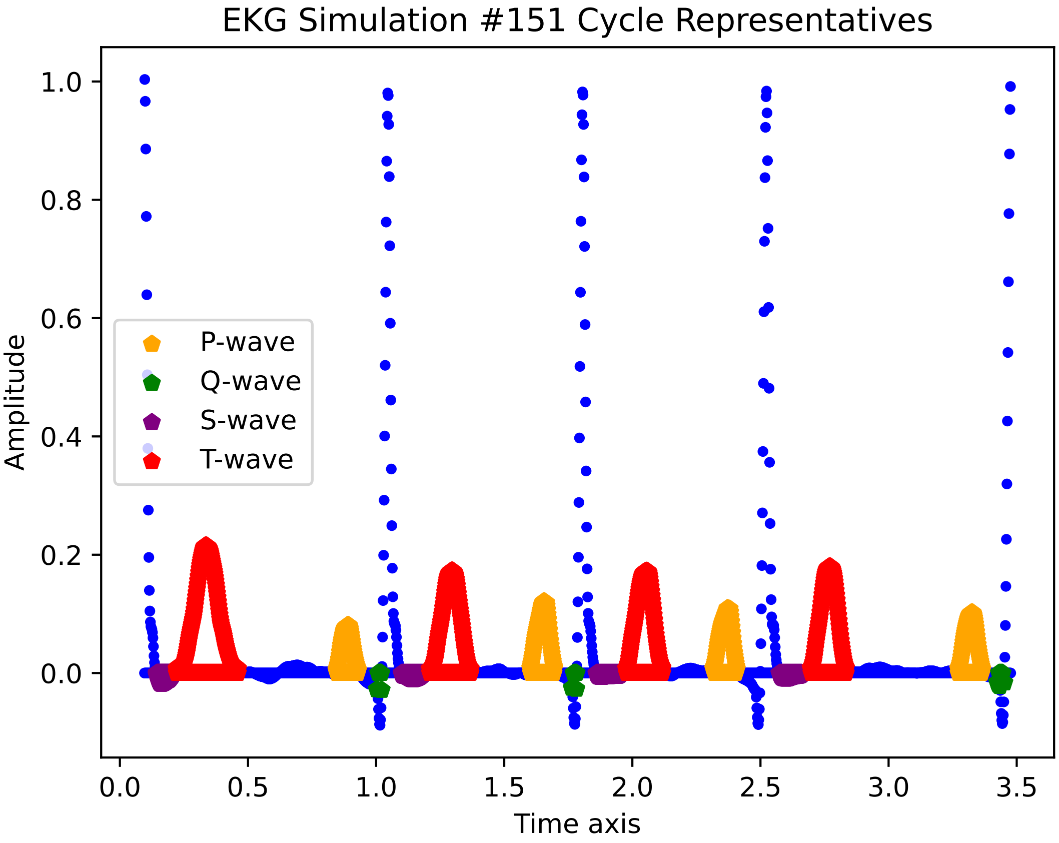
EKG Simulation #151 Cycle Representatives
1.0
0.8
0.6
0.4
0.2
0.0
Amplitude
P-wave
Q-wave
S-wave
T-wave
0.0
0.5
1.0
1.5
2.0
2.5
3.0
3.5
Time axis

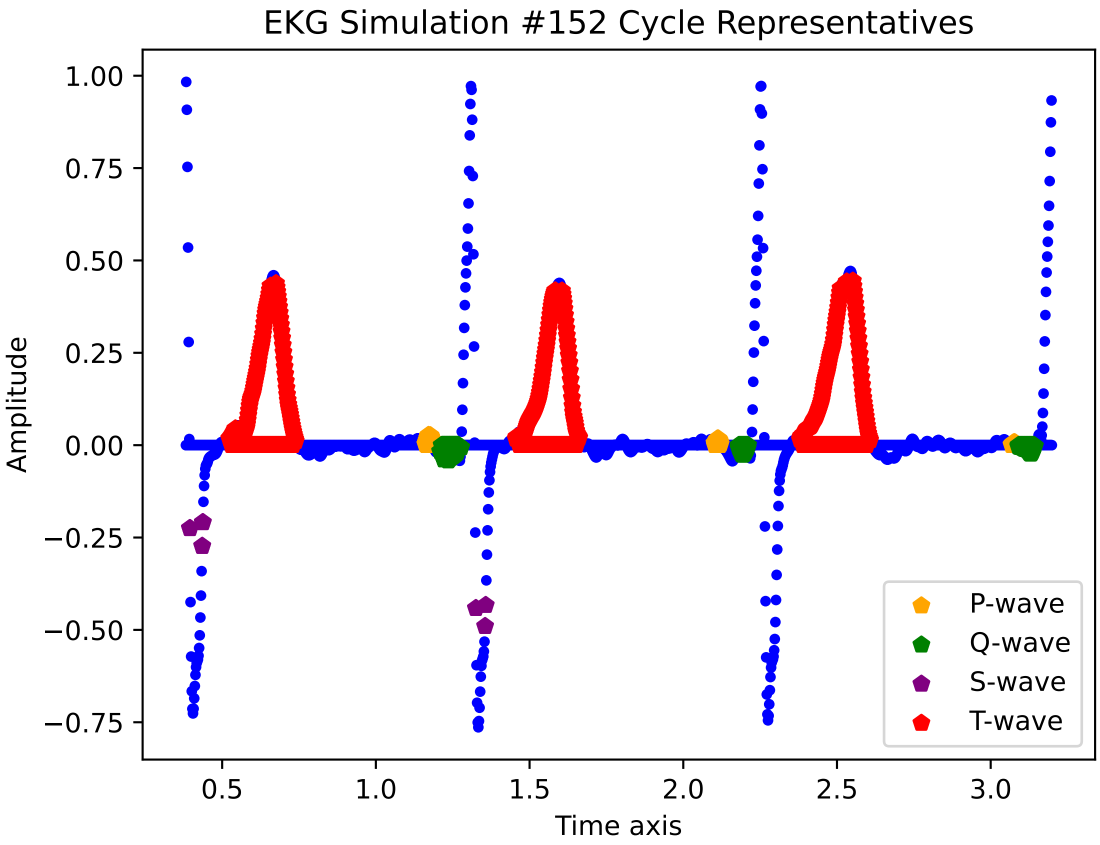

EKG Simulation #152 Cycle Representatives
Amplitude
Time axis
1.00
0.75
0.50
0.25
0.00
−0.25
−0.50
−0.75
0.5
1.0
1.5
2.0
2.5
3.0
P-wave
Q-wave
S-wave
T-wave

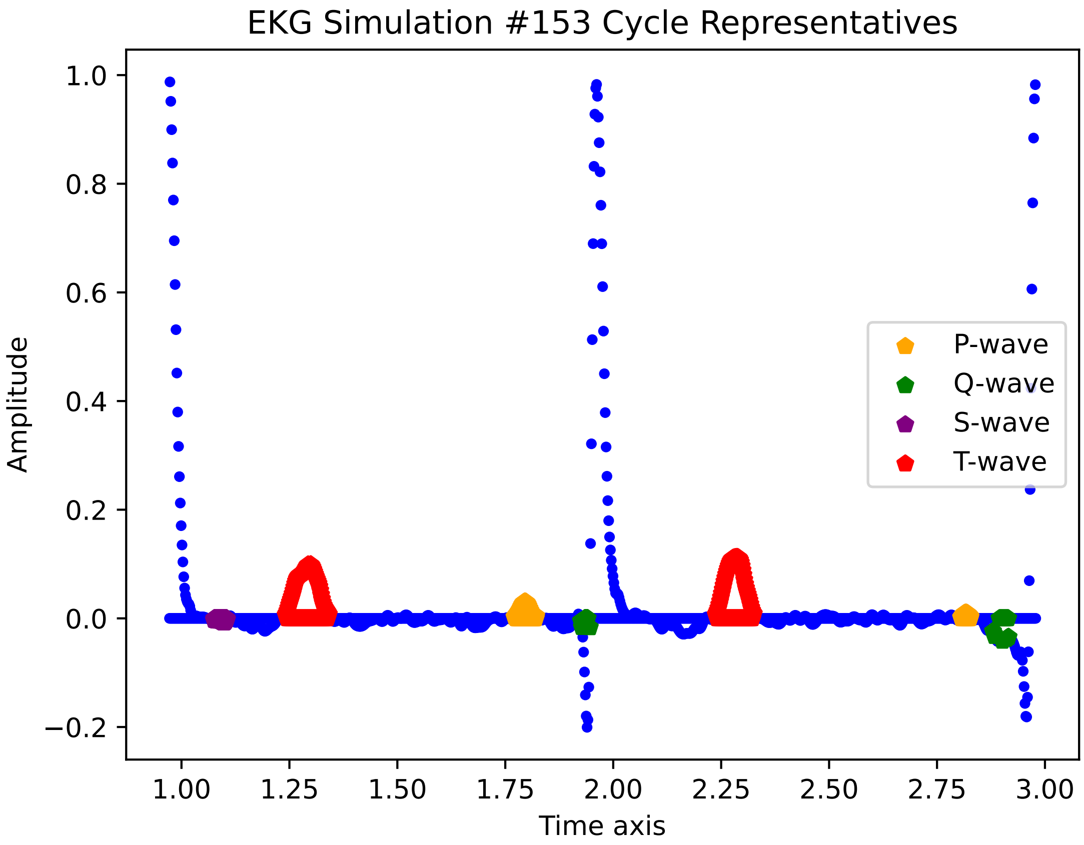
EKG Simulation #153 Cycle Representatives
1.0
0.8
0.6
0.4
0.2
0.0
−0.2
1.00
1.25
1.50
1.75
2.00
2.25
2.50
2.75
3.00
Time axis
Amplitude
P-wave
Q-wave
S-wave
T-wave

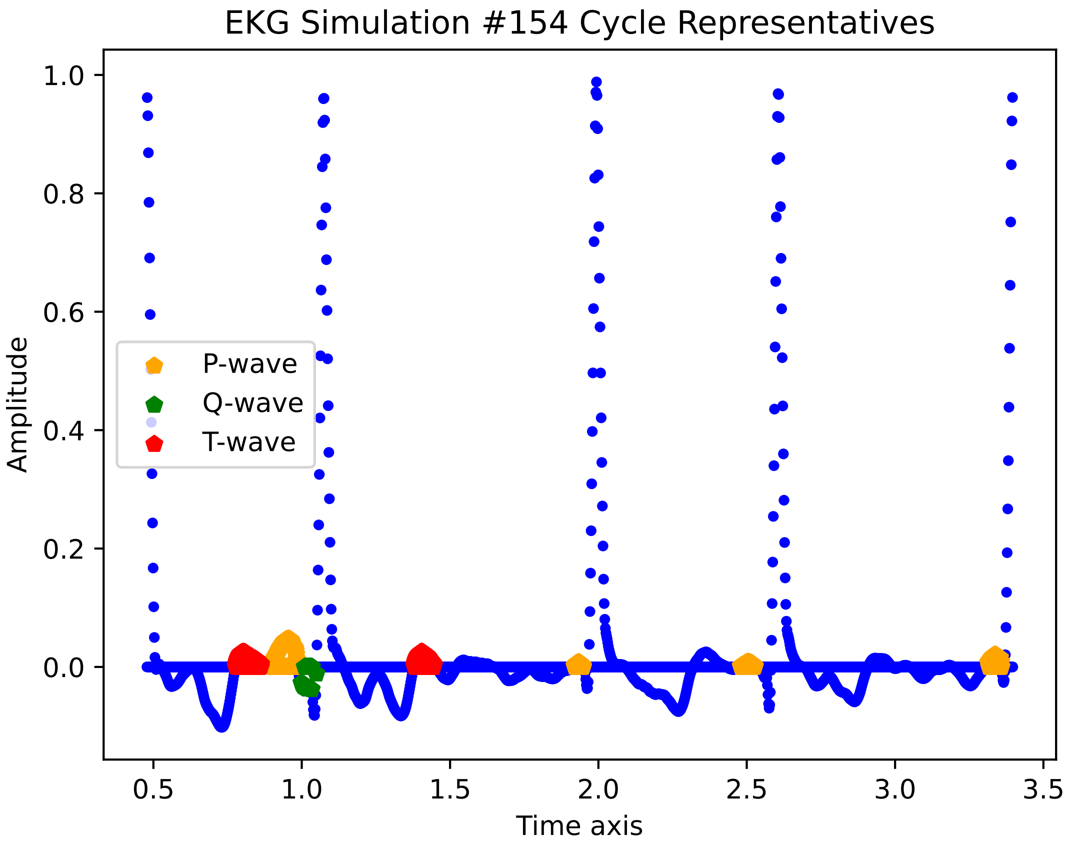

EKG Simulation #154 Cycle Representatives
Amplitude
Time axis
P-wave
Q-wave
T-wave
1.0
0.8
0.6
0.4
0.2
0.0
0.5
1.0
1.5
2.0
2.5
3.0
3.5

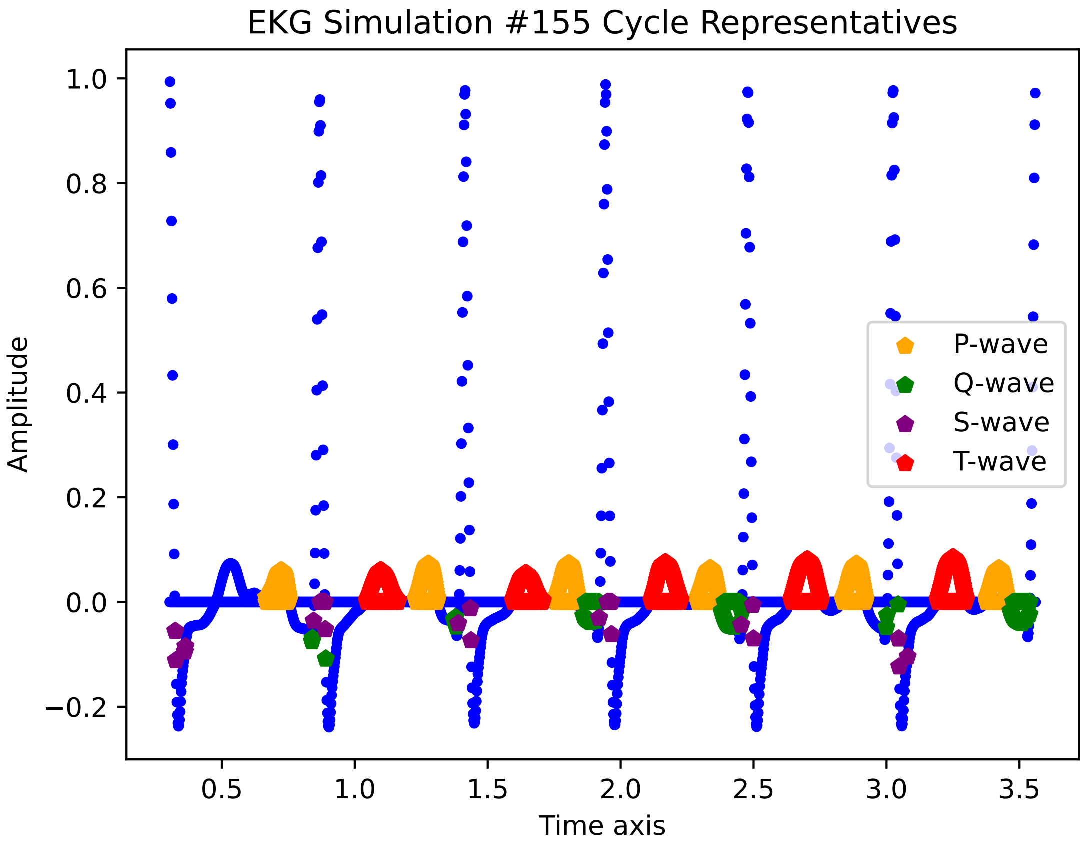

EKG Simulation #155 Cycle Representatives
Amplitude
Time axis
1.0
0.8
0.6
0.4
0.2
0.0
−0.2
0.5
1.0
1.5
2.0
2.5
3.0
3.5
P-wave
Q-wave
S-wave
T-wave

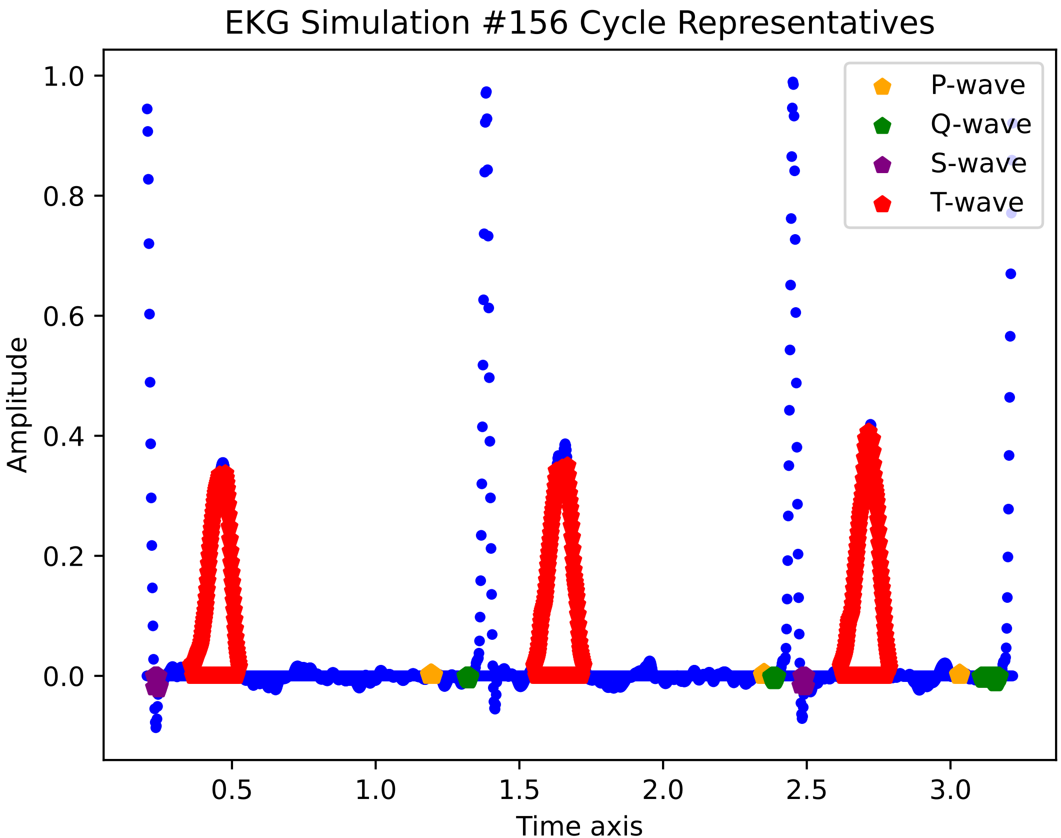
EKG Simulation #156 Cycle Representatives
P-wave
Q-wave
S-wave
T-wave
Amplitude
Time axis
1.0
0.8
0.6
0.4
0.2
0.0
0.5
1.0
1.5
2.0
2.5
3.0

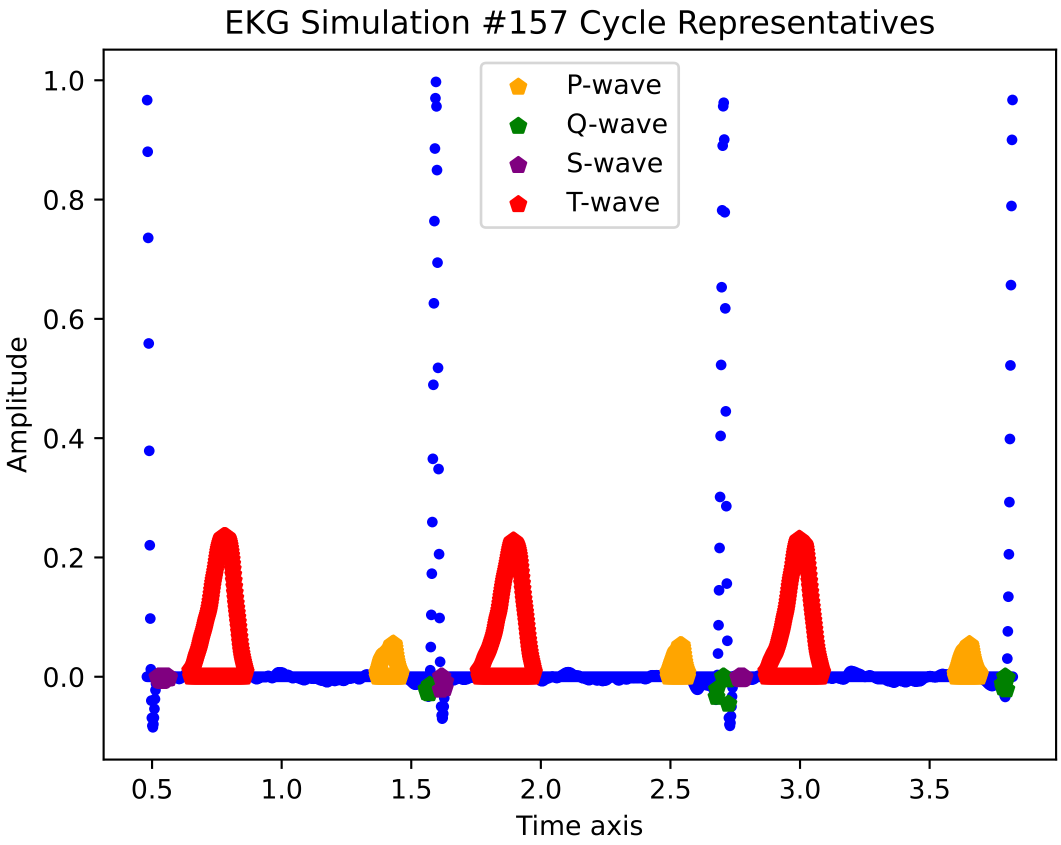

EKG Simulation #157 Cycle Representatives
P-wave
Q-wave
S-wave
T-wave
Amplitude
Time axis
1.0
0.8
0.6
0.4
0.2
0.0
0.5
1.0
1.5
2.0
2.5
3.0
3.5

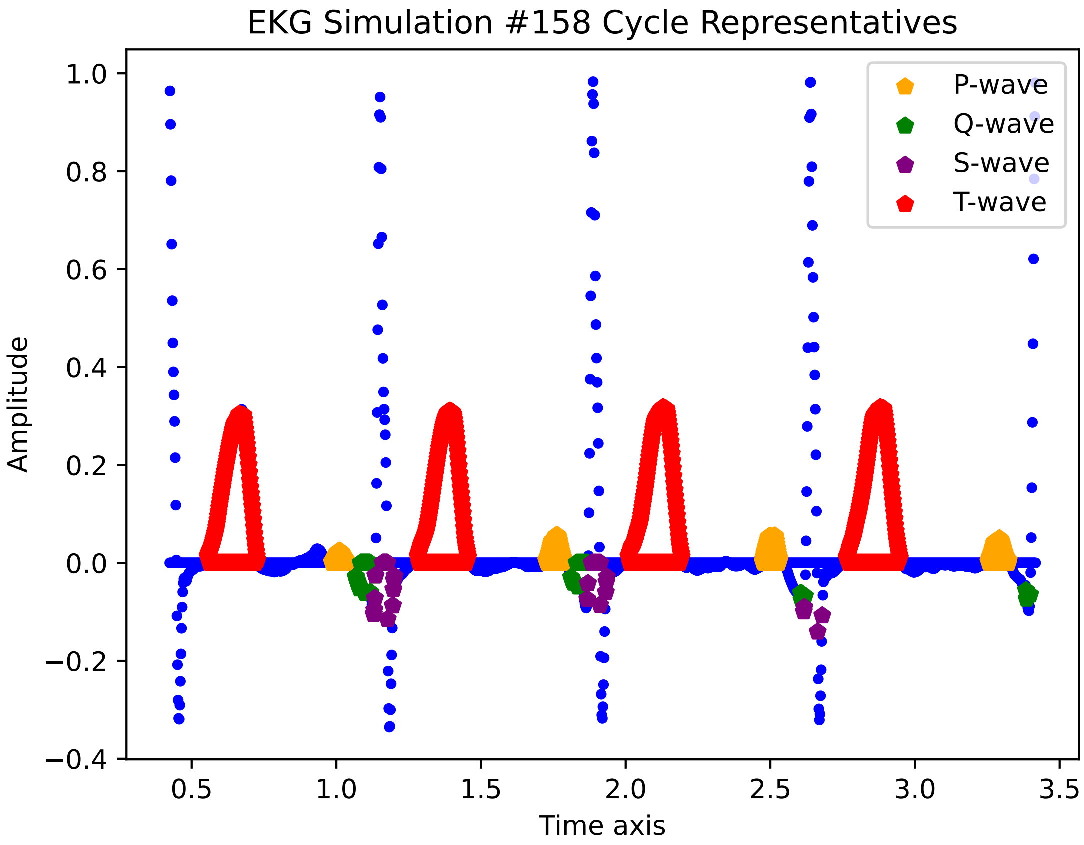
EKG Simulation #158 Cycle Representatives
Amplitude
Time axis
P-wave
Q-wave
S-wave
T-wave
1.0
0.8
0.6
0.4
0.2
0.0
−0.2
−0.4
0.5
1.0
1.5
2.0
2.5
3.0
3.5

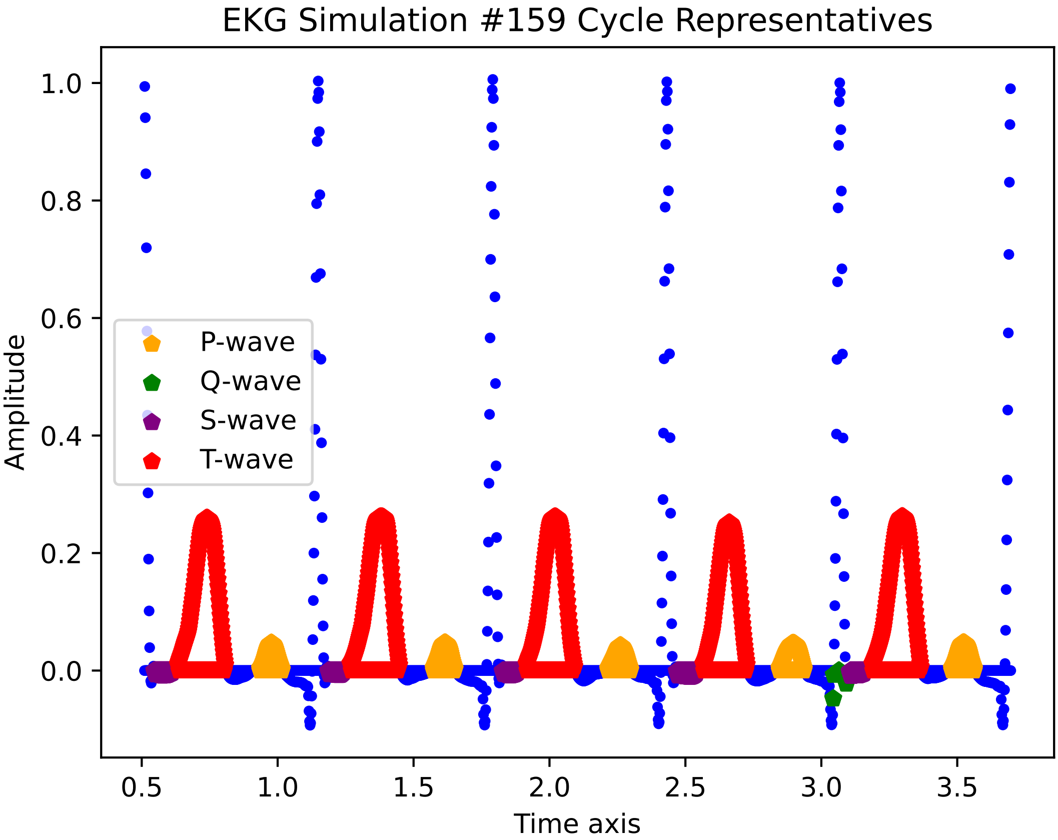

EKG Simulation #159 Cycle Representatives
Amplitude
Time axis
P-wave
Q-wave
S-wave
T-wave
1.0
0.8
0.6
0.4
0.2
0.0
0.5
1.0
1.5
2.0
2.5
3.0
3.5

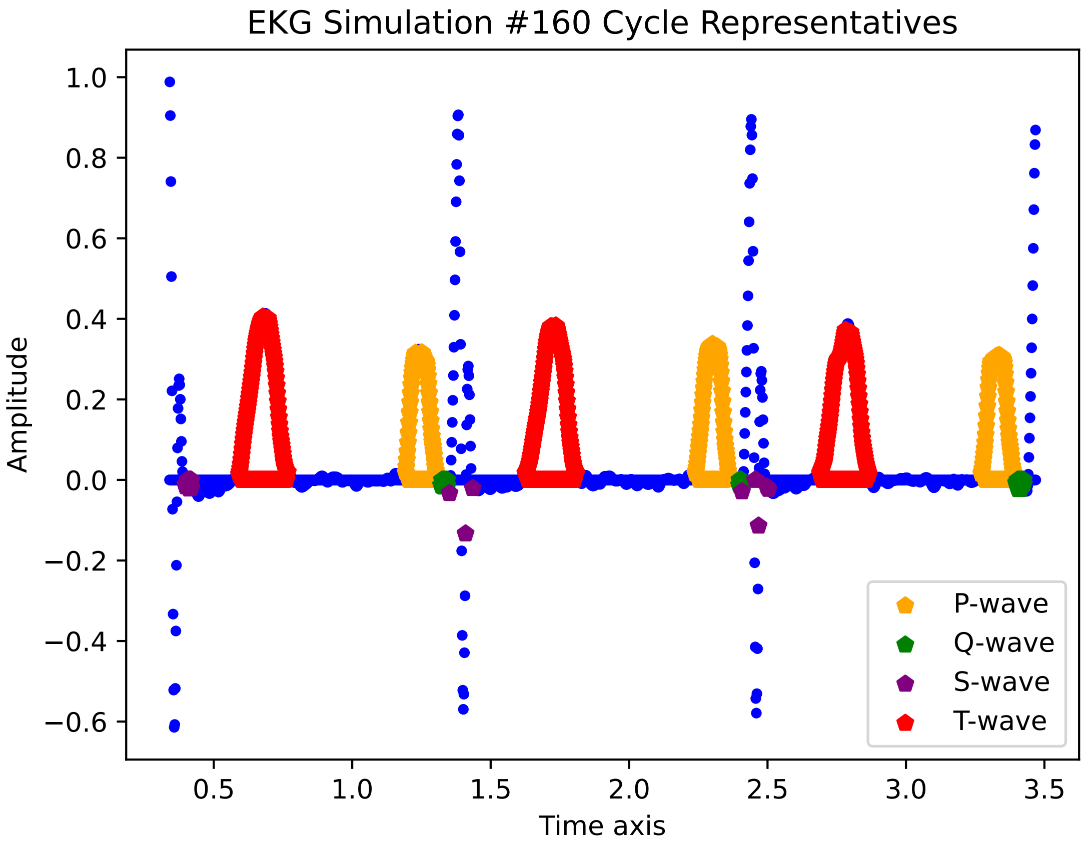
EKG Simulation #160 Cycle Representatives
Amplitude
Time axis
1.0
0.8
0.6
0.4
0.2
0.0
−0.2
−0.4
−0.6
0.5
1.0
1.5
2.0
2.5
3.0
3.5
P-wave
Q-wave
S-wave
T-wave

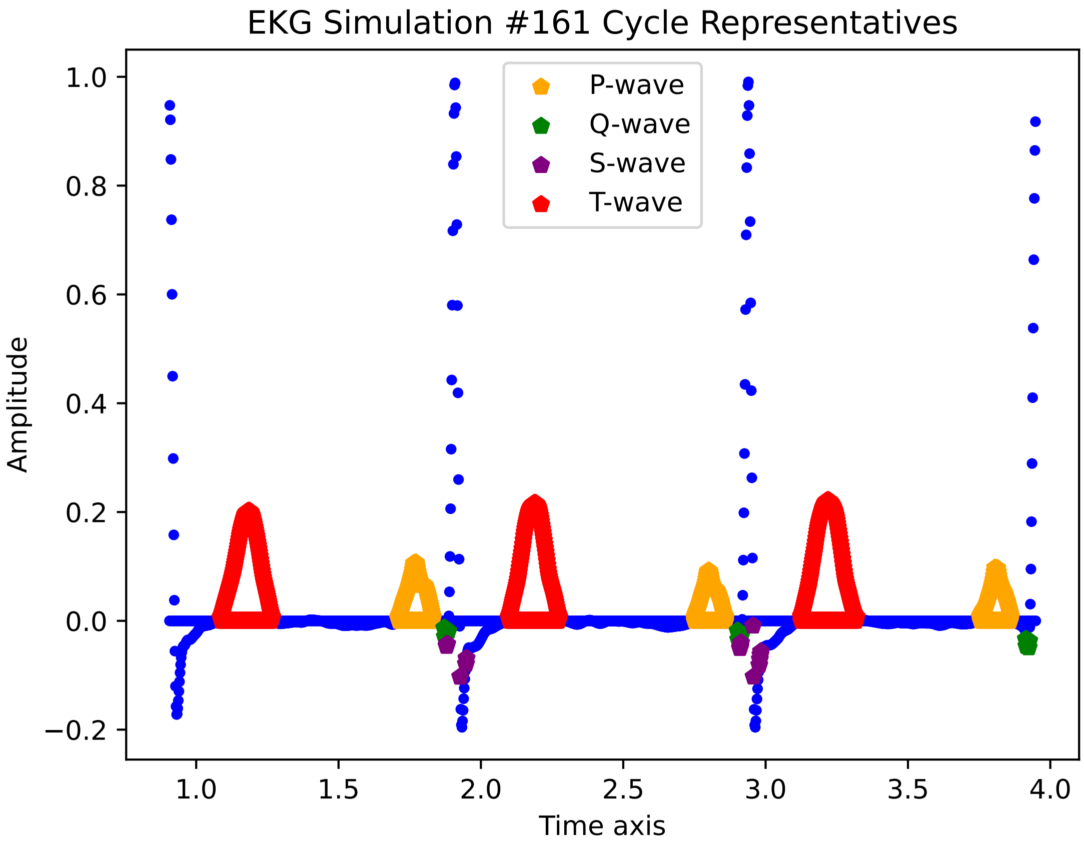

EKG Simulation #161 Cycle Representatives
P-wave
Q-wave
S-wave
T-wave
Amplitude
Time axis
1.0
0.8
0.6
0.4
0.2
0.0
−0.2
1.0
1.5
2.0
2.5
3.0
3.5
4.0

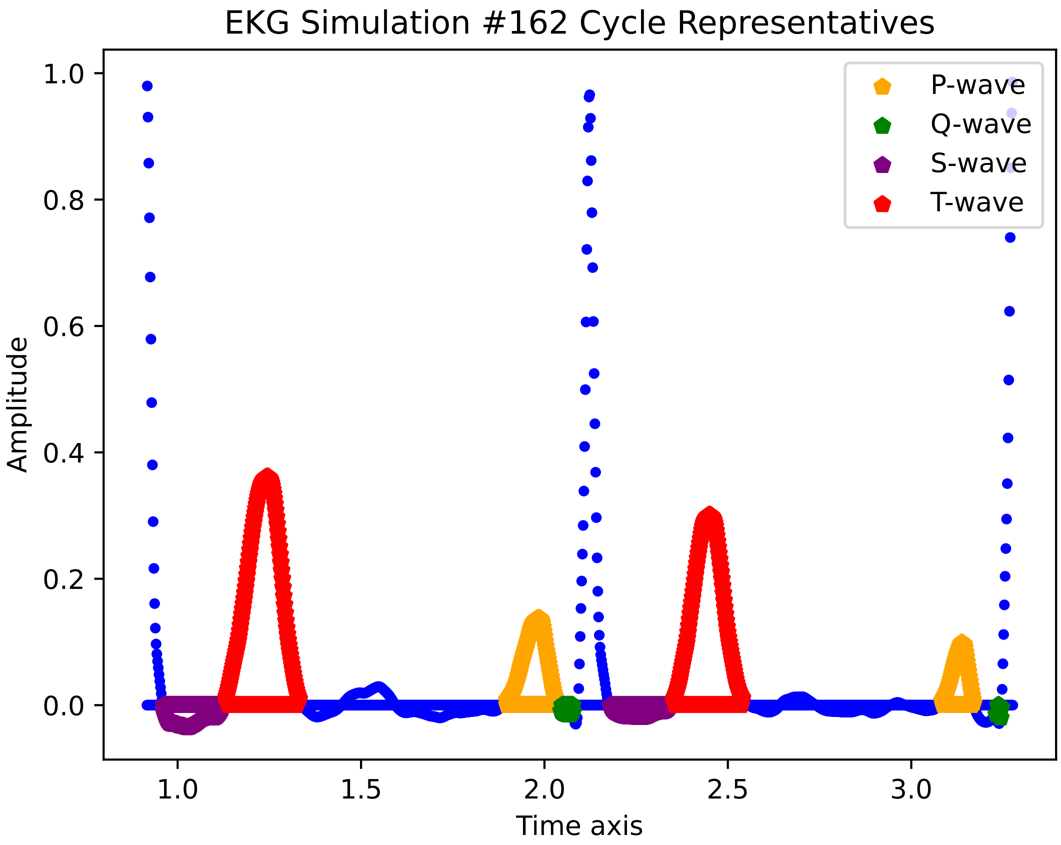
EKG Simulation #162 Cycle Representatives
P-wave
Q-wave
S-wave
T-wave
Amplitude
Time axis
1.0
0.8
0.6
0.4
0.2
0.0
1.0
1.5
2.0
2.5
3.0

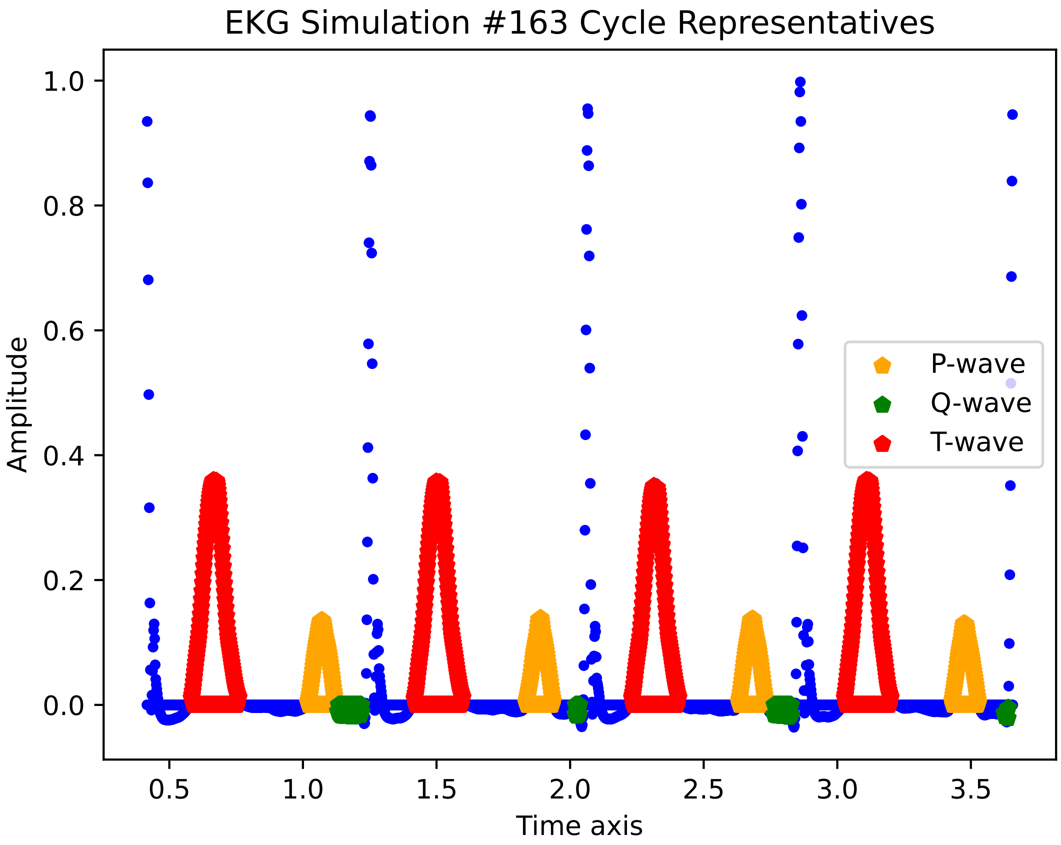

EKG Simulation #163 Cycle Representatives
Amplitude
Time axis
1.0
0.8
0.6
0.4
0.2
0.0
0.5
1.0
1.5
2.0
2.5
3.0
3.5
P-wave
Q-wave
T-wave

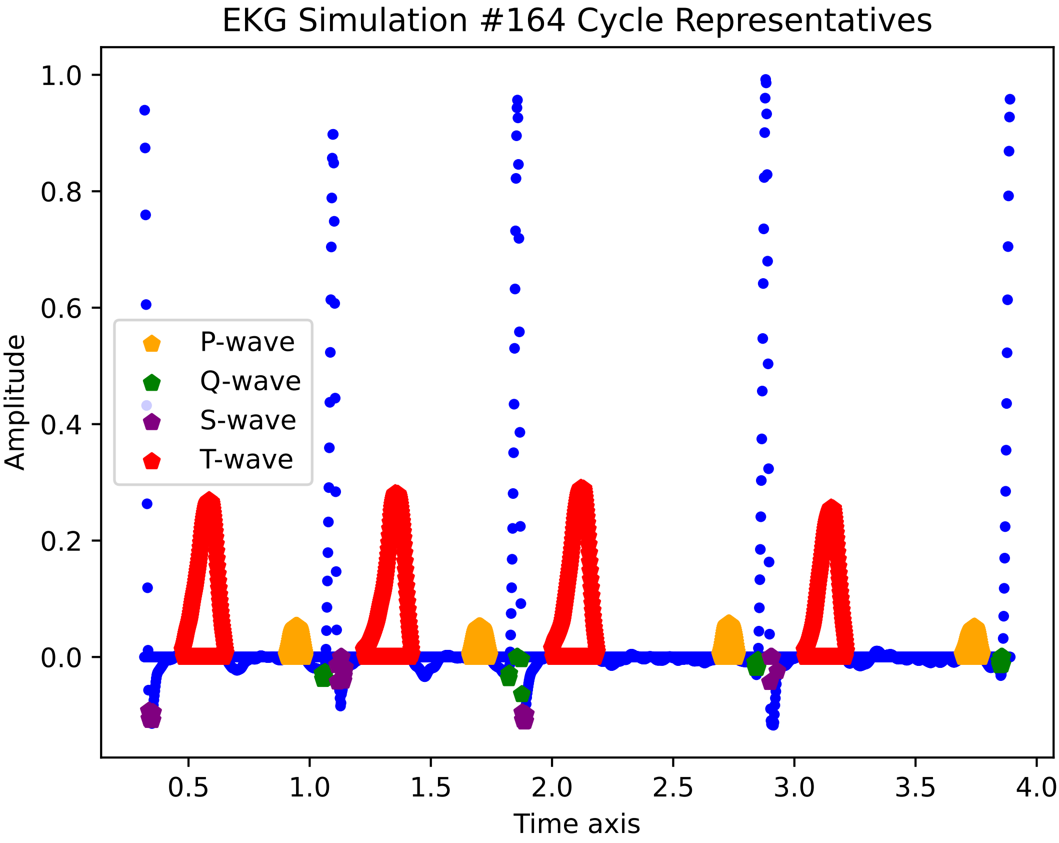

EKG Simulation #164 Cycle Representatives
1.0
0.8
0.6
0.4
0.2
0.0
Amplitude
P-wave
Q-wave
S-wave
T-wave
0.5
1.0
1.5
2.0
2.5
3.0
3.5
4.0
Time axis

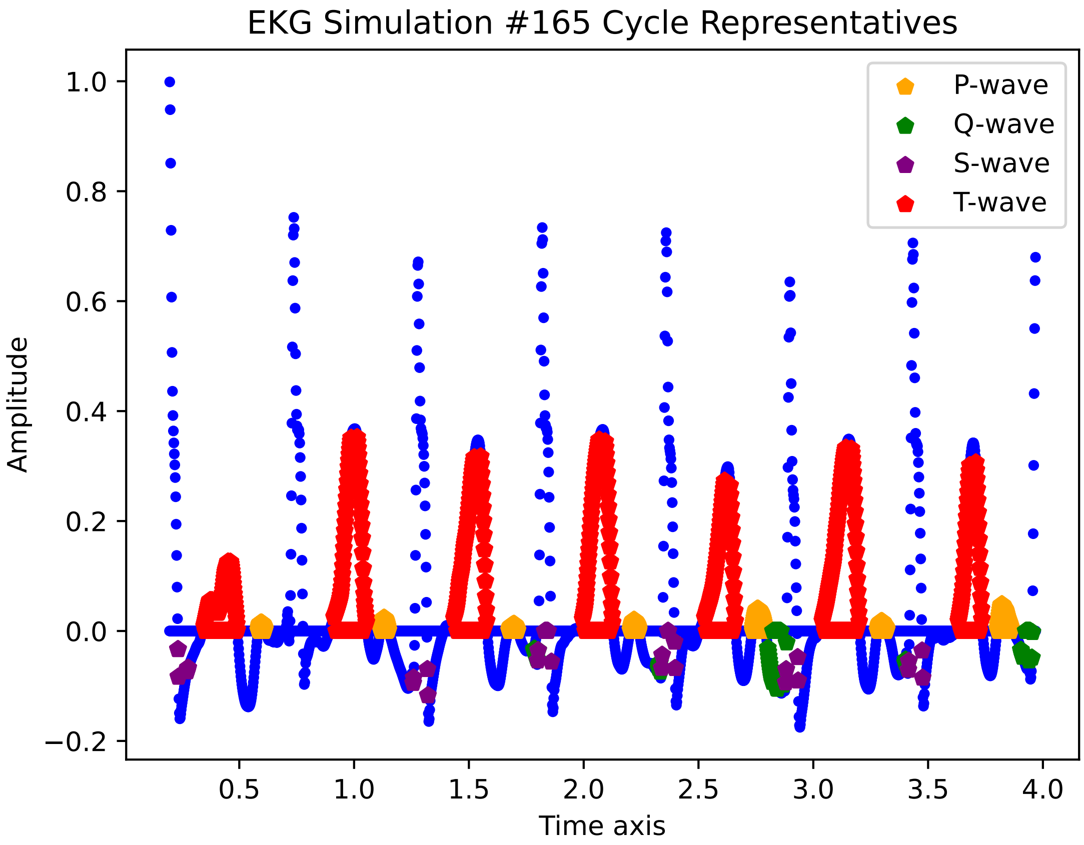
EKG Simulation #165 Cycle Representatives
P-wave
Q-wave
S-wave
T-wave
Amplitude
Time axis
1.0
0.8
0.6
0.4
0.2
0.0
−0.2
0.5
1.0
1.5
2.0
2.5
3.0
3.5
4.0

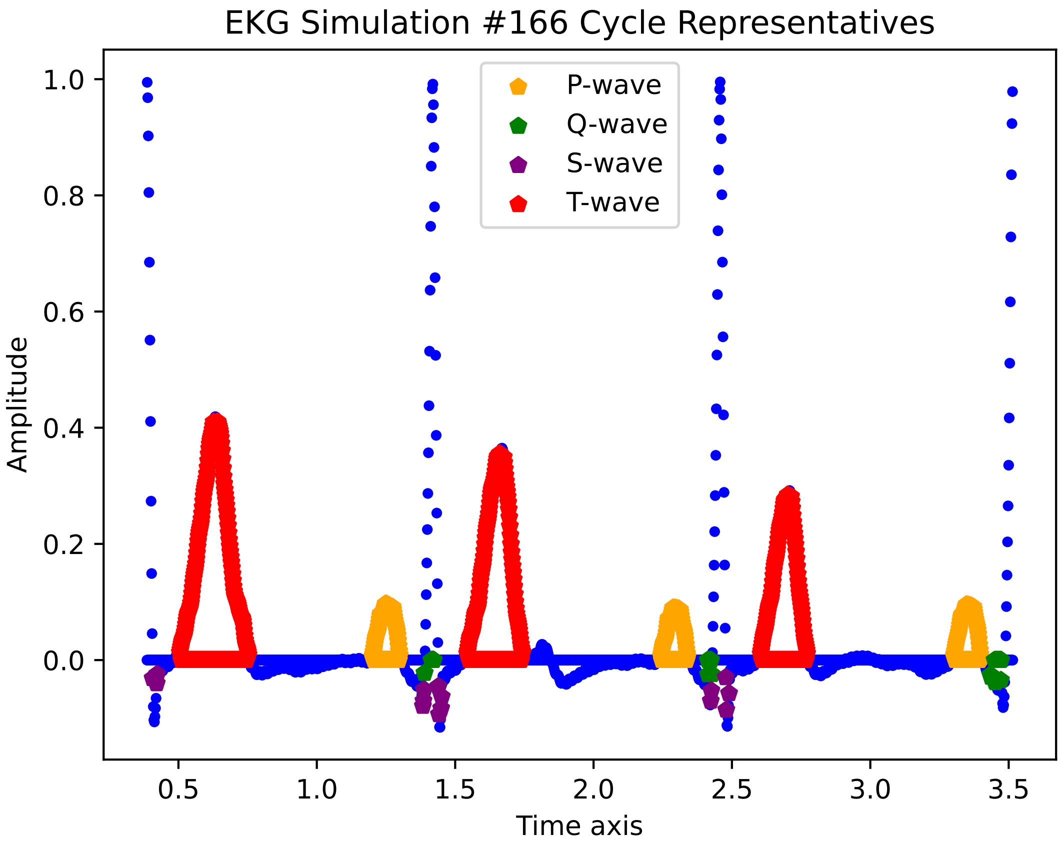
EKG Simulation #166 Cycle Representatives
P-wave
Q-wave
S-wave
T-wave
Amplitude
Time axis
1.0
0.8
0.6
0.4
0.2
0.0
0.5
1.0
1.5
2.0
2.5
3.0
3.5

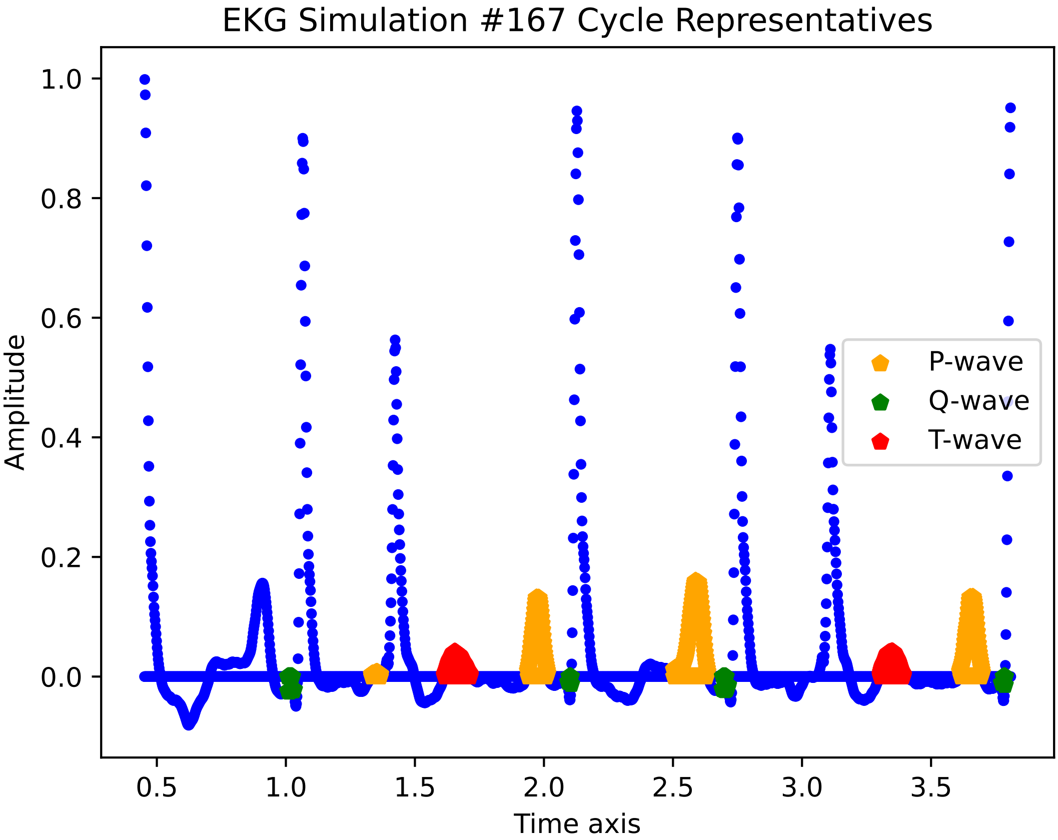

EKG Simulation #167 Cycle Representatives
Amplitude
Time axis
1.0
0.8
0.6
0.4
0.2
0.0
0.5
1.0
1.5
2.0
2.5
3.0
3.5
P-wave
Q-wave
T-wave

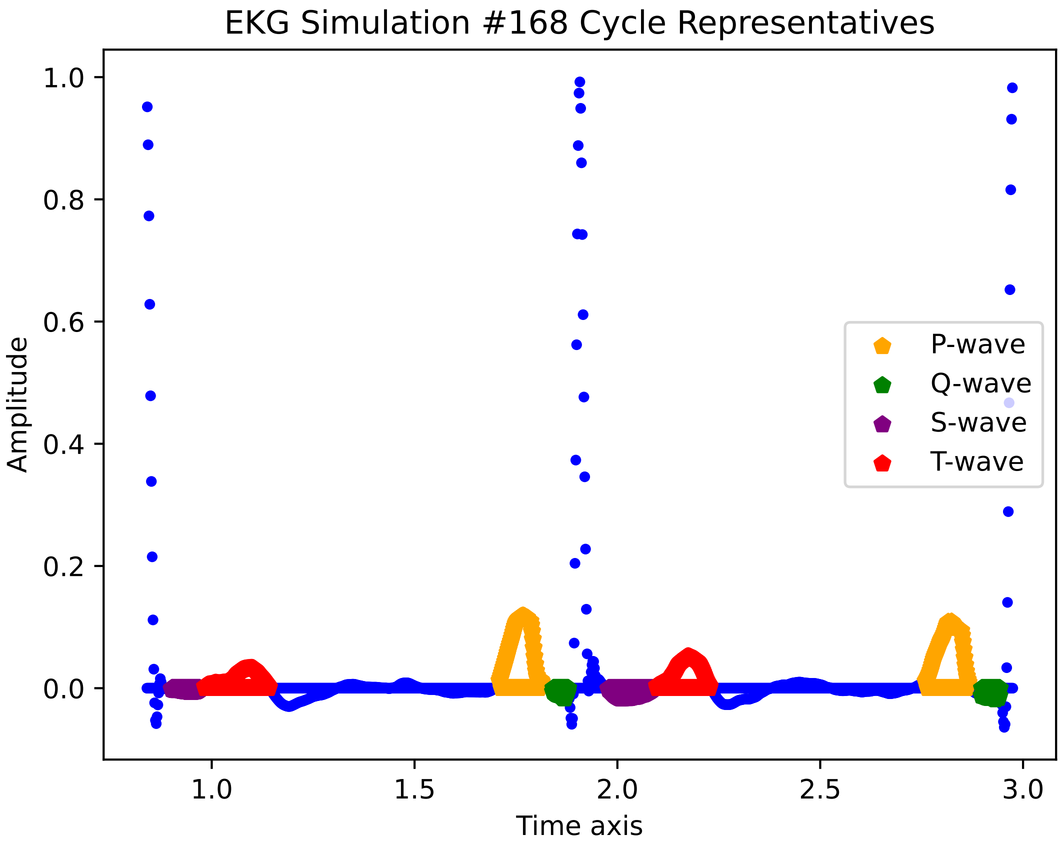

EKG Simulation #168 Cycle Representatives
Amplitude
Time axis
P-wave
Q-wave
S-wave
T-wave
1.0
0.8
0.6
0.4
0.2
0.0
1.0
1.5
2.0
2.5
3.0

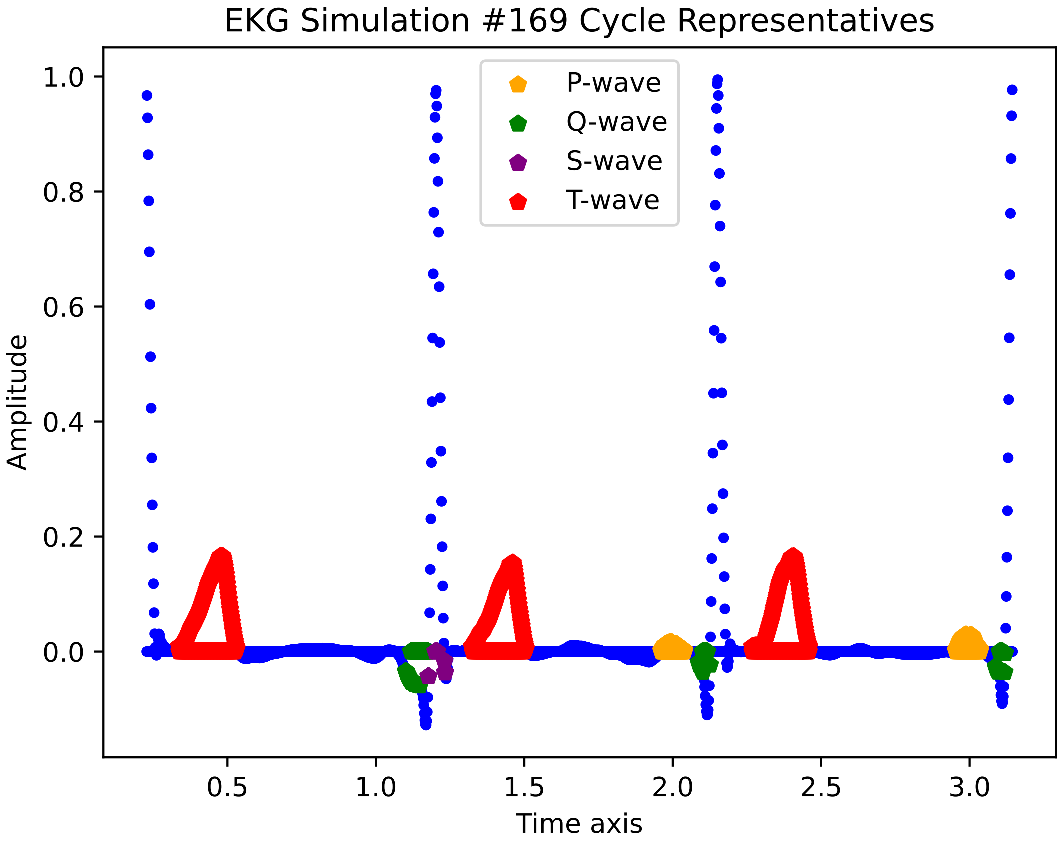
EKG Simulation #169 Cycle Representatives
P-wave
Q-wave
S-wave
T-wave
Amplitude
1.0
0.8
0.6
0.4
0.2
0.0
0.5
1.0
1.5
2.0
2.5
3.0
Time axis

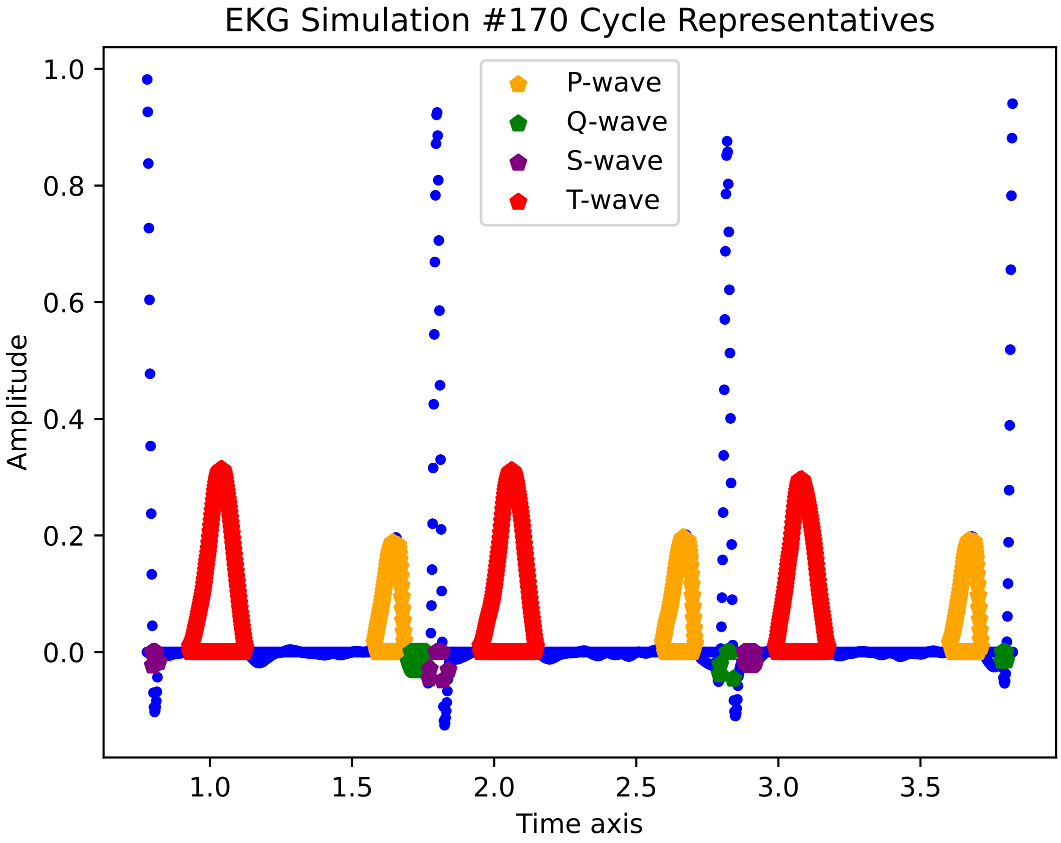
EKG Simulation #170 Cycle Representatives
P-wave
Q-wave
S-wave
T-wave
Amplitude
Time axis
1.0
0.8
0.6
0.4
0.2
0.0
1.0
1.5
2.0
2.5
3.0
3.5

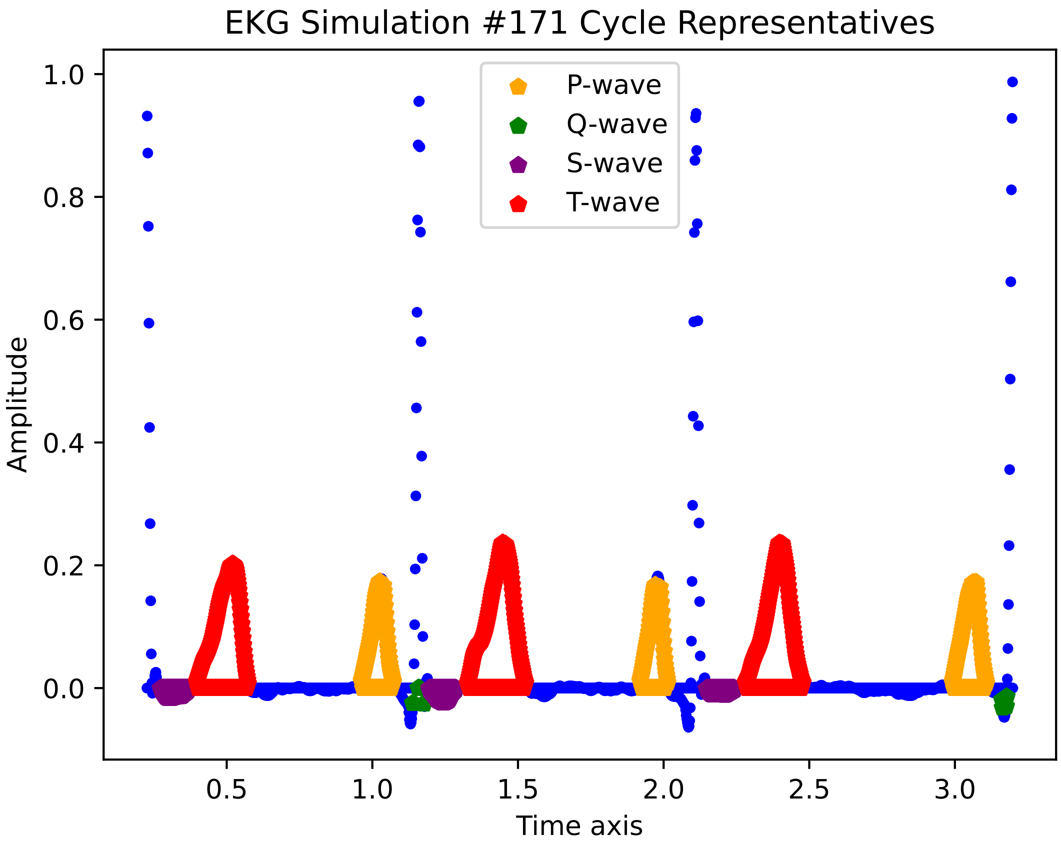
EKG Simulation #171 Cycle Representatives
P-wave
Q-wave
S-wave
T-wave
Amplitude
Time axis
1.0
0.8
0.6
0.4
0.2
0.0
0.5
1.0
1.5
2.0
2.5
3.0

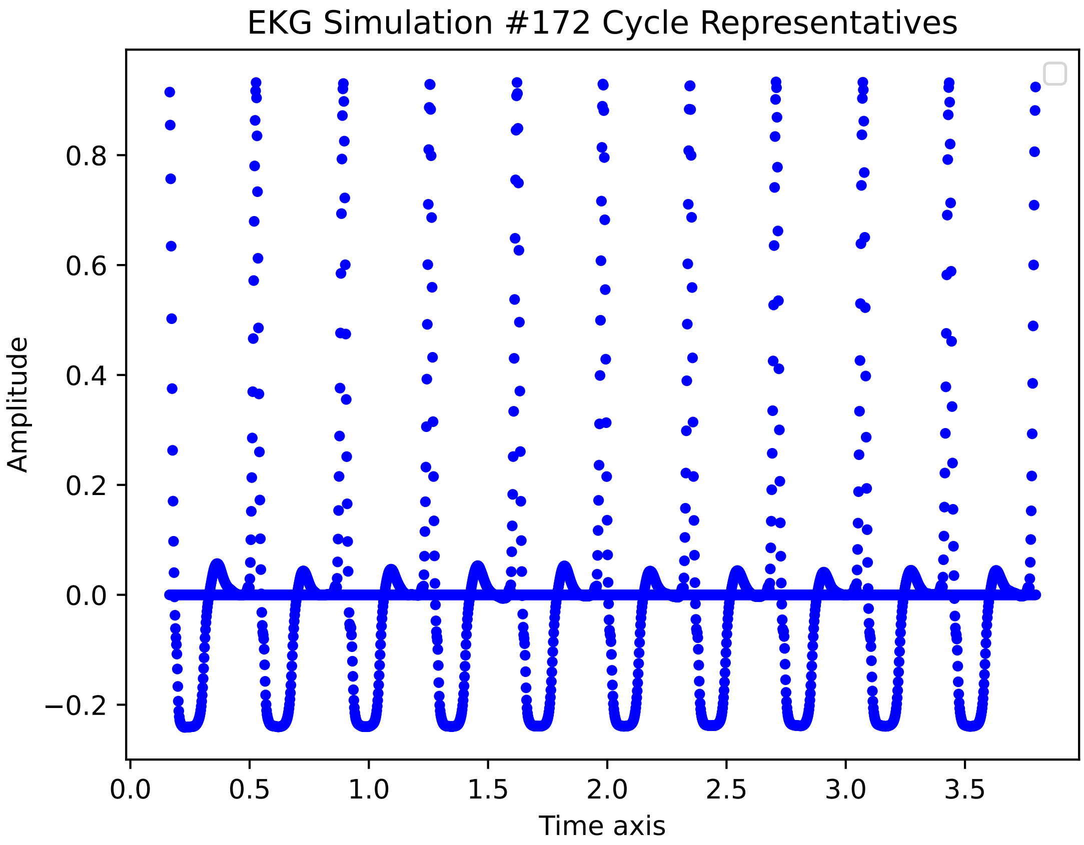
EKG Simulation #172 Cycle Representatives
Amplitude
Time axis
0.8
0.6
0.4
0.2
0.0
−0.2
0.0
0.5
1.0
1.5
2.0
2.5
3.0
3.5

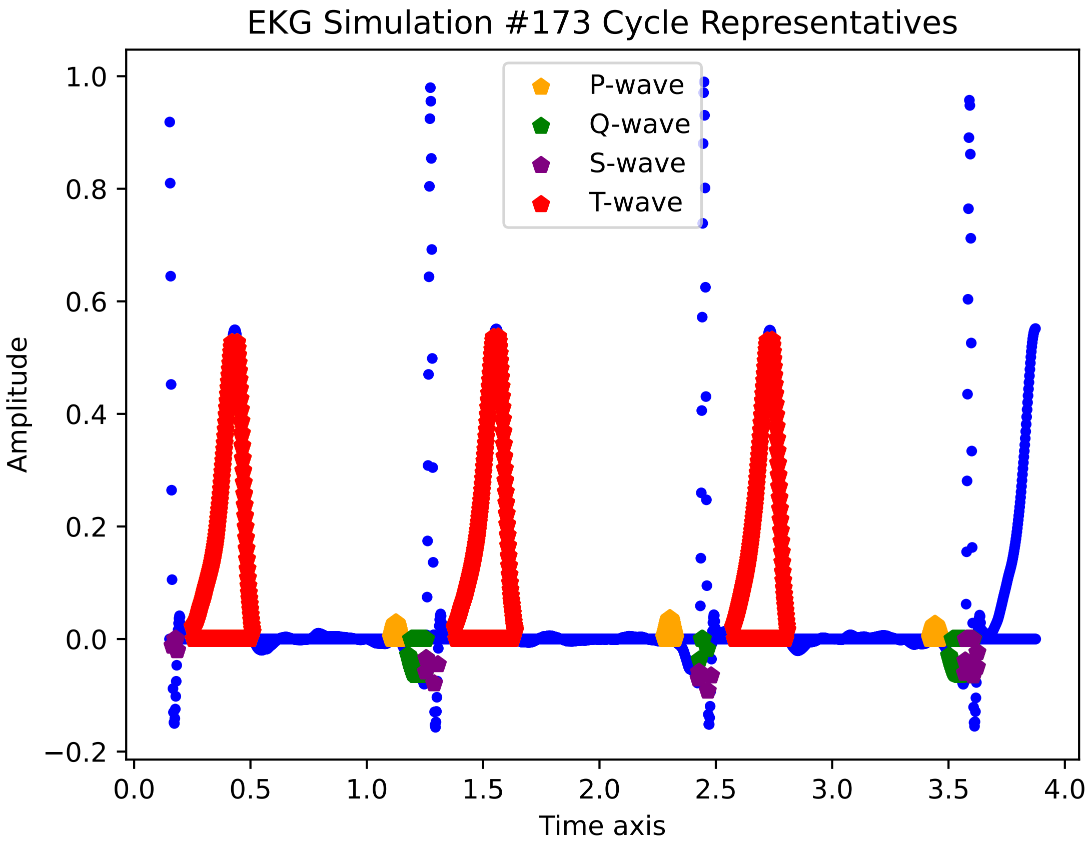
EKG Simulation #173 Cycle Representatives
P-wave
Q-wave
S-wave
T-wave
Amplitude
Time axis
1.0
0.8
0.6
0.4
0.2
0.0
−0.2
0.0
0.5
1.0
1.5
2.0
2.5
3.0
3.5
4.0

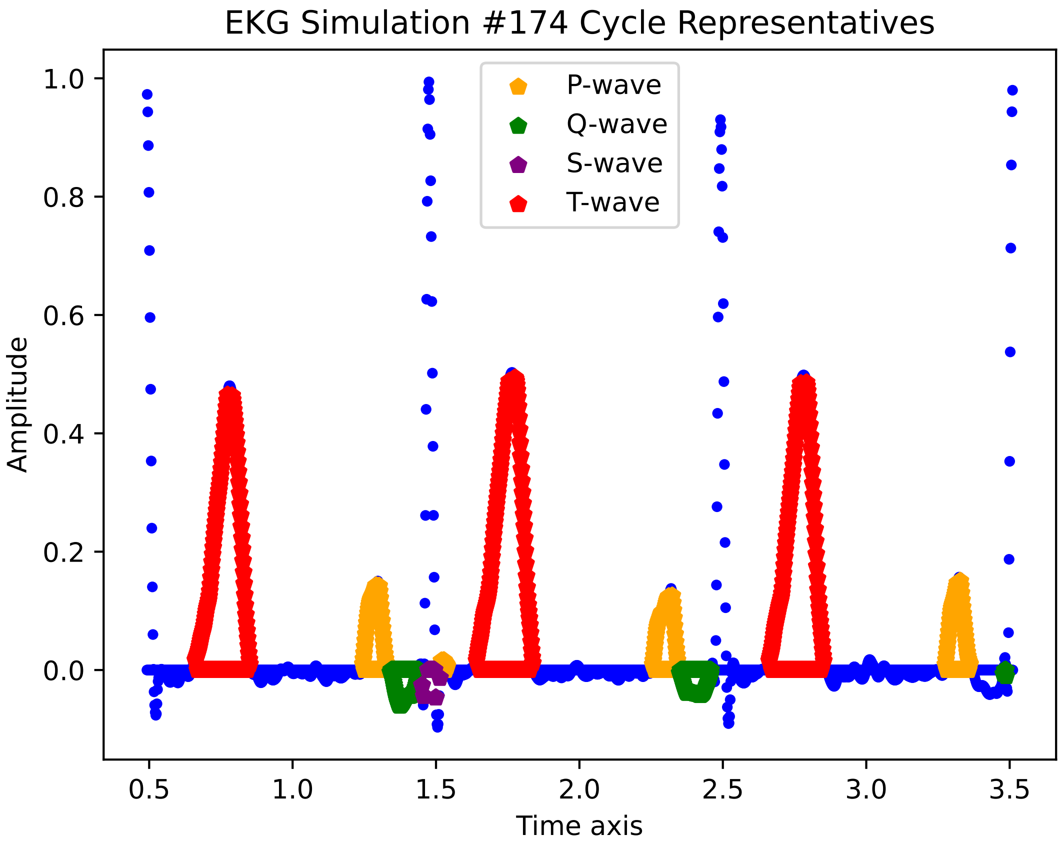
EKG Simulation #174 Cycle Representatives
P-wave
Q-wave
S-wave
T-wave
Amplitude
1.0
0.8
0.6
0.4
0.2
0.0
0.5
1.0
1.5
2.0
2.5
3.0
3.5
Time axis

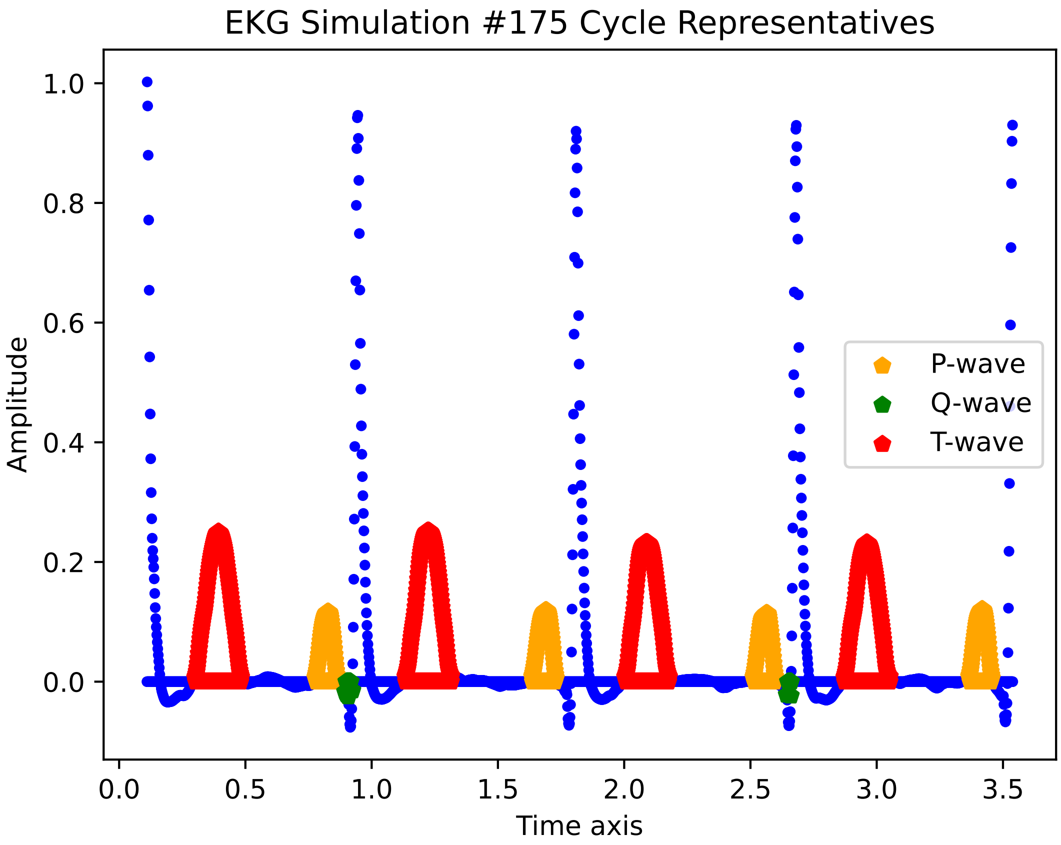
EKG Simulation #175 Cycle Representatives
Amplitude
Time axis
P-wave
Q-wave
T-wave
1.0
0.8
0.6
0.4
0.2
0.0
0.0
0.5
1.0
1.5
2.0
2.5
3.0
3.5

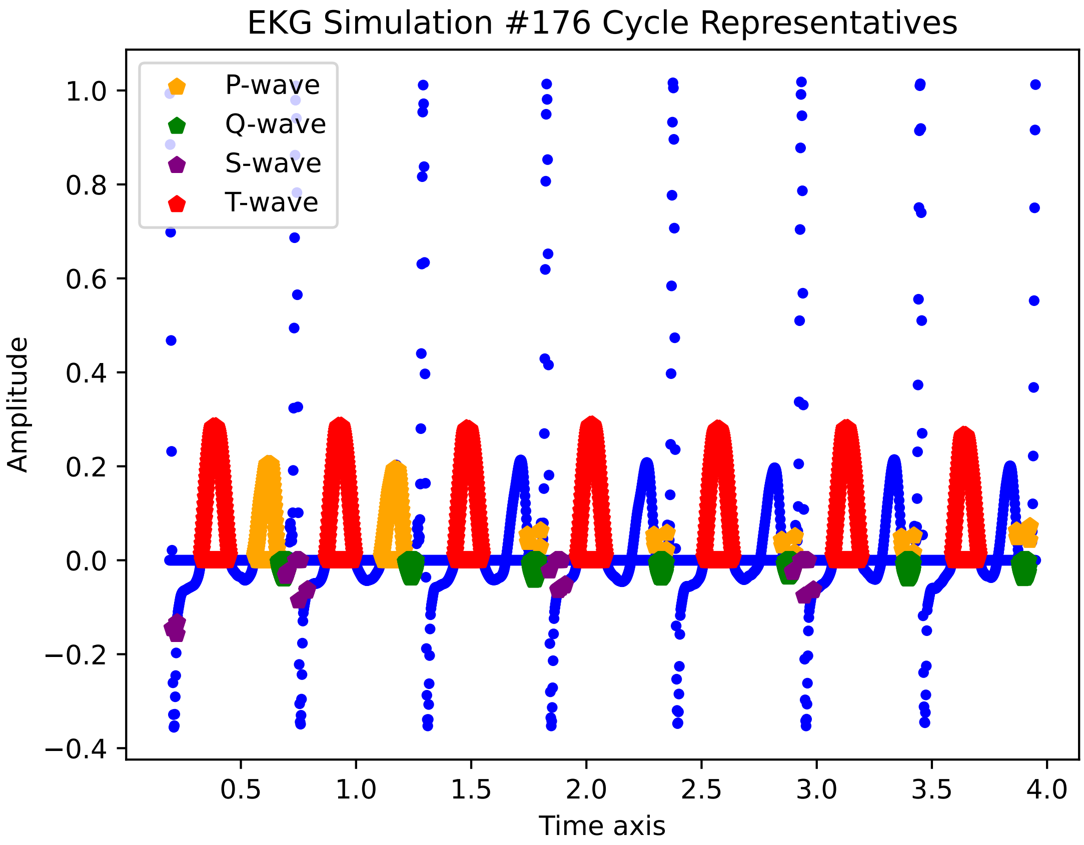
EKG Simulation #176 Cycle Representatives
P-wave
Q-wave
S-wave
T-wave
Amplitude
Time axis
1.0
0.8
0.6
0.4
0.2
0.0
−0.2
−0.4
0.5
1.0
1.5
2.0
2.5
3.0
3.5
4.0

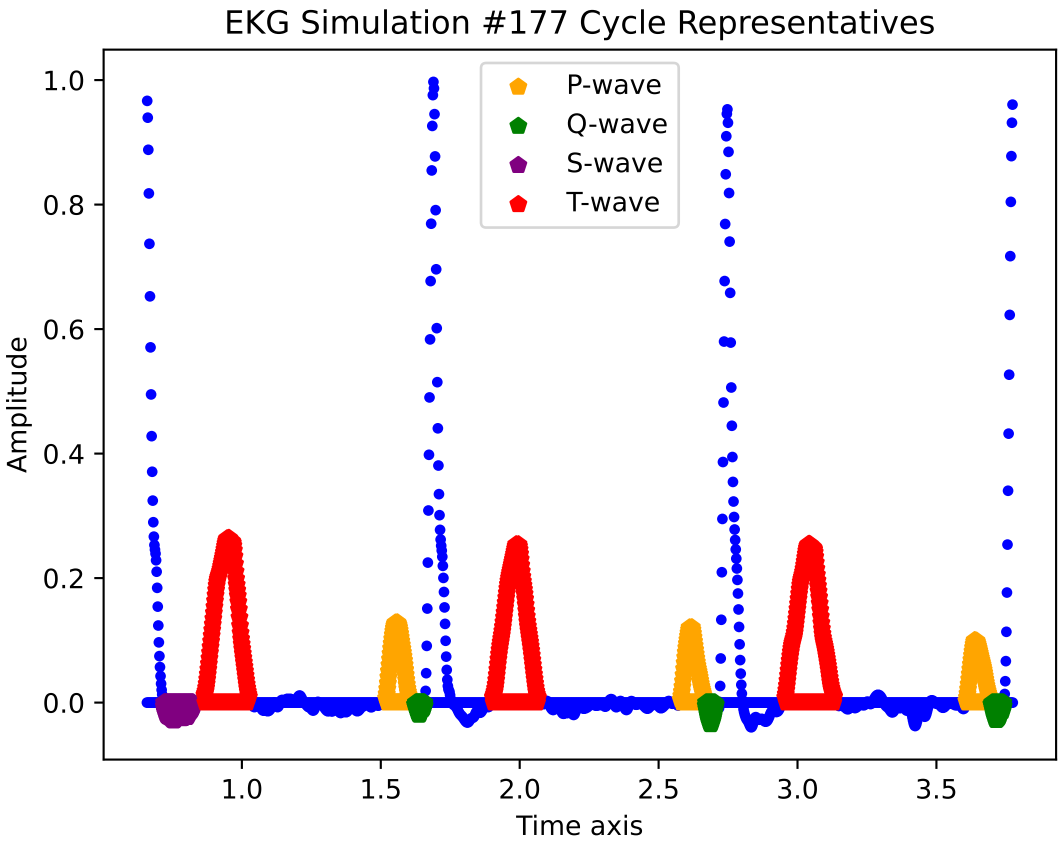

EKG Simulation #177 Cycle Representatives
Amplitude
Time axis
P-wave
Q-wave
S-wave
T-wave
1.0
0.8
0.6
0.4
0.2
0.0
1.0
1.5
2.0
2.5
3.0
3.5

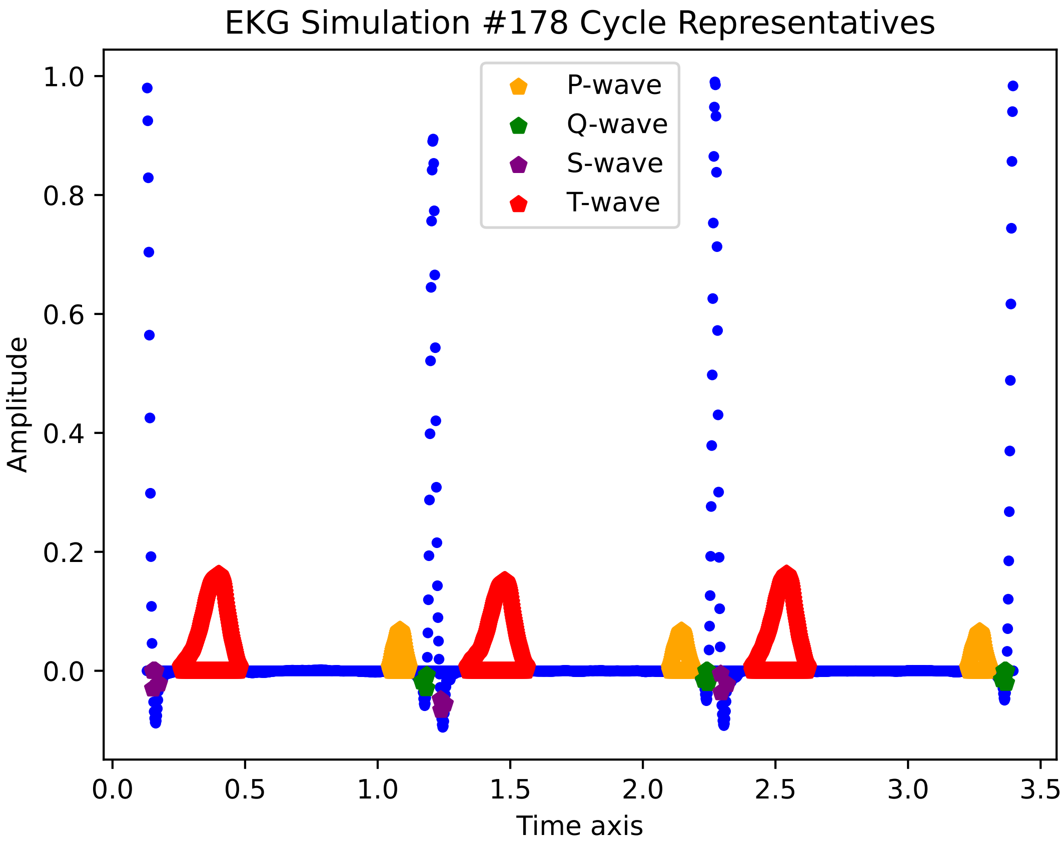
EKG Simulation #178 Cycle Representatives
P-wave
Q-wave
S-wave
T-wave
Amplitude
Time axis
1.0
0.8
0.6
0.4
0.2
0.0
0.0
0.5
1.0
1.5
2.0
2.5
3.0
3.5

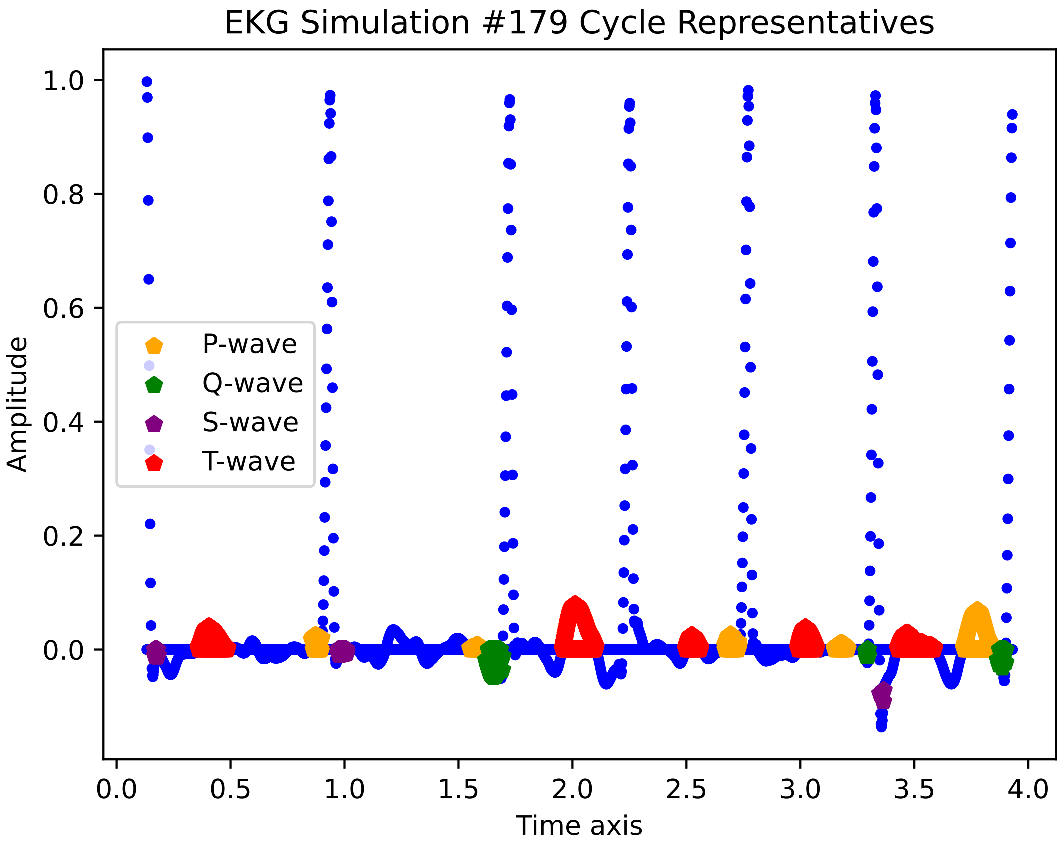

EKG Simulation #179 Cycle Representatives
Amplitude
Time axis
P-wave
Q-wave
S-wave
T-wave
1.0
0.8
0.6
0.4
0.2
0.0
0.0
0.5
1.0
1.5
2.0
2.5
3.0
3.5
4.0

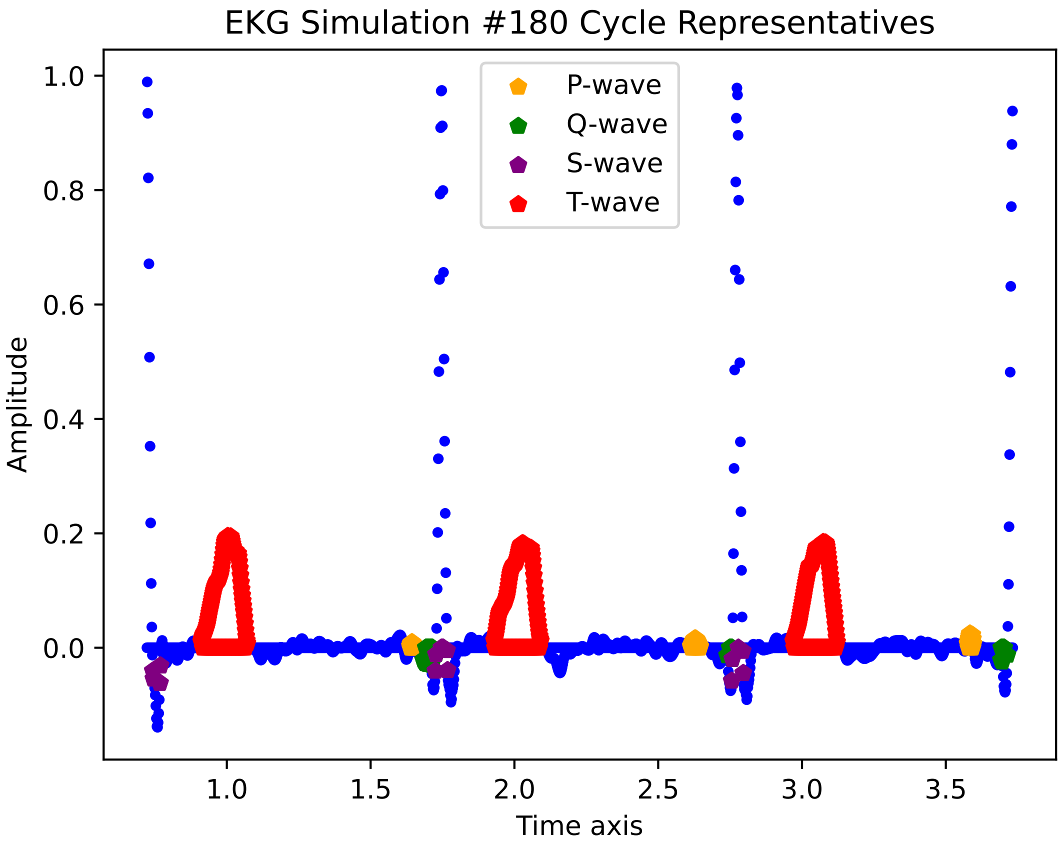
EKG Simulation #180 Cycle Representatives
P-wave
Q-wave
S-wave
T-wave
Amplitude
Time axis
1.0
0.8
0.6
0.4
0.2
0.0
1.0
1.5
2.0
2.5
3.0
3.5

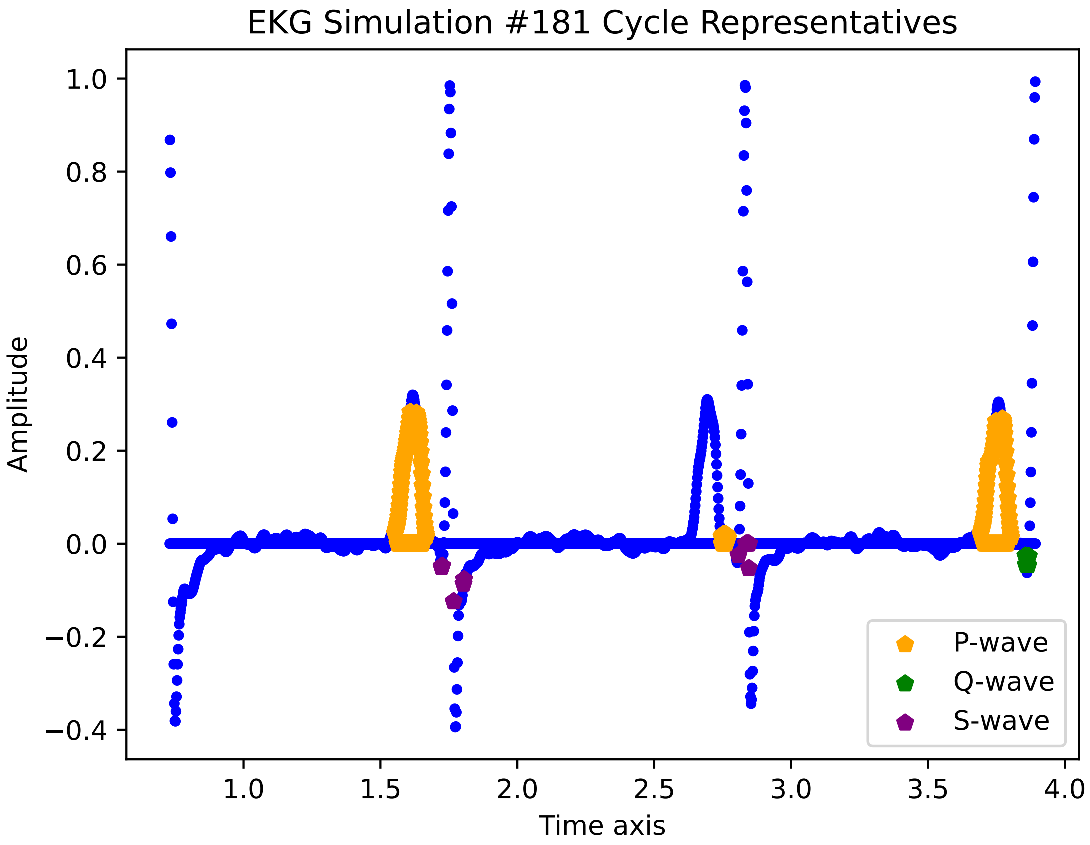
EKG Simulation #181 Cycle Representatives
Amplitude
Time axis
1.0
0.8
0.6
0.4
0.2
0.0
−0.2
−0.4
1.0
1.5
2.0
2.5
3.0
3.5
4.0
P-wave
Q-wave
S-wave

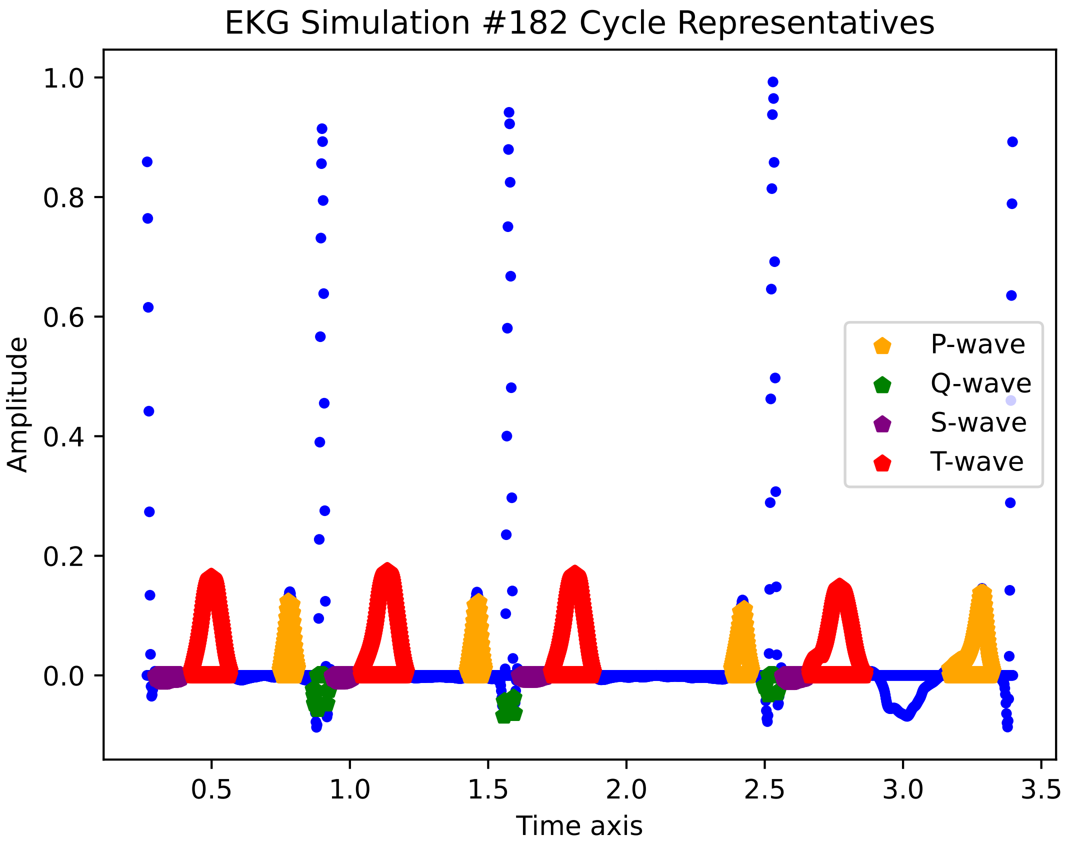
EKG Simulation #182 Cycle Representatives
Amplitude
Time axis
1.0
0.8
0.6
0.4
0.2
0.0
0.5
1.0
1.5
2.0
2.5
3.0
3.5
P-wave
Q-wave
S-wave
T-wave

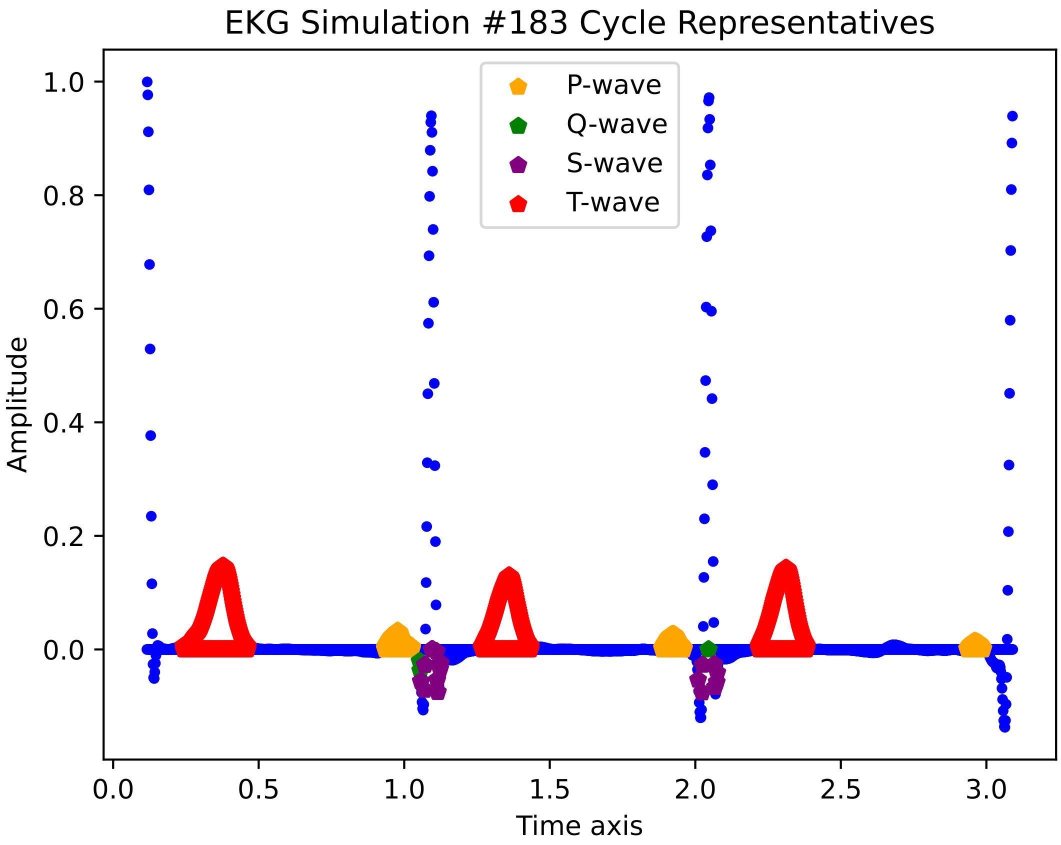

EKG Simulation #183 Cycle Representatives
P-wave
Q-wave
S-wave
T-wave
Amplitude
Time axis
1.0
0.8
0.6
0.4
0.2
0.0
0.0
0.5
1.0
1.5
2.0
2.5
3.0

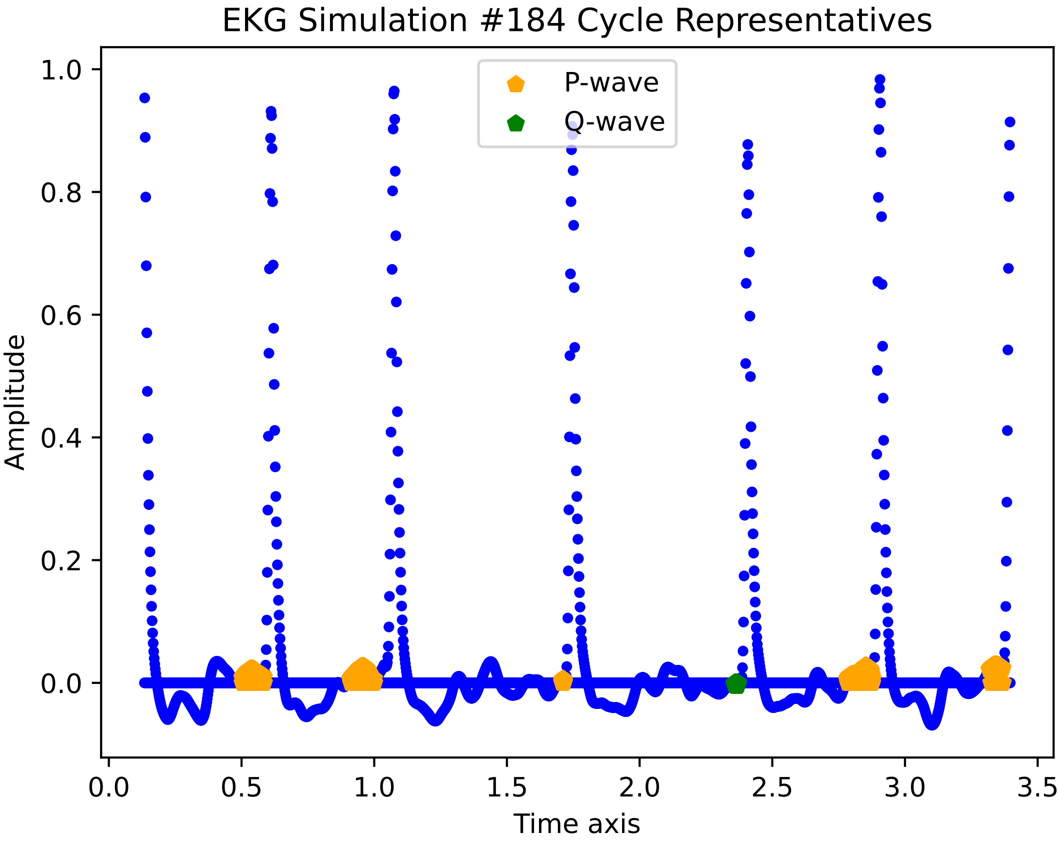

EKG Simulation #184 Cycle Representatives
P-wave
Q-wave
Amplitude
Time axis
1.0
0.8
0.6
0.4
0.2
0.0
0.0
0.5
1.0
1.5
2.0
2.5
3.0
3.5

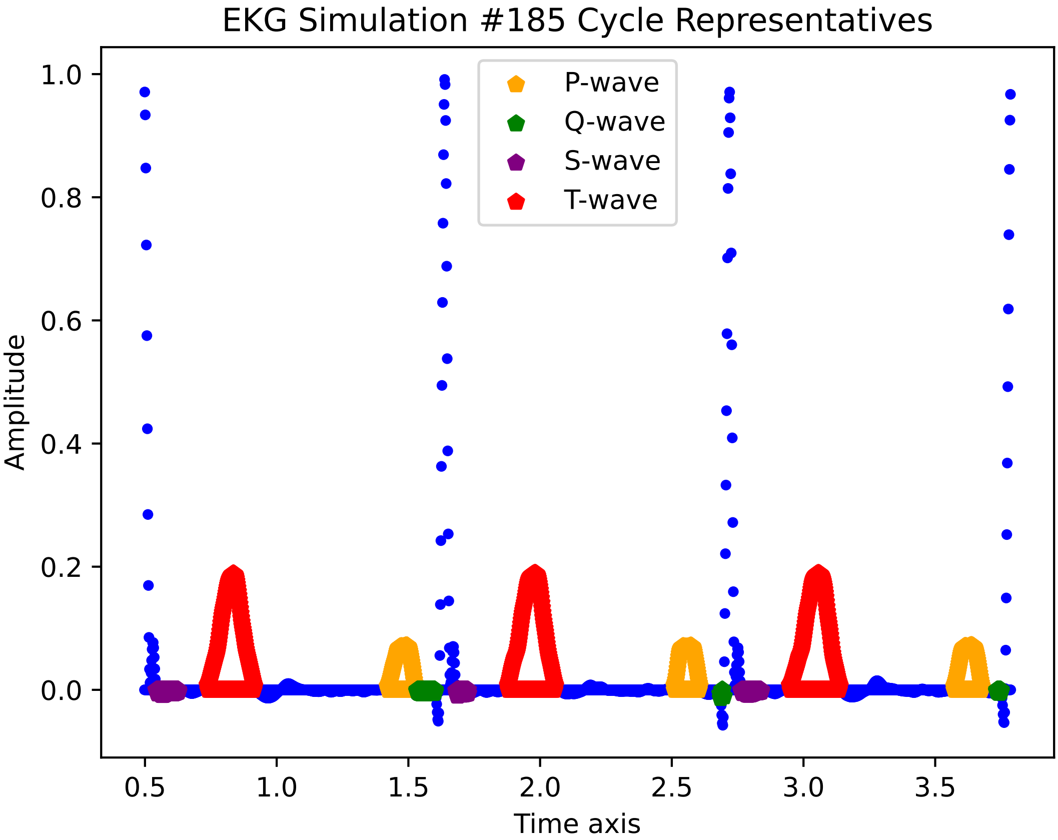
EKG Simulation #185 Cycle Representatives
P-wave
Q-wave
S-wave
T-wave
Amplitude
Time axis
1.0
0.8
0.6
0.4
0.2
0.0
0.5
1.0
1.5
2.0
2.5
3.0
3.5

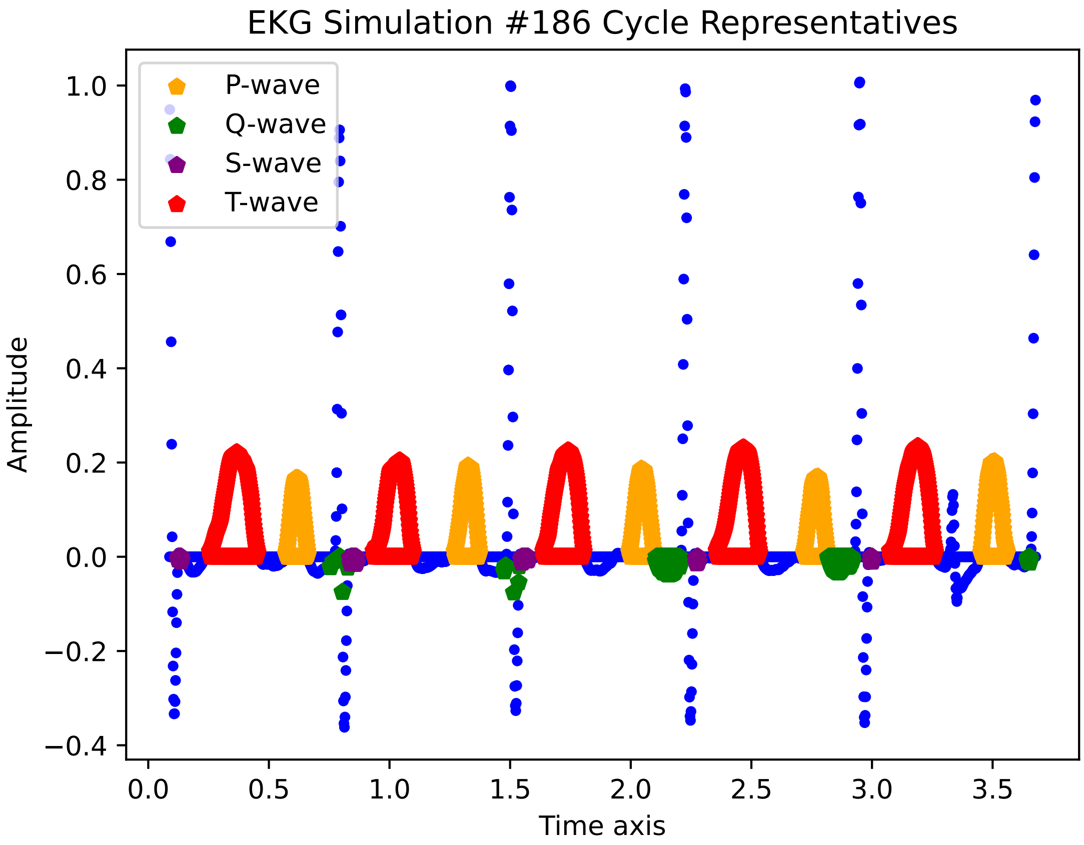
EKG Simulation #186 Cycle Representatives
P-wave
Q-wave
S-wave
T-wave
Amplitude
Time axis
1.0
0.8
0.6
0.4
0.2
0.0
−0.2
−0.4
0.0
0.5
1.0
1.5
2.0
2.5
3.0
3.5

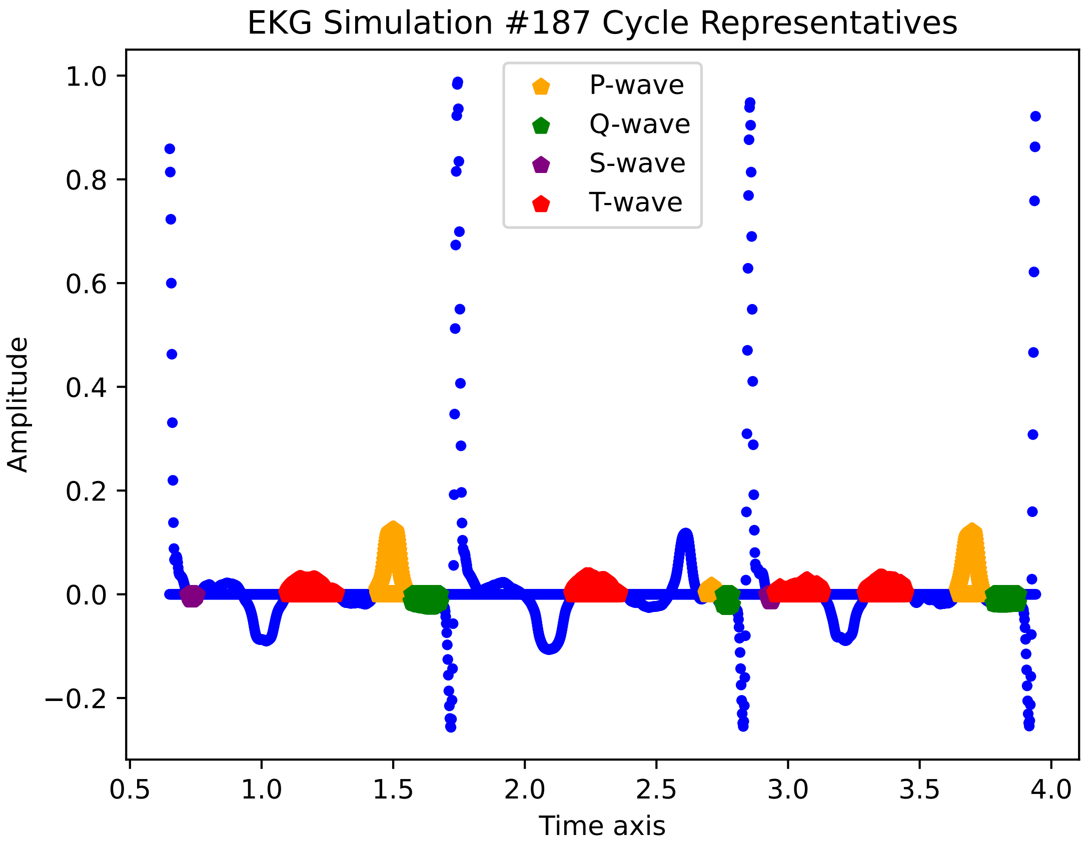
EKG Simulation #187 Cycle Representatives
P-wave
Q-wave
S-wave
T-wave
Amplitude
1.0
0.8
0.6
0.4
0.2
0.0
−0.2
0.5
1.0
1.5
2.0
2.5
3.0
3.5
4.0
Time axis

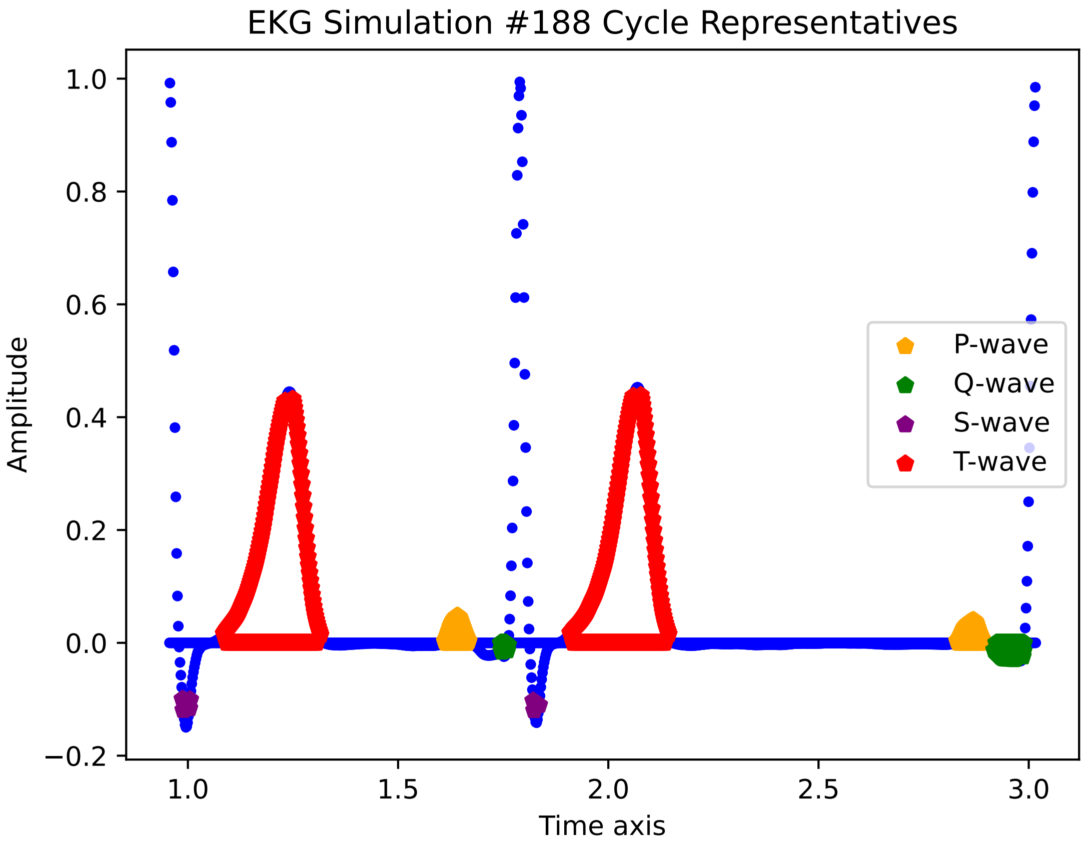
EKG Simulation #188 Cycle Representatives
Amplitude
Time axis
1.0
0.8
0.6
0.4
0.2
0.0
−0.2
1.0
1.5
2.0
2.5
3.0
P-wave
Q-wave
S-wave
T-wave

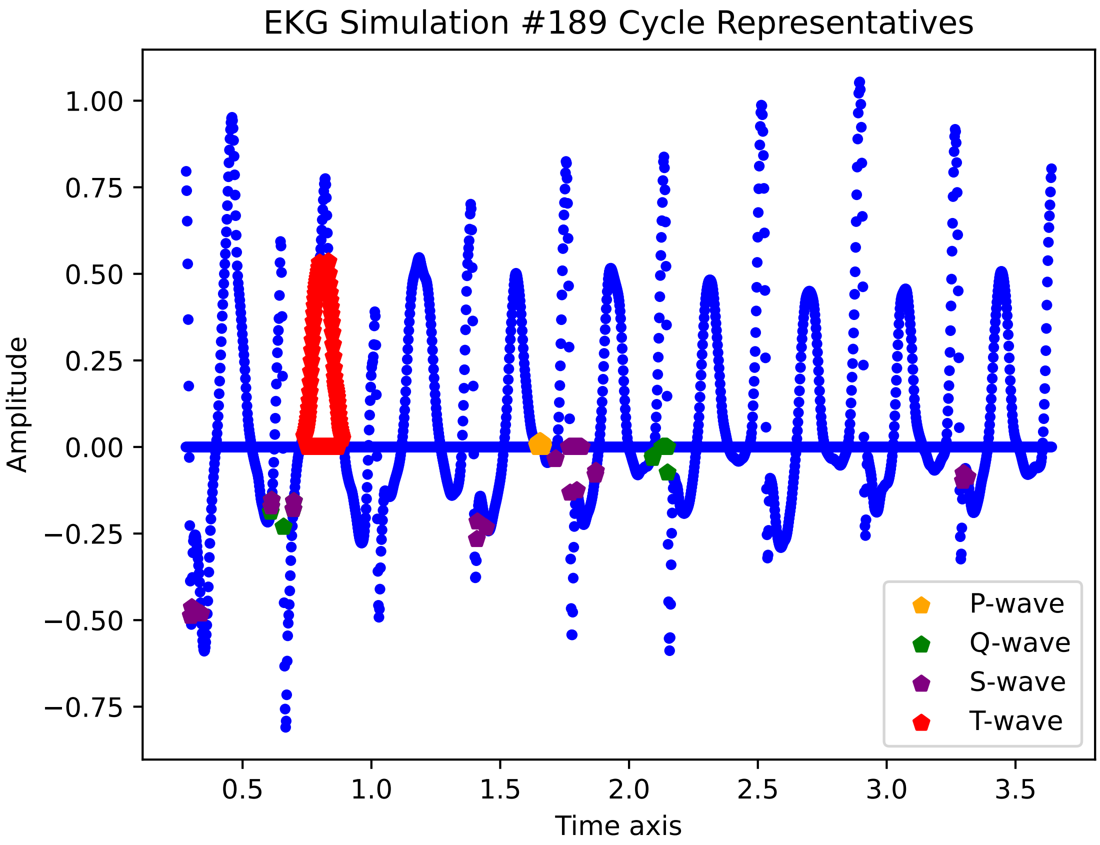
EKG Simulation #189 Cycle Representatives
Amplitude
Time axis
1.00
0.75
0.50
0.25
0.00
−0.25
−0.50
−0.75
0.5
1.0
1.5
2.0
2.5
3.0
3.5
P-wave
Q-wave
S-wave
T-wave

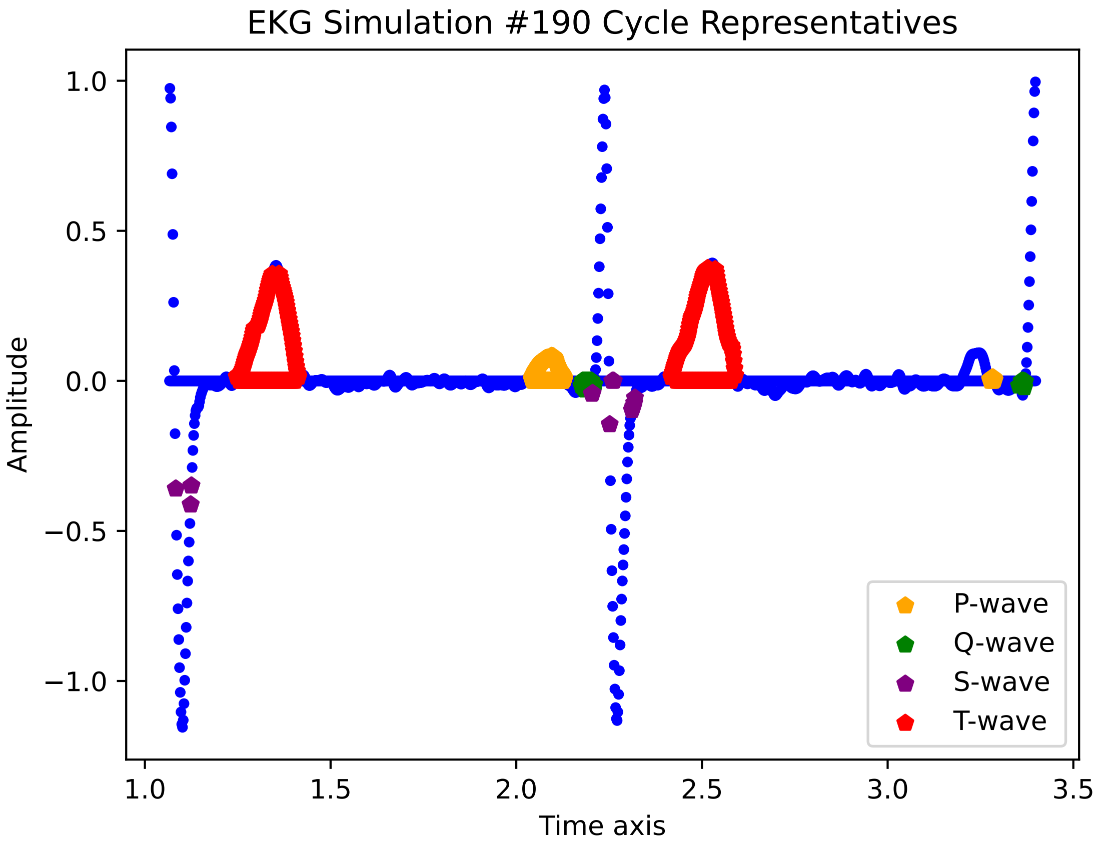
EKG Simulation #190 Cycle Representatives
1.0
0.5
0.0
−0.5
−1.0
Amplitude
1.0
1.5
2.0
2.5
3.0
3.5
Time axis
P-wave
Q-wave
S-wave
T-wave

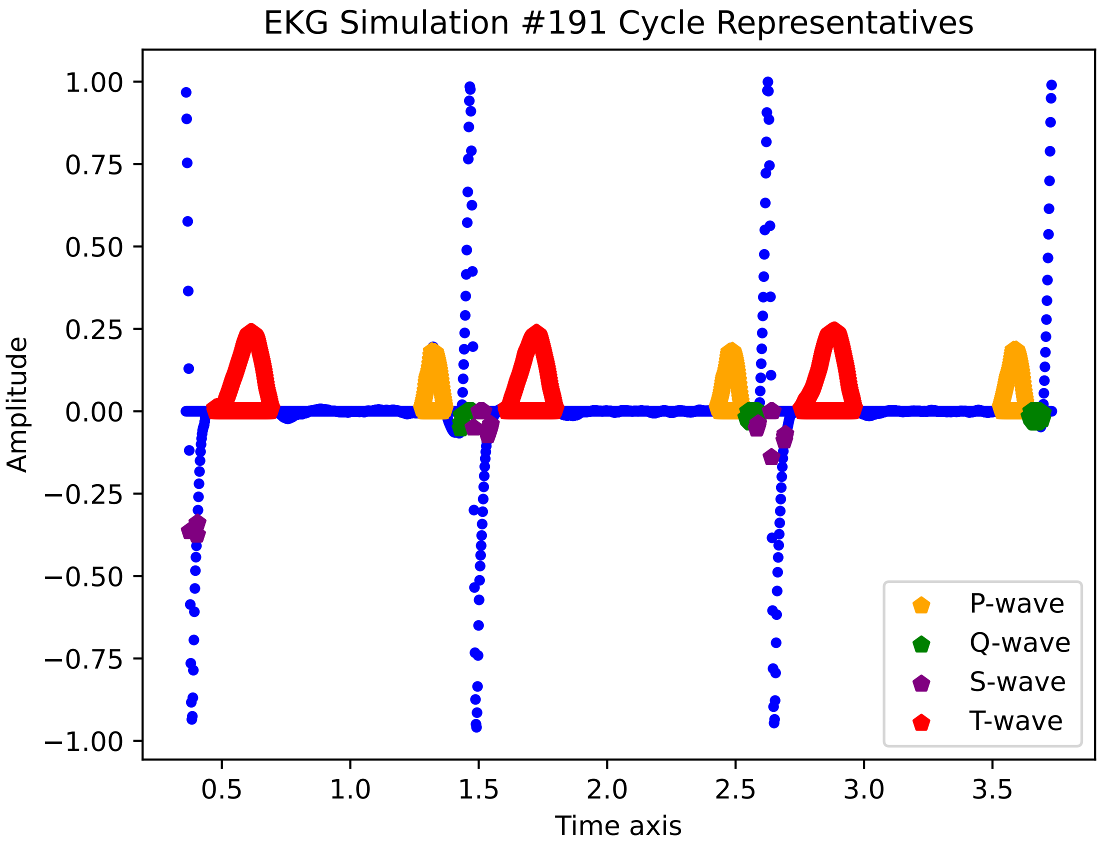
EKG Simulation #191 Cycle Representatives
Amplitude
Time axis
1.00
0.75
0.50
0.25
0.00
−0.25
−0.50
−0.75
−1.00
0.5
1.0
1.5
2.0
2.5
3.0
3.5
P-wave
Q-wave
S-wave
T-wave

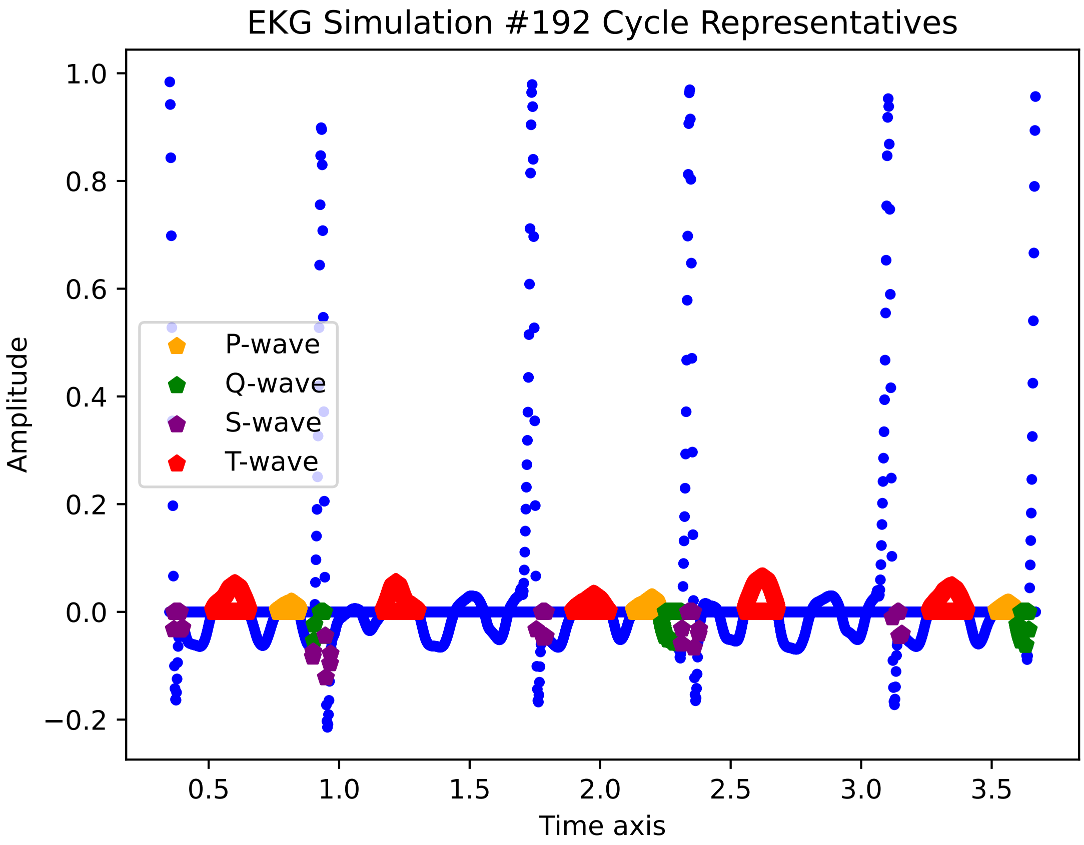

EKG Simulation #192 Cycle Representatives
Amplitude
Time axis
P-wave
Q-wave
S-wave
T-wave
1.0
0.8
0.6
0.4
0.2
0.0
−0.2
0.5
1.0
1.5
2.0
2.5
3.0
3.5

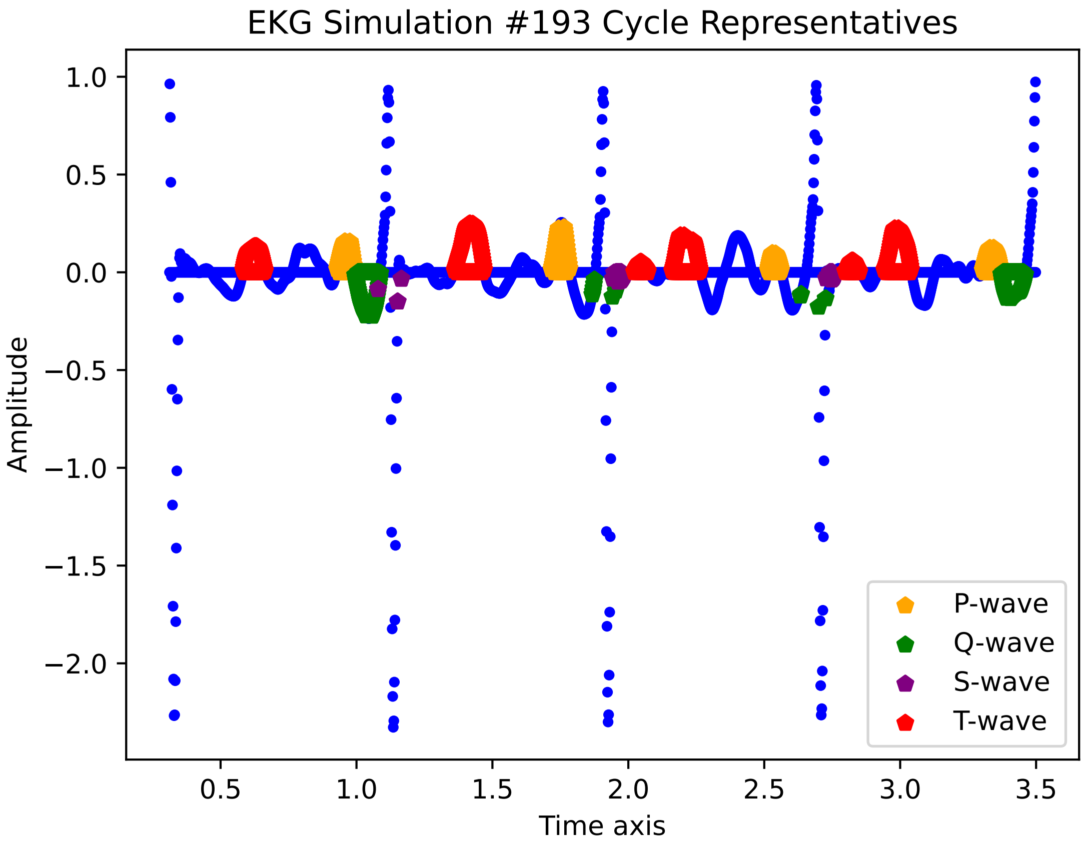

EKG Simulation #193 Cycle Representatives
Amplitude
Time axis
1.0
0.5
0.0
−0.5
−1.0
−1.5
−2.0
0.5
1.0
1.5
2.0
2.5
3.0
3.5
P-wave
Q-wave
S-wave
T-wave

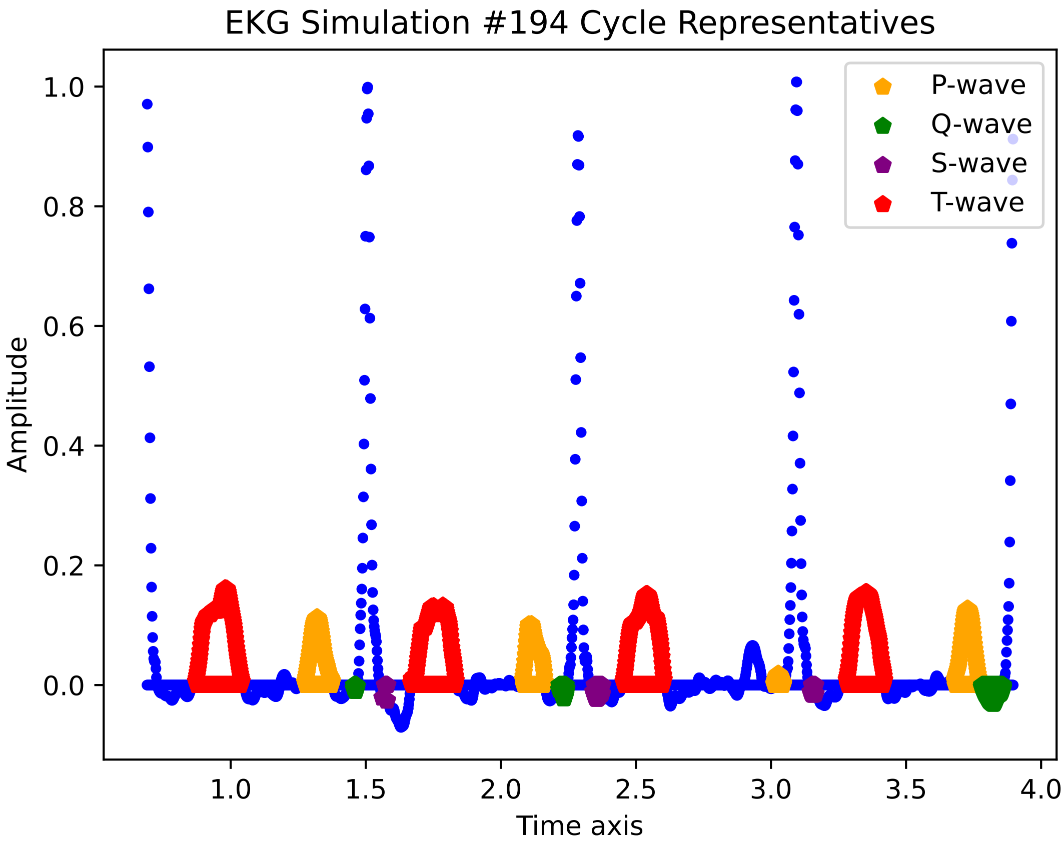

EKG Simulation #194 Cycle Representatives
Amplitude
Time axis
P-wave
Q-wave
S-wave
T-wave
1.0
0.8
0.6
0.4
0.2
0.0
1.0
1.5
2.0
2.5
3.0
3.5
4.0

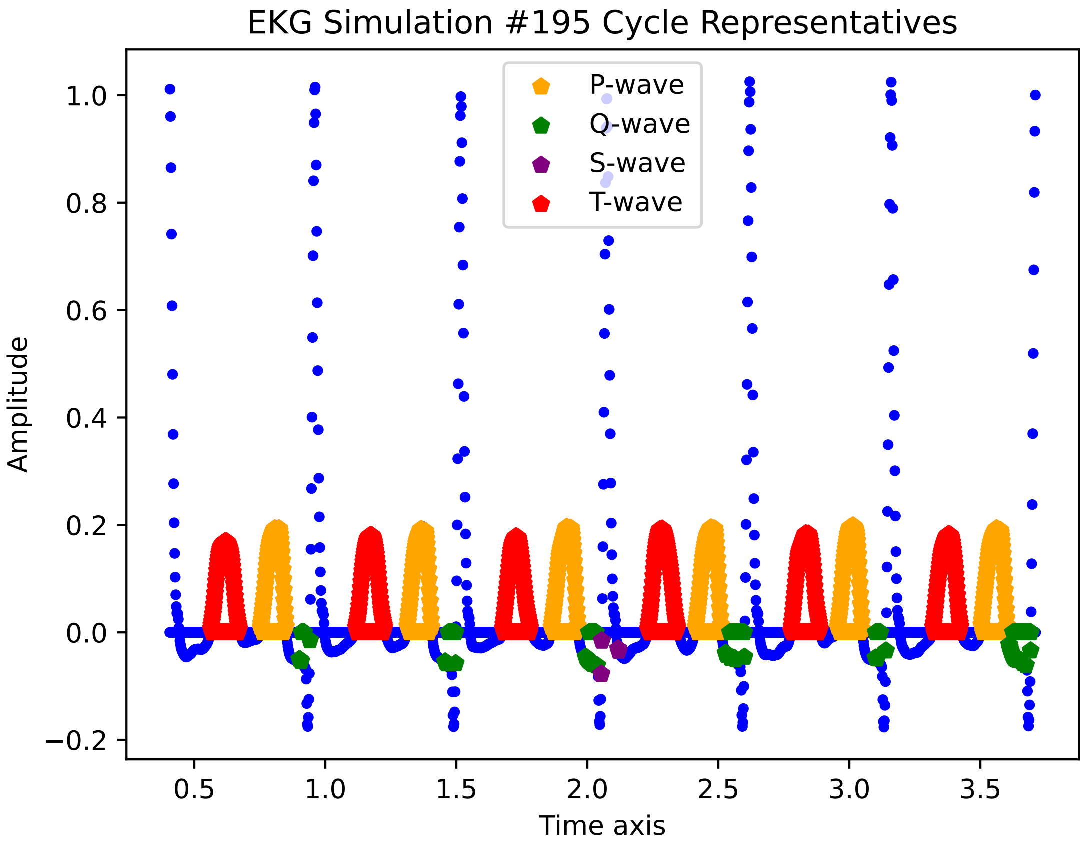
EKG Simulation #195 Cycle Representatives
P-wave
Q-wave
S-wave
T-wave
Amplitude
Time axis
1.0
0.8
0.6
0.4
0.2
0.0
−0.2
0.5
1.0
1.5
2.0
2.5
3.0
3.5

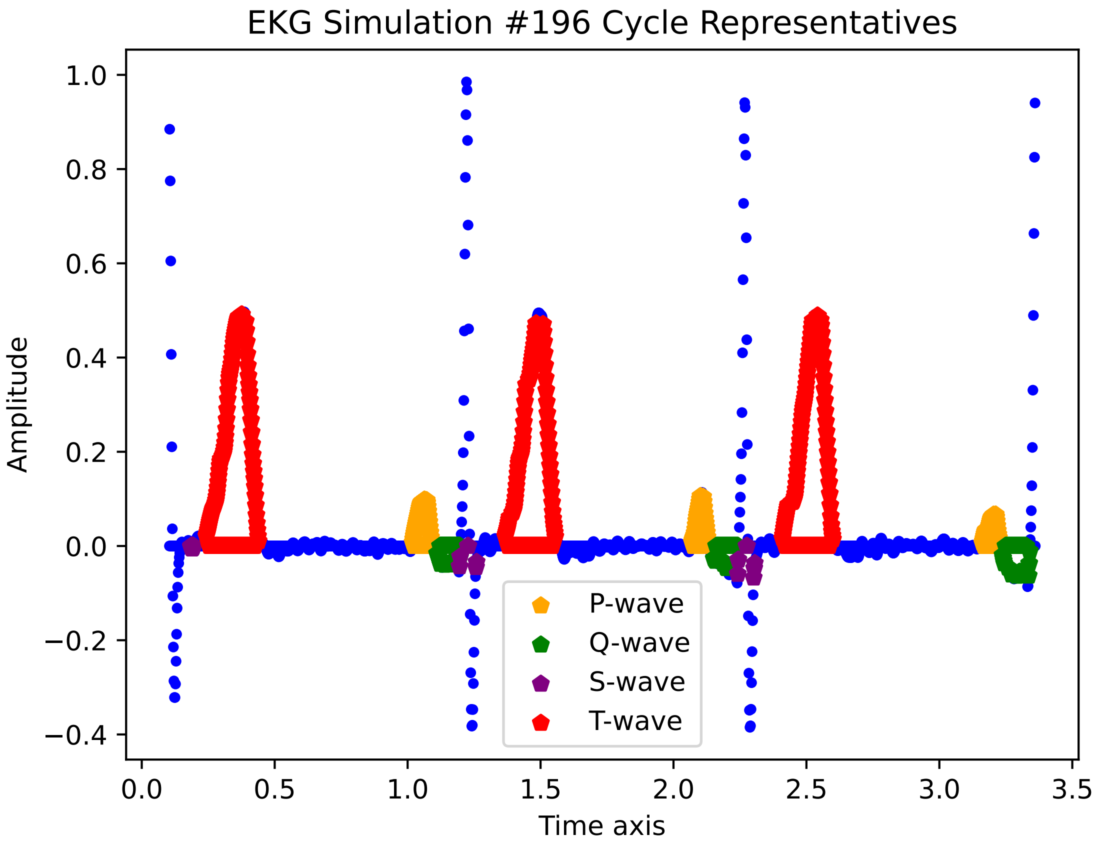
EKG Simulation #196 Cycle Representatives
Amplitude
Time axis
1.0
0.8
0.6
0.4
0.2
0.0
−0.2
−0.4
0.0
0.5
1.0
1.5
2.0
2.5
3.0
3.5
P-wave
Q-wave
S-wave
T-wave

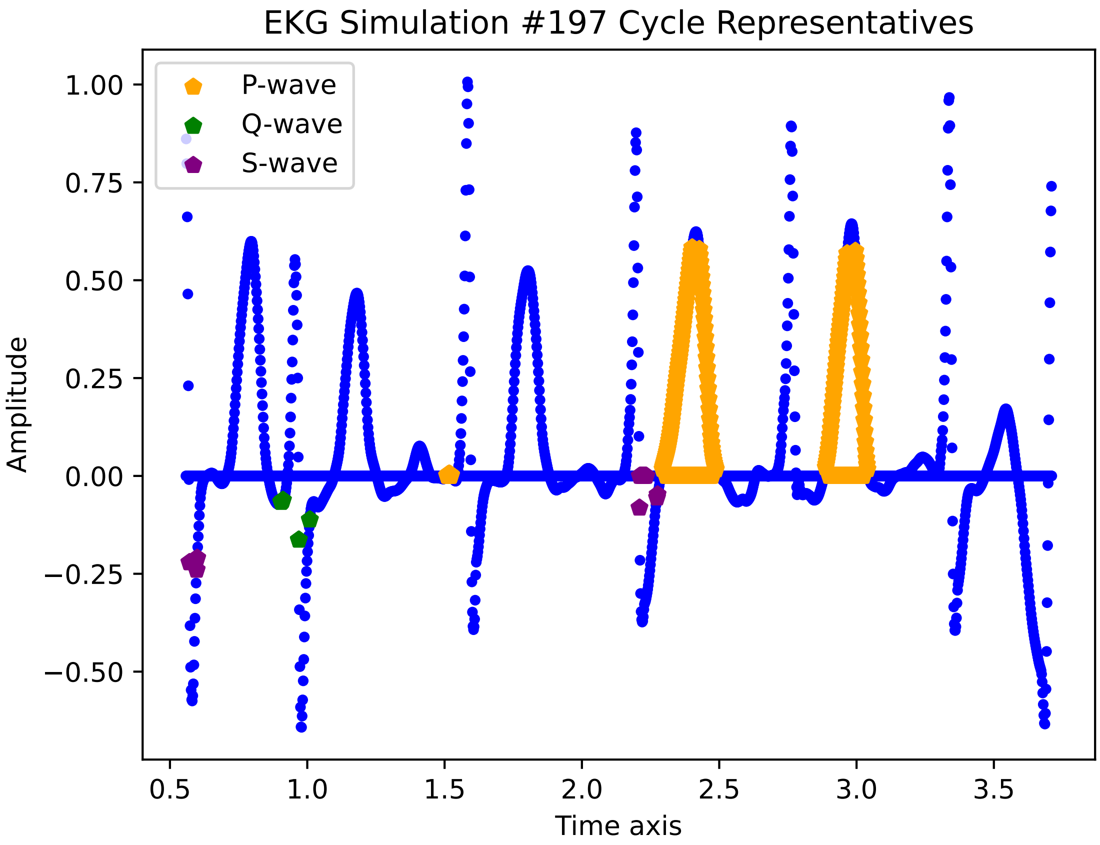
EKG Simulation #197 Cycle Representatives
P-wave
Q-wave
S-wave
1.00
0.75
0.50
0.25
0.00
−0.25
−0.50
Amplitude
0.5
1.0
1.5
2.0
2.5
3.0
3.5
Time axis

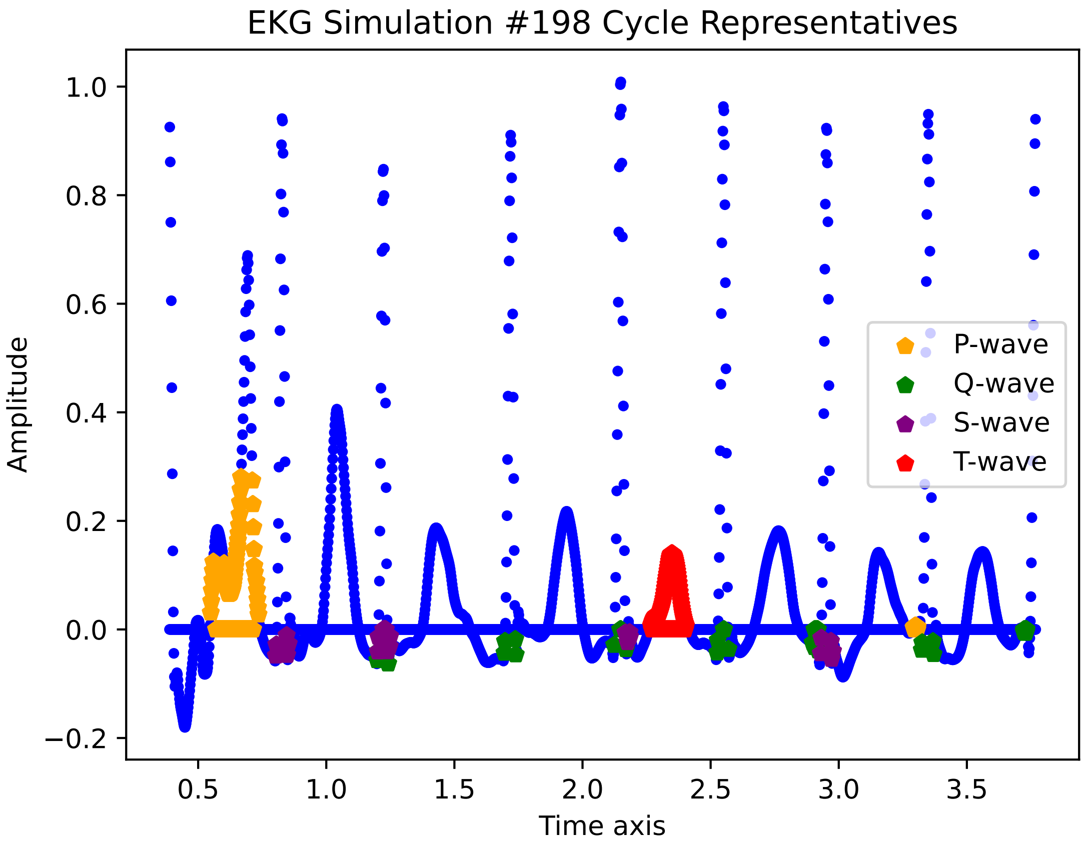

EKG Simulation #198 Cycle Representatives
Amplitude
Time axis
P-wave
Q-wave
S-wave
T-wave
1.0
0.8
0.6
0.4
0.2
0.0
−0.2
0.5
1.0
1.5
2.0
2.5
3.0
3.5

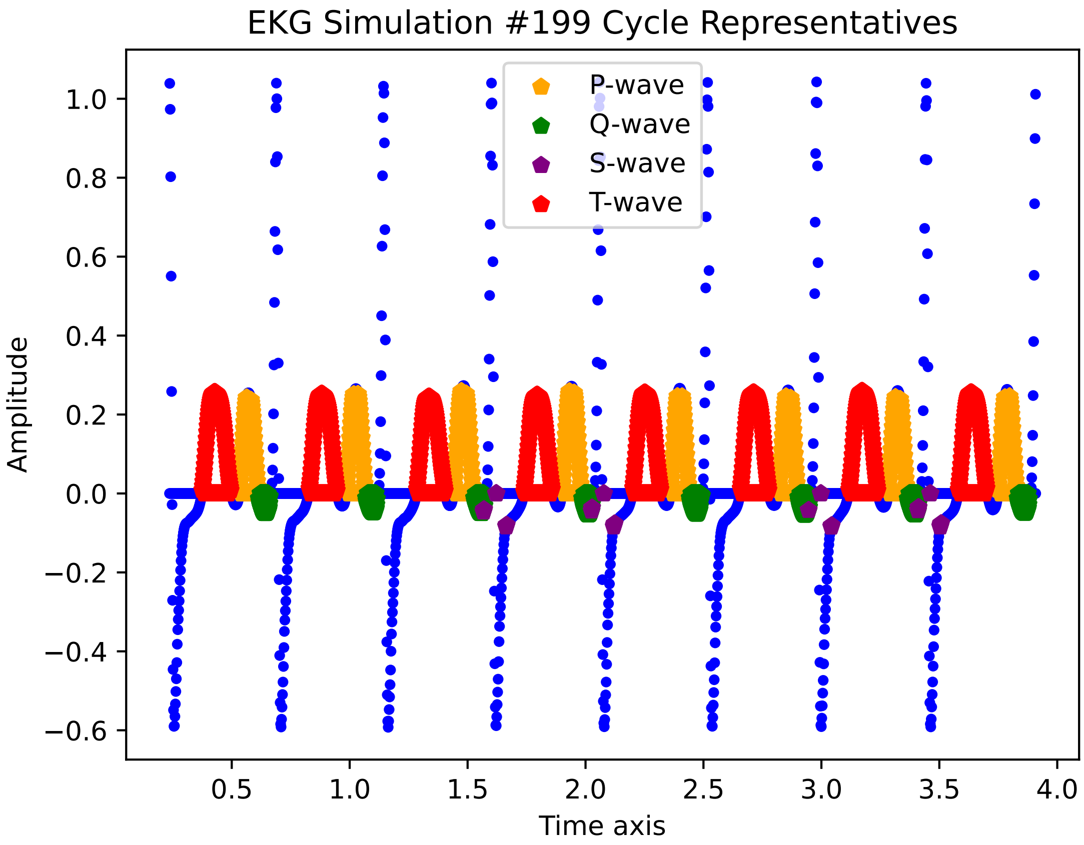

EKG Simulation #199 Cycle Representatives
P-wave
Q-wave
S-wave
T-wave
Amplitude
Time axis
1.0
0.8
0.6
0.4
0.2
0.0
−0.2
−0.4
−0.6
0.5
1.0
1.5
2.0
2.5
3.0
3.5
4.0

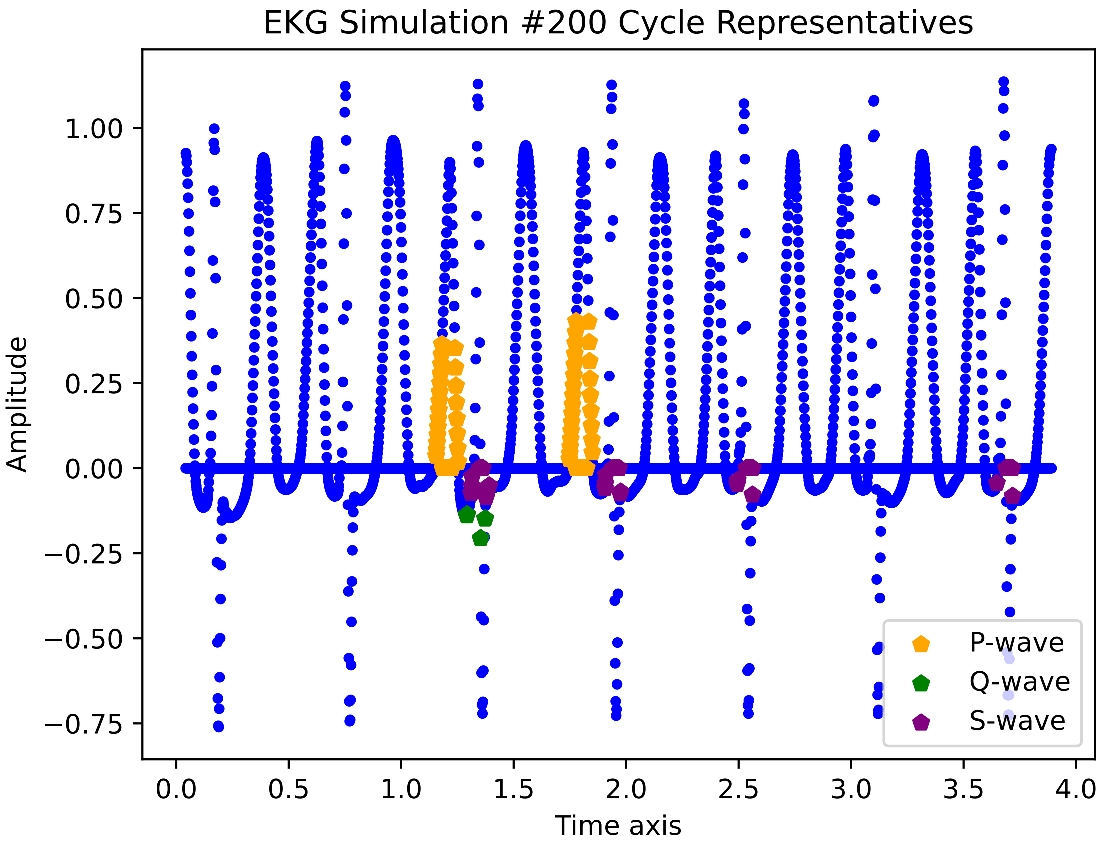
EKG Simulation #200 Cycle Representatives
Amplitude
Time axis
1.00
0.75
0.50
0.25
0.00
−0.25
−0.50
−0.75
0.0
0.5
1.0
1.5
2.0
2.5
3.0
3.5
4.0
P-wave
Q-wave
S-wave
